# Molecular epidemiology of peste des petits ruminants virus emergence in critically endangered Mongolian saiga antelope and other wild ungulates

**DOI:** 10.1101/2021.05.08.443216

**Authors:** Camilla T. O. Benfield, Sarah Hill, Munkduuren Shatar, Enkhtuvshin Shiilegdamba, Batchuluun Damdinjav, Amanda Fine, Brian Willett, Richard Kock, Arnaud Bataille

**Author notes:** Corresponding author, (CTOB).

## Abstract

Peste des petits ruminants virus (PPRV) causes disease in domestic and wild ungulates, is the target of a global eradication programme and threatens biodiversity. Understanding the epidemiology and evolution of PPRV in wildlife is important, but hampered by the paucity of wildlife-origin PPRV genomes. In this study, full PPRV genomes were generated from three Mongolian saiga antelope, one Siberian ibex and one goitered gazelle from the 2016-2017 PPRV outbreak. Phylogenetic analysis showed that for Mongolian and Chinese PPRV since 2013, the wildlife and livestock-origin genomes were closely related and interspersed. There was strong phylogenetic support for a monophyletic group of PPRV from Mongolian wildlife and livestock, belonging to clade of lineage IV PPRV from livestock and wildlife from China since 2013. Discrete diffusion analysis found strong support for PPRV spread into Mongolia from China and phylogeographic analysis indicated Xinjiang Province as the most likely origin, although genomic surveillance for PPRV is poor and lack of sampling from other regions could bias this result. Times of most recent common ancestor (TMRCA) were June 2015 (95% HPD: August 2014 – March 2016) for all Mongolian PPRV genomes and May 2016 (95% HPD: October 2015 – October 2016) for Mongolian wildlife-origin PPRV. This suggests that PPRV was circulating undetected in Mongolia for at least six months before the first reported outbreak in August 2016, and that wildlife were likely infected before livestock vaccination began in October 2016. Finally, genetic variation and positively-selected sites were identified that might be related to PPRV emergence in Mongolian wildlife. This study is the first to sequence multiple PPRV genomes from a wildlife outbreak, across several host species. Additional full PPRV genomes and associated metadata from the livestock-wildlife interface are needed to enhance the power of molecular epidemiology, support PPRV eradication and safeguard the health of the whole ungulate community.

**Author Summary:** Recent mass mortality of critically endangered Mongolian saiga antelope due to peste des petits ruminants virus (PPRV) has dramatically highlighted the threat this viral disease represents for biodiversity. The genome of viruses such as PPRV evolve fast, so virus genetic data gathered from infected animals can be used to trace disease spread between livestock and wildlife, and to determine if the virus is adapting to infect wildlife more efficiently. Here we obtained PPRV virus genomes from Mongolian wildlife and compared them with other published PPRV genomes. Using a molecular clock, we estimated that the disease was circulating in Mongolia well before it was first reported. Genetic analyses support the hypothesis of virus spread from livestock to wildlife, with genetic changes potentially helping infection in Asian wild ungulates. However, more PPR virus genomes and epidemiology data are needed from disease outbreaks in areas shared between livestock and wildlife to confirm these results and take efficient actions to safeguard the health of the whole ungulate community.

## Introduction

Peste des petits ruminants (PPR) is a contagious viral disease of sheep and goats with high morbidity and mortality rates, which is a major barrier to sustainable small ruminant production, dependent livelihoods and economies. Consequently, PPR is the only livestock disease currently targeted by a Global Eradication Programme (GEP), which aims to rid the world of PPR by 2030 through vaccination of livestock, and thereby contribute to achieving the Sustainable Development Goals. The etiological agent, peste des petits ruminants virus (PPRV), has a broad host range, with serological or virological evidence of natural infection in a growing list of wild species within the order Artiodactyla [1–4]. PPRV infection of both captive and free-ranging wildlife may result in severe outbreaks and mortality, threatening species’ survival and ecosystem integrity. PPRV has caused mass mortality of mountain caprine species categorised as vulnerable by the IUCN [5, 6], with > 1000 deaths of wild goats (*Capra aegagrus*) and sheep (*Ovis orientalis*) in Iran [7] and > 750 wild goats in Iraq [8]. Fatal PPR outbreaks have also been reported in free-ranging Sindh ibex (*Capra aegagrus blythi*) in Pakistan [9] and in ibex (*Capra ibex*) [10–12], bharal (*Pseudois nayaur*) [10–14], argali sheep (*Ovis ammon*) [10], goitered gazelle (*Gazella subgutturosa*) [10] and Przewalski’s gazelle (*Procapra przewalskii*) [15] in China. To date, the most devastating impact of PPRV on biodiversity was its emergence in the critically endangered Mongolian saiga antelope (*Saiga tatarica mongolica*) in 2016-2017, which caused a mass mortality event and contributed to loss of ~80% of the population [16, 17]. In contrast, clinical disease has not been confirmed in free-ranging wildlife in Africa, despite high apparent PPRV seropositivity in wildlife populations in East Africa [18, 19]. The only published disease outbreak in free-ranging African wildlife in Africa occurred in Dorcas gazelles (*Gazella dorcas*) in Dinder National Park, Sudan [20]. However, this was not supported by field data to confirm the nature of the epidemic or event, and so whether this represents true wildlife disease remains equivocal, whilst African species in captivity have been shown to express PPR disease in zoological collections in the Middle East [21–23]. Therefore, while it is now clear that PPRV poses a threat to biodiversity, the determinants of differential disease expression among wildlife hosts are not understood. There are also significant knowledge gaps regarding the role of wildlife in the epidemiology and evolution of PPRV. It remains unclear whether wildlife can maintain or transmit the virus to livestock, and thereby pose a threat to the PPR GEP.

It is important to assess the genetic diversity of PPRV to understand whether host range plasticity and viral virulence are linked to genetic changes in the virus. PPRV is a morbillivirus with a negative sense single-stranded RNA genome of approximately 16 kilobases, which encodes six structural proteins, the nucleocapsid (N), phosphoprotein (P), matrix (M), fusion (F), hemagglutinin (H), and polymerase (L) proteins, and two non-structural proteins, V and C. The infectivity of PPRV is mediated by its envelope glycoproteins, H and F, which are therefore key viral determinants of cellular and host tropism. H binds the morbillivirus receptors SLAM and nectin-4 on immune and epithelial host cells, respectively, while F mediates the subsequent membrane fusion events to enable cell entry. The efficiency of receptor usage and entry into target cells are likely to be critical barriers to the emergence of morbilliviruses in atypical hosts. A recent study showed that a single amino acid substitution in PPRV H enabled it to use human SLAM as an entry receptor [24]. Studies on the related morbillivirus canine distemper virus have also shown that only one or two amino acid changes in H are associated with host range expansion in nature [25, 26] or via *in vitro* adaptation [27]. The crystal structure of measles virus (MeV) H protein in complex with marmoset SLAM has been solved [28] and shows that the receptor binding domain (RBD) comprises four sites on MeV H which interact with SLAM and which are well conserved in PPRV H [24]. Several recent mutagenesis studies have also identified amino acid residues in PPRV H important for its ability to bind SLAM [29] and induce cell fusion [30]. In addition to cell entry, PPRV evidently requires efficient replicative and immune-evasive abilities for successful infection of atypical hosts, but the role of other viral genes in host range remains obscure.

PPRV is classified into 4 genetically distinct lineages, which can be discriminated based on phylogenetic analysis of short gene regions, often a few hundred nucleotides of the N gene [31, 32]. Lineage IV viruses have dominated both the host range and geographic expansion of PPRV seen in recent years and are now replacing other lineages in many African countries [33–36]. Understanding this expansion is critical to mitigate challenges to the PPR GEP and to understand the threat of PPRV to biodiversity. To do so necessitates the phylogenetic resolution provided by full genome sequencing using high coverage high throughput sequencing technologies [37], which is particularly important since such limited molecular epidemiological data on PPRV in wildlife exists at the global level. Earlier molecular evolutionary studies of PPRV based on full genomes have included a few wildlife-origin sequences [38, 39]. However, no studies have hitherto used phylogenomic approaches to address inter-species transmission patterns of PPRV.

In Mongolia, PPR was first confirmed in August 2016 (https://wahis.oie.int/#/report-info?reportld=8043), and a full PPRV genome was generated from livestock sampled in September 2016 [40]. The outbreak in Mongolian wildlife was laboratory-confirmed in December 2016 and led to mortalities of Mongolian saiga antelope (*Saiga tatarica mongolica*), goitered gazelle (*Gazella subgutturosa*), Siberian ibex (*Capra ibex sibirica*) and Argali (*Ovis ammon*), thought to have been caused by spillover of the virus from livestock and subsequent spread among wild ungulates (https://wahis.oie.int/#/report-info?reportId=10463, [17]). Previously, the only molecular data for PPRV from Mongolian wildlife were partial N gene sequences from two saiga antelope [17]. Here, we generated five novel full genome sequences for the PPRV which emerged in three species of Mongolian wildlife: saiga antelope, goitered gazelle and Siberian ibex. Using these sequences and all other PPRV genomes available in GenBank from both wildlife and livestock hosts, we performed phylogenetic and molecular evolutionary analyses to address PPRV emergence in Mongolian wildlife and dynamics at the livestock-wildlife interface.

## Results

### Tissue distribution of PPRV in Mongolian wildlife

Total RNA was extracted from tissue samples collected at necropsy from four saiga antelope, one goitered gazelle and one Siberian ibex (Table 1). To determine the tissue distribution of PPRV replication in the wildlife hosts, and select samples for whole genome sequencing, RT-PCR for a 350 nucleotide region of the N gene was performed on all available samples. Every tissue tested was RT PCR-positive (S1 Fig), with an amplicon of the expected size, namely liver and ocular swab from Siberian ibex; tongue, soft palate and lung samples from goitered gazelle, and tongue, soft palate, ocular and nasal swabs, gum scurf, mesenteric lymph node, spleen, liver, lung, heart and blood from saiga antelope. Nucleic acid sequencing showed that this N gene region was identical in all six wildlife hosts, and to the two published partial N gene sequences from saiga [17], and differed from the Mongolian livestock PPRV (KY888168.1) by two nucleotides (data not shown).

**Table 1.**
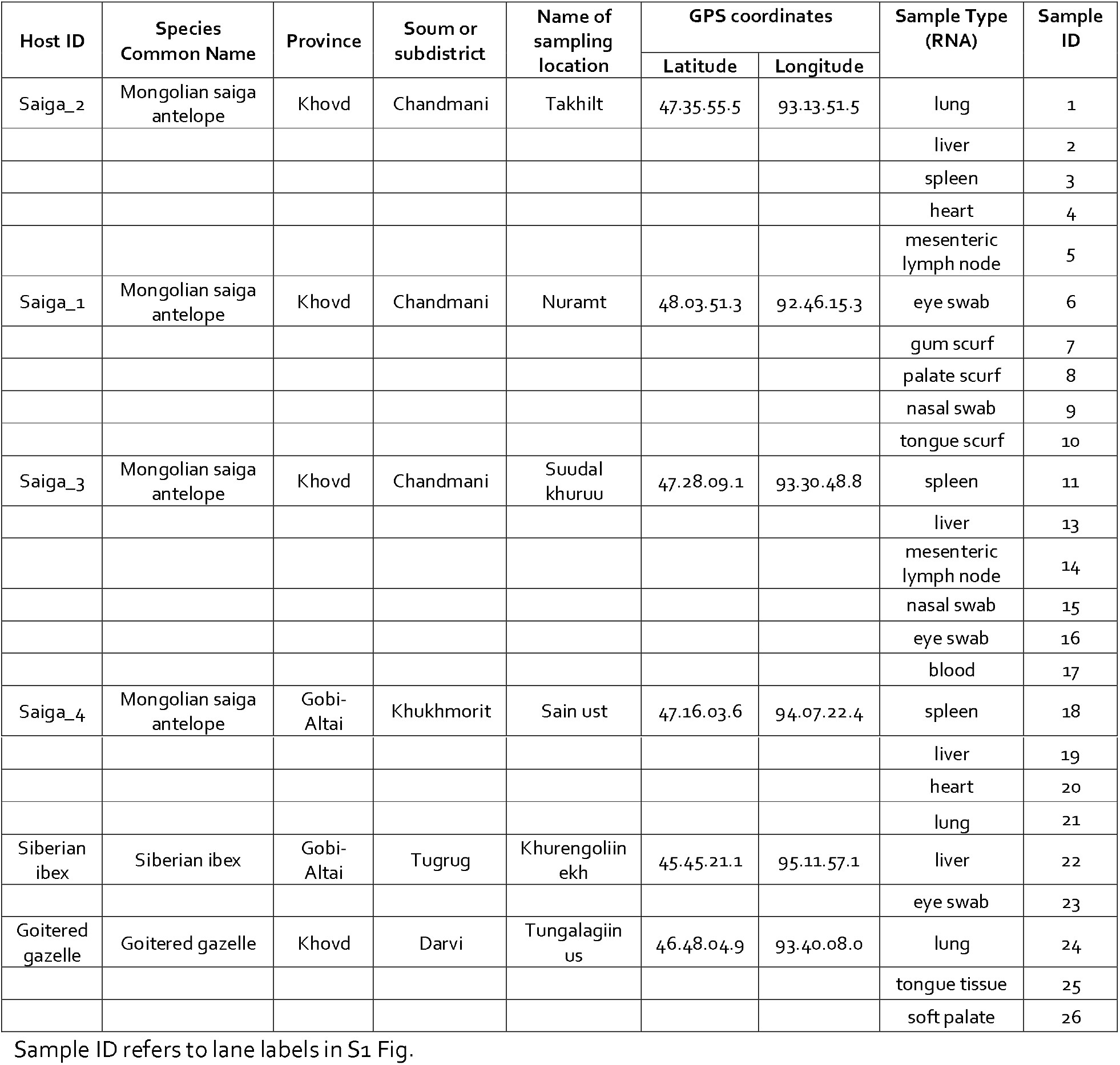
Sampling locations and sample types for PPRV-infected wildlife.

### PPRV genome sequences from Mongolian wildlife hosts

Using the Illumina NextSeq sequencing platform, five new PPRV genomes were obtained for three wildlife species: three individuals of the Mongolian saiga antelope, one goitered gazelle, and one Siberian ibex (S1 Table). The genomes were 15,954 nucleotides in length, and contained a 6-nucleotide insertion within the 5’ UTR of F gene (at position 5216 in the alignment), shared by PPRV from Mongolian livestock (KY888168.1) and Chinese lineage IV strains after 2013 [41], but not observed in other published PPRV genomes. Two of the PPRV genomes from saiga had complete nucleotide coverage across the entire genome (saiga3 and saiga4) whereas another sequence from saiga contained sequence gaps totalling 1133 nucleotides, the goitered gazelle sequence had a 33 nucleotide gap (in the M-F intergenic region) and the ibex sequence a 759 nucleotide gap (in the M-F intergenic region, plus 39nt of the F gene which was later confirmed by F gene RT-PCR). The two complete PPRV sequences from different saiga individuals differed from each other at only three nucleotide sites. Aligning the PPRV sequences from saiga with the only full PPRV genome available for Mongolian livestock (KY888168.1), showed 99.7% nucleotide identity, i.e. out of 15,954 nt sites in the genome, there were 42 (saiga 4) or 45 (saiga 3) nucleotide differences.

### Evolutionary rates and lineage divergence of PPRV

Following sequence curation as described above, 76 PPRV genomes from Genbank were added to the five novel PPRV genomes generated in this study, yielding a total of 81 sequences for phylogenetic analysis. These spanned 49 years from 1969 to December 2018, and included isolates from 24 countries. Previous phylogenomic analyses have included sequences found to be unreliable by our recombination analysis [38, 39, 41, 42], which we excluded (see methods section and S2 Table). We therefore first analysed the evolutionary rates and global lineage diversification of PPRV using our Bayesian time-scaled phylogeny of 81 genomes.

The mean evolutionary rate across the phylogeny, under an uncorrelated relaxed clock model found to be the best fit for the data, was 9.22E-4 nucleotide substitutions/site/year (95% highest posterior density (HPD) interval: 6.78E-4 - 1.17E-3) (Fig 1A). Table 2 gives the countries of origins and divergence times of PPRV lineages inferred from the Bayesian phylogenetic analysis. For lineage IV, which is expanding its geographic and host range, the median TMRCA was estimated to be 1975 (95% HPD: 1961-1985) and its country of origin was inferred as Nigeria, with moderate support (67% root state posterior probability).

**Fig 1.**
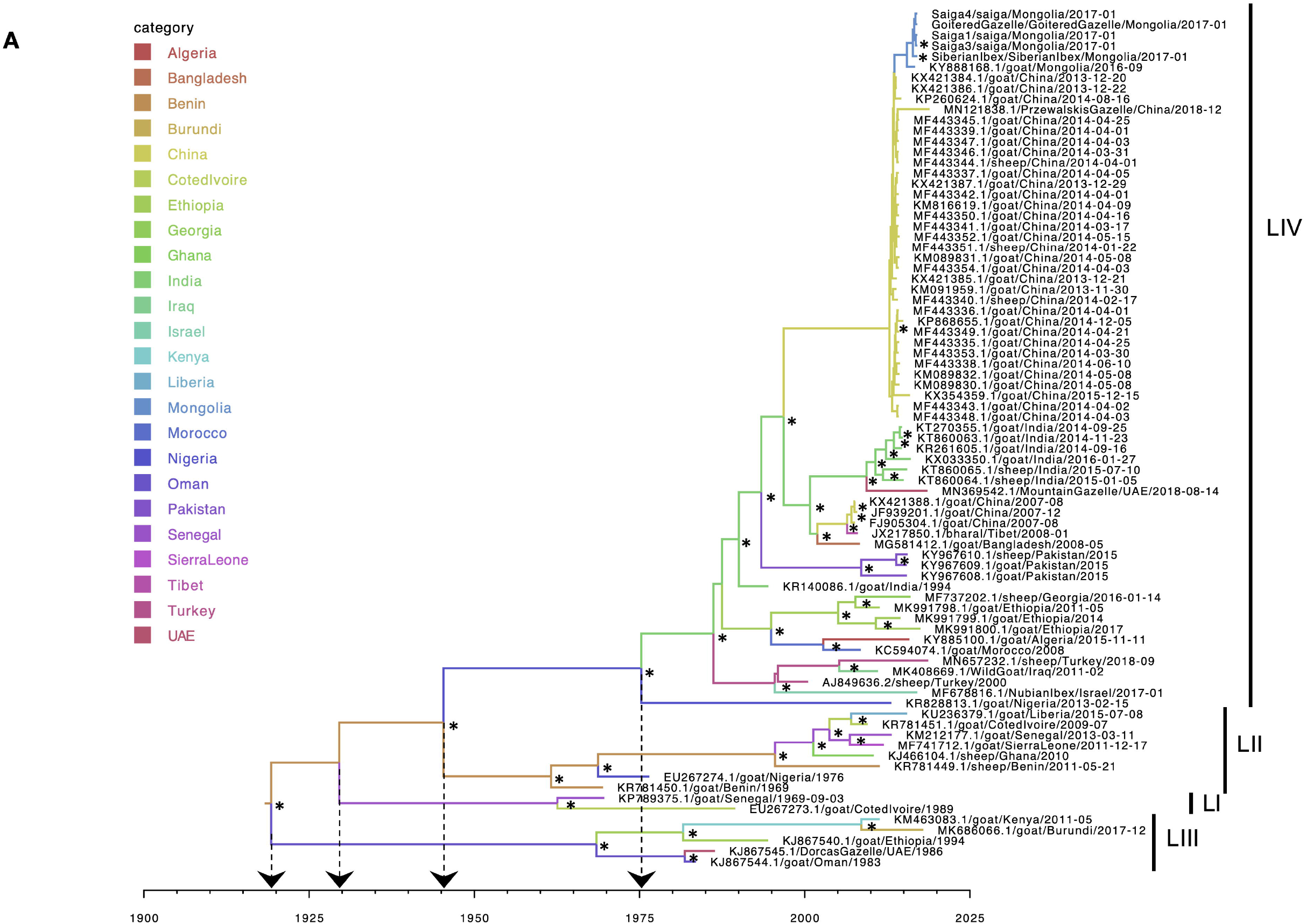
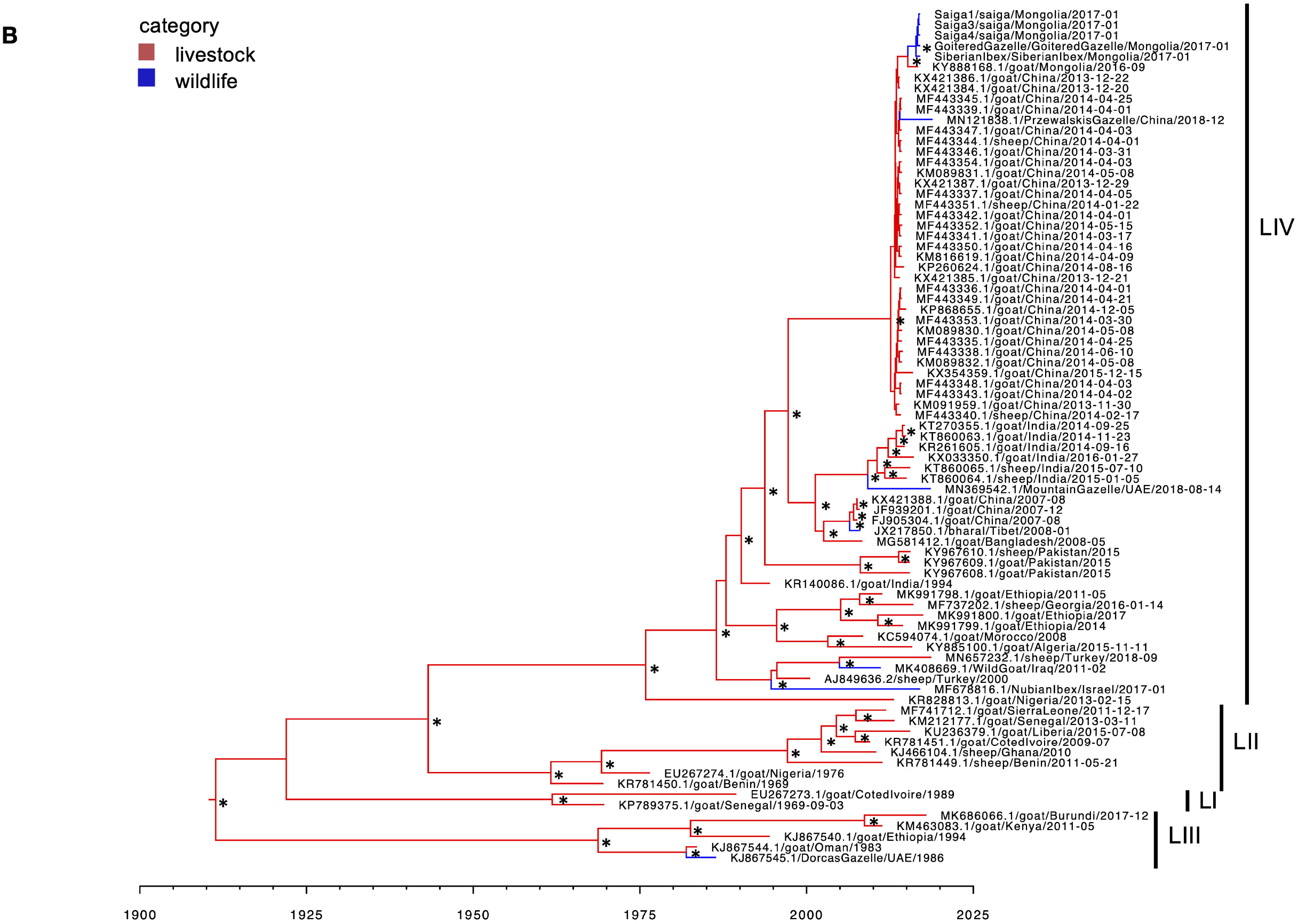
Bayesian time-scaled Maximum Clade Credibility Trees using country (A) or host category (B) partitions. Bayesian phylogenetic analysis (n=81 genomes) was run using BEAST v1.10.4 and trees are the combined output of three (A) or two (B) independent MCMC chains, visualized in FigTree. x-axis shows date. Branches are colour-coded by country (A) or host (B) as inferred using discrete trait analyses. Lineages, referred to as LI, LII, LIII or LIV, are shown. Arrows to the x-axis in (A) show ancestral nodes and corresponding TMRCAs for different lineages. * indicates posterior probability > 0.9 at the node opposite.

**Table 2.**
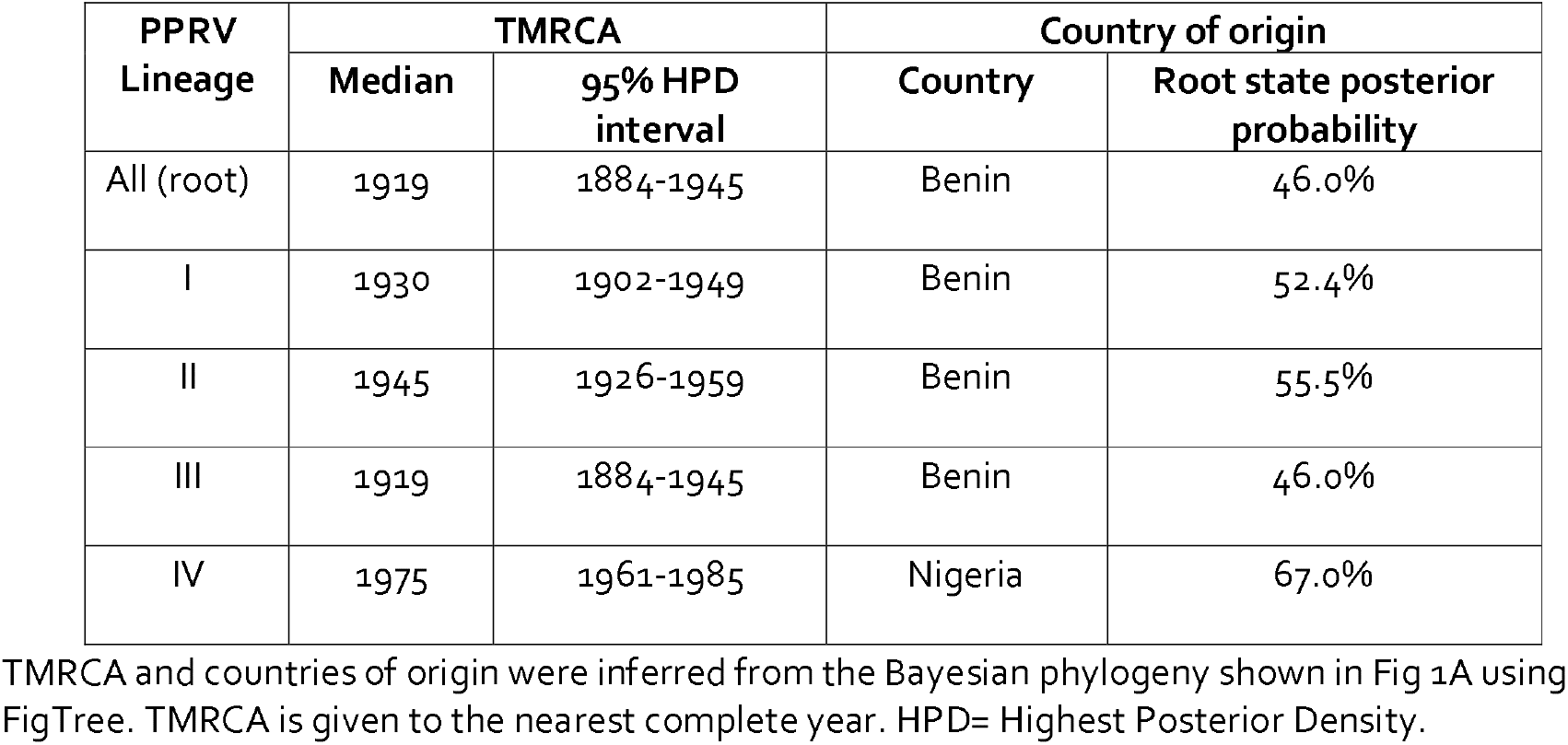
Time to the most recent common ancestor (TMRCA) and country of origin of PPRV lineages.

### Phylogenetic analysis of PPRV at the livestock-wildlife interface

Including the five novel genomes from this study, twelve wildlife-origin PPRV genomes are currently available (Table 3), although one of these (KT633939.1/ibex/China/2015-01-20) was these was excluded from our phylogenetic analysis. Two cases occurred in zoological collections, and the other ten PPRV genomes were from infections of free-ranging wildlife. Eleven of the wildlife PPRV genomes belong to lineage IV and one to lineage III. The only countries having both wildlife and livestock sequences were Mongolia and China. In contrast, three Middle Eastern countries, Israel, Iraq and UAE, had full PPRV genomes available in GenBank from wildlife hosts, but no genomes from infected livestock were available for these countries. Table 3 summarises the key epidemiological data for the 2016-2017 outbreak in Mongolian wildlife [17, 43] and the disease events associated with the other available wildlife-origin PPRV genomes.

**Table 3.**
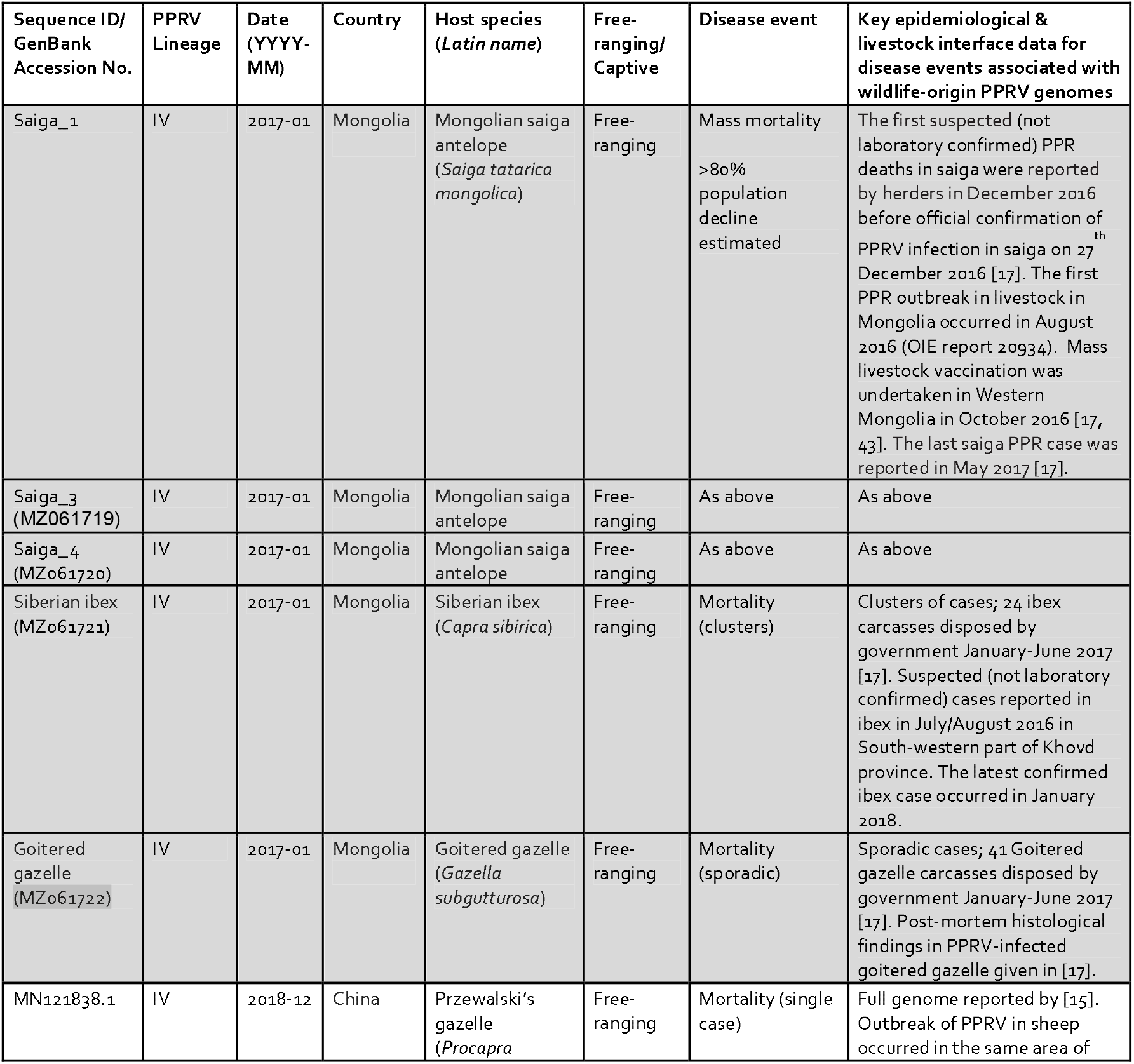
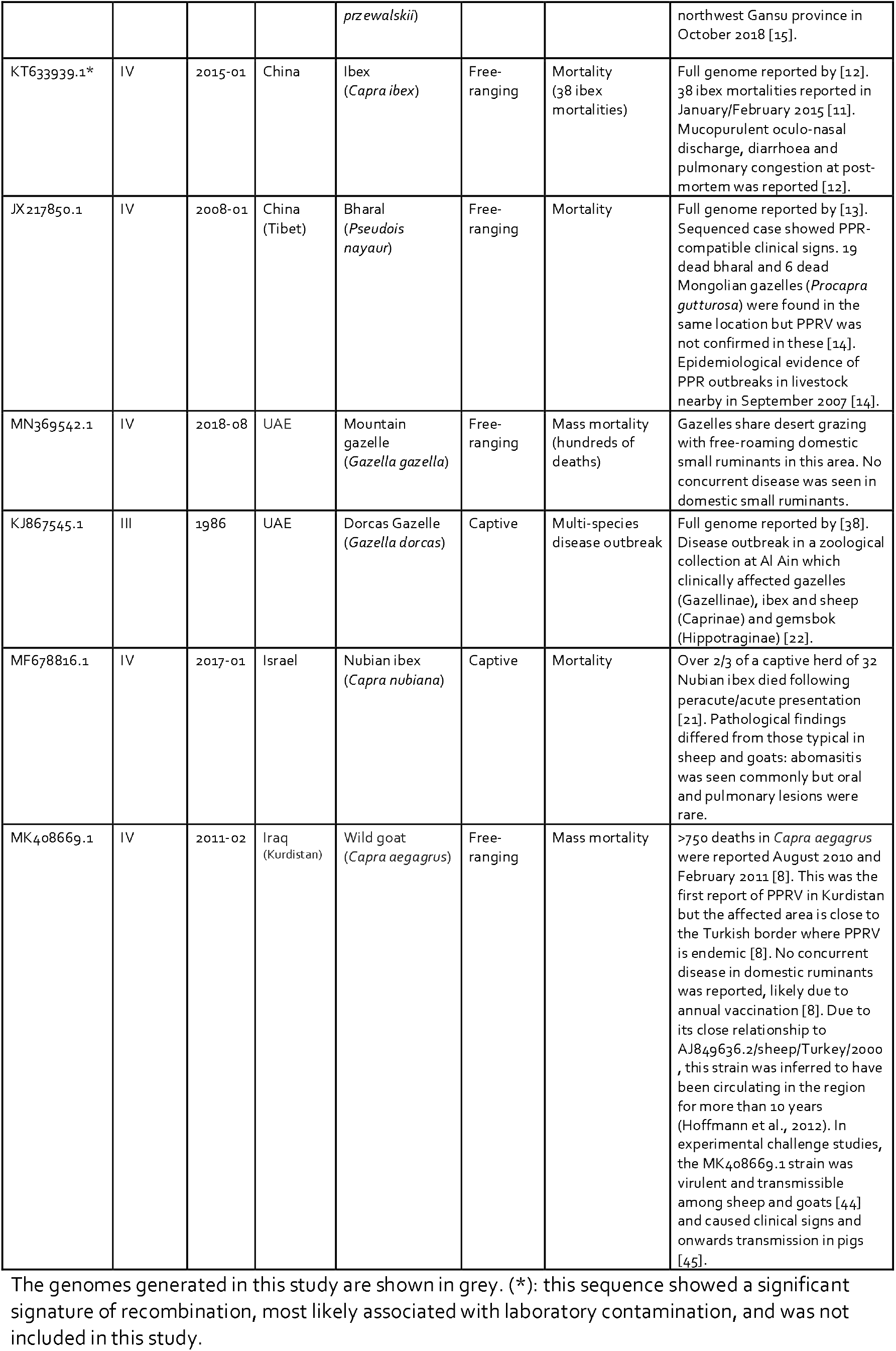
Twelve PPRV full genomes available from wildlife host species and associated metadata.

Assessing the host-traited MCC tree shows that the wildlife-origin PPRV from China after 2013, i.e. MN121838.1 (Przewalski’s gazelle), lies within the clade of livestock PPRV (Fig 1B). In contrast, several other wildlife PPRV genomes lie on branches that are basal to clades circulating in nearby locations. For example, PPRV from a bharal (Tibet, 2008) was basal to Chinese livestock sequences from 2007, PPRV from a mountain gazelle (UAE, 2018) was basal to a clade of six livestock sequences from India in 2014-2016, and PPRV from a Nubian ibex (Israel, 2017) was on a long branch basal to isolates from Turkey and Iraqi Kurdistan close to the Turkish border (Fig 1B). Similar phylogenetic relationships for wildlife-origin PPRV genomes were seen using ML phylogenetic reconstruction (Fig 2).

**Fig 2.**
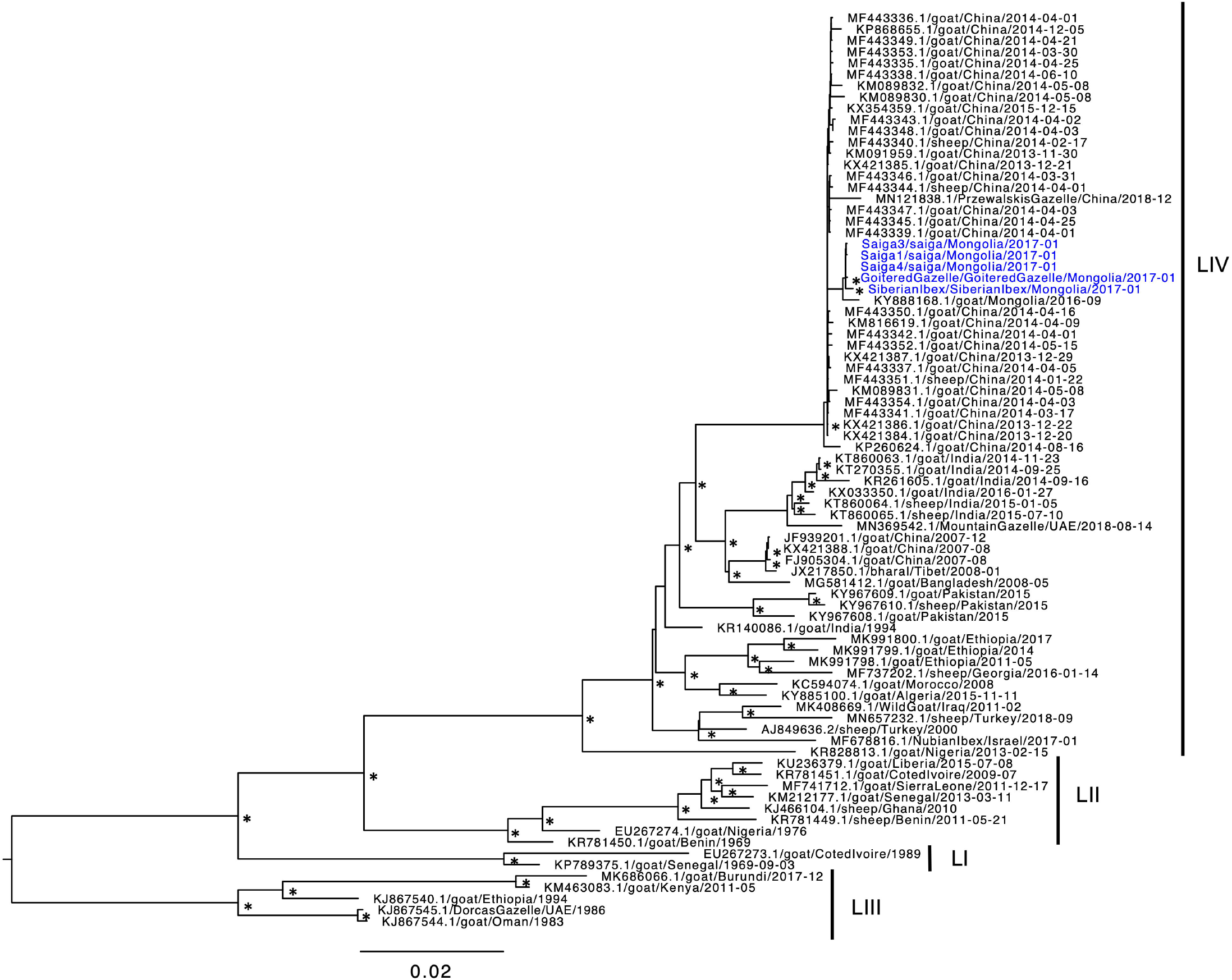
Maximum likelihood phylogeny of PPRV genomes. 81 PPRV genomes were analysed using PhyML with a GTR nucleotide substitution model and 100 bootstrap replicates. The novel genomes from this study are shown in blue. Lineages, referred to as LI, LII, LIII or LIV, are shown. Scale bar shows nucleotide substitutions per site. * indicates bootstrap proportion > 0.9 at the node opposite.

### Phylogenetic analysis and TMRCA for PPRV emergence in Mongolian wildlife

The five novel genomes from wild Mongolian ungulates were most closely related to PPRV from a Mongolian goat (KY888168.1) sampled in September 2016, the only PPRV genome sequence available from Mongolian livestock. There was strong support for monophyletic grouping of the Mongolian wildlife and livestock sequences using both ML and Bayesian inference methods (Figs 1 and 2). There was also strong support for the grouping of all five Mongolian wildlife sequences and, although there was poor support for the clade structure within the Mongolian wildlife clade, in every analysis the Siberian ibex formed a sister branch to the four other wildlife sequences from saiga and the goitered gazelle (Figs 1, 2, S3). The Mongolian sequences lie within a strongly supported clade of lineage IV sequences from China, that includes livestock sequences from 2013-2015, and a Przewalski’s gazelle sequence from 2018.

The dates of PPRV emergence in Mongolia and its wildlife were inferred using TMRCA analysis of the Bayesian time-scaled MCC tree shown in Fig 1A. The median TMRCA of the six Mongolian PPRV genomes was June 2015 (95% HPD: August 2014 – March 2016). The median date for the MRCA for the five Mongolian wildlife PPRV sequences was May 2016 (95% HPD: October 2015 – October 2016). The ancestor at the node linking all the Mongolian PPRV sequences with the most closely related Chinese sequences was dated to July 2013 (95% HPD: March 2013 – November 2013). To check that the data partitioning for the traited analysis did not substantially alter the TMCRA analysis, an untraited phylogeny was also analysed, and gave very similar results (S2 Fig).

### Phylogeographic analysis of PPRV emergence in Mongolian wildlife

SpreaD3 was used to visualise geographic spread inferred through discrete phylogeographic analysis and identify well supported rates using Bayes Factor tests. This found that PPRV spread from China into Mongolia was very strongly supported with a Bayes Factor of 494 and associated posterior probability of 0.96 (S3 Table).

To further explore and visualise phylogeographic patterns in the data, samples were geocoded and analysed using Microreact (www.microreact.org). Samples were geocoded at the highest resolution possible. GPS sampling locations were available for all Mongolian wildlife samples (Table 1), the Mongolian livestock sample and the 2008 bharal sequence from Tibet. Of the other 33 PPRV genomes within the Chinese 2013-2018 clade, 28 samples had province-level location data, hence region centroids were used for geocoding, while country-level location only was available for five sequences, in which case the China centroid was used. An open-access interactive dynamic visualisation of our global PPRV dataset, integrating phylogenetic, spatial, temporal and host (wildlife/livestock host) data is available at the permanent link https://microreact.org/project/5WNeX14MRFvwe8YLhn5a1S/e2d5dafd (and will be updated as further genomes become publicly-available).

The map highlights the proximity of the sampling locations for the wildlife and livestock PPRV genomes in the Western Mongolian provinces of Khovd and Gobi Altai, with <40km between the livestock sample and one of the saiga antelope sampled in Khovd near the Khar-Us lake four months later (Fig 3). The Siberian ibex sample, from Tugrug soum of Gobi-Altai (Table 1), was furthest from the sampling location of the Mongolian livestock (~355 km) and marginally closer to the Chinese border than other detected cases.

**Fig 3.**
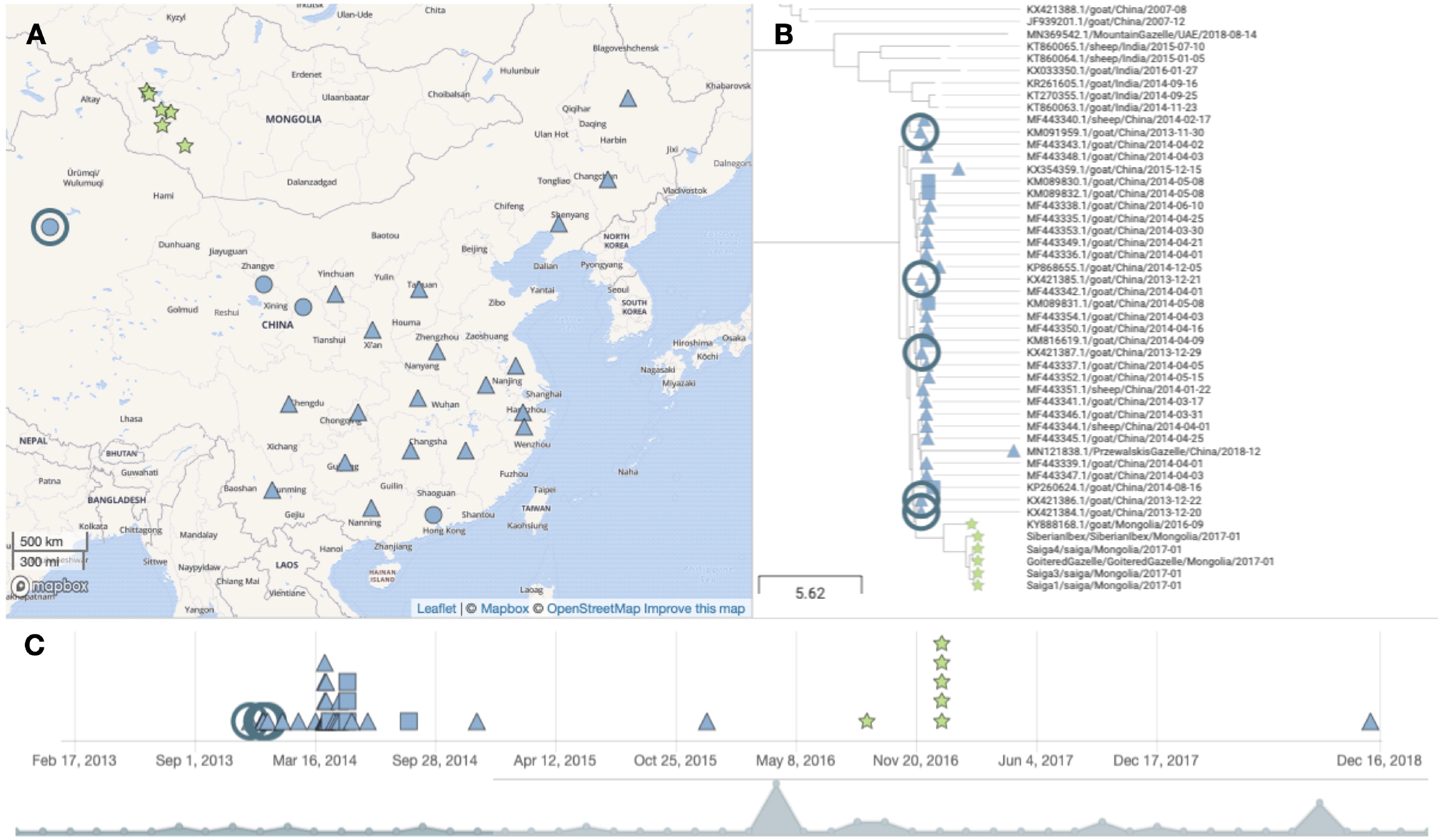
Phylogeographic visualization. An MCC phylogenetic tree (81 sequences) was uploaded to Microreact together with geocoded locations of PPRV genomes and metadata to produce the dynamic visualization of phylogenetic, spatial and temporal relationships of the global PPRV dataset. The figure shows one view available at the Microreact project link (https://microreact.org/project/5WNeX14MRFvwe8YLhn5a1S/e2d5dafd). showing the location (A), phylogenetic relationship (B) and timeline (C) for the clade of PPRV genomes from Mongolia and China since 2013. Symbol colour denotes country of origin (Mongolia in green and China in blue). The symbol shape denotes different resolutions of geocoding for genomes: stars denotes samples with GPS coordinates; triangles denote region centroids; squares denote country centroids and a circle on the map view indicates multiple genomes from the same location. The ringed samples in panels (A)-(C) show samples from Xinjiang province, including KX421386.1 and KX421384.1. The map shown in (A) uses base map and data from OpenStreetMap and OpenStreetMap Foundation, available under the <u>Open Database License</u>,with tiles from the <u>Mapbox mapping platform</u>, used via the freely-available Microreact application.

The PPRV genomes KX421386.1 and KX421384.1, from December 2013, are closely phylogenetically related to the Mongolian sequences, and also the geographically closest sequences, from Xinjiang province of Northwest China, which borders Mongolia (Fig 3).

### Amino acid polymorphisms associated with host range expansion

The five new PPRV sequences were compared with the 76 other genomes in our dataset in order to identify polymorphisms of interest that might be associated with PPRV emergence in Mongolian wildlife (Table 4).

**Table 4.**
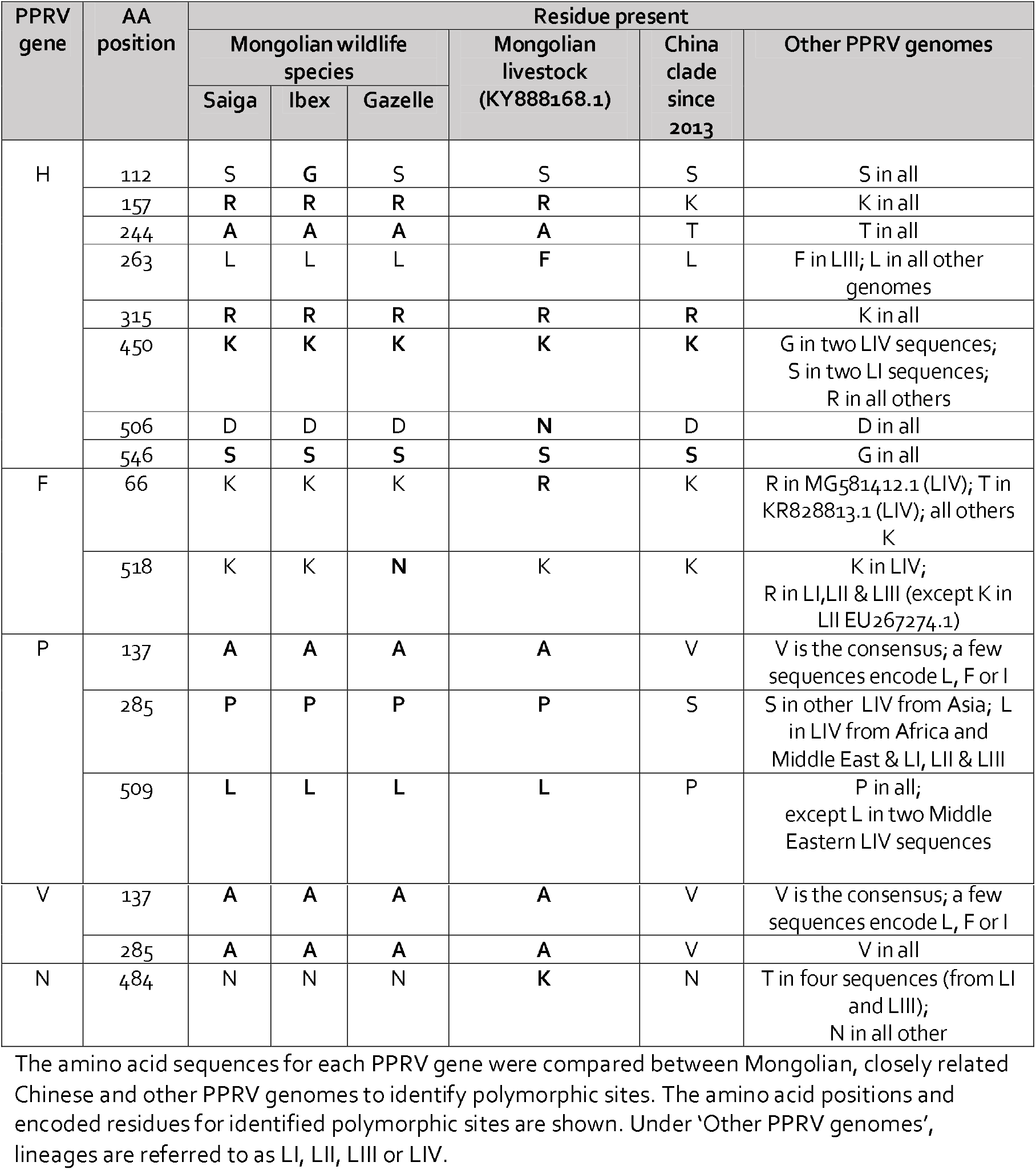
Amino acid polymorphisms.

Among the polymorphisms observed, the Mongolian livestock PPRV (KY888168.1) encodes asparagine (N) at position 506 of PPRV H, whereas every other sequence encodes aspartate (D) as part of the completely conserved ^505^DDD^507^ motif. The PPRV H from Siberian ibex has a glycine (G) at amino acid 112, whereas the other 80 sequences in the dataset encode serine (S). In addition, two residues in H are unique to the Mongolian PPRV sequences, including the five wildlife and one livestock sequence: arginine (R) instead of lysine (K) at amino acid 157, and hydrophobic alanine (A) instead of hydrophilic threonine (T) at position 244. We also noted several sites within the H gene where the monophyletic clade of 39 Chinese/Mongolian sequences encode common signature residues compared other PPRV sequences, namely amino acids 315 (R), 450 (K) and 546 (S) (Table 4). For the F protein, PPRV from goitered gazelle was the only genome to encode asparagine (N) at position 518, a significantly different residue from lysine (K) (in all other lineage IV genomes) or R (lineages I, II and III).

Mongolian PPRV sequences encode alanine (A) at position 137 of their P and V proteins, a residue not seen in any other PPRV isolate in the database (otherwise valine (V) is the consensus, with a few sequences encoding L, F and I). Another polymorphism was seen at amino acid 285, where proline (P) is seen only in the P proteins of Mongolian sequences instead of either Serine (S) in other lineage IV PPRV from East and South Asia or leucine (L) in lineage IV viruses from Africa and the Middle East as well as lineages I, II and III. This mutation lies after the RNA editing site, leading to a different substitution in the V protein, with Mongolian sequences encoding alanine (A) and all other sequences encoding valine (V) at amino acid 285, which lies close to the zinc-binding domain comprising amino acids 237-280.

### Molecular modelling of polymorphic amino acids in Mongolian PPRV H

To determine the location of the polymorphic sites identified in PPRV H from Mongolian wildlife and livestock, structural homology modelling was performed, based the solved crystal structure of the head domain of MeV H in complex with SLAM, the host cell entry receptor for morbilliviruses [28]. Of the polymorphic sites identified between H sequences, sites 112 and 157 were not captured by the crystal structure. Amino acid residues 244 and 263 were predicted to be distant from the SLAM binding interface and surface exposed (Fig 4A,B). Residues 506 and 546 were predicted to lie within the region that forms the SLAM binding interface (Fig 4A,B). While amino acid site 546 of H is not thought to form a direct contact with SLAM, amino acid 506 lies between two residues (D505 and D507) which in MeV H form salt bridges to K77 and R90 of marmoset SLAM, comprising site 1 of the RBD (Fig 4C) [28]. Modelling caprine SLAM in place of marmoset SLAM reveals K78 in place of K77, whereas R90 in marmoset SLAM is replaced by R91 in caprine SLAM, with the preceding P90 facing away from PPRV H (Fig 4D). Replacing G506 of MeV H with D506 (i.e. the consensus residue in PPRV H) or N506 (i.e. the substitution seen in PPRV from Mongolian livestock) could affect the interaction with SLAM due to its proximity (Fig 4D,E,F).

**Fig 4.**
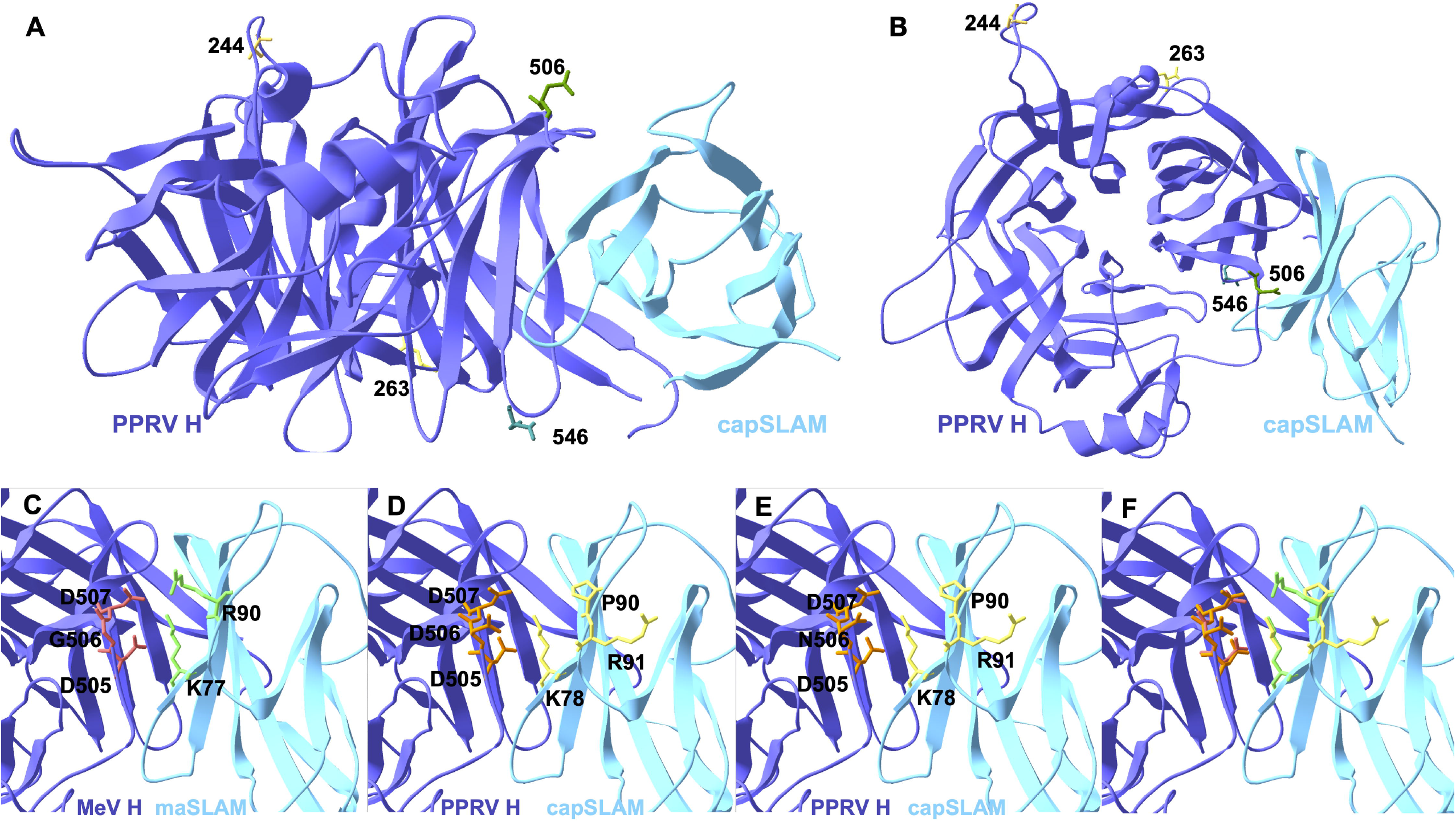
Homology modelling of the PPRV H-SLAM complex. Modelling was performed using SWISS_MODEL using the published crystal structure of MeV H bound to marmoset SLAM to show positions of Mongolian-specific amino acid polymorphisms in PPRV H. Side (A) and top (B) orthogonal views of PPRV H bound to caprine SLAM (capSLAM) showing residues for PPRV H from saiga antelope (244A, 263L, 506D, 546S). Residues at site 1 of the RBD for (C) MeV H and marmoset SLAM (maSLAM), (D) PPRV H 506D and capSLAM, (E) PPRV H 506N and capSLAM, or (F) overlayed image of panels C and D. H proteins are shown in purple and SLAM proteins in turquoise.

### Selection pressure analysis

To test for positive selection, methods were used that assess the numbers of non-synonymous (dN) to synonymous (dS) nucleotide substitutions per site, with dN > dS indicative of positive selection (i.e. adaptive evolution). To assess positive selection acting on individual codons, the CDS of each PPRV gene was analysed using MEME, FUBAR, FEL and CodeML. Sites under positive selection were identified in all genes with all four methods, except for the F gene (three methods), the C gene (two methods) and the M gene (one method) (Table 5, S4 Table). The amino acid positions identified as evolving under positive selection by all four methods were amino acid 246 of the H protein, amino acid 616 of the L protein and amino acids 52 and 101 of the P and V proteins. Of note, all methods except CodeML also identified positive selection acting on amino acid 137 of the P and V proteins.

**Table 5.**
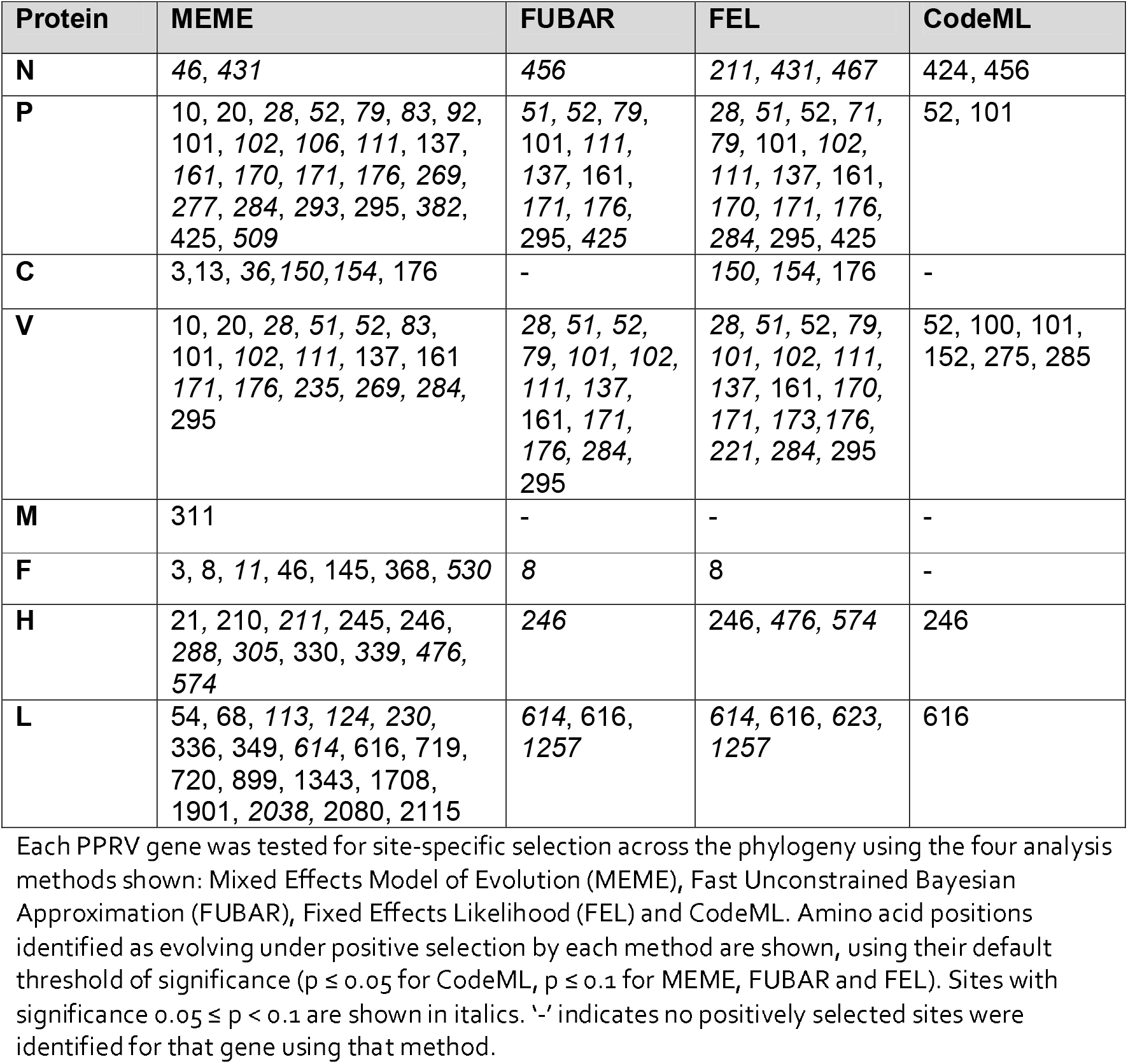
Amino acid sites under positive selection

Methods for detecting lineage specific selection were also used to test whether the clade of Mongolian and Chinese PPRV sequences since 2013 showed evidence of positive selection. Both BUSTED and aBSREL found no evidence of selection for N, P, C, V, M, F or H genes but did find evidence of positive selection acting on the L gene of the Mongolian/Chinese PPRV clade (p<0.05) (Table 6). The BUSTED result indicates L gene-wide episodic diversifying selection, i.e. evidence that at least one site on at least one test branch within the clade has experienced diversifying selection. aBSREL identified episodic diversifying selection acting on two branches, namely the single Mongolian livestock PPRV sequence (KY888168.1) and the PPRV L gene from Chinese goat (KP260624) (Table S5). FEL identified positively selected sites within each gene in the Chinese/Mongolian clade (Table 6). Some sites were identified both by lineage specific FEL and earlier by MEME in all but the C proteins, including position 137 in the P and V proteins (Table 5 and 6).

**Table 6.**
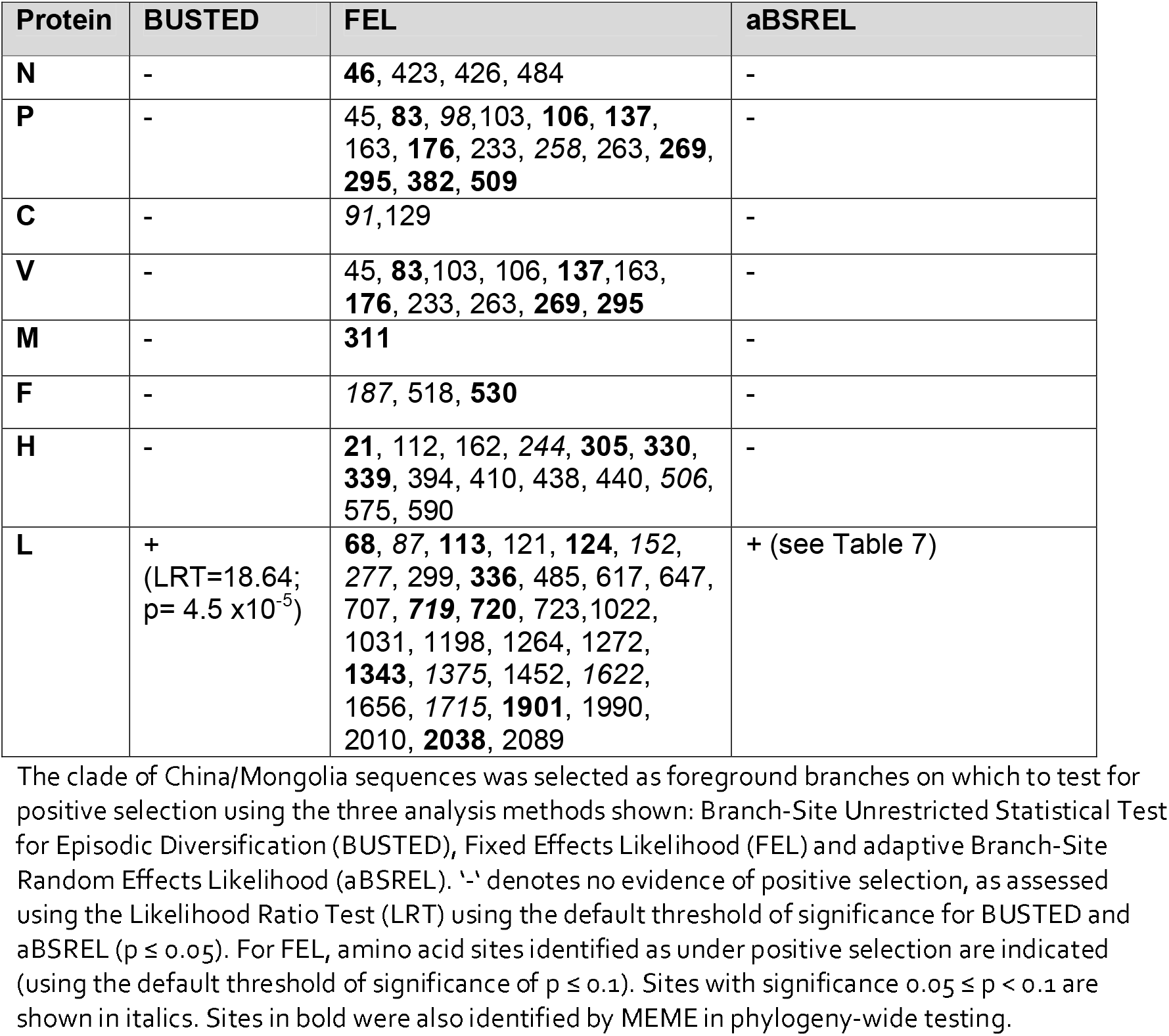
Lineage specific selection tests.

Eight sites at which Mongolian PPRV-specific amino acid polymorphisms had been noted in either wildlife or livestock (Table 4) were also identified as positively selected by the lineage-specific FEL analysis (Table 6), suggesting functional significance of these sites, namely H_112, H_244, H_506, F_518, P_137, P_509, V_137 and N_484.

## Discussion

Despite the broad host tropism and impact of PPRV across wild ungulate species, there is a paucity of wildlife-origin PPRV genomes which, along with the lack of field epidemiological data, has hindered understanding of viral evolution and dynamics at the wildlife-livestock interface. This understanding is critical in order to design effective disease control and eradication strategies and thereby support the success of the PPR Global Eradication Programme (GEP) and mitigate the threat of PPRV to both domestic and wild ungulates. In this study, full PPRV genomes were generated from three Mongolian saiga antelope, one goitered gazelle and one Siberian ibex that were part of a major mortality event in Mongolian wildlife in 2016-2017. These were analysed together with a curated set of all other genomes available in GenBank, to examine PPRV evolution and cross-species transmission. This showed strong support for monophyletic grouping of genomes from Mongolian wildlife and livestock, and for incursion of PPRV into Mongolian from China. Our TMRCA analysis also indicated that PPRV emerged in Mongolia’s endangered wildlife populations before livestock vaccination was initiated, and sequence polymorphisms and signatures of adaptive evolution were identified.

Prior to our phylogenetic analysis, recombination detection algorithms were applied to our dataset and identified several potential recombinant PPRV genomes. Recombination has never been reported for PPRV and is rare among negative sense RNA viruses [46, 47]. However, bioinformatics errors or laboratory contamination can lead to false signals of recombination in PPRV [39, 48]. Previous reports have also highlighted the issue of contamination in published PPRV sequences [33, 39, 48, 49], so the most probable explanation is that the recombinant PPRV genomes we identified are the result of laboratory contamination during the process of genome sequencing. However, PPRV recombination cannot be totally ruled out and should be further explored in dedicated infection experiments. The inclusion of dubious and/or recombinant genomes in earlier phylogenomic studies may have influenced their results [38, 39, 41, 42], and we therefore first analysed PPRV molecular evolution globally.

The genome-wide mean evolutionary rate inferred from the Bayesian MCC phylogeny of 81 PPRV genomes was 9.22E-4 nucleotide substitutions/site/year (95% highest posterior density (HPD) interval: 6.78E-4 - 1.17E-3), which is comparable to previous estimates [38, 39]. The TMRCA and country of origin of each of the PPRV lineages were inferred from our Bayesian MCC phylogeny (Table 2). In the case of lineage IV, which is becoming the predominant lineage globally, the median TMRCA was estimated as 1975 (95% HPD: 1961-1985). This median date is slightly earlier than that reported by Muniraju et al. using a more limited dataset of twelve genomes (1987), although the HPD intervals overlap, or from partial N or F gene datasets [38, 50], and broadly equivalent to estimates obtained from analysis of 53 full genomes (1968) [42].

All phylogeographic analyses are affected by sampling bias. Our analysis indicates with moderate support that the country of origin of lineage IV was Nigeria, with the second most supported origin in Benin. Previous analyses identified India as the most likely origin of lineage IV, but recent sequences from Nigeria were not included in that analysis [38]. The extremely poor sampling prior to the 1990s means that whilst an origin of lineage IV in Africa seems plausible, we do not believe that there is sufficient resolution of genomic sequences to robustly establish the country of origin. However, the detection of lineage IV PPRV in livestock in Cameroon in 1997, in the Central African Republic in 2004 and in several other Northern and Eastern African countries in the early 2000s, in the absence of known animal movement from Asia [33, 34, 51, 52], also supports the hypothesis that lineage IV may have emerged in Africa before spreading to the Middle East and Asia and then re-emerging in African countries. Additional full genomes for lineage IV viruses from Africa, both historical and contemporary, would improve the phylogenetic power for more robust testing of this hypothesis.

Phylogenetic relationships between wildlife and livestock origin PPRV were assessed across the global phylogeny. With the exception of the Mongolian wildlife clade, all wildlife sequences occur as isolated sequences within the broader livestock diversity. This is consistent with repeated dead-end spillovers of PPRV from livestock into wildlife, but the lack of sequences from wildlife, long branches and geographic distances between the most related livestock and wildlife sequences makes it impossible to rule out transmission from largely undetected outbreaks in wildlife into livestock. No additional PPRV genomes were available from UAE, Iraq or Israel, although 18 full genomes from livestock in Israel between 1997-2004 were sequenced previously, but unfortunately not made publicly available [42]. More dense sampling of PPRV genomes globally would likely correct sampling bias and help reveal that viruses from wild and domestic hosts cluster together according to geographic location, as seen for China and Mongolia (Figs 1 and 2), and indicated by partial N gene analysis [53].

The novel PPRV genomes from Mongolian wildlife were most closely phylogenetically related to the PPRV genome from Mongolian livestock, strongly supported in both Bayesian (Fig 1) and ML (Fig 2) phylogenies. Although this might be expected, it was not evident in earlier analysis, which showed closer phylogenetic proximity of saiga-origin PPRV to Chinese livestock as opposed to Mongolian livestock sequences, owing to insufficient phylogenetic resolution of partial N gene sequence data used (255 nucleotides) [17]. China is the only other country for which both wildlife and livestock genomes are available, with phylogenetic analysis showing PPRV from Przewalski’s gazelle embedded, albeit with poor node resolution, within a well-supported clade of 32 livestock viruses sampled since 2013.

Our study is the first to sequence multiple PPRV genomes from an outbreak in wildlife. Taking advantage of this, the full PPRV genome available for Mongolian livestock [40] and the dense sampling of PPRV genomes in China since 2013 [41] we examined PPRV emergence in Mongolian wildlife. The six Mongolian PPRV genomes formed a monophyletic group within a large clade (n=39) that includes lineage IV PPRV from China sampled since 2013 (Figs 1 and 2). In China, the first PPR outbreak was in Tibet in 2007-2008 and there were subsequently no reports of PPR until its re-emergence in Xinjiang province in November 2013 [54]. PPRV from China in 2007/2008 (n=4 genomes) was more closely phylogenetically related to viruses from India (spanning 2014-2016) than to the PPRV in China since 2013, consistent with earlier reports [41, 54]. Statistical testing of the Bayesian analysis of full genomes provided very strong phylogenetic support that PPRV spread into Mongolia from China (Bayes Factor: 494; posterior probability: 0.96), although the lack of surveillance or genomic sequencing from other neighbouring countries (e.g., Kazakhstan, Russia) means that we cannot exclude introduction via an intermediary location. Phylogeographic visualisation using Microreact showed that PPRV genomes closely related to Mongolian PPRV were from livestock in Xinjiang province. Incursion of PPRV from Xinjiang province would be consistent with epidemiological reports that initial livestock cases and the earliest suspected wildlife cases clustered at the southwestern Mongolia-China border [17, 43]. Between 2013 and 2016, PPRV cases in Argali sheep, *Capra ibex* and goitered gazelle have been confirmed across six different counties within Xinjiang province, where shared grazing provides opportunity for cross-species transmission [10, 12]. Owing to the presence of multiple PPRV-susceptible wild ungulate species in Xinjiang [55] and its extensive national borders (with Mongolia, Kazakhstan, Kyrgyzstan, India, Pakistan, Russia, Afghanistan and Tajikistan and Afghanistan), Xinjiang should be a focus for increased surveillance and sampling of PPRV across the wild and domestic ungulate community.

The TMRCA of the clade of Mongolian PPRV genomes (one livestock and five wildlife) in our analysis was June 2015 (95% HPD: August 2014 – March 2016). The TMRCA linking the Mongolian to the Chinese genomes was inferred as July 2013 (95% HPD: March 2013 – November 2013). The introduction of PPRV into Mongolia likely occurred between the TMRCA estimates for these two nodes. Therefore, the very latest date of emergence in the country that is consistent with the phylogenetic analysis is March 2016 (with 95% probability). This suggests that PPRV was circulating undetected for minimum of six months before the first reported PPR outbreak in Mongolia, which was reported in livestock in Khovd province in August 2016 (notified to OIE in September 2016, Notification report REF OIE 20934). The TMRCA analysis indicates a median date for PPRV emergence in Mongolian wildlife of May 2016 (95% HPD: October 2015 – October 2016). This suggests that PPRV infections in wildlife may have pre-dated the first confirmed wildlife case in late December 2016, as proposed earlier based on interview data and undiagnosed wildlife mortalities [17, 43]. However, more genomic sequencing of PPRV detections in livestock during 2015 to 2016 in this region is required to confirm this, as it is possible that ‘missing’ livestock sequences would intersperse with the (currently monophyletic) wildlife clade, thereby changing this interpretation. Pruvot et al. reported that earliest suspected unconfirmed cases in Mongolian wildlife occurred in Siberian ibex in July/August 2016 [17]. All wildlife PPRV genomes in our analysis date from January 2017 and the clade branching structure is poorly supported in our analyses, thus no inference can be made regarding the relative timing of emergence among different wildlife species. Interestingly, however, phylogenetic reconstruction consistently showed PPRV from Siberian ibex located on a sister branch to the saiga and goitered gazelle viruses (Figs 1, 2, S2). If the phylogenetic separation observed correctly captures true patterns of ancestry, phylogenetic separation could be related to (i) some structuring of the virus between wild species, potentially related to a longer period of transmission and evolution in ibex consistent with earlier emergence in this species, and/or (ii) the phylogenetic structure seen could be related to geographical separation of the ibex sample, which was further southeast than the other sampling sites.

Based on epidemiological and ecological evidence, multiple spillover events from livestock to different wildlife populations during the Mongolian outbreak have been proposed [17]. The lack of additional livestock-origin genomes means that this cannot currently be confirmed using phylogenetic approaches, although our data are compatible with the hypothesis of spillover from domestic into wild ungulates in Mongolia. Our TMRCA analyses suggest that transmission of PPRV at the livestock-wildlife interface occurred prior to, or at the latest contemporaneous with, the livestock vaccination campaign which began in October 2016. This could explain emergence in wildlife despite vaccination of ~10.4 million small ruminant livestock in Mongolia’s Western provinces, without inferring inadequacies in vaccination coverage or seroconversion [43].

At the time of the disease outbreak, it is clear that the PPRV infecting wildlife was not only virulent but already well-adapted for infection and transmission in these hosts, at least in saiga antelope. It is possible that mutations present in the wildlife-infective PPRV strains, or their ancestral viruses, contributed to this, and therefore genomic sequences were analysed for notable sequence features and signatures of adaptive evolution. The genomes from Mongolian wildlife, as well as from Mongolian livestock and all PPRV from China since 2013, have a six nucleotide insertion in the 5’UTR of the F gene, which renders the genome 15,954 nucleotides long, instead of 15,948 nucleotides in all other PPRV strains [12, 40, 41, 56]. This insertion maintains the ‘rule of six’, i.e. that genome length is a multiple of six, which is necessary for efficient replication of members of the *Paramyxoviridae* family [57]. Comparing the nucleotide sequence at the insertion site shows that this is much more cytosine rich in lineage IV viruses and that a tract of six cytosines is present directly upstream of the insertion site in PPRV from sister clades to the Mongolia/China clade (S1 file alignment). Insertions in measles virus genomes are thought to arise via polymerase slippage errors when transcribing homopolymeric regions [58], suggesting a similar mechanism may have occurred in an ancestor to the 2013 China PPRV clade, and the resulting insertion has been maintained in the Mongolian PPRV. Interestingly, six nucleotide insertions in the 5’UTR of the F gene were also reported in goat-adapted rinderpest virus vaccine strains [59]. Similarly, several variants of measles virus have been identified with a net gain of six nucleotides within the M-F UTR and possibly associated with lineage emergence [60, 61]. In light of these reports, and the ability of the PPRV-F 5’UTR to enhance F gene translation [62], experiments to address the functional consequences, and potential selective advantage, of the observed insertion in the Mongolian and Chinese PPRV should be prioritised.

The CDS of each PPRV gene was assessed for non-synonymous nucleotide changes and these were correlated with the results of selection pressure analyses using multiple methods. Positive selection in viral genomes indicates adaptive evolution in response to changing fitness or functional requirements [63, 64], including infection and replication in novel hosts and evasion of host innate and adaptive immune responses, and is most likely at interaction interfaces between viral and host cell molecules.

For the H and F envelope entry proteins, polymorphisms specific to Mongolian wildlife PPRV were present at amino acid 112 in H from Siberian ibex and at amino acid 518 in F from goitered gazelle, with both sites under lineage-specific positive selection (Table 4, Table 6). More PPRV genomes would clearly be needed to assess the possibility that these are host species-specific mutations. Of note, however, are two polymorphisms in H which are specific to the six PPRV genomes from Mongolian livestock and wildlife, namely R157 (instead of K157 in all 76 other genomes) and A244 (instead of T244 in all 76 other genomes), raising the possibility that these substitutions may have helped PPRV spillover to wildlife. Whereas K157R is not a major change in physiochemical or steric properties of the amino acid, the change from the polar, uncharged amino acid threonine (T) to hydrophobic alanine (A) at position 244 of the Mongolian PPRV H protein could have more significant effects on inter-molecular interactions. The adjacent residues 234-243 form a predicted T cell epitope [65] and therefore one possibility is that T244A influences interactions with MHC I. This region of H is also subject to adaptive evolution: site 244 is detected as under positive selection in the China/Mongolia linage using FEL methods, site 245 is detected by MEME as under episodic positive selection and site 246 is identified by all four methods in phylogeny-wide testing (Table 5), and was also reported previously as a positively selected site [66]. The single genome from Mongolian livestock has two substitutions in H, L263F and D506N, which are not shared by the five wildlife-origin PPRV and should be assessed in other livestock samples. If consistently seen, D506N is particularly noteworthy since it lies at the SLAM binding interface (Fig 4) and all other PPRV strains encode aspartate (D) at this site [24]. Neighbouring residues 505N and 507N are completely conserved among morbilliviruses [24], and comprise site 1 of the RBD (Fig 4), forming salt bridges to SLAM in the crystal structure for MeV H [28] and in homology models for PPRV H [4, 66]. Therefore this residue could be functionally important for infectivity and viral tropism. Substitutions already present in the Chinese lineage of PPRV since 2013 may also have contributed to emergence in Mongolian wildlife, including unique residues at sites 315, 450 and 546 of the H protein. At position 546, all sequences from the Mongolia/China clade encode serine, as does MeV at the equivalent position, whereas every other PPRV strain in the phylogeny encodes glycine. This residue is located adjacent to hydrophobic site 4 of the RBD (Fig 4) but is not thought to form a direct contact with SLAM [28].

The P and V proteins antagonise the interferon (IFN) response via intermolecular interactions with host STAT1, STAT2, Tyk2 and Jak1 proteins [67–69]. Potentially, Mongolian PPRV-specific residues in the P (137, 285, 509) and V proteins (137, 285), particularly those under positive selection (i.e. 137 in P and V; 509 in P) might affect evasion of host innate immunity and emergence. In addition to its role in genome encapsidation and replication, the N protein was shown recently also to inhibit type I IFN by binding IRF3 [70]. We identified residues 46 and 426 in N as positively selected, with both MEME and FEL (Tables 5 and 6), as well as a Mongolian livestock-specific residue and lineage-specific positive selection at site 484 (Tables 4 and 6). The L gene, although lacking any Mongolian-PPRV specific polymorphisms, was the only gene identified as positively selected by all three methods when the China/Mongolia clade was tested (Table 6), suggesting this catalytic polymerase protein has undergone adaptive evolution in this clade. Functional studies including pseudotyping (to assess receptor usage), minigenome (to assess replication), immune reporter and intermolecular binding assays are beyond the scope of the current study but should be planned to define the role of the mutations identified here in lineage IV PPRV expansion.

In summary, our study has provided insights into PPRV transmission at the livestock-wildlife interface. More PPRV full genomes are required in order to strengthen molecular epidemiological studies. The occurrence of cases in wildlife should serve as a trigger to initiate local surveillance/sampling in livestock and the surveillance and opportunistic sampling for wildlife disease events should be increased. However, quantifying the direction and extent of pathogen transmission in multi-host systems is challenging, even in the case of extremely detailed longitudinal study systems with pathogen genomic and host life history data [71]. There is thus a need for other types of data, including interspecific contact rates and pathways, and enhanced epidemiological and serological data, to enable approaches which integrate these with genomics [71–73] and can lead to improved understanding of interspecific PPRV transmission dynamics and the epidemiological roles of wildlife species.

## Materials and Methods

### Sample collection

Samples from five wild Mongolian ungulates suspected of being infected with PPRV were all collected in January 2017 during an emergency field mission to urgently respond, assess and advise the Mongolian Government through the National Emergency Committee. The mission involved FAO’s Crisis Management Centre-Animal Health (CMC-AH), the Wildlife Conservation Society, the Royal Veterinary College (RVC), the Veterinary and Animal Breeding Agency (VABA) and the State Central Veterinary Laboratory (SCVL) [43]. Tissues were collected from fresh carcass necropsies from three saiga antelope, one goitered gazelle and one Siberian ibex. GPS locations of sampled animals are provided in Table 1. Total RNA was extracted from tissues using a NucleoSpin RNA Virus mini kit (Macherey-Nagel, 740956) at SCVL, Ulaanbaatar. Samples were then imported to RVC under APHA and CITES import permits.

### RT-PCR

One-step RT-PCR was performed on 2ul total RNA using the SuperScript IV One-Step PCR kit (ThermoFisher, Catalog No. 12594025) either to amplify nucleotides 1232-1583 of the PPRV N gene using published primers NP3 and NP4 (Couacy-Hymann et al 2002) or to amplify the full length H gene (with primers 5’-CTCCACGCTCCACCACAC-3’ and 5’-CTCGGTGGCGACTCAAGG-3’) or the full length F gene (with primers 5’-GCTATGCGGCCGCACCATGACGCGGGTCGCAATYTT-3’ and 5’-GGTGAGGATCCCTACAGTGATCTTACGTACGAC-3’).

### Whole genome next generation sequencing

Total RNA from tissues was converted to cDNA with SuperScript IV and sequencing libraries prepared with the Nextera XT DNA Library Preparation Kit (Illumina, FC-131-1024). Sequencing was performed using the Illumina NextSeq platform with 150bp paired-end reads. Sequencing data was mapped to a reference PPRV genome using BWA [74] and then consensus calling was performed using SAMtools [75]. This was corroborated by removing host reads and then undertaking *de novo* assembly with SPAdes [76], and then both outputs were aligned to confirm PPRV genome sequences.

### PPRV full genome dataset curation and recombination analysis

In addition to the five novel PPRV genomes, all full PPRV genomes available in Genbank were downloaded (last accessed 22/11/2020). Vaccine sequences were removed from the dataset, as well as one sequence noted in GenBank as multiply passaged in cell culture (MN369543.1). Sequence alignment was performed using MAFFT [77] in the Geneious software package, followed by manual editing. TempEst was run to assess temporal signal in the data and evidenced clock-like evolution (S3 Fig) [78]. TempEst also identified an outlier sequence (KJ867543.1, lineage III) that was removed prior to phylogenetic analysis. The alignment of 84 PPRV genomes was then analysed using the Recombination Detection Program (RDP) version 4.101 [79], using seven different recombination detection methods (RDP, GENECONV, BootScan, MaxChi, Chimaera, SiScan and 3Seq) and default settings. Signatures of recombination were detected in three sequences (KR828814.1/goat/Nigeria/2012-05-09, KJ867541.1/goat/Ethiopia/2010 and KT633939.1/ibex/China/2015-01-20) using all seven detection algorithms (p<0.01) (S2 Table). These are most likely the result of laboratory contamination [47] and these sequences were considered unreliable and removed from our dataset, leaving a total of 81 PPRV genomes (alignment provided in S1 File).

### Phylogenetic analysis

The General Time Reversible (GTR) nucleotide substitution model with gamma-distributed variable rates (G) and some invariable sites (+I) best fitted our dataset, according to the Baysian Information Criterion (BIC) values calculated using MEGA7 [80]. Maximum likelihood (ML) phylogenetic reconstruction was performed using PhyML with a GTR nucleotide substitution model and 100 bootstrap replicates. Phylogenetic analysis was also performed using a Bayesian Markov Chain Monte Carlo (MCMC) framework using BEAUti and BEAST v1.10.4 [81] and run via the CIPRES server. Prior to traiting the sequences, marginal likelihood estimation was performed using path sampling/stepping-stone sampling to choose the most appropriate combination of tree model (coalescent constant size or coalescent GMRF Bayesian skyride) and clock models (strict or uncorrelated relaxed) using a GTR nucleotide substitution model (4 gamma categories, estimated base frequencies and no codon partitioning). MCMC outputs from different runs were evaluated and convergence confirmed using Tracer v1.7.1 [82]. A model with a coalescent constant size tree prior and uncorrelated relaxed clock (lognormal distribution) [83] was determined to be the best fit for the data, based on the log marginal likelihood estimates from path sampling/stepping-stone sampling. This model of nucleotide evolution was used for subsequent analysis of the discrete traits, ‘host category (livestock or wildlife)’ and ‘country’. Asymmetric substitution models were selected for the discrete traits since these are the biologically more plausible scenario of virus transmission. Bayesian Stochastic Search Variable Selection (BSSVS) was also used, which limits the number of rates to those which adequately explain the phylogenetic diffusion process. At least two independent MCMC chains, of 40 million steps each, were performed for each analysis, and Tracer v1.7.1 was used to confirm that the MCMC chains converged at the same level and assess effective sample sizes (ESS). LogCombiner was used to combine the output of the independent BEAST runs to generate tree and log files for analysis. Maximum clade credibility (MCC) trees were generated using TreeAnnotator v1.10.4. and Figtree v1.4.4 was used to visualise and interpret MCC trees and derive TMRCA estimates. Evolutionary rates (ucld.mean) were taken from the combined log files analysed in Tracer v1.7.1. MCC trees and geocoded metadata were imported into Microreact to visualise temporal and geographic spread [84]. The MCC tree and log files from the BEAST analysis were uploaded to the SpreaD3 software (Spatial Phylogenetic Reconstruction of EvolutionAry Dynamics using Data-Driven Documents (D3) [85], in order to visualize the output from the BSSVS procedure and compute Bayes Factors for transitions. For country transitions, each country was assigned one latitude and longitude coordinate, either the precise sampling location for those countries with a single PPRV sequence in the dataset (Kenya, Tibet, Ghana), the GPS location of the saiga3 sample for Mongolia, or the country centroid coordinates (worldmap.harvard.edu) for other countries with >1 sequence.

### Detection of selection pressures

Multiple analysis methods were implemented to detect positive selection in our phylogeny. Fast Unconstrained Bayesian Approximation (FUBAR, [86]) and Fixed Effects Likelihood (FEL, [87]) both available in datamonkey.org [88] were used to detect positive selection at individual sites across the whole PPRV phylogeny. Mixed Effects Model of Evolution (MEME, [89]) analysis was also performed on datamonkey.org, with the capability to identify sites under episodic selection (i.e. in a subset of branches) as well as under pervasive selection [64, 89]. In addition, the ratio of non-synonymous to synonymous nucleotide substitutions (ω = dN/dS) was estimated for different selection models (in which the ω ratio varies among codons) using CodeML as implemented in EasyCodeML [90]. The model M7 (beta; no positive selection) was compared to the model M8 (beta&ω; positive selection) for each gene using likelihood ratio tests (LRTs) [91, 92] (S4 Table). If model M8 was more likely than M7, the Bayes empirical Bayes (BEB) method [93] was used to calculate the posterior probabilities for site classes and identify sites under positive selection. Finally, three methods were implemented in datamonkey.org to detect linage-specific selection, using user-defined PhyML maximum likelihood trees and alignments for the CDS of each gene. The monophyletic clade of Mongolian/Chinese sequences was selected *a priori* to test for selection acting on these branches. Adaptive Branch-Site Random Effects Likelihood (aBSREL, [94]) was used to detect branches under positive selection. Branch-Site Unrestricted Statistical Test for Episodic Diversification (BUSTED, [95]) was also used to test for gene wide lineage-specific positive selection. Specific sites under selection in the selected clade were identified using Fixed Effects Likelihood (FEL, [87]).

### Protein homology modelling

The predicted structures of PPRV Hs were modelled by submitting the amino acid sequences of each H to the SWISS_MODEL automated protein structure homology modelling server in “alignment” mode [96]. Structures were visualised from the .pdb files using Swiss-pdb viewer.

## Acknowledgements

Support for field sample collection and multi-sectoral coordination was provided by the Wildlife Conservation Society, the Morris Animal Foundation’s Betty White Wildlife Rapid Response Fund, the FAO Crisis Management Centre-Animal Health and through the Science for Nature and People Partnership (SNAPP), a partnership of The Nature Conservancy, the Wildlife Conservation Society (WCS), and the National Center for Ecological Analysis and Synthesis (NCEAS) at University of California, Santa Barbara.

Special thanks to Khovd and Gobi-Altai Province Veterinary Agencies for their generous support for wildlife field investigations and response, and to Dr. Munkhbaatar Batsukh and Dr. Burenbayasakh Ravdan for their support.

## Supporting information

**S1 Fig. Molecular detection of PPRV N gene in different tissues from wild Mongolian ungulates.** RT-PCR for PPRV N gene was performed on 2ul RNA extracted from different tissue samples from the indicated host (RNA concentration ranged from 26-1125 ng/ul) prior to gel electrophoresis and UV transillumination. Lane numbers refer to sample IDs for different tissues given in Table 1. DNA ladder markers of different base pair (bp) lengths are shown. ‘x’ denotes an empty lane; ‘-’ denotes a no template control and ‘+’ denotes a positive control RNA template. Samples used subsequently for whole genome NGS are marked with *.

**S2 Fig. Untraited Bayesian time-scaled Maximum Clade Credibility Tree.** MCC tree from the combined output of three MCMC chains run in BEAST v1.10.4, inferred without partitioning the data by traits, and visualized in FigTree. The novel genomes from this study are shown in blue. x-axis shows date. Lineages, referred to as LI, LII, LIII or LIV, are marked. * indicates posterior probability > 0.9 at the node opposite.

**S3 Fig. TempEst analysis of temporal signal in the PPRV genome dataset.** Plots from TempEst showing root-to-tip genetic distance against sampling date (A) and residuals (B)for a ML phylogeny of 85 PPRV genomes, using the best-fitting root. The correlation coefficient for the regression was 0.9362 and R^2^ was 0.8764. An outlier sequence, shown in blue, was removed from the alignment before phylogenetic analysis.

**S1 Table. Illumina NGS read summary for wildlife samples.** Total read number, PPRV-specific read number and % total reads which were PPRV are shown for the five novel PPRV genomes from wildlife hosts. Average genome coverage was calculated as (read count * read length) / total genome size.

**S2 Table. Recombination analysis using RPD4.** The PPRV genome alignment (n=84) was analysed using Recombination Detection Program v4.101 (RPD4) using default settings. Genomes unambiguously identified as recombinant sequences are shown, for which at least 5 of 7 of the detection methods found significant evidence of recombination. Recombinant: genome sequence identified as potential recombinant; Lin-r: PPRV genetic lineage of Recombinant; Minor parental sequence: genome sequence identified as most likely minor parent of the recombinant, i.e. most closely related to the genome portion inserted; * indicates that other potential minor parents were also identified by RPD4 for that recombination event; Lin-mp: PPRV genetic lineage of minor parent; Begin: average genome position (in alignment) of the beginning breakpoint point of recombination; End: average genome position (in alignment) of the end breakpoint point of recombination; NS: not significant. The p-values for analysis with each of the 7 different recombination detection algorithms are shown, after Bonferroni correction for multiple comparisons.

**S3 Table. Bayes factors for spread of PPRV between countries.** Using SpreaD3, Bayes Factors (BFs) were calculated using the log file from the BEAST BSSVS analysis and a discrete set of longitude and latitude coordinates for each country, coupled to a geoJSON formatted world map. The output gave BFs for all possible transitions between locations. The table shows the transitions with posterior probabilities >0.7. The transition from China to Mongolia, which had the highest BF of any transition, is highlighted in grey.

**S4 Table. Values of Log-likelihood (lnL) for PPRV genes using different selection models in the CodeML analysis, and LRT comparing the two models.** Two different site selection models (in which the ω ratio varies among codons) were implemented in CodeML: M7 (beta; no positive selection) and M8 (beta&ω; positive selection). For each gene, Log-likelihood (lnL) values are shown and the likelihood ratio test (LRT) to show the significance of model comparison. Bayes empirical Bayes (BEB) were used to calculate the posterior probabilities for site classes and identify sites under positive selection.

**S5 Table: Evidence from aBSREL for episodic diversifying selection acting on the PPRV L gene.** The PPRV L gene was analysed using aBSREL with the China/Mongolia clade selected the set of foreground branches on which to test for episodic diversifying selection. Significance was assessed using the likelihood ratio test statistic for selection (LRT) at a threshold of p ≤ 0.05, after correcting for multiple testing. The two branches shown were identified as under positive selection, while all other branches were best described by a single ω rate category (ω1). The ω distribution shows inferred estimates for ω1 and ω2 and proportion of sites in each category.

**S1 File: Alignment of PPRV genomes (n=81)**

