## Supplementary figures and images for "Molecular epidemiology of peste des petits ruminants virus emergence in critically endangered Mongolian saiga antelope and other wild ungulates"

### S1 Fig

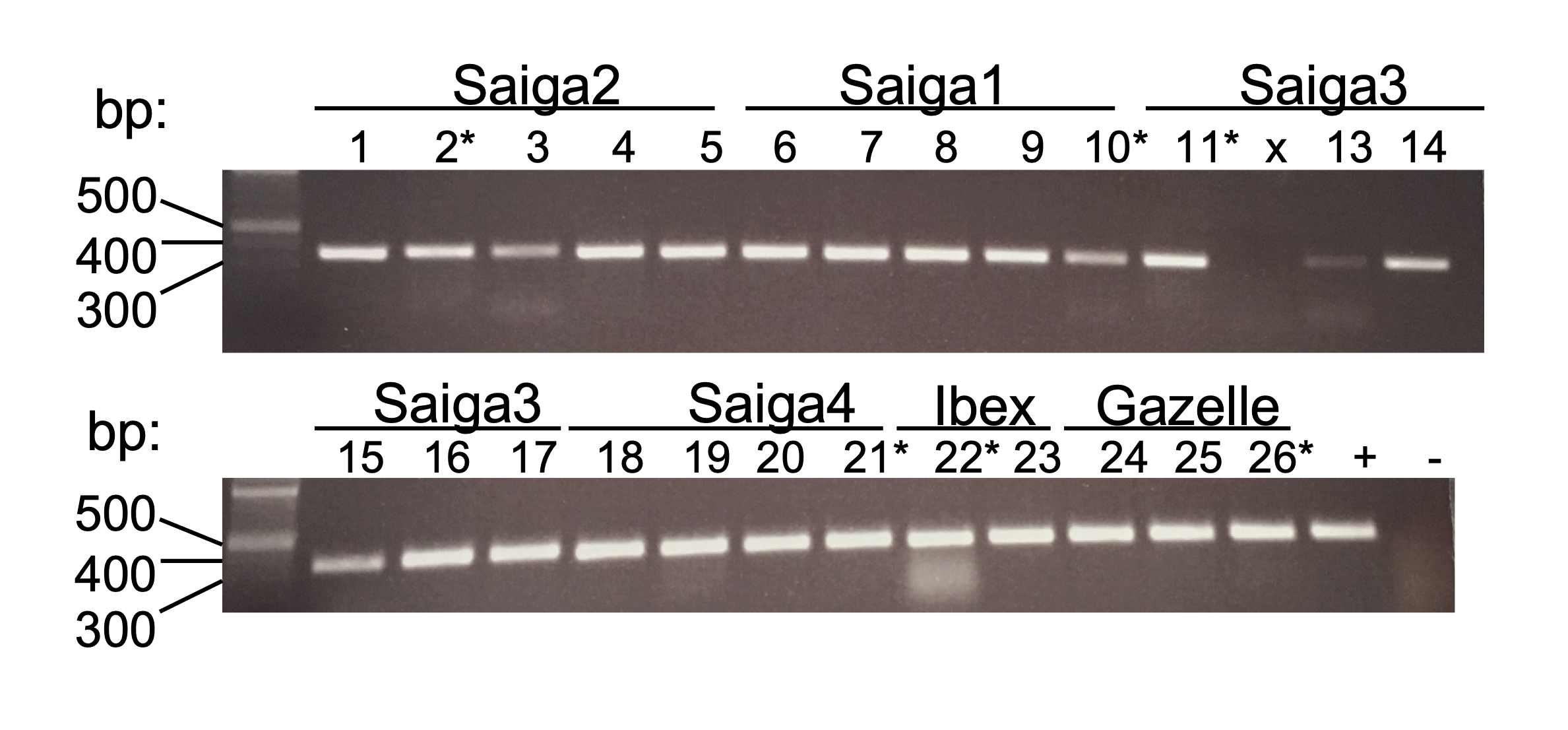

### S2 Fig

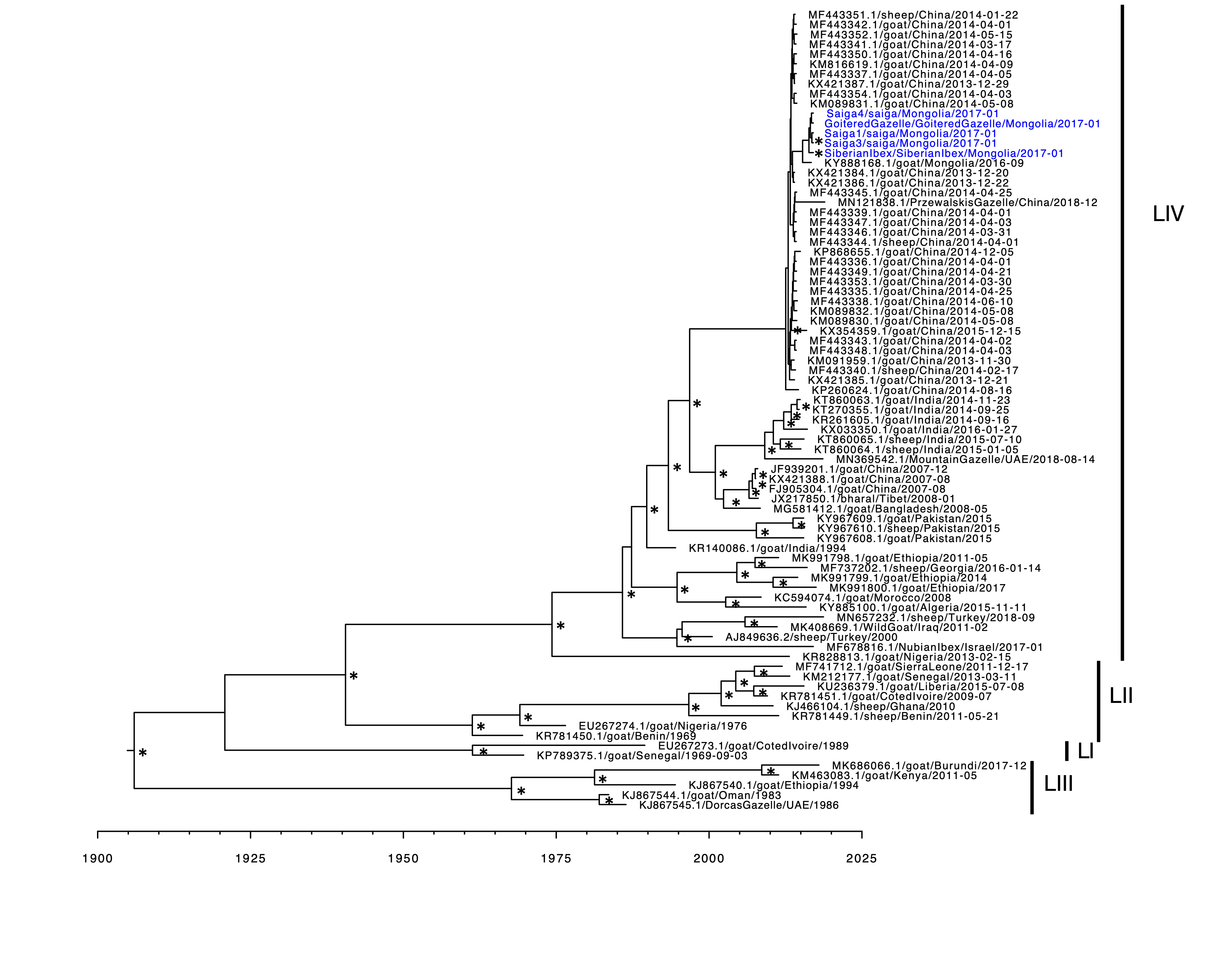

### S3 Fig

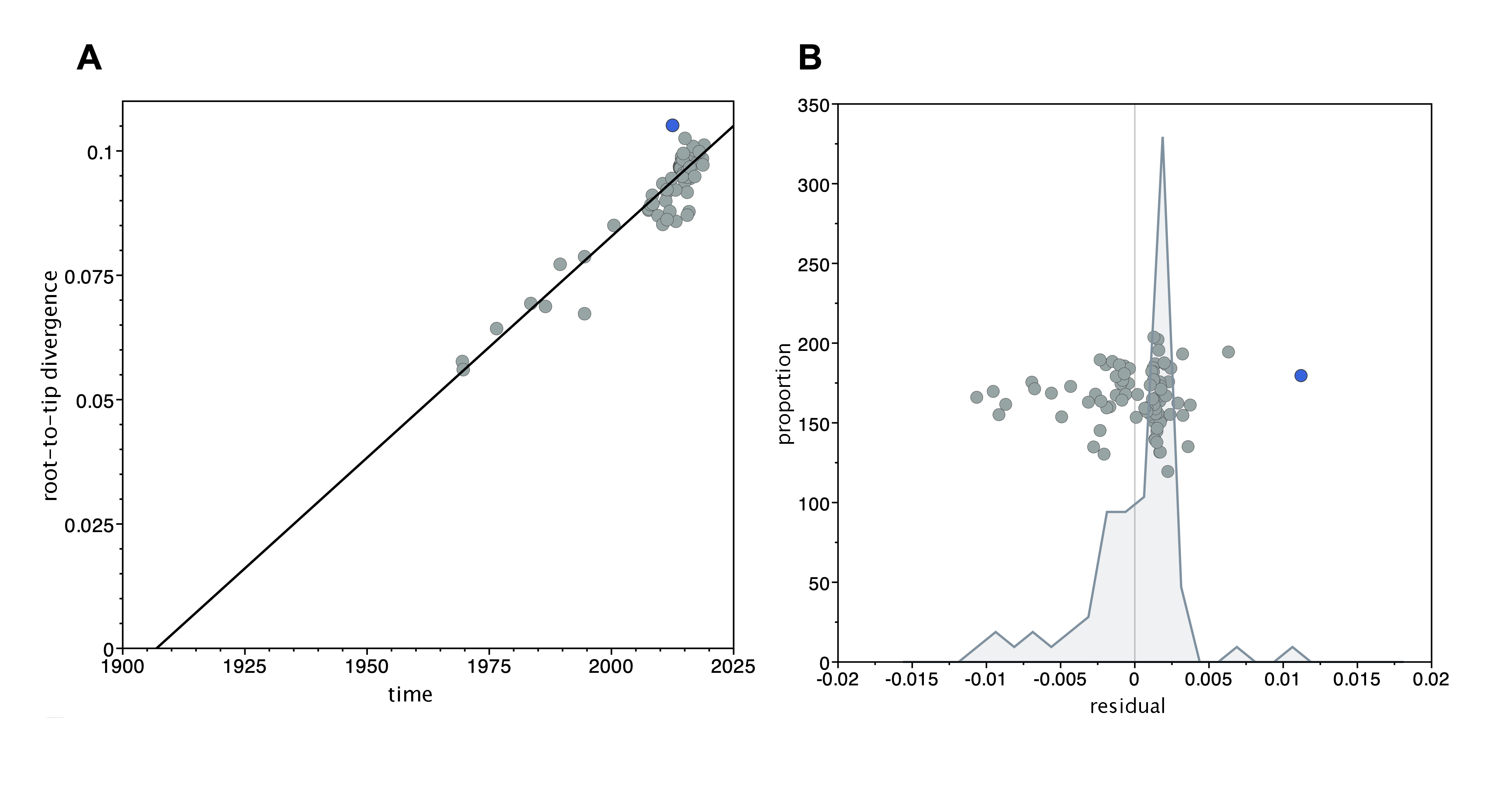
