## Supplementary material for "Molecular epidemiology of peste des petits ruminants virus emergence in critically endangered Mongolian saiga antelope and other wild ungulates": S1 Table

| Sample | Total reads | PPRV-specific reads | % PPRV | PPRV genome coverage |
|---|---|---|---|---|
| Saiga_1 | 34,322,736 | 1008 | <0.003 | x 9.5 |
| Saiga_3 | 20,500,220 | 318,857 | 1.56 | x 2998 |
| Saiga_4 | 33,389,334 | 122,645 | 0.37 | x 1153 |
| Goitered gazelle | 36,678,534 | 124,090 | 0.34 | x 1167 |
| Siberian ibex | 39,462,531 | 2,908 | 0.01 | x 27 |
