## Supplementary material for "Molecular epidemiology of peste des petits ruminants virus emergence in critically endangered Mongolian saiga antelope and other wild ungulates": S2 Table

| Recombinant Sequence | Minor Parental Sequence | Lin-r | Lin-mp | Breakpoint Positions In Alignment | | Detection Methods | | | | | | |
|---|---|---|---|---|---|---|---|---|---|---|---|---|
| | | | | Begin | End | RDP | GENECONV | Bootscan | Maxchi | Chimaera | SiSscan | 3Seq |
| KR828814.1/goat/Nigeria/2012-05-09 | KR828813.1/goat/Nigeria/2013-02-15 | II | IV | 3096 | 4001 | 1.38E-55 | 7.05E-55 | 1.12E-55 | 7.39E-14 | 4.71E-14 | 2.98E-15 | 1.37E-11 |
| KR828814.1/goat/Nigeria/2012-05-09 | KR828813.1/goat/Nigeria/2013-02-15 | II | IV | 6388 | 6848 | 4.74E-29 | 1.31E-24 | 4.81E-19 | 5.41E-08 | 2.50E-07 | 1.22E-06 | 1.37E-11 |
| KR828814.1/goat/Nigeria/2012-05-09 | KJ867541.1/goat/Ethiopia/2010 | II | IV | 550 | 1073 | 9.97E-13 | NS | 3.57E-15 | 3.57E-05 | 0.033456 | NS | 8.63E-07 |
| KJ867541.1/goat/Ethiopia/2010 | KC594074.1/goat/Morocco/2008 | IV | IV | 4138 | 5544 | 8.72E-41 | 1.47E-37 | 3.72E-39 | 1.09E-16 | 3.78E-16 | 7.72E-21 | 0.004423 |
| KT633939.1/ibex/China/2015-01-20 | KR781450.1/goat/Benin/1969 | IV | II | 9084 | 9556 | 3.20E-40 | 1.03E-32 | 1.56E-38 | 9.99E-10 | 5.40E-09 | 2.71E-09 | 1.03E-11 |
| KT633939.1/ibex/China/2015-01-20 | KR781450.1/goat/Benin/1969 | IV | II | 6623 | 6922 | 1.14E-21 | 5.33E-19 | 1.17E-20 | 0.00012 | 0.0009816 | 4.54E-05 | 2.16E-10 |
