## Supplementary material for "Molecular epidemiology of peste des petits ruminants virus emergence in critically endangered Mongolian saiga antelope and other wild ungulates": S3 Table

| From | To | Bayes Factor | Posterior Probability |
|---|---|---|---|
| China | Mongolia | 494 | 0.96 |
| Ethiopia | Georgia | 103 | 0.82 |
| Kenya | Burundi | 65 | 0.75 |
| Turkey | Iraq | 64 | 0.74 |
| China | Tibet | 56 | 0.72 |
| India | UAE | 51 | 0.70 |
