## Supplementary material for "Molecular epidemiology of peste des petits ruminants virus emergence in critically endangered Mongolian saiga antelope and other wild ungulates": S4 Table

| Protein | LnL M7 | LnL M8 | LRT |
|---|---|---|---|
| **N** | -6569.74 | -6554.43 | <0.0001 |
| **P** | -6830.54 | -6823.22 | <0.001 |
| **C** | -2877.59 | -2875.13 | 0.09 |
| **V** | -4191.50 | -4179.66 | <0.0001 |
| **M** | -4106.74 | -4106.74 | 1.00 |
| **F** | -5750.98 | -5749.07 | 0.15 |
| **H** | -8619.88 | -8613.37 | 0.002 |
| **L** | -25535.46 | -25513.47 | <0.0001 |
