## Supplementary material for "Molecular epidemiology of peste des petits ruminants virus emergence in critically endangered Mongolian saiga antelope and other wild ungulates": S5 Table

| Branch name | LRT | Test p value (corrected for multiple testing) | ω distribution |
| --- | --- | --- | --- |
| KP260624_1_GOAT_CHINA_2014_08_16 | 35.5617 | $2.91 \times 10^{-7}$ | ω1 = 0.327 (100%)<br>ω2 = 4150 (0.20%) |
| KY888168_1_GOAT_MONGOLIA_2016_09 | 12.3338 | 0.0330 | ω1 = 0.824 (100%)<br>ω2 = 1970 (0.21% |
