## Supplementary material for "Molecular epidemiology of peste des petits ruminants virus emergence in critically endangered Mongolian saiga antelope and other wild ungulates": S1 File

>KR781450.1/goat/Benin/1969

ACCAAACAAAGTTGGGTAAGGATAGATCTTATAATAACTATAGACTGGCAAACTTAGGAGTAAAGATCCTACTGTCGGGG

AGAGGAGGAGGAGCAAGATCTTTGACCATGGCGACTCTCCTTAAAAGCTTGGCATTGTTCAAGAGGAACAAAGACAAAGC

GCCTACTGCGTCGGGTTCAGGAGGGGCCATCCGGGGGATTAAGAATGTTATCATAGTCCCTATTCCCGGGGACTCATCCA

TCATTACCCGTTCAAGACTGCTCGACAGGCTTGTTAGATTGGCCGGAGATCCTGACATCAACGGGTCAAAGCTGACCGGC

GTGATGATCAGCATGTTATCTTTGTTCGTGGAGTCACCCGGGCAATTGATACAGCGGATCACAGATGATCCAGATGTTAG

CATCCGCCTTGTTGAGGTAGTTCAAAGTACTAGGTCCCAGTCCGGGTTGACCTTTGCATCACGTGGTGCTGATTTGGACA

ATGAGGCAGATATGTATTTTTCAACTGAAGGACCCTCGAGTGGAAGTAAGAAAAGGATCAACTGGTTTGAGAACAGAGAA

ATAATAGACATAGAGGTGCAAGATGCAGAAGAGTTCAATATGTTGTTAGCCTCCATCTTAGCACAAGTTTGGATCCTCCT

GGCCAAGGCGGTTACGGCACCGGATACGGCAGCTGACTCAGAACTGAGAAGGTGGGTTAAATACACACAACAAAGGAGAG

TGATTGGGGAATTTCGCCTTGACAAAGGGTGGCTGGACGCAGTCCGCAACAGGATTGCAGAAGATCTATCACTTCGGCGG

TTCATGGTATCTCTCATACTTGACATCAAAAGGACCCCCGGCAACAAGCCAAGGATTGCAGAAATGATCTGCGACATTGA

CAACTATATTGTCGAAGCCGGACTCGCCAGTTTCATTCTTACTATCAAGTTTGGTATTGAAACCATGTATCCTGCATTAG

GCCTTCACGAGTTCGCCGGGGAATTGTCCACTATTGAATCCTTGATGAACTTGTATCAACAGCTAGGAGAGGTTGCACCC

TACATGGTAATTCTAGAGAACTCAATTCAGAACAAGTTTAGTGCAGGAGCCTATCCCCTCCTCTGGAGCTATGCGATGGG

TGTCGGAGTCGAGTTGGAGAACTCAATGGGGGGCCTGAACTTTGGCAGGTCATATTTTGACCCGGCCTATTTCCGTCTCG

GACAGGAGATGGTCAGAAGATCTGCAGGAAAGGTCAGCTCTGTAATCGCGGCTGAGCTTGGCATCACAGCAGAGGAAGCC

AAACTAGTCTCGGAAATCGCCTCGCAGACTGGAGATGAACGAACCGTTAGAGGGACTGGGCCTCGACAGGCGCAGGTTTC

CTTCCTCCAGCATAAAATAGATGAGGGAGAGTCGCCTACACCAGCGACCAGAGAAGAAGTCAAAGCTGCGATCCCAAATG

GGTCTGAAGGAAGGGACATAAAGCGAACACGCTCAGGAAAGCCCAGAGGAGAAACTCCCGGCCAACTGCTTCTGGAGATC

ATGCCAGAGGATGAAGTCTCGCGAGAGTCTAGTCAAAACCCTCGTGAGGCTCAAAGATCGGCTGAGGCACTCTTCAGGCT

GCAGGCCATGGCCAAGATTCTGGAGGACCAGGAGGAGGGAGAAGACAACAGTCAGATCTACAACGACAAGGATCTCCTCA

GCTGAGCAGACGCACCCTCCGTCTAAATCAGTGACAAGACATCGCCCGCCAGTATTATAAAAAACTTAGGACCCAGGTCC

AAGCAACCACACATCGACACCCCAGTCAATCGAGCAGAGACCACCGATGGCAGAAGAACAAGCATACCATGTCAACAAGG

GACTGGAATGTATCAAGTCTCTCAAAGCCTCTCCCCCGGATCTATCCACCATCAAAGATGCCCTTGAGAGCTGGAGAGAG

GGGCTTAGCCCCTCAGGCCGTGCAACACCGAACCCTGATACGTCCGAGGGAGACCATCAGAATATCAACCAATCATGCTC

ACCAGCAATCGGATCAGACAAAGTCGACATGTCTCCTGAAGATAATCTCGGATTTAGAGAGATCACTTGTAATGACAGTG

AGGCTGGGCTCGGAGGAGTTCAGGACAAAGGATCCAACTCTCAAGTACAGCGTTACTATGTTTATAGCCACGGGGGTGAA

GAGATTGAAGGACTCGAGGATGCTGACTCTCTCGTGGTTCAAGCAAATCCTCCAGTTGCTAACACCTTCAATGGAGGAGA

GGATGGATCTGACGACAGCGATGTGGACTCTGGCCCAGATGATCCCGGCAGAGATCCTCTATCTGACCGGGGATCTGCTG

CCGGCAATGATGTCTCTAGGTCAACAGATGTCGAAAAATTAGAAGGTGATGACATTCAAGAAGTTCTTAACTCCCAGAAG

AGTAAAGGAGGAAGATTCCAAGGCGGGAAAATCTTGCGGGTCCCGGAAATACCCGATGTCAAGAACTCCAGACCATCAGC

CCAATCAATTAAAAAGGGCACAGACGGGAACTCAGTCTTATCTGGAACGGTGACAGAGTGTTCATCGATAAGTGGTGCAA

CCCAAGCTGTGCCAGAGTCCAGATGGGAGTCATCAGGGCGAAATGCGTCTGTGGGGAGTGTCCCCAAATCTGCGAGGAGT

GCAAAGACGATCCAGGGGTTGACACAAGAATCTGGTACCATAGCATCACTGACTCAGCCTAAAGAGAATGACTCCGAGTA

TGAGTATGAGGATGATCTATTCACAGAGATGCAGGAGATTCGTGCAAGCATTGCTAAGATCCATGATGACAACAAAACTA

TCCTCTCAAAACTTGATTCTCTACTGTTATTGAAAGGAGAAATCGATACTATCAAGAAACAAATCAGCAAACAAAATATA

AGTATATCTACCATTGAGGGGCATCTATCCAGTATAATGATAGCCATCCCGGGCTTTGGGAAGGACATCAAGGACCCAAC

ATCTGAGGTTGAGTTGAACCCGGATTTGAGACCTATAATCAGCCGTGATTCTGGCAGGGCTCTTGCGGAGGTCCTCAAGA

AACCCGCTGTTGATAGGTCTCAGAAAAGCGGAATCAAAGTCAACTCCGGTTCAAAGGGTCAGCTCCTTAAGGATCTCCAG

CTAAAACCTGTCGACAAACAGGCAAGCTCTGCAATCGGGTTTGTTCCATCTGACCATGAATCATCCAGGAGTGTCATCCG

CTCCATAATCAAGTCGAGCAAGCTTAACATTGATCACAAGGACTATCTTCTAGATTTACTGAATGATGTGAAAGGCTCTA

AGGATCTTAAGGAATTCCACAAGATGCTAACAGCCATTCTTGCCAAGCAGCCGTAACACATCCTATAATCAACATCTCAT

ACTCGGTTGAAAACATCCTCTCAATCAGGCTATTACAAAAAACTTAGGAGCAAGGGCAACTGAGCTTCACAGACAAGATG

ACCGAGATCTACGATTTCGATAAATCAGCATGGGATGTCAAAGGGTCAATTGCTCGCATAGAACCCACCACCTATCACGA

CGGCCGACTGATACCCCAGGTGAGGGTCATCGATCCTGGTCTGGGAGACAGAAAAGATGAGTGCTTCATGTACCTGTTTC

TTCTAGGAGTGATTGAGGATAACGACCCCCTGTCTCCCCCAGTCGGGAGAACCTTCGGGTCTTTACCTCTAGGGGTCGGT

AGATCAACCGCTAAGCCAGAAGAACTACTAAGGGAGGCCACAGAACTAGATATAGTGGTGAGGCGCACGGCAGGACTAAA

TGAGAAACTGGTATTTTACAACAACACTCCGCTGTCTTTGTTAACACCTTGGAGGAAAGTCTTGACAACCGGAAGTGTGT

TTAGCGCTAACCAGGTTTGCAATGCAGTCAACCTAGTCCCACTTGATACTCCCCAGAGATTCAGGGTTGTGTACATGAGT

ATAACTAGATTGTCAGATAATGGTTATTATAGTGTCCCCAGAAGAATGTTGGAGTTCCGCTCAGCCAATGCAGTCGCCTT

CAATATCTTGGTTACACTGAGAATTGAAAATGGCACAAACCCTAGAAGATACATAGTCGGCTCATGGGAGAATCCAGAAG

TCACATTTATGGTACACGTGGGCAACTTTAGAAGAAAGAAGAACGAAGTATACTCTGCTGATTATTGCAAAATGAAGATT

GAAAAGATGGGTCTAGTTTTTGCCTTGGGAGGAATAGGTGGAACAAGTCTCCATATTCGAAGCACAGGGAAAATGAGCAA

GACCCTCCATGCACAGCTGGGGTTCAAGAAAATCCTATGCTACCCCCTAATGGATATTAATGAGGATCTTAACCGATATC

TCTGGCGGGCAGAGTGCCGAATAGTCAAAATCCAGGCCGTCTTACAGCCATCAGTACCCCAAGAATTCCGTGTCTATGAT

GATGTCATCATCAACGATGATCAAGGCCTGTTCAAGATCCTGTAATTCACTTGCAGGGCCACGGACCCACCTCCAGGGAG

AGGCCCTCCTCTCTCCCGTCCAAAACATACACCCCAAGGTTCCCCGAAACAGCACTGACACAGCCACTCTCCCGGCGGGA

CGAGGCCGACT-CCCCTAACAAACTCGCTCCGGGGGGGGGCCCCCCCCCACGACCCGCCGGGCGGCGACCGCGCGAGCCG

GCGAGACACCCCCAAGAGGTCCCGACGAAGACCGGGCAACC--CCCCCCA-GAAAAGGCCCCCC-AACACACCCGAGGCC

AGGCGCCGGGCACCCCCAGCCCCGCCACCGGCGGGACAGGAGGACAAGACGGGGTCCCCCCAACCAAAGGGACCACCAGG

GAAGGCTCTGCCGACCCAGCACAAACCCGGCCCAAACATGTCAGACCCCCAAGACGAAATGCCCCAGCACACCCCATTCA

TCATCAAACAAAACTTAGGGGCCAAGTCCAGACGATAGACCCCAGTCCGCCAGGCAGCAAGCCGCAGAAAGGAAGGAGAC

AAGACCGGCCCGGAAGGAGGACACCCAACCACAGAAACCCCCAACCCCAGGGGAGGAG-CCCCCCCTCCGCACGGCGGAG

GAACGGGCGAGGGACACCGAGGAGAACCCGATCGGGAGCCAGGAGCCGGAGACGCCAGCGGCAAACAGGCCGGGCACACA

CCCGACACCACCGGAGGACCAGAGAAGAGCCCGACCCCGCGTGGACCCGACGGCAGGGGACCCAACCCCGCCACAAAGAG

AGGCGAGAACCCTC-------CCCCCCCGCACACAACCGGCAGACAGACCGAGGGAGGGCCGCCTCCCAACAAGACAGGC

GGCTCCCCCGCAGCCGAGCCGCAGGTCCCAAGAAGAGGACACCCCCGAAACCTCCCAGAACCCCTCCGCTCCCGGCCGCC

GGGACAGGAGCTCCGCCGCCAAAGACCAGAGCCACCGAGAGCCAACCAGGGGCCCCCCCCAGTCCCCCCCGGCGCCCTCC

CACAGAGGCCCCAGCAAAGTCCCCCGGCCCGACGACCCCTGCATGCGCCGAAAGAGAAACCCACCAAAGGACGTCCAGCA

CATCCATGTATAAACATCATGACACGGGTCGCAACCTTGGTATTTCTGTTTCTTTTCCCAAACACTGTCGCGTGCCAGAT

TCACTGGGGCAATCTATCCAAGATCGGGATTGTAGGAACAGGGAGTGCCAGTTACAAGGTGATGACTAGGCCAAGCCACC

AAACTCTGGTCATAAAGTTGATGCCAAATATAACAGCCATCGACAATTGTACGAAATCAGAGATTTCAGAGTACAAAAGA

TTGCTGATCACAGTGTTAAAGCCTGTAGAGGATGCCCTGTCAGTGATAACCAAGAATGTAAGACCAATTCAAACTCTAAC

ACCTGGGCGCAGGACCCGCCGTTTTGCCGGAGCTGTTCTGGCCGGAGTAGCACTTGGAGTCGCGACGGCTGCTCAAATAA

CTGCCGGAGTCGCACTCCATCAGTCATTGATGAATTCCCAAGCAATTGAAAGTTTAAAAACCAGTCTTGAGAAGTCGAAT

CAGGCAATAGAAGAAATCAGACTTGCAAATAAGGAGACCATACTGGCGGTACAGGGCGTCCAAGATTATATCAACAATGA

GCTTGTCCCCTCTGTTCATAGAATGTCATGTGAGCTGGTAGGTCACAAACTCAGTCTCAAGCTCCTTAGGTATTATACCG

AGATCCTGTCTATATTCGGGCCTAGCCTTCGAGACCCGATAGCTGCTGAAATATCAATCCAGGCACTCAGCTATGCATTA

GGCGGAGACATCAATAAAATTCTGGACAAGCTTGGGTATAGCGGCGGGGATTTCCTTGCTATCCTAGAAAGCAAGGGGAT

AAAGGCCCGGGTCACATATGTGGACACAAGAGATTACTTTATAATTCTTAGCATAGCCTACCCAACCTTATCTGAGATCA

AAGGGGTGATAGTTCATAAGATAGAAGCTATATCATACAACATTGGGGCACAGGAATGGTATACTACTATCCCTAGATAT

GTAGCCACTCAGGGATATCTGATATCGAATTTCGATGAGACGTCATGTGTCTTCACTCCAGAGGGGACAGTCTGCAGCCA

GAATGCGTTGTATCCAATGAGCCCATTGCTTCAGGAATGCTTCAGGGGGTCGACAAAATCGTGCGCCAGAACCCTAGTTT

CAGGGACCACAAGTAATAGATTTATCCTATCAAAAGGGAACTTGATTGCAAATTGTGCGTCAGTTTTGTGCAAGTGTTAC

ACAACGGAGACAGTTATCAACCAAGATCCTGATAAACTACTAACTGTTATAGCCTCCGATAAGTGTCCCGTAGTCGAGGT

GGATGGAGTGACAATACAGGTCGGCAGTCGAGAGTACCCAGATTCTGTATACCTACATAAAATAGACTTAGGCCCAGCCA

TCTCCCTGGAAAAACTGGATGTAGGCACCAATTTAGGCAATGCAGTCACAAGACTGGAGAATGCAAAGGAGCTACTAGAT

GCATCAGACCAGATACTGAAGACTGTTAAAGGGGTACCTTTCAGTGGCAATATGTACATAGCACTGGCAGCTTGCATTGG

GGTATCCCTAGGGCTTGTCACATTGATATGCTGCTGTAAGGGGAGATGTAGGAACAAGGAGATTCCTGCCTCCAAAATCA

ACCCAGGGCTCAAACCCGACCTGACCGGGACTTCAAAGTCGTACGTGAGATCACTGTAGTCAGAATAACCCGAATCATCC

AGCATCACACATATACATGTGCGACACAAGCAGTCAGAGGACGCAGAAGATTCAACTTCCGATCACCGACCAGACCCCAC

TCTACGCCCTATTACACATTGGTCATCAAACAAAACTTAGGACGAAAGGTCAATCACCATGTCCGCACAAAGGGAAAGGA

TCAATGCCTTCTACAAAGACAATCTTCATAATAAGAACCATAGGGTAATCCTGGATAGGGAACGCTTAACTATTGAAAGA

CCCTACATCTTACTTGGAGTCCTGCTGGTAATGTTCCTGAGTCTAATCGGGCTGCTGGCCATTGCAGGGATCAGGCTTCA

CCGGGCCACCGTTGGAACTGCGGAGATCCAGAGTCGGCTGAATACCAACATTGAGTTGACCGAATCCATTGATCATCAAA

CTAAGGATGTCTTAACTCCCCTGTTTAAAATCATTGGTGATGAAGTCGGCATCAGAATTCCACAGAAGTTCAGTGATCTT

GTCAAGTTCATCTCCGATAAGATTAAGTTCCTCAACCCTGACAGAGAATATGATTTTAGGGATCTCCGGTGGTGTATGAA

CCCCCCTGAGAGAGTCAAAATTAACTTTGACCAGTTTTGTGAATACAAAGCCGCGGTCAAGTCAGTTGAACATATATTTG

AGTCATCACTCAACAGGTCAGAAAGGTTGCGATTATTGACTCTTGGGCCCGGAACAGGCTGTCTCGGCAGGACAGTAACA

AGAGCTCAGTTCTCAGAGCTTACGCTGACCCTGATGGACCTGGATCTCGAGATGAAGCACAACGTGTCCTCAGTGCTTAC

CGTAGTCGAAGAGGGATTATTCGGAAGAACATATACTGTCTGGAGATCCGACACCGGAAAACCGAGCACCAGTCCAGGTA

TTGGCCATTTTTTAAGAGTCTTCGAGATCGGGCTGGTGAGAGATCTCGAGCTGGGTGCCCCTATTTTCCATATGACCAAC

TACCTCACAGTGAACATGAGTGATGACTATCGGAGCTGTCTTTTAGCAGTAGGGGAGTTGAAGCTGACAGCCCTATGCAC

CCCATCTGAGACTGTGACTCTGAGTGAGAGAGGAGTTCCAAAGAGAGAGCCTCTTGTGGTTGTGATACTCAACCTAGCTG

GGCCTACTCTAGGGGGCGAACTATACAGTGTATTGCCTACCTCTGACCCCACGGTGGAGAAACTCTATTTATCCTCACAT

AGGGGGATTATCAAAGATAACGAGGCCAATTGGGTAGTACCGTCTACCGATGTTCGTGATCTTCAAAACAAAGGAGAATG

TCTGGTGGAAGCATGCAAGACTCGACCTCCTTCATTTTGCAATGGCACAGGAATAGGCCCATGGTCAGAGGGGAGAATCC

CTGCCTACGGGGTGATCAGGGTCAGTCTTGACTTAGCTAGTGACCCGGGTGTAGTTATCACTTCAGTGTTTGGCCCATTG

ATACCTCACCTATCCGGCATGGATCTTTACAACAATCCGTTTTCAAGAGCTGCATGGTTGGCTGTACCACCTTATGAGCA

GTCATTTCTAGGAATGATAAATACAATTGGCTTCCCGGACAGAGTAGAGGTTATGCCGCACATTTTGACCACAGAGATCA

GAGGGCCTCGGGGTCGTTGTCATGTTCCTATAGAGTTGTCCCGCAGGATTGATGATGATATCAAGATCGGGTCCAACATG

GTTGTATTGCCGACGAAGGACCTGAGGTACATAACAGCCACTTATGATGTTTCCAGGAGCGAGCATGCAATCGTGTACTA

TATCTATGACACGGGTCGCTCATCATCTTACTTCTACCCAGTTCGATTGAATTTCAAGGGCAATCCTCTCTCTCTGAGGA

TAGAGTGTTTTCCCTGGTATCATAAGGTGTGGTGCTACCATGATTGTCTTATATACAACACCATAACAAACGAAGAAGTC

CATACGAGAGGGCTGACCGGTATAGAGGTAACATGTAATCCAGTCTGAGTAGAGCTGCAACCATCGCTCAAGCAGGCCTC

CGAGCTATCCCCTAGTCCAAGCAGCATAGCCTGGGACACTCAGCAGCACAACCCAGCCAACAATGTTATAAAAAACTTAG

GAGCCAAGGTTGTAGGAGCCATGGACTCACTATCAGTCAATCAGGTCTTGTACCCTGAGGTCCATCTAGATAGCCCTATT

GTCACAAACAAACTAGTTGCCATCCTTGAATACTCGGGGATCGACCACAACTATGTTCTTGAAGACCAGACCCTTATCAA

GAATATTAGATATAGACTGGGGTGCGGTTTTTCAAATCAAATGATCATCAATAATAGGGGGGTAGGTGAAACAGTCAATT

CCAAACTTAAAAGTTACCCCCGTAATTGTCATATCATATACCCAGACTGCAATAAGGATTTGTTTTGTATCAAAGATAGC

TGCATATCTAGGAAGCTCTCGGAGCTATTCAAGAAGGGTAATTCCTTGTACTCTAAGATAAGTCACCAGGTACTGGATTG

TCTTAAGAGAGTCAACGGGAAATTAGGCCTGGGCACAGATCTTACTCACGGCCTGAAGGAGGGTATCCTCGACTTGGGGT

TGCACATGCATAGCTCTCAATGGTTCGAGACCTTTCTGTTCTGGTTCACTATCAAGACAGAGATGAGATCAATGATCAAA

GAACAGTCCCATATATGCCACAAGAGGAGGTATAACCCAATTTTTGTGTCGGGGGATGCCTTCGAGGTGCTCGTATCACG

AGACCTCGTTGTGATAATTGATAAGAACACCCAGTATGTCTTCTACCTGACATTTGAGCTGGTCCTTATGTATTGTGATG

TCATAGAGGGCAGACTTATGACGGAGACAGCCATGGCTATAGACCAGAGATATTCAGAGCTTCTAAACCGGGTCAGATAC

TTGTGGGATCTTATTGATGGGTTCTTCCCAACACTGGGTAACACCACATACCAAGTTGTTGCTCTGCTTGAACCACTGTC

ATTGGCTTATCTTCAACTTCAGGATGTCACTCTAGAGTTAAGAGGTGCTTTTTTGGACCACTGCTTCAAAGAACTTTATG

AGATACTGGAGCATTGTGGCATTGACACGGAAGGTACCTACAATTCCATCACTGAAGGATTGGATTACGTATTTATCACC

CACGATATACACTTAACTGGGGAGATTTTTTCATTTTTTCGGAGTTTCGGACACCCCCGCCTCGAAGCGGTCACCGCTGC

AGAGAATGTCAGAAAACATATGAACCAACCGAAGGTAATCAGTTATGAGACTATGATGAAAGGGCATGCAGTATTTTGCG

GGATAATCATAAATGGTTTTAGGGACCGGCACGGCGGCAGCTGGCCCCCTGTTGCATTGCCAGAACATGCTTCTGCTGCG

ATCCGGAATGCGCAGGCATCCGGTGAAGGACTGACCCATGACCTGTGTATAGACAACTGGAAGTCCTTTGTTGGATTCAG

ATTTGGCTGCTTCATGCCGCTCAGCCTAGATAGTGATTTGACCATGTACCTCAAAGACAAAGCACTGGCTGCACTGAAGA

ATGAGTGGGATTCAGTTTACCCGAAAGAATACCTCCGTTATAATCCACCTAGAGGGACAGAGTCAAGGCGACTGGTAGAG

GTGTTCCTGAATGACTCCAGCTTTGATCCTTATAACATGATAATGTACGTGGTGAATGGCTCCTACCTTAAAGACCCTGA

GTTTAATCTCTCATACAGCTTAAAGGAGAAGGAGATAAAAGAGACAGGGCGATTGTTTGCCAAAATGACCTACAAGATGC

GGGCCTGTCAGGTAATTGCTGAAAATCTGATATCAAATGGTGTTGGGAAGTATTTCCGAGACAATGGGATGGCAAAAGAC

GAGCATGACCTAACAAAAGCCCTTCACACTCTGGCAGTCTCAGGTGTTCCTAAGAATAACAAAGATAACCACCGAGGTGG

ACCTCCCAGAAGGACCACAAGCCGAGAGATGAGATCAAGCCAAGACATCAACACACAAAATAGAGACAAGATCCAAGGGG

GCCCTATGTACAACTACTTGCGATGCCAACCGACCAGCCCTGATCAGGGTGAGTCATACGAGACTGTTAGTGCATTCATC

ACCGCTGACCTTAAGAAGTATTGCCTAAATTGGAGATACGAGACAATCAGCATATTTGCACAGAGACTGAATGAAATATA

TGGGTTGCCATCCTTCTTTCAATGGTTACACAGGATATTGGAAAAATCCGTGCTCTACGTCAGTGATCCGCATTGCCCTC

CCGACTTAGATGATCATATCCCTCTGGACAGTGTCCCTAATGCCCAAATATTCATCAAGTACCCAATGGGCGGAATAGAA

GGTTATTGTCAAAAACTATGGACAATCAGTACTATACCTTACTTGTATCTGGCAGCCTATGAGAGCGGAGTAAGAATCGC

CTCACTAGTGCAAGGTGACAATCAGACAATTGCAGTGACAAAAAGAGTTCCAAGTTCTTGGCCTTATTCACTAAAAAAGA

GGGAGGCATCCAAAGCAGCTCAAAATTACTTCGTGGTCTTAAGGCAAAGGTTGCACGATGTAGGTCATCACTTAAAGGCT

AATGAGACCATAGTATCTTCTCATTTTTTTGTATACTCTAAAGGGATTTATTATGACGGCCTGTTAGTCTCACAATCACT

AAAGAGCATCGCCAGATGTGTCTTCTGGTCCGAGACCATTGTGGATGAAACCAGAGCGGCCTGCAGCAATATTGCAACAA

CTATCGCCAAGAGTATAGAGAGGGGTTATGATAGGTACCTCGCATACTCTTTGAATATCCTCAAAATTTTCCAACAGATC

CTTATATCTCTTGACTTCACGATTAACACAACAATGACTCAAGATGTCGTGGCACCGATCATCGAGAACGGTGATTTACT

AATAAGGATGGCACTCTTGCCAGCACCCATTGGGGGTCTCAATTATCTTAACATGAGCAGGTTATTTGTGAGAAATATCG

GTGACCCGGTCACTTCCTCCATAGCCGACCTGAAGAGGATGATAGACGCTGGGCTAATGCCAGAAGAAACATTGCATCAA

GTGATGACCCAGACCCCGGGAGAATCATCCTACCTCGATTGGGCAAGTGACCCTTATTCTGCCAACCTAACCTGCGTACA

GAGTATAACTCGCCTTCTCAAGAACATCACTGCACGGTATATCTTAATCAGCAGCCCAAATCCGATGCTGAAAGGATTGT

TTCATGAGGGGAGTAGAGATGAAGACGAGGAGCTTGCGAGTTTCTTGATGGATCGGCATATAATTGTTCCGAGAGCTGCA

CATGAAATCTTAGACCATAGCATAACCGGAGCAAGAGAAGCTATAGCCGGGATGTTGGACACCACCAAGGGTCTGATTAG

AACAAGTATGAAGCGGGGTGGCCTCACCCCTCGAGTATTAGCCCGCCTTTCCAATTATGATTATGAACAATTCAGATCCG

GGATAACGTTATTGACAAAGAAAGGGCAGTGTTATCTCATTGACAAGGACTCGTGCTCGGTGCAGCTCGCTATAGCTCTG

AGGGGCCACATGTGGGCCAGGTTAGCTCGCGGGAGACCTATCTATGGGTTGGAGGTGCCTGATATACTGGAATCGATGAA

CGGCTACCTTATCAAACGCCACGAGTCTTGTGCCATCTGTGAAACGGGCTCAAGTCACTACGGGTGGTTTTTCGTCCCTG

CAGGGTGCCAGCTTGACGATGTCTCAAGAGAGACTTCGGCTCTTCGTGTACCTTATGTCGGATCAACCACTGAGGAAAGG

ACAGATATGAAACTTGCTTTCGTTAGATCTCCAAGCCGATCCCTCAAATCAGCAGTCAGAATTGCCACAGTTTACTCATG

GGCCTACGGGGATGATGAGAAATCATGGAGTGAAGCTTGGATGCTAGCTAGGCAGAGAGCTGATATCACCTTAGATGAAT

TGAGAATGATCACTCCAGTCTCTACATCCACCAACCTAGCCCATCGGTTGAGGGACCGGAGCACCCAGGTGAAATATTCG

GGGACATCCCTTGTGAGGGTTGCAAGATACACAACCATCTCCAATGACAATTTGTCATTTGTGATATCTGAGAAAAAAGT

AGATACCAACTTCATTTACCAGCAAGGGATGCTGCTCGGTCTTGGGATCCTTGAGAATCTCTTCAGGTTAGAGGCCACCA

CAGGGGTATCCAACACAGTGCTACACCTGCACGTGGAAACAGAATGTTGTGTTGTACCCATGGTTGATCACCCAAGGATA

CCGAGTCTCCGTAATATTAAAGTTACGAATGAGCTATGCACAAACCCTCTGATCTACGACAGGTCCCCCATCATAGAACA

CGATGCAACTCGATTATACTCACAAAGCCACAGGAGACATTTGGTGGAGTTTGTTACCTGGTCAACAAGCCAGCTTTATC

ATATACTGGCCAAATCTACAGCAATGTCCATGATTGAGTTGATCACAAGATTTGAGAAAGATCACATGAATGAAATAGCC

GCCCTGATTGGCGATGACGACATCAACAGTTTCATCACAGAATTTTTGCTAGTAGAGCCCAGACTGTTTATAGTTTACCT

TGGCCAGTGTGCTGCCATCAATTGGGCTTTTGATATACATTATCATCGGCCCTCGGGCAAGTACCAGATGGGGGAACTCC

TCTACTCTTTACTCTCTCGGATGAGCAAAGGAGTATATAAGATCTTCACTAATGCTTTGAGCCACCCCAAAGTTTACAAG

AAATTTTGGCGAAGTGGTGTAATTGAGCCGATTCATGGCCCATCCCTAGATACACAGAATTTACATGTCACTGTCTGTGA

CATGATATACGGATCATACGTCACCTATCTGGATCTTTTGCTGAATGATGAGCTAGATGATTACCCGTATTTGCTCTGCG

AGAGTGATGAGGACGTGGTCACAGACAGGTTCGACAACATTCAAGCCAAACATCTCTGTGTACTGGCCGATGTATACTGC

AGCTCCAAGAGATGTCCCTCGATAATTGGGATGTCTCCTATAGAAAAATGCACCATCCTCACACATTACATCAAGGGAGA

ATCGGTACAATCCCCGTCCGGGATCTCATGGAACACTGACCCCCTTGTAGTAGATCATTACTCATGCTCTCTGACCTACC

TTCGCCGCGGTTCCATCAAACAAATCAGGTTGAGAGTGGATCCTGGGTTTGTATTCGAGGCGTTGACAGACATCGACTTC

AAACAGCCTCGCAAGGCTAAGTTGGATGTATCGGTTGTGGGGTTGACTGATTTTTCTCCCCCTTGGGATAACGTCGGTGA

TTTTCTAGGGACTATCAACACATTGAGGCACAATCTGCCCGTCACCGGAACCGGGGTCTCGAACTATGAAGTCCACGCTT

ATCGTAGAATCGGCCTGAATTCATCAGCATGTTATAAAGCTGTAGAGATCTCCACGTTAATCAAGTCATCTTTAGAAGTC

GGAGAGAATGGATTGTTCTTAGGAGAAGGTTCCGGTTCTATGCTGGCTGCGTACAAGGAAGTTCTTAAATTGGCAAGCTG

TTATTACAACAGCGGGGTAACAGCGGAGGGCAGAGCCGGACAGAGGGAAATCTCTCCCTATCCCTCAGAGATGAGCCTAG

TAGAGAATCAGATGGGGATAGAGAGGAGTGTTAAAGTGCTGTTCAATGGCAAACCTGAAGTAACCTGGGTAGGGACCACC

GATTGCTACAAGTATATAATCAGTAACATTCAGACCTCTAGTCTGGGTTTCATACACTCAGATATTGAGACACTCCCAAC

CAAGGATGCCGTTGAGAAGTTAGAAGAATTTGCCTCTATCCTATCCCTATCCCTAATTTTGGGAAAAATCGGCTCTATTA

CAGTTGTCAAAATTATGCCCATTAGCGGAGATTTTACCCAAGGCTTCATAGCCTATGCCATTCAATATTTTAGGGAGAGC

CTGCTTGCCTACCCGAGATATAGTAACTTCATCTCGACTGAGTGTTACCTTATTATGATAGGATTAAAGGCCAATCGGTT

GATAAACCCAGAAGCCATTAAGCAAAGCATAATCAGAGTGGGGACTAGGACTGCACCAGGACTTGTGAGCCACATATTAT

CAGAGAAACAGAAAGGTTGTATTCAATCTTTTCTGGGTGATCCTTATATCCAAGGAGACTTCAATAAGCACCTTAAAGCT

CTAACCCCTATTGAGAAAATCCTAGTAAATTGTGGTCTCTCGATCAATGGCACAAAAATCTGTAGGGATCTAATCCACCA

TGATATCGCCTCCGGTCCAGACGGTCTGATGAGCTCCACAATTATTTTATATAGGGAACTGGCCCATTTCAAAGACAATA

TAAGAAGTCAGCACGGTATGTTCCACCCCTACCCAGTATTGGCCAGTAGCAGGCAACGTGAATTAATCCTTCGAATAGCC

AAGAAATTCTGGGGGTATGTCTTGCTATATTCTGATGACCCGGCACTAATCAAACAAACGATCAAGAACTTGAAGCGGAA

TCACCTAACCTTTGACTTACACAGTAATCCGTTTATTAAGGGCTTATCCAAAGCTGAGAAACTGCTAGTGCGGACAAGTT

CACTCAGAAGGGAATGGTTGTTCACTCTCGATACGAAAGAAGTGAAAGAGTGGTTCAAATTGGTGGGTTACAGTGCACTC

GTCAGAGGTTAATTCGCGATACATCTGCCCCCTTCTCCTCCTCCATGAGACTCTACTGGCAATCTAAAAGATTAAAGAAA

ACTACATATTGGATAAGTATCTATTCCCAGCTTTGTCTGGT

>EU267274.1/goat/Nigeria/1976

ACCAAACAAAGTTGGGTAAGGATAGATCTTATAATAACTATGGACTGGCAAACTTAGGAGTAAAGATCCTACTGTCGGGG

AGAGGAGGAGGAGCAAGATCTTTGACCATGGCGACTCTCCTTAAAAGCTTAGCATTGTTCAAGAGGAACAAAGACAAAGC

GCCTACTGCGTCGGGTTCAGGAGGGGCCATCCGGGGGATTAAGAATGTTATCATAGTCCCTATCCCCGGGGACTCATCCA

TCATTACCCGTTCAAGACTGCTCGACAGGCTTGTCAGATTGGCCGGAGATCCTGACATCAACGGGTCAAAGCTGACCGGC

GTGATGATCAGCATGTTATCTTTGTTCGTGGAGTCACCCGGGCAATTGATACAGCGGATCACAGATGATCCAGATGTCAG

CATCCGCCTTGTTGAGGTAGTTCAAAGTACTAGGTCCCAGTCCGGGTTGACCTTTGCATCACGTGGTGCTGACTTGGACA

ATGAGGCAGATATGTATTTTTCAACTGAAGGACCCTCGAGTGGATGTAAGAAAAGGATCAACTGGTTTGAGAACAGAGAA

ATAATAGACATAGAAGTGCAAGATGCAGAAGAGTTCAATATGTTGTTAGCCTCCATCTTAGCACAAGTTTGGATCCTCCT

GGCCAAGGCGGTTACGGCACCAGATACGGCAGCCGACTCAGAACCGAGAAGGTGGGTTAAATACACACAACAAAGGAGAG

TGATTGGGGAATTTCGCCTTGACAAAGGGTGGCTGGACGCAGTCCGCAACAGGATTGCAGAAGATCTATCACTCCGGCGG

TTCATGGTATCTCTCATACTTGACATCAAGAGGACCCCCGGTAACAAGCCAAGGATTGCAGAAATGATCTGCGACATTGA

CAACTATATTGTCGAAGCCGGACTCGCCAGTTTCATTCTTACTATCAAGTTTGGTATTGAAACCATGTATCCTGCACTGG

GCCTTCACGAGTTCGCCGGGGAATTGTCCACTATTGAATCCTTGATGAACTTGTATCAACAGCTAGGAGAGGTTGCACCC

TACATGGTAATTCTAGAGAACTCAATTCAGAACAAGTTTAGTGCAGGAGCCTATCCCCTCCTCTGGAGCTATGCGATGGG

TGTCGGAGTCGAGTTGGAGAACTCAATGGGGGGCTTGAACTTTGGCAGGTCATATTTTGACCCGGCCTATTTTCGTCTCG

GACAGGAGATGGTCAGAAGATCTGCAGGAAAGGTCAGCTCCGTAATCGCGGCTGAGCTTGGCATCACAGCAGAGGAAGCC

AAACTAGTCTCGGAAATCGCCTCGCAGACTGGGGAAGAACGAACCGTTAGAGGGACTGGGCCTCGTCAGGCGCAGGTTTC

CTTCCTCCAGCATAAAATGGATGAGGGAGAGTCGCCTACACCAGCGACCAAAGAAGAAGTCAAAGCTGCGATTCCAAATG

GGTCTGAAGGAAGGGACATAAAGCGAACACGCTCAGGGAAGCCCAGAGGAGAAACTCCCGGCCAACTGCTTCCGGAGATC

ATGCCAGAGGATGAAGTCTCGCGAGAGTCCAGTCAAAACCCTCGTGAGGCTCAAAGATCGGCTGAGGCACTCTTCAGGCT

GCAGGCCATGGCCAAGATTCTGGAGGACCAGGAGGAGGGAGAAGACAACAGTCAGATCCACAACGACAAGGATCTCCTCA

GTTGAGCAGATGCACCCTCCGTCTGAATCAGTGACAAGACATCACCTGCCAGTATTATAAAAAACTTAGGACCCAGGTCC

AAGCAACCGCACATCGACACCCCAGTCAATCGAGCGGAGACCACCGATGGCAGAAGAACAAGCATACCATGTCAACAAGG

GACTGGAATGTATCAAGTCTCTCAAAGCCTCTCCCCCGGATCTATCCACCATCAAAGATGCCCTTGAGAGCTGGAGAGAG

GGGCCTAGCCCCCCAGGCCGTGCAACACCGAACCCTGATACGCCCGAGGGAGACCATCAGAATATCAACCAATCATGCTC

ACCAGCAATCGGATCAGACAAAGTCGACATGTCTCCTGAAGATAATCTCGGATTTAGAGAGATCACTTGTACTGACAGTG

AGGCCGGGCTCGGAGGAGTTCAGGACAAACGATCCGACTCTCAAGTACAGCGTTACTATGTTTATAGCCACGGGGGTGAA

GAGATTGAAGGACTCGAGGATGCTGACTCTCTCGTGGTTCAAGCAAATCCTCCGGTTGCTAACACCTTCAATGGAGGAGA

GGATGGATCTGACAACAGCGATGTGGACTCTGGCCCAGATGATCCCGGCAGAGATCCTCTATATGACCGGGGACCTGCTG

CCGGCAATGATGTCTCTAGGTCAACAGATGTCGAAAAATTAGAAGGTGATGACATTCAAGAAGTTCTTAACTCCCAGAAA

AGTAAAGGAGGAAGATTCCAAGGCGGGAAAATCTTGCGAGTCCCGGAAATACCCGATGTCAAGAACTCCAGACCATCAGC

CCAATCAATTAAAAAGGGCACAGACGGGAGCTCAGCCTTATCTGGAACGGTGACAGAGTGTTCATCGATAAGTGGTGCAA

CCCAAGCTGTGCCAGAGTCCAGATGGGAGTCATCAGAGCGAAATGCGTCTGTGGGGAGTGTCCTCAAATCTGCGAGGAGT

GCAAAGACGATCCCGGGGTTAACACAAGAATCTGGTACCATAGCATCACTGACTCAGCCTAAAGAGAATGACTCCGAGTA

TGAGTATGAGGATGACCTATTCACTGAGATGCAGGACATTCGTGCAAGCATTGCTAAGATCCATGATGACAACAAAACTA

TCCTCTCTAGACTTGATTCTCTACTGTTATTGAAAGGAGAAATCGATACTATCAAGAAACAAATCAGCAAACAAAATATA

AGTATATCTACCATTGAGGGCCATCTATCCAGTATAATGATAGCCATCCCAGGCTTTGGGAAGGACATCAAGGACCCAAC

ATCTGAGGTTGAGTTGAACCCAGATTTGAGACCTATAATCAGCCGTGATTCTGGCAGGGCTCTTGCGGAGGTCCTCAAGA

AACCCGCTGTTGATAGGTCTCAGAAAAGCGGAATCAAAGTCAACTCCGGTTCAAAGGGTCAGCTCCTTAAGGATCTCCAG

CTAAAACCTGTCGACAAACAGGCAAGCTCTGCAATCGGGTTTGTTCCATCCGACCATGAATCATCCAGGAGTGTCATCCG

CTCCATAATCAAGTCGAGCAAGCTTAACATTGATCACAAGGACTATCTTCTAGATTTACTGAATGATGTGAAAGGCTCTA

AGGATCTTAAGGAATTCCACAAGATGCTAACAGCCATTCTTGCGAAGCAGCCGTAACACATCCCATAATCAACATCTCAT

ACTCGGTTGAAAACATCCCCTCAATCAGGCTATTACAAAAAACTTAGGAGCAAGGGCAACTGAGCTTCACAGACAAGATG

ACCGAGATCTACGATTTCGATAAATCAGCATGGGATGTCAAAGGGTCAATTGCTCGCATAGAACCCACCACCTATCACGA

CGGCCGACTGATACCCCAGGTGAGGGTCATCGACCCTGGTCTGGGAGACAGAAAAGATGAGTGCTTCATGTACCTGTTTC

TTCTAGGAGTGATTGAGGATAACGACCCCCTGTCTCCCCCAGTCGGGAGAACCTTCGGGTCTTTACCTCTAGGGGTCGGT

AGGTCAACTGCTAAGCCAGAGGAACTACTAAGGGAGGCCACAGAACTAGATATAGTGGTGAGGCGCACGGCAGGACTAAA

TGAGAAACTGGTATTTTACAACAACACTCCGCTGTCTTTGTTAACACCCTGGAGGAAAGTCTTGACAACCGGAAGTGTGT

TCAGCGCCAACCAGGTTTGCAATGCAGTCAACCTAGTCCCACTTGATACTCCCCAGAGATTCAGGGTTGTGTACATGAGT

ATAACTAGATTGTCAGATAATGGTTATTATAGTGTTCCCAGAAGAATGTTGGAGTTCCGCTCAGCCAATGCAGTCGCCTT

CAATATCTTGGTTACACTGAGAATTGAAAATGGCACAAACCCTAGGAGATACATAGTCGGCTCATGGGAGAATCCAGAAG

TCACATTTATGGTACACGTGGGCAACTTTAGAAGAAAGAAGAACGAAGTATACTCTGCTGATTATTGCAAAATGAAGATT

GAAAAGATGGGTCTAGTTTTTGCCTTGGGAGGAATAGGTGGAACAAGTCTCCATATTCGAAGCACAGGGAAAATGAGCAA

GACCCTCCATGCACAGCTGGGGTTCAAGAAAATCCTATGCTACCCCCTAATGGATATCAATGAGGATCTTAACCGATATC

TCTGGCGGGCAGAGTGCCGAATAGTTAAAATCCAGGCCGTCTTACAGCCATCAGTACCCCAAGAATTCCGTGTCTATGAT

GATGTCATCATCAACGATGATCAAGGCCTGTTCAAGATCCTGTAATTCACCTGCGGGGCCACGGACCCACCTCCAGGAAG

AGGCCCTCTCCTCTCCCGTCCAAAACACACACCCCAAGGCTCCCCGAAACAGCGCCGACACAGCCACTCTCCCGGCGGGA

CGAGGCCGACT-CCCCCAACAAACTCGCTCCGGGGGGGGG-CCCCCCCCACGACCCGCCGGGCGTTGACCGCGCGAGCCG

GCGAGACACCCCCAAGAGGCCCCGACGAAGACCGGGCAACCCTCCCCCCA-AAAAAAGCCCCCC-AACACACCCAAGGCC

AGGCGCCGGGCACCCCCAGCCCCGCCATCGGCGGGACAGGAGGCCAAGACGGGGTCCCCCCAACCAAAGGGACCACCAGG

GAAGGCTCTGCCGACCCAGCACAGACCCGGCCCAAACAAGGCAGACCCCCAAGACGAAATCTCCCAGAGCACCCCATTCA

TCATCAAACAAAACTTAGGGGCCAAGTTCAGACGACAGACCCCAGTCCGCCAGGCAGCAAGCCGCAGAAAGGAAGGAGAC

AAGACCGGCCCGGAAAGAGGACACCCAGCCACAGAAACCCCCAACTCCAGGGGAGGAG-CCCCCCCTCCACACGGCGGAG

GAACGGGCGAGAGACACCGGGGAGAACCCGATCGGGAGCCAGGAGCCGGAGACGCCAGCGGCAAACAGGCCGGGCACACA

ACCGACACCACCGGAGGATCAGAGAAGAGCCCGACCCCGCGTAGACCCGACGGCAGAGGACCCAACCCCGCCACTAAGAG

GGGCGAGAACCCT--------CCCCCCCGCACACAACCGGCAAGCAGACCGAGGGAGGACCACCTCCCAGCAAGACAGAC

GGCTCCCCCGCAGCCGAGCCGCAGCTCCCCAGAAGAGGACACCCCCGAAACCTCCCAGAACCCCTCCGCTCCCGGCCGCC

GGGACAGGAGCTCCGCCGCCAAAGACTAGAACCACCGAGAGTCAACCAGGGGTCCCCCCCAGTCCCTCCCGGCGTTCTCC

CGCAGAGGCCCCAGCAAGGTCCCCCGGCCCGACGACCCCTGCATGCGCCGAAAGAGAAACCCACCAAAGGACGTCCAGCA

CATCCATGTATAAACATCATGACACGGGTCGCAACCTTGGTATTTCTGTTTCTTTTCCCAAACACTGTCGCGTGCCAGAT

TCACTGGGGCAATCTATCCAAGATCGGGATTGTAGGAACAGGGAGTGCCAGTTACAAGGTGATGACTAGGCCAAGCCACC

AAACCCTGGTCATAAAGTTGATGCCAAATATAACAGCCATTGACAATTGTACGAAATCAGAGATTTCAGAGTACAAAAGA

TTGCTGATCACAGTGTTAAAGCCTGTAGAGGATGCTCTGTCAGTGATAACCAAGAATGTAAGACCAATTCAAACTCTAAC

ACCTGGGCGCAGGACCCGCCGTTTTGCCGGAGCTGTTCTGGCCGGAGTAGCACTTGGAGTCGCGACGGCTGCTCAAATAA

CTGCCGGAGTCGCACTTCATCAGTCATTGATGAATTCCCAAGCAATTGAAAGTTTAAAAACCAGTCTTGAGAAGTCGAAT

CAGGCAATAGAAGAAATCAGACTTGCAAATAAGGAGACCATACTGGCAGTACAGGGCGTCCAAGATTATATCAACAATGA

GCTTGTCCCCTCTGTTCATAGAATGTCATGTGAGCTGGTAGGTCACAAACTCAGTCTCAAGCTCCTTAGGTATTATACCG

AGATCCTGTCTATATTCGGGCCTAGCCTTCGAGACCCGATAGCTGCTGAAATATCAATCCAGGCACTCAGCTATGCACTA

GGCGGAGACATCAATAAAATCCTGGACAAGCTTGGGTATAGCGGCGGGGATTTCCTTGCTATTCTAGAAAGCAAGGGGAT

AAAGGCCCGGGTGACATATGTGGACACAAGAGATTACTTTATAATTCTTAGCATAGCCTACCCAACCTTATCTGAGATCA

AGGGGGTGATAGTTCATAAGATAGAAGCTATATCATACAACATTGGGGCACAGGAATGGTATACTACTATCCCTAAATAT

GTAGCCACTCAGGGATATCTGATATCGAATTTCGATGAGACGTCATGCGTCTTCACTCCAGAGGGGACAGTCTGCAGCCA

GAATGCGTTGTATCCAATGAGCCCATTGCTTCAGGAATGTTTCAGGGGGTCGACAAAATCGTGCGCCAGAACCCTAGTTT

CAGGGACCACAAGTAATAGATTTATCCTATCAAAAGGGAACTTGATTGCAAATTGTGCGTCAGTTTTGTGCAAGTGTTAC

ACAACGGAGACAGTCATCAACCAAGATCCTGATAAACTACTAACTGTTATAGCCTCCGATAAGTGTCCCGTAGTTGAGGT

GGATGGAGTGACAATACAGGTCGGCAGTCGAGAGTACCCAGATTCTGTATACCTACATAAAATAGACTTAGGCCCAGCCA

TCTCTCTGGAAAAACTGGATGTAGGCACCAATTTAGGCAATGCAGTCACAAGACTGGAGAATGCAAAGGAGCTCCTAGAT

GCATCAGACCAGATACTGAAGACTGTTAAAGGGGTACCCTTCAGTGGCAATATGTACATAGCACTGGCAGCTTGCATTGG

GGTATCCCTAGGGCTTGTCACATTAATATGCTGCTGTAAGGGGAGATGTAAGAACAAGGAGATTCCTGCCTCCAAAATCA

ATCCAGGGCTCAAACCCGACCTGACCGGGACTTCAAAGTCGTACGTGAGATCACTGTAGTCAGAGTAACCCGAATCATCC

AGCATCACACGTATACATGTGCGACACAAGCAGTCAGAGGACACAGAAGACCCAACTTCCGATCACCGATCAGACCCTTC

TCTACGCCCTACTACACATTGGTCATCAAACAAAACTTAGGACGAAAGGTCAATCACCATGTCCGCTCAAAGGGAGAGGA

TCAATGCCTTCTACAAAGACAATCCTCACAATAAGAACCATAGGATAATCCTGGATAGGGAACGCTTAACTATTGAAAGA

CCCTACATCTTACTTGGAGTCCTGCTGGTAATGTTCCTGAGTCTAATCGGGCTGCTGGCCATTGCAGGGATCAGGCTTCA

CCGGGCCACCGTTGGAACTGCGGAGATCCAGAGTCGGCTGAAGACCAACATTGAGTTGACCGAATCCATTGATCATCAAA

CTAAGGATGTCTTAACTCCCCTGTTTAAAATCATTGGTGATGAAGTCGGCATCAGAATTCCACAGAAGTTCAGTGATCTT

GTCAAGTTCATCTCCGATAAGATTAAGTTCCTCAACCCTGACAGGGAATACGATTTTAGGGATCTCCGGTGGTGTATGAA

CCCCCCTGAGAGAGTCAAAATTAACTTTGATCAGTTTTGTGAATACAAAGCCGCGGTCAAATCAGTTGAACATATATTTG

AGTCATCACTCAACAGGTCAGAAATGTTGCGACTATTGACTCTTGGGCCCGGAACAGGCTGTCTCGGCAGGACAGTAACA

AGAGCTCAGTTCTCAGAGCTCACGCTGACCTTGATGGACCTGGATCTCGAGATGAAGCACAACGTGTCCTCAGTGTTTAC

CGTAGTCGAAGAGGGATTATTCGGAAGAACATATACTGTCTGGAGATCCGACACCGGGAAACCGAGCACCAGTCCAGGTA

TTGGCCATTTTTTAAGAGTCTTCGAGATCGGGCTGGTGAGAGATCTCGAGCTGGGTGCCCCTATTTTCCATATGACCAAC

TACCTCACAGTGAACATGAGTGATGACTATCGGAGCTGTCTTTTAGCGGTAGGGGAGTTGAAGCTGACAGCCCTATGCAC

CCCATCTGAGACTGTGACTCTGAGTGAGAGAGGAATTCCAAAGAGAGAGCCTCTTGTGGTTGTGATACTCAACCTGGCTG

GGCCTACTCTAGGGGGCGAACTATACAGTGTCTTGCCTACCTCTGACCTCATGGTGGAGAAACTCTATTTATCCTCACAT

AGAGGGATTATCAAAGATAACGAGGCCAATTGGGTAGTACCGTCTACCGATGTTCGTGATCTTCAAAACAAAGGAGAATG

TCTGGTGGAAGCATGCAAGACTCGACCTCCTTCATTTTGCAATGGCACAGGAATAGGCCCATGGTCAGAGGGGAGAATCC

CTGCCTACGGAGTGATCAGGGTCAGTCTTGACTTAGCTAGTGACCCAGGTGTGGTTATCACTTCAGTGTTTGGCCCATTG

ATACCTCACCTATCCGGCATGGATCTTTACAACAATCCGTTTTCAAGAGCTGCATGGTTAGCTGTACCACCTTATGAGCA

GTCATTTCTAGGAATGATAAATACAATTGGCTTCCCGGACAGAGTAGAGGTTATGCCGCACATTTTGACCACAGAGATCA

GAGGGCCTCGAGGTCGTTGTCATATTCCTATAGAGTTGTCCCGCAGGATTGATGATGATATCAAGATCGGGTCCAACATG

GTTGTATTGCCGACGAAGGACCTGAGGTACATAACAGCCACTTATGATGTTTCCAGGAGCGAGCATGCAATCGTGTACTA

TATCTATGACACGGGTCGCTCATCATCTTACTTCTACCCAGTTCGATTGAATTTCAGGGGCAATCCTCTCTCTCTGAGGA

TAGAGTGTTTTCCCTGGTATCATAAGGTGTGGTGCTACCATGATTGCCTTATATACAACACCATAACAAACGAAGAAGTC

CATATGAGAGGGCTGACCGGTATAGAGGTAACATGTAATCCAGTCTGAGTAGAGCTACAACCATCGCTCAAGCAGGCCTC

CGAGCCATCCCCTAGTCCAAGAAGCATAGCCTGGGACACTCAGCAGCACAACCCAGCCAACAATGTTATAAAAAACTTAG

GAGCCAAGGTTGTAGGAGCCATGGACTCACTATCAGTCAATCAGGTCTTGTACCCTGAGGTCCATCTAGATAGCCCTATT

GTCACAAACAAACTAGTTGCCATCCTTGAATACTCGGGGATCGACCACAACTATGTTCTTGAAGACCAGACCCTTATCAA

GAATATTAGATATAGACTGGGGTGCGGTTTTTCAAATCAAATGATCATCAATAATAGGGGGGTAGGCGAAACAGTCAATT

CCAAACTTAAAAGTTACCCCCGTAATTGTCATATCATATACCCAGACTGCAATAAGGATTTGTTTTGTATCAAAGATAGC

TGCATATCTAGAAAGCTCTCGGAGCTATTCAAGAAGGGTAATTCCTTGTACTCTAAGATAAGTCACCAGGTACTGGATTG

TCTTAAGAGAGTCAACGGGAAATTAGGCCTGGGCACAGATCTTACTCACGGCCTGAAGGAGGGTATCATCGACTTGGGGT

TGCACATGCATAGCTCTCAATGGTTCGAGACCTTTCTGTTCTGGTTCACTATCAAGACAGAGATGAGATCAATGATTAAA

GAACAGTCCCATATATGCCACAAGAGGAGGTATAACCCAATTTTTGTGTCGGGAGATGCATTCGAGGTGCTCGTATCACG

AGACCTCGTTGTGATAATTGATAAGAACACCCAGTATGTCTTCTACCTGACATTTGAGCTGGTCCTTATGTATTGTGATG

TCATAGAGGGCAGACTTATGACGGAGACAGCCATGGCTATAGACCAGAGATATTCAGAGCTTCTAAACCGGGTCAGATAC

TTGTGGGATCTCATTGACGGGTTCTTCCCAACACTGGGTAACACCACATACCAAGTTGTTGCTCTGCTTGAACCACTGTC

GTTAGCTTATCTTCAACTTCAGGATGTCACTCTAGAGTTAAGAGGTGCTTTTTTGGACCACTGCTTCAAAGAACTTTATG

AGATACTGGAGCATTGTGGCATTGACACGGAAGGTACCTACAATTCCATCACTGAAGGATTGGATTACGTATTTATCACC

CACGATATACACTTAACTGGGGAGATTTTTTCATTTTTCCGGAGTTTCGGACACCCCCGCCTCGAAGCGGTCACCGCTGC

AGAGAATGTCAGAAAACATATGAACCAACCGAAGGTAATCAGTTATGAGACTATGATGAAAGGGCATGCAGTATTTTGCG

GGATAATCATAAATGGTTTTAGGGACCGGCACGGCGGCAGCTGGCCCCCTGTTGCATTGCCAGAACATGCTTCTGCTGCG

ATCCGGAATGCGCAGGCATCCGGTGAAGGACTGACCCATGACCTGTGTATAGACAACTGGAAGTCCTTTGTTGGATTCAA

ATTTGGCTGCTTCATGCCGCTCAGCCTGGATAGTGATTTGACCATGTACCTCAAAGACAAAGCACTGGCTGCACTGAAGA

ATGAGTGGGATTCAGTTTACCCGAAAGAATACCTCCGTTATAATCCACCTAGAGGGACAGAGTCAAGGCGACTGGTGGAG

GTGTTCCTGAATGACTCCAGCTTTGATCCTTATAACATGATAATGTACGTGGTGAATGGCTCCTACCTTAAAGACCCTGA

GTTTAACCTCTCATACAGCTTAAAAGAGAAGGAGATAAAAGAGACAGGGCGGTTGTTTGCCAAAATGACCTACAAGATGC

GGGCCTGTCAGGTAATTGCTGAAAATCTAATATCAAATGGTGTCGGGAAGTATTTCCGAGACAATGGGATGGCAAAAGAC

GAGCATGACCTAACAAAGGCCCTTCACACTCTGGCAGTCTCAGGTGTTCCCAAGAACAACAAAGATAACCACCGAGGTGG

GCCTCCCAGAAGGACCGCAAGCCGAGAGATGAGATCAAGCCAAGCCATCAACACACAAAATAGAGACAGGATCCAAGGGG

GCCCTATGTACAACTACTTGCGATGCCAACCGACCAGCTCTGATCAGGGTGAGTCATACGAGACTGTTAGTGCATTCATC

ACCGCTGACCTTAAGAAATATTGCCTAAATTGGAGATACGAGACAATCAGCATATTTGCACAGAGACTGAATGAAATATA

TGGGTTGCCATCCTTCTTTCAATGGTTACACAGGGTATTGGAAAAATCCGTGCTCTACGTCAGTGACCCGCATTGCCCTC

CCGACTTAGATGATCATATCCCCCTGGACAGTGTCCCTAATGCCCAAATATTCATCAAGTACCCAATGGGCGGAATAGAA

GGTTATTGTCAAAAACTATGGACAATCAGTACTATACCATACCTGTATCTGGCAGCCTATGAGAGCGGAGTAAGAATTGC

CTCACTAGTGCAAGGTGACAATCAGACAATTGCAGTGACAAAAAGAGTTCCAAGTTCTTGGCCTTATTCACTAAAAAAGA

GGGAGGCATCTAAAGCAGCTCAAAATTACTTCGTGGTCTTAAGGCAAAGGTTGCACGATGTAGGTCATCACTTAAAGGCT

AATGAGACCATAGTATCTTCTCACTTTTTTGTATACTCTAAAGGGATTTATTATGACGGCCTGTTAGTCTCACAATCACT

AAAGAGCATCGCCAGATGTGTCTTCTGGTCCGAGACTATTGTAGATGAAACCAGAGCGGCCTGCAGCAATATTGCAACAA

CTATCGCCAAGAGCATAGAGAGGGGTTATGATAGGTACCTCGCATACTCTTTGAATATCCTCAAAATTTTCCAACAGATC

CTTATATCCCTTGACTTCACGATTAACACAACTATGACTCAAGATGTCGTGGCACCGATCATCGAGAACGGTGATTTACT

AATAAGGATGGCACTCTTGCCAGCACCCATTGGGGGTCTCAATTATCTTAACATGAGCAGGTTGTTTGTGAGAAATATCG

GTGACCCGGTCACTTCCTCCATAGCCGACCTGAAGAGGATGATAGACGCTGGGCTAATGCCAGAAGAAACATTGCATCAA

GTGATGACCCAGACCCCGGGAGAATCATCTTACCTCGATTGGGCAAGTGACCCTTATTCTGCCAACCTAACCTGCGTACA

GAGTATAACTCGCCTTCTCAAGAACATCACTGCACGGTACATCTTAATCAGCAGCCCAAATCCGATGCTGAAAGGATTGT

TTCATGAGGGGAGTAGAGATGAAGACGAGGAGCTTGCGAGTTTCTTGATGGATCGGCATATAATTGTTCCGAGAGCCGCA

CATGAAATCTTAGACCACAGCATCACCGGAGCAAGAGAAGCTATAGCCGGGATGTTGGACACCACCAAAGGTCTGATTAG

AACAAGTATGAAACGGGGTGGCCTCACCCCTCGAGTATTAGCCCGCCTTTCCAATTATGATTATGAACAATTCAGATCCG

GAATAACACTATTGACAAAGAAAGGGCAATGTTATCTCATTGACAAGGACTCGTGCTCGGTGCAGCTCGCTATAGCTCTG

AGGGGCCACATGTGGGCCAGGTTAGCTCGCGGGAGACCTATCTATGGGTTGGAGGTGCCTGATGTACTGGAATCGATGAA

CGGCTACCTTATCAAACGCCACGAGTCTTGTGCCATCTGCGAAACGGGCTCAAGTCACTACGGGTGGTTTTTCGTCCCTG

CAGGGTGCCAGCTTGACGATGTCTCAAGAGAGACTTCGGCTCTTCGTGTACCTTATGTCGGATCAACCACTGAGGAAAGG

ACAGATATGAAACTTGCTTTCGTTAGATCTCCAAGCCGATCCCTCAAATCAGCAGTCAGGATTGCCACAGTTTACTCATG

GGCCTACGGGGATGATGAGAAATCATGGAGTGAAGCTTGGATGCTAGCTAGGCAGAGAGCTGATATCACCTTAGATGAAC

TGAGAATGATCACTCCAGTCTCTACATCCACCAACCTAGCCCATCGGTTGAGGGATCGGAGCACCCAGGTGAAATATTCG

GGGACATCCCTTGTGAGGGTTGCAAGATACACAACCATCTCCAATGACAATTTGTCATTTGTGATATCTGAGAAAAAAGT

AGATACCAACTTCATTTACCAGCAAGGGATGCTGCTCGGTCTTGGGATCCTTGAGAATCTCTTCAGGTTAGAGGCCACCA

CAGGGGTATCCAACACAGTGCTACACCTGCACGTGGAAACAGAATGTTGTGTTGTACCTATGGTTGATCACCCAAGGATA

CCGAGTCTCCGTAATATTAAAGTTACGAATGAGCTATGCACAAACCCTCTGATCTACGACAGGTCCCCCATCATAGAACA

CGATGCAACTCGATTATACTCACAAAGCCACAGGAGACATTTGGTGGAGTTTGTTACCTGGTCAACAAGCCAGCTTTATC

ATATACTGGCCAAATCTACAGCAATGTCCATGATTGAGTTGATCACAAGATTTGAGAAAGATCACATGAATGAAATAGCC

GCCCTGATTGGCGATGACGACATCAACAGTTTCATCACAGAATTTTTGCTCGTGGAGCCCAGACTGTTTATAGTCTACCT

TGGCCAGTGTGCTGCCATCAATTGGGCTTTTGATATACATTATCATCGGCCCTCGGGCAAGTACCAGATGGGGGAACTCC

TCTACTCTTTACTCTCTCGGATGAGCAAAGGAGTATATAAGATCTTCACTAATGCTTTGAGCCACCCCAAAGTTTACAAG

AAATTTTGGCGAAGTGGTGTAATTGAGCCGATTCATGGCCCATCCCTAGATACACAGAATTTACATGTCACTGTCTGTGA

CATGATATACGGATCATACGTCACCTATCTGGATCTTTTGCTGAATGATGAGCTAGATGCTTACCCGTACTTGCTCTGCG

AGAGTGATGAGGACGTGGTCACAGACAGGTTCGACAACATTCAAGCCAAACATCTCTGTGTATTGGCCGATGTATACTGC

AGCTCCAAGAGATGTCCCTCGATAATCGGGATGTCTCCTATAGAAAAATGTACCATCCTCACACATTACATCAAGGGAGA

ATCGGTACAATCCCCGTCCGGGATCTCATGGAACACTGATCCCCTTGTAGTAGATCATTACTCATGCTCTCTGACCTACC

TTCGCCGCGGTTCCATCAAACAAATCAGGTTGAGAGTGGATCCTGGGTTTGTATTCGAGGCGTTGACAGACATCGACTTC

AAACAGCCTCGCAAGGCTAAGTTGGATGCATCGATTGTGGGGTTGACTGATTTTTCTCCCCCTTGGGATAACGTCGGTGA

TTTTCTAGGGACTATCAACACATTGAGGCACAATCTGCCCGTCACCGGAACCGGGGTCTCGAACTATGAAGTCCACGCTT

ATCGTAGAATTGGTCTGAATTCATCAGCATGTTACAAAGCTGTAGAGATCTCCACGTTAATCAAGTCATCCTTAGAAGTC

GGAGAGGATGGATTGTTCTTAGGAGAAGGTTCCGGTTCTATGCTGGCTGCGTACAAGGAAGTTCTTAAATTGGCAAACTG

TTATTACAACAGCGGGGTAACAGCGGAGGGTAGAGCCGGACAGAGGGAAATCTCTCCCTATCCCTCAGAGGTGAGCCTGG

TAGAGAGTCAGATGGGGGTAGAGAGAAGTGTTAAAGTGCTGTTCAATGGCAAACCTGAAGTAACCTGGGTAGGGACCACC

GATTGCTACAAGTATATAATCAGTAACATTCAGACCTCTAGTCTGGGTTTCATACACTCAGATATTGAGACACTCCCAAC

CAAGGATGCCGTTGAGAAGTTAGAAGAATTTGCCTCTATCCTATCCCTATCCCTAATTTTGGGAAAAATCGGCTCTATTA

CAGTTGTCAAAATTATGCCCATTAGCGGAGATTTTACCCAAGGCTTCATAGCCTATGCCATTCAATATTTTAGGGAGAGC

CTGCTTGCCTATCCGAGATATAGTAACTTCATCTCGACTGAGTGTTACCTTATTATGATAGGATTAAAGGCCAATCGGTT

GATAAACCCAGAAGCCATTAAGCAAAGCATAATCAGAGTGGGGACTAGGACTGCACCAGGGCTTGTGAGCCACATATTAT

CAGAGAAACAGAAAGGTTGTATTCAATCTTTTCTGGGTGATCCTTATATCCAAGGAGACTTCAATAAGCACCTTAAAGCT

CTAACTCCTATTGAGAAAATCCTAGTAAATTGTGGTCTCTCGATCAATGGCACAAAAATCTGTAGGGATCTAATCCACCA

TGATATCGCCTCCGGTCCAGACGGTCTGATGAGCTCCACAATTATTTTATATAGGGAACTGGCCCATTTCAAAGACAATA

TAAGAAGTCAGCACGGTATGTTCCACCCCTACCCAGTATTGGCCAGTAGCAGGCAACGTGAATTAATCCTTCGAATAGCC

AAGAAATTCTGGGGGTATGTCTTGCTATATTCTGATGACCCGGCACTAATCAAACAAACGATCAAGAACTTGAAGCGGAA

TCACCTAACCTTTGACTTACACAGTAATCCGTTTATTAAGGGCTTATCCAAAGCTGAGAAACTGCTAGTGCGGACAAGTT

CACTCAGAAGGGAATGGTTGTTCACTCTCGATACGAAAGAAGTGAAAGAGTGGTTCAAATTGGTGGGTTACAGTGCACTC

GTCAGAGGTTAATTCGCGATACATCTGCCCCCTTCTCCTCCACCATGAGACTCTACTGGCAATCTAAAAGATTAAAGAAA

ACTACATATTGGATAAGTTTCTATTCCCAGCTTTGTCTGGT

>MF741712.1/goat/SierraLeone/2011-12-17

ACCAAACAAAGTTGGGTAAGGATAGATCTTACAATGACTATAGACTAGCAAACTTAGGAGTAAAGATCCTACTGTCGGGG

AGAGGAAGAGGAGCAGGATCTTTGACCATGGCGACTCTCCTTAAAAGCTTAGCACTGTTCAAGAGGAACAAAGACAAAGC

GCCTACGGCGTCAGGTTCAGGAGGGGCCATCCGGGGGATTAAGAATGTTATCATAGTCCCTATCCCCGGGGACTCATCCA

TCATTACCCGTTCAAGACTGCTCGACAGGCTTGTCAGACTGGCCGGAGATCCTGACATCAACGGGTCAAAGCTGACTGGC

GTGATGATCAGCATGTTATCTTTGTTCGTGGAGTCACCCGGGCAATTGATACAGCGGATCACAGATGATCCAGATGTTAG

CATCCGTCTTGTTGAGGTAGTTCAAAGTACCAGGTCCCAGTCCGGATTGACCTTTGCATCACGTGGTGCTGATTTGGACA

ATGAGGCAGATATGTATTTTTCAACTGAAGGACCCTCGAGTGGAAGTAAGAAAAGGATCAACTGGTTTGAGAACAGAGAA

ATAATAGACATAGAAGTGCAAGATGCAGAAGAGTTCAATATGTTGTTAGCCTCCATCTTAGCACAAGTTTGGATCCTCCT

GGCCAAGGCGGTTACGGCACCAGATACAGCAGCCGACTCAGAACTGAGAAGGTGGGTTAAATACACACAACAAAGGAGAG

TGATTGGGGAATTTCGCTTTGACAAAGGGTGGCTGGACGCAGTCCGCAACAGGATTGCAGAAGATCTATCACTCCGGCGG

TTCATGGTATCCCTGATACTTGACATCAAGAGGACCCCCGGCAACAAGCCAAGGATTGCAGAAATGATCTGCGACATTGA

CAACTATATTGTCGAGGCCGGACTCGCAAGTTTCATTCTTACTATCAAGTTTGGTATTGAAACCATGTATCCTGCACTAG

GCCTTCACGAGTTCGCTGGGGAATTGTCCACTATTGAATCCTTGATGAACTTGTATCAACAGCTAGGAGAAGTTGCACCC

TATATGGTAATTCTAGAGAACTCAATTCAGAACAAGTTTAGTGCAGGAGCCTATCCTCTCCTCTGGAGCTATGCGATGGG

TGTCGGAGTCGAGCTGGAGAACTCAATGGGGGGCTTGAACTTTGGCAGGTCATATTTTGACCCGGCCTATTTTCGTCTCG

GACAGGAGATGGTCAGAAGATCTGCAGGAAAGGTCAGCTCTGTAATTGCGGCTGAGCTTGGCATTACAGCAGAGGAAGCC

AAACTAGTCTCGGAAATCGCCTCGCAGACTGGGGAAGAACGAACCGTTAGAGGGACTGGGCCTCGACAGGCGCAGGTTTC

CTTCCTCCAGCACAAAATAGGTGAGGGAGAGTCGCCTACACCAGCGACCAAAGAAGAAGTCAAAGCTGTGATCCCAAACG

GATCTGAAGGAAGAGACATAAAACGAACACGCCCAGGGAAGCCCAGAGGAGAAACTCCCGGGCAACTGCTTCTGGAGATC

ATGCCAGAGGATGGAGTCTCGCGAGAGTCTGGTCAAAACCCTCGTGAGGCTCAAAGATCGGCTGAGGCACTCTTCAGGCT

GCAGGCCATGGCCAAAATTCTGGAGGACCAGGAGGAGGGAGAAGACAACAGTCAGATCTACAACGACAAGGATCTCCTCA

GCTGAGCAGATACACCCTCTGTCCAAATCAGTGACAAGACATCACCTGCCAGTATTATAAAAAACTTAGGACCCAGGTCC

AACCAACCGCACATCGACACCCCAGCCAATCGAGCGGAGACCACCGATGGCAGAAGAACAAGCATACCATGTCAACAAGG

GGCTGGAATGTATCAAGTCCCTCAAAGCCTCTCCCCCGGATCTATCCACCATCCAAGATGCCCTTGAGAGCTGGAGAGAG

GGGTTCAACCCCTCAGGCCGTGCAACACCGAACCATGATATGTCCGAGGGGGACCATCAGAATATCAACCAATCATGCTC

ATCAGCAATCGGATCAGACAAAGTCGACATGTCTCCTGAAGGTAATCTCGGATTTAGAGAGATCACTTGTGATGACAATG

AGGCTGGGCTCGGAGGAGTTCAGGACAAAAGATCCGACTCTCAAGTACAGCGTTACTATGTTTATAACCACGGGGGTGAA

GAGATTGAAGGACTCGAGGATGCTGACTCTCTCGTGGTTCAAGCAAATCCTCCGGTTGCTAACACCTTCGATGGAGGAGA

GGATGGATCTGACAACAGCGATGTGGACTCTGGCCCGGATGATCCCGGCAGAGATCCTCTATATGACCGGGGACCTGCTG

CCGGCAATGATGTCTCTAGGTCAACAGATGTCGAAAAATTAGAAGGTGATGACATTCAAGAAGTTCTTAACTCCCAGAAG

AGTAAAGGAGGAAGATTCCAGGGCGGGAAAATCTTGCGAGTCCCGGAAATACCCGATGTCAAGAACTCCGGACCGTCGGC

CCAATCAATTAAAAAGGGCACAGACGGGAGCTCAGTCTTATCTGGAATGGTGACAGCGTGTTCATCGATAAGTGGTGCAA

CCCAAGCTGTGCTAGAGTCCAGATGGGAGTCATCAGAGCGAAATGCATCTGTGGGGAGTGTCCCCAAGTCTGTGAGGAGT

GCAAAGACGATCCAGGGGTTAACACAAGAATCTGGTACCATAGCATCACTGACTCAGCCTAAAGAAAATGACTCCGAGTA

TGAGTATGAGGATGATCTATTCACTGAGATGCAGGACATTCGTGCAAGCATTGCTAAGATCCATGATGACAACAAAACTG

TCCTCTCAAAACTTGATTCTCTACTGTTATTGAAAGGAGAAATCGATACTATCAAGAAACAAATCAGCAAACAAAATATA

AGTATATCTACCATTGAGGGCCATCTATCCAGTATAATGATAGCCATCCCAGGTTTTGGGAAGGACATCAAGGACCCAAC

ATCTGAGGTTGAGCTGAACCCAGATTTGAGACCTATAATCAGCCGCGATTCTGGCAGGGCTCTTGCGGAGGTCCTTAAGA

AACCCGCTGTTGATAGGTCTCAGAAGAGCGGAATCAAAGTCAACTCCGGTTCAAAGGGTCAGCTCCTCAAGGATCTCCAG

CTAAAACCCGTCGACAAACAAGCAAGCTCTGCAATGGGGTTTGTTCCATCCGACCATGAATCATCCAGAAGTGTCATCCG

CTCCATAATCAAGTCGAGCAAGCTTAACATTGATCACAAAGACTATCTTCTAGATTTACTGAATGATGTGAAAGGCTCTA

AGGATCTCAAGGAATTCCACAAGATGCTAACAGCCATTCTTGCCAAGCAGCCGTAACACATCCCATAATTTACATCTCAT

ACTCGGTTGAAAGCATCCTCTCAATCAGGCTATTACAAAAAACTTAGGAGCAAGGGCAACTGAGCTTCACAGACAGGATG

ACCGAGATTTACGATTTCGATAAATCAGCATGGGATGTCAAAGGGTCAATTGCTCGCATAGAACCCACCACCTATCACGA

CGGCCGACTGGTACCCCAGGTGAGGGTCATCGACCCTGGTCTGGGAGACAGAAAAGATGAGTGCTTCATGTACCTGTTCC

TTCTAGGAGTGATTGAGGATAACGACCCCCTTTCTCCCCCAGTCGGGAGAACCTTCGGGTCTTTACCTCTAGGTGTCGGC

AGGTCAACTGCTAAGCCGGAGGAACTACTAAGGGAGGCTACAGAACTAGATATAGTGGTGAGGCGCACGGCAGGACTAAA

TGAGAAACTGGTGTTTTACAACAACACTCCGCTATCTTTGTTAACACCCTGGAGGAAAGTCTTGACGACCGGAAGTGTGT

TCAGCGCCAACCAGGTTTGCAATGCGGTCAACCTAGTCCCACTCGATACTCCCCAGAGATTCAGGGTTGTGTACATGAGT

ATAACTAGATTGTCAGATAATGGTTATTATAGTGTCCCCAGAAGAATGTTGGAGTTCCGCTCAGCCAATGCAGTCGCTTT

CAATATCTTGGTTACACTGAGAATTGAAAATGGCACAAACCCTAGAAGATACATAGTCGGCTCATGGGGAAATCCAGAAG

TCACATTTATGGTACACGTGGGCAACTTTAGAAGAAAGAAGAACGAAGTATACTCTGCTGATTATTGCAAAATGAAGATT

GAAAAGATGGGTCTAGTTTTTGCCTTGGGAGGAATAGGTGGAACAAGTCTCCATATTCGAAGCACAGGGAAAATGAGCAA

GACCCTCCATGCACAGCTGGGGTTCAAGAAAATTCTATGCTACCCTCTGATGGATATCAATGAGGATCTTAACCGATATC

TCTGGCGGGCAGAGTGCCGTATAGTCAAAATCCAGGCCGTCTTACAGCCATCAGTACCCCAAGAATTCCGTGTCTATGAT

GATGTCATCATCAACGATGACCAAGGCCTGTTCAAGATCCTGTAAGTCACCTGCAGGGCCATGGACCCACCTCCAGGAAG

AGGCCCTCCCACCTCCCGCCCAAAACACACACCCCAAGACTCCCCGAAACAGCGCCGGCAAAACCACTCTCCCGGCGGAA

CGAGGCCGACC-CCCCCAACAAACCCGCTCCGGAAGGGGG-CCCCACCCGCGACCCACCGGGCGCCGACCGCGCGAGCCG

GCGAGACACCCCCAAGAGGTCTCGACGAAGACCGGGGAACCCTTTCCCCA-AAAATGGCCCCCC-AACACACCCAAGGCC

ATGCGCTGAGCACCCCCAGCCCCCCCACTGGCGGGACAGGAGGCCAGGACGGGGTTCCCCAGAACGGGGGGACCACCAGG

GAGGACTCCGCCGACCCAGCACGGACCCGGCCCAAACAAAGCAGACCCCCAAGACGAAATCTCCCAGAGCACCCTACTCA

TCATCAAACAAAACTTAGGGGCCAAGTCCAGACGACAGACCCCGGCCCGCCAGGCAGCAAGCCACAGAAAAGAAGGAGAC

AAGATCGGCCCGGGAAGAGGACACCCAACCACAGAAACCCCCAGCCCCAGGGGAAGAG-CCCCCCCTCCACACAACGGAG

GAACGGGCGAGAGACACCGAGGAGAACACGATCGGGAGCCGAGAGCCGGAGACGCCGGCGGCAAACAGGCCAGGCACTCA

ACCGACACCACCGGAGGACCCGAGGAGAGCCCGACCCCGCGCAGACTCGACGGCAGAGGACCCAATCCCGCCACAAAGAG

AGGTGAGAACCCT--------CCCTCCTGCACACAACCGGCAAACAGACCGCGGGAGGACCACCTCCCAGCAAGGCTGAC

GTCTCCCCCGCAGCCGAGCCGCAGCTCCTCAGAAGAGGGCACCCCCGAAACCTCCCAGAATCCCTCCGCTCACGGCCGCC

GGGACAAGGGCTCCGCCGCCAAAGACCAGAGCCACCGAGAGTCAACCAGGGATCCCCCCCAGTCCCTCCCGGCGATCCCC

CACAGAGGCCCCAGCAAGGTCCCCCGGCCCGACGACCCCTGCATGCGCCGAAAGGGAAACCAACCAGAGGACGTCCAGCA

CATCCGTGTACAGACATCATGACACGGGTCGCAACCTTGGTATTGCTGGTTCTTTTCCCAAACACTGTCGCGTGCCAAAT

TCACTGGGGCAATCTATCCAAGATAGGGATTGTCGGAACAGGGAGTGCCAGTTACAAGGTGATGACCAGACCAAGCCACC

AAACTCTGGTTATAAAGTTGATGCCAAATATAACAGCCATCGACAATTGTACAAAATCAGAGATTTCAGAGTACAAAAGA

TTGCTAATCACAGTGTTAAAGCCTGTAGAGGATGCTCTGTCAGTAATAACCAAGAATGTAAGACCAATTCAAGCTCTAAC

ACCTGGGCGCAGGACCCGCCGTTTCGCCGGAGCGGTTCTGGCCGGAGTAGCACTTGGAGTCGCGACGGCTGCTCAAATAA

CTGCCGGAGTCGCACTCCATCAGTCATTGATGAATTCCCAAGCAATTGAAAGTTTAAAAACCAGTCTTGAGAAATCAAAT

CAGGCAATAGAAGAAATCAGACTTGCAAATAAGGAGACCATACTGGCAGTACAGGGCGTCCAAGATTATATCAACAGCGA

GCTTGTCCCCTCTGTTCATAGAATGTCATGTGAACTAATAGGTCACAAACTCAGCCTCAAACTCCTTAGGTATTATACCG

AGATCCTGTCTATATTCGGGCCTAGCCTTCGAGACCCGATAGCTGCTGAAATATCAATCCAGGCACTCAGCTATGCACTA

GGCGGAGACATCAATAAAATTCTGGACAAGCTTGGGTATAGCGGCGGGGATTTCCTTGCTATTCTAGAAAGCAAGGGGAT

AAAGGCCCGGGTCACATATGTGGACACAAGAGATTACTTTATAATTCTTAGCATAGCCTACCCAACCTTATCTGAGATCA

AAGGGGTGATAGTTCATAAGATAGAAGCTATATCATACAACATTGGGGCACAGGAATGGTATACTACTATCCCTAAATAT

GTAGCCACTCAGGGATATTTGATATCGAATTTCGATGAGACATCATGTGTCTTCACTCCAGAGGGGACAGTCTGCAGCCA

GAATGCGCTGTATCCAATGAGCCCATTGCTTCAGGAATGCTTCAGGGGGTCGACAAAATCGTGTGCCAGAACCCTAGTTT

CAGGGACCACAAGTAATAGATTTATCCTATCAAAAGGGAACTTGATTGCAAATTGTGCGTCAGTTCTGTGCAAGTGTTAC

ACAACGGAGACAGTCATCAACCAAGATCCTGATAAACTACTAACTGTTATAGCCTCAGATAAGTGTCCCGTAGTTGAGGT

GGATGGAGTGACAATACAAGTCGGCAGTCGAGAGTACCCAGATTCTGTATACCTACATAAAATAGACTTAGGCCCAGCCA

TCTCTCTGGAAAAACTGGATGTAGGCACCAATTTAGGCAATGCAGTCACAAGACTGGAGAATGCAAAGGAGCTCCTAGAC

GCATCAGACCAGATACTGAAGACTGTTAAAGGGGTACCCTTCAGTGGCAATATGTACATAGCACTGGCAGCTTGCATTGG

GGTATCCCTAGGGCTTGTCACATTAATATGCTGCTGTAAGGGGAGATGTAGGAACAAGGAGATTCCTGCCTCCAAAATCA

ACCCAGGGCTCAAACCCGACCTGACCGGGACTTCAAAGTCTTACGTGAGATCACTGTAGTCAGAATAACCCGAATCACCC

GGCATCACGCGTATACATGTGCGACACAACCAGTCAGAGGACGCAGAAGACTCAACCTCCGATCACCGACCAGACCCCAA

TCTACGCCCTACTGCACATTGGTCATCAAACAAAACTTAGGACGAAAGGTCAATCACCATGTCCGCACAAAGGGAGAGGA

TCAATGCCTTCTACAAAGACAACCCTCACAATAAAAACCATAGGATAATCCTGGATAGGGAACGCTTAACTATTGAAAGA

CCCTACATCTTACTTGGAGTCCTGCTGGTAATGTTCCTGAGTCTAATCGGGCTGCTAGCCATTGCAGGGATCAGGCTTCA

CCGGGCCACCGTTGGAACTGCGGAGATCCAGAGTCGGCTGAATACCAACATTGAGTTGACCGAATCCATTGATCATCAAA

CTAAGGATGTCTTAACACCCCTGTTCAAAATCATTGGTGATGAAGTCGGTATCAGAATTCCACAGAAGTTCAGTGATCTT

GTCAAGTTCATCTCCGATAAGATTAAGTTCCTCAACCCTGACAGAGAATACGATTTTAGGGATCTCCGGTGGTGTATGAA

CCCCCCTGAGAGAGTCAAAATTAACTTTGATCAATTCTGTGAATACAAAGCCGCGGTCAAGTCAGTTGAACATATATTTG

AGTCATCATTCAACAGGTCAGAAAGATTGCGACTATTGACTCTTGGGCCCGGAACAGGCTGTCTCGGCAGGACAGTGACA

AGAGCTCAGTTCTCAGAACTTACGCTGACCCTGAGGGACCTGGATTCCGAGATGAAGCACAACGTGTCCTCAGTGTTTAC

CGTAGTTGAAGAGGGATTATTCGGAAGAACATATATTGTCTGGAGATCTGACACCGGGAAACCGAGCACCAGTCTAGATA

TTGGCCAGTTTTTAAGAGTCTTCGAGATCGGGTTGGTGAGGGATCTCGAGCTGGGTGCCCCCATTTTCCATATGACCAAC

TACCTCACAGTGAACATGAGTGATGACTATCGGAATTGCCTTTTAGCGGTAGGGGAGTTGAAGCTGACAGCCCTATGCAC

CCCATCTGAGACTGTGACTCTGAGTGAGAGAGGAGTTCCAAAGAGAGAGCCTCTTGTGGTTGTGATACTCAACCTAGTTG

GGCCTACTCTAGGGGGCGAACTATACAGTGTCTTGCCTGCCTCTGACCTCATGGTGGAGAAACTCCATCTATCCTCACAT

AGAGGGATCATCAAAGACAACGAGGCCAATTGGGTAGTACCGTCTACCGATGTTCGTGATCTCCAAAACAAGGGAGAATG

TCTGGTGGAAGCATGCAAAACTCGACCTCCTTCATTTTGCAATGGCACAGGAATAGGCCCATGGTCAGAGGGGAGAATCC

CTGCCTACGGAGTGATCAGGGTCAGTCTTGACTTAACTAGTGACCCAGGTGTGGTTATCACTTCAGTGTTTGGCCCACTG

ATACCTCACCTATCTGGCATGGATCTTTACAACAATCCGTTTTCAAGAGCTGCATGGCTGGCTGTACCACCCTACGAGCA

GTCATTTCTAGGAATGATAAATACAATTGGCTTCCCGGACAGAGCAGAGGTCATGCCGCACATTTTGACCACAGAGATCA

AAGGGCCTCGAGGTCGTTGTCATGTTCCTATAGAGTTGTCCCGCAGGATTGATGATGATATCAAGATCGGGTCCAATATG

GTTGTATTGCCGACGAGGGATCTGAGGTACATAACAGCCACTTATGATGTTTCCAGGAGCGAGCATGCAATCGTGTACTA

TATCTATGACACGGGTCGCTCATCATCTTACTTCTACCCAGTTCGATTGAATTTCAAGGGCAATCCTCTCTCTCTGAGGA

TAGAGTGTTTCCCCTGGTCTCATAAGGTGTGGTGCTACCATGATTGTCTTATATACAACACCATGACAAACGAAGAAGTC

CATACGAGAGGGCTGACCGGTATAGAGGTAACATGCAATCCAGTCTGAGTCGAGCTGAAACCATCGCTCAAGCAGGCTTC

CGAGCCATCCCCCAGTTCAAACAGCATAGTCTGGGACACTCAGCAGCACAACCTAGCCAACAATGTTATAAAAAACTTAG

GAGCCAAGGTTGTAGGAGCCATGGACTCACTATCAGTCAATCAGGTTTTGTACCCTGAGGTCCATCTAGATAGCCCTATT

GTCACAAACAAACTAGTTGCTATCCTTGAATACTCGGGGATCAACCACAACTATGTTCTTGAGGACCAGACTCTTATCAA

GAATATTAGATATAGATTGAGTTGCGGTTTCTCAAATCAAATGATCATCAATAATAGGGGGTTAGGTGAAACAGTCAATT

CCAAGCTTAAAAGTTACCCCCGTAACTGTCATATCATATACCCAGACTGCAATAAGGAGTTGTTTTGTATCAAAGATAGC

TGCATATCTAGAAAGCTCTCGGAGCTATTCAAGAAGGGTAATTCCTTGTACTCTAAGGTAAGTCACCAGGTACTAGATTG

TCTTAAGAGAGTCAACGGGAAATTAGGGCTGGGCACAGATCTTAATCACGGCCTGAAGGATGGTATCCTCGACTTGGGGT

TGCACATGCATAGCTCTCAGTGGTTCGAGACCTTTCTATTCTGGTTCACTATCAAGACAGAGATGAGATCAATGATCAAA

GAACAGTCCCATATATGCCACAAGAGGAGGTATAACCCAATTTTTGTGTCGGGGGATGCATTCGAGGTGCTTGTATCACG

AGACCTCGTAGTGATAATTGATAAGAACACCCAGTATGTCTTCTACCTGACGTTTGAGCTGGTCCTTATGTATTGTGATG

TCATAGAGGGTAGACTTATGACGGAGACAGCCATGGCTATTGACCAGAGATATTCAGAGCTTCTAAACCGGGTCAGATAC

TTGTGGGACCTTATCGATGGGTTCTTCCCAACACTGGGTAACACCACATACCAAGTTGTTGCTCTGCTTGAACCACTGTC

GTTGGCTTATCTTCAACTTCAGGATGTCACTCTAGAGTTAAGAGGTGCCTTTTTGGACCACTGCTTCAAAGAACTTTATG

AGATACTGGAGCATTGTGGCATTGACACGGAAGGTACCTACAATTCCATCACTGAGGGATTGGATTACGTATTTATCACT

CACGATATACACTTAACTGGGGAGATTTTTTCATTTTTCCGGAGTTTCGGACACCCCCGCCTCGAAGCGGTCACCGCTGC

AGAGAATGTCAGAAAACATATGAACCAACCGAAGGTAATCAGTTATGAGACTATGATGAAAGGGCATGCAGTATTTTGCG

GGATAATCATAAATGGTTTTAGGGACCGGCACGGCGGCAGCTGGCCCCCTGTTGCATTGCCAGAACATGCTTCTGCTGCG

ATCCGGAATGCGCAGGCATCCGGCGAAGGACTGACCCACGACCTGTGTATAGACAACTGGAAGTCCTTTGTAGGATTCAG

ATTTGGCTGCTTCATGCCGCTCAGCCTAGATAGTGATTTGACCATGTACCTCAAAGACAAAGCATTGGCTGCATTGAAGA

ATGAGTGGGATTCAGTTTACCCGAAAGAATACCTCCGTTATAATCCACCTAGAGGGACAGAGTCAAGGCGACTAGTAGAG

GTGTTCCTGAATGACTCCAGCTTTGATCCTTATAACATGATAATGTACGTGGTGAATGGCTCCTACCTTAAAGACCCTGA

GTTTAATCTCTCATACAGCTTAAAAGAAAAGGAGATAAAAGAGACAGGGCGGTTGTTTGCCAAGATGACCTACAAGATGC

GGGCCTGTCAGGTGATTGCTGAAAATCTGATATCAAATGGTGTTGGGAAGTATTTCCGAGACAATGGGATGGCAAAAGAC

GAGCATGACCTAACAAAAGCCCTTCACACTCTGGCAGTCTCAGGGGTTCCCAAGAACAACAAAGACAACCACCGAGGTGG

GCCCCCCAGAAGGAACGCAAGCCGAGAGATGAGATCAAGCCAAGCCGCCAACACACAAAATAGGGACAAGATCCAAGGGG

GCCCTATGTACAACTACTTGCAATGCCAACCAACCAGCCCTGATCAAGGTGAGTCATACGAGACTGTTAGTGCATTCATC

ACCGCTGACCTTAAGAAGTATTGCCTAAATTGGAGATACGAGACAATCAGCATATTTGCACAGAGATTGAATGAAATATA

TGGGTTGCCATCCTTCTTTCAATGGTTACATAAGGTATTGGAAAAATCCGTCCTCTACGTCAGTGATCCGCATTGCCCTC

CCGACTTAGATGATCATATCCCTCTGGACAGTGTCCCTAATGCCCAAATATTCATCAAGTACCCAATGGGCGGAATAGAA

GGTTATTGTCAAAAGCTATGGACAATCAGTACTATACCATACTTGTATCTAGCAGCCTATGAGAGTGGAGTAAGAATCGC

CTCACTAGTGCAAGGTGACAATCAGACAATTGCAGTGACAAAAAGAGTTCCAAGTTCTTGGCCTTATTCACTAAAAAAGA

GGGAGGCATCTAAAGCAGCTCAAAATTACTTTGTGGTCTTAAGGCAAAGGTTGCACGATGTAGGTCATCACTTGAAGGCT

AATGAGACCATAGTATCTTCTCACTTTTTTGTATACTCTAAAGGGATCTATTATGACGGCCTGTTAGTTTCACAATCACT

AAAGAGCATCGCCAGATGTGTCTTCTGGTCCGAGACTATCGTGGATGAAACTAGAGCGGCCTGCAGCAATATTGCAACAA

CTATCGCTAAGAGTATAGAGAGGGGTTATGATAGGTACCTCGCATACTCTTTGAATATCCTCAAAATTTTCCAACAGATC

CTTGTATCCCTTGACTTCACGATTAATACAACAATGACTCAAGATGTCGTGGCACCGATCATCGAGAACGGTGATTTACT

GATAAGGATGGCACTCTTGCCAGCACCCATTGGGGGTCTCAATTATCTCAACATGAGCAGGTTGTTTGTGAGAAATATCG

GTGACCCGGTCACTTCCTCTATAGCCGACCTGAAGAGGATGATAGACGCTGGGCTAATGCCAGAAGAAACATTGCATCAA

GTGATGACCCAGACCCCGGGAGAATCATCCTACCTTGATTGGGCAAGTGATCCTTATTCTGCCAACCTACCCTGCGTACA

GAGTATAACTCGACTTCTCAAGAACATCACTGCACGGTATATTTTAATCAGCAGCCCAAATCCGATGCTGAAAGGGTTGT

TCCACGAGGGGAGCAGAGATGAAGACGAGGAGCTTGCGAGTTTCTTAATGGATCGGCATATAATTGTTCCGAGAGCTGCA

CATGAAATCTTAGACCACAGCATAACCGGGGCAAGAGAAGCTATAGCCGGGATGTTGGACACCACCAAGGGTCTGATTAG

AACAAGTATGAAACGGGGTGGCCTCACCCCTCGAGTATTAGCCCGCCTTTCCAATTATGATTATGAACAATTCAGATCCG

GGATAACACTATTGACAAAGAAAGGGCAGTGTTATCTCATTGACAAGGACTCGTGCTCGGTGCAGCTCGCTATAGCTCTG

AGGAGCCATATGTGGGCAAGGTTGGCTCGCGGGAGACCTATCTATGGCTTGGAGGTGCCTGATGTACTGGAATCAATGAA

CGGCTACCTTATCAAACGTCATGAGTCTTGTGCCATCTGTGAAACGGGCTCAAGTCACTACGGGTGGTTCTTCGTCCCTG

CAGGGTGTCAGCTTGACGATGTCTCAAGAGAGACTTCAGCTCTTCGTGTGCCTTATGTCGGATCAACCACTGAGGAAAGG

ACAGATATGAAACTTGCTTTCGTTAGATCTCCAAGCCGATCCCTCAAATCAGCAGTCAGGATTGCCACAGTCTACTCATG

GGCCTACGGGGATGATGAGAAATCATGGAGTGAAGCTTGGATGCTAGCTAGGCAGAGAGCCGATATCACCTTAGATGAAC

TGAGAATGATCACTCCAGTCTCTACATCCACCAACCTAGCTCATCGGTTAAGGGATCGGAGCACTCAGGTGAAATATTCG

GGGACATCCCTTGTGAGGGTTGCAAGGTACACAACCATCTCCAATGACAATTTGTCATTTGTAATATCTGAGAAAAAAGT

AGATACCAACTTCATTTACCAGCAAGGGATGCTGCTCGGTCTTGGGATCCTTGAGAATCTATTCAGGTTAGAGGCCACCA

CAGGGGTATCCAACACAGTGCTACACCTGCACGTAGAGACAGAATGTTGTGTTGTACCTATGGTTGATCACCCAAGGATA

CCGAGTCTCCGCAATATCAAAGTCACGAACGAGCTATGCACAAACCCTCTTATCTACGACAAGTCCCCCATCATAGAACA

CGATGCAACTCGATTATACTCACAAAGCCACAGGAGACATTTGGTGGAGTTTGTTACCTGGTCAACAAGCCAGCTCTATC

ATATACTGGCCAAATCTACAGCAATGTCCATGATTGAGCTGATCACGAGATTTGAGAAAGATCACATGAATGAAATAGCC

GCTCTGATTGGCGATGACGACATCAACAGTTTCATCACAGAATTTTTACTTGTGGAGCCCAGACTGTTTATAGTTTACCT

CGGCCAGTGTGCTGCCATCAATTGGGCTTTTGATATACATTATCATCGGCCCTCGGGCAAGTACCAGATGGGGGAACTCC

TCTACTCTTTACTCTCTCGGATGAGCAAGGGAGTATATAAGATCTTCACTAATGCTCTGAGCCACCCCAAAGTTTACAAG

AAATTTTGGCGAAGTGGTATAATTGAGCCGGTTCATGGCCCATCCCTAGATACACAAAATCTACATGTCACTGTCTGTGA

CATGATATACGGATCATACGTCACCTATCTGGATCTTTTGCTGAATGATGAGCTGGATGCTTACCCGTATTTGCTCTGCG

AGAGTGATGAGGACGTGGTCACAGACAGGTTCGACAACATTCAAGCCAAACATCTCTGTGTACTGGCCGATGTATACTGC

AGCTCCAAGAGGTGTCCCTCGATAATCGGGATGTCCCCTATAGAAAAATGTACCATCCTCACACATTACATCAAGGGAGA

ATCGGTACAATCCCCGTCCGGGATCTCATGGAACACTGATCCCCTTGTAGTAGATCATTACTCATGCTCTCTGACCTACC

TTCGCCGTGGTTCCATCAAACAAATCAGATTGAGAGTGGATCCTGGGTTTGTGTTCGAGGCGTTGACAGACATCGACTTC

AAACAGCCTCGCAAGGCTAAGTTGGACGTATCGATTGTGGGATTGACTGATTTTTCTCCCCCTTGGGATAACGTCGGTGA

TTTTCTAGGGACTATCAACACATTGAGGCACAATCTGCCCGTCACCGGGACCGGGGTCTCGAACTATGAAGTCCACGCTT

ATCGTAGAATTGGTCTGAATTCATCAGCATGTTATAAAGCTGTAGAGATCTCCACGTTAATAAAGTCATCCTTAGAAGCC

GGGGAGAATGGATTGTTCTTAGGAGAAGGCTCCGGCTCGATGCTGGCTGCGTACAAGGAAGTTCTTAAATTGGCAAACTG

TTATTACAACAGTGGGGTAACAGCAGAGGGTAGAGCCGGACAGAGGGAAATCTCTCCCTATCCCTCAGAGGTGAGCCTGG

TAGAGAGTCAGATGGGGATAGAGAGAAGTGTTAAAGTCCTGTTCAATGGCAAACCTGAAGTAACCTGGGTAGGGACCACC

GATTGCTACAAGTATATAATCAGTAACATTCAGACCTCAAGTCTGGGTTTCATACACTCAGATATCGAGACACTCCCAAC

CAAGGATGCCGTTGAGAAGTTGGAGGAATTTGCCTCTATTCTATCTCTATCCCTAATTTTGGGAAAAATCGGCTCTATTA

CAGTTGTCAAAATTATGCCCATTAGCGGAGATTTTACCCAAGGCTTCATAGCCTATGCCATTCAATATTTCAGAGAGAGC

CTGCTTGCCTATCCGAGATATAGTAACTTCATCTCGACTGAGTGTTACCTTATTATGATAGGATTAAAGGCCAATCGGTT

GATAAACCCAGAAGCCATTAAGCAAAGCATAATCAGGGTGGGGACTAGGACTGCACCAGGACTTGTGAGCCACATATTAT

CAGAGAAACAGAAAGGTTGTATTCAATCTTTTCTGGGTGATCCTTATGTCCAAGGAGACTTCAATAAGCACCTTAAAGCT

CTAACTCCTATTGAGAAAATCCTTGTAAATTGTGGTCTCTCGATCAATGGTACAAAAATCTGTAGGGATCTAATTCACCA

TGATATTGCCTCCGGTCCAGACGGTCTGATGAGCTCCACAATTATTTTATACAGGGAACTGGCCCATTTCAAAGACAATA

TAAGAAGTCAGCACGGCATGTTCCACCCCTACCCAGTATTGGCCAGTAGCAGGCAACGTGAATTGATCCTTCGAATAGCC

AAGAAGTTTTGGGGGTATGTCTTGCTATATTCTGATGACCCGGCACTAATCAAACAAACGATCAAGAACTTGAAGCGGAA

TTACCTAACCTTTGACTTACACAGTAATCCGTTTATTAAGGGCTTATCCAAAGCTGAGAAACTGCTAGTGCGGACAAGTT

CACTCAGAAGGGAATGGTTGTTCACTCTCGATACGAAAGAAGTGAAAGAGTGGTTCAAATTGGTGGGTTACAGTGCACTC

GTCAGAGGTTAATTCGCGATACATCTGCCCCCTTCTCCTCCACCATGGGACTCTACTGGCAATCCAAAAGATTAAAGAAA

ACTACATATTGGATAAGTATCTATTCCCAGCTTTGTCTGGT

>KM212177.1/goat/Senegal/2013-03-11

ACCAAACAAAGTTGGGTAAGGATAGATCTTACAATGACTATAGACTAGCAAACTTAGGAGTAAAGATCCTACTGTCGGGG

AGAGGAGGAGGAGCAAGATCTTTGACCATGGCGACTCTCCTTAAAAGCTTAGCACTGTTCAAGAGGAACAAAGACAAAGC

GCCTACTGCGTCAGGTTCAGGAGGGGCCATCCGGGGGATTAAGAATGTTATCATAGTCCCTATCCCCGGGGACTCATCCA

TCATTACCCGCTCAAGACTGCTCGACAGGCTTGTCAGACTGGCCGGAGATCCTGACATCAACGGGTCAAAGCTGACCGGC

GTGATGATCAGCATGTTATCTTTGTTCGTGGAGTCACCCGGGCAATTGATACAGCGGATCACAGATGATCCAGATGTTAG

CATCCGTCTTGTTGAGGTAGTTCAAAGTACCAGGTCCCAGTCCGGATTGACCTTTGCATCACGTGGTGCTGATTTGGACA

ATGAGGCAGATATGTATTTTTCAACTGAAGGACCCTCGAGTGGAAGTAAGAAAAGGATCAACTGGTTTGAGAACAGAGAA

ATAATAGACATAGAAGTGCAAGATGCAGAAGAGTTCAATATGTTGTTAGCCTCCATCTTAGCACAAGTTTGGATCCTCCT

GGCCAAGGCGGTTACGGCACCAGATACAGCAGCCGACTCAGAACTGAGAAGGTGGGTTAAATACACACAACAAAGGAGAG

TGATTGGGGAATTTCGCTTTGACAAAGGGTGGTTGGACGCAGTCCGCAACAGGATTGCAGAAGATCTATCACTCCGGCGG

TTCATGGTATCCCTGATACTTGACATCAAGAGGACCCCCGGCAACAAGCCAAGGATTGCAGAAATGATCTGCGACATTGA

CAACTATATTGTCGAGGCCGGACTCGCAAGTTTCATTCTTACTATCAAGTTTGGTATTGAAACCATGTATCCTGCACTAG

GCCTTCACGAGTTCGCTGGGGAATTGTCCACTATTGAATCCTTGATGAACCTGTATCAACAGCTAGGAGAAGTTGCACCC

TATATGGTAATTCTAGAGAACTCAATTCAGAACAAGTTTAGTGCAGGAGCCTATCCTCTCCTCTGGAGCTATGCGATGGG

TGTCGGAGTCGAGTTGGAGAACTCAATGGGGGGCTTGAACTTTGGCAGGTCATATTTTGACCCGGCCTATTTTCGTCTCG

GACAGGAGATGGTCAGAAGATCTGCAGGAAAGGTCAGCTCTGTAATTGCGGCTGAGCTTGGCATCACAGCAGAGGAAGCC

AAACTAGTCTCGGAAATCGCCTCGCAGACTGGGGAAGAACGGACCGTTAGAGGGACTGGGCCTCGACAGGCGCAGGTTTC

CTTCCTCCAGCACAAAATAGGTGAGGGAGAGTCGCCTACACCAGCGACCAAAGAAGAAGTCAAAGCTGCGATCCCAAACG

GATCCGAAGGAAGAGACATAAAGCGAACACGCCCAGGGAAGCCCAGAGGAGAAACTCCCGGGCAACTGCTTCTGGAGATC

ATGCCAGAGGATGGAGTCTCGCGAGAGTCTGGTCAAAACCCTCGTGAGGCTCAAAGATCGGCTGAGGCACTCTTCAGGCT

GCAGGCCATGGCCAAAATTCTGGAGGACCAGGAGGAGGGAGAAGACAACAGTCAGATCTACAACGACAAGGATCTCCTCA

GCTGAGCAGACACACCCTCCGTCCAAATCAGCGACAAGACATCACCTGTCAGTATTATAAAAAACTTAGGACCCAGGTCC

AACCAACCGCACATCGACACCCCAGTCAATCGAGCGGAGACCACCGATGGCAGAAGAACAAGCATACCATGTCAACAAGG

GGCTGGAATGTATCAAGTCCCTCAAAGCCTCTCCCCCGGATCTATCCACCATCCAAGATGCCCTTGAGAGCTGGAGAGAG

GGGTTCAACCCCTCAGGCCGTGCAACACCGAACCATGATACGTCCGAGGGGGACCATCAGAATATCAACCAATCATGCTC

ATCAGCAATCGGATCAGACAAAGTCGACATGTCTCCTGAAGGTAATCTCGGATTTAGAGAAATCACTTGTGATGACAATG

AGGCTGGGCTCGGAGGAGTTCAGGACAAAAGATCCGACTCTCAAGTACAGCGTTATTATGTTTATAACCACGGGGGTGAA

GAGATTGAAGGACTCGAGGATGCTGACTCTCTCGTGGTTCAAGCAAATCCTCCGATTGCTAACACCTTCGATGGAGGAGA

GGATGGATCTGACAACAGCGATGTGGACTCTGGCCCAGATGATCCCGGCAGAGATCCTCTATATGACCGGGGACCTGCTG

CCGGCAATGATGTCTCTAGGTCAACAGATGTCGAAAAATTAGAAGGTGATGACATTCAAGAAGTTCTTAACTCCCAGAAG

AGTAAAGGAGGAAGATTCCAGGGCGGGAAAATATTGCGAGTCCCGGAAATACCCGATGTCAAGAACTCCAGACCATCGGC

CCAATCAATTAAAAAGGGCACAGACGGGAGCTCAGTCTTATCTGGAATGGTGACAGAGTGTTCATCGATAAGTGGTGCAA

CCCAAGCTGTGCTAGAGTCCAGATGGGAGTCATCAGAGCGAAATGCATCTGTGGGGAGTGTCCCCAAATCTGTGAGGAGT

GCAAAGACGATCCAGGGGTTAACACAAGAATCTGGTACCATAGCATCACTGACTCAGCCTAAAGAAAATGACTCCGAGTA

TGAGTATGAGGATGATCTATTCACTGAGATGCAGGACATTCGTGCAAGCATTGCTAAGATCCACGATGACAACAAAACTC

TCCTCTCAAAACTTGATTCTCTACTGTTATTGAAAGGAGAAATCGATACTATCAAGAAACAAATCAGCAAACAAAATATA

AGTATATCTACCATTGAGGGCCATCTATCCAGTATAATGATAGCCATCCCAGGTTTTGGGAAGGACATCAAGGACCCAAC

ATCTGAGGTTGAGCTGAACCCAGATTTGAGACCTATAATCAGCCGCGATTCTGGCAGGGCTCTTGCGGAGGTCCTTAAGA

AACCCGCTGTTGATAGGTCTCAGAAGAGCGGAATCAAAGTCAACTCCGGTTCAAAGGGTCAGCTCCTCAAGGATCTCCAG

CTAAAACCCGTCGACAAGCAGGCAAGCTCTGCAATCGGGTTTGTTCCATCCGACCATGAATCATCCAGAAGTGTCATCCG

CTCCATAATCAAGTCGAGCAAGCTTAACATTGATCACAAAGACTATCTTCTAGATTTACTGAATGATGTGAAAGGCTCTA

AGGATCTCAAGGAATTCCACAAGATGCTGACAGCCATTCTTGCCAAGCAGCCGTAACACATCCCATAATCTACATCTCAT

ACTCGGTTGAAAGCATCCTCTCAATCAGGCTATTACAAAAAACTTAGGAGCAAGGGCAACTGAGCTTCACAGACAGGATG

ACCGAGATTTACGATTTCGACAAATCAGCATGGGATGTCAAAGGGTCAATTGCTCGCATAGAACCCACCACCTATCACGA

CGGCCGACTGGTACCCCAGGTGAGGGTCATCGACCCTGGTCTGGGAGACAGAAAAGATGAGTGCTTCATGTACCTGTTCC

TTCTAGGAGTGATTGAGGATAACGACCCCCTTTCTCCCCCAGTCGGGAGAACCTTCGGGTCTTTACCTCTAGGTGTCGGC

AGGTCAACTGCTAAGCCAGAGGAACTACTAAGGGAGGCCACAGAACTAGATATAGTGGTGAGGCGCACGGCAGGACTAAA

TGAGAAATTGGTGTTTTACAACAACACTCCGCTATCTTTGTTAACACCATGGAGGAAAGTCTTGACGACCGGAAGTGTGT

TCAGCGCCAACCAGGTTTGCAATGCGGTCAACCTAGTCCCACTTGATACTCCCCAGAGATTCAGGGTTGTGTACATGAGT

ATAACTAGATTGTCAGATAATGGTTATTATAGTGTCCCCAGAAGAATGTTGGAGTTCCGCTCAGCCAATGCAGTCGCTTT

CAATATCTTGGTTACACTGAGAATTGAAAATGGCACAAACCCTAGAAGATACATAGTCGGCTCATGGGAGAATCCAGAAG

TCACATTTATGGTACACGTGGGCAACTTTAGAAGAAAGAAGAACGAAGTATACTCTGCTGATTATTGCAAAATGAAGATT

GAAAAGATGGGTCTAGTTTTTGCCTTGGGAGGAATAGGTGGAACAAGTCTCCATATTCGAAGCACAGGGAAAATGAGCAA

GACCCTCCATGCACAGCTGGGGTTCAAGAAAATTCTATGCTACCCTCTGATGGATATCAATGAGGATCTTAACCGATATC

TCTGGCGGGCAGAGTGCCGAATAGTCAAAATCCAGGCCGTCTTACAGCCATCAGTACCCCAAGAATTCCGTGTCTATGAT

GATGTCATCATCAACGATGACCAAGGCCTGTTCAAGATCCTGTAAGTCACCTGCAGGGCCATAGACCCACCTCCAGGAAG

AGGCCCTCCCACCTCCCGCCCAAAACACACACCCCAAGACTCCCCGAAACAGCGCCGGCAAAACCACTCTCCTGGCGGGA

CGAGGCCGACC-CCCCCAACAAACCCGCTCCGGAGGGGGG-CCCCCCCCGCGACCCGCCGGGCGCCGACCGCGCGAGCTG

GCGAGACACCCCCAAGAGGCCTCGACGAAGATCGGGGAACCCTTTCCCCA-AAAATGGCCCCCC-AACACACCCAAGGCC

ATGCGCTGAGCACCCCCAGCCCCCCAACCGGCGGGACAGGAGGCCAGGACGGGGCTCCCCAAAACAAAGGGACCACCAGG

GAAGGCTCTGCCGACCCAGCACAGACCCGGCCCAAACAAGGCAGACCCCCAAGACGAAATCCCCCAGAGCACCCCATTCA

TCATCAAACAAAACTTAGGGGCCAAGTCCAGACGACAGACCCCGGCCCGCCGGGCAGCAAGCCACAGAAAAGAAGGAGAC

AAGATCGGCCCGGGAAGAGGACACCCAACCACAGGAACCCCCAGCTCCAGGGGAAGAG-CCCCCCCTCCACACAACGGGG

GAACGGGCGAGAGACACCGAGGAGAGCACGATCGGGAGCCGGGAGCCGGAGACGCCGGCGGCAAGCAGGCCGGGCACTCG

GCCGACACCACCGGAGGACCCGAGGAGAGCCCGACTCCGCGCAGACTCGACGGCAGAGGACCCAGTCCCGCCACAGAGAG

AGGCGAGAGCCCT--------CCCTCCTGCACACAACCGGCAGACAGACCGCGGGAGGACCACCTCCCAGCAAGACTGAC

GTCTCCCCCGCAGCCGAGCCGCAGCTCCTCAGAAGAGGGCACCCCCGAAATCTCCCAGAACCCCTCCGCTCACGGCCGCC

GGGACAAGAGCTCCGCCGCCAAAGACCAGAGCCACCGAGAGTCAACCAGGGGTCCCCCCCAGTCCCTCCCGGCGATCCCC

CACAGAGGCCCCAGCAAGGTCCCCCGGCCCGACTACCCCTGCATGCGCCGAAAGGGAAACCCACCAGAGGACGTCCAGCA

CATCCGTGTATAGACATCATGACACGGGTCGCAACCTTGGTATTGCTGGTTCTTTTCCCTAACACTGTCGCGTGCCAGAT

TCACTGGGGCAATCTATCCAAGATAGGGATTGTCGGAACAGGGAGTGCCAGTTACAAGGTGATGACCAGACCAAGCCACC

AAACTCTGGTTATAAAGTTGATGCCAAATATAACAGCCATCGACAATTGTACAAAATCAGAGATTTCAGAGTACAAAAGA

TTGCTAATCACAGTGTTAAAGCCTGTAGAGGATGCTCTGTCAGTAATAACCAAGAATGTAAGACCAATTCAAGCTCTAAC

ACCTGGGCGCAGGACCCGCCGTTTCGCCGGAGCTGTTCTGGCCGGAGTAGCACTTGGAGTCGCGACGGCTGCTCAAATAA

CTGCCGGAGTCGCACTCCATCAGTCATTGATGAATTCCCAAGCAATTGAAAGTTTAAAAACCAGTCTTGAGAAGTCAAAT

CAGGCAATAGAAGAAATCAGACTTGCAAATAAGGAGACCATACTGGCAGTACAGGGCGTCCAAGATTATATCAACAACGA

GCTTGTCCCCTCTGTTCATAGAATGTCATGTGAGCTAATAGGTCACAAACTCAGTCTCAAACTCCTTAGGTATTATACCG

AGATCCTGTCTATATTCGGGCCTAGCCTTCGAGACCCGATAGCTGCTGAAATATCAATCCAGGCACTCAGCTATGCACTA

GGCGGAGACATCAATAAAATTCTGGACAAGCTTGGGTATAGCGGCGGGGATTTCCTTGCTATTCTAGAAAGCAAGGGGAT

AAAGGCCCGGGTCACATATGTGGACACAAGAGATTACTTTATAATTCTTAGCATAGCCTACCCAACCTTATCTGAGATCA

AAGGGGTGATAGTTCATAAGATAGAAGCTATATCATACAACATTGGGGCACAGGAATGGTATACTACTATCCCTAAATAT

GTAGCCACTCAGGGATATTTGATATCAAATTTCGATGAGACATCATGTGTCTTCACTCCAGAGGGGACAGTCTGCAGCCA

GAATGCGCTGTATCCAATGAGCCCATTGCTTCAGGAATGTTTCAGGGGGTCGACAAAATCGTGTGCCAGAACCCTAGTTT

CAGGGACCACAAGTAATAGATTTATCCTATCAAAAGGGAACTTGATTGCAAATTGTGCGTCAGTTTTGTGCAAGTGTTAC

ACAACGGAGACAGTCATCAACCAAGATCCTGATAAACTACTAACTGTTATAGCCTCAGATAAGTGTCCCGTAGTTGAGGT

GGATGGAGTGACAATACAAGTCGGCAGTCGAGAGTACCCAGATTCTGTATACCTACATAAAATAGACTTAGGCCCAGCCA

TCTCTCTGGAAAAACTGGATGTAGGCACCAATTTAGGCAATGCAGTCACAAGACTGGAGAATGCAAAGGAGCTCCTAGAC

GCATCAGACCAGATACTGAAGACTGTCAAAGGGGTACCCTTCAGTGGCAATATGTACATAGCACTGGCAGCTTGCATTGG

GGTATCCCTAGGGCTTGTCACATTAATATGCTGCTGTAAGGGGAGATGTAGGAACAAGGAGATTCCTGCCTCCAAAATCA

ACCCAGGGCTCAAACCCGACCTGACCGGGACTTCAAAGTCTTACGTGAGATCACTGTAGTCAGAATAACCCGAATCACCC

GGCATCACGCGTATACATGTGCGACACAAGCAGTCAGAGGACGCAGAAGACTCAACCTCCGATCACCGACCAGACCCCAC

TCTACGCCCTACTGCACATTGGTCATCAAACAAAACTTAGGACGAAAGGTCAATCACCATGTCCGCACAAAGGGAGAGGA

TCAATGCCTTCTACAAAGACAACCCTCACAATAAAAACCATAGGATAATCCTGGATAGGGAACGCTTAACTATTGAAAGA

CCCTACATCTTACTTGGAGTCCTGCTGGTAATGTTCTTGAGTCTAATCGGGCTGCTAGCCATTGCAGGGATCAGGCTTCA

CCGGGCCACCGTTGGAACTGCGGAGATCCAGAGTCGGCTGAATACCAACATTGAGTTGACCGAATCCATTGATCATCAAA

CTAAGGATGTCTTAACACCCCTGTTCAAAATCATTGGTGATGAAGTCGGTATCAGGATTCCACAGAAGTTCAGTGATCTT

GTCAAGTTCATCTCCGATAAGATTAAGTTCCTCAACCCTGACAGAGAATACGATTTTAGGGATCTCCGGTGGTGTATGAA

CCCCCCTGAGAGAGTCAAAATTAACTTTGATCAATTCTGTGAATACAAAGCCGCGGTCAAGTCAGTTGAACATATATTTG

AGTCATCATTCAACAGGTCAGAAAGATTGCGACTATTGACTCTTGGGCCCGGAACAGGCTGTCTCGGCAGGACAGTAACA

AGAGCTCAGTTCTCAGAACTTACGATGACCCTGATGGACCTGGATCTCGAGACGAAGCACAACGTGTCCTCAGTGTTTAC

CGTAGTTGAAGAGGGATTATTCGGAAGAACATATATTGTCTGGAGATCTGACACCGGGAAACCGAGCACCAGTCTAGATA

TTGGCCAGTTTTTAAGAGTCTTCGAGATCGGGTTGGTGAGGGATCTCGAGCTGGGTGCCCCCATTTTCCATATGACCAAC

TACCTCACAGTGAACATGAGTGATGACTATCGGAATTGCCTTTTAGCGGTAGGGGAGTTGAAGCTGACAGCCCTATGCAC

CCCATCTGAGACTGTGACTCTGAGTGAGAGAGGAGTTCCAAAGAGAGAGCCTCTTGTGGTTGTGATACTCAACCTAGTTG

GGCCTACTCTAGGGGGCGAACTATACAGTGTCTTGCCTACCTCTGACCTCATGGTGGAGAAACTCCATCTATCCTCACAC

AGAGGGATCATCAAAGACAACGAGGCCAATTGGGTAGTACCGTCTACTGATGTTCGTGATCTCCAAAACAAAGGAGAATG

TCTGGTGGAAGCATGCAAAACACGACCTCCTTCATTTTGCAATGGCACAGGAATAGGCCCATGGTCAGAGGGGAGAATCC

CTGCCTACGGAGTGATCAGGGTCAGTCTTGACTTAGCTAGTGACCCAGGTGTGGTTATCACCTCAGTGTTTGGCCCACTG

ATACCTCACCTATCTGGCATGGATCTTTACAACAATCCGTTTTCAAGAGCTGCATGGCTGGCTGTACCACCCTACGAGCA

GTCATTTCTAGGAATGATAAATACAATTGGCTTCCCGGACAGAGCAGAGGTCATGCCGCACATCTTGACCACAGAGATCA

AAGGGCCTCGAGGTCGTTGTCATGTTCCTATAGAGTTGTCCCGCAGGATTGATGATGATATCAAGATCGGGTCCAATATG

GTTGTATTGCCGACGAGGGATCTGAGGTACATAACAGCCACTTATGATGTTTCCAGGAGCGAGCATGCAATCGTGTACTA

TATCTATGACACGGGTCGCTCATCATCTTACTTCTACCCAGTTCGATTGAATTTCAAGGGCAATCCTCTCTCTCTGAGGA

TAGAGTGTTTCCCCTGGTCTCATAAGGTGTGGTGCTACCATGATTGTCTTATATACAACACCATGACAAACGAAGAAGTC

CATACGAGAGGGCTGACCGGTATAGAGGTAACATGCAATCCAGTCTGAGTCGAGCTGAAACCATCGCTCAAGCAGGCTTT

CGAGCCATCCCCTAGTTCAAGCAGCATAGTCTGGGACACTCAGCAGCACAACCCAGCCAACAATGTTATAAAAAACTTAG

GAGCCAAGGTTGTAGGAGCCATGGACTCACTATCAGTCAATCAGGTTTTGTACCCTGAGGTCCATCTAGATAGCCCTATT

GTCACAAACAAACTAGTTGCTATCCTTGAATACTCGGGGATCAACCACAACTATGTTCTTGAGGACCAGACTCTTATCAA

GAATATTAGATATAGATTGGGGTGCGGTTTCTCAAATCAAATGATCATCAATAATAGGGGGGTAGGTGAAACAGTCAATT

CCAAGCTTAAAAGTTACCCCCGTAACTGTCATATCATATACCCAGACTGCAATAAGGAGTTGTTTTGTATCAAAGATAGC

TGCATATCTAGAAAGCTCTCGGAGCTATTCAAGAAGGGTAATTCCTTGTACTCTAAGGTAAGTCACCAGGTACTGGATTG

TCTTAAGAGAGTCAACGGGAAATTAGGGCTGGGCACAGATCTTAATCACGGCCTGAAGGATGGTATCCTCGACTTGGGGT

TGCACATGCATAGCTCTCAGTGGTTCGAGACCTTTCTGTTCTGGTTCACTATCAAGACAGAGATGAGATCAATGATCAAA

GAACAGTCCCATATATGCCACAAGAGGAGGTATAACCCAATTTTTGTGTCGGGGGATGCATTCGAGGTGCTTGTATCACG

AGACCTCGTAGTGATAATTGATAAGAACACCCAGTATGTCTTCTACCTGACGTTTGAGCTGGTCCTTATGTATTGTGATG

TCATAGAGGGTAGACTTATGACGGAGACAGCCATGGCTATAGACCAGAGATATTCAGAGCTTCTAAACCGGGTCAGATAC

TTGTGGGACCTTATCGATGGGTTCTTCCCAACACTGGGTAACACCACATACCAAGTTGTTGCTCTGCTTGAACCACTGTC

GTTGGCTTATCTTCAACTTCAGGATGTCACTCTAGAGTTAAGAGGTGCCTTTTTGGACCACTGCTTTAAAGAACTTTATG

AGATACTGGAGCATTGTGGCATTGACACGGAAGGTACCTACAATTCCATCACTGAGGGATTGGATTACGTATTTATCACT

CACGATATACACTTAACTGGGGAGATTTTTTCATTTTTCCGGAGTTTCGGACACCCCCGCCTCGAAGCGGTCACCGCTGC

AGAGAATGTCAGAAAACATATGAACCAACCGAAGGTAATCAGCTATGAGACTATGATGAAAGGGCATGCAGTATTTTGCG

GGATAATCATAAATGGTTTTAGGGACCGGCACGGCGGCAGCTGGCCCCCTGTTGCATTGCCAGAACATGCTTCTGCTGCG

ATCCGGAATGCGCAGGCATCCGGCGAAGGACTGACCCACGATCTGTGTATAGACAACTGGAAGTCCTTTGTAGGATTCAG

ATTTGGCTGCTTCATGCCGCTCAGCCTAGATAGTGATTTGACCATGTACCTCAAAGACAAAGCATTGGCTGCATTGAAGA

ATGAGTGGGATTCAGTTTACCCGAAAGAATACCTCCGTTATAATCCACCTAGAGGGACAGAGTCAAGGCGACTAGTAGAG

GTGTTCCTGAATGACTCCAGCTTTGATCCTTATAACATGATAATGTACGTGGTGAATGGCTCCTACCTTAAAGACCCTGA

GTTTAATCTCTCATACAGCTTAAAAGAAAAGGAGATAAAAGAGACAGGGCGGTTGTTTGCCAAGATGACCTACAAGATGC

GGGCCTGTCAGGTGATTGCTGAAAATCTGATATCAAATGGTGTTGGGAAGTATTTCCGAGACAATGGGATGGCAAAAGAC

GAGCATGACCTAACAAAAGCCCTTCACACTCTGGCAGTCTCAGGCGTTCCCAAGAACAACAAAGACAACCACCGAGGTGG

GCCCCCCAGAAGGAACGCAAGCCGAGAGATGAGATCAAGCCAAGCCGCCAACACACAAAATAGGGACAAGATCCAAGGGG

GCCCCATGTACAACTACTTGCGATGCCAACCAACCAGCCCTGATCAGGGTGAGTCATACGAGACTGTTAGTGCATTCATC

ACCGCTGACCTTAAGAAGTATTGCCTAAATTGGAGATACGAGACAATCAGCATATTTGCACAGAGACTGAATGAAATATA

TGGGTTGCCATCCTTCTTTCAATGGTTACATAAGGTATTGGAAAAATCCGTGCTCTACGTCAGTGATCCGCATTGCCCTC

CCGACTTAGATGATCATATCCCTCTGGACAGTGTCCCTAATGCCCAAATATTCATCAAGTACCCAATGGGCGGAATAGAA

GGTTATTGTCAAAAACTATGGACAATCAGTACTATACCATACTTGTATCTAGCAGCCTATGAGAGTGGAGTAAGAATCGC

CTCACTAGTGCAAGGTGACAATCAGACAATTGCAGTGACAAAAAGAGTTCCAAGTTCTTGGCCTTATTCACTAAAAAAGA

GGGAGGCATCTAAAGCAGCTCAAAATTACTTTGTGGTCTTAAGGCAAAGGTTGCACGATGTAGGTCATCACTTAAAGGCT

AATGAGACCATAGTATCTTCTCACTTTTTTGTATACTCTAAAGGGATCTATTATGACGGCCTGTTAGTTTCACAATCACT

AAAGAGCATCGCCAGATGTGTCTTCTGGTCCGAGACTATCGTGGATGAAACTAGAGCGGCCTGCAGCAATATTGCAACAA

CTATCGCTAAGAGTATAGAGAGGGGTTATGATAGGTACCTCGCATACTCTTTGAATATCCTCAAAATTTTCCAACAGATC

CTTGTATCCCTTGACTTCACGATTAATACAACAATGACTCAAGATGTCGTGGCACCGATCATCGAGAACGGTGATTTACT

GATAAGGATGGCACTCTTGCCAGCACCCATTGGGGGTCTCAATTATCTCAACATGAGCAGGTTGTTTGTGAGAAATATCG

GTGACCCGGTCACTTCCTCCATAGCCGACCTGAAGAGGATGATAGACGCTGGGCTAATGCCAGAAGAAACATTGCATCAA

GTGATGACCCAGACCCCGGGAGAATCATCCTACCTTGATTGGGCAAGTGATCCTTATTCTGCCAACCTACCCTGCGTACA

GAGTATAACTCGACTTCTCAAGAACATCACTGCACGGTATATTTTAATCAGCAGCCCAAATCCGATGCTGAAAGGGTTGT

TCCACGAGGGGAGCAGAGATGAAGACGAGGAGCTTGCGAGTTTCTTAATGGATCGGCATATAATTGTTCCGAGAGCTGCA

CATGAAATCTTAGACCACAGCATAACCGGGGCAAGAGAAGCTATAGCCGGGATGTTGGACACCACCAAGGGTCTGATTAG

AACAAGTATGAAACGGGGTGGCCTCACCCCTCGAGTATTAGCCCGCCTTTCCAATTATGATTATGAACAATTCAGATCCG

GGATAACACTATTGACAAAGAAAGGGCAGTGTTATCTCATTGACAAGGACTCGTGCTCGGTGCAGCTCGCTATAGCTCTG

AGGAGCCATATGTGGGCAAGGTTGGCTCGCGGGAGACCTATCTATGGCTTGGAGGTGCCTGATGTACTGGAATCGATGAA

CGGCTACCTTATCAAACGTCACGAGTCTTGTGCCATCTGTGAAACGGGCTCAAGTCACTACGGGTGGTTTTTCGTCCCTG

CAGGGTGTCAGCTTGACGATGTCTCAAGAGAGACTTCAGCTCTTCGTGTGCCTTATGTCGGATCAACCACTGAGGAAAGG

ACAGATATGAAACTTGCTTTTGTTAGATCTCCAAGCCGATCCCTCAAATCAGCAGTCAGGATTGCCACAGTCTACTCATG

GGCCTACGGGGATGATGAGAAATCATGGAGTGAAGCTTGGATGCTAGCTAGGCAGAGAGCCGATATCACCTTAGATGAAC

TGAGAATGATCACTCCAGTCTCTACATCCACCAACCTAGCTCATCGGTTAAGGGATCGGAGCACTCAGGTGAAATATTCG

GGGACATCCCTTGTGAGGGTTGCAAGGTACACAACCATCTCCAATGACAATTTGTCATTTGTAATATCTGAGAAAAAAGT

AGATACCAACTTCATTTACCAGCAAGGGATGCTGCTCGGTCTTGGGATCCTTGAGAATCTATTCAGGTTAGAGGCCACCA

CAGGGGTATCCAACACAGTGCTACACCTGCACGTGGAAACAGAATGTTGTGTTGTACCTATGGTTGATCACCCAAGGATA

CCGAGTCTCCGCAATATCAAAGTTACGAACGAGCTATGCACAAACCCTCTTATCTACGACAAGTCCCCTATCATAGAACA

CGATGCAACTCGATTATACTCACAAAGCCACAGGAGACATTTGGTGGAGTTTGTTACCTGGTCAACAAGCCAGCTCTATC

ATATACTGGCCAAATCTACAGCAATGTCCATGATTGAGCTGATCACAAGATTTGAGAAAGATCACATGAATGAAATAGCC

GCTCTGATTGGCGATGACGACATCAACAGTTTCATCACAGAATTTTTGCTTGTGGAGCCCAGACTGTTTATAGTTTACCT

CGGCCAGTGTGCTGCCATCAATTGGGCTTTTGATATACATTATCATCGGCCCTCGGGCAAGTACCAGATGGGGGAACTCC

TCTACTCTTTACTCTCTCGGATGAGCAAGGGAGTATATAAGATCTTCACTAATGCTCTGAGCCACCCCAAAGTTTACAAG

AAATTTTGGCGAAGTGGTATAATTGAGCCGGTTCATGGCCCATCCCTAGATACACAGAATTTACATGTCACTGTCTGTGA

CATGATATACGGATCATACGTCACCTATCTGGATCTTTTGCTGAATGATGAGCTAGATGCTTACCCGTATTTGCTCTGCG

AGAGTGATGAGGACGTGGTCACAGATAGGTTCGACAACATTCAAGCCAAACATCTCTGTGTACTGGCCGATGTATACTGC

AGCTCCAAGAGGTGTCCCTCGATAATCGGGATGTCCCCTATAGAAAAATGTACCATCCTCACACATTACATCAAGGGAGA

ATCGGTACAATCCCCGTCCGGGATCTCATGGAACACTGATCCCCTTGTAGTAGATCATTACTCATGCTCTCTGACCTACC

TTCGCCGTGGTTCCATCAAACAAATCAGATTGAGAGTGGATCCTGGGTTTGTGTTCGAGGCGTTGACAGACATCGACTTC

AAACAGCCTCGCAAGGCTAAGTTGGACGTATCGATTGTGGGATTGACTGATTTTTCTCCCCCTTGGGATAACGTCGGTGA

TTTTCTAGGGACTATCAACACATTGAGGCACAATCTGCCCGTCACCGGGACCGGGGTCTCGAACTATGAAGTCCACGCTT

ATCGTAGAGTTGGTCTGAATTCATCAGCATGTTATAAAGCTGTAGAGATCTCCACGTTAATAAAGGCATCCTTAGAAGCC

GGGGAGAATGGATTGTTCTTAGGAGAAGGCTCCGGCTCGATGCTGGCTGCGTACAAGGAAGTTCTTAAATTGGCAAACTG

TTATTACAACAGTGGGGTAACAGCAGAGGGTAGAGCCGGACAGAGGGAAATCTCTCCCTATCCCTCAGAGGTGAGCCTGG

TAGAGAGTCAGATGGGGATAGAGAGAAGTGTTAAAGTCCTGTTCAATGGCAAACCTGAAGTAACCTGGGTAGGGACCACC

GATTGCTACAAGTATATAATCAGTAACATTCAGACCTCAAGTCTGGGTTTCATACACTCAGATATCGAGACACTCCCAAC

CAAGGATGCTGTTGAGAAGTTGGAGGAATTTGCCTCTATTCTATCTCTATCCCTAATTTTGGGAAAAATCGGCTCTATTA

CAGTTGTCAAAATTATGCCCATTAGCGGAGATTTTACCCAAGGCTTCATAGCCTATGCCATTCAATATTTCAGAGAGAGC

CTGCTTGCCTATCCGAGATATAGTAACTTCATCTCGACTGAGTGTTACCTTATTATGATAGGATTAAAGGCCAATCGGTT

GATAAACCCAGAAGCCATTAAGCAAAGCATAATCAGGGTGGGGACTAGGACTGCACCAGGACTTGTGAGCCACATATTAT

CAGAGAAACAGAAAGGTTGTATTCAATCTTTTCTGGGTGATCCTTATGTCCAAGGAGACTTCAATAAGCACCTTAAAGCT

CTAACTCCTATTGAGAAAATCCTTGTAAATTGTGGTCTCTCGATCAATGGTACAAAAATCTGTAGGGATCTAATCCACCA

TGATATTGCCTCCGGTCCAGACGGTCTGATGAGCTCCACAATTATTCTATACAGGGAACTGGCCCATTTCAAAGACAATA

TAAGAAGTCAGCACGGCATGTTCCACCCCTACCCAGTATTGGCCAGTAGCAGGCAACGTGAATTGATCCTTCGAATAGCC

AAGAAGTTTTGGGGGTATGTCTTGCTATATTCTGATGACCCGGCACTAATCAAACAAACGATCAAGAACTTGAAGCGGAA

TCACCTAACCTTTGACTTACACAGTAATCCGTTTATTAAGGGCTTATCCAAAGCTGAGAAACTGCTAGTGCGGACAAGTT

CACTCAGAAGGGAATGGTTGTTCACTCTCGATACGAAAGAAGTGAAAGAGTGGTTCAAATTGGTGGGTTACAGTGCACTC

GTCAGAGGTTAATTAGCGATACATCTGCCCCCTTCTCCTCCACCATGAGACTCTACTGGCAATCCAAAAGATTAAAGAAA

ACTACATATTGGATAAGTATCTATTCCCAGCTTTGTCTGGT

>KU236379.1/goat/Liberia/2015-07-08

ACCAAACAAAGTTGGGTAAGGATAGATCTTACAATGACTATAGACTAGCAAACTTAGGAGTAAAGATCCTACTGTCGGGG

AGAGGAGGAGGAGCAAGATCTTTGACCATGGCGACTCTCCTTAAAAGCTTAGCACTGTTCAAGAGGAACAAAGACAAGGC

GCCTACTGCGTCAGGTTCAGGAGGGGCCATCCGGGGGATTAAGAATGTTATCATAGTCCCTATCCCCGGGGACTCATCCA

TCATTACCCGTTCAAGACTGCTCGACAGGCTTGTCAGACTGGCCGGAGACCCTGACATCAACGGGTCAAAGTTGACCGGC

GTGATGATCAGCATGTTATCTTTGTTCGTGGAGTCACCTGGACAATTGATACAGCGGATCACAGATGATCCAGATGTTAG

CATCCGTCTTGTTGAGGTAGTTCAAAGTACCAGGTCCCAGTCCGGATTGACCTTTGCATCACGTGGTGCTGATTTGGACA

ATGAGGCAGATATGTATTTTTCAACTGAAGGGCCCTCGAGTGGAAGTAAGAAAAGGATCAACTGGTTTGAGAACAGAGAA

ATAATAGACATAGAAGTGCAAGATGCAGAAGAGTTCAATATGTTGTTAGCCTCCATCTTAGCACAAGTTTGGATCCTCCT

GGCCAAGGCGGTTACGGCACCAGATACAGCAGCCGACTCAGAACTGAGAAGGTGGGTTAAATACACACAACAAAGGAGAG

TGATTGGAGAATTTCGCTTTGACAAAGGGTGGCTGGACGCAGTCCGTAACAGGATTGCAGAAGATCTATCACTCCGGCGG

TTCATGGTATCCCTGATACTTGACATCAAGAGGACCCCCGGCAACAAGCCAAGGATTGCAGAAATGATCTGCGACATTGA

CAACTATATTGTCGAGGCCGGACTCGCAAGTTTCATTCTTACTATCAAGTTTGGTATTGAAACTATGTATCCTGCACTAG

GCCTTCACGAGTTCGCTGGGGAATTGTCCACTATTGAATCCTTGATGAACTTGTATCAACAGCTAGGAGAAGTTGCACCC

TATATGGTAATTCTAGAGAACTCAATTCAGAACAAGTTTAGTGCAGGAGCCTATCCTCTCCTCTGGAGCTATGCGATGGG

TGTCGGAGTCGAGTTGGAGAACTCAATGGGGGGCTTGAACTTTGGCAGGTCATATTTTGACCCGGCCTATTTTCGTCTCG

GACAGGAGATGGTCAGAAGATCTGCAGGAAGGGTCAGCTCTGTAATTGCGGCTGAGCTTGGCATCACAGCAGAGGAAGCC

AAACTGGTCTCGGAAATCGCCTCGCAGACTGGGGAAGAACGAACAGTTAGAGGGACTGGGCCTCGACAGGCGCAGGTTTC

CTTCCTCCAGCACAAAATAGGTGAGGGAGAGTCGCCTACACCAGCAACCAAAGAAGAAGTCAAAGCTGCGATCCCAAACG

GATCTGAAGGAAGGGACATAAAGCGAACACGCCCAGGGAAGCCCAGAGGAGAAACTCCCGGGCAACTGCTTCTGGAGATC

ATGCCAGAGGATGGAGTCTCGCGAGAGTCTGGTCAGAACCCTCGTGAGGCTCAAAGATCGGCTGAGGCACTCTTCAGGCT

GCAGGCCATGGCCAAAATTTTGGAGGACCAGGAGGAGGGAGAAGACAACAGTCAGATCTACAACGACAAGGATCTCCTCA

GCTGAGCAGATACACCCTCTGTCCAAATCAGTGACAAGACATCACCTGCCAGTATTATAAAAAACTTAGGACCCAGGTCC

AACCAACCGCACATCGACACCCCAGTCAATCGAGCGGAGACCACCGATGGCAGAAGAACAAGCATACCATGTCAACAAGG

GGCTGGAATGTATCAAGTCCCTCAAAGCCTCTCCCCCGGATCTATCCACCATCCAAGATGCCCTTGAGAGCTGGAGAGAG

GGGTTCAACCCCTCAGGCCGTGCAACACCGAACCATGATACGTCCAAGGGGGACCATCAGAATATCAACCAATCATGCTC

ATCAGCAATCGGATCAGACAAAGTCGACATGTCTCCTGAAGGTAATCTCGGATTTAGAGAGATCACTTGTGATGACAATG

AGGCTGGGCTCGGAGGAGTTCAGGACAAAGGATCCGACTCTCAAGTACAGCGTTACCATGTTTATAACCACGGGGGTGAA

GAGATTGAAGGACTCGAGGATGCTGACTCTCTCGTGGTTCAAGCAAATCCTCCGGTTGCTCACACCTTCGATGGAGGAGA

GGATGGATCTGACAACAGCGATGTGGACTTTGGCCCAGATGATCCCGACAGAGATCCTCTATATGACCGGGGACCTGCTG

CCGGCAATGATGTCTCTAGGTCAACAGATGTCGAAAAATTAGAAGGTGATGACATTCAAGAAGTTCTTAACTCCCAGAAG

AGTAAAGGAGGAAGATTCCAGGGCGGGAAAATCTTGCGAGTCCCGGAAATACCCGATGTCAAGAACTCCAGACCATCGGC

CCAATCAATTAAAAAGGGCACAGACGGGAGCTCAGTCTTATCTGGAATGGTGACAGAGTGTTCATCGATAAGTGGTGCAA

CCCAAGCTGTGCTAGAGTCCAGATGGGAGTCATCAGAGCGAAATGCATCTGTGGGGAGTGTCCCCAAATCTGTGAGGAGT

GCAAAGACGATCCAGGGGTTAACACAAGAATCTGGTACCATAGCATCACTGACTCAGCCTAAAGAAAATGACTCCGAGTA

TGAGTATGAGGATGATCTATTCGCTGAGATGCAGGACATTCGTGCAAGCATTGCTAAGATCCATGATGACAACAAAACTA

TCCTCTCAAAACTTGATTCTCTACTGTTATTGAAAGGAGAAATCGATACTATCAAGAAACAAATCAGCAAACAAAATATA

AGTATATCTACCATTGAGGGCCATCTATCCAGTATAATGATAGCCATCCCGGGTTTTGGGAAGGACATCAAGGACCCAAC

ATCTGAGGTTGAGCTGAACCCAGATTTGAGACCTATAATCAGCCGCGATTCTGGCAGGGCTCTTGCGGAGGTCCTTAAGA

AACCCGCTGTTGATAGGTCTCAGAAGAGCGGAATCAAAGTCAACTCCGGTTCAAAGGGTCAGCTCCTCAAGGATCTCCAG

CTAAAACCCGTCGACAAGCAGGCAAGCTCTGCAATCGGGTTTGTTCCATCCGACCATGAATCATCCAGAAGTGTCATCCG

CTCCATAATCAAGTCGAGCAAGCTTAACATTGATCACAAAGACTATCTTCTGGATTTACTGAATGATGTGAAAGGCTCTA

AGGATCTCAAGGAATTCCACAAGATGCTAACAGCCATTCTTGCCAAGCAGCCGTAACACATCCCACAATCTACATCTTAT

ACTCGGTTGAAAGCATCCTCTCAATCAGGCTATTACAAAAAACTTAGGAGCAAGGGCAACTGAGCTTCACAGACAGGATG

ACCGAGATTTACGATTTCGATAAATCAGCATGGGATGTCAAAGGGTCAATTGCTCGCATAGAACCCACCACCTATCACGA

CGGCCGACTGGTACCCCAGGTGAGGGTCATCGACCCCGGTCTGGGAGACAGAAAAGATGAATGCTTCATGTACCTGTTTC

TTCTAGGAGTGATTGAGGATAACGACCCACTTTCTCCCCCAGTCGGGAGAACCTTCGGGTCTTTACCCCTAGGTGTCGGC

AGGTCAACTGCTAAGCCAGAGGAACTACTAAGGGAGGCCACTGAACTAGATATAGTGGTGAGGCGCACGGCAGGACTAAA

TGAGAAACTGGTATTTTACAACAACACTCCGCTATCTTTGTTAACACCCTGGAGGAAAGTCTTGACGACCGGAAGTGTGT

TCAGCGCCAACCAGGTTTGCAATGCGGTCAACCTAGTCCCACTCGATACTCCCCAAAGATTCAGGGTTGTGTACATGAGT

ATAACTAGATTGTCAGATAATGGTTATTATAGTGTCCCCAGAAGAATGTTGGAGTTCCGCTCAGCCAATGCAGTCGCTTT

CAATATCTTGGTTACACTGAGAATTGAAAATGGCACAAACCCTAGAAGATACATAGTCGGCTCATGGGAGAATCCAGAAG

TCACATTTATGGTACACGTGGGCAACTTTAGAAGAAAGAAGAACGAAGTATACTCTGCTGATTATTGCAAAATGAAGATT

GAAAAGATGGGTCTAGTTTTTGCCTTGGGAGGAATAGGTGGAACAAGTCTCCATATTCGAAGCACAGGGAAAATGAGCAA

GACCCTCCATGCACAGCTGGGGTTCAAGAAAATTCTATGCTACCCTCTGATGGATATCAATGAGGATCTTAACCGATATC

TCTGGCGGGCAGAGTGCCGAATAGTCAAAATCCAGGCCGTCTTACAGCCATCAGTACCCCAAGAATTCCGTGTCTATGAT

GATGTCATCATCAACGATGACCAAGGCCTGTTCAAGATCCTGTAAGTCACCTGCAGGGCCATGGACCCACCTCCAGGAAG

AGGCCCTCCCACCTCCCGCCCAAAACACACACCCCAAGACTCCCCGAAACAGCGCCGGCAAAACCACTCTCCCGGCGGGA

CGAGACCGCCC-CCCCCAACAAACCCGCTCCGGAGGGGGG-CCCCCCCCGCGACCCGCCGGGCGCCGACCGCGCGAGCCG

GCGAGACACCCCCAAGAGGCCTCGACGAAGACCGGGGAACCCCTTCCCCA-AAAATGGCCCCCC-AACACACCCAAGGCC

ATGCGCCTAGCACCCCCAGTCCCCCAACCGGCGGGACAGGAGGCCAGGAAGGGGTCCCCCAAAACAAAGGGACCACCAGG

GAAGGCTCCGCCGACCCAGCACAGACCCGGCCCAAACAAAGCAGACCCCCAAGACGAAATCTCCCAGAGCACCCCATTCA

TCATCAAACAAAACTTAGGGGCCAAGTCCAGACGACAGACCCCGGCCCGCCAGGCAGCAAGCCACAGAAAAGAAGGAGAC

AAGATCGGCCCGGGGAGAGGACACCCAACCACAGAAACCCCCAGCTCCAGGGGAAGAG-CCCCCCCTCCACACGACGGAG

GAACGGGCGAGAGACACCGAGGAGAACACGATCGGGAGCCGAGAGCCGGAGACGCCGGAGGCAAACAGGCCGGGCACTCA

ACCGACACCACCGGAGGACCCGAGGAGAGCCCGACTCCGCGCAGACTCGACGGCAGAGGACCTAGTCCCGCCATAAAGAG

AGGTGAGAACCCT--------CCCTCCTGCTCACAACCGGCAAACAGACCGCGGGAGGACCACCTCCCAGCAAGACAGAC

GTCTCCCCTGCAGCCGAGCCGCAGCTCCTCAGAAGAGGGCACCCCCGAAACCCCCCAGAACCCCTCCGCCCACGGCCGCC

GGGACCAGAGCTCCGCCGCCAAAGATCAGAGCCGCCGAGAGTCAACCGGGGGTCCCCCCCAGTCCCTCCCGGCGATCCGC

CACAGAGGCCCCAGCAAGGTCCCCCGGCCCGACGACCCCTGCATGCGCCGAAAGGGAAACCCACCAGAGGACGTCCAGCA

CATCCGTGTATAAACATCATGACACGGGTCGCAACCTTGGTATTGCTGTTTCTTTTCCCAAACACTGTCGCGTGCCAGAT

TCACTGGGGCAATCTATCCAAGATAGGGATTGTCGGAACAGGGAGTGCCAGTTACAAGGTGATGACCAGACCAAGCCACC

AAACTCTGGTTATAAAGTTGATGCCAAATATAACAGCCATCGACAATTGTACGAAATCAGAGATTTCAGAGTACAAAAGA

TTGCTAATCACAGTGTTAAAGCCTGTAGAGGATGCTTTGTCAGTAATAACCAAGAATGTAAGACCAATTCAAGCTCTAAC

ACCTGGGCGCAGGACCCGCCGTTTCGCCGGAGCTGTTCTGGCCGGAGTAGCACTTGGAGTCGCGACGGCTGCTCAAATAA

CTGCCGGAGTCGCACTCCATCAGTCATTGATGAATTCCCAAGCAATTGAAAGTTTAAAAACCAGTCTTGAGAAGTCAAAT

CAGGCAATAGAAGAAATCAGACTTGCAAATAAGGAGACCATACTGGCAGTACAGGGCGTCCAAGATTATATCAACAACGA

GCTTGTCCCCTCTGTTCATAGAATGTCATGTGAGCTAATAGGTCACAAACTCAGTCTCAAACTCCTTAGGTATTATACCG

AGATCTTGTCTATATTCGGGCCTAGCCTTCGAGACCCGATAGCTGCTGAAATATCAATCCAGGCACTCAGCTATGCACTA

GGCGGAGACATCAATAAAATTCTGGACAAGCTTGGGTATAGCGGCGGGGATTTCCTTGCTATTCTAGAAAGCAAGGGGAT

AAAGGCCCGGGTCACATATGTGGACACAAGAGATTACTTTATAATTCTTAGCATAGCCTACCCAACCTTATCTGAGATCA

AAGGGGTGATAGTTCATAAGATAGAAGCTATATCATACAACATTGGGGCACAGGAATGGTATACTACTATCCCTAAATAT

GTAGCCACTCAGGGATATTTGATATCGAATTTCGATGAGACATCATGTGTATTCACTCCAGAGGGGACAGTCTGCAGCCA

GAATGCGCTGTATCCAATGAGCCCATTGCTTCAGGAATGTTTCAGGGGGTCGACAAAATCGTGTGCCAGAACCCTAGTTT

CAGGGACCACAAGTAATAGATTTATCCTATCAAAAGGGAACTTGATTGCAAATTGTGCGTCAGTTTTGTGCAAGTGTTAC

ACAACGGAGACAGTCATCAACCAAGATCCTGATAAACTACTAACTGTTATAGCCTCAGATAAGTGTCCCGTAGTTGAGGT

GGATGGAGTGACAATACAAGTCGGCAGTCGAGAGTACCCAGATTCTGTATACCTACATAAAATAGACTTAGGCCCAGCCA

TCTCTCTGGAAAAACTGGATGTAGGCACCAATTTAGGCAATGCAGTCACAAGACTGGAGAATGCAAAGGAGCTCCTAGAC

GCATCAGACCAGGTACTGAAGACTGTCAAAGGGGTACCCTTCAGTGGCAATATGTACATAGCACTGGCAGCTTGCATTGG

GGTATCCCTAGGGCTTGTCACATTAATATGCTGCTGTAAGGGGAGATGTAGGAACAAGGAGATTCCTGCCTCCAAAATCA

ACCCAGGGCTCAAACCCGACCTGACCGGGACTTCAAAGTCTTACGTGAGATCACTGTAGTCAGAGTAACCCGAATCACCC

GGCATCACGCGTATACATGTGTGACACCAGCAGTCAGAGGACGCAGAAGACTCAACCTCCGATCACCGACCAGACCCCAC

TCTACGCCCTATTACACATTGGTCATCAAACAAAACTTAGGACGAAAGGTCAATCACCATGTCCGCACAAAAGGAGAGGA

TCAATGCCTTCTACAAAGACAACCCTCACAATAAAAACCATAGGATAATCCTGGATAGGGAACGCTTAACTATTGAAAGA

CCCTACATCTTACTTGGAGTCCTGCTGGTAATGTTCCTGAGTCTAATCGGGCTGCTAGCCATTGCAGGGATCAGGCTTCA

CCGGGCCACCGTTGGAACTGCGGAGATCCAGAGTCGGCTGAATACCAACATTGAGTTGACCGAATCCATTGATCATCAAA

CTAAGGATGTCTTAACACCCCTGTTCAAAATCATAGGTGATGAAGTCGGTATCAGAATTCCACAGAAGTTCAGTGATCTT

GTCAAGTTCATCTCCGATAAGATTAAGTTCCTCAACCCTGACAGAGAATACGATTTTAGGGATCTCCGGTGGTGTATGAA

CCCCCCTGAGAGAGTCAAAATTAACTTTGATCAATTCTGTGAATACAAAGCCACAGTCAAGTCAGTTGAACATATATTTG

AGTCATCATTCAACAGGTCAGAAAGATTGCGACTATTGACTCTTGGGCCCGGAACAGGCTGTCTCGGCAGGACAGTAACA

AGAGCTCAGTTCTCAGAACTTACGCTGACCCTGATGGACCTGGATCTCGAGATGAAGCACAACGTGTCCTCAGTGTTTAC

CGTAATTGAAGAGGGATTATTCGGAAGAACATATATTGTTTGGAGATCTGACACCGGGAAACCGAGCACCAGTCTAGATA

TTGGCCAGTTTTTAAGAGTTTTTGAGATCGGGTTGGTGAGGGATCTCGAGCTGGGTGCCCCCATTTTCCATATGACCAAC

TACCTCACAGTGAACATGAGTGATGACTATCGGAATTGCCTTTTAGCGGTAGGGGAGTTGAAGCTGACAGCCCTATGCAC

CCCATCTGAGACTGTGACTCTGAGTGAGAGAGGAGTTCCAAAGAGAGAGCCTCTTGTGGTTGTGATACTCAACCTAGTTG

GGCCTACTCTAGGGGGCGAACTATACAGTGTCTTGCCTACCTCTGACCTCATGGTGGAGAAACTCCATCTATCCTCACAC

AGAGGGATCATCAAAGACAACGAGGCCAATTGGGTAGTACCGTCTACCGATGTTCGTGATCTCCAAAACAAAGGAGAATG

TCTGGTGGAAGCATGCAAAACCCGACCTCCTTCATTTTGCAATGGCACAGGAATAGGCCCATGGTCAGAGGGGAGAATCC

CTGCCTACGGAGTGATCAGGGTCAGTCTTGACTTAGCTAGTGACCCAGGTGTGGTTATCACTTCAGTGTTTGGCCCACTG

ATACCTCACCTATCTGGCATGGATCTTTACAACAATCCGTTTTCAAGAGCTGCATGGCTGGCTGTGCCACCCTACGAGCA

GTCATTTCTAGGAATGATAAATACAATTGGCTTCCCGGACAGAGCAGAGGTCATGCCGCACATTTTGACCACAGAGATCA

AAGGGCCTCGAGGTCGTTGTCATGTTCCTATAGAGTTGTCCCGCAGGATTGATGATGATATCAAGATCGGGTCCAATATG

GTTGTATTGCCGACGAGGGATCTGAGGTACATAACAGCCACTTATGATGTTTCCAGGAGCGAGCATGCAATCGTGTACTA

TATCTATGACACGGGTCGCTCATCATCTTACTTCTACCCAGTTCGATTGAATTTCAAGGGCAATCCTCTCTCTCTGAGGA

TAGAGTGTTTCCCCTGGTCTCATAAGGTGTGGTGCTACCATGATTGTCTTATATACAACACCATGACAAACGAAGAAGTC

CGTACGAGAGGGCTGACCGGTATAGAGGTAACATGCAATCCAGTCTGAGTCGAGCTGAAACCATCGCTCAAGCAGGCTTC

CGAGCCATCCCCTAGTTCAAGCAGCATAGTCTGGGACACTCAGCAGCACAACCCAGCCAACAATGTTATAAAAAACTTAG

GAGCCAAGGTTGTAGGAGCCATGGACTCACTATCAGTCAATCAGGTTTTGTACCCTGAGGTCCATCTAGATAGCCCTATT

GTCACAAACAAACTAGTTGCTATCCTTGAATACTCGGGGATCAACCACAACTATGTTCTTGAAGACCAGACTCTTATCAA

AAATATTAGATATAGACTGGGGTGCGGTTTCTCAAATCAAATGATCATCAATAATAGGGGGGTAGGTGAAACAGTCAATT

CCAAGCTTAAAAGTTACCCCCGTAACTGTCATATCATATACCCAGACTGCAATAAGGAGTTGTTTTGTATCAAAGATAGC

TGCATATCTAGAAAGCTCTCGGAGCTATTCAAGAAGGGTAATTCCTTGTACTCTAAGATAAGTCACCAGGTACTGGATTG

TCTTAAGAGAGTCAACGGGAAATTAGGGCTGGGCACAGATCTTAATCACGGCCTGAAGGATGGTATCCTCGACTTGGGGT

TGCACATGCATAGCTCTCAGTGGTTCGAGACCTTTCTGTTCTGGTTCACTATCAAGACAGAGATGAGATCAATGATCAAA

GAACAGTCCCATATATGCCACAAGAGGAGGTATAATCCAATTTTTGTGTCGGGGGATGCATTCGAGGTGCTTGTATCACG

AGACCTCGTAGTAATAATTGATAAGAACACCCAGTATGTCTTCTACCTGACGTTTGAGCTGGTCCTTATGTATTGTGATG

TCATAGAGGGTAGACTTATGACGGAGACAGCCATGGCTATAGACCAGAGATATTCAGAGCTTCTAAACCGGGTCAGATAC

TTGTGGGACCTTATCGATGGGTTCTTCCCAACTCTGGGTAACACCACATACCAAGTTGTTGCTCTGCTTGAACCACTGTC

GTTGGCTTATCTTCAACTTCAGGATGTCACTCTAGAGTTAAGAGGTGCCTTTTTGGACCACTGCTTCAAAGAACTTTATG

AGATACTGGAGCATTGTGGCATTGACACGGAAGGTACCTACAATTCTATCACTGAGGGATTGGATTACGTATTTATCACT

CACGATATACACTTAACTGGGGAGATTTTTTCATTTTTCCGGAGTTTCGGACACCCCCGCCTCGAAGCGGTCACCGCTGC

AGAGAATGTCAGAAAACATATGAACCAACCGAAGGTAATCAGTTATGAGACTATGATGAAAGGGCATGCAGTATTTTGCG

GGATAATCATAAATGGTTTTAGGGACCGGCACGGCGGCAGCTGGCCCCCTGTTGCATTGCCAGAACATGCTTCTGCTGCG

ATCCGGAATGCGCAGGCATCCGGCGAAGGACTGACCCACGACCTGTGTATAGACAACTGGAAGTCCTTTGTAGGGTTCAG

ATTTGGCTGCTTCATGCCGCTCAGCCTGGATAGTGATTTGACCATGTACCTCAAAGACAAAGCATTGGCTGCATTAAAGA

ATGAGTGGGATTCAGTTTACCCGAAAGAATACCTCCGTTATAATCCACCTAGAGGGACAGAGTCAAGGCGACTAGTAGAG

GTGTTCCTGAATGACTCCAGCTTTGATCCCTATAACATGATAATGTACGTGGTGAATGGCTCCTACCTTAAAGACCCTGA

GTTTAATCTCTCATACAGCTTAAAAGAAAAGGAGATAAAAGAGACAGGGCGGTTGTTTGCCAAGATGACCTACAAGATGC

GGGCCTGTCAGGTGATTGCTGAAAATCTGATATCAAATGGTGTTGGGAAGTATTTCCGAGACAATGGGATGGCAAAAGAC

GAGCATGACCTAACAAAAGCCCTTCACACTCTGGCAGTCTCAGGCGTTCCCAAGAACAACAAAGACAACCACCGAGGTGG

GCCCCCCAGAAGGAACGCAAGGCGAGAGATGAGATCAAGCCAAGCCGTCAACACACAAAATAGAGACAAGATCCAGGGGG

GCCCTATGTACAACTACTTGCGATGCCAACCAACCAGCCCTGATCAGGGTGAGTCATACGAGACTGTTAGTGCATTCATC

ACCGCTGACCTTAAGAAGTATTGCCTAAATTGGAGATACGAGACAATCAGCATATTTGCACAGAGACTGAATGAAATATA

TGGGTTGCCATCCTTCTTTCAATGGTTACATAAGGTATTGGAAAAATCCGTGCTCTACGTCAGTGATCCGCATTGCCCTC

CCGACTTAGATGATCATATCCCTCTGGACAGTGTCCCTAATGCCCAAATATTCATCAAGTACCCAATGGGCGGAATAGAA

GGTTATTGTCAAAAACTATGGACAATCAGTACTATACCATACTTGTATCTAGCAGCCTATGAGAGTGGAGTAAGAATCGC

CTCACTAGTGCAAGGTGACAATCAGACAATTGCAGTGACAAAAAGAGTTCCAAGTTCTTGGCCTTATTCACTAAAAAAGA

GGGAGGCATCTAAAGCAGCTCAAAATTACTTTGTGGTCTTAAGGCAAAGGTTGCACGATGTAGGTCATCACTTAAAGGCT

AATGAGACCATAGTATCTTCTCACTTTTTTGTATACTCTAAAGGGATCTATTATGACGGCCTGTTAGTTTCACAATCACT

AAAGAGCATCGCCAGATGTGTCTTCTGGTCTGAGACTATCGTGGATGAAACTAGAGCGGCCTGCAGCAATATTGCAACAA

CTATCGCTAAGAGTATAGAGAGGGGTTATGATAGGTACCTCGCATACTCTTTAAATATCCTCAAAATTCTTCAACAGATC

CTTGTATCCCTTGACTTCACGATTAATACAACAATGACTCAAGATGTCGTGGCACCGATCATCGAGAACGGTGATTTACT

GATAAGGATGGCACTCTTGCCAGCACCCATTGGGGGTCTCAATTATCTCAACATGAGCAGGTTGTTTGTGAGAAATATCG

GGGACCCGGTCACTTCCTCCATAGCCGACCTGAAGAGGATGATAGACGCTGGGCTAATGCCAGAAGAAACATTGCATCAA

GTGATGACCCAGACCCCGGGAGAATCATCCTACCTTGATTGGGCAAGTGATCCTTATTCTGCTAACCTACCCTGCGTACA

GAGTATAACTCGCCTTCTCAAAAACATCACTGCACGGTATATTTTAATCAGCAGCCCAAATCCGATGCTGAAAGGGTTGT

TTCATGAGGGGAGTAGAGATGAAGACGAGGAGCTTGCGAGTTTCTTAATGGATCGGCATATAATTATTCCGAGAGCTGCA

CATGAAATCTTAGACCACAGCATAACCGGGGCAAGGGAAGCTATAGCCGGGATGTTGGACACCACCAAGGGTCTGATTAG

AACAAGTATGAAACGGGGTGGCCTCACCCCTCGAGTATTAGCCCGCCTTTCCAATTATGATTATGAACAATTCAGATCCG

GAATAACACTATTGACAAAGAAAGGGCAGTGTTATCTCATTGACAAGGACTCGTGCTCGGTGCAGCTCGCTATAGCTCTG

AGGAGCCATATGTGGGCAAGGTTGGCTCGCGGGAGACCTATCTATGGCTTGGAGGTGCCTGATGTACTGGAATCGATGAA

CGGCTACCTTATCAAACGTCACGAGTCTTGTGCCATCTGTGAAACGGGCTCAAGTCACTACGGGTGGTTTTTCGTCCCTG

CAGGGTGTCAGCTTGACGATGTCTCAAGAGAGACTTCAGCTCTTCGTGTGCCTTATGTCGGATCAACCACTGAGGAAAGG

ACAGATATGAAACTTGCTTTCGTTAGATCTCCAAGCCGATCCCTCAAATCAGCAGTCAGGATTGCCACAGTCTACTCATG

GGCCTACGGGGATGATGAGAAATCATGGAGTGAAGCTTGGATGCTAGCTAGGCAGAGAGCCGATATCACCTTAGATGAAC

TGAGAATGATCACTCCAGTCTCTACATCCACCAACCTAGCTCATCGGTTAAGGGATCGGAGCACTCAGGTGAAATATTCG

GGGACATCCCTTGTGAGGGTTGCAAGGTACACAACCATCTCCAATGACAATTTGTCATTTGTGATATCTGAGAAAAAAGT

AGATACCAACTTCATTTACCAGCAAGGGATGCTGCTCGGTCTTGGGATCCTTGAGAATCTTTTCAGGTTAGAGGCCACCA

CAGGGGTATCCAACACAGTGCTACACCTGCACGTGGAAACAGAATGTTGTGTTGTACCTATGGTTGACCACCCAAGGATA

CCGAGTCTCCGCAATATCAAAGTTACGAACGAGCTATGCACAAACCCTCTTATCTACGACAAGTCCCCCATCAAAGAACA

CGATGCAACTCGATTATACTCACAAAGCCACAGGAGACATTTGGTGGAGTTTGTTACCTGGTCAACAAGCCAGCTCTATC

ATATACTGGCCAAATCTACAGCAATGTCCATGATTGAGCTGATCACAAGATTTGAGAAAGATCACATGAATGAAATAGCC

GCTCTGATTGGCGATGACGACATCAACAGTTTCATCACAGAATTTTTGCTTGTGGAGCCCAGACTGTTTATAGTTTACCT

CGGCCAGTGTGCTGCCATCAATTGGGCTTTTGATATACATTATCATCGGCCCTCGGGCAAGTACCAGATGGGGGAACTCC

TCTACTCTTTACTCTCTCGGATGAGCAAGGGAGTATATAAGATCTTCACTAATGCTCTGAGCCACCCCAAAGTTTACAAG

AAATTCTGGCGGAGTGGTATAATTGAGCCGGTTCATGGCCCATCCCTAGATACACAGAATTTACATATCACTGTCTGTGA

CATGATATACGGATCATACGTCACCTATCTGGATCTTTTGCTGAATGATGAGCTGGATGCTTACCCGTATTTGCTCTGCG

AGAGTGATGAGGACGTGGTCACAGACAGGTTCGACAACATTCAAGCCAAACATCTCTGTGTACTGGCCGATGTATACTGC

AGCTCCAAGAGGTGTCCCTCGATAATCGGGATGTCCCCTATAGAAAAATGTACCATCCTCACACATTACATCAAGGGAGA

ATCAGTACAATCCCCGTCCGGGATCTCATGGAACACCGATCCTCTTGTAGTAGATCATTACTCATGCTCTCTGACCTACC

TCCGCCGTGGTTCCATCAAACAAATCAGATTGAGAGTGGATCCTGGGTTTGTGTTCGAGGCGTTGACAGACATCGACTTC

AAACAGCCTCGCAAGGCTAAGTTGGACGTATCGATTGTGGGATTGACTGATTTTTCTCCCCCTTGGGATAACGTCGGTGA

TTTTCTAGGGACTATCAACACATTGAGGCACAATCTGCCCGTCACCGGGACCGGGGTCTCGAACTATGAAGTCCACGCTT

ATCGTAGAATTGGTCTGAATTCATCAGCATGTTATAAAGCTGTAGAGATCTCCACGTTAATAAAGTCATCCTTAGAAGCC

GGGGAGAATGGATTGTTCTTAGGAGAAGGCTCCGGCTCGATGCTGGCTGCGTACAAGGAAGTTCTTAAATTGGCAAACTG

TTATTACAACAGTGGGGTAACAGCAGAGGGTAGAGCCGGACAGAGGGAAATCTCTCCCTATCCCTCAGAGGTGAGCCTGG

TAGAGAGTCAGATGGGGATAGAGAGAAGTGTTAAAGTCCTGTTCAATGGCAAACCTGAAGTAACCTGGGTAGGGACCACC

GATTGCTACAAGTATATAATCAGTAACATTCAGACCTCTAGTCTGGGTTTCATACACTCAGATATCGAGACACTCCCAAC

CAAGGATGCCGTTGAGAAGTTGGAGGAATTTGCCTCTATTCTATCTCTATCCCTAATTTTGGGAAAAATCGGCTCTATTA

CAGTTGTCAAAATTATGCCCATTAGCGGAGATTTTACCCAAGGCTTCATAGCCTATGCCATTCAATATTTCAGGGAGAGC

CTGCTTGCCTATCCGAGATATAGTAACTTCATCTCGACTGAGTGTTACCTTATTATGATAGGATTAAAGGCCAATCGGTT

GATAAACCCAGAAGCCATTAAGCAAAGCATAATCAGGGTGGGGACTAGGACCGCACCAGGACTTGTAAGCCACATATTAT

CAGAGAAACAGAAAGGTTGTATTCAATCTTTACTGGGTGATCCTTATGTCCAAGGAGACTTCAATAAGCACCTTAAAGCT

CTAACTCCTATTGAGAAAATCCTTGTAAATTGTGGTCTCTCGATCAATGGTACAAAAATCTGTAGGGATCTAATCCACCA

TGATATTGCCTCCGGTCCAGACGGTCTGATGAGCTCCACAATTATTTTATACAGGGAACTGGCCCATTTCAAAGACAATA

TAAGAAGTCAGCACGGCATGTTCCACCCCTACCCAGTATTGGCCAGTAGCAGGCAACGTGAATTGATCCTTCGAATAGCC

AAGAAGTTTTGGGGTTATGTCTTGCTATATTCTGATGACCCGGCACTAATCAAACAAACGATCAAGAACTTGAAGCGGAA

TCACCTAACCTTTGACTTACACAGTAATCCGTTTATTAAGGGCTTATCCAAAGCTGAGAAACTGCTAGTGCGGACAAGTT

CACTCAGAAGGGAATGGTTGTTCACTCTCGATACGAAAGAAGTAAAAGAGTGGTTCAAATTGGTGGGTTACAGTGCACTC

GTCAGAGGTTAATTCGCGATACATCTGCCCCCTTCTCCTCCACCATGAGACTCTACTGGCAATCCAAAAGATTAAAGAAA

ACTACATATTGGATAAGTATCTATTCCCAGCTTTGTCTGGT

>KR781451.1/goat/CotedIvoire/2009-07

ACCAAACAAAGTTGGGTAAGGATAGATCTTACAATGACTATAGACTAGCAAACTTAGGAGTAAAGATCCTACTGTCGGGG

AGAGGAGGAGGAGCAAGATCTTTGACCATGGCGACTCTCCTTAAAAGCTTAGCACTGTTCAAGAGGAACAAAGACAAGGC

GCCAACTGCGTCAGGTTCAGGAGGGGCCATCCGGGGGATTAAGAATGTTATCATAGTCCCTATCCCCGGGGACTCATCCA

TCATTACCCGTTCAAGACTGCTCGACAGGCTTGTCAGACTGGCCGGAGACCCTGACATCAACGGGTCAAAGTTGACCGGC

GTGATGATCAGCATGTTATCTTTGTTCGTGGAGTCACCCGGGCAATTGATACAGCGGATCACAGATGATCCAGATGTTAG

CATCCGTCTTGTTGAGGTAGTTCAAAGTACCAGGTCCCAGTCCGGATTGACCTTTGCATCACGTGGTGCTGATTTGGACA

ATGAGGCAGATATGTATTTTTCAACTGAAGGACCCTCGAGTGGAAGTAAGAAAAGGATCAACTGGTTTGAGAACAGAGAA

ATAATAGACATAGAAGTGCAAGATGCAGAAGAGTTCAATATGTTGTTAGCCTCCATCTTAGCACAAGTTTGGATCCTCCT

GGCCAAGGCGGTTACGGCACCAGATACAGCAGCCGACTCAGAACTGAGAAGGTGGGTTAAATACACACAACAAAGGAGAG

TGATTGGGGAATTTCGCTTTGACAAAGGGTGGCTGGACGCAGTCCGTAACAGGATTGCAGAAGATCTATCACTCCGGCGG

TTCATGGTATCCCTGATACTTGACATCAAGAGGACCCCCGGCAACAAGCCAAGGATTGCAGAAATGATCTGCGACATTGA

CAACTATATTGTCGAGGCCGGACTCGCAAGTTTCATTCTTACTATCAAGTTCGGTATTGAAACTATGTACCCTGCACTAG

GCCTTCACGAGTTCGCTGGGGAATTGTCCACTATTGAATCCTTGATGAACTTGTATCAACAGCTAGGAGAAGTTGCACCC

TACATGGTAATTCTAGAGAACTCAATTCAGAACAAGTTTAGTGCAGGAGCCTATCCTCTCCTCTGGAGCTATGCGATGGG

TGTCGGAGTCGAGCTGGAGAACTCAATGGGGGGCTTGAACTTTGGCAGGTCATATTTTGACCCGGCCTATTTTCGTCTCG

GACAGGAGATGGTCAGAAGATCTGCAGGAAGGGTCAGCTCTGTAATTGCGGCTGAGCTTGGCATCACAGCAGAGGAAGCC

AAACTGGTCTCGGAAATCGCCTCGCAGACTGGGGAAGAACGAACAGTTAGAGGGACTGGGCCTCGACAGGCGCAGGTTTC

CTTCCTCCAGCACAAAATAGGTGAGGGAGAGTCGCCTACACCAGCAACCAAAGAAGAAGTCAAAGCTGCGATCCCAAACG

GATCTGAAGGAAGGGACATAAAGCGAACACGCCCAGGGAAGCCCAGAGGAGAAACTCCCGGGCAACTGCTTCTGGAGATC

ATGCCAGAGGATGGAGTCTCGCGAGAGTCTGGTCAAAACCCTCGTGAGGCTCAAAGATCGGCTGAGGCACTTTTCAGGCT

GCAGGCCATGGCCAAAATTTTGGAGGACCAGGAGGAGGGAGAAGACAACAGTCAGATCTACAACGACAAGGATCTCCTCA

GCTGAGCAGATACACCCTCTGCCCAAATCAGTGACAAGACATCACCTGCCAGTATTATAAAAAACTTAGGACCCAGGTCC

AACCAACCGCACATCGACACCCCAGTCAATCGAGCGGAGACCACCGATGGCAGAAGAACAAGCATACCATGTCAACAAGG

GGCTGGAATGTATCAAGTCCCTCAAAGCCTCTCCCCCGGATCTATCCACCATCCAAGATGCCCTTGAGAGCTGGAGAGAG

GGGTTCAACCTCTCAGGTCGTGCAACACCGAACCATGATACGTCCAAGGGGGACCATCAGAATATCAACCAATCATGCTC

ATCAGCAATCGGATCAGACAAAGTCGACATGTCTCCTGAAGGTAATCTCGGATTTAGAGAGATCACTTGTGATGACAATG

AGGCTGGGCTCGGAGGAGTTCAGGACAAAAGATCCGACTCTCAAGTACAGCGTTACTATGTTTATAACCACGGGGGTGAA

GAGATTGAAGGACTCGAGGATGCTGACTCTCTCGTGGTTCAAGCAAATCCTCCGGTTGCTAACACCTTCGATGGAGGAGA

GGATGGATCTGACAACAGCGATGTGGACTTTGGCCCAGATGATCCCGACAGAGATCCTCTATATGACCGGGGACCTGCTG

CCGGCAATGATGTCTCTAGGTCAACAGATGTCGAAAAATTAGAAGGTGATGACATTCAAGAAGTTCTTAACTCCCAGAAG

AGTAAAGGAGGAAGATTCCAGGGCGGGAAAATCTTGCGAGTCCCGGAAATACCCGATGTCAAGAACTCCAGACCATCGGC

CCAATCAATTAAAAAGGGCACAGACGGGAGCTCAGTCTTATCTGGAATGGTGACAGAGTGTTCATCGATAAGTGGTGCAA

CCCAAGCTGTGCTAGAGTCCAGATGGGAGTCATCAGAGCGAAATGCATCTGTGGGGAGTGTCCCCAAATCTGTGAGGAGT

GCAAAGACGATCCAGGGGTTAACACAAGAATCTGGTACCATAGCATCACTGACTCAGCCTAAAGAAAATGACTCCGAGTA

TGAGTATGAGGATGATCTATTCGCTGAGATGCAGGACATTCGTGCAAGCATTGCTAAGATCCATGATGACAACAAAACTA

TCCTCTCAAAACTTGATTCTCTACTGTTATTGAAAGGAGAAATCGATACTATCAAGAAACAAATCAGCAAACAAAATATA

AGTATATCTACCATTGAGGGCCATCTATCCAGTATAATGATAGCCATCCCAGGTTTTGGGAAGGACATCAAGGACCCAAC

ATCTGAGGTTGAGCTGAACCCAGATTTGAGACCTATAATCAGCCGCGATTCTGGCAGGGCTCTTGCGGAGGTCCTTAAGA

AACCCGCTGTTGATAGGTCTCAGAAGAGCGGAATCAAAGTCAACTCCGGTTCAAAGGGTCAGCTCCTCAAGGATCTCCAG

CTAAAACCCGTCGACAAGCAGGCAAGCTCTGCAATCGGGTTTGTTCCATCCGACCATGAATCATCCAGAAGTGTCATCCG

CTCCATAATCAAGTCGAGCAAGCTTAACAATGATCACAAAGACTATCTTCTAGATTTACTGAATGATGTGAAAGGCTCTA

AGGATCTCAAGGAATTCCACAAGATGCTAACAGCCATTCTTGCCAAGCAGCCGTAACACATCCCACAATCTACATCCCAT

ACTCGGTTGAAAGCATCCTCTCAATCAGGCTATTACAAAAAACTTAGGAGCAAGGGCAACTGAGCTTCACAGACAGGATG

ACCGAGATTTACGATTTCGATAAATCAGCATGGGATGTCAAAGGGTCAATTGCTCGCATAGAACCCACCACCTATCACGA

CGGCCGACTGGTACCCCAGGTGAGGGTCATCGACCCTGGTCTGGGAGACAGAAAAGATGAATGCTTCATGTACCTGTTTC

TTCTAGGAGTGATTGAGGATAACGACCCCCTTTCTCCCCCAGTCGGGAGAACCTTCGGGTCTTTACCTCTAGGTGTCGGC

AGGTCAACTGCTAAGCCAGAGGAACTACTAAGGGAGGCCACTGAACTAGATATAGTGGTGAGGCGCACGGCAGGACTAAA

TGAGAAACTGGTATTTTACAACAACACTCCGCTATCTTTGTTAACACCCTGGAGGAAAGTCTTGACGACCGGAAGTGTGT

TCAGCGCCAACCAGGTTTGCAATGCGGTCAACCTAGTCCCACTCGATACTCCCCAAAGATTCAGGGTTGTGTACATGAGT

ATAACTAGATTGTCAGATAATGGTTATTATAGTGTCCCCAGAAGAATGTTGGAGTTCCGCTCAGCCAATGCAGTCGCTTT

CAATATCTTGGTTACACTGAGAATTGAAAATGGCACAAACCCTAGAAGATACATAGTCGGCTCATGGGAGAATCCAGAGG

TCACATTTATGGTACACGTGGGCAACTTTAGAAGAAAGAAGAACGAAGTATACTCTGCTGATTATTGCAAAATGAAGATT

GAAAAGATGGGTCTAGTTTTTGCCTTGGGAGGAATAGGTGGAACAAGTCTCCATATTCGAAGCACAGGGAAAATGAGCAA

GACCCTCCATGCACAGCTGGGGTTCAAGAAAATTCTATGCTACCCTCTGATGGATATCAATGAGGATCTTAACCGATATT

TCTGGCGGGCAGAGCGCCGAATAGTCAAAATCCAGGCCGTCTTACAGCCATCAGTACCCCAAGAATTCCGTGTCTACGAT

GATGTCATCATCAACGATGACCAAGGCCTGTTCAAGATCCTGTAAGTCACCTGCAGGGCCATGGACCCACCTCCAGGAAG

AGGCCCTCCCACCTCCCGCCCAAAACACACACCCCAAGACTCCCCGAAACAGCGCCGGCAAAACCACTCTCCCGGCGGGA

CGAGACCGACC-CCCCCAACAAACCCGCTCCGAGGGGGGG-CCCCCCCCGCGACCCGCCGGGCGCCGACCGCGCGAGCCG

GCGAGACACCCCCAAGAGGCTTCGACGAAGACCGGGGAACCCCTTCCCCA-AAAATGGCCCCCA-AACACACCCAAGACC

ATGCGCCTAGCACCCCCAGCCCCCCAACCGGCGGGACAGGGGGCCAGGAAGGGGTCCCCCAAAACAAAGGGACCACCAGG

GAAGGCTCCGCCGACCCAGCACAGACCCGGCCCAAACAAAGCAGACCCCCAAGACGAAATCTCCCAGAGCACCCCATTCA

TCATCAAACAAAACTTAGGGGCCAAGTCCAGACGACAGACCCTGGCCCGCCAGGCAGCAAGCCACAGAAAAGAAGGAGAC

AAGATCTGCCCGGGAAGAGGACACCCAACCACAGAAACCCCCAGCTCCAGGGGAAGAG-CCCCCCCTCCACACGACGGAG

GAACGGGCGAGAGACACCGAGGAGAACACGATCGGGAGCCGAGAGCCGGAGACGCCGGCGGCAAACAGGCCGGGCACCCA

ACCGACACCACCGGAGGACCCGAGGAGAGCCCGACTCCGCGCAGACTCGACGGCAGAGGACCCAGTCCCGCCACAAAGAG

AGGTGAGAACCCT--------CCCTCCTGCACACAACCGGCAAACAGACCGCGGGAGGACCACCTCCCAGCAAGACAGAC

GTCTCCCCCGCAGCCGAGCCGCAGCTCCTCAGAAGAGGGCACCCCCGAAGCCCCCCAGAACCCCTCCGCTCACGGCCGCC

GGGACAAGGGCTCCGCCGCCAAAGATCAGAGCCACCGAGAGTCAACCGGGGGTCCCCCCCAGTCCCTCCCGGCGATCCCC

CACAGAGGCCCCAGCAAGGTCCCCCGGCCCGACGACCCCTGCATGCGCCGAAAGGGAAACCCACCAGAGGACGTCCAGCA

CATCCGTGCATAAACATCATGACACGGGTCGCAACCTTGGTATTGCTGTTTCTTTTCCCAAACACTGTCGCGTGCCAGAT

TCACTGGGGCAATCTATCCAAGATAGGGATTGTCGGAACAGGGAGTGCCAGTTACAAGGTGATGACCAGGCCAAGCCACC

AAACTCTGGTTATAAAGTTGATGCCAAATATAACAGCCATCGACAATTGTACGAAATCAGAGATTTCAGAGTACAAAAGA

TTGCTAATCACAGTGTTAAAGCCTGTAGAGGATGCTTTGTCAGTAATAACCAAGAATGTAAGACCAATTCAAGCTCTAAC

ACCTGGGCGCAGGACCCGCCGTTTCGCCGGAGCTGTTCTGGCCGGAGTAGCACTTGGAGTCGCGACGGCTGCTCAAATAA

CTGCCGGAGTCGCACTCCATCAGTCATTGATGAATTCCCAAGCAATTGAAAGTTTAAAAACCAGTCTTGAGAAGTCAAAT

CAGGCAATAGAAGAAATCAGACTTGCAAATAAGGAGACCATACTGGCGGTACAGGGCGTCCAAGATTATATCAACAACGA

GCTTGTCCCCTCTGTTCATAGAATGTCATGTGAGCTAATAGGTCACAAACTCAGTCTCAAACTCCTTAGGTATTATACCG

AGATCCTGTCTATATTCGGGCCTAGCCTTCGAGACCCGATAGCTGCTGAAATATCAATCCAGGCACTCAGCTATGCACTA

GGCGGAGACATCAATAAAATTCTGGACAAGCTTGGGTATAGCGGCGGGGATTTCCTTGCTATTCTAGAAAGCAAGGGGAT

AAAGGCCCGGGTCACATATGTGGACACAAGAGATTACTTTATAATTCTTAGCATAGCCTACCCAACCTTATCTGAGATCA

AAGGGGTGATAGTTCATAAGATAGAAGCTATATCATACAACATTGGGGCACAGGAATGGTATACTACTATCCCTAAATAT

GTAGCCACTCAAGGATATTTGATATCGAATTTCGATGAGACATCATGTGTATTCACTCCAGAGGGGACAGTCTGCAGCCA

GAATGCGCTGTATCCAATGAGCCCATTGCTTCAGGAATGTTTCAGGGGGTCGACAAAATCGTGTGCCAGAACCCTAGTTT

CAGGGACCACAAGTAACAGATTTATCCTATCAAAAGGGAACTTGATTGCAAATTGTGCGTCAGTTTTGTGCAAGTGTTAC

ACAACGGAGACAGTCATCAACCAAGATCCTGATAAACTACTAACTGTTATAGCCTCAGATAAGTGTCCCGTAGTTGAGGT

GGATGGAGTGACAATACAAGTCGGCAGTCGAGAGTACCCAGATTCTGTATACCTACATAAAATAGACTTAGGCCCAGCCA

TCTCTCTGGAAAAACTGGATGTAGGCACCAATTTAGGCAATGCAGTCACAAGACTGGAGAATGCAAAGGAGCTCCTAGAC

GCATCAGACCAGATACTGAAGACTGTTAAAGGGGTACCCTTCAGTGGCAATATGTACATAGCACTGGCAGCTTGCATTGG

GGTATCCCTAGGGCTTGTCACATTAGTATGCTGCTGTAAGGGGAGATGTAGGAACAAGGAGATTCCTGCCTCCAAAATCA

ACCCAGGGCTCAAACCCGACCTGACCGGGACTTCAAAGTCTTACGTGAGATCACTGTAGTCAGAATAACCCGAATCACCC

GGCATCACGCGTATACATGTGTGACACCAGCAGTCAGAGGACGCAGAAGACTCAACCTCCGATCACCGACCAGACCCCAC

TCTACGCCCTATTACACATTGGTCATCAAACAAAACTTAGGACGAAAGGTCAATCACCATGTCCGCACAAAGGGAGAGGA

TCAATGCCTTCTACAAAGACAACCCTCACAATAAAAACCATAGGATAATCCTGGATAGGGAACGCTTAACTATTGAAAGA

CCCTACATCTTACTTGGAGTCCTGCTGGTAATGTTCCTTAGTCTAATCGGGCTGCTAGCCATTGCAGGGATCAGGCTTCA

CCGGGCCACCGTTGGAACTGCGGAGATCCAGAGTCGGCTGAATACCAACATTGAGTTGACCGAATCCATTGATCATCAAA

CTAAGGATGTCTTAACACCCCTGTTCAAAATCATTGGTGATGAAGTCGGTATCAGAATTCCACAGAAGTTCAGTGATCTT

GTCAAGTTCATCTCCGATAAGATTAAGTTCCTCAACCCTGACAGAGAATACGATTTTAGGGATCTCCGGTGGTGTATGAA

CCCCCCTGAGAGAGTCAAAATTAACTTTGATCAATTCTGTGAATACAAAGCCACGGTCAAGTCAGTTGAACATATATTTG

AGTCATCATTCAACAGGTCAGAAAGATTGCGACTATTGACTCTTGGACCCGGAACAGGCTGTCTCGGCAGGACAGTAACA

AGAGCTCAGTTCTCAGAACTTACGCTGACCCTGATGGACCTGGATCTCGAGATGAAGCACAACGTGTCCTCAGTGTTTAC

CGTAGTTGAAGAGGGATTATTCGGAAGAACATATATTGTTTGGAGATCTGACACCGGGAAACCGAGCACCAGTCTAGATA

TTGGCCAGTTTTTAAGAGTTTTCGAGATCGGGTTGGTGAGGGATCTCGAGCTGGGTGCCCCCATTTTCCATATGACCAAC

TACCTCACAGTGAACATGAGTGATGACTATCGGAATTGCCATTTAGCGGTAGGGGAGTTGAAGCTGACAGCCCTATGCAC

CCCATCTGAGACTGTGACTCTGAGTGAGAGAGGAGTTCCAAAGAGAGAGCCTCTTGTGGTTGTGATACTCAACCTAGTTG

GGCCTACTCTAGGGGGCGAACTATACAGTGTCTTGCCTACCTCTGACCTCATGGTGGAGAAACTCCATCTATCCTCACAT

AGAGGGATCATCAAAGACAACGAGGCCAATTGGGTAGTACCGTCTACCGATGTTCGTGATCTCCAAAACAAGGGAGAATG

TCTGGTGGAAGCATGCAAAACCCGACCTCCTTCATTTTGCAATGGCACAGGAATAGGCCCATGGTCAGAGGGGAGAATCC

CTGCCTACGGAGTGATCAGGGTCAGTCTTGACTTTGCTAGTGACCCGGGTGTGGTTATCACTTCAGTGTTTGGCCCACTG

ATACCTCACCTATCTGGCATGGATCTTTACAACAATCCGTTTTCAAGAGCTGCATGGCTGGCTGTACCACCCTACGAGCA

GTCATTTCTAGGAATGATAAATACAATTGGCTTCCCGGACAGAGCAGAGGTCATGCCGCACATTTTGACCACAGAGATCA

AAGGGCCTCGAGGTCGTTGTCATGTTCCTATAGAGTTGTCCCGCAGGATTGATGATGATATCAAGATCGGGTCCAATATG

GTTGTATTGCCGACGAGGGATCTGAGGTACATAACAGCCACTTATGATGTTTCCAGGAGCGAGCATGCAATTGTGTACTA

TATCTATGACACGGGTCGATCATCATCTTACTTCTACCCAGTTCGATTGAATTTCAAGGGCAATCCTCTCTCTCTGAGGA

TAGAGTGTTTCCCGTGGTCTCATAAGGTGTGGTGCTACCATGATTGTCTTATATACAACACCATGACAAACGAAGAAGTC

CGTACGAGAGGGCTGACCGGTATAGAGGTAACATGCAATCCAGTCTGAGTCGAGCTGAAACCATCGCTCAAGCAGGCTTC

CGAGCCATCCCCTAGTTCAAGCAGCATAGTCTGGGACACGCAGCAGCACAACCCAGCCAACAATGTTATAAAAAACTTAG

GAGCCAAGGTTGTAGGAGCCATGGACTCACTATCAGTCAATCAGGTTTTGTACCCTGAGGTCCATCTAGATAGCCCTATT

GTCACAAACAAACTAGTTGCTATCCTTGAATACTCGGGGATCAACCACAACTATGTTCTTGAAGACCAGACTCTTATCAA

AAATATTAGATATAGACTGGGGTGTGGTTTCTCAAATCAAATGATCATCAATAATAGGGGGGTAGGTGAAACAGTCAATT

CCAAGCTTAAAAGTTACCCCCGTAACTGTCATATCATATACCCAGACTGCAATAAGGAGTTGTTTTGTATCAAAGATAGC

TGCATATCTAGAAAGCTCTCGGAGCTATTCAAGAAGGGTAATTCCTTGTACTCTAAGATAAGTCACCAGGTACTGGATTG

TCTTAAGAGAGTCAACGGGAAATTAGGGCTGGGCACAGATCTTAATCACGGCCTGAAGGATGGTATCCTCGACTTGGGGT

TGCACATGCATAGCTCTCAGTGGTTCGAGACCTTTCTGTTCTGGTTCACTATCAAGACAGAGATGAGATCAATGATCAAA

GAACAGTCCCATATATGCCACAAGAGGAGGTATAACCCAATTTTTGTGTCGGGGGATGCATTCGAGGTGCTTGTATCACG

AGACCTCGTAGTGATAATTGATAAGAACACCCAGTATGTCTTCTACCTGACGTTTGAGCTGGTCCTTATGTATTGTGATG

TCATAGAGGGTAGACTTATGACGGAGACAGCCATGGCTATAGACCAGAGATATTCAGAGCTTCTAAACCGAGTCAGATAC

TTGTGGGACCTTATCGATGGGTTCTTCCCAACACTGGGTAACACCACATACCAAGTTGTTGCTCTGCTTGAACCACTGTC

GTTGGCTTATCTTCAACTTCAGGATGTCACTCTAGAGTTAAGAGGTGCCTTTTTGGACCACTGCTTCAAAGAACTTTATG

AGATACTGGAGCATTGTGGCATTGACACGGAAGGTACCTACAATTCTATCACTGAGGGATTGGATTACGTATTTATCACT

CATGATATACACTTAACTGGGGAGATTTTTTCATTTTTCCGGAGTTTCGGACACCCCCGCCTCGAAGCGGTCACCGCTGC

AGAGAATGTCAGAAAACATATGAACCAACCGAAGGTAATCAGTTATGAGACTATGATGAAAGGGCATGCAGTATTTTGCG

GGATAATCATAAATGGTTTTAGGGACCGGCACGGCGGCAGCTGGCCCCCTGTTGCATTGCCAGAACATGCTTCTGCTGCG

ATCCGGAATGCGCAGGCATCCGGCGAAGGACTGACCCACGACCTGTGTATAGACAACTGGAAGTCCTTTGTAGGATTCAG

ATTTGGCTGCTTCATGCCGCTCAGCCTAGATAGTGATTTGACCATGTACCTCAAAGACAAAGCATTGGCTGCATTAAAGA

ATGAGTGGGATTCAGTTTACCCGAAAGAATACCTCCGTTATAATCCACCTAGAGGGACAGAGTCAAGGCGACTAGTAGAG

GTGTTCCTGAATGACTCCAGCTTTGATCCTTATAACATGATAATGTACGTGGTGAATGGCTCCTACCTTAAAGACCCTGA

GTTTAATCTCTCATACAGCTTAAAAGAAAAGGAGATAAAAGAGACAGGGCGGTTGTTTGCCAAGATGACCTACAAGATGC

GGGCCTGTCAGGTGATTGCTGAAAATCTGATATCAAATGGTGTTGGGAAGTATTTCCGAGACAATGGGATGGCAAAAGAC

GAGCATGACCTAACAAAAGCCCTTCACACTCTGGCAGTCTCAGGCGTTCCCAAGAACAACAAAGACAACCACCGAGGTGG

GCCCCCCAGAAGGAACGCAAGCCGAGAGACGAGACCAAGCCAAGCCGTCAACACACAAAATAGAGACAAGATCCAGGGGG

GCCCTATGTACAACTACTTGCGATGCCAACCAACCAGCCCTGATCAGGGTGAGTCATACGAGACTGTTAGTGCATTCATC

ACCGCTGACCTTAAGAAGTATTGCCTAAATTGGAGATACGAGACAATCAGCATATTTGCACAGAGACTGAATGAAATATA

CGGGTTGCCATCCTTCTTTCAATGGTTACATAAGGTATTGGAAAAATCCGTGCTCTACGTCAGTGATCCGCATTGCCCTC

CCGACTTAGATGATCATATCCCTCTGGACAGTGTCCCTAATGCCCAAATATTCATCAAGTACCCAATGGGCGGAATAGAA

GGTTATTGTCAAAAACTATGGACAATCAGTACTATACCATACTTGTATCTAGCAGCCTATGAGAGTGGAGTAAGAATCGC

CTCACTAGTGCAAGGTGACAATCAGACAATTGCAGTGACAAAAAGAGTTCCAAGTTCTTGGCCTTATTCACTAAAAAAGA

GGGAGGCATCTAAAGCAGCTCAAAATTACTTTGTGGTCTTAAGGCAAAGGTTGCATGATGTAGGTCATCACTTAAAGGCT

AATGAGACCATAGTATCTTCTCACTTTTTTGTGTACTCTAAAGGGATCTATTATGACGGCCTGTTAGTTTCACAATCACT

AAAGAGCATCGCCAGATGTGTCTTCTGGTCTGAGACTATCGTGGATGAAACTAGAGCAGCCTGCAGCAATATTGCAACAA

CTATCGCTAAGAGTATAGAGAGGGGTTATGATAGGTACCTCGCATACTCTTTGAATATCCTCAAAATTTTCCAACAGATC

CTTGTATCCCTTGACTTCACGATTAATACAACAATGACTCAAGATGTCGTGGCACCGATCATCGAGAACGGTGATTTACT

GATAAGGATGGCACTCTTGCCAGCACCCATTGGGGGTCTCAATTATCTCAACATGAGCAGGTTGTTTGTGAGAAATATCG

GGGACCCGGTCACTTCCTCCATAGCCGACCTGAAGAGGATGATAGACGCTGGGCTAATGCCAGAAGAAACATTGCATCAA

GTGATGACCCAGACCCCGGGAGAATCATCCTACCTTGATTGGGCAAGTGATCCTTATTCTGCCAACCTACCCTGCGTACA

GAGTATAACTCGCCTTCTCAAGAACATCACTGCACGGTATATTTTAATCAGCAGCCCAAATCCGATGCTGAAAGGGTTGT

TTCACGAGGGGAGTAGAGATGAAGACGAGGAGCTTGCGAGTTTCTTAATGGATCGGCATATAATTGTTCCAAGAGCTGCA

CATGAAATCTTAGACCACAGCATAACCGGGGCAAGGGAAGCTATAGCCGGGATGTTGGACACCACCAAGGGTCTGATTAG

AACAAGTATGAAACGGGGTGGCCTCACCCCTCGAGTATTAGCCCGCCTTTCCAATTATGATTATGAACAATTCAGATCCG

GGATAACACTATTGACAAAGAAAGGGCAGTGTTATCTCATTGACAAGGACTCGTGCTCGGTGCAGCTCGCTATAGCTCTG

AGGAGCCATATGTGGGCAAGGTTGGCTCGCGGGAGACCTATCTATGGCTTGGAGGTGCCTGATGTACTGGAATCGATGAA

CGGCTACCTTATCAAACGTCACGAGTCTTGTGCCATCTGTGAAACGGGCTCAAGTCACTACGGGTGGTTTTTCGTCCCTG

CAGGGTGTCAGCTTGACGATGTCTCAAGAGAGACTTCAGCTCTTCGTGTGCCTTATGTCGGATCAACCACTGAGGAAAGG

ACAGATATGAAACTTGCTTTCGTTAGATCTCCAAGCCGATCCCTCAAATCAGCAGTCAGGATTGCCACAGTCTACTCATG

GGCCTACGGGGATGATGAGAAATCATGGAGTGAAGCTTGGATGCTAGCTAGGCAGAGAGCCGATGTCACCTTAGATGAAC

TGAGAATGATCACTCCAGTCTCTACATCCACCAACCTAGCTCATCGGTTAAGGGATCGGAGCACTCAGGTTAAATATTCG

GGGACATCCCTTGTGAGGGTTGCAAGGTACACAACCATCTCCAATGACAATTTGTCATTTGTGATATCTGAGAAAAAAGT

AGATACCAACTTCATTTACCAGCAAGGGATGCTGCTCGGTCTTGGGATCCTTGAGAATCTCTTCAGGTTAGAGGCCACCA

CAGGGGTATCCAACACAGTGCTACACCTGCACGTGGAAACAGAATGTTGTGTTGTACCTATGGTTGACCACCCAAGGATA

CCGAGTCTCCGCAATATCAAAGTTACGAACGAGCTATGCACAAACCCTCTTATCTACGACAAGTCCCCCATCATAGAACA

CGATGCAACTCGATTATACTCACAAAGCCACAGGAGACATTTGGTGGAGTTTGTTACCTGGTCAACAAGCCAGCTCTATC

ATATACTGGCCAAATCTACAGCAATGTCCATGATTGAGCTGATCACAAGATTTGAGAAAGATCACATGAATGAAATAGCC

GCTCTGATTGGCGATGACGACATCAACAGTTTCATCACAGAATTTTTGCTTGTGGAGCCCAGACTGTTTATAGTTTACCT

CGGCCAGTGTGCTGCCATCAATTGGGCCTTTGATATACATTATCATCGGCCCTCGGGCAAGTACCAGATGGGGGAACTCC

TCTACTCTTTACTCTCTCGGATGAGCAAGGGAGTATATAAGATCTTCACTAATGCTCTGAGCCACCCCAAAGTTTACAAG

AAATTCTGGCGAAGTGGTATAATTGAGCCGGTTCATGGCCCATCCCTAGATACACAGAATTTACATGTCACTGTCTGTGA

CATGATATTCGGATCATACGTCACCTATCTGGATCTTTTGCTGAATGATGAGCTGGATGCTTACCCGTATTTGCTCTGCG

AGAGTGATGAGGACGTGGTCACAGACAGGTTCGACAACATTCAAGCCAAACATCTCTGTGTACTGGCCGATGTATACTGC

AGCTCCAAGAGGTGTCCCTCGATAATCGGGATGTCCCCTATAGAAAAATGTACCATCCTCACACATTACATCAAGGGAGA

ATCGGTACAATCCCCGTCCGGGATCTCATGGAACACCGATCCCCTTGTAGTAGATCATTACTCATGCTCTCTGACCTACC

TCCGCCGTGGTTCCATCAAACAAATCAGATTGAGAGTGGATCCTGGGTTTGTGTTCGAGGCGTTGACAGACATCGACTTC

AAACAGCCTCGCAAGGCTAAGTTGGACGTATCGATTGTGGGATTGACTGATTTTTCTCCCCCTTGGGATAACGTCGGTGA

TTTTCTAGGGACTATCAACACATTGAGGCACAATCTGCCCGTCACCGGGACCGGGGTCTCGAACTATGAAGTCCACGCTT

ATCGTAGAATTGGTCTGAATTCATCAGCATGTTATAAAGCTGTAGAGATCTCCACGTTAATAAAGTCATCCTTAGAAGCT

GGGGAGAATGGATTGTTCTTAGGAGAAGGCTCCGGCTCGATGCTGGCTGCGTACAAGGAAGTTCTTAAATTGGCTAACTG

TTATTACAACAGTGGGGTAACAGCAGAGGGTAGAGCCGGACAGAGGGAAATCTCTCCCTATCCCTCAGAGGTGAGCCTGG

TAGAGAGTCAGATGGGGATAGAGAGAAGTGTTAAAGTCCTGTTCAATGGCAAACCTGAAGTAACCTGGGTAGGGACCACC

GATTGCTACAAGTATATAATCAGTAACATTCAGACCTCTAGTCTGGGTTTCATACACTCAGATATCGAGACACTCCCAAC

CAAGGATGCCGTTGAGAAGTTGGAGGAATTTGCCTCTATTCTATCTCTATCCCTAATTTTGGGAAAAATCGGCTCTATTA

CAGTTGTCAAAATTATGCCCATTAGCGGAGATTTTACCCAAGGCTTCATAGCCTATGCCATTCAATATTTCAGGGAGAGC

CTGCTTGCCTATCCGAGATATAGTAACTTCATCTCGACTGAGTGTTACCTTATTATGATAGGATTAAAGGCCAATCGGTT

GATAAACCCAGAAGCCATTAAGCAAAGCATAATCAGGATGGGGACTAGGACTGCACCAGGGCTTGTGAGCCACATATTAT

CAGAGAAACAGAAAGGTTGTATTCAATCTTTTCTGGGTGATCCTTATGTCCAAGGAGACTTCAATAAGCACCTTAAAGCT

CTAACTCCTATTGAGAAAATCCTTGTAAATTGTGGTCTCTCGATCAATGGTACAAAAATCTGTAGGGATCTAATCCACCA

TGATATTGCCTCCGGTCCAGACGGTCTGATGAGCTCCACAATTATTTTATACAGGGAACTGGCCCATTTCAAAGACAATA

TAAGAAGTCAGCACGGCATGTTCCACCCCTACCCAGTATTGGCCAGTAGCAGGCAACGTGAATTGATCCTTCGAATAGCC

AAGAAGTTTTGGGGGTATGTCTTGCTATATTCTGATGACCCGGCACTAATCAAACAAACGATCAAGAACTTGAAGCGGAA

TCACCTAACCTTTGACTTACACAGTAATCCGTTTATTAAGGGCTTATCCAAAGCTGAGAAACTGCTAGTGCGGACAAGTT

CACTCAGAAGGGAATGGTTGTTCACTCTCGATACGAAAGAAGTGAAAGAGTGGTTCAAATTGGTGGGTTACAGTGCACTC

GTCAGAGGTTAATTCGCGATACATCTGCCCCCTTCCCCTCCACCATGAGACTCTACTGGCAATCCAAAAGATTAAAGAAA

ACTACATATTGGATAAGTATCTATTCCCAGCTTTGTCTGGT

>KJ466104.1/sheep/Ghana/2010

ACCAAACAAAGTTGGGTAAGGATAGATCTTACAATGACTATAGACTAGCAAACTTAGGAGTAAAGATCCTACTGTCGGGG

AGAGGAGGAGGAGCAAGATCTTTGACCATGGCGACTCTCCTTAAAAGCTTAGCACTGTTCAAGAGGAACAAAGACAAAGC

GCCTACTGCGTCAGGTTCAGGAGGGGCCATCCGGGGGATTAAGAATGTTATCATAGTCCCTATCCCCGGGGACTCATCCA

TCATTACCCGTTCAAGACTGCTCGACAGGCTTGTCAGACTGGCCGGAGATCCTGACATCAACGGGTCAAAGCTGACCGGC

GTGATGATCAGCATGTTATCTTTGTTCGTGGAGTCACCCGGGCAATTGATACAGCGGATCACAGATGATCCAGATGTTAG

CATCCGTCTTGTTGAGGTAGTTCAAAGTACCAGGTCCCAGTCAGGATTAACCTTTGCATCACGTGGTGCTGATTTGGACA

ATGAGGCAGATATGTATTTTTCAACTGAAGGACCCTCGAGTGGAAGTAAGAAAAGGATCAACTGGTTTGAGAACAGAGAA

ATAATAGATATAGAAGTGCAAGATGCAGAAGAGTTCAATATGTTGTTAGCCTCCATCTTAGCACAAGTTTGGATCCTCCT

GGCCAAGGCGGTCACGGCACCAGATACAGCAGCCGACTCAGAACTGAGAAGGTGGGTTAAATACACACAACAAAGGAGAG

TGATTGGGGAATTTCGCTTTGACAAAGGGTGGCTGGACGCAATCCGCAACAGGATTGCAGAAGATCTATCACTCCGGCGG

TTCATGGTATCCCTGATACTTGACATCAAGAGGACCCCCGGCAACAAGCCAAGGATTGCAGAAATGATCTGCGACATTGA

CAACTATATTGTCGAAGCCGGACTTGCAAGTTTCATTCTCACTATCAAGTTTGGTATTGAAACCATGTATCCTGCATTAG

GCCTTCACGAGTTCGCCGGGGAATTGTCCACTATTGAATCCTTGATGAACTTGTATCAACAGCTAGGAGAAGTTGCACCC

TATATGGTAATTCTAGAGAACTCAATTCAGAACAAGTTTAGTGCAGGAGCCTATCCTCTCCTCTGGAGCTATGCGATGGG

TGTCGGAGTCGAGTTGGAGAACTCAATGGGGGGCTTGAACTTTGGCAGGTCATATTTTGACCCGGCCTATTTTCGTCTCG

GACAGGAGATGGTCAGAAGATCTGCAGGAAAGGTCAGCTCTGTAATTGCGGCTGAGCTTGGCATCACAGCAGAGGAAGCC

AAACTAGTCTCGGAAATCGCCTCGCAGACTGGGGAAGAACGAACCGTTAGAGGGACTGGGCCTCGACAGGCGCAGGTTTC

CTTCCTCCAGCACAAAATTGGTGAGGGAGAGTCGCCTACACCAGCGACCAAAGAAGAAGTTAAAGCTGCGATCCCAAACG

GATCTGAAGGAAGGGACATAAAGCGAACACGCCCAAGGAAGCCCAGAGGAGAAACTCCCGGGCAACTGCTTCTGGAGATC

ATGCCAGAGGATGAAGCCTCGCGAGAGTCTGGTCAAAACCCTCGTGAGGCTCAAAGATCGGCTGAGGCACTCTTCAGGCT

GCAGGCCATGGCCAAGATTCTGGAGGACCAGGAGGAGGGAGAAGACAACAGTCAGATCTACAACGACAAGGATCTCCTCA

GCTGAGCAGATACACCCTCTGTCCAAATCAGTGACAAGACATCACCTACCAGTATTATAAAAAACTTAGGACCCAGGTCC

AACCAACCGCACATCGACACCCTAGTCAATCGAGCGGAGACCACCGATGGCAGAAGAACAAGCATACCATGTCAACAAGG

GGCTGGAATGTATCAAGTCCCTCAAAGCCTCTCCCCCGGATCTATCCACCATCCAAGATGCCCTTGAGAGCTGGAGAGAG

GGGTTCAACCCCTCAGGCCGTGCAACACCGAACCATGATACGTCCGAGGGGGACCATCAGAATATCAACCAATCATGCTC

TTCAGCAATCGGATCAGACAAAGTCGACATGTCTCCTGAAGGTAATCTCGGATTTAGAGAGATCACTTGTGATGACAATG

AGACTGGGCTCGGAGGAGTTCAGGACAAAGGATCCGACTCTCAAGTACAGCGTTACTATGTTTATAACCACGGGGGTGAA

GAGATTGAAGGACTCGAGGATGCTGACTCTCTCGTGGTTCAAGCAAATCCTCCGGTTGCTAACACCTTCGATGGAGGAGA

GGATGGATCTGACAACAGCGATGTGGACTCTGGCCCAGATGATACCGGCAGAGATCCTCTATATGACCGGGGACCTGCTG

CCGGCAATGATGTCTCTAGGTCAACAGATGTCGAAAAATTAGAAGGTGATGACATTCAAGAAGTTCTTAACTCCCAGAAG

AGTAAAGGAGGAAGATTCCAGGGCGGGAAAATCCTGCGAGTCCCGGAAATACCCGATGTCAAGAACTCCAGACCATCGGC

CCAATCAATTAAAAAGGGCACAGACGGGAGCTCAGTCTTATCTGGAATGGTGACAGAGTGTTCATCGATAAGTGGTGCAA

CCCAAGCTGTGCTAGAGTCCAGATGGGAGTCATCAGAGCGAAATGCATCTGTGGGGAGTGTCCCCAAATCTGCGAGGAGT

GCAAAGACGATCCAGGGGTTAACACAAGAATCTGGTACCATAGCATCACTGACTCAGCCTAAAGAAAATGACTCCGAGTA

TGAGTATGAGGATGATCTATTCACTGAGATGCAGGACATTCGTGCAAGCATTGCTAAGATCCATGATGACAACAAAACTA

TCCTCTCAAAACTTGATTCTCTACTGTTATTGAAAGGAGAAATCGATACTATCAAGAAACAAATCAGCAAACAAAATATA

AGTATATCTACCATTGAGGGCCATCTATCCAGTATAATGATAGCCATCCCAGGTTTTGGGAAGGACATCAAGGATCCAAC

ATCTGAGGTTGAGTTGAACCCAGATTTGAGACCTATAATCAGCCGCGATTCTGGCAGGGCTCTTGCGGAGGTCCTTAAGA

AACCCGCTGTTGATAGGTCTCAGAAGAGCGGAATCAAAGTCAACTCCGGTTCAAAGGGTCAGCTCCTCAAGGATCTCCAG

CTAAAACCCGTCGACAAGCAGGCAAGCTCTGCAATCGGGTTTGTTCCATCCGACCATGAATCATCCAGAAGTGTCATCCG

CTCCATAATCAAGTCGAGCAAGCTTAACATTGATCACAAAGACTATCTTCTAGATTTACTGAATGATGTGAAAGGCTCTA

AGGATCTCAAGGAATTCCACAAGATGCTAACAGCCATTCTTGCCAAGCAGCCGTAACACATCCCATAATCTACATCTCAT

ACTCGGTTGAAAGCATCCTCTCAATCAGGATATTACAAAAAACTTAGGAGCAAGGGCAACTGAGCTTCACAGACAGGATG

ACCGAGATTTACGATTTCGATAAATCAGCATGGGATGTCAAAGGGTCAATTGCTCGCATAGAACCCACCACCTATCACGA

CGGCCGACTGGTACCCCAGGTGAGGGTCATCGACCCTGGTCTGGGAGACAGAAAAGATGAGTGCTTCATGTACCTGTTTC

TTCTAGGAGTGATTGAGGATAACGACCCCCTTTCTCCCCCAGTCGGGAGAACCTTCGGGTCTTTACCTCTAGGTGTCGGT

AAGTCAACTGCTAAGCCAGAGGAACTACTAAGGGAGGCCACAGAACTAGATATAGTGGTGAGGCGCACGGCAGGATTAAA

TGAGAAACTGGTATTTTACAACAACACTCCGCTATCTTTGTTAACACCCTGGAGGAAAGTCTTGACGACCGGAAGTGTAT

TCAGCGCCAACCAGGTTTGCAATGCGGTCAACCTAGTCCCACTCGATACTCCCCAGAGATTCAGGGTTGTGTACATGAGT

ATAACTAGATTGTCAGATAATGGTTATTATAGTGTCCCCAGAAGAATGTTGGAGTTCCGCTCAGCCAATGCAGTCGCTTT

TAATATCTTGGTTACACTGAGAATTGAGAATGGCACAAACCCTAGAAGATACATAGTCGGCTCATGGGAGAATCCAGAAG

TCACATTTATGGTACACGTGGGCAACTTTAGAAGAAAGAAGAACGAAGTATACTCTGCTGATTATTGCAAAATGAAGATT

GAAAAGATGGGTCTAGTTTTTGCCTTGGGAGGAATAGGTGGAACAAGTCTCCATATTCGAAGCACAGGGAAAATGAGCAA

GACCCTCCATGCACAGCTGGGGTTCAAGAAAATTCTATGCTACCCTCTGATGGATATCAATGAGGATCTTAACCGATATC

TCTGGCGGGCAGAGTGCCGAATAGTCAAAATCCAGGCCGTCTTACAGCCATCAGTACCCCAAGAATTCCGTGTCTATGAT

GATGTCATCATCAACGATGACCAAGGCCTGTTCAAGATCCTGTAAGTCACCTGCAGGGCCATGGACCCACCTCCAGGAAG

AGGCCCTCCCACCTCCCGCCCAAAACACACACCCCAAGACTCCCCGAAACAGCGCCGGCACAACCACTCTCCCGGCGGGA

CGAGGCCGACC-CCCCCAACAAACCCGCTCCGGAGGGGGG-CCCCCCCCGCGACCCGCCGGGCGCCAACCGCGCGAGCCG

GCGAGACACCCCCAAGAGGCCTCGACGGAGACCGGGCAACCCCTTCCCCA-AAAATGGCCCCCC-AACACACCCAAGGCC

ATGCGCTGAGCACCCCCAGCCCTCCAATCGGCAGGACAAGAGGCCAGGACGGGGTCCCCCAAAACAAAGGGACCACCATG

GAAGGCTCCGCCGACCCAGCACAGACCCGGCTTAAACAAAGCAGACCCCCAAGACGAAATCTCCCAGAGCACCCCATTCA

TCATCAAACAAAACTTAGGGGCCAAGTCCAGACGACAGACCCCAGCCCGCCAGGCAGCAAGCCACAGAAAAGAAGGAGAT

AAGATCGGCCCGGGAAGAGGACACCCAACCACAGAAACCCCCAGCTCCAGAGGAAGAG-CCCCCCCTCCACACGACGGAG

GAACGGGCGAGAGACACCGAGGAGAGCACGATCGGGAGCCAAGAGCCGGAGACGCCGGCGGCAAACAGGCCGGGCACTCA

ACCGACACCACCGGAGGACCCGAGGAGAGCCCGACTCCGCGCAGACTCGACGGAAGAGGACCCAGTCCCGCCACAAGGAG

AGGTGAGAACCCT--------CCCTCCTGCACACAACCGGCAAACAGACCGCGGGAGGACCACCTCCCAGCAAGACAGAC

GTCTCCCCCGCAACTGAGCCGCAGCTCCTCAGAAGAGGGCACCCCCGAAACCTCCCAGAACCCCTCCGCTCACGGCCGCA

GGGACAAGAGTTCCGCCGCCAAAGACCAGAGCCACCGAGAGCCAACCAGGGGTCCCCCCCAGTCCCTCCCGGTGATCCCC

CACAGAGGCCCCAGCAAGGTCCCCCGGCCCGACGACCCCCGCATGCGCCGGAAGGGAAACCCACCAGAGGACGTCCAGCA

CATCCGTGTATAAACATCATGACACGGGTCGCAACCTTGGTATTGCTGCTTCTCTTCCCAAACACCGTCGCGTGCCAGAT

TCACTGGGGCAATCTATCCAAGATAGGGATTGTCGGAACAGGGAGTGCCAGTTACAAGGTGATGACCAGGCCAAGCCACC

AAACTCTGGTTATAAAGTTGATGCCAAATATAACAGCCATCGACAATTGTACGAAATCAGAGATTTCAGAGTACAAAAGA

TTGCTAATCACAGTGTTAAAGCCTGTAGAGGATGCTCTGTCAGTAATAACCAAGAATGTAAGACCAATTCAAGCTCTAAC

ACCTGGGCGCAGGACCCGCCGTTTCGCCGGAGCTGTTCTGGCCGGAGTAGCACTTGGAGTCGCGACGGCTGCTCAAATAA

CTGCCGGAGTCGCACTCCATCAGTCATTGATGAATTCCCAAGCAATTGAAAGTTTAAAAACCAGTCTTGAGAAGTCAAAT

CAGGCAATAGAAGAAATCAGACTTGCAAATAAGGAGACCATACTGGCAGTACAGGGCGTCCAAGATTATATCAACAACGA

GCTTGTCCCCTCTGTTCATAGAATGTCATGTGAGCTAATAGGTCACAAACTCAGTCTCAAACTCCTTAGGTATTATACCG

AGATCCTGTCTATATTCGGGCCTAGCCTTCGAGACCCGATAGCAGCTGAAATATCAATCCAGGCACTCAGCTATGCACTA

GGCGGAGACATCAATAAAATTCTGGACAAGCTTGGGTATAGCGGCGGGGATTTCCTTGCTATTCTAGAAAGCAAGGGGAT

AAAGGCTCGGGTCACATATGTGGACACAAGAGATTACTTTATAATTCTTAGCATAGCCTACCCGACCTTATCTGAGATCA

AAGGGGTGATAGTTCATAAGATAGAAGCTATATCATACAACATTGGGGCACAGGAATGGTATACTACTATCCCTAAATAT

GTAGCCACTCAGGGATATTTGATATCGAATTTCGATGAGACGTCATGCGTCTTCACTCCAGAGGGGACAGTCTGCAGCCA

GAATGCGTTGTATCCAATGAGCCCATTGCTTCAGGAATGTTTCAGGGGGTCGACCAAATCGTGTGCCAGAACCCTAGTTT

CAGGGACCACAAGTAATAGATTTATCCTGTCAAAAGGGAACTTGATTGCAAATTGTGCGTCAGTTTTGTGCAAGTGTTAC

ACAACGGAGACAGTCATCAACCAAGATCCTGATAAACTACTAACTGTTATAGCCTCAGATAAGTGTCCCGTAGTTGAGGT

GGATGGAGTGACAATACAAGTCGGCAGTCGAGAGTACCCAGATTCTGTATACCTACATAAAATAGACTTAGGCCCAGCCA

TCTCTCTGGAAAAACTGGATGTGGGCACCAATTTAGGCAATGCAGTCACGAGACTGGAGAATGCAAAGGAGCTCCTAGAC

GCATCAGACCAGATACTGAAGACTGTGAAAGGGGTACCCTTCAGTGGCAATATGTACATAGCACTGGCAGCTTGCATTGG

GGTATCCCTAGGGCTTGTCACATTAATATGCTGCTGTAAGGGGAGATGTAGGAACAAGGAGATTCCTGCCTCCAAAATCA

ACCCAGGGCTCAAACCCGACCTGACCGGGACTTCAAAGTCTTACGTGAGATCACTGTAGTCAGAACAACCCGAATCATCC

GGCATCACGCGTATACATGTGCGACACAAGCAGTCAGAGGACGCAGAAGAGTCAACCTCCGATCACCGACCAGACCCCAC

TCTACGCCCTACTACACATTGGTCATCAAACAAAACTTAGGACGAAAGGTCAATCACCATGTCCGCACAAAGGGAGAGGA

TCAATGCCTTCTACAAAGACAACCCTCACAATAAAAACCATAGGATAATCCTGGATAGGGAACGCTTAACTATTGAAAGA

CCCTACATCTTACTTGGAGTCCTGCTGGTAATGTTCCTGAGTCTAATCGGGCTGCTAGCCATTGCAGGGATCAGGCTTCA

CCGGGCCACCGTTGGAACTGCGGAGATCCAGAGTCGGCTGAATACCAACATTGAGTTGACCGAATCCATTGATCATCAAA

CTAAGGATGTCTTAACACCCCTGTTTAAAATCATTGGTGATGAAGTCGGTATCAGAATTCCACAGAAGTTCAGTGATCTT

GTCAAGTTCATCTCCGATAAGATTAAGTTCCTCAACCCTGACAGAGAATACGATTTTAGGGATCTCCGGTGGTGTATGAA

CCCCCCTGAGAGAGTCAAAATTAACTTTGATCAATTCTGTGAATACAAAGCCGCGGTCAAGTCAGTTGAACATATATTTG

AGTCATCATTCAACAGGTCAGAAAGATTGCGACTATTGACTCTTGGGCCCGGAACAGGCTGTCTCGGCAGGACAGTAACA

AGAGCTCAGTTCTCAGAACTTACGCTGACCCTGATGGACCTGGATCTCGAGATGAAGCACAACGTGTCCTCAGTGTTTAC

CGTAGTTGAAGAGGGATTATTCGGAAGAACATATATTGTCTGGAGATCTGACACCGGGAAACCGAGCACCAGTCTAGATA

TTGGCCAGTTTTTAAGAGTCTTTGAGATCGGGTTGGTGAGGGATCTCGAGCTGGGTGCCCCCATTTTCCATATGACCAAC

TACCTCACGGTGAACATGAGTGATGACTATCGGAGTTGCCTTTTAGCGGTAGGGGAGTTGAAGCTAACAGCCCTATGCAC

CCCATCTGAGACTGTGACTCTGAGTGAGAGAGGAGTTCCAAAGAGAGAGCCTCTTGTGGTTGTGATACTCAACCTAGTTG

GGCCTACTCTAGGGGGCGAACTATACAGTGTCTTGCCTACCTCTGACCTCATGGTAGAGAAACTCCATCTATCCTCACAC

AGAGGGATCATCAAAGACAACGAGGCCAATTGGGTAGTACCGTCTACCGATGTTCGTGATCTCCAAAACAAGGGAGAATG

TCTGGTGGAAGCATGCAAAACTCGACCTCCTTCATTTTGCAATGGCACAGGAATAGGCCCATGGTCAGAGGGGAGAATCC

CTGCCTACGGAGTGATCAGGGTCAGTCTTGACTTAGCTAGTGACCCAGGTGTGGTTATCACTTCAGTGTTTGGCCCACTG

ATACCTCACCTATCTGGCATGGATCTTTACAACAATCCGTTTTCAAGAGCTGCATGGCTGGCTGTACCACCCTACGAGCA

GTCATTTCTAGGAATGATAAATACAATTGGCTTCCCGGACAGAGCAGAGGTCATGCCGCACATTTTGACCACAGAGATCA

AAGGGCCTCGAGGTCGTTGTCATGTTCCTATAGAGTTGTCCCGCAGGATTGATGATGATATCAAGATCGGGTCCAATATG

GTTGTATTGCCGACGAGGGATCTGAGGTACATAACAGCCGCTTATGATGTTTCCAGGAGCGAGCATGCAATCGTGTACTA

TATCTATGACACGGGTCGCTCATCATCTTACTTCTACCCAGTTCGATTGAATTTCAAGGGCAATCCTCTCTCTCTGAGGA

TAGAGTGTTTCCCCTGGTCTCATAAGGTGTGGTGCTACCATGATTGTCTTATATACAACACCATGACAAACGAAGAAGTC

CATACGAGAGGGCTGACCGGTATAGAGGTAACATGCAATCCAGTCTGAGTCGAGCTGAAACCATCGCTCAAGCAGGCTTC

CGAGCCATCCCCTAGTTCAAGCAGCATAGTCTGGGACACTCAGCAGCACAACCCAGCCAACAATGTTATAAAAAACTTAG

GAGCCAAGGTTGTAGGAGCCATGGACTCACTATCAGTCAATCAGGTTTTGTACCCTGAGGTCCATCTAGATAGCCCTATT

GTCACAAACAAACTAGTTGCTATCCTTGAATACTCGGGGATCAACCACAACTATGTTCTTGAAGACCAGACTCTTATCAA

GAATATCAGATATAGACTGGGGTGCGGTTTTTCAAATCAAATGATCATCAATAATAGGGGGGTAGGTGAAACAGTCAATT

CCAAACTTAAAAGTTACCCCCGTAACTGTCATATCATATACCCAGACTGCAATAAGGAGTTGTTTTGTATCAAAGATAGC

TGCATATCTAGAAAGCTCTCGGAGCTATTCAAGAAGGGTAATTCCTTGTACTCTAAGGTAAGTCACCAGGTACTGGATTG

TCTTAAGAGAGTCAACGGGAAATTAGGGCTGGGCACAGATCTTAATCACGGCCTGAAGGATGGTATCCTCGACTTGGGGT

TGCACATGCATAGCTCTCAATGGTTCGAGACCTTTCTGTTCTGGTTCACTATCAAGACAGAGATGAGATCAATGATCAAA

GAACAGTCCCATATATGCCACAAGAGGAGGTATAACCCAATTTTTGTGTCAGGGGATGCATTCGAGGTGCTTGTATCACG

AGACCTCGTAGTGATAATTGATAAGAACACCCAGTATGTCTTCTACCTGACGTTTGAGCTGGTCCTTATGTATTGTGATG

TCATAGAGGGTAGACTTATGACGGAGACAGCCATGGCTATAGACCAGAGATATTCAGAGCTTCTAAACCGGGTCAGATAC

TTGTGGGACCTTATCGATGGGTTCTTCCCAACACTGGGTAACACCACATACCAAGTTGTTGCTCTGCTTGAACCGCTGTC

GTTGGCTTATCTTCAACTTCAGGATGTCACTCTAGAGTTAAGAGGTGCCTTTTTGGACCACTGCTTCAAAGAACTTTATG

AGATACTGGAGCATTGTGGCATTGACACGGAAGGTACCTACAATTCCATCACTGAGGGATTGGATTACGTATTTATCACT

CACGATATACACTTAACTGGGGAGATTTTTTCATTTTTTCGGAGTTTCGGACACCCCCGCCTCGAAGCGGTCACCGCTGC

AGAGAATGTCAGAAAACATATGAACCAGCCGAAGGTAATCAGTTATGAGACTATGATGAAAGGGCATGCAGTATTTTGCG

GGATAATCATAAATGGTTTTAGGGACCGGCACGGCGGCAGCTGGCCCCCTGTTGCATTGCCAGAACATGCTTCTGCTGCG

ATCCGGAATGCGCAGGCATCCGGCGAAGGACTGACCCACGACCTGTGTATAGACAACTGGAAGTCTTTTGTAGGATTCAG

ATTTGGCTGCTTCATGCCGCTCAGCCTAGATAGTGATTTGACCATGTACCTCAAAGACAAAGCATTGGCTGCACTTAAGA

ATGAGTGGGATTCAGTTTACCCGAAAGAATACCTCCGTTATAATCCACCTAGAGGGACAGAGTCAAGGCGACTAGTAGAG

GTGTTCCTGAATGACTCCAGCTTTGATCCTTATAACATGATAATGTACGTGGTGAATGGCTCCTACCTTAAAGACCCTGA

GTTTAATCTCTCATACAGCTTAAAAGAAAAGGAGATAAAAGAGACAGGGCGGTTGTTTGCCAAGATGACCTACAAGATGC

GGGCCTGTCAGGTGATTGCTGAAAATCTGATATCAAATGGTGTTGGGAAGTATTTCCGAGACAATGGGATGGCGAAAGAC

GAGCATGACCTAACAAAAGCCCTTCACACTCTGGCAGTCTCAGGCGTTCCCAAGAACAACAAAGACAACCACCGAGGTGG

GCCCCCCAGAAGGAACGCAAGCCGAGAGATGAGATCAAGCCAAGCCGTCAACACACAAAATAGAGACAAGATCCAGGGGG

GCCCTATGTACAACTACTTGCGATGCCAACCAACCAGCCCTGATCAGGGTGAGTCATACGAGACTGTTAGTGCATTCATC

ACCGCTGACCTTAAGAAGTATTGCCTAAATTGGAGATACGAGACAATCAGCATATTTGCACAGAGACTGAATGAAATATA

TGGGTTGCCATCCTTCTTTCAATGGTTACATAAGGTATTGGAGAAATCCGTGCTCTACGTCAGTGATCCGCATTGCCCTC

CCGACTTAGATGATCACATCCCTCTGGACAGTGTCCCTAATGCCCAAATATTCATCAAGTACCCGATGGGCGGAATAGAA

GGTTATTGTCAAAAACTATGGACAATCAGTACTATACCATATTTGTATCTAGCAGCCTATGAGAGTGGAGTAAGAATCGC

CTCACTAGTGCAAGGTGACAATCAGACAATTGCAGTGACAAAAAGAGTTCCAAGTTCTTGGCCTTATTCACTAAAAAAGA

GGGAGGCATCTAAAGCAGCTCAAAATTACTTTGTGGTCTTAAGGCAGAGGTTGCACGATGTAGGTCATCACTTAAAGGCT

AATGAGACCATAGTATCTTCTCACTTTTTTGTATACTCTAAAGGGATCTATTATGACGGCCTGTTAGTTTCACAATCACT

AAAAAGCATCGCCAGATGTGTCTTCTGGTCCGAGACCATCGTGGATGAAACCAGAGCGGCCTGCAGCAACATTGCAACAA

CTATCGCCAAGAGTATAGAGAGGGGTTACGATAGGTACCTCGCATACTCTTTGAATATCCTAAAAATTTTTCAACAGATC

CTTGTATCCCTTGACTTCACGATTAATACAACAATGACTCAAGATGTCGTGGCACCGATCATCGAGAACGGCGATTTACT

GATAAGGATGGCACTCTTGCCAGCACCCATTGGGGGTCTCAATTATCTCAACATGAGCAGGTTGTTTGTGAGAAATATCG

GTGACCCGGTCACTTCCTCCATAGCCGACCTGAAGAGGATGATAGACGCTGGGCTAATGCCAGAAGAAACATTGCATCAA

GTGATGACCCAGACCCCGGGAGAATCATCCTACCTTGATTGGGCAAGTGATCCTTATTCTGCCAACCTACCCTGCGTACA

GAGTATAACCCGCCTTCTCAAGAACATCACTGCACGGTATATTTTAATCAGCAGCCCAAATCCGATGCTGAAAGGGTTGT

TTCACGAGGGGAGTAGAGATGAAGACGAGGAGCTTGCGAGTTTCTTAATGGATCGGCATATAATTGTTCCGAGAGCTGCA

CATGAAATCTTAGACCACAGCATAACCGGGGCAAGAGAAGCTATAGCCGGGATGTTGGACACCACCAAGGGTCTGATTAG

AACAAGTATGAAACGGGGTGGCCTCACCCCTCGAGTATTAGCCCGCCTTTCCAATTATGATTATGAACAATTCAGATCCG

GGATCACACTATTGACAAAGAAAGGGCAGTGTTATCTCATTGACAAGGACTCGTGCTCGGTGCAGCTCGCTATAGCTCTG

AGGAGCCATATGTGGGCAAGGTTGGCTCGCGGGAGACCTATCTATGGCTTGGAGGTGCCTGATGTACTGGAATCGATGAA

CGGCTACCTTATCAAACGTCACGAGTCTTGTGCCATCTGTGAAACGGGCTCAAGTCACTACGGGTGGTTTTTCGTCCCTG

CAGGGTGTCAGCTTGACGATGTCTCAAGAGAGACCTCAGCTCTTCGTGTGCCTTATGTCGGATCAACCACTGAGGAAAGG

ACAGATATGAAACTTGCTTTCGTTAGATCTCCAAGCCGATCCCTCAAATCAGCAGTCAGGATTGCCACAGTCTACTCATG

GGCCTACGGGGATGATGAGAAATCATGGAGTGAAGCTTGGATGCTAGCTAGGCAGAGAGCCGATATCACCTTAGATGAAC

TGAGAATGATCACTCCAGTCTCTACATCCACCAACCTAGCTCATCGGTTAAGGGATCGGAGCACTCAGGTGAAATATTCG

GGGACATCCCTTGTGAGGGTTGCAAGGTACACAACCATCTCCAATGACAATTTGTCATTTGTGATATCTGAGAAAAAAGT

AGATACCAACTTCATTTACCAGCAAGGGATGCTGCTCGGTCTTGGGATCCTTGAGAATCTCTTCAGGTTAGAGGCCACCA

CAGGGGTATCCAACACAGTGCTACACCTGCACGTGGAAACAGAATGTTGCGTTGTACCTATGGTTGATCACCCAAGGATA

CCGAGTCTCCGCAATATCAAAGTTACGAACGAGCTATGCACAAACCCTCTTATCTACGACAAGTCCCCCATCATAGAACA

CGATGCAACTCGATTATACTCACAAAGCCACAGGAGACATTTGGTGGAGTTTGTTACCTGGTCAACAAGCCAGCTCTATC

ATATACTGGCCAAATCTACAGCAATGTCCATGATTGAGCTGATCACAAGATTTGAGAAAGATCACATGAATGAAATAGCC

GCTCTGATTGGCGATGACGACATCAACAGTTTCATCACAGAATTTTTGCTTGTGGAGCCCAGACTGTTTATAGTTTACCT

CGGCCAGTGTGCTGCCATTAATTGGGCTTTTGATATACATTATCATCGGCCCTCGGGCAAGTACCAGATGGGGGAACTCC

TCTACTCTTTACTCTCTCGGATGAGCAAGGGAGTATATAAGATCTTCACCAATGCTCTGAGCCACCCCAAAGTTTACAAG

AAATTTTGGCGAAGTGGTATAATTGAGCCGGTTCATGGCCCATCCCTAGATACACAGAATTTACATGTCACTGTCTGTGA

CATGATATACGGATCATACGTCACCTATCTGGATCTTTTGCTGAATGATGAGCTGGATGCTTACCCGTATTTGCTCTGCG

AGAGTGATGAGGACGTGGTCACAGACAGGTTCGACAACATTCAAGCCAAACATCTCTGTGTACTGGCCGATGTATACTGC

AGCTCCAAGAGGTGTCCCTCGATAATCGGGATGTCCCCGATAGAAAAATGTACCATCCTCACACATTACATCAAGGGAGA

ATCGGTACAATCCCCGTCCGGGATCTCATGGAACACTGATCCCCTTGTAGTAGATCATTACTCATGCTCTCTGACCTACC

TTCGCCGTGGTTCCATCAAACAAATCAGATTGAGAGTGGATCCTGGGTTTGTGTTCGAGGCGTTGACAGACATCGACTTC

AAACAGCCTCGCAAGGCTAAGTTGGATGTATCGATTGTGGGATTGACTGATTTTTCTCCCCCTTGGGATAACGTCGGTGA

TTTTCTAGGGACTATCAACACATTGAGGCACAATCTGCCCGTCACCGGGACCGGGGTCTCGAACTATGAAGTCCACGCTT

ATCGTAGAATTGGTCTGAATTCATCAGCATGTTATAAAGCTGTAGAGATCTCCACGTTAATAAAGTCATCCTTAGAAGCC

GGGGAGAATGGATTGTTCTTAGGAGAAGGCTCCGGCTCGATGCTGGCTGCGTACAAGGAAGTTCTTAAATTAGCAAACTG

TTATTACAACAGTGGGGTAACAGCAGAGGGTAGGGCCGGACAGAGGGAAATCTCTCCCTATCCCTCAGAGGTGAGCCTGG

TAGAGAGTCAGATGGGGATAGAGAGAAGTGTTAAAGTCTTGTTCAATGGCAAACCTGAAGTAACCTGGGTAGGGACCACC

GATTGCTACAAGTATATAATCAGTAACATTCAGACCTCTAGTCTGGGTTTCATACACTCAGATATCGAGACACTCCCAAC

CAAGGATGCCGTTGAGAAGTTGGAGGAATTTGCCTCTATTCTATCTCTATCCCTAATTTTGGGAAAAATCGGCTCTATTA

CAGTTGTCAAAATTATGCCCATTAGCGGAGATTTTACCCAAGGCTTCATAGCCTATGCCATTCAATATTTCAGGGAGAGC

CTGCTTGCCTATCCGAGATATAGTAACTTCATCTCGACTGAGTGTTACCTTATTATGATAGGATTAAAGGCCAATCGGTT

GATAAACCCAGAAGCCATTAAGCAAAGCATAATCAGGGTGGGGACTAGGACTGCACCAGGACTTGTGAGCCACATATTAT

CAGAGAAACAGAAAGGTTGTATTCAATCTTTTCTGGGTGATCCTTATGTCCAAGGAGACTTCAATAAGCACCTTAAAGCT

CTAACTCCTATTGAGAAAATCCTTGTAAATTGTGGTCTCTCGATCAATGGTACAAAAATCTGTAGGGATCTAATCCACCA

TGATATTGCCTCCGGTCCAGACGGTCTGATGAGCTCCACAATTATTTTATACAGGGAACTGGCCCATTTCAAAGACAATA

TAAGAAGTCAGCACGGCATGTTCCACCCCTACCCAGTATTGGCCAGTAGCAGGCAACGTGAATTGATCCTTCGAATAGCC

AAGAAGTTTTGGGGGTATGTCTTGCTATATTCTGATGACCCGGCACTAATCAAACAAACGATCAAGAGCTTGAAGCGGAA

TCACCTAACCTTTGACTTACACAGCAATCCGTTTATTAAGGGCTTATCTAAAGCTGAGAAACTGCTAGTGCGGACAAGTT

CACTCAGAAGGGAATGGTTGTTCACTCTCGATACGAAAGAAGTGAAAGAGTGGTTCAAATTGGTGGGTTACAGTGCACTC

GTCAGAGGTTAATTCGCGATACATCTGCCCCCTTCTCCTCCACCATGAGACTCTACTGGCAATCCAAAAGATTAAAGAAA

ACTACATATTGGATAAGTATCTATTCCCAGCTTTGTCTGGT

>KR781449.1/sheep/Benin/2011-05-21

ACCAAACAAAGTTGGGTAAGGATAGATCTTACAATGACTATAGACTGGCAAACTTAGGAGTAAAGATCCTACTGTCGGGG

AGAGGAGGAGGAGCAAGATCTTTGACCATGGCGACTCTCCTCAAAAGCTTAGCACTGTTCAAGAGGAACAAAGACAAAGC

GCCTACTGCGTCAGGTTCAGGAGGGGCCATCCGGGGGATTAAGAATGTTATCATAGTCCCTATCCCCGGGGACTCATCCA

TCATTACCCGTTCAAGACTGCTCGACAGGCTTGTCAGACTGGCCGGAGATCCTGACATCAACGGGTCAAAGCTGACCGGC

GTGATGATCAGCATGTTATCTTTGTTCGTGGAGTCACCCGGGCAATTGATACAGCGGATCACAGATGATCCAGATGTTAG

CATCCGTCTTGTTGAGGTAGTTCAAAGTACCAGGTCCCAGTCCGGATTGACCTTTGCATCACGTGGTGCTGATTTGGATA

ATGAGGCAGATATGTATTTTTCAACTGAAGGACCCTCGAGTGGAAGTAAGAAAAGGATCAACTGGTTTGAGAACAGAGAA

ATAATAGACATAGAAGTGCAAGATGCAGAAGAGTTCAATATGTTGTTAGCCTCCATCTTAGCACAAGTTTGGATCCTCCT

GGCCAAGGCGGTTACGGCACCAGATACGGCAGCCGACTCAGAACTGAGAAGGTGGGTTAAATACACACAGCAAAGGAGAG

TGATTGGGGAATTTCGCTTTGACAAAGGGTGGCTGGACGCAGTCCGTAACAGGATTGCAGAAGATCTATCACTCCGGCGG

TTCATGGTATCCCTGATACTTGACATCAAGAGGACCCCCGGCAACAAGCCAAGGATTGCAGAAATGATCTGCGACATTGA

CAACTATATTGTCGAGGCCGGACTCGCAAGTTTCATTCTTACTATCAAGTTTGGTATTGAAACCATGTATCCTGCACTAG

GCCTTCACGAGTTCGCAGGGGAATTGTCCACTATTGAATCCTTGATGAACTTGTATCAACAGCTAGGAGAAGTTGCACCC

TATATGGTAATTCTAGAGAACTCAATTCAGAACAAGTTTAGTGCAGGAGCCTATCCTCTCCTCTGGAGCTATGCGATGGG

TGTCGGAGTCGAGTTGGAGAACTCAATGGGGGGCTTGAACTTTGGCAGGTCATATTTTGACCCGGCCTATTTTCGTCTCG

GACAGGAGATGGTTAGGAGATCTGCAGGAAAGGTCAGCTCTGTAATTGCGGCTGAGCTTGGCATCACAGCAGAGGAAGCC

AAACTAGTCTCGGAAATCGCCTCGCAGACTGGGGAAGAACGAACCGTTAGAGGGACTGGGCCTCGGCAGGCGCAGGTTTC

CTTCCTCCAGCACAAAATGGGTGAGGGAGAGTCGCCTACACCAGCGACCAAAGAAGAAGTCAAAGCTGCGATCCCAAACG

GGTCCGAAGGAAGGGACATAAAGCGAACACGCCCAGGGAAGCCCAGAGGAGAAACTCCCGGCCAACTGCTTCTGGAGATC

ATGCCAGAGGATGAAGTCTCGCGAGAGTCTGGTCAAAACCCTCGTGAGGCTCAAAGATCGGCTGAGGCACTTTTCAGGCT

GCAGGCCATGGCCAAGATTCTGGAGGACCAGGGGGAGGGAGAAGACAACAGTCAGATCTACAACGACAAGGATCTCCTCA

GCTGAGCAGATACACCCTCCGTCCGAATCAGTGACAAGACATCACCTGCCAGTATTATAAAAAACTTAGGACCCAGGTCC

AACCAACCGCACATCGACACCCCAGTCAATCGAGCGGAGACCACCGATGGCAGAAGAACAAGCATATCATGTCAACAAGG

GGCTGGAATGTATCAAGTCCCTCAAAGCCTCTCCCCCGGATCTATCCACCATCCAAGATGCCCTTGAGAGCTGGAGAGAG

GGGTTTAACCCCTCAGGCCGTGCAACACCGAACCCTGATACGTCCGAGGGGGACCATCAGAATATCAACCAATCATGCTC

ATCAGCAATCGGATCAGACAAAGTCGACATGTCTCCTGAAGGTAATCTCGGATTTAGAGAGATCACTTGTGATGACAATG

AGACTGGGCTCGGAGGAGTTCAGGACAAAGGATCCGACTCTCAAGTACAGCGTTACTATGTTTATAACCACGGGGGTGAA

GAGATTGAAGGACTCGAGGATGCTGACTCTCTCGTGGTTCAAGCAAATCCTCCGGTTGCTAACACCTTCGATGGAGGAGA

GGATGGATCTGACAACAGCGATGTGGACTCTGGCCCAGATGATCCCGGCAGAGATCCTCTATATGACCGGGGACCTGCTG

CCGGCAATGATGTCTCTAGGTCAACAGATGTCGAAAAATTAGAAGGTGATGACATTCAAGAAGTTCTTAACTCCCAGAGG

AGTAAAGGAGGAAGATTCCAGGGAGGGAAAATCTTGCGAGTCCCGGAAATACCCGATGTCAAGAACTCCAGACCATCGGC

CCAATCAATTAAAAAGGGCACAGACGGGAGCTCAGTCTTATCTGGAACGGTGACAGAGTATTCATCGATAAGTGGTGCAA

CCCAAGCTGTGCTAGAGTCCAGATGGGAGTCATCAGAGCGAAATGCATCTGTGGGGAGTGTCCCCAAATCTGCGAGGAGT

GCAGAGACGATCCAGGGGTTAACACAAGAATCTGGTACCATAGCATCACTGACTCAGCCTAAAGAAAATGACTCCGAGTA

TGAGTATGAGGATGAGTTATTCACTGAGATGCAGGAGATTCGTGCAAGCATTGCTAAGATCCATGATGACAACAAAACTA

TCCTCTCAAAACTTGACTCTCTGCTGTTACTGAAAGGAGAAATCGACACTATCAAGAAACAAATCAGCAAACAAAATATA

AGTATATCTACCATTGAGGGCCATCTATCCAGTATAATGATAGCCATCCCAGGTTTTGGGAAGGACATCAAGGACCCAAC

ATCTGAGGTTGAGTTGAACCCAGATTTGAGACCTATAATCAGCCGTGATTCTGGCAGGGCTCTTGCGGAGGTCCTTAAGA

AACCCGCTGTTGATAGGTCTCAGAAGAGCGGAATCAAAGCCAACTCCGGTTCAAAGGGTCAGATCCTCAAGGATCTCCAG

CTAAAGCCCGTCGACAAGCAGGCAAGCTCTGCAATCGGGTTCGTTCCATCCGACCATGAATCATCCAGAAGTGTCATCCG

CTCCATAATCAAGTCAAGCAAGCTTAACATTGATCACAAAGACTATCTTCTAGATTTACTGAATGATGTGAAAGGATCTA

AGGATCTCAAGGAATTCCACAAGATGCTAACAGCCATTCTTGCCAAGCAGCCGTAACACATTCCACAATCTATATCTCAT

ACTCGGTTGAAAGCATCCTCTCAACCAGGCTATTACAAAAAACTTAGGAGCAAGGGCAACTGAGCTTCACAGACAGGATG

ACCGAGATCTATGATTTCGATAAATCAGCATGGGATGTCAAAGGGTCAATTGCTCGCATAGAACCCACCACCTATCACGA

CGGCCGACTGGTACCCCAGGTGAGGGTCATCGACCCTGGTCTGGGAGACAGAAAAGATGAGTGCTTCATGTATCTGTTTC

TTCTAGGAGTGATTGAGGATAACGACCCCCTTTCTCCCCCAGTCGGGAGAACCTTCGGGTCTTTACCTCTAGGTGTCGGT

AGGTCAACTGCTAAGCCAGAGGAACTACTAAGGGAGGCCACAGAACTAGATATAGTGGTGAGGCGCACGGCAGGACTAAA

TGAGAAACTGGTATTTTACAACAACACTCCGCTATCTTTGTTAACACCCTGGAGGAAAGTCTTGACGACCGGGAGTGTGT

TCAGCGCCAACCAGGTTTGCAATGCGGTCAACCTAGTCCCACTCGATACTCCCCAGAGATTCAGGGTTGTGTACATGAGT

ATAACTAGATTGTCAGATAATGGTTATTATAGTGTCCCCAGAAGAATGTTGGAGTTCCGCTCAGCCAATGCAGTCGCTTT

CAATATCTTGGTTACACTGAGAATTGAAAATGGCACAAACCCTAGAAGATACATAGTCGGCTCATGGGAGAATCCGGAGG

TCACATTTATGGTACACGTGGGCAACTTTAGAAGAAAGAAGAACGAAGTATACTCTGCTGATTATTGCAAAATGAAGATT

GAAAAGATGGGTCTAGTTTTTGCCTTGGGAGGAATAGGCGGAACAAGTCTCCATATTCGAAGCACAGGGAAAATGAGCAA

GACCCTCCATGCACAGCTGGGGTTCAAGAAAATTCTATGCTATCCTCTGATGGATATCAATGAGGATCTTAACCGATATC

TCTGGCGGGCAGAGTGCCGAATAGTCAAAATCCAGGCCGTCTTACAGCCGTCAGTACCCCAAGAATTCCGTGTCTATGAT

GATGTCATCATCAACGATGACCAAGGCCTGTTCAAGATCCTGTAAGTCACCTGCAGGGCCACGGACCCACCCCCAGGAAG

AGGTCCTCCCACCTCCCGCCCAAAACACACACTCCAAGACTCCCCGAAACAGCGCCGGCACAGCCACTCTCCCGGCGGGA

CGAGGCCGACC-CCCCCAACAAACCCGCCCCGGAGGGGGG-CCCC-CCCACGACCCGCCGGGCGCCGACCGTGCGAGCCG

GCGAGACACCCTCAAGAGGCCTCCACGAAGACCGGGCAACCCCCTCCCCA-GAAACGGCCCCCC-AACACACCCAAGACC

ACGCGCTGAGCACCCCCAGCCCCCCAACCGGCGGGACAGGAGGCCAGGACGGGGCTCCCCCAAACAAGGGGACCACCAAG

AAAGGCTCCGCCGACCCAGCACAAACCCGGGCCAAACAAGGCAGACCCCCAAGACGAAATCCCCCAGAGCACCCCATTCA

TCATCAAACAAAACTTAGGGGCCAGGTCCAGACGACAGACCCCAGCCCGCCAGGCTGCAAGCCACAGAAAAGAAGGAGAC

AAGATCGGCCCGGGAAGAGGACACCCAACCACAGAAACCCCCAGCTCCAGGGGAGGAG-CCCCCCCTCCACACGGCGGAG

GAACGGGCAAGAGACACCGAGGAGAACCCGATCGGGGGCCAAGAACCGGAGACGCCGGTGGCAAATAGGCCGGGCACTCA

ACCGACACCACCGGAGGACCCGGAGAGAGCCCGACTCCGCGCAGACTCGACGGCAGAGGACCCAGTCCCGCCACAAAGAG

AGGCGAGAACCCT--------CCCCTCCGCCCACAACCGGCGAACAGACCGCGGGAGGACCAACCCCCAGCAAGACAGAC

GCCTCCCCCGCAGCCGAGCCGCAGCTCCCCAGGAGAGGGCACCCCCGAAACCCCCCAGAACCCCTCCGCCCACGGCCGCC

GGGACAAGAGATCCGCCGCCAAAGACCAGAACCACCGAGAGTCAACCGGGGGCCCCCCCCAGTCCCCCCCGGCGACCCCC

CACAGAGGCCCCAGCAAGGCCCCCCGGCCCGACGACCCCCGCATGCGCCGAAGGGGAAACCCACTAGAGAACGTCCAGCA

CATCCGTGTATAAACATCATGACACGGGTCGTAACCTTGGTATTTCTGTTTCTTTTCCCAAACACTGTCTCGTGCCAGAT

TCACTGGGGCAATCTATCTAAGATAGGGATTGTAGGAACAGGGAGTGCCAGTTACAAGGTGATGACTAGGCCAAGCCACC

AAACTCTGGTTATAAAGTTGATGCCAAATATAACAGCCATCGACAATTGTACGAAATCAGAGATTTCAGAGTACAAAAGA

TTGCTAATCACAGTGTTAAAGCCTGTAGAGGATGCTCTGTCAGTAATAACCAAGAATGTAAGACCAATTCAAGCTCTAAC

ACCTGGGCGCAGGACCCGCCGTTTCGCCGGAGCTGTTCTGGCCGGAGTAGCACTTGGAGTCGCGACGGCTGCTCAAATAA

CTGCCGGAGTCGCACTCCATCAGTCATTGATGAATTCCCAAGCAATTGAAAGTTTAAAAACCAGTCTTGAGAAGTCAAAT

CAGGCAATAGAAGAAATCAGACTTGCAAATAAGGAGACCATACTGGCAGTACAGGGCGTCCAAGACTATATCAACAACGA

GCTTGTCCCCTCTGTTCATAGAATGTCATGTGAGCTAATAGGTCACAAACTCAGTCTCAAACTCCTTAGGTATTATACCG

AGATCCTGTCTATATTCGGGCCTAGCCTTCGAGACCCGATAGCTGCTGAAATATCAATCCAGGCACTCAGCTATGCACTA

GGCGGAGACATCAATAAAATTCTGGACAAGCTTGGGTATAGCGGCGGGGATTTCCTTGCTATTCTAGAAAGCAAGGGGAT

AAAGGCCCGTGTCACATATGTGGACACAAGAGATTACTTTATAATTCTTAGCATAGCCTACCCAACCTTATCTGAGATCA

AAGGGGTGATAGTTCATAAGATAGAAGCTATATCATACAACATTGGGGCACAGGAATGGTATACTACTATCCCTAAATAT

GTAGCCACTCAGGGATATCTGATATCAAATTTCGATGAGACGTCATGCGTCTTCACTCCAGAGGGGACAGTCTGCAGCCA

GAATGCGTTGTATCCAATGAGCCCATTGCTTCAGGAATGTTTCAGGGGGTCGACAAAATCGTGTGCCAGAACCCTAGTTT

CAGGGACCACAAGCAATAGATTTATCCTATCAAAAGGGAACTTGATTGCAAATTGTGCGTCAGTTTTGTGCAAGTGTTAC

ACAACGGAGACAGTCATCAACCAAGATCCTGATAAACTACTAACTGTTATAGCCTCAGATAAGTGTCCCGTAGTTGAGGT

GGATGGAGTGACAATACAAGTCGGCAGTCGAGAGTACCCAGATTCTGTATACCTACATAAAATAGACTTAGGCCCAGCCA

TCTCTCTGGAAAAACTGGACGTAGGCACCAATTTAGGCAATGCAGTCACAAGACTGGAGAATGCAAAGGAGCTCCTAGAC

GCATCAGACCAGATACTGAAGACTGTTAAAGGGGTACCCTTCAGTGGCAATATGTACATAGCACTGGCAGCTTGCATTGG

GGTATCCCTAGGGCTTGTCACATTAATATGCTGCTGTAAGGGGAGATGTAGGAACAAGGAGATTCCTGCCTCCAAAATCA

ACCCAGGGCTCAAACCCGACCTGACCGGGACTTCAAAGTCGTACGTGAGATCACTGTAGTCAGAATAACCCGAATCATCC

GGCATCACGCGTATACATGTGCGACACAAGCAGTCAGAGGACGCAGAAGACTCAACCTCCGATCACCGACCAGACCCCAC

TCTACGCCCTACTACACATTGGTCATCAAACAAAACTTAGGACGAAAGGTCAATCACCATGTCCGCACAAAGGGAGAGGA

TCAATGCCTTCTACAAAGACAACCCTCACAATAAAAACCATAGGATAATCCTGGATAGGGAACGCTTAACTATTGAAAGA

CCCTACATCTTACTTGGAGTCCTGCTGGTAATGTTCCTGAGTCTAATCGGGCTTCTAGCCATTGCAGGGATCAGGCTTCA

CCGGGCCACCGTTGGGACTGCGGAGATCCAGAGTCGGCTGAATACCAACATTGAGTTGACCGAATCCATTGATCATCAAA

CTAAGGATGTCTTAACACCCCTGTTTAAAATCATTGGTGATGAAGTCGGCATCAGAATTCCACAGAAGTTTAGTGATCTT

GTCAAGTTCATCTCCGATAAGATTAAGTTCCTCAACCCTGACAGAGAATACGATTTTAGGGATCTCCGGTGGTGTATGAA

CCCTCCTGAGAGAGTCAAAATTAACTTTGATCAATTCTGTGAATACAAAGCCGCGGTCAAGTCAGTTGAACATATATTTG

AGTCATCATTCAACAGGTCAGAAAGATTGCGACTATTGACTCTTGGGCCCGGAACAGGCTGTCTCGGCAGGGCAGTAACA

AGAGCTCAGTTCTCAGAGCTTACGCTGACCCTGATGGACCTGGATCTCGAGATGAAGCACAACGTGTCCTCAGTGTTTAC

CGTAGTTGAAGAGGGGTTATTCGGAAGAACATATATTGTCTGGAGATCTGACACCAGGAAACCGAGCACCAGTCCAGTTA

TTGGCCAGTTTTTAAGAGTCTTCGAGATCGGGTTGGTGAGAGATCTCGAGCTGGGTGCCCCCATTTTCCATATGACCAAC

TACCTCACAGTGAACATGAGTGATGACTATCGGAGTTGCCTTTTAGCGGTAGGGGAGTTGAAGCTGACAGCCCTATGCAC

TCCATCTGAGACTGTGACTCTGAGTGAGAGAGGAGTTCCAAAGAGAGAGCCTCTTGTGGTTGTGATACTCAACCTAGTTG

GGCCTACTCTAGGGGGCGAACTATACAGTGTCTTGCCTACCTCTGACCTCATGGTGGAGAAACTCTATTTATCCTCACAT

AGAGGGATAATCAAAGATAACGAGGCCAATTGGGTAGTGCCGTCTACCGATGTTCGTGATCTCCAAAACAAAGGGGAATG

TCTGGTGGAAGCATGCAAAACTCGACCTCCTTCATTTTGCAATGGCACAGGAATAGGCCCATGGTCAGAGGGGAGAATCC

CTGCCTACGGAGTGATCAGGGTCAGTCTTGACTTAGCTAGTGACCCAGGTGTGGTTATCACTTCAGTGTTTGGCCCACTG

ATACCTCACCTATCTGGCATGGATCTTTACAACAATCCGTTTTCAAGAGGTGCATGGCTGGCTGTACCACCCTATGAGCA

GTCATTTCTAGGAATGATAAATACAATTGACTTCCCGGACAGAGCAGAGGTCATGCCGCACATTTTGACCACAGAGATCA

GAGGGCCTCGAGGTCGTTGCCATGTTCCTATAGAGTTGTCCCGCAGGATTGATGATGATATCAAGATCGGGTCCAATATG

GTTGTATTGCCGACGAGGGATCTGAGGTACGTAACAGCCACTTATGATGTTTCCAGGAGCGAGCATGCAATCGTGTACTA

TATCTATGACACGGGTCGCTCATCATCTTACTTCTACCCAGTTCGATTGAATTTCAGGGGCAATCCTCTCTCTCTGAGGA

TAGAGTGTTTCCCCTGGTCTCATAAGGTGTGGTGCTACCATGATTGTCTTATATACAACACCATGACAAACGAAGAAGTC

CATACGAGAGGGCTGACCGGTATAGAGGTAACATGCAATCCAGTCTGAGTCGAGCTGAAACCATCGCTCAAGCAGGCTTC

CGAGCCATCCCCTAGTCCAAGCAGCATAGTCTGGGACACTCAGCAGCACAACCCAGCCAACAATGTTATAAAAAACTTAG

GAGCCAAGGTTGTATGAGCCATGGACTCACTATCAGTCAATCAGGTTTTGTACCCTGAGGTCCATCTAGATAGCCCTATT

GTTACAAACAAACTAGTTGCTATCCTTGAATACTCGGGGATCAACCACAACTATGTTCTTGAAGACCAGACCCTTATCAA

GAATATTAGATATAGACTGGGGTGCGGTTTTTCAAATCAAATGATCATCAATAATAGGGGGGTAGGTGAAACAGTCAATT

CCAAACTTAAAAGTTACCCCCGTAACTGTCATATCATATACCCAGACTGCAATAAGGAGTTGTTTTGTATCAAAGATAGC

TGCATATCTAGAAAGCTCTCGGAGCTATTCAAGAAGGGTAATTCCTTGTACTCTAAGGTGAGTCACCAGGTACTGGATTG

TCTCAAGAGAGTCAACGGGAAACTAGGGCTGGGCACAGATCTTACTCACGGCCTGAAGGATGGTATCCTCGACTTGGGGT

TGCACATGCATAGCTCTCAATGGTTCGAGACCTTTCTGTTCTGGTTCACTATCAAGACAGAGATGAGATCAATGATCAAA

GAACAGTCCCATATATGCCACAAGAGGAGGTATAACCCAATTTTTGTGTCGGGGGATGCATTCGAGGTGCTTGTATCACG

AGACCTCGTAGTGATAATTGATAAGAACACCCAGTATGTCTTCTACCTGACATTTGAGCTGGTCCTTATGTATTGTGATG

TCATAGAGGGCAGACTTATGACGGAGACAGCCATGGCTATAGACCAGAGATATTCAGAGCTTCTAAACCGGGTCAGATAC

TTGTGGGACCTTATCGATGGGTTTTTCCCAACACTGGGTAACACCACATACCAAGTTGTTGCTCTGCTTGAACCACTGTC

GTTGGCTTATCTTCAACTTCAGGATGTCACTCTAGAGTTAAGAGGTGCCTTTTTGGACCACTGCTTCAAAGAACTTTATG

AGATACTGGAGCATTGTGGCATTGACACGGAAGGTACCTACAATTCCATCACTGAGGGATTGGATTACGTATTTATCACT

CACGATATACACTTAACTGGGGAGATTTTTTCATTTTTCCGGAGTTTCGGACACCCCCGTCTCGAAGCGGTCACCGCTGC

AGAGAATGTCAGAAAACATATGAACCAACCGAAGGTAATCAGTTATGAGACTATGATGAAAGGGCATGCAGTATTTTGCG

GGATAATCATAAATGGTTTTAGGGACCGGCACGGCGGCAGCTGGCCCCCTGTTGCATTGCCAGAACATGCTTCTGCTGCG

ATCCGGAATGCGCAGGCATCCGGCGAAGGACTGACCCACGACCTGTGTATAGACAACTGGAAGTCCTTTGTAGGATTCAG

ATTTGGCTGCTTCATGCCACTCAGCCTAGATAGTGATTTGACCATGTACCTCAAAGACAAAGCACTGGCTGCACTGAAGA

ATGAGTGGGATTCAGTTTACCCGAAAGAATACCTCCGTTATAATCCACCTAGAGGGACAGAGTCAAGGCGACTAGTAGAG

GTGTTCCTGAATGACTCCAGCTTTGATCCTTATAACATGATAATGTACGTGGTGAATGGCTCCTACCTTAAAGACCCTGA

GTTTAATCTCTCATACAGCTTAAAAGAGAAGGAGATAAAAGAGACAGGGCGGTTGTTTGCCAAGATGACCTACAAGATGC

GGGCCTGTCAGGTGATTGCTGAAAATCTGATATCAAATGGTGTTGGGAAGTATTTCCGAGACAATGGGATGGCAAAAGAC

GAGCATGACCTAACAAAAGCCCTTCACACTCTGGCAGTCTCGGGCGTTCCCAAGAACAACAAAGACAACCACCGAGGCGG

GCCCCCCAGAAGGAACGCAAGCCGAGAGACGAGATCAAGTCAAGCCGTCAACACACAAAATAGAGACAAGATCCAAGGGG

GCCCTATGTACAACTACATGCGATGCCAACCAACCAGCCCTGATCAGGGTGAGTCATACGAGACTGTTAGTGCATTCATC

ACCGCTGACCTTAAGAAGTATTGCCTAAATTGGAGATACGAGACAATCAGCATATTTGCACAGAGACTGAATGAAATATA

TGGGTTGCCATCCTTCTTTCAATGGTTACATAAGGTATTGGAAAAATCCGTGCTCTACGTCAGTGATCCGCATTGCCCTC

CCGACTTAGATGATCATATCCCTCTGGACAGTGTCCCTAATGCCCAAATATTCATCAAGTACCCAATGGGCGGAATAGAA

GGTTATTGTCAAAAATTATGGACAATCAGTACTATACCATACTTGTATCTAGCAGCCTATGAGAGTGGAGTAAGAATCGC

CTCACTAGTGCAAGGTGACAATCAGACAATTGCAGTGACAAAAAGAGTTCCAAGTTCTTGGCCTTATTCACTAAAAAAGA

GGGAGGCATCTAAAGCAGCTCAAAATTACTTTGTGGTCTTAAGGCAAAGGTTGCACGATGTAGGTCATCACTTAAAGGCT

AATGAGACCATAGTATCTTCTCACTTTTTTGTATACTCTAAAGGGATTTATTATGACGGCCTGTTAGTTTCACAATCACT

AAAGAGCATCGCCAGATGTGTCTTCTGGTCCGAGACTATCGTGGATGAAACTAGAGCGGCCTGCAGCAATATTGCAACAA

CTATCGCTAAGAGTATAGAGAGGGGTTATGATCGGTACCTCGCATACTCTTTGAATATCCTCAAAATTTTCCAACAGATC

CTTGTATCCCTTGACTTCACGATTAATACAACAATGACTCAAGATGTCGTGGCACCGATCATTGAGAACGGTGATTTACT

GATAAGAATGGCACTCTTGCCAGCACCCATTGGGGGTCTCAATTATCTCAACATGAGCAGGTTGTTTGTGAGAAATATCG

GTGACCCGGTCACTTCCTCTATAGCCGACCTGAAGAGGATGATAGACGCTGGGCTAATGCCAGAAGAAACATTGCATCAA

GTGATGACCCAGACCCCGGGAGAATCATCCTACCTTGATTGGGCAAGTGATCCTTATTCTGCCAACCTACCCTGCGTACA

GAGTATAACTCGCCTTCTCAAGAACATCACTGCACGGTACATTTTAATCAGCAGCCCAAACCCGATGCTGAAAGGGTTGT

TTCACGAGGGGAGTAGGGATGAAGACGAGGAGCTTGCGAGTTTCTTAATGGATCGGCATATAATTGTTCCGAGAGCTGCA

CATGAAATCTTAGACCACAGCATAACCGGGGCAAGGGAAGCTATAGCCGGGATGTTGGACACCACCAAGGGTCTGATTAG

AACAAGTATGAAACGGGGTGGCCTCACTCCTCGAGTATTGGCCCGCCTTTCCAATTATGATTATGAACAATTCAGATCCG

GGATAACACTATTGACAAAGAAAGGGCAGTGTTATCTCATTGACAAGGACTCGTGCTCGGTGCAGCTCGCTATAGCTCTG

AGAGGCCATATGTGGGCAAGATTGGCCCGCGGGAGACCTATCTATGGCTTGGAGGTGCCTGATGTACTGGAATCGATGAA

CGGCTACCTTATCAAACGTCACGAGTCTTGTGCCATCTGTGAAACGGGCTCAAGTCACTACGGGTGGTTTTTCGTCCCTG

CAGGGTGTCAGCTTGACGATGTTTCAAGAGAGACTTCAGCTCTTCGTGTACCTTATGTCGGATCAACCACTGAGGAAAGG

ACTGATATGAAACTTGCTTTCGTTAGATCTCCAAGCCGATCCCTCAAATCAGCAGTCAGGATTGCCACAGTCTACTCATG

GGCCTACGGGGATGATGAGAAATCATGGAGTGAAGCTTGGATGCTAGCTAGGCAGAGAGCCGATATCACCTTAGATGAAC

TGAGAATGATCACTCCGGTCTCTACATCCACCAACCTAGCTCATCGGTTAAGGGATCGGAGCACTCAGGTGAAATATTCG

GGGACATCCCTTGTGAGGGTTGCAAGGTACACAACCATCTCCAATGACAATTTGTCATTTGTGATATCTGAGAAAAAAGT

AGATACTAACTTCATTTACCAACAAGGGATGCTGCTCGGTCTTGGGATCCTTGAGAATCTCTTCAGGTTAGAGGCCACCA

CAGGGGTATCCAACACAGTGCTACACCTGCACGTGGAAACAGAATGTTGTGTTGTACCTATGATTGATCACCCAAGGATA

CCGAGTCTCCGCAATATCAAAGTTACAAACGAGCTATGCACAAACCCTCTTATCTACGACAAGTCCCCCATCATAGAACA

CGATGCAACTCGATTATACTCACAAAGCCATAGGAGACATTTGGTAGAGTTTGTTACCTGGTCAACAAGCCAGCTCTACC

ATATACTGGCCAAATCTACAGCAATGTCCATGATTGAGCTGATCACAAGATTTGAGAAAGATCACATGAATGAAATAGCC

GCTCTGATTGGTGATGACGACATCAACAGTTTCATCACAGAATTTTTGCTTGTGGAGCCCAGACTGTTTATAGTTTACCT

CGGCCAGTGTGCCGCCATCAATTGGGCTTTTGATATACATTATCATCGGCCCTCGGGCAAGTACCAGATGGGGGAACTCC

TCTACTCTTTACTCTCTCGGATGAGCAAGGGAGTATATAAGATCTTCACTAATGCTCTGAGCCACCCCAAAGTTTACAAG

AAATTTTGGCGAAGTGGTATAATTGAGCCGGTTCATGGCCCATCCCTAGATACACAGAATTTACATGTCACTGTCTGTGA

CATGATATACGGATCATACGTCACCTATCTGGATCTTTTGCTGAATGATGAGCTGGATGCTTACCCGTATTTGCTCTGCG

AGAGTGATGAGGACGTGGTCACAGACAGGTTCGACAACATTCAAGCCAAACATCTCTGTGTACTGGCCGATGTATACTGC

AGCTCCAAGAGATGTCCCTCGATAATCGGGATGTCCCCTATAGAAAAATGTACCATCCTCACACATTACATCAAGGGAGA

ATCGGTACAATCCCCGTCCGGGATCTCATGGAACACTGATCCCCTTGTAGTAGATCATTACTCATGCTCTCTGACCTACC

TTCGCCGTGGTTCCATCAAACAAATCAGATTGAGAGTGGATCCTGGGTTTGTGTTCGAGGCCTTGACAGACATCGACTCC

AAACAGCCTCGCAAGGCTAAGTTGGACGTATCGATTGTGGGGTTGACTGATTTTTCCCCCCCTTGGGATAACGTCGGTGA

TTTTCTAGGGACTATCAACACATTGAGGCACAATCTGCCCGTCACCGGGACCGGGGTCTCGAACTATGAAGTCCACGCTT

ATCGTAGAATTGGTCTGAATTCATCAGCATGTTATAAAGCTGTAGAGATCTCCACGTTAATAAAGTCATCCTTAGAGGTC

GGGGAGAATGGATTGTTCTTAGGAGAAGGCTCCGGCTCGATGCTGGCTGCGTACAAGGAAGTTCTTAAATTGGCAAACTG

TTATTACAACAGTGGGGTAACAGCAGAGGGTAGAGCCGGACAGAGGGAAATCTCTCCCTATCCCTCAGAGGTGAGCCTGG

TAGAGAGTCAGATGGGGATAGAGAGAAGTGTTAAAGTCCTGTTCAATGGCAAACCTGAAGTAACCTGGGTAGGGACCACC

GATTGCTACAAGTATATAATCAGTAACATTCAGACCTCTAGTCTGGGTTTCATACACTCAGATATCGAGACACTCCCAAC

CAAGGATGCCGTTGAGAAGTTGGAGGAATTTGCCTCTATTCTATCTCTATCCCTAATTTTGGGAAAAATCGGCTCTATTG

CAGTTGTCAAAATTATGCCCATTAGCGGAGATTTTACCCAAGGCTTCATAGCCTATGCCATTCAATATTTCAGGGAGAGC

CTGCTTGCCTATCCGAGATATAGTAACTTCATCTCGACTGAGTGTTACCTTATTATGATAGGATTAAAGGCCAATCGGTT

GATAAACCCAGAAACCATTAAGCAAAGCATAATCAGGGTGGGGACTAGGACTGCACCAGGACTTGTGAGCCACATATTAT

CAGAGAAACAGAAAGGTTGTATTCAATCTTTTCTGGGTGATCCTTATGTCCAAGGAGACTTCAATAAGCACCTTAAAGCT

CTAACTCCTATTGAGAAAATCCTTGTAAATTGTGGTCTCTCGATCAATGGTACAAAAATCTGTAGGGATCTAATCCACCA

TGATATTGCCTCCGGTCCAGACGGTCTGATGAGCTCCACAATTATTTTATACAGGGAACTGGCCCATTTCAAAGACAATA

TAAGAAGTCAGCACGGCATGTTCCACCCCTACCCAGTATTGGCCAGTAGCAGGCAACGTGAATTGATCCTTCGAATAGCC

AAGAAGTTTTGGGGGTATGTCTTGCTATATTCTGATGACCCGGCACTAATCAAACAAACGATCAGGAACTTGAAGCGGAA

TCACCTAACCTTTGACTTACACAGTAATCCGTTTATTAAGGGGTTATCCAAAGCTGAGAAACTGCTAGTGCGGACAAGTT

CACTCAGAAGGGAATGGTTGTTCACTCTCGACACGAAAGAAGTGAAAGAGTGGTTCAAATTGGTGGGTTACAGTGCACTC

GTCAGAGGTTAATTCGCGATACATCTGCCCCCTTCTCCTCCACCATGAGACTTTACTGGCAATCCAAAAGATTAAAGAAA

ACTACATATTGGATAAGGATCTATTCCCAGCTTTGTCTGGT

>Saiga4/saiga/Mongolia/2017-01

ACCAAACAAAGTTGGGTAAGGATAGATCTTATAATAACTATGGACTGGCAAACTTAGGAGTAAAGATCCTACTGTCGGGG

GGAGGAGGAGGAGCAAGATCCTTGATCATGGCGACTCTCCTTAAAAGCTTAGCATTGTTCAAAAGGAACAAAGACAAAGC

GCCGACAGCATCAGGTTCAGGAGGGGCCATCCGGGGGATTAAGAATGTTATCATAGTCCCGATTCCCGGAGACTCGTCCA

TCACTACCCGTTCAAGACTGCTCGACAGGCTTGTCAGATTGGCCGGAGATCCTGACATCAACGGGTCAAAGCTGACCGGG

GTGATGATCAGCATGTTATCCTTGTTCGTAGAGTCACCCGGGCAATTGATACAGCGAATCACAGATGATCCGGATGTTAG

TATCCGCCTTGTTGAGGTAGTTCAAAGTACTAGGTCCCAGTCCGGGTTGACCTTTGCATCACGTGGTGCTGATTTAGACA

ACGAGGCAGACATGTATTTTTCAACTGAGGGGCCCTCGAGTGGAGGTAAGAAAAGGATCAACTGGTTTGAGAACAGAGAA

ATAATAGACATAGAGGTGCAGGATCCAGAAGAGTTCAATATGTTGTTAGCCTCCATACTAGCACAAGTCTGGATCCTTCT

GGCCAAGGCTGTTACGGCACCAGATACGGCAGCTGACTCAGAACTGAGAAGGTGGGTTAAATACACACAACAAAGGAGAG

TGATTGGGGAATTTCGCCTTGACAAGGGGTGGCTGGACGCAGTCCGCAACAGAATTGCAGAAGATCTATCACTCCGGCGG

TTCATGGTATCTCTCATACTTGACATCAAGAGGACCCCAGGCAACAAGCCAAGGATTGCAGAAATGATCTGCGACATTGA

CAACTATATTGTCGAAGCAGGACTCGCCAGTTTCATCCTTACTATCAAATTTGGTATCGAAACCATGTATCCTGCATTAG

GGCTCCACGAGTTTGCCGGAGAATTGTCCACTATTGAGTCCTTGATGAACTTGTATCAACAGCTAGGAGAGGTTGCACCC

TACATGGTAATTCTAGAGAATTCAATTCAGAACAAGTTTAGTGCAGGAGCTTATCCCCTCCTCTGGAGCTATGCGATGGG

TGTCGGAGTCGAGCTGGAGAACTCAATGGGGGGCTTGAACTTTGGCAGGTCATATTTTGACCCTGCCTATTTTCGTCTCG

GACAGGAGATGGTCAGAAGATCTGCAGGAAAGGTCAGCTCTGTGATTGCGGCTGAGCTCGGCATCACAGCAGAGGAAGCT

AAACTAGTCTCGGAAATCGCCTCGCAGGCCGGGGACGAAAGAACCGCCAGAGGGACTGGGCCTCGACAGGCGCAGGTTTC

CTTCCTTCAGCACCAAACAGGAGGGGGAGAGTCGTCCGCACCAGCGACCAGAGAAGGGGTCAAAGCTGCGATCCCAAACG

GAGCTGAAGAAAGGGACAGAAAACAAACACGCCCAGGAAGGCCCAGAGGAGAGACCTCCGGCCAACTGCTCCTGGACATC

ATGCCAGAGGATGAGATCTTGCGAGAGTCTGGTCAAAACCCTCGTGAGGCTCAAAGATCGGCCGAGGCACTCTTCAGGCT

GCAGGCCATGGCCAAGATTCTGGAGGACCAGGAGGAGGGAGAAGACAACAGTCAGGTCTACAACGACAAGGATCTCCTCG

GCTAAGCAGACGCACCCTCTGTCGAAATCAGTGACAAGACATCTCCTACCAGTATTATAAAAAACTTAGGACCCAGGTCC

AAGCAAGCACACATCGACACTCCAACCAGTCGAGCGGAGACCACCGATGGCAGAAGAACAAGCATACCATGTNAACAAGG

GGCTGGAATGTATCAAGTCTCTCAAAGCCTCACCCCCGGATCTATCCACCATCAGAGATACCATCGAGAGCTGGAGAGAG

GGGCTTAGCCCCTCGGGCCGTGCAACACCGAACCCTGATACGTCCGAGGGAGATCATCAGAATATCAACCAATCATGCTC

ACCAGCAATCGGACCAAACAAAGTCTACTTGTCTCCTGAAGATAATCTCGGATTTAGAGAGATCACTGGCAACGACTGTG

AGGCTGGGCTCGGAGGGGTCCAGAGAGAAGGATCCAACTCTCAAGTACAGCGTTACCATGTTTATAGTCACGGGGGTGAA

GAGATTGAAGGACTCGAGGATGCTGACTCTCTCGTGGTTCAAGCAGATCCTCCAGCTGCTAACGTCTTCAATGGAGGAGA

GGATGGATCTGACGACAGCGATGTGGACTCTGGCCCAGATGATCCCGGCAGAGATACTCTATATGACCGGGGATCTGTTG

CCGGCAATGGTATCGCTAGGTCCACAGATGTCGAAAAACTAGAAGGTGCTGATATTCAAGAAGTTCTTAACTCCCAGAAA

GGCAAAGGAGGAAGATTCCAGGGCGGGAAAACCTTGCGAGTCCCGGAAATACCCGATGTCAAGCACTCCAGACCATCAGC

CCAATCAATTAAAAAGGGCACAGACGGGAACTCAGTCTCATCTGGAACGGTGATAGAGTGTTTATCGATAAGTGGTGCAA

CCCAAACTGTGCCAGAGTCAAGATGGGAGTCATCAGAGCAAAATGCGTCTGTGGGGAGTGTCCTCAAGTCTGCGAGGAGT

GCAAAGACGATCCAGGGGCCGACACAAGAATCTGGTACCATAGCATCACTGACTCAGCCTAAAGAGAATGACTCCGAGTA

TGAGTATGAGGATGACCTATTTACAGAGATTCAGGACATCCGTGCAAGCATTGCCAAGATCCATGATGACAATAAAACTA

TCCTCTCAAAGCTTGATTCTATACTGTTATTGAAAGGGGAAGTCGACACTATCAAGAAACAAATCAGCAAGCAGAATATA

AGTATATCCACCATTGAGGGCCATCTCTCCAGTATAATGATAGCCATCCCTGGCTTTGGGAAGGACATCAAGGACCCAAC

ATCCGAAGTCGAGTTGAACCCAGATCTAAGACCTATAATAAGCCGTGACTCTGGCAGAGCTCTCGCGGAGGTCCTCAAGA

AACCCGCTGTGGATAGGTCTCAGAAAATTGGAACCAAAGCCAACTCCAGCTCAAAGGGTCAGCTTCTTAAGGATCTCCAG

CTAAAACCTGTCGACAAGCAGGCAAGCTCTGCAATCGGGTTTGTCCCGTCCGACCATGAATCATCCAGAAATGTCATCCG

CTCCATAATCAAGTCGAGCAAGCTAAATATTGATCACAAGGACTATCTTCTAGATTTACTAAATGATGTGAAAGGCTCCA

AGGATCTTAAGGAATTTCACAAGATGCTAACAGCAATTCTCGCCAAGCACCTGTAACACATCCCCCAGTCATCATCTCAT

ACTCGACTAAAAACATCCTTTCAATCAGGCTATTACAAAAAACTTAGGAGCAAGGGCAACTGAGCTTCGCAGACAGGATG

ACCGAGATCTACGACTTCGATAAATCAGCATGGGATGTCAAAGGGTCAATTGCCCGCATAGAACCCACCACCTACCACGA

CGGCCGACTGATACCCCAGGTGAGGGTCATCGATCCTGGTCTGGGAGACAGAAAAGATGAGTGCTTTATGTACCTGTTTC

TCCTAGGAGTGATTGAGGATAACGACCCCCTGTCTCCCCCCGTCGGAAGAACCTTTGGCTCTCTACCTCTAGGGGTTGGT

AGGTCAACTGCCAAGCCAGAAGAACTACTAAGGGAGGCCACAGAATTAGATATAGTGGTGAGGCGCACTGCAGGAGTAAA

TGAGAAACTGGTATTTTACAACAACACTCCGCTGTCCTTGTTAACGCCATGGAAGAAAGTTCTGACAACCGGAAGTGTGT

TTAGCGCTAACCAGGTTTGCAATGCAGTCAACCTAGTCCCGCTTGATACTCCCCAGAGATTCAGGGTTGTGTACATGAGC

ATAACTAGATTATCAGACAATGGTTACTATAGTGTGCCCAGAAGAATGCTGGAGTTCCGCTCGGCCAATGCAGTAGCCTT

CAACATCTTGGTTACACTAAGAATTGAAAATGGCACAAACCCTAGAAGATACATAGTCGGCTCATGGGAGAATTCAGAGG

TCACATTTATGGTACACGTGGGCAACTTTAGGAGAAAGAAGAACGAAGTATACTCTGCTGATTACTGCAAAATGAAGATT

GAAAAGATGGGTTTAGTTTTTGCCCTGGGTGGAATAGGCGGAACAAGTCTCCATATTAGAAGCACCGGGAAAATGAGCAA

AACCCTCCATGCACAGCTGGGGTTCAAGAAAATCTTATGTTACCCCCTAATGGATGTCAATGAGGATCTTAACCGATATC

TCTGGCGGGCAGAGTGCCGAATAGTCAAAATCCAAGCTGTCTTACAGCCATCAGTACCCCAAGAGTTCCGTGTCTACGAT

GATGTCATCATCAACGATGATCAAGGCTTGTTCAAGATCTTGTAGTTCATTTGCAAAGGAACGAACCCACCCCCAGGAAG

GGGCCCCGCCCTCCCCCG-CCAAAAAACACACTCCAAGGCTCCCCGGCACAGCGCCAACACAGCTACTCCCGCGATGGAA

CGAGGTCGGCC-CCCGAACAAATCCGGTCCAAAGAGGGGGGCCCCCCCCCTGACCAACCGGGCGCCGACCGAGCAGGCCG

GCGAGAAGCCCCCAAGAGGCTCCGACGAAGACCGGGCACCCCATCCCCCC-AAAAACGTCCCCC-AACACGCCCGAGGCT

AGCCACCGGACACCCCCAGACCCGCCACCGGCGAGACAGAACCCCAAGGGGGGACCCTCCAAACCAAAGGAACCACCGGG

GTAGGCTCCGCCGAACCAGCACAGACCCGACCCAAGCAAGGCGGACCCCCAACACGAAACTGCCCAGCACACCCTACTCA

TCATCAAACAAAACTTAGGGGCCAAGTCCACACGAGAGGCCCCAGTCCGTCAGGCGGAGGGACGCAGAAAGGAAGGAGAC

ACGACCGTCCCAGAAGGAGGACACCCAACCACCGAAACGCCCGCCCCCAGGGAAGGAG--CCA-CCCCCACGCAGCGAAG

GAGCGGGCGGGAGACCCCGGGGAGGGCCCGATCGGGACCCGGGGGCCGGCGGCGCCGGCGGTGAGCAGGCCGGGCGCACA

CCCGACCCCACCGGAGGATCCGAGGAGGGCCCGACCCCGCGCGGACCAAACGGCAGGGAGCCCAACCCCACCAAAAAGAG

AGGCGAGGCCCCCCCTCCCTCCCCCCCCGCACACAACCCGCAGACAGACCGAGGGAGGGCCATCCCCCAGCGAGACAGGC

GGCTCCCTCGCAGCCGAGCCGCAGCTCCCCAGAATAGGACACCCCCGCACACTCCGAACACCCCCCCGCCCCCCGCCACC

GGGACAAGGGCTCCACCACCACAGACCGGAGCCACCGACAGCCAGCCAGAGGTCCCCTCCAGTCCCGCCCGGCGCCCCCC

CGAAGAGGCCCCGGCAAGGCCCCCCAGCCCGACGTCCCCCGCACGCACCGAAAGAAGGACCCGCCCGAGAACGTGCGGCA

CATCCATGCGCAAACATCATGACGCGGGTCGCAATTTTGACATTTCTGTTTCTTTTCCCAAATGTTGTTGCGTGTCAGAT

TCACTGGGGCAATCTATCCAAGATCGGGATTGTAGGAACAGGGAGTGCCAGCTACAAGGTGATGACTAGGCCAAGCCACC

AGACTCTGGTTATAAAGTTAATGCCAAATATAACGGCCATCGACAATTGTACAAAGTCAGAGATTGCAGAGTACAAGAGA

TTGCTGATCACAGTGTTAAAGCCTGTAGAGGATGCTCTGTCGGTGATAACCAAGAATGTAAGACCAATTCAAACTCTAAC

ACCTGGGCGTAGAACCCGCCGTTTTGCTGGAGCTGTTCTGGCCGGGGTAGCACTTGGAGTTGCGACAGCCGCTCAGATAA

CTGCAGGAGTCGCCCTTCATCAATCATTGATGAACTCCCAAGCAATTGAGAGTTTAAAAACCAGTCTTGAGATGTCGAAT

CAGGCAATAGAAGAAATCAGACTTGCAAATAAGGAGACCATACTGGCAGTACAGGGCGTCCAGGATTATATCAACAATGA

GCTCGTCCCTTCTGTTCATAGAATGTCATGCGAGCTGGTAGGTCACAAGCTCGGCCTCAAGCTCCTTAGGTACTACACCG

AGATCCTGTCCATATTCGGGCCCAGTCTTCGAGACCCGATATCTGCCGAAATATCAATCCAGGCACTTAGTTATGCATTA

GGCGGAGACATTAATAAAATCCTGGACAAGCTTGGGTATAGCGGTGGGGATTTCCTTGCCATCCTAGAAAGCAAGGGAAT

AAAGGCCCGGGTTACATATGTGGACACAAGAGATTACTTTATAATCCTTAGCATCGCCTACCCAACCTTATCTGAGATCA

AGGGAGTGATAGTTCACAAGATAGAAGCTATAACATACAACATTGGGGCACAGGAGTGGTATACTACTATCCCTAAATAT

GTAGCCACTCAGGGGTATCTGATATCGAACTTTGATGAGACGTCATGCGTATTCACTCCAGAGGGGACAGTTTGCAGCCA

GAATGCGTTGTACCCAATGAGCCCATTGCTTCAGGAATGTTTCAGGGGGTCAACAAAATCGTGCGCCAGAACCTTAGTTT

CAGGGACCATAAGTAATAGATTTATCCTATCAAAAGGGAACCTGATTGCAAATTGTGCGTCAGTTTTGTGCAAATGTTAC

ACAACAGAGACAGTTATCAGCCAAGATCCTGACAAACTACTAACTGTTGTAGCATCCGACAAGTGTCCTGTAGTTGAGGT

GGATGGAGTGACAATACAGGTCGGCAGTCGAGAGTATCCGGATTCTGTATACTTACACAAAATAGACTTAGGTCCAGCCA

TCTCCCTAGAAAAACTGGATGTAGGCACCAATTTAGGCAATGCAGTCACAAGACTGGAGAATGCAAAGGAGCTCCTAGAT

GCATCAGACCAAATACTGAAGACTGTTAAAGGGGTACCTTTCAGTGGGAATATGTACATAGCACTGGCAGCTTGCATAGG

AGTATCCCTGGGCCTTGTCACATTAATATGCTGCTGTAAGGGGAGGTGTAAGAACAAGGAAATCCCTATCTCCAAAATCA

ACCCAGGGCTCAAACCCGACCTGACCGGGACCTCAAAGTCGTACGTAAGATCACTGTAGTCAGAATCACCTGGATCATCT

GGCATCACACACATACATGCACGACACAGGCAGTCCGATGACGCAAGAAACCCAACCTCCGGTCACCCACCCGACCCCAC

TCCACGCTCCACCACACATTAGTCATCAAACAAAACTTAGGACGAAAGGTCAATCACCATGTCCGCACAAAGGGAGAGGA

TCAATGCCTTCTACAAAGACAATCCTCACAATAAGAACCATAGGGTGATCCTGGATAGAGAACGCTTGGTCATTGAAAGA

CCCTACATCTTGCTTGGAGTCCTGCTGGTAATGTTCCTGAGTCTAATCGGACTGCTGGCCATTGCAGGGATCAGGCTTCA

CCGGGCCACCGTTGGAACTTCAGAGATCCAGAGTCGGCTGAATACCAATATTAAGTTGACCGAATCTATTGATCACCAGA

CTAAGGATGTCTTAACTCCCCTTTTTAAAATCATTGGCGATGAAGTCGGCATCAGAATTCCACAGAAATTCAGTGATCTT

GTCAAGTTCATCTCCGATAAGATTAAATTCCTCAACCCTGATAGAGAGTATGATTTCAGAGATCTCCGGTGGTGTATGAA

TCCTCCCGAGAGAGTCAAAATTAATTTTGATCAGTTTTGTGAGTACAGGGCTGCGGTTAAGTCAATTGAACATATATTTG

AGTCACCACTCAACAAGTCAAAAAAGCTGCAATCTTTGACTCTCGGGCCCGGAACAGGCTGTCTAGGCAGGACAGTAACA

AGAGCCCATTTCTCAGAACTTACAATGACCTTAATGGACCTGGATCTAGAGATGAAGCACAACGTGTCCTCAGTGTTTAC

CGTAGTTGAAGAGGGATTATTCGGAAGAACATATACCGTCTGGAGATCCGATGCCAGGGATCCGAGCGCCGATCTAGGTA

TTGGCCATTTTTTAAGAGTCTTCGAGATTGGACTGGTAAGAGATCTTGGGCTGGGTCCCCCTGTTTTTCATATGACCAAC

TATCTCACGGTGAACATGAGTGATGACTATCGGAGATGTCTTTTAGCGGTAGGGGAGTTGAAGTTGACAGCCCTATGCAC

CTCATCTGAGACTGTGACACTGAGTGAGAGAGGAGTTCCAAGGAGGGAACCTCTTGTGGTTGTGATACTTAATCTAGCTG

GACCCACTCTAGGGGGAGAGCTATACAGTGTCTTGCCTACCTCTGATCTCATGGTGGAGAAACTCTATTTGTCTTCACAT

AGAGGGATCATCAAAGATGATGAGGCCAATTGGGTAGTGCCGTCTACCGATGTTCGTGATCTTCAGAACAAAGGTGAATG

TCTGGTGGAAGCATGCAAGACTCGACCTCCTTCATTTTGCAATGGCACAGGATCAGGCCCGTGGTCAGAAGGGAGAATCC

CTGCCTACGGGGTGATCAGGGTCAGTCTTGACTTAGCGAGTGACCCGGATGTAGTTATCACTTCAGTGTTTGGCCCACTG

ATACCTCACCTATCCGGCATGGATCTTTACAACAACCCGTTTTCAAAAGCTGTATGGTTGGCTGTACCACCTTATGAGCA

GTCATTTCTAGGAATGATAAATACAATTGGATTCCCTAACAGAGCAGAGGTTATGCCGCACATTTTGACCACAGAGATCA

GAGGCCCTCGGGGTCGTTGCCATGTTCCCATAGAATTGTCCCGCCGGGTTGATGACGATATCAAGATCGGGTCCAACATG

GTCATATTGCCGACGATGGACCTGAGGTATATTACAGCCACTTATGATGTTTCCAGGAGCGAGCATGCAATCGTGTACTA

TATCTATGACACGAGTCGCTCATCATCTTACTTCTACCCAGTTCGACTGAATTTCAAAGGCAATCCTCTCTCTTTGAGGA

TAGAGTGTTTCCCCTGGCGTCATAAGGTGTGGTGCTACCATGATTGTCTTATATACAACACCATAACAGGTGAAGAGGTC

CATACGAGAGGGCTGACCGGCATAGAGGTAACATGCAATCCAGTCTGAGCAGAGCTACGACCGTCACTCAGGCAGTCCCT

TGAGTCGCCACCGAGTCCAAGCAGCACAGCCTGGGACAATCAACGGCACAGCCCAGCCAACAATGTTATAAAAAACTTAG

GAGCCAGGGTTGTAGGGGCCATGGACTCTCTATCAGTCAATCAGGTCTTGTACCCTGAGGTTCATCTAGACAGCCCTATT

GTCACAAACAAACTAGTTGCCATCCTTGAGTACTCGGGTATCGACCATAACTATGTTCTTGAAGACCAGACTCTTGTCAA

GAATATTAGGTATAGGCTGGGGTGCGGTTTTTCAAATCAAATGATCATCAATAACAGGGGGGTGGGTGAAACAGTCAATT

CTAAACTTAAAAGTTACCCCCATAATCGTCATATCATATACCCGGATTGCAATAAGGAGTTGTTTTGTATCAAGGATAGC

TGCATATCTAAGAAGCTCTCGGAGCTATTCAAGAAGGGTAATTCCTTGTACTCTAAGATAAGTCACCAGGTACTGGATTG

TCTTAAGAGAGTCAATGGGAAATTAGGTCTGGGCACAGATCTTACCCATGGTCTGAAGGAGGGTATCCTTGACCTAGGGT

TGCACATGCATAGCTCTCAATGGTTCGAGACTTTCCTGTTCTGGTTCACTATCAAGACAGAGATGAGATCAATGATCAAA

GAACAGTCCCATATATGCCACAAGAGGAGGTATAACCCGACTTTTGTGTCGGGGGATGCATTCGAGGTGCTCGTATCGCG

AGACCTCGTTGTGATAATTGATAAGAATACCCAGTATGTCTTCTACTTAACATTTGAGCTGGTCCTTATGTACTGTGATG

TCATAGAGGGTAGACTTATGACGGAGACAGCCATGGCCATAGACCAGAGATATTCAGAACTTCTAAGCCGGGTCAGATAC

TTGTGGGATCTTATTGATGGGTTCTTCCCAACACTAGGCAACACCACATACCAAATTGTTGCTCTGCTTGAACCATTATC

GTTGGCTTATCTTCAACTTCAGGATGTCACTCTGGAGTTAAGAGGTGCTTTTTTAGACCATTGCTTCAAAGAACTCTATG

AGATACTGGAACATTGCGGCATTGACACAGCAGGCACTTACAATTCCATCACTGAAGGATTGGATTACGTATTCATCACC

CACGATATACATCTAACTGGGGAGATTTTTTCATTTTTCCGGAGTTTCGGACACCCCCGTCTAGAAGCAGTCACCGCTGC

GGAAAATGTCAGAAAACATATGAACCAACCGAAGGTAATCAGTTATGAGACTATGATGAAGGGCCATGCGGTATTTTGCG

GGATAATCATAAATGGTTTCCGGGACCGACATGGCGGCAGCTGGCCCCCTGTTGCACTGCCAGAACATGCCTCTGCTGCG

ATCCGGAATGCGCAGGCATCAGGTGAGGGACTGACCCATGACCTGTGTATAGACAACTGGAAATCCTTTGTTGGATTCAA

ATTTGGCTGCTTTATGCCACTCAGCCTAGATAGTGATTTGACCATGTATCTCAAAGACAAGGCATTGGCTGCACTGAAGA

ATGAGTGGGATTCAGTTTACCCGAAAGAATACCTCCGTTATAACCCACCTAGAGGGACAGAGTCAAGGCGATTGGTAGAG

GTATTCCTGAATGACTCCAGCTTTGATCCTTATAACATGATAATGTACGTGGTGAATGGCTCCTACCTTAAAGACCCTGA

GTTTAACCTCTCATACAGCCTAAAAGAGAAGGAGATAAAGGAGACAGGGCGGTTGTTTGCCAAAATGACCTATAAGATGC

GGGCCTGTCAGGTAATCGCTGAAAATCTGATATCGAATGGTGTTGGGAAGTATTTCCGAGACAATGGGATGGCAAAGGAC

GAGCATGACCTAACAAAAGCCCTTCACACCCTGGCAGTCTCAGGTGTTCCCAAAAACAACAAAGATAACCACCGAGGTGG

GCCTCCCAGAAGGACAACAAACCGAGGGGTGAGATCAAGCCAAGGCACCAAAACACAAGATAGAGACAAGGTTCAAGGGG

GACCTATGTACAACTATTTGCGATGCCAACCGATCAGCCCTGATCAGGGTGAGTCATACGAGACTGTTAGTGCATTCATC

ACCGCTGACCTTAAGAAGTATTGCCTGAATTGGAGATACGAGACAATCAGCATATTTGCACAGAGGCTGAATGAAATATA

TGGACTACCATCCTTCTTTCAATGGTTACACAGGATATTGGAAAAATCCGTACTCTACGTCAGTGACCCACATTGTCCTC

CCGATCTAGATAATCATATCCCTCTGGACAGTGTCCCCAATGCCCAAATATTCATCAAGTACCCAATGGGCGGAATAGAA

GGTTATTGCCAAAAACTATGGACAATCAGCACTATACCATACTTGTATCTGGCAGCCTATGAGAGCGGAGTAAGAATCGC

CTCACTGGTGCAGGGTGACAATCAGACAATCGCAGTGACAAAAAGAGTTCCAAGTTCTTGGCCTTATTCACTAAAAAAGA

GGGAGGCCTCCAAGGCCGCTCAAAATTACTTCGTAGTCTTAAGGCAAAGATTGCACGATGTAGGTCATCACTTAAAGGCT

AATGAGACCATAGTATCTTCTCACTTTTTTGTATATTCCAAAGGGATTTATTATGACGGCCTGCTAGTCTCACAATCACT

AAAGAGCATCGCCAGATGTGTCTTCTGGTCCGAGACTATAGTGGATGAAACCAGAGCGGCGTGCAGTAATATTGCAACAA

CTGTCGCTAAGAGTATAGAGAGGGGTTATGATAGGTACCTTGCATATTCTTTGAATATCCTCAAGATTTTCCAACAGATC

CTTATATCCCTTAACTTCACTATTAACACAACAATGACTCAGGATGTCGTGGCACCGATCATCGAGAACGGTGATTTGCT

GATAAGGATGGCACTCTTGCCAGCACCCATCGGGGGTCTCAATTATCTTAACATGAGCAGGTTATTTGTGAGAAATATCG

GTGACCCGGTCACCTCCTCTATAGCCGACCTGAAGAGGATGATAGATGCCGGGCTAATGCCAGAAGAAACATTGCATCAA

GTGATGACCCAGACCCCGGGAGAATCATCTTACCTAGATTGGGCAAGTGACCCTTATTCTGCCAACCTACCCTGCGTACA

GAGTATCACTCGCCTTCTCAAAAACATCACTGCACGATATATTTTAATCAGCAGCCCAAACCCGATGCTGAAAGGGTTAT

TTCATGAGGGGAGTAGAGATGAAGACGAGGAGCTTGCAAGTTTCCTAATGGATCGGCATATAATTGTTCCAAGAGCTGCA

CATGAAATCTTAGACCATAGTATAACCGGAGCAAGAGAAGCTATAGCCGGGATGTTGGACACCACCAAGGGTCTGATTAG

AACAAGTATGAAACGGGGCGGCCTCACCCCTCGAGTCTTAGCCCGCCTTTCCAATTATGATTATGAACAATTCCGATCCG

GGATAACACTATTAACAAAGAAAGGGCAGTGTTATCTCATTGACAAGGACTCGTGCTCGGTGCAGCTCGCTATAGCTCTG

AGGGGCCACATGTGGGCTAAGTTGGCTCGCGGGAGACCAATCTATGGGTTGGAGGTGCCTGATGTATTAGAATCGATGAA

CGGCTACCTTATTAAACGCCACGAGTCGTGTGCTATCTGTGAAACGGGCTCAAGTCACTACGGATGGTTTTTCGTCCCTG

CAGGGTGCCAGCTTGACGATGTATCTAGAGAGACTTCGGCTCTTCGTGTACCTTATGTCGGATCAACAACTGAGGAAAGG

ACAGACATGAAACTTGCATTTGTTAGGTCTCCAAGTAGATCCCTCAAATCAGCAGTTAGAATTGCAACAGTTTACTCATG

GGCTTACGGGGATGATGAGAAATCATGGAGTGAAGCCTGGATGCTAGCTAGGCAGAGAGCCAATATCACCTTAGATGAAT

TGAGAATGATCACTCCAGTCTCCACATCCACCAACTTAGCCCACCGGTTGAGGGATCGGAGTACCCAGGTGAAGTACTCG

GGGACATCCCTTGTGAGGGTTGCAAGATACACAACCATCTCCAATGATAACTTGTCGTTTGTGATATCTGAGAAAAAAGT

GGATACCAACTTCATTTATCAGCAAGGGATGTTGCTTGGTCTTGGGATCCTTGAGAACCTCTTTAGGCTAGAGGCCACCA

CCGGAGTATCCAACACAGTGCTACACCTGCACGTGGAAACAGAATGTTGTGTTGTACCTATGGTGGATCACCCAAGGATA

CCGAGTCTCCGTAATATAAAAGTTACGGATGAGCTATGCACAAACCCTTTGATCTACGACAGGTCCCCCATCATAGAGCA

CGATGCAACCCGGCTATATTCACAGAGCCACAGGAGACATTTGGTGGAGTTTGTTACCTGGTCAACAAGCCAGCTTTACC

ATATACTGGCTAAGTCTACAGCAATGTCCATGATTGAATTGATTACGAGATTCGAGAAAGATCACATGAATGAAATAGCC

GCCCTGATTGGCGATGACGACATCAACAGCTTCATCACAGAATTTTTGCTAGTGGAGCCTAGATTATTTATAGTTTACCT

TGGTCAGTGTGCTGCCATCAATTGGGCCTTTGATATACATTATCATCGGCCCTCGGGCAAGTACCAGATGGGGGAACTTC

TCTACTCTCTGCTCTCTCGGATGAGCAAAGGAGTATATAAGATCTTTACTAATGCTTTGAGCCACCCCAAAGTTTACAAG

AAATTTTGGCGAAGTGGCATCATTGAGCCGATTCATGGACCATCCTTGGATACACAGAATTTACATGTCACTGTCTGTGA

CATGATATATGGGTCCTATGTCACCTATTTGGATCTTTTGCTGAATGATGAGCTAGATGATTACCCGTATTTGCTCTGCG

AGAGTGATGAGGACGTGGTCACAGACAGGTTCGACAACATTCAAGCCAAACATCTCTGTGTATTGGCCGATGTATATTGC

AGCTCCAAGAGGTGTCCTTCAATAATCGGGATGTCTCCTATAGAAAAATGTACCATTCTCACACATTACATCAAGGGAGA

ATCAGTACAATCCCCGTCTGGGACCTCATGGAACACTGATCCCCTTGTAGTAGATCATTACTCATGCTCTCTGACCTACC

TTCGCCGCGGTTCCATCAAACAAATCAGGTTGAGGGTGGATCCTGGGTTTGTATTCGAGGCGTTGACAGACGTCGACTTT

AAACAACCTCGCAAAGCTAAGTTAGATATATCGGTCGTTGGGTTGACTGATTTCTCTCCCCCTTACGACAACGTCGGTGA

TTTTTTAGGAACTATCAACACATTGAGGCACGATCTGCCTGTCACCGGAACCGGGGTCTCGAACTATGAAGTCCACGCTT

ATCGTAGAATTGGCCTGAATTCATCAGCATGTTATAAAGCCGTAGAGATCTCCACGTTAATCAAGCCATCCTTAGAAGTC

GGAGAGCATGGATTGTTCTTAGGAGAAGGTTCCGGTTCAATGCTGGCTGCGTACAAAGAAGTTCTTAAATTAGCAAATTG

TTACTACAACAGCGGAGTAACAGCGGAGGGCAGAGCCGGACAGAGGGAAATATCTCCCTATCCTTCAGAGATGAGCCTAG

TAGAGAATCAAATGGGGATAGAGCGGAGTGTTAAAGTGCTGTTCAACGGGAAACCTGAAGTAACTTGGGTGGGGACCACA

GATTGCTACAAGTACATAATCAGTAACATCCAAACCTCTAGTCTGGGTTTCATACATTCAGATATTGAGACGCTCCCAAC

CAAAGATGCTGTTGAGAAATTGGAGGAATTTGCCTCTATCCTATCCTTATCCTTGATTTTGGGGAAAATCGGCTCTATTA

CAGTTGTCAAAATTATGCCCATTAGCGGGGATTTTACCCAAGGCTTCATAGCCTATGCCATCCAATATTTTAGGGAGAGC

CTGCTTGCCTATCCGAGATATAGTAACTTCATCTCGACTGAGTGTTACCTTATTATGATAGGATTAAAGGCCAATCGATT

AATAAACCCAGAAGCCATTAAGCAAAGCATAATCAGAGCGGGGGTCAGGACTGCACCAGGACTTGTGAGCCATGTATTAT

CAGAGAAACAAAAAGGTTGTATTCAATCTTTCTTGGGTGATCCTTATATCCAAGGAGACTTCAATAAGCACCTTAAATCT

CTAACCTCTATTGAGAAAATCCTGGTAAATTGTGGTCTCTCGATCAATGGCATAAAAATCTGTAGAGATCTAATCCACCA

TGATATCGCCTCCGGTCCAGACGGCCTTATGAGCTCTACAATTATTTTATATAGGGAATTGGCTCATTTCAAAGATAATA

TAAGGAGTCAGCACGGAATGTTCCACCCCTACCCGGTACTGGCCAACAGCAGGCAACGTGAATTAATCCTTCGAATAGCC

AAGAAATTCTGGGGGTATGTCTTGCTATATTCTGATGACCCGGCACTAATCAGACAAACAATCAAGAACTTGAAACGGAA

TCATCTAACCTTTGACTTACATAGTAACCCATTTATTAAAGGGCTATCCAAGGCTGAGAAACTGCTGGTGCGGACGAGTT

CACTAAGAAGAGAATGGTTGTTCACTCTCGAAACGAAAGAAGTAAAAGAGTGGTTCAAATTGGTAGGTTACAGTGCACTC

ATCAGAGGCTAGTCAGCTATGCATCTGCCCCCTTCTCCTCCGCCATGAGACCCCACTGGCGATCCAGAAGATTAAAGAAA

ACTACATATTGGATAAGTATCTATTCCCAGCTTTGTCTGGT

>Saiga3/saiga/Mongolia/2017-01

ACCAAACAAAGTTGGGTAAGGATAGATCTTATAATAACTATGGACTGGCAAACTTAGGAGTAAAGATCCTACTGTCGGGG

GGAGGAGGAGGAGCAAGATCCTTGATCATGGCGACTCTCCTTAAAAGCTTAGCATTGTTCAAAAGGAACAAGGACAAAGC

GCCGACAGCATCAGGTTCAGGAGGGGCCATCCGGGGGATTAAGAATGTTATCATAGTCCCGATTCCCGGAGACTCGTCCA

TCACTACCCGTTCAAGACTGCTCGACAGGCTTGTCAGATTGGCCGGAGATCCTGACATCAACGGGTCAAAGCTGACCGGG

GTGATGATCAGCATGTTATCCTTGTTCGTAGAGTCACCCGGGCAATTGATACAGCGAATCACAGATGATCCGGATGTTAG

TATCCGCCTTGTTGAGGTAGTTCAAAGTACTAGGTCCCAGTCCGGGTTGACCTTTGCATCACGTGGTGCTGATTTAGACA

ACGAGGCAGACATGTATTTTTCAACTGAGGGGCCCTCGAGTGGAGGTAAGAAAAGGATCAACTGGTTTGAGAACAGAGAA

ATAATAGACATAGAGGTGCAGGATCCAGAAGAGTTCAATATGTTGTTAGCCTCCATACTAGCACAAGTCTGGATCCTTCT

GGCCAAGGCTGTTACGGCACCAGATACGGCAGCTGACTCAGAACTGAGAAGGTGGGTTAAATACACACAACAAAGGAGAG

TGATTGGGGAATTTCGCCTTGACAAGGGGTGGCTGGACGCAGTCCGCAACAGAATTGCAGAAGATCTATCACTCCGGCGG

TTCATGGTATCTCTCATACTTGACATCAAGAGGACCCCAGGCAACAAGCCAAGGATTGCAGAAATGATCTGCGACATTGA

CAACTATATTGTCGAAGCAGGACTCGCCAGTTTCATCCTTACTATCAAATTTGGTATCGAAACCATGTATCCTGCATTAG

GGCTCCACGAGTTTGCCGGAGAATTGTCCACTATTGAGTCCTTGATGAACTTGTATCAACAGCTAGGAGAGGTTGCACCC

TACATGGTAATTCTAGAGAATTCAATTCAGAACAAGTTTAGTGCAGGAGCTTATCCCCTCCTCTGGAGCTATGCGATGGG

TGTCGGAGTCGAGCTGGAGAACTCAATGGGGGGCTTGAACTTTGGCAGGTCATATTTTGACCCTGCCTATTTTCGTCTCG

GACAGGAGATGGTCAGAAGATCTGCAGGAAAGGTCAGCTCTGTGATTGCGGCTGAGCTCGGCATCACAGCAGAGGAAGCT

AAACTAGTCTCGGAAATCGCCTCGCAGGCCGGGGACGAAAGAACCGCCAGAGGGACTGGGCCTCGACAGGCGCAGGTTTC

CTTCCTTCAGCACCAAACAGGAGGGGGAGAGTCGTCCGCACCAGCGACCAGAGAAGGGGTCAAAGCTGCGATCCCAAACG

GAGCTGAAGAAAGGGACAGAAAACAAACACGCCCAGGAAGGCCCAGAGGAGAGACCTCCGGCCAACTGCTCCTGGACATC

ATGCCAGAGGATGAGATCTTGCGAGAGTCTGGTCAAAACCCTCGTGAGGCTCAAAGATCGGCCGAGGCACTCTTCAGGCT

GCAGGCCATGGCCAAGATTCTGGAGGACCAGGAGGAGGGAGAAGACAACAGTCAGGTCTACAACGACAAGGATCTCCTCG

GCTAAGCAGACGCACCCTCTGTCGAAATCAGTGACAAGACATCTCCTACCAGTATTATAAAAAACTTAGGACCCAGGTCC

AAGCAAGCACACATCGACACTCCAACCAGTCGAGCGGAGACCACCGATGGCAGAAGAACAAGCATACCATGTNAACAAGG

GGCTGGAATGTATCAAGTCTCTCAAAGCCTCACCCCCGGATCTATCCACCATCAGAGATACCATCGAGAGCTGGAGAGAG

GGGCTTAGCCCCTCGGGCCGTGCAACACCGAACCCTGATACGTCCGAGGGAGATCATCAGAATATCAACCAATCATGCTC

ACCAGCAATCGGACCAAACAAAGTCTACTTGTCTCCTGAAGATAATCTCGGATTTAGAGAGATCACTGGCAACGACTGTG

AGGCTGGGCTCGGAGGGGTCCAGAGAGAAGGATCCAACTCTCAAGTACAGCGTTACCATGTTTATAGTCACGGGGGTGAA

GAGATTGAAGGACTCGAGGATGCTGACTCTCTCGTGGTTCAAGCAGATCCTCCAGCTGCTAACGTCTTCAATGGAGGAGA

GGATGGATCTGACGACAGCGATGTGGACTCTGGCCCAGATGATCCCGGCAGAGATACTCTATATGACCGGGGATCTGTTG

CCGGCAATGGTATCGCTAGGTCCACAGATGTCGAAAAACTAGAAGGTGCTGATATTCAAGAAGTTCTTAACTCCCAGAAA

GGCAAAGGAGGAAGATTCCAGGGCGGGAAAACCTTGCGAGTCCCGGAAATACCCGATGTCAAGCACTCCAGACCATCAGC

CCAATCAATTAAAAAGGGCACAGACGGGAACTCAGTCTCATCTGGAACGGTGATAGAGTGTTTATCGATAAGTGGTGCAA

CCCAAACTGTGCCAGAGTCAAGATGGGAGTCATCAGAGCAAAATGCGTCTGTGGGGAGTGTCCTCAAGTCTGCGAGGAGT

GCAAAGACGATCCAGGGGCCGACACAAGAATCTGGTACCATAGCATCACTGACTCAGCCTAAAGAGAATGACTCCGAGTA

TGAGTATGAGGATGACCTATTTACAGAGATTCAGGACATCCGTGCAAGCATTGCCAAGATCCATGATGACAATAAAACTA

TCCTCTCAAAGCTTGATTCTATACTGTTATTGAAAGGGGAAGTCGACACTATCAAGAAACAAATCAGCAAGCAGAATATA

AGTATATCCACCATTGAGGGCCATCTCTCCAGTATAATGATAGCCATCCCTGGCTTTGGGAAGGACATCAAGGACCCAAC

ATCCGAAGTCGAGTTGAACCCAGATCTAAGACCTATAATAAGCCGTGACTCTGGCAGAGCTCTCGCGGAGGTCCTCAAGA

AACCCGCTGTGGATAGGTCTCAGAAAATTGGAACCAAAGCCAACTCCAGCTCAAAGGGTCAGCTTCTTAAGGATCTCCAG

CTAAAACCTGTCGACAAGCAGGCAAGCTCTGCAATCGGGTTTGTCCCGTCCGACCATGAATCATCCAGAAATGTCATCCG

CTCCATAATCAAGTCGAGCAAGCTAAATATTGATCACAAGGACTATCTTCTAGATTTACTAAATGATGTGAAAGGCTCCA

AGGATCTTAAGGAATTTCACAAGATGCTAACAGCAATTCTCGCCAAGCACCTGTAACACATCCCCCAGTCATCATCTCAT

ACTCGACTAAAAACATCCTTTCAATCAGGCTATTACAAAAAACTTAGGAGCAAGGGCAACTGAGCTTCGCAGACAGGATG

ACCGAGATCTACGACTTCGATAAATCAGCATGGGATGTCAAAGGGTCAATTGCCCGCATAGAACCCACCACCTACCACGA

CGGCCGACTGATACCCCAGGTGAGGGTCATCGATCCTGGTCTGGGAGACAGAAAAGATGAGTGCTTTATGTACCTGTTTC

TCCTAGGAGTGATTGAGGATAACGACCCCCTGTCTCCCCCCGTCGGAAGAACCTTTGGCTCTCTACCTCTAGGGGTTGGT

AGGTCAACTGCCAAGCCAGAAGAACTACTAAGGGAGGCCACAGAATTAGATATAGTGGTGAGGCGCACTGCAGGAGTAAA

TGAGAAACTGGTATTTTACAACAACACTCCGCTGTCCTTGTTAACGCCATGGAAGAAAGTTCTGACAACCGGAAGTGTGT

TTAGCGCTAACCAGGTTTGCAATGCAGTCAACCTAGTCCCGCTTGATACTCCCCAGAGATTCAGGGTTGTGTACATGAGC

ATAACTAGGTTATCAGACAATGGTTACTATAGTGTGCCCAGAAGAATGCTGGAGTTCCGCTCGGCCAATGCAGTAGCCTT

CAACATCTTGGTTACACTAAGAATTGAAAATGGCACAAACCCTAGAAGATACATAGTCGGCTCATGGGAGAATTCAGAGG

TCACATTTATGGTACACGTGGGCAACTTTAGGAGAAAGAAGAACGAAGTATACTCTGCTGATTACTGCAAAATGAAGATT

GAAAAGATGGGTTTAGTTTTTGCCCTGGGTGGAATAGGCGGAACAAGTCTCCATATTAGAAGCACCGGGAAAATGAGCAA

AACCCTCCATGCACAGCTGGGGTTCAAGAAAATCTTATGTTACCCCCTAATGGATGTCAATGAGGATCTTAACCGATATC

TCTGGCGGGCAGAGTGCCGAATAGTCAAAATCCAAGCTGTCTTACAGCCATCAGTACCCCAAGAGTTCCGTGTCTACGAT

GATGTCATCATCAACGATGATCAAGGCTTGTTCAAGATCTTGTAGTTCATTTGCAAAGGAACGAACCCACCCCCAGGAAG

GGGCCCCGCCCTCCCCCG-CCAAAAAACACACTCCAAGGCTCCCCGGCACAGCGCCAACACAGCTACTCCCGCGATGGAA

CGAGGTCGGCC-CCCGAACAAATCCGGTCCAAAGAGGGGGGCCCCCCCCCTGACCAACCGGGCGCCGACCGAGCAGGCCG

GCGAGAAGCCCCCAAGAGGCTCCGACGAAGACCGGGCACCCCATCCCCCC-AAAAACGTCCCCC-AACACGCCCGAGGCT

AGCCACCGGACACCCCCAGACCCGCCACCGGCGAGACAGAACCCCAAGGGGGGACCCTCCAAACCAAAGGAACCACCGGG

GTAGGCTCCGCCGAACCAGCACAGACCCGACCCAAGCAAGGCGGACCCCCAACACGAAACTGCCCAGCACACCCTACTCA

TCATCAAACAAAACTTAGGGGCCAAGTCCACACGAGAGGCCCCAGTCCGTCAGGCGGAGGGACGCAGAAAGGAAGGAGAC

ACGACCGTCCCAGAAGGAGGACACCCAACCACCGAAACGCCCGCCCCCAGGGAAGGAG--CCA-CCCCCACGCAGCGAAG

GAGCGGGCGGGAGACCCCGGGGAGGGCCCGATCGGGACCCGGGGGCCGGCGGCGCCGGCGGTGAGCAGGCCGGGCGCACA

CCCGACCCCACCGGAGGATCCGAGGAGGGCCCGACCCCGCGCGGACCAAACGGCAGGGAGCCCAACCCCACCAAAAAGAG

AGGCGAGGCCCCCCCTCCCTCCCCCCCCGCACACAACCCGCAGACAGACCGAGGGAGGGCCATCCCCCAGCGAGACAGGC

GGCTCCCTCGCAGCCGAGCCGCAGCTCCCCAGAATAGGACACCCCCGCACACTCCGAACACCCCCCCGCCCCCCGCCACC

GGGACAAGGGCTCCACCACCACAGACCGGAGCCACCGACAGCCAGCCAGAGGTCCCCTCCAGTCCCGCCCGGCGCCCCCC

CGAAGAGGCCCCGGCAAGGCCCCCCAGCCCGACGTCCCCCGCACGCACCGAAAGAAGGACCCGCCCGAGAACGTGCGGCA

CATCCATGCGCAAACATCATGACGCGGGTCGCAATTTTGACATTTCTGTTTCTTTTCCCAAATGTTGTTGCGTGTCAGAT

TCACTGGGGCAATCTATCCAAGATCGGGATTGTAGGAACAGGGAGTGCCAGCTACAAGGTGATGACTAGGCCAAGCCACC

AGACTCTGGTTATAAAGTTAATGCCAAATATAACGGCCATCGACAATTGTACAAAGTCAGAGATTGCAGAGTACAAGAGA

TTGCTGATCACAGTGTTAAAGCCTGTAGAGGATGCTCTGTCGGTGATAACCAAGAATGTAAGACCAATTCAAACTCTAAC

ACCTGGGCGTAGAACCCGCCGTTTTGCTGGAGCTGTTCTGGCCGGGGTAGCACTTGGAGTTGCGACAGCCGCTCAGATAA

CTGCAGGAGTCGCCCTTCATCAATCATTGATGAACTCCCAAGCAATTGAGAGTTTAAAAACCAGTCTTGAGATGTCGAAT

CAGGCAATAGAAGAAATCAGACTTGCAAATAAGGAGACCATACTGGCAGTACAGGGCGTCCAGGATTATATCAACAATGA

GCTCGTCCCTTCTGTTCATAGAATGTCATGCGAGCTGGTAGGTCACAAGCTCGGCCTCAAGCTCCTTAGGTACTACACCG

AGATCCTGTCCATATTCGGGCCCAGTCTTCGAGACCCGATATCTGCCGAAATATCAATCCAGGCACTTAGTTATGCATTA

GGCGGAGACATTAATAAAATCCTGGACAAGCTTGGGTATAGCGGTGGGGATTTCCTTGCCATCCTAGAAAGCAAGGGAAT

AAAGGCCCGGGTTACATATGTGGACACAAGAGATTACTTTATAATCCTTAGCATCGCCTACCCAACCTTATCTGAGATCA

AGGGAGTGATAGTTCACAAGATAGAAGCTATAACATACAACATTGGGGCACAGGAGTGGTATACTACTATCCCTAAATAT

GTAGCCACTCAGGGGTATCTGATATCGAACTTTGATGAGACGTCATGCGTATTCACTCCAGAGGGGACAGTTTGCAGCCA

GAATGCGTTGTACCCAATGAGCCCATTGCTTCAGGAATGTTTCAGGGGGTCAACAAAATCGTGCGCCAGAACCTTAGTTT

CAGGGACCATAAGTAATAGATTTATCCTATCAAAAGGGAACCTGATTGCAAATTGTGCGTCAGTTTTGTGCAAATGTTAC

ACAACAGAGACAGTTATCAGCCAAGATCCTGACAAACTACTAACTGTTGTAGCATCCGACAAGTGTCCTGTAGTTGAGGT

GGATGGAGTGACAATACAGGTCGGCAGTCGAGAGTATCCGGATTCTGTATACTTACACAAAATAGACTTAGGTCCAGCCA

TCTCCCTAGAAAAACTGGATGTAGGCACCAATTTAGGCAATGCAGTCACAAGACTGGAGAATGCAAAGGAGCTCCTAGAT

GCATCAGACCAAATACTGAAGACTGTTAAAGGGGTACCTTTCAGTGGGAATATGTACATAGCACTGGCAGCTTGCATAGG

AGTATCCCTGGGCCTTGTCACATTAATATGCTGCTGTAAGGGGAGGTGTAAGAACAAGGAAATCCCTATCTCCAAAATCA

ACCCAGGGCTCAAACCCGACCTGACCGGGACCTCAAAGTCGTACGTAAGATCACTGTAGTCAGAATCACCTGGATCATCT

GGCATCACACACATACATGCACGACACAGGCAGTCCGATGACGCAAGAAACCCAACCTCCGGTCACCCACCCGACCCCAC

TCCACGCTCCACCACACATTAGTCATCAAACAAAACTTAGGACGAAAGGTCAATCACCATGTCCGCACAAAGGGAGAGGA

TCAATGCCTTCTACAAAGACAATCCTCACAATAAGAACCATAGGGTGATCCTGGATAGAGAACGCTTGGTCATTGAAAGA

CCCTACATCTTGCTTGGAGTCCTGCTGGTAATGTTCCTGAGTCTAATCGGACTGCTGGCCATTGCAGGGATCAGGCTTCA

CCGGGCCACCGTTGGAACTTCAGAGATCCAGAGTCGGCTGAATACCAATATTAAGTTGACCGAATCTATTGATCACCAGA

CTAAGGATGTCTTAACTCCCCTTTTTAAAATCATTGGCGATGAAGTCGGCATCAGAATTCCACAGAAATTCAGTGATCTT

GTCAAGTTCATCTCCGATAAGATTAAATTCCTCAACCCTGATAGAGAGTATGATTTCAGAGATCTCCGGTGGTGTATGAA

TCCTCCCGAGAGAGTCAAAATTAATTTTGATCAGTTTTGTGAGTACAGGGCTGCGGTTAAGTCAATTGAACATATATTTG

AGTCACCACTCAACAAGTCAAAAAAGCTGCAATCTTTGACTCTCGGGCCCGGAACAGGCTGTCTAGGCAGGACAGTAACA

AGAGCCCATTTCTCAGAACTTACAATGACCTTAATGGACCTGGATCTAGAGATGAAGCACAACGTGTCCTCAGTGTTTAC

CGTAGTTGAAGAGGGATTATTCGGAAGAACATATACCGTCTGGAGATCCGATGCCAGGGATCCGAGCGCCGATCTAGGTA

TTGGCCATTTTTTAAGAGTCTTCGAGATTGGACTGGTAAGAGATCTTGGGCTGGGTCCCCCTGTTTTTCATATGACCAAC

TATCTCACGGTGAACATGAGTGATGACTATCGGAGATGTCTTTTAGCGGTAGGGGAGTTGAAGTTGACAGCCCTATGCAC

CTCATCTGAGACTGTGACACTGAGTGAGAGAGGAGTTCCAAGGAGGGAACCTCTTGTGGTTGTGATACTTAATCTAGCTG

GACCCACTCTAGGGGGAGAGCTATACAGTGTCTTGCCTACCTCTGATCTCATGGTGGAGAAACTCTATTTGTCTTCACAT

AGAGGGATCATCAAAGATGATGAGGCCAATTGGGTAGTGCCGTCTACCGATGTTCGTGATCTTCAGAACAAAGGTGAATG

TCTGGTGGAAGCATGCAAGACTCGACCTCCTTCATTTTGCAATGGCACAGGATCAGGCCCGTGGTCAGAAGGGAGAATCC

CTGCCTACGGGGTGATCAGGGTCAGTCTTGACTTAGCGAGTGACCCGGATGTAGTTATCACTTCAGTGTTTGGCCCACTG

ATACCTCACCTATCCGGCATGGATCTTTACAACAACCCGTTTTCAAAAGCTGTATGGTTGGCTGTACCACCTTATGAGCA

GTCATTTCTAGGAATGATAAATACAATTGGATTCCCTAACAGAGCAGAGGTTATGCCGCACATTTTGACCACAGAGATCA

GAGGCCCTCGGGGCCGTTGCCATGTTCCCATAGAATTGTCCCGCCGGGTTGATGACGATATCAAGATCGGGTCCAACATG

GTCATATTGCCGACGATGGACCTGAGGTATATTACAGCCACTTATGATGTTTCCAGGAGCGAGCATGCAATCGTGTACTA

TATCTATGACACGAGTCGCTCATCATCTTACTTCTACCCAGTTCGACTGAATTTCAAAGGCAATCCTCTCTCTTTGAGGA

TAGAGTGTTTCCCCTGGCGTCATAAGGTGTGGTGCTACCATGATTGTCTTATATACAACACCATAACAGGTGAAGAGGTC

CATACGAGAGGGCTGACCGGCATAGAGGTAACATGCAATCCAGTCTGAGCAGAGCTACGACCGTCACTCAGGCAGTCCCT

TGAGTCGCCACCGAGTCCAAGCAGCACAGCCTGGGACAATCAACGGCACAGCCCAGCCAACAATGTTATAAAAAACTTAG

GAGCCAGGGTTGTAGGGGCCATGGACTCTCTATCAGTCAATCAGGTCTTGTACCCTGAGGTTCATCTAGACAGCCCTATT

GTCACAAACAAACTAGTTGCCATCCTTGAGTACTCGGGTATCGACCATAACTATGTTCTTGAAGACCAGACTCTTGTCAA

GAATATTAGGTATAGGCTGGGGTGCGGTTTTTCAAATCAAATGATCATCAATAACAGGGGGGTGGGTGAAACAGTCAATT

CTAAACTTAAAAGTTACCCCCATAATCGTCATATCATATACCCGGATTGCAATAAGGAGTTGTTTTGTATCAAGGATAGC

TGCATATCTAAGAAGCTCTCGGAGCTATTCAAGAAGGGTAATTCCTTGTACTCTAAGATAAGTCACCAGGTACTGGATTG

TCTTAAGAGAGTCAATGGGAAATTAGGTCTGGGCACAGATCTTACCCATGGTCTGAAGGAGGGTATCCTTGACCTAGGGT

TGCACATGCATAGCTCTCAATGGTTCGAGACTTTCCTGTTCTGGTTCACTATCAAGACAGAGATGAGATCAATGATCAAA

GAACAGTCCCATATATGCCACAAGAGGAGGTATAACCCGACTTTTGTGTCGGGGGATGCATTCGAGGTGCTCGTATCGCG

AGACCTCGTTGTGATAATTGATAAGAATACCCAGTATGTCTTCTACTTAACATTTGAGCTGGTCCTTATGTACTGTGATG

TCATAGAGGGTAGACTTATGACGGAGACAGCCATGGCCATAGACCAGAGATATTCAGAACTTCTAAGCCGGGTCAGATAC

TTGTGGGATCTTATTGATGGGTTCTTCCCAACACTAGGCAACACCACATACCAAATTGTTGCTCTGCTTGAACCATTATC

GTTGGCTTATCTTCAACTTCAGGATGTCACTCTGGAGTTAAGAGGTGCTTTTTTAGACCATTGCTTCAAAGAACTCTATG

AGATACTGGAACATTGCGGCATTGACACAGCAGGCACTTACAATTCCATCACTGAAGGATTGGATTACGTATTCATCACC

CACGATATACATCTAACTGGGGAGATTTTTTCATTTTTCCGGAGTTTCGGACACCCCCGTCTAGAAGCAGTCACCGCTGC

GGAAAATGTCAGAAAACATATGAACCAACCGAAGGTAATCAGTTATGAGACTATGATGAAGGGCCATGCGGTATTTTGCG

GGATAATCATAAATGGTTTCCGGGACCGACATGGCGGCAGCTGGCCCCCTGTTGCACTGCCAGAACATGCCTCTGCTGCG

ATCCGGAATGCGCAGGCATCAGGTGAGGGACTGACCCATGACCTGTGTATAGACAACTGGAAATCCTTTGTTGGATTCAA

ATTTGGCTGCTTTATGCCACTCAGCCTAGATAGTGATTTGACCATGTATCTCAAAGACAAGGCATTGGCTGCACTGAAGA

ATGAGTGGGATTCAGTTTACCCGAAAGAATACCTCCGTTATAACCCACCTAGAGGGACAGAGTCAAGGCGATTGGTAGAG

GTATTCCTGAATGACTCCAGCTTTGATCCTTATAACATGATAATGTACGTGGTGAATGGCTCCTACCTTAAAGACCCTGA

GTTTAACCTCTCATACAGCCTAAAAGAGAAGGAGATAAAGGAGACAGGGCGGTTGTTTGCCAAAATGACCTATAAGATGC

GGGCCTGTCAGGTAATCGCTGAAAATCTGATATCGAATGGTGTTGGGAAGTATTTCCGAGACAATGGGATGGCAAAGGAC

GAGCATGACCTAACAAAAGCCCTTCACACCCTGGCAGTCTCAGGTGTTCCCAAAAACAACAAAGATAACCACCGAGGTGG

GCCTCCCAGAAGGACAACAAACCGAGGGGTGAGATCAAGCCAAGGCACCAAAACACAAGATAGAGACAAGGTTCAAGGGG

GACCTATGTACAACTATTTGCGATGCCAACCGATCAGCCCTGATCAGGGTGAGTCATACGAGACTGTTAGTGCATTCATC

ACCGCTGACCTTAAGAAGTATTGCCTGAATTGGAGATACGAGACAATCAGCATATTTGCACAGAGGCTGAATGAAATATA

TGGACTACCATCCTTCTTTCAATGGTTACACAGGATATTGGAAAAATCCGTACTCTACGTCAGTGACCCACATTGTCCTC

CCGATCTAGATAATCATATCCCTCTGGACAGTGTCCCCAATGCCCAAATATTCATCAAGTACCCAATGGGCGGAATAGAA

GGTTATTGCCAAAAACTATGGACAATCAGCACTATACCATACTTGTATCTGGCAGCCTATGAGAGCGGAGTAAGAATCGC

CTCACTGGTGCAGGGTGACAATCAGACAATCGCAGTGACAAAAAGAGTTCCAAGTTCTTGGCCTTATTCACTAAAAAAGA

GGGAGGCCTCCAAGGCCGCTCAAAATTACTTCGTAGTCTTAAGGCAAAGATTGCACGATGTAGGTCATCACTTAAAGGCT

AATGAGACCATAGTATCTTCTCACTTTTTTGTATATTCCAAAGGGATTTATTATGACGGCCTGCTAGTCTCACAATCACT

AAAGAGCATCGCCAGATGTGTCTTCTGGTCCGAGACTATAGTGGATGAAACCAGAGCGGCGTGCAGTAATATTGCAACAA

CTGTCGCTAAGAGTATAGAGAGGGGTTATGATAGGTACCTTGCATATTCTTTGAATATCCTCAAGATTTTCCAACAGATC

CTTATATCCCTTAACTTCACTATTAACACAACAATGACTCAGGATGTCGTGGCACCGATCATCGAGAACGGTGATTTGCT

GATAAGGATGGCACTCTTGCCAGCACCCATCGGGGGTCTCAATTATCTTAACATGAGCAGGTTATTTGTGAGAAATATCG

GTGACCCGGTCACCTCCTCTATAGCCGACCTGAAGAGGATGATAGATGCCGGGCTAATGCCAGAAGAAACATTGCATCAA

GTGATGACCCAGACCCCGGGAGAATCATCTTACCTAGATTGGGCAAGTGACCCTTATTCTGCCAACCTACCCTGCGTACA

GAGTATCACTCGCCTTCTCAAAAACATCACTGCACGATATATTTTAATCAGCAGCCCAAACCCGATGCTGAAAGGGTTAT

TTCATGAGGGGAGTAGAGATGAAGACGAGGAGCTTGCAAGTTTCCTAATGGATCGGCATATAATTGTTCCAAGAGCTGCA

CATGAAATCTTAGACCATAGTATAACCGGAGCAAGAGAAGCTATAGCCGGGATGTTGGACACCACCAAGGGTCTGATTAG

AACAAGTATGAAACGGGGCGGCCTCACCCCTCGAGTCTTAGCCCGCCTTTCCAATTATGATTATGAACAATTCCGATCCG

GGATAACACTATTAACAAAGAAAGGGCAGTGTTATCTCATTGACAAGGACTCGTGCTCGGTGCAGCTCGCTATAGCTCTG

AGGGGCCACATGTGGGCTAAGTTGGCTCGCGGGAGACCAATCTATGGGTTGGAGGTGCCTGATGTATTAGAATCGATGAA

CGGCTACCTTATTAAACGCCACGAGTCGTGTGCTATCTGTGAAACGGGCTCAAGTCACTACGGATGGTTTTTCGTCCCTG

CAGGGTGCCAGCTTGACGATGTATCTAGAGAGACTTCGGCTCTTCGTGTACCTTATGTCGGATCAACAACTGAGGAAAGG

ACAGACATGAAACTTGCATTTGTTAGGTCTCCAAGTAGATCCCTCAAATCAGCAGTTAGAATTGCAACAGTTTACTCATG

GGCTTACGGGGATGATGAGAAATCATGGAGTGAAGCCTGGATGCTAGCTAGGCAGAGAGCCAATATCACCTTAGATGAAT

TGAGAATGATCACTCCAGTCTCCACATCCACCAACTTAGCCCACCGGTTGAGGGATCGGAGTACCCAGGTGAAGTACTCG

GGGACATCCCTTGTGAGGGTTGCAAGATACACAACCATCTCCAATGATAACTTGTCGTTTGTGATATCTGAGAAAAAAGT

GGATACCAACTTCATTTATCAGCAAGGGATGTTGCTTGGTCTTGGGATCCTTGAGAACCTCTTTAGGCTAGAGGCCACCA

CCGGAGTATCCAACACAGTGCTACACCTGCACGTGGAAACAGAATGTTGTGTTGTACCTATGGTGGATCACCCAAGGATA

CCGAGTCTCCGTAATATAAAAGTTACGGATGAGCTATGCACAAACCCTTTGATCTACGACAGGTCCCCCATCATAGAGCA

CGATGCAACCCGGCTATATTCACAGAGCCACAGGAGACATTTGGTGGAGTTTGTTACCTGGTCAACAAGCCAGCTTTACC

ATATACTGGCTAAGTCTACAGCAATGTCCATGATTGAATTGATTACGAGATTCGAGAAAGATCACATGAATGAAATAGCC

GCCCTGATTGGCGATGACGACATCAACAGCTTCATCACAGAATTTTTGCTAGTGGAGCCTAGATTATTTATAGTTTACCT

TGGTCAGTGTGCTGCCATCAATTGGGCCTTTGATATACATTATCATCGGCCCTCGGGCAAGTACCAGATGGGGGAACTTC

TCTACTCTCTGCTCTCTCGGATGAGCAAAGGAGTATATAAGATCTTTACTAATGCTTTGAGCCACCCCAAAGTTTACAAG

AAATTTTGGCGAAGTGGCATCATTGAGCCGATTCATGGACCATCCTTGGATACACAGAATTTACATGTCACTGTCTGTGA

CATGATATATGGGTCCTATGTCACCTATTTGGATCTTTTGCTGAATGATGAGCTAGATGATTACCCGTATTTGCTCTGCG

AGAGTGATGAGGACGTGGTCACAGACAGGTTCGACAACATTCAAGCCAAACATCTCTGTGTATTGGCCGATGTATATTGC

AGCTCCAAGAGGTGTCCTTCAATAATCGGGATGTCTCCTATAGAAAAATGTACCATTCTCACACATTACATCAAGGGAGA

ATCAGTACAATCCCCGTCTGGGACCTCATGGAACACTGATCCCCTTGTAGTAGATCATTACTCATGCTCTCTGACCTACC

TTCGCCGCGGTTCCATCAAACAAATCAGGTTGAGGGTGGATCCTGGGTTTGTATTCGAGGCGTTGACAGACGTCGACTTT

AAACAACCTCGCAAAGCTAAGTTAGATATATCGGTCGTTGGGTTGACTGATTTCTCTCCCCCTTACGACAACGTCGGTGA

TTTTTTAGGAACTATCAACACATTGAGGCACGATCTGCCTGTCACCGGAACCGGGGTCTCGAACTATGAAGTCCACGCTT

ATCGTAGAATTGGCCTGAATTCATCAGCATGTTATAAAGCCGTAGAGATCTCCACGTTAATCAAGCCATCCTTAGAAGTC

GGAGAGCATGGATTGTTCTTAGGAGAAGGTTCCGGTTCAATGCTGGCTGCGTACAAAGAAGTTCTTAAATTAGCAAATTG

TTACTACAACAGCGGAGTAACAGCGGAGGGCAGAGCCGGACAGAGGGAAATATCTCCCTATCCTTCAGAGATGAGCCTAG

TAGAGAATCAAATGGGGATAGAGCGGAGTGTTAAAGTGCTGTTCAACGGGAAACCTGAAGTAACTTGGGTGGGGACCACA

GATTGCTACAAGTACATAATCAGTAACATCCAAACCTCTAGTCTGGGTTTCATACATTCAGATATTGAGACGCTCCCAAC

CAAAGATGCTGTTGAGAAATTGGAGGAATTTGCCTCTATCCTATCCTTATCCTTGATTTTGGGGAAAATCGGCTCTATTA

CAGTTGTCAAAATTATGCCCATTAGCGGGGATTTTACCCAAGGCTTCATAGCCTATGCCATCCAATATTTTAGGGAGAGC

CTGCTTGCCTATCCGAGATATAGTAACTTCATCTCGACTGAGTGTTACCTTATTATGATAGGATTAAAGGCCAATCGATT

AATAAACCCAGAAGCCATTAAGCAAAGCATAATCAGAGCGGGGGTCAGGACTGCACCAGGACTTGTGAGCCATGTATTAT

CAGAGAAACAAAAAGGTTGTATTCAATCTTTCTTGGGTGATCCTTATATCCAAGGAGACTTCAATAAGCACCTTAAATCT

CTAACCTCTATTGAGAAAATCCTGGTAAATTGTGGTCTCTCGATCAATGGCATAAAAATCTGTAGAGATCTAATCCACCA

TGATATCGCCTCCGGTCCAGACGGCCTTATGAGCTCTACAATTATTTTATATAGGGAATTGGCTCATTTCAAAGATAATA

TAAGGAGTCAGCACGGAATGTTCCACCCCTACCCGGTACTGGCCAACAGCAGGCAACGTGAATTAATCCTTCGAATAGCC

AAGAAATTCTGGGGGTATGTCTTGCTATATTCTGATGACCCGGCACTAATCAGACAAACAATCAAGAACTTGAAACGGAA

TCATCTAACCTTTGACTTACATAGTAACCCATTTATTAAAGGGCTATCCAAGGCTGAGAAACTGCTGGTGCGGACGAGTT

CACTAAGAAGAGAATGGTTGTTCACTCTCGAAACGAAAGAAGTAAAAGAGTGGTTCAAATTGGTAGGTTACAGTGCACTC

ATCAGAGGCTAGTCAGCTATGCATCTGCCCCCTTCTCCTCCGCCATGAGACCCCACTGGCGATCCAGAAGATTAAAGAAA

ACTACATATTGGATAAGTATCTATTCCCAGCTTTGTCTGGT

>Saiga1/saiga/Mongolia/2017-01

NNNNNNNNNNNNNNNNNNNNNNNNNNNNNNNNNNNNACTATGGACTGGCAAACTTAGGAGTAAAGATCCTACTGTCGGGG

GGAGGAGGAGGAGCAAGATCCTTGATCATGGCGACTCTCCTTAAAAGCTTAGCATTGTTCAAAAGGAACAAGGACAAAGC

GCCGACAGCATCAGGTTCAGGAGGGGCCATCCGGGGGATTAAGAATGTTATCATAGTCCCGATTCCCGGAGACTCGTCCA

TCACTACCCGTTCAAGACTGCTCGACAGGCTTGTCAGATTGGCCGGAGATCCTGACATCAACGGGTCAAAGCTGACCGGG

GTGATGATCAGCATGTTATCCTTGTTCGTAGAGTCACCCGGGCAATTGATACAGCGAATCACAGATGATCCGGATGTTAG

TATCCGCCTTGTTGAGGTAGTTCAAAGTACTAGGTCCCAGTCCGGGTTGACCTTTGCATCACGTGGTGCTGATTTAGACA

ACGAGGCAGACATGTATTTTTCAACTGAGGGGCCCTCGAGTGGAGGTAAGAAAAGGATCAACTGGTTTGAGAACAGAGAA

ATAATAGACATAGAGGTGCAGGATCCAGAAGAGTTCAATATGTTGTTAGCCTCCATACTAGCACAAGTCTGGATCCTTCT

GGCCAAGGCTGTTACGGCACCAGATACGGCAGCTGACTCAGAACTGAGAAGGTGGGTTAAATACACACAACAAAGGAGAG

TGATTGGGGAATTTCGCCTTGACAAGGGGTGGCTGGACGCAGTCCGCAACAGAATTGCAGAAGATCTATCACTCCGGCGG

TTCATGGTATCTCTCATACTTGACATCAAGAGGACCCCAGGCAACAAGCCAAGGATTGCAGAAATGATCTGCGACATTGA

CAACTATATTGTCGAAGCAGGACTCGCCAGTTTCATCCTTACTATCAAATTTGGTATCGAAACCATGTATCCTGCATTAG

GGCTCCACGAGTTTGCCGGAGAATTGTCCACTATTGAGTCCTTGATGAACTTGTATCAACAGCTAGGAGAGGTTGCACCC

TACATGGTAATTCTAGAGAATTCAATTCAGAACAAGTTTAGTGCAGGAGCTTATCCCCTCCTCTGGAGCTATGCGATGGG

TGTCGGAGTCGAGCTGGAGAACTCAATGGGGGGCTTGAACTTTGGCAGGTCATATTTTGACCCTGCCTATTTTCGTCTCG

GACAGGAGATGGTCAGAAGATCTGCAGGAAAGGTCAGCTCTGTGATTGCGGCTGAGCTCGGCATCACAGCAGAGGAAGCT

AAACTAGTCTCGGAAATCGCCTCGCAGGCCGGGGACGAAAGAACCGCCAGAGGGACTGGGCCTCGACAGGCGCAGGTTTC

CTTCCTTCAGCACCAAACAGGAGGGGGAGAGTCGTCCGCACCAGCGACCAGAGAAGGGGTCAAAGCTGCGATCCCAAACG

GAGCTGAAGAAAGGGACAGAAAACAAACACGCCCAGGAAGGCCCAGAGGAGAGACCTCCGGCCAACTGCTCCTGGACATC

ATGCCAGAGGATGAGATCTTGCGAGAGTCTGGTCAAAACCCTCGTGAGGCTCAAAGATCGGCCGAGGCACTCTTCAGGCT

GCAGGCCATGGCCAAGATTCTGGAGGACCAGGAGGAGGGAGAAGACAACAGTCAGGTCTACAACGACAAGGATCTCCTCG

GCTAAGCAGACGCACCCTCTGTCGAAATCAGTGACAAGACATCTCCTACCAGTATTATAAAAAACTTAGGACCCAGGTCC

AAGCAAGCACACATCGACACTCCAACCAGTCGAGCGGAGACCACCGATGGCAGAAGAACAAGCATACCATGTNAACAAGG

GGCTGGAATGTATCAAGTCTCTCAAAGCCTCACCCCCGGATCTATCCACCATCAGAGATACCATCGAGAGCTGGAGAGAG

GGGCTTAGCCCCTCGGGCCGTGCAACACCGAACCCTGATACGTCCGAGGGAGATCATCAGAATATCAACCAATCATGCTC

ACCAGCAATCGGACCAAACAAAGTCTACTTGTCTCCTGAAGATAATCTCGGATTTAGAGAGATCACTGGCAACGACTGTG

AGGCTGGGCTCGGAGGGGTCCAGAGAGAAGGATCCAACTCTCAAGTACNNNNNNNNNNNNNNNNNNNNNNNNGGGGTGAA

GAGATTGAAGGACTCGAGGATGCTGACTCTCTCGTGGTTCAAGCAGATCCTCCAGCTGCTAACGTCTTCAATGGAGGAGA

GGATGGATCTGACGACAGCGATGTGGACTCTGGCCCAGATGATCCCGGCAGAGATACTCTATATGACCGGGGATCTGTTG

CCGGCAATGGTATCGCTAGGTCCACAGATGTCGAAAAACTAGAAGGTGCTGATATTCAAGAAGTTCTTAACTCCCAGAAA

GGCAAAGGAGGAAGATTCCAGGGCGGGAAAACCTTGCGAGTCCCGGAAATACCCGATGTCAAGCACTCCAGACCATCAGC

CCAATCAATTAAAAAGGGCACAGACGGGAACTCAGTCTCATCTGGAACGGTGATAGAGTGTTTATCGATAAGTGGTGCAA

CCCAAACTGTGCCAGAGTCAAGATGGGAGTCATCAGAGCAAAATGCGTCTGTGGGGAGTGTCCTCAAGTCTGCGAGGAGT

GCAAAGACGATCCAGGGGCCGACACAAGAATCTGGTACCATAGCATCACTGACTCAGCCTAAAGAGAATGACTCCGAGTA

TGAGTATGAGGATGACCTATTTACAGAGATTCAGGACATCCGTGCAAGCATTGCCAAGATCCATGATGACAATAAAACTA

TCCTCTCAAAGCTTGATTCTATACTGTTATTGAAAGGGGAAGTCGACACTATCAAGAAACAAATCAGCAAGCAGAATATA

AGTATATCCACCATTGAGGGCCATCTCTCCAGTATAATGATAGCCATCCCTGGCTTTGGGAAGGACATCATGGACCCAAC

ATCCGAAGTCGAGTTGAACCCAGATCTAAGACCTATAATAAGCCGTGACTCTGGCAGAGCTCTCGCGGAGGTCCTCAAGA

AACCCGCTGTGGATAGGTCTCAGAAAATTGGAACCAAAGCCAACTCCAGCTCAAAGGGTCAGCTTCTTAAGGATCTCCAG

CTAAAACCTGTCGACAAGCAGGCAAGCTCTGCAATCGGGTTTGTCCCGTCCGACCATGAATCATCCAGAAATGTCATCCG

CTCCATAATCAAGTCGAGCAAGCTAAATATTGATCACAAGGACTATCTTCTAGATTTACTAAATGATGTGAAAGGCTCCA

AGGATCTTAAGGAATTTCACAAGATGCTAACAGCAATTCTCGCCAAGCACCTGTAACACATCCCCCAGTCATCATCTCAT

ACTCGACTAAAAACATCCTTTCAATCAGGCTATTACAAAAAACTTAGGAGCAAGGGCAACTGAGCTTCGCAGACAGGANN

NNNNNNNNNNNNNNNNNNNNNNNNNNNNNATGGGATGTCAAAGGGTCAATTGCCCGCATAGAACCCACCACCTACCACGA

CGGCCGACTGATACCCCAGGTGAGGGTCATCGATCCTGGTCTGGGAGACAGAAAAGATGAGTGCTTTATGTACCTGTTTC

TCCTAGGAGTGATTGAGGATAACGACCCCCTGTCTCCCCCCGTCGGAAGAACCTTTGGCTCTCTACCTCTAGGGGTTGGT

AGGTCAACTGCCAAGCCAGAAGAACTACTAAGGGAGGCCACAGAATTAGATATAGTGGTGAGGCGCACTGCAGGAGTAAA

TGAGAAACTGGTATTTTACAACAACACTCCGCTGTCCTTGTTAACGCCATGGAAGAAAGTTCTGACAACCGGAAGTGTGT

TTAGCGCTAACCAGGTTTGCAATGCAGTCAACCTAGTCCCGCTTGATACTCCCCAGAGATTCAGGGTTGTGTACATGAGC

ATAACTAGATTATCAGACAATGGTTACTATAGTGTGCCCAGAAGAATGCTGGAGTTCCGCTCGGCCAATGCAGTAGCCTT

CAACATCTTGGTTACACTAAGAATTGAAAATGGCACAAACCCTAGAAGATACATAGTCGGCTCATGGGAGAATTCAGAGG

TCACATTTATGGTACACGTGGGCAACTTTAGGAGAAAGAAGAACGAAGTATACTCTGCTGATTACTGCAAAATGAAGATT

GAAAAGATGGGTTTAGTTTTTGCCCTGGGTGGAATAGGCGGAACAAGTCTCCATATTAGAAGCACCGGGAAAATGAGCAA

AACCCTCCATGCACAGCTGGGGTTCAAGAAAATCTTATGTTACCCCCTAATGGATGTCAATGAGGATCTTAACCGATATC

TCTGGCGGGCAGAGTGCCGAATAGTCAAAATCCAAGCTGTCTTACAGCCATCAGTACCCCAAGAGTTCCGTGTCTACGAT

GATGTCATCATCAACGATGATCAAGGCNNNNNNNNNNNNNNNNNNNNNNNNNNNNNNNNNNNNNNNNNNNNNNNNNNNNN

NNNNNNNNNNNNNNNNNN-NNNNNNNNNNNNNNNNNNNNNNNNNNNNNNNNNNNNNNNNNNNNNNNNNNNNNNNNNNNNN

NNNNNNNNNNN-NNNNNNNNNNNNNNNNNNNNNNNNNNNNNNNNNNNNNNNNNNNNNNNNNNNNNNNNNNNNNNNNNNNN

NNNNNNNNNNNNNNNNNNNNNNNNNNNNNNNNNNNNNNNNNNNNNNNNNN-NNNNNNNNNNNNN-NNNNNNNNNNNNNNN

NNNNNNNNNNNNNNNNNNNNNNNNNNNNNNNNNNNNNNNNACCCCAAGGGGGGACCCTCCAAACCAAAGGAACCACCGGG

GTAGGCTCCGCCGAACCAGCACAGACCCGACCCAAGCAAGGCGGACCCCCAACACGAAACTGCCCAGCACACCCTACTCA

TCATCAAACAAAACTTAGGGGCCAAGTCCACACGAGAGGCCCCAGTCCGTCAGGCGGAGGGACGCAGAAAGGAAGGAGAC

ACGACCGTCCCAGAAGGNNNNNNNNNNNNNNNNNNNNNNNNNNNNNNNNNNNNNNNNN--NNN-NNNNNNNNNNNNNNNN

NNNNNNNNNNNNNNNNNNNNNNNNNNNNNNNNNNNNNNNNNNNNNNNNNNNNNNNNNNNNNNNNNNNNNNNNNNNNNNNN

NNNNNNNNNNNNNNNNNNNNNNNNNNNNNNNNNNNNNNNNNNNNNNNNNNNNNNNNNNNNNNNNNNNNNNNNNNNNNNNN

NNNNNNNNNNNNNNNNNNNNNNNNNNNNNNNNNNNNNNNNNNNNNNNNNNNNNNNNNNNNNNNNNNNNNNNNNNNNNNNN

NNNNNNNNNNNNNNNNNNNNNNNNNNNNNNNNNNNNNNNNNNNNNNNNNNNNNNNNNNNNNNNNNNNNNNNNNNNNNNNN

NNNNNNNNNNNNNNNNNNNNNNNNNNNNNNNNNNNNNNNNNNNNNNNNNNNNNNNNNNNNNNNNNNNNNNNNNNNNNNNN

NNNNNNNNNNNNNNNNNNNNNNNNNNNNNNNNNNNNNNNNNNNNNNNNNNNNNNNNNNNNNNNNNNNNNNNNNNNNNNNN

NNNNNNNNNNNNNNNNNNNNNNNNNNNNNNNNNNNNNNNNNNNNNNNNNNNNNNNNNNNNNNNNNNNNNNNNNNNNNNNN

NNNNNNNNNNNNNNNNNNNNNNNNNNNNNNNNNNGGAACAGGGAGTGCCAGCTACAAGGTGATGACTAGGCCAAGCCACC

AGACTCTGGTTATAAAGTTAATGCCAAATATAACGGCCATCGACAATTGTACAAAGTCAGAGATTGCAGAGTACAAGAGA

TTGCTGATCACAGTGTTAAAGCCTGTAGAGGATGCTCTGTCGGTGATAACCAAGAATGTAAGACCAATTCAAACTCTAAC

ACCTGGGCGTAGAACCCGCCGTTTTGCTGGAGCTGTTCTGGCCGGGGTAGCACTTGGAGTTGCGACAGCCGCTCAGATAA

CTGCAGGAGTCGCCCTTCATCAATCATTGATGAACTCCCAAGCAATTGAGAGTTTAAAAACCAGTCTTGAGATGTCGAAT

CAGGCAATAGAAGAAATCAGACTTGCAAATAAGGAGACCATACTGGCAGTACAGGGCGTCCAGGATTATATCAACAATGA

GCTCGTCCCTTCTGTTCATAGAATGTCATGCGAGCTGGTAGGTCACAAGCTCGGCCTCAAGCTCCTTAGGTACTACACCG

AGATCCTGTCCATATTCGGGCCCAGTCTTCGAGACCCGATATCTGCCGAAATATCAATCCAGGCACTTAGTTATGCATTA

GGCGGAGACATTAATAAAATCCTGGACAAGCTTGGGTATAGCGGTGGGGATTTCCTTGCCATCCTAGAAAGCAAGGGAAT

AAAGGCCCGGGTTACATATGTGGACACAAGAGATTACTTTATAATCCTTAGCATCGCCTACCCAACCTTATCTGAGATCA

AGGGAGTGATAGTTCACAAGATAGAAGCTATAACATACAACATTGGGGCACAGGAGTGGTATACTACTATCCCTAAATAT

GTAGCCACTCAGGGGTATCTGATATCGAACTTTGATGAGACGTCATGCGTATTCACTCCAGAGGGGACAGTTTGCAGCCA

GAATGCGTTGTACCCAATGAGCCCATTGCTTCAGGAATGTTTCAGGGGGTCAACAAAATCGTGCGCCAGAACCTTAGTTT

CAGGGACCATAAGTAATAGATTTATCCTATCAAAAGGGAACCTGATTGCAAATTGTGCGTCAGTTTTGTGCAAATGTTAC

ACAACAGAGACAGTTATCAGCCAAGATCCTGACAAACTACTAACTGTTGTAGCATCCGACAAGTGTCCTGTAGTTGAGGT

GGATGGAGTGACAATACAGGTCGGCAGTCGAGAGTNNNNNNNNNNNNNNNNNNNNNNNNNNNNNNNNNNNNNNNNNNNNN

NNNNNNNNNNNNNNNNNGATGTAGGCACCAATTTAGGCAATGCAGTCACAAGACTGGAGAATGCAAAGGAGCTCCTAGAT

GCATCAGACCAAATACTGAAGACTGTTAAAGGGGTACCTTTCAGTGGGAATATGTACATAGCACTGGCAGCTTGCATAGG

AGTATCCCTGGGCCTTGTCACATTAATATGCTGCTGTAAGGGGAGGTGTAAGAACAAGGAAATCCCTATCTCCAAAATCA

ACCCAGGGCTCAAACCCGACCTGACCGGGACCTCAAAGTCGTACGTAAGATCACTGTAGTCAGAATCACCTGGATCATCT

GGCATCACACACATACATGCACGACACAGGCAGTCCGATGACGCAAGAAACCCAACCTCCGGTCACCCACCCGACCCCAC

TCCACGCTCCACCACACATTAGTCATCAAACAAAACTTAGGACGAAAGGTCAATCACCATGTCCGCACAAAGGGAGAGGA

TCAATGCCTTCTACAAAGACAATCCTCACAATAAGAACCATAGGGTGATCCTGGATAGAGAACGCTTGGTCATTGAAAGA

CCCTACATCTTGCTTGGAGTCCTGCTGGTAATGTTCCTGAGTCTAATCGGACTGCTGGCCATTGCAGGGATCAGGCTTCA

CCGGGCCACCGTTGGAACTTCAGAGATCCAGAGTCGGCTGAATACCAATATTAAGTTGACCGAATCTATTGATCACCAGA

CTAAGGATGTCTTAACTCCCCTTTTTAAAATCATTGGCGATGAAGTCGGCATCAGAATTCCACAGAAATTCAGTGATCTT

GTCAAGTTCATCTCCGATAAGATTAAATTCCTCAACCCTGATAGAGAGTATGATTTCAGAGATCTCCGGTGGTGTATGAA

TCCTCCCGAGAGAGTCAAAATTAATTTTGATCAGTTTTGTGAGTACAGGGCTGCGGTTAAGTCAATTGAACATATATTTG

AGTCACCACTCAACAAGTCAAAAAAGCTGCAATCTTTGACTCTCGGGCCCGGAACAGGCTGTCTAGGCAGGACAGTAACA

AGAGCCCATTTCTCAGAACTTACAATGACCTTAATGGACCTGGATCTAGAGATGAAGCACAACGTGTCCTCAGTGTTTAC

CGTAGTTGAAGAGGGATTATTCGGAAGAACATATACCGTCTGGAGATCCGATGCCAGGGATCCGAGCGCCGATCTAGGTA

TTGGCCATTTTTTAAGAGTCTTCGAGATTGGACTGGTAAGAGATCTTGGGCTGGGTCCCCCTGTTTTTCATATGACCAAC

TATCTCACGGTGAACATGAGTGATGACTATCGGAGATGTCTTTTAGCGGTAGGGGAGTTGAAGTTGACAGCCCTATGCAC

CTCATCTGAGACTGTGACACTGAGTGAGAGAGGAGTTCCAAGGAGGGAACCTCTTGTGGTTGTGATACTTAATCTAGCTG

GACCCACTCTAGGGGGAGAGCTATACAGTGTCTTGCCTACCTCTGATCTCATGGTGGAGAAACTCTATTTGTCTTCACAT

AGAGGGATCATCAAAGATGATGAGGCCAATTGGGTAGTGCCGTCTACCGATGTTCGTGATCTTCAGAACAAAGGTGAATG

TCTGGTGGAAGCATGCAAGACTCGACCTCCTTCATTTTGCAATGGCACAGGATCAGGCCCGTGGTCAGAAGGGAGAATCC

CTGCCTACGGGGTGATCAGGGTCAGTCTTGACTTAGCGAGTGACCCGGATGTAGTTATCACTTCAGTGTTTGGCCCACTG

ATACCTCACCTATCCGGCATGGATCTTTACAACAACCCGTTTTCAAAAGCTGTATGGTTGGCTGTACCACCTTATGAGCA

GTCATTTCTAGGAATGATAAATACAATTGGATTCCCTAACAGAGCAGAGGTTATGCCGCACATTTTGACCACAGAGATCA

GAGGCCCTCGGGGTCGTTGCCATGTTCCCATAGAATTGTCCCGCCGGGTTGATGACGATATCAAGATCGGGTCCAACATG

GTCATATTGCCGACGATGGACCTGAGGTATATTACAGCCACTTATGATGTTTCCAGGAGCGAGCATGCAATCGTGTACTA

TATCTATGACACGAGTCGCTCATCATCTTACTTCTACCCAGTTCGACTGAATTTCAAAGGCAATCCTCTCTCTTTGAGGA

TAGAGTGTTTCCCCTGGCGTCATAAGGTGTGGTGCTACCATGATTGTCTTATATACAACACCATAACAGGTGAAGAGGTC

CATACGAGAGGGCTGACCGGCATAGAGGTAACATGCAATCCAGTCTGAGCAGAGCTACGACCGTCACTCAGGCAGTCCCT

TGAGTCGCCACCGAGTCCAAGCAGCACAGCCTGGGACAATCAACGGCACAGCCCAGCCAACAATGTTATAAAAAACTTAG

GAGCCAGGGTTGTAGGGGCCATGGACTCTCTATCAGTCAATCAGGTCTTGTACCCTGAGGTTCATCTAGACAGCCCTATT

GTCACAAACAAACTAGTTGCCATCCTTGAGTACTCGGGTATCGACCATAACTATGTTCTTGAAGACCAGACTCTTGTCAA

GAATATTAGGTATAGGCTGGGGTGCGGTTTTTCAAATCAAATGATCATCAATAACAGGGGGGTGGGTGAAACAGTCAATT

CTAAACTTAAAAGTTACCCCCATAATCGTCATATCATATACCCGGATTGCAATAAGGAGTTGTTTTGTATCAAGGATAGC

TGCATATCTAAGAAGCTCTCGGAGCTATTCAAGAAGGGTAATTCCTTGTACTCTAAGATAAGTCACCAGGTACTGGATTG

TCTTAAGAGAGTCAATGGGAAATTAGGTCTGGGCACAGATCTTACCCATGGTCTGAAGGAGGGTATCCTTGACCTAGGGT

TGCACATGCATAGCTCTCAATGGTTCGAGACTTTCCTGTTCTGGTTCACTATCAAGACAGAGATGAGATCAATGATCAAA

GAACAGTCCCATATATGCCACAAGAGGAGGTATAACCCGACTTTTGTGTCGGGGGATGCATTCGAGGTGCTCGTATCGCG

AGACCTCGTTGTGATAATTGATAAGAATACCCAGTATGTCTTCTACTTAACATTTGAGCTGGTCCTTATGTACTGTGATG

TCATAGAGGGTAGACTTATGACGGAGACAGCCATGGCCATAGACCAGAGATATTCAGAACTTCTAAGCCGGGTCAGATAC

TTGTGGGATCTTATTGATGGGTTCTTCCCAACNNNNNNNNNNNNNNNNNNNNNNNNNNNNNNNNNGCTTGAACCATTATC

GTTGGCTTATCTTCAACTTCAGGATGTCACTCTGGAGTTAAGAGGTGCTTTTTTAGACCATTGCTTCAAAGAACTCTATG

AGATACTGGAACATTGCGGCATTGACACAGCAGGCACTTACAATTCCATCACTGAAGGATTGGATTACGTATTCATCACC

CACGATATACATCTAACTGGGGAGATTTTTTCATTTTTCCGGAGTTTCGGACACCCCCGTCTAGAAGCAGTCACCGCTGC

GGAAAATGTCAGAAAACATATGAACCAACCGAAGGTAATCAGTTATGAGANNNNNNNNNNNNNCCATGCGGTATTTTGCG

GGATAATCATAAATGGTTTCCGGGACCGACATGGCGGCAGCTGGCCCCCTGTTGCACTGCCAGAACATGCCTCTGCTGCG

ATCCGGAATGCGCAGGCATCAGGTGAGGGACTGACCCATGACCTGTGTATAGACAACTGGAAATCCTTTGTTGGATTCAA

ATTTGGCTGCTTTATGCCACTCAGCCTAGATAGTGATTTGACCATGTATCTCAAAGACAAGGCATTGGCTGCACTGAAGA

ATGAGTGGGATTCAGTTTACCCGAAAGAATACCTCCGTTATAACCCACCTAGAGGGACAGAGTCAAGGCGATTGGTAGAG

GTATTCCTGAATGACTCCAGCTTTGATCCTTATAACATGATAATGTACGTGGTGAATGGCTCCTACCTTAAAGACCCTGA

GTTTAACCTCTCATACAGCCTAAAAGAGAAGGAGATAAAGGAGACAGGGCGGTTGTTTGCCAAAATGACCTATAAGATGC

GGGCCTGTCAGGTAATCGCTGAAAATCTGATATCGAATGGTGTTGGGAAGTATTTCCGAGACAATGGGATGGCAAAGGAC

GAGCATGACCTAACAAAAGCCCTTCACACCCTGGCAGTCTCAGGTGTTCCCAAAAACAACAAAGATAACCACCGAGGTGG

GCCTCCCAGAAGGACAACAAACCGAGGGGTGAGATCAAGCCAAGGCACCAAAACACAAGATAGAGACAAGGTTCAAGGGG

GACCTATGTACAACTATTTGCGATGCCAACCGATCAGCCCTGATCAGGGTGAGTCATACGAGACTGTTAGTGCATTCATC

ACCGCTGACCTTAAGAAGTATTGCCTGAATTGGAGATACGAGACAATCAGCATATTTGCACAGAGGCTGAATGAAATATA

TGGACTACCATCCTTCTTTCAATGGTTACACAGGATATTGGAAAAATCCGTACTCTACGTCAGTGACCCACATTGTCCTC

CCGATCTAGATAATCATATCCCTCTGGACAGTGTCCCCAATGCCCAAATATTCATCAAGTACCCAATGGGCGGAATAGAA

GGTTATTGCCAAAAACTATGGACAATCAGCACTATACCATACTTGTATCTGGCAGCCTATGAGAGCGGAGTAAGAATCGC

CTCACTGGTGCAGGGTGACAATCAGACAATCGCAGTGACAAAAAGAGTTCCAAGTTCTTGGCCTTATTCACTAAAAAAGA

GGGAGGCCTCCAAGGCCGCTCAAAATTACTTCGTAGTCTTAAGGCAAAGATTGCACGATGTAGGTCATCACTTAAAGGCT

AATGAGACCATAGTATCTTCTCACTTTTTTGTATATTCCAAAGGGATTTATTATGACGGCCTGCTAGTCTCACAATCACT

AAAGAGCATCGCCAGATGTGTCTTCTGGTCCGAGACTATAGTGGATGAAACCAGAGCGGCGTGCAGTAATATTGCAACAA

CTGTCGCTAAGAGTATAGAGAGGGGTTATGATAGGTACCTTGCATATTCTTTGAATATCCTCAAGATTTTCCAACAGATC

CTTATATCCCTTAACTTCACTATTAACACAACAATGACTCAGGATGTCGTGGCACCGATCATCGAGAACGGTGATTTGCT

GATAAGGATGGCACTCTTGCCAGCACCCATCGGGGGTCTCAATTATCTTAACATGAGCAGGTTATTTGTGAGAAATATCG

GTGACCCGGTCACCTCCTCTATAGCCGACCTGAAGAGGATGATAGATGCCGGGCTAATGCCAGAAGAAACATTGCATCAA

GTGATGACCCAGACCCCGGGAGAATCATCTTACCTAGATTGGGCAAGTGACCCTTATTCTGCCAACCTACCCTGCGTACA

GAGTATCACTCGCCTTCTCAAAAACATCACTGCACGATATATTTTAATCAGCAGCCCAAACCCGATGCTGAAAGGGTTAT

TTCATGAGGGGAGTAGAGATGAAGACGAGGAGCTTGCAAGTTTCCTAATGGATCGGCATATAATTGTTCCAAGAGCTGCA

CATGAAATCTTAGACCATAGTATAACCGGAGCAAGAGAAGCTATAGCCGGGATGTTGGACACCACCAAGGGTCTGATTAG

AACAAGTATGAAACGGGGCGGCCTCACCCCTCGAGTCTTAGCCCGCCTTTCCAATTATGATTATGAACAATTCCGATCCG

GGATAACACTATTAACAAAGAAAGGGCAGTGTTATCTCATTGACAAGGACTCGTGCTCGGTGCAGCTCGCTATAGCTCTG

AGGGGCCACATGTGGGCTAAGTTGGCTCGCGGGAGACCAATCTATGGGTTGGAGGTGCCTGATGTATTAGAATCGATGAA

CGGCTACCTTATTAAACGCCACGAGTCGTGTGCTATCTGTGAAACGGGCTCAAGTCACTACGGATGGTTTTTCGTCCCTG

CAGGGTGCCAGCTTGACGATGTATCTAGAGAGACTTCGGCTCTTCGTGTACCTTATGTCGGATCAACAACTGAGGAAAGG

ACAGACATGAAACTTGCATTTGTTAGGTCTCCAAGTAGATCCCTCAAATCAGCAGTTAGAATTGCAACAGTTTACTCATG

GGCTTACGGGGATGATGAGAAATCATGGAGTGAAGCCTGGATGCTAGCTAGGCAGAGAGCCAATATCACCTTAGATGAAT

TGAGAATGATCACTCCAGTCTCCACATCCACCAACTTAGCCCACCGGTTGAGGGATCGGAGTACCCAGGTGAAGTACTCG

GGGACATCCCTTGTGAGGGTTGCAAGATACACAACCATCTCCAATGATAACTTGTCGTTTGTGATATCTGAGAAAAAAGT

GGATACCAACTTCATTTATCAGCAAGGGATGTTGCTTGGTCTTGGGATCCTTGAGAACCTCTTTAGGCTAGAGGCCACCA

CCGGAGTATCCAACACAGTGCTACACCTGCACGTGGAAACAGAATGTTGTGTTGTACCTATGGTGGATCACCCAAGGATA

CCGAGTCTCCGTAATATAAAAGTTACGGATGAGCTATGCACAAACCCTTTGATCTACGACAGGTCCCCCATCATAGAGCA

CGATGCAACCCGGCTATATTCACAGAGCCACAGGAGACATTTGGTGGAGTTTGTTACCTGGTCAACAAGCCAGCTTTACC

ATATACTGGCTAAGTCTACAGCAATGTCCATGATTGAATTGATTACGAGATTCGAGAAAGATCACATGAATGAAATAGCC

GCCCTGATTGGCGATGACGACATCAACAGCTTCATCACAGAATTTTTGCTAGTGGAGCCTAGATTATTTATAGTTTACCT

TGGTCAGTGTGCTGCCATCAATTGGGCCTTTGATATACATTATCATCGGCCCTCGGGCAAGTACCAGATGGGGGAACTTC

TCTACTCTCTGCTCTCTCGGATGAGCAAAGGAGTATATAAGATCTTTACTAATGCTTTGAGCCACCCCAAAGTTTACAAG

AAATTTTGGCGAAGTGGCATCATTGAGCCGATTCATGGACCATCCTTGGATACACAGAATTTACATGTCACTGTCTGTGA

CATGATATATGGGTCCTATGTCACCTATTTGGATCTTTTGCTGAATGATGAGCTAGATGATTACCCGTATTTGCTCTGCG

AGAGTGATGAGGACGTGGTCACAGACAGGTTCGACAACATTCAAGCCAAACATCTCTGTGTATTGGCCGATGTATATTGC

AGCTCCAAGAGGTGTCCTTCAATAATCGGGATGTCTCCTATAGAAAAATGTACCATTCTCACACATTACATCAAGGGAGA

ATCAGTACAATCCCCGTCTGGGACCTCATGGAACACTGATCCCCTTGTAGTAGATCATTACTCATGCTCTCTGACCTACC

TTCGCCGCGGTTCCATCAAACAAATCAGGTTGAGGGTGGATCCTGGGTTTGTATTCGAGGCGTTGACAGACGTCGACTTT

AAACAACCTCGCAAAGCTAAGTTAGATATATCGGTCGTTGGGTTGACTGATTTCTCTCCCCCTTACGACAACGTCGGTGA

TTTTTTAGGAACTATCAACACATTGAGGCACGATCTGCCTGTCACCGGAACCGGGGTCTCGAACTATGAAGTCCACGCTT

ATCGTAGAATTGGCCTGAATTCATCAGCATGTTATAAAGCCGTAGAGATCTCCACGTTAATCAAGCCATCCTTAGAAGTC

GGAGAGCATGGATTGTTCTTAGGAGAAGGTTCCGGTTCAATGCTGGCTGCGTACAAAGAAGTTCTTAAATTAGCAAATTG

TTACTACAACAGCGGAGTAACAGCGGAGGGCAGAGCCGGACAGAGGGAAATATCTCCCTATCCTTCAGAGATGAGCCTAG

TAGAGAATCAAATGGGGATAGAGCGGAGTGTTAAAGTGCTGTTCAACGGGAAACCTGAAGTAACTTGGGTGGGGACCACA

GATTGCTACAAGTACATAATCAGTAACATCCAAACCTCTAGTCTGGGTTTCATACATTCAGATATTGAGACGCTCCCAAC

CAAAGATGCTGTTGAGAAATTGGAGGAATTTGCCTCTATCCTATCCTTATCCTTGATTTTGGGAAAAATCGGCTCTATTA

CAGTTGTCAAAATTATGCCCATTAGCGGGGATTTTACCCAAGGCTTCATAGCCTATGCCATCCAATATTTTAGGGAGAGC

CTGCTTGCCTATCCGAGATATAGTAACTTCATCTCGACTGAGTGTTACCTTATTATGATAGGATTAAAGGCCAATCGATT

AATAAACCCAGAAGCCATTAAGCAAAGCATAATCAGAGCGGGGGTCAGGACTGCACCAGGACTTGTGAGCCATGTATTAT

CAGAGAAACAAAAAGGTTGTATTCAATCTTTCTTGGGTGATCCTTATATCCAAGGAGACTTCAATAAGCACCTTAAATCT

CTAACCTCTATTGAGAAAATCCTGGTAAATTGTGGTCTCTCGATCAATGGCATAAAAATCTGTAGAGATCTAATCCACCA

TGATATCGCCTCCGGTCCAGACGGCCTTATGAGCTCTACAATTATTTTATATAGGGAATTGGCTCATTTCAAAGATAATA

TAAGGAGTCAGCACGGAATGTTCCACCCCTACCCGGTACTGGCCAACAGCAGGCAACGTGAATTAATCCTTCGAATAGCC

AAGAAATTCTGGGGGTATGTCTTGCTATATTCTGATGACCCGGCACTAATCAGACAAACAATCAAGAACTTGAAACGGAA

TCATCTAACCTTTGACTTACATAGTARCCCATTTATTAAAGGGCTATCCAAGGCTGAGAAACTGCTGGTGCGGACGAGTT

CACTAAGAAGAGAATGGTTGTTCACTCTCGAAACGAAAGAAGTAAAAGAGTGGTTCAAATTGGTAGGTTACAGNNNNNNN

NNNNNNNNNNNNNNNNNNNNNNNNNNNNNNNNNNNNNNNNNNNNNNNNNNNNNNNNNNNNNNNNNNNNNNNNNNNNNNNN

NNNNNNNNNNNNNNNNNNNNNNNNNNNNNNNNNNNNNNNNN

>GoiteredGazelle/GoiteredGazelle/Mongolia/2017-01

ACCAAACAAAGTTGGGTAAGGATAGATCTTATAATAACTATGGACTGGCAAACTTAGGAGTAAAGATCCTACTGTCGGGG

GGAGGAGGAGGAGCAAGATCCTTGATCATGGCGACTCTCCTTAAAAGCTTAGCATTGTTCAAAAGGAACAAAGACAAAGC

GCCGACAGCATCAGGTTCAGGAGGGGCCATCCGGGGGATTAAGAATGTTATCATAGTCCCGATTCCCGGAGACTCGTCCA

TCACTACCCGTTCAAGACTGCTCGACAGGCTTGTCAGATTGGCCGGAGATCCTGACATCAACGGGTCAAAGCTGACCGGG

GTGATGATCAGCATGTTATCCTTGTTCGTAGAGTCACCCGGGCAATTGATACAGCGAATCACAGATGATCCGGATGTTAG

TATCCGCCTTGTTGAGGTAGTTCAAAGTACTAGGTCCCAGTCCGGGTTGACCTTTGCATCACGTGGTGCTGATTTAGACA

ACGAGGCAGACATGTATTTTTCAACTGAGGGGCCCTCGAGTGGAGGTAAGAAAAGGATCAACTGGTTTGAGAACAGAGAA

ATAATAGACATAGAGGTGCAGGATCCAGAAGAGTTCAATATGTTGTTAGCCTCCATACTAGCACAAGTCTGGATCCTTCT

GGCCAAGGCTGTTACGGCACCAGATACGGCAGCTGACTCAGAACTGAGAAGGTGGGTTAAATACACACAACAAAGGAGAG

TGATTGGGGAATTTCGCCTTGACAAGGGGTGGCTGGACGCAGTCCGCAACAGAATTGCAGAAGATCTATCACTCCGGCGG

TTCATGGTATCTCTCATACTTGACATCAAGAGGACCCCAGGCAACAAGCCAAGGATTGCAGAAATGATCTGCGACATTGA

CAACTATATTGTCGAAGCAGGACTCGCCAGTTTCATCCTTACTATCAAATTTGGTATCGAAACCATGTATCCTGCATTAG

GGCTCCACGAGTTTGCCGGAGAATTGTCCACTATTGAGTCCTTGATGAACTTGTATCAACAGCTAGGAGAGGTTGCACCC

TACATGGTAATTCTAGAGAATTCAATTCAGAACAAGTTTAGTGCAGGAGCTTATCCCCTCCTCTGGAGCTATGCGATGGG

TGTCGGAGTCGAGCTGGAGAACTCAATGGGGGGCTTGAACTTTGGCAGGTCATATTTTGACCCTGCCTATTTTCGTCTCG

GACAGGAGATGGTCAGAAGATCTGCAGGAAAGGTCAGCTCTGTGATTGCGGCTGAGCTCGGCATCACAGCAGAGGAAGCT

AAACTAGTCTCGGAAATCGCCTCGCAGGCCGGGGACGAAAGAACCGCCAGAGGGACTGGGCCTCGACAGGCGCAGGTTTC

CTTCCTTCAGCACCAAACAGGAGGGGGAGAGTCGTCCGCACCAGCGACCAGAGAAGGGGTCAAAGCTGCGATCCCAAACG

GAGCTGAAGAAAGGGACAGAAAACAAACACGCCCAGGAAGGCCCAGAGGAGAGACCTCCGGCCAACTGCTCCTGGACATC

ATGCCAGAGGATGAGATCTTGCGAGAGTCTGGTCAAAACCCTCGTGAGGCTCAAAGATCGGCCGAGGCACTCTTCAGGCT

GCAGGCCATGGCCAAGATTCTGGAGGACCAGGAGGAGGGAGAAGACAACAGTCAGGTCTACAACGACAAGGATCTCCTCG

GCTAAGCAGACGCACCCTCTGTCGAAATCAGTGACAAGACATCTCCTACCCGTATTATAAAAAACTTAGGACCCAGGTCC

AAGCAAGCACACATCGACACTCCAACCAGTCGAGCGGAGACCACCGATGGCAGAAGAACAAGCATACCATGTNAACAAGG

GGCTGGAATGTATCAAGTCTCTCAAAGCCTCACCCCCGGATCTATCCACCATCAGAGATACCATCGAGAGCTGGAGAGAG

GGGCTTAGCCCCTCGGGCCGTGCAACACCGAACCCTGATACGTCCGAGGGAGATCATCAGAATATCAACCAATCATGCTC

ACCAGCAATCGGACCAAACAAAGTCTACTTGTCTCCTGAAGATAATCTCGGATTTAGAGAGATCACTGGCAACGACTGTG

AGGCTGGGCTCGGAGGGGTCCAGAGAGAAGGATCCAACTCTCAAGTACAGCGTTACCATGTTTATAGTCACGGGGGTGAA

GAGATTGAAGGACTCGAGGATGCTGACTCTCTCGTGGTTCAAGCAGATCCTCCAGCTGCTAACGTCTTCAATGGAGGAGA

GGATGGATCTGACGACAGCGATGTGGACTCTGGCCCAGATGATCCCGGCAGAGATACTCTATATGACCGGGGATCTGTTG

CCGGCAATGGTATCGCTAGGTCCACAGATGTCGAAAAACTAGAAGGTGCTGATATTCAAGAAGTTCTTAACTCCCAGAAA

GGCAAAGGAGGAAGATTCCAGGGCGGGAAAACCTTGCGAGTCCCGGAAATACCCGATGTCAAGCACTCCAGACCATCAGC

CCAATCAATTAAAAAGGGCACAGACGGGAACTCAGTCTCATCTGGAACGGTGATAGAGTGTTTATCGATAAGTGGTGCAA

CCCAAACTGTGCCAGAGTCAAGATGGGAGTCATCAGAGCAAAATGCGTCTGTGGGGAGTGTCCTCAAGTCTGCGAGGAGT

GCAAAGACGATCCAGGGGCCGACACAAGAATCTGGTACCATAGCATCACTGACTCAGCCTAAAGAGAATGACTCCGAGTA

TGAGTATGAGGATGACCTATTTACAGAGATTCAGGACATCCGTGCAAGCATTGCCAAGATCCATGATGACAATAAAACTA

TCCTCTCAAAGCTTGATTCTATACTGTTATTGAAAGGGGAAGTCGACACTATCAAGAAACAAATCAGCAAGCAGAATATA

AGTATATCCACCATTGAGGGCCATCTCTCCAGTATAATGATAGCCATCCCTGGCTTTGGGAAGGACATCAAGGACCCAAC

ATCCGAAGTCGAGTTGAACCCAGATCTAAGACCTATAATAAGCCGTGACTCTGGCAGAGCTCTCGCGGAGGTCCTCAAGA

AACCCGCTGTGGATAGGTCTCAGAAAATTGGAACCAAAGCCAACTCCAGCTCAAAGGGTCAGCTTCTTAAGGATCTCCAG

CTAAAACCTGTCGACAAGCAGGCAAGCTCTGCAATCGGGTTTGTCCCGTCCGACCATGAATCATCCAGAAATGTCATCCG

CTCCATAATCAAGTCGAGCAAGCTAAATATTGATCACAAGGACTATCTTCTAGATTTACTAAATGATGTGAAAGGCTCCA

AGGATCTTAAGGAATTTCACAAGATGCTAACAGCAATTCTCGCCAAGCACCTGTAACACATCCCCCAGTCATCATCTCAT

ACTCGACTAAAAACATCCTTTCAATCAGGCTATTACAAAAAACTTAGGAGCAAGGGCAACTGAGCTTCGCAGACAGGATG

ACCGAGATCTACGACTTCGATAAATCAGCATGGGATGTCAAAGGGTCAATTGCCCGCATAGAACCCACCACCTACCACGA

CGGCCGACTGATACCCCAGGTGAGGGTCATCGATCCTGGTCTGGGAGACAGAAAAGATGAGTGCTTTATGTACCTGTTTC

TCCTAGGAGTGATTGAGGATAACGACCCCCTGTCTCCCCCCGTCGGAAGAACCTTTGGCTCTCTACCTCTAGGGGTTGGT

AGGTCAACTGCCAAGCCAGAAGAACTACTAAGGGAGGCCACAGAATTAGATATAGTGGTGAGGCGCACTGCAGGAGTAAA

TGAGAAACTGGTATTTTACAACAACACTCCGCTGTCCTTGTTAACGCCATGGAAGAAAGTTCTGACAACCGGAAGTGTGT

TTAGCGCTAACCAGGTTTGCAATGCAGTCAACCTAGTCCCGCTTGATACTCCCCAGAGATTCAGGGTTGTGTACATGAGC

ATAACTAGATTATCAGACAATGGTTACTATAGTGTGCCCAGAAGAATGCTGGAGTTCCGCTCGGCCAATGCAGTAGCCTT

CAACATCTTGGTTACACTAAGAATTGAAAATGGCACAAACCCTAGAAGATACATAGTCGGCTCATGGGAGAATTCAGAGG

TCACATTTATGGTACACGTGGGCAACTTTAGGAGAAAGAAGAACGAAGTATACTCTGCTGATTACTGCAAAATGAAGATT

GAAAAGATGGGTTTAGTTTTTGCCCTGGGTGGAATAGGCGGAACAAGTCTCCATATTAGAAGCACCGGGAAAATGAGCAA

AACCCTCCATGCACAGCTGGGGTTCAAGAAAATCTTATGTTACCCCCTAATGGATGTCAATGAGGATCTTAACCGATATC

TCTGGCGGGCAGAGTGCCGAATAGTCAAAATCCAAGCTGTCTTACAGCCATCAGTACCCCAAGAGTTCCGTGTCTACGAT

GATGTCATCATCAACGATGATCAAGGCTTGTTCAAGATCTTGTAGTTCATTTGCAAAGGAACGAACCCACCCCCAGGAAG

GGGCCCCGCCCTCCCCCG-CCAAAAAACACACTCCAAGGCTCCCCGGCACAGCGCCAACACAGCTACTCCCGCGATGGAA

CGAGGTCGGCC-CCCGAACAAATCCGGTCCAAAGAGGGGGGCCCCCCCCCTGACCAACCGGGCGCCGACCGAGCAGGCCG

GCGAGAAGCCCCCAAGAGGCTCCGACGAAGACCGGGCACCCCATCCCCCC-AAAAACGTCCCCC-AACACGCCCGAGGCT

AGCCACCGGACACCCCCAGACCCGCCACCGGCGAGACAGAACCCCAAGGGGGGACCCTCCAAACCAAAGGAACCACCGGG

GTAGGCTCCGCCGAACCAGCACAGACCCGACCCAAGCAAGGCGGACCCCCAACACGAAACTGCCCAGCACACCCTACTCA

TCATCAAACAAAACTTAGGGGCCAAGTCCACACGAGAGGCCCCAGTCCGTCAGGCGGAGGGACGCAGAAAGGAAGGAGAC

ACGACCGTCCCAGAAGGAGGACACCCAACCACCGAAACGCCCGCCCCCAGGGAAGGAG--CCA-CCCCCACGCAGCGAAG

GAGCGGGCGGGAGACCCCGGGGAGGGCCCGATCGGGACCCGGGGGCCGGCGGCGCCGGCGGTGAGCAGGCCGGGCGCACA

CCCGACCCCACCGGAGGATCCGAGGAGGGCCCGACCCCGCGCGGACCAAACGGCAGGGAGCCCAACCCCACCAAAAAGAG

AGGCGAGGCCCCCCCTCCCTCCCCCCCCGCACACAACCCGCAGACAGACCGAGGGAGGGCCATCCCCCAGCGAGACAGGC

GGCTCCCTCGCAGCCGAGCCGCAGCTCCCCAGAATAGGACNNNNNNNNNNNNNNNNNNNNNNCCCCCGCCCCCCGCCACC

GGGACAAGGGCTCCACCACCACAGACCGGAGCCACCGACAGCCAGCCAGAGGTCCCCTCCAGTCCNNNNNNNNNNNCCCC

CGAAGAGGCCCCGGCAAGGCCCCCCAGCCCGACGTCCCCCGCACGCACCGAAAGAAGGACCCGCCCGAGAACGTGCGGCA

CATCCATGCGCAAACATCATGACGCGGGTCGCAATTTTGACATTTCTGTTTCTTTTCCCAAATGTTGTTGCGTGTCAGAT

TCACTGGGGCAATCTATCCAAGATCGGGATTGTAGGAACAGGGAGTGCCAGCTACAAGGTGATGACTAGGCCAAGCCACC

AGACTCTGGTTATAAAGTTAATGCCAAATATAACGGCCATCGACAATTGTACAAAGTCAGAGATTGCAGAGTACAAGAGA

TTGCTGATCACAGTGTTAAAGCCTGTAGAGGATGCTCTGTCGGTGATAACCAAGAATGTAAGACCAATTCAAACTCTAAC

ACCTGGGCGTAGAACCCGCCGTTTTGCTGGAGCTGTTCTGGCCGGGGTAGCACTTGGAGTTGCGACAGCCGCTCAGATAA

CTGCAGGAGTCGCCCTTCATCAATCATTGATGAACTCCCAAGCAATTGAGAGTTTAAAAACCAGTCTTGAGATGTCGAAT

CAGGCAATAGAAGAAATCAGACTTGCAAATAAGGAGACCATACTGGCAGTACAGGGCGTCCAGGATTATATCAACAATGA

GCTCGTCCCTTCTGTTCATAGAATGTCATGCGAGCTGGTAGGTCACAAGCTCGGCCTCAAGCTCCTTAGGTACTACACCG

AGATCCTGTCCATATTCGGGCCCAGTCTTCGAGACCCGATATCTGCCGAAATATCAATCCAGGCACTTAGTTATGCATTA

GGCGGAGACATTAATAAAATCCTGGACAAGCTTGGGTATAGCGGTGGGGATTTCCTTGCCATCCTAGAAAGCAAGGGAAT

AAAGGCCCGGGTTACATATGTGGACACAAGAGATTACTTTATAATCCTTAGCATCGCCTACCCAACCTTATCTGAGATCA

AGGGAGTGATAGTTCACAAGATAGAAGCTATAACATACAACATTGGGGCACAGGAGTGGTATACTACTATCCCTAAATAT

GTAGCCACTCAGGGGTATCTGATATCGAACTTTGATGAGACGTCATGCGTATTCACTCCAGAGGGGACAGTTTGCAGCCA

GAATGCGTTGTACCCAATGAGCCCATTGCTTCAGGAATGTTTCAGGGGGTCAACAAAATCGTGCGCCAGAACCTTAGTTT

CAGGGACCATAAGTAATAGATTTATCCTATCAAAAGGGAACCTGATTGCAAATTGTGCGTCAGTTTTGTGCAAATGTTAC

ACAACAGAGACAGTTATCAGCCAAGATCCTGACAAACTACTAACTGTTGTAGCATCCGACAAGTGTCCTGTAGTTGAGGT

GGATGGAGTGACAATACAGGTCGGCAGTCGAGAGTATCCGGATTCTGTATACTTACACAAAATAGACTTAGGTCCAGCCA

TCTCCCTAGAAAAACTGGATGTAGGCACCAATTTAGGCAATGCAGTCACAAGACTGGAGAATGCAAAGGAGCTCCTAGAT

GCATCAGACCAAATACTGAAGACTGTTAAAGGGGTACCTTTCAGTGGGAATATGTACATAGCACTGGCAGCTTGCATAGG

AGTATCCCTGGGCCTTGTCACATTAATATGCTGCTGTAAGGGGAGGTGTAATAACAAGGAAATCCCTATCTCCAAAATCA

ACCCAGGGCTCAAACCCGACCTGACCGGGACCTCAAAGTCGTACGTAAGATCACTGTAGTCAGAATCACCTGGATCATCT

GGCATCACACACATACATGCACGACACAGGCAGTCCGATGACGCAAGAAACCCAACCTCCGGTCACCCACCCGACCCCAC

TCCACGCTCCACCACACATTAGTCATCAAACAAAACTTAGGACGAAAGGTCAATCACCATGTCCGCACAAAGGGAGAGGA

TCAATGCCTTCTACAAAGACAATCCTCACAATAAGAACCATAGGGTGATCCTGGATAGAGAACGCTTGGTCATTGAAAGA

CCCTACATCTTGCTTGGAGTCCTGCTGGTAATGTTCCTGAGTCTAATCGGACTGCTGGCCATTGCAGGGATCAGGCTTCA

CCGGGCCACCGTTGGAACTTCAGAGATCCAGAGTCGGCTGAATACCAATATTAAGTTGACCGAATCTATTGATCACCAGA

CTAAGGATGTCTTAACTCCCCTTTTTAAAATCATTGGCGATGAAGTCGGCATCAGAATTCCACAGAAATTCAGTGATCTT

GTCAAGTTCATCTCCGATAAGATTAAATTCCTCAACCCTGATAGAGAGTATGATTTCAGAGATCTCCGGTGGTGTATGAA

TCCTCCCGAGAGAGTCAAAATTAATTTTGATCAGTTTTGTGAGTACAGGGCTGCGGTTAAGTCAATTGAACATATATTTG

AGTCACCACTCAACAAGTCAAAAAAGCTGCAATCTTTGACTCTCGGGCCCGGAACAGGCTGTCTAGGCAGGACAGTAACA

AGAGCCCATTTCTCAGAACTTACAATGACCTTAATGGACCTGGATCTAGAGATGAAGCACAACGTGTCCTCAGTGTTTAC

CGTAGTTGAAGAGGGATTATTCGGAAGAACATATACCGTCTGGAGATCCGATGCCAGGGATCCGAGCGCCGATCTAGGTA

TTGGCCATTTTTTAAGAGTCTTCGAGATTGGACTGGTAAGAGATCTTGGGCTGGGTCCCCCTGTTTTTCATATGACCAAC

TATCTCACGGTGAACATGAGTGATGACTATCGGAGATGTCTTTTAGCGGTAGGGGAGTTGAAGTTGACAGCCCTATGCAC

CTCATCTGAGACTGTGACACTGAGTGAGAGAGGAGTTCCAAGGAGGGAACCTCTTGTGGTTGTGATACTTAATCTAGCTG

GACCCACTCTAGGGGGAGAGCTATACAGTGTCTTGCCTACCTCTGATCTCATGGTGGAGAAACTCTATTTGTCTTCACAT

AGAGGGATCATCAAAGATGATGAGGCCAATTGGGTAGTGCCGTCTACCGATGTTCGTGATCTTCAGAACAAAGGTGAATG

TCTGGTGGAAGCATGCAAGACTCGACCTCCTTCATTTTGCAATGGCACAGGATCAGGCCCGTGGTCAGAAGGGAGAATCC

CTGCCTACGGGGTGATCAGGGTCAGTCTTGACTTAGCGAGTGACCCGGATGTAGTTATCACTTCAGTGTTTGGCCCACTG

ATACCTCACCTATCCGGCATGGATCTTTACAACAACCCGTTTTCAAAAGCTGTATGGTTGGCTGTACCACCTTATGAGCA

GTCATTTCTAGGAATGATAAATACAATTGGATTCCCTAACAGAGCAGAGGTTATGCCGCACATTTTGACCACAGAGATCA

GAGGCCCTCGGGGTCGTTGCCATGTTCCCATAGAATTGTCCCGCCGGGTTGATGACGATATCAAGATCGGGTCCAACATG

GTCATATTGCCGACGATGGACCTGAGGTATATTACAGCCACTTATGATGTTTCCAGGAGCGAGCATGCAATCGTGTACTA

TATCTATGACACGAGTCGCTCATCATCTTACTTCTACCCAGTTCGACTGAATTTCAAAGGCAATCCTCTCTCTTTGAGGA

TAGAGTGTTTCCCCTGGCGTCATAAGGTGTGGTGCTACCATGATTGTCTTATATACAACACCATAACAGGTGAAGAGGTC

CATACGAGAGGGCTGACCGGCATAGAGGTAACATGCAATCCAGTCTGAGCAGAGCTACGACCGTCACTCAGGCAGTCCCT

TGAGTCGCCACCGAGTCCAAGCAGCACAGCCTGGGACAATCAACGGCACAGCCCAGCCAACAATGTTATAAAAAACTTAG

GAGCCAGGGTTGTAGGGGCCATGGACTCTCTATCAGTCAATCAGGTCTTGTACCCTGAGGTTCATCTAGACAGCCCTATT

GTCACAAACAAACTAGTTGCCATCCTTGAGTACTCGGGTATCGACCATAACTATGTTCTTGAAGACCAGACTCTTGTCAA

GAATATTAGGTATAGGCTGGGGTGCGGTTTTTCAAATCAAATGATCATCAATAACAGGGGGGTGGGTGAAACAGTCAATT

CTAAACTTAAAAGTTACCCCCATAATCGTCATATCATATGCCCGGATTGCAATAAGGAGTTGTTTTGTATCAAGGATAGC

TGCATATCTAAGAAGCTCTCGGAGCTATTCAAGAAGGGTAATTCCTTGTACTCTAAGATAAGTCACCAGGTACTGGATTG

TCTTAAGAGAGTCAATGGGAAATTAGGTCTGGGCACAGATCTTACCCATGGTCTGAAGGAGGGTATCCTTGACCTAGGGT

TGCACATGCATAGCTCTCAATGGTTCGAGACTTTCCTGTTCTGGTTCACTATCAAGACAGAGATGAGATCAATGATCAAA

GAACAGTCCCATATATGCCACAAGAGGAGGTATAACCCGACTTTTGTGTCGGGGGATGCATTCGAGGTGCTCGTATCGCG

AGACCTCGTTGTGATAATTGATAAGAATACCCAGTATGTCTTCTACTTAACATTTGAGCTGGTCCTTATGTACTGTGATG

TCATAGAGGGTAGACTTATGACGGAGACAGCCATGGCCATAGACCAGAGATATTCAGAACTTCTAAGCCGGGTCAGATAC

TTGTGGGATCTTATTGATGGGTTCTTCCCAACACTAGGCAACACCACATACCAAATTGTTGCTCTGCTTGAACCATTATC

GTTGGCTTATCTTCAACTTCAGGATGTCACTCTGGAGTTAAGAGGTGCTTTTTTAGACCATTGCTTCAAAGAACTCTATG

AGATACTGGAACATTGCGGCATTGACACAGCAGGCACTTACAATTCCATCACTGAAGGATTGGATTACGTATTCATCACC

CACGATATACATCTAACTGGGGAGATTTTTTCATTTTTCCGGAGTTTCGGACACCCCCGTCTAGAAGCAGTCACCGCTGC

GGAAAATGTCAGAAAACATATGAACCAACCGAAGGTAATCAGTTATGAGACTATGATGAAGGGCCATGCGGTATTTTGCG

GGATAATCATAAATGGTTTCCGGGACCGACATGGCGGCAGCTGGCCCCCTGTTGCACTGCCAGAACATGCCTCTGCTGCG

ATCCGGAATGCGCAGGCATCAGGTGAGGGACTGACCCATGACCTGTGTATAGACAACTGGAAATCCTTTGTTGGATTCAA

ATTTGGCTGCTTTATGCCACTCAGCCTAGATAGTGATTTGACCATGTATCTCAAAGACAAGGCATTGGCTGCACTGAAGA

ATGAGTGGGATTCAGTTTACCCGAAAGAATACCTCCGTTATAACCCACCTAGAGGGACAGAGTCAAGGCGATTGGTAGAG

GTATTCCTGAATGACTCCAGCTTTGATCCTTATAACATGATAATGTACGTGGTGAATGGCTCCTACCTTAAAGACCCTGA

GTTTAACCTCTCATACAGCCTAAAAGAGAAGGAGATAAAGGAGACAGGGCGGTTGTTTGCCAAAATGACCTATAAGATGC

GGGCCTGTCAGGTAATCGCTGAAAATCTGATATCGAATGGTGTTGGGAAGTATTTCCGAGACAATGGGATGGCAAAGGAC

GAGCATGACCTAACAAAAGCCCTTCACACCCTGGCAGTCTCAGGTGTTCCCAAAAACAACAAAGATAACCACCGAGGTGG

GCCTCCCAGAAGGACAACAAACCGAGGGGTGAGATCAAGCCAAGGCACCAAAACACAAGATAGAGACAAGGTTCAAGGGG

GACCTATGTACAACTATTTGCGATGCCAACCGATCAGCCCTGATCAGGGTGAGTCATACGAGACTGTTAGTGCATTCATC

ACCGCTGACCTTAAGAAGTATTGCCTGAATTGGAGATACGAGACAATCAGCATATTTGCACAGAGGCTGAATGAAATATA

TGGACTACCATCCTTCTTTCAATGGTTACACAGGATATTGGAAAAATCCGTACTCTACGTCAGTGACCCACATTGTCCTC

CCGATCTAGATAATCATATCCCTCTGGACAGTGTCCCCAATGCCCAAATATTCATCAAGTACCCAATGGGCGGAATAGAA

GGTTATTGCCAAAAACTATGGACAATCAGCACTATACCATACTTGTATCTGGCAGCCTATGAGAGCGGAGTAAGAATCGC

CTCACTGGTGCAGGGTGACAATCAGACAATCGCAGTGACAAAAAGAGTTCCAAGTTCTTGGCCTTATTCACTAAAAAAGA

GGGAGGCCTCCAAGGCCGCTCAAAATTACTTCGTAGTCTTAAGGCAAAGATTGCACGATGTAGGTCATCACTTAAAGGCT

AATGAGACCATAGTATCTTCTCACTTTTTTGTATATTCCAAAGGGATTTATTATGACGGCCTGCTAGTCTCACAATCACT

AAAGAGCATCGCCAGATGTGTCTTCTGGTCCGAGACTATAGTGGATGAAACCAGAGCGGCGTGCAGTAATATTGCAACAA

CTGTCGCTAAGAGTATAGAGAGGGGTTATGATAGGTACCTTGCATATTCTTTGAATATCCTCAAGATTTTCCAACAGATC

CTTATATCCCTTAACTTCACTATTAACACAACAATGACTCAGGATGTCGTGGCACCGATCATCGAGAACGGTGATTTGCT

GATAAGGATGGCACTCTTGCCAGCACCCATCGGGGGTCTCAATTATCTTAACATGAGCAGGTTATTTGTGAGAAATATCG

GTGACCCGGTCACCTCCTCTATAGCCGACCTGAAGAGGATGATAGATGCCGGGCTAATGCCAGAAGAAACATTGCATCAA

GTGATGACCCAGACCCCGGGAGAATCATCTTACCTAGATTGGGCAAGTGACCCTTATTCTGCCAACCTACCCTGCGTACA

GAGTATCACTCGCCTTCTCAAAAACATCACTGCACGATATATTTTAATCAGCAGCCCAAACCCGATGCTGAAAGGGTTAT

TTCATGAGGGGAGTAGAGATGAAGACGAGGAGCTTGCAAGTTTCCTAATGGATCGGCATATAATTGTTCCAAGAGCTGCA

CATGAAATCTTAGACCATAGTATAACCGGAGCAAGAGAAGCTATAGCCGGGATGTTGGACACCACCAAGGGTCTGATTAG

AACAAGTATGAAACGGGGCGGCCTCACCCCTCGAGTCTTAGCCCGCCTTTCCAATTATGATTATGAACAATTCCGATCCG

GGATAACACTATTAACAAAGAAAGGGCAGTGTTATCTCATTGACAAGGACTCGTGCTCGGTGCAGCTCGCTATAGCTCTG

AGGGGCCACATGTGGGCTAAGTTGGCTCGCGGGAGACCAATCTATGGGTTGGAGGTGCCTGATGTATTAGAATCGATGAA

CGGCTACCTTATTAAACGCCACGAGTCGTGTGCTATCTGTGAAACGGGCTCAAGTCACTACGGATGGTTTTTCGTCCCTG

CAGGGTGCCAGCTTGACGATGTATCTAGAGAGACTTCGGCTCTTCGTGTACCTTATGTCGGATCAACAACTGAGGAAAGG

ACAGACATGAAACTTGCATTTGTTAGGTCTCCAAGTAGATCCCTCAAATCAGCAGTTAGAATTGCAACAGTTTACTCATG

GGCTTACGGGGATGATGAGAAATCATGGAGTGAAGCCTGGATGCTAGCTAGGCAGAGAGCCAATATCACCTTAGATGAAT

TGAGAATGATCACTCCAGTCTCCACATCCACCAACTTAGCCCACCGGTTGAGGGATCGGAGTACCCAGGTGAAGTACTCG

GGGACATCCCTTGTGAGGGTTGCAAGATACACAACCATCTCCAATGATAACTTGTCGTTTGTGATATCTGAGAAAAAAGT

GGATACCAACTTCATTTATCAGCAAGGGATGTTGCTTGGTCTTGGGATCCTTGAGAACCTCTTTAGGCTAGAGGCCACCA

CCGGAGTATCCAACACAGTGCTACACCTGCACGTGGAAACAGAATGTTGTGTTGTACCTATGGTGGATCACCCAAGGATA

CCGAGTCTCCGTAATATAAAAGTTACGGATGAGCTATGCACAAACCCTTTGATCTACGACAGGTCCCCCATCATAGAGCA

CGATGCAACCCGGCTATATTCACAGAGCCACAGGAGACATTTGGTGGAGTTTGTTACCTGGTCAACAAGCCAGCTTTACC

ATATACTGGCTAAGTCTACAGCAATGTCCATGATTGAATTGATTACGAGATTCGAGAAAGATCACATGAATGAAATAGCC

GCCCTGATTGGCGATGACGACATCAACAGCTTCATCACAGAATTTTTGCTAGTGGAGCCTAGATTATTTATAGTTTACCT

TGGTCAGTGTGCTGCCATCAATTGGGCCTTTGATATACATTATCATCGGCCCTCGGGCAAGTACCAGATGGGGGAACTTC

TCTACTCTCTGCTCTCTCGGATGAGCAAAGGAGTATATAAGATCTTTACTAATGCTTTGAGCCACCCCAAAGTTTACAAG

AAATTTTGGCGAAGTGGCATCATTGAGCCGATTCATGGACCATCCTTGGATACACAGAATTTACATGTCACTGTCTGTGA

CATGATATATGGGTCCTATGTCACCTATTTGGATCTTTTGCTGAATGATGAGCTAGATGATTACCCGTATTTGCTCTGCG

AGAGTGATGAGGACGTGGTCACAGACAGGTTCGACAACATTCAAGCCAAACATCTCTGTGTATTGGCCGATGTATATTGC

AGCTCCAAGAGGTGTCCTTCAATAATCGGGATGTCTCCTATAGAAAAATGTACCATTCTCACACATTACATCAAGGGAGA

ATCAGTACAATCCCCGTCTGGGACCTCATGGAACACTGATCCCCTTGTAGTAGATCATTACTCATGCTCTCTGACCTACC

TTCGCCGCGGTTCCATCAAACAAATCAGGTTGAGGGTGGATCCTGGGTTTGTATTCGAGGCGTTGACAGACGTCGACTTT

AAACAACCTCGCAAAGCTAAGTTAGATATATCGGTCGTTGGGTTGACTGATTTCTCTCCCCCTTACGACAACGTCGGTGA

TTTTTTAGGAACTATCAACACATTGAGGCACGATCTGCCTGTCACCGGAACCGGGGTCTCGAACTATGAAGTCCACGCTT

ATCGTAGAATTGGCCTGAATTCATCAGCATGTTATAAAGCCGTAGAGATCTCCACGTTAATCAAGCCATCCTTAGAAGTC

GGAGAGCATGGATTGTTCTTAGGAGAAGGTTCCGGTTCAATGCTGGCTGCGTACAAAGAAGTTCTTAAATTAGCAAATTG

TTACTACAACAGCGGAGTAACAGCGGAGGGCAGAGCCGGACAGAGGGAAATATCTCCCTATCCTTCAGAGATGAGCCTAG

TAGAGAATCAAATGGGGATAGAGCGGAGTGTTAAAGTGCTGTTCAACGGGAAACCTGAAGTAACTTGGGTGGGGACCACA

GATTGCTACAAGTACATAATCAGTAACATCCAAACCTCTAGTCTGGGTTTCATACATTCAGATATTGAGACGCTCCCAAC

CAAAGATGCTGTTGAGAAATTGGAGGAATTTGCCTCTATCCTATCCTTATCCTTGATTTTGGGAAAAATCGGCTCTATTA

CAGTTGTCAAAATTATGCCCATTAGCGGGGATTTTACCCAAGGCTTCATAGCCTATGCCATCCAATATTTTAGGGAGAGC

CTGCTTGCCTATCCGAGATATAGTAACTTCATCTCGACTGAGTGTTACCTTATTATGATAGGATTAAAGGCCAATCGATT

AATAAACCCAGAAGCCATTAAGCAAAGCATAATCAGAGCGGGGGTCAGGACTGCACCAGGACTTGTGAGCCATGTATTAT

CAGAGAAACAAAAAGGTTGTATTCAATCTTTCTTGGGTGATCCTTATATCCAAGGAGACTTCAATAAGCACCTTAAATCT

CTAACCTCTATTGAGAAAATCCTGGTAAATTGTGGTCTCTCGATCAATGGCATAAAAATCTGTAGAGATCTAATCCACCA

TGATATCGCCTCCGGTCCAGACGGCCTTATGAGCTCTACAATTATTTTATATAGGGAATTGGCTCATTTCAAAGATAATA

TAAGGAGTCAGCACGGAATGTTCCACCCCTACCCGGTACTGGCCAACAGCAGGCAACGTGAATTAATCCTTCGAATAGCC

AAGAAATTCTGGGGGTATGTCTTGCTATATTCTGATGACCCGGCACTAATCAGACAAACAATCAAGAACTTGAAACGGAA

TCATCTAACCTTTGACTTACATAGTAACCCATTTATTAAAGGGCTATCCAAGGCTGAGAAACTGCTGGTGCGGACGAGTT

CACTAAGAAGAGAATGGTTGTTCACTCTCGAAACGAAAGAAGTAAAAGAGTGGTTCAAATTGGTAGGTTACAGTGCACTC

ATCAGAGGCTAGTCAGCTATGCATCTGCCCCCTTCTCCTCCGCCATGAGACCCCACTGGCGATCCAGAAGATTAAAGAAA

ACTACATATTGGATAAGTATCTATTCCCAGCTTTGTCTGGN

>SiberianIbex/SiberianIbex/Mongolia/2017-01

NNNNNNNNNNNNNNNNNNNNNATAGATCTTATAATAACTATGGACTGGCAAACTTAGGAGTAAAGATCCTACTGTCGGGG

GGAGGAGGAGGAGCAAGGTCCTTGATCATGGCGACTCTCCTTAAAAGCTTAGCATTGTTCAAAAGGAACAAAGACAAAGC

GCCGACAGCATCAGGTTCAGGAGGGGCCATCCGGGGGATTAAGAATGTTATCATAGTCCCGATTCCCGGAGACTCGTCCA

TCACTACCCGTTCAAGACTGCTCGACAGGCTTGTCAGATTGGCCGGAGATCCTGACATCAACGGGTCAAAGCTGACCGGG

GTGATGATCAGCATGTTATCCTTGTTCGTAGAGTCACCCGGGCAATTGATACAGCGAATCACAGATGATCCGGATGTTAG

TATCCGCCTTGTTGAGGTAGTTCAAAGTACTAGGTCCCAGTCCGGGTTGACCTTTGCATCACGTGGTGCTGATTTAGACA

ACGAGGCAGACATGTATTTTTCAACTGAGGGGCCCTCGAGTGGAGGTAAGAAAAGGATCAACTGGTTTGAGAACAGAGAA

ATAATAGACATAGAGGTGCAGGATCCAGAAGAGTTCAATATGTTGTTAGCCTCCATACTAGCACAAGTCTGGATCCTTCT

GGCCAAGGCTGTTACGGCACCAGATACGGCAGCTGACTCAGAACTGAGAAGGTGGGTTAAATACACACAACAAAGGAGAG

TGATTGGGGAATTTCGCCTTGACAAGGGGTGGCTGGACGCAGTCCGCAACAGAATTGCAGAAGATCTATCACTCCGGCGG

TTCATGGTATCTCTCATACTTGACATCAAGAGGACCCCAGGCAACAAGCCAAGGATTGCAGAAATGATCTGCGACATTGA

CAACTATATTGTCGAAGCAGGACTCGCCAGTTTCATCCTTACTATCAAATTTGGTATCGAAACCATGTATCCTGCATTAG

GGCTCCACGAGTTTGCCGGAGAATTGTCCACTATTGAGTCCTTGATGAACTTGTATCAACAGCTAGGAGAGGTTGCACCC

TACATGGTAATTCTAGAGAATTCAATTCAGAACAAGTTTAGTGCAGGAGCTTATCCCCTCCTCTGGAGCTATGCGATGGG

TGTCGGAGTCGAGCTGGAGAACTCAATGGGGGGCTTGAACTTTGGCAGGTCATATTTTGACCCTGCCTATTTTCGTCTCG

GACAGGAGATGGTCAGAAGATCTGCAGGAAAGGTCAGCTCTGTGATTGCGGCTGAGCTCGGCATCACAGCAGAGGAAGCT

AAACTAGTCTCGGAAATCGCCTCGCAGGCCGGGGACGAAAGAACCGCCAGAGGGACTGGGCCTCGACAGGCGCAGGTTTC

CTTCCTTCAGCACCAAACAGGAGGGGGAGAGTCGTCCGCACCAGCGACCAGAGAAGGGGTCAAAGCTGCGATCCCAAACG

GAGCTGAAGAAAGGGACAGAAAACAAACACGCCCAGGAAGGCCCAGAGGAGAGACCTCCGGCCAACTGCTCCTGGACATC

ATGCCAGAGGATGAGATCTTGCGAGAGTCTGGTCAAAACCCTCGTGAGGCTCAAAGATCGGCCGAGGCACTCTTCAGGCT

GCAGGCCATGGCCAAGATTCTGGAGGACCAGGAGGAGGGAGAAGACAACAGTCAGGTCTACAACGACAAAGATCTCCTCG

GCTAAGCAGACGCACCCTCTGTCGAAATCAGTGACAAGACATCTCCTACCAGTATTATAAAAAACTTAGGACCCAGGTCC

AAGCAAGCACACATCGACACTCCAACCAGTCGAGCGGAGACCACCGATGGCAGAAGAACAAGCATACCATGTNAACAAGG

GGCTGGAATGTATCAAGTCTCTCAAAGCCTCACCCCCGGATCTATCCACCATCAGAGATACCATCGAGAGCTGGAGAGAG

GGGCTWAGCCCCTCGGGCCGTGCAACACCGAACCCTGATACGTCCGAGGGAGATCATCAGAATATCAACCAATCATGCTC

ACCAGCAATCGGACCAAACAAAGTCTACTTGTCTCCTGAAGATAATCTCGGATTTAGAGAGATCACTGGCAACGACTGTG

AGGCTGGGCTCGGAGGGGTCCAGAGAGAAGGATCCAACTCTCAAGTACAGCGTTACCATGTTTATAGTCACGGGGGTGAA

GAGATTGAAGGACTCGAGGATGCTGACTCTCTCGTGGTTCAAGCAGATCCTCCAGCTGCTAACGTCTTCAATGGAGGAGA

GGATGGATCTGACGACAGCGATGTGGACTCTGGCCCAGATGATCCCGGCAGAGATACTCTATATGACCGGGGATCTGTTG

CCGGCAATGGTATCGCTAGGTCCACAGATGTCGAAAAACTAGAAGGTGCTGATATTCAAGAAGTTCTTAACTCCCAGAAA

GGCAAAGGAGGAAGATTCCAGGGCGGGAAAACCTTGCGAGTCCCGGAAATACCCGATGTCAAGCACTCCAGACCATCAGC

CCAATCAATTAAAAAGGGCACAGACGGGAACTCAGTCTCATCTGGAACGGTGATAGAGTGTTTATCGATAAGTGGTGCAA

CCCAAACTGTGCCAGAGTCAAGATGGGAGTCATCAGAGCAAAATGCGTCTGCGGGGAGTGTCCTCAAGTCTGCGAGGAGT

GCAAAGACGATCCAGGGGCCGACACAAGAATCTGGTACCATAGCATCACTGACTCAGCCTAAAGAGAATGACTCCGAGTA

TGAGTATGAGGATGACCTATTTACAGAGATTCAGGACATCCGTGCAAGCATTGCCAAGATCCATGATGACAATAAAACTA

TCCTCTCAAAGCTTGATTCTATACTGTTATTGAAAGGGGAAGTCGACACTATCAAGAAACAAATCAGCAAGCAGAATATA

AGTATATCCACCATTGAGGGCCATCTCTCCAGTATAATGATAGCCATCCCTGGCTTTGGGAAGGACATCAAGGACCCAAC

ATCCGAAGTCGAGTTGAACCCAGATCTAAGACCTATAATAAGCCGTGACTCTGGCAGAGCTCTCGCGGAGGTCCTCAAGA

AACCCGCTGTGGATAGGTCTCAGAAAATTGGAACCAAAGCCAACTCCAGCTCAAAGGGTCAGCTTCTTAAGGATCTCCAG

CTAAAACCTGTCGACAAGCAGGCAAGCTCTGCAATCGGGTTTGTCCCGTCCGACCATGAATCATCCAGAAATGTCATCCG

CTCCATAATCAAGTCGAGCAAGCTAAATATTGATCACAAGGACTATCTTCTAGATTTACTAAATGATGTGAAAGGCTCCA

AGGATCTTAAGGAATTTCACAAGATGCTAACAGCAATTCTCGCCAAGCACCTGTAACACATCCCCCAGTCATCATCTCAT

ACTCGACTAAAAACATCCTTTCAATCAGGCTATTACAAAAAACTTAGGAGCAAGGGCAACTGAGCTTCGCAGACAGGATG

ACCGAGATCTACGACTTCGATAAATCAGCATGGGATGTCAAAGGGTCAATTGCCCGCATAGAACCCACCACCTACCACGA

CGGCCGACTGATACCCCAGGTGAGGGTCATCGATCCTGGTCTGGGAGACAGAAAAGATGAGTGCTTTATGTACCTGTTTC

TCCTAGGAGTGATTGAGGATAACGACCCCCTGTCTCCCCCCGTCGGAAGAACCTTTGGCTCTCTACCTCTAGGGGTTGGT

AGGTCAACTGCCAAGCCAGAAGAACTACTAAGGGAGGCCACAGAATTAGATATAGTGGTGAGGCGCACTGCAGGAGTAAA

TGAGAAACTGGTATTTTACAACAACACTCCGCTGTCCTTGTTAACGCCATGGAAGAAAGTTCTGACAACCGGAAGTGTGT

TTAGCGCTAACCAGGTTTGCAATGCAGTCAACCTAGTCCCGCTTGATACTCCCCAGAGATTCAGGGTTGTGTACATGAGC

ATAACTAGATTATCAGACAATGGTTACTATAGTGTGCCCAGAAGAATGCTGGAGTTCCGCTCGGCCAATGCAGTAGCCTT

CAACATCTTGGTTACACTAAGAATTGAAAATGGCACAAACCCTAGAAGATACATAGTCGGCTCATGGGAGAATTCAGAGG

TCACATTTATGGTACACGTGGGCAACTTTAGGAGAAAGAAGAACGAAGTATACTCTGCTGATTACTGCAAAATGAAGATT

GAAAAGATGGGTTTAGTTTTTGCCCTGGGTGGAATAGGCGGAACAAGTCTCCATATTAGAAGCACCGGGAAAATGAGCAA

AACCCTCCATGCACAGCTGGGGTTCAAGAAAATCTTATGTTACCCCCTAATGGATGTCAATGAGGATCTTAACCGATATC

TCTGGCGGGCAGAGTGCCGAATAGTCAAAATCCAAGCTGTCTTACAGCCATCAGTACCCCAAGAGTTCCGTGTCTACGAT

GATGTCATCATCAACGATGATCAAGGCTTGTTCAAGATCTTGTAGTTCATTTGCAAAGGAACGAACCCACCCCCAGGNNN

NNNNNNNNNNNNNNNNNN-NNNNNNNNNNNNNNNNNNNNNNNNNNNNNNNNNNNNNNNNNNNNNNNNNNNNNNNNNNNNN

NNNNNNNNNNN-NNNNNNNNNNNNNNNNNNNNNNNNNNNNNNNNNNNNNNNNNNNNNNNNNNNNNNNNNNNNNNNNNNNN

NNNNNNNNNNNNNNNNNNNNNNNNNNNNNNNNNNNNNNNNNNNNNNNNNN-NNNNNNNNNNNNN-NNNNNNNNNNAGGCT

AGCCACCGGACACCCCCAGACCCGCCACCGGCGAGACAGAACCCCAAGGGGGGACCCTCCAAACCAAAGGAACCACCGGG

GTAGGCTCTGCCGAACCAGCACAGACCCGACCCAAGCAAGGCGGACCCCCAACACGAAACTGCCCAGCACACCCTACTCA

TCATCAAACAAAACTTAGGGGCCAAGTCCACACGAGAGGCCCCAGTCCGTCAGGCAGAGGGACGCAGAAAGGAAGGAGAC

ACGACCGTCCCAGAAGGAGGANNNNNNNNNNNNNNNNNNNNNNNNNNNNNNNNNNNNN--NNN-NNNNNNNNNNNNNNNN

NNNNNNNNNNNNNNNNNNNNNNNNNNNNNNNNNNNNNNNNNNNNNNNNNNNNNNNNNNNNNNNNNNNNNNNNNNNNNNNN

NNNNNNNNNNNNNNNNNNNNNNNNNNNNNNNNNNNNNNNNNNNNNNNNNNNNNNNNNNNNCCCAACCCCACCAAAAAGAN

NNNNNNNNNNNNNNNNNNNNNNNNNNNNNNNNNNNNNNNNNNNNNNNNNNNNNNNNNNNNNNNNNNNNNNNNNNNNNNNN

NNNNNNNNNNNNNNNNNNNNNNNNNNNNNNNNNNNNNNNNNNNNNNNNNNNNNNNNNNNNNCCCCCCGCCCCCCGCCACC

NNNNNNNNNNNNNNNNNNNNNNNNNNNNNNNNNNNNNNNNNNNNNNNNNNNNNNNNNNNNNNNNNNNNNNNNNNNNNNNN

NNNNNNNNNNNNNNNNNNNNNNNNNNNNNNNNNNNNNNNNNNNNNNNCCGAAAGAAGGACCCGCCCGAGAACGTGCGGCA

CATCCATGCGCAAACATCATGACGCGGGTCGCAATTTTGACATTTCTGTTTCTTTTCCCAAATGTTGTTGCGTGTCAGAT

TCACTGGGGCAATCTATCCAAGATCGGGATTGTAGGAANNNNNNNNNNNNNNNNNNNNNNNNNNNNNNNNNNNNNNNACC

AGACTCTGGTTATAAAGTTAATGCCAAATATAACGGCCATCGACAATTGTACAAAGTCAGAGATTGCAGAGTACAAGAGA

TTGCTGATCACAGTGTTAAAGCCTGTAGAGGATGCTCTGTCGGTGATAACCAAGAATGTAAGACCAATTCAAACTCTAAC

ACCTGGGCGTAGAACCCGCCGTTTTGCTGGAGCTGTTCTGGCCGGGGTAGCACTTGGAGTTGCGACAGCCGCTCAGATAA

CTGCAGGAGTCGCCCTTCATCAATCATTGATGAACTCCCAAGCAATTGAGAGTTTAAAAACCAGTCTTGAGATGTCGAAT

CAGGCAATAGAAGAAATCAGACTTGCAAATAAGGAGACCATACTGGCAGTACAGGGCGTCCAGGATTATATCAACAATGA

GCTCGTCCCTTCTGTTCATAGAATGTCATGCGAGCTGGTAGGTCACAAGCTCGGCCTCAAGCTCCTTAGGTACTACACCG

AGATCCTGTCCATATTCGGGCCCAGTCTTCGAGACCCGATATCTGCCGAAATATCAATCCAGGCACTTAGTTATGCATTA

GGCGGAGACATTAATAAAATCCTGGACAAGCTTGGGTATAGCGGTGGGGATTTCCTTGCCATCCTAGAAAGCAAGGGAAT

AAAGGCCCGGGTTACATATGTAGACACAAGAGATTACTTTATAATCCTTAGCATCGCCTACCCAACCTTATCTGAGATCA

AGGGAGTGATAGTTCACAAGATAGAAGCTATAACATACAACATTGGGGCACAGGAGTGGTATACTACTATCCCTAAATAT

GTAGCCACTCAGGGGTATCTGATATCGAACTTTGATGAGACGTCATGCGTATTCACTCCAGAGGGGACAGTTTGCAGCCA

GAATGCGTTGTACCCAATGAGCCCATTGCTTCAGGAATGTTTCAGGGGGTCAACAAAATCGTGCGCCAGAACCTTAGTTT

CAGGGACCATAAGTAATAGATTTATCCTATCAAAAGGGAACCTGATTGCAAATTGTGCGTCAGTTTTGTGCAAATGTTAC

ACAACAGRGACAGTTATCAGCCAAGATCCTGACAAACTACTAACTGTTGTAGCATCCGACAAGTGTCCTGTAGTTGAGGT

GGATGGAGTGACAATACAGGTCGGCAGTCGAGAGTATCCGGATTCTGTATACTTACACAAAATAGACTTAGGTCCAGCCA

TCTCCCTAGAAAAACTGGATGTAGGCACCAATTTAGGCAATGCAGTCACAAGACTGGAGAATGCAAAGGAGCTCCTAGAT

GCATCAGACCAAATACTGAAGACTGTTAAAGGGGTACCTTTCAGTGGGAACATGTACATAGCACTGGCAGCTTGCATAGG

AGTATCCCTGGGCCTTGTCACATTAATATGCTGCTGTAAGGGGAGGTGTAAGAACAAGGAAATCCCTATCTCCAAAATCA

ACCCAGGGCTCAAACCCGACCTGACCGGGACCTCAAAGTCGTACGTAAGATCACTGTAGTCAGAATCACCTGGATCATCT

GGCATCACACACATACATGCACGACACAGGCAGTCCGATGACGCAAGAAACCCAACCTCCGGTCACCCACCCGACCCCAC

TCCACGCTCCACCACACATTAGTCATCAAACAAAACTTAGGACGAAAGGTCAATCACCATGTCCGCACAAAGGGAGAGGA

TCAATGCCTTCTACAAAGACAATCCTCACAATAAGAACCATAGGGTGATCCTGGATAGAGAACGCTTGGTCATTGAAAGA

CCCTACATCTTGCTTGGAGTCCTGCTGGTAATGTTCCTGAGTCTAATCGGACTGCTGGCCATTGCAGGGATCAGGCTTCA

CCGGGCCACCGTTGGAACTTCAGAGATCCAGAGTCGGCTGAATACCAATATTAAGTTGACCGAATCTATTGATCACCAGA

CTAAGGATGTCTTAACTCCCCTTTTTAAAATCATTGGCGATGAAGTCGGCATCAGAATTCCACAGAAATTCGGTGATCTT

GTCAAGTTCATCTCCGATAAGATTAAATTCCTCAACCCTGATAGAGAGTATGATTTCAGAGATCTCCGGTGGTGTATGAA

TCCTCCCGAGAGAGTCAAAATTAATTTTGATCAGTTTTGTGAGTACAGGGCTGCTGTTAAGTCAATTGAACATATATTTG

AGTCACCACTCAACAAGTCAAAAAAGCTGCAATCTTTGACTCTCGGGCCCGGAACAGGCTGTCTAGGCAGGACAGTAACA

AGAGCCCATTTCTCAGAACTTACAATGACCTTAATGGACCTGGATCTAGAGATGAAGCACAACGTGTCCTCAGTGTTTAC

CGTAGTTGAAGAGGGATTATTCGGAAGAACATATACCGTCTGGAGATCCGATGCCAGGGATCCGAGCGCCGATCTAGGTA

TTGGCCATTTTTTAAGAGTCTTCGAGATTGGACTGGTAAGAGATCTTGGGCTGGGTCCCCCTGTTTTTCATATGACCAAC

TATCTCACGGTGAACATGAGTGATGACTATCGGAGATGTCTTTTAGCGGTAGGGGAGTTGAAGTTGACAGCCCTATGCAC

CTCATCTGAGACTGTGACACTGAGTGAGAGAGGAGTTCCAAGGAGGGAACCTCTTGTGGTTGTGATACTTAATCTAGCTG

GACCCACTCTAGGGGGAGAGCTATACAGTGTCTTGCCTACCTCTGATCTCATGGTGGAGAAACTCTATTTGTCTTCACAT

AGAGGGATCATCAAAGATGATGAGGCCAATTGGGTAGTGCCGTCTACCGATGTTCGTGATCTTCAGAACAAAGGTGAATG

TCTGGTGGAAGCATGCAAGACTCGACCTCCTTCATTTTGCAATGGCACAGGATCAGGCCCGTGGTCAGAAGGGAGAATCC

CTGCCTACGGGGTGATCAGGGTCAGTCTTGACTTAGCGAGTGACCCGGATGTAGTTATCACTTCAGTGTTTGGCCCACTG

ATACCTCACCTATCCGGCATGGATCTTTACAACAACCCGTTTTCAAAAGCTGTATGGTTGGCTGTACCACCTTATGAGCA

GTCATTTCTAGGAATGATAAATACAATTGGATTCCCTAACAGAGCAGAGGTTATGCCGCACATTTTGACCACAGAGATCA

GAGGCCCTCGGGGTCGTTGCCATGTTCCCATAGAATTGTCCCGCCGGGTTGATGACGATATCAAGATCGGGTCCAACATG

GTCATATTGCCGACGATGGACCTGAGGTATATTACAGCCACTTATGATGTTTCCAGGAGCGAGCATGCAATCGTGTACTA

TATCTATGACACGAGTCGCTCATCATCTTACTTCTACCCAGTTCGACTGAATTTCAAAGGCAATCCTCTCTCTTTGAGGA

TAGAGTGTTTCCCCTGGCGTCATAAGGTGTGGTGCTACCATGATTGTCTTATATACAACACCATAACAGGTGAAGAGGTC

CATACGAGAGGGCTGACCGGCATAGAGGTAACATGCAATCCAGTCTGAGCAGAGCTACGACCGTCACTCAGGCAGTCCCT

TGAGTCGCCACCGAGTCCAAGCAGCACAGCCTGGGACAATCAACGGCACAGCCCAGCCAACAATGTTATAAAAAACTTAG

GAGCCAGGGTTGTAGGGGCCATGGACTCTCTATCAGTCAATCAGGTCTTGTACCCTGAGGTTCATCTAGACAGCCCTATT

GTCACAAACAAACTAGTTGCCATCCTTGAGTACTCGGGTATCGACCATAACTATGTTCTTGAAGACCAGACTCTTGTCAA

GAATATTAGGTATAGGCTGGGGTGCGGTTTTTCAAATCAAATGATCATCAATAACAGGGGGGTGGGTGAAACAGTCAATT

CTAAACTTAAAAGTTACCCCCATAATCGTCATATCATATACCCGGATTGCAATAAGGAGTTGTTTTGTATCAAGGATAGC

TGCATATCTAAGAAGCTCTCGGAGCTATTCAAGAAGGGTAATTCCTTGTACTCTAAGATAAGTCACCAGGTACTGGATTG

TCTTAAGAGAGTCAATGGGAAATTAGGTCTGGGCACAGATCTTACCCATGGTCTGAAGGAGGGTATCCTTGACCTAGGGT

TGCACATGCATAGCTCTCAATGGTTCGAGACTTTCCTGTTCTGGTTCACTATCAAGACAGAGATGAGATCAATGATCAAA

GAACAGTCCCATATATGCCACAAGAGGAGGTATAACCCGACTTTTGTGTCGGGGGATGCATTCGAGGTGCTCGTATCGCG

AGACCTCGTTGTGATAATTGATAAGAATACCCAGTATGTCTTCTACTTAACATTTGAGCTGGTCCTTATGTACTGTGATG

TCATAGAGGGTAGACTTATGACGGAGACAGCCATGGCCATAGACCAGAGATATTCAGAACTTCTAAGCCGGGTCAGATAC

TTGTGGGATCTTATTGATGGGTTCTTCCCAACACTAGGCAACACCACATACCAAATTGTTGCTCTGCTTGAACCATTATC

GTTGGCTTATCTTCAACTTCAGGATGTCACTCTGGAGTTAAGAGGTGCTTTTTTAGACCATTGCTTCAAAGAACTCTATG

AGATACTGGAACATTGCGGCATTGACACAGCAGGCACTTACAATTCCATCACTGAAGGATTGGATTACGTATTCATCACC

CACGATATACATCTAACTGGGGAGATTTTTTCATTTTTCCGGAGTTTCGGACACCCCCGTCTAGAAGCAGTCACCGCTGC

GGAAAATGTCAGAAAACATATGAACCAACCGAAGGTAATCAGTTATGAGACTATGATGAAGGGCCATGCGGTATTTTGCG

GGATAATCATAAATGGTTTCCGGGACCGACATGGCGGCAGCTGGCCCCCTGTTGCACTGCCAGAACATGCCTCTGCTGCG

ATCCGGAATGCGCAGGCATCAGGTGAGGGACTGACCCATGACCTGTGTATAGACAACTGGAAATCCTTTGTTGGATTCAA

GTTTGGCTGCTTTATGCCACTCAGCCTAGATAGTGATTTGACCATGTATCTCAAAGACAAGGCATTGGCTGCACTGAAGA

ATGAGTGGGATTCAGTTTACCCGAAAGAATACCTCCGTTATAACCCACCTAGAGGGACAGAGTCAAGGCGATTGGTAGAG

GTATTCCTGAATGACTCCAGCTTTGATCCTTATAACATGATAATGTACGTGGTGAATGGCTCCTACCTTAAAGACCCTGA

GTTTAACCTCTCATACAGCCTAAAAGAGAAGGAGATAAAGGAGACAGGGCGGTTGTTTGCCAAAATGACCTATAAGATGC

GGGCCTGTCAGGTAATCGCTGAAAATCTGATATCGAATGGTGTTGGGAAGTATTTCCGAGACAATGGGATGGCAAAGGAC

GAGCATGACCTAACAAAAGCCCTTCACACCCTGGCAGTCTCAGGTGTTCCCAAAAACAACAAAGATAACCACCGAGGTGG

GCCTCCCAGAAGGACAACAAACCGAGGGGTGAGATCAAGCCAAGGCACCAAAACACAAGATAGAGACAAGGTTCAAGGGG

GACCTATGTACAACTATTTGCGATGCCAACCGATCAGCCCTGATCAGGGTGAGTCATACGAGACTGTTAGTGCATTCATC

ACCGCTGACCTTAAGAAGTATTGCCTGAATTGGAGATACGAGACAATCAGCATATTTGCACAGAGGCTGAATGAAATATA

TGGACTACCATCCTTCTTTCAATGGTTACACAGGATATTGGAAAAATCCGTACTCTACGTCAGTGACCCACATTGTCCTC

CCGATCTAGATAATCATATCCCTCTGGACAGTGTCCCCAATGCCCAAATATTCATCAAGTACCCAATGGGCGGAATAGAA

GGTTATTGCCAAAAACTATGGACAATCAGCACTATACCATACTTGTATCTGGCAGCCTATGAGAGCGGAGTAAGAATCGC

CTCACTGGTGCAGGGTGACAATCAGACAATCGCAGTGACAAAAAGAGTTCCAAGTTCTTGGCCTTATTCACTAAAAAAGA

GGGAGGCCTCCAAGGCCGCTCAAAATTACTTCGTAGTCTTAAGGCAAAGATTGCACGATGTAGGTCATCACTTAAAGGCT

AATGAGACCATAGTATCTTCTCACTTTTTTGTATATTCCAAAGGGATTTATTATGACGGCCTGCTAGTCTCACAATCACT

AAAGAGCATCGCCAGATGTGTCTTCTGGTCCGAGACTATAGTGGATGAAACCAGAGCGGCGTGCAGTAATATTGCAACAA

CTGTCGCTAAGAGTATAGAGAGGGGTTATGATAGGTACCTTGCATATTCTTTGAATATCCTCAAGATTTTCCAACAGATC

CTTATATCCCTTAACTTCACTATTAACACAACAATGACTCAGGATGTCGTGGCACCGATCATCGAGAACGGTGATTTGCT

GATAAGGATGGCACTCTTGCCAGCACCCATCGGGGGTCTCAATTATCTTAACATGAGCAGGTTATTTGTGAGAAATATCG

GTGACCCGGTCACCTCCTCTATAGNCGACCTGAAGAGGATGATAGATGCTGGGCTAATGCCAGAAGAAACATTGCATCAA

GTGATGACCCAGACCCCGGGAGAATCATCTTACCTAGATTGGGCAAGTGACCCTTATTCTGCCAACCTACCCTGCGTACA

GAGTATCACTCGCCTTCTCAAAAACATCACTGCACGATATATTTTAATCAGCAGCCCAAACCCGATGCTGAAAGGGTTAT

TTCATGAGGGGAGTAGAGATGAAGACGAGGAGCTTGCAAGTTTCCTAATGGATCGGCATATAATTGTTCCAAGAGCTGCA

CATGAAATCTTAGACCATAGTATAACCGGAGCAAGAGAAGCTATAGCCGGGATGTTGGACACCACCAAGGGTCTGATTAG

AACAAGTATGAAACGGGGCGGCCTCACCCCTCGAGTCTTAGCCCGCCTTTCCAATTATGATTATGAACAATTCCGATCCG

GGATAACACTATTAACAAAGAAAGGGCAGTGTTATCTCATTGACAAGGACTCGTGCTCGGTGCAGCTCGCTATAGCTCTG

AGGGGCCACATGTGGGCTAAGTTGGCTCGCGGGAGACCAATCTATGGGTTGGAGGTGCCTGATGTATTAGAATCGATGAA

CGGCTACCTTATTAAACGCCACGAGTCGTGTGCTATCTGTGAAACGGGCTCAAGTCACTACGGATGGTTTTTCGTCCCTG

CAGGGTGCCAGCTTGACGATGTATCTAGAGAGACTTCGGCTCTTCGTGTACCTTATGTCGGATCAACAACTGAGGAAAGG

ACAGACATGAAACTTGCATTTGTTAGGTCTCCAAGTAGATCCCTCAAATCAGCAGTTAGAATTGCAACAGTTTACTCATG

GGCTTACGGGGATGATGAGAAATCATGGAGTGAAGCCTGGATGCTAGCTAGGCAGAGAGCCAATATCACCTTAGATGAAT

TGAGAATGATCACTCCAGTCTCCACATCCACCAACTTAGCCCACCGGTTGAGGGATCGGAGTACCCAGGTGAAGTACTCG

GGGACATCCCTTGTGAGGGTTGCAAGATACACAACCATCTCGAATGATAACTTGTCGTTTGTGATATCTGAGAAAAAAGT

GGATACCAACTTCATTTATCAGCAAGGGATGTTGCTTGGTCTTGGGATCCTTGAGAACCTCTTTAGGCTAGAGGCCACCA

CCGGAGTATCCAACACAGTGCTACACCTGCACGTGGAAACAGAATGTTGTGTTGTACCTATGGTGGATCACCCAAGGATA

CCGAGTCTCCGTAATATAAAAGTTACGGATGAGCTATGCACAAACCCTTTGATCTACGACAGGTCCCCCATCATAGAGCA

CGATGCAACCCGGCTATATTCACAGAGCCACAGGAGACATTTGGTGGAGTTTGTTACCTGGTCAACAAGCCAGCTTTACC

ATATACTGGCTAAGTCTACAGCAATGTCCATGATTGAATTGATTACGAGATTCGAGAAAGATCACATGAATGAAATAGCC

GCCCTGATTGGCGATGACGACATCAACAGCTTCATCACAGAATTTTTGCTAGTGGAGCCTAGATTATTTATAGTTTACCT

TGGTCAGTGTGCTGCCATCAATTGGGCCTTTGATATACATTATCATCGGCCCTCGGGCAAGTACCAGATGGGGGAACTTC

TCTACTCTCTGCTCTCTCGGATGAGCAAAGGAGTATATAAGATCTTTACTAATGCTTTGAGCCACCCCAAAGTTTACAAG

AAATTTTGGCGAAGTGGCATCATTGAGCCGATTCATGGACCATCCTTGGATACACAGAATTTACATGTCACTGTCTGTGA

CATGATATATGGGTCCTATGTCACCTATTTGGATCTTTTGCTGAATGATGAGCTAGATGATTACCCGTATTTGCTCTGCG

AGAGTGATGAGGACGTGGTCACAGACAGGTTCGACAACATTCAAGCCAAACATCTCTGTGTATTGGCCGATGTATATTGC

AGCTCCAAGAGGTGTCCTTCAATAATCGGGATGTCTCCTATAGAAAAATGTACCATTCTCACACATTACATCAAGGGAGA

ATCAGTACAATCCCCGTCTGGGACCTCATGGAACACTGATCCCCTTGTTGTAGATCATTACTCATGCTCTCTGACCTACC

TTCGCCGCGGTTCCATCAAACAAATCAGGTTGAGGGTGGATCCTGGGTTTGTATTCGAGGCGTTGACAGACGTCGACTTT

AAACAACCTCGCAAAGCTAAGTTAGATATATCGGTCGTTGGGTTGACTGATTTCTCTCCCCCTTACGACAACGTCGGTGA

TTTTTTAGGAACTATCAACACATTGAGGCACGATCTGCCTGTCACCGGAACCGGGGTCTCGAACTATGAAGTCCACGCTT

ATCGTAGAATTGGCCTGAATTCATCAGCATGTTATAAAGCCGTAGAGATCTCCACGTTAATCAAGCCATCCTTAGAAGTC

GGAGAGCATGGATTGTTCTTAGGAGAAGGTTCCGGTTCAATGCTGGCTGCGTACAAAGAAGTTCTTAAATTAGCAAATTG

TTACTACAACAGCGGAGTAACAGCGGAGGGCAGAGCCGGACAGAGGGAAATATCTCCCTATCCTTCAGAGATGAGCCTAG

TAGAGAATCAAATGGGGATAGAGCGGAGTGTTAAAGTGCTGTTCAACGGGAAACCTGAAGTAACTTGGGTGGGGACCACA

GATTGCTACAAGTACATAATCAGTAACATCCAAACTTCTAGTCTGGGTTTCATACATTCAGATATTGAGACGCTCCCAAC

CAAAGATGCTGTTGAGAAATTGGAGGAATTTGCCTCTATCCTATCCTTATCCTTGATTTTGGGGAAAATCGGCTCTATTA

CAGTTGTCAAAATTATGCCCATTAGCGGGGATTTTACCCAAGGCTTCATAGCCTATGCCATCCAATATTTTAGGGAGAGC

CTGCTTGCCTATCCGAGATATAGTAACTTCATCTCGACTGAGTGTTACCTTATTATGATAGGATTAAAGGCCAATCGATT

AATAAACCCAGAAGCCATTAAGCAAAGCATAATCAGAGCGGGGGTCAGGACTGCACCAGGACTTGTGAGCCATGTATTAT

CAGAGAAACAAAAAGGTTGTATTCAATCTTTCTTGGGTGATCCTTATATCCAAGGAGACTTCAATAAGCACCTTAAATCT

CTAACCTCTATTGAGAAAATCCTGGTAAATTGTGGTCTCTCGATCAATGGCATAAAAATCTGTAGAGATCTAATCCACCA

TGATATCGCCTCCGGTCCAGACGGCCTTATGAGCTCTACAATTATTTTATATAGGGAATTGGCTCATTTCAAAGATAATA

TAAGGAGTCAGCACGGAATGTTCCACCCCTACCCGGTACTGGCCAACAGCAGGCAACGTGAATTAATCCTTCGAATAGCC

AAGAAATTCTGGGGGTATGTCTTGCTATATTCTGATGACCCGGCATTAATCAGACAAACAATCAAGAACTTGAAACGGAA

TCATCTAACCTTTGACTTACATAGTAACCCATTTATTAAAGGGCTATCCAAGGCTGAGAAACTGCTGGTGCGGACGAGTT

CACTAAGAAGAGAATGGTTGTTCACTCTCGAAACGAAAGAAGTAAAAGAGTGGTTCAAATTGGTAGGTTACAGTGCACTC

ATCAGAGGCTAGTCAGCTATGCATCTGCCCCCTTCTCCTCCGCCATGAGACCCCACTGGCGATCCAGAAGATTAAAGAAA

ACTACATATTGGATAAGTATCTATTCCCAGCTTTGTCTGGN

>MF443354.1/goat/China/2014-04-03

ACCAAACAAAGTTGGGTAAGGATAGATCTTATAATAACTATGGACTGGCAAACTTAGGAGTAAAGATCCTACTGTCGGGG

GGAGGAGGAGGAGCAAGATCCTTGATCATGGCGACTCTCCTTAAAAGCTTAGCATTGTTCAAAAGGAACAAAGACAAAGC

GCCGACAGCATCAGGTTCAGGAGGGGCCATCCGGGGGATTAAGAATGTTATCATAGTCCCGATTCCCGGAGACTCGTCCA

TCACTACCCGTTCAAGACTGCTCGACAGGCTTGTCAGATTGGCCGGAGATCCTGACATCAACGGGTCAAAGCTGACCGGG

GTGATGATCAGCATGTTATCCTTGTTCGTAGAGTCACCCGGGCAATTGATACAGCGAATCACAGATGATCCGGATGTTAG

TATCCGCCTTGTTGAGGTAGTTCAAAGTACTAGGTCCCAGTCCGGGTTGACCTTTGCATCACGTGGTGCTGATTTAGACA

ACGAGGCAGACATGTATTTTTCAACTGAGGGGCCCTCGAGTGGAGGTAAGAAAAGGATCAACTGGTTTGAGAACAGAGAA

ATAATAGACATAGAGGTGCAGGATCCAGAAGAGTTCAATATGTTGTTAGCCTCCATACTAGCACAAGTCTGGATCCTTCT

GGCCAAGGCTGTTACGGCACCAGATACGGCAGCTGACTCAGAACTGAGAAGGTGGGTTAAATACACACAACAAAGGAGAG

TGATTGGGGAATTTCGCCTTGACAAGGGGTGGCTGGACGCAGTCCGCAACAGAATTGCAGAAGATCTATCACTCCGGCGG

TTCATGGTATCTCTCATACTTGACATCAAGAGGACCCCAGGCAACAAGCCAAGGATTGCAGAAATGATCTGCGACATTGA

CAACTATATTGTCGAAGCAGGACTCGCCAGTTTCATCCTTACTATCAAATTTGGTATCGAAACCATGTATCCTGCATTAG

GGCTCCACGAGTTTGCCGGAGAATTGTCCACTATTGAGTCCTTGATGAACTTGTATCAACAGCTAGGAGAGGTTGCACCC

TACATGGTAATTCTAGAGAATTCAATTCAGAACAAGTTTAGTGCAGGAGCTTATCCCCTCCTCTGGAGCTATGCGATGGG

TGTCGGAGTCGAGCTGGAGAACTCAATGGGGGGCTTGAACTTTGGCAGGTCATATTTTGACCCTGCCTATTTTCGTCTCG

GACAGGAGATGGTCAGAAGATCTGCAGGAAAGGTCAGCTCTGTGATTGCGGCTGAGCTCGGCATCACAGCAGAGGAAGCT

AAACTAGTCTCGGAAATCGCCTCGCAGGCTGGGGACGAAAGAACCGCCAGAGGGACTGGGCCTCGACAGGCGCAGGTTTC

CTTCCTTCAGCACCAAACAGGAGGGGGAGAGTCGTCCGCACCAGCGACCAGAGAAGGGGTCAAAGCTGCGATCCCAAACG

GAGCTGAAGAAAGGGACAGAAAGCAAACACGCCCAGGAAGGCCCAGAGGAGAGACCTCCGGCCAACTGCTCCTGGACATC

ATGCCAGAGGATGAGATCTTGCGAGAGTCTGGTCAAAACCCTCGTGAGGCTCAAAGATCGGCCGAGGCACTCTTCAGGCT

GCAGGCCATGGCCAAGATTCTGGAGGACCAGGAGGAGGGAGAAGACAACAGTCAGGTCTACAACGACAAGGATCTCCTCG

GCTAAGCAGACGCACCCTCTGTCGAAATCAGTGACAAGACATCTCCTACCAGTATTATAAAAAACTTAGGACCCAGGTCC

AAGCAAGCACACATCGACACTCCAACCAGTCGAGCGGAGACCACCGATGGCAGAAGAACAAGCATACCATGTCAACAAGG

GGCTGGAATGTATCAAGTCTCTCAAAGCCTCACCCCCGGATCTATCCACCATCAGAGATACCATCGAGAGCTGGAGAGAG

GGGCTTAGCCCCTCGGGCCGTGCAACACCGAACCCTGATACGTCCGAGGGAGATCATCAGAATATCAACCAATCATGCTC

ACCAGCAATCGGACCAAACAAAGTCTACTTGTCTCCTGAAGATAATCTCGGATTTAGAGAGATCACTGGCAACGACTGTG

AGGCTGGGCTCGGAGGGGTCCAGAGAGAAGGATCCAACTCTCAAGTACAGCGTTACCATGTTTATAGTCACGGGGGTGAA

GAGATTGAAGGACTCGAGGATGCTGACTCTCTCGTGGTTCAAGCAGATCCTCCATTTGCTAACGTCTTCAATGGAGGAGA

GGATGGATCTGACGACAGCGATGTGGACTCTGGCCCAGATGATCCCGGCAGAGATACTCTATATGACCGGGGATCTGTTG

CCGGCAATGGTATCGCTAGGTCCACAGATGTCGAAAAACTAGAAGGTGCTGATATTCAAGAAGTTCTTAACTCCCAGAAA

GGCAAAGGAGGAAGATTCCAGGGCGGGAAAACCTTGCGAGTCCCGGAAATACCCGATGTCAAGCACTCCAGACCATCAGC

CCAATCAATTAAAAAGGGCACAGACGGGAACTCAGTCTCATCTGGAACGGTGATAGAGTGTTTATCGATAAGTGGTGCAA

CCCAAACTGTGCCAGAGTCAAGATGGGAGTCATCAGAGCAAAATGCGTCTGTGGGGAGTGTCCTCAAGTCTGCGAGGAGT

GCAAAGACGATCCAGGGGTCGACACAAGAATCTGGTACCATAGCATCACTGACTCAGCCTAAAGAGAATGACTCCGAGTA

TGAGTATGAGGATGACCTATTTACAGAGATTCAGGACATCCGTGCAAGCATTGCCAAGATCCATGATGACAATAAAACTA

TCCTCTCAAAGCTTGATTCTATACTGTTATTGAAAGGGGAAGTCGACACTATCAAGAAACAAATCAGCAAGCAGAATATA

AGTATATCCACCATTGAGGGCCATCTCTCCAGTATAATGATAGCCATCCCTGGCTTTGGGAAGGACATCAAGGACCCAAC

ATCCGAAGTCGAGTTGAACCCAGATCTAAGACCTATAATAAGCCGTGACTCTGGCAGAGCTCTCGCGGAGGTCCTCAAGA

AACCCGCTGTGGATAGGTCTCAGAAAATTGGAACCAAAGCCAACTCCAGCTCAAAGGGTCAGCTTCTTAAGGATCTCCAG

CTAAAACCTGTCGACAAGCAGGCAAGCTCTGCAATCGGGTTTGTCCCGTCCGACCATGAATCATCCAGAAATGTCATCCG

CTCCATAATCAAGTCGAGCAAGCTAAATATTGATCACAAGGACTATCTTCTAGATTTACTAAATGATGTGAAAGGCTCCA

AGGATCTTAAGGAATTTCACAAGATGCTAACAGCAATTCTCGCCAAGCACCCGTAACACATCCTCCAGTCATCATCTCAT

ACTCGACTAAAAACATCCTTTCAATCAGGCTATTACAAAAAACTTAGGAGCAAGGGCAACTGAGCTTCGCAGACAGGATG

ACCGAGATCTACGACTTCGATAAATCAGCATGGGATGTCAAAGGGTCAATTGCCCGCATAGAACCCACCACCTACCACGA

CGGCCGACTGATACCCCAGGTGAGGGTCATCGATCCTGGTCTGGGAGACAGAAAAGATGAGTGCTTTATGTACCTGTTTC

TCCTAGGAGTGATTGAGGATAACGACCCCCTGTCTCCCCCCGTCGGAAGAACCTTTGGCTCTCTACCTCTAGGGGTTGGT

AGGTCAACTGCCAAGCCAGAAGAACTACTAAGGGAGGCCACAGAATTAGATATAGTGGTGAGGCGCACTGCAGGAGTAAA

TGAGAAACTGGTATTTTACAACAACACTCCGCTGTCCTTGTTAACGCCCTGGAAGAAAGTTCTGACAACCGGAAGTGTGT

TTAGCGCTAACCAGGTTTGCAATGCAGTCAACCTAGTCCCGCTTGATACTCCCCAGAGATTCAGGGTTGTGTACATGAGC

ATAACTAGATTATCAGACAATGGTTACTATAGTGTGCCCAGAAGAATGCTGGAGTTCCGCTCGGCCAATGCAGTAGCCTT

CAACATCTTGGTTACACTAAGAATTGAAAATGGCACAAACCCTAGAAGATACATAGTCGGCTCATGGGAGAATTCAGAGG

TCACATTTATGGTACACGTGGGCAACTTTAGGAGAAAGAAGAACGAAGTATACTCTGCTGATTACTGCAAAATGAAGATT

GAAAAGATGGGTTTAGTTTTTGCCCTGGGTGGAATAGGCGGAACAAGTCTCCATATTAGAAGCACCGGGAAAATGAGCAA

AACCCTCCATGCACAGCTGGGGTTCAAGAAAATCTTATGTTACCCCCTAATGGATGTCAATGAGGATCTTAACCGATATC

TCTGGCGGGCAGAGTGCCGAATAGTCAAAATCCAAGCTGTCTTACAGCCATCAGTACCCCAAGAGTTCCGTGTCTACGAT

GATGTCATCATCAACGATGATCAAGGCTTGTTCAAGATCTTGTAGTTCATTTGCAAAGGAACGAACCCACCCCCAGGAAG

GGGCCCCACCCTCCCCCG-CCAAAAAACACACTCCAAGGCTCCCCGGCACAGCGCCAACACAGCTACTCCCGCGATGGAA

CGAGGTCGGCC-CCCGAACAAATCCGGTCCAAGGGGGGGGGCCCCCCCCCTGACCAACCGGGCGCCGACCGAGCAGGCCG

GCGAGAAGCCCCCAAGAGGCCCCGACGAAGACCGAGCACCCCATCCCCCC-AAAAACGTCCCCC-AACACGCCCGAGGCT

AGCCACCGGACACCCCCAGACCCGCCACCGGCGAGACAGAACCCCAAGGGGGGACCCTCCAAACCAAAGGAATCACCGGG

GTAGGCTCCGCCGAACCAGCACAGACCCGACCCAAGCAAGGCGGACCCCCAACACGAAACTGCCCAGCACACCCTACTCA

TCATCAAACAAAACTTAGGGGCCAAGTCCATACGAGAGGCCCTAGTCCGTCAGGCGGAGGGACGCAGAAAGGAAGGAGAC

ACGACCGTCCCAGAAGGAGGACACCCAACCACCGAAACGCCCGCCCCCAGGGAAGGAG--CCA-CCCCCACGCAGCGAAG

GAGCGGGTGGGAGACCTCGGGGAGGGCCCGACCGGGACCCGGGGGCCGGCGGCGCCGGCGGTGAGCAGGCCGGGCGCATA

CCTGACCCCACCGGAGGATCCGAGGAGGGCCCGACCCCGCGTGGACCAAACGGCAGGGAGCCCAACCCCACCAAAAAGAG

AGGCGAGGCCCCCCCTCCCTCCCCCCCCGCACACAACCCGCAGACAGACCGAGGGAGGGCCATCCCCCAGCGAGACAGGC

GGCTCCCTCGCAGCCGAGCCGCAGCTCCCCAGAATAGGACACCCCCGCACACTCCGAACACCCCCTCGCCCCCCGCCACC

GGGACAGGGGCTCCACCACCACAGACCGGAGCCACCGACAGCCAGCCAGAGGTCCCCTCCAGTCCCGCCCGGCGCCCCCC

CGAAGAGGCCCCGGCAAGGCCCCCCAGCCCGACGTCCCCCGCACGCACCGAAAGAAGGACCCGCCCGAGAACGTGCGGCA

CATCCATGCGCAAACATCATGACGCGGGTCGCAATTTTGACATTTCTGTTTCTTTTCCCAAATGTTGTTGCGTGTCAGAT

TCACTGGGGCAATCTATCCAAGATCGGGATTGTAGGAACAGGGAGTGCCAGCTACAAGGTGATGACTAGGCCAAGCCACC

AGACTCTGGTTATAAAGTTAATGCCAAATATAACGGCCATCGACAATTGTACAAAGTCAGAGATTGCAGAGTACAAGAGA

TTGCTGATCACAGTGTTAAAGCCTGTAGAGGATGCTCTGTCGGTGATAACCAAGAATGTAAGACCAATTCAAACTCTAAC

ACCTGGGCGTAGAACCCGCCGTTTTGCTGGAGCTGTTCTGGCCGGGGTAGCACTTGGAGTTGCGACAGCCGCTCAGATAA

CTGCAGGAGTCGCCCTTCATCAATCATTGATGAACTCCCAAGCAATTGAGAGTTTAAAAACCAGTCTTGAGATGTCGAAT

CAGGCAATAGAAGAAATCAGACTTGCAAATAAGGAGACCATACTGGCAGTACAGGGCGTCCAGGATTATATCAACAATGA

GCTCGTCCCTTCTGTTCATAGAATGTCATGCGAGCTGGTAGGTCACAAGCTCGGCCTCAAGCTCCTTAGGTACTACACCG

AGATCCTGTCCATATTCGGGCCCAGTCTTCGAGACCCGATATCTGCCGAAATATCAATCCAGGCACTTAGTTATGCATTA

GGCGGAGACATTAATAAAATCCTGGACAAGCTTGGGTATAGCGGTGGGGATTTCCTTGCCATCCTAGAAAGCAAGGGAAT

AAAGGCCCGGGTTACATATGTGGACACAAGAGATTACTTTATAATCCTTAGCATCGCCTACCCAACCTTATCTGAGATCA

AGGGAGTGATAGTTCACAAGATAGAAGCTATAACATACAACATTGGGGCACAGGAGTGGTATACTACTATCCCTAAATAT

GTAGCCACTCAGGGGTATCTGATATCGAACTTTGATGAGACGTCATGCGTATTCACTCCAGAGGGGACAGTTTGCAGCCA

GAATGCGTTGTATCCAATGAGCCCATTGCTTCAGGAATGTTTCAGGGGGTCAACAAAATCGTGCGCCAGAACCTTAGTTT

CAGGGACCATAAGTAATAGATTTATCCTATCAAAAGGGAACCTGATTGCAAATTGTGCGTCAGTTTTGTGCAAATGTTAC

ACAACAGAGACAGTTATCAGCCAAGATCCTGACAAACTACTAACTGTTGTAGCATCCGACAAGTGTCCTGTAGTTGAGGT

GGATGGAGTGACAATACAGGTCGGCAGTCGAGAGTATCCGGATTCTGTATACTTACACAAAATAGACTTAGGTCCAGCCA

TCTCCCTAGAAAAACTGGATGTAGGCACCAATTTAGGCAATGCAGTCACAAGACTGGAGAATGCAAAGGAGCTCCTAGAT

GCATCAGACCAAATACTGAAGACTGTTAAAGGGGTACCTTTCAGTGGGAATATGTACATAGCACTGGCAGCTTGCATAGG

AGTATCCCTGGGCCTTGTCACATTAATATGCTGCTGTAAGGGGAGGTGTAAGAACAAGGAAATCCCTATCTCCAAAATCA

ACCCAGGGCTCAAACCCGACCTGACCGGGACCTCAAAGTCGTACGTAAGATCACTGTAGTCAGAATCACCTGGATCATCT

GGCATCACACACATACATGCACGACACAGGCAGTCCGAGGACGCAAGAAACCCAGCCTCCGGTCACCCACCCGACCCCAC

TCCACGCTCCACCACACATTAGTCATCAAACAAAACTTAGGACGAAAGGTCAATCACCATGTCCGCACAAAGGGAGAGGA

TCAATGCCTTCTACAAAGACAATCCTCACAATAAGAACCATAGGGTGATCCTGGATAGAGAACGCTTGGTCATTGAAAGA

CCCTACATCTTGCTTGGAGTCCTGCTGGTAATGTTCCTGAGTCTAATCGGACTGCTGGCCATTGCAGGGATCAGGCTTCA

CCGGGCCACCGTTGGAACTTCAGAGATCCAGAGTCGGCTGAATACCAATATTAAGTTGACCGAATCTATTGATCACCAGA

CTAAGGATGTCTTAACTCCCCTTTTTAAAATCATTGGCGATGAAGTCGGCATCAGAATTCCACAGAAATTCAGTGATCTT

GTCAAGTTCATCTCCGATAAGATTAAATTCCTCAACCCTGATAGAGAGTATGATTTCAGAGATCTCCGGTGGTGTATGAA

TCCCCCCGAGAGAGTCAAAATTAATTTTGATCAGTTTTGTGAGTACAAGGCTGCGGTTAAGTCAATTGAACATATATTTG

AGTCACCACTCAACAAGTCAAAAAAGCTGCAATCTTTGACTCTCGGGCCCGGAACAGGCTGTCTAGGCAGGACAGTAACA

AGAGCCCATTTCTCAGAACTTACAATGACCTTAATGGACCTGGATCTAGAGATGAAGCACAACGTGTCCTCAGTGTTTAC

CGTAGTTGAAGAGGGATTATTCGGAAGAACATATACCGTCTGGAGATCCGATGCCAGGGATCCGAGCACCGATCTAGGTA

TCGGCCATTTTTTAAGAGTCTTCGAGATTGGACTGGTAAGAGATCTTGGGCTGGGTCCCCCTGTTTTTCATATGACCAAC

TATCTCACGGTGAACATGAGTGATGACTATCGGAGATGTCTTTTAGCGGTAGGGGAGTTGAAGTTGACAGCCCTATGCAC

CTCATCTGAGACTGTGACACTGAGTGAGAGAGGAGTTCCAAGGAGGGAACCTCTTGTGGTTGTGATACTTAATCTAGCTG

GACCCACTCTAGGGGGAGAGCTATACAGTGTCTTGCCTACCTCTGATCTCATGGTGGAGAAACTCTATTTGTCTTCACAT

AGAGGGATCATCAAAGATGATGAGGCCAATTGGGTAGTGCCGTCTACCGATGTTCGTGATCTTCAGAATAAAGGTGAATG

TCTGGTGGAAGCATGCAAGACTCGACCTCCTTCATTTTGCAATGGCACAGGATCAGGCCCGTGGTCAGAAGGGAGAATCC

CTGCCTACGGGGTGATCAGGGTCAGTCTTGACTTAGCGAGTGACCCGGATGTAGTTATCACTTCAGTGTTTGGCCCACTG

ATACCTCACCTATCCGGCATGGATCTTTACAACAACCCGTTTTCAAAAGCTGTATGGTTGGCTGTACCACCTTATGAGCA

GTCATTTCTAGGAATGATAAATACAATTGGATTCCCTAACAGAGCAGAGGTTATGCCGCACATTTTGACCACAGAGATCA

GAGGCCCTCGGGGTCGTTGCCATGTTCCCATAGAATTGTCCCGCCGGGTTGATGACGATATCAAGATCGGGTCCAACATG

GTCATATTGCCGACGATGGACCTGAGGTATATTACAGCCACTTATGATGTTTCCAGGAGCGAGCATGCAATCGTGTACTA

TATCTATGACACGAGTCGCTCATCATCTTACTTCTACCCAGTTCGACTGAATTTCAAAGGCAATCCTCTCTCTTTGAGGA

TAGAGTGTTTCCCCTGGCGTCATAAGGTGTGGTGCTACCATGATTGTCTTATATACAACACCATAACAGGTGAAGAGGTC

CATACGAGAGGGCTGACCGGCATAGAGGTAACATGCAATCCAGTCTGAGCAGAGCTACGACCGTCACTCAGGCAGTCCCT

TGAGTCGCCACCGAGTCCAAGCAGCACAGCCTGGGACAATCAACGGCACAGCCCAGCCAACAATGTTATAAAAAACTTAG

GAGCCAGGGTTGTAGGGGCCATGGACTCTCTATCAGTCAATCAGGTCTTGTACCCTGAGGTTCATCTAGACAGCCCTATT

GTCACAAACAAACTAGTTGCCATCCTTGAGTACTCGGGTATCGACCATAACTATGTTCTTGAAGACCAGACTCTTGTCAA

GAATATTAGGTATAGGCTGGGGTGCGGTTTTTCAAATCAAATGATCATCAATAACAGGGGGGTGGGTGAAACAGTCAATT

CTAAACTTAAAAGTTACCCCCATAATCGTCATATCATATACCCGGATTGCAATAAGGAGTTGTTTTGTATCAAGGATAGC

TGCATATCTAAGAAGCTCTCGGAGCTATTCAAGAAGGGTAATTCCTTGTACTCTAAGATAAGTCACCAGGTACTGGATTG

TCTTAAGAGAGTCAATGGGAAATTAGGTCTGGGCACAGATCTTACCCATGGTCTGAAGGAGGGTATCCTTGACCTAGGGT

TGCACATGCATAGCTCTCAATGGTTCGAGACTTTCCTGTTCTGGTTCACTATCAAGACAGAGATGAGATCAATGATCAAA

GAACAGTCCCATATATGCCACAAGAGGAGGTATAACCCGACTTTTGTGTCGGGGGATGCATTCGAGGTGCTCGTATCGCG

AGACCTCGTTGTGATAATTGATAAGAATACCCAGTATGTCTTCTACTTAACATTTGAGCTGGTCCTTATGTATTGTGATG

TCATAGAGGGTAGACTTATGACGGAGACAGCCATGGCCATAGACCAGAGATATTCAGAACTTCTAAGCCGGGTCAGATAC

TTGTGGGATCTTATTGATGGGTTCTTCCCAACACTAGGCAACACCACATACCAAATTGTTGCTCTGCTTGAACCATTATC

GTTGGCTTATCTTCAACTTCAGGATGTCACTCTGGAGTTAAGAGGTGCTTTTTTAGACCATTGCTTCAAAGAACTCTATG

AGATACTGGAACATTGCGGCATTGACACAGCAGGCACTTACAATTCCATCACTGAAGGATTGGATTACGTATTCATCACC

CACGATATACATCTAACTGGGGAGATTTTTTCATTTTTCCGGAGTTTCGGACACCCCCGTCTAGAAGCAGTCACCGCTGC

GGAAAATGTCAGAAAACATATGAACCAACCGAAGGTAATCAGTTATGAGACTATGATGAAAGGCCATGCGGTATTTTGCG

GGATAATCATAAATGGTTTTCGGGACCGACATGGCGGCAGCTGGCCCCCTGTTGCACTGCCAGAACATGCCTCTGCTGCG

ATCCGGAATGCGCAGGCATCAGGTGAGGGACTGACCCATGACCTGTGTATAGACAACTGGAAATCCTTTGTTGGATTCAA

ATTTGGCTGCTTTATGCCACTCAGCCTAGATAGTGATTTGACCATGTATCTCAAAGACAAGGCATTGGCTGCACTGAAGA

ATGAGTGGGATTCAGTTTACCCGAAAGAATACCTCCGTTATAACCCACCTAGAGGGACAGAGTCAAGGCGATTGGTAGAG

GTATTCCTGAATGACTCCAGCTTTGATCCTTATAACATGATAATGTACGTGGTGAATGGCTCCTACCTTAAAGACCCTGA

GTTTAACCTCTCATACAGCCTAAAAGAGAAGGAGATAAAGGAGACAGGGCGGTTGTTTGCCAAAATGACCTATAAGATGC

GGGCCTGTCAGGTAATCGCTGAAAATCTGATATCGAATGGTGTTGGGAAGTATTTCCGAGACAATGGGATGGCAAAGGAC

GAGCATGACCTAACAAAAGCCCTTCACACCCTGGCAGTCTCAGGTGTTCCCAAAAACAACAAAGATAACCACCGAGGTGG

GCCTCCCAGAAGGACAACAAACCGAGGGGTGAGATCAAGCCAAGGCACCAAAACACAAGATAGAGACAAGGTTCAAGGGG

GACCTATGTACAACTATTTGCGATGCCAACCGATCAGCCCTGATCAGGGTGAGTCATACGAGACTGTTAGTGCATTCATC

ACCGCTGACCTTAAGAAGTATTGCCTGAATTGGAGATACGAGACAATCAGCATATTTGCACAGAGGCTGAATGAAATATA

TGGACTACCATCCTTCTTTCAATGGTTACACAGGATATTGGAAAAATCCGTACTCTACGTCAGTGACCCACATTGTCCTC

CCGATCTAGATAATCATATCCCTCTGGACAGTGTCCCCAATGCCCAAATATTCATCAAGTACCCAATGGGCGGAATAGAA

GGTTATTGCCAAAAACTATGGACAATCAGCACTATACCATACTTGTATCTGGCAGCCTATGAGAGCGGAGTAAGAATCGC

CTCACTGGTGCAGGGTGACAATCAGACAATCGCAGTGACAAAAAGAGTTCCAAGTTCTTGGCCTTATTCACTAAAAAAGA

GGGAGGCCTCCAAGGCCGCTCAAAATTACTTCGTAGTCTTAAGGCAAAGATTGCACGATGTAGGTCATCACTTAAAGGCT

AATGAGACCATAGTATCTTCTCACTTTTTTGTATATTCCAAAGGGATTTATTATGACGGCCTGCTAGTCTCACAATCACT

AAAGAGCATCGCCAGATGTGTCTTCTGGTCCGAGACTATAGTGGATGAAACCAGAGCGGCGTGCAGTAATATTGCAACAA

CTGTCGCTAAGAGTATAGAGAGGGGTTATGATAGGTACCTTGCATATTCTTTGAATATCCTCAAGATTTTCCAACAGATC

CTTATATCCCTTAACTTCACTATTAACACAACAATGACTCAGGATGTCGTGGCACCGATCATCGAGAACGGTGATTTGCT

GATAAGGATGGCACTCTTGCCAGCACCCATCGGGGGTCTCAATTATCTTAACATGAGCAGGTTATTTGTGAGAAATATCG

GTGACCCGGTCACCTCCTCTATAGCCGACCTGAAGAGGATGATAGATGCCGGGCTAATGCCAGAAGAAACATTGCATCAA

GTGATGACCCAGACCCCGGGAGAATCATCCTACCTAGATTGGGCAAGTGACCCTTATTCTGCCAACCTACCCTGCGTACA

GAGTATCACTCGCCTTCTCAAAAACATCACTGCACGATATATTTTAATCAGCAGCCCAAACCCGATGCTGAAAGGGTTAT

TTCATGAGGGGAGTAGAGATGAAGACGAGGAGCTTGCAAGTTTCCTAATGGATCGGCATATAATTGTTCCAAGAGCTGCA

CATGAAATCTTAGACCATAGTATAACCGGAGCAAGAGAAGCTATAGCCGGGATGTTGGACACCACCAAGGGTCTGATTAG

AACAAGTATGAAACGGGGCGGCCTCACCCCTCGAGTCTTAGCCCGCCTTTCCAATTATGATTATGAACAATTCCGATCCG

GGATAACACTATTAACAAAGAAAGGGCAGTGTTATCTCATTGACAAGGACTCGTGCTCGGTGCAGCTCGCTATAGCTCTG

AGGGGCCACATGTGGGCTAAGTTGGCTCGCGGGAGACCAATCTATGGGTTGGAGGTGCCTGATGTATTAGAATCGATGAA

CGGCTACCTTATCAAACGCCACGAGTCGTGTGCTATCTGTGAAACGGGCTCAAGTCACTACGGATGGTTTTTCGTCCCTG

CAGGGTGCCAGCTTGACGATGTATCTAGAGAGACTTCGGCTCTTCGTGTACCTTATGTCGGATCAACAACTGAGGAAAGG

ACAGACATGAAACTTGCATTTGTTAGGTCTCCAAGTAGATCCCTCAAATCAGCAGTTAGAATTGCAACAGTTTACTCATG

GGCTTACGGGGATGATGAGAAATCATGGAGTGAAGCCTGGATGCTAGCTAGGCAGAGAGCCAATATCACCTTAGATGAAT

TGAGAATGATCACTCCAGTCTCCACATCCACCAACTTAGCCCACCGGTTGAGGGATCGGAGTACCCAGGTGAAATACTCG

GGGACATCCCTTGTGAGGGTTGCAAGATACACAACCATCTCCAATGATAACTTGTCGTTTGTGATATCTGAGAAAAAAGT

GGATACCAACTTCATTTATCAGCAAGGGATGTTGCTTGGTCTTGGGATCCTTGAGAACCTCTTTAGGCTAGAGGCCACCA

CCGGAGTATCCAACACAGTGCTACACCTGCACGTGGAAACAGAATGTTGTGTTGTACCTATGGTGGATCACCCAAGGATA

CCGAGTCTCCGTAATATAAAAGTTACGGATGAGCTATGCACAAACCCTTTGATCTACGACAGGTCCCCCATCATAGAGCA

CGATGCAACCCGGCTATATTCACAGAGCCACAGGAGACATTTGGTGGAGTTTGTTACCTGGTCAACAAGCCAGCTTTACC

ATATACTGGCTAAGTCTACAGCAATGTCCATGATTGAATTGATCACGAGATTCGAGAAAGATCACATGAATGAAATAGCC

GCCCTGATTGGCGATGACGACATCAACAGCTTCATCACAGAATTTTTGCTAGTGGAGCCTAGATTATTTATAGTTTACCT

TGGTCAGTGTGCTGCCATCAATTGGGCCTTTGATATACATTATCATCGGCCCTCGGGCAAGTACCAGATGGGGGAACTTC

TCTACTCTCTGCTCTCTCGGATGAGCAAAGGAGTATATAAGATCTTTACTAATGCTTTGAGCCACCCCAAAGTTTACAAG

AAATTTTGGCGAAGTGGCATCATTGAGCCGATTCATGGCCCATCCTTGGATACACAGAATTTACATGTCACTGTCTGTGA

CATGATATATGGGTCCTATGTCACCTATTTGGATCTTTTGCTGAATGATGAGCTAGATGATTACCCGTATTTGCTCTGCG

AGAGTGATGAGGACGTGGTCACAGACAGGTTCGACAACATTCAAGCCAAACATCTCTGTGTATTGGCCGATGTATATTGC

AGCTCCAAGAGGTGTCCTTCAATAATCGGGATGTCTCCTATAGAAAAATGTACCATTCTCACACATTACATCAAGGGAGA

ATCAGTACAATCCCCGTCTGGGACCTCATGGAACACTGATCCCCTTGTAGTAGATCATTACTCATGCTCTCTGACCTACC

TTCGCCGCGGTTCCATCAAACAAATCAGGTTGAGGGTGGATCCTGGGTTTGTATTCGAGGCGTTGACAGACGTCGACTTT

AAACAACCTCGCAAAGCTAAGTTAGATATATCGGTCGTTGGGTTGACTGATTTCTCTCCCCCTTACGACAACGTCGGTGA

TTTTTTAGGGACTATCAACACATTGAGGCACGATCTGCCTGTCACCGGAACCGGGGTCTCGAACTATGAAGTCCACGCTT

ATCGTAGAATTGGCCTGAATTCATCAGCATGTTATAAAGCCGTAGAGATCTCCACGTTAATCAAGCCATCCTTAGAAGTC

GGAGAGCATGGATTGTTCTTAGGAGAAGGTTCCGGTTCAATGCTGGCTGCGTACAAAGAAGTTCTTAAATTAGCAAATTG

TTACTACAACAGCGGAGTAACAGCGGAGGGCAGAGCCGGACAGAGGGAAATATCTCCCTATCCTTCAGAGATGAGCCTAG

TAGAGAATCAAATGGGGATAGAGCGGAGTGTTAAAGTGCTGTTCAACGGGAAACCTGAAGTAACTTGGGTGGGGACCACA

GATTGCTACAAGTACATAATCAGTAACATCCAAACCTCTAGTCTGGGTTTCATACATTCAGATATTGAGACGCTCCCAAC

CAAAGATGCTGTTGAGAAATTGGAGGAATTTGCCTCTATCCTATCCTTATCCTTGATTTTGGGGAAAATCGGCTCTATTA

CAGTTGTCAAAATTATGCCCATTAGCGGGGATTTTACCCAAGGCTTCATAGCCTATGCCATCCAATATTTTAGGGAGAGC

CTGCTTGCCTATCCGAGATATAGTAACTTCATCTCGACTGAGTGTTACCTTATTATGATAGGATTAAAGGCCAATCGATT

GATAAACCCAGAAGCCATTAAGCAAAGCATAATCAGAGCGGGGGTCAGGACTGCACCAGGACTTGTGAGCCATGTATTAT

CAGAGAAACAAAAAGGTTGTATTCAATCTTTCTTGGGTGATCCTTATATCCAAGGAGACTTCAATAAGCACCTTAAATCT

CTAACCTCTATTGAGAAAATCCTGGTAAATTGTGGTCTCTCGATCAATGGCATAAAAATCTGTAGAGATCTAATCCACCA

TGATATCGCCTCCGGTCCAGATGGCCTTATGAGCTCTACAATTATTTTATATAGGGAATTGGCTCATTTCAAAGATAATA

TAAGGAGTCAGCACGGAATGTTCCACCCCTACCCGGTACTGGCCAACAGCAGGCAACGTGAATTAATCCTTCGAATAGCC

AAGAAATTCTGGGGGTATGTCTTGCTATATTCTGATGACCCGGCACTAATCAGACAAACAATCAAGAACTTGAAACGGAA

TCATCTAACCTTTGACTTACATAGTAACCCATTTATTAAAGGGCTATCCAAGGCTGAGAAACTGCTGGTGCGGACGAGTT

CACTAAGAAGAGAATGGTTGTTCACTCTCGAAACGAAAGAAGTAAAAGAGTGGTTCAAATTGGTAGGTTACAGTGCACTC

ATCAGAGGCTAGTCAGCTATGCATCTGCCCCCTTCTCCTCCGCCATGAGACCCCACTGGCGATCCAGAAGATTAAAGAAA

ACTACATATTGGATAAGTATCTATTCCCAGCTTTGTCTGGT

>MF443351.1/sheep/China/2014-01-22

ACCAAACAAAGTTGGGTAAGGATAGATCTTATAATAACTATGGACTGGCAAACTTAGGAGTAAAGATCCTACTGTCGGGG

GGAGGAGGAGGAGCAAGATCCTTGATCATGGCGACTCTCCTTAAAAGCTTAGCATTGTTCAAAAGGAACAAAGACAAAGC

GCCGACAGCATCAGGTTCAGGAGGGGCCATCCGGGGGATTAAGAATGTTATCATAGTCCCGATTCCCGGAGACTCGTCCA

TCACTACCCGTTCAAGACTGCTCGACAGGCTTGTCAGATTGGCCGGAGATCCTGACATCAACGGGTCAAAGCTGACCGGG

GTGATGATCAGCATGTTATCCTTGTTCGTAGAGTCACCCGGGCAATTGATACAGCGAATCACAGATGATCCGGATGTTAG

TATCCGCCTTGTTGAGGTAGTTCAAAGTACTAGGTCCCAGTCCGGGTTGACCTTTGCATCACGTGGTGCTGATTTAGACA

ACGAGGCAGACATGTATTTTTCAACTGAGGGGCCCTCGAGTGGAGGTAAGAAAAGGATCAACTGGTTTGAGAACAGAGAA

ATAATAGACATAGAGGTGCAGGATCCAGAAGAGTTCAATATGTTGTTAGCCTCCATACTAGCACAAGTCTGGATCCTTCT

GGCCAAGGCTGTTACGGCACCAGATACGGCAGCTGACTCAGAACTGAGAAGGTGGGTTAAATACACACAACAAAGGAGAG

TGATTGGGGAATTTCGCCTTGACAAGGGGTGGCTGGACGCAGTCCGCAACAGAATTGCAGAAGATCTATCACTCCGGCGG

TTCATGGTATCTCTCATACTTGACATCAAGAGGACCCCAGGCAACAAGCCAAGGATTGCAGAAATGATCTGCGACATTGA

CAACTATATTGTCGAAGCAGGACTCGCCAGTTTCATCCTTACTATCAAATTTGGTATCGAAACCATGTATCCTGCATTAG

GGCTCCACGAGTTTGCCGGAGAATTGTCCACTATTGAGTCCTTGATGAACTTGTATCAACAGCTAGGAGAGGTTGCACCC

TACATGGTAATTCTAGAGAATTCAATTCAGAACAAGTTTAGTGCAGGAGCTTATCCCCTCCTCTGGAGCTATGCGATGGG

TGTCGGAGTCGAGCTGGAGAACTCAATGGGGGGCTTGAACTTTGGCAGGTCATATTTTGACCCTGCCTATTTTCGTCTCG

GACAGGAGATGGTCAGAAGATCTGCAGGAAAGGTCAGCTCTGTGATTGCGGCTGAGCTCGGCATCACAGCAGAGGAAGCT

AAACTAGTCTCGGAAATCGCCTCGCAGGCTGGGGACGAAAGAACCGCCAGAGGGACTGGGCCTCGACAGGCGCAGGTTTC

CTTCCTTCAGCACCAAACAGGAGGGGGAGAGTCGTCCGCACCAGCGACCAGAGAAGGGGTCAAAGCTGCGATCCCAAACG

GAGCTGAAGAAAGGGACAGAAAGCAAACACGCCCAGGAAGGCCCAGAGGAGAGACCTCCGGCCAACTGCTCCTGGACATC

ATGCCAGAGGATGAGATCTTGCGAGAGTCTGGTCAAAACCCTCGTGAGGCTCAAAGATCGGCCGAGGCACTCTTCAGGCT

GCAGGCCATGGCCAAGATTCTGGAGGACCAGGAGGAGGGAGAAGACAACAGTCAGGTCTACAACGACAAGGATCTCCTCG

GCTAAGCAGACGCACCCTCTGTCGAAATCAGTGACAAGACATCTCCTACCAGTATTATAAAAAACTTAGGACCCAGGTCC

AAGCAAGCACACATCGACACTCCAACCAGTCGAGCGGAGACCACCGATGGCAGAAGAACAAGCATACCATGTCAACAAGG

GGCTGGAATGTATCAAGTCTCTCAAAGCCTCACCCCCGGATCTATCCACCATCAGAGATACCATCGAGAGCTGGAGAGAG

GGGCTTAGCCCCTCGGGCCGTGCAACACCGAACCCTGATACGTCCGAGGGAGATCATCAGAATATCAACCAATCATGCTC

ACCAGCAATCGGACCAAACAAAGTCTACTTGTCTCCTGAAGATAATCTCGGATTTAGAGAGATCACTGGCAACGACTGTG

AGGCTGGGCTCGGAGGGGTCCAGAGAGAAGGATCCAACTCTCAAGTACAGCGTTACCATGTTTATAGTCACGGGGGTGAA

GAGATTGAAGGACTCGAGGATGCTGACTCTCTCGTGGTTCAAGCAGATCCTCCAGTTGCTAACGTCTTCAATGGAGGAGA

GGATGGATCTGACGACAGCGATGTGGACTCTGGCCCAGATGATCCCGGCAGAGATACTCTATATGACCGGGGATCTGTTG

CCGGCAATGGTATCGCTAGGTCCACAGATGTCGAAAAACTAGAAGGTGCTGATATTCAAGAAGTTCTTAACTCCCAGAAA

GGCAAAGGAGGAAGATTCCAGGGCGGGAAAACCTTGCGAGTCCCGGAAATACCCGATGTCAAGCACTCCAGACCATCAGC

CCAATCAATTAAAAAGGGCACAGACGGGAACTCAGTCTCATCTGGAACGGTGATAGAGTGTTTATCGATAAGTGGTGCAA

CCCAAACTGTGCCAGAGTCAAGATGGGAGTCATCAGAGCAAAATGCGTCTGTGGGGAGTGTCCTCAAGTCTGCGAGGAGT

GCAAAGACGATCCAGGGGTCGACACAAGAATCTGGTACCATAGCATCACTGACTCAGCCTAAAGAGAATGACTCCGAGTA

TGAGTATGAGGATGACCTATTTACAGAGATTCAGGACATCCGTGCAAGCATTGCCAAGATCCATGATGACAATAAAACTA

TCCTCTCAAAGCTTGATTCTATACTGTTATTGAAAGGGGAAGTCGACACTATCAAGAAACAAATCAGCAAGCAGAATATA

AGTATATCCACCATTGAGGGCCATCTCTCCAGTATAATGATAGCCATCCCTGGCTTTGGGAAGGACATCAAGGACCCAAC

ATCCGAAGTCGAGTTGAACCCAGATCTAAGACCTATAATAAGCCGTGACTCTGGCAGAGCTCTCGCGGAGGTCCTCAAGA

AACCCGCTGTGGATAGGTCTCAGAAAATTGGAACCAAAGCCAACTCCAGCTCAAAGGGTCAGCTTCTTAAGGATCTCCAG

CTAAAACCTGTCGACAAGCAGGCAAGCTCTGCAATCGGGTTTGTCCCGTCCGACCATGAATCATCCAGAAATGTCATCCG

CTCCATAATCAAGTCGAGCAAGCTAAATATTGATCACAAGGACTATCTTCTAGATTTACTAAATGATGTGAAAGGCTCCA

AGGATCTTAAGGAATTTCACAAGATGCTAACAGCAATTCTCGCCAAGCACCCGTAACACATCCTCCAGTCATCATCTCAT

ACTCGACTAAAAACATCCTTTCAATCAGGCTATTACAAAAAACTTAGGAGCAAGGGCAACTGAGCTTCGCAGACAGGATG

ACCGAGATCTACGACTTCGATAAATCAGCATGGGATGTCAAAGGGTCAATTGCCCGCATAGAACCCACCACCTACCACGA

CGGCCGACTGATACCCCAGGTGAGGGTCATCGATCCTGGTCTGGGAGACAGAAAAGATGAGTGCTTTATGTACCTGTTTC

TCCTAGGAGTGATTGAGGATAACGACCCCCTGTCTCCCCCCGTCGGAAGAACCTTTGGCTCTCTACCTCTAGGGGTTGGT

AGGTCAACTGCCAAGCCAGAAGAACTACTAAGGGAGGCCACAGAATTAGATATAGTGGTGAGGCGCACTGCAGGAGTAAA

TGAGAAACTGGTATTTTACAACAACACTCCGCTGTCCTTGTTAACGCCCTGGAAGAAAGTTCTGACAACCGGAAGTGTGT

TTAGCGCTAACCAGGTTTGCAATGCAGTCAACCTAGTCCCGCTTGATACTCCCCAGAGATTCAGGGTTGTGTACATGAGC

ATAACTAGATTATCAGACAATGGTTACTATAGTGTGCCCAGAAGAATGCTGGAGTTCCGCTCGGCCAATGCAGTAGCCTT

CAACATCTTGGTTACACTAAGAATTGAAAATGGCACAAACCCTAGAAGATACATAGTCGGCTCATGGGAGAATTCAGAGG

TCACATTTATGGTACACGTGGGCAACTTTAGGAGAAAGAAGAACGAAGTATACTCTGCTGATTACTGCAAAATGAAGATT

GAAAAGATGGGTTTAGTTTTTGCCCTGGGTGGAATAGGCGGAACAAGTCTCCATATTAGAAGCACCGGGAAAATGAGCAA

AACCCTCCATGCACAGCTGGGGTTCAAGAAAATCTTATGTTACCCCCTAATGGATGTCAATGAGGATCTTAACCGATATC

TCTGGCGGGCAGAGTGCCGAATAGTCAAAATCCAAGCTGTCTTACAGCCATCAGTACCCCAAGAGTTCCGTGTCTACGAT

GATGTCATCATCAACGATGATCAAGGCTTGTTCAAGATCTTGTAGTTCATTTGCAAAGGAACGAACCCACCCCCAGGAAG

GGGCCCCACCCTCCCCCG-CCAAAAAACACACTCCAAGGCTCCCCGGCACAGCGCCAACACAGCTACTCCCGCGATGGAA

CGAGGTCGGCC-CCCGAACAAATCCGGTCCAGGGGGGGGGGCCCCCCCCCTGACCAACCGGGCGCCGACCGAGCAGGCCG

GCGAGAAGCCCCCAAGAGGCCCCGACGAAGACCGAGCACCCCATCCCCCC-AAAAACGTCCCCC-AACACGCCCGAGGCT

AGCCACCGGACACCCCCAGACCCGCCACCGGCGAGACAGAACCCCAAGGGGGGACCCTCCAAACCAAAGGAATCACCGGG

GTAGGCTCCGCCGAACCAGCACAGACCCGACCCAAGCAAGGCGGACCCCCAACACGAAACTGCCCAGCACACCCTACTCA

TCATCAAACAAAACTTAGGGGCCAAGTCCATACGAGAGGCCCTAGTCCGTCAGGCGGAGGGACGCAGAAAGGAAGGAGAC

ACGACCGTCCCAGAAGGAGGACACCCAACCACCGAAACGCCCGCCCCCAGGGAAGGAG--CCA-CCCCCACGCAGCGAAG

GAGCGGGTGGGAGACCTCGGGGAGGGCCCGACCGGGACCCGGGGGCCGGCGGCGCCGGCGGTGAGCAGGCCGGGCGCATA

CCTGACCCCACCGGAGGATCCGAGGAGGGCCCGACCCCGCGTGGACCAAACGGCAGGGAGCCCAACCCCACCAAAAAGAG

AGGCGAGGCCCCCCCTCCCTCCCCCCCCGCACACAACCCGCAGACAGACCGAGGGAGGGCCATCCCCCAGCGAGACAGGC

GGCTCCCTCGCAGCCGAGCCGCAGCTCCCCAGAATAGGACACCCCCGCACACTCCGAACACCCCCTCGCCCCCCGCCACC

GGGACAGGGGCTCCACCACCACAGACCGGAGCCACCGACAGCCAGCCAGAGGTCCCCTCCAGTCCCGCCCGGCGCCCCCC

CGAAGAGGCCCCGGCAAGGCCCCCCAGCCCGACGTCCCCCGCACGCACCGAAAGAAGGACCCGCCCGAGAACGTGCGGCA

CATCCATGCGCAAACATCATGACGCGGGTCGCAATTTTGACATTTCTGTTTCTTTTCCCAAATGTTGTTGCGTGTCAGAT

TCACTGGGGCAATCTATCCAAGATCGGGATTGTAGGAACAGGGAGTGCCAGCTACAAGGTGATGACTAGGCCAAGCCACC

AGACTCTGGTTATAAAGTTAATGCCAAATATAACGGCCATCGACAATTGTACAAAGTCAGAGATTGCAGAGTACAAGAGA

TTGCTGATCACAGTGTTAAAGCCTGTAGAGGATGCTCTGTCGGTGATAACCAAGAATGTAAGACCAATTCAAACTCTAAC

ACCTGGGCGTAGAACCCGCCGTTTTGCTGGAGCTGTTCTGGCCGGGGTAGCACTTGGAGTTGCGACAGCCGCTCAGATAA

CTGCAGGAGTCGCCCTTCATCAATCATTGATGAACTCCCAAGCAATTGAGAGTTTAAAAACCAGTCTTGAGATGTCGAAT

CAGGCAATAGAAGAAATCAGACTTGCAAATAAGGAGACCATACTGGCAGTACAGGGCGTCCAGGATTATATCAACAATGA

GCTCGTCCCTTCTGTTCATAGAATGTCATGCGAGCTGGTAGGTCACAAGCTCGGCCTCAAGCTCCTTAGGTACTACACCG

AGATCCTGTCCATATTCGGGCCCAGTCTTCGAGACCCGATATCTGCCGAAATATCAATCCAGGCACTTAGTTATGCATTA

GGCGGAGACATTAATAAAATCCTGGACAAGCTTGGGTATAGCGGTGGGGATTTCCTTGCCATCCTAGAAAGCAAGGGAAT

AAAGGCCCGGGTTACATATGTGGACACAAGAGATTACTTTATAATCCTTAGCATCGCCTACCCAACCTTATCTGAGATCA

AGGGAGTGATAGTTCACAAGATAGAAGCTATAACATACAACATTGGGGCACAGGAGTGGTATACTACTATCCCTAAATAT

GTAGCCACTCAGGGGTATCTGATATCGAACTTTGATGAGACGTCATGCGTATTCACTCCAGAGGGGACAGTTTGCAGCCA

GAATGCGTTGTACCCAATGAGCCCATTGCTTCAGGAATGTTTCAGGGGGTCAACAAAATCGTGCGCCAGAACCTTAGTTT

CAGGGACCATAAGTAATAGATTTATCCTATCAAAAGGGAACCTGATTGCAAATTGTGCGTCAGTTTTGTGCAAATGTTAC

ACAACAGAGACAGTTATCAGCCAAGATCCTGACAAACTACTAACTGTTGTAGCATCCGACAAGTGTCCTGTAGTTGAGGT

GGATGGAGTGACAATACAGGTCGGCAGTCGAGAGTATCCGGATTCTGTATACTTACACAAAATAGACTTAGGTCCAGCCA

TCTCCCTAGAAAAACTGGATGTAGGCACCAATTTAGGCAATGCAGTCACAAGACTGGAGAATGCAAAGGAGCTCCTAGAT

GCATCAGACCAAATACTGAAGACTGTTAAAGGGGTACCTTTCAGTGGGAATATGTACATAGCACTGGCAGCTTGCATAGG

AGTATCCCTGGGCCTTGTCACATTAATATGCTGCTGTAAGGGGAGGTGTAAGAACAAGGAAATCCCTATCTCCAAAATCA

ACCCAGGGCTCAAACCCGACCTGACCGGGACCTCAAAGTCGTACGTAAGATCACTGTAGTCAGAATCACCTGGATCATCT

GGCATCACACACATACATGCACGACACAGGCAGTCCGAGGACGCAAGAAACCCAGCCTCCGGTCACCCACCCGACCCCAC

TCCACGCTCCACCACACATTAGTCATCAAACAAAACTTAGGACGAAAGGTCAATCACCATGTCCGCACAAAGGGAGAGGA

TCAATGCCTTCTACAAAGACAATCCTCACAATAAGAACCATAGGGTGATCCTGGATAGAGAACGCTTGGTCATTGAAAGA

CCCTACATCTTGCTTGGAGTCCTGCTGGTAATGTTCCTGAGTCTAATCGGACTGCTGGCCATTGCAGGGATCAGGCTTCA

CCGGGCCACCGTTGGAACTTCAGAGATCCAGAGTCGGCTGAATACCAATATTAAGTTGACCGAATCTATTGATCACCAGA

CTAAGGATGTCTTAACTCCCCTTTTTAAAATCATTGGCGATGAAGTCGGCATCAGAATTCCACAGAAATTCAGTGATCTT

GTCAAGTTCATCTCCGATAAGATTAAATTCCTCAACCCTGATAGAGAGTATGATTTCAGAGATCTCCGGTGGTGTATGAA

TCCCCCCGAGAGAGTCAAAATTAATTTTGATCAGTTTTGTGAGTACAAGGCTGCGGTTAAGTCAATTGAACATATATTTG

AGTCACCACTCAACAAGTCAAAAAAGCTGCAATCTTTGACTCTCGGGCCCGGAACAGGCTGTCTAGGCAGGACAGTAACA

AGAGCCCATTTCTCAGAACTTACAATGACCTTAATGGACCTGGATCTAGAGATGAAGCACAACGTGTCCTCAGTGTTTAC

CGTAGTTGAAGAGGGATTATTCGGAAGAACATATACCGTCTGGAGATCCGATGCCAGGGATCCGAGCACCGATCTAGGTA

TCGGCCATTTTTTAAGAGTCTTCGAGATTGGACTGGTAAGAGATCTTGGGCTGGGTCCCCCTGTTTTTCATATGACCAAC

TATCTCACGGTGAACATGAGTGATGACTATCGGAGATGTCTTTTAGCGGTAGGGGAGTTGAAGTTGACAGCCCTATGCAC

CTCATCTGAGACTGTGACACTGAGTGAGAGAGGAGTTCCAAGGAGGGAACCTCTTGTGGTTGTGATACTTAATCTAGCTG

GACCCACTCTAGGGGGAGAGCTATACAGTGTCTTGCCTACCTCTGATCTCATGGTGGAGAAACTCTATTTGTCTTCACAT

AGAGGGATCATCAAAGATGATGAGGCCAATTGGGTAGTGCCGTCTACCGATGTTCGTGATCTTCAGAATAAAGGTGAATG

TCTGGTGGAAGCATGCAAGACTCGACCTCCTTCATTTTGCAATGGCACAGGATCAGGCCCGTGGTCAGAAGGGAGAATCC

CTGCCTACGGGGTGATCAGGGTCAGTCTTGACTTAGCGAGTGACCCGGATGTAGTTATCACTTCAGTGTTTGGCCCACTG

ATACCTCACCTATCCGGCATGGATCTTTACAACAACCCGTTTTCAAAAGCTGTATGGTTGGCTGTACCACCTTATGAGCA

GTCATTTCTAGGAATGATAAATACAATTGGATTCCCTAACAGAGCAGAGGTTATGCCGCACATTTTGACCACAGAGATCA

GAGGCCCTCGGGGTCGTTGCCATGTTCCCATAGAATTGTCCCGCCGGGTTGATGACGATATCAAGATCGGGTCCAACATG

GTCATATTGCCGACGATGGACCTGAGGTATATTACAGCCACTTATGATGTTTCCAGGAGCGAGCATGCAATCGTGTACTA

TATCTATGACACGAGTCGCTCATCATCTTACTTCTACCCAGTTCGACTGAATTTCAAAGGCAATCCTCTCTCTTTGAGGA

TAGAGTGTTTCCCCTGGCGTCATAAGGTGTGGTGCTACCATGATTGTCTTATATACAACACCATAACAGGTGAAGAGGTC

CATACGAGAGGGCTGACCGGCATAGAGGTAACATGCAATCCAGTCTGAGCAGAGCYACGACCGTCACTCAGGCAGTCCCT

TGAGTCGCCACCGAGTCCAAGCAGCACAGCCTGGGACAATCAACGGCACAGCCCAGCCAACAATGTTATAAAAAACTTAG

GAGCCAGGGTTGTAGGGGCCATGGACTCTCTATCAGTCAATCAGGTCTTGTACCCTGAGGTTCATCTAGACAGCCCTATT

GTCACAAACAAACTAGTTGCCATCCTTGAGTACTCGGGTATCGACCATAACTATGTTCTTGAAGACCAGACTCTTGTCAA

GAATATTAGGTATAGGCTGGGGTGCGGTTTTTCAAATCAAATGATCATCAATAACAGGGGGGTGGGTGAAACAGTCAATT

CTAAACTTAAAAGTTACCCCCATAATCGTCATATCATATACCCGGATTGCAATAAGGAGTTGTTTTGTATCAAGGATAGC

TGCATATCTAAGAAGCTCTCGGAGCTATTCAAGAAGGGTAATTCCTTGTACTCTAAGATAAGTCACCAGGTACTGGATTG

TCTTAAGAGAGTCAATGGGAAATTAGGTCTGGGCACAGATCTTACCCATGGTCTGAAGGAGGGTATCCTTGACCTAGGGT

TGCACATGCATAGCTCTCAATGGTTCGAGACTTTCCTGTTCTGGTTCACTATCAAGACAGAGATGAGATCAATGATCAAA

GAACAGTCCCATATATGCCACAAGAGGAGGTATAACCCGACTTTTGTGTCGGGGGATGCATTCGAGGTGCTCGTATCGCG

AGACCTCGTTGTGATAATTGATAAGAATACCCAGTATGTCTTCTACTTAACATTTGAGCTGGTCCTTATGTATTGTGATG

TCATAGAGGGTAGACTTATGACGGAGACAGCCATGGCCATAGACCAGAGATATTCAGAACTTCTAAGCCGGGTCAGATAC

TTGTGGGATCTTATTGATGGGTTCTTCCCAACACTAGGCAACACCACATACCAAATTGTTGCTCTGCTTGAACCATTATC

GTTGGCTTATCTTCAACTTCAGGATGTCACTCTGGAGTTAAGAGGTGCTTTTTTAGACCATTGCTTCAAAGAACTCTATG

AGATACTGGAACATTGCGGCATTGACACAGCAGGCACTTACAATTCCATCACTGAAGGATTGGATTACGTATTCATCACC

CACGATATACATCTAACTGGGGAGATTTTTTCATTTTTCCGGAGTTTCGGACACCCCCGTCTAGAAGCAGTCACCGCTGC

GGAAAATGTCAGAAAACATATGAACCAACCGAAGGTAATCAGTTATGAGACTATGATGAAAGGCCATGCGGTATTTTGCG

GGATAATCATAAATGGTTTTCGGGACCGACATGGCGGCAGCTGGCCCCCTGTTGCACTGCCAGAACATGCCTCTGCTGCG

ATCCGGAATGCGCAGGCATCAGGTGAGGGACTGACCCATGACCTGTGTATAGACAACTGGAAATCCTTTGTTGGATTCAA

ATTTGGCTGCTTTATGCCACTCAGCCTAGATAGTGATTTGACCATGTATCTCAAAGACAAGGCATTGGCTGCACTGAAGA

ATGAGTGGGATTCAGTTTACCCGAAAGAATACCTCCGTTATAACCCACCTAGAGGGACAGAGTCAAGGCGATTGGTAGAG

GTATTCCTGAATGACTCCAGCTTTGATCCTTATAACATGATAATGTACGTGGTGAATGGCTCCTACCTTAAAGACCCTGA

GTTTAACCTCTCATACAGCCTAAAAGAGAAGGAGATAAAGGAGACAGGGCGGTTGTTTGCCAAAATGACCTATAAGATGC

GGGCCTGTCAGGTAATCGCTGAAAATCTGATATCGAATGGTGTTGGGAAGTATTTCCGAGACAATGGGATGGCAAAGGAC

GAGCATGACCTAACAAAAGCCCTTCACACCCTGGCAGTCTCAGGTGTTCCCAAAAACAACAAAGATAACCACCGAGGTGG

GCCTCCCAGAAGGACAACAAACCGAGGGGTGAGATCAAGCCAAGGCACCAAAACACAAGATAGAGACAAGGTTCAAGGGG

GACCTATGTACAACTATTTGCGATGCCAACCGATCAGCCCTGATCAGGGTGAGTCATACGAGACTGTTAGTGCATTCATC

ACCGCTGACCTTAAGAAGTATTGCCTGAATTGGAGATACGAGACAATCAGCATATTTGCACAGAGGCTGAATGAAATATA

TGGACTACCATCCTTCTTTCAATGGTTACACAGGATATTGGAAAAATCCGTACTCTACGTCAGTGACCCACATTGTCCTC

CCGATCTAGATAATCATATCCCTCTGGACAGTGTCCCCAATGCCCAAATATTCATCAAGTACCCAATGGGCGGAATAGAA

GGTTATTGCCAAAAACTATGGACAATCAGCACTATACCATACTTGTATCTGGCAGCCTATGAGAGCGGAGTAAGAATCGC

CTCACTGGTGCAGGGTGACAATCAGACAATCGCAGTGACAAAAAGAGTTCCAAGTTCTTGGCCTTATTCACTAAAAAAGA

GGGAGGCCTCCAAGGCCGCTCAAAATTACTTCGTAGTCTTAAGGCAAAGATTGCACGATGTAGGTCATCACTTAAAGGCT

AATGAGACCATAGTATCTTCTCACTTTTTTGTATATTCCAAAGGGATTTATTATGACGGCCTGCTAGTCTCACAATCACT

AAAGAGCATCGCCAGATGTGTCTTCTGGTCCGAGACTATAGTGGATGAAACCAGAGCGGCGTGCAGTAATATTGCAACAA

CTGTCGCTAAGAGTATAGAGAGGGGTTATGATAGGTACCTTGCATATTCTTTGAATATCCTCAAGATTTTCCAACAGATC

CTTATATCCCTTAACTTCACTATTAACACAACAATGACTCAGGATGTCGTGGCACCGATCATCGAGAACGGTGATTTGCT

GATAAGGATGGCACTCTTGCCAGCACCCATCGGGGGTCTCAATTATCTTAACATGAGCAGGTTATTTGTGAGAAATATCG

GTGACCCGGTCACCTCCTCTATAGCCGACCTGAAGAGGATGATAGATGCCGGGCTAATGCCAGAAGAAACATTGCATCAA

GTGATGACCCAGACCCCGGGAGAATCATCCTACCTAGATTGGGCAAGTGACCCTTATTCTGCCAACCTACCCTGCGTACA

GAGTATCACTCGCCTTCTCAAAAACATCACTGCACGATATATTTTAATCAGCAGCCCAAACCCGATGCTGAAAGGGTTAT

TTCATGAGGGGAGTAGAGATGAAGACGAGGAGCTTGCAAGTTTCCTAATGGATCGGCATATAATTGTTCCAAGAGCTGCA

CATGAAATCTTAGACCATAGTATAACCGGAGCAAGAGAAGCTATAGCCGGGATGTTGGACACCACCAAGGGTCTGATTAG

AACAAGTATGAAACGGGGCGGCCTCACCCCTCGAGTCTTAGCCCGCCTTTCCAATTATGATTATGAACAATTCCGATCCG

GGATAACACTATTAACAAAGAAAGGGCAGTGTTATCTCATTGACAAGGACTCGTGCTCGGTGCAGCTCGCTATAGCTCTG

AGGGGCCACATGTGGGCTAAGTTGGCTCGCGGGAGACCAATCTATGGGTTGGAGGTGCCTGATGTATTAGAATCGATGAA

CGGCTACCTTATCAAACGCCACGAGTCGTGTGCTATCTGTGAAACGGGCTCAAGTCACTACGGATGGTTTTTCGTCCCTG

CAGGGTGCCAGCTTGACGATGTATCTAGAGAGACTTCGGCTCTTCGTGTACCTTATGTCGGATCAACAACTGAGGAAAGG

ACAGACATGAAACTTGCATTTGTTAGGTCTCCAAGTAGATCCCTCAAATCAGCAGTTAGAATTGCAACAGTTTACTCATG

GGCTTACGGGGATGATGAGAAATCATGGAGTGAAGCCTGGATGCTAGCTAGGCAGAGAGCCAATATCACCTTAGATGAAT

TGAGAATGATCACTCCAGTCTCCACATCCACCAACTTAGCCCACCGGTTGAGGGATCGGAGTACCCAGGTGAAATACTCG

GGGACATCCCTTGTGAGGGTTGCAAGATACACAACCATCTCCAATGATAACTTGTCGTTTGTGATATCTGAGAAAAAAGT

GGATACCAACTTCATTTATCAGCAAGGGATGTTGCTTGGTCTTGGGATCCTTGAGAACCTCTTTAGGCTAGAGGCCACCA

CCGGAGTATCCAACACAGTGCTACACCTGCACGTGGAAACAGAATGTTGTGTTGTACCTATGGTGGATCACCCAAGGATA

CCGAGCCTCCGTAATATAAAAGTTACGGATGAGCTATGCACAAACCCTTTGATCTACGACAGGTCCCCCATCATAGAGCA

CGATGCAACCCGGCTATATTCACAGAGCCACAGGAGACATTTGGTGGAGTTTGTTACCTGGTCAACAAGCCAGCTTTACC

ATATACTGGCTAAGTCTACAGCAATGTCCATGATTGAATTGATCACGAGATTCGAGAAAGATCACATGAATGAAATAGCC

GCCCTGATTGGCGATGACGACATCAACAGCTTCATCACAGAATTTTTGCTAGTGGAGCCTAGATTATTTATAGTTTACCT

TGGTCAGTGTGCTGCCATCAATTGGGCCTTTGATATACATTATCATCGGCCCTCGGGCAAGTACCAGATGGGGGAACTTC

TCTACTCTCTGCTCTCTCGGATGAGCAAAGGAGTATATAAGATCTTTACTAATGCTTTGAGCCACCCCAAAGTTTACAAG

AAATTTTGGCGAAGTGGCATCATTGAGCCGATTCATGGCCCATCCTTGGATACACAGAATTTACATGTCACTGTCTGTGA

CATGATATATGGGTCCTATGTCACCTATTTGGATCTTTTGCTGAATGATGAGCTAGATGATTACCCGTATTTGCTCTGCG

AGAGTGATGAGGACGTGGTCACAGACAGGTTCGACAACATTCAAGCCAAACATCTCTGTGTATTGGCCGATGTATATTGC

AGCTCCAAGAGGTGTCCTTCAATAATCGGGATGTCTCCTATAGAAAAATGTACCATTCTCACACATTACATCAAGGGAGA

ATCAGTACAATCCCCGTCTGGGACCTCATGGAACACTGATCCCCTTGTAGTAGATCATTACTCATGCTCTCTGACCTACC

TTCGCCGCGGTTCCATCAAACAAATCAGGTTGAGGGTGGATCCTGGGTTTGTATTCGAGGCGTTGACAGACGTCGACTTT

AAACAACCTCGCAAAGCTAAGTTAGATATATCGGTCGTTGGGTTGACTGATTTCTCTCCCCCTTACGACAACGTCGGTGA

TTTTTTAGGGACTATCAACACATTGAGGCACGATCTGCCTGTCACCGGAACCGGGGTCTCGAACTATGAAGTCCACGCTT

ATCGTAGAATTGGCCTGAATTCATCAGCATGTTATAAAGCCGTAGAGATCTCCACGTTAATCAAGCCATCCTTAGAAGTC

GGAGAGCATGGATTGTTCTTAGGAGAAGGTTCCGGTTCAATGCTGGCTGCGTACAAAGAAGTTCTTAAATTAGCAAATTG

TTACTACAACAGCGGAGTAACAGCGGAGGGCAGAGCCGGACAGAGGGAAATATCTCCCTATCCTTCAGAGATGAGCCTAG

TAGAGAATCAAATGGGGATAGAGCGGAGTGTTAAAGTGCTGTTCAACGGGAAACCTGAAGTAACTTGGGTGGGGACCACA

GATTGCTACAAGTACATAATCAGTAACATCCAAACCTCTAGTCTGGGTTTCATACATTCAGATATTGAGACGCTCCCAAC

CAAAGATGCTGTTGAGAAATTGGAGGAATTTGCCTCTATCCTATCCTTATCCTTGATTTTGGGGAAAATCGGCTCTATTA

CAGTTGTCAAAATTATGCCCATTAGCGGGGATTTTACCCAAGGCTTCATAGCCTATGCCATCCAATATTTTAGGGAGAGC

CTGCTTGCCTATCCGAGATATAGTAACTTCATCTCGACTGAGTGTTACCTTATTATGATAGGATTAAAGGCCAATCGATT

GATAAACCCAGAAGCCATTAAGCAAAGCATAATCAGAGCGGGGGTCAGGACTGCACCAGGACTTGTGAGCCATGTATTAT

CAGAGAAACAAAAAGGTTGTATTCAATCTTTCTTGGGTGATCCTTATATCCAAGGAGACTTCAATAAGCACCTTAAATCT

CTAACCTCTATTGAGAAAATCCTGGTAAATTGTGGTCTCTCGATCAATGGCATAAAAATCTGTAGAGATCTAATCCACCA

TGATATCGCCTCCGGTCCAGACGGCCTTATGAGCTCTACAATTATTTTATATAGGGAATTGGCTCATTTCAAAGATAATA

TAAGGAGTCAGCACGGAATGTTCCACCCCTACCCGGTACTGGCCAACAGCAGGCAACGTGAATTAATCCTTCGAATAGCC

AAGAAATTCTGGGGGTATGTCTTGCTATATTCTGATGACCCGGCACTAATCAGACAAACAATCAAGAACTTGAAACGGAA

TCATCTAACCTTTGACTTACATAGTAACCCATTTATTAAAGGGCTATCCAAGGCTGAGAAACTGCTGGTGCGGACGAGTT

CACTAAGAAGAGAATGGTTGTTCACTCTCGAAACGAAAGAAGTAAAAGAGTGGTTCAAATTGGTAGGTTACAGTGCACTC

ATCAGAGGCTAGTCAGCTATGCATCTGCCCCCTTCTCCTCCGCCATGAGACCCCACTGGCGATCCAGAAGATTAAAGAAA

ACTACATATTGGATAAGTATCTATTCCCAGCTTTGTCTGGT

>KX421387.1/goat/China/2013-12-29

ACCAAACAAAGTTGGGTAAGGATAGATCTTATAATAACTATGGACTGGCAAACTTAGGAGTAAAGATCCTACTGTCGGGG

GGAGGAGGAGGAGCAAGATCCTTGATCATGGCGACTCTCCTTAAAAGCTTAGCATTGTTCAAAAGGAACAAAGACAAAGC

GCCGACAGCATCAGGTTCAGGAGGGGCCATCCGGGGGATTAAGAATGTTATCATAGTCCCGATTCCCGGAGACTCGTCCA

TCACTACCCGTTCAAGACTGCTCGACAGGCTTGTCAGATTGGCCGGAGATCCTGACATCAACGGGTCAAAGCTGACCGGG

GTGATGATCAGCATGTTATCCTTGTTCGTAGAGTCACCCGGGCAATTGATACAGCGAATCACAGATGATCCGGATGTTAG

TATCCGCCTTGTTGAGGTAGTTCAAAGTACTAGGTCCCAGTCCGGGTTGACCTTTGCATCACGTGGTGCTGATTTAGACA

ACGAGGCAGACATGTATTTTTCAACTGAGGGGCCCTCGAGTGGAGGTAAGAAAAGGATCAACTGGTTTGAGAACAGAGAA

ATAATAGACATAGAGGTGCAGGATCCAGAAGAGTTCAATATGTTGTTAGCCTCCATACTAGCACAAGTCTGGATCCTTCT

GGCCAAGGCTGTTACGGCACCAGATACGGCAGCTGACTCAGAACTGAGAAGGTGGGTTAAATACACACAACAAAGGAGAG

TGATTGGGGAATTTCGCCTTGACAAGGGGTGGCTGGACGCAGTCCGCAACAGAATTGCAGAAGATCTATCACTCCGGCGG

TTCATGGTATCTCTCATACTTGACATCAAGAGGACCCCAGGCAACAAGCCAAGGATTGCAGAAATGATCTGCGACATTGA

CAACTATATTGTCGAAGCAGGACTCGCCAGTTTCATCCTTACTATCAAATTTGGTATCGAAACCATGTATCCTGCATTAG

GGCTCCACGAGTTTGCCGGAGAATTGTCCACTATTGAGTCCTTGATGAACTTGTATCAACAGCTAGGAGAGGTTGCACCC

TACATGGTAATTCTAGAGAATTCAATTCAGAACAAGTTTAGTGCAGGAGCTTATCCCCTCCTCTGGAGCTATGCGATGGG

TGTCGGAGTCGAGCTGGAGAACTCAATGGGGGGCTTGAACTTTGGCAGGTCATATTTTGACCCTGCCTATTTTCGTCTCG

GACAGGAGATGGTCAGAAGATCTGCAGGAAAGGTCAGCTCTGTGATTGCGGCTGAGCTCGGCATCACAGCAGAGGAAGCT

AAACTAGTCTCGGAAATCGCCTCGCAGGCTGGGGACGAAAGAACCGCCAGAGGGACTGGGCCTCGACAGGCGCAGGTTTC

CTTCCTTCAGCACCAAACAGGAGGGGGAGAGTCGTCCGCACCAGCGACCAGAGAAGGGGTCAAAGCTGCGATCCCAAACG

GAGCTGAAGAAAGGGACAGAAAGCAAACACGCCCAGGAAGGCCCAGAGGAGAGACCTCCGGCCAACTGCTCCTGGACATC

ATGCCAGAGGATGAGATCTTGCGAGAGTCTGGTCAAAACCCTCGTGAGGCTCAAAGATCGGCCGAGGCACTCTTCAGGCT

GCAGGCCATGGCCAAGATTCTGGAGGACCAGGAGGAGGGAGAAGACAACAGTCAGGTCTACAACGACAAGGATCTCCTCG

GCTAAGCAGACGCACCCTCTGTCGAAATCAGTGACAAGACATCTCCTACCAGTATTATAAAAAACTTAGGACCCAGGTCC

AAGCAAGCACACATCGACACTCCAACCAGTCGAGCGGAGACCACCGATGGCAGAAGAACAAGCATACCATGTCAACAAGG

GGCTGGAATGTATCAAGTCTCTCAAAGCCTCACCCCCGGATCTATCCACCATCAGAGATACCATCGAGAGCTGGAGAGAG

GGGCTTAGCCCCTCGGGCCGTGCAACACCGAACCCTGATACGTCCGAGGGAGATCATCAGAATATCAACCAATCATGCTC

ACCAGCAATCGGACCAAACAAAGTCTACTTGTCTCCTGAAGATAATCTCGGATTTAGAGAGATCACTGGCAACGACTGTG

AGGCTGGGCTCGGAGGGGTCCAGAGAGAAGGATCCAACTCTCAAGTACAGCGTTACCATGTTTATAGTCACGGGGGTGAA

GAGATTGAAGGACTCGAGGATGCTGACTCTCTCGTGGTTCAAGCAGATCCTCCAGTTGCTAACGTCTTCAATGGAGGAGA

GGATGGATCTGACGACAGCGATGTGGACTCTGGCCCAGATGATCCCGGCAGAGATACTCTATATGACCGGGGATCTGTTG

CCGGCAATGGTATCGCTAGGTCCACAGATGTCGAAAAACTAGAAGGTGCTGATATTCAAGAAGTTCTTAACTCCCAGAAA

GGCAAAGGAGGAAGATTCCAGGGCGGGAAAACCTTGCGAGTCCCGGAAATACCCGATGTCAAGCACTCCAGACCATCAGC

CCAATCAATTAAAAAGGGCACAGACGGGAACTCAGTCTCATCTGGAACGGTGATAGAGTGTTTATCGATAAGTGGTGCAA

CCCAAACTGTGCCAGAGTCAAGATGGGAGTCATCAGAGCAAAATGCGTCTGTGGGGAGTGTCCTCAAGTCTGCGAGGAGT

GCAAAGACGATCCAGGGGTCGACACAAGAATCTGGTACCATAGCATCACTGACTCAGCCTAAAGAGAATGACTCCGAGTA

TGAGTATGAGGATGACCTATTTACAGAGATTCAGGACATCCGTGCAAGCATTGCCAAGATCCATGATGACAATAAAACTA

TCCTCTCAAAGCTTGATTCTATACTGTTATTGAAAGGGGAAGTCGACACTATCAAGAAACAAATCAGCAAGCAGAATATA

AGTATATCCACCATTGAGGGCCATCTCTCCAGTATAATGATAGCCATCCCTGGCTTTGGGAAGGACATCAAGGACCCAAC

ATCCGAAGTCGAGTTGAACCCAGATCTAAGACCTATAATAAGCCGTGACTCTGGCAGAGCTCTCGCGGAGGTCCTCAAGA

AACCCGCTGTGGATAGGTCTCAGAAAATTGGAACCAAAGCCAACTCCAGCTCAAAGGGTCAGCTTCTTAAGGATCTCCAG

CTAAAACCTGTCGACAAGCAGGCAAGCTCTGCAATCGGGTTTGTCCCGTCCGACCATGAATCATCCAGAAATGTCATCCG

CTCCATAATCAAGTCGAGCAAGCTAAATATTGATCACAAGGACTATCTTCTAGATTTACTAAATGATGTGAAAGGCTCCA

AGGATCTTAAGGAATTTCACAAGATGCTAACAGCAATTCTCGCCAAGCACCCGTAACACATCCTCCAGTCATCATCTCAT

ACTCGACTAAAAACATCCTTTCAATCAGGCTATTACAAAAAACTTAGGAGCAAGGGCAACTGAGCTTCGCAGACAGGATG

ACCGAGATCTACGACTTCGATAAATCAGCATGGGATGTCAAAGGGTCAATTGCCCGCATAGAACCCACCACCTACCACGA

CGGCCGACTGATACCCCAGGTGAGGGTCATCGATCCTGGTCTGGGAGACAGAAAAGATGAGTGCTTTATGTACCTGTTTC

TCCTAGGAGTGATTGAGGATAACGACCCCCTGTCTCCCCCCGTCGGAAGAACCTTTGGCTCTCTACCTCTAGGGGTTGGT

AGGTCAACTGCCAAGCCAGAAGAACTACTAAGGGAGGCCACAGAATTAGATATAGTGGTGAGGCGCACTGCAGGAGTAAA

TGAGAAACTGGTATTTTACAACAACACTCCGCTGTCCTTGTTAACGCCCTGGAAGAAAGTTCTGACAACCGGAAGTGTGT

TTAGCGCTAACCAGGTTTGCAATGCAGTCAACCTAGTCCCGCTTGATACTCCCCAGAGATTCAGGGTTGTGTACATGAGC

ATAACTAGATTATCAGACAATGGTTACTATAGTGTGCCCAGAAGAATGCTGGAGTTCCGCTCGGCCAATGCAGTAGCCTT

CAACATCTTGGTTACACTAAGAATTGAAAATGGCACAAACCCTAGAAGATACATAGTCGGCTCATGGGAGAATTCAGAGG

TCACATTTATGGTACACGTGGGCAACTTTAGGAGAAAGAAGAACGAAGTATACTCTGCTGATTACTGCAAAATGAAGATT

GAAAAGATGGGTTTAGTTTTTGCCCTGGGTGGAATAGGCGGAACAAGTCTCCATATTAGAAGCACCGGGAAAATGAGCAA

AACCCTCCATGCACAGCTGGGGTTCAAGAAAATCTTATGTTACCCCCTAATGGATGTCAATGAGGATCTTAACCGATATC

TCTGGCGGGCAGAGTGCCGAATAGTCAAAATCCAAGCTGTCTTACAGCCATCAGTACCCCAAGAGTTCCGTGTCTACGAT

GATGTCATCATCAACGATGATCAAGGCTTGTTCAAGATCTTGTAGTTCATTTGCAAAGGAACGAACCCACCCCCAGGAAG

GGGCCCCACCCTCCCCCG-CCAAAAAACACACTCCAAGGCTCCCCGGCACAGCGCCAACACAGCTACTCCCGCGATGGAA

CGAGGTCGGCC-CCCGAACAAATCCGGTCCAGGGGGGGGGGCCCCCCCCCTGACCAACCGGGCGCCGACCGAGCAGGCCG

GCGAGAAGCCCCCAAGAGGCCCCGACGAAGACCGAGCACCCCATCCCCCC-AAAAACGTCCCCC-AACACGCCCGAGGCT

AGCCACCGGACACCCCCAGACCCGCCACCGGCGAGACAGAACCCCAAGGGGGGACCCTCCAAACCAAAGGAATCACCGGG

GTAGGCTCCGCCGAACCAGCACAGACCCGACCCAAGCAAGGCGGACCCCCAACACGAAACTGCCCAGCACACCCTACTCA

TCATCAAACAAAACTTAGGGGCCAAGTCCATACGAGAGGCCCTAGTCCGTCAGGCGGAGGGACGCAGAAAGGAAGGAGAC

ACGACCGTCCCAGAAGGAGGACACCCAACCACCGAAACGCCCGCCCCCAGGGAAGGAG--CCA-CCCCCACGCAGCGAAG

GAGCGGGTGGGAGACCTCGGGGAGGGCCCGACCGGGACCCGGGGGCCGGCGGCGCCGGCGGTGAGCAGGCCGGGCGCATA

CCTGACCCCACCGGAGGATCCGAGGAGGGCCCGACCCCGCGTGGACCAAACGGCAGGGAGCCCAACCCCACCAAAAAGAG

AGGCGAGGCCCCCCCTCCCTCCCCCCCCGCACACAACCCGCAGACAGACCGAGGGAGGGCCATCCCCCAGCGAGACAGGC

GGCTCCCTCGCAGCCGAGCCGCAGCTCCCCAGAATAGGACACCCCCGCACACTCCGAACACCCCCTCGCCCCCCGCCACC

GGGACAGGGGCTCCACCACCACAGACCGGAGTCACCGACAGCCAGCCAGAGGTCCCCTCCAGTCCCGCCCGGCGCCCCCC

CGAAGAGGCCCCGGCAAGGCCCCCCAGCCCGACGTCCCCCGCACGCACCGAAAGAAGGACCCGCCCGAGAACGTGCGGCA

CATCCATGCGCAAACATCATGACGCGGGTCGCAATTTTGACATTTCTGTTTCTTTTCCCAAATGTTGTTGTGTGTCAGAT

TCACTGGGGCAATCTATCCAAGATCGGGATTGTAGGAACAGGGAGTGCCAGCTACAAGGTGATGACTAGGCCAAGCCACC

AGACTCTGGTTATAAAGTTAATGCCAAATATAACGGCCATCGACAATTGTACAAAGTCAGAGATTGCAGAGTACAAGAGA

TTGCTGATCACAGTGTTAAAGCCTGTAGAGGATGCTCTGTCGGTGATAACCAAGAATGTAAGACCAATTCAAACTCTAAC

ACCTGGGCGTAGAACCCGCCGTTTTGCTGGAGCTGTTCTGGCCGGGGTAGCACTTGGAGTTGCGACAGCCGCTCAGATAA

CTGCAGGAGTCGCCCTTCATCAATCATTGATGAACTCCCAAGCAATTGAGAGTTTAAAAACCAGTCTTGAGATGTCGAAT

CAGGCAATAGAAGAAATCAGACTTGCAAATAAGGAGACCATACTGGCAGTACAGGGCGTCCAGGATTATATCAACAATGA

GCTCGTCCCTTCTGTTCATAGAATGTCATGCGAGCTGGTAGGTCACAAGCTCGGCCTCAAGCTCCTTAGGTACTACACCG

AGATCCTGTCCATATTCGGGCCCAGTCTTCGAGACCCGATATCTGCCGAAATATCAATCCAGGCACTTAGTTATGCATTA

GGCGGAGACATTAATAAAATCCTGGACAAGCTTGGGTATAGCGGTGGGGATTTCCTTGCCATCCTAGAAAGCAAGGGAAT

AAAGGCCCGGGTTACATATGTGGACACAAGAGATTACTTTATAATCCTTAGCATCGCCTACCCAACCTTATCTGAGATCA

AGGGAGTGATAGTTCACAAGATAGAAGCTATAACATACAACATTGGGGCACAGGAGTGGTATACTACTATCCCTAAATAT

GTAGCCACTCAGGGGTATCTGATATCGAACTTTGATGAGACGTCATGCGTATTCACTCCAGAGGGGACAGTTTGCAGCCA

GAATGCGTTGTACCCAATGAGCCCATTGCTTCAGGAATGTTTCAGGGGGTCAACAAAATCGTGCGCCAGAACCTTAGTTT

CAGGGACCATAAGTAATAGATTTATCCTATCAAAAGGGAACCTGATTGCAAATTGTGCGTCAGTTTTGTGCAAATGTTAC

ACAACAGAGACAGTTATCAGCCAAGATCCTGACAAACTACTAACTGTTGTAGCATCCGACAAGTGTCCTGTAGTTGAGGT

GGATGGAGTGACAATACAGGTCGGCAGTCGAGAGTATCCGGATTCTGTATACTTACACAAAATAGACTTAGGTCCAGCCA

TCTCCCTAGAAAAACTGGATGTAGGCACCAATTTAGGCAATGCAGTCACAAGACTGGAGAATGCAAAGGAGCTCCTAGAT

GCATCAGACCAAATACTGAAGACTGTTAAAGGGGTACCTTTCAGTGGGAATATGTACATAGCACTGGCAGCTTGCATAGG

AGTATCCCTGGGCCTTGTCACATTAATATGCTGCTGTAAGGGGAGGTGTAAGAACAAGGAAATCCCTATCTCCAAAATCA

ACCCAGGGCTCAAACCCGACCTGACCGGGACCTCAAAGTCGTACGTAAGATCACTGTAGTCAGAATCACCTGGATCATCT

GGCATCACACACATACATGCACGACACAGGCAGTCCGAGGACGCAAGAAACCCAGCCTCCGGTCACCCACCCGACCCCAC

TCCACGCTCCACCACACATTAGTCATCAAACAAAACTTAGGACGAAAGGTCAATCACCATGTCCGCACAAAGGGAGAGGA

TCAATGCCTTCTACAAAGACAATCCTCACAATAAGAACCATAGGGTGATCCTGGATAGAGAACGCTTGGTCATTGAAAGA

CCCTACATCTTGCTTGGAGTCCTGCTGGTAATGTTCCTGAGTCTAATCGGACTGCTGGCCATTGCAGGGATCAGGCTTCA

CCGGGCCACCGTTGGAACTTCAGAGATCCAGAGTCGGCTGAATACCAATATTAAGTTGACCGAATCTATTGATCACCAGA

CTAAGGATGTCTTAACTCCCCTTTTTAAAATCATTGGCGATGAAGTCGGCATCAGAATTCCACAGAAATTCAGTGATCTT

GTCAAGTTCATCTCCGATAAGATTAAATTCCTCAACCCTGATAGAGAGTATGATTTCAGAGATCTCCGGTGGTGTATGAA

TCCCCCCGAGAGAGTCAAAATTAATTTTGATCAGTTTTGTGAGTACAAGGCTGCGGTTAAGTCAATTGAACATATATTTG

AGTCACCACTCAACAAGTCAAAAAAGCTGCAATCTTTGACTCTCGGGCCCGGAACAGGCTGTCTAGGCAGGACAGTAACA

AGAGCCCATTTCTCAGAACTTACAATGACCTTAATGGACCTGGATCTAGAGATGAAGCACAACGTGTCCTCAGTGTTTAC

CGTAGTTGAAGAGGGATTATTCGGAAGAACATATACCGTCTGGAGATCCGATGCCAGGGATCCGAGCACCGATCTAGGTA

TCGGCCATTTTTTAAGAGTCTTCGAGATTGGACTGGTAAGAGATCTTGGGCTGGGTCCCCCTGTTTTTCATATGACCAAC

TATCTCACGGTGAACATGAGTGATGACTATCGGAGATGTCTTTTAGCGGTAGGGGAGTTGAAGTTGACAGCCCTATGCAC

CTCATCTGAGACTGTGACACTGAGTGAGAGAGGAGTTCCAAGGAGGGAACCTCTTGTGGTTGTGATACTTAATCTAGCTG

GACCCACTCTAGGGGGAGAGCTATACAGTGTCTTGCCTACCTCTGATCTCATGGTGGAGAAACTCTATTTGTCTTCACAT

AGAGGGATCATCAAAGATGATGAGGCCAATTGGGTAGTGCCGTCTACCGATGTTCGTGATCTTCAGAATAAAGGTGAATG

TCTGGTGGAAGCATGCAAGACTCGACCTCCTTCATTTTGCAATGGCACAGGATCAGGCCCGTGGTCAGAAGGGAGAATCC

CTGCCTACGGGGTGATCAGGGTCAGTCTTGACTTAGCGAGTGACCCGGATGTAGTTATCACTTCAGTGTTTGGCCCACTG

ATACCTCACCTATCCGGCATGGATCTTTACAACAACCCGTTTTCAAAAGCTGTATGGTTGGCTGTACCACCTTATGAGCA

GTCATTTCTAGGAATGATAAATACAATTGGATTCCCTAACAGAGCAGAGGTTATGCCGCACATTTTGACCACAGAGATCA

GAGGCCCTCGGGGTCGTTGCCATGTTCCCATAGAATTGTCCCGCCGGGTTGATGACGATATCAAGATCGGGTCCAACATG

GTCATATTGCCGACGATGGACCTGAGGTATATTACAGCCACTTATGATGTTTCCAGGAGCGAGCATGCAATCGTGTACTA

TATCTATGACACGAGTCGCTCATCATCTTACTTCTACCCAGTTCGACTGAATTTCAAAGGCAATCCTCTCTCTTTGAGGA

TAGAGTGTTTCCCCTGGCGTCATAAGGTGTGGTGCTACCATGATTGTCTTATATACAACACCATAACAGGTGAAGAGGTC

CATACGAGAGGGCTGACCGGCATAGAGGTAACATGCAATCCAGTCTGAGCAGAGCTACGACCGTCACTCAGGCAGTCCCT

TGAGTCGCCACCGAGTCCAAGCAGCACAGCCTGGGACAATCAACGGCACAGCCCAGCCAACAATGTTATAAAAAACTTAG

GAGCCAGGGTTGTAGGGGCCATGGACTCTCTATCAGTCAATCAGGTCTTGTACCCTGAGGTTCATCTAGACAGCCCTATT

GTCACAAACAAACTAGTTGCCATCCTTGAGTACTCGGGTATCGACCATAACTATGTTCTTGAAGACCAGACTCTTGTCAA

GAATATTAGGTATAGGCTGGGGTGCGGTTTTTCAAATCAAATGATCATCAATAACAGGGGGGTGGGTGAAACAGTCAATT

CTAAACTTAAAAGTTACCCCCATAATCGTCATATCATATACCCGGATTGCAATAAGGAGTTGTTTTGTATCAAGGATAGC

TGCATATCTAAGAAGCTCTCGGAGCTATTCAAGAAGGGTAATTCCTTGTACTCTAAGATAAGTCACCAGGTACTGGATTG

TCTTAAGAGAGTCAATGGGAAATTAGGTCTGGGCACAGATCTTACCCATGGTCTGAAGGAGGGTATCCTTGACCTAGGGT

TGCACATGCATAGCTCTCAATGGTTCGAGACTTTCCTGTTCTGGTTCACTATCAAGACAGAGATGAGATCAATGATCAAA

GAACAGTCCCATATATGCCACAAGAGGAGGTATAACCCGACTTTTGTGTCGGGGGATGCATTCGAGGTGCTCGTATCGCG

AGACCTCGTTGTGATAATTGATAAGAATACCCAGTATGTCTTCTACTTAACATTTGAGCTGGTCCTTATGTATTGTGATG

TCATAGAGGGTAGACTTATGACGGAGACAGCCATGGCCATAGACCAGAGATATTCAGAACTTCTAAGCCGGGTCAGATAC

TTGTGGGATCTTATTGATGGGTTCTTCCCAACACTAGGCAACACCACATACCAAATTGTTGCTCTGCTTGAACCATTATC

GTTGGCTTATCTTCAACTTCAGGATGTCACTCTGGAGTTAAGAGGTGCTTTTTTAGACCATTGCTTCAAAGAACTCTATG

AGATACTGGAACATTGCGGCATTGACACAGCAGGCACTTACAATTCCATCACTGAAGGATTGGATTACGTATTCATCACC

CACGATATACATCTAACTGGGGAGATTTTTTCATTTTTCCGGAGTTTCGGACACCCCCGTCTAGAAGCAGTCACCGCTGC

GGAAAATGTCAGAAAACATATGAACCAACCGAAGGTAATCAGTTATGAGACTATGATGAAAGGCCATGCGGTATTTTGCG

GGATAATCATAAATGGTTTTCGGGACCGACATGGCGGCAGCTGGCCCCCTGTTGCACTGCCAGAACATGCCTCTGCTGCG

ATCCGGAATGCGCAGGCATCAGGTGAGGGACTGACCCATGACCTGTGTATAGACAACTGGAAATCCTTTGTTGGATTCAA

ATTTGGCTGCTTTATGCCACTCAGCCTAGATAGTGATTTGACCATGTATCTCAAAGACAAGGCATTGGCTGCACTGAAGA

ATGAGTGGGATTCAGTTTACCCGAAAGAATACCTCCGTTATAACCCACCTAGAGGGACAGAGTCAAGGCGATTGGTAGAG

GTATTCCTGAATGACTCCAGCTTTGATCCTTATAACATGATAATGTACGTGGTGAATGGCTCCTACCTTAAAGACCCTGA

GTTTAACCTCTCATACAGCCTAAAAGAGAAGGAGATAAAGGAGACAGGGCGGTTGTTTGCCAAAATGACCTATAAGATGC

GGGCCTGTCAGGTAATCGCTGAAAATCTGATATCGAATGGTGTTGGGAAGTATTTCCGAGACAATGGGATGGCAAAGGAC

GAGCATGACCTAACAAAAGCCCTTCACACCCTGGCAGTCTCAGGTGTTCCCAAAAACAACAAAGATAACCACCGAGGTGG

GCCTCCCAGAAGGACAACAAACCGAGGGGTGAGATCAAGCCAAGGCACCAAAACACAAGATAGAGACAAGGTTCAAGGGG

GACCTATGTACAACTATTTGCGATGCCAACCGATCAGCCCTGATCAGGGTGAGTCATACGAGACTGTTAGTGCATTCATC

ACCGCTGACCTTAAGAAGTATTGCCTGAATTGGAGATACGAGACAATCAGCATATTTGCACAGAGGCTGAATGAAATATA

TGGACTACCATCCTTCTTTCAATGGTTACACAGGATATTGGAAAAATCCGTACTCTACGTCAGTGACCCACATTGTCCTC

CCGATCTAGATAATCATATCCCTCTGGACAGTGTCCCCAATGCCCAAATATTCATCAAGTACCCAATGGGCGGAATAGAA

GGTTATTGCCAAAAACTATGGACAATCAGCACTATACCATACTTGTATCTGGCAGCCTATGAGAGCGGAGTAAGAATCGC

CTCACTGGTGCAGGGTGACAATCAGACAATCGCAGTGACAAAAAGAGTTCCAAGTTCTTGGCCTTATTCACTAAAAAAGA

GGGAGGCCTCCAAGGCCGCTCAAAATTACTTCGTAGTCTTAAGGCAAAGATTGCACGATGTAGGTCATCACTTAAAGGCT

AATGAGACCATAGTATCTTCTCACTTTTTTGTATATTCCAAAGGGATTTATTATGACGGCCTGCTAGTCTCACAATCACT

AAAGAGCATCGCCAGATGTGTCTTCTGGTCCGAGACTATAGTGGATGAAACCAGAGCGGCGTGCAGTAATATTGCAACAA

CTGTCGCTAAGAGTATAGAGAGGGGTTATGATAGGTACCTTGCATATTCTTTGAATATCCTCAAGATTTTCCAACAGATC

CTTATATCCCTTAACTTCACTATTAACACAACAATGACTCAGGATGTCGTGGCACCGATCATCGAGAACGGTGATTTGCT

GATAAGGATGGCACTCTTGCCAGCACCCATCGGGGGTCTCAATTATCTTAACATGAGCAGGTTATTTGTGAGAAATATCG

GTGACCCGGTCACCTCCTCTATAGCCGACCTGAAGAGGATGATAGATGCCGGGCTAATGCCAGAAGAAACATTGCATCAA

GTGATGACCCAGACCCCGGGAGAATCATCCTACCTAGATTGGGCAAGTGACCCTTATTCTGCCAACCTACCCTGCGTACA

GAGTATCACTCGCCTTCTCAAAAACATCACTGCACGATATATTTTAATCAGCAGCCCAAACCCGATGCTGAAAGGGTTAT

TTCATGAGGGGAGTAGAGATGAAGACGAGGAGCTTGCAAGTTTCCTAATGGATCGGCATATAATTGTTCCAAGAGCTGCA

CATGAAATCTTAGACCATAGTATAACCGGAGCAAGAGAAGCTATAGCCGGGATGTTGGACACCACCAAGGGTCTGATTAG

AACAAGTATGAAACGGGGCGGCCTCACCCCTCGAGTCTTAGCCCGCCTTTCCAATTATGATTATGAACAATTCCGATCCG

GGATAACACTATTAACAAAGAAAGGGCAGTGTTATCTCATTGACAAGGACTCGTGCTCGGTGCAGCTCGCTATAGCTCTG

AGGGGCCACATGTGGGCTAAGTTGGCTCGCGGGAGACCAATCTATGGGTTGGAGGTGCCTGATGTATTAGAATCGATGAA

CGGCTACCTTATCAAACGCCACGAGTCGTGTGCTATCTGTGAAACGGGCTCAAGTCACTACGGATGGTTTTTCGTCCCTG

CAGGGTGCCAGCTTGACGATGTATCTAGAGAGACTTCGGCTCTTCGTGTACCTTATGTCGGATCAACAACTGAGGAAAGG

ACAGACATGAAACTTGCATTTGTTAGGTCTCCAAGTAGATCCCTCAAATCAGCAGTTAGAATTGCAACAGTTTACTCATG

GGCTTACGGGGATGATGAGAAATCATGGAGTGAAGCCTGGATGCTAGCTAGGCAGAGAGCCAATATCACCTTAGATGAAT

TGAGAATGATCACTCCAGTCTCCACATCCACCAACTTAGCCCACCGGTTGAGGGATCGGAGTACCCAGGTGAAATACTCG

GGGACATCCCTTGTGAGGGTTGCAAGATACACAACCATCTCCAATGATAACTTGTCGTTTGTGATATCTGAGAAAAAAGT

GGATACCAACTTCATTTATCAGCAAGGGATGTTGCTTGGTCTTGGGATCCTTGAGAACCTCTTTAGGCTAGAGGCCACCA

CCGGAGTATCCAACACAGTGCTACACCTGCACGTGGAAACAGAATGTTGTGTTGTACCTATGGTGGATCACCCAAGGATA

CCGAGTCTCCGTAATATAAAAGTTACGGATGAGCTATGCACAAACCCTTTGATCTACGACAGGTCCCCCATCATAGAGCA

CGATGCAACCCGGCTATATTCACAGAGCCACAGGAGACATTTGGTGGAGTTTGTTACCTGGTCAACAAGCCAGCTTTACC

ATATACTGGCTAAGTCTACAGCAATGTCCATGATTGAATTGATCACGAGATTCGAGAAAGATCACATGAATGAAATAGCC

GCCCTGATTGGCGATGACGACATCAACAGCTTCATCACAGAATTTTTGCTAGTGGAGCCTAGATTATTTATAGTTTACCT

TGGTCAGTGTGCTGCCATCAATTGGGCCTTTGATATACATTATCATCGGCCCTCGGGCAAGTACCAGATGGGGGAACTTC

TCTACTCTCTGCTCTCTCGGATGAGCAAAGGAGTATATAAGATCTTTACTAATGCTTTGAGCCACCCCAAAGTTTACAAG

AAATTTTGGCGAAGTGGCATCATTGAGCCGATTCATGGCCCATCCTTGGATACACAGAATTTACATGTCACTGTCTGTGA

CATGATATATGGGTCCTATGTCACCTATTTGGATCTTTTGCTGAATGATGAGCTAGATGATTACCCGTATTTGCTCTGCG

AGAGTGATGAGGACGTGGTCACAGACAGGTTCGACAACATTCAAGCCAAACATCTCTGTGTATTGGCCGATGTATATTGC

AGCTCCAAGAGGTGTCCTTCAATAATCGGGATGTCTCCTATAGAAAAATGTACCATTCTCACACATTACATCAAGGGAGA

ATCAGTACAATCCCCGTCTGGGACCTCATGGAACACTGATCCCCTTGTAGTAGATCATTACTCATGCTCTCTGACCTACC

TTCGCCGCGGTTCCATCAAACAAATCAGGTTGAGGGTGGATCCTGGGTTTGTATTCGAGGCGTTGACAGACGTCGACTTT

AAACAACCTCGCAAAGCTAAGTTAGATATATCGGTCGTTGGGTTGACTGATTTCTCTCCCCCTTACGACAACGTCGGTGA

TTTTTTAGGGACTATCAACACATTGAGGCACGATCTGCCTGTCACCGGAACCGGGGTCTCGAACTATGAAGTCCACGCTT

ATCGTAGAATTGGCCTGAATTCATCAGCATGTTATAAAGCCGTAGAGATCTCCACGTTAATCAAGCCATCCTTAGAAGTC

GGAGAGCATGGATTGTTCTTAGGAGAAGGTTCCGGTTCAATGCTGGCTGCGTACAAAGAAGTTCTTAAATTAGCAAATTG

TTACTACAACAGCGGAGTAACAGCGGAGGGCAGAGCCGGACAGAGGGAAATATCTCCCTATCCTTCAGAGATGAGCCTAG

TAGAGAATCAAATGGGGATAGAGCGGAGTGTTAAAGTGCTGTTCAACGGGAAACCTGAAGTAACTTGGGTGGGGACCACA

GATTGCTACAAGTACATAATCAGTAACATCCAAACCTCTAGTCTGGGTTTCATACATTCAGATATTGAGACGCTCCCAAC

CAAAGATGCTGTTGAGAAATTGGAGGAATTTGCCTCTATCCTATCCTTATCCTTGATTTTGGGGAAAATCGGCTCTATTA

CAGTTGTCAAAATTATGCCCATTAGCGGGGATTTTACCCAAGGCTTCATAGCCTATGCCATCCAATATTTTAGGGAGAGC

CTGCTTGCCTATCCGAGATATAGTAACTTCATCTCGACTGAGTGTTACCTTATTATGATAGGATTAAAGGCCAATCGATT

GATAAACCCAGAAGCCATTAAGCAAAGCATAATCAGAGCGGGGGTCAGGACTGCACCAGGACTTGTGAGCCATGTATTAT

CAGAGAAACAAAAAGGTTGTATTCAATCTTTCTTGGGTGATCCTTATATCCAAGGAGACTTCAATAAGCACCTTAAATCT

CTAACCTCTATTGAGAAAATCCTGGTAAATTGTGGTCTCTCGATCAATGGCATAAAAATCTGTAGAGATCTAATCCACCA

TGATATCGCCTCCGGTCCAGACGGCCTTATGAGCTCTACAATTATTTTATATAGGGAATTGGCTCATTTCAAAGATAATA

TAAGGAGTCAGCACGGAATGTTCCACCCCTACCCGGTACTGGCCAACAGCAGGCAACGTGAATTAATCCTTCGAATAGCC

AAGAAATTCTGGGGGTATGTCTTGCTATATTCTGATGACCCGGCACTAATCAGACAAACAATCAAGAACTTGAAACGGAA

TCATCTAACCTTTGACTTACATAGTAACCCATTTATTAAAGGGCTATCCAAGGCTGAGAAACTGCTGGTGCGGACGAGTT

CACTAAGAAGAGAATGGTTGTTCACTCTCGAAACGAAAGAAGTAAAAGAGTGGTTCAAATTGGTAGGTTACAGTGCACTC

ATCAGAGGCTAGTCAGCTATGCATCTGCCCCCTTCTCCTCCGCCATGAGACCCCACTGGCGATCCAGAAGATTAAAGAAA

ACTACATATTGGATAAGTATCTATTCCCAGCTTTGTCTGGT

>MF443341.1/goat/China/2014-03-17

ACCAAACAAAGTTGGGTAAGGATAGATCTTATAATAACTATGGACTGGCAAACTTAGGAGTAAAGATCCTACTGTCGGGG

GGAGGAGGAGGAGCAAGATCCTTGATCATGGCGACTCTCCTTAAAAGCTTAGCATTGTTCAAAAGGAACAAAGACAAAGC

GCCGACAGCATCAGGTTCAGGAGGGGCCATCCGGGGGATTAAGAATGTTATCATAGTCCCGATTCCCGGAGACTCGTCCA

TCACTACCCGTTCAAGACTGCTCGACAGGCTTGTCAGATTGGCCGGAGATCCTGACATCAACGGGTCAAAGCTGACCGGG

GTGATGATCAGCATGTTATCCTTGTTCGTAGAGTCACCCGGGCAATTGATACAGCGAATCACAGATGATCCGGATGTTAG

TATCCGCCTTGTTGAGGTAGTTCAAAGTACTAGGTCCCAGTCCGGGTTGACCTTTGCATCACGTGGTGCTGATTTAGACA

ACGAGGCAGACATGTATTTTTCAACTGAGGGGCCCTCGAGTGGAGGTAAGAAAAGGATCAACTGGTTTGAGAACAGAGAA

ATAATAGACATAGAGGTGCAGGATCCAGAAGAGTTCAATATGTTGTTAGCCTCCATACTAGCACAAGTCTGGATCCTTCT

GGCCAAGGCTGTTACGGCACCAGATACGGCAGCTGACTCAGAACTGAGAAGGTGGGTTAAATACACACAACAAAGGAGAG

TGATTGGGGAATTTCGCCTTGACAAGGGGTGGCTGGACGCAGTCCGCAACAGAATTGCAGAAGATCTATCACTCCGGCGG

TTCATGGTATCTCTCATACTTGACATCAAGAGGACCCCAGGCAACAAGCCAAGGATTGCAGAAATGATCTGCGACATTGA

CAACTATATTGTCGAAGCAGGACTCGCCAGTTTCATCCTTACTATCAAATTTGGTATCGAAACCATGTATCCTGCATTAG

GGCTCCACGAGTTTGCCGGAGAATTGTCCACTATTGAGTCCTTGATGAACTTGTATCAACAGCTAGGAGAGGTTGCACCC

TACATGGTAATTCTAGAGAATTCAATTCAGAACAAGTTTAGTGCAGGAGCTTATCCCCTCCTCTGGAGCTATGCGATGGG

TGTCGGAGTCGAGCTGGAGAACTCAATGGGGGGCTTGAACTTTGGCAGGTCATATTTTGACCCTGCCTATTTTCGTCTCG

GACAGGAGATGGTCAGAAGATCTGCAGGAAAGGTCAGCTCTGTGATTGCGGCTGAGCTCGGCATCACAGCAGAGGAAGCT

AAACTAGTCTCGGAAATCGCCTCGCAGGCTGGGGACGAAAGAACCGCCAGAGGGACTGGGCCTCGACAGGCGCAGGTTTC

CTTCCTTCAGCACCAAACAGGAGGGGGAGAGTCGTCCGCACCAGCGACCAGAGAAGGGGTCAAAGCTGCGATCCCAAACG

GAGCTGAAGAAAGGGACAGAAAGCAAACACGCCCAGGAAGGCCCAGAGGAGAGACCTCCGGCCAACTGCTCCTGGACATC

ATGCCAGAGGATGAGATCTTGCGAGAGTCTGGTCAAAACCCTCGTGAGGCTCAAAGATCGGCCGAGGCACTCTTCAGGCT

GCAGGCCATGGCCAAGATTCTGGAGGACCAGGAGGAGGGAGAAGACAACAGTCAGGTCTACAACGACAAGGATCTCCTCG

GCTAAGCAGACGCACCCTCTGTCGAAATCAGTGACAAGACATCTCCTACCAGTATTATAAAAAACTTAGGACCCAGGTCC

AAGCAAGCACACATCGACACTCCAACCAGTCGAGCGGAGACCACCGATGGCAGAAGAACAAGCATACCATGTCAACAAGG

GGCTGGAATGTATCAAGTCTCTCAAAGCCTCACCCCCGGATCTATCCACCATCAGAGATACCATCGAGAGCTGGAGAGAG

GGGCTTAGCCCCTCGGGCCGTGCAACACCGAACCCTGATACGTCCGAGGGAGATCATCAGAATATCAACCAATCATGCTC

ACCAGCAATCGGACCAAACAAAGTCTACTTGTCTCCTGAAGATAATCTCGGATTTAGAGAGATCACTGGCAACGACTGTG

AGGCTGGGCTCGGAGGGGTCCAGAGAGAAGGATCCAACTCTCAAGTACAGCGTTACCATGTTTATAGTCACGGGGGTGAA

GAGATTGAAGGACTCGAGGATGCTGACTCTCTCGTGGTTCAAGCAGATCCTCCAGTTGCTAACGTCTTCAATGGAGGAGA

GGATGGATCTGACGACAGCGATGTGGACTCTGGCCCAGATGATCCCGGCAGAGATACTCTATATGACCGGGGATCTGTTG

CCGGCAATGGTATCGCTAGGTCCACAGATGTCGAAAAACTAGAAGGTGCTGATATTCAAGAAGTTCTTAACTCCCAGAAA

GGCAAAGGAGGAAGATTCCAGGGCGGGAAAACCTTGCGAGTCCCGGAAATACCCGATGTCAAGCACTCCAGACCATCAGC

CCAATCAATTAAAAAGGGCACAGACGGGAACTCAGTCTCATCTGGAACGGTGATAGAGTGTTTATCGATAAGTGGTGCAA

CCCAAACTGTGCCAGAGTCAAGATGGGAGTCATCAGAGCAAAATGCGTCTGTGGGGAGTGTCCTCAAGTCTGCGAGGAGT

GCAAAGACGATCCAGGGGTCGACACAAGAATCTGGTACCATAGCATCACTGACTCAGCCTAAAGAGAATGACTCCGAGTA

TGAGTATGAGGATGACCTATTTACAGAGATTCAGGACATCCGTGCAAGCATTGCCAAGATCCATGATGACAATAAAACTA

TCCTCTCAAAGCTTGATTCTATACTGTTATTGAAAGGGGAAGTCGACACTATCAAGAAACAAATCAGCAAGCAGAATATA

AGTATATCCACCATTGAGGGCCATCTCTCCAGTATAATGATAGCCATCCCTGGCTTTGGGAAGGACATCAAGGACCCAAC

ATCCGAAGTCGAGTTGAACCCAGATCTAAGACCTATAATAAGCCGTGACTCTGGCAGAGCTCTCGCGGAGGTCCTCAAGA

AACCCGCTGTGGATAGGTCTCAGAAAATTGGAACCAAAGCCAACTCCAGCTCAAAGGGTCAGCTTCTTAAGGATCTCCAG

CTAAAACCTGTCGACAAGCAGGCAAGCTCTGCAATCGGGTTTGTCCCGTCCGACCATGAATCATCCAGAAATGTCATCCG

CTCCATAATCAAGTCGAGCAAGCTAAATATTGATCACAAGGACTATCTTCTAGATTTACTAAATGATGTGAAAGGCTCCA

AGGATCTTAAGGAATTTCACAAGATGCTAACAGCAATTCTCGCCAAGCACCCGTAACACATCCTCCAGTCATCATCTCAT

ACTCGACTAAAAACATCCTTTCAATCAGGCTATTACAAAAAACTTAGGAGCAAGGGCAACTGAGCTTCGCAGACAGGATG

ACCGAGATCTACGACTTCGATAAATCAGCATGGGATGTCAAAGGGTCAATTGCCCGCATAGAACCCACCACCTACCACGA

CGGCCGACTGATACCCCAGGTGAGGGTCATCGATCCTGGTCTGGGAGACAGAAAAGATGAGTGCTTTATGTACCTGTTTC

TCCTAGGAGTGATTGAGGATAACGACCCCCTGTCTCCCCCCGTCGGAAGAACCTTTGGCTCTCTACCTCTAGGGGTTGGT

AGGTCAACTGCCAAGCCAGAAGAACTACTAAGGGAGGCCACAGAATTAGATATAGTGGTGAGGCGCACTGCAGGAGTAAA

TGAGAAACTGGTATTTTACAACAACACTCCGCTGTCCTTGTTAACGCCCTGGAAGAAAGTTCTGACAACCGGAAGTGTGT

TTAGCGCTAACCAGGTTTGCAATGCAGTCAACCTAGTCCCGCTTGATACTCCCCAGAGATTCAGGGTTGTGTACATGAGC

ATAACTAGATTATCAGACAATGGTTACTATAGTGTGCCCAGAAGAATGCTGGAGTTCCGCTCGGCCAATGCAGTAGCCTT

CAACATCTTGGTTACACTAAGAATTGAAAATGGCACAAACCCTAGAAGATACATAGTCGGCTCATGGGAGAATTCAGAGG

TCACATTTATGGTACACGTGGGCAACTTTAGGAGAAAGAAGAACGAAGTATACTCTGCTGATTACTGCAAAATGAAGATT

GAAAAGATGGGTTTAGTTTTTGCCCTGGGTGGAATAGGCGGAACAAGTCTCCATATTAGAAGCACCGGGAAAATGAGCAA

AACCCTCCATGCACAGCTGGGGTTCAAGAAAATCTTATGTTACCCCCTAATGGATGTCAATGAGGATCTTAACCGATATC

TCTGGCGGGCAGAGTGCCGAATAGTCAAAATCCAAGCTGTCTTACAGCCATCAGTACCCCAAGAGTTCCGTGTCTACGAT

GATGTCATCATCAACGATGATCAAGGCTTGTTCAAGATCTTGTAGTTCATTTGCAAAGGAACGAACCCACCCCCAGGAAG

GGGCCCCACCCTCCCCCG-CCAAAAAACACACTCCAAGGCTCCCCGGCACAGCGCCAACACAGCTACTCCCGCGATGGAA

CGAGGTCGGCC-CCCGAACAAATCCGGTCCAGGGGGGGGGGCCCCCCCCCTGACCAACCGGGCGCCGACCGAGCAGGCCG

GCGAGAAGCCCCCAAGAGGCCCCGACGAAGACCGAGCACCCCATCCCCCC-AAAAACGTCCCCC-AACACGCCCGAGGCT

AGCCACCGGACACCCCCAGACCCGCCACCGGCGAGACAGAACCCCAAGGGGGGACCCTCCAAACCAAAGGAATCACCGGG

GTAGGCTCCGCCGAACCAGCACAGACCCGACCCAAGCAAGGCGGACCCCCAACACGAAACTGCCCAGCACACCCTACTCA

TCATCAAACAAAACTTAGGGGCCAAGTCCATACGAGAGGCCCTAGTCCGTCAGGCGGAGGGACGCAGAAAGGAAGGAGAC

ACGACCGTCCCAGAAGGAGGACACCCAACCACCGAAACGCCCGCCCCCAGGGAAGGAG--CCA-CCCCCACGCAGCGAAG

GAGCGGGTGGGAGACCTCGGGGAGGGSCCGACCGGGACCCGGGGGCCGGCGGCGCCGGCGGTGAGCAGGCCGGGCGCATA

CCTGACCCCACCGGAGGATCCGAGGAGGGCCCGACCCCGCGTGGACCAAACGGCAGGGAGCCCAACCCCACCAAAAAGAG

AGGCGAGGCCCCCCCTCCCTCCCCCCCCGCACACAACCCGCAGACAGACCGAGGGAGGGCCATCCCCCAGCGAGACAGGC

GGCTCCCTCGCAGCCGAGCCGCAGCTCCCCAGAATAGGACACCCCCGCACACTCCGAACACCCCCTCGCCCCCCGCCACC

GGGACAGGGGCTCCACCACCACAGACCGGAGCCACCGACAGCCAGCCAGAGGTCCCCTCCAGTCCCGCCCGGCGCCCCCC

CGAAGAGGCCCCGGCAAGGCCCCCCAGCCCGACGTCCCCCGCACGCACCGAAAGAAGGACCCGCCCGAGAACGTGCGGCA

CATCCATGCGCAAACATCATGACGCGGGTCGCAATTTTGACATTTCTGTTTCTTTTCCCAAATGTTGTTGCGTGTCAGAT

TCACTGGGGCAATCTATCCAAGATCGGGATTGTAGGAACAGGGAGTGCCAGCTACAAGGTGATGACTAGGCCAAGCCACC

AGACTCTGGTTATAAAGTTAATGCCAAATATAACGGCCATCGACAATTGTACAAAGTCAGAGATTGCAGAGTACAAGAGA

TTGCTGATCACAGTGTTAAAGCCTGTAGAGGATGCTCTGTCGGTGATAACCAAGAATGTAAGACCAATTCAAACTCTAAC

ACCTGGGCGTAGAACCCGCCGTTTTGCTGGAGCTGTTCTGGCCGGGGTAGCACTTGGAGTTGCGACAGCCGCTCAGATAA

CTGCAGGAGTCGCCCTTCATCAATCATTGATGAACTCCCAAGCAATTGAGAGTTTAAAAACCAGTCTTGAGATGTCGAAT

CAGGCAATAGAAGAAATCAGACTTGCAAATAAGGAGACCATACTGGCAGTACAGGGCGTCCAGGATTATATCAACAATGA

GCTCGTCCCTTCTGTTCATAGAATGTCATGCGAGCTGGTAGGTCACAAGCTCGGCCTCAAGCTCCTTAGGTACTACACCG

AGATCCTGTCCATATTCGGGCCCAGTCTTCGAGACCCGATATCTGCCGAAATATCAATCCAGGCACTTAGTTATGCATTA

GGCGGAGACATTAATAAAATCCTGGACAAGCTTGGGTATAGCGGTGGGGATTTCCTTGCCATCCTAGAAAGCAAGGGAAT

AAAGGCCCGGGTTACATATGTGGACACAAGAGATTACTTTATAATCCTTAGCATCGCCTACCCAACCTTATCTGAGATCA

AGGGAGTGATAGTTCACAAGATAGAAGCTATAACATACAACATTGGGGCACAGGAGTGGTATACTACTATCCCTAAATAT

GTAGCCACTCAGGGGTATCTGATATCGAACTTTGATGAGACGTCATGCGTATTCACTCCAGAGGGGACAGTTTGCAGCCA

GAATGCGTTGTACCCAATGAGCCCATTGCTTCAGGAATGTTTCAGGGGGTCAACAAAATCGTGCGCCAGAACCTTAGTTT

CAGGGACCATAAGTAATAGATTTATCCTATCAAAAGGGAACCTGATTGCAAATTGTGCGTCAGTTTTGTGCAAATGTTAC

ACAACAGAGACAGTTATCAGCCAAGATCCTGACAAACTACTAACTGTTGTAGCATCCGACAAGTGTCCTGTAGTTGAGGT

GGATGGAGTGACAATACAGGTCGGCAGTCGAGAGTATCCGGATTCTGTATACTTACACAAAATAGACTTAGGTCCAGCCA

TCTCCCTAGAAAAACTGGATGTAGGCACCAATTTAGGCAATGCAGTCACAAGACTGGAGAATGCAAAGGAGCTCCTAGAT

GCATCAGACCAAATACTGAAGACTGTTAAAGGGGTACCTTTCAGTGGGAATATGTACATAGCACTGGCAGCTTGCATAGG

AGTATCCCTGGGCCTTGTCACATTAATATGCTGCTGTAAGGGGAGGTGTAAGAACAAGGAAATCCCTATCTCCAAAATCA

ACCCAGGGCTCAAACCCGACCTGACCGGGACCTCAAAGTCGTACGTAAGATCACTGTAGTCAGAATCACCTGGATCATCT

GGCATCACACACATACATGCACGACACAGGCAGTCCGAGGACGCAAGAAACCCAGCCTCCGGTCACCCACCCGACCCCAC

TCCACGCTCCACCACACATTAGTCATCAAACAAAACTTAGGACGAAAGGTCAATCACCATGTCCGCACAAAGGGAGAGGA

TCAATGCCTTCTACAAAGACAATCCTCACAATAAGAACCATAGGGTGATCCTGGATAGAGAACGCTTGGTCATTGAAAGA

CCCTACATCTTGCTTGGAGTCCTGCTGGTAATGTTCCTGAGTCTAATCGGACTGCTGGCCATTGCAGGGATCAGGCTTCA

CCGGGCCACCGTTGGAACTTCAGAGATCCAGAGTCGGCTGAATACCAATATTAAGTTGACCGAATCTATTGATCACCAGA

CTAAGGATGTCTTAACTCCCCTTTTTAAAATCATTGGCGATGAAGTCGGCATCAGAATTCCACAGAAATTCAGTGATCTT

GTCAAGTTCATCTCCGATAAGATTAAATTCCTCAACCCTGATAGAGAGTATGATTTCAGAGATCTCCGGTGGTGTATGAA

TCCCCCCGAGAGAGTCAAAATTAATTTTGATCAGTTTTGTGAGTACAAGGCTGCGGTTAAGTCAATTGAACATATATTTG

AGTCACCACTCAACAAGTCAAAAAAGCTGCAATCTTTGACTCTCGGGCCCGGAACAGGCTGTCTAGGCAGGACAGTAACA

AGAGCCCATTTCTCAGAACTTACAATGACCTTAATGGACCTGGATCTAGAGATGAAGCACAACGTGTCCTCAGTGTTTAC

CGTAGTTGAAGAGGGATTATTCGGAAGAACATATACCGTCTGGAGATCCGATGCCAGGGATCCGAGCACCGATCTAGGTA

TCGGCCATTTTTTAAGAGTCTTCGAGATTGGACTGGTAAGAGATCTTGGGCTGGGTCCCCCTGTTTTTCATATGACCAAC

TATCTCACGGTGAACATGAGTGATGACTATCGGAGATGTCTTTTAGCGGTAGGGGAGTTGAAGTTGACAGCCCTATGCAC

CTCATCTGAGACTGTGACACTGAGTGAGAGAGGAGTTCCAAGGAGGGAACCTCTTGTGGTTGTGATACTTAATCTAGCTG

GACCCACTCTAGGGGGAGAGCTATACAGTGTCTTGCCTACCTCTGATCTCATGGTGGAGAAACTCTATTTGTCTTCACAT

AGAGGGATCATCAAAGATGATGAGGCCAATTGGGTAGTGCCGTCTACCGATGTTCGTGATCTTCAGAATAAAGGTGAATG

TCTGGTGGAAGCATGCAAGACTCGACCTCCTTCATTTTGCAATGGCACAGGATCAGGCCCGTGGTCAGAAGGGAGAATCC

CTGCCTACGGGGTGATCAGGGTCAGTCTTGACTTAGCGAGTGACCCGGATGTAGTTATCACTTCAGTGTTTGGCCCACTG

ATACCTCACCTATCCGGCATGGATCTTTACAACAACCCGTTTTCAAAAGCTGTATGGTTGGCTGTACCACCTTATGAGCA

GTCATTTCTAGGAATGATAAATACAATTGGATTCCCTAACAGAGCAGAGGTTATGCCGCACATTTTGACCACAGAGATCA

GAGGCCCTCGGGGTCGTTGCCATGTTCCCATAGAATTGTCCCGCCGGGTTGATGACGATATCAAGATCGGGTCCAACATG

GTCATATTGCCGACGATGGACCTGAGGTATATTACAGCCACTTATGATGTTTCCAGGAGCGAGCATGCAATCGTGTACTA

TATCTATGACACGAGTCGCTCATCATCTTACTTCTACCCAGTTCGACTGAATTTCAAAGGCAATCCTCTCTCTTTGAGGA

TAGAGTGTTTCCCCTGGCGTCATAAGGTGTGGTGCTACCATGATTGTCTTATATACAACACCATAACAGGTGAAGAGGTC

CATACGAGAGGGCTGACCGGCATAGAGGTAACATGCAATCCAGTCTGAGCAGAGCTACGACCGTCACTCAGGCAGTCCCT

TGAGTCGCCACCGAGTCCAAGCAGCACAGCCTGGGACAATCAACGGCACAGCCCAGCCAACAATGTTATAAAAAACTTAG

GAGCCAGGGTTGTAGGGGCCATGGACTCTCTATCAGTCAATCAGGTCTTGTACCCTGAGGTTCATCTAGACAGCCCTATT

GTCACAAACAAACTAGTTGCCATCCTTGAGTACTCGGGTATCGACCATAACTATGTTCTTGAAGACCAGACTCTTGTCAA

GAATATTAGGTATAGGCTGGGGTGCGGTTTTTCAAATCAAATGATCATCAATAACAGGGGGGTGGGTGAAACAGTCAATT

CTAAACTTAAAAGTTACCCCCATAATCGTCATATCATATACCCGGATTGCAATAAGGAGTTGTTTTGTATCAAGGATAGC

TGCATATCTAAGAAGCTCTCGGAGCTATTCAAGAAGGGTAATTCCTTGTACTCTAAGATAAGTCACCAGGTACTGGATTG

TCTTAAGAGAGTCAATGGGAAATTAGGTCTGGGCACAGATCTTACCCATGGTCTGAAGGAGGGTATCCTTGACCTAGGGT

TGCACATGCATAGCTCTCAATGGTTCGAGACTTTCCTGTTCTGGTTCACTATCAAGACAGAGATGAGATCAATGATCAAA

GAACAGTCCCATATATGCCACAAGAGGAGGTATAACCCGACTTTTGTGTCGGGGGATGCATTCGAGGTGCTCGTATCGCG

AGACCTCGTTGTGATAATTGATAAGAATACCCAGTATGTCTTCTACTTAACATTTGAGCTGGTCCTTATGTATTGTGATG

TCATAGAGGGTAGACTTATGACGGAGACAGCCATGGCCATAGACCAGAGATATTCAGAACTTCTAAGCCGGGTCAGATAC

TTGTGGGATCTTATTGATGGGTTCTTCCCAACACTAGGCAACACCACATACCAAATTGTTGCTCTGCTTGAACCATTATC

GTTGGCTTATCTTCAACTTCAGGATGTCACTCTGGAGTTAAGAGGTGCTTTTTTAGACCATTGCTTCAAAGAACTCTATG

AGATACTGGAACATTGCGGCATTGACACAGCAGGCACTTACAATTCCATCACTGAAGGATTGGATTACGTATTCATCACC

CACGATATACATCTAACTGGGGAGATTTTTTCATTTTTCCGGAGTTTCGGACACCCCCGTCTAGAAGCAGTCACCGCTGC

GGAAAATGTCAGAAAACATATGAACCAACCGAAGGTAATCAGTTATGAGACTATGATGAAAGGCCATGCGGTATTTTGCG

GGATAATCATAAATGGTTTTCGGGACCGACATGGCGGCAGCTGGCCCCCTGTTGCACTGCCAGAACATGCCTCTGCTGCG

ATCCGGAATGCGCAGGCATCAGGTGAGGGACTGACCCATGACCTGTGTATAGACAACTGGAAATCCTTTGTTGGATTCAA

ATTTGGCTGCTTTATGCCACTCAGCCTAGATAGTGATTTGACCATGTATCTCAAAGACAAGGCATTGGCTGCACTGAAGA

ATGAGTGGGATTCAGTTTACCCGAAAGAATACCTCCGTTATAACCCACCTAGAGGGACAGAGTCAAGGCGATTGGTAGAG

GTATTCCTGAATGACTCCAGCTTTGATCCTTATAACATGATAATGTACGTGGTGAATGGCTCCTACCTTAAAGACCCTGA

GTTTAACCTCTCATACAGCCTAAAAGAGAAGGAGATAAAGGAGACAGGGCGGTTGTTTGCCAAAATGACCTATAAGATGC

GGGCCTGTCAGGTAATCGCTGAAAATCTGATATCGAATGGTGTTGGGAAGTATTTCCGAGACAATGGGATGGCAAAGGAC

GAGCATGACCTAACAAAAGCCCTTCACACCCTGGCAGTCTCAGGTGTTCCCAAAAACAACAAAGATAACCACCGAGGTGG

GCCTCCCAGAAGGACAACAAACCGAGGGGTGAGATCAAGCCAAGGCACCAAAACACAAGATAGAGACAAGGTTCAAGGGG

GACCTATGTACAACTATTTGCGATGCCAACCGATCAGCCCTGATCAGGGTGAGTCATACGAGACTGTTAGTGCATTCATC

ACCGCTGACCTTAAGAAGTATTGCCTGAATTGGAGATACGAGACAATCAGCATATTTGCACAGAGGCTGAATGAAATATA

TGGACTACCATCCTTCTTTCAATGGTTACACAGGATATTGGAAAAATCCGTACTCTACGTCAGTGACCCACATTGTCCTC

CCGATCTAGATAATCATATCCCTCTGGACAGTGTCCCCAATGCCCAAATATTCATCAAGTACCCAATGGGCGGAATAGAA

GGTTATTGCCAAAAACTATGGACAATCAGCACTATACCATACTTGTATCTGGCAGCCTATGAGAGCGGAGTAAGAATCGC

CTCACTGGTGCAGGGTGACAATCAGACAATCGCAGTGACAAAAAGAGTTCCAAGTTCTTGGCCTTATTCACTAAAAAAGA

GGGAGGCCTCCAAGGCCGCTCAAAATTACTTCGTAGTCTTAAGGCAAAGATTGCACGATGTAGGTCATCACTTAAAGGCT

AATGAGACCATAGTATCTTCTCACTTTTTTGTATATTCCAAAGGGATTTATTATGACGGCCTGCTAGTCTCACAATCACT

AAAGAGCATCGCCAGATGTGTCTTCTGGTCCGAGACTATAGTGGATGAAACCAGAGCGGCGTGCAGTAATATTGCAACAA

CTGTCGCTAAGAGTATAGAGAGGGGTTATGATAGGTACCTTGCATATTCTTTGAATATCCTCAAGATTTTCCAACAGATC

CTTATATCCCTTAACTTCACTATTAACACAACAATGACTCAGGATGTCGTGGCACCGATCATCGAGAACGGTGATTTGCT

GATAAGGATGGCACTCTTGCCAGCACCCATCGGGGGTCTCAATTATCTTAACATGAGCAGGTTATTTGTGAGAAATATCG

GTGACCCGGTCACCTCCTCTATAGCCGACCTGAAGAGGATGATAGATGCCGGGCTAATGCCAGAAGAAACATTGCATCAA

GTGATGACCCAGACCCCGGGAGAATCATCCTACCTAGATTGGGCAAGTGACCCTTATTCTGCCAACCTACCCTGCGTACA

GAGTATCACTCGCCTTCTCAAAAACATCACTGCACGATATATTTTAATCAGCAGCCCAAACCCGATGCTGAAAGGGTTAT

TTCATGAGGGGAGTAGAGATGAAGACGAGGAGCTTGCAAGTTTCCTAATGGATCGGCATATAATTGTTCCAAGAGCTGCA

CATGAAATCTTAGACCATAGTATAACCGGAGCAAGAGAAGCTATAGCCGGGATGTTGGACACCACCAAGGGTCTGATTAG

AACAAGTATGAAACGGGGCGGCCTCACCCCTCGAGTCTTAGCCCGCCTTTCCAATTATGATTATGAACAATTCCGATCCG

GGATAACACTATTAACAAAGAAAGGGCAGTGTTATCTCATTGACAAGGACTCGTGCTCGGTGCAGCTCGCTATAGCTCTG

AGGGGCCACATGTGGGCTAAGTTGGCTCGCGGGAGACCAATCTATGGGTTGGAGGTGCCTGATGTATTAGAATCGATGAA

CGGCTACCTTATCAAACGCCACGAGTCGTGTGCTATCTGTGAAACGGGCTCAAGTCACTACGGATGGTTTTTCGTCCCTG

CAGGGTGCCAGCTTGACGATGTATCTAGAGAGACTTCGGCTCTTCGTGTACCTTATGTCGGATCAACAACTGAGGAAAGG

ACAGACATGAAACTTGCATTTGTTAGGTCTCCAAGTAGATCCCTCAAATCAGCAGTTAGAATTGCAACAGTTTACTCATG

GGCTTACGGGGATGATGAGAAATCATGGAGTGAAGCCTGGATGCTAGCTAGGCAGAGAGCCAATATCACCTTAGATGAAT

TGAGAATGATCACTCCAGTCTCCACATCCACCAACTTAGCCCACCGGTTGAGGGATCGGAGTACCCAGGTGAAATACTCG

GGGACATCCCTTGTGAGGGTTGCAAGATACACAACCATCTCCAATGATAACTTGTCGTTTGTGATATCTGAGAAAAAAGT

GGATACCAACTTCATTTATCAGCAAGGGATGTTGCTTGGTCTTGGGATCCTTGAGAACCTCTTTAGGCTAGAGGCCACCA

CCGGAGTATCCAACACAGTGCTACACCTGCACGTGGAAACAGAATGTTGTGTTGTACCTATGGTGGATCACCCAAGGATA

CCGAGTCTCCGTAATATAAAAGTTACGGATGAGCTATGCACAAACCCTTTGATCTACGACAGGTCCCCCATCATAGAGCA

CGATGCAACCCGGCTATATTCACAGAGCCACAGGAGACATTTGGTGGAGTTTGTTACCTGGTCAACAAGCCAGCTTTACC

ATATACTGGCTAAGTCTACAGCAATGTCCATGATTGAATTGATCACGAGATTCGAGAAAGATCACATGAATGAAATAGCC

GCCCTGATTGGCGATGACGACATCAACAGCTTCATCACAGAATTTTTGCTAGTGGAGCCTAGATTATTTATAGTTTACCT

TGGTCAGTGTGCTGCCATCAATTGGGCCTTTGATATACATTATCATCGGCCCTCGGGCAAGTACCAGATGGGGGAACTTC

TCTACTCTCTGCTCTCTCGGATGAGCAAAGGAGTATATAAGATCTTTACTAATGCTTTGAGCCACCCCAAAGTTTACAAG

AAATTTTGGCGAAGTGGCATCATTGAGCCGATTCATGGCCCATCCTTGGATACACAGAATTTACATGTCACTGTCTGTGA

CATGATATATGGGTCCTATGTCACCTATTTGGATCTTTTGCTGAATGATGAGCTAGATGATTACCCGTATTTGCTCTGCG

AGAGTGATGAGGACGTGGTCACAGACAGGTTCGACAACATTCAAGCCAAACATCTCTGTGTATTGGCCGATGTATATTGC

AGCTCCAAGAGGTGTCCTTCAATAATCGGGATGTCTCCTATAGAAAAATGTACCATTCTCACACATTACATCAAGGGAGA

ATCAGTACAATCCCCGTCTGGGACCTCATGGAACACTGATCCCCTTGTAGTAGATCATTACTCATGCTCTCTGACCTACC

TTCGCCGCGGTTCCATCAAACAAATCAGGTTGAGGGTGGATCCTGGGTTTGTATTCGAGGCGTTGACAGACGTCGACTTT

AAACAACCTCGCAAAGCTAAGTTAGATATATCGGTCGTTGGGTTGACTGATTTCTCTCCCCCTTACGACAACGTCGGTGA

TTTTTTAGGAACTATCAACACATTGAGGCACGATCTGCCTGTCACCGGAACCGGGGTCTCGAACTATGAAGTCCACGCTT

ATCGTAGAATTGGCCTGAATTCATCAGCATGTTATAAAGCCGTAGAGATCTCCACGTTAATCAAGCCATCCTTAGAAGTC

GGAGAGCATGGATTGTTCTTAGGAGAAGGTTCCGGTTCAATGCTGGCTGCGTACAAAGAAGTTCTTAAATTAGCAAATTG

TTACTACAACAGCGGAGTAACAGCGGAGGGCAGAGCCGGACAGAGGGAAATATCTCCCTATCCTTCAGAGATGAGCCTAG

TAGAGAATCAAATGGGGATAGAGCGGAGTGTTAAAGTGCTGTTCAACGGGAAACCTGAAGTAACTTGGGTGGGGACCACA

GATTGCTACAAGTACATAATCAGTAACATCCAAACCTCTAGTCTGGGTTTCATACATTCAGATATTGAGACGCTCCCAAC

CAAAGATGCTGTTGAGAAATTGGAGGAATTTGCCTCTATCCTATCCTTATCCTTGATTTTGGGGAAAATCGGCTCTATTA

CAGTTGTCAAAATTATGCCCATTAGCGGGGATTTTACCCAAGGCTTCATAGCCTATGCCATCCAATATTTTAGGGAGAGC

CTGCTTGCCTATCCGAGATATAGTAACTTCATCTCGACTGAGTGTTACCTTATTATGATAGGATTAAAGGCCAATCGATT

GATAAACCCAGAAGCCATTAAGCAAAGCATAATCAGAGCGGGGGTCAGGACTGCACCAGGACTTGTGAGCCATGTATTAT

CAGAGAAACAAAAAGGTTGTATTCAATCTTTCTTGGGTGATCCTTATATCCAAGGAGACTTCAATAAGCACCTTAAATCT

CTAACCTCTATTGAGAAAATCCTGGTAAATTGTGGTCTCTCGATCAATGGCATAAAAATCTGTAGAGATCTAATCCACCA

TGATATCGCCTCCGGTCCAGACGGCCTTATGAGCTCTACAATTATTTTATATAGGGAATTGGCTCATTTCAAAGATAATA

TAAGGAGTCAGCACGGAATGTTCCACCCCTACCCGGTACTGGCCAACAGCAGGCAACGTGAATTAATCCTTCGAATAGCC

AAGAAATTCTGGGGGTATGTCTTGCTATATTCTGATGACCCGGCACTAATCAGACAAACAATCAAGAACTTGAAACGGAA

TCATCTAACCTTTGACTTACATAGTAACCCATTTATTAAAGGGCTATCCAAGGCTGAGAAACTGCTGGTGCGGACGAGTT

CACTAAGAAGAGAATGGTTGTTCACTCTCGAAACGAAAGAAGTAAAAGAGTGGTTCAAATTGGTAGGTTACAGTGCACTC

ATCAGAGGCTAGTCAGCTATGCATCTGCCCCCTTCTCCTCCGCCATGAGACCCCACTGGCGATCCAGAAGATTAAAGAAA

ACTACATATTGGATAAGTATCTATTCCCAGCTTTGTCTGGT

>KX421386.1/goat/China/2013-12-22

ACCAAACAAAGTTGGGTAAGGATAGATCTTATAATAACTATGGACTGGCAAACTTAGGAGTAAAGATCCTACTGTCGGGG

GGAGGAGGAGGAGCAAGATCCTTGATCATGGCGACTCTCCTTAAAAGCTTAGCATTGTTCAAAAGGAACAAAGACAAAGC

GCCGACAGCATCAGGTTCAGGAGGGGCCATCCGGGGGATTAAGAATGTTATCATAGTCCCGATTCCCGGAGACTCGTCCA

TCACTACCCGTTCAAGACTGCTCGACAGGCTTGTCAGATTGGCCGGAGATCCTGACATCAACGGGTCAAAGCTGACCGGG

GTGATGATCAGCATGTTATCCTTGTTCGTAGAGTCACCCGGGCAATTGATACAGCGAATCACAGATGATCCGGATGTTAG

TATCCGCCTTGTTGAGGTAGTTCAAAGTACTAGGTCCCAGTCCGGGTTGACCTTTGCATCACGTGGTGCTGATTTAGACA

ACGAGGCAGACATGTATTTTTCAACTGAGGGGCCCTCGAGTGGAGGTAAGAAAAGGATCAACTGGTTTGAGAACAGAGAA

ATAATAGACATAGAGGTGCAGGATCCAGAAGAGTTCAATATGTTGTTAGCCTCCATACTAGCACAAGTCTGGATCCTTCT

GGCCAAGGCTGTTACGGCACCAGATACGGCAGCTGACTCAGAACTGAGAAGGTGGGTTAAATACACACAACAAAGGAGAG

TGATTGGGGAATTTCGCCTTGACAAGGGGTGGCTGGACGCAGTCCGCAACAGAATTGCAGAAGATCTATCACTCCGGCGG

TTCATGGTATCTCTCATACTTGACATCAAGAGGACCCCAGGCAACAAGCCAAGGATTGCAGAAATGATCTGCGACATTGA

CAACTATATTGTCGAAGCAGGACTCGCCAGTTTCATCCTTACTATCAAATTTGGTATCGAAACCATGTATCCTGCATTAG

GGCTCCACGAGTTTGCCGGAGAATTGTCCACTATTGAGTCCTTGATGAACTTGTATCAACAGCTAGGAGAGGTTGCACCC

TACATGGTAATTCTAGAGAATTCAATTCAGAACAAGTTTAGTGCAGGAGCTTATCCCCTCCTCTGGAGCTATGCGATGGG

TGTCGGAGTCGAGCTGGAGAACTCAATGGGGGGCTTGAACTTTGGCAGGTCATATTTTGACCCTGCCTATTTTCGTCTCG

GACAGGAGATGGTCAGAAGATCTGCAGGAAAGGTCAGCTCTGTGATTGCGGCTGAGCTCGGCATCACAGCAGAGGAAGCT

AAACTAGTCTCGGAAATCGCCTCGCAGGCTGGGGACGAAAGAACCGCCAGAGGGACTGGGCCTCGACAGGCGCAGGTTTC

CTTCCTTCAGCACCAAACAGGAGGGGGAGAGTCGTCCGCACCAGCGACCAGAGAAGGGGTCAAAGCTGCGATCCCAAACG

GAGCTGAAGAAAGGGACAGAAAGCAAACACGCCCAGGAAGGCCCAGAGGAGAGACCTCCGGCCAACTGCTCCTGGACATC

ATGCCAGAGGATGAGATCTTGCGAGAGTCTGGTCAAAACCCTCGTGAGGCTCAAAGATCGGCCGAGGCACTCTTCAGGCT

GCAGGCCATGGCCAAGATTCTGGAGGACCAGGAGGAGGGAGAAGACAACAGTCAGGTCTACAACGACAAGGATCTCCTCG

GCTAAGCAGACGCACCCTCTGTCGAAATCAGTGACAAGACATCTCCTACCAGTATTATAAAAAACTTAGGACCCAGGTCC

AAGCAAGCACACATCGACACTCCAACCAGTCGAGCGGAGACCACCGATGGCAGAAGAACAAGCATACCATGTCAACAAGG

GGCTGGAATGTATCAAGTCTCTCAAAGCCTCACCCCCGGATCTATCCACCATCAGAGATACCATCGAGAGCTGGAGAGAG

GGGCTTAGCCCCTCGGGCCGTGCAACACCGAACCCTGATACGTCCGAGGGAGATCATCAGAATATCAACCAATCATGCTC

ACCAGCAATCGGACCAAACAAAGTCTACTTGTCTCCTGAAGATAATCTCGGATTTAGAGAGATCACTGGCAACGACTGTG

AGGCTGGGCTCGGAGGGGTCCAGAGAGAAGGATCCAACTCTCAAGTACAGCGTTACCATGTTTATAGTCACGGGGGTGAA

GAGATTGAAGGACTCGAGGATGCTGACTCTCTCGTGGTTCAAGCAGATCCTCCAGTTGCTAACGTCTTCAATGGAGGAGA

GGATGGATCTGACGACAGCGATGTGGACTCTGGCCCAGATGATCCCGGCAGAGATACTCTATATGACCGGGGATCTGTTG

CCGGCAATGGTATCGCTAGGTCCACAGATGTCGAAAAACTAGAAGGTGCTGATATTCAAGAAGTTCTTAACTCCCAGAAA

GGCAAAGGAGGAAGATTCCAGGGCGGGAAAACCTTGCGAGTCCCGGAAATACCCGATGTCAAGCACTCCAGACCATCAGC

CCAATCAATTAAAAAGGGCACAGACGGGAACTCAGTCTCATCTGGAACGGTGATAGAGTGTTTATCGATAAGTGGTGCAA

CCCAAACTGTGCCAGAGTCAAGATGGGAGTCATCAGAGCAAAATGCGTCTGTGGGGAGTGTCCTCAAGTCTGCGAGGAGT

GCAAAGACGATCCAGGGGTCGACACAAGAATCTGGTACCATAGCATCACTGACTCAGCCTAAAGAGAATGACTCCGAGTA

TGAGTATGAGGATGACCTATTTACAGAGATTCAGGACATCCGTGCAAGCATTGCCAAGATCCATGATGACAATAAAACTA

TCCTCTCAAAGCTTGATTCTATACTGTTATTGAAAGGGGAAGTCGACACTATCAAGAAACAAATCAGCAAGCAGAATATA

AGTATATCCACCATTGAGGGCCATCTCTCCAGTATAATGATAGCCATCCCTGGCTTTGGGAAGGACATCAAGGACCCAAC

ATCCGAAGTCGAGTTGAACCCAGATCTAAGACCTATAATAAGCCGTGACTCTGGCAGAGCTCTCGCGGAGGTCCTCAAGA

AACCCGCTGTGGATAGGTCTCAGAAAATTGGAACCAAAGCCAACTCCAGCTCAAAGGGTCAGCTTCTTAAGGATCTCCAG

CTAAAACCTGTCGACAAGCAGGCAAGCTCTGCAATCGGGTTTGTCCCGTCCGACCATGAATCATCCAGAAATGTCATCCG

CTCCATAATCAAGTCGAGCAAGCTAAATATTGATCACAAGGACTATCTTCTAGATTTACTAAATGATGTGAAAGGCTCCA

AGGATCTTAAGGAATTTCACAAGATGCTAACAGCAATTCTCGCCAAGCACCCGTAACACATCCTCCAGTCATCATCTCAT

ACTCGACTAAAAACATCCTTTCAATCAGGCTATTACAAAAAACTTAGGAGCAAGGGCAACTGAGCTTCGCAGACAGGATG

ACCGAGATCTACGACTTCGATAAATCAGCATGGGATGTCAAAGGGTCAATTGCCCGCATAGAACCCACCACCTACCACGA

CGGCCGACTGATACCCCAGGTGAGGGTCATCGATCCTGGTCTGGGAGACAGAAAAGATGAGTGCTTTATGTACCTGTTTC

TCCTAGGAGTGATTGAGGATAACGACCCCCTGTCTCCCCCCGTCGGAAGAACCTTTGGCTCTCTACCTCTAGGGGTTGGT

AGGTCAACTGCCAAGCCAGAAGAACTACTAAGGGAGGCCACAGAATTAGATATAGTGGTGAGGCGCACTGCAGGAGTAAA

TGAGAAACTGGTATTTTACAACAACACTCCGCTGTCCTTGTTAACGCCCTGGAAGAAAGTTCTGACAACCGGAAGTGTGT

TTAGCGCTAACCAGGTTTGCAATGCAGTCAACCTAGTCCCGCTTGATACTCCCCAGAGATTCAGGGTTGTGTACATGAGC

ATAACTAGATTATCAGACAATGGTTACTATAGTGTGCCCAGAAGAATGCTGGAGTTCCGCTCGGCCAATGCAGTAGCCTT

CAACATCTTGGTTACACTAAGAATTGAAAATGGCACAAACCCTAGAAGATACATAGTCGGCTCATGGGAGAATTCAGAGG

TCACATTTATGGTACACGTGGGCAACTTTAGGAGAAAGAAGAACGAAGTATACTCTGCTGATTACTGCAAAATGAAGATT

GAAAAGATGGGTTTAGTTTTTGCCCTGGGTGGAATAGGCGGAACAAGTCTCCATATTAGAAGCACCGGGAAAATGAGCAA

AACCCTCCATGCACAGCTGGGGTTCAAGAAAATCTTATGTTACCCCCTAATGGATGTCAATGAGGATCTTAACCGATATC

TCTGGCGGGCAGAGTGCCGAATAGTCAAAATCCAWGCTGTCTTACAGCCATCAGTACCCCAAGAGTTCCGTGTCTACGAT

GATGTCATCATCAACGATGATCAAGGCTTGTTCAAGATCTTGTAGTTCATTTGCAAAGGAACGAACCCACCCCCAGGAAG

GGGCCCCACCCTCCCCCG-CCAAAAAACACACTCCAAGGCTCCCCGGCACAGCGCCAACACAGCTACTCCCGCGATGGAA

CGAGGTCGGCC-CCCGAACAAATCCGGTCCAGGGGGGGGGGCCCCCCCCCTGACCAACCGGGCGCCGACCGAGCAGGCCG

GCGAGAAGCCCCCAAGAGGCCCCGACGAAGACCGAGCACCCCATCCCCCC-AAAAACGTCCCCC-AACACGCCCGAGGCT

AGCCACCGGACACCCCCAGACCCGCCACCGGCGAGACAGAACCCCAAGGGGGGACCCTCCAAACCAAAGGAATCACCGGG

GTAGGCTCCGCCGAACCAGCACAGACCCGACCCAAGCAAGGCGGACCCCCAACACGAAACTGCCCAGCACACCCTACTCA

TCATCAAACAAAACTTAGGGGCCAAGTCCATACGAGAGGCCCTAGTCCGTCAGGCGGAGGGACGCAGAAAGGAAGGAGAC

ACGACCGTCCCAGAAGGAGGACACCCAACCACCGAAACGCCCGCCCCCAGGGAAGGAG--CCA-CCCCCACGCAGCGAAG

GAGCGGGTGGGAGACCTCGGGGAGGGCCCGACCGGGACCCGGGGGCCGGCGGCGCCGGCGGTGAGCAGGCCGGGCGCATA

CCTGACCCCACCGGAGGATCCGAGGAGGGCCCGACCCCGCGTGGACCAAACGGCAGGGAGCCCAACCCCACCAAAAAGAG

AGGCGAGGCCCCCCCTCCCTCCCCCCCCGCACACAACCCGCAGACAGACCGAGGGAGGGCCATCCCCCAGCGAGACAGGC

GGCTCCCTCGCAGCCGAGCCGCAGCTCCCCAGAATAGGACACCCCCGCACACTCCGAACACCCCCTCGCCCCCCGCCACC

GGGACAGGGGCTCCACCACCACAGACCGGAGCCACCGACAGCCAGCCAGAGGTCCCCTCCAGTCCCGCCCGGCGCCCCCC

CGAAGAGGCCCCGGCAAGGCCCCCCAGCCCGACGTCCCCCGCACGCACCGAAAGAAGGACCCGCCCGAGAACGTGCGGCA

CATCCATGCGCAAACATCATGACGCGGGTCGCAATTTTGACATTTCTGTTTCTTTTCCCAAATGTTGTTGCGTGTCAGAT

TCACTGGGGCAATCTATCCAAGATCGGGATTGTAGGAACAGGGAGTGCCAGCTACAAGGTGATGACTAGGCCAAGCCACC

AGACTCTGGTTATAAAGTTAATGCCAAATATAACGGCCATCGACAATTGTACAAAGTCAGAGATTGCAGAGTACAAGAGA

TTGCTGATCACAGTGTTAAAGCCTGTAGAGGATGCTCTGTCGGTGATAACCAAGAATGTAAGACCAATTCAAACTCTAAC

ACCTGGGCGTAGAACCCGCCGTTTTGCTGGAGCTGTTCTGGCCGGGGTAGCACTTGGAGTTGCGACAGCCGCTCAGATAA

CTGCAGGAGTCGCCCTTCATCAATCATTGATGAACTCCCAAGCAATTGAGAGTTTAAAAACCAGTCTTGAGATGTCGAAT

CAGGCAATAGAAGAAATCAGACTTGCAAATAAGGAGACCATACTGGCAGTACAGGGCGTCCAGGATTATATCAACAATGA

GCTCGTCCCTTCTGTTCATAGAATGTCATGCGAGCTGGTAGGTCACAAGCTCGGCCTCAAGCTCCTTAGGTACTACACCG

AGATCCTGTCCATATTCGGGCCCAGTCTTCGAGACCCGATATCTGCCGAAATATCAATCCAGGCACTTAGTTATGCATTA

GGCGGAGACATTAATAAAATCCTGGACAAGCTTGGGTATAGCGGTGGGGATTTCCTTGCCATCCTAGAAAGCAAGGGAAT

AAAGGCCCGGGTTACATATGTGGACACAAGAGATTACTTTATAATCCTTAGCATCGCCTACCCAACCTTATCTGAGATCA

AGGGAGTGATAGTTCACAAGATAGAAGCTATAACATACAACATTGGGGCACAGGAGTGGTATACTACTATCCCTAAATAT

GTAGCCACTCAGGGGTATCTGATATCGAACTTTGATGAGACGTCATGCGTATTCACTCCAGAGGGGACAGTTTGCAGCCA

GAATGCGTTGTACCCAATGAGCCCATTGCTTCAGGAATGTTTCAGGGGGTCAACAAAATCGTGCGCCAGAACCTTAGTTT

CAGGGACCATAAGTAATAGATTTATCCTATCAAAAGGGAACCTGATTGCAAATTGTGCGTCAGTTTTGTGCAAATGTTAC

ACAACAGAGACAGTTATCAGCCAAGATCCTGACAAACTACTAACTGTTGTAGCATCCGACAAGTGTCCTGTAGTTGAGGT

GGATGGAGTGACAATACAGGTCGGCAGTCGAGAGTATCCGGATTCTGTATACTTACACAAAATAGACTTAGGTCCAGCCA

TCTCCCTAGAAAAACTGGATGTAGGCACCAATTTAGGCAATGCAGTCACAAGACTGGAGAATGCAAAGGAGCTCCTAGAT

GCATCAGACCAAATACTGAAGACTGTTAAAGGGGTACCTTTCAGTGGGAATATGTACATAGCACTGGCAGCTTGCATAGG

AGTATCCCTGGGCCTTGTCACATTAATATGCTGCTGTAAGGGGAGGTGTAAGAACAAGGAAATCCCTATCTCCAAAATCA

ACCCAGGGCTCAAACCCGACCTGACCGGGACCTCAAAGTCGTACGTAAGATCACTGTAGTCAGAATCACCTGGATCATCT

GGCATCACACACATACATGCACGACACAGGCAGTCCGAGGACGCAAGAAACCCAGCCTCCGGTCACCCACCCGACCCCAC

TCCACGCTCCACCACACATTAGTCATCAAACAAAACTTAGGACGAAAGGTCAATCACCATGTCCGCACAAAGGGAGAGGA

TCAATGCCTTCTACAAAGACAATCCTCACAATAAGAACCATAGGGTGATCCTGGATAGAGAACGCTTGGTCATTGAAAGA

CCCTACATCTTGCTTGGAGTCCTGCTGGTAATGTTCCTGAGTCTAATCGGACTGCTGGCCATTGCAGGGATCAGGCTTCA

CCGGGCCACCGTTGGAACTTCAGAGATCCAGAGTCGGCTGAATACCAATATTAAGTTGACCGAATCTATTGATCACCAGA

CTAAGGATGTCTTAACTCCCCTTTTTAAAATCATTGGCGATGAAGTCGGCATCAGAATTCCACAGAAATTCAGTGATCTT

GTCAAGTTCATCTCCGATAAGATTAAATTCCTCAACCCTGATAGAGAGTATGATTTCAGAGATCTCCGGTGGTGTATGAA

TCCCCCCGAGAGAGTCAAAATTAATTTTGATCAGTTTTGTGAGTACAAGGCTGCGGTTAAGTCAATTGAACATATATTTG

AGTCACCACTCAACAAGTCAAAAAAGCTGCAATCTTTGACTCTCGGGCCCGGAACAGGCTGTCTAGGCAGGACAGTAACA

AGAGCCCATTTCTCAGAACTTACAATGACCTTAATGGACCTGGATCTAGAGATGAAGCACAACGTGTCCTCAGTGTTTAC

CGTAGTTGAAGAGGGATTATTCGGAAGAACATATACCGTCTGGAGATCCGATGCCAGGGATCCGAGCACCGATCTAGGTA

TCGGCCATTTTTTAAGAGTCTTCGAGATTGGACTGGTAAGAGATCTTGGGCTGGGTCCCCCTGTTTTTCATATGACCAAC

TATCTCACGGTGAACATGAGTGATGACTATCGGAGATGTCTTTTAGCGGTAGGGGAGTTGAAGTTGACAGCCCTATGCAC

CTCATCTGAGACTGTGACACTGAGTGAGAGAGGAGTTCCAAGGAGGGAACCTCTTGTGGTTGTGATACTTAATCTAGCTG

GACCCACTCTAGGGGGAGAGCTATACAGTGTCTTGCCTACCTCTGATCTCATGGTGGAGAAACTCTATTTGTCTTCACAT

AGAGGGATCATCAAAGATGATGAGGCCAATTGGGTAGTGCCGTCTACCGATGTTCGTGATCTTCAGAACAAAGGTGAATG

TCTGGTGGAAGCATGCAAGACTCGACCTCCTTCATTTTGCAATGGCACAGGATCAGGCCCGTGGTCAGAAGGGAGAATCC

CTGCCTACGGGGTGATCAGGGTCAGTCTTGACTTAGCGAGTGACCCGGATGTAGTTATCACTTCAGTGTTTGGCCCACTG

ATACCTCACCTATCCGGCATGGATCTTTACAACAACCCGTTTTCAAAAGCTGTATGGTTGGCTGTACCACCTTATGAGCA

GTCATTTCTAGGAATGATAAATACAATTGGATTCCCTAACAGAGCAGAGGTTATGCCGCACATTTTGACCACAGAGATCA

GAGGCCCTCGGGGTCGTTGCCATGTTCCCATAGAATTGTCCCGCCGGGTTGATGACGATATCAAGATCGGGTCCAACATG

GTCATATTGCCGACGATGGACCTGAGGTATATTACAGCCACTTATGATGTTTCCAGGAGCGAGCATGCAATCGTGTACTA

TATCTATGACACGAGTCGCTCATCATCTTACTTCTACCCAGTTCGACTGAATTTCAAAGGCAATCCTCTCTCTTTGAGGA

TAGAGTGTTTCCCCTGGCGTCATAAGGTGTGGTGCTACCATGATTGTCTTATATACAACACCATAACAGGTGAAGAGGTC

CATACGAGAGGGCTGACCGGCATAGAGGTAACATGCAATCCAGTCTGAGCAGAGCTACGACCGTCACTCAGGCAGTCCCT

TGAGTCGCCACCGAGTCCAAGCAGCACAGCCTGGGACAATCAACGGCACAGCCCAGCCAACAATGTTATAAAAAACTTAG

GAGCCAGGGTTGTAGGGGCCATGGACTCTCTATCAGTCAATCAGGTCTTGTACCCTGAGGTTCATCTAGACAGCCCTATT

GTCACAAACAAACTAGTTGCCATCCTTGAGTACTCGGGTATCGACCATAACTATGTTCTTGAAGACCAGACTCTTGTCAA

GAATATTAGGTATAGGCTGGGGTGCGGTTTTTCAAATCAAATGATCATCAATAACAGGGGGGTGGGTGAAACAGTCAATT

CTAAACTTAAAAGTTACCCCCATAATCGTCATATCATATACCCGGATTGCAATAAGGAGTTGTTTTGTATCAAGGATAGC

TGCATATCTAAGAAGCTCTCGGAGCTATTCAAGAAGGGTAATTCCTTGTACTCTAAGATAAGTCACCAGGTACTGGATTG

TCTTAAGAGAGTCAATGGGAAATTAGGTCTGGGCACAGATCTTACCCATGGTCTGAAGGAGGGTATCCTTGACCTAGGGT

TGCACATGCATAGCTCTCAATGGTTCGAGACTTTCCTGTTCTGGTTCACTATCAAGACAGAGATGAGATCAATGATCAAA

GAACAGTCCCATATATGCCACAAGAGGAGGTATAACCCGACTTTTGTGTCGGGGGATGCATTCGAGGTGCTCGTATCGCG

AGACCTCGTTGTGATAATTGATAAGAATACCCAGTATGTCTTCTACTTAACATTTGAGCTGGTCCTTATGTATTGTGATG

TCATAGAGGGTAGACTTATGACGGAGACAGCCATGGCCATAGACCAGAGATATTCAGAACTTCTAAGCCGGGTCAGATAC

TTGTGGGATCTTATTGATGGGTTCTTCCCAACACTAGGCAACACCACATACCAAATTGTTGCTCTGCTTGAACCATTATC

GTTGGCTTATCTTCAACTTCAGGATGTCACTCTGGAGTTAAGAGGTGCTTTTTTAGACCATTGCTTCAAAGAACTCTATG

AGATACTGGAACATTGCGGCATTGACACAGCAGGCACTTACAATTCCATCACTGAAGGATTGGATTACGTATTCATCACC

CACGATATACATCTAACTGGGGAGATTTTTTCATTTTTCCGGAGTTTCGGACACCCCCGTCTAGAAGCAGTCACCGCTGC

GGAAAATGTCAGAAAACATATGAACCAACCGAAGGTAATCAGTTATGAGACTATGATGAAAGGCCATGCGGTATTTTGCG

GGATAATCATAAATGGTTTTCGGGACCGACATGGCGGCAGCTGGCCCCCTGTTGCACTGCCAGAACATGCCTCTGCTGCG

ATCCGGAATGCGCAGGCATCAGGTGAGGGACTGACCCATGACCTGTGTATAGACAACTGGAAATCCTTTGTTGGATTCAA

ATTTGGCTGCTTTATGCCACTCAGCCTAGATAGTGATTTGACCATGTATCTCAAAGACAAGGCATTGGCTGCACTGAAGA

ATGAGTGGGATTCAGTTTACCCGAAAGAATACCTCCGTTATAACCCACCTAGAGGGACAGAGTCAAGGCGATTGGTAGAG

GTATTCCTGAATGACTCCAGCTTTGATCCTTATAACATGATAATGTACGTGGTGAATGGCTCCTACCTTAAAGACCCTGA

GTTTAACCTCTCATACAGCCTAAAAGAGAAGGAGATAAAGGAGACAGGGCGGTTGTTTGCCAAAATGACCTATAAGATGC

GGGCCTGTCAGGTAATCGCTGAAAATCTGATATCGAATGGTGTTGGGAAGTATTTCCGAGACAATGGGATGGCAAAGGAC

GAGCATGACCTAACAAAAGCCCTTCACACCCTGGCAGTCTCAGGTGTTCCCAAAAACAACAAAGATAACCACCGAGGTGG

GCCTCCCAGAAGGACAACAAACCGAGGGGTGAGATCAAGCCAAGGCACCAAAACACAAGATAGAGACAAGGTTCAAGGGG

GACCTATGTACAACTATTTGCGATGCCAACCGATCAGCCCTGATCAGGGTGAGTCATACGAGACTGTTAGTGCATTCATC

ACCGCTGACCTTAAGAAGTATTGCCTGAATTGGAGATACGAGACAATCAGCATATTTGCACAGAGGCTGAATGAAATATA

TGGACTACCATCCTTCTTTCAATGGTTACACAGGATATTGGAAAAATCCGTACTCTACGTCAGTGACCCACATTGTCCTC

CCGATCTAGATAATCATATCCCTCTGGACAGTGTCCCCAATGCCCAAATATTCATCAAGTACCCAATGGGCGGAATAGAA

GGTTATTGCCAAAAACTATGGACAATCAGCACTATACCATACTTGTATCTGGCAGCCTATGAGAGCGGAGTAAGAATCGC

CTCACTGGTGCAGGGTGACAATCAGACAATCGCAGTGACAAAAAGAGTTCCAAGTTCTTGGCCTTATTCACTAAAAAAGA

GGGAGGCCTCCAAGGCCGCTCAAAATTACTTCGTAGTCTTAAGGCAAAGATTGCACGATGTAGGTCATCACTTAAAGGCT

AATGAGACCATAGTATCTTCTCACTTTTTTGTATATTCCAAAGGGATTTATTATGACGGCCTGCTAGTCTCACAATCACT

AAAGAGCATCGCCAGATGTGTCTTCTGGTCCGAGACTATAGTGGATGAAACCAGAGCGGCGTGCAGTAATATTGCAACAA

CTGTCGCTAAGAGTATAGAGAGGGGTTATGATAGGTACCTTGCATATTCTTTGAATATCCTCAAGATTTTCCAACAGATC

CTTATATCCCTTAACTTCACTATTAACACAACAATGACTCAGGATGTCGTGGCACCGATCATCGAGAACGGTGATTTGCT

GATAAGGATGGCACTCTTGCCAGCACCCATCGGGGGTCTCAATTATCTTAACATGAGCAGGTTATTTGTGAGAAATATCG

GTGACCCGGTCACCTCCTCTATAGCCGACCTGAAGAGGATGATAGATGCCGGGCTAATGCCAGAAGAAACATTGCATCAA

GTGATGACCCAGACCCCGGGAGAATCATCCTACCTAGATTGGGCAAGTGACCCTTATTCTGCCAACCTACCCTGCGTACA

GAGTATCACTCGCCTTCTCAAAAACATCACTGCACGATATATTTTAATCAGCAGCCCAAACCCGATGCTGAAAGGGTTAT

TTCATGAGGGGAGTAGAGATGAAGACGAGGAGCTTGCAAGTTTCCTAATGGATCGGCATATAATTGTTCCAAGAGCTGCA

CATGAAATCTTAGACCATAGTATAACCGGAGCAAGAGAAGCTATAGCCGGGATGTTGGACACCACCAAGGGTCTGATTAG

AACAAGTATGAAACGGGGCGGCCTCACCCCTCGAGTCTTAGCCCGCCTTTCCAATTATGATTATGAACAATTCCGATCCG

GGATAACACTATTAACAAAGAAAGGGCAGTGTTATCTCATTGACAAGGACTCGTGCTCGGTGCAGCTCGCTATAGCTCTG

AGGGGCCACATGTGGGCTAAGTTGGCTCGCGGGAGACCAATCTATGGGTTGGAGGTGCCTGATGTATTAGAATCGATGAA

CGGCTACCTTATCAAACGCCACGAGTCGTGTGCTATCTGTGAAACGGGCTCAAGTCACTACGGATGGTTTTTCGTCCCTG

CAGGGTGCCAGCTTGACGATGTATCTAGAGAGACTTCGGCTCTTCGTGTACCTTATGTCGGATCAACAACTGAGGAAAGG

ACAGACATGAAACTTGCATTTGTTAGGTCTCCAAGTAGATCCCTCAAATCAGCAGTTAGAATTGCAACAGTTTACTCATG

GGCTTACGGGGATGATGAGAAATCATGGAGTGAAGCCTGGATGCTAGCTAGGCAGAGAGCCAATATCACCTTAGATGAAT

TGAGAATGATCACTCCAGTCTCCACATCCACCAACTTAGCCCACCGGTTGAGGGATCGGAGTACCCAGGTGAAATACTCG

GGGACATCCCTTGTGAGGGTTGCAAGATACACAACCATCTCCAATGATAACTTGTCGTTTGTGATATCTGAGAAAAAAGT

GGATACCAACTTCATTTATCAGCAAGGGATGTTGCTTGGTCTTGGGATCCTTGAGAACCTCTTTAGGCTAGAGGCCACCA

CCGGAGTATCCAACACAGTGCTACACCTGCACGTGGAAACAGAATGTTGTGTTGTACCTATGGTGGATCACCCAAGGATA

CCGAGTCTCCGTAATATAAAAGTTACGGATGAGCTATGCACAAACCCTTTGATCTACGACAGGTCCCCCATCATAGAGCA

CGATGCAACCCGGCTATATTCACAGAGCCACAGGAGACATTTGGTGGAGTTTGTTACCTGGTCAACAAGCCAGCTTTACC

ATATACTGGCTAAGTCTACAGCAATGTCCATGATTGAATTGATCACGAGATTCGAGAAAGATCACATGAATGAAATAGCC

GCCCTGATTGGCGATGACGACATCAACAGCTTCATCACAGAATTTTTGCTAGTGGAGCCTAGATTATTTATAGTTTACCT

TGGTCAGTGTGCTGCCATCAATTGGGCCTTTGATATACATTATCATCGGCCCTCGGGCAAGTACCAGATGGGGGAACTTC

TCTACTCTCTGCTCTCTCGGATGAGCAAAGGAGTATATAAGATCTTTACTAATGCTTTGAGCCACCCCAAAGTTTACAAG

AAATTTTGGCGAAGTGGCATCATTGAGCCGATTCATGGCCCATCCTTGGATACACAGAATTTACATGTCACTGTCTGTGA

CATGATATATGGGTCCTATGTCACCTATTTGGATCTTTTGCTGAATGATGAGCTAGATGATTACCCGTATTTGCTCTGCG

AGAGTGATGAGGACGTGGTCACAGACAGGTTCGACAACATTCAAGCCAAACATCTCTGTGTATTGGCCGATGTATATTGC

AGCTCCAAGAGGTGTCCTTCAATAATCGGGATGTCTCCTATAGAAAAATGTACCATTCTCACACATTACATCAAGGGAGA

ATCAGTACAATCCCCGTCTGGGACCTCATGGAACACTGATCCCCTTGTAGTAGATCATTACTCATGCTCTCTGACCTACC

TTCGCCGCGGTTCCATCAAACAAATCAGGTTGAGGGTGGATCCTGGGTTTGTATTCGAGGCGTTGACAGACGTCGACTTT

AAACAACCTCGCAAAGCTAAGTTAGATATATCGGTCGTTGGGTTGACTGATTTCTCTCCCCCTTACGACAACGTCGGTGA

TTTTTTAGGGACTATCAACACATTGAGGCACGATCTGCCTGTCACCGGAACCGGGGTCTCGAACTATGAAGTCCACGCTT

ATCGTAGAATTGGCCTGAATTCATCAGCATGTTATAAAGCCGTAGAGATCTCCACGTTAATCAAGCCATCCTTAGAAGTC

GGAGAGCATGGATTGTTCTTAGGAGAAGGTTCCGGTTCAATGCTGGCTGCGTACAAAGAAGTTCTTAAATTAGCAAATTG

TTACTACAACAGCGGAGTAACAGCGGAGGGCAGAGCCGGACAGAGGGAAATATCTCCCTATCCTTCAGAGATGAGCCTAG

TAGAGAATCAAATGGGGATAGAGCGGAGTGTTAAAGTGCTGTTCAACGGGAAACCTGAAGTAACTTGGGTGGGGACCACA

GATTGCTACAAGTACATAATCAGTAACATCCAAACCTCTAGTCTGGGTTTCATACATTCAGATATTGAGACGCTCCCAAC

CAAAGATGCTGTTGAGAAATTGGAGGAATTTGCCTCTATCCTATCCTTATCCTTGATTTTGGGGAAAATCGGCTCTATTA

CAGTTGTCAAAATTATGCCCATTAGCGGGGATTTTACCCAAGGCTTCATAGCCTATGCCATCCAATATTTTAGGGAGAGC

CTGCTTGCCTATCCGAGATATAGTAACTTCATCTCGACTGAGTGTTACCTTATTATGATAGGATTAAAGGCCAATCGATT

GATAAACCCAGAAGCCATTAAGCARAGCATAATCAGAGCGGGGGTCAGGACTGCACCAGGACTTGTGAGCCATGTATTAT

CAGAGAAACAAAAAGGTTGTATTCAATCTTTCTTGGGTGATCCTTATATCCAAGGAGACTTCAATAAGCACCTTAAATCT

CTAACCTCTATTGAGAAAATCCTGGTAAATTGTGGTCTCTCGATCAATGGCATAAAAATCTGTAGAGATCTAATCCACCA

TGATATCGCCTCCGGTCCAGACGGCCTTATGAGCTCTACAATTATTTTATATAGGGAATTGGCTCATTTCAAAGATAATA

TAAGGAGTCAGCACGGAATGTTCCACCCCTACCCGGTACTGGCCAACAGCAGGCAACGTGAATTAATCCTTCGAATAGCC

AAGAAATTCTGGGGGTATGTCTTGCTATATTCTGATGACCCGGCACTAATCAGACAAACAATCAAGAACTTGAAACGGAA

TCATCTAACCTTTGACTTACATAGTAACCCATTTATTAAAGGGCTATCCAAGGCTGAGAAACTGCTGGTGCGGACGAGTT

CACTAAGAAGAGAATGGTTGTTCACTCTCGAAACGAAAGAAGTAAAAGAGTGGTTCAAATTGGTAGGTTACAGTGCACTC

ATCAGAGGCTAGTCAGCTATGCATCTGCCCCCTTCTCCTCCGCCATGAGACCCCACTGGCGATCCGGAAGATTAAAGAAA

ACTACATATTGGATAAGTATCTATTCCCAGCTTTGTCTGGT

>MF443337.1/goat/China/2014-04-05

ACCAAACAAAGTTGGGTAAGGATAGATCTTATAATAACTATGGACTGGCAAACTTAGGAGTAAAGATCCTACTGTCGGGG

GGAGGAGGAGGAGCAAGATCCTTGATCATGGCGACTCTCCTTAAAAGCTTAGCATTGTTCAAAAGGAACAAAGACAAAGC

GCCGACAGCATCAGGTTCAGGAGGGGCCATCCGGGGGATTAAGAATGTTATCATAGTCCCGATTCCCGGAGACTCGTCCA

TCACTACCCGTTCAAGACTGCTCGACAGGCTTGTCAGATTGGCCGGAGATCCTGACATCAACGGGTCAAAGCTGACCGGG

GTGATGATCAGCATGTTATCCTTGTTCGTAGAGTCACCCGGGCAATTGATACAGCGAATCACAGATGATCCGGATGTTAG

TATCCGCCTTGTTGAGGTAGTTCAAAGTACTAGGTCCCAGTCCGGGTTGACCTTTGCATCACGTGGTGCTGATTTAGACA

ACGAGGCAGACATGTATTTTTCAACTGAGGGGCCCTCGAGTGGAGGTAAGAAAAGGATCAACTGGTTTGAGAACAGAGAA

ATAATAGACATAGAGGTGCAGGATCCAGAAGAGTTCAATATGTTGTTAGCCTCCATACTAGCACAAGTCTGGATCCTTCT

GGCCAAGGCTGTTACGGCACCAGATACGGCAGCTGACTCAGAACTGAGAAGGTGGGTTAAATACACACAACAAAGGAGAG

TGATTGGGGAATTTCGCCTTGACAAGGGGTGGCTGGACGCAGTCCGCAACAGAATTGCAGAAGATCTATCACTCCGGCGG

TTCATGGTATCTCTCATACTTGACATCAAGAGGACCCCAGGCAACAAGCCAAGGATTGCAGAAATGATCTGCGACATTGA

CAACTATATTGTCGAAGCAGGACTCGCCAGTTTCATCCTTACTATCAAATTTGGTATCGAAACCATGTATCCTGCATTAG

GGCTCCACGAGTTTGCCGGAGAATTGTCCACTATTGAGTCCTTGATGAACTTGTATCAACAGCTAGGAGAGGTTGCACCC

TACATGGTAATTCTAGAGAATTCAATTCAGAACAAGTTTAGTGCAGGAGCTTATCCCCTCCTCTGGAGCTATGCGATGGG

TGTCGGAGTCGAGCTGGAGAACTCAATGGGGGGCTTGAACTTTGGCAGGTCATATTTTGACCCTGCCTATTTTCGTCTCG

GACAGGAGATGGTCAGAAGATCTGCAGGAAAGGTCAGCTCTGTGATTGCGGCTGAGCTCGGCATCACAGCAGAGGAAGCT

AAACTAGTCTCGGAAATCGCCTCGCAGGCTGGGGACGAAAGAACCGCCAGAGGGACTGGGCCTCGACAGGCGCAGGTTTC

CTTCCTTCAGCACCAAACAGGAGGGGGAGAGTCGTCCGCACCAGCGACCAGAGAAGGGGTCAAAGCTGCGATCCCAAACG

GAGCTGAAGAAAGGGACAGAAAGCAAACACGCCCAGGAAGGCCCAGAGGAGAGACCTCCGGCCAACTGCTCCTGGACATC

ATGCCAGAGGATGAGATCTTGCGAGAGTCTGGTCAAAACCCTCGTGAGGCTCAAAGATCGGCCGAGGCACTCTTCAGGCT

GCAGGCCATGGCCAAGATTCTGGAGGACCAGGAGGAGGGAGAAGACAACAGTCAGGTCTACAACGACAAGGATCTCCTCG

GCTAAGCAGACGCACCCTCTGTCGAAATCAGTGACAAGACATCTCCTACCAGTATTATAAAAAACTTAGGACCCAGGTCC

AAGCAAGCACACATCGACACTCCAACCAGTCGAGCGGAGACCACCGATGGCAGAAGAACAAGCATACCATGTCAACAAGG

GGCTGGAATGTATCAAGTCTCTCAAAGCCTCACCCCCGGATCTATCCACCATCAGAGATACCATCGAGAGCTGGAGAGAG

GGGCTTAGCCCCTCGGGCCGTGCAACACCGAACCCTGATACGTCCGAGGGAGATCATCAGAATATCAACCAATCATGCTC

ACCAGCAATCGGACCAAACAAAGTCTACTTGTCTCCTGAAGATAATCTCGGATTTAGAGAGATCACTGGCAACGACTGTG

AGGCTGGGCTCGGAGGGGTCCAGAGAGAAGGATCCAACTCTCAAGTACAGCGTTACCATGTTTATAGTCACGGGGGTGAA

GAGATTGAAGGACTCGAGGATGCTGACTCTCTCGTGGTTCAAGCAGATCCTCCAGTTGCTAACGTCTTCAATGGAGGAGA

GGATGGATCTGACGACAGCGATGTGGACTCTGGCCCAGATGATCCCGGCAGAGATACTCTATATGACCGGGGATCTGTTG

CCGGCAATGGTATCGCTAGGTCCACAGATGTCGAAAAACTAGAAGGTGCTGATATTCAAGAAGTTCTTAACTCCCAGAAA

GGCAAAGGAGGAAGATTCCAGGGCGGGAAAACCTTGCGAGTCCCGGAAATACCCGATGTCAAGCACTCCAGACCATCAGC

CCAATCAATTAAAAAGGGCACAGACGGGAACTCAGTCTCATCTGGAACGGTGATAGAGTGTTTATCGATAAGTGGTGCAA

CCCAAACTGTGCCAGAGTCAAGATGGGAGTCATCAGAGCAAAATGCGTCTGTGGGGAGTGTCCTCAAGTCTGCGAGGAGT

GCAAAGACGATCCAGGGGTCGACACAAGAATCTGGTACCATAGCATCACTGACTCAGCCTAAAGAGAATGACTCCGAGTA

TGAGTATGAGGATGACCTATTTACAGAGATTCAGGACATCCGTGCAAGCATTGCCAAGATCCATGATGACAATAAAACTA

TCCTCTCAAAGCTTGATTCTATACTGTTATTGAAAGGGGAAGTCGACACTATCAAGAAACAAATCAGCAAGCAGAATATA

AGTATATCCACCATTGAGGGCCATCTCTCCAGTATAATGATAGCCATCCCTGGCTTTGGGAAGGACATCAAGGACCCAAC

ATCCGAAGTCGAGTTGAACCCAGATCTAAGACCTATAATAAGCCGTGACTCTGGCAGAGCTCTCGCGGAGGTCCTCAAGA

AACCCGCTGTGGATAGGTCTCAGAAAATTGGAACCAAAGCCAACTCCAGCTCAAAGGGTCAGCTTCTTAAGGATCTCCAG

CTAAAACCTGTCGACAAGCAGGCAAGCTCTGCAATCGGGTTTGTCCCGTCCGACCATGAATCATCCAGAAATGTCATCCG

CTCCATAATCAAGTCGAGCAAGCTAAATATTGATCACAAGGACTATCTTCTAGATTTACTAAATGATGTGAAAGGCTCCA

AGGATCTTAAGGAATTTCACAAGATGCTAACAGCAATTCTCGCCAAGCACCCGTAACACATCCTCCAGTCATCATCTCAT

ACTCGACTAAAAACATCCTTTCAATCAGGCTATTACAAAAAACTTAGGAGCAAGGGCAACTGAGCTTCGCAGACAGGATG

ACCGAGATCTACGACTTCGATAAATCAGCATGGGATGTCAAAGGGTCAATTGCCCGCATAGAACCCACCACCTACCACGA

CGGCCGACTGATACCCCAGGTGAGGGTCATCGATCCTGGTCTGGGAGACAGAAAAGATGAGTGCTTTATGTACCTGTTTC

TCCTAGGAGTGATTGAGGATAACGACCCCCTGTCTCCCCCCGTCGGAAGAACCTTTGGCTCTCTACCTCTAGGGGTTGGT

AGGTCAACTGCCAAGCCAGAAGAACTACTAAGGGAGGCCACAGAATTAGATATAGTGGTGAGGCGCACTGCAGGAGTAAA

TGAGAAACTGGTATTTTACAACAACACTCCGCTGTCCTTGTTAACGCCCTGGAAGAAAGTTCTGACAACCGGAAGTGTGT

TTAGCGCTAACCAGGTTTGCAATGCAGTCAACCTAGTCCCGCTTGATACTCCCCAGAGATTCAGGGTTGTGTACATGAGC

ATAACTAGATTATCAGACAATGGTTACTATAGTGTGCCCAGAAGAATGCTGGAGTTCCGCTCGGCCAATGCAGTAGCCTT

CAACATCTTGGTTACACTAAGAATTGAAAATGGCACAAACCCTAGAAGATACATAGTCGGCTCATGGGAGAATTCAGAGG

TCACATTTATGGTACACGTGGGCAACTTTAGGAGAAAGAAGAACGAAGTATACTCTGCTGATTACTGCAAAATGAAGATT

GAAAAGATGGGTTTAGTTTTTGCCCTGGGTGGAATAGGCGGAACAAGTCTCCATATTAGAAGCACCGGGAAAATGAGCAA

AACCCTCCATGCACAGCTGGGGTTCAAGAAAATCTTATGTTACCCCCTAATGGATGTCAATGAGGATCTTAACCGATATC

TCTGGCGGGCAGAGTGCCGAATAGTCAAAATCCAAGCTGTCTTACAGCCATCAGTACCCCAAGAGTTCCGTGTCTACGAT

GATGTCATCATCAACGATGATCAAGGCTTGTTCAAGATCTTGTAGTTCATTTGCAAAGGAACGAACCCACCCCCAGGAAG

GGGCCCCACCCTCCCCCG-CCAAAAAACACACTCCAAGGCTCCCCGGCACAGCGCCAACACAGCTACTCCCGCGATGGAA

CGAGGTCGGCC-CCCGAACAAATCCGGTCCAGGGGGGGGGGCCCCCCCCCTGACCAACCGGGCGCCGACCGAGCAGGCCG

GCGAGAAGCCCCCAAGAGGCCCCGACGAAGACCGAGCACCCCATCCCCCC-AAAAACGTCCCCC-AACACGCCCGAGGCT

AGCCACCGGACACCCCCAGACCCGCCACCGGCGAGACAGAACCCCAAAGGGGGACCCTCCAAACCAAAGGAATCACCGGG

GTAGGCTCCGCCGAACCAGCACAGACCCGACCCAAGCAAGGCGGACCCCCAACACGAAACTGCCCAGCACACCCTACTCA

TCATCAAACAAAACTTAGGGGCCAAGTCCATACGAGAGGCCCTAGTCCGTCAGGCGGAGGGACGCAGAAAGGAAGGAGAC

ACGACCGTCCCAGAAGGAGGACACCCAACCACCGAAACGCCCGCCCCCAGGGAAGGAG--CCA-CCCCCACGCAGCGAAG

GAGCGGGTGGGAGACCTCGGGGAGGGCCCGACCGGGACCCGGGGGCCGGCGGCGCCGGCGGTGAGCAGGCCGGGCGCATA

CCTGACCCCACCGGAGGATCCGAGGAGGGCCCGACCCCGCGTGGACCAAACGGCAGGGAGCCCAACCCCACCAAAAAGAG

AGGCGAGGCCCCCCCTCCCTCCCCCCCCGCACACAACCTGCAGACAGACCGAGGGAGGGCCATCCCCCAGCGAGACAGGC

GGCTCCCTCGCAGCCGAGCCGCAGCTCCCCAGAATAGGACACCCCCGCACACTCCGAACACCCCCTCGCCCCCCGCCACC

GGGACAGGGGCTCCACCACCACAGACCGGAGTCACCGACAGCCAGCCAGAGGTCCCCTCCAGTCCCGCCCGGCGCCCCCC

CGAAGAGGCCCCGGCAAGGCCCCCCAGCCCGACGTCCCCCGCACGCACCGAAAGAAGGACCCGCCCGAGAACGTGCGGCA

CATCCATGCGCAAACATCATGACGCGGGTCGCAATTTTGACATTTCTGTTTCTTTTCCCAAATGTTGTTGTGTGTCAGAT

TCACTGGGGCAATCTATCCAAGATCGGGATTGTAGGAACAGGGAGTGCCAGCTACAAGGTGATGACTAGGCCAAGCCACC

AGACTCTGGTTATAAAGTTAATGCCAAATATAACGGCCATCGACAATTGTACAAAGTCAGAGATTGCAGAGTACAAGAGA

TTGCTGATCACAGTGTTAAAGCCTGTAGAGGATGCTCTGTCGGTGATAACCAAGAATGTAAGACCAATTCAAACTCTAAC

ACCTGGGCGTAGAACCCGCCGTTTTGCTGGAGCTGTTCTGGCCGGGGTAGCACTTGGAGTTGCGACAGCCGCTCAGATAA

CTGCAGGAGTCGCCCTTCATCAATCATTGATGAACTCCCAAGCAATTGAGAGTTTAAAAACCAGTCTTGAGATGTCGAAT

CAGGCAATAGAAGAAATCAGACTTGCAAATAAGGAGACCATACTGGCAGTACAGGGCGTCCAGGATTATATCAACAATGA

GCTCGTCCCTTCTGTTCATAGAATGTCATGCGAGCTGGTAGGTCACAAGCTCGGCCTCAAGCTCCTTAGGTACTACACCG

AGATCCTGTCCATATTCGGGCCCAGTCTTCGAGACCCGATATCTGCCGAAATATCAATCCAGGCACTTAGTTATGCATTA

GGCGGAGACATTAATAAAATCCTGGACAAGCTTGGGTATAGCGGTGGGGATTTCCTTGCCATCCTAGAAAGCAAGGGAAT

AAAGGCCCGGGTTACATATGTGGACACAAGAGATTACTTTATAATCCTTAGCATCGCCTACCCAACCTTATCTGAGATCA

AGGGAGTGATAGTTCACAAGATAGAAGCTATAACATACAACATTGGGGCACAGGAGTGGTATACTACTATCCCTAAATAT

GTAGCCACTCAGGGGTATCTGATATCGAACTTTGATGAGACGTCATGCGTATTCACTCCAGAGGGGACAGTTTGCAGCCA

GAATGCGTTGTACCCAATGAGCCCATTGCTTCAGGAATGTTTCAGGGGGTCAACAAAATCGTGCGCCAGAACCTTAGTTT

CAGGGACCATAAGTAATAGATTTATCCTATCAAAAGGGAACCTGATTGCAAATTGTGCGTCAGTTTTGTGCAAATGTTAC

ACAACAGAGACAGTTATCAGCCAAGATCCCGACAAACTACTAACTGTTGTAGCATCCGACAAGTGTCCTGTAGTTGAGGT

GGATGGAGTGACAATACAGGTCGGCAGTCGAGAGTATCCGGATTCTGTATACTTACACAAAATAGACTTAGGTCCAGCCA

TCTCCCTAGAAAAACTGGATGTAGGCACCAATTTAGGCAATGCAGTCACAAGACTGGAGAATGCAAAGGAGCTCCTAGAT

GCATCAGACCAAATACTGAAGACTGTTAAAGGGGTACCTTTCAGTGGGAATATGTACATAGCACTGGCAGCTTGCATAGG

AGTATCCCTGGGCCTTGTCACATTAATATGCTGCTGTAAGGGGAGGTGTAAGAACAAGGAAATCCCTATCTCCAAAATCA

ACCCAGGGCTCAAACCCGACCTGACCGGGACCTCAAAGTCGTACGTAAGATCACTGTAGTCAGAATCACCTGGATCATCT

GGCATCACACACATACATGCACGACACAGGCAGTCCGAGGACGCAAGAAACCCAGCCTCCGGTCACCCACCCGACCCCAC

TCCACGCTCCACCACACATTAGTCATCAAACAAAACTTAGGACGAAAGGTCAATCACCATGTCCGCACAAAGGGAGAGGA

TCAATGCCTTCTACAAAGACAATCCTCACAATAAGAACCATAGGGTGATCCTGGATAGAGAACGCTTGGTCATTGAAAGA

CCCTACATCTTGCTTGGAGTCCTGCTGGTAATGTTCCTGAGTCTAATCGGACTGCTGGCCATTGCAGGGATCAGGCTTCA

CCGGGCCACCGTTGGAACTTCAGAGATCCAGAGTCGGCTGAATACCAATATTAAGTTGACCGAATCTATTGATCACCAGA

CTAAGGATGTCTTAACTCCCCTTTTTAAAATCATTGGCGATGAAGTCGGCATCAGAATTCCACAGAAATTCAGTGATCTT

GTCAAGTTCATCTCCGATAAGATTAAATTCCTCAACCCTGATAGAGAGTATGATTTCAGAGATCTCCGGTGGTGTATGAA

TCCCCCCGAGAGAGTCAAAATTAATTTTGATCAGTTTTGTGAGTACAAGGCTGCGGTTAAGTCAATTGAACATATATTTG

AGTCACCACTCAACAAGTCAAAAAAGCTGCAATCTTTGACTCTCGGGCCCGGAACAGGCTGTCTAGGCAGGACAGTAACA

AGAGCCCATTTCTCAGAACTTACAATGACCTTAATGGACCTGGATCTAGAGATGAAGCACAACGTGTCCTCAGTGTTTAC

CGTAGTTGAAGAGGGATTATTCGGAAGAACATATACCGTCTGGAGATCCGATGCCAGGGATCCGAGCACCGATCTAGGTA

TCGGCCATTTTTTAAGAGTCTTCGAGATTGGACTGGTAAGAGATCTTGGGCTGGGTCCCCCTGTTTTTCATATGACCAAC

TATCTCACGGTGAACATGAGTGATGACTATCGGAGATGTCTTTTAGCGGTAGGGGAGTTGAAGTTGACAGCCCTATGCAC

CTCATCTGAGACTGTGACACTGAGTGAGAGAGGAGTTCCAAGGAGGGAACCTCTTGTGGTTGTGATACTTAATCTAGCTG

GACCCACTCTAGGGGGAGAGCTATACAGTGTCTTGCCTACCTCTGATCTCATGGTGGAGAAACTCTATTTGTCTTCACAT

AGAGGGATCATCAAAGATGATGAGGCCAATTGGGTAGTGCCGTCTACCGATGTTCGTGATCTTCAGAATAAAGGTGAATG

TCTGGTGGAAGCATGCAAGACTCGACCTCCTTCATTTTGCAATGGCACAGGATCAGGCCCGTGGTCAGAAGGGAGAATCC

CTGCCTACGGGGTGATCAGGGTCAGTCTTGACTTAGCGAGTGACCCGGATGTAGTTATCACTTCAGTGTTTGGCCCACTG

ATACCTCACCTATCCGGCATGGATCTTTACAACAACCCGTTTTCAAAAGCTGTATGGTTGGCTGTACCACCTTATGAGCA

GTCATTTCTAGGAATGATAAATACAATTGGATTCCCTAACAGAGCAGAGGTTATGCCGCACATTTTGACCACAGAGATCA

GAGGCCCTCGGGGTCGTTGCCATGTTCCCATAGAATTGTCCCGCCGGGTTGATGACGATATCAAGATCGGGTCCAACATG

GTCATATTGCCGACGATGGACCTGAGGTATATTACAGCCACTTATGATGTTTCCAGGAGCGAGCATGCAATCGTGTACTA

TATCTATGACACGAGTCGCTCATCATCTTACTTCTACCCAGTTCGACTGAATTTCAAAGGCAATCCTCTCTCTTTGAGGA

TAGAGTGTTTCCCCTGGCGTCATAAGGTGTGGTGCTACCATGATTGTCTTATATACAACACCATAACAGGTGAAGAGGTC

CATACGAGAGGGCTGACCGGCATAGAGGTAACATGCAATCCAGTCTGAGCAGAGCTACGACCGTCACTCAGGCAGTCCCT

TGAGTCGCCACCGAGTCCAAGCAGCACAGCCTGGGACAATCAACGGCACAGCCCAGCCAACAATGTTATAAAAAACTTAG

GAGCCAGGGTTGTAGGGGCCATGGACTCTCTATCAGTCAATCAGGTCTTGTACCCTGAGGTTCATCTAGACAGCCCTATT

GTCACAAACAAACTAGTTGCCATCCTTGAGTACTCGGGTATCGACCATAACTATGTTCTTGAAGACCAGACTCTTGTCAA

GAATATTAGGTATAGGCTGGGGTGCGGTTTTTCAAATCAAATGATCATCAATAACAGGGGGGTGGGTGAAACAGTCAATT

CTAAACTTAAAAGTTACCCCCATAATCGTCATATCATATACCCGGATTGCAATAAGGAGTTGTTTTGTATCAAGGATAGC

TGCATATCTAAGAAGCTCTCGGAGCTATTCAAGAAGGGTAATTCCTTGTACTCTAAGATAAGTCACCAGGTACTGGATTG

TCTTAAGAGAGTCAATGGGAAATTAGGTCTGGGCACAGATCTTACCCATGGTCTGAAGGAGGGTATCCTTGACCTAGGGT

TGCACATGCATAGCTCTCAATGGTTCGAGACTTTCCTGTTCTGGTTCACTATCAAGACAGAGATGAGATCAATGATCAAA

GAACAGTCCCATATATGCCACAAGAGGAGGTATAACCCGACTTTTGTGTCGGGGGATGCATTCGAGGTGCTCGTATCGCG

AGACCTCGTTGTGATAATTGATAAGAATACCCAGTATGTCTTCTACTTAACATTTGAGCTGGTCCTTATGTATTGTGATG

TCATAGAGGGTAGACTTATGACGGAGACAGCCATGGCCATAGACCAGAGATATTCAGAACTTCTAAGCCGGGTCAGATAC

TTGTGGGATCTTATTGATGGGTTCTTCCCAACACTAGGCAACACCACATACCAAATTGTTGCTCTGCTTGAACCATTATC

GTTGGCTTATCTTCAACTTCAGGATGTCACTCTGGAGTTAAGAGGTGCTTTTTTAGACCATTGCTTCAAAGAACTCTATG

AGATACTGGAACATTGCGGCATTGACACAGCAGGCACTTACAATTCCATCACTGAAGGATTGGATTACGTATTCATCACC

CACGATATACATCTAACTGGGGAGATTTTTTCATTTTTCCGGAGTTTCGGACACCCCCGTCTAGAAGCAGTCACCGCTGC

GGAAAATGTCAGAAAACATATGAACCAACCGAAGGTAATCAGTTATGAGACTATGATGAAAGGCCATGCGGTATTTTGCG

GGATAATCATAAATGGTTTTCGGGACCGACATGGCGGCAGCTGGCCCCCTGTTGCACTGCCAGAACATGCCTCTGCTGCG

ATCCGGAATGCGCAGGCATCAGGTGAGGGACTGACCCATGACCTGTGTATAGACAACTGGAAATCCTTTGTTGGATTCAA

ATTTGGCTGCTTTATGCCACTCAGCCTAGATAGTGATTTGACCATGTATCTCAAAGACAAGGCATTGGCTGCACTGAAGA

ATGAGTGGGATTCAGTTTACCCGAAAGAATACCTCCGTTATAACCCACCTAGAGGGACAGAGTCAAGGCGATTGGTAGAG

GTATTCCTGAATGACTCCAGCTTTGATCCTTATAACATGATAATGTACGTGGTGAATGGCTCCTACCTTAAAGACCCTGA

GTTTAACCTCTCATACAGCCTAAAAGAGAAGGAGATAAAGGAGACAGGGCGGTTGTTTGCCAAAATGACCTATAAGATGC

GGGCCTGTCAGGTAATCGCTGAAAATCTGATATCGAATGGTGTTGGGAAGTATTTCCGAGACAATGGGATGGCAAAGGAC

GAGCATGACCTAACAAAAGCCCTTCACACCCTGGCAGTCTCAGGTGTTCCCAAAAACAACAAAGATAACCACCGAGGTGG

GCCTCCCAGAAGGACAACAAACCGAGGGGTGAGATCAAGCCAAGGCACCAAAACACAAGATAGAGACAAGGTTCAAGGGG

GACCTATGTACAACTATTTGCGATGCCAACCGATCAGCCCTGATCAGGGTGAGTCATACGAGACTGTTAGTGCATTCATC

ACCGCTGACCTTAAGAAGTATTGCCTGAATTGGAGATACGAGACAATCAGCATATTTGCACAGAGGCTGAATGAAATATA

TGGACTACCATCCTTCTTTCAATGGTTACACAGGATATTGGAAAAATCCGTACTCTACGTCAGTGACCCACATTGTCCTC

CCGATCTAGATAATCATATCCCTCTGGACAGTGTCCCCAATGCCCAAATATTCATCAAGTACCCAATGGGCGGAATAGAA

GGTTATTGCCAAAAACTATGGACAATCAGCACTATACCATACTTGTATCTGGCAGCCTATGAGAGCGGAGTAAGAATCGC

CTCACTGGTGCAGGGTGACAATCAGACAATCGCAGTGACAAAAAGAGTTCCAAGTTCTTGGCCTTATTCACTAAAAAAGA

GGGAGGCCTCCAAGGCCGCTCAAAATTACTTCGTAGTCTTAAGGCAAAGATTGCACGATGTAGGTCATCACTTAAAGGCT

AATGAGACCATAGTATCTTCTCACTTTTTTGTATATTCCAAAGGGATTTATTATGACGGCCTGCTAGTCTCACAATCACT

AAAGAGCATCGCCAGATGTGTCTTCTGGTCCGAGACTATAGTGGATGAAACCAGAGCGGCGTGCAGTAATATTGCAACAA

CTGTCGCTAAGAGTATAGAGAGGGGTTATGATAGGTACCTTGCATATTCTTTGAATATCCTCAAGATTTTCCAACAGATC

CTTATATCCCTTAACTTCACTATTAACACAACAATGACTCAGGATGTCGTGGCACCGATCATCGAGAACGGTGATTTGCT

GATAAGGATGGCACTCTTGCCAGCACCCATCGGGGGTCTCAATTATCTTAACATGAGCAGGTTATTTGTGAGAAATATCG

GTGACCCGGTCACCTCCTCTATAGCCGACCTGAAGAGGATGATAGATGCCGGGCTAATGCCAGAAGAAACATTGCATCAA

GTGATGACCCAGACCCCGGGAGAATCATCCTACCTAGATTGGGCAAGTGACCCTTATTCTGCCAACCTACCCTGCGTACA

GAGTATCACTCGCCTTCTCAAAAACATCACTGCACGATATATTTTAATCAGCAGCCCAAACCCGATGCTGAAAGGGTTAT

TTCATGAGGGGAGTAGAGATGAAGACGAGGAGCTTGCAAGTTTCCTAATGGATCGGCATATAATTGTTCCAAGAGCTGCA

CATGAAATCTTAGACCATAGTATAACCGGAGCAAGAGAAGCTATAGCCGGGATGTTGGACACCACCAAGGGTCTGATTAG

AACAAGTATGAAACGGGGCGGCCTCACCCCTCGAGTCTTAGCCCGCCTTTCCAATTATGATTATGAACAATTCCGATCCG

GGATAACACTATTAACAAAGAAAGGGCAGTGTTATCTCATTGACAAGGACTCGTGCTCGGTGCAGCTCGCTATAGCTCTG

AGGGGCCACATGTGGGCTAAGTTGGCTCGCGGGAGACCAATCTATGGGTTGGAGGTGCCTGATGTATTAGAATCGATGAA

CGGCTACCTTATCAAACGCCACGAGTCGTGTGCTATCTGTGAAACGGGCTCAAGTCACTACGGATGGTTTTTCGTCCCTG

CAGGGTGCCAGCTTGACGATGTATCTAGAGAGACTTCGGCTCTTCGTGTACCTTATGTCGGATCAACAACTGAGGAAAGG

ACAGACATGAAACTTGCATTTGTTAGGTCTCCAAGTAGATCCCTCAAATCAGCAGTTAGAATTGCAACAGTTTACTCATG

GGCTTACGGGGATGATGAGAAATCATGGAGTGAAGCCTGGATGCTAGCTAGGCAGAGAGCCAATATCACCTTAGATGAAT

TGAGAATGATCACTCCAGTCTCCACATCCACCAACTTAGCCCACCGGTTGAGGGATCGGAGTACCCAGGTGAAATACTCG

GGGACATCCCTTGTGAGGGTTGCAAGATACACAACCATCTCCAATGATAACTTGTCGTTTGTGATATCTGAGAAAAAAGT

GGATACCAACTTCATTTATCAGCAAGGGATGTTGCTTGGTCTTGGGATCCTTGAGAACCTCTTTAGGCTAGAGGCCACCA

CCGGAGTATCCAACACAGTGCTACACCTGCACGTGGAAACAGAATGTTGTGTTGTACCTATGGTGGATCACCCAAGGATA

CCGAGTCTCCGTAATATAAAAGTTACGGATGAGCTATGCACAAACCCTTTGATCTACGACAGGTCCCCCATCATAGAGCA

CGATGCAACCCGGCTATATTCACAGAGCCACAGGAGACATTTGGTGGAGTTTGTTACCTGGTCAACAAGCCAGCTTTACC

ATATACTGGCTAAGTCTACAGCAATGTCCATGATTGAATTGATCACGAGATTCGAGAAAGATCACATGAATGAAATAGCC

GCCCTGATTGGCGATGACGACATCAACAGCTTCATCACAGAATTTTTGCTAGTGGAGCCTAGATTATTTATAGTTTACCT

TGGTCAGTGTGCTGCCATCAATTGGGCCTTTGATATACATTATCATCGGCCCTCGGGCAAGTACCAGATGGGGGAACTTC

TCTACTCTCTGCTCTCTCGGATGAGCAAAGGAGTATATAAGATCTTTACTAATGCTTTGAGCCACCCCAAAGTTTACAAG

AAATTTTGGCGAAGTGGCATCATTGAGCCGATTCATGGCCCATCCTTGGATACACAGAATTTACATGTCACTGTCTGTGA

CATGATATATGGGTCCTATGTCACCTATTTGGATCTTTTGCTGAATGATGAGCTAGATGATTACCCGTATTTGCTCTGCG

AGAGTGATGAGGACGTGGTCACAGACAGGTTCGACAACATTCAAGCCAAACATCTCTGTGTATTGGCCGATGTATATTGC

AGCTCCAAGAGGTGTCCTTCAATAATCGGGATGTCTCCTATAGAAAAATGTACCATTCTCACACATTACATCAAGGGAGA

ATCAGTACAATCCCCGTCTGGGACCTCATGGAACACTGATCCCCTTGTAGTAGATCATTACTCATGCTCTCTGACCTACC

TTCGCCGCGGTTCCATCAAACAAATCAGGTTGAGGGTGGATCCTGGGTTTGTATTCGAGGCGTTGACAGACGTCGACTTT

AAACAACCTCGCAAAGCTAAGTTAGATATATCGGTCGTTGGGTTGACTGATTTCTCTCCCCCTTACGACAACGTCGGTGA

TTTTTTAGGGACTATCAACACATTGAGGCACGATCTGCCTGTCACCGGAACCGGGGTCTCGAACTATGAAGTCCACGCTT

ATCGTAGAATTGGCCTGAATTCATCAGCATGTTATAAAGCCGTAGAGATCTCCACGTTAATCAAGCCATCCTTAGAAGTC

GGAGAGCATGGATTGTTCTTAGGAGAAGGTTCCGGTTCAATGCTGGCTGCGTACAAAGAAGTTCTTAAATTAGCAAATTG

TTACTACAACAGCGGAGTAACAGCGGAGGGCAGAGCCGGACAGAGGGAAATATCTCCCTATCCTTCAGAGATGAGCCTAG

TAGAGAATCAAATGGGGATAGAGCGGAGTGTTAAAGTGCTGTTCAACGGGAAACCTGAAGTAACTTGGGTGGGGACCACA

GATTGCTACAAGTACATAATCAGTAACATCCAAACCTCTAGTCTGGGTTTCATACATTCAGATATTGAGACGCTCCCAAC

CAAAGATGCTGTTGAGAAATTGGAGGAATTTGCCTCTATCCTATCCTTATCCTTGATTTTGGGGAAAATCGGCTCTATTA

CAGTTGTCAAAATTATGCCCATTAGCGGGGATTTTACCCAAGGCTTCATAGCCTATGCCATCCAATATTTTAGGGAGAGC

CTGCTTGCCTATCCGAGATATAGTAACTTCATCTCGACTGAGTGTTACCTTATTATGATAGGATTAAAGGCCAATCGATT

GATAAACCCAGAAGCCATTAAGCAAGGCATAATCAGAGCGGGGGTCAGGACTGCACCAGGACTTGTGAGCCATGTATTAT

CAGAGAAACAAAAAGGTTGTATTCAATCTTTCTTGGGTGATCCTTATATCCAAGGAGACTTCAATAAGCACCTTAAATCT

CTAACCTCTATTGAGAAAATCCTGGTAAATTGTGGTCTCTCGATCAATGGCATAAAAATCTGTAGAGATCTAATCCACCA

TGATATCGCCTCCGGTCCAGACGGCCTTATGAGCTCTACAATTATTTTATATAGGGAATTGGCTCATTTCAAAGATAATA

TAAGGAGTCAGCACGGAATGTTCCACCCCTACCCGGTACTGGCCAACAGCAGGCAACGTGAATTAATCCTTCGAATAGCC

AAGAAATTCTGGGGGTATGTCTTGCTATATTCTGATGACCCGGCACTAATCAGACAAACAATCAAGAACTTGAAACGGAA

TCATCTAACCTTTGACTTACATAGTAACCCATTTATTAAAGGGCTATCCAAGGCTGAGAAACTGCTGGTGCGGACGAGTT

CACTAAGAAGAGAATGGTTGTTCACTCTCGAAACGAAAGAAGTAAAAGAGTGGTTCAAATTGGTAGGTTACAGTGCACTC

ATCAGAGGCTAGTCAGCTATGCATCTGCCCCCTTCTCCTCCGCCATGAGACCCCACTGGCGATCCAGAAGATTAAAGAAA

ACTACATATTGGATAAGTATCTATTCCCAGCTTTGTCTGGT

>KX421385.1/goat/China/2013-12-21

ACCAAACAAAGTTGGGTAAGGATAGATCTTATAATAACTATGGACTGGCAAACTTAGGAGTAAAGATCCTACTGTCGGGG

GGAGGAGGAGGAGCAAGATCCTTGATCATGGCGACTCTCCTTAAAAGCTTAGCATTGTTCAAAAGGAACAAAGACAAAGC

GCCGACAGCATCAGGTTCAGGAGGGGCCATCCGGGGGATTAAGAATGTTATCATAGTCCCGATTCCCGGAGACTCGTCCA

TCACTACCCGTTCAAGACTGCTCGACAGGCTTGTCAGATTGGCCGGAGATCCTGACATCAACGGGTCAAAGCTGACCGGG

GTGATGATCAGCATGTTATCCTTGTTCGTAGAGTCACCCGGGCAATTGATACAGCGAATCACAGATGATCCGGATGTTAG

TATCCGCCTTGTTGAGGTAGTTCAAAGTACTAGGTCCCAGTCCGGGTTGACCTTTGCATCACGTGGTGCTGATTTAGACA

ACGAGGCAGACATGTATTTTTCAACTGAGGGGCCCTCGAGTGGAGGTAAGAAAAGGATCAACTGGTTTGAGAACAGAGAA

ATAATAGACATAGAGGTGCAGGATCCAGAAGAGTTCAATATGTTGTTAGCCTCCATACTAGCACAAGTCTGGATCCTTCT

GGCCAAGGCTGTTACGGCACCAGATACGGCAGCTGACTCAGAACTGAGAAGGTGGGTTAAATACACACAACAAAGGAGAG

TGATTGGGGAATTTCGCCTTGACAAGGGGTGGCTGGACGCAGTCCGCAACAGAATTGCAGAAGATCTATCACTCCGGCGG

TTCATGGTATCTCTCATACTTGACATCAAGAGGACCCCAGGCAACAAGCCAAGGATTGCAGAAATGATCTGCGACATTGA

CAACTATATTGTCGAAGCAGGACTCGCCAGTTTCATCCTTACTATCAAATTTGGTATCGAAACCATGTATCCTGCATTAG

GGCTCCACGAGTTTGCCGGAGAATTGTCCACTATTGAGTCCTTGATGAACTTGTATCAACAGCTAGGAGAGGTTGCACCC

TACATGGTAATTCTAGAGAATTCAATTCAGAACAAGTTTAGTGCAGGAGCTTATCCCCTCCTCTGGAGCTATGCGATGGG

TGTCGGAGTCGAGCTGGAGAACTCAATGGGGGGCTTGAACTTTGGCAGGTCATATTTTGACCCTGCCTATTTTCGTCTCG

GACAGGAGATGGTCAGAAGATCTGCAGGAAAGGTCAGCTCTGTGATTGCGGCTGAGCTCGGCATCACAGCAGAGGAAGCT

AAACTAGTCTCGGAAATCGCCTCGCAGGCTGGGGACGAAAGAACCGCCAGAGGGACTGGGCCTCGACAGGCGCAGGTTTC

CTTCCTTCAGCACCAAACAGGAGGGGGAGAGTCGTCCGCACCAGCGACCAGAGAAGGGGTCAAAGCTGCGATCCCAAACG

GAGCTGAAGAAAGGGACAGAAAGCAAACACGCCCAGGAAGGCCCAGAGGAGAGACCTCCGGCCAACTGCTCCTGGACATC

ATGCCAGAGGATGAGATCTTGCGAGAGTCTGGTCAAAACCCTCGTGAGGCTCAAAGATCGGCCGAGGCACTCTTCAGGCT

GCAGGCCATGGCCAAGATTCTGGAGGACCAGGAGGAGGGAGAAGACAACAGTCAGGTCTACAACGACAAGGATCTCCTCG

GCTAAGCAGACGCACCCTCTGTCGAAATCAGTGACAAGACATCTCCTACCAGTATTATAAAAAACTTAGGACCCAGGTCC

AAGCAAGCACACATCGACACTCCAACCAGTCGAGCGGAGACCACCGATGGCAGAAGAACAAGCATACCATGTCAACAAGG

GGCTGGAATGTATCAAGTCTCTCAAAGCCTCACCCCCGGATCTATCCACCATCAGAGATACCATCGAGAGCTGGAGAGAG

GGGCTTAGCCCCTCGGGCCGTGCAACACCGAACCCTGATACGTCCGAGGGAGATCATCAGAATATCAACCAATCATGCTC

ACCAGCAATCGGACCAAACAAAGTCTACTTGTCTCCTGAAGATAATCTCGGATTTAGAGAGATCACTGGCAACGACTGTG

AGGCTGGGCTCGGAGGGGTCCAGAGAGAAGGATCCAACTCTCAAGTACAGCGTTACCATGTTTATAGTCACGGGGGTGAA

GAGATTGAAGGACTCGAGGATGCTGACTCTCTCGTGGTTCAAGCAGATCCTCCAGTTGCTAACGTCTTCAATGGAGGAGA

GGATGGATCTGACGACAGCGATGTGGACTCTGGCCCAGATGATCCCGGCAGAGATACTCTATATGACCGGGGATCTGTTG

CCGGCAATGGTATCGCTAGGTCCACAGATGTCGAAAAACTAGAAGGTGCTGATATTCAAGAAGTTCTTAACTCCCAGAAA

GGCAAAGGAGGAAGATTCCAGGGCGGGAAAACCTTGCGAGTCCCGGAAATACCCGATGTCAAGCACTCCAGACCATCAGC

CCAATCAATTAAAAAGGGCACAGACGGGAACTCAGTCTCATCTGGAACGGTGATAGAGTGTTTATCGATAAGTGGTGCAA

CCCAAACTGTGCCAGAGTCAAGATGGGAGTCATCAGAGCAAAATGCGTCTGTGGGGAGTGTCCTCAAGTCTGCGAGGAGT

GCAAAGACGATCCAGGGGTCGACACAAGAATCTGGTACCATAGCATCACTGACTCAGCCTAAAGAGAATGACTCCGAGTA

TGAGTATGAGGATGACCTATTTACAGAGATTCAGGACATCCGTGCAAGCATTGCCAAGATCCATGATGACAATAAAACTA

TCCTCTCAAAGCTTGATTCTATACTGTTATTGAAAGGGGAAGTCGACACTATCAAGAAACAAATCAGCAAGCAGAATATA

AGTATATCCACCATTGAGGGCCATCTCTCCAGTATAATGATAGCCATCCCTGGCTTTGGGAAGGACATCAAGGACCCAAC

ATCCGAAGTCGAGTTGAACCCAGATCTAAGACCTATAATAAGCCGTGACTCTGGCAGAGCTCTCGCGGAGGTCCTCAAGA

AACCCGCTGTGGATAGGTCTCAGAAAATTGGAACCAAAGCCAACTCCAGCTCAAAGGGTCAGCTTCTTAAGGATCTCCAG

CTAAAACCTGTCGACAAGCAGGCAAGCTCTGCAATCGGGTTTGTCCCGTCCGACCATGAATCATCCAGAAATGTCATCCG

CTCCATAATCAAGTCGAGCAAGCTAAATATTGATCACAAGGACTATCTTCTAGATTTACTAAATGATGTGAAAGGCTCCA

AGGATCTTAAGGAATTTCACAAGATGCTAACAGCAATTCTCGCCAAGCACCCGTAACACATCCTCCAGTCATCATCTCAT

ACTCGACTAAAAACATCCTTTCAATCAGGCTATTACAAAAAACTTAGGAGCAAGGGCAACTGAGCTTCGCAGACAGGATG

ACCGAGATCTACGACTTCGATAAATCAGCATGGGATGTCAAAGGGTCAATTGCCCGCATAGAACCCACCACCTACCACGA

CGGCCGACTGATACCCCAGGTGAGGGTCATCGATCCTGGTCTGGGAGACAGAAAAGATGAGTGCTTTATGTACCTGTTTC

TCCTAGGAGTGATTGAGGATAACGACCCCCTGTCTCCCCCCGTCGGAAGAACCTTTGGCTCTCTACCTCTAGGGGTTGGT

AGGTCAACTGCCAAGCCAGAAGAACTACTAAGGGAGGCCACAGAATTAGATATAGTGGTGAGGCGCACTGCAGGAGTAAA

TGAGAAACTGGTATTTTACAACAACACTCCGCTGTCCTTGTTAACGCCCTGGAAGAAAGTTCTGACAACCGGAAGTGTGT

TTAGCGCTAACCAGGTTTGCAATGCAGTCAACCTAGTCCCGCTTGATACTCCCCAGAGATTCAGGGTTGTGTACATGAGC

ATAACTAGATTATCAGACAATGGTTACTATAGTGTGCCCAGAAGAATGCTGGAGTTCCGCTCGGCCAATGCAGTAGCCTT

CAACATCTTGGTTACACTAAGAATTGAAAATGGCACAAACCCTAGAAGATACATAGTCGGCTCATGGGAGAATTCAGAGG

TCACATTTATGGTACACGTGGGCAACTTTAGGAGAAAGAAGAACGAAGTATACTCTGCTGATTACTGCAAAATGAAGATT

GAAAAGATGGGTTTAGTTTTTGCCCTGGGTGGAATAGGCGGAACAAGTCTCCATATTAGAAGCACCGGGAAAATGAGCAA

AACCCTCCATGCACAGCTGGGGTTCAAGAAAATCTTATGTTACCCCCTAATGGATGTCAATGAGGATCTTAACCGATATC

TCTGGCGGGCAGAGTGCCGAATAGTCAAAATCCAAGCTGTCTTACAGCCATCAGTACCCCAAGAGTTCCGTGTCTACGAT

GATGTCATCATCAACGATGATCAAGGCTTGTTCAAGATCTTGTAGTTCATTTGCAAAGGAACGAACCCACCCCCAGGAAG

GGGCCCCACCCTCCCCCG-CCAAAAAACACACTCCAAGGCTCCCCGGCACAGCGCCAACACAGCTACTCCCGCGATGGAA

CGAGGTCGGCC-CCCGAACAAATCCGGTCCAGGGGGGGGGGCCCCCCCCCTGACCAACCGGGCGCCGACCGAGCAGGCCG

GCGAGAAGCCCCCAAGAGGCCCCGACGAAGACCGAGCACCCCATCCCCCC-AAAAACGTCCCCC-AACACGCCCGAGGCT

AGCCACCGGACACCCCCAGACCCGCCACCGGCGAGACAGAACCCCAAGGGGGGACCCTCCAAACCAAAGGAATCACCGGG

GTAGGCTCCGCCGAACCAGCACAGACCCGACCCAAGCAAGGCGGACCCCCAACACGAAACTGCCCAGCACACCCTACTCA

TCATCAAACAAAACTTAGGGGCCAAGTCCATACGAGAGGCCCTAGTCCGTCAGGCGGAGGGACGCAGAAAGGAAGGAGAC

ACGACCGTCCCAGAAGGAGGACACCCAACCACCGAAACGCCCGCCCCCAGGGAAGGAG--CCA-CCCCCACGCAGCGAAG

GAGCGGGTGGGAGACCTCGGGGAGGGCCCGACCGGGACCCGGGGGCCGGCGGCGCCGGCGGTGAGCAGGCCGGGCGCATA

CCTGACCCCACCGGAGGATCCGAGGAGGGCCCGACCCCGCGTGGACCAAACGGCAGGGAGCCCAACCCCACCAAAAAAAG

AGGCGAGGCCCCCCCTCCCTCCCCCCCCGCACACAACCCGCAGACAGACCGAGGGAGGGCCATCCCCCAGCGAGACAGGC

GGCTCCCTCGCAGCCGAGCCGCAGCTCCCCAGAATAGGACACCCCCGCACACTCCGAACACCCCCTCGCCCCCCGCCACC

GGGACAGGGGCTCCACCACCACAGACCGGAGCCACCGACAGCCAGCCAGAGGTCCCCTCCAGTCCCGCCCGGCGCCCCCC

CGAAGAGGCCCCGGCAAGGCCCCCCAGCCCGACGTCCCCCGCACGCACCGAAAGAAGGACCCGCCCGAGAACGTGCGGCA

CATCCATGCGCAAACATCATGACGCGGGTCGCAATTTTGACATTTCTGTTTCTTTTCCCAAATGTTGTTGCGTGTCAGAT

TCACTGGGGCAATCTATCCAAGATCGGGATTGTAGGAACAGGGAGTGCCAGCTACAAGGTGATGACTAGGCCAAGCCACC

AGACTCTGGTTATAAAGTTAATGCCAAATATAACGGCCATCGACAATTGTACAAAGTCAGAGATTGCAGAGTACAAGAGA

TTGCTGATCACAGTGTTAAAGCCTGTAGAGGATGCTCTGTCGGTGATAACCAAGAATGTAAGACCAATTCAAACTCTAAC

ACCTGGGCGTAGAACCCGCCGTTTTGCTGGAGCTGTTCTGGCCGGGGTAGCACTTGGAGTTGCGACAGCCGCTCAGATAA

CTGCAGGAGTCGCCCTTCATCAATCATTGATGAACTCCCAAGCAATTGAGAGTTTAAAAACCAGTCTTGAGATGTCGAAT

CAGGCAATAGAAGAAATCAGACTTGCAAATAAGGAGACCATACTGGCAGTACAGGGCGTCCAGGATTATATCAACAATGA

GCTCGTCCCTTCTGTTCATAGAATGTCATGCGAGCTGGTAGGTCACAAGCTCGGCCTCAAGCTCCTTAGGTACTACACCG

AGATCCTGTCCATATTCGGGCCCAGTCTTCGAGACCCGATATCTGCCGAAATATCAATCCAGGCACTTAGTTATGCATTA

GGCGGAGACATTAATAAAATCCTGGACAAGCTTGGGTATAGCGGTGGGGATTTCCTTGCCATCCTAGAAAGCAAGGGAAT

AAAGGCCCGGGTTACATATGTGGACACAAGAGATTACTTTATAATCCTTAGCATCGCCTACCCAACCTTATCTGAGATCA

AGGGAGTGATAGTTCACAAGATAGAAGCTATAACATACAACATTGGGGCACAGGAGTGGTATACTACTATCCCTAAATAT

GTAGCCACTCAGGGGTATCTGATATCGAACTTTGATGAGACGTCATGCGTATTCACTCCAGAGGGGACAGTTTGCAGCCA

GAATGCGTTGTACCCAATGAGCCCATTGCTTCAGGAATGTTTCAGGGGGTCAACAAAATCGTGCGCCAGAACCTTAGTTT

CAGGGACCATAAGTAATAGATTTATCCTATCAAAAGGGAACCTGATTGCAAATTGTGCGTCAGTTTTGTGCAAATGTTAC

ACAACAGAGACAGTTATCAGCCAAGATCCTGACAAACTACTAACTGTTGTAGCATCCGACAAGTGTCCTGTAGTTGAGGT

GGATGGAGTGACAATACAGGTCGGCAGTCGAGAGTATCCGGATTCTGTATACTTACACAAAATAGACTTAGGTCCAGCCA

TCTCCCTAGAAAAACTGGATGTAGGCACCAATTTAGGCAATGCAGTCACAAGACTGGAGAATGCAAAGGAGCTCCTAGAT

GCATCAGACCAAATACTGAAGACTGTTAAAGGGGTACCTTTCAGTGGGAATATGTACATAGCACTGGCAGCTTGCATAGG

AGTATCCCTGGGCCTTGTCACATTAATATGCTGCTGTAAGGGGAGGTGTAAGAACAAGGAAATCCCTATCTCCAAAATCA

ACCCAGGGCTCAAACCCGACCTGACCGGGACCTCAAAGTCGTACGTAAGATCACTGTAGTCAGAATCACCTGGATCATCT

GGCATCACACACATACATGCACGACACAGGCAGTCCGAGGACGCAAGAAACCCAGCCTCCGGTCACCCACCCGACCCCAC

TCCACGCTCCACCACACATTAGTCATCAAACAAAACTTAGGACGAAAGGTCAATCACCATGTCCGCACAAAGGGAGAGGA

TCAATGCCTTCTACAAAGACAATCCTCACAATAAGAACCATAGGGTGATCCTGGATAGAGAACGCTTGGTCATTGAAAGA

CCCTACATCTTGCTTGGAGTCCTGCTGGTAATGTTCCTGAGTCTAATCGGACTGCTGGCCATTGCAGGGATCAGGCTTCA

CCGGGCCACCGTTGGAACTTCAGAGATCCAGAGTCGGCTGAATACCAATATTAAGTTGACCGAATCTATTGATCACCAGA

CTAAGGATGTCTTAACTCCCCTTTTTAAAATCATTGGCGATGAAGTCGGCATCAGAATTCCACAGAAATTCAGTGATCTT

GTCAAGTTCATCTCCGATAAGATTAAATTCCTCAACCCTGATAGAGAGTATGATTTCAGAGATCTCCGGTGGTGTATGAA

TCCCCCCGAGAGAGTCAAAATTAATTTTGATCAGTTTTGTGAGTACAAGGCTGCGGTTAAGTCAATTGAACATATATTTG

AGTCACCACTCAACAAGTCAAAAAAGCTGCAATCTTTGACTCTCGGGCCCGGAACAGGCTGTCTAGGCAGGACAGTAACA

AGAGCCCATTTCTCAGAACTTACAATGACCTTAATGGACCTGGATCTAGAGATGAAGCACAACGTGTCCTCAGTGTTTAC

CGTAGTTGAAGAGGGATTATTCGGAAGAACATATACCGTCTGGAGATCCGATGCCAGGGATCCGAGCACCGATCTAGGTA

TCGGCCATTTTTTAAGAGTCTTCGAGATTGGACTGGTAAGAGATCTTGGGCTGGGTCCCCCTGTTTTTCATATGACCAAC

TATCTCACGGTGAACATGAGTGATGACTATCGGAGATGTCTTTTAGCGGTAGGGGAGTTGAAGTTGACAGCCCTATGCAC

CTCATCTGAGACTGTGACACTGAGTGAGAGAGGAGTTCCAAGGAGGGAACCTCTTGTGGTTGTGATACTTAATCTAGCTG

GACCCACTCTAGGGGGAGAGCTATACAGTGTCTTGCCTACCTCTGATCTCATGGTGGAGAAACTCTATTTGTCTTCACAT

AGAGGGATCATCAAAGATGATGAGGCCAATTGGGTAGTGCCGTCTACCGATGTTCGTGATCTTCAGAACAAAGGTGAATG

TCTGGTGGAAGCATGCAAGACTCGACCTCCTTCATTTTGCAATGGCACAGGATCAGGCCCGTGGTCAGAAGGGAGAATCC

CTGCCTACGGGGTGATCAGGGTCAGTCTTGACTTAGCGAGTGACCCGGATGTAGTTATCACTTCAGTGTTTGGCCCACTG

ATACCTCACCTATCCGGCATGGATCTTTACAACAACCCGTTTTCAAAAGCTGTATGGTTGGCTGTACCACCTTATGAGCA

GTCATTTCTAGGAATGATAAATACAATTGGATTCCCTAACAGAGCAGAGGTTATGCCGCACATTTTGACCACAGAGATCA

GAGGCCCTCGGGGTCGTTGCCATGTTCCCATAGAATTGTCCCGCCGGGTTGATGACGATATCAAGATCGGGTCCAACATG

GTCATATTGCCGACGATGGACCTGAGGTATATTACAGCCACTTATGATGTTTCCAGGAGCGAGCATGCAATCGTGTACTA

TATCTATGACACGAGTCGCTCATCATCTTACTTCTACCCAGTTCGACTGAATTTCAAAGGCAATCCTCTCTCTTTGAGGA

TAGAGTGTTTCCCCTGGCGTCATAAGGTGTGGTGCTACCATGATTGTCTTATATACAACACCATAACAGGTGAAGAGGTC

CATACGAGAGGGCTGACCGGCATAGAGGTAACATGCAATCCAGTCTGAGCAGAGCTACGACCGTCACTCAGGCAGTCCCT

TGAGTCGCCACCGAGTCCAAGCAGCACAGCCTGGGACAATCAACGGCACAGCCCAGCCAACAATGTTATAAAAAACTTAG

GAGCCAGGGTTGTAGGGGCCATGGACTCTCTATCAGTCAATCAGGTCTTGTACCCTGAGGTTCATCTAGACAGCCCTATT

GTCACAAACAAACTAGTTGCCATCCTTGAGTACTCGGGTATCGACCATAACTATGTTCTTGAAGACCAGACTCTTGTCAA

GAATATTAGGTATAGGCTGGGGTGCGGTTTTTCAAATCAAATGATCATCAATAACAGGGGGGTGGGTGAAACAGTCAATT

CTAAACTTAAAAGTTACCCCCATAATCGTCATATCATATACCCGGATTGCAATAAGGAGTTGTTTTGTATCAAGGATAGC

TGCATATCTAAGAAGCTCTCGGAGCTATTCAAGAAGGGTAATTCCTTGTACTCTAAGATAAGTCACCAGGTACTGGATTG

TCTTAAGAGAGTCAATGGGAAATTAGGTCTGGGCACAGATCTTACCCATGGTCTGAAGGAGGGTATCCTTGACCTAGGGT

TGCACATGCATAGCTCTCAATGGTTCGAGACTTTCCTGTTCTGGTTCACTATCAAGACAGAGATGAGATCAATGATCAAA

GAACAGTCCCATATATGCCACAAGAGGAGGTATAACCCGACTTTTGTGTCGGGGGATGCATTCGAGGTGCTCGTATCGCG

AGACCTCGTTGTGATAATTGATAAGAATACCCAGTATGTCTTCTACTTAACATTTGAGCTGGTCCTTATGTATTGTGATG

TCATAGAGGGTAGACTTATGACGGAGACAGCCATGGCCATAGACCAGAGATATTCAGAACTTCTAAGCCGGGTCAGATAC

TTGTGGGATCTTATTGATGGGTTCTTCCCAACACTAGGCAACACCACATACCAAATTGTTGCTCTGCTTGAACCATTATC

GTTGGCTTATCTTCAACTTCAGGATGTCACTCTGGAGTTAAGAGGTGCTTTTTTAGACCATTGCTTCAAAGAACTCTATG

AGATACTGGAACATTGCGGCATTGACACAGCAGGCACTTACAATTCCATCACTGAAGGATTGGATTACGTATTCATCACC

CACGATATACATCTAACTGGGGAGATTTTTTCATTTTTCCGGAGTTTCGGACACCCCCGTCTAGAAGCAGTCACCGCTGC

GGAAAATGTCAGAAAACATATGAACCAACCGAAGGTAATCAGTTATGAGACTATGATGAAAGGCCATGCGGTATTTTGCG

GGATAATCATAAATGGTTTTCGGGACCGACATGGCGGCAGCTGGCCCCCTGTTGCACTGCCAGAACATGCCTCTGCTGCG

ATCCGGAATGCGCAGGCATCAGGTGAGGGACTGACCCATGACCTGTGTATAGACAACTGGAAATCCTTTGTTGGATTCAA

ATTTGGCTGCTTTATGCCACTCAGCCTAGATAGTGATTTGACCATGTATCTCAAAGACAAGGCATTGGCTGCACTGAAGA

ATGAGTGGGATTCAGTTTACCCGAAAGAATACCTCCGTTATAACCCACCTAGAGGGACAGAGTCAAGGCGATTGGTAGAG

GTATTCCTGAATGACTCCAGCTTTGATCCTTATAACATGATAATGTACGTGGTGAATGGCTCCTACCTTAAAGACCCTGA

GTTTAACCTCTCATACAGCCTAAAAGAGAAGGAGATAAAGGAGACAGGGCGGTTGTTTGCCAAAATGACCTATAAGATGC

GGGCCTGTCAGGTAATCGCTGAAAATCTGATATCGAATGGTGTTGGGAAGTATTTCCGAGACAATGGGATGGCAAAGGAC

GAGCATGACCTAACAAAAGCCCTTCACACCCTGGCAGTCTCAGGTGTTCCCAAAAACAACAAAGATAACCACCGAGGTGG

GCCTCCCAGAAGGACAACAAACCGAGGGGTGAGATCAAGCCAAGGCACCAAAACACAAGATAGAGACAAGGTTCAAGGGG

GACCTATGTACAACTATTTGCGATGCCAACCGATCAGCCCTGATCAGGGTGAGTCATACGAGACTGTTAGTGCATTCATC

ACCGCTGACCTTAAGAAGTATTGCCTGAATTGGAGATACGAGACAATCAGCATATTTGCACAGAGGCTGAATGAAATATA

TGGACTACCATCCTTCTTTCAATGGTTACACAGGATATTGGAAAAATCCGTACTCTACGTCAGTGACCCACATTGTCCTC

CCGATCTAGATAATCATATCCCTCTGGACAGTGTCCCCAATGCCCAAATATTCATCAAGTACCCAATGGGCGGAATAGAA

GGTTATTGTCAAAAACTATGGACAATCAGCACTATACCATACTTGTATCTGGCAGCCTATGAGAGCGGAGTAAGAATCGC

CTCACTGGTGCAGGGTGACAATCAGACAATCGCAGTGACAAAAAGAGTTCCAAGTTCTTGGCCTTATTCACTAAAAAAGA

GGGAGGCCTCCAAGGCCGCTCAAAATTACTTCGTAGTCTTAAGGCAAAGATTGCACGATGTAGGTCATCACTTAAAGGCT

AATGAGACCATAGTATCTTCTCACTTTTTTGTATATTCCAAAGGGATTTATTATGACGGCCTGCTAGTCTCACAATCACT

AAAGAGCATCGCCAGATGTGTCTTCTGGTCCGAGACTATAGTGGATGAAACCAGAGCGGCGTGCAGTAATATTGCAACAA

CTGTCGCTAAGAGTATAGAGAGGGGTTATGATAGGTACCTTGCATATTCTTTGAATATCCTCAAGATTTTCCAACAGATC

CTTATATCCCTTAACTTCACTATTAACACAACAATGACTCAGGATGTCGTGGCACCGATCATCGAGAACGGTGATTTGCT

GATAAGGATGGCACTCTTGCCAGCACCCATCGGGGGTCTCAATTATCTTAACATGAGCAGGTTATTTGTGAGAAATATCG

GTGACCCGGTCACCTCCTCTATAGCCGACCTGAAGAGGATGATAGATGCCGGGCTAATGCCAGAAGAAACATTGCATCAA

GTGATGACCCAGACCCCGGGAGAATCATCCTACCTAGATTGGGCAAGTGACCCTTATTCTGCCAACCTACCCTGCGTACA

GAGTATCACTCGCCTTCTCAAAAACATCACTGCACGATATATTTTAATCAGCAGCCCAAACCCGATGCTGAAAGGGTTAT

TTCATGAGGGGAGTAGAGATGAAGACGAGGAGCTTGCAAGTTTCCTAATGGATCGGCATATAATTGTTCCAAGAGCTGCA

CATGAAATCTTAGACCATAGTATAACCGGAGCAAGAGAAGCTATAGCCGGGATGTTGGACACCACCAAGGGTCTGATTAG

AACAAGTATGAAACGGGGCGGCCTCACCCCTCGAGTCTTAGCCCGCCTTTCCAATTATGATTATGAACAATTCCGATCCG

GGATAACACTATTAACAAAGAAAGGGCAGTGTTATCTCATTGACAAGGACTCGTGCTCGGTGCAGCTCGCTATAGCTCTG

AGGGGCCACATGTGGGCTAAGTTGGCTCGCGGGAGACCAATCTATGGGTTGGAGGTGCCTGATGTATTAGAATCGATGAA

CGGCTACCTTATCAAACGCCACGAGTCGTGTGCTATCTGTGAAACGGGCTCAAGTCACTACGGATGGTTTTTCGTCCCTG

CAGGGTGCCAGCTTGACGATGTATCTAGAGAGACTTCGGCTCTTCGTGTACCTTATGTCGGATCAACAACTGAGGAAAGG

ACAGACATGAAACTTGCATTTGTTAGGTCTCCAAGTAGATCCCTCAAATCAGCAGTTAGAATTGCAACAGTTTACTCATG

GGCTTACGGGGATGATGAGAAATCATGGAGTGAAGCCTGGATGCTAGCTAGGCAGAGAGCCAATATCACCTTAGATGAAT

TGAGAATGATCACTCCAGTCTCCACATCCACCAACTTAGCCCACCGGTTGAGGGATCGGAGTACCCAGGTGAAATACTCG

GGGACATCCCTTGTGAGGGTTGCAAGATACACAACCATCTCCAATGATAACTTGTCGTTTGTGATATCTGAGAAAAAAGT

GGATACCAACTTCATTTATCAGCAAGGGATGTTGCTTGGTCTTGGGATCCTTGAGAACCTCTTTAGGCTAGAGGCCACCA

CCGGAGTATCCAACACAGTGCTACACCTGCACGTGGAAACAGAATGTTGTGTTGTACCTATGGTGGATCACCCAAGGATA

CCGAGTCTCCGTAATATAAAAGTTACGGATGAGCTATGCACAAACCCTTTGATCTACGACAGGTCCCCCATCATAGAGCA

CGATGCAACCCGGCTATATTCACAGAGCCACAGGAGACATTTGGTGGAGTTTGTTACCTGGTCAACAAGCCAGCTTTACC

ATATACTGGCTAAGTCTACAGCAATGTCCATGATTGAATTGATCACGAGATTCGAGAAAGATCACATGAATGAAATAGCC

GCCCTGATTGGCGATGACGACATCAACAGCTTCATCACAGAATTTTTGCTAGTGGAGCCTAGATTATTTATAGTTTACCT

TGGTCAGTGTGCTGCCATCAATTGGGCCTTTGATATACATTATCATCGGCCCTCGGGCAAGTACCAGATGGGGGAACTTC

TCTACTCTCTGCTCTCTCGGATGAGCAAAGGAGTATATAAGATCTTTACTAATGCTTTGAGCCACCCCAAAGTTTACAAG

AAATTTTGGCGAAGTGGCATCATTGAGCCGATTCATGGCCCATCCTTGGATACACAGAATTTACATGTCACTGTCTGTGA

CATGATATATGGGTCCTATGTCACCTATTTGGATCTTTTGCTGAATGATGAGCTAGATGATTACCCGTATTTGCTCTGCG

AGAGTGATGAGGACGTGGTCACAGACAGGTTCGACAACATTCAAGCCAAACATCTCTGTGTATTGGCCGATGTATATTGC

AGCTCCAAGAGGTGTCCTTCAATAATCGGGATGTCTCCTATAGAAAAATGTACCATTCTCACACATTACATCAAGGGAGA

ATCAGTACAATCCCCGTCTGGGACCTCATGGAACACTGATCCCCTTGTAGTAGATCATTACTCATGCTCTCTGACCTACC

TTCGCCGCGGTTCCATCAAACAAATCAGGTTGAGGGTGGATCCTGGGTTTGTATTCGAGGCGTTGACAGACGTCGACTTT

AAACAACCTCGCAAAGCTAAGTTAGATATATCGGTCGTTGGGTTGACTGATTTCTCTCCCCCTTACGACAACGTCGGTGA

TTTTTTAGGGACTATCAACACATTGAGGCACGATCTGCCTGTCACCGGAACCGGGGTCTCGAACTATGAAGTCCACGCTT

ATCGTAGAATTGGCCTGAATTCATCAGCATGTTATAAAGCCGTAGAGATCTCCACGTTAATCAAGCCATCCTTAGAAGTC

GGAGAGCATGGATTGTTCTTAGGAGAAGGTTCCGGTTCAATGCTGGCTGCGTACAAAGAAGTTCTTAAATTAGCAAATTG

TTACTACAACAGCGGAGTAACAGCGGAGGGCAGAGCCGGACAGAGGGAAATATCTCCCTATCCTTCAGAGATGAGCCTAG

TAGAGAATCAAATGGGGATAGAGCGGAGTGTTAAAGTGCTGTTCAACGGGAAACCTGAAGTAACTTGGGTGGGGACCACA

GATTGCTACAAGTACATAATCAGTAACATCCAAACCTCTAGTCTGGGTTTCATACATTCAGATATTGAGACGCTCCCAAC

CAAAGATGCTGTTGAGAAATTGGAGGAATTTGCCTCTATCCTATCCTTATCCTTGATTTTGGGGAAAATCGGCTCTATTA

CAGTTGTCAAAATTATGCCCATTAGCGGGGATTTTACCCAAGGCTTCATAGCCTATGCCATCCAATATTTTAGGGAGAGC

CTGCTTGCCTAYCCGAGATATAGTAACTTCATCTCGACTGAGTGTTACCTTATTATGATAGGATTAAAGGCCAATCGATT

GATAAACCCAGAAGCCATTAAGCAAAGCATAATCAGAGCGGGGGTCAGGACTGCACCAGGACTTGTGAGCCATGTATTAT

CAGAGAAACAAAAAGGTTGTATTCAATCTTTCTTGGGTGATCCTTATATCCAAGGAGACTTCAATAAGCACCTTAAATCT

CTAACCTCTATTGAGAAAATCCTGGTAAATTGTGGTCTCTCGATCAATGGCATAAAAATCTGTAGAGATCTAATCCACCA

TGATATCGCCTCCGGTCCAGACGGCCTTATGAGCTCTACAATTATTTTATATAGGGAATTGGCTCATTTCAAAGATAATA

TAAGGAGTCAGCACGGAATGTTCCACCCCTACCCGGTACTGGCCAACAGCAGGCAACGTGAATTAATCCTTCGAATAGCC

AAGAAATTCTGGGGGTATGTCTTGCTATATTCTGATGACCCGGCACTAATCAGACAAACAATCAAGAACTTGAAACGGAA

TCATCTAACCTTTGACTTACATAGTAACCCATTTATTAAAGGGCTATCCAAGGCTGAGAAACTGCTGGTGCGGACGAGTT

CACTAAGAAGAGAATGGTTGTTCACTCTCGAAACGAAAGAAGTAAAAGAGTGGTTCAAATTGGTAGGTTACAGTGCACTC

ATCAGAGGCTAGTCAGCTATGCATCTGCCCCCTTCTCCTCCGCCATGAGACCCCACTGGCGATCCAGAAGATTAAAGAAA

ACTACATATTGGATAAGTATCTATTCCCAGCTTTGTCTGGT

>KX421384.1/goat/China/2013-12-20

ACCAAACAAAGTTGGGTAAGGATAGATCTTATAATAACTATGGACTGGCAAACTTAGGAGTAAAGATCCTACTGTCGGGG

GGAGGAGGAGGAGCAAGATCCTTGATCATGGCGACTCTCCTTAAAAGCTTAGCATTGTTCAAAAGGAACAAAGACAAAGC

GCCGACAGCATCAGGTTCAGGAGGGGCCATCCGGGGGATTAAGAATGTTATCATAGTCCCGATTCCCGGAGACTCGTCCA

TCACTACCCGTTCAAGACTGCTCGACAGGCTTGTCAGATTGGCCGGAGATCCTGACATCAACGGGTCAAAGCTGACCGGG

GTGATGATCAGCATGTTATCCTTGTTCGTAGAGTCACCCGGGCAATTGATACAGCGAATCACAGATGATCCGGATGTTAG

TATCCGCCTTGTTGAGGTAGTTCAAAGTACTAGGTCCCAGTCCGGGTTGACCTTTGCATCACGTGGTGCTGATTTAGACA

ACGAGGCAGACATGTATTTTTCAACTGAGGGGCCCTCGAGTGGAGGTAAGAAAAGGATCAACTGGTTTGAGAACAGAGAA

ATAATAGACATAGAGGTGCAGGATCCAGAAGAGTTCAATATGTTGTTAGCCTCCATACTAGCACAAGTCTGGATCCTTCT

GGCCAAGGCTGTTACGGCACCAGATACGGCAGCTGACTCAGAACTGAGAAGGTGGGTTAAATACACACAACAAAGGAGAG

TGATTGGGGAATTTCGCCTTGACAAGGGGTGGCTGGACGCAGTCCGCAACAGAATTGCAGAAGATCTATCACTCCGGCGG

TTCATGGTATCTCTCATACTTGACATCAAGAGGACCCCAGGCAACAAGCCAAGGATTGCAGAAATGATCTGCGACATTGA

CAACTATATTGTCGAAGCAGGACTCGCCAGTTTCATCCTTACTATCAAATTTGGTATCGAAACCATGTATCCTGCATTAG

GGCTCCACGAGTTTGCCGGAGAATTGTCCACTATTGAGTCCTTGATGAACTTGTATCAACAGCTAGGAGAGGTTGCACCC

TACATGGTAATTCTAGAGAATTCAATTCAGAACAAGTTTAGTGCAGGAGCTTATCCCCTCCTCTGGAGCTATGCGATGGG

TGTCGGAGTCGAGCTGGAGAACTCAATGGGGGGCTTGAACTTTGGCAGGTCATATTTTGACCCTGCCTATTTTCGTCTCG

GACAGGAGATGGTCAGAAGATCTGCAGGAAAGGTCAGCTCTGTGATTGCGGCTGAGCTCGGCATCACAGCAGAGGAAGCT

AAACTAGTCTCGGAAATCGCCTCGCAGGCTGGGGACGAAAGAACCGCCAGAGGGACTGGTCCTCGACAGGCGCAGGTTTC

CTTCCTTCAGCACCAAACAGGAGGGGGAGAGTCGTCCGCACCAGCGACCAGAGAAGGGGTCAAAGCTGCGATCCCAAACG

GAGCTGAAGAAAGGGACAGAAAGCAAACACGCCCAGGAAGGCCCAGAGGAGAGACCTCCGGCCAACTGCTCCTGGACATC

ATGCCAGAGGATGAGATCTTGCGAGAGTCTGGTCAAAACCCTCGTGAGGCTCAAAGATCGGCCGAGGCACTCTTCAGGCT

GCAGGCCATGGCCAAGATTCTGGAGGACCAGGAGGAGGGAGAAGACAACAGTCAGGTCTACAACGACAAGGATCTCCTCG

GCTAAGCAGACGCACCCTCTGTCGAAATCAGTGACAAGACATCTCCTACCAGTATTATAAAAAACTTAGGACCCAGGTCC

AAGCAAGCACACATCGACACTCCAACCAGTCGAGCGGAGACCACCGATGGCAGAAGAACAAGCATACCATGTCAACAAGG

GGCTGGAATGTATCAAGTCTCTCAAAGCCTCACCCCCGGATCTATCCACCATCAGAGATACCATCGAGAGCTGGAGAGAG

GGGCTTAGCCCCTCGGGCCGTGCAACACCGAACCCTGATACGTCCGAGGGAGATCATCAGAATATCAACCAATCATGCTC

ACCAGCAATCGGACCAAACAAAGTCTACTTGTCTCCTGAAGATAATCTCGGATTTAGAGAGATCACTGGCAACGACTGTG

AGGCTGGGCTCGGAGGGGTCCAGAGAGAAGGATCCAACTCTCAAGTACAGCGTTACCATGTTTATAGTCACGGGGGTGAA

GAGATTGAAGGACTCGAGGATGCTGACTCTCTCGTGGTTCAAGCAGATCCTCCAGTTGCTAACGTCTTCAATGGAGGAGA

GGATGGATCTGACGACAGCGATGTGGACTCTGGCCCAGATGATCCCGGCAGAGATACTCTATATGACCGGGGATCTGTTG

CCGGCAATGGTATCGCTAGGTCCACAGATGTCGAAAAACTAGAAGGTGCTGATATTCAAGAAGTTCTTAACTCCCAGAAA

GGCAAAGGAGGAAGATTCCAGGGCGGGAAAACCTTGCGAGTCCCGGAAATACCCGATGTCAAGCACTCCAGACCATCAGC

CCAATCAATTAAAAAGGGCACAGACGGGAACTCAGTCTCATCTGGAACGGTGATAGAGTGTTTATCGATAAGTGGTGCAA

CCCAAACTGTGCCAGAGTCAAGATGGGAGTCATCAGAGCAAAATGCGTCTGTGGGGAGTGTCCTCAAGTCTGCGAGGAGT

GCAAAGACGATCCAGGGGTCGACACAAGAATCTGGTACCATAGCATCACTGACTCAGCCTAAAGAGAATGACTCCGAGTA

TGAGTATGAGGATGACCTATTTACAGAGATTCAGGACATCCGTGCAAGCATTGCCAAGATCCATGATGACAATAAAACTA

TCCTCTCAAAGCTTGATTCTATACTGTTATTGAAAGGGGAAGTCGACACTATCAAGAAACAAATCAGCAAGCAGAATATA

AGTATATCCACCATTGAGGGCCATCTCTCCAGTATAATGATAGCCATCCCTGGCTTTGGGAAGGACATCAAGGACCCAAC

ATCCGAAGTCGAGTTGAACCCAGATCTAAGACCTATAATAAGCCGTGACTCTGGCAGAGCTCTCGCGGAGGTCCTCAAGA

AACCCGCTGTGGATAGGTCTCAGAAAATTGGAACCAAAGCCAACTCCAGCTCAAAGGGTCAGCTTCTTAAGGATCTCCAG

CTAAAACCTGTCGACAAGCAGGCAAGCTCTGCAATCGGGTTTGTCCCGTCCGACCATGAATCATCCAGAAATGTCATCCG

CTCCATAATCAAGTCGAGCAAGCTAAATATTGATCACAAGGACTATCTTCTAGATTTACTAAATGATGTGAAAGGCTCCA

AGGATCTTAAGGAATTTCACAAGATGCTAACAGCAATTCTCGCCAAGCACCCGTAACACATCCTCCAGTCATCATCTCAT

ACTCGACTAAAAACATCCTTTCAATCAGGCTATTACAAAAAACTTAGGAGCAAGGGCAACTGAGCTTCGCAGACAGGATG

ACCGAGATCTACGACTTCGATAAATCAGCATGGGATGTCAAAGGGTCAATTGCCCGCATAGAACCCACCACCTACCACGA

CGGCCGACTGATACCCCAGGTGAGGGTCATCGATCCTGGTCTGGGAGACAGAAAAGATGAGTGCTTTATGTACCTGTTTC

TCCTAGGAGTGATTGAGGATAACGACCCCCTGTCTCCCCCCGTCGGAAGAACCTTTGGCTCTCTACCTCTAGGGGTTGGT

AGGTCAACTGCCAAGCCAGAAGAACTACTAAGGGAGGCCACAGAATTAGATATAGTGGTGAGGCGCACTGCAGGAGTAAA

TGAGAAACTGGTATTTTACAACAACACTCCGCTGTCCTTGTTAACGCCCTGGAAGAAAGTTCTGACAACCGGAAGTGTGT

TTAGCGCTAACCAGGTTTGCAATGCAGTCAACCTAGTCCCGCTTGATACTCCCCAGAGATTCAGGGTTGTGTACATGAGC

ATAACTAGATTATCAGACAATGGTTACTATAGTGTGCCCAGAAGAATGCTGGAGTTCCGCTCGGCCAATGCAGTAGCCTT

CAACATCTTGGTTACACTAAGAATTGAAAATGGCACAAACCCTAGAAGATACATAGTCGGCTCATGGGAGAATTCAGAGG

TCACATTTATGGTACACGTGGGCAACTTTAGGAGAAAGAAGAACGAAGTATACTCTGCTGATTACTGCAAAATGAAGATT

GAAAAGATGGGTTTAGTTTTTGCCCTGGGTGGAATAGGCGGAACAAGTCTCCATATTAGAAGCACCGGGAAAATGAGCAA

AACCCTCCATGCACAGCTGGGGTTCAAGAAAATCTTATGTTACCCCCTAATGGATGTCAATGAGGATCTTAACCGATATC

TCTGGCGGGCAGAGTGCCGAATAGTCAAAATCCAAGCTGTCTTACAGCCATCAGTACCCCAAGAGTTCCGTGTCTACGAT

GATGTCATCATCAACGATGATCAAGGCTTGTTCAAGATCTTGTAGTTCATTTGCAAAGGAACGAACCCACCCCCAGGAAG

GGGCCCCACCCTCCCCCG-CCAAAAAACACACTCCAAGGCTCCCCGGCACAGCGCCAACACAGCTACTCCCGCGATGGAA

CGAGGTCGGCC-CCCGAACAAATCCGGTCCAGGGGGGGGGGCCCCCCCCCTGACCAACCGGGCGCCGACCGAGCAGGCCG

GCGAGAAGCCCCCAAGAGGCCCCGACGAAGACCGAGCACCCCATCCCCCC-AAAAACGTCCCCC-AACACGCCCGAGGCT

AGCCACCGGACACCCCCAGACCCGCCACCGGCGAGACAGAACCCCAAGGGGGGACCCTCCAAACCAAAGGAATCACCGGG

GTAGGCTCCGCCGAACCAGCACAGACCCGACCCAAGCAAGGCGGACCCCCAACACGAAACTGCCCAGCACACCCTACTCA

TCATCAAACAAAACTTAGGGGCCAAGTCCATACGAGAGGCCCTAGTCCGTCAGGCGGAGGGACGCAGAAAGGAAGGAGAC

ACGACCGTCCCAGAAGGAGGACACCCAACCACCGAAACGCCCGCCCCCAGGGAAGGAG--CCA-CCCCCACGCAGCGAAG

GAGCGGGTGGGAGACCTCGGGGAGGGCCCGACCGGGACCCGGGGGCCGGCGGCGCCGGCGGTGAGCAGGCCGGGCGCATA

CCTGACCCCACCGGAGGATCCGAGGAGGGCCCGACCCCGCGTGGACCAAACGGCAGGGAGCCCAACCCCACCAAAAAGAG

AGGCGAGGCCCCCCCTCCCTCCCCCCCCGCACACAACCCGCAGACAGACCGAGGGAGGGCCATCCCCCAGCGAGACAGGC

GGCTCCCTCGCAGCCGAGCCGCAGCTCCCCAGAATAGGACACCCCCGCACACTCCGAACACCCCCTCGCCCCCCGCCACC

GGGACAGGGGCTCCACCACCACAGACCGGAGCCACCGACAGCCAGCCAGAGGTCCCCTCCAGTCCCGCCCGGCGCCCCCC

CGAAGAGGCCCCGGCAAGGCCCCCCAGCCCGACGTCCCCCGCACGCACCGAAAGAAGGACCCGCCCGAGAACGTGCGGCA

CATCCATGCGCAAACATCATGACGCGGGTCGCAATTTTGACATTTCTGTTTCTTTTCCCAAATGTTGTTGCGTGTCAGAT

TCACTGGGGCAATCTATCCAAGATCGGGATTGTAGGAACAGGGAGTGCCAGCTACAAGGTGATGACTAGGCCAAGCCACC

AGACTCTGGTTATAAAGTTAATGCCAAATATAACGGCCATCGACAATTGTACAAAGTCAGAGATTGCAGAGTACAAGAGA

TTGCTGATCACAGTGTTAAAGCCTGTAGAGGATGCTCTGTCGGTGATAACCAAGAATGTAAGACCAATTCAAACTCTAAC

ACCTGGGCGTAGAACCCGCCGTTTTGCTGGAGCTGTTCTGGCCGGGGTAGCACTTGGAGTTGCGACAGCCGCTCAGATAA

CTGCAGGAGTCGCCCTTCATCAATCATTGATGAACTCCCAAGCAATTGAGAGTTTAAAAACCAGTCTTGAGATGTCGAAT

CAGGCAATAGAAGAAATCAGACTTGCAAATAAGGAGACCATACTGGCAGTACAGGGCGTCCAGGATTATATCAACAATGA

GCTCGTCCCTTCTGTTCATAGAATGTCATGCGAGCTGGTAGGTCACAAGCTCGGCCTCAAGCTCCTTAGGTACTACACCG

AGATCCTGTCCATATTCGGGCCCAGTCTTCGAGACCCGATATCTGCCGAAATATCAATCCAGGCACTTAGTTATGCATTA

GGCGGAGACATTAATAAAATCCTGGACAAGCTTGGGTATAGCGGTGGGGATTTCCTTGCCATCCTAGAAAGCAAGGGAAT

AAAGGCCCGGGTTACATATGTGGACACAAGAGATTACTTTATAATCCTTAGCATCGCCTACCCAACCTTATCTGAGATCA

AGGGAGTGATAGTTCACAAGATAGAAGCTATAACATACAACATTGGGGCACAGGAGTGGTATACTACTATCCCTAAATAT

GTAGCCACTCAGGGGTATCTGATATCGAACTTTGATGAGACGTCATGCGTATTCACTCCAGAGGGGACAGTTTGCAGCCA

GAATGCGTTGTACCCAATGAGCCCATTGCTTCAGGAATGTTTCAGGGGGTCAACAAAATCGTGCGCCAGAACCTTAGTTT

CAGGGACCATAAGTAATAGATTTATCCTATCAAAAGGGAACCTGATTGCAAATTGTGCGTCAGTTTTGTGCAAATGTTAC

ACAACAGAGACAGTTATCAGCCAAGATCCTGACAAACTACTAACTGTTGTAGCATCCGACAAGTGTCCTGTAGTTGAGGT

GGATGGAGTGACAATACAGGTCGGCAGTCGAGAGTATCCGGATTCTGTATACTTACACAAAATAGACTTAGGTCCAGCCA

TCTCCCTAGAAAAACTGGATGTAGGCACCAATTTAGGCAATGCAGTCACAAGACTGGAGAATGCAAAGGAGCTCCTAGAT

GCATCAGACCAAATACTGAAGACTGTTAAAGGGGTACCTTTCAGTGGGAATATGTACATAGCACTGGCAGCTTGCATAGG

AGTATCCCTGGGCCTTGTCACATTAATATGCTGCTGTAAGGGGAGGTGTAAGAACAAGGAAATCCCTATCTCCAAAATCA

ACCCAGGGCTCAAACCCGACCTGACCGGGACCTCAAAGTCGTACGTAAGATCACTGTAGTCAGAATCACCTGGATCATCT

GGCATCACACACATACATGCACGACACAGGCAGTCCGAGGACGCAAGAAACCCAGCCTCCGGTCACCCACCCGACCCCAC

TCCACGCTCCACCACACATTAGTCATCAAACAAAACTTAGGACGAAAGGTCAATCACCATGTCCGCACAAAGGGAGAGGA

TCAATGCCTTCTACAAAGACAATCCTCACAATAAGAACCATAGGGTGATCCTGGATAGAGAACGCTTGGTCATTGAAAGA

CCCTACATCTTGCTTGGAGTCCTGCTGGTAATGTTCCTGAGTCTAATCGGACTGCTGGCCATTGCAGGGATCAGGCTTCA

CCGGGCCACCGTTGGAACTTCAGAGATCCAGAGTCGGCTGAATACCAATATTAAGTTGACCGAATCTATTGATCACCAGA

CTAAGGATGTCTTAACTCCCCTTTTTAAAATCATTGGCGATGAAGTCGGCATCAGAATTCCACAGAAATTCAGTGATCTT

GTCAAGTTCATCTCCGATAAGATTAAATTCCTCAACCCTGATAGAGAGTATGATTTCAGAGATCTCCGGTGGTGTATGAA

TCCCCCCGAGAGAGTCAAAATTAATTTTGATCAGTTTTGTGAGTACAAGGCTGCGGTTAAGTCAATTGAACATATATTTG

AGTCACCACTCAACAAGTCAAAAAAGCTGCAATCTTTGACTCTCGGGCCCGGAACAGGCTGTCTAGGCAGGACAGTAACA

AGAGCCCATTTCTCAGAACTTACAATGACCTTAATGGACCTGGATCTAGAGATGAAGCACAACGTGTCCTCAGTGTTTAC

CGTAGTTGAAGAGGGATTATTCGGAAGAACATATACCGTCTGGAGATCCGATGCCAGGGATCCGAGCACCGATCTAGGTA

TCGGCCATTTTTTAAGAGTCTTCGAGATTGGACTGGTAAGAGATCTTGGGCTGGGTCCCCCTGTTTTTCATATGACCAAC

TATCTCACGGTGAACATGAGTGATGACTATCGGAGATGTCTTTTAGCGGTAGGGGAGTTGAAGTTGACAGCCCTATGCAC

CTCATCTGAGACTGTGACACTGAGTGAGAGAGGAGTTCCAAGGAGGGAACCTCTTGTGGTTGTGATACTTAATCTAGCTG

GACCCACTCTAGGGGGAGAGCTATACAGTGTCTTGCCTACCTCTGATCTCATGGTGGAGAAACTCTATTTGTCTTCACAT

AGAGGGATCATCAAAGATGATGAGGCCAATTGGGTAGTGCCGTCTACCGATGTTCGTGATCTTCAGAACAAAGGTGAATG

TCTGGTGGAAGCATGCAAGACTCGACCTCCTTCATTTTGCAATGGCACAGGATCAGGCCCGTGGTCAGAAGGGAGAATCC

CTGCCTACGGGGTGATCAGGGTCAGTCTTGACTTAGCGAGTGACCCGGATGTAGTTATCACTTCAGTGTTTGGCCCACTG

ATACCTCACCTATCCGGCATGGATCTTTACAACAACCCGTTTTCAAAAGCTGTATGGTTGGCTGTACCACCTTATGAGCA

GTCATTTCTAGGAATGATAAATACAATTGGATTCCCTAACAGAGCAGAGGTTATGCCGCACATTTTGACCACAGAGATCA

GAGGCCCTCGGGGTCGTTGCCATGTTCCCATAGAATTGTCCCGCCGGGTTGATGACGATATCAAGATCGGGTCCAACATG

GTCATATTGCCGACGATGGACCTGAGGTATATTACAGCCACTTATGATGTTTCCAGGAGCGAGCATGCAATCGTGTACTA

TATCTATGACACGAGTCGCTCATCATCTTACTTCTACCCAGTTCGACTGAATTTCAAAGGCAATCCTCTCTCTTTGAGGA

TAGAGTGTTTCCCCTGGCGTCATAAGGTGTGGTGCTACCATGATTGTCTTATATACAACACCATAACAGGTGAAGAGGTC

CATACGAGAGGGCTGACCGGCATAGAGGTAACATGCAATCCAGTCTGAGCAGAGCTACGACCGTCACTCAGGCAGTCCCT

TGAGTCGCCACCGAGTCCAAGCAGCACAGCCTGGGACAATCAACGGCACAGCCCAGCCAACAATGTTATAAAAAACTTAG

GAGCCAGGGTTGTAGGGGCCATGGACTCTCTATCAGTCAATCAGGTCTTGTACCCTGAGGTTCATCTAGACAGCCCTATT

GTCACAAACAAACTAGTTGCCATCCTTGAGTACTCGGGTATCGACCATAACTATGTTCTTGAAGACCAGACTCTTGTCAA

GAATATTAGGTATAGGCTGGGGTGCGGTTTTTCAAATCAAATGATCATCAATAACAGGGGGGTGGGTGAAACAGTCAATT

CTAAACTTAAAAGTTACCCCCATAATCGTCATATCATATACCCGGATTGCAATAAGGAGTTGTTTTGTATCAAGGATAGC

TGCATATCTAAGAAGCTCTCGGAGCTATTCAAGAAGGGTAATTCCTTGTACTCTAAGATAAGTCACCAGGTACTGGATTG

TCTTAAGAGAGTCAATGGGAAATTAGGTCTGGGCACAGATCTTACCCATGGTCTGAAGGAGGGTATCCTTGACCTAGGGT

TGCACATGCATAGCTCTCAATGGTTCGAGACTTTCCTGTTCTGGTTCACTATCAAGACAGAGATGAGATCAATGATCAAA

GAACAGTCCCATATATGCCACAAGAGGAGGTATAACCCGACTTTTGTGTCGGGGGATGCATTCGAGGTGCTCGTATCGCG

AGACCTCGTTGTGATAATTGATAAGAATACCCAGTATGTCTTCTACTTAACATTTGAGCTGGTCCTTATGTATTGTGATG

TCATAGAGGGTAGACTTATGACGGAGACAGCCATGGCCATAGACCAGAGATATTCAGAACTTCTAAGCCGGGTCAGATAC

TTGTGGGATCTTATTGATGGGTTCTTCCCAACACTAGGCAACACCACATACCAAATTGTTGCTCTGCTTGAACCATTATC

GTTGGCTTATCTTCAACTTCAGGATGTCACTCTGGAGTTAAGAGGTGCTTTTTTAGACCATTGCTTCAAAGAACTCTATG

AGATACTGGAACATTGCGGCATTGACACAGCAGGCACTTACAATTCCATCACTGAAGGATTGGATTACGTATTCATCACC

CACGATATACATCTAACTGGGGAGATTTTTTCATTTTTCCGGAGTTTCGGACACCCCCGTCTAGAAGCAGTCACCGCTGC

GGAAAATGTCAGAAAACATATGAACCAACCGAAGGTAATCAGTTATGAGACTATGATGAAAGGCCATGCGGTATTTTGCG

GGATAATCATAAATGGTTTTCGGGACCGACATGGCGGCAGCTGGCCCCCTGTTGCACTGCCAGAACATGCCTCTGCTGCG

ATCCGGAATGCGCAGGCATCAGGTGAGGGACTGACCCATGACCTGTGTATAGACAACTGGAAATCCTTTGTTGGATTCAA

ATTTGGCTGCTTTATGCCACTCAGCCTAGATAGTGATTTGACCATGTATCTCAAAGACAAGGCATTGGCTGCACTGAAGA

ATGAGTGGGATTCAGTTTACCCGAAAGAATACCTCCGTTATAACCCACCTAGAGGGACAGAGTCAAGGCGATTGGTAGAG

GTATTCCTGAATGACTCCAGCTTTGATCCTTATAACATGATAATGTACGTGGTGAATGGCTCCTACCTTAAAGACCCTGA

GTTTAACCTCTCATACAGCCTAAAAGAGAAGGAGATAAAGGAGACAGGGCGGTTGTTTGCCAAAATGACCTATAAGATGC

GGGCCTGTCAGGTAATCGCTGAAAATCTGATATCGAATGGTGTTGGGAAGTATTTCCGAGACAATGGGATGGCAAAGGAC

GAGCATGACCTAACAAAAGCCCTTCACACCCTGGCAGTCTCAGGTGTTCCCAAAAACAACAAAGATAACCACCGAGGTGG

GCCTCCCAGAAGGACAACAAACCGAGGGGTGAGATCAAGCCAAGGCACCAAAACACAAGATAGAGACAAGGTTCAAGGGG

GACCTATGTACAACTATTTGCGATGCCAACCGATCAGCCCTGATCAGGGTGAGTCATACGAGACTGTTAGTGCATTCATC

ACCGCTGACCTTAAGAAGTATTGCCTGAATTGGAGATACGAGACAATCAGCATATTTGCACAGAGGCTGAATGAAATATA

TGGACTACCATCCTTCTTTCAATGGTTACACAGGATATTGGAAAAATCCGTACTCTACGTCAGTGACCCACATTGTCCTC

CCGATCTAGATAATCATATCCCTCTGGACAGTGTCCCCAATGCCCAAATATTCATCAAGTACCCAATGGGCGGAATAGAA

GGTTATTGCCAAAAACTATGGACAATCAGCACTATACCATACTTGTATCTGGCAGCCTATGAGAGCGGAGTAAGAATCGC

CTCACTGGTGCAGGGTGACAATCAGACAATCGCAGTGACAAAAAGAGTTCCAAGTTCTTGGCCTTATTCACTAAAAAAGA

GGGAGGCCTCCAAGGCCGCTCAAAATTACTTCGTAGTCTTAAGGCAAAGATTGCACGATGTAGGTCATCACTTAAAGGCT

AATGAGACCATAGTATCTTCTCACTTTTTTGTATATTCCAAAGGGATTTATTATGACGGCCTGCTAGTCTCACAATCACT

AAAGAGCATCGCCAGATGTGTCTTCTGGTCCGAGACTATAGTGGATGAAACCAGAGCGGCGTGCAGTAATATTGCAACAA

CTGTCGCTAAGAGTATAGAGAGGGGTTATGATAGGTACCTTGCATATTCTTTGAATATCCTCAAGATTTTCCAACAGATC

CTTATATCCCTTAACTTCACTATTAACACAACAATGACTCAGGATGTCGTGGCACCGATCATCGAGAACGGTGATTTGCT

GATAAGGATGGCACTCTTGCCAGCACCCATCGGGGGTCTCAATTATCTTAACATGAGCAGGTTATTTGTGAGAAATATCG

GTGACCCGGTCACCTCCTCTATAGCCGACCTGAAGAGGATGATAGATGCCGGGCTAATGCCAGAAGAAACATTGCATCAA

GTGATGACCCAGACYCCGGGAGAATCATCCTACCTAGATTGGGCAAGTGACCCTTATTCTGCCAACCTACCCTGCGTACA

GAGTATCACTCGCCTTCTCAAAAACATCACTGCACGATATATTTTAATCAGCAGCCCAAACCCGATGCTGAAAGGGTTAT

TTCATGAGGGGAGTAGAGATGAAGACGAGGAGCTTGCAAGTTTCCTAATGGATCGGCATATAATTGTTCCAAGAGCTGCA

CATGAAATCTTAGACCATAGTATAACCGGAGCAAGAGAAGCTATAGCCGGGATGTTGGACACCACCAAGGGTCTGATTAG

AACAAGTATGAAACGGGGCGGCCTCACCCCTCGAGTCTTAGCCCGCCTTTCCAATTATGATTATGAACAATTCCGATCCG

GGATAACACTATTAACAAAGAAAGGGCAGTGTTATCTCATTGACAAGGACTCGTGCTCGGTGCAGCTCGCTATAGCTCTG

AGGGGCCACATGTGGGCTAAGTTGGCTCGCGGGAGACCAATCTATGGGTTGGAGGTGCCTGATGTATTAGAATCGATGAA

CGGCTACCTTATCAAACGCCACGAGTCGTGTGCTATCTGTGAAACGGGCTCAAGTCACTACGGATGGTTTTTCGTCCCTG

CAGGGTGCCAGCTTGACGATGTATCTAGAGAGACTTCGGCTCTTCGTGTACCTTATGTCGGATCAACAACTGAGGAAAGG

ACAGACATGAAACTTGCATTTGTTAGGTCTCCAAGTAGATCCCTCAAATCAGCAGTTAGAATTGCAACAGTTTACTCATG

GGCTTACGGGGATGATGAGAAATCATGGAGTGAAGCCTGGATGCTAGCTAGGCAGAGAGCCAATATCACCTTAGATGAAT

TGAGAATGATCACTCCAGTCTCCACATCCACCAACTTAGCCCACCGGTTGAGGGATCGGAGTACCCAGGTGAAATACTCG

GGGACATCCCTTGTGAGGGTTGCAAGATACACAACCATCTCCAATGATAACTTGTCGTTTGTGATATCTGAGAAAAAAGT

GGATACCAACTTCATTTATCAGCAAGGGATGTTGCTTGGTCTTGGGATCCTTGAGAACCTCTTTAGGCTAGAGGCCACCA

CCGGAGTATCCAACACAGTGCTACACCTGCACGTGGAAACAGAATGTTGTGTTGTACCTATGGTGGATCACCCAAGGATA

CCGAGTCTCCGTAATATAAAAGTTACGGATGAGCTATGCACAAACCCTTTGATCTACGACAGGTCCCCCATCATAGAGCA

CGATGCAACCCGGCWATATTCACAGAGCCACAGGAGACATTTGGTGGAGTTTGTTACCTGGTCAACAAGCCAGCTTTACC

ATATACTGGCTAAGTCTACAGCAATGTCCATGATTGAATTGATCACGAGATTCGAGAAAGATCACATGAATGAAATAGCC

GCCCTGATTGGCGATGACGACATCAACAGCTTCATCACAGAATTTTTGCTAGTGGAGCCTAGATTATTTATAGTTTACCT

TGGTCAGTGTGCTGCCATCAATTGGGCCTTTGATATACATTATCATCGGCCCTCGGGCAAGTACCAGATGGGGGAACTTC

TCTACTCTCTGCTCTCTCGGATGAGCAAAGGAGTATATAAGATCTTTACTAATGCTTTGAGCCACCCCAAAGTTTACAAG

AAATTTTGGCGAAGTGGCATCATTGAGCCGATTCATGGCCCATCCTTGGATACACAGAATTTACATGTCACTGTCTGTGA

CATGATATATGGGTCCTATGTCACCTATTTGGATCTTTTGCTGAATGATGAGCTAGATGATTACCCGTATTTGCTCTGCG

AGAGTGATGAGGACGTGGTCACAGACAGGTTCGACAACATTCAAGCCAAACATCTCTGTGTATTGGCCGATGTATATTGC

AGCTCCAAGAGGTGTCCTTCAATAATCGGGATGTCTCCTATAGAAAAATGTACCATTCTCACACATTACATCAAGGGAGA

ATCAGTACAATCCCCGTCTGGGACCTCATGGAACACTGATCCCCTTGTAGTAGATCATTACTCATGCTCTCTGACCTACC

TTCGCCGCGGTTCCATCAAACAAATCAGGTTGAGGGTGGATCCTGGGTTTGTATTCGAGGCGTTGACAGACGTCGACTTT

AAACAACCTCGCAAAGCTAAGTTAGATATATCGGTCGTTGGGTTGACTGATTTCTCTCCCCCTTACGACAACGTCGGTGA

TTTTTTAGGGACTATCAACACATTGAGGCACGATCTGCCTGTCACCGGAACCGGGGTCTCGAACTATGAAGTCCACGCTT

ATCGTAGAATTGGCCTGAATTCATCAGCATGTTATAAAGCCGTAGAGATCTCCACGTTAATCAAGCCATCCTTAGAAGTC

GGAGAGCATGGATTGTTCTTAGGAGAAGGTTCCGGTTCAATGCTGGCTGCGTACAAAGAAGTTCTTAAATTAGCAAATTG

TTACTACAACAGCGGAGTAACAGCGGAGGGCAGAGCCGGACAGAGGGAAATATCTCCCTATCCTTCAGAGATGAGCCTAG

TAGAGAATCAAATGGGGATAGAGCGGAGTGTTAAAGTGCTGTTCAACGGGAAACCTGAAGTAACTTGGGTGGGGACCACA

GATTGCTACAAGTACATAATCAGTAACATCCAAACCTCTAGTCTGGGTTTCATACATTCAGATATTGAGACGCTCCCAAC

CAAAGATGCTGTTGAGAAATTGGAGGAATTTGCCTCTATCCTATCCTTATCCTTGATTTTGGGGAAAATCGGCTCTATTA

CAGTTGTCAAAATTATGCCCATTAGCGGGGATTTTACCCAAGGCTTCATAGCCTATGCCATCCAATATTTTAGGGAGAGC

CTGCTTGCCTATCCGAGATATAGTAACTTCATCTCGACTGAGTGTTACCTTATTATGATAGGATTAAAGGCCAATCGATT

GATAAACCCAGAAGCCATTAAGCAAAGCATAATCAGAGCGGGGGTCAGGACTGCACCAGGACTTGTGAGCCATGTATTAT

CAGAGAAACAAAAAGGTTGTATTCAATCTTTCTTGGGTGATCCTTATATCCAAGGAGACTTCAATAAGCACCTTAAATCT

CTAACCTCTATTGAGAAAATCCTGGTAAATTGTGGTCTCTCGATCAATGGCATAAAAATCTGTAGAGATCTAATCCACCA

TGATATCGCCTCCGGTCCAGACGGCCTTATGAGCTCTACAATTATTTTATATAGGGAATTGGCTCATTTCAAAGATAATA

TAAGGAGTCAGCACGGAATGTTCCACCCCTACCCGGTACTGGCCAACAGCAGGCAACGTGAATTAATCCTTCGAATAGCC

AAGAAATTCTGGGGGTATGTCTTGCTATATTCTGATGACCCGGCACTAATCAGACAAACAATCAAGAACTTGAAACGGAA

TCATCTAACCTTTGACTTACATAGTAACCCATTTATTAAAGGGCTATCCAAGGCTGAGAAACTGCTGGTGCGGACGAGTT

CACTAAGAAGAGAATGGTTGTTCACTCTCGAAACGAAAGAAGTAAAAGAGTGGTTCAAATTGGTAGGTTACAGTGCACTC

ATCAGAGGCTAGTCAGCTATGCATCTGCCCCCTTCTCCTCCGCCATGAGACCCCACTGGCGATCCAGAAGATTAAAGAAA

ACTACATATTGGATAAGTATCTATTCCCAGCTTTGTCTGGT

>MF443350.1/goat/China/2014-04-16

ACCAAACAAAGTTGGGTAAGGATAGATCTTATAATAACTATGGACTGGCAAACTTAGGAGTAAAGATCCTACTGTCGGGG

GGAGGAGGAGGAGCAAGATCCTTGATCATGGCGACTCTCCTTAAAAGCTTAGCATTGTTCAAAAGGAACAAAGACAAAGC

GCCGACAGCATCAGGTTCAGGAGGGGCCATCCGGGGGATTAAGAATGTTATCATAGTCCCGATTCCCGGAGACTCGTCCA

TCACTACCCGTTCAAGACTGCTCGACAGGCTTGTCAGATTGGCCGGAGATCCTGACATCAACGGGTCAAAGCTGACCGGG

GTGATGATCAGCATGTTATCCTTGTTCGTAGAGTCACCCGGGCAATTGATACAGCGAATCACAGATGATCCGGATGTTAG

TATCCGCCTTGTTGAGGTAGTTCAAAGTACTAGGTCCCAGTCCGGGTTGACCTTTGCATCACGTGGTGCTGATTTAGACA

ACGAGGCAGACATGTATTTTTCAACTGAGGGGCCCTCGAGTGGAGGTAAGAAAAGGATCAACTGGTTTGAGAACAGAGAA

ATAATAGACATAGAGGTGCAGGATCCAGAAGAGTTCAATATGTTGTTAGCCTCCATACTAGCACAAGTCTGGATCCTTCT

GGCCAAGGCTGTTACGGCACCAGATACGGCAGCTGACTCAGAACTGAGAAGGTGGGTTAAATACACACAACAAAGGAGAG

TGATTGGGGAATTTCGCCTTGACAAGGGGTGGCTGGACGCAGTCCGCAACAGAATTGCAGAAGATCTATCACTCCGGCGG

TTCATGGTATCTCTCATACTTGACATCAAGAGGACCCCAGGCAACAAGCCAAGGATTGCAGAAATGATCTGCGACATTGA

CAACTATATTGTCGAAGCAGGACTCGCCAGTTTCATCCTTACTATCAAATTTGGTATCGAAACCATGTATCCTGCATTAG

GGCTCCACGAGTTTGCCGGAGAATTGTCCACTATTGAGTCCTTGATGAACTTGTATCAACAGCTAGGAGAGGTTGCACCC

TACATGGTAATTCTAGAGAATTCAATTCAGAACAAGTTTAGTGCAGGAGCTTATCCCCTCCTCTGGAGCTATGCGATGGG

TGTCGGAGTCGAGCTGGAGAACTCAATGGGGGGCTTGAACTTTGGCAGGTCATATTTTGACCCTGCCTATTTTCGTCTCG

GACAGGAGATGGTCAGAAGATCTGCAGGAAAGGTCAGCTCTGTGATTGCGGCTGAGCTCGGCATCACAGCAGAGGAAGCT

AAACTAGTCTCGGAAATCGCCTCGCAGGCTGGGGACGAAAGAACCGCCAGAGGGACTGGGCCTCGACAGGCGCAGGTTTC

CTTCCTTCAGCACCAAACAGGAGGGGGAGAGTCGTCCGCACCAGCGACCAGAGAAGGGGTCAAAGCTGCGATCCCAAACG

GAGCTGAAGAAAGGGATAGAAAGCAAACACGCCCAGGAAGGCCCAGAGGAGAGACCTCCGGCCAACTGCTCCTGGACATC

ATGCCAGAGGATGAGATCTTGCGAGAGTCTGGTCAAAACCCTCGTGAGGCTCAAAGATCGGCCGAGGCACTCTTCAGGCT

GCAGGCCATGGCCAAGATTCTGGAGGACCAGGAGGAGGGAGAAGACAACAGTCAGGTCTACAACGACAAGGATCTCCTCG

GCTAAGCAGACGCACCCTCTGTCGAAATCAGTGACAAGACATCTCCTACCAGTATTATAAAAAACTTAGGACCCAGGTCC

AAGCAAGCACACATCGACACTCCAACCAGTCGAGCGGAGACCACCGATGGCAGAAGAACAAGCATACCATGTCAACAAGG

GGCTGGAATGTATCAAGTCTCTCAAAGCCTCACCCCCGGATCTATCCACCATCAGAGATACCATCGAGAGCTGGAGAGAG

GGGCTTAGCCCCTCGGGCCGTGCAACACCGAACCCTGATACGTCCGAGGGAGATCATCAGAATATCAACCAATCATGCTC

ACCAGCAATCGGACCAAACAAAGTCTACTTGTCTCCTGAAGATAATCTCGGATTTAGAGAGATCACTGGCAACGACTGTG

AGGCTGGGCTCGGAGGGGTCCAGAGAGAAGGATCCAACTCTCAAGTACAGCGTTACCATGTTTATAGTCACGGGGGTGAA

GAGATTGAAGGACTCGAGGATGCTGACTCTCTCGTGGTTCAAGCAGATCCTCCAGTTGCTAACGTCTTCAATGGAGGAGA

GGATGGATCTGACGACAGCGATGTGGACTCTGGCCCAGATGATCCCGGCAGAGATACTCTATATGACCGGGGATCTGTTG

CCGGCAATGGTCTCGCTAGGTCCACAGATGTCGAAAAACTAGAAGGTGCTGATATTCAAGAAGTTCTTAACTCCCAGAAA

GGCAAAGGAGGAAGATTCCAGGGCGGGAAAACCTTGCGAGTCCCGGAAATACCCGATGTCAAGCACTCCAGACCATCAGC

CCAATCAATTAAAAAGGGCACAGACGGGAACTCAGTCTCATCTGGAACGGTGATAGAGTGTTTATCGATAAGTGGTGCAA

CCCAAACTGTGCCAGAGTCAAGATGGGAGTCATCAGAGCAAAATGCGTCTGTGGGGAGTGTCCTCAAGTCTGCGAGGAGT

GCAAAGACGATCCAGGGGTCGACACAAGAATCTGGTACCATAGCATCACTGACTCAGCCTAAAGAGAATGACTCCGAGTA

TGAGTATGAGGATGACCTATTTACAGAGATTCAGGACATCCGTGCAAGCATTGCCAAGATCCATGATGACAATAAAACTA

TCCTCTCAAAGCTTGATTCTATACTGTTATTGAAAGGGGAAGTCGACACTATCAAGAAACAAATCAGCAAGCAGAATATA

AGTATATCCACCATTGAGGGCCATCTCTCCAGTATAATGATAGCCATCCCTGGCTTTGGGAAGGACATCAAGGACCCAAC

ATCCGAAGTCGAGTTGAACCCAGATCTAAGACCTATAATAAGCCGTGACTCTGGCAGAGCTCTCGCGGAGGTCCTCAAGA

AACCCGCTGTGGATAGGTCTCAGAAAATTGGAACCAAAGCCAACTCCAGCTCAAAGGGTCAGCTTCTTAAGGATCTCCAG

CTAAAACCTGTCGACAAGCAGGCAAGCTCTGCAATCGGGTTTGTCCCGTCCGACCATGAATCATCCAGAAATGTCATCCG

CTCCATAATCAAGTCGAGCAAGCTAAATATTGATCACAAGGACTATCTTCTAGATTTACTAAATGATGTGAAAGGCTCCA

AGGATCTTAAGGAATTTCACAAGATGCTAACAGCAATTCTCGCCAAGCACCCGTAACACATCCTCCAGTCATCATCTCAT

ACTCGACTAAAAACATCCTTTCAATCAGGCTATTACAAAAAACTTAGGAGCAAGGGCAACTGAGCTTCGCAGACAGGATG

ACCGAGATCTACGACTTCGATAAATCAGCATGGGATGTCAAAGGGTCAATTGCCCGCATAGAACCCACCACCTACCACGA

CGGCCGACTGATACCCCAGGTGAGGGTCATCGATCCTGGTCTGGGAGACAGAAAAGATGAGTGCTTTATGTACCTGTTTC

TCCTAGGAGTGATTGAGGATAACGACCCCCTGTCTCCCCCCGTCGGAAGAACCTTTGGCTCTCTACCTCTAGGGGTTGGT

AGGTCAACTGCCAAGCCAGAAGAACTACTAAGGGAGGCCACAGAATTAGATATAGTGGTGAGGCGCACTGCAGGAGTAAA

TGAGAAACTGGTATTTTACAACAACACTCCGCTGTCCTTGTTAACGCCCTGGAAGAAAGTTCTGACAACCGGAAGTGTGT

TTAGCGCTAACCAGGTTTGCAATGCAGTCAACCTAGTCCCGCTTGATACTCCCCAGAGATTCAGGGTTGTGTACATGAGC

ATAACTAGATTATCAGACAATGGTTACTATAGTGTGCCCAGAAGAATGCTGGAGTTCCGCTCGGCCAATGCAGTAGCCTT

CAACATCTTGGTTACACTAAGAATTGAAAATGGCACAAACCCTAGAAGATACATAGTCGGCTCATGGGAGAATTCAGAGG

TCACATTTATGGTACACGTGGGCAACTTTAGGAGAAAGAAGAACGAAGTATACTCTGCTGATTACTGCAAAATGAAGATT

GAAAAGATGGGTTTAGTTTTTGCCCTGGGTGGAATAGGCGGAACAAGTCTCCATATTAGAAGCACCGGGAAAATGAGCAA

AACCCTCCATGCACAGCTGGGGTTCAAGAAAATCTTATGTTACCCCCTAATGGATGTCAATGAGGATCTTAACCGATATC

TCTGGCGGGCAGAGTGCCGAATAGTCAAAATCCAAGCTGTCTTACAGCCATCAGTACCCCAAGAGTTCCGTGTCTACGAT

GATGTCATCATCAACGATGATCAAGGCTTGTTCAAGATCTTGTAGTTCATTTGCAAAGGAACGAACCCACCCCCAGGAAG

GGGCCCCACCCTCCCCCG-CCAAAAAACACACTCCAAGGCTCCCCGGCACAGCGCCAACACAGCTACTCCCGCGATGGAA

CGAGGTCGGCC-CCCGAACAAATCCGGTCCAGGGGGGGGGGCCCCCCCCCTGACCAACCGGGCGCCGACCGAGCAGGCCG

GCGAGAAGCCCCCAAGAGGCCCCGACGAAGACCGAGCACCCCATCCCCCC-AAAAACGTCCCCC-AACACGCCCGAGGCT

AGCCACCGGACACCCCCAGACCCGCCACCGGCGAGACAGAACCCCAAGGGGGGACCCTCCAAACCAAAGGAATCACCGGG

GTAGGCTCCGCCGAACCAGCACAGACCCGACCCAAGCAAGGCGGACCCCCAACACGAAACTGCCCAGCACACCCTACTCA

TCATCAAACAAAACTTAGGGGCCAAGTCCATACGAGAGGCCCTAGTCCGTCAGGCGGAGGGACGCAGAAAGGAAGGAGAC

ACGACCGTCCCAGAAGGAGGACACCCAACCACCGAAACGCCCGCCCCCAGGGAAGGAG--CCA-CCCCCACGCAGCGAAG

GAGCGGGTGGGAGACCTCGGGGAGGGCCCGACCGGGACCCGGGGGCCGGCGGCGCCGGCGGTGAGCAGGCCGGGCGCATA

CCTGACCCCACCGGAGGATCCGAGGAGGGCCCGACCCCGCGTGGACCAAACGGCAGGGAGCCCAACCCCACCAAAAAGAG

AGGCGAGGCCCCCCCTCCCTCCCCCCCCGCACACAACCCGCAGACAGACCGAGGGAGGGCCATCCCCCAGCGAGACAGGC

GGCTCCCTCGCAGCCGAGCCGCAGCTCCCCAGAATAGGACACCCCCGCACACTCCGAACACCCCCTCGCCCCCCGCCACC

GGGACAGGGGCTCCACCACCACAGACCGGAGCCACCGACAGCCAGCCAGAGGTCCCCTCCAGTCCCGCCCGGCGCCCCCC

CGAAGAGGCCCCGGCAAGGCCCCCCAGCCCGACGTCCCCCGCACGCACCGAAAGAAGGACCCGCCCGAGAACGTGCGGCA

CATCCATGCGCAAACATCATGACGCGGGTCGCAATTTTGACATTTCTGTTTCTTTTCCCAAATGTTGTTGCGTGTCAGAT

TCACTGGGGCAATCTATCCAAGATCGGGATTGTAGGAACAGGGAGTGCCAGCTACAAGGTGATGACTAGGCCAAGCCACC

AGACTCTGGTTATAAAGTTAATGCCAAATATAACGGCCATCGACAATTGTACAAAGTCAGAGATTGCAGAGTACAAGAGA

TTGCTGATCACAGTGTTAAAGCCTGTAGAGGATGCTCTGTCGGTGATAACCAAGAATGTAAGACCAATTCAAACTCTAAC

ACCTGGGCGTAGAACCCGCCGTTTTGCTGGAGCTGTTCTGGCCGGGGTAGCACTTGGAGTTGCGACAGCCGCTCAGATAA

CTGCAGGAGTCGCCCTTCATCAATCATTGATGAACTCCCAAGCAATTGAGAGTTTAAAAACCAGTCTTGAGATGTCGAAT

CAGGCAATAGAAGAAATCAGACTTGCAAATAAGGAGACCATACTGGCAGTACAGGGCGTCCAGGATTATATCAACAATGA

GCTCGTCCCTTCTGTTCATAGAATGTCATGCGAGCTGGTAGGTCACAAGCTCGGCCTCAAGCTCCTTAGGTACTACACCG

AGATCCTGTCCATATTCGGGCCCAGTCTTCGAGACCCGATATCTGCCGAAATATCAATCCAGGCACTTAGTTATGCATTA

GGCGGAGACATTAATAAAATCCTGGACAAGCTTGGGTATAGCGGTGGGGATTTCCTTGCCATCCTAGAAAGCAAGGGAAT

AAAGGCCCGGGTTACATATGTGGACACAAGAGATTACTTTATAATCCTTAGCATCGCCTACCCAACCTTATCTGAGATCA

AGGGAGTGATAGTTCACAAGATAGAAGCTATAACATACAACATTGGGGCACAGGAGTGGTATACTACTATCCCTAAATAT

GTAGCCACTCAGGGGTATCTGATATCGAACTTTGATGAGACGTCATGCGTATTCACTCCAGAGGGGACAGTTTGCAGCCA

GAATGCGTTGTACCCAATGAGCCCATTGCTTCAGGAATGTTTCAGGGGGTCAACAAAATCGTGCGCCAGAACCTTAGTTT

CAGGGACCATAAGTAATAGATTTATCCTATCAAAAGGGAACCTGATTGCAAATTGTGCGTCAGTTTTGTGCAAATGTTAC

ACAACAGAGACAGTTATCAGCCAAGATCCTGACAAACTACTAACTGTTGTAGCATCCGACAAGTGTCCTGTAGTTGAGGT

GGATGGAGTGACAATACAGGTCGGCAGTCGAGAGTATCCGGATTCTGTATACTTACACAAAATAGACTTAGGTCCAGCCA

TCTCCCTAGAAAAACTGGATGTAGGCACCAATTTAGGCAATGCAGTCACAAGACTGGAGAATGCAAAGGAGCTCCTAGAT

GCATCAGACCAAATACTGAAGACTGTTAAAGGGGTACCTTTCAGTGGGAATATGTACATAGCACTGGCAGCTTGCATAGG

AGTATCCCTGGGCCTTGTCACATTAATATGCTGCTGTAAGGGGAGGTGTAAGAACAAGGAAATCCCTATCTCCAAAATCA

ACCCAGGGCTCAAACCCGACCTGACCGGGACCTCAAAGTCGTACGTAAGATCACTGTAGTCAGAATCACCTGGATCATCT

GGCATCACACACATACATGCACGACACAGGCAGTCCGAGGACGCAAGAAACCCAGCCTCCGGTCACCCACCCGACCCCAC

TCCACGCTCCACCACACATTAGTCATCAAACAAAACTTAGGACGAAAGGTCAATCACCATGTCCGCACAAAGGGAGAGGA

TCAATGCCTTCTACAAAGACAATCCTCACAATAAGAACTATAGGGTGATCCTGGATAGAGAACGCTTGGTCATTGAAAGA

CCCTACATCTTGCTTGGAGTCCTGCTGGTAATGTTCCTGAGTCTAATCGGACTGCTGGCCATTGCAGGGATCAGGCTTCA

CCGGGCCACCGTTGGAACTTCAGAGATCCAGAGTCGGCTGAATACCAATATTAAGTTGACCGAATCTATTGATCACCAGA

CTAAGGATGTCTTAACTCCCCTTTTTAAAATCATTGGCGATGAAGTCGGCATCAGAATTCCACAGAAATTCAGTGATCTT

GTCAAGTTCATCTCCGATAAGATTAAATTCCTCAACCCTGATAGAGAGTATGATTTCAGAGATCTCCGGTGGTGTATGAA

TCCCCCCGAGAGAGTCAAAATTAATTTTGATCAGTTTTGTGAGTACAAGGCTGCGGTTAAGTCAATTGAACATATATTTG

AGTCACCACTCAACAAGTCAAAAAAGCTGCAATCTTTGACTCTCGGGCCCGGAACAGGCTGTCTAGGCAGGACAGTAACA

AGAGCCCATTTCTCAGAACTTACAATGACCTTAATGGACCTGGATCTAGAGATGAAGCACAACGTGTCCTCAGTGTTTAC

CGTAGTTGAAGAGGGATTATTCGGAAGAACATATACCGTCTGGAGATCCGATGCCAGGGATCCGAGCACCGATCTAGGTA

TCGGCCATTTTTTAAGAGTCTTCGAGATTGGACTGGTAAGAGATCTTGGGCTGGGTCCCCCTGTTTTTCATATGACCAAC

TATCTCACGGTGAACATGAGTGATGACTATCGGAGATGTCTTTTAGCGGTAGGGGAGTTGAAGTTGACAGCCCTATGCAC

CTCATCTGAGACTGTGACACTGAGTGAGAGAGGAGTTCCAAGGAGGGAACCTCTTGTGGTTGTGATACTTAATCTAGCTG

GACCCACTCTAGGGGGAGAGCTATACAGTGTCTTGCCTACCTCTGATCTCATGGTGGAGAAACTCTATTTGTCTTCACAT

AGAGGGATCATCAAAGATGATGAGGCCAATTGGGTAGTGCCGTCTACCGATGTTCGTGATCTTCAGAATAAAGGTGAATG

TCTGGTGGAAGCATGCAAGACTCGACCTCCTTCATTTTGCAATGGCACAGGATCAGGCCCGTGGTCAGAAGGGAGAATCC

CTGCCTACGGGGTGATCAGGGTCAGTCTTGACTTAGCGAGTGACCCGGATGTAGTTATCACTTCAGTGTTTGGCCCACTG

ATACCTCACCTATCCGGCATGGATCTTTACAACAACCCGTTTTCAAAAGCTGTATGGTTGGCTGTACCACCTTATGAGCA

GTCATTTCTAGGAATGATAAATACAATTGGATTCCCTAACAGAGCAGAGGTTATGCCGCACATTTTGACCACAGAGATCA

GAGGCCCTCGGGGTCGTTGCCATGTTCCCATAGAATTGTCCCGCCGGGTTGATGACGATATCAAGATCGGGTCCAACATG

GTCATATTGCCGACGATGGACCTGAGGTATATTACAGCCACTTATGATGTTTCCAGGAGCGAGCATGCAATCGTGTACTA

TATCTATGACACGAGTCGCTCATCATCTTACTTCTACCCAGTTCGACTGAATTTCAAAGGCAATCCTCTCTCTTTGAGGA

TAGAGTGTTTCCCCTGGCGTCATAAGGTGTGGTGCTACCATGATTGTCTTATATACAACACCATAACAGGTGAAGAGGTC

CATACGAGAGGGCTGACCGGCATAGAGGTAACATGCAATCCAGTCTGAGCAGAGCTACGACCGTCACTCAGGCAGTCCCT

TGAGTCGCCACCGAGTCCAAGCAGCACAGCCTGGGACAATCAACGGCACAGCCCAGCCAACAATGTTATAAAAAACTTAG

GAGCCAGGGTTGTAGGGGCCATGGACTCTCTATCAGTCAATCAGGTCTTGTACCCTGAGGTTCATCTAGACAGCCCTATT

GTCACAAACAAACTAGTTGCCATCCTTGAGTACTCGGGTATCGACCATAACTATGTTCTTGAAGACCAGACTCTTGTCAA

GAATATTAGGTATAGGCTGGGGTGCGGTTTTTCAAATCAAATGATCATCAATAACAGGGGGGTGGGTGAAACAGTCAATT

CTAAACTTAAAAGTTACCCCCATAATCGTCATATCATATACCCGGATTGCAATAAGGAGTTGTTTTGTATCAAGGATAGC

TGCATATCTAAGAAGCTCTCGGAGCTATTCAAGAAGGGTAATTCCTTGTACTCTAAGATAAGTCACCAGGTACTGGATTG

TCTTAAGAGAGTCAATGGGAAATTAGGTCTGGGCACAGATCTTACCCATGGTCTGAAGGAGGGTATCCTTGACCTAGGGT

TGCACATGCATAGCTCTCAATGGTTCGAGACTTTCCTGTTCTGGTTCACTATCAAGACAGAGATGAGATCAATGATCAAA

GAACAGTCCCATATATGCCACAAGAGGAGGTATAACCCGACTTTTGTGTCGGGGGATGCATTCGAGGTGCTCGTATCGCG

AGACCTCGTTGTGATAATTGATAAGAATACCCAGTATGTCTTCTACTTAACATTTGAGCTGGTCCTTATGTATTGTGATG

TCATAGAGGGTAGACTTATGACGGAGACAGCCATGGCCATAGACCAGAGATATTCAGAACTTCTAAGCCGGGTCAGATAC

TTGTGGGATCTTATTGATGGGTTCTTCCCAACACTAGGCAACACCACATACCAAATTGTTGCTCTGCTTGAACCATTATC

GTTGGCTTATCTTCAACTTCAGGATGTCACTCTGGAGTTAAGAGGTGCTTTTTTAGACCATTGCTTCAAAGAACTCTATG

AGATACTGGAACATTGCGGCATTGACACAGCAGGCACTTACAATTCCATCACTGAAGGATTGGATTACGTATTCATCACC

CACGATATACATCTAACTGGGGAGGTTTTTTCATTTTTCCGGAGTTTCGGACACCCCCGTCTAGAAGCAGTCACCGCTGC

GGAAAATGTCAGAAAACATATGAACCAACCGAAGGTAATCAGTTATGAGACTATGATGAAAGGCCATGCGGTATTTTGCG

GGATAATCATAAATGGTTTTCGGGACCGACATGGCGGCAGCTGGCCCCCTGTTGCACTGCCAGAACATGCCTCTGCTGCG

ATCCGGAATGCGCAGGCATCAGGTGAGGGACTGACCCATGACCTGTGTATAGACAACTGGAAATCCTTTGTTGGATTCAA

ATTTGGCTGCTTTATGCCACTCAGCCTAGATAGTGATTTGACCATGTATCTCAAAGACAAGGCATTGGCTGCACTGAAGA

ATGAGTGGGATTCAGTTTACCCGAAAGAATACCTCCGTTATAACCCACCTAGAGGGACAGAGTCAAGGCGATTGGTAGAG

GTATTCCTGAATGACTCCAGCTTTGATCCTTATAACATGATAATGTACGTGGTGAATGGCTCCTACCTTAAAGACCCTGA

GTTTAACCTCTCATACAGCCTAAAAGAGAAGGAGATAAAGGAGACAGGGCGGTTGTTTGCCAAAATGACCTATAAGATGC

GGGCCTGTCAGGTAATCGCTGAAAATCTGATATCGAATGGTGTTGGGAAGTATTTCCGAGACAATGGGATGGCAAAGGAC

GAGCATGACCTAACAAAAGCCCTTCACACCCTGGCAGTCTCAGGTGTTCCCAAAAACAACAAAGATAACCACCGAGGTGG

GCCTCCCAGAAGGACAACAAACCGAGGGGTGAGATCAAGCCAAGGCACCAAAACACAAGATAGAGACAAGGTTCAAGGGG

GACCTATGTACAACTATTTGCGATGCCAACCGATCAGCCCTGATCAGGGTGAGTCATACGAGACTGTTAGTGCATTCATC

ACCGCTGACCTTAAGAAGTATTGCCTGAATTGGAGATACGAGACAATCAGCATATTTGCACAGAGGCTGAATGAAATATA

TGGACTACCATCCTTCTTTCAATGGTTACACAGGATATTGGAAAAATCCGTACTCTACGTCAGTGACCCACATTGTCCTC

CCGATCTAGATAATCATATCCCTCTGGACAGTGTCCCCAATGCCCAAATATTCATCAAGTACCCAATGGGCGGAATAGAA

GGTTATTGCCAAAAACTATGGACAATCAGCACTATACCATACTTGTATCTGGCAGCCTATGAGAGCGGAGTAAGAATCGC

CTCACTGGTGCAGGGTGACAATCAGACAATCGCAGTGACAAAAAGAGTTCCAAGTTCTTGGCCTTATTCACTAAAAAAGA

GGGAGGCCTCCAAGGCCGCTCAAAATTACTTCGTAGTCTTAAGGCAAAGATTGCACGATGTAGGTCATCACTTAAAGGCT

AATGAGACCATAGTATCTTCTCACTTTTTTGTATATTCCAAAGGGATTTATTATGACGGCCTGCTAGTCTCACAATCACT

AAAGAGCATCGCCAGATGTGTCTTCTGGTCCGAGACTATAGTGGATGAAACCAGAGCGGCGTGCAGTAATATTGCAACAA

CTGTCGCTAAGAGTATAGAGAGGGGTTATGATAGGTACCTTGCATATTCTTTGAATATCCTCAAGATTTTCCAACAGATC

CTTATATCCCTTAACTTCACTATTAACACAACAATGACTCAGGATGTCGTGGCACCGATCATCGAGAACGGTGATTTGCT

GATAAGGATGGCACTCTTGCCAGCACCCATCGGGGGTCTCAATTATCTTAACATGAGCAGGTTATTTGTGAGAAATATCG

GTGACCCGGTCACCTCCTCTATAGCCGACCTGAAGAGGATGATAGATGCCGGGCTAATGCCAGAAGAAACATTGCATCAA

GTGATGACCCAGACCCCGGGAGAATCATCCTACCTAGATTGGGCAAGTGACCCTTATTCTGCCAACCTACCCTGCGTACA

GAGTATCACTCGCCTTCTCAAAAACATCACTGCACGATATATTTTAATCAGCAGCCCAAACCCGATGCTGAAAGGGTTAT

TTCATGAGGGGAGTAGAGATGAAGACGAGGAGCTTGCAAGTTTCCTAATGGATCGGCATATAATTGTTCCAAGAGCTGCA

CATGAAATCTTAGACCATAGTATAACCGGAGCAAGAGAAGCTATAGCCGGGATGTTGGACACCACCAAGGGTCTGATTAG

AACAAGTATGAAACGGGGCGGCCTCACCCCTCGAGTCTTAGCCCGCCTTTCCAATTATGATTATGAACAATTCCGATCCG

GGATAACACTATTAACAAAGAAAGGGCAGTGTTATCTCATTGACAAGGACTCGTGCTCGGTGCAGCTCGCTATAGCTCTG

AGGGGCCACATGTGGGCTAAGTTGGCTCGCGGGAGACCAATCTATGGGTTGGAGGTGCCTGATGTATTAGAATCGATGAA

CGGCTACCTTATCAAACGCCACGAGTCGTGTGCTATCTGTGAAACGGGCTCAAGTCACTACGGATGGTTTTTCGTCCCTG

CAGGGTGCCAGCTTGACGATGTATCTAGAGAGACTTCGGCTCTTCGTGTACCTTATGTCGGATCAACAACTGAGGAAAGG

ACAGACATGAAACTTGCATTTGTTAGGTCTCCAAGTAGATCCCTCAAATCAGCAGTTAGAATTGCAACAGTTTACTCATG

GGCTTACGGGGATGATGAGAAATCATGGAGTGAAGCCTGGATGCTAGCTAGGCAGAGAGCCAATATCACCTTAGATGAAT

TGAGAATGATCACTCCAGTCTCCACATCCACCAACTTAGCCCACCGGTTGAGGGATCGGAGTACCCAGGTGAAATACTCG

GGGACATCCCTTGTGAGGGTTGCAAGATACACAACCATCTCCAATGATAACTTGTCGTTTGTGATATCTGAGAAAAAAGT

GGATACCAACTTCATTTATCAGCAAGGGATGTTGCTTGGTCTTGGGATCCTTGAGAACCTCTTTAGGCTAGAGGCCACCA

CCGGAGTATCCAACACAGTGCTACACCTGCACGTGGAAACAGAATGTTGTGTTGTACCTATGGTGGATCACCCAAGGATA

CCGAGTCTCCGTAATATAAAAGTTACGGATGAGCTATGCACAAACCCTTTGATCTACGACAGGTCCCCCATCATAGAGCA

CGATGCAACCCGGCTATATTCACAGAGCCACAGGAGACATTTGGTGGAGTTTGTTACCTGGTCAACAAGCCAGCTTTACC

ATATACTGGCTAAGTCTACAGCAATGTCCATGATTGAATTGATCACGAGATTCGAGAAAGATCACATGAATGAAATAGCC

GCCCTGATTGGCGATGACGACATCAACAGCTTCATCACAGAATTTTTGCTAGTGGAGCCTAGATTATTTATAGTTTACCT

TGGTCAGTGTGCTGCCATCAATTGGGCCTTTGATATACATTATCATCGGCCCTCGGGCAAGTACCAGATGGGGGAACTTC

TCTACTCTCTGCTCTCTCGGATGAGCAAAGGAGTATATAAGATCTTTACTAATGCTTTGAGCCACCCCAAAGTTTACAAG

AAATTTTGGCGAAGTGGCATCATTGAGCCGATTCATGGCCCATCCTTGGATACACAGAATTTACATGTCACTGTCTGTGA

CATGATATATGGGTCCTATGTCACCTATTTGGATCTTTTGCTGAATGATGAGCTAGATGATTACCCGTATTTGCTCTGCG

AGAGTGATGAGGACGTGGTCACAGACAGGTTCGACAACATTCAAGCCAAACATCTCTGTGTATTGGCCGATGTATATTGC

AGCTCCAAGAGGTGTCCTTCAATAATCGGGATGTCTCCTATAGAAAAATGTACCATTCTCACACATTACATCAAGGGAGA

ATCAGTACAATCCCCGTCTGGGACCTCATGGAACACTGATCCCCTTGTAGTAGATCATTACTCATGCTCTCTGACCTACC

TTCGCCGCGGTTCCATCAAACAAATCAGGTTGAGGGTGGATCCTGGGTTTGTATTCGAGGCGTTGACAGACGTCGACTTT

AAACAACCTCGCAAAGCTAAGTTAGATATATCGGTCGTTGGGTTGACTGATTTCTCTCCCCCTTACGACAACGTCGGTGA

TTTTTTAGGGACTATCAACACATTGAGGCACGATCTGCCTGTCACCGGAACCGGGGTCTCGAACTATGAAGTCCACGCTT

ATCGTAGAATTGGCCTGAATTCATCAGCATGTTATAAAGCCGTAGAGATCTCCACGTTAATCAAGCCATCCTTAGAAGTC

GGAGAGCATGGATTGTTCTTAGGAGAAGGTTCCGGTTCAATGCTGGCTGCGTACAAAGAAGTTCTTAAATTAGCAAATTG

TTACTACAACAGCGGAGTAACAGCGGAGGGCAGAGCCGGACAGAGGGAAATATCTCCCTATCCTTCAGAGATGAGCCTAG

TAGAGAATCAAATGGGGATAGAGCGGAGTGTTAAAGTGCTGTTCAACGGGAAACCTGAAGTAACTTGGGTGGGGACCACA

GATTGCTACAAGTACATAATCAGTAACATCCAAACCTCTAGTCTGGGTTTCATACATTCAGATATTGAGACGCTCCCAAC

CAAAGATGCTGTTGAGAAATTGGAGGAATTTGCCTCTATCCTATCCTTATCCTTGATTTTGGGGAAAATCGGCTCTATTA

CAGTTGTCAAAATTATGCCCATTAGCGGGGATTTTACCCAAGGCTTCATAGCCTATGCCATCCAATATTTTAGGGAGAGC

CTGCTTGCCTATCCGAGATATAGTAACTTCATCTCGACTGAGTGTTACCTTATTATGATAGGATTAAAGGCCAATCGATT

GATAAACCCAGAAGCCATTAAGCAAAGCATAATCAGAGCGGGGGTCAGGACTGCACCAGGACTTGTGAGCCATGTATTAT

CAGAGAAACAAAAAGGTTGTATTCAATCTTTCTTGGGTGATCCTTATATCCAAGGAGACTTCAATAAGCACCTTAAATCT

CTAACCTCTATTGAGAAAATCCTGGTAAATTGTGGTCTCTCGATCAATGGCATAAAAATCTGTAGAGATCTAATCCACCA

TGATATCGCCTCCGGTCCAGACGGCCTTATGAGCTCTACAATTATTTTATATAGGGAATTGGCTCATTTCAAAGATAATA

TAAGGAGTCAGCACGGAATGTTCCACCCCTACCCGGTACTGGCCAACAGCAGGCAACGTGAATTAATCCTTCGAATAGCC

AAGAAATTCTGGGGGTATGTCTTGCTATATTCTGATGACCCGGCACTAATCAGACAAACAATCAAGAACTTGAAACGGAA

TCATCTAACCTTTGACTTACATAGTAACCCATTTATTAAAGGGCTATCCAAGGCTGAGAAACTGCTGGTGCGGACGAGTT

CACTAAGAAGAGAATGGTTGTTCACTCTCGAAACGAAAGAAGTAAAAGAGTGGTTCAAATTGGTAGGTTACAGTGCACTC

ATCAGAGGCTAGTCAGCTATGCATCTGCCCCCTTCTCCTCCGCCATGAGACCCCACTGGCGATCCAGAAGATTAAAGAAA

ACTACATATTGGATAAGTATCTATTCCCAGCTTTGTCTGGT

>MF443342.1/goat/China/2014-04-01

ACCAAACAAAGTTGGGTAAGGATAGATCTTATAATAACTATGGACTGGCAAACTTAGGAGTAAAGATCCTACTGTCGGGG

GGAGGAGGAGGAGCAAGATCCTTGATCATGGCGACTCTCCTTAAAAGCTTAGCATTGTTCAAAAGGAACAAAGACAAAGC

GCCGACAGCATCAGGTTCAGGAGGGGCCATCCGGGGGATTAAGAATGTTATCATAGTCCCGATTCCCGGAGACTCGTCCA

TCACTACCCGTTCAAGACTGCTCGACAGGCTTGTCAGATTGGCCGGAGATCCTGACATCAACGGGTCAAAGCTGACCGGG

GTGATGATCAGCATGTTATCCTTGTTCGTAGAGTCACCCGGGCAATTGATACAGCGAATCACAGATGATCCGGATGTTAG

TATCCGCCTTGTTGAGGTAGTTCAAAGTACTAGGTCCCAGTCCGGGTTGACCTTTGCATCACGTGGTGCTGATTTAGACA

ACGAGGCAGACATGTATTTTTCAACTGAGGGGCCCTCGAGTGGAGGTAAGAAAAGGATCAACTGGTTTGAGAACAGAGAA

ATAATAGACATAGAGGTGCAGGATCCAGAAGAGTTCAATATGTTGTTAGCCTCCATACTAGCACAAGTCTGGATCCTTCT

GGCCAAGGCTGTTACGGCACCAGATACGGCAGCTGACTCAGAACTGAGAAGGTGGGTTAAATACACACAACAAAGGAGAG

TGATTGGGGAATTTCGCCTTGACAAGGGGTGGCTGGACGCAGTCCGCAACAGAATTGCAGAAGATCTATCACTCCGGCGG

TTCATGGTATCTCTCATACTTGACATCAAGAGGACCCCAGGCAACAAGCCAAGGATTGCAGAAATGATCTGCGACATTGA

CAACTATATTGTCGAAGCAGGACTCGCCAGTTTCATCCTTACTATCAAATTTGGTATCGAAACCATGTATCCTGCATTAG

GGCTCCACGAGTTTGCCGGAGAATTGTCCACTATTGAGTCCTTGATGAACTTGTATCAACAGCTAGGAGAGGTTGCACCC

TACATGGTAATTCTAGAGAATTCAATTCAGAACAAGTTTAGTGCAGGAGCTTATCCCCTCCTCTGGAGCTATGCGATGGG

TGTCGGAGTCGAGCTGGAGAACTCAATGGGGGGCTTGAACTTTGGCAGGTCATATTTTGACCCTGCCTATTTTCGTCTCG

GACAGGAGATGGTCAGAAGATCTGCAGGAAAGGTCAGCTCTGTGATTGCGGCTGAGCTCGGCATCACAGCAGAGGAAGCT

AAACTAGTCTCGGAAATCGCCTCGCAGGCTGGGGACGAAAGAACCGCCAGAGGGACTGGGCCTCGACAGGCGCAGGTTTC

CTTCCTTCAGCACCAAACAGGAGGGGGAGAGTCGTCCGCACCAGCGACCAGAGAAGGGGTCAAAGCTGCGATCCCAAACG

GAGCTGAAGAAAGGGACAGAAAGCAAACACGCCCAGGAAGGCCCAGAGGAGAGACCTCCGGCCAACTGCTCCTGGACATC

ATGCCAGAGGATGAGATCTTGCGAGAGTCTGGTCAAAACCCTCGTGAGGCTCAAAGATCGGCCGAGGCACTCTTCAGGCT

GCAGGCCATGGCCAAGATTCTGGAGGACCAGGAGGAGGGAGAAGACAACAGTCAGGTCTACAACGACAAGGATCTCCTCG

GCTAAGCAGACGCACCCTCTGTCGAAATCAGTGACAAGACATCTCCTACCAGTATTATAAAAAACTTAGGACCCAGGTCC

AAGCAAGCACACATCGACACTCCAACCAGTCGAGCGGAGACCACCGATGGCAGAAGAACAAGCATACCATGTCAACAAGG

GGCTGGAATGTATCAAGTCTCTCAAAGCCTCACCCCCGGATCTATCCACCATCAGAGATACCATCGAGAGCTGGAGAGAG

GGGCTTAGCCCCTCGGGCTGTGCAACACCGAACCCTGATACGTCCGAGGGAGATCATCAGAATATCAACCAATCATGCTC

ACCAGCAATCGGACCAAACAAAGTCTACTTGTCTCCTGAAGATAATCTCGGACTTAGAGAGATCACTGGCAACGACTGTG

AGGCTGGGCTCGGAGGGGTCCAGAGAGAAGGATCCAACTCTCAAGTACAGCGTTACCATGTTTATAGTCACGGGGGTGAA

GAGATTGAAGGACTCGAGGATGCTGACTCTCTCGTGGTTCAAGCAGATCCTCCAGTTGCTAACGTCTTCAATGGAGGAGA

GGATGGATCTGACGACAGCGATGTGGACTCTGGCCCAGATGATCCCGGCAGAGATACTCTATATGACCGGGGATCTGTTG

CCGGCAATGGTATCGCTAGGTCCACAGATGTCGAAAAACTAGAAGGTGCTGATATTCAAGAAGTTCTTAACTCCCAGAAA

GGCAAAGGAGGAAGATTCCAGGGCGGGAAAACCTTGCGAGTCCCGGAAATACCCGATGTCAAGCACTCCAGACCATCAGC

CCAATCAATTAAAAAGGGCACAGACGGGAACTCAGTCTCATCTGGAACGGTGATAGAGTGTTTATCGATAAGTGGTGCAA

CCCAAACTGTGCCAGAGTCAAGATGGGAGTCATCAGAGCAAAATGCGTCTGTGGGGAGTGTCCTCAAGTCTGCGAGGAGT

GCAAAGACGATCCAGGGGTCGACACAAGAATCTGGTACCATAGCATCACTGACTCAGCCTAAAGAGAATGACTCCGAGTA

TGAGTATGAGGATGACCTATTTACAGAGATTCAGGACATCCGTGCAAGCATTGCCAAGATCCATGATGACAATAAAACTA

TCCTCTCAAAGCTTGATTCTATACTGTTATTGAAAGGGGAAGTCGACACTATCAAGAAACAAATCAGCAAGCAGAATATA

AGTATATCCACCATTGAGGGCCATCTCTCCAGTATAATGATAGCCATCCCTGGCTTTGGGAAGGACATCAAGGACCCAAC

ATCCGAAGTCGAGTTGAACCCAGATCTAAGACCTATAATAAGCCGTGACTCTGGCAGAGCTCTCGCGGAGGTCCTCAAGA

AACCCGCTGTGGATAGGTCTCAGAAAATTGGAACCAAAGCCAACTCCAGCTCAAAGGGTCAGCTTCTTAAGGATCTCCAG

CTAAAACCTGTCGACAAGCAGGCAAGCTCTGCAATCGGGTTTGTCCCGTCCGACCATGAATCATCCAGAAATGTCATCCG

CTCCATAATCAAGTCGAGCAAGCTAAATATTGATCACAAGGACTATCTTCTAGATTTACTAAATGATGTGAAAGGCTCCA

AGGATCTTAAGGAATTTCACAAGATGCTAACAGCAATTCTCGCCAAGCACCCGTAACACATCCTCCAGTCATCATCTCAT

ACTCGACTAAAAACATCCTTTCAATCAGGCTATTACAAAAAACTTAGGAGCAAGGGCAACTGAGCTTCGCAGACAGGATG

ACCGAGATCTACGACTTCGATAAATCAGCATGGGATGTCAAAGGGTCAATTGCCCGCATAGAACCCACCACCTACCACGA

CGGCCGACTGATACCCCAGGTGAGGGTCATCGATCCTGGTCTGGGAGACAGAAAAGATGAGTGCTTTATGTACCTGTTTC

TCCTAGGAGTGATTGAGGATAACGACCCCCTGTCTCCCCCCGTCGGAAGAACCTTTGGCTCTCTACCTCTAGGGGTTGGT

AGGTCAACTGCCAAGCCAGAAGAACTACTAAGGGAGGCCACAGAATTAGATATAGTGGTGAGGCGCACTGCAGGAGTAAA

TGAGAAACTGGTATTTTACAACAACACTCCGCTGTCCTTGTTAACGCCCTGGAAGAAAGTTCTGACAACCGGAAGTGTGT

TTAGCGCTAACCAGGTTTGCAATGCAGTCAACCTAGTCCCGCTTGATACTCCCCAGAGATTCAGGGTTGTGTACATGAGC

ATAACTAGATTATCAGACAATGGTTACTATAGTGTGCCCAGAAGAATGCTGGAGTTCCGCTCGGCCAATGCAGTAGCCTT

CAACATCTTGGTTACACTAAGAATTGAAAATGGCACAAACCCTAGAAGATACATAGTCGGCTCATGGGAGAATTCAGAGG

TCACATTTATGGTACACGTGGGCAACTTTAGGAGAAAGAAGAACGAAGTATACTCTGCTGATTACTGCAAAATGAAGATT

GAAAAGATGGGTTTAGTTTTTGCCCTGGGTGGAATAGGCGGAACAAGTCTCCATATTAGAAGCACCGGGAAAATGAGCAA

AACCCTCCATGCACAGCTGGGGTTCAAGAAAATCTTATGTTACCCCCTAATGGATGTCAATGAGGATCTTAACCGATATC

TCTGGCGGGCAGAGTGCCGAATAGTCAAAATCCAAGCTGTCTTACAGCCATCAGTACCCCAAGAGTTCCGTGTCTACGAT

GATGTCATCATCAACGATGATCAAGGCTTGTTCAAGATCTTGTAGTTCATTTGCAAAGGAACGAACCCACCCCCAGGAAG

GGGCCCCACCCTCCCCCG-CCAAAAAACACACTCCAAGGCTCCCCGGCACAGCGCCAACACAGCTACTCCCGCGATGGAA

CGAGGTCGGCC-CCCGAACAAATCCGGTCCAGGGGGGGGGGCCCCCCCCCTGACCAACCGGGCGCCGACCGAGCAGGCCG

GCGAGAAGCCCCCAAGAGGCCCCGACGAAGACCGAGCACCCCATCCCCCC-AAAAACGTCCCCC-AACACGCCCGAGGCT

AGCCACCGGACACCCCCAGACCCGCCACCGGCGAGACAGAACCCCAAGGGGGGACCCTCCAAACCAAAGGAATCACCGGG

GTAGGCTCCGCCGAACCAGCACAGACCCGACCCAAGCAAGGCGGACCCCCAACACGAAACTGCCCAGCACACCCTACTCA

TCATCAAACAAAACTTAGGGGCCAAGTCCATACGAGAGGCCCTAGTCCGTCAGGCGGAGGGACGCAGAAAGGAAGGAGAC

ACGACCGTCCCAGAAGGAGGACACCCAACCACCGAAACGCCCGCCCCCAGGGAAGGAG--CCA-CCCCCACGCAGCGAAG

GAGCGGGTGGGAGACCTCGGGGAGGGCCCGACCGGGACCCGGGGGCCGGCGGCGCCGGCGGTGAGCAGGCCGGGCGCATA

CCTGACCCCACCGGAGGATCCGAGGAGGGCCCGACCCCGCGTGGACCAAACGGCAGGGAGCCCAACCCCACCAAAAAGAG

AGGCGAGGCCCCCCCTCCCTCCCCCCCCGCACACAACCCGCAGACAGACCGAGGGAGGGCCATCCCCCAGCGAGACAGGC

GGCTCCCTCGCAGCCGAGCCGCAGCTCCCCAGAACAGGACACCCCCGCACACCCCGAACACCCCCCCGCCCCCCGCCACC

GGGACAGGGGCTCCACCACCACAGACCGGAGCCACCGACAGCCAGCCAGAGGTCCCCTCCAGTCCCGCCCGGCGCCCCCC

CGAAGAGGCCCCGGCAAGGCCCCCCAGCCCGACGTCCCCCGCACGCACCGAAAGAAGGACCCGCCCGAGAACGTGCGGCA

CATCCATGCGCAAACATCATGACGCGGGTCGCAATTTTGACATTTCTGTTTCTTTTCCCAAATGTTGTTGCGTGTCAGAT

TCACTGGGGCAATCTATCCAAGATCGGGATTGTAGGAACAGGGAGTGCCAGCTACAAGGTGATGACTAGGCCAAGCCACC

AGACTCTGGTTATAAAGTTAATGCCAAATATAACGGCCATCGACAATTGTACAAAGTCAGAGATTGCAGAGTACAAGAGA

TTGCTGATCACAGTGTTAAAGCCTGTAGAGGATGCTCTGTCGGTGATAACCAAGAATGTAAGACCAATTCAAACTCTAAC

ACCTGGGCGTAGAACCCGCCGTTTTGCTGGAGCTGTTCTGGCCGGGGTAGCACTTGGAGTTGCGACAGCCGCTCAGATAA

CTGCAGGAGTCGCCCTTCATCAATCATTGATGAACTCCCAAGCAATTGAGAGTTTAAAAACCAGTCTTGAGATGTCGAAT

CAGGCAATAGAAGAAATCAGACTTGCAAATAAGGAGACCATACTGGCAGTACAGGGCGTCCAGGATTATATCAACAATGA

GCTCGTCCCTTCTGTTCATAGAATGTCATGCGAGCTGGTAGGTCACAAGCTCGGCCTCAAGCTCCTTAGGTACTACACCG

AGATCCTGTCCATATTCGGGCCCAGTCTTCGAGACCCGATATCTGCCGAAATATCAATCCAGGCACTTAGTTATGCATTA

GGCGGAGACATTAATAAAATCCTGGACAAGCTTGGGTATAGCGGTGGGGATTTCCTTGCCATCCTAGAAAGCAAGGGAAT

AAAGGCCCGGGTTACATATGTGGACACAAGAGATTACTTTATAATCCTTAGCATCGCCTACCCAACCTTATCTGAGATCA

AGGGAGTGATAGTTCACAAGATAGAAGCTATAACATACAACATTGGGGCACAGGAGTGGTATACTACTATCCCTAAATAT

GTAGCCACTCAGGGGTATCTGATATCGAACTTTGATGAGACGTCATGCGTATTCACTCCAGAGGGGACAGTTTGCAGCCA

GAATGCGTTGTACCCAATGAGCCCATTGCTTCAGGAATGTTTCAGGGGGTCAACAAAATCGTGCGCCAGAACCTTAGTTT

CAGGGACCATAAGTAATAGATTTATCCTATCAAAAGGGAACCTGATTGCAAATTGTGCGTCAGTTTTGTGCAAATGTTAC

ACAACAGAGACAGTTATCAGCCAAGATCCTGACAAACTACTAACTGTTGTAGCATCCGACAAGTGTCCTGTAGTTGAGGT

GGATGGAGTGACAATACAGGTCGGCAGTCGAGAGTATCCGGATTCTGTATACTTACACAAAATAGACTTAGGTCCAGCCA

TCTCCCTAGAAAAACTGGATGTAGGCACCAATTTAGGCAATGCAGTCACAAGACTGGAGAATGCAAAGGAGCTCCTAGAT

GCATCAGACCAAATACTGAAGACTGTTAAAGGGGTACCTTTCAGTGGGAATATGTACATAGCACTGGCAGCTTGCATAGG

AGTATCCCTGGGCCTTGTCACATTAATATGCTGCTGTAAGGGGAGGTGTAAGAACAAGGAAATCCCTATCTCCAAAATCA

ACCCAGGGCTCAAACCCGACCTGACCGGGACCTCAAAGTCGTACGTAAGATCACTGTAGTCAGAATCACCTGGATCATCT

GGCATCACACACATACATGCACGACACAGGCAGTCCGAGGACGCAAGAAACCCAGCCTCCGGTCACCCACCCGACCCCAC

TCCACGCTCCACCACACATTAGTCATCAAACAAAACTTAGGACGAAAGGTCAATCACCATGTCCGCACAAAGGGAGAGGA

TCAATGCCTTCTACAAAGACAATCCTCACAATAAGAACCATAGGGTGATCCTGGATAGAGAACGCTTGGTCATTGAAAGA

CCCTACATCTTGCTTGGAGTCCTGCTGGTAATGTTCCTGAGTCTAATCGGACTGCTGGCCATTGCAGGGATCAGGCTTCA

CCGGGCCACCGTTGGAACTTCAGAGATCCAGAGTCGGCTGAATACCAATATTAAGTTGACCGAATCTATTGATCACCAGA

CTAAGGATGTCTTAACTCCCCTTTTTAAAATCATTGGCGATGAAGTCGGCATCAGAATTCCACAGAAATTCAGTGATCTT

GTCAAGTTCATCTCCGATAAGATTAAATTCCTCAACCCTGATAGAGAGTATGATTTCAGAGATCTCCGGTGGTGTATGAA

TCCCCCCGAGAGAGTCAAAATTAATTTTGATCAGTTTTGTGAGTACAAGGCTGCGGTTAAGTCAATTGAACATATATTTG

AGTCACCACTCAACAAGTCAAAAAAGCTGCAATCTTTGACTCTCGGGCCCGGAACAGGCTGTCTAGGCAGGACAGTAACA

AGAGCCCATTTCTCAGAACTTACAATGACCTTAATGGACCTGGATCTAGAGATGAAGCACAACGTGTCCTCAGTGTTTAC

CGTAGTTGAAGAGGGATTATTCGGAAGAACATATACCGTCTGGAGATCCGATGCCAGGGATCCGAGCACCGATCTAGGTA

TCGGCCATTTTTTAAGAGTCTTCGAGATTGGACTGGTAAGAGATCTTGGGCTGGGTCCCCCTGTTTTTCATATGACCAAC

TATCTCACGGTGAACATGAGTGATGACTATCGGAGATGTCTTTTAGCGGTAGGGGAGTTGAAGTTGACAGCCCTATGCAC

CTCATCTGAGGCTGTGACACTGAGTGAGAGAGGAGTTCCAAGGAGGGAACCTCTTGTGGTTGTGATACTTAATCTAGCTG

GACCCACTCTAGGAGGAGAGCTATACAGTGTCTTGCCTACCTCTGATCTCATGGTGGAGAAACTCTATTTGTCTTCACAT

AGAGGGATCATCAAAGATGATGAGGCCAATTGGGTAGTGCCGTCTACCGATGTTCGTGATCTTCAGAATAAAGGTGAATG

TCTGGTGGAAGCATGCAAGACTCGACCTCCTTCATTTTGCAATGGCACAGGATCAGGCCCGTGGTCAGAAGGGAGAATCC

CTGCCTACGGGGTGATCAGGGTCAGTCTTGACTTAGCGAGTGACCCGGATGTAGTTATCACTTCAGTGTTTGGCCCACTG

ATACCTCACCTATCCGGCATGGATCTTTACAACAACCCGTTTTCAAAAGCTGTATGGTTGGCTGTACCACCTTATGAGCA

GTCATTTCTAGGAATGATAAATACAATTGGATTCCCTAACAGAGCAGAGGTTATGCCGCACATTTTGACCACAGAGATCA

GAGGCCCTCGGGGTCGTTGCCATGTTCCCATAGAATTGTCCCGCCGGGTTGATGACGATATCAAGATCGGGTCCAACATG

GTCATATTGCCGACGATGGACCTGAGGTATATTACAGCCACTTATGATGTTTCCAGGAGCGAGCATGCAATCGTGTACTA

TATCTATGACACGAGTCGCTCATCATCTTACTTCTACCCAGTTCGACTGAATTTCAAAGGCAATCCTCTCTCTTTGAGGA

TAGAGTGTTTCCCCTGGCGTCATAAGGTGTGGTGCTACCATGATTGTCTTATATACAACACCATAACAGGTGAAGAGGTC

CATACGAGAGGGCTGACCGGCATAGAGGTAACATGCAATCCAGTCTGAGCAGAGCTACGACCGTCACTCAGGCAGTCCCT

TGAGTCGCCACCGAGTCCAAGCAGCACAGCCTGGGACAATCAACGGCACAGCCCAGCCAACAATGTTATAAAAAACTTAG

GAGCCAGGGTTGTAGGGGCCATGGACTCTCTATCAGTCAATCAGGTCTTGTACCCTGAGGTTCATCTAGACAGCCCTATT

GTCACAAACAAACTAGTTGCCATCCTTGAGTACTCGGGTATCGACCATAACTATGTTCTTGAAGACCAGACTCTTGTCAA

GAATATTAGGTATAGGCTGGGGTGCGGTTTTTCAAATCAAATGATCATCAATAACAGGGGGGTGGGTGAAACAGTCAATT

CTAAACTTAAAAGTTACCCCCATAATCGTCATATCATATACCCGGATTGCAATAAGGAGTTGTTTTGTATCAAGGATAGC

TGCATATCTAAGAAGCTCTCGGAGCTATTCAAGAAGGGTAATTCCTTGTACTCTAAGATAAGTCACCAGGTACTGGATTG

TCTTAAGAGAGTCAATGGGAAATTAGGTCTGGGCACAGATCTTACCCATGGTCTGAAGGAGGGTATCCTTGACCTAGGGT

TGCACATGCATAGCTCTCAATGGTTCGAGACTTTCCTGTTCTGGTTCACTATCAAGACAGAGATGAGATCAATGATCAAA

GAACAGTCCCATATATGCCACAAGAGGAGGTATAACCCGACTTTTGTGTCGGGGGATGCATTCGAGGTGCTCGTATCGCG

AGACCTCGTTGTGATAATTGATAAGAATACCCAGTATGTCTTCTACTTAACATTTGAGCTGGTCCTTATGTATTGTGATG

TCATAGAGGGTAGACTTATGACGGAGACAGCCATGGCCATAGACCAGAGATATTCAGAACTTCTAAGCCGGGTCAGATAC

TTGTGGGATCTTATTGATGGGTTCTTCCCAACACTAGGCAACACCACATACCAAATTGTTGCTCTGCTTGAACCATTATC

GTTGGCTTATCTTCAACTTCAGGATGTCACTCTGGAGTTAAGAGGTGCTTTTTTAGACCATTGCTTCAAAGAACTCTATG

AGATACTGGAACATTGCGGCATTGACACAGCAGGCACTTACAATTCCATCACTGAAGGATTGGATTACGTATTCATCACC

CACGATATACATCTAACTGGGGAGATTTTTTCATTTTTCCGGAGTTTCGGACACCCCCGTCTAGAAGCAGTCACCGCTGC

GGAAAATGTCAGAAAACATATGAACCAACCGAAGGTAATCAGTTATGAGACTATGATGAAAGGCCATGCGGTATTTTGCG

GGATAATCATAAATGGTTTTCGGGACCGACATGGCGGCAGCTGGCCCCCTGTTGCACTGCCAGAACATGCCTCTGCTGCG

ATCCGGAATGCGCAGGCATCAGGTGAGGGACTGACCCATGACCTGTGTATAGACAACTGGAAATCCTTTGTTGGATTCAA

ATTTGGCTGCTTTATGCCACTCAGCCTAGATAGTGATTTGACCATGTATCTCAAAGACAAGGCATTGGCTGCACTGAAGA

ATGAGTGGGATTCAGTTTACCCGAAAGAATACCTCCGTTATAACCCACCTAGAGGGACAGAGTCAAGGCGATTGGTAGAG

GTATTCCTGAATGACTCCAGCTTTGATCCTTATAACATGATAATGTACGTGGTGAATGGCTCCTACCTTAAAGACCCTGA

GTTTAACCTCTCATACAGCCTAAAAGAGAAGGAGATAAAGGAGACAGGGCGGTTGTTTGCCAAAATGACCTATAAGATGC

GGGCCTGTCAGGTAATCGCTGAAAATCTGATATCGAATGGTGTTGGGAAGTATTTCCGAGACAATGGGATGGCAAAGGAC

GAGCATGACCTAACAAAAGCCCTTCACACCCTGGCAGTCTCAGGTGTTCCCAAAAACAACAAAGATAACCACCGAGGTGG

GCCTCCCAGAAGGACAACAAACCGAGGGGTGAGATCAAGCCAAGGCACCAAAACACAAGATAGAGACAAGGTTCAAGGGG

GACCTATGTACAACTATTTGCGATGCCAACCGATCAGCCCTGATCAGGGTGAGTCATACGAGACTGTTAGTGCATTCATC

ACCGCTGACCTTAAGAAGTATTGCCTGAATTGGAGATACGAGACAATCAGCATATTTGCACAGAGGCTGAATGAAATATA

TGGACTACCATCCTTCTTTCAATGGTTACACAGGATATTGGAAAAATCCGTACTCTACGTCAGTGACCCACATTGTCCTC

CCGATCTAGATAATCATATCCCTCTGGACAGTGTCCCCAATGCCCAAATATTCATCAAGTACCCAATGGGCGGAATAGAA

GGTTATTGCCAAAAACTATGGACAATCAGCACTATACCATACTTGTATCTGGCAGCCTATGAGAGCGGAGTAAGAATCGC

CTCACTGGTGCAGGGTGACAATCAGACAATCGCAGTGACAAAAAGAGTTCCAAGTTCTTGGCCTTATTCACTAAAAAAGA

GGGAGGCCTCCAAGGCCGCTCAAAATTACTTCGTAGTCTTAAGGCAAAGATTGCACGATGTAGGTCATCACTTAAAGGCT

AATGAGACCATAGTATCTTCTCACTTTTTTGTATATTCCAAAGGGATTTATTATGACGGCCTGCTAGTCTCACAATCACT

AAAGAGCATCGCCAGATGTGTCTTCTGGTCCGAGACTATAGTGGATGAAACCAGAGCGGCGTGCAGTAATATTGCAACAA

CTGTCGCTAAGAGTATAGAGAGGGGTTATGATAGGTACCTTGCATATTCTTTGAATATCCTCAAGATTTTCCAACAGATC

CTTATATCCCTTAACTTCACTATTAACACAACAATGACTCAGGATGTCGTGGCACCGATCATCGAGAACGGTGATTTGCT

GATAAGGATGGCACTCTTGCCAGCACCCATCGGGGGTCTCAATTATCTTAACATGAGCAGGTTATTTGTGAGAAATATCG

GTGACCCGGTCACCTCCTCTATAGCCGACCTGAAGAGGATGATAGATGCCGGGCTAATGCCAGAAGAAACATTGCATCAA

GTGATGACCCAGACCCCGGGAGAATCATCCTACCTAGATTGGGCAAGTGACCCTTATTCTGCCAACCTACCCTGCGTACA

GAGTATCACTCGCCTTCTCAAAAACATCACTGCACGATATATTTTAATCAGCAGCCCAAACCCGATGCTGAAAGGGTTAT

TTCATGAGGGGAGTAGAGATGAAGACGAGGAGCTTGCAAGTTTCCTAATGGATCGGCATATAATTGTTCCAAGAGCTGCA

CATGAAATCTTAGACCATAGTATAACCGGAGCAAGAGAAGCTATAGCCGGGATGTTGGACACCACCAAGGGTCTGATTAG

AACAAGTATGAAACGGGGCGGCCTCACCCCTCGAGTCTTAGCCCGCCTTTCCAATTATGATTATGAACAATTCCGATCCG

GGATAACACTATTAACAAAGAAAGGGCAGTGTTATCTCATTGACAAGGACTCGTGCTCGGTGCAGCTCGCTATAGCTCTG

AGGGGCCACATGTGGGCTAAGTTGGCTCGCGGGAGACCAATCTATGGGTTGGAGGTGCCTGATGTATTAGAATCGATGAA

CGGCTACCTTATCAAACGCCACGAGTCGTGTGCTATCTGTGAAACGGGCTCAAGTCACTACGGATGGTTTTTCGTCCCTG

CAGGGTGCCAGCTTGACGATGTATCTAGAGAGACTTCGGCTCTTCGTGTACCTTATGTCGGATCAACAACTGAGGAAAGG

ACAGACATGAAACTTGCATTTGTTAGGTCTCCAAGTAGATCCCTCAAATCAGCAGTTAGAATTGCAACAGTTTACTCATG

GGCTTACGGGGATGATGAGAAATCATGGAGTGAAGCCTGGATGCTAGCTAGGCAGAGAGCCAATATCACCTTAGATGAAT

TGAGAATGATCACTCCAGTCTCCACATCCACCAACTTAGCCCACCGGTTGAGGGATCGGAGTACCCAGGTGAAATACTCG

GGGACATCCCTTGTGAGGGTTGCAAGATACACAACCATCTCCAATGATAACTTGTCGTTTGTGATATCTGAGAAAAAAGT

GGATACCAACTTCATTTATCAGCAAGGGATGTTGCTTGGTCTTGGGATCCTTGAGAACCTCTTTAGGCTAGAGGCCACCA

CCGGAGTATCCAACACAGTGCTACACCTGCACGTGGAAACAGAATGTTGTGTTGTACCTATGGTGGATCACCCAAGGATA

CCGAGTCTCCGTAATATAAAAGTTACGGATGAGCTATGCACAAACCCTTTGATCTACGACAGGTCCCCCATCATAGAGCA

CGATGCAACCCGGCTATATTCACAGAGCCACAGGAGACATTTGGTGGAGTTTGTTACCTGGTCAACAAGCCAGCTTTACC

ATATACTGGCTAAGTCTACAGCAATGTCCATGATTGAATTGATCACGAGATTCGAGAAAGATCACATGAATGAAATAGCC

GCCCTGATTGGCGATGACGACATCAACAGCTTCATCACAGAATTTTTGCTAGTGGAGCCTAGATTATTTATAGTTTACCT

TGGTCAGTGTGCTGCCATCAATTGGGCCTTTGATATACATTATCATCGGCCCTCGGGCAAGTACCAGATGGGGGAACTTC

TCTACTCTCTGCTCTCTCGGATGAGCAAAGGAGTATATAAGATCTTTACTAATGCTTTGAGCCACCCCAAAGTTTACAAG

AAATTTTGGCGAAGTGGCATCATTGAGCCGATTCATGGCCCATCCTTGGATACACAGAATTTACATGTCACTGTCTGTGA

CATGATATATGGGTCCTATGTCACCTATTTGGATCTTTTGCTGAATGATGAGCTAGATGATTACCCGTATTTGCTCTGCG

AGAGTGATGAGGACGTGGTCACAGACAGGTTCGACAACATTCAAGCCAAACATCTCTGTGTATTGGCCGATGTATATTGC

AGCTCCAAGAGGTGTCCTTCAATAATCGGGATGTCTCCTATAGAAAAATGTACCATTCTCACACATTACATCAAGGGAGA

ATCAGTACAATCCCCGTCTGGGACCTCATGGAACACTGATCCCCTTGTAGTAGATCATTACTCATGCTCTCTGACCTACC

TTCGCCGCGGTTCCATCAAACAAATCAGGTTGAGGGTGGATCCTGGGTTTGTATTCGAGGCGTTGACAGACGTCGACTTT

AAACAACCTCGCAAAGCTAAGTTAGATATATCGGTCGTTGGGTTGACTGATTTCTCTCCCCCTTACGACAACGTCGGTGA

TTTTTTAGGGACTATCAACACATTGAGGCACGATCTGCCTGTCACCGGAACCGGGGTCTCGAACTATGAAGTCCACGCTT

ATCGTAGAATTGGCCTGAATTCATCAGCATGTTATAAAGCCGTAGAGATCTCCACGTTAATCAAGCCATCCTTAGAAGTC

GGAGAGCATGGATTGTTCTTAGGAGAAGGTTCCGGTTCAATGCTGGCTGCGTACAAAGAAGTTCTTAAATTAGCAAATTG

TTACTACAACAGCGGAGTAACAGCGGAGGGCAGAGCCGGACAGAGGGAAATATCTCCCTATCCTTCAGAGATGAGCCTAG

TAGAGAATCAAATGGGGATAGAGCGGAGTGTTAAAGTGCTGTTCAACGGGAAACCTGAAGTAACTTGGGTGGGGACCACA

GATTGCTACAAGTACATAATCAGTAACATCCAAACCTCTAGTCTGGGTTTCATACATTCAGATATTGAGACGCTCCCAAC

CAAAGATGCTGTTGAGAAATTGGAGGAATTTGCCTCTATCCTATCCTTATCCTTGATTTTGGGGAAAATCGGCTCTATTA

CAGTTGTCAAAATTATGCCCATTAGCGGGGATTTTACCCAAGGCTTCATAGCCTATGCCATCCAATATTTTAGGGAGAGC

CTGCTTGCCTATCCGAGATATAGTAACTTCATCTCGACTGAGTGTTACCTTATTATGATAGGATTAAAGGCCAATCGATT

GATAAACCCAGAAGCCATTAAGCAAAGCATAATCAGAGCGGGGGTCAGGACTGCACCAGGACTTGTGAGCCATGTACTAT

CAGAGAAACAAAAAGGTTGTATTCAATCTTTCTTGGGTGATCCTTATATCCAAGGAGACTTCAATAAGCACCTTAAATCT

CTAACCTCTATTGAGAAAATCCTGGTAAATTGTGGTCTCTCGATCAATGGCATAAAAATCTGTAGAGATCTAATCCACCA

TGATATCGCCTCCGGTCCAGACGGCCTTATGAGCTCTACAATTATTTTATATAGGGAATTGGCTCATTTCAAAGATAATA

TAAGGAGTCAGCACGGAATGTTCCACCCCTACCCGGTACTGGCCAACAGCAGGCAACGTGAATTAATCCTTCGAATAGCC

AAGAAATTCTGGGGGTATGTCTTGCTATATTCTGATGACCCGGCACTAATCAGACAAACAATCAAGAACTTGAAACGGAA

TCATCTAACCTTTGACTTACATAGTAACCCATTTATTAAAGGGCTATCCAAGGCTGAGAAACTGCTGGTGCGGACGAGTT

CACTAAGAAGAGAATGGTTGTTCACTCTCGAAACGAAAGAAGTAAAAGAGTGGTTCAAATTGGTAGGTTACAGTGCACTC

ATCAGAGGCTAGTCAGCTATGCATCTGCCCCCTTCTCCTCCGCCATGAGACCCCACTGGCGATCCAGAAGATTAAAGAAA

ACTACATATTGGATAAGTATCTATTCCCAGCTTTGTCTGGT

>KM091959.1/goat/China/2013-11-30

ACCAAACAAAGTTGGGTAAGGATCGATCTTATAATAACTATGGACTGGCAAACTTAGGAGTAAAGATCCTACTGTCGGGG

GGAGGAGGAGGAGCAAGATCTTTGATTATGGCGACTCTCCTTAAAAGCTTAGCATTGTTCAAAAGGAACAAAGACAAAGC

GCCGACAGCATCAGGTTCAGGAGGGGCCATCCGGGGGATTAAGAATGTTATCATAGTCCCGATTCCCGGAGACTCGTCCA

TCACTACCCGTTCAAGACTGCTCGACAGGCTTGTCAGATTGGCCGGAGATCCTGACATCAACGGGTCAAAGCTGACCGGG

GTGATGATCAGCATGTTATCCTTGTTCGTAGAGTCACCCGGGCAATTGATACAGCGAATCACAGATGATCCGGATGTTAG

TATCCGCCTTGTTGAGGTAGTTCAAAGTACTAGGTCCCAGTCCGGGTTGACCTTTGCATCACGTGGTGCTGATTTAGACA

ACGAGGCAGACATGTATTTTTCAACTGAGGGGCCCTCGAGTGGAGGTAAGAAAAGGATCAACTGGTTTGAGAACAGAGAA

ATAATAGACATAGAGGTGCAGGATCCAGAAGAGTTCAATATGTTGTTAGCCTCCATACTAGCACAAGTCTGGATCCTTCT

GGCCAAGGCTGTTACGGCACCAGATACGGCAGCTGACTCAGAACTGAGAAGGTGGGTTAAATACACACAACAAAGGAGAG

TGATTGGGGAATTTCGCCTTGACAAGGGGTGGCTGGACGCAGTCCGCAACAGAATTGCAGAAGATCTATCACTCCGGCGG

TTCATGGTATCTCTCATACTTGACATCAAGAGGACCCCAGGCAACAAGCCAAGGATTGCAGAAATGATCTGCGACATTGA

CAACTATATTGTCGAAGCAGGACTCGCCAGTTTCATCCTTACTATCAAATTTGGTATCGAAACCATGTATCCTGCATTAG

GGCTCCACGAGTTTGCCGGAGAATTGTCCACTATTGAGTCCTTGATGAACTTGTATCAACAGCTAGGAGAGGTTGCACCC

TACATGGTAATTCTAGAGAATTCAATTCAGAACAAGTTTAGTGCAGGAGCTTATCCCCTCCTCTGGAGCTATGCGATGGG

TGTCGGAGTCGAGCTGGAGAACTCAATGGGGGGCTTGAACTTTGGCAGGTCATATTTTGACCCTGCCTATTTTCGTCTCG

GACAGGAGATGGTCAGAAGATCTGCAGGAAAGGTCAGCTCTGTGATTGCGGCTGAGCTCGGCATCACAGCAGAGGAAGCT

AAACTAGTCTCGGAAATCGCCTCGCAGGCTGGGGACGAAAGAACCGCCAGAGGGACTGGGCCTCGACAGGCGCAGGTTTC

CTTCCTTCAGCACCAAACAGGAGGGGGAGAGTCGTCCGCACCAGCGACCAGAGAAGGGGTCAAAGCTGCGATCCCAAACG

GAGCTGAAGAAAGGGACAGAAAGCAAACACGCCCAGGAAGGCCCAGAGGAGAGACCTCCGGCCAACTGCTCCTGGACATC

ATGCCAGAGGATGAGATCTTGCGAGAGTCTGGTCAAAACCCTCGTGAGGCTCAAAGATCGGCCGAGGCACTCTTCAGGCT

GCAGGCCATGGCCAAGATTCTGGAGGACCAGGAGGAGGGAGAAGACAACAGTCAGGTCTACAACGACAAGGATCTCCTCG

GCTAAGCAGACGCACCCTCTGTCGAAATCAGTGACAAGACATCTCCTACCAGTATTATAAAAAACTTAGGACCCAGGTCC

AAGCAAGCACACATCGACACTCCAACCAGTCGAGCGGAGACCACCGATGGCAGAAGAACAAGCATACCATGTCAACAAGG

GGCTGGAATGTATCAAGTCTCTCAAAGCCTCACCCCCGGATCTATCCACCATCAGAGATACCATCGAGAGCTGGAGAGAG

GGGCTTAGCCCCTCGGGCCGTGCAACACCGAACCCTGATACGTCCGAGGGAGATCATCAGAATATCAACCAATCATGCTC

ACCAGCAATCGGACCAAACAAAGTCTACTTGTCTCCTGAAGATAATCTCGGATTTAGAGAGATCACTGGCAACGACTGTG

AGGCTGGGCTCGGAGGGGTCCAGAGAGAAGGATCCAACTCTCAAGTACAGCGTTACCATGTTTATAGTCACGGGGGTGAA

GAGATTGAAGGACTCGAGGATGCTGACTCTCTCGTGGTTCAAGCAGATCCTCCAGTTGCTAACGTCTTCAATGGAGGAGA

GGATGGATCTGACGACAGCGATGTGGACTCTGGCCCAGATGATCCCGGCAGAGATACTCTATATGACCGGGGATCTGTTG

CCGGCAATGGTATCGCTAGGTCCACAGATGTCGAAAAACTAGAAGGTGCTGATATTCAAGAAGTTCTTAACTCCCAGAAA

GGCAAAGGAGGAAGATTCCAGGGCGGGAAAACCTTGCGAGTCCCGGAAATACCCGATGTCAAGCACTCCAGACCATCAGC

CCAATCAATTAAAAAGGGCACAGACGGGAACTCAGTCTCATCTGGAACGGTGATAGAGTGTTTATCGATAAGTGGTGCAA

CCCAAACTGTGCCAGAGTCAAGATGGGAGTCATCAGAGCAAAATGCGTCTGTGGGGAGTGTCCTCAAGTCTGCGAGGAGT

GCAAAGACGATCCAGGGGTCGACACAAGAATCTGGTACCATAGCATCACTGACTCAGCCTAAAGAGAATGACTCCGAGTA

TGAGTATGAGGATGACCTATTTACAGAGATTCAGGACATCCGTGCAAGCATTGCCAAGATCCATGATGACAATAAAACTA

TCCTCTCAAAGCTTGATTCTATACTGTTATTGAAAGGGGAAGTCGACACTATCAAGAAACAAATCAGCAAGCAGAATATA

AGTATATCCACCATTGAGGGCCATCTCTCCAGTATAATGATAGCCATCCCTGGCTTTGGGAAGGACATCAAGGACCCAAC

ATCCGAAGTCGAGTTGAACCCAGATCTAAGACCTATAATAAGCCGTGACTCTGGCAGAGCTCTCGCGGAGGTCCTCAAGA

AACCCGCTGTGGATAGGTCTCAGAAAATTGGAACCAAAGCCAACTCCAGCTCAAAGGGTCAGCTTCTTAAGGATCTCCAG

CTAAAACCTGTCGACAAGCAGGCAAGCTCTGCAATCGGGTTTGTCCCGTCCGACCATGAATCATCCAGAAATGTCATCCG

CTCCATAATCAAGTCGAGCAAGCTAAATATTGATCACAAGGACTATCTTCTAGATTTACTAAATGATGTGAAAGGCTCCA

AGGATCTTAAGGAATTTCACAAGATGCTAACAGCAATTCTCGCCAAGCACCCGTAACACATCCTCCAGTCATCATCTCAT

ACTCGACTAAAAACATCCTTTCAATCAGGCTATTACAAAAAACTTAGGAGCAAGGGCAACTGAGCTTCGCAGACAGGATG

ACCGAGATCTACGACTTCGATAAATCAGCATGGGATGTCAAAGGGTCAATTGCCCGCATAGAACCCACCACCTACCACGA

CGGCCGACTGATACCCCAGGTGAGGGTCATCGATCCTGGTCTGGGAGACAGAAAAGATGAGTGCTTTATGTACCTGTTTC

TCCTAGGAGTGATTGAGGATAACGACCCCCTGTCTCCCCCCGTCGGAAGAACCTTTGGCTCTCTACCTCTAGGGGTTGGT

AGGTCAACTGCCAAGCCAGAAGAACTACTAAGGGAGGCCACAGAATTAGATATAGTGGTGAGGCGCACTGCAGGAGTAAA

TGAGAAACTGGTATTTTACAACAACACTCCGCTGTCCTTGTTAACGCCCTGGAAGAAAGTTCTGACAACCGGAAGTGTGT

TTAGCGCTAACCAGGTTTGCAATGCAGTCAACCTAGTCCCGCTTGATACTCCCCAGAGATTCAGGGTTGTGTACATGAGC

ATAACTAGATTATCAGACAATGGTTACTATAGTGTGCCCAGAAGAATGCTGGAGTTCCGCTCGGCCAATGCAGTAGCCTT

CAACATCTTGGTTACACTAAGAATTGAAAATGGCACAAACCCTAGAAGATACATAGTCGGCTCATGGGAGAATTCAGAGG

TCACATTTATGGTACACGTGGGCAACTTTAGGAGAAAGAAGAACGAAGTATACTCTGCTGATTACTGCAAAATGAAGATT

GAAAAGATGGGTTTAGTTTTTGCCCTGGGTGGAATAGGCGGAACAAGTCTCCATATTAGAAGCACCGGGAAAATGAGCAA

AACCCTCCATGCACAGCTGGGGTTCAAGAAAATCTTATGTTACCCCCTAATGGATGTCAATGAGGATCTTAACCGATATC

TCTGGCGGGCAGAGTGCCGAATAGTCAAAATCCAAGCTGTCTTACAGCCATCAGTACCCCAAGAGTTCCGTGTCTACGAT

GATGTCATCATCAACGATGATCAAGGCTTGTTCAAGATCTTGTAGTTCATTTGCAAAGGAACGAACCCACCCCCAGGAAG

GGGCCCCACCCTCCCCCG-CCAAAAAACACACTCCAAGGCTCCCCGGCACAGCGCCAACACAGCTACTCCCGCGATGGAA

CGAGGTCGGCC-CCCGAACAAATCCGGTCCAGGGGGGGGGGCCCCCCCCCTGACCAACCGGGCGCCGACCGAGCAGGCCG

GCGAGAAGCCCCCAAGAGGCCCCGACGAAGACCGAGCACCCCATCCCCCC-AAAAACGTCCCCC-AACACGCCCGAGGCT

AGCCACCGGACACCCCCAGACCCGCCACCGGCGAGACAGAACCCCAAGGGGGGACCCTCCAAACCAAAGGAATCACCGGG

GTAGGCTCCGCCGAACCAGCACAGACCCGACCCAAGCAAGGCGGACCCCCAACACGAAACTGCCCAGCACACCCTACTCA

TCATCAAACAAAACTTAGGGGCCAAGTCCATACGAGAGGCCCTAGTCCGTCAGGCGGAGGGACGCAGAAAGGAAGGAGAC

ACGACCGTCCCAGAAGGAGGACACCCAACCACCGAAACGCCCGCCCCCAGGGAAGGAG--CCA-CCCCCACGCAGCGAAG

GAGCGGGTGGGAGACCTCGGGGAGGGCCCGACCGGGACCCGGGGGCCGGCGGCGCCGGCGGTGAGCAGGCCGGGCGCATA

CCTGACCCCACCGGAGGATCCGAGGAGGGCCCGACCCCGCGTGGACCAAACGGCAGGGAGCCCAACCCCACCAAAAAAAG

AGGCGAGGCCCCCCCTCCCTCCCCCCCCGCACACAACCCGCAGACAGACCGAGGGAGGGCCATCCCCCAGCGAGACAGGC

GGCTCCCTCGCAGCCGAGCCGCAGCTCCCCAGAATAGGACACCCCCGCACACTCCGAACACCCCCTCGCCCCCCGCCACC

GGGACAGGGGCTCCACCACCACAGACCGGAGCCACCGACAGCCAGCCAGAGGTCCCCTCCAGTCCCGCCCGGCGCCCCCC

CGAAGAGGCCCCGGCAAGGCCCCCCAGCCCGACGTCCCCCGCACGCACCGAAAGAAGGACCCGCCCGAGAACGTGCGGCA

CATCCATGCGCAAACATCATGACGCGGGTCGCAATTTTGACATTTCTGTTTCTTTTCCCAAATGTTGTTGCGTGTCAGAT

TCACTGGGGCAATCTATCCAAGATCGGGATTGTAGGAACAGGGAGTGCCAGCTACAAGGTGATGACTAGGCCAAGCCACC

AGACTCTGGTTATAAAGTTAATGCCAAATATAACGGCCATCGACAATTGTACAAAGTCAGAGATTGCAGAGTACAAGAGA

TTGCTGATCACAGTGTTAAAGCCTGTAGAGGATGCTCTGTCGGTGATAACCAAGAATGTAAGACCAATTCAAACTCTAAC

ACCTGGGCGTAGAACCCGCCGTTTTGCTGGAGCTGTTCTGGCCGGGGTAGCACTTGGAGTTGCGACAGCCGCTCAGATAA

CTGCAGGAGTCGCCCTTCATCAATCATTGATGAACTCCCAAGCAATTGAGAGTTTAAAAACCAGTCTTGAGATGTCGAAT

CAGGCAATAGAAGAAATCAGACTTGCAAATAAGGAGACCATACTGGCAGTACAGGGCGTCCAGGATTATATCAACAATGA

GCTCGTCCCTTCTGTTCATAGAATGTCATGCGAGCTGGTAGGTCACAAGCTCGGCCTCAAGCTCCTTAGGTACTACACCG

AGATCCTGTCCATATTCGGGCCCAGTCTTCGAGACCCGATATCTGCCGAAATATCAATCCAGGCACTTAGTTATGCATTA

GGCGGAGACATTAATAAAATCCTGGACAAGCTTGGGTATAGCGGTGGGGATTTCCTTGCCATCCTAGAAAGCAAGGGAAT

AAAGGCCCGGGTTACATATGTGGACACAAGAGATTACTTTATAATCCTTAGCATCGCCTACCCAACCTTATCTGAGATCA

AGGGAGTGATAGTTCACAAGATAGAAGCTATAACATACAACATTGGGGCACAGGAGTGGTATACTACTATCCCTAAATAT

GTAGCCACTCAGGGGTATCTGATATCGAACTTTGATGAGACGTCATGCGTATTCACTCCAGAGGGGACAGTTTGCAGCCA

GAATGCGTTGTACCCAATGAGCCCATTGCTTCAGGAATGTTTCAGGGGGTCAACAAAATCGTGCGCCAGAACCTTAGTTT

CAGGGACCATAAGTAATAGATTTATCCTATCAAAAGGGAACCTGATTGCAAATTGTGCGTCAGTTTTGTGCAAATGTTAC

ACAACAGAGACAGTTATCAGCCAAGATCCTGACAAACTACTAACTGTTGTAGCATCCGACAAGTGTCCTGTAGTTGAGGT

GGATGGAGTGACAATACAGGTCGGCAGTCGAGAGTATCCGGATTCTGTATACTTACACAAAATAGACTTAGGTCCAGCCA

TCTCCCTAGAAAAACTGGATGTAGGCACCAATTTAGGCAATGCAGTCACAAGACTGGAGAATGCAAAGGAGCTCCTAGAT

GCATCAGACCAAATACTGAAGACTGTTAAAGGGGTACCTTTCAGTGGGAATATGTACATAGCACTGGCAGCTTGCATAGG

AGTATCCCTGGGCCTTGTCACATTAATATGCTGCTGTAAGGGGAGGTGTAAGAACAAGGAAATCCCTATCTCCAAAATCA

ACCCAGGGCTCAAACCCGACCTGACCGGGACCTCAAAGTCGTACGTAAGATCACTGTAGTCAGAATCACCTGGATCATCT

GGCATCACACACATACATGCACGACACAGGCAGTCCGAGGACGCAAGAAACCCAGCCTCCGGTCACCCACCCGACCCCAC

TCCACGCTCCACCACACATTAGTCATCAAACAAAACTTAGGACGAAAGGTCAATCACCATGTCCGCACAAAGGGAGAGGA

TCAATGCCTTCTACAAAGACAATCCTCACAATAAGAACCATAGGGTGATCCTGGATAGAGAACGCTTGGTCATTGAAAGA

CCCTACATCTTGCTTGGAGTCCTGCTGGTAATGTTCCTGAGTCTAATCGGACTGCTGGCCATTGCAGGGATCAGGCTTCA

CCGGGCCACCGTTGGAACTTCAGAGATCCAGAGTCGGCTGAATACCAATATTAAGTTGACCGAATCTATTGATCACCAGA

CTAAGGATGTCTTAACTCCCCTTTTTAAAATCATTGGCGATGAAGTCGGCATCAGAATTCCACAGAAATTCAGTGATCTT

GTCAAGTTCATCTCCGATAAGATTAAATTCCTCAACCCTGATAGAGAGTATGATTTCAGAGATCTCCGGTGGTGTATGAA

TCCCCCCGAGAGAGTCAAAATTAATTTTGATCAGTTTTGTGAGTACAAGGCTGCGGTTAAGTCAATTGAACATATATTTG

AGTCACCACTCAACAAGTCAAAAAAGCTGCAATCTTTGACTCTCGGGCCCGGAACAGGCTGTCTAGGCAGGACAGTAACA

AGAGCCCATTTCTCAGAACTTACAATGACCTTAATGGACCTGGATCTAGAGATGAAGCACAACGTGTCCTCAGTGTTTAC

CGTAGTTGAAGAGGGATTATTCGGAAGAACATATACCGTCTGGAGATCCGATGCCAGGGATCCGAGCACCGATCTAGGTA

TCGGCCATTTTTTAAGAGTCTTCGAGATTGGACTGGTAAGAGATCTTGGGCTGGGTCCCCCTGTTTTTCATATGACCAAC

TATCTCACGGTGAACATGAGTGATGACTATCGGAGATGTCTTTTAGCGGTAGGGGAGTTGAAGTTGACAGCCCTATGCAC

CTCATCTGAGACTGTGACACTGAGTGAGAGAGGAGTTCCAAGGAGGGAACCTCTTGTGGTTGTGATACTTAATCTAGCTG

GACCCACTCTAGGGGGAGAGCTATACAGTGTCTTGCCTACCTCTGATCTCATGGTGGAGAAACTCTATTTGTCTTCACAT

AGAGGGATCATCAAAGATGATGAGGCCAATTGGGTAGTGCCGTCTACCGATGTTCGTGATCTTCAGAACAAAGGTGAATG

TCTGGTGGAAGCATGCAAGACTCGACCTCCTTCATTTTGCAATGGCACAGGATCAGGCCCGTGGTCAGAAGGGAGAATCC

CTGCCTACGGGGTGATCAGGGTCAGTCTTGACTTAGCGAGTGACCCGGATGTAGTTATCACTTCAGTGTTTGGCCCACTG

ATACCTCACCCATCCGGCATGGATCTTTACAACAACCCGTTTTCAAAAGCTGTATGGTTGGCTGTACCACCTTATGAGCA

GTCATTTCTAGGAATGATAAATACAATTGGATTCCCTAACAGAGCAGAGGTTATGCCGCACATTTTGACCACAGAGATCA

GAGGCCCTCGGGGTCGTTGCCATGTTCCCATAGAATTGTCCCGCCGGGTTGATGACGATATCAAGATCGGGTCCAACATG

GTCATATTGCCGACGATGGACCTGAGGTATATTACAGCCACTTATGATGTTTCCAGGAGCGAGCATGCAATCGTGTACTA

TATCTATGACACGAGTCGCTCATCATCTTACTTCTACCCAGTTCGACTGAATTTCAAAGGCAATCCTCTCTCTTTGAGGA

TAGAGTGTTTCCCCTGGCGTCATAAGGTGTGGTGCTACCATGATTGTCTTATATACAACACCATAACAGGTGAAGAGGTC

CATACGAGAGGGCTGACCGGCATAGAGGTAACATGCAATCCAGTCTGAGCAGAGCTACGACCGTCACTCAGGCAGTCCCT

TGAGTCGCCACCGAGTCCAAGCAGCACAGCCTGGGACAATCAACGGCACAGCCCAGCCAACAATGTTATAAAAAACTTAG

GAGCCAGGGTTGTAGGGGCCATGGACTCTCTATCAGTCAATCAGGTCTTGTACCCTGAGGTTCATCTAGACAGCCCTATT

GTCACAAACAAACTAGTTGCCATCCTTGAGTACTCGGGTATCGACCATAACTATGTTCTTGAAGACCAGACTCTTGTCAA

GAATATTAGGTATAGGCTGGGGTGCGGTTTTTCAAATCAAATGATCATCAATAACAGGGGGGTGGGTGAAACAGTCAATT

CTAAACTTAAAAGTTACCCCCATAATCGTCATATCATATACCCGGATTGCAATAAGGAGTTGTTTTGTATCAAGGATAGC

TGCATATCTAAGAAGCTCTCGGAGCTATTCAAGAAGGGTAATTCCTTGTACTCTAAGATAAGTCACCAGGTACTGGATTG

TCTTAAGAGAGTCAATGGGAAATTAGGTCTGGGCACAGATCTTACCCATGGTCTGAAGGAGGGTATCCTTGACCTAGGGT

TGCACATGCATAGCTCTCAATGGTTCGAGACTTTCCTGTTCTGGTTCACTATCAAGACAGAGATGAGATCAATGATCAAA

GAACAGTCCCATATATGCCACAAGAGGAGGTATAACCCGACTTTTGTGTCGGGGGATGCATTCGAGGTGCTCGTATCGCG

AGACCTCGTTGTGATAATTGATAAGAATACCCAGTATGTCTTCTACTTAACATTTGAGCTGGTCCTTATGTATTGTGATG

TCATAGAGGGTAGACTTATGACGGAGACAGCCATGGCCATAGACCAGAGATATTCAGAACTTCTAAGCCGGGTCAGATAC

TTGTGGGATCTTATTGATGGGTTCTTCCCAACACTAGGCAACACCACATACCAAATTGTTGCTCTGCTTGAACCATTATC

GTTGGCTTATCTTCAACTTCAGGATGTCACTCTGGAGTTAAGAGGTGCTTTTTTAGACCATTGCTTCAAAGAACTCTATG

AGATACTGGAACATTGCGGCATTGACACAGCAGGCACTTACAATTCCATCACTGAAGGATTGGATTACGTATTCATCACC

CACGATATACATCTAACTGGGGAGATTTTTTCATTTTTCCGGAGTTTCGGACACCCCCGTCTAGAAGCAGTCACCGCTGC

GGAAAATGTCAGAAAACATATGAACCAACCGAAGGTAATCAGTTATGAGACTATGATGAAAGGCCATGCGGTATTTTGCG

GGATAATCATAAATGGTTTTCGGGACCGACATGGCGGCAGCTGGCCCCCTGTTGCACTGCCAGAACATGCCTCTGCTGCG

ATCCGGAATGCGCAGGCATCAGGTGAGGGACTGACCCATGACCTGTGTATAGACAACTGGAAATCCTTTGTTGGATTCAA

ATTTGGCTGCTTTATGCCACTCAGCCTAGATAGTGATTTGACCATGTATCTCAAAGACAAGGCATTGGCTGCACTGAAGA

ATGAGTGGGATTCAGTTTACCCGAAAGAATACCTCCGTTATAACCCACCTAGAGGGACAGAGTCAAGGCGATTGGTCGAG

GTATTCCTGAATGACTCCAGCTTTGATCCTTATAACATGATAATGTACGTGGTGAATGGCTCCTACCTTAAAGACCCTGA

GTTTAACCTCTCATACAGCCTAAAAGAGAAGGAGATAAAGGAGACAGGGCGGTTGTTTGCCAAAATGACCTATAAGATGC

GGGCCTGTCAGGTAATCGCTGAAAATCTGATATCGAATGGTGTTGGGAAGTATTTCCGAGACAATGGGATGGCAAAGGAC

GAGCATGACCTAACAAAAGCCCTTCACACCCTGGCAGTCTCAGGTGTTCCCAAAAACAACAAAGATAACCACCGAGGTGG

GCCTCCCAGAAGGACAACAAACCGAGGGGTGAGATCAAGCCAAGGCACCAAAACACAAGATAGAGACAAGGTTCAAGGGG

GACCTATGTACAACTATTTGCGATGCCAACCGATCAGCCCTGATCAGGGTGAGTCATACGAGACTGTTAGTGCATTCATC

ACCGCTGACCTTAAGAAGTATTGCCTGAATTGGAGATACGAGACAATCAGCATATTTGCACAGAGGCTGAATGAAATATA

TGGACTACCATCCTTCTTTCAATGGTTACACAGGATATTGGAAAAATCCGTACTCTACGTCAGTGACCCACATTGTCCTC

CCGATCTAGATAATCATATCCCTCTGGACAGTGTCCCTAATGCCCAAATATTCATCAAGTACCCAATGGGCGGAATAGAA

GGTTATTGCCAAAAACTATGGACAATCAGCACTATACCATACTTGTATCTGGCAGCCTATGAGAGCGGAGTAAGAATCGC

CTCACTGGTGCAGGGTGACAATCAGACAATCGCAGTGACAAAAAGAGTTCCAAGTTCTTGGCCTTATTCACTAAAAAAGA

GGGAGGCCTCCAAGGCCGCTCAAAATTACTTCGTAGTCTTAAGGCAAAGATTGCACGATGTAGGTCATCACTTAAAGGCT

AATGAGACCATAGTATCTTCTCACTTTTTTGTATATTCCAAAGGGATTTATTATGACGGCCTGCTAGTCTCACAATCACT

AAAGAGCATCGCCAGATGTGTCTTCTGGTCCGAGACTATAGTGGATGAAACCAGAGCGGCGTGCAGTAATATTGCAACAA

CTGTCGCTAAGAGTATAGAGAGGGGTTATGATAGGTACCTTGCATATTCTTTGAATATCCTCAAGATTTTCCAACAGATC

CTTATATCCCTTAACTTCACTATTAACACAACAATGACTCAGGATGTCGTGGCACCGATCATCGAGAACGGTGATTTGCT

GATAAGGATGGCACTCTTGCCAGCACCCATCGGGGGTCTCAATTATCTTAACATGAGCAGGTTATTTGTGAGAAATATCG

GTGACCCGGTCACCTCCTCTATAGCCGACCTGAAGAGGATGATAGATGCCGGGCTAATGCCAGAAGAAACATTGCATCAA

GTGATGACCCAGACCCCGGGAGAATCATCCTACCTAGATTGGGCAAGTGACCCTTATTCTGCCAACCTACCCTGCGTACA

GAGTATCACTCGCCTTCTCAAAAACATCACTGCACGATATATTTTAATCAGCAGCCCAAACCCGATGCTGAAAGGGTTAT

TTCATGAGGGGAGTAGAGATGAAGACGAGGAGCTTGCAAGTTTCCTAATGGATCGGCATATAATTGTTCCAAGAGCTGCA

CATGAAATCTTAGACCATAGTATAACCGGAGCAAGAGAAGCTATAGCCGGGATGTTGGACACCACCAAGGGTCTGATTAG

AACAAGTATGAAACGGGGCGGCCTCACCCCTCGAGTCTTAGCCCGCCTTTCCAATTATGATTATGAACAATTCCGATCCG

GGATAACACTATTAACAAAGAAAGGGCAGTGTTATCTCATTGACAAGGACTCGTGCTCGGTGCAGCTCGCTATAGCTCTG

AGGGGCCACATGTGGGCTAAGTTGGCTCGCGGGAGACCAATCTATGGGTTGGAGGTGCCTGATGTATTAGAATCGATGAA

CGGCTACCTTATCAAACGCCACGAGTCGTGTGCTATCTGTGAAACGGGCTCAAGTCACTACGGATGGTTTTTCGTCCCTG

CAGGGTGCCAGCTTGACGATGTATCTAGAGAGACTTCGGCTCTTCGTGTACCTTATGTCGGATCAACAACTGAGGAAAGG

ACAGACATGAAACTTGCATTTGTTAGGTCTCCAAGTAGATCCCTCAAATCAGCAGTTAGAATTGCAACAGTTTACTCATG

GGCTTACGGGGATGATGAGAAATCATGGAGTGAAGCCTGGATGCTAGCTAGGCAGAGAGCCAATATCACCTTAGATGAAT

TGAGAATGATCACTCCAGTCTCCACATCCACCAACTTAGCCCACCGGTTGAGGGATCGGAGTACCCAGGTGAAATACTCG

GGGACATCCCTTGTGAGGGTTGCAAGATACACAACCATCTCCAATGATAACTTGTCGTTTGTGATATCTGAGAAAAAAGT

GGATACCAACTTCATTTATCAGCAAGGGATGTTGCTTGGTCTTGGGATCCTTGAGAACCTCTTTAGGCTAGAGGCCACCA

CCGGAGTATCCAACACAGTGCTACACCTGCACGTGGAAACAGAATGTTGTGTTGTACCTATGGTGGATCACCCAAGGATA

CCGAGTCTCCGTAATATAAAAGTTACGGATGAGCTATGCACAAACCCTTTGATCTACGACAGGTCCCCCATCATAGAGCA

CGATGCAACCCGGCTATATTCACAGAGCCACAGGAGACATTTGGTGGAGTTTGTTACCTGGTCAACAAGCCAGCTTTACC

ATATACTGGCTAAGTCTACAGCAATGTCCATGATTGAATTGATCACGAGATTCGAGAAAGATCACATGAATGAAATAGCC

GCCCTGATTGGCGATGACGACATCAACAGCTTCATCACAGAATTTTTGCTAGTGGAGCCTAGATTATTTATAGTTTACCT

TGGTCAGTGTGCTGCCATCAATTGGGCCTTTGATATACATTATCATCGGCCCTCGGGCAAGTACCAGATGGGGGAACTTC

TCTACTCTCTGCTCTCTCGGATGAGCAAAGGAGTATATAAGATCTTTACTAATGCTTTGAGCCACCCCAAAGTTTACAAG

AAATTTTGGCGAAGTGGCATCATTGAGCCGATTCATGGCCCATCCTTGGATACACAGAATTTACATGTCACTGTCTGTGA

CATGATATATGGGTCCTATGTCACCTATTTGGATCTTTTGCTGAATGATGAGCTAGATGATTACCCGTATTTGCTCTGCG

AGAGTGATGAGGACGTGGTCACAGACAGGTTCGACAACATTCAAGCCAAACATCTCTGTGTATTGGCCGATGTATATTGC

AGCTCCAAGAGGTGTCCTTCAATAATCGGGATGTCTCCTATAGAAAAATGTACCATTCTCACACATTACATCAAGGGAGA

ATCAGTACAATCCCCGTCTGGGACCTCATGGAACACTGATCCCCTTGTAGTAGATCATTACTCATGCTCTCTGACCTACC

TTCGCCGCGGTTCCATCAAACAAATCAGGTTGAGGGTGGATCCTGGGTTTGTATTCGAGGCGTTGACAGACGTCGACTTT

AAACAACCTCGCAAAGCTAAGTTAGATATATCGGTCGTTGGGTTGACTGATTTCTCTCCCCCTTACGACAACGTCGGTGA

TTTTTTAGGGACTATCAACACATTGAGGCACGATCTGCCTGTCACCGGAACCGGGGTCTCGAACTATGAAGTCCACGCTT

ATCGTAGAATTGGCCTGAATTCATCAGCATGTTATAAAGCCGTAGAGATCTCCACGTTAATCAAGCCATCCTTAGAAGTC

GGAGAGCATGGATTGTTCTTAGGAGAAGGTTCCGGTTCAATGCTGGCTGCGTACAAAGAAGTTCTTAAATTAGCAAATTG

TTACTACAACAGCGGAGTAACAGCGGAGGGCAGAGCCGGACAGAGGGAAATATCTCCCTATCCTTCAGAGATGAGCCTAG

TAGAGAATCAAATGGGGATAGAGCGGAGTGTTAAAGTGCTGTTCAACGGGAAACCTGAAGTAACTTGGGTGGGGACCACA

GATTGCTACAAGTACATAATCAGTAACATCCAAACCTCTAGTCTGGGTTTCATACATTCAGATATTGAGACGCTCCCAAC

CAAAGATGCTGTTGAGAAATTGGAGGAATTTGCCTCTATCCTATCCTTATCCTTGATTTTGGGGAAAATCGGCTCTATTA

CAGTTGTCAAAATTATGCCCATTAGCGGGGATTTTACCCAAGGCTTCATAGCCTATGCCATCCAATATTTTAGGGAGAGC

CTGCTTGCCTATCCGAGATATAGTAACTTCATCTCGACTGAGTGTTACCTTATTATGATAGGATTAAAGGCCAATCGATT

GATAAACCCAGAAGCCATTAAGCAAAGCATAATCAGAGCGGGGGTCAGGACTGCACCAGGACTTGTGAGCCATGTATTAT

CAGAGAAACAAAAAGGTTGTATTCAATCTTTCTTGGGTGATCCTTATATCCAAGGAGACTTCAATAAGCACCTTAAATCT

CTAACCTCTATTGAGAAAATCCTGGTAAATTGTGGTCTCTCGATCAATGGCATAAAAATCTGTAGAGATCTAATCCACCA

TGATATCGCCTCCGGTCCAGACGGCCTTATGAGCTCTACAATTATTTTATATAGGGAATTGGCTCATTTCAAAGATAATA

TAAGGAGTCAGCACGGAATGTTCCACCCCTACCCGGTACTGGCCAACAGCAGGCAACGTGAATTAATCCTTCGAATAGCC

AAGAAATTCTGGGGGTATGTCTTGCTATATTCTGATGACCCGGCACTAATCAGACAAACAATCAAGAACTTGAAACGGAA

TCATCTAACCTTTGACTTACATAGTAACCCATTTATTAAAGGGCTATCCAAGGCTGAGAAACTGCTGGTGCGGACGAGTT

CACTAAGAAGAGAATGGTTGTTCACTCTCGAAACGAAAGAAGTAAAAGAGTGGTTCAAATTGGTAGGTTACAGTGCACTC

ATCAGAGGCTAGTCAGCTATGCATCTGCCCCCTTCTCCTCCGCCATGAGACCCCACTGGCGATCCAGAAGATTAAAGAAA

ACTACATATTGGATAAGTATCTATTCCCAGCTTTGTCTGGT

>MF443353.1/goat/China/2014-03-30

ACCAAACAAAGTTGGGTAAGGATAGATCTTATAATAACTATGGACTGGCAAACTTAGGAGTAAAGATCCTACTGTCGGGG

GGAGGAGGAGGAGCAAGATCTTTGATTATGGCGACTCTCCTTAAAAGCTTAGCATTGTTCAAAAGGAACAAAGACAAAGC

GCCGACAGCATCAGGTTCAGGAGGGGCCATCCGGGGGATTAAGAATGTTATCATAGTCCCGATTCCCGGAGACTCGTCCA

TCACTACCCGTTCAAGACTGCTCGACAGGCTTGTCAGATTGGCCGGAGATCCTGACATCAACGGGTCAAAGCTGACCGGG

GTGATGATCAGCATGTTATCCTTGTTCGTAGAGTCACCCGGGCAATTGATACAGCGAATCACAGATGATCCGGATGTTAG

TATCCGCCTTGTTGAGGTAGTTCAAAGTACTAGGTCCCAGTCCGGGTTGACCTTTGCATCACGTGGTGCTGATTTAGACA

ACGAGGCAGACATGTATTTTTCAACTGAGGGGCCCTCGAGTGGAGGTAAGAAAAGGATCAACTGGTTTGAGAACAGAGAA

ATAATAGACATAGAGGTGCAGGATCCAGAAGAGTTCAATATGTTGTTAGCCTCCATACTAGCACAAGTCTGGATCCTTCT

GGCCAAGGCTGTTACGGCACCAGATACGGCAGCTGACTCAGAACTGAGAAGGTGGGTTAAATACACACAACAAAGGAGAG

TGATTGGGGAATTTCGCCTTGACAAGGGGTGGCTGGACGCAGTCCGCAACAGAATTGCAGAAGATCTATCACTCCGGCGG

TTCATGGTATCTCTCATACTTGACATCAAGAGGACCCCAGGCAACAAGCCAAGGATTGCAGAAATGATCTGCGACATTGA

CAACTATATTGTCGAAGCAGGACTCGCCAGTTTCATCCTTACTATCAAATTTGGTATCGAAACCATGTATCCTGCATTAG

GGCTCCACGAGTTTGCCGGAGAATTGTCCACTATTGAGTCCTTGATGAACTTGTATCAACAGCTAGGAGAGGTTGCACCC

TACATGGTAATTCTAGAGAATTCAATTCAGAACAAGTTTAGTGCAGGAGCTTATCCCCTCCTCTGGAGCTATGCGATGGG

TGTCGGAGTCGAGCTGGAGAACTCAATGGGGGGCTTGAACTTTGGCAGGTCATATTTTGACCCTGCCTATTTTCGTCTCG

GACAGGAGATGGTCAGAAGATCTGCAGGAAAGGTCAGCTCTGTGATTGCGGCTGAGCTCGGCATCACAGCAGAGGAAGCT

AAACTAGTCTCGGAAATCGCCTCGCAGGCTGGGGACGAAAGAACCGCCAGAGGGACTGGGCCTCGACAGGCGCAGGTTTC

CTTCCTTCAGCACCAAACAGGAGGGGGAGAGTCGTCCGCACCAGCGACCAGAGAAGGGGTCAAAGCTGCGATCCCAAACG

GAGCTGAAGAAAGGGACAGAAAGCAAACACGCCCAGGAAGGCCCAGAGGAGAGACCTCCGGCCAACTGCTCCTGGACATC

ATGCCAGAGGATGAGATCTTGCGAGAGTCTGGTCAAAACCCTCGTGAGGCTCAAAGATCGGCCGAGGCACTCTTCAGGCT

GCAGGCCATGGCCAAGATTCTGGAGGACCAGGAGGAGGGAGAAGACAACAGTCAGGTCTACAACGACAAGGATCTCCTCG

GCTAAGCAGACGCACCCTCTGTCGAAATCAGTGACAAGACATCTCTTACCAGTATTATAAAAAACTTAGGACCCAGGTCC

AAGCAAGCACACATCGACACTCCAACCAGTCGAGCGGAGACCACCGATGGCAGAAGAACAAGCATACCATGTCAACAAGG

GGCTGGAATGTATCAAGTCTCTCAAAGCCTCACCCCCGGATCTATCCACCATCAGAGATACCATCGAGAGCTGGAGAGAG

GGGCTTAGCCCCTCGGGCCGTGCAACACCGAACCCTGATACGTCCGAGGGAGATCATCAGAATATCAACCAATCATGCTC

ACCAGCAATCGGACCAAACAAAGTCTACTTGTCTCCTGAAGATAATCTCGGATTTAGAGAGATCACTGGCAACGACTGTG

AGGCTGGGCTCGGAGGGGTCCAGAGAGAAGGATCCAACTCTCAAGTACAGCGTTACCATGTTTATAGTCACGGGGGTGAA

GAGATTGAAGGACTCGAGGATGCTGACTCTCTCGTGGTTCAAGCAGATCCTCCAGTTGCTAACGTCTTCAATGGAGGAGA

GGATGGATCTGACGACAGCGATGTGGACTCTGGCCCAGATGATCCCGGCAGAGATACTCTATATGACCGGGGATCTGTTG

CCGGCAATGGTATCGCTAGGTCCACAGATGTCGAAAAACTAGAAGGTGCTGATATTCAAGAAGTTCTTAACTCCCAGAAA

GGCAAAGGAGGAAGATTCCAGGGCGGGAAAACCTTGCGAGTCCCGGAAATACCCGATGTCAAGCACTCCAGACCATCAGC

CCAATCAATTAAAAAGGGCACAGACGGGAACTCAGTCTCATCTGGAACGGTGATAGAGTGTTTATCGATAAGTGGTGCAA

CCCAAACTGTGCCAGAGTCAAGATGGGAGTCATCAGAGCAAAATGCGTCTGTGGGGAGTGTCCTCAAGTCTGCGAGGAGT

GCAAAGACGATCCAGGGGTCGACACAAGAATCTGGTACCATAGCATCACTGACTCAGCCTAAAGAGAATGACTCCGAGTA

TGAGTATGAGGATGACCTATTTACAGAGATTCAGGACATCCGTGCAAGCATTGCCAAGATCCATGATGACAATAAAACTA

TCCTCTCAAAGCTTGATTCTATACTGTTATTGAAAGGGGAAGTCGACACTATCAAGAAACAAATCAGCAAGCAGAATATA

AGTATATCCACCATTGAGGGCCATCTCTCCAGTATAATGATAGCCATCCCTGGCTTTGGGAAGGACATCAAGGACCCAAC

ATCCGAAGTCGAGTTGAACCCAGATCTAAGACCTATAATAAGCCGTGACTCTGGCAGAGCTCTCGCGGAGGTCCTCAAGA

AACCCGCTGTGGATAGGTCTCAGAAAATTGGAACCAAAGCCAACTCCAGCTCAAAGGGTCAGCTTCTTAAGGATCTCCAG

CTAAAACCTGTCGACAAGCAGGCAAGCTCTGCAATCGGGTTTGTCCCGTCCGACCATGAATCATCCAGAAATGTCATCCG

CTCCATAATCAAGTCGAGCAAGCTAAATATTGATCACAAGGACTATCTTCTAGATTTACTAAATGATGTGAAAGGCTCCA

AGGATCTTAAGGAATTTCACAAGATGCTAACAGCAATTCTCGCCAAGCACCCGTAACACATCCTCCAGTCATCATCTCAT

ACTCGACTAAAAACATCCTTTCAATCAGGCTATTACAAAAAACTTAGGAGCAAGGGCAACTGAGCTTCGCAGACAGGATG

ACCGAGATCTACGACTTCGATAAATCAGCATGGGATGTCAAAGGGTCAATTGCCCGCATAGAACCCACCACCTACCACGA

CGGCCGACTGATACCCCAGGTGAGGGTCATCGATCCTGGTCTGGGAGACAGAAAAGATGAGTGCTTTATGTACCTGTTTC

TCCTAGGAGTGATTGAGGATAACGACCCCCTGTCTCCCCCCGTCGGAAGAACCTTTGGCTCTCTACCTCTAGGGGTTGGT

AGGTCAACTGCCAAGCCAGAAGAACTACTAAGGGAGGCCACAGAATTAGATATAGTGGTGAGGCGCACTGCAGGAGTAAA

TGAGAAACTGGTATTTTACAACAACACTCCGCTGTCCTTGTTAACGCCCTGGAAGAAAGTTCTGACAACCGGAAGTGTGT

TTAGCGCTAACCAGGTTTGCAATGCAGTCAACCTAGTCCCGCTTGATACTCCCCAGAGATTCAGGGTTGTGTACATGAGC

ATAACTAGATTATCAGACAATGGTTACTATAGTGTGCCCAGAAGAATGCTGGAGTTCCGCTCGGCCAATGCAGTAGCCTT

CAACATCTTGGTTACACTAAGAATTGAAAATGGCACAAACCCTAGAAGATACATAGTCGGCTCATGGGAGAATTCAGAGG

TCACATTTATGGTACACGTGGGCAACTTTAGGAGAAAGAAGAACGAAGTATACTCTGCTGATTACTGCAAAATGAAGATT

GAAAAGATGGGTTTAGTTTTTGCCCTGGGTGGAATAGGCGGAACAAGTCTCCATATTAGAAGCACCGGGAAAATGAGCAA

AACCCTCCATGCACAGCTGGGGTTCAAGAAAATCTTATGTTACCCCCTAATGGATGTCAATGAGGATCTTAACCGATATC

TCTGGCGGGCAGAGTGCCGAATAGTCAAAATCCAWGCTGTCTTACAGCCATCAGTACCCCAAGAGTTCCGTGTCTACGAT

GATGTCATCATCAACGATGATCAAGGCTTGTTCAAGATCTTGTAGTTCATTTGCAAAGGAACGAACCCACCCCCAGGAAG

GGGCCCCACCCTCCCCCG-CCCAAAAACACACTCCAAGGCTCCCCGGCACAGCGCCAACACAGCTACTCCCGCGATGGAA

CGAGGTCGGCC-CCCGAACAAATCCGGTCCAGGGGGGGGGGCCCCCCCCCTGACCAACCGGGCGCCGACCGAGCAGGCCG

GCGAGAAGCCCCCAAGAGGCCCCGACGAAGACCGAGCACCCCATCCCCCC-AAAAACGTCCCCC-AACACGCCCGAGGCT

AGCCACCGGACACCCCCAGACCCGCCACCGGCGAGACAGAACCCCAAGGGGGRACCCTYCARACTAAAGGAATCACCGGG

GTAGGCTCCGCCGAACCAGCACAGACCCGACCCAAGCAAGGCGGACCCCCAACACGAAACTGCCCAGCACACCCTACTCA

TCATCAAACAAAACTTAGGGGCCAAGTCCATACGAGAGGCCCTAGTCCGTCAGGCGGAGGGACGCAGAAAGGAAGGAGAC

ACGACCGTCCCAGAAGGAGGACACCCAACCACCGAAACGCCCGCCCCCAGGGAAGGAG--CCA-CCCCCACGCAGCGAAG

GAGCGGGTGGGAGACCTCGGGGAGGGCCCGACCGGGACCCGGGGGCCGGCGGCGCCGGCGGTGAGCAGGCCGGGCGCATA

CCTGACCCCACCGGAGGATCCGAGGAGGGCCCGACCCCGCGTGGACCAAACGGCAGGGAGCCCAACCCCACCAAAAAAAG

AGGCGAGGCCCCCCCTCCCTCCCCCCCCGCACACAACCCGCAGACAGACCGAGGGAGGGCCATCCCCCAGCGAGACAGGC

GGCTCCCTCGCAGCCGAGCCGCAGCTCCCCAGAATAGGACACCCCCGCACACTCCGAACACCCCCTCGCCCCCCGCCACC

GGGACAGGGGCTCCACCACCACAGACCGGAGCCACCGACAGCCAGCCAGAGGTCCCCTCCAGTCCCGCCCGGCGCCCCCC

CGAAGAGGCCCCGGCAAGGCCCCCCAGCCCGACGTCCCCCGCACGCACCGAAAGAAGGACCCGCCCGAGAACGTGCGGCA

CATCCATGCGCAAACATCATGACGCGGGTCGCAATTTTGACATTTCTGTTTCTTTTCCCAAATGTTGTTGCGTGTCAGAT

TCACTGGGGCAATCTATCCAAGATCGGGATTGTAGGAACAGGGAGTGCCAGCTACAAGGTGATGACTAGGCCAAGCCACC

AGACTCTGGTTATAAAGTTAATGCCAAATATAACGGCCATCGACAATTGTACAAAGTCAGAGATTGCAGAGTACAAGAGA

TTGCTGATCACAGTGTTAAAGCCTGTAGAGGATGCTCTGTCGGTGATAACCAAGAATGTAAGACCAATTCAAACTCTAAC

ACCTGGGCGTAGAACCCGCCGTTTTGCTGGAGCTGTTCTGGCCGGGGTAGCACTTGGAGTTGCGACAGCCGCTCAGATAA

CTGCAGGAGTCGCCCTTCATCAATCATTGATGAACTCCCAAGCAATTGAGAGTTTAAAAACCAGTCTTGAGATGTCGAAT

CAGGCAATAGAAGAAATCAGACTTGCAAATAAGGAGACCATACTGGCAGTACAGGGCGTCCAGGATTATATCAACAATGA

GCTCGTCCCTTCTGTTTATAGAATGTCATGCGAGCTGGTAGGTCACAAGCTCGGCCTCAAGCTCCTTAGGTACTACACCG

AGATCCTGTCCATATTCGGGCCCAGTCTTCGAGACCCGATATCTGCCGAAATATCAATCCAGGCACTTAGTTATGCATTA

GGCGGAGACATTAATAAAATCCTGGACAAGCTTGGGTATAGCGGTGGGGATTTCCTTGCCATCCTAGAAAGCAAGGGAAT

AAAGGCCCGGGTTACATATGTGGACACAAGAGATTACTTTATAATCCTTAGCATCGCCTACCCAACCTTATCTGAGATCA

AGGGAGTGATAGTTCACAAGATAGAAGCTATAACATACAACATTGGGGCACAGGAGTGGTATACTACTATCCCTAAATAT

GTAGCCACTCAGGGGTATCTGATATCGAACTTTGATGAGACGTCATGCGTATTCACTCCAGAGGGGACAGTTTGCAGCCA

GAATGCGTTGTACCCAATGAGCCCATTGCTTCAGGAATGTTTCAGGGGGTCAACAAAATCGTGCGCCAGAACCTTAGTTT

CAGGGACCATAAGTAATAGATTTATCCTATCAAAAGGGAACCTGATTGCAAATTGTGCGTCAGTTTTGTGCAAATGTTAC

ACAACAGAGACAGTTATCAGCCAAGATCCTGACAAACTACTAACTGTTGTAGCATCCGACAAGTGTCCTGTAGTTGAGGT

GGATGGAGTGACAATACAGGTCGGCAGTCGAGAGTATCCGGATTCTGTATACTTACACAAAATAGACTTAGGTCCAGCCA

TCTCCCTAGAAAAACTGGATGTAGGCACCAATTTAGGCAATGCAGTCACAAGACTGGAGAATGCAAAGGAGCTCCTAGAT

GCATCAGACCAAATACTGAAGACTGTTAAAGGGGTACCTTTCAGTGGGAATATGTACATAGCACTGGCAGCTTGCATAGG

AGTATCCCTGGGCCTTGTCACATTAATATGCTGCTGTAAGGGGAGGTGTAAGAACAAGGAAATCCCTATCTCCAAAATCA

ACCCAGGGCTCAAACCCGACCTGACCGGGACCTCAAAGTCGTACGTAAGATCACTGTAGTCAGAATCACCTGGATCATCT

GGCATCGCACACATACATGCACGACACAGGCAGTTCGAGGACGCAAGAAACCCAGCCTCCGGTCACCCACCCGACCCCAC

TCCACGCTCCACCACACATTAGTCATCAAACAAAACTTAGGACGAAAGGTCAATCACCATGTCCGCACAAAGGGAGAGGA

TCAATGCCTTCTACAAAGACAATCCTCACAATAAGAACCATAGGGTGATCCTGGATAGAGAACGCTTGGTCATTGAAAGA

CCCTACATCTTGCTTGGAGTCCTGCTGGTAATGTTCCTGAGTCTAATCGGACTGCTGGCCATTGCAGGGATCAGGCTTCA

CCGGGCCACCGTTGGAACTTCAGAGATCCAGAGTCGGCTGAATACCAATATTAAGTTGACCGAATCTATTGATCACCAGA

CTAAGGATGTCTTAACTCCCCTTTTTAAAATCATTGGCGATGAAGTCGGCATCAGAATTCCACAGAAATTCAGTGATCTT

GTCAAGTTCATCTCCGATAAGATTAAATTCCTCAACCCTGATAGAGAGTATGATTTCAGAGATCTCCGGTGGTGTATGAA

TCCCCCCGAGAGAGTCAAAATTAATTTTGATCAGTTTTGTGAGTACAAGGCTGCGGTTAAGTCAATTGAACATATATTTG

AGTCACCACTCAACAAGTCAAAAAAGCTGCAATCTTTGACTCTCGGGCCCGGAACAGGCTGTCTAGGCAGGACAGTAACA

AGAGCCCATTTCTCAGAACTTACAATGACCTTAATGGACCTGGATCTAGAGATGAAGCACAACGTGTCCTCAGTGTTTAC

CGTAGTTGAAGAGGGATTATTCGGAAGAACATATACCGTCTGGAGATCCGATGCCAGGGATCCGAGCACCGATCTAGGTA

TCGGCCATTTTTTAAGAGTCTTCGAGATTGGACTGGTAAGAGATCTTGGGCTGGGTCCCCCTGTTTTTCATATGACCAAC

TATCTCACGGTGAACATGAGTGATGACTATCGGAGATGTCTTTTAGCGGTAGGGGAGTTGAAGTTGACAGCCCTATGCAC

CTCATCTGAGACTGTGACACTGAGTGAGAGAGGAGTTCCAAGGAGGGAACCTCTTGTGGTTGTGATACTTAATCTAGCTG

GACCCACTCTAGGGGGAGAGCTATACAGTGTCTTGCCTACCTCTGATCTCATGGTGGAGAAACTCTATTTGTCTTCACAT

AGAGGGATCATCAAAGATGATGAGGCCAATTGGGTAGTGCCGTCTACCGATGTTCGTGATCTTCAGAACAAAGGTGAATG

TCTGGTGGAAGCATGCAAGACTCGACCTCCTTCATTTTGCAATGGCACAGGATCAGGCCCGTGGTCAGAAGGGAGAATCC

CTGCCTACGGGGTGATCAGGGTCAGTCTTGACTTAGCGAGTGACCCGGATGTAGTTATCACTTCAGTGTTTGGCCCACTG

ATACCTCACCTATCCGGCATGGATCTTTACAACAACCCGTTTTCAAAAGCTGTATGGTTGGCTGTACCACCTTATGAGCA

GTCATTTCTAGGAATGATAAATACAATTGGATTCCCTAACAGAGCAGAGGTTATGCCGCACATTTTGACCACAGAGATCA

GAGGCCCTCGGGGTCGTTGCCATGTTCCCATAGAATTGTCCCGCCGGGTTGATGACGATATCAAGATCGGGTCCAACATG

GTCATATTGCCGACGATGGACCTGAGGTATATTACAGCCACTTATGATGTTTCCAGGAGCGAGCATGCAATCGTGTACTA

TATCTATGACACGAGTCGCTCATCATCTTACTTCTACCCAGTTCGACTGAATTTCAAAGGCAATCCTCTCTCTTTGAGGA

TAGAGTGTTTCCCCTGGCGTCATAAGGTGTGGTGCTACCATGATTGTCTTATATACAACACCATAACAGGTGAAGAGGTC

CATACGAGAGGGCTGACCGGCATAGAGGTAACATGCAATCCAGTCTGAGCAGAGCTACGACCGTCACTCAGGCAGTCCCT

TGAGTCGCCACCGAGTCCAAGCAGCACAGCCTGGGACAATCAACGGCACAGCCCAGCCAACAATGTTATAAAAAACTTAG

GAGCCAGGGTTGTAGGGGCCATGGACTCTCTATCAGTCAATCAGGTCTTGTACCCTGAGGTTCATCTAGACAGCCCTATT

GTCACAAACAAACTAGTTGCCATCCTTGAGTACTCGGGTATCGACCATAACTATGTTCTTGAAGACCAGACTCTTGTCAA

GAATATTAGGTATAGGCTGGGGTGCGGTTTTTCAAATCAAATGATCATCAATAACAGGGGGGTGGGTGAAACAGTCAATT

CTAAACTTAAAAGTTACCCCCATAATCGTCATATCATATACCCGGATTGCAATAAGGAGTTGTTTTGTATCAAGGATAGC

TGCATATCTAAGAAGCTCTCGGAGCTATTCAAGAAGGGTAATTCCTTGTACTCTAAGATAAGTCACCAGGTACTGGATTG

TCTTAAGAGAGTCAATGGGAAATTAGGTCTGGGCACAGATCTTACCCATGGTCTGAAGGAGGGTATCCTTGACCTAGGGT

TGCACATGCATAGCTCTCAATGGTTCGAGACTTTCCTGTTCTGGTTCACTATCAAGACAGAGATGAGATCAATGATCAAA

GAACAGTCCCATATATGCCACAAGAGGAGGTATAACCCGACTTTTGTGTCGGGGGATGCATTCGAGGTGCTCGTATCGCG

AGACCTCGTTGTGATAATTGATAAGAATACCCAGTATGTCTTCTACTTAACATTTGAGCTGGTCCTTATGTATTGTGATG

TCATAGAGGGTAGACTTATGACGGAGACAGCCATGGCCATAGACCAGAGATATTCAGAACTTCTAAGCCGGGTCAGATAC

TTGTGGGATCTTATTGATGGGTTCTTCCCAACACTAGGCAACACCACATACCAAATTGTTGCTCTGCTTGAACCATTATC

GTTGGCTTATCTTCAACTTCAGGATGTCACTCTGGAGTTAAGAGGTGCTTTTTTAGACCATTGCTTCAAAGAACTCTATG

AGATACTGGAACATTGCGGCATTGACACAGCAGGCACTTACAATTCCATCACTGAAGGATTGGATTACGTATTCATCACC

CACGATATACATCTAACTGGGGAGATTTTTTCATTTTTCCGGAGTTTCGGACACCCCCGTCTAGAAGCAGTCACCGCTGC

GGAAAATGTCAGAAAACATATGAACCAACCGAAGGTAATCAGTTATGAGACTATGATGAAAGGCCATGCGGTATTTTGCG

GGATAATCATAAATGGTTTTCGGGACCGACATGGCGGCAGCTGGCCCCCTGTTGCACTGCCAGAACATGCCTCTGCTGCG

ATCCGGAATGCGCAGGCATCAGGTGAGGGACTGACCCATGACCTGTGTATAGACAACTGGAAATCCTTTGTTGGATTCAA

ATTTGGCTGCTTTATGCCACTCAGCCTAGATAGTGATTTGACCATGTATCTCAAAGACAAGGCATTGGCTGCACTGAAGA

ATGAGTGGGATTCAGTTTACCCGAAAGAATACCTCCGTTATAACCCACCTAGAGGGACAGAGTCAAGGCGATTGGTAGAG

GTATTCCTGAATGACTCCAGCTTTGATCCTTATAACATGATAATGTACGTGGTGAATGGCTCCTACCTTAAAGACCCTGA

GTTTAACCTCTCATACAGCCTAAAAGAGAAGGAGATAAAGGAGACAGGGCGGTTGTTTGCCAAAATGACCTATAAGATGC

GGGCCTGTCAGGTAATCGCTGAAAATCTGATATCGAATGGTGTTGGGAAGTATTTCCGAGACAATGGGATGGCAAAGGAC

GAGCATGACCTAACAAAAGCCCTTCACACCCTGGCAGTCTCAGGTGTTCCCAAAAACAACAAAGATAACCACCGAGGTGG

GCCTCCCAGAAGGACAACAAACCGAGGGGTGAGATCAAGCCAAGGCACCAAAACACAAGATAGAGACAAGGTTCAAGGGG

GACCTATGTACAACTATTTGCGATGCCAACCGATCAGCCCTGATCAGGGTGAGTCATACGAGACTGTTAGTGCATTCATC

ACCGCTGACCTTAAGAAGTATTGCCTGAATTGGAGATACGAGACAATCAGCATATTTGCACAGAGGCTGAATGAAATATA

TGGACTACCATCCTTCTTTCAATGGTTACACAGGATATTGGAAAAATCCGTACTCTACGTCAGTGACCCACATTGTCCTC

CCGATCTAGATAATCATATCCCTCTGGACAGTGTCCCCAATGCCCAAATATTCATCAAGTACCCAATGGGCGGAATAGAA

GGTTATTGCCAAAAACTATGGACAATCAGCACTATACCATACTTGTATCTGGCAGCCTATGAGAGCGGAGTAAGAATCGC

CTCACTGGTGCAGGGTGACAATCAGACAATCGCAGTGACAAAAAGAGTTCCAAGTTCTTGGCCTTATTCACTAAAAAAGA

GGGAGGCCTCCAAGGCCGCTCAAAATTACTTCGTAGTCTTAAGGCAAAGATTGCACGATGTAGGTCATCACTTAAAGGCT

AATGAGACCATAGTATCTTCTCACTTTTTTGTATATTCCAAAGGGATTTATTATGACGGCCTGCTAGTCTCACAATCACT

AAAGAGCATCGCCAGATGTGTCTTCTGGTCCGAGACTATAGTGGATGAAACCAGAGCGGCGTGCAGTAATATTGCAACAA

CTGTCGCTAAGAGTATAGAGAGGGGTTATGATAGGTACCTTGCATATTCTTTGAATATCCTCAAGATTTTCCAACAGATC

CTTATATCCCTTAACTTCACTATTAACACAACAATGACTCAGGATGTCGTGGCACCGATCATCGAGAACGGTGATTTGCT

GATAAGGATGGCACTCTTGCCAGCACCCATCGGGGGTCTCAATTATCTTAACATGAGCAGGTTATTTGTGAGAAATATCG

GTGACCCGGTCACCTCCTCTATAGCCGACCTGAAGAGGATGATAGATGCCGGGCTAATGCCAGAAGAAACATTGCATCAA

GTGATGACCCAGACCCCGGGAGAATCATCCTACCTAGATTGGGCAAGTGACCCTTATTCTGCCAACCTACCCTGCGTACA

GAGTATCACTCGCCTTCTCAAAAACATCACTGCACGATATATTTTAATCAGCAGCCCAAACCCGATGCTGAAAGGGTTAT

TTCATGAGGGGAGTAGAGATGAAGACGAGGAGCTTGCAAGTTTCCTAATGGATCGGCATATAATTGTTCCAAGAGCTGCA

CATGAAATCTTAGACCATAGTATAACCGGAGCAAGAGAAGCTATAGCCGGGATGTTGGACACCACCAAGGGTCTGATTAG

AACAAGTATGAAACGGGGCGGCCTCACCCCTCGAGTCTTAGCCCGCCTTTCCAATTATGATTATGAACAATTCCGATCCG

GGATAACACTATTAACAAAGAAAGGGCAGTGTTATCTCATTGACAAGGACTCGTGCTCGGTGCAGCTCGCTATAGCTCTG

AGGGGCCACATGTGGGCTAAGTTGGCTCGCGGGAGACCAATCTATGGGTTGGAGGTGCCTGATGTATTAGAATCGATGAA

CGGCTACCTTATCAAACGCCACGAGTCGTGTGCTATCTGTGAAACGGGCTCAAGTCACTACGGATGGTTTTTCGTCCCTG

CAGGGTGCCAGCTTGACGATGTATCTAGAGAGACTTCGGCTCTTCGTGTACCTTATGTCGGATCAACAACTGAGGAAAGG

ACAGACATGAAACTTGCATTTGTTAGGTCTCCAAGTAGATCCCTCAAATCAGCAGTTAGAATTGCAACAGTTTACTCATG

GGCTTACGGGGATGATGAGAAATCATGGAGTGAAGCCTGGATGCTAGCTAGGCAGAGAGCCAATATCACCTTAGATGAAT

TGAGAATGATCACTCCAGTCTCCACATCCACCAACTTAGCCCACCGGTTGAGGGATCGGAGTACCCAGGTGAAATACTCG

GGGACATCCCTTGTGAGGGTTGCAAGATACACAACCATCTCCAATGATAACTTGTCGTTTGTGATATCTGAGAAAAAAGT

GGATACCAACTTCATTTATCAGCAAGGGATGTTGCTTGGTCTTGGGATCCTTGAGAACCTCTTTAGGCTAGAGGCCACCA

CCGGAGTATCCAACACAGTGCTACACCTGCACGTGGAAACAGAATGTTGTGTTGTACCTATGGTGGATCACCCAAGGATA

CCGAGTCTCCGTAATATAAAAGTTACGGATGAGCTATGCACAAACCCTTTGATCTACGACAGGTCCCCCATCATAGAGCA

CGATGCAACCCGGCTATATTCACAGAGCCACAGGAGACATTTGGTGGAGTTTGTTACCTGGTCAACAAGCCAGCTTTACC

ATATACTGGCTAAGTCTACAGCAATGTCCATGATTGAATTGATCACGAGATTCGAGAAAGATCACATGAATGAAATAGCC

GCCCTGATTGGCGATGACGACATCAACAGCTTCATCACAGAATTTTTGCTAGTGGAGCCTAGATTATTTATAGTTTACCT

TGGTCAGTGTGCTGCCATCAATTGGGCCTTTGATATACATTATCATCGGCCCTCGGGCAAGTACCAGATGGGGGAACTTC

TCTACTCTCTGCTCTCTCGGATGAGCAAAGGAGTATATAAGATCTTTACTAATGCTTTGAGCCACCCCAAAGTTTACAAG

AAATTTTGGCGAAGTGGCATCATTGAGCCGATTCATGGCCCATCCTTGGATACACAGAATTTACATGTCACTGTCTGTGA

CATGATATATGGGTCCTATGTCACCTATTTGGATCTTTTGCTGAATGATGAGCTAGATGATTACCCGTATTTGCTCTGCG

AGAGTGATGAGGACGTGGTCACAGACAGGTTCGACAACATTCAAGCCAAACATCTCTGTGTATTGGCCGATGTATATTGC

AGCTCCAAGAGGTGTCCTTCAATAATCGGGATGTCTCCTATAGAAAAATGTACCATTCTCACACATTACATCAAGGGAGA

ATCAGTACAATCCCCGTCTGGGACCTCATGGAACACTGATCCCCTTGTAGTAGATCATTACTCATGCTCTCTGACCTACC

TTCGCCGCGGTTCCATCAAACAAATCAGGTTGAGGGTGGATCCTGGGTTTGTATTCGAGGCGTTGACAGACGTCGACTTT

AAACAACCTCGCAAAGCTAAGTTAGATATATCGGTCGTTGGGTTGACTGATTTCTCTCCCCCTTACGACAACGTCGGTGA

TTTTTTAGGGACTATCAACACATTGAGGCACGATCTGCCTGTCACCGGAACCGGGGTCTCGAACTATGAAGTCCACGCTT

ATCGTAGAATTGGCCTGAATTCATCAGCATGTTATAAAGCCGTAGAGATCTCCACGTTAATCAAGCCATCCTTAGAAGTC

GGAGAGCATGGATTGTTCTTAGGAGAAGGTTCCGGTTCAATGCTGGCTGCGTACAAAGAAGTTCTTAAATTAGCAAATTG

TTACTACAACAGCGGAGTAACAGCGGAGGGCAGAGCCGGACAGAGGGAAATATCTCCCTATCCTTCAGAGATGAGCCTAG

TAGAGAATCAAATGGGGATAGAGCGGAGTGTTAAAGTGCTGTTCAACGGGAAACCTGAAGTAACTTGGGTGGGGACCACA

GATTGCTACAAGTACATAATCAGTAACATCCAAACCTCTAGTCTGGGTTTCATACATTCAGATATTGAGACGCTCCCAAC

CAAAGATGCTGTTGAGAAATTGGAGGAATTTGCCTCTATCCTATCCTTATCCTTGATTTTGGGGAAAATCGGCTCTATTA

CAGTTGTCAAAATTATGCCCATTAGCGGGGATTTTACCCAAGGCTTCATAGCCTATGCCATCCAATATTTTAGGGAGAGC

CTGCTTGCCTACCCGAGATATAGTAACTTCATCTCGACTGAGTGTTACCTTATTATGATAGGATTAAAGGCCAATCGATT

GATAAACCCAGAAGCCATTAAGCAAAGCATAATCAGAGCGGGGGTCAGGACTGCACCAGGACTTGTGAGCCATGTATTAT

CAGAGAAACAAAAAGGTTGTATTCAATCTTTCTTGGGTGATCCTTATATCCAAGGAGACTTCAATAAGCACCTTAAATCT

CTAACCTCTATTGAGAAAATCCTGGTAAATTGTGGTCTCTCGATCAATGGCATAAAAATCTGTAGAGATCTAATCCACCA

TGATATCGCCTCCGGTCCAGACGGCCTTATGAGCTCTACAATTATTTTATATAGGGAATTGGCTCATTTCAAAGATAATA

TAAGGAGTCAGCACGGAATGTTCCACCCCTACCCGGTACTGGCCAACAGCAGGCAACGTGAATTAATCCTTCGAATAGCC

AAGAAATTCTGGGGGTATGTCTTGCTATATTCTGATGACCCGGCACTAATCAGACAAACAATCAAGAACTTGAAACGGAA

TCATCTAACCTTTGACTTACATAGTAACCCATTTATTAAAGGGCTATCCAAGGCTGAGAAACTGCTGGTGCGGACGAGTT

CACTAAGAAGAGAATGGTTGTTCACTCTCGAAACGAAAGAAGTAAAAGAGTGGTTCAAATTGGTAGGTTACAGTGCACTC

ATCAGAGGCTAGTCAGCTATGCATCTGCCCCCTTCTCCTCCGCCATGAGACCCCACTGGCGATCCAGAAGATTAAAGAAA

ACTACATATTGGATAAGTATCTATTCCCAGCTTTGTCTGGT

>MF443349.1/goat/China/2014-04-21

ACCAAACAAAGTTGGGTAAGGATAGATCTTATAATAACTATGGACTGGCAAACTTAGGAGTAAAGATCCTACTGTCGGGG

GGAGGAGGAGGAGCAAGATCTTTGATTATGGCGACTCTCCTTAAAAGCTTAGCATTGTTCAAAAGGAACAAAGACAAAGC

GCCGACAGCATCAGGTTCAGGAGGGGCCATCCGGGGGATTAAGAATGTTATCATAGTCCCGATTCCCGGAGACTCGTCCA

TCACTACCCGTTCAAGACTGCTCGACAGGCTTGTCAGATTGGCCGGAGATCCTGACATCAACGGGTCAAAGCTGACCGGG

GTGATGATCAGCATGTTATCCTTGTTCGTAGAGTCACCCGGGCAATTGATACAGCGAATCACAGATGATCCGGATGTTAG

TATCCGCCTTGTTGAGGTAGTTCAAAGTACTAGGTCCCAGTCCGGGTTGACCTTTGCATCACGTGGTGCTGATTTAGACA

ACGAGGCAGACATGTATTTTTCAACTGAGGGGCCCTCGAGTGGAGGTAAGAAAAGGATCAACTGGTTTGAGAACAGAGAA

ATAATAGACATAGAGGTGCAGGATCCAGAAGAGTTCAATATGTTGTTAGCCTCCATACTAGCACAAGTCTGGATCCTTCT

GGCCAAGGCTGTTACGGCACCAGATACGGCAGCTGACTCAGAACTGAGAAGGTGGGTTAAATACACACAACAAAGGAGAG

TGATTGGGGAATTTCGCCTTGACAAGGGGTGGCTGGACGCAGTCCGCAACAGAATTGCAGAAGATCTATCACTCCGGCGG

TTCATGGTATCTCTCATACTTGACATCAAGAGGACCCCAGGCAACAAGCCAAGGATTGCAGAAATGATCTGCGACATTGA

CAACTATATTGTCGAAGCAGGACTCGCCAGTTTCATCCTTACTATCAAATTTGGTATCGAAACCATGTATCCTGCATTAG

GGCTCCACGAGTTTGCCGGAGAATTGTCCACTATTGAGTCCTTGATGAACTTGTATCAACAGCTAGGAGAGGTTGCACCC

TACATGGTAATTCTAGAGAATTCAATTCAGAACAAGTTTAGTGCAGGAGCTTATCCCCTCCTCTGGAGCTATGCGATGGG

TGTCGGAGTCGAGCTGGAGAACTCAATGGGGGGCTTGAACTTTGGCAGGTCATATTTTGACCCTGCCTATTTTCGTCTCG

GACAGGAGATGGTCAGAAGATCTGCAGGAAAGGTCAGCTCTGTGATTGCGGCTGAGCTCGGCATCACAGCAGAGGAAGCT

AAACTAGTCTCGGAAATCGCCTCGCAGGCTGGGGACGAAAGAACCGCCAGAGGGACTGGGCCTCGACAGGCGCAGGTTTC

CTTCCTTCAGCACCAAACAGGAGGGGGAGAGTCGTCCGCACCAGCGACCAGAGAAGGGGTCAAAGCTGCGATCCCAAACG

GAGCTGAAGAAAGGGACAGAAAGCAAACACGCCCAGGAAGGCCCAGAGGAGAGACCTCCGGCCAACTGCTCCTGGACATC

ATGCCAGAGGATGAGATCTTGCGAGAGTCTGGTCAAAACCCTCGTGAGGCTCAAAGATCGGCCGAGGCACTCTTCAGGCT

GCAGGCCATGGCCAAGATTCTGGAGGACCAGGAGGAGGGAGAAGACAACAGTCAGGTCTACAACGACAAGGATCTCCTCG

GCTAAGCAGACGCACCCTCTGTCGAAATCAGTGACAAGACATCTCTTACCAGTATTATAAAAAACTTAGGACCCAGGTCC

AAGCAAGCACACATCGACACTCCAACCAGTCGAGCGGAGACCACCGATGGCAGAAGAACAAGCATACCATGTCAACAAGG

GGCTGGAATGTATCAAGTCTCTCAAAGCCTCACCCCCGGATCTATCCACCATCAGAGATACCATCGAGAGCTGGAGAGAG

GGGCTTAGCCCCTCGGGCCGTGCAACACCGAACCCTGATACGTCCGAGGGAGATCATCAGAATATCAACCAATCATGCTC

ACCAGCAATCGGACCAAACAAAGTCTACTTGTCTCCTGAAGATAATCTCGGATTTAGAGAGATCACTGGCAACGACTGTG

AGGCTGGGCTCGGAGGGGTCCAGAGAGAAGGATCCAACTCTCAAGTACAGCGTTACCATGTTTATAGTCACGGGGGTGAA

GAGATTGAAGGACTCGAGGATGCTGACTCTCTCGTGGTTCAAGCAGATCCTCCAGTTGCTAACGTCTTCAATGGAGGAGA

GGATGGATCTGACGACAGCGATGTGGACTCTGGCCCAGATGATCCCGGCAGAGATACTCTATATGACCGGGGATCTGTTG

CCGGCAATGGTATCGCTAGGTCCACAGATGTCGAAAAACTAGAAGGTGCTGATATTCAAGAAGTTCTTAACTCCCAGAAA

GGCAAAGGAGGAAGATTCCAGGGCGGGAAAACCTTGCGAGTCCCGGAAATACCCGATGTCAAGCACTCCAGACCATCAGC

CCAATCAATTAAAAAGGGCACAGACGGGAACTCAGTCTCATCTGGAACGGTGATAGAGTGTTTATCGATAAGTGGTGCAA

CCCAAACTGTGCCAGAGTCAAGATGGGAGTCATCAGAGCAAAATGCGTCTGTGGGGAGTGTCCTCAAGTCTGCGAGGAGT

GCAAAGACGATCCAGGGGTCGACACAAGAATCTGGTACCATAGCATCACTGACTCAGCCTAAAGAGAATGACTCCGAGTA

TGAGTATGAGGATGACCTATTTACAGAGATTCAGGACATCCGTGCAAGCATTGCCAAGATCCATGATGACAATAAAACTA

TCCTCTCAAAGCTTGATTCTATACTGTTATTGAAAGGGGAAGTCGACACTATCAAGAAACAAATCAGCAAGCAGAATATA

AGTATATCCACCATTGAGGGCCATCTCTCCAGTATAATGATAGCCATCCCTGGCTTTGGGAAGGACATCAAGGACCCAAC

ATCCGAAGTCGAGTTGAACCCAGATCTAAGACCTATAATAAGCCGTGACTCTGGCAGAGCTCTCGCGGAGGTCCTCAAGA

AACCCGCTGTGGATAGGTCTCAGAAAATTGGAACCAAAGCCAACTCCAGCTCAAAGGGTCAGCTTCTTAAGGATCTCCAG

CTAAAACCTGTCGACAAGCAGGCAAGCTCTGCAATCGGGTTTGTCCCGTCCGACCATGAATCATCCAGAAATGTCATCCG

CTCCATAATCAAGTCGAGCAAGCTAAATATTGATCACAAGGACTATCTTCTAGATTTACTAAATGATGTGAAAGGCTCCA

AGGATCTTAAGGAATTTCACAAGATGCTAACAGCAATTCTCGCCAAGCACCCGTAACACATCCTCCAGTCATCATCTCAT

ACTCGACTAAAAACATCCTTTCAATCAGGCTATTACAAAAAACTTAGGAGCAAGGGCAACTGAGCTTCGCAGACAGGATG

ACCGAGATCTACGACTTCGATAAATCAGCATGGGATGTCAAAGGGTCAATTGCCCGCATAGAACCCACCACCTACCACGA

CGGCCGACTGATACCCCAGGTGAGGGTCATCGATCCTGGTCTGGGAGACAGAAAAGATGAGTGCTTTATGTACCTGTTTC

TCCTAGGAGTGATTGAGGATAACGACCCCCTGTCTCCCCCCGTCGGAAGAACCTTTGGCTCTCTACCTCTAGGGGTTGGT

AGGTCAACTGCCAAGCCAGAAGAACTACTAAGGGAGGCCACAGAATTAGATATAGTGGTGAGGCGCACTGCAGGAGTAAA

TGAGAAACTGGTATTTTACAACAACACTCCGCTGTCCTTGTTAACGCCCTGGAAGAAAGTTCTGACAACCGGAAGTGTGT

TTAGCGCTAACCAGGTTTGCAATGCAGTCAACCTAGTCCCGCTTGATACTCCCCAGAGATTCAGGGTTGTGTACATGAGC

ATAACTAGATTATCAGACAATGGTTACTATAGTGTGCCCAGAAGAATGCTGGAGTTCCGCTCGGCCAATGCAGTAGCCTT

CAACATCTTGGTTACACTAAGAATTGAAAATGGCACAAACCCTAGAAGATACATAGTCGGCTCATGGGAGAATTCAGAGG

TCACATTTATGGTACACGTGGGCAACTTTAGGAGAAAGAAGAACGAAGTATACTCTGCTGATTACTGCAAAATGAAGATT

GAAAAGATGGGTTTAGTTTTTGCCCTGGGTGGAATAGGCGGAACAAGTCTCCATATTAGAAGCACCGGGAAAATGAGCAA

AACCCTCCATGCACAGCTGGGGTTCAAGAAAATCTTATGTTACCCCCTAATGGATGTCAATGAGGATCTTAACCGATATC

TCTGGCGGGCAGAGTGCCGAATAGTCAAAATCCAAGCTGTCTTACAGCCATCAGTACCCCAAGAGTTCCGTGTCTACGAT

GATGTCATCATCAACGATGATCAAGGCTTGTTCAAGATCTTGTAGTTCATTTGCAAAGGAACGAACCCACCCCCAGGAAG

GGGCCCCACCCTCCCCCG-CCCAAAAACACACTCCAAGGCTCCCCGGCACAGCGCCAACACAGCTACTCCCGCGATGGAA

CGAGGTCGGCC-CCCGAACAAATCCGGTCCAGGGGGGGGGGCCCCCCCCCTGACCAACCGGGCGCCGACCGAGCAGGCCG

GCGAGAAGCCCCCAAGAGGCCCCGAGGAAGACCGAGCACCCCATCCCCCC-AAAAACGTCCCCC-AACACGCCCGAGGCT

AGCCACCGGACACCCCCAGACCCGCCACCGGCGAGACAGAACCCCAAGGGGGGACCCTCCAAACCAAAGGAATCACCGGG

GTAGGCTCCGCCGAACCAGCACAGACCCGACCCAAGCAAGGCGGACCCCCAACACGAAACTGCCCAGCACACCCTACTCA

TCATCAAACAAAACTTAGGGGCCAAGTCCATACGAGAGGCCCTAGTCCGTCAGGCGGAGGGACGCAGAAAGGAAGGAGAC

ACGACCGTCCCAGAAGGAGGACACCCAACCACCGAAACGCCCGCCCCCAGGGAAGGAG--CCA-CCCCCACGCAGCGAAG

GAGCGGGTGGGAGACCTCGGGGAGGGCCCGACCGGGACCCGGGGGCCGGCGGCGCCGGCGGTGAGCAGGCCGGGCGCATA

CCTGACCCCACCGGAGGATCCGAGGAGGGCCCGACCCCGCGTGGACCAAACGGCAGGGAGCCCAACCCCACCAAAAAAAG

AGGCGAGGCCCCCCCTCCCCCCCCCCCCGCACACAACCCGCAGACAGACCGAGGGAGGGCCATCCCCCAGCGAGACAGGC

GGCTCCCTCGCAGCCGAGCCGCAGCTCCCCAGAATAGGACACCCCCGCACACTCCGAACACCCCCTCGCCCCCCGCCACC

GGGACAGGGGCTCCACCACCACAGACCGGAGCCACCGACAGCCAGCCAGAGGTCCCCTCCAGTCCCGCCCGGCGCCCCCC

CGAAGAGGCCCCGGCAAGGCCCCCCAGCCCGACGTCCCCCGCACGCACCGAAAGAAGGACCCGCCCGAGAACGTGCGGCA

CATCCATGCGCAAACATCATGACGCGGGTCGCAATTTTGACATTTCTGTTTCTTTTCCCAAATGTTGTTGCGTGTCAGAT

TCACTGGGGCAATCTATCCAAGATCGGGATTGTAGGAACAGGGAGTGCCAGCTACAAGGTGATGACTAGGCCAAGCCACC

AGACTCTGGTTATAAAGTTAATGCCAAATATAACGGCCATCGACAATTGTACAAAGTCAGAGATTGCAGAGTACAAGAGA

TTGCTGATCACAGTGTTAAAGCCTGTAGAGGATGCTCTGTCGGTGATAACCAAGAATGTAAGACCAATTCAAACTCTAAC

ACCTGGGCGTAGAACCCGCCGTTTTGCTGGAGCTGTTCTGGCCGGGGTAGCACTTGGAGTTGCGACAGCCGCTCAGATAA

CTGCAGGAGTCGCCCTTCATCAATCATTGATGAACTCCCAAGCAATTGAGAGTTTAAAAACCAGTCTTGAGATGTCGAAT

CAGGCAATAGAAGAAATCAGACTTGCAAATAAGGAGACCATACTGGCAGTACAGGGCGTCCAGGATTATATCAACAATGA

GCTCGTCCCTTCTGTTCATAGAATGTCATGCGAGCTGGTAGGTCACAAGCTCGGCCTCAAGCTCCTTAGGTACTACACCG

AGATCCTGTCCATATTCGGGCCCAGTCTTCGAGACCCGATATCTGCCGAAATATCAATCCAGGCACTTAGTTATGCATTA

GGCGGAGACATTAATAAAATCCTGGACAAGCTTGGGTATAGCGGTGGGGATTTCCTTGCCATCCTAGAAAGCAAGGGAAT

AAAGGCCCGGGTTACATATGTGGACACAAGAGATTACTTTATAATCCTTAGCATCGCCTACCCAACCTTATCTGAGATCA

AGGGAGTGATAGTTCACAAGATAGAAGCTATAACATACAACATTGGGGCACAGGAGTGGTATACTACTATCCCTAAATAT

GTAGCCACTCAGGGGTATCTGATATCGAACTTTGATGAGACGTCATGCGTATTCACTCCAGAGGGGACAGTTTGCAGCCA

GAATGCGTTGTACCCAATGAGCCCATTGCTTCAGGAATGTTTCAGGGGGTCAACAAAATCGTGCGCCAGAACCTTAGTTT

CAGGGACCATAAGTAATAGATTTATCCTATCAAAAGGGAACCTGATTGCAAATTGTGCGTCAGTTTTGTGCAAATGTTAC

ACAACAGAGACAGTTATCAGCCAAGATCCTGACAAACTACTAACTGTTGTAGCATCCGACAAGTGTCCTGTAGTTGAGGT

GGATGGAGTGACAATACAGGTCGGCAGTCGAGAGTATCCGGATTCTGTATACTTACACAAAATAGACTTAGGTCCAGCCA

TCTCCCTAGAAAAACTGGATGTAGGCACCAATTTAGGCAATGCAGTCACAAGACTGGAGAATGCAAAGGAGCTCCTAGAT

GCATCAGACCAAATACTGAAGACTGTTAAAGGGGTACCTTTCAGTGGGAATATGTACATAGCACTGGCAGCTTGCATAGG

AGTATCCCTGGGCCTTGTCACATTAATATGCTGCTGTAAGGGGAGGTGTAAGAACAAGGAAATCCCTATCTCCAAAATCA

ACCCAGGGCTCAAACCCGACCTGACCGGGACCTCAAAGTCGTACGTAAGATCACTGTAGTCAGAATCACCTGGATCATCT

GGCATCGCACACATACATGCACGACACAGGCAGTCCGAGGACGCAAGAAACCCAGCCTCCGGTCACCCACCCGACCCCAC

TCCACGCTCCACCACACATTAGTCATCAAACAAAACTTAGGACGAAAGGTCAATCACCATGTCCGCACAAAGGGAGAGGA

TCAATGCCTTCTACAAAGACAATCCTCACAATAAGAACCATAGGGTGATCCTGGATAGAGAACGCTTGGTCATTGAAAGA

CCCTACATCTTGCTTGGAGTCCTGCTGGTAATGTTCCTGAGTCTAATCGGACTGCTGGCCATTGCAGGGATCAGGCTTCA

CCGGGCCACCGTTGGAACTTCAGAGATCCAGAGTCGGCTGAATACCAATATTAAGTTGACCGAATCTATTGATCACCAGA

CTAAGGATGTCTTAACTCCCCTTTTTAAAATCATTGGCGATGAAGTCGGCATCAGAATTCCACAGAAATTCAGTGATCTT

GTCAAGTTCATCTCCGATAAGATTAAATTCCTCAACCCTGATAGAGAGTATGATTTCAGAGATCTCCGGTGGTGTATGAA

TCCCCCCGAGAGAGTCAAAATTAATTTTGATCAGTTTTGTGAGTACAAGGCTGCGGTTAAGTCAATTGAACATATATTTG

AGTCACCACTCAACAAGTCAAAAAAGCTGCAATCTTTGACTCTCGGGCCCGGAACAGGCTGTCTAGGCAGGACAGTAACA

AGAGCCCATTTCTCAGAACTTACAATGACCTTAATGGACCTGGATCTAGAGATGAAGCACAACGTGTCCTCAGTGTTTAC

CGTAGTTGAAGAGGGATTATTCGGAAGAACATATACCGTCTGGAGATCCGATGCCAGGGATCCGAGCACCGATCTAGGTA

TCGGCCATTTTTTAAGAGTCTTCGAGATTGGACTGGTAAGAGATCTTGGGCTGGGTCCCCCTGTTTTTCATATGACCAAC

TATCTCACGGTGAACATGAGTGATGACTATCGGAGATGTCTTTTAGCGGTAGGGGAGTTGAAGTTGACAGCCCTATGCAC

CTCATCTGAGACTGTGACACTGAGTGAGAGAGGAGTTCCAAGGAGGGAACCTCTTGTGGTTGTGATACTTAATCTAGCTG

GACCCACTCTAGGGGGAGAGCTATACAGTGTCTTGCCTACCTCTGATCTCATGGTGGAGAAACTCTATTTGTCTTCACAT

AGAGGGATCATCAAAGATGATGAGGCCAATTGGGTAGTGCCGTCTACCGATGTTCGTGATCTTCAGAACAAAGGTGAATG

TCTGGTGGAAGCATGCAAGACTCGACCTCCTTCATTTTGCAATGGCACAGGATCAGGCCCGTGGTCAGAAGGGAGAATCC

CTGCCTACGGGGTGATCAGGGTCAGTCTTGACTTAGCGAGTGACCCGGATGTAGTTATCACTTCAGTGTTTGGCCCACTG

ATACCTCACCTATCCGGCATGGATCTTTACAACAACCCGTTTTCAAAAGCTGTATGGTTGGCTGTACCACCTTATGAGCA

GTCATTTCTAGGAATGATAAATACAATTGGATTCCCTAACAGAGCAGAGGTTATGCCGCACATTTTGACCACAGAGATCA

GAGGCCCTCGGGGTCGTTGCCATGTTCCCATAGAATTGTCCCGCCGGGTTGATGACGATATCAAGATCGGGTCCAACATG

GTCATATTGCCGACGATGGACCTGAGGTATATTACAGCCACTTATGATGTTTCCAGGAGCGAGCATGCAATCGTGTACTA

TATCTATGACACGAGTCGCTCATCATCTTACTTCTACCCAGTTCGACTGAATTTCAAAGGCAATCCTCTCTCTTTGAGGA

TAGAGTGTTTCCCCTGGCGTCATAAGGTGTGGTGCTACCATGATTGTCTTATATACAACACCATAACAGGTGAAGAGGTC

CATACGAGAGGGCTGACCGGCATAGAGGTAACATGCAATCCAGTCTGAGCAGAGCTACGACCGTCACTCAGGCAGTCCCT

TGAGTCGCCACCGAGTCCAAGCAGCACAGCCTGGGACAATCAACGGCACAGCCCAGCCAACAATGTTATAAAAAACTTAG

GAGCCAGGGTTGTAGGGGCCATGGACTCTCTATCAGTCAATCAGGTCTTGTACCCTGAGGTTCATCTAGACAGCCCTATT

GTCACAAACAAACTAGTTGCCATCCTTGAGTACTCGGGTATCGACCATAACTATGTTCTTGAAGACCAGACTCTTGTCAA

GAATATTAGGTATAGGCTGGGGTGCGGTTTTTCAAATCAAATGATCATCAATAACAGGGGGGTGGGTGAAACAGTCAATT

CTAAACTTAAAAGTTACCCCCATAATCGTCATATCATATACCCGGATTGCAATAAGGAGTTGTTTTGTATCAAGGATAGC

TGCATATCTAAGAAGCTCTCGGAGCTATTCAAGAAGGGTAATTCCTTGTACTCTAAGATAAGTCACCAGGTACTGGATTG

TCTTAAGAGAGTCAATGGGAAATTAGGTCTGGGCACAGATCTTACCCATGGTCTGAAGGAGGGTATCCTTGACCTAGGGT

TGCACATGCATAGCTCTCAATGGTTCGAGACTTTCCTGTTCTGGTTCACTATCAAGACAGAGATGAGATCAATGATCAAA

GAACAGTCCCATATATGCCACAAGAGGAGGTATAACCCGACTTTTGTGTCGGGGGATGCATTCGAGGTGCTCGTATCGCG

AGACCTCGTTGTGATAATTGATAAGAATACCCAGTATGTCTTCTACTTAACATTTGAGCTGGTCCTTATGTATTGTGATG

TCATAGAGGGTAGACTTATGACGGAGACAGCCATGGCCATAGACCAGAGATATTCAGAACTTCTAAGCCGGGTCAGATAC

TTGTGGGATCTTATTGATGGGTTCTTCCCAACACTAGGCAACACCACATACCAAATTGTTGCTCTGCTTGAACCATTATC

GTTGGCTTATCTTCAACTTCAGGATGTCACTCTGGAGTTAAGAGGTGCTTTTTTAGACCATTGCTTCAAAGAACTCTATG

AGATACTGGAACATTGCGGCATTGACACAGCAGGCACTTACAATTCCATCACTGAAGGATTGGATTACGTATTCATCACC

CACGATATACATCTAACTGGGGAGATTTTTTCATTTTTCCGGAGTTTCGGACACCCCCGTCTAGAAGCAGTCACCGCTGC

GGAAAATGTCAGAAAACATATGAACCAACCGAAGGTAATCAGTTATGAGACTATGATGAAAGGCCATGCGGTATTTTGCG

GGATAATCATAAATGGTTTTCGGGACCGACATGGCGGCAGCTGGCCCCCTGTTGCACTGCCAGAACATGCCTCTGCTGCG

ATCCGGAATGCGCAGGCATCAGGTGAGGGACTGACCCATGACCTGTGTATAGACAACTGGAAATCCTTTGTTGGATTCAA

ATTTGGCTGCTTTATGCCACTCAGCCTAGATAGTGATTTGACCATGTATCTCAAAGACAAGGCATTGGCTGCACTGAAGA

ATGAGTGGGATTCAGTTTACCCGAAAGAATACCTCCGTTATAACCCACCTAGAGGGACAGAGTCAAGGCGATTGGTAGAG

GTATTCCTGAATGACTCCAGCTTTGATCCTTATAACATGATAATGTACGTGGTGAATGGCTCCTACCTTAAAGACCCTGA

GTTTAACCTCTCATACAGCCTAAAAGAGAAGGAGATAAAGGAGACAGGGCGGTTGTTTGCCAAAATGACCTATAAGATGC

GGGCCTGTCAGGTAATCGCTGAAAATCTGATATCGAATGGTGTTGGGAAGTATTTCCGAGACAATGGGATGGCAAAGGAC

GAGCATGACCTAACAAAAGCCCTTCACACCCTGGCAGTCTCAGGTGTTCCCAAAAACAACAAAGATAACCACCGAGGTGG

GCCTCCCAGAAGGACAACAAACCGAGGGGTGAGATCAAGCCAAGGCACCAAAACACAAGATAGAGACAAGGTTCAAGGGG

GACCTATGTACAACTATTTGCGATGCCAACCGATCAGCCCTGATCAGGGTGAGTCATACGAGACTGTTAGTGCATTCATC

ACCGCTGACCTTAAGAAGTATTGCCTGAATTGGAGATACGAGACAATCAGCATATTTGCACAGAGGCTGAATGAAATATA

TGGACTACCATCCTTCTTTCAATGGTTACACAGGATATTGGAAAAATCCGTACTCTACGTCAGTGACCCACATTGTCCCC

CCGATCTAGATAATCATATCCCTCTGGACAGTGTCCCCAATGCCCAAATATTCATCAAGTACCCAATGGGCGGAATAGAA

GGTTATTGCCAAAAACTATGGACAATCAGCACTATACCATACTTGTATCTGGCAGCCTATGAGAGCGGAGTAAGAATCGC

CTCACTGGTGCAGGGTGACAATCAGACAATCGCAGTGACAAAAAGAGTTCCAAGTTCTTGGCCTTATTCACTAAAAAAGA

GGGAGGCCTCCAAGGCCGCTCAAAATTACTTCGTAGTCTTAAGGCAAAGATTGCACGATGTAGGTCATCACTTAAAGGCT

AATGAGACCATAGTATCTTCTCACTTTTTTGTATATTCCAAAGGGATTTATTATGACGGCCTGCTAGTCTCACAATCACT

AAAGAGCATCGCCAGATGTGTCTTCTGGTCCGAGACTATAGTGGATGAAACCAGAGCGGCGTGCAGTAATATTGCAACAA

CTGTCGCTAAGAGTATAGAGAGGGGTTATGATAGGTACCTTGCATATTCTTTGAATATCCTCAAGATTTTCCAACAGATC

CTTATATCCCTTAACTTCACTATTAACACAACAATGACTCAGGATGTCGTGGCACCGATCATCGAGAACGGTGATTTGCT

GATAAGGATGGCACTCTTGCCAGCACCCATCGGGGGTCTCAATTATCTTAACATGAGCAGGTTATTTGTGAGAAATATCG

GTGACCCGGTCACCTCCTCTATAGCCGACCTGAAGAGGATGATAGATGCCGGGCTAATGCCAGAAGAAACATTGCATCAA

GTGATGACCCAGACCCCGGGAGAATCATCCTACCTAGATTGGGCAAGTGACCCTTATTCTGCCAACCTACCCTGCGTACA

GAGTATCACTCGCCTTCTCAAAAACATCACTGCACGATATATTTTAATCAGCAGCCCAAACCCGATGCTGAAAGGGTTAT

TTCATGAGGGGAGTAGAGATGAAGACGAGGAGCTTGCAAGTTTCCTAATGGATCGGCATATAATTGTTCCAAGAGCTGCA

CATGAAATCTTAGACCATAGTATAACCGGAGCAAGAGAAGCTATAGCCGGGATGTTGGACACCACCAAGGGTCTGATTAG

AACAAGTATGAAACGGGGCGGCCTCACCCCTCGAGTCTTAGCCCGCCTTTCCAATTATGATTATGAACAATTCCGATCCG

GGATAACACTATTAACAAAGAAAGGGCAGTGTTATCTCATTGACAAGGACTCGTGCTCGGTGCAGCTCGCTATAGCTCTG

AGGGGCCACATGTGGGCTAAGTTGGCTCGCGGGAGACCAATCTATGGGTTGGAGGTGCCTGATGTATTAGAATCGATGAA

CGGCTACCTTATCAAACGCCACGAGTCGTGTGCTATCTGTGAAACGGGCTCAAGTCACTACGGATGGTTTTTCGTCCCTG

CAGGGTGCCAGCTTGACGATGTATCTAGAGAGACTTCGGCTCTTCGTGTACCTTATGTCGGATCAACAACTGAGGAAAGG

ACAGACATGAAACTTGCATTTGTTAGGTCTCCAAGTAGATCCCTCAAATCAGCAGTTAGAATTGCAACAGTTTACTCATG

GGCTTACGGGGATGATGAGAAATCATGGAGTGAAGCCTGGATGCTAGCTAGGCAGAGAGCCAATATCACCTTAGATGAAT

TGAGAATGATCACTCCAGTCTCCACATCCACCAACTTAGCCCACCGGTTGAGGGATCGGAGTACCCAGGTGAAATACTCG

GGGACATCCCTTGTGAGGGTTGCAAGATACACAACCATCTCCAATGATAACTTGTCGTTTGTGATATCTGAGAAAAAAGT

GGATACCAACTTCATTTATCAGCAAGGGATGTTGCTTGGTCTTGGGATCCTTGAGAACCTCTTTAGGCTAGAGGCCACCA

CCGGAGTATCCAACACAGTGCTACACCTGCACGTGGAAACAGAATGTTGTGTTGTACCTATGGTGGATCACCCAAGGATA

CCGAGTCTCCGTAATATAAAAGTTACGGATGAGCTATGCACAAACCCTTTGATCTACGACAGGTCCCCCATCATAGAGCA

CGATGCAACCCGGCTATATTCACAGAGCCACAGGAGACATTTGGTGGAGTTTGTTACCTGGTCAACAAGCCAGCTTTACC

ATATACTGGCTAAGTCTACAGCAATGTCCATGATTGAATTGATCACGAGATTCGAGAAAGATCACATGAATGAAATAGCC

GCCCTGATTGGCGATGACGACATCAACAGCTTCATCACAGAATTTTTGCTAGTGGAGCCTAGATTATTTATAGTTTACCT

TGGTCAGTGTGCTGCCATCAATTGGGCCTTTGATATACATTATCATCGGCCCTCGGGCAAGTACCAGATGGGGGAACTTC

TCTACTCTCTGCTCTCTCGGATGAGCAAAGGAGTATATAAGATCTTTACTAATGCTTTGAGCCACCCCAAAGTTTACAAG

AAATTTTGGCGAAGTGGCATCATTGAGCCGATTCATGGCCCATCCTTGGATACACAGAATTTACATGTCACTGTCTGTGA

CATGATATATGGGTCCTATGTCACCTATTTGGATCTTTTGCTGAATGATGAGCTAGATGATTACCCGTATTTGCTCTGCG

AGAGTGATGAGGACGTGGTCACAGACAGGTTCGACAACATTCAAGCCAAACATCTCTGTGTATTGGCCGATGTATATTGC

AGCTCCAAGAGGTGTCCTTCAATAATCGGGATGTCTCCTATAGAAAAATGTACCATTCTCACACATTACATCAAGGGAGA

ATCAGTACAATCCCCGTCTGGGACCTCATGGAACACTGATCCCCTTGTAGTAGATCATTACTCATGCTCTCTGACCTACC

TTCGCCGCGGTTCCATCAAACAAATCAGGTTGAGGGTGGATCCTGGGTTTGTATTCGAGGCGTTGACAGACGTCGACTTT

AAACAACCTCGCAAAGCTAAGTTAGATATATCGGTCGTTGGGTTGACTGATTTCTCTCCCCCTTACGACAACGTCGGTGA

TTTTTTAGGGACTATCAACACATTGAGGCACGATCTGCCTGTCACCGGAACCGGGGTCTCGAACTATGAAGTCCACGCTT

ATCGTAGAATTGGCCTGAATTCATCAGCATGTTATAAAGCCGTAGAGATCTCCACGTTAATCAAGCCATCCTTAGAAGTC

GGAGAGCATGGATTGTTCTTAGGAGAAGGTTCCGGTTCAATGCTGGCTGCGTACAAAGAAGTTCTTAAATTAGCAAATTG

TTACTACAACAGCGGAGTAACAGCGGAGGGCAGAGCCGGACAGAGGGAAATATCTCCCTATCCTTCAGAGATGAGCCTAG

TAGAGAATCAAATGGGGATAGAGCGGAGTGTTAAAGTGCTGTTCAACGGGAAACCTGAAGTAACTTGGGTGGGGACCACA

GATTGCTACAAGTACATAATCAGTAACATCCAAACCTCTAGTCTGGGTTTCATACATTCAGATATTGAGACGCTCCCAAC

CAAAGATGCTGTTGAGAAATTGGAGGAATTTGCCTCTATCCTATCCTTATCCTTGATTTTGGGGAAAATCGGCTCTATTA

CAGTTGTCAAAATTATGCCCATTAGCGGGGATTTTACCCAAGGCTTCATAGCCTATGCCATCCAATATTTTAGGGAGAGC

CTGCTTGCCTACCCGAGATATAGTAACTTCATCTCGACTGAGTGTTACCTTATTATGATAGGATTAAAGGCCAATCGATT

GATAAACCCAGAAGCCATTAAGCAAAGCATAATCAGAGCGGGGGTCAGGACTGCACCAGGACTTGTGAGCCATGTATTAT

CAGAGAAACAAAAAGGTTGTATTCAATCTTTCTTGGGTGATCCTTATATCCAAGGAGACTTCAATAAGCACCTTAAATCT

CTAACCTCTATTGAGAAAATCCTGGTAAATTGTGGTCTCTCGATCAATGGCATAAAAATCTGTAGAGATCTAATCCACCA

TGATATCGCCTCCGGTCCAGACGGCCTTATGAGCTCTACAATTATTTTATATAGGGAATTGGCTCATTTCAAAGATAATA

TAAGGAGTCAGCACGGAATGTTCCACCCCTACCCGGTACTGGCCAACAGCAGGCAACGTGAATTAATCCTTCGAATAGCC

AAGAAATTCTGGGGGTATGTCTTGCTATATTCTGATGACCCGGCACTAATCAGACAAACAATCAAGAACTTGAAACGGAA

TCATCTAACCTTTGACTTACATAGTAACCCATTTATTAAAGGGCTATCCAAGGCTGAGAAACTGCTGGTGCGGACGAGTT

CACTAAGAAGAGAATGGTTGTTCACTCTCGAAACGAAAGAAGTAAAAGAGTGGTTCAAATTGGTAGGTTACAGTGCACTC

ATCAGAGGCTAGTCAGCTATGCATCTGCCCCCTTCTCCTCCGCCATGAGACCCCACTGGCGATCCAGAAGATTAAAGAAA

ACTACATATTGGATAAGTATCTATTCCCAGCTTTGTCTGGT

>MF443338.1/goat/China/2014-06-10

ACCAAACAAAGTTGGGTAAGGATAGATCTTATAATAACTATGGACTGGCAAACTTAGGAGTAAAGATCCTACTGTCGGGG

GGAGGAGGAGGAGCAAGATCTTTGATTATGGCGACTCTCCTTAAAAGCTTAGCATTGTTCAAAAGGAACAAAGACAAAGC

GCCGACAGCATCAGGTTCAGGAGGGGCCATCCGGGGGATTAAGAATGTTATCATAGTCCCGATTCCCGGAGACTCGTCCA

TCACTACCCGTTCAAGACTGCTCGACAGGCTTGTCAGATTGGCCGGAGATCCTGACATCAACGGGTCAAAGCTGACCGGG

GTGATGATCAGCATGTTATCCTTGTTCGTAGAGTCACCCGGGCAATTGATACAGCGAATCACAGATGATCCGGATGTTAG

TATCCGCCTTGTTGAGGTAGTTCAAAGTACTAGGTCCCAGTCCGGGTTGACCTTTGCATCACGTGGTGCTGATTTAGACA

ACGAGGCAGACATGTATTTTTCAACTGAGGGGCCCTCGAGTGGAGGTAAGAAAAGGATCAACTGGTTTGAGAACAGAGAA

ATAATAGACATAGAGGTGCAGGATCCAGAAGAGTTCAATATGTTGTTAGCCTCCATACTAGCACAAGTCTGGATCCTTCT

GGCCAAGGCTGTTACGGCACCAGATACGGCAGCTGACTCAGAACTGAGAAGGTGGGTTAAATACACACAACAAAGGAGAG

TGATTGGGGAATTTCGCCTTGACAAGGGGTGGCTGGACGCAGTCCGCAACAGAATTGCAGAAGATCTATCACTCCGGCGG

TTCATGGTATCTCTCATACTTGACATCAAGAGGACCCCAGGCAACAAGCCAAGGATTGCAGAAATGATCTGCGACATTGA

CAACTATATTGTCGAAGCAGGACTCGCCAGTTTCATCCTTACTATCAAATTTGGTATCGAAACCATGTATCCTGCATTAG

GGCTCCACGAGTTTGCCGGAGAATTGTCCACTATTGAGTCCTTGATGAACTTGTATCAACAGCTAGGAGAGGTTGCACCC

TACATGGTAATTCTAGAGAATTCAATTCAGAACAAGTTTAGTGCAGGAGCTTATCCCCTCCTCTGGAGCTATGCGATGGG

TGTCGGAGTCGAGCTGGAGAACTCAATGGGGGGCTTGAACTTTGGCAGGTCATATTTTGACCCTGCCTATTTTCGTCTCG

GACAGGAGATGGTCAGAAGATCTGCAGGAAAGGTCAGCTCTGTGATTGCGGCTGAGCTCGGCATCACAGCAGAGGAAGCT

AAACTAGTCTCGGAAATCGCCTCGCAGGCTGGGGACGAAAGAACCGCCAGAGGGACTGGGCCTCGACAGGCGCAGGTTTC

CTTCCTTCAGCACCAAACAGGAGGGGGAGAGTCGTCCGCACCAGCGACCAGAGAAGGGGTCAAAGCTGCGATCCCAAACG

GAGCTGAAGAAAGGGACAGAAAGCAAACACGCCCAGGAAGGCCCAGAGGAGAGACCTCCGGCCAACTGCTCCTGGACATC

ATGCCAGAGGATGAGATCTTGCGAGAGTCTGGTCAAAACCCTCGTGAGGCTCAAAGATCGGCCGAGGCACTCTTCAGGCT

GCAGGCCATGGCCAAGATTCTGGAGGACCAGGAGGAGGGAGAAGACAACAGTCAGGTCTACAACGACAAGGATCTCCTCG

GCTAAGCAGACGCACCCTCTGTCGAAATCAGTGACAAGACATCTCTTACCAGTATTATAAAAAACTTAGGACCCAGGTCC

AAGCAAGCACACATCGACACTCCAACCAGTCGAGCGGAGACCACCGATGGCAGAAGAACAAGCATACCATGTCAACAAGG

GGCTGGAATGTATCAAGTCTCTCAAAGCCTCACCCCCGGATCTATCCACCATCAGAGATACCATCGAGAGCTGGAGAGAG

GGGCTTAGCCCCTCGGGCCGTGCAACACCGAACCCTGATACGTCCGAGGGAGATCATCAGAATATCAACCAATCATGCTC

ACCAGCAATCGGACCAAACAAAGTCTACTTGTCTCCTGAAGATAATCTCGGATTTAGAGAGATCACTGGCAACGACTGTG

AGGCTGGGCTCGGAGGGGTCCAGAGAGAAGGATCCAACTCTCAAGTACAGCGTTACCATGTTTATAGTCACGGGGGTGAA

GAGATTGAAGGACTCGAGGATGCTGACTCTCTCGTGGTTCAAGCAGATCCTCCAGTTGCTAACGTCTTCAATGGAGGAGA

GGATGGATCTGACGACAGCGATGTGGACTCTGGCCCAGATGATCCCGGCAGAGATACTCTATATGACCGGGGATCTGTTG

CCGGCAATGGTATCGCTAGGTCCACAGATGTCGAAAAACTAGAAGGTGCTGATATTCAAGAAGTTCTTAACTCCCAGAAA

GGCAAAGGAGGAAGATTCCAGGGCGGGAAAACCTTGCGAGTCCCGGAAATACCCGATGTCAAGCACTCCAGACCATCAGC

CCAATCAATTAAAAAGGGCACAGACGGGAACTCAGTCTCATCTGGAACGGTGATAGAGTGTTTATCGATAAGTGGTGCAA

CCCAAACTGTGCCAGAGTCAAGATGGGAGTCATCAGAGCAAAATGCGTCTGTGGGGAGTGTCCTCAAGTCTGCGAGGAGT

GCAAAGACGATCCAGGGGTCGACACAAGAATCTGGTACCATAGCATCACTGACTCAGCCTAAAGAGAATGACTCCGAGTA

TGAGTATGAGGATGACCTATTTACAGAGATTCAGGACATCCGTGCAAGCATTGCCAAGATCCATGATGACAATAAAACTA

TCCTCTCAAAGCTTGATTCTATACTGTTATTGAAAGGGGAAGTCGACACTATCAAGAAACAAATCAGCAAGCAGAATATA

AGTATATCCACCATTGAGGGCCATCTCTCCAGTATAATGATAGCCATCCCTGGCTTTGGGAAGGACATCAAGGACCCAAC

ATCCGAAGTCGAGTTGAACCCAGATCTAAGACCTATAATAAGCCGTGACTCTGGCAGAGCTCTCGCGGAGGTCCTCAAGA

AACCCGCTGTGGATAGGTCTCAGAAAATTGGAACCAAAGCCAACTCCAGCTCAAAGGGTCAGCTTCTTAAGGATCTCCAG

CTAAAACCTGTCGACAAGCAGGCAAGCTCTGCAATCGGGTTTGTCCCGTCCGACCATGAATCATCCAGAAATGTCATCCG

CTCCATAATCAAGTCGAGCAAGCTAAATATTGATCACAAGGACTATCTTCTAGATTTACTAAATGATGTGAAAGGCTCCA

AGGATCTTAAGGAATTTCACAAGATGCTAACAGCAATTCTCGCCAAGCACCCGTAACACATCCTCCAGTCATCATCTCAT

ACTCGACTAAAAACATCCTTTCAATCAGGCTATTACAAAAAACTTAGGAGCAAGGGCAACTGAGCTTCGCAGACAGGATG

ACCGAGATCTACGACTTCGATAAATCAGCATGGGATGTCAAAGGGTCAATTGCCCGCATAGAACCCACCACCTACCACGA

CGGCCGACTGATACCCCAGGTGAGGGTCATCGATCCTGGTCTGGGAGACAGAAAAGATGAGTGCTTTATGTACCTGTTTC

TCCTAGGAGTGATTGAGGATAACGACCCCCTGTCTCCCCCCGTCGGAAGAACCTTTGGCTCTCTACCTCTAGGGGTTGGT

AGGTCAACTGCCAAGCCAGAAGAACTACTAAGGGAGGCCACAGAATTAGATATAGTGGTGAGGCGCACTGCAGGAGTAAA

TGAGAAACTGGTATTTTACAACAACACTCCGCTGTCCTTGTTAACGCCCTGGAAGAAAGTTCTGACAACCGGAAGTGTGT

TTAGCGCTAACCAGGTTTGCAATGCAGTCAACCTAGTCCCGCTTGATACTCCCCAGAGATTCAGGGTTGTGTACATGAGC

ATAACTAGATTATCAGACAATGGTTACTATAGTGTGCCCAGAAGAATGCTGGAGTTCCGCTCGGCCAATGCAGTAGCCTT

CAACATCTTGGTTACACTAAGAATTGAAAATGGCACAAACCCTAGAAGATACATAGTCGGCTCATGGGAGAATTCAGAGG

TCACATTTATGGTACACGTGGGCAACTTTAGGAGAAAGAAGAACGAAGTATACTCTGCTGATTACTGCAAAATGAAGATT

GAAAAGATGGGTTTAGTTTTTGCCCTGGGTGGAATAGGCGGAACAAGTCTCCATATTAGAAGCACCGGGAAAATGAGCAA

AACCCTCCATGCACAGCTGGGGTTCAAGAAAATCTTATGTTACCCCCTAATGGATGTCAATGAGGATCTTAACCGATATC

TCTGGCGGGCAGAGTGCCGAATAGTCAAAATCCAAGCTGTCTTACAGCCATCAGTACCCCAAGAGTTCCGTGTCTACGAT

GATGTCATCATCAACGATGATCAAGGCTTGTTCAAGATCTTGTAGTTCATTTGCAAAGGAACGAACCCACCCCCAGGAAG

GGGCCCCACCCTCCCCCG-CCCAAAAACACACTCCAAGGCTCCCCGGCACAGCGCCAACACAGCTACTCCCGCGATGGAA

CGAGGTCGGCC-CCCGAACAAATCCGGTCCAGGGGGGGGGGCCCCCCCCCTGACCAACCGGGCGCCGACCGAGCAGGCCG

GCGAGAAGCCCCCAAGAGGCCCCGACGAAGACCGAGCACCCCATCCCCCC-AAAAACGTCCCCC-AACACGCCCGAGGCT

AGCCACCGGACACCCCCAGACCCGCCACCGGCGAGACAGAACCCCAAGGGGGGACCCTCCAAACCAAAGGAATCACCGGG

GTAGGCTCCGCCGAACCAGCACAGACCCGACCCAAGCAAGGCGGACCCCCAACACGAAACTGCCCAGCACACCCTACTCA

TCATCAAACAAAACTTAGGGGCCAAGTCCATACGAGAGGCCCTAGTCCGTCAGGCGGAGGGACGCAGAAAGGAAGGAGAC

ACGACCGTCCCAGAAGGAGGACACCCAACCACCGAAACGCCCGCCCCCAGGGAAGGAG--CCA-CCCCCACGCAGCGAAG

GAGCGGGTGGGAGACCTCGGGGAGGGCCCGACCGGGACCCGGGGGCCGGCGGCGCCGGCGGTGAGCAGGCCGGGCGCATA

CCTGACCCCACCGGAGGATCCGAGGAGGGCCCGACCCCGCGTGGACCAAACGGCAGGGAGCCCAACCCCACCAAAAAAAG

AGGCGAGGCCCCCCCTCCCTCCCCCCCCGCACACAACCCGCAGACAGACCGAGGGAGGGCCATCCCCCAGCGAGACAGGC

GGCTCCCTCGCAGCCGAGCCGCAGCTCCCCAGAATAGGACACCCCCGCACACTCCGAACACCCCCTCGCCCCCCGCCACC

GGGACAGGGGCTCCACCACCACAGACCGGAGCCACCGACAGCCAGCCAGAGGTCCCCTCCAGTCCCGCCCGGCGCCCCCC

CGAAGAGGCCCCGGCAAGGCCCCCCAGCCCGACGTCCCCCGCACGCACCGAAAGAAGGACCCGCCCGAGAACGTGCGGCA

CATCCATGCGCAAACATCATGACGCGGGTCGCAATTTTGACATTTCTGTTTCTTTTCCCAAATGTTGTTGCGTGTCAGAT

TCACTGGGGCAATCTATCCAAGATCGGGATTGTAGGAACAGGGAGTGCCAGCTACAAGGTGATGACTAGGCCAAGCCACC

AGACTCTGGTTATAAAGTTAATGCCAAATATAACGGCCATCGACAATTGTACAAAGTCAGAGATTGCAGAGTACAAGAGA

TTGCTGATCACAGTGTTAAAGCCTGTAGAGGATGCTCTGTCGGTGATAACCAAGAATGTAAGACCAATTCAAACTCTAAC

ACCTGGGCGTAGAACCCGCCGTTTTGCTGGAGCTGTTCTGGCCGGGGTAGCACTTGGAGTTGCGACAGCCGCTCAGATAA

CTGCAGGAGTCGCCCTTCATCAATCATTGATGAACTCCCAAGCAATTGAGAGTTTAAAAACCAGTCTTGAGATGTCGAAT

CAGGCAATAGAAGAAATCAGACTTGCAAATAAGGAGACCATACTGGCAGTACAGGGCGTCCAGGATTATATCAACAATGA

GCTCGTCCCTTCTGTTCATAGAATGTCATGCGAGCTGGTAGGTCACAAGCTCGGCCTCAAGCTCCTTAGGTACTACACCG

AGATCCTGTCCATATTCGGGCCCAGTCTTCGAGACCCGATATCTGCCGAAATATCAATCCAGGCACTTAGTTATGCATTA

GGCGGAGACATTAATAAAATCCTGGACAAGCTTGGGTATAGCGGTGGGGATTTCCTTGCCATCCTAGAAAGCAAGGGAAT

AAAGGCCCGGGTTACATATGTGGACACAAGAGATTACTTTATAATCCTTAGCATCGCCTACCCAACCTTATCTGAGATCA

AGGGAGTGATAGTTCACAAGATAGAAGCTATAACATACAACATTGGGGCACAGGAGTGGTATACTACTATCCCTAAATAT

GTAGCCACTCAGGGGTATCTGATATCGAACTTTGATGAGACGTCATGCGTATTCACTCCAGAGGGGACAGTTTGCAGCCA

GAATGCGTTGTACCCAATGAGCCCATTGCTTCAGGAATGTTTCAGGGGGTCAACAAAATCGTGCGCCAGAACCTTAGTTT

CAGGGACCATAAGTAATAGATTTATCCTATCAAAAGGGAACCTGATTGCAAATTGTGCGTCAGTTTTGTGCAAATGTTAC

ACAACAGAGACAGTTATCAGCCAAGATCCTGACAAACTACTAACTGTTGTAGCATCCGACAAGTGTCCTGTAGTTGAGGT

GGATGGAGTGACAATACAGGTCGGCAGTCGAGAGTATCCGGATTCTGTATACTTACACAAAATAGACTTAGGTCCAGCCA

TCTCCCTAGAAAAACTGGATGTAGGCACCAATTTAGGCAATGCAGTCACAAGACTGGAGAATGCAAAGGAGCTCCTAGAT

GCATCAGACCAAATACTGAAGACTGTTAAAGGGGTACCTTTCAGTGGGAATATGTACATAGCACTGGCAGCTTGCATAGG

AGTATCCCTGGGCCTTGTCACATTAATATGCTGCTGTAAGGGGAGGTGTAAGAACAAGGAAATCCCTATCTCCAAAATCA

ACCCAGGGCTCAAACCCGACCTGACCGGGACCTCAAAGTCGTACGTAAGATCACTGTAGTCAGAATCACCTGGATCATCT

GGCATCGCACACATACATGCACGACACAGGCAGTCCGAGGACGCAAGAAACCCAGCCTCCGGTCACCCACCCGACCCCAC

TCCACGCTCCACCACACATTAGTCATCAAACAAAACTTAGGACGAAAGGTCAATCACCATGTCCGCACAAAGGGAGAGGA

TCAATGCCTTCTACAAAGACAATCCTCACAATAAGAACCATAGGGTGATCCTGGATAGAGAACGCTTGGTCATTGAAAGA

CCCTACATCTTGCTTGGAGTCCTGCTGGTAATGTTCCTGAGTCTAATCGGACTGCTGGCCATTGCAGGGATCAGGCTTCA

CCGGGCCACCGTTGGAACTTCAGAGATCCAGAGTCGGCTGAATACCAATATTAAGTTGACCGAATCTATTGATCACCAGA

CTAAGGATGTCTTAACTCCCCTTTTTAAAATCATTGGCGATGAAGTCGGCATCAGAATTCCACAGAAATTCAGTGATCTT

GTCAAGTTCATCTCCGATAAGATTAAATTCCTCAACCCTGATAGAGAGTATGATTTCAGAGATCTCCGGTGGTGTATGAA

TCCCCCCGAGAGAGTCAAAATTAATTTTGATCAGTTTTGTGAGTACAAGGCTGCGGTTAAGTCAATTGAACATATATTTG

AGTCACCACTCAACAAGTCAAAAAAGCTGCAATCTTTGACTCTCGGGCCCGGAACAGGCTGTCTAGGCAGGACAGTAACA

AGAGCCCATTTCTCAGAACTTACAATGACCTTAATGGACCTGGATCTAGAGATGAAGCACAACGTGTCCTCAGTGTTTAC

CGTAGTTGAAGAGGGATTATTCGGAAGAACATATACCGTCTGGAGATCCGATGCCAGGGATCCGAGCACCGATCTAGGTA

TCGGCCATTTTTTAAGAGTCTTCGAGATTGGACTGGTAAGAGATCTTGGGCTGGGTCCCCCTGTTTTTCATATGACCAAC

TATCTCACGGTGAACATGAGTGATGACTATCGGAGATGTCTTTTAGCGGTAGGGGAGTTGAAGTTGACAGCCCTATGCAC

CTCATCTGAGACTGTGACACTGAGTGAGAGAGGAGTTCCAAGGAGGGAACCTCTTGTGGTTGTGATACTTAATCTAGCTG

GACCCACTCTAGGGGGAGAGCTATACAGTGTCTTGCCTACCTCTGATCTCATGGTGGAGAAACTCTATTTGTCTTCACAT

AGAGGGATCATCAAAGATGATGAGGCCAATTGGGTAGTGCCGTCTACCGATGTTCGTGATCTTCAGAACAAAGGTGAATG

TCTGGTGGAAGCATGCAAGACTCGACCTCCTTCATTTTGCAATGGCACAGGATCAGGCCCGTGGTCAGAAGGGAGAATCC

CTGCCTACGGGGTGATCAGGGTCAGTCTTGACTTAGCGAGTGACCCGGATGTAGTTATCACTTCAGTGTTTGGCCCACTG

ATACCTCACCTATCCGGCATGGATCTTTACAACAACCCGTTTTCAAAAGCTGTATGGTTGGCTGTACCACCTTATGAGCA

GTCATTTCTAGGAATGATAAATACAATTGGATTCCCTAACAGAGCAGAGGTTATGCCGCACATTTTGACCACAGAGATCA

GAGGCCCTCGGGGTCGTTGCCATGTTCCCATAGAATTGTCCCGCCGGGTTGATGACGATATCAAGATCGGGTCCAACATG

GTCATATTGCCGACGATGGACCTGAGGTATATTACAGCCACTTATGATGTTTCCAGGAGCGAGCATGCAATCGTGTACTA

TATCTATGACACGAGTCGCTCATCATCTTACTTCTACCCAGTTCGACTGAATTTCAAAGGCAATCCTCTCTCTTTGAGGA

TAGAGTGTTTCCCCTGGCGTCATAAGGTGTGGTGCTACCATGATTGTCTTATATACAACACCATAACAGGTGAAGAGGTC

CATACGAGAGGGCTGACCGGCATAGAGGTAACATGCAATCCAGTCTGAGCAGAGCTACGACCGTCACTCAGGCAGTCCCT

TGAGTCGCCACCGAGTCCAAGCAGCACAGCCTGGGACAATCAACGGCACAGCCCAGCCAACAATGTTATAAAAAACTTAG

GAGCCAGGGTTGTAGGGGCCATGGACTCTCTATCAGTCAATCAGGTCTTGTACCCTGAGGTTCATCTGGACAGCCCTATT

GTCACAAACAAACTAGTTGCCATCCTTGAGTACTCGGGTATCGACCATAACTATGTTCTTGAAGACCAGACTCTTGTCAA

GAATATTAGGTATAGGCTGGGGTGCGGTTTTTCAAATCAAATGATCATCAATAACAGGGGGGTGGGTGAAACAGTCAATT

CTAAACTTAAAAGTTACCCCCATAATCGTCATATCATATACCCGGATTGCAATAAGGAGTTGTTTTGTATCAAGGATAGC

TGCATATCTAAGAAGCTCTCGGAGCTATTCAAGAAGGGTAATTCCTTGTACTCTAAGATAAGTCACCAGGTACTGGATTG

TCTTAAGAGAGTCAATGGGAAATTAGGTCTGGGCACAGATCTTACCCATGGTCTGAAGGAGGGTATCCTTGACCTAGGGT

TGCACATGCATAGCTCTCAATGGTTCGAGACTTTCCTGTTCTGGTTCACTATCAAGACAGAGATGAGATCAATGATCAAA

GAACAGTCCCATATATGCCACAAGAGGAGGTATAACCCGACTTTTGTGTCGGGGGATGCATTCGAGGTGCTCGTATCGCG

AGACCTCGTTGTGATAATTGATAAGAATACCCAGTATGTCTTCTACTTAACATTTGAGCTGGTCCTTATGTATTGTGATG

TCATAGAGGGTAGACTTATGACGGAGACAGCCATGGCCATAGACCAGAGATATTCAGAACTTCTAAGCCGGGTCAGATAC

TTRTGGGATCTTATTGATGGGTTCTTCCCAACACTAGGCAACACCACATACCAAATTGTTGCTCTGCTTGAACCATTATC

GTTGGCTTATCTTCAACTTCAGGATGTCACTCTGGAGTTAAGAGGTGCTTTTTTAGACCATTGCTTCAAAGAACTCTATG

AGATACTGGAACATTGCGGCATTGACACAGCAGGCACTTACAATTCCATCACTGAAGGATTGGATTACGTATTCATCACC

CACGATATACATCTAACTGGGGAGATTTTTTCATTTTTCCGGAGTTTCGGACACCCCCGTCTAGAAGCAGTCACCGCTGC

GGAAAATGTCAGAAAACATATGAACCAACCGAAGGTAATCAGTTATGAGACTATGATGAAAGGCCATGCGGTATTTTGCG

GGATAATCATAAATGGTTTTCGGGACCGACATGGCGGCAGCTGGCCCCCTGTTGCACTGCCAGAACATGCCTCTGCTGCG

ATCCGGAATGCGCAGGCATCAGGTGAGGGACTGACCCATGACCTGTGTATAGACAACTGGAAATCCTTTGTTGGATTCAA

ATTTGGCTGCTTTATGCCACTCAGCCTAGATAGTGATTTGACCATGTATCTCAAAGACAAGGCATTGGCTGCACTGAAGA

ATGAGTGGGATTCAGTTTACCCGAAAGAATACCTCCGTTATAACCCACCTAGAGGGACAGAGTCAAGGCGATTGGTAGAG

GTATTCCTGAATGACTCCAGCTTTGATCCTTATAACATGATAATGTACGTGGTGAATGGCTCCTACCTTAAAGACCCTGA

GTTTAACCTCTCATACAGCCTAAAAGAGAAGGAGATAAAGGAGACAGGGCGGTTGTTTGCCAAAATGACCTATAAGATGC

GGGCCTGTCAGGTAATCGCTGAAAATCTGATATCGAATGGTGTTGGGAAGTATTTCCGAGACAATGGGATGGCAAAGGAC

GAGCATGACCTAACAAAAGCCCTTCACACCCTGGCAGTCTCAGGTGTTCCCAAAAACAACAAAGATAACCACCGAGGTGG

GCCTCCCAGAAGGACAACAAACCGAGGGGTGAGATCAAGCCAAGGCACCAAAACACAAGATAGAGACAAGGTTCAAGGGG

GACCTATGTACAACTATTTGCGATGCCAACCGATCAGCCTTGATCAGGGTGAGTCATACGAGACTGTTAGTGCATTCATC

ACCGCTGACCTTAAGAAGTATTGCCTGAATTGGAGATACGAGACAATCAGCATATTTGCACAGAGGCTGAATGAAATATA

TGGACTACCATCCTTCTTTCAATGGTTACACAGGATATTGGAAAAATCCGTACTCTACGTCAGTGACCCACATTGTCCTC

CCGATCTAGATAATCATATCCCTCTGGACAGTGTCCCCAATGCCCAAATATTCATCAAGTACCCAATGGGCGGAATAGAA

GGTTATTGCCAAAAACTATGGACAATCAGCACTATACCATACTTGTATCTGGCAGCCTATGAGAGCGGAGTAAGAATCGC

CTCACTGGTGCAGGGTGACAATCAGACAATCGCAGTGACAAAAAGAGTTCCAAGTTCTTGGCCTTATTCACTAAAAAAGA

GGGAGGCCTCCAAGGCCGCTCAAAATTACTTCGTAGTCTTAAGGCAAAGATTGCACGATGTAGGTCATCACTTAAAGGCT

AATGAGACCATAGTATCTTCTCACTTTTTTGTATATTCCAAAGGGATTTATTATGACGGCCTGCTAGTCTCACAATCACT

AAAGAGCATCGCCAGATGTGTCTTCTGGTCCGAGACTATAGTGGATGAAACCAGAGCGGCGTGCAGTAATATTGCAACAA

CTGTCGCTAAGAGTATAGAGAGGGGTTATGATAGGTACCTTGCATATTCTTTGAATATCCTCAAGATTTTCCAACAGATC

CTTATATCCCTTAACTTCACTATTAACACAACAATGACTCAGGATGTCGTGGCACCGATCATCGAGAACGGTGATTTGCT

GATAAGGATGGCACTCTTGCCAGCACCCATCGGGGGTCTCAATTATCTTAACATGAGCAGGTTATTTGTGAGAAATATCG

GTGACCCGGTCACCTCCTCTATAGCCGACCTGAAGAGGATGATAGATGCCGGGCTAATGCCAGAAGAAACATTGCATCAA

GTGATGACCCAGACCCCGGGAGAATCATCCTACCTAGATTGGGCAAGTGACCCTTATTCTGCCAACCTACCCTGCGTACA

GAGTATCACTCGCCTTCTCAAAAACATCACTGCACGATATATTTTAATCAGCAGCCCAAACCCGATGCTGAAAGGGTTAT

TTCATGAGGGGAGTAGAGATGAAGACGAGGAGCTTGCAAGTTTCCTAATGGATCGGCATATAATTGTTCCAAGAGCTGCA

CATGAAATCTTAGACCATAGTATAACCGGAGCAAGAGAAGCTATAGCCGGGATGTTGGACACCACCAAGGGTCTGATTAG

AACAAGTATGAAACGGGGCGGCCTCACCCCTCGAGTCTTAGCCCGCCTTTCCAATTATGATTATGAACAATTCCGATCCG

GGATAACACTATTAACAAAGAAAGGGCAGTGTTATCTCATTGACAAGGACTCRTGCTCGGTGCAGCTCGCTATAGCTCTG

AGGGGCCACATGTGGGCTAAGTTGGCTCGCGGGAGACCAATCTATGGGTTGGAGGTGCCTGATGTATTAGAATCGATGAA

CGGCTACCTTATCAAACGCCACGAGTCGTGTGCTATCTGTGAAACGGGCTCAAGTCACTACGGATGGTTTTTCGTCCCTG

CAGGGTGCCAGCTTGACGATGTATCTAGAGAGACTTCGGCTCTTCGTGTACCTTATGTCGGATCAACAACTGAGGAAAGG

ACAGACATGAAACTTGCATTTGTTAGGTCTCCAAGTAGATCCCTCAAATCAGCAGTTAGAATTGCAACAGTTTACTCATG

GGCTTACGGGGATGATGAGAAATCATGGAGTGAAGCCTGGATGCTAGCTAGGCAGAGAGCCAATATCACCTTAGATGAAT

TGAGAATGATCACTCCAGTCTCCACATCCACCAACTTAGCCCACCGGTTGAGGGATCGGAGTACCCAGGTGAAATACTCG

GGGACATCCCTTGTGAGGGTTGCAAGATACACAACCATCTCCAATGATAACTTGTCGTTTGTGATATCTGAGAAAAAAGT

GGATACCAACTTCATTTATCAGCAAGGGATGTTGCTTGGTCTTGGGATCCTTGAGAACCTCTTTAGGCTAGAGGCCACCA

CCGGAGTATCCAACACAGTGCTACACCTGCACGTGGAAACAGAATGTTGTGTTGTACCTATGGTGGATCACCCAAGGATA

CCGAGTCTCCGTAATATAAAAGTTACGGATGAGCTATGCACAAACCCTTTGATCTACGACAGGTCCCCCATCATAGAGCA

CGATGCAACCCGGCTATATTCACAGAGCCACAGGAGACAYTTGGTGGAGTTTGTTACCTGGTCAACAAGCCAGCTTTACC

ATATACTGGCTAAGTCTACAGCAATGTCCATGATTGAATTGATCACGAGATTCGAGAAAGATCACATGAATGAAATAGCC

GCCCTGATTGGCGATGACGACATCAACAGCTTCATCACAGAATTTTTGCTAGTGGAGCCTAGATTATTTATAGTTTACCT

TGGTCAGTGTGCTGCCATCAATTGGGCCTTTGATATACATTATCATCGGCCCTCGGGCAAGTACCAGATGGGGGAACTTC

TCTACTCTCTGCTCTCTCGGATGAGCAAAGGAGTATATAAGATCTTTACTAATGCTTTGAGCCACCCCAAAGTTTACAAG

AAATTTTGGCGAAGTGGCATCATTGAGCCGATTCATGGCCCATCCTTGGATACACAGAATTTACATGTCACTGTCTGTGA

CATGATATATGGGTCCTATGTCACCTATTTGGATCTTTTGCTGAATGATGAGCTAGATGATTACCCGTATTTGCTCTGCG

AGAGTGATGAGGACGTGGTCACAGACAGGTTCGACAACATTCAAGCCAAACATCTCTGTGTATTGGCCGATGTATATTGC

AGCTCCAAGAGGTGTCCTTCAATAATCGGGATGTCTCCTATAGAAAAATGTACCATTCTCACACATTACATCAAGGGAGA

ATCAGTACAATCCCCGTCTGGGACCTCATGGAACACTGATCCCCTTGTAGTAGATCATTACTCATGCTCTCTGACCTACC

TTCGCCGCGGTTCCATCAAACAAATCAGGTTGAGGGTGGATCCTGGGTTTGTATTCGAGGCGTTGACAGACGTCGACTTT

AAACAACCTCGCAAAGCTAAGTTAGATATATCGGTCGTTGGGTTGACTGATTTCTCTCCCCCTTACGACAACGTCGGTGA

TTTTTTAGGGACTATCAACACATTGAGGCACGATCTGCCTGTCACCGGAACCGGGGTCTCGAACTATGAAGTCCACGCTT

ATCGTAGAATTGGCCTGAATTCATCAGCATGTTATAAAGCCGTAGAGATCTCCACGTTAATCAAGCCATCCTTAGAAGTC

GGAGAGCATGGATTGTTCTTAGGAGAAGGTTCCGGTTCAATGCTGGCTGCGTACAAAGAAGTTCTTAAATTAGCAAATTG

TTACTACAACAGCGGAGTAACAGCGGAGGGCAGAGCCGGACAGAGGGAAATATCTCCCTATCCTTCAGAGATGAGCCTAG

TAGAGAATCAAATGGGGATAGAGCGGAGTGTTAAAGTGCTGTTCAACGGGAAACCTGAAGTAACTTGGGTGGGGACCACA

GATTGCTACAAGTACATAATCAGTAACATCCAAACCTCTAGTCTGGGTTTCATACATTCAGATATTGAGACGCTCCCAAC

CAAAGATGCTGTTGAGAAATTGGAGGAATTTGCCTCTATCCTATCCTTATCCTTGATTTTGGGGAAAATCGGCTCTATTA

CAGTTGTCAAAATTATGCCCATTAGCGGGGATTTTACCCAAGGCTTCATAGCCTATGCCATCCAATATTTTAGGGAGAGC

CTGCTTGCCTACCCGAGATATAGTAACTTCATCTCGACTGAGTGTTACCTTATTATGATAGGATTAAAGGCCAATCGATT

GATAAACCCAGAAGCCATTAAGCAAAGCATAATCAGAGCGGGGGTCAGGACTGCACCAGGACTTGTGAGCCATGTATTAT

CAGAGAAACAAAAAGGTTGTATTCAATCTTTCTTGGGTGATCCTTATATCCAAGGAGACTTCAATAAGCACCTTAAATCT

CTAACCTCTATTGAGAAAATCCTGGTAAATTGTGGTCTCTCGATCAATGGCATAAAAATCTGTAGAGATCTAATCCACCA

TGATATCGCCTCCGGTCCAGACGGCCTTATGAGCTCTACAATTATTTTATATAGGGAATTGGCTCATTTCAAAGATAATA

TAAGGAGTCAGCACGGAATGTTCCACCCCTACCCGGTACTGGCCAACAGCAGGCAACGTGAATTAATCCTTCGAATAGCC

AAGAAATTCTGGGGGTATGTCTTGCTATATTCTGATGACCCGGCACTAATCAGACAAACAATCAAGAACTTGAAACGGAA

TCATCTAACCTTTGACTTACATAGTAACCCATTTATTAAAGGGCTATCCAAGGCTGAGAAACTGCTGGTGCGGACGAGTT

CACTAAGAAGAGAATGGTTGTTCACTCTCGAAACGAAAGAAGTAAAAGAGTGGTTCAAATTGGTAGGTTACAGTGCACTC

ATCAGAGGCTAGTCAGCTATGCATCTGCCCCCTTCTCCTCCGCCATGAGACCCCACTGGCGATCCAGAAGATTAAAGAAA

ACTACATATTGGATAAGTATCTATTCCCAGCTTTGTCTGGT

>MF443336.1/goat/China/2014-04-01

ACCAAACAAAGTTGGGTAAGGATAGATCTTATAATAACTATGGACTGGCAAACTTAGGAGTAAAGATCCTACTGTCGGGG

GGAGGAGGAGGAGCAAGATCTTTGATTATGGCGACTCTCCTTAAAAGCTTAGCATTGTTCAAAAGGAACAAAGACAAAGC

GCCGACAGCATCAGGTTCAGGAGGGGCCATCCGGGGGATTAAGAATGTTATCATAGTCCCGATTCCCGGAGACTCGTCCA

TCACTACCCGTTCAAGACTGCTCGACAGGCTTGTCAGATTGGCCGGAGATCCTGACATCAACGGGTCAAAGCTGACCGGG

GTGATGATCAGCATGTTATCCTTGTTCGTAGAGTCACCCGGGCAATTGATACAGCGAATCACAGATGATCCGGATGTTAG

TATCCGCCTTGTTGAGGTAGTTCAAAGTACTAGGTCCCAGTCCGGGTTGACCTTTGCATCACGTGGTGCTGATTTAGACA

ACGAGGCAGACATGTATTTTTCAACTGAGGGGCCCTCGAGTGGAGGTAAGAAAAGGATCAACTGGTTTGAGAACAGAGAA

ATAATAGACATAGAGGTGCAGGATCCAGAAGAGTTCAATATGTTGTTAGCCTCCATACTAGCACAAGTCTGGATCCTTCT

GGCCAAGGCTGTTACGGCACCAGATACGGCAGCTGACTCAGAACTGAGAAGGTGGGTTAAATACACACAACAAAGGAGAG

TGATTGGGGAATTTCGCCTTGACAAGGGGTGGCTGGACGCAGTCCGCAACAGAATTGCAGAAGATCTATCACTCCGGCGG

TTCATGGTATCTCTCATACTTGACATCAAGAGGACCCCAGGCAACAAGCCAAGGATTGCAGAAATGATCTGCGACATTGA

CAACTATATTGTCGAAGCAGGACTCGCCAGTTTCATCCTTACTATCAAATTTGGTATCGAAACCATGTATCCTGCATTAG

GGCTCCACGAGTTTGCCGGAGAATTGTCCACTATTGAGTCCTTGATGAACTTGTATCAACAGCTAGGAGAGGTTGCACCC

TACATGGTAATTCTAGAGAATTCAATTCAGAACAAGTTTAGTGCAGGAGCTTATCCCCTCCTCTGGAGCTATGCGATGGG

TGTCGGAGTCGAGCTGGAGAACTCAATGGGGGGCTTGAACTTTGGCAGGTCATATTTTGACCCTGCCTATTTTCGTCTCG

GACAGGAGATGGTCAGAAGATCTGCAGGAAAGGTCAGCTCTGTGATTGCGGCTGAGCTCGGCATCACAGCAGAGGAAGCT

AAACTAGTCTCGGAAATCGCCTCGCAGGCTGGGGACGAAAGAACCGCCAGAGGGACTGGGCCTCGACAGGCGCAGGTTTC

CTTCCTTCAGCACCAAACAGGAGGGGGAGAGTCGTCCGCACCAGCGACCAGAGAAGGGGTCAAAGCTGCGATCCCAAACG

GAGCTGAAGAAAGGGACAGAAAGCAAACACGCCCAGGAAGGCCCAGAGGAGAGACCTCCGGCCAACTGCTCCTGGACATC

ATGCCAGAGGATGAGATCTTGCGAGAGTCTGGTCAAAACCCTCGTGAGGCTCAAAGATCGGCCGAGGCACTCTTCAGGCT

GCAGGCCATGGCCAAGATTCTGGAGGACCAGGAGGAGGGAGAAGACAACAGTCAGGTCTACAACGACAAGGATCTCCTCG

GCTAAGCAGACGCACCCTCTGTCGAAATCAGTGACAAGACATCTCTTACCAGTATTATAAAAAACTTAGGACCCAGGTCC

AAGCAAGCACACATCGACACTCCAACCAGTCGAGCGGAGACCACCGATGGCAGAAGAACAAGCATACCATGTCAACAAGG

GGCTGGAATGTATCAAGTCTCTCAAAGCCTCACCCCCGGATCTATCCACCATCAGAGATACCATCGAGAGCTGGAGAGAG

GGGCTTAGCCCCTCGGGCCGTGCAACACCGAACCCTGATACGTCCGAGGGAGATCATCAGAATATCAACCAATCATGCTC

ACCAGCAATCGGACCAAACAAAGTCTACTTGTCTCCTGAAGATAATCTCGGATTTAGAGAGATCACTGGCAACGACTGTG

AGGCTGGGCTCGGAGGGGTCCAGAGAGAAGGATCCAACTCTCAAGTACAGCGTTACCATGTTTATAGTCACGGGGGTGAA

GAGATTGAAGGACTCGAGGATGCTGACTCTCTCGTGGTTCAAGCAGATCCTCCAGTTGCTAACGTCTTCAATGGAGGAGA

GGATGGATCTGACGACAGCGATGTGGACTCTGGCCCAGATGATCCCGGCAGAGATACTCTATATGACCGGGGATCTGTTG

CCGGCAATGGTATCGCTAGGTCCACAGATGTCGAAAAACTAGAAGGTGCTGATATTCAAGAAGTTCTTAACTCCCAGAAA

GGCAAAGGAGGAAGATTCCAGGGCGGGAAAACCTTGCGAGTCCCGGAAATACCCGATGTCAAGCACTCCAGACCATCAGC

CCAATCAATTAAAAAGGGCACAGACGGGAACTCAGTCTCATCTGGAACGGTGATAGAGTGTTTATCGATAAGTGGTGCAA

CCCAAACTGTGCCAGAGTCAAGATGGGAGTCATCAGAGCAAAATGCGTCTGTGGGGAGTGTCCTCAAGTCTGCGAGGAGT

GCAAAGACGATCCAGGGGTCGACACAAGAATCTGGTACCATAGCATCACTGACTCAGCCTAAAGAGAATGACTCCGAGTA

TGAGTATGAGGATGACCTATTTACAGAGATTCAGGACATCCGTGCAAGCATTGCCAAGATCCATGATGACAATAAAACTA

TCCTCTCAAAGCTTGATTCTATACTGTTATTGAAAGGGGAAGTCGACACTATCAAGAAACAAATCAGCAAGCAGAATATA

AGTATATCCACCATTGAGGGCCATCTCTCCAGTATAATGATAGCCATCCCTGGCTTTGGGAAGGACATCAAGGACCCAAC

ATCCGAAGTCGAGTTGAACCCAGATCTAAGACCTATAATAAGCCGTGACTCTGGCAGAGCTCTCGCGGAGGTCCTCAAGA

AACCCGCTGTGGATAGGTCTCAGAAAATTGGAACCAAAGCCAACTCCAGCTCAAAGGGTCAGCTTCTTAAGGATCTCCAG

CTAAAACCTGTCGACAAGCAGGCAAGCTCTGCAATCGGGTTTGTCCCGTCCGACCATGAATCATCCAGAAATGTCATCCG

CTCCATAATCAAGTCGAGCAAGCTAAATATTGATCACAAGGACTATCTTCTAGATTTACTAAATGATGTGAAAGGCTCCA

AGGATCTTAAGGAATTTCACAAGATGCTAACAGCAATTCTCGCCAAGCACCCGTAACACATCCTCCAGTCATCATCTCAT

ACTCGACTAAAAACATCCTTTCAATCAGGCTATTACAAAAAACTTAGGAGCAAGGGCAACTGAGCTTCGCAGACAGGATG

ACCGAGATCTACGACTTCGATAAATCAGCATGGGATGTCAAAGGGTCAATTGCCCGCATAGAACCCACCACCTACCACGA

CGGCCGACTGATACCCCAGGTGAGGGTCATCGATCCTGGTCTGGGAGACAGAAAAGATGAGTGCTTTATGTACCTGTTTC

TCCTAGGAGTGATTGAGGATAACGACCCCCTGTCTCCCCCCGTCGGAAGAACCTTTGGCTCTCTACCTCTAGGGGTTGGT

AGGTCAACTGCCAAGCCAGAAGAACTACTAAGGGAGGCCACAGAATTAGATATAGTGGTGAGGCGCACTGCAGGAGTAAA

TGAGAAACTGGTATTTTACAACAACACTCCGCTGTCCTTGTTAACGCCCTGGAAGAAAGTTCTGACAACCGGAAGTGTGT

TTAGCGCTAACCAGGTTTGCAATGCAGTCAACCTAGTCCCGCTTGATACTCCCCAGAGATTCAGGGTTGTGTACATGAGC

ATAACTAGATTATCAGACAATGGTTACTATAGTGTGCCCAGAAGAATGCTGGAGTTCCGCTCGGCCAATGCAGTAGCCTT

CAACATCTTGGTTACACTAAGAATTGAAAATGGCACAAACCCTAGAAGATACATAGTCGGCTCATGGGAGAATTCAGAGG

TCACATTTATGGTACACGTGGGCAACTTTAGGAGAAAGAAGAACGAAGTATACTCTGCTGATTACTGCAAAATGAAGATT

GAAAAGATGGGTTTAGTTTTTGCCCTGGGTGGAATAGGCGGAACAAGTCTCCATATTAGAAGCACCGGGAAAATGAGCAA

AACCCTCCATGCACAGCTGGGGTTCAAGAAAATCTTATGTTACCCCCTAATGGATGTCAATGAGGATCTTAACCGATATC

TCTGGCGGGCAGAGTGCCGAATAGTCAAAATCCAAGCTGTCTTACAGCCATCAGTACCCCAAGAGTTCCGTGTCTACGAT

GATGTCATCATCAACGATGATCAAGGCTTGTTCAAGATCTTGTAGTTCATTTGCAAAGGAACGAACCCACCCCCAGGAAG

GGGCCCCACCCTCCCCCG-CCCAAAAACACACTCCAAGGCTCCCCGGCACAGCGCCAACACAGCTACTCCCGCGATGGAA

CGAGGTCGGCC-CCCGAACAAATCCGGTCCAGGGGGGGGGGCCCCCCCCCTGACCAACCGGGCGCCGACCGAGCAGGCCG

GCGAGAAGCCCCCAAGAGGCCCCGACGAAGACCGAGCACCCCATCCCCCC-AAAAACGTCCCCC-AACACGCCCGAGGCT

AGCCACCGGACACCCCCAGACCCGCCACCGGCGAGACAGAACCCCAAGGGGGGACCCTCCAAACCAAAGGAATCACCGGG

GTAGGCTCCGCCGAACCAGCACAGACCCGACCCAAGCAAGGCGGACCCCCAACACGAAACTGCCCAGCACACCCTACTCA

TCATCAAACAAAACTTAGGGGCCAAGTCCATACGAGAGGCCCTAGTCCGTCAGGCGGAGGGACGCAGAAAGGAAGGAGAC

ACGACCGTCCCAGAAGGAGGACACCCAACCACCGAAACGCCCGCCCCCAGGGAAGGAG--CCA-CCCCCACGCAGCGAAG

GAGCGGGTGGGAGACCTCGGGGAGGGCCCGACCGGGACCCGGGGGCCGGCGGCGCCGGCGGTGAGCAGGCCGGGCGCATA

CCTGACCCCACCGGAGGATCCGAGGAGGGCCCGACCCCGCGTGGACCAAACGGCAGGGAGCCCAACCCCACCAAAAAAAG

AGGCGAGGCCCCCCCTCCCTCCCCCCCCGCACACAACCCGCAGACAGACCGAGGGAGGGCCATCCCCCAGCGAGACAGGC

GGCTCCCTCGCAGCCGAGCCGCAGCTCCCCAGAATAGGACACCCCCGCACACTCCGAACACCCCCTCGCCCCCCGCCACC

GGGACAGGGGCTCCACCACCACAGACCGGAGCCACCGACAGCCAGCCAGAGGTCCCCTCCAGTCCCGCCCGGCGCCCCCC

CGAAGAGGCCCCGGCAAGGCCCCCCAGCCCGACGTCCCCCGCACGCACCGAAAGAAGGACCCGCCCGAGAACGTGCGGCA

CATCCATGCGCAAACATCATGACGCGGGTCGCAATTTTGACATTTCTGTTTCTTTTCCCAAATGTTGTTGCGTGTCAGAT

TCACTGGGGCAATCTATCCAAGATCGGGATTGTAGGAACAGGGAGTGCCAGCTACAAGGTGATGACTAGGCCAAGCCACC

AGACTCTGGTTATAAAGTTAATGCCAAATATAACGGCCATCGACAATTGTACAAAGTCAGAGATTGCAGAGTACAAGAGA

TTGCTGATCACAGTGTTAAAGCCTGTAGAGGATGCTCTGTCGGTGATAACCAAGAATGTAAGACCAATTCAAACTCTAAC

ACCTGGGCGTAGAACCCGCCGTTTTGCTGGAGCTGTTCTGGCCGGGGTAGCACTTGGAGTTGCGACAGCCGCTCAGATAA

CTGCAGGAGTCGCCCTTCATCAATCATTGATGAACTCCCAAGCAATTGAGAGTTTAAAAACCAGTCTTGAGATGTCGAAT

CAGGCAATAGAAGAAATCAGACTTGCAAATAAGGAGACCATACTGGCAGTACAGGGCGTCCAGGATTATATCAACAATGA

GCTCGTCCCTTCTGTTCATAGAATGTCATGCGAGCTGGTAGGTCACAAGCTCGGCCTCAAGCTCCTTAGGTACTACACCG

AGATCCTGTCCATATTCGGGCCCAGTCTTCGAGACCCGATATCTGCCGAAATATCAATCCAGGCACTTAGTTATGCATTA

GGCGGAGACATTAATAAAATCCTGGACAAGCTTGGGTATAGCGGTGGGGATTTCCTTGCCATCCTAGAAAGCAAGGGAAT

AAAGGCCCGGGTTACATATGTGGACACAAGAGATTACTTTATAATCCTTAGCATCGCCTACCCAACCTTATCTGAGATCA

AGGGAGTGATAGTTCACAAGATAGAAGCTATAACATACAACATTGGGGCACAGGAGTGGTATACTACTATCCCTAAATAT

GTAGCCACTCAGGGGTATCTGATATCGAACTTTGATGAGACGTCATGCGTATTCACTCCAGAGGGGACAGTTTGCAGCCA

GAATGCGTTGTACCCAATGAGCCCATTGCTTCAGGAATGTTTCAGGGGGTCAACAAAATCGTGCGCCAGAACCTTAGTTT

CAGGGACCATAAGTAATAGATTTATCCTATCAAAAGGGAACCTGATTGCAAATTGTGCGTCAGTTTTGTGCAAATGTTAC

ACAACAGAGACAGTTATCAGCCAAGATCCTGACAAACTACTAACTGTTGTAGCATCCGACAAGTGTCCTGTAGTTGAGGT

GGATGGAGTGACAATACAGGTCGGCAGTCGAGAGTATCCGGATTCTGTATACTTACACAAAATAGACTTAGGTCCAGCCA

TCTCCCTAGAAAAACTGGATGTAGGCACCAATTTAGGCAATGCAGTCACAAGACTGGAGAATGCAAAGGAGCTCCTAGAT

GCATCAGACCAAATACTGAAGACTGTTAAAGGGGTACCTTTCAGTGGGAATATGTACATAGCACTGGCAGCTTGCATAGG

AGTATCCCTGGGCCTTGTCACATTAATATGCTGCTGTAAGGGGAGGTGTAAGAACAAGGAAATCCCTATCTCCAAAATCA

ACCCAGGGCTCAAACCCGACCTGACCGGGACCTCAAAGTCGTACGTAAGATCACTGTAGTCAGAATCACCTGGATCATCT

GGCATCGCACACATACATGCACGACACAGGCAGTCCGAGGACGCAAGAAACCCAGCCTCCGGTCACCCACCCGACCCCAC

TCCACGCTCCACCACACATTAGTCATCAAACAAAACTTAGGACGAAAGGTCAATCACCATGTCCGCACAAAGGGAGAGGA

TCAATGCCTTCTACAAAGACAATCCTCACAATAAGAACCATAGGGTGATCCTGGATAGAGAACGCTTGGTCATTGAAAGA

CCCTACATCTTGCTTGGAGTCCTGCTGGTAATGTTCCTGAGTCTAATCGGACTGCTGGCCATTGCAGGGATCAGGCTTCA

CCGGGCCACCGTTGGAACTTCAGAGATCCAGAGTCGGCTGAATACCAATATTAAGTTGACCGAATCTATTGATCACCAGA

CTAAGGATGTCTTAACTCCCCTTTTTAAAATCATTGGCGATGAAGTCGGCATCAGAATTCCACAGAAATTCAGTGATCTT

GTCAAGTTCATCTCCGATAAGATTAAATTCCTCAACCCTGATAGAGAGTATGATTTCAGAGATCTCCGGTGGTGTATGAA

TCCCCCCGAGAGAGTCAAAATTAATTTTGATCAGTTTTGTGAGTACAAGGCTGCGGTTAAGTCAATTGAACATATATTTG

AGTCACCACTCAACAAGTCAAAAAAGCTGCAATCTTTGACTCTCGGGCCCGGAACAGGCTGTCTAGGCAGGACAGTAACA

AGAGCCCATTTCTCAGAACTTACAATGACCTTAATGGACCTGGATCTAGAGATGAAGCACAACGTGTCCTCAGTGTTTAC

CGTAGTTGAAGAGGGATTATTCGGAAGAACATATACCGTCTGGAGATCCGATGCCAGGGATCCGAGCACCGATCTAGGTA

TCGGCCATTTTTTAAGAGTCTTCGAGATTGGACTGGTAAGAGATCTTGGGCTGGGTCCCCCTGTTTTTCATATGACCAAC

TATCTCACGGTGAACATGAGTGATGACTATCGGAGATGTCTTTTAGCGGTAGGGGAGTTGAAGTTGACAGCCCTATGCAC

CTCATCTGAGACTGTGACACTGAGTGAGAGAGGAGTTCCAAGGAGGGAACCTCTTGTGGTTGTGATACTTAATCTAGCTG

GACCCACTCTAGGGGGAGAGCTATACAGTGTCTTGCCTACCTCTGATCTCATGGTGGAGAAACTCTATTTGTCTTCACAT

AGAGGGATCATCAAAGATGATGAGGCCAATTGGGTAGTGCCGTCTACCGATGTTCGTGATCTTCAGAACAAAGGTGAATG

TCTGGTGGAAGCATGCAAGACTCGACCTCCTTCATTTTGCAATGGCACAGGATCAGGCCCGTGGTCAGAAGGGAGAATCC

CTGCCTACGGGGTGATCAGGGTCAGTCTTGACTTAGCGAGTGACCCGGATGTAGTTATCACTTCAGTGTTTGGCCCACTG

ATACCTCACCTATCCGGCATGGATCTTTACAACAACCCGTTTTCAAAAGCTGTATGGTTGGCTGTACCACCTTATGAGCA

GTCATTTCTAGGAATGATAAATACAATTGGATTCCCTAACAGAGCAGAGGTTATGCCGCACATTTTGACCACAGAGATCA

GAGGCCCTCGGGGTCGTTGCCATGTTCCCATAGAATTGTCCCGCCGGGTTGATGACGATATCAAGATCGGGTCCAACATG

GTCATATTGCCGACGATGGACCTGAGGTATATTACAGCCACTTATGATGTTTCCAGGAGCGAGCATGCAATCGTGTACTA

TATCTATGACACGAGTCGCTCATCATCTTACTTCTACCCAGTTCGACTGAATTTCAAAGGCAATCCTCTCTCTTTGAGGA

TAGAGTGTTTCCCCTGGCGTCATAAGGTGTGGTGCTACCATGATTGTCTTATATACAACACCATAACAGGTGAAGAGGTC

CATACGAGAGGGCTGACCGGCATAGAGGTAACATGCAATCCAGTCTGAGCAGAGCTACGACCGTCACTCAGGCAGTCCCT

TGAGTCGCCACCGAGTCCAAGCAGCACAGCCTGGGACAATCAACGGCACAGCCCAGCCAACAATGTTATAAAAAACTTAG

GAGCCAGGGTTGTAGGGGCCATGGACTCTCTATCAGTCAATCAGGTCTTGTACCCTGAGGTTCATCTAGACAGCCCTATT

GTCACAAACAAACTAGTTGCCATCCTTGAGTACTCGGGTATCGACCATAACTATGTTCTTGAAGACCAGACTCTTGTCAA

GAATATTAGGTATAGGCTGGGGTGCGGTTTTTCAAATCAAATGATCATCAATAACAGGGGGGTGGGTGAAACAGTCAATT

CTAAACTTAAAAGTTACCCCCATAATCGTCATATCATATACCCGGATTGCAATAAGGAGTTGTTTTGTATCAAGGATAGC

TGCATATCTAAGAAGCTCTCGGAGCTATTCAAGAAGGGTAATTCCTTGTACTCTAAGATAAGTCACCAGGTACTGGATTG

TCTTAAGAGAGTCAATGGGAAATTAGGTCTGGGCACAGATCTTACCCATGGTCTGAAGGAGGGTATCCTTGACCTAGGGT

TGCACATGCATAGCTCTCAATGGTTCGAGACTTTCCTGTTCTGGTTCACTATCAAGACAGAGATGAGATCAATGATCAAA

GAACAGTCCCATATATGCCACAAGAGGAGGTATAACCCGACTTTTGTGTCGGGGGATGCATTCGAGGTGCTCGTATCGCG

AGACCTCGTTGTGATAATTGATAAGAATACCCAGTATGTCTTCTACTTAACATTTGAGCTGGTCCTTATGTATTGTGATG

TCATAGAGGGTAGACTTATGACGGAGACAGCCATGGCCATAGACCAGAGATATTCAGAACTTCTAAGCCGGGTCAGATAC

TTGTGGGATCTTATTGATGGGTTCTTCCCAACACTAGGCAACACCACATACCAAATTGTTGCTCTGCTTGAACCATTATC

GTTGGCTTATCTTCAACTTCAGGATGTCACTCTGGAGTTAAGAGGTGCTTTTTTAGACCATTGCTTCAAAGAACTCTATG

AGATACTGGAACATTGCGGCATTGACACAGCAGGCACTTACAATTCCATCACTGAAGGATTGGATTACGTATTCATCACC

CACGATATACATCTAACTGGGGAGATTTTTTCATTTTTCCGGAGTTTCGGACACCCCCGTCTAGAAGCAGTCACCGCTGC

GGAAAATGTCAGAAAACATATGAACCAACCGAAGGTAATCAGTTATGAGACTATGATGAAAGGCCATGCGGTATTTTGCG

GGATAATCATAAATGGTTTTCGGGACCGACATGGCGGCAGCTGGCCCCCTGTTGCACTGCCAGAACATGCCTCTGCTGCG

ATCCGGAATGCGCAGGCATCAGGTGAGGGACTGACCCATGACCTGTGTATAGACAACTGGAAATCCTTTGTTGGATTCAA

ATTTGGCTGCTTTATGCCACTCAGCCTAGATAGTGATTTGACCATGTATCTCAAAGACAAGGCATTGGCTGCACTGAAGA

ATGAGTGGGATTCAGTTTACCCGAAAGAATACCTCCGTTATAACCCACCTAGAGGGACAGAGTCAAGGCGATTGGTAGAG

GTATTCCTGAATGACTCCAGCTTTGATCCTTATAACATGATAATGTACGTGGTGAATGGCTCCTACCTTAAAGACCCTGA

GTTTAACCTCTCATACAGCCTAAAAGAGAAGGAGATAAAGGAGACAGGGCGGTTGTTTGCCAAAATGACCTATAAGATGC

GGGCCTGTCAGGTAATCGCTGAAAATCTGATATCGAATGGTGTTGGGAAGTATTTCCGAGACAATGGGATGGCAAAGGAC

GAGCATGACCTAACAAAAGCCCTTCACACCCTGGCAGTCTCAGGTGTTCCCAAAAACAACAAAGATAACCACCGAGGTGG

GCCTCCCAGAAGGACAACAAACCGAGGGGTGAGATCAAGCCAAGGCACCAAAACACAAGATAGAGACAAGGTTCAAGGGG

GACCTATGTACAACTATTTGCGATGCCAACCGATCAGCCCTGATCAGGGTGAGTCATACGAGACTGTTAGTGCATTCATC

ACCGCTGACCTTAAGAAGTATTGCCTGAATTGGAGATACGAGACAATCAGCATATTTGCACAGAGGCTGAATGAAATATA

TGGACTACCATCCTTCTTTCAATGGTTACACAGGATATTGGAAAAATCCGTACTCTACGTCAGTGACCCACATTGTCCCC

CCGATCTAGATAATCATATCCCTCTGGACAGTGTCCCCAATGCCCAAATATTCATCAAGTACCCAATGGGCGGAATAGAA

GGTTATTGCCAAAAACTATGGACAATCAGCACTATACCATACTTGTATCTGGCAGCCTATGAGAGCGGAGTAAGAATCGC

CTCACTGGTGCAGGGTGACAATCAGACAATCGCAGTGACAAAAAGAGTTCCAAGTTCTTGGCCTTATTCACTAAAAAAGA

GGGAGGCCTCCAAGGCCGCTCAAAATTACTTCGTAGTCTTAAGGCAAAGATTGCACGATGTAGGTCATCACTTAAAGGCT

AATGAGACCATAGTATCTTCTCACTTTTTTGTATATTCCAAAGGGATTTATTATGACGGCCTGCTAGTCTCACAATCACT

AAAGAGCATCGCCAGATGTGTCTTCTGGTCCGAGACTATAGTGGATGAAACCAGAGCGGCGTGCAGTAATATTGCAACAA

CTGTCGCTAAGAGTATAGAGAGGGGTTATGATAGGTACCTTGCATATTCTTTGAATATCCTCAAGATTTTCCAACAGATC

CTTATATCCCTTAACTTCACTATTAACACAACAATGACTCAGGATGTCGTGGCACCGATCATCGAGAACGGTGATTTGCT

GATAAGGATGGCACTCTTGCCAGCACCCATCGGGGGTCTCAATTATCTTAACATGAGCAGGTTATTTGTGAGAAATATCG

GTGACCCGGTCACCTCCTCTATAGCCGACCTGAAGAGGATGATAGATGCCGGGCTAATGCCAGAAGAAACATTGCATCAA

GTGATGACCCAGACCCCGGGAGAATCATCCTACCTAGATTGGGCAAGTGACCCTTATTCTGCCAACCTACCCTGCGTACA

GAGTATCACTCGCCTTCTCAAAAACATCACTGCACGATATATTTTAATCAGCAGCCCAAACCCGATGCTGAGAGGGTTAT

TTCATGAGGGGAGTAGAGATGAAGACGAGGAGCTTGCAAGTTTCCTAATGGATCGGCATATAATTGTTCCAAGAGCTGCA

CATGAAATCTTAGACCATAGTATAACCGGAGCAAGAGAAGCTATAGCCGGGATGTTGGACACCACCAAGGGTCTGATTAG

AACAAGTATGAAACGGGGCGGCCTCACCCCTCGAGTCTTAGCCCGCCTTTCCAATTATGATTATGAACAATTCCGATCCG

GGATAACACTATTAACAAAGAAAGGGCAGTGTTATCTCATTGACAAGGACTCGTGCTCGGTGCAGCTCGCTATAGCTCTG

AGGGGCCACATGTGGGCTAAGTTGGCTCGCGGGAGACCAATCTATGGGTTGGAGGTGCCTGATGTATTAGAATCGATGAA

CGGCTACCTTATCAAACGCCACGAGTCGTGTGCTATCTGTGAAACGGGCTCAAGTCACTACGGATGGTTTTTCGTCCCTG

CAGGGTGCCAGCTTGACGATGTATCTAGAGAGACTTCGGCTCTTCGTGTACCTTATGTCGGATCAACAACTGAGGAAAGG

ACAGACATGAAACTTGCATTTGTTAGGTCTCCAAGTAGATCCCTCAAATCAGCAGTTAGAATTGCAACAGTTTACTCATG

GGCTTACGGGGATGATGAGAAATCATGGAGTGAAGCCTGGATGCTAGCTAGGCAGAGAGCCAATATCACCTTAGATGAAT

TGAGAATGATCACTCCAGTCTCCACATCCACCAACTTAGCCCACCGGTTGAGGGATCGGAGTACCCAGGTGAAATACTCG

GGGACATCCCTTGTGAGGGTTGCAAGATACACAACCATCTCCAATGATAACTTGTCGTTTGTGATATCTGAGAAAAAAGT

GGATACCAACTTCATTTATCAGCAAGGGATGTTGCTTGGTCTTGGGATCCTTGAGAACCTCTTTAGGCTAGAGGCCACCA

CCGGAGTATCCAACACAGTGCTACACCTGCACGTGGAAACAGAATGTTGTGTTGTACCTATGGTGGATCACCCAAGGATA

CCGAGTCTCCGTAATATAAAAGTTACGGATGAGCTATGCACAAACCCTTTGATCTACGACAGGTCCCCCATCATAGAGCA

CGATGCAACCCGGCTATATTCACAGAGCCACAGGAGACATTTGGTGGAGTTTGTTACCTGGTCAACAAGCCAGCTTTACC

ATATACTGGCTAAGTCTACAGCAATGTCCATGATTGAATTGATCACGAGGTTCGAGAAAGATCACATGAATGAAATAGCC

GCCCTGATTGGCGATGACGACATCAACAGCTTCATCACAGAATTTTTGCTAGTGGAGCCTAGATTATTTATAGTTTACCT

TGGTCAGTGTGCTGCCATCAATTGGGCCTTTGATATACATTATCATCGGCCCTCGGGCAAGTACCAGATGGGGGAACTTC

TCTACTCTCTGCTCTCTCGGATGAGCAAAGGAGTATATAAGATCTTTACTAATGCTTTGAGCCACCCCAAAGTTTACAAG

AAATTTTGGCGAAGTGGCATCATTGAGCCGATTCATGGCCCATCCTTGGATACACAGAATTTACATGTCACTGTCTGTGA

CATGATATATGGGTCCTATGTCACCTATTTGGATCTTTTGCTGAATGATGAGCTAGATGATTACCCGTATTTGCTCTGCG

AGAGTGATGAGGACGTGGTCACAGACAGGTTCGACAACATTCAAGCCAAACATCTCTGTGTATTGGCCGATGTATATTGC

AGCTCCAAGAGGTGTCCTTCAATAATCGGGATGTCTCCTATAGAAAAATGTACCATTCTCACACATTACATCAAGGGAGA

ATCAGTACAATCCCCGTCTGGGACCTCATGGAACACTGATCCCCTTGTAGTAGATCATTACTCATGCTCTCTGACCTACC

TTCGCCGCGGTTCCATCAAACAAATCAGGTTGAGGGTGGATCCTGGGTTTGTATTCGAGGCGTTGACAGACGTCGACTTT

AAACAACCTCGCAAAGCTAAGTTAGATATATCGGTCGTTGGGTTGACTGATTTCTCTCCCCCTTACGACAACGTCGGTGA

TTTTTTAGGGACTATCAACACATTGAGGCACGATCTGCCTGTCACCGGAACCGGGGTCTCGAACTATGAAGTCCACGCTT

ATCGTAGAATTGGCCTGAATTCATCAGCATGTTATAAAGCCGTAGAGATCTCCACGTTAATCAAGCCATCCTTAGAAGTC

GGAGAGCATGGATTGTTCTTAGGAGAAGGTTCCGGTTCAATGCTGGCTGCGTACAAAGAAGTTCTTAAATTAGCAAATTG

TTACTACAACAGCGGAGTAACAGCGGAGGGCAGAGCCGGACAGAGGGAAATATCTCCCTATCCTTCAGAGATGAGCCTAG

TAGAGAATCAAATGGGGATAGAGCGGAGTGTTAAAGTGCTGTTCAACGGGAAACCTGAAGTAACTTGGGTGGGGACCACA

GATTGCTACAAGTACATAATCAGTAACATCCAAACCTCTAGTCTGGGTTTCATACATTCAGATATTGAGACGCTCCCAAC

CAAAGATGCTGTTGAGAAATTGGAGGAATTTGCCTCTATCCTATCCTTATCCTTGATTTTGGGGAAAATCGGCTCTATTA

CAGTTGTCAAAATTATGCCCATTAGCGGGGATTTTACCCAAGGCTTCATAGCCTATGCCATCCAATATTTTAGGGAGAGC

CTGCTTGCCTACCCGAGATATAGTAACTTCATCTCGACTGAGTGTTACCTTATTATGATAGGATTAAAGGCCAATCGATT

GATAAACCCAGAAGCCATTAAGCAAAGCATAATCAGAGCGGGGGTCAGGACTGCACCAGGACTTGTGAGCCATGTATTAT

CAGAGAAACAAAAAGGTTGTATTCAATCTTTCTTGGGTGATCCTTATATCCAAGGAGACTTCAATAAGCACCTTAAATCT

CTAACCTCTATTGAGAAAATCCTGGTAAATTGTGGTCTCTCGATCAATGGCATAAAAATCTGTAGAGATCTAATCCACCA

TGATATCGCCTCCGGTCCAGACGGCCTTATGAGCTCTACAATTATTTTATATAGGGAATTGGCTCATTTCAAAGATAATA

TAAGGAGTCAGCACGGACTGTTCCACCCCTACCCGGTACTGGCCAACAGCAGGCAACGTGAATTAATCCTTCGAATAGCC

AAGAAATTCTGGGGGTATGTCTTGCTATATTCTGATGACCCGGCACTAATCAGACAAACAATCAAGAACTTGAAACGGAA

TCATCTAACCTTTGACTTACATAGTAACCCATTTATTAAAGGGCTATCCAAGGCTGAGAAACTGCTGGTGCGGACGAGTT

CACTAAGAAGAGAATGGTTGTTCACTCTCGAAACGAAAGAAGTAAAAGAGTGGTTCAAATTGGTAGGTTACAGTGCACTC

ATCAGAGGCTAGTCAGCTATGCATCTGCCCCCTTCTCCTCCGCCATGAGACCCCACTGGCGATCCAGAAGATTAAAGAAA

ACTACATATTGGATAAGTATCTATTCCCAGCTTTGTCTGGT

>MF443335.1/goat/China/2014-04-25

ACCAAACAAAGTTGGGTAAGGATAGATCTTATAATAACTATGGACTGGCAAACTTAGGAGTAAAGATCCTACTGTCGGGG

GGAGGAGGAGGAGCAAGATCTTTGATTATGGCGACTCTCCTTAAAAGCTTAGCATTGTTCAAAAGGAACAAAGACAAAGC

GCCGACAGCATCAGGTTCAGGAGGGGCCATCCGGGGGATTAAGAATGTTATCATAGTCCCGATTCCCGGAGACTCGTCCA

TCACTACCCGTTCAAGACTGCTCGACAGGCTTGTCAGATTGGCCGGAGATCCTGACATCAACGGGTCAAAGCTGACCGGG

GTGATGATCAGCATGTTATCCTTGTTCGTAGAGTCACCCGGGCAATTGATACAGCGAATCACAGATGATCCGGATGTTAG

TATCCGCCTTGTTGAGGTAGTTCAAAGTACTAGGTCCCAGTCCGGGTTGACCTTTGCATCACGTGGTGCTGATTTAGACA

ACGAGGCAGACATGTATTTTTCAACTGAGGGGCCCTCGAGTGGAGGTAAGAAAAGGATCAACTGGTTTGAGAACAGAGAA

ATAATAGACATAGAGGTGCAGGATCCAGAAGAGTTCAATATGTTGTTAGCCTCCATACTAGCACAAGTCTGGATCCTTCT

GGCCAAGGCTGTTACGGCACCAGATACGGCAGCTGACTCAGAACTGAGAAGGTGGGTTAAATACACACAACAAAGGAGAG

TGATTGGGGAATTTCGCCTTGACAAGGGGTGGCTGGACGCAGTCCGCAACAGAATTGCAGAAGATCTATCACTCCGGCGG

TTCATGGTATCTCTCATACTTGACATCAAGAGGACCCCAGGCAACAAGCCAAGGATTGCAGAAATGATCTGCGACATTGA

CAACTATATTGTCGAAGCAGGACTCGCCAGTTTCATCCTTACTATCAAATTTGGTATCGAAACCATGTATCCTGCATTAG

GGCTCCACGAGTTTGCCGGAGAATTGTCCACTATTGAGTCCTTGATGAACTTGTATCAACAGCTAGGAGAGGTTGCACCC

TACATGGTAATTCTAGAGAATTCAATTCAGAACAAGTTTAGTGCAGGAGCTTATCCCCTCCTCTGGAGCTATGCGATGGG

TGTCGGAGTCGAGCTGGAGAACTCAATGGGGGGCTTGAACTTTGGCAGGTCATATTTTGACCCTGCCTATTTTCGTCTCG

GACAGGAGATGGTCAGAAGATCTGCAGGAAAGGTCAGCTCTGTGATTGCGGCTGAGCTCGGCATCACAGCAGAGGAAGCT

AAACTAGTCTCGGAAATCGCCTCGCAGGCTGGGGACGAAAGAACCGCCAGAGGGACTGGGCCTCGACAGGCGCAGGTTTC

CTTCCTTCAGCACCAAACAGGAGGGGGAGAGTCGTCCGCACCAGCGACCAGAGAAGGGGTCAAAGCTGCGATCCCAAACG

GAGCTGAAGAAAGGGACAGAAAGCAAACACGCCCAGGAAGGCCCAGAGGAGAGACCTCCGGCCAACTGCTCCTGGACATC

ATGCCAGAGGATGAGATCTTGCGAGAGTCTGGTCAAAACCCTCGTGAGGCTCAAAGATCGGCCGAGGCACTCTTCAGGCT

GCAGGCCATGGCCAAGATTCTGGAGGACCAGGAGGAGGGAGAAGACAACAGTCAGGTCTACAACGACAAGGATCTCCTCG

GCTAAGCAGACGCACCCTCTGTCGAAATCAGTGACAAGACATCTCTTACCAGTATTATAAAAAACTTAGGACCCAGGTCC

AAGCAAGCACACATCGACACCCCAACCAGTCGAGCGGAGACCACCGATGGCAGAAGAACAAGCATACCATGTCAACAAGG

GGCTGGAATGTATCAAGTCTCTCAAAGCCTCACCCCCGGATCTATCCACCATCAGAGATACCATCGAGAGCTGGAGAGAG

GGGCTTAGCCCCTCGGGCCGTGCAACACCGAACCCTGATACGTCCGAGGGAGATCATCAGAATATCAACCAATCATGCTC

ACCAGCAATCGGACCAAACAAAGTCTACTTGTCTCCTGAAGATAATCTCGGATTTAGAGAGATCACTGGCAACGACTGTG

AGACTGGGCTCGGAGGGGTCCAGAGAGAAGGATCCAACTCTCAAGTACAGCGTTACCATGTTTATAGTCACGGGGGTGAA

GAGATTGAAGGACTCGAGGATGCTGACTCTCTCGTGGTTCAAGCAGATCCTCCAGTTGCTAACGTCTTCAATGGAGGAGA

GGATGGATCTGACGACAGCGATGTGGACTCTGGCCCAGATGATCCCGGCAGAGATACTCTATATGACCGGGGATCTGTTG

CCGGCAATGGTATCGCTAGGTCCACAGATGTCGAAAAACTAGAAGGTGCTGATATTCAAGAAGTTCTTAACTCCCAGAAA

GGCAAAGGAGGAAGATTCCAGGGCGGGAAAACCTTGCGAGTCCCGGAAATACCCGATGTCAAGCACTCCAGACCATCAGC

CCAATCAATTAAAAAGGGCACAGACGGGAACTCAGTCTCATCTGGAACGGTGATAGAGTGTTTATCGATAAGTGGTGCAA

CCCAAACTGTGCCAGAGTCAAGATGGGAGTCATTAGAGCAAAATGCGTCTGTGGGGAGTGTCCTCAAGTCTGCGAGGAGT

GCAAAGACGATCCAGGGGTCGACACAAGAATCTGGTACCATAGCATCACTGACTCAGCCTAAAGAGAATGACTCCGAGTA

TGAGTATGAGGATGACCTATTTACAGAGATTCAGGACATCCGTGCAAGCATTGCCAAGATCCATGATGACAATAAAACTA

TCCTCTCAAAGCTTGATTCTATACTGTTATTGAAAGGGGAAGTCGACACTATCAAGAAACAAATCAGCAAGCAGAATATA

AGTATATCCACCATTGAGGGCCATCTCTCCAGTATAATGATAGCCATCCCTGGCTTTGGGAAGGACATCAAGGACCCAAC

ATCCGAAGTCGAGTTGAACCCAGATCTAAGACCTATAATAAGCCGTGACTCTGGCAGAGCTCTCGCGGAGGTCCTCAAGA

AACCCGCTGTGGATAGGTCTCAGAAAATTGGAACCAAAGCCAACTCCAGCTCAAAGGGTCAGCTTCTTAAGGATCTCCAG

CTAAAACCTGTCGACAAGCAGGCAAGCTCTGCAATCGGGTTTGTCCCGTCCGACCATGAATCATCCAGAAATGTCATCCG

CTCCATAATCAAGTCGAGCAAGCTAAATATTGATCACAAGGACTATCTTCTAGATTTACTAAATGATGTGAAAGGCTCCA

AGGATCTTAAGGAATTTCACAAGATGCTAACAGCAATTCTCGCCAAGCACCCGTAACACATCCTCCAGTCATCATCTCAT

ACTCGACTAAAAACATCCTTTCAATCAGGCTATTACAAAAAACTTAGGAGCAAGGGCAACTGAGCTTCGCAGACAGGATG

ACCGAGATCTACGACTTCGATAAATCAGCATGGGATGTCAAAGGGTCAATTGCCCGCATAGAACCCACCACCTACCACGA

CGGCCGACTGATACCCCAGGTGAGGGTCATCGATCCTGGTCTGGGAGACAGAAAAGATGAGTGCTTTATGTACCTGTTTC

TCCTAGGAGTGATTGAGGATAACGACCCCCTGTCTCCCCCCGTCGGAAGAACCTTTGGCTCTCTACCTCTAGGGGTTGGT

AGGTCAACTGCCAAGCCAGAAGAACTACTAAGGGAGGCCACAGAATTAGATATAGTGGTGAGGCGCACTGCAGGAGTAAA

TGAGAAACTGGTATTTTACAACAACACTCCGCTGTCCTTGTTAACGCCCTGGAAGAAAGTTCTGACAACCGGAAGTGTGT

TTAGCGCTAACCAGGTTTGCAATGCAGTCAACCTAGTCCCGCTTGATACTCCCCAGAGATTCAGGGTTGTGTACATGAGC

ATAACTAGATTATCAGACAATGGTTACTATAGTGTGCCCAGAAGAATGCTGGAGTTCCGCTCGGCCAATGCAGTAGCCTT

CAACATCTTGGTTACACTAAGAATTGAAAATGGCACAAACCCTAGAAGATACATAGTCGGCTCATGGGAGAATTCAGAGG

TCACATTTATGGTACACGTGGGCAACTTTAGGAGAAAGAAGAACGAAGTATACTCTGCTGATTACTGCAAAATGAAGATT

GAAAAGATGGGTTTAGTTTTTGCCCTGGGTGGAATAGGCGGAACAAGTCTCCATATTAGAAGCACCGGGAAAATGAGCAA

AACCCTCCATGCACAGCTGGGGTTCAAGAAAATCTTATGTTACCCCCTAATGGATGTCAATGAGGATCTTAACCGATATC

TCTGGCGGGCAGAGTGCCGAATAGTCAAAATCCAAGCTGTCTTACAGCCATCAGTACCCCAAGAGTTCCGTGTCTACGAT

GATGTCATCATCAACGATGATCAAGGCTTGTTCAAGATCTTGTAGTTCATTTGCAAAGGAACGAACCCACCCCCAGGAAG

GGGCCCCACCCTCCCCCG-CCCAAAAACACACTCCAAGGCTCCCCGGCACAGCGCCAACACAGCTACTCCCGCGATGGAA

CGAGGTCGGCC-CCCGAACAAATCCGGTCCAGGGGGGGGGGCCCCCCCCCTGACCAACCGGGCGCCGACCGAGCAGGCCG

GCGAGAAGCCCCCAAGAGGCCCCGACGAAGACCGAGCACCCCATCCCCCC-AAAAACGTCCCCC-AACACGCCCGAGGCT

AGCCACCGGACACCCCCAGACCCGCCACCGGCGAGACAGAACCCCAAGGGGGGACCCTCCAAACCAAAGGAATCACCGGG

GTAGGCTCCGCCGAACCAGCACAGACCCGACCCAAGCAAGGCGGACCCCCAACACGAAACTGCCCAGCACACCCTACTCA

TCATCAAACAAAACTTAGGGGCCAAGTCCATACGAGAGGCCCTAGTCCGTCAGGCGGAGGGACGCAGAAAGGAAGGAGAC

ACGACCGTCCCAGAAGGAGGACACCCAACCACCGAAACGCCCGCCCCCAGGGAAGGAG--CCA-CCCCCACGCAGCGAAG

GAGCGGGTGGGAGACCTCGGGGAGGGCCCGACCGGGACCCGGGGGCCGGCGGCGCCGGCGGTGAGCAGGCCGGGCGCATA

CCTGACCCCACCGGAGGATCCGAGGAGGGCCCGACCCCGCGTGGACCAAACGGCAGGGAGCCCAACCCCACCAAAAAAAG

AGGCGAGGCCCCCCCTCCCTCCCCCCCCGCACACAACCCGCAGACAGACCGAGGGAGGGCCATCCCCCAGCGAGACAGGC

GGCTCCCTCGCAGCCGAGCCGCAGCTCCCCAGAATAGGACACCCCCGCACACTCCGAACACCCCCTCGCCCCCCGCCACC

GGGACAGGGGCTCCACCACCACAGACCGGAGCCACCGACAGCCAGCCAGAGGTCCCCTCCAGTCCCGCCCGGCGCCCCCC

CGAAGAGGCCCCGGCAAGGCCCCCCAGCCCGACGTCCCCCGCACGCACCGAAAGAAGGACCCGCCCGAGAACGTGCGGCA

CATCCATGCGCAAACATCATGACGCGGGTCGCAATTTTGACATTTCTGTTTCTTTTCCCAAATGTTGTTGCGTGTCAGAT

TCACTGGGGCAATCTATCCAAGATCGGGATTGTAGGAACAGGGAGTGCCAGCTACAAGGTGATGACTAGGCCAAGCCACC

AGACTCTGGTTATAAAGTTAATGCCAAATATAACGGCCATCGACAATTGTACAAAGTCAGAGATTGCAGAGTACAAGAGA

TTGCTGATCACAGTGTTAAAGCCTGTAGAGGATGCTCTGTCGGTGATAACCAAGAATGTAAGACCAATTCAAACTCTAAC

ACCTGGGCGTAGAACCCGCCGTTTTGCTGGAGCTGTTCTGGCCGGGGTAGCACTTGGAGTTGCGACAGCCGCTCAGATAA

CTGCAGGAGTCGCCCTTCATCAATCATTGATGAACTCCCAAGCAATTGAGAGTTTAAAAACCAGTCTTGAGATGTCGAAT

CAGGCAATAGAAGAAATCAGACTTGCAAATAAGGAGACCATACTGGCAGTACAGGGCGTCCAGGATTATATCAACAATGA

GCTCGTCCCTTCTGTTCATAGAATGTCATGCGAGCTGGTAGGTCACAAGCTCGGCCTCAAGCTCCTTAGGTACTACACCG

AGATCCTGTCCATATTCGGGCCCAGTCTTCGAGACCCGATATCTGCCGAAATATCAATCCAGGCACTTAGTTATGCATTA

GGCGGAGACATTAATAAAATCCTGGACAAGCTTGGGTATAGCGGTGGGGATTTCCTTGCCATCCTAGAAAGCAAGGGAAT

AAAGGCCCGGGTTACATATGTGGACACAAGAGATTACTTTATAATCCTTAGCATCGCCTACCCAACCTTATCTGAGATCA

AGGGAGTGATAGTTCACAAGATAGAAGCTATAACATACAACATTGGGGCACAGGAGTGGTATACTACTATCCCTAAATAT

GTAGCCACTCAGGGGTATCTGATATCGAACTTTGATGAGACGTCATGCGTATTCACTCCAGAGGGGACAGTTTGCAGCCA

GAATGCGTTGTACCCAATGAGCCCATTGCTTCAGGAATGTTTCAGGGGGTCAACAAAATCGTGCGCCAGAACCTTAGTTT

CAGGGACCATAAGTAATAGATTTATCCTATCAAAAGGGAACCTGATTGCAAATTGTGCGTCAGTTTTGTGCAAATGTTAC

ACAACAGAGACAGTTATCAGCCAAGATCCTGACAAACTACTAACTGTTGTAGCATCCGACAAGTGTCCTGTAGTTGAGGT

GGATGGAGTGACAATACAGGTCGGCAGTCGAGAGTATCCGGATTCTGTATACTTACACAAAATAGACTTAGGTCCAGCCA

TCTCCCTAGAAAAACTGGATGTAGGCACCAATTTAGGCAATGCAGTCACAAGACTGGAGAATGCAAAGGAGCTCCTAGAT

GCATCAGACCAAATACTGAAGACTGTTAAAGGGGTACCTTTCAGTGGGAATATGTACATAGCACTGGCAGCTTGCATAGG

AGTATCCCTGGGCCTTGTCACATTAATATGCTGCTGTAAGGGGAGGTGTAAGAACAAGGAAATCCCTATCTCCAAAATCA

ACCCAGGGCTCAAACCCGACCTGACCGGGACCTCAAAGTCGTACGTAAGATCACTGTAGTCAGAATCACCTGGATCATCT

GGCATCGCACACATACATGCACGACACAGGCAGTCCGAGGACGCAAGAAACCCAGCCTCCGGTCACCCACCCGACCCCAC

TCCACGCTCCACCACACATTAGTCATCAAACAAAACTTAGGACGAAAGGTCAATCACCATGTCCGCACAAAGGGAGAGGA

TCAATGCCTTCTACAAAGACAATCCTCACAATAAGAACCATAGGGTGATCCTGGATAGAGAACGCTTGGTCATTGAAAGA

CCCTACATCTTGCTTGGAGTCCTGCTGGTAATGTTCCTGAGTCTAATCGGACTGCTGGCCATTGCAGGGATCAGGCTTCA

CCGGGCCACCGTTGGAACTTCAGAGATCCAGAGTCGGCTGAATACCAATATTAAGTTGACCGAATCTATTGATCACCAGA

CTAAGGATGTCTTAACTCCCCTTTTTAAAATCATTGGCGATGAAGTCGGCATCAGAATTCCACAGAAATTCAGTGATCTT

GTCAAGTTCATCTCCGATAAGATTAAATTCCTCAACCCTGATAGAGAGTATGATTTCAGAGATCTCCGGTGGTGTATGAA

TCCCCCCGAGAGAGTCAAAATTAATTTTGATCAGTTTTGTGAGTACAAGGCTGCGGTTAAGTCAATTGAACATATATTTG

AGTCACCACTCAACAAGTCAAAAAAGCTGCAATCTTTGACTCTCGGGCCCGGAACAGGCTGTCTAGGCAGGACAGTAACA

AGAGCCCATTTCTCAGAACTTACAATGACCTTAATGGACCTGGATCTAGAGATGAAGCACAACGTGTCCTCAGTGTTTAC

CGTAGTTGAAGAGGGATTATTCGGAAGAACATATACCGTCTGGAGATCCGATGCCAGGGATCCGAGCACCGATCTAGGTA

TCGGCCATTTTTTAAGAGTCTTCGAGATTGGACTGGTAAGAGATCTTGGGCTGGGTCCCCCTGTTTTTCATATGACCAAC

TATCTCACGGTGAACATGAGTGATGACTATCGGAGATGTCTTTTAGCGGTAGGGGAGTTGAAGTTGACAGCCCTATGCAC

CTCATCTGAGACTGTGACACTGAGTGAGAGAGGAGTTCCAAGGAGGGAACCTCTTGTGGTTGTGATACTTAATCTAGCTG

GACCCACTCTAGGGGGAGAGCTATACAGTGTCTTGCCTACCTCTGATCTCATGGTGGAGAAACTCTATTTGTCTTCACAT

AGAGGGATCATCAAAGATGATGAGGCCAATTGGGTAGTGCCGTCTACCGATGTTCGTGATCTTCAGAACAAAGGTGAATG

TCTGGTGGAAGCATGCAAGACTCGACCTCCTTCATTTTGCAATGGCACAGGATCAGGCCCGTGGTCAGAAGGGAGAATCC

CTGCCTACGGGGTGATCAGGGTCAGTCTTGACTTAGCGAGTGACCCGGATGTAGTTATCACTTCAGTGTTTGGCCCACTG

ATACCTCACCTATCCGGCATGGATCTTTACAACAACCCGTTTTCAAAAGCTGTATGGTTGGCTGTACCACCTTATGAGCA

GTCATTTCTAGGAATGATAAATACAATTGGATTCCCTAACAGAGCAGAGGTTATGCCGCACATTTTGACCACAGAGATCA

GAGGCCCTCGGGGTCGTTGCCATGTTCCCATAGAATTGTCCCGCCGGGTTGATGACGATATCAAGATCGGGTCCAACATG

GTCATATTGCCGACGATGGACCTGAGGTATATTACAGCCACTTATGATGTTTCCAGGAGCGAGCATGCAATCGTGTACTA

TATCTATGACACGAGTCGCTCATCATCTTACTTCTACCCAGTTCGACTGAATTTCAAAGGCAATCCTCTCTCTTTGAGGA

TAGAGTGTTTCCCCTGGCGTCATAAGGTGTGGTGCTACCATGATTGTCTTATATACAACACCATAACAGGTGAAGAGGTC

CATACGAGAGGGCTGACCGGCATAGAGGTAACATGCAATCCAGTCTGAGCAGAGCTACGACCGTCACTCAGGCAGTCCCT

TGAGTCGCCACCGAGTCCAAGCAGCACAGCCTGGGACAATCAACGGCACAGCCCAGCCAACAATGTTATAAAAAACTTAG

GAGCCAGGGTTGTAGGGGCCATGGACTCTCTATCAGTCAATCAGGTCTTGTACCCTGAGGTTCATCTAGACAGCCCTATT

GTCACAAACAAACTAGTTGCCATCCTTGAGTACTCGGGTATCGACCATAACTATGTTCTTGAAGACCAGACTCTTGTCAA

GAATATTAGGTATAGGCTGGGGTGCGGTTTTTCAAATCAAATGATCATCAATAACAGGGGGGTGGGTGAAACAGTCAATT

CTAAACTTAAAAGTTACCCCCATAATCGTCATATCATATACCCGGATTGCAATAAGGAGTTGTTTTGTATCAAGGATAGC

TGCATATCTAAGAAGCTCTCGGAGCTATTCAAGAAGGGTAATTCCTTGTACTCTAAGATAAGTCACCAGGTACTGGATTG

TCTTAAGAGAGTCAATGGGAAATTAGGTCTGGGCACAGATCTTACCCATGGTCTGAAGGAGGGTATCCTTGACCTAGGGT

TGCACATGCATAGCTCTCAATGGTTCGAGACTTTCCTGTTCTGGTTCACTATCAAGACAGAGATGAGATCAATGATCAAA

GAACAGTCCCATATATGCCACAAGAGGAGGTATAACCCGACTTTTGTGTCGGGGGATGCATTCGAGGTGCTCGTATCGCG

AGACCTCGTTGTGATAATTGATAAGAATACCCAGTATGTCTTCTACTTAACATTTGAGCTGGTCCTTATGTATTGTGATG

TCATAGAGGGTAGACTTATGACGGAGACAGCCATGGCCATAGACCAGAGATATTCAGAACTTCTAAGCCGGGTCAGATAC

TTGTGGGATCTTATTGATGGGTTCTTCCCAACACTAGGCAACACCACATACCAAATTGTTGCTCTGCTTGAACCATTATC

GTTGGCTTATCTTCAACTTCAGGATGTCACTCTGGAGTTAAGAGGTGCTTTTTTAGACCATTGCTTCAAAGAACTCTATG

AGATACTGGAACATTGCGGCATTGACACAGCAGGCACTTACAATTCCATCACTGAAGGATTGGATTACGTATTCATCACC

CACGATATACATCTAACTGGGGAGATTTTTTCATTTTTCCGGAGTTTCGGACACCCCCGTCTAGAAGCAGTCACCGCTGC

GGAAAATGTCAGAAAACATATGAACCAACCGAAGGTAATCAGTTATGAGACTATGATGAAAGGCCATGCGGTATTTTGCG

GGATAATCATAAATGGTTTTCGGGACCGACATGGCGGCAGCTGGCCCCCTGTTGCACTGCCAGAACATGCCTCTGCTGCG

ATCCGGAATGCGCAGGCATCAGGTGAGGGACTGACCCATGACCTGTGTATAGACAACTGGAAATCCTTTGTTGGATTCAA

ATTTGGCTGCTTTATGCCACTCAGCCTAGATAGTGATTTGACCATGTATCTCAAAGACAAGGCATTGGCTGCACTGAAGA

ATGAGTGGGATTCAGTTTACCCGAAAGAATACCTCCGTTATAACCCACCTAGAGGGACAGAGTCAAGGCGATTGGTAGAG

GTATTCCTGAATGACTCCAGCTTTGATCCTTATAACATGATAATGTACGTGGTGAATGGCTCCTACCTTAAAGACCCTGA

GTTTAACCTCTCATACAGCCTAAAAGAGAAGGAGATAAAGGAGACAGGGCGGTTGTTTGCCAAAATGACCTATAAGATGC

GGGCCTGTCAGGTAATCGCTGAAAATCTGATATCGAATGGTGTTGGGAAGTATTTCCGAGACAATGGGATGGCAAAGGAC

GAGCATGACCTAACAAAAGCCCTTCACACCCTGGCAGTCTCAGGTGTTCCCAAAAACAACAAAGATAACCACCGAGGTGG

GCCTCCCAGAAGGACAACAAACCGAGGGGTGAGATCAAGCCAAGGCACCAAAACACAAGATAGAGACAAGGTTCAAGGGG

GACCTATGTACAACTATTTGCGATGCCAACCGATCAGCCCTGATCAGGGTGAGTCATACGAGACTGTTAGTGCATTCATC

ACCGCTGACCTTAAGAAGTATTGCCTGAATTGGAGATACGAGACAATCAGCATATTTGCACAGAGGCTGAATGAAATATA

TGGACTACCATCCTTCTTTCAATGGTTACACAGGATATTGGAAAAATCCGTACTCTACGTCAGTGACCCACATTGTCCTC

CCGACCTAGATAATCATATCCCTCTGGACAGTGTCCCCAATGCCCAAATATTCATCAAGTACCCAATGGGCGGAATAGAA

GGTTATTGCCAAAAACTATGGACAATCAGCACTATACCATACTTGTATCTGGCAGCCTATGAGAGCGGAGTAAGAATCGC

CTCACTGGTGCAGGGTGACAATCAGACAATCGCAGTGACAAAAAGAGTTCCAAGTTCTTGGCCTTATTCACTAAAAAAGA

GGGAGGCCTCCAAGGCCGCTCAAAATTACTTCGTAGTCTTAAGGCAAAGATTGCACGATGTAGGTCATCACTTAAAGGCT

AATGAGACCATAGTATCTTCTCACTTTTTTGTATATTCCAAAGGGATTTATTATGACGGCCTGCTAGTCTCACAATCACT

AAAGAGCATCGCCAGATGTGTCTTCTGGTCCGAGACTATAGTGGATGAAACCAGAGCGGCGTGCAGTAATATTGCAACAA

CTGTCGCTAAGAGTATAGAGAGGGGTTATGATAGGTACCTTGCATATTCTTTGAATATCCTCAAGATTTTCCAACAGATC

CTTATATCCCTTAACTTCACTATTAACACAACAATGACTCAGGATGTCGTGGCACCGATCATCGAGAACGGTGATTTGCT

GATAAGGATGGCACTCTTGCCAGCACCCATCGGGGGTCTCAATTATCTTAACATGAGCAGGTTATTTGTGAGAAATATCG

GTGACCCGGTCACCTCCTCTATAGCCGACCTGAAGAGGATGATAGATGCCGGGCTAATGCCAGAAGAAACATTGCATCAA

GTGATGACCCAGACCCCGGGAGAATCATCCTACCTAGATTGGGCAAGTGACCCTTATTCTGCCAACCTACCCTGCGTACA

GAGTATCACTCGCCTTCTCAAAAACATCACTGCACGATATATTTTAATCAGCAGCCCAAACCCGATGCTGAAAGGGTTAT

TTCATGAGGGGAGTAGAGATGAAGACGAGGAGCTTGCAAGTTTCCTAATGGATCGGCATATAATTGTTCCAAGAGCTGCA

CATGAAATCTTAGACCATAGTATAACCGGAGCAAGAGAAGCTATAGCCGGGATGTTGGACACCACCAAGGGTCTGATTAG

AACAAGTATGAAACGGGGCGGCCTCACCCCTCGAGTCTTAGCCCGCCTTTCCAATTATGATTATGAACAATTCCGATCCG

GGATAACACTATTAACAAAGAAAGGGCAGTGTTATCTCATTGACAAGGACTCGTGCTCGGTGCAGCTCGCTATAGCTCTG

AGGGGCCACATGTGGGCTAAGTTGGCTCGCGGGAGACCAATCTATGGGTTGGAGGTGCCTGATGTATTAGAATCGATGAA

CGGCTACCTTATCAAACGCCACGAGTCGTGTGCTATCTGTGAAACGGGCTCAAGTCACTACGGATGGTTTTTCGTCCCTG

CAGGGTGCCAGCTTGACGATGTATCTAGAGAGACTTCGGCTCTTCGTGTACCTTATGTCGGATCAACAACTGAGGAAAGG

ACAGACATGAAACTTGCATTTGTTAGGTCTCCAAGTAGATCCCTCAAATCAGCAGTTAGAATTGCAACAGTTTACTCATG

GGCTTACGGGGATGATGAGAAATCATGGAGTGAAGCCTGGATGCTAGCTAGGCAGAGAGCCAATATCACCTTAGATGAAT

TGAGAATGATCACTCCAGTCTCCACATCCACCAACTTAGCCCACCGGTTGAGGGATCGGAGTACCCAGGTGAAATACTCG

GGGACATCCCTTGTGAGGGTTGCAAGATACACAACCATCTCCAATGATAACTTGTCGTTTGTGATATCTGAGAAAAAAGT

GGATACCAACTTCATTTATCAGCAAGGGATGTTGCTTGGTCTTGGGATCCTTGAGAACCTCTTTAGGCTAGAGGCCACCA

CCGGAGTATCCAACACAGTGCTACACCTGCACGTGGAAACAGAATGTTGTGTTGTACCTATGGTGGATCACCCAAGGATA

CCGAGTCTCCGTAATATAAAAGTTACGGATGAGCTATGCACAAACCCTTTGATCTACGACAGGTCCCCCATCATAGAGCA

CGATGCAACCCGGCTATATTCACAGAGCCACAGGAGACATTTGGTGGAGTTTGTTACCTGGTCAACAAGCCAGCTTTACC

ATATACTGGCTAAGTCTACAGCAATGTCCATGATTGAATTGATCACGAGATTCGAGAAAGATCACATGAATGAAATAGCC

GCCCTGATTGGCGATGACGACATCAACAGCTTCATCACAGAATTTTTGCTAGTGGAGCCTAGATTATTTATAGTTTACCT

TGGTCAGTGTGCTGCCATCAATTGGGCCTTTGATATACATTATCATCGGCCCTCGGGCAAGTACCAGATGGGGGAACTTC

TCTACTCTCTGCTCTCTCGGATGAGCAAAGGAGTATATAAGATCTTTACTAATGCTTTGAGCCACCCCAAAGTTTACAAG

AAATTTTGGCGAAGTGGCATCATTGAGCCGATTCATGGCCCATCCTTGGATACACAGAATTTACATGTCACTGTCTGTGA

CATGATATATGGGTCCTATGTCACCTATTTGGATCTTTTGCTGAATGATGAGCTAGATGATTACCCGTATTTGCTCTGCG

AGAGTGATGAGGACGTGGTCACAGACAGGTTCGACAACATTCAAGCCAAACATCTCTGTGTATTGGCCGATGTATATTGC

AGCTCCAAGAGGTGTCCTTCAATAATCGGGATGTCTCCTATAGAAAAATGTACCATTCTCACACATTACATCAAGGGAGA

ATCAGTACAATCCCCGTCTGGGACCTCATGGAACACTGATCCCCTTGTAGTAGATCATTACTCATGCTCTCTGACCTACC

TTCGCCGCGGTTCCATCAAACAAATCAGGTTGAGGGTGGATCCTGGGTTTGTATTCGAGGCGTTGACAGACGTCGACTTT

AAACAACCTCGCAAAGCTAAGTTAGATATATCGGTCGTTGGGTTGACTGATTTCTCTCCCCCTTACGACAACGTCGGTGA

TTTTTTAGGGACTATCAACACATTGAGGCACGATCTGCCTGTCACCGGAACCGGGGTCTCGAACTATGAAGTCCACGCTT

ATCGTAGAATTGGCCTGAATTCATCAGCATGTTATAAAGCCGTAGAGATCTCCACGTTAATCAAGCCATCCTTAGAAGTC

GGAGAGCATGGATTGTTCTTAGGAGAAGGTTCCGGTTCAATGCTGGCTGCGTACAAAGAAGTTCTTAAATTAGCAAATTG

TTACTACAACAGCGGAGTAACAGCGGAGGGCAGAGCCGGACAGAGGGAAATATCTCCCTATCCTTCAGAGATGAGCCTAG

TAGAGAATCAAATGGGGATAGAGCGGAGTGTTAAAGTGCTGTTCAACGGGAAACCTGAAGTAACTTGGGTGGGGACCACA

GATTGCTACAAGTACATAATCAGTAACATCCAAACCTCTAGTCTGGGTTTCATACATTCAGATATTGAGACGCTCCCAAC

CAAAGATGCTGTTGAGAAATTGGAGGAATTTGCCTCTATCCTATCCTTATCCTTGATTTTGGGGAAAATCGGCTCTATTA

CAGTTGTCAAAATTATGCCCATTAGCGGGGATTTTACCCAAGGCTTCATAGCCTATGCCATCCAATATTTTAGGGAGAGC

CTGCTTGCCTACCCGAGATATAGTAACTTCATCTCGACTGAGTGTTACCTTATTATGATAGGATTAAAGGCCAATCGATT

GATAAACCCAGAAGCCATTAAGCAAAGCATAATCAGAGCGGGGGTCAGGACTGCACCAGGACTTGTGAGCCATGTATTAT

CAGAGAAACAAAAAGGTTGTATTCAATCTTTCTTGGGTGATCCTTATATCCAAGGAGACTTCAATAAGCACCTTAAATCT

CTAACCTCTATTGAGAAAATCCTGGTAAATTGTGGTCTCTCGATCAATGGCATAAAAATCTGTAGAGATCTAATCCACCA

TGATATCGCCTCCGGTCCAGACGGCCTTATGAGCTCTACAATTATTTTATATAGGGAATTGGCTCATTTCAAAGATAATA

TAAGGAGTCAGCACGGAATGTTCCACCCCTACCCGGTACTGGCCACCAGCAGGCAACGTGAATTAATCCTTCGAATAGCC

AAGAAATTCTGGGGGTATGTCTTGCTATATTCTGATGACCCGGCACTAATCAGACAAACAATCAAGAACTTGAAACGGAA

TCATCTAACCTTTGACTTACATAGTAACCCATTTATTAAAGGGCTATCCAAGGCTGAGAAACTGCTGGTGCGGACGAGTT

CACTAAGAAGAGAATGGTTGTTCACTCTCGAAACGAAAGAAGTAAAAGAGTGGTTCAAATTGGTAGGTTACAGTGCACTC

ATCAGAGGCTAGTCAGCTATGCATCTGCCCCCTTCTCCTCCGCCATGAGACCCCACTGGCGATCCAGAAGATTAAAGAAA

ACTACATATTGGATAAGTATCTATTCCCAGCTTTGTCTGGT

>MF443347.1/goat/China/2014-04-03

ACCAAACAAAGTTGGGTAAGGATAGATCTTATAATAACTATGGACTGGCAAACTTAGGAGTAAAGATCCTACTGTCGGGG

GGAGGAGGAGGAGCAAGATCCTTGATCATGGCGACTCTCCTTAAAAGCTTAGCATTGTTCAAAAGGAACAAAGACAAAGC

GCCGACAGCATCAGGTTCAGGAGGGGCCATCCGGGGGATTAAGAATGTTATCATAGTCCCGATTCCCGGAGACTCGTCCA

TCACTACCCGTTCAAGACTGCTCGACAGGCTTGTCAGATTGGCCGGAGATCCTGACATCAACGGGTCAAAGCTGACCGGG

GTGATGATCAGCATGTTATCCTTGTTCGTAGAGTCACCCGGGCAATTGATACAGCGAATCACAGATGATCCGGATGTTAG

TATCCGCCTTGTTGAGGTAGTTCAAAGTACTAGGTCCCAGTCCGGGTTGACCTTTGCATCACGTGGTGCTGATTTAGACA

ACGAGGCAGACATGTATTTTTCAACTGAGGGGCCCTCGAGTGGAGGTAAGAAAAGGATCAACTGGTTTGAGAACAGAGAA

ATAATAGACATAGAGGTGCAGGATCCAGAAGAGTTCAATATGTTGTTAGCCTCCATACTAGCACAAGTCTGGATCCTTCT

GGCCAAGGCTGTTACGGCACCAGATACGGCAGCTGACTCAGAACTGAGAAGGTGGGTTAAATACACACAACAAAGGAGAG

TGATTGGGGAATTTCGCCTTGACAAGGGGTGGCTGGACGCAGTCCGCAACAGAATTGCAGAAGATCTATCACTCCGGCGG

TTCATGGTATCTCTCATACTTGACATCAAGAGGACCCCAGGCAACAAGCCAAGGATTGCAGAAATGATCTGCGACATTGA

CAACTATATTGTCGAAGCAGGACTCGCCAGTTTCATCCTTACTATCAAATTTGGTATCGAAACCATGTATCCTGCATTAG

GGCTCCACGAGTTTGCCGGAGAATTGTCCACTATTGAGTCCTTGATGAACTTGTATCAACAGCTAGGAGAGGTTGCACCC

TACATGGTAATTCTAGAGAATTCAATTCAGAACAAGTTTAGTGCAGGAGCTTATCCCCTCCTCTGGAGCTATGCGATGGG

TGTCGGAGTCGAGCTGGAGAACTCAATGGGGGGCTTGAACTTTGGCAGGTCATATTTTGACCCTGCCTATTTTCGTCTCG

GACAGGAGATGGTCAGAAGATCTGCAGGAAAGGTCAGCTCTGTGATTGCGGCTGAGCTCGGCATCACAGCAGAGGAAGCT

AAACTAGTCTCGGAAATCGCCTCGCAGGCTGGGGACGAAAGAACCGCCAGAGGGACTGGGCCTCGACAGGCGCAGGTTTC

CTTCCTTCAGCACCAAACAGGAGGGGGAGAGTCGTCCGCACCAGCGACCAGAGAAGGGGTCAAAGCTGCGATCCCAAACG

GAGCTGAAGAAAGGGACAGAAAGCAAACACGCCCAGGAAGGCCCAGAGGAGAGACCTCCGGCCAACTGCTCCTGGACATC

ATGCCAGAGGATGAGATCTTGCGAGAGTCTGGTCAAAACCCTCGTGAGGCTCAAAGATCGGCCGAGGCACTCTTCAGGCT

GCAGGCCATGGCCAAGATTCTGGAGGACCAGGAGGAGGGAGAAGACAACAGTCAGGTCTACAACGACAAGGATCTCCTCG

GCTAAGCAGACGCACCCTCTGTCGAAATCAGTGACAAGACATCCCCTACCAGTATTATAAAAAACTTAGGACCCAGGTCC

AAGCAAGCACACATCGACACTCCAACCAGTCGAGCGGAGACCACCGATGGCAGAAGAACAAGCATACCATGTCAACAAGG

GGCTGGAATGTATCAAGTCTCTCAAAGCCTCACCCCCGGATCTATCCACCATCAGAGATACCATCGAGAGCTGGAGAGAG

GGGCTTAGCCCCTCGGGCCGTGCAACACCGAACCCTGATACGTCCGAGGGAGATCATCAGAATATCAACCAATCATGCTC

ACCAGCAATCGGACCAAACAAAGTCTACTTGTCTCCTGAAGATAATCTCGGATTTAGAGAGATCACTGGCAACGACTGTG

AGGCTGGGCTCGGAGGGGTCCAGAGAGAAGGATCCAACTCTCAAGTACAGCGTTACCATGTTTATAGTCACGGGGGTGAA

GAGATTGAAGGACTCGAGGATGCTGACTCTCTCGTGGTTCAAGCAGATCCTCCAGTTGCTAACGTCTTCAATGGAGGAGA

GGATGGATCTGACGACAGCGATGTGGACTCTGGCCCAGATGATCCCGGCAGAGATACTCTATATGACCGGGGATCTGTTG

CCGGCAATGGTATCGCTAGGTCCACAGATGTCGAAAAACTAGAAGGTGCTGATATTCAAGAAGTTCTTAACTCCCAGAAA

GGCAAAGGAGGAAGATTCCAGGGCGGGAAAACCTTGCGAGTCCCGGAAATACCCGATGTCAAGCACTCCAGACCATCAGC

CCAATCAATTAAAAAGGGCACAGACGGGAACTCAGTCTCATCTGGAACGGTGATAGAGTGTTTATCGATAAGTGGTGCAA

CCCAAACTGTGCCAGAGTCAAGATGGGAGTCATCAGAGCAAAATGCGTCTGTGGGGAGTGTCCTCAAGTCTGCGAGGAGT

GCAAAGACGATCCAGGGGTCGACACAAGAATCTGGTACCATAGCATCACTGACTCAGCCTAAAGAGAATGACTCCGAGTA

TGAGTATGAGGATGACCTATTTACAGAGATTCAGGACATCCGTGCAAGCATTGCCAAGATCCATGATGACAATAAAACTA

TCCTCTCAAAGCTTGATTCTATACTGTTATTGAAAGGGGAAGTCGACACTATCAAGAAACAAATCAGCAAGCAGAATATA

AGTATATCCACCATTGAGGGCCATCTCTCCAGTATAATGATAGCCATCCCTGGCTTTGGGAAGGACATCAAGGACCCAAC

ATCCGAAGTCGAGTTGAACCCAGATCTAAGACCTATAATAAGCCGTGACTCTGGCAGAGCTCTCGCGGAGGTCCTCAAGA

AACCCGCTGTGGATAGGTCTCAGAAAATTGGAACCAAAGCCAACTCCAGCTCAAAGGGTCAGCTTCTTAAGGATCTCCAG

CTAAAACCTGTCGACAAGCAGGCAAGCTCTGCAATCGGGTTTGTCCCGTCCGACCATGAATCATCCAGAAATGTCATCCG

CTCCATAATCAAGTCGAGCAAGCTAAATATTGATCACAAGGACTATCTTCTAGATTTACTAAATGATGTGAAAGGCTCCA

AGGATCTTAAGGAATTTCACAAGATGCTAACAGCAATTCTCGCCAAGCACCCGTAACACATCCTCCAGTCATCATCTCAT

ACTCGACTAAAAACATCCTTTCAATCAGGCTATTACAAAAAACTTAGGAGCAAGGGCAACTGAGCTTCGCAGACAGGATG

ACCGAGATCTACGACTTCGATAAATCAGCATGGGATGTCAAAGGGTCAATTGCCCGCATAGAACCCACCACCTACCACGA

CGGCCGACTGATACCCCAGGTGAGGGTCATCGATCCTGGTCTGGGAGACAGGAAAGATGAGTGCTTTATGTACCTGTTTC

TCCTAGGAGTGATTGAGGATAACGACCCCCTGTCTCCCCCCGTCGGAAGAACCTTTGGCTCTCTACCTCTAGGGGTTGGT

AGGTCAACTGCCAAGCCAGAAGAACTACTAAGGGAGGCCACAGAATTAGATATAGTGGTGAGGCGCACTGCAGGAGTAAA

TGAGAAACTGGTATTTTACAACAACACTCCGCTGTCCTTGTTAACGCCCTGGAAGAAAGTTCTGACAACCGGAAGTGTGT

TTAGCGCTAACCAGGTTTGCAATGCAGTCAACCTAGTCCCGCTTGATACTCCCCAGAGATTCAGGGTTGTGTACATGAGC

ATAACTAGATTATCAGACAATGGTTACTATAGTGTGCCCAGAAGAATGCTGGAGTTCCGCTCGGCCAATGCAGTAGCCTT

CAACATCTTGGTTACACTAAGAATTGAAAATGGCACAAACCCTAGAAGATACATAGTCGGCTCATGGGAGAATTCAGAGG

TCACATTTATGGTACACGTGGGCAACTTTAGGAGAAAGAAGAACGAAGTATACTCTGCTGATTACTGCAAAATGAAGATT

GAAAAGATGGGTTTAGTTTTTGCCCTGGGTGGAATAGGCGGAACAAGTCTCCATATTAGAAGCACCGGGAAAATGAGCAA

AACCCTCCATGCACAGCTGGGGTTCAAGAAAATCTTATGTTACCCCCTAATGGATGTCAATGAGGATCTTAACCGATATC

TCTGGCGGGCAGAGTGCCGAATAGTCAAAATCCAAGCTGTCTTACAGCCATCAGTACCCCAAGAGTTCCGTGTCTACGAT

GATGTCATCATCAACGATGATCAAGGCTTGTTCAAGATCTTGTAGTTCATTTGCAAAGGAACGAACCCACCCCCAGGAAG

GGGCCCCACCCTCCCCCG-CCAAAAAACACACTCCAAGGCTCCCCGGCACAGCGCCAACACAGCTACTCCCGCGATGGAA

CGAGGTCGGCC-CCCGAACAAATCCGGTCCAGGGGGGGGGGCCCCCCCCCTGACCAACCGGGCGCCGACCGAGCAGGCCG

GCGAGAAGCCCCCAAGAGGCCCCGACGAAGACCGAGCACCCCATCCCCCC-AAAAACGTCCCCC-AACACGCCCGAGGCT

AGCCACCGGACACCCCCAGACCCGCCACCGGCGAGACAGAACCCCAAGGGGGGACCCTCCAAACCAAAGGAATCACCGGG

GTAGGCTCCGCCGAACCAGCACAGACCCGACCCAAGCAAGGCGGACCCCCAACACGAAACTGCCCAGCACACCCTACTCA

TCATCAAACAAAACTTAGGGGCCAAGTCCATACGAGAGGCCCTAGTCCGTCAGGCGGAGGGACGCAGAAAGGAAGGAGAC

ACGACCGTCCCAGAAGGAGGACACCCAACCACCGAAACGCCCGCCCCCAGGGAAGGAG--CCA-CCCCCACGCAGCGAAG

GAGCGGGTGGGAGACCTCGGGGAGGGGCCGACCGGGACCCGGGGGCCGGCGGCGCCGGCGGTGAGCAGGCCGGGCGCATA

CCTGACCCCACCGGAGGATCCGAGGAGGGCCCGACCCCGCGTGGACCAAACGGCAGGGAGCCCAACCCCACCAAAAAGAG

AGGCGAGGCCCCCCCTCCCTCCCCCCCCGCACACAACCCGCAGACAGACCGAGGGAGGGCCATCCCCCAGCGAGACAGGC

GGCTCCCTCGCAGCCGAGCCGCAGCTCCCCAGAATAGGACACCCCCGCACACTCCGAACACCCCCTCGCCCCCCGCCACC

GGGACAGGGGCTCCACCACCACAGACCGGAGCCACCGACAGCCAGCCAGAGGTCCCCTCCAGTCCCGCCCGGCGCCCCCC

CGAAGAGGCCCCGGCAAGGCCCCCCAGCCCGACGTCCCCCGCACGCACCGAAAGAAGGACCCGCCCGAGAACGTGCGGCA

CATCCATGCGCAAACATCATGACGCGGGTCGCAATTTTGACATTTCTGTTTCTTTTCCCAAATGTTGTTGCGTGTCAGAT

TCACTGGGGCAATCTATCCAAGATCGGGATTGTAGGAACAGGGAGTGCCAGCTACAAGGTGATGACTAGGCCAAGCCACC

AGACTCTGGTTATAAAGTTAATGCCAAATATAACGGCCATCGACAATTGTACAAAGTCAGAGATTGCAGAGTACAAGAGA

TTGCTGATCACAGTGTTAAAGCCTGTAGAGGATGCTCTGTCGGTGATAACCAAGAATGTAAGACCAATTCAAACTCTAAC

ACCTGGGCGTAGAACCCGCCGTTTTGCTGGAGCTGTTCTGGCCGGGGTAGCACTTGGAGTTGCGACAGCCGCTCAGATAA

CTGCAGGAGTCGCCCTTCATCAATCATTGATGAACTCCCAAGCAATTGAGAGTTTAAAAACCAGTCTTGAGATGTCGAAT

CAGGCAATAGAAGAAATCAGACTTGCAAATAAGGAGACCATACTGGCAGTACAGGGCGTCCAGGATTATATCAACAATGA

GCTCGTCCCTTCTGTTCATAGAATGTCATGCGAGCTGGTAGGTCACAAGCTCGGCCTCAAGCTCCTTAGGTACTACACCG

AGATCCTGTCCATATTCGGGCCCAGTCTTCGAGACCCGATATCTGCCGAAATATCAATCCAGGCACTTAGTTATGCATTA

GGCGGAGACATTAATAAAATCCTGGACAAGCTTGGGTATAGCGGTGGGGATTTCCTTGCCATCCTAGAAAGCAAGGGAAT

AAAGGCCCGGGTTACATATGTGGACACAAGAGATTACTTTATAATCCTTAGCATCGCCTACCCAACCTTATCTGAGATCA

AGGGAGTGATAGTTCACAAGATAGAAGCTATAACATACAACATTGGGGCACAGGAGTGGTATACTACTATCCCTAAATAT

GTAGCCACTCAGGGGTATCTGATATCGAACTTTGATGAGACGTCATGCGTATTCACTCCAGAGGGGACAGTTTGCAGCCA

GAATGCGTTGTACCCAATGAGCCCATTGCTTCAGGAATGTTTCAGGGGGTCAACAAAATCGTGCGCCAGAACCTTAGTTT

CAGGGACCATAAGTAATAGATTTATCCTATCAAAAGGGAACCTGATTGCAAATTGTGCGTCAGTTTTGTGCAAATGTTAC

ACAACAGAGACAGTTATCAGCCAAGATCCTGACAAACTACTAACTGTTGTAGCATCCGACAAGTGTCCTGTAGTTGAGGT

GGATGGAGTGACAATACAGGTCGGCAGTCGAGAGTATCCGGATTCTGTATACTTACACAAAATAGACTTAGGTCCAGCCA

TCTCCCTAGAAAAACTGGATGTAGGCACCAATTTAGGCAATGCAGTCACAAGACTGGAGAATGCAAAGGAGCTCCTAGAT

GCATCAGACCAAATACTGAAGACTGTTAAAGGGGTACCTTTCAGTGGGAATATGTACATAGCACTGGCAGCTTGCATAGG

AGTATCCCTGGGCCTTGTCACATTAATATGCTGCTGTAAGGGGAGGTGTAAGAACAAGGAAATCCCTATCTCCAAAATCA

ACCCAGGGCTCAAACCCGACCTGACCGGGACCTCAAAGTCGTACGTAAGATCACTGTAGTCAGAATCACCTGGATCATCT

GACATCACACACAGACATGCACGACACAGGCAGTCCGAGGACGCAAGAAACCCAGCCTCCGGTCACCCACCCGACCCCAC

TCCACGCTCCACCACACATTAGTCATCAAACAAAACTTAGGACGAAAGGTCAATCACCATGTCCGCACAAAGGGAGAGGA

TCAATGCCTTCTACAAAGACAATCCTCACAATAAGAACCATAGGGTGATCCTGGATAGAGAACGCTTGGTCATTGAAAGA

CCCTACATCTTGCTTGGAGTCCTGCTGGTAATGTTCCTGAGTCTAATCGGACTGCTGGCCATTGCAGGGATCAGGCTTCA

CCGGGCCACCGTTGGAACTTCAGAGATCCAGAGTCGGCTGAATACCAATATTAAGTTGACCGAATCTATTGATCACCAGA

CTAAGGATGTCTTAACTCCCCTTTTTAAAATCATTGGCGATGAAGTCGGCATCAGAATTCCACAGAAATTCAGTGATCTT

GTCAAGTTCATCTCCGATAAGATTAAATTCCTCAACCCTGATAGAGAGTATGATTTCAGAGATCTCCGGTGGTGTATGAA

TCCCCCCGAGAGAGTCAAAATTAATTTTGATCAGTTTTGTGAGTACAAGGCTGCGGTTAAGTCAATTGAACATATATTTG

AGTCACCACTCAACAAGTCAAAAAAGCTGCAATCTTTGACTCTCGGGCCCGGAACAGGCTGTCTAGGCAGGACAGTAACA

AGAGCCCATTTCTCAGAACTTACAATGACCTTAATGGACCTGGATCTAGAGATGAAGCACAACGTGTCCTCAGTGTTTAC

CGTAGTTGAAGAGGGATTATTCGGAAGAACATATACCGTCTGGAGATCCGATGCCAGGGATCCGAGCACCGATCTAGGTA

TCGGCCATTTTTTAAGAGTCTTCGAGATTGGACTGGTAAGAGATCTTGGGCTGGGTCCCCCTGTTTTTCATATGACCAAC

TATCTCACGGTGAACATGAGTGATGACTATCGGAGATGTCTTTTAGCGGTAGGGGAGTTGAAGTTGACAGCCCTATGCAC

CTCATCTGAGACTGTGACACTGAGTGAGAGAGGAGTTCCAAGGAGGGAACCTCTTGTGGTTGTGATACTTAATCTAGCTG

GACCCACTCTAGGGGGAGAGCTATACAGTGTCTTGCCTACCTCTGATCTCATGGTGGAGAAACTCTATTTGTCTTCACAT

AGAGGGATCATCAAAGATGATGAGGCCAATTGGGTAGTGCCGTCTACCGATGTTCGTGATCTTCAGAACAAAGGTGAATG

TCTGGTGGAAGCATGCAAGACTCGACCTCCTTCATTTTGCAATGGCACAGGATCAGGCCCGTGGTCAGAAGGGAGAATCC

CTGCCTACGGGGTGATCAGGGTCAGTCTTGACTTAGCGAGTGACCCGGATGTAGTTATCACTTCAGTGTTTGGCCCACTG

ATACCTCACCTATCCGGCATGGATCTTTACAACAACCCGTTTTCAAAAGCTGTATGGTTGGCTGTACCACCTTATGAGCA

GTCATTTCTAGGAATGATAAATACAATTGGATTCCCTAACAGAGCAGAGGTTATGCCGCACATTTTGACCACAGAGATCA

GAGGCCCTCGGGGTCGTTGCCATGTTCCCATAGAATTGTCCCGCCGGGTTGATGACGATATCAAGATCGGGTCCAACATG

GTCATATTGCCGACGATGGACCTGAGGTATATTACAGCCACTTATGATGTTTCCAGGAGCGAGCATGCAATCGTGTACTA

TATCTATGACACGAGTCGCTCATCATCTTACTTCTACCCAGTTCGACTGAATTTCAAAGGCAATCCTCTCTCTTTGAGGA

TAGAGTGTTTCCCCTGGCGTCATAAGGTGTGGTGCTACCATGATTGTCTTATATACAACACCATAACAGGTGAAGAGGTC

CATACGAGAGGGCTGACCGGCATAGAGGTAACATGCAATCCAGTCTGAGCAGAGCTACGACCGTCACTCAGGCAGTCCCT

TGAGTCGCCACCGAGTCCAAGCAGCACAGCCTGGGACAATCAACGGCACAGCCCAGCCAACAATGTTATAAAAAACTTAG

GAGCCAGGGTTGTAGGGGCCATGGACTCTCTATCAGTCAATCAGGTCTTGTACCCTGAGGTTCATCTAGACAGCCCTATT

GTCACAAACAAACTAGTTGCCATCCTTGAGTACTCGGGTATCGACCATAACTATGTTCTTGAAGACCAGACTCTTGTCAA

GAATATTAGGTATAGGCTGGGGTGCGGTTTTTCAAATCAAATGATCATCAATAACAGGGGGGTGGGTGAAACAGTCAATT

CTAAACTTAAAAGTTACCCCCATAATCGTCATATCATATACCCGGATTGCAATAAGGAGTTGTTTTGTATCAAGGATAGC

TGCATATCTAAGAAGCTCTCGGAGCTATTCAAGAAGGGTAATTCCTTGTACTCTAAGATAAGTCACCAGGTACTGGATTG

TCTTAAGAGAGTCAATGGGAAATTAGGTCTGGGCACAGATCTTACCCATGGTCTGAAGGAGGGTATCCTTGACCTAGGGT

TGCACATGCATAGCTCTCAATGGTTCGAGACTTTCCTGTTCTGGTTCACTATCAAGACAGAGATGAGATCAATGATCAAA

GAACAGTCCCATATATGCCACAAGAGGAGGTATAACCCGACTTTTGTGTCGGGGGATGCATTCGAGGTGCTCGTATCGCG

AGACCTCGTTGTGATAATTGATAAGAATACCCAGTATGTCTTCTACTTAACATTTGAGCTGGTCCTTATGTATTGTGATG

TCATAGAGGGTAGACTTATGACGGAGACAGCCATGGCCATAGACCAGAGATATTCAGAACTTCTAAGCCGGGTCAGATAC

TTGTGGGATCTTATTGATGGGTTCTTCCCAACACTAGGCAACACCACATACCAAATTGTTGCTCTGCTTGAACCATTATC

GTTGGCTTATCTTCAACTTCAGGATGTCACTCTGGAGTTAAGAGGTGCTTTTTTAGACCATTGCTTCAAAGAACTCTATG

AGATACTGGAACATTGCGGCATTGACACAGCAGGCACTTACAATTCCATCACTGAAGGATTGGATTACGTATTCATCACC

CACGATATACATCTAACTGGGGAGATTTTTTCATTTTTCCGGAGTTTCGGACACCCCCGTCTAGAAGCAGTCACCGCTGC

GGAAAATGTCAGAAAACATATGAACCAACCGAAGGTAATCAGTTATGAGACTATGATGAAAGGCCATGCGGTATTTTGCG

GGATAATCATAAATGGTTTTCGGGACCGACATGGCGGCAGCTGGCCCCCTGTTGCACTGCCAGAACATGCCTCTGCTGCG

ATCCGGAATGCGCAGGCATCAGGTGAGGGACTGACCCATGACCTGTGTATAGACAACTGGAAATCCTTTGTTGGATTCAA

ATTTGGCTGCTTTATGCCACTCAGCCTAGATAGTGATTTGACCATGTATCTCAAAGACAAGGCATTGGCTGCACTGAAGA

ATGAGTGGGATTCAGTTTACCCGAAAGAATACCTCCGTTATAACCCACCTAGAGGGACAGAGTCAAGGCGATTGGTAGAG

GTATTCCTGAATGACTCCAGCTTTGATCCTTATAACATGATAATGTACGTGGTGAATGGCTCCTACCTTAAAGACCCTGA

GTTTAACCTCTCATACAGCCTAAAAGAGAAGGAGATAAAGGAGACAGGGCGGTTGTTTGCCAAAATGACCTATAAGATGC

GGGCCTGTCAGGTAATCGCTGAAAATCTGATATCGAATGGTGTTGGGAAGTATTTCCGAGACAATGGGATGGCAAAGGAC

GAGCATGACCTAACAAAAGCCCTTCACACCCTGGCAGTCTCAGGTGTTCCCAAAAACAACAAAGATAACCACCGAGGTGG

GCCTCCCAGAAGGACAACAAACCGAGGGGTGAGATCAAGCCAAGGCACCAAAACACAAGATAGAGACAAGGTTCAAGGGG

GACCTATGTACAACTATTTGCGATGCCAACCGATCAGCCCTGATCAGGGTGAGTCATACGAGACTGTTAGTGCATTCATC

ACCGCTGACCTTAAGAAGTATTGCCTGAATTGGAGATACGAGACAATCAGCATATTTGCACAGAGGCTGAATGAAATATA

TGGACTACCATCCTTCTTTCAATGGTTACACAGGATATTGGAAAAATCCGTACTCTACGTCAGTGACCCACATTGTCCTC

CCGATCTAGATAATCATATCCCTCTGGACAGTGTCCCCAATGCCCAAATATTCATCAAGTACCCAATGGGCGGAATAGAA

GGTTATTGCCAAAAACTATGGACAATCAGCACTATACCATACTTGTATCTGGCAGCCTATGAGAGCGGAGTAAGAATCGC

CTCACTGGTGCAGGGTGACAATCAGACAATCGCAGTGACAAAAAGAGTTCCAAGTTCTTGGCCTTATTCACTAAAAAAGA

GGGAGGCCTCCAAGGCCGCTCAAAATTACTTCGTAGTCTTAAGGCAAAGATTGCACGATGTAGGTCATCACTTAAAGGCT

AATGAGACCATAGTATCTTCTCACTTTTTTGTATATTCCAAAGGGATTTATTATGACGGCCTGCTAGTCTCACAATCACT

AAAGAGCATCGCCAGATGTGTCTTCTGGTCCGAGACTATAGTGGATGAAACCAGAGCGGCGTGCAGTAATATTGCAACAA

CTGTCGCTAAGAGTATAGAGAGGGGTTATGATAGGTACCTTGCATATTCTTTGAATATCCTCAAGATTTTCCAACAGATC

CTTATATCCCTTAACTTCACTATTAACACAACAATGACTCAGGATGTCGTGGCACCGATCATAGAGAACGGTGATTTGCT

GATAAGGATGGCACTCTTGCCAGCACCCATCGGGGGTCTCAATTATCTTAACATGAGCAGGTTATTTGTGAGAAATATCG

GTGACCCGGTCACCTCCTCTATAGCCGACCTGAAGAGGATGATAGATGCCGGGCTAATGCCAGAAGAAACATTGCATCAA

GTGATGACCCAGACCCCGGGAGAATCATCCTACCTAGATTGGGCAAGTGACCCTTATTCTGCCAACCTACCCTGCGTACA

GAGTATCACTCGCCTTCTCAAAAACATCACTGCACGATATATTTTAATCAGCAGCCCAAACCCGATGCTGAAAGGGTTAT

TTCATGAGGGGAGTAGAGATGAAGACGAGGAGCTTGCAAGTTTCCTAATGGATCGGCATATAATTGTTCCAAGAGCTGCA

CATGAAATCTTAGACCATAGTATAACCGGAGCAAGAGAAGCTATAGCCGGGATGTTGGACACCACCAAGGGTCTGATTAG

AACAAGTATGAAACGGGGCGGCCTCACCCCTCGAGTCTTAGCCCGCCTTTCCAATTATGATTATGAACAATTCCGATCCG

GGATAACACTATTAACAAAGAAAGGGCAGTGTTATCTCATTGACAAGGACTCGTGCTCGGTGCAGCTCGCTATAGCTCTG

AGGGGCCACATGTGGGCTAAGTTGGCTCGCGGGAGACCAATCTATGGGTTGGAGGTGCCTGATGTATTAGAATCGATGAA

CGGCTACCTTATCAAACGCCACGAGTCGTGTGCTATCTGTGAAACGGGCTCAAGTCACTACGGATGGTTTTTCGTCCCTG

CAGGGTGCCAGCTTGACGATGTATCTAGAGAGACTTCGGCTCTTCGTGTACCTTATGTCGGATCAACAACTGAGGAAAGG

ACAGACATGAAACTTGCATTTGTTAGGTCTCCAAGTAGATCCCTCAAATCAGCAGTTAGAATTGCAACAGTTTACTCATG

GGCTTACGGGGATGATGAGAAATCATGGAGTGAAGCCTGGATGCTAGCTAGGCAGAGAGCCAATATCACCTTAGATGAAT

TGAGAATGATCACTCCAGTCTCCACATCCACCAACTTAGCCCACCGGTTGAGGGATCGGAGTACCCAGGTGAAATACTCG

GGGACATCCCTTGTGAGGGTTGCAAGATACACAACCATCTCCAATGATAACTTGTCGTTTGTGATATCTGAGAAAAAAGT

GGATACCAACTTCATTTATCAGCAAGGGATGTTGCTCGGTCTTGGGGTCCTTGAGAACCTCTTTAGGCTAGAGGCCACCA

CCGGAGTATCCAACACAGTGCTACACCTGCACGTGGAAACAGAATGTTGTGTTGTACCTATGGTGGATCACCCAAGGATA

CCGAGTCTCCGTAATATAAAAGTTACGGATGAGCTATGCACAAACCCTTTGATCTACGACAGGTCCCCCATCATAGAGCA

CGATGCAACCCGGCTATATTCACAGAGCCACAGGAGACATTTGGTGGAGTTTGTTACCTGGTCAACAAGCCAGCTTTACC

ATATACTGGCTAAGTCTACAGCAATGTCCATGATTGAATTGATCACGAGATTCGAGAAAGATCACATGAATGAAATAGCC

GCCCTGATTGGCGATGACGACATCAACAGCTTCATCACAGAATTTTTGCTAGTGGAGCCTAGATTATTTATAGTTTACCT

TGGTCAGTGTGCTGCCATCAATTGGGCCTTTGATATACATTATCATCGGCCCTCGGGCAAGTACCAGATGGGGGAACTTC

TCTACTCTCTGCTCTCTCGGATGAGCAAAGGAGTATATAAGATCTTTACTAATGCTTTGAGCCACCCCAAAGTTTACAAG

AAATTTTGGCGAAGTGGCATCATTGAGCCGATTCATGGCCCATCCTTGGATACACAGAATTTACATGTCACTGTCTGTGA

CATGATATATGGGTCCTATGTCACCTATTTGGATCTTTTGCTGAATGATGAGCTAGATGATTACCCGTATTTGCTCTGCG

AGAGTGATGAGGACGTGGTCACAGACAGGTTCGACAACATTCAAGCCAAACATCTCTGTGTATTGGCCGATGTATATTGC

AGCTCCAAGAGGTGTCCTTCAATAATCGGGATGTCTCCTATAGAAAAATGTACCATTCTCACACATTACATCAAGGGAGA

ATCAGTACAATCCCCGTCTGGGACCTCATGGAACACTGATCCCCTTGTAGTAGATCATTACTCATGCTCTCTGACCTACC

TTCGCCGCGGTTCCATCAAACAAATCAGGTTGAGGGTGGATCCTGGGTTTGTATTCGAGGCGTTGACAGACGTCGACTTT

AAACAACCTCGCAAAGCTAAGTTAGATATATCGGTCGTTGGGTTGACTGATTTCTCTCCCCCTTACGACAACGTCGGTGA

TTTTTTAGGGACTATCAACACATTGAGGCACGATCTGCCTGTCACCGGAACCGGGGTCTCGAACTATGAAGTCCACGCTT

ATCGTAGAATTGGCCTGAATTCATCAGCATGTTATAAAGCCGTAGAGATCTCCACGTTAATCAAGCCATCCTTAGAAGTC

GGAGAGCATGGATTGTTCTTAGGAGAAGGTTCCGGTTCAATGCTGGCTGCGTACAAAGAAGTTCTTAAATTAGCAAATTG

TTACTACAACAGCGGAGTAACAGCGGAGGGCAGAGCCGGACAGAGGGAAATATCTCCCTATCCTTCAGAGATGAGCCTAG

TAGAGAATCAAATGGGGATAGAGCGGAGTGTTAAAGTGCTGTTCAACGGGAAACCTGAAGTAACTTGGGTGGGGACCACA

GATTGCTACAAGTACATAATCAGTAACATCCAAACCTCTAGTCTGGGTTTCATACATTCAGATATTGAGACGCTCCCAAC

CAAAGATGCTGTTGAGAAATTGGAGGAATTTGCCTCTATCCTATCCTTATCCTTGATTTTGGGGAAAATCGGCTCTATTA

CAGTTGTCAAAATTATGCCCATTAGCGGGGATTTTACCCAAGGCTTCATAGCCTATGCCATCCAATATTTTAGGGAGAGC

CTGCTTGCCTATCCGAGATATAGTAACTTCATCTCGACTGAGTGTTACCTTATTATGATAGGATTAAAGGCCAATCGATT

GATAAACCCAGAAGCCATTAAGCAAAGCATAATCAGAGCGGGGGTCAGGACTGCACCAGGACTTGTGAGCCATGTATTAT

CAGAGAAACAAAAAGGTTGTATTCAATCTTTCTTGGGTGATCCTTATATCCAAGGAGACTTCAATAAGCACCTTAAATCT

CTAACCTCTATTGAGAAAATCCTGGTAAATTGTGGTCTCTCGATCAATGGCATAAAAATCTGTAGAGATCTAATCCACCA

TGATATCGCCTCCGGTCCAGACGGCCTTATGAGCTCTACAATTATTTTATATAGGGAATTGGCTCATTTCAAAGATAATA

TAAGGAGTCAGCACGGAATGTTCCACCCCTACCCGGTACTGGCCAACAGCAGGCAACGTGAATTAATCCTTCGAATAGCC

AAGAAATTCTGGGGGTATGTCTTGCTATATTCTGATGACCCGGCACTAATCAGACAAACAATCAAGAACTTGAAACGGAA

TCATCTAACCTTTGACTTACATAGTAACCCATTTATTAAAGGGCTATCCAAGGCTGAGAAACTGCTGGTGCGGACGAGTT

CACTAAGAAGAGAATGGTTGTTCACTCTCGAAACGAAAGAAGTAAAAGAGTGGTTCAAATTGGTAGGTTACAGTGCACTC

ATCAGAGGCTAGTCAGCTATGCATCTGCCCCCTTCTCCTCCGCCATGAGACCCCACTGGCGATCCAGAAGATTAAAGAAA

ACTACATATTGGATAAGTATCTATTCCCAGCTTTGTCTGGT

>MF443345.1/goat/China/2014-04-25

ACCAAACAAAGTTGGGTAAGGATAGATCTTATAATAACTATGGACTGGCAAACTTAGGAGTAAAGATCCTACTGTCGGGG

GGAGGAGGAGGAGCAAGATCCTTGATCATGGCGACTCTCCTTAAAAGCTTAGCATTGTTCAAAAGGAACAAAGACAAAGC

GCCGACAGCATCAGGTTCAGGAGGGGCCATCCGGGGGATTAAGAATGTTATCATAGTCCCGATTCCCGGAGACTCGTCCA

TCACTACCCGTTCAAGACTGCTCGACAGGCTTGTCAGATTGGCCGGAGATCCTGACATCAACGGGTCAAAGCTGACCGGG

GTGATGATCAGCATGTTATCCTTGTTCGTAGAGTCACCCGGGCAATTGATACAGCGAATCACAGATGATCCGGATGTTAG

TATCCGCCTTGTTGAGGTAGTTCAAAGTACTAGGTCCCAGTCCGGGTTGACCTTTGCATCACGTGGTGCTGATTTAGACA

ACGAGGCAGACATGTATTTTTCAACTGAGGGGCCCTCGAGTGGAGGTAAGAAAAGGATCAACTGGTTTGAGAACAGAGAA

ATAATAGACATAGAGGTGCAGGATCCAGAAGAGTTCAATATGTTGTTAGCCTCCATACTAGCACAAGTCTGGATCCTTCT

GGCCAAGGCTGTTACGGCACCAGATACGGCAGCTGACTCAGAACTGAGAAGGTGGGTTAAATACACACAACAAAGGAGAG

TGATTGGGGAATTTCGCCTTGACAAGGGGTGGCTGGACGCAGTCCGCAACAGAATTGCAGAAGATCTATCACTCCGGCGG

TTCATGGTATCTCTCATACTTGACATCAAGAGGACCCCAGGCAACAAGCCAAGGATTGCAGAAATGATCTGCGACATTGA

CAACTATATTGTCGAAGCAGGACTCGCCAGTTTCATCCTTACTATCAAATTTGGTATCGAAACCATGTATCCTGCATTAG

GGCTCCACGAGTTTGCCGGAGAATTGTCCACTATTGAGTCCTTGATGAACTTGTATCAACAGCTAGGAGAGGTTGCACCC

TACATGGTAATTCTAGAGAATTCAATTCAGAACAAGTTTAGTGCAGGAGCTTATCCCCTCCTCTGGAGCTATGCGATGGG

TGTCGGAGTCGAGCTGGAGAACTCAATGGGGGGCTTGAACTTTGGCAGGTCATATTTTGACCCTGCCTATTTTCGTCTCG

GACAGGAGATGGTCAGAAGATCTGCAGGAAAGGTCAGCTCTGTGATTGCGGCTGAGCTCGGCATCACAGCAGAGGAAGCT

AAACTAGTCTCGGAAATCGCCTCGCAGGCTGGGGACGAAAGAACCGCCAGAGGGACTGGGCCTCGACAGGCGCAGGTTTC

CTTCCTTCAGCACCAAACAGGAGGGGGAGAGTCGTCCGCACCAGCGACCAGAGAAGGGGTCAAAGCTGCGATCCCAAACG

GAGCTGAAGAAAGGGACAGAAAGCAAACACGCCCAGGAAGGCCCAGAGGAGAGACCTCCGGCCAACTGCTCCTGGACATC

ATGCCAGAGGATGAGATCTTGCGAGAGTCTGGTCAAAACCCTCGTGAGGCTCAAAGATCGGCCGAGGCACTCTTCAGGCT

GCAGGCCATGGCCAAGATTCTGGAGGACCAGGAGGAGGGAGAAGACAACAGTCAGGTCTACAACGACAAGGATCTCCTCG

GCTAAGCAGACGCACCCTCTGTCGAAATCAGTGACAAGACATCCCCTACCAGTATTATAAAAAACTTAGGACCCAGGTCC

AAGCAAGCACACATCGACACTCCAACCAGTCGAGCGGAGACCACCGATGGCAGAAGAACAAGCATACCATGTCAACAAGG

GGCTGGAATGTATCAAGTCTCTCAAAGCCTCACCCCCGGATCTATCCACCATCAGAGATACCATCGAGAGCTGGAGAGAG

GGGCTTAGCCCCTCGGGCCGTGCAACACCGAACCCTGATACGTCCGAGGGAGATCATCAGAATATCAACCAATCATGCTC

ACCAGCAATCGGACCAAACAAAGTCTACTTGTCTCCTGAAGATAATCTCGGATTTAGAGAGATCACTGGCAACGACTGTG

AGGCTGGGCTCGGAGGGGTCCAGAGAGAAGGATCCAACTCTCAAGTACAGCGTTACCATGTTTATAGTCACGGGGGTGAA

GAGATTGAAGGACTCGAGGATGCTGACTCTCTCGTGGTTCAAGCAGATCCTCCAGTTGCTAACGTCTTCAATGGAGGAGA

GGATGGATCTGACGACAGCGATGTGGACTCTGGCCCAGATGATCCCGGCAGAGATACTCTATATGACCGGGGATCTGTTG

CCGGCAATGGTATCGCTAGGTCCACAGATGTCGAAAAACTAGAAGGTGCTGATATTCAAGAAGTTCTTAACTCCCAGAAA

GGCAAAGGAGGAAGATTCCAGGGCGGGAAAACCTTGCGAGTCCCGGAAATACCCGATGTCAAGCACTCCAGACCATCAGC

CCAATCAATTAAAAAGGGCACAGACGGGAACTCAGTCTCATCTGGAACGGTGATAGAGTGTTTATCGATAAGTGGTGCAA

CCCAAACTGTGCCAGAGTCAAGATGGGAGTCATCAGAGCAAAATGCGTCTGTGGGGAGTGTCCTCAAGTCTGCGAGGAGT

GCAAAGACGATCCAGGGGTCGACACAAGAATCTGGTACCATAGCATCACTGACTCAGCCTAAAGAGAATGACTCCGAGTA

TGAGTATGAGGATGACCTATTTACAGAGATTCAGGACATCCGTGCAAGCATTGCCAAGATCCATGATGACAATAAAACTA

TCCTCTCAAAGCTTGATTCTATACTGTTATTGAAAGGGGAAGTCGACACTATCAAGAAACAAATCAGCAAGCAGAATATA

AGTATATCCACCATTGAGGGCCATCTCTCCAGTATAATGATAGCCATCCCTGGCTTTGGGAAGGACATCAAGGACCCAAC

ATCCGAAGTCGAGTTGAACCCAGATCTAAGACCTATAATAAGCCGTGACTCTGGCAGAGCTCTCGCGGAGGTCCTCAAGA

AACCCGCTGTGGATAGGTCTCAGAAAATTGGAACCAAAGCCAACTCCAGCTCAAAGGGTCAGCTTCTTAAGGATCTCCAG

CTAAAACCTGTCGACAAGCAGGCAAGCTCTGCAATCGGGTTTGTCCCGTCCGACCATGAATCATCCAGAAATGTCATCCG

CTCCATAATCAAGTCGAGCAAGCTAAATATTGATCACAAGGACTATCTTCTAGATTTACTAAATGATGTGAAAGGCTCCA

AGGATCTTAAGGAATTTCACAAGATGCTAACAGCAATTCTCGCCAAGCACCCGTAACACATCCTCCAGTCATCATCTCAT

ACTCGACTAAAAACATCCTTTCAATCAGGCTATTACAAAAAACTTAGGAGCAAGGGCAACTGAGCTTCGCAGACAGGATG

ACCGAGATCTACGACTTCGATAAATCAGCATGGGATGTCAAAGGGTCAATTGCCCGCATAGAACCCACCACCTACCACGA

CGGCCGACTGATACCCCAGGTGAGGGTCATCGATCCTGGTCTGGGAGACAGAAAAGATGAGTGCTTTATGTACCTGTTTC

TCCTAGGAGTGATTGAGGATAACGACCCCCTGTCTCCCCCCGTCGGAAGAACCTTTGGCTCTCTACCTCTAGGGGTTGGT

AGGTCAACTGCCAAGCCAGAAGAACTACTAAGGGAGGCCACAGAATTAGATATAGTGGTGAGGCGCACTGCAGGAGTAAA

TGAGAAACTGGTATTTTACAACAACACTCCGCTGTCCTTGTTAACGCCCTGGAAGAAAGTTCTGACAACCGGAAGTGTGT

TTAGCGCTAACCAGGTTTGCAATGCAGTCAACCTAGTCCCGCTTGATACTCCCCAGAGATTCAGGGTTGTGTACATGAGC

ATAACTAGATTATCAGACAATGGTTACTATAGTGTGCCCAGAAGAATGCTGGAGTTCCGCTCGGCCAATGCAGTAGCCTT

CAACATCTTGGTTACACTAAGAATTGAAAATGGCACAAACCCTAGAAGATACATAGTCGGCTCATGGGAGAATTCAGAGG

TCACATTTATGGTACACGTGGGCAACTTTAGGAGAAAGAAGAACGAAGTATACTCTGCTGATTACTGCAAAATGAAGATT

GAAAAGATGGGTTTAGTTTTTGCCCTGGGTGGAATAGGCGGAACAAGTCTCCATATTAGAAGCACCGGGAAAATGAGCAA

AACCCTCCATGCACAGCTGGGGTTCAAGAAAATCTTATGTTACCCCCTAATGGATGTCAATGAGGATCTTAACCGATATC

TCTGGCGGGCAGAGTGCCGAATAGTCAAAATCCAAGCTGTCTTACAGCCATCAGTACCCCAAGAGTTCCGTGTCTACGAT

GATGTCATCATCAACGATGATCAAGGCTTGTTCAAGATCTTGTAGTTCATTTGCAAAGGAACGAACCCACCCCCAGGAAG

GGGCCCCACCCTCCCCCG-CCAAAAAACACACTCCAAGGCTCCCCGGCACAGCGCCAACACAGCTACTCCCGCGATGGAA

CGAGGTCGGCC-CCCGAACAAATCCGGTCCAGGGGGGGGGGCCCCCCCCCTGACCAACCGGGCGCCGACCGAGCAGGCCG

GCGAGAAGCCCCCAAGAGGCCCCGACGAAGACCGAGCACCCCATCCCCCC-AAAAACGTCCCCC-AACACGCCCGAGGCT

AGCCACCGGACACCCCCAGACCCGCCACCGGCGAGACAGAACCCCAAGGGGGGACCCTCCAAACCAAAGGAATCACCGGG

GTAGGCTCCGCCGAACCAGCACAGACCCGACCCAAGCAAGGCGGACCCCCAACACGAAACTGCCCAGCACACCCTACTCA

TCATCAAACAAAACTTAGGGGCCAAGTCCATACGAGATGCCCTAGTCCGTCAGGCGGAGGGACGCAGAAAGGAAGGAGAC

ACGACCGTCCCAGAAGGAGGACACCCAACCACCGAAACGCCCGCCCCCAGGGAAGGAG--CCA-CCCCCACGCAGCGAAG

GAGCGGGTGGGAGACCTCGGGGAGGGGCCGACCGGGACCCGGGGGCCGGCGGCGCCGGCGGTGAGCAGGCCGGGCGCATA

CCTGACCCCACCGGAGGATCCGAGGAGGGCCCGACCCCGCGTGGACCAAACGGCAGGGAGCCCAACCCCACCAAAAAGAG

AGGCGAGGCCCCCCCTCCCTCCCCCCCCGCACACAACCCGCAGACAGACCGAGGGAGGGCCATCCCCCAGCGAGACAGGC

GGCTCCCTCGCAGCCGAGCCGCAGCTCCCCAGAATAGGACACCCCCGCACACTCCGAACACCCCCTCGCCCCCCGCCACC

GGGACAGGGGCTCCACCACCACAGACCGGAGCCACCGACAGCCAGCCAGAGGTCCCCTCCAGTCCCGCCCGGCGCCCCCC

CGAAGAGGCCCCGGCAAGGCCCCCCAGCCCGACGTCCCCCGCACGCACCGAAAGAAGGACCCGCCCGAGAACGTGCGGCA

CATCCATGCGCAAACATCATGACGCGGGTCGCAATTTTGACATTTCTGTTTCTTTTCCCAAATGTTGTTGCGTGTCAGAT

TCACTGGGGCAATCTATCCAAGATCGGGATTGTAGGAACAGGGAGTGCCAGCTACAAGGTGATGACTAGGCCAAGCCACC

AGACTCTGGTTATAAAGTTAATGCCAAATATAACGGCCATCGACAATTGTACAAAGTCAGAGATTGCAGAGTACAAGAGA

TTGCTGATCACAGTGTTAAAGCCTGTAGAGGATGCTCTGTCGGTGATAACCAAGAATGTAAGACCAATTCAAACTCTAAC

ACCTGGGCGTAGAACCCGCCGTTTTGCTGGAGCTGTTCTGGCCGGGGTAGCACTTGGAGTTGCGACAGCCGCTCAGATAA

CTGCAGGAGTCGCCCTTCATCAATCATTGATGAACTCCCAAGCAATTGAGAGTTTAAAAACCAGTCTTGAGATGTCGAAT

CAGGCAATAGAAGAAATCAGACTTGCAAATAAGGAGACCATACTGGCAGTACAGGGCGTCCAGGATTATATCAACAATGA

GCTCGTCCCTTCTGTTCATAGAATGTCATGCGAGCTGGTAGGTCACAAGCTCGGCCTCAAGCTCCTTAGGTACTACACCG

AGATCCTGTCCATATTCGGGCCCAGTCTTCGAGACCCGATATCTGCCGAAATATCAATCCAGGCACTTAGTTATGCATTA

GGCGGAGACATTAATAAAATCCTGGACAAGCTTGGGTATAGCGGTGGGGATTTCCTTGCCATCCTAGAAAGCAAGGGAAT

AAAGGCCCGGGTTACATATGTGGACACAAGAGATTACTTTATAATCCTTAGCATCGCCTACCCAACCTTATCTGAGATCA

AGGGAGTGATAGTTCACAAGATAGAAGCTATAACATACAACATTGGGGCACAGGAGTGGTATACTACTATCCCTAAATAT

GTAGCCACTCAGGGGTATCTGATATCGAACTTTGATGAGACGTCATGCGTATTCACTCCAGAGGGGACAGTTTGCAGCCA

GAATGCGTTGTACCCAATGAGCCCATTGCTTCAGGAATGTTTCAGGGGGTCAACAAAATCGTGCGCCAGAACCTTAGTTT

CAGGGACCATAAGTAATAGATTTATCCTATCAAAAGGGAACCTGATTGCAAATTGTGCGTCAGTTTTGTGCAAATGTTAC

ACAACAGAGACAGTTATCAGCCAAGATCCTGACAAACTACTAACTGTTGTAGCATCCGACAAGTGTCCTGTAGTTGAGGT

GGATGGAGTGACAATACAGGTCGGCAGTCGAGAGTATCCGGATTCTGTATACTTACACAAAATAGACTTAGGTCCAGCCA

TCTCCCTAGAAAAACTGGATGTAGGCACCAATTTAGGCAATGCAGTCACAAGACTGGAGAATGCAAAGGAGCTCCTAGAT

GCATCAGACCAAATACTGAAGACTGTTAAAGGGGTACCTTTCAGTGGGAATATGTACATAGCACTGGCAGCTTGCATAGG

AGTATCCCTGGGCCTTGTCACATTAATATGCTGCTGTAAGGGGAGGTGTAAGAACAAGGAAATCCCTATCTCCAAAATCA

ACCCAGGGCTCAAACCCGACCTGACCGGGACCTCAAAGTCGTACGTAAGATCACTGTAGTCAGAATCACCTGGATCATCT

GGCATCACACACAGACATGCACGACACAGGCAGTCCGAGGACGCAAGAAACCCAGCCTCCGGTCACCCACCCGACCCCAC

TCCACGCTCCACCACACATTAGTCATCAAACAAAACTTAGGACGAAAGGTCAATCACCATGTCCGCACAAAGGGAGAGGA

TCAATGCCTTCTACAAAGACAATCCTCACAATAAGAACCATAGGGTGATCCTGGATAGAGAACGCTTGGTCATTGAAAGA

CCCTACATCTTGCTTGGAGTCCTGCTGGTAATGTTCCTGAGTCTAATCGGACTGCTGGCCATTGCAGGGATCAGGCTTCA

CCGGGCCACCGTTGGAACTTCAGAGATCCAGAGTCGGCTGAATACCAATATTAAGTTGACCGAATCTATTGATCACCAGA

CTAAGGATGTCTTAACTCCCCTTTTTAAAATCATTGGCGATGAAGTCGGCATCAGAATTCCACAGAAATTCAGTGATCTT

GTCAAGTTCATCTCCGATAAGATTAAATTCCTCAACCCTGATAGAGAGTATGATTTCAGAGATCTCCGGTGGTGTATGAA

TCCCCCCGAGAGAGTCAAAATTAATTTTGATCAGTTTTGTGAGTACAAGGCTGCGGTTAAGTCAATTGAACATATATTTG

AGTCACCACTCAACAAGTCAAAAAAGCTGCAATCTTTGACTCTCGGGCCCGGAACAGGCTGTCTAGGCAGGACAGTAACA

AGAGCCCATTTCTCAGAACTTACAATGACCTTAATGGACCTGGATCTAGAGATGAAGCACAACGTGTCCTCAGTGTTTAC

CGTAGTTGAAGAGGGATTATTCGGAAGAACATATACCGTCTGGAGATCCGATGCCAGGGATCCGAGCACCGATCTAGGTA

TCGGCCATTTTTTAAGAGTCTTCGAGATTGGACTGGTAAGAGATCTTGGGCTGGGTCCCCCTGTTTTTCATATGACCAAC

TATCTCACGGTGAACATGAGTGATGACTATCGGAGATGTCTTTTAGCGGTAGGGGAGTTGAAGTTGACAGCCCTATGCAC

CTCATCTGAGACTGTGACACTGAGTGAGAGAGGAGTTCCAAGGAGGGAACCTCTTGTGGTTGTGATACTTAATCTAGCTG

GACCCACTCTAGGGGGAGAGCTATACAGTGTCTTGCCTACCTCTGATCTCATGGTGGAGAAACTCTATTTGTCTTCACAT

AGAGGGATCATCAAAGATGATGAGGCCAATTGGGTAGTGCCGTCTACCGATGTTCGTGATCTTCAGAACAAAGGTGAATG

TCTGGTGGAAGCATGCAAGACTCGACCTCCTTCATTTTGCAATGGCACAGGATCAGGCCCGTGGTCAGAAGGGAGAATCC

CTGCCTACGGGGTGATCAGGGTCAGTCTTGACTTAGCGAGTGACCCGGATGTAGTTATCACTTCAGTGTTTGGCCCACTG

ATACCTCACCTATCCGGCATGGATCTTTACAACAACCCGTTTTCAAAAGCTGTATGGTTGGCTGTACCACCTTATGAGCA

GTCATTTCTAGGAATGATAAATACAATTGGATTCCCTAACAGAGCAGAGGTTATGCCGCACATTTTGACCACAGAGATCA

GAGGCCCTCGGGGTCGTTGCCATGTTCCCATAGAATTGTCCCGCCGGGTTGATGACGATATCAAGATCGGGTCCAACATG

GTCATATTGCCGACGATGGACCTGAGGTATATTACAGCCACTTATGATGTTTCCAGGAGCGAGCATGCAATCGTGTACTA

TATCTATGACACGAGTCGCTCATCATCTTACTTCTACCCAGTTCGACTGAATTTCAAAGGCAATCCTCTCTCTTTGAGGA

TAGAGTGTTTCCCCTGGCGTCATAAGGTGTGGTGCTACCATGATTGTCTTATATACAACACCATAACAGGTGAAGAGGTC

CATACGAGAGGGCTGACCGGCATAGAGGTAACATGCAATCCAGTCTGAGCAGAGCTACGACCGTCACTCAGGCAGTCCCT

TGAGTCGCCACCGAGTCCAAGCAGCACAGCCTGGGACAATCAACGGCACAGCCCAGCCAACAATGTTATAAAAAACTTAG

GAGCCAGGGTTGTAGGGGCCATGGACTCTCTATCAGTCAATCAGGTCTTGTACCCTGAGGTTCATCTAGACAGCCCTATT

GTCACAAACAAACTAGTTGCCATCCTTGAGTACTCGGGTATCGACCATAACTATGTTCTTGAAGACCAGACTCTTGTCAA

GAATATTAGGTATAGGCTGGGGTGCGGTTTTTCAAATCAAATGATCATCAATAACAGGGGGGTGGGTGAAACAGTCAATT

CTAAACTTAAAAGTTACCCCCATAATCGTCATATCATATACCCGGATTGCAATAAGGAGTTGTTTTGTATCAAGGATAGC

TGCATATCTAAGAAGCTCTCGGAGCTATTCAAGAAGGGTAATTCCTTGTACTCTAAGATAAGTCACCAGGTACTGGATTG

TCTTAAGAGAGTCAATGGGAAATTAGGTCTGGGCACAGATCTTACCCATGGTCTGAAGGAGGGTATCCTTGACCTAGGGT

TGCACATGCATAGCTCTCAATGGTTCGAGACTTTCCTGTTCTGGTTCACTATCAAGACAGAGATGAGATCAATGATCAAA

GAACAGTCCCATATATGCCACAAGAGGAGGTATAACCCGACTTTTGTGTCGGGGGATGCATTCGAGGTGCTCGTATCGCG

AGACCTCGTTGTGATAATTGATAAGAATACCCAGTATGTCTTCTACTTAACATTTGAGCTGGTCCTTATGTATTGTGATG

TCATAGAGGGTAGACTTATGACGGAGACAGCCATGGCCATAGACCAGAGATATTCAGAACTTCTAAGCCGGGTCAGATAC

TTGTGGGATCTTATTGATGGGTTCTTCCCAACACTAGGCAACACCACATACCAAATTGTTGCTCTGCTTGAACCATTATC

GTTGGCTTATCTTCAACTTCAGGATGTCACTCTGGAGTTAAGAGGTGCTTTTTTAGACCATTGCTTCAAAGAACTCTATG

AGATACTGGAACATTGCGGCATTGACACAGCAGGCACTTACAATTCCATCACTGAAGGATTGGATTACGTATTCATCACC

CACGATATACATCTAACTGGGGAGATTTTTTCATTTTTCCGGAGTTTCGGACACCCCCGTCTAGAAGCAGTCACCGCTGC

GGAAAATGTCAGAAAACATATGAACCAACCGAAGGTAATCAGTTATGAGACTATGATGAAAGGCCATGCGGTATTTTGCG

GGATAATCATAAATGGTTTTCGGGACCGACATGGCGGCAGCTGGCCCCCTGTTGCACTGCCAGAACATGCCTCTGCTGCG

ATCCGGAATGCGCAGGCATCAGGTGAGGGACTGACCCATGACCTGTGTATAGACAACTGGAAATCCTTTGTTGGATTCAA

ATTTGGCTGCTTTATGCCACTCAGCCTAGATAGTGATTTGACCATGTATCTCAAAGACAAGGCATTGGCTGCACTGAAGA

ATGAGTGGGATTCAGTTTACCCGAAAGAATACCTCCGTTATAACCCACCTAGAGGGACAGAGTCAAGGCGATTGGTAGAG

GTATTCCTGAATGACTCCAGCTTTGATCCTTATAACATGATAATGTACGTGGTGAATGGCTCCTACCTTAAAGACCCTGA

GTTTAACCTCTCATACAGCCTAAAAGAGAAGGAGATAAAGGAGACAGGGCGGTTGTTTGCCAAAATGACCTATAAGATGC

GGGCCTGTCAGGTAATCGCTGAAAATCTGATATCGAATGGTGTTGGGAAGTATTTCCGAGACAATGGGATGGCAAAGGAC

GAGCATGACCTAACAAAAGCCCTTCACACCCTGGCAGTCTCAGGTGTTCCCAAAAACAACAAAGATAACCACCGAGGTGG

GCCTCCCAGAAGGACAACAAACCGAGGGGTGAGATCAAGCCAAGGCACCAAAACACAAGATAGAGACAAGGTTCAAGGGG

GACCTATGTACAACTATTTGCGATGCCAACCGATCAGCCCTGATCAGGGTGAGTCATACGAGACTGTTAGTGCATTCATC

ACCGCTGACCTTAAGAAGTATTGCCTGAATTGGAGATACGAGACAATCAGCATATTTGCACAGAGGCTGAATGAAATATA

TGGACTACCATCCTTCTTTCAATGGTTACACAGGATATTGGAAAAATCCGTACTCTACGTCAGTGACCCACATTGTCCTC

CCGATCTAGATAATCATATCCCTCTGGACAGTGTCCCCAATGCCCAAATATTCATCAAGTACCCAATGGGCGGAATAGAA

GGTTATTGCCAAAAACTATGGACAATCAGCACTATACCGTACTTGTATCTGGCAGCCTATGAGAGCGGAGTAAGAATCGC

CTCACTGGTGCAGGGTGACAATCAGACAATCGCAGTGACAAAAAGAGTTCCAAGTTCTTGGCCTTATTCACTAAAAAAGA

GGGAGGCCTCCAAGGCCGCTCAAAATTACTTCGTAGTCTTAAGGCAAAGATTGCACGATGTAGGTCATCACTTAAAGGCT

AATGAGACCATAGTATCTTCTCACTTTTTTGTATATTCCAAAGGGATTTATTATGACGGCCTGCTAGTCTCACAATCACT

AAAGAGCATCGCCAGATGTGTCTTCTGGTCCGAGACTATAGTGGATGAAACCAGAGCGGCGTGCAGTAATATTGCAACAA

CTGTCGCTAAGAGTATAGAGAGGGGTTATGATAGGTACCTTGCATATTCTTTGAATATCCTCAAGATTTTCCAACAGATC

CTTATATCCCTTAACTTCACTATTAACACAACAATGACTCAGGATGTCGTGGCACCGATCATAGAGAACGGTGATTTGCT

GATAAGGATGGCACTCTTGCCAGCACCCATCGGGGGTCTCAATTATCTTAACATGAGCAGGTTATTTGTGAGAAATATCG

GTGACCCGGTCACCTCCTCTATAGCCGACCTGAAGAGGATGATAGATGCCGGGCTAATGCCAGAAGAAACATTGCATCAA

GTGATGACCCAGACCCCGGGAGAATCATCCTACCTAGATTGGGCAAGTGACCCTTATTCTGCCAACCTACCCTGCGTACA

GAGTATCACTCGCCTTCTCAAAAACATCACTGCACGATATATTTTAATCAGCAGCCCAAACCCGATGCTGAAAGGGTTAT

TTCATGAGGGGAGTAGAGATGAAGACGAGGAGCTTGCAAGTTTCCTAATGGATCGGCATATAATTGTTCCAAGAGCTGCA

CATGAAATCTTAGACCATAGTATAACCGGAGCAAGAGAAGCTATAGCCGGGATGTTGGACACCACCAAGGGTCTGATTAG

AACAAGTATGAAACGGGGCGGCCTCACCCCTCGAGTCTTAGCCCGCCTTTCCAATTATGATTATGAACAATTCCGATCCG

GGATAACACTATTAACAAAGAAAGGGCAGTGTTATCTCATTGACAAGGACTCGTGCTCGGTGCAGCTCGCTATAGCTCTG

AGGGGCCACATGTGGGCTAAGTTGGCTCGCGGGAGACCAATCTATGGGTTGGAGGTGCCTGATGTATTAGAATCGATGAA

CGGCTACCTTATCAAACGCCACGAGTCGTGTGCTATCTGTGAAACGGGCTCAAGTCACTACGGATGGTTTTTCGTCCCTG

CAGGGTGCCAGCTTGACGATGTATCTAGAGAGACTTCGGCTCTTCGTGTACCTTATGTCGGATCAACAACTGAGGAAAGG

ACAGACATGAAACTTGCATTTGTTAGGTCTCCAAGTAGATCCCTCAAATCAGCAGTTAGAATTGCAACAGTTTACTCATG

GGCTTACGGGGATGATGAGAAATCATGGAGTGAAGCCTGGATGCTAGCTAGGCAGAGAGCCAATATCACCTTAGATGAAT

TGAGAATGATCACTCCAGTCTCCACATCCACCAACTTAGCCCACCGGTTGAGGGATCGGAGTACCCAGGTGAAATACTCG

GGGACATCCCTTGTGAGGGTTGCAAGATACACAACCATCTCCAATGATAACTTGTCGTTTGTGATATCTGAGAAAAAAGT

GGATACCAACTTCATTTATCAGCAAGGGATGTTGCTTGGTCTTGGGATCCTTGAGAACCTCTTTAGGCTAGAGGCCACCA

CCGGAGTATCCAACACAGTGCTACACCTGCACGTGGAAACAGAATGTTGTGTTGTACCTATGGTGGATCACCCAAGGATA

CCGAGTCTCCGTAATATAAAAGTTACGGATGAGCTATGCACAAACCCTTTGATCTACGACAGGTCCCCCATCATAGAGCA

CGATGCAACCCGGCTATATTCACAGAGCCACAGGAGACATTTGGTGGAGTTTGTTACCTGGTCAACAAGCCAGCTTTACC

ATATACTGGCTAAGTCTACAGCAATGTCCATGATTGAATTGATCACGAGATTCGAGAAAGATCACATGAATGAAATAGCC

GCCCTGATTGGCGATGACGACATCAACAGCTTCATCACAGAATTTTTGCTAGTGGAGCCTAGATTATTTATAGTTTACCT

TGGTCAGTGTGCTGCCATCAATTGGGCCTTTGATATACATTATCATCGGCCCTCGGGCAAGTACCAGATGGGGGAACTTC

TCTACTCTCTGCTCTCTCGGATGAGCAAAGGAGTATATAAGATCTTTACTAATGCTTTGAGCCACCCCAAAGTTTACAAG

AAATTTTGGCGAAGTGGCATCATTGAGCCGATTCATGGCCCATCCTTGGATACACAGAATTTACATGTCACTGTCTGTGA

CATGATATATGGGTCCTATGTCACCTATTTGGATCTTTTGCTGAATGATGAGCTAGATGATTACCCGTATTTGCTCTGCG

AGAGTGATGAGGACGTGGTCACAGACAGGTTCGACAACATTCAAGCCAAACATCTCTGTGTATTGGCCGATGTATATTGC

AGCTCCAAGAGGTGTCCTTCAATAATCGGGATGTCTCCTATAGAAAAATGTACCATTCTCACACATTACATCAAGGGAGA

ATCAGTACAATCCCCGTCTGGGACCTCATGGAACACTGATCCCCTTGTAGTAGATCATTACTCATGCTCTCTGACCTACC

TTCGCCGCGGTTCCATCAAACAAATCAGGTTGAGGGTGGATCCTGGGTTTGTATTCGAGGCGTTGACAGACGTCGACTTT

AAACAACCTCGCAAAGCTAAGTTAGATATATCGGTCGTTGGGTTGACTGATTTCTCTCCCCCTTACGACAACGTCGGTGA

TTTTTTAGGGACTATCAACACATTGAGGCACGATCTGCCTGTCACCGGAACCGGGGTCTCGAACTATGAAGTCCACGCTT

ATCGTAGAATTGGCCTGAATTCATCAGCATGTTATAAAGCCGTAGAGATCTCCACGTTAATCAAGCCATCCTTAGAAGTC

GGAGAGCATGGATTGTTCTTAGGAGAAGGTTCCGGTTCAATGCTGGCTGCGTACAAAGAAGTTCTTAAATTAGCAAATTG

TTACTACAACAGCGGAGTAACAGCGGAGGGCAGAGCCGGACAGAGGGAAATATCTCCCTATCCTTCAGAGATGAGCCTAG

TAGAGAATCAAATGGGGATAGAGCGGAGTGTTAAAGTGCTGTTCAACGGGAAACCTGAAGTAACTTGGGTGGGGACCACA

GATTGCTACAAGTACATAATCAGTAACATCCAAACCTCTAGTCTGGGTTTCATACATTCAGATATTGAGACGCTCCCAAC

CAAAGATGCTGTTGAGAAATTGGAGGAATTTGCCTCTATCCTATCCTTATCCTTGATTTTGGGGAAAATCGGCTCTATTA

CAGTTGTCAAAATTATGCCCATTAGCGGGGATTTTACCCAAGGCTTCATAGCCTATGCCATCCAATATTTTAGGGAGAGC

CTGCTTGCCTATCCGAGATATAGTAACTTCATCTCGACTGAGTGTTACCTTATTATGATAGGATTAAAGGCCAATCGATT

GATAAACCCAGAAGCCATTAAGCAAAGCATAATCAGAGCGGGGGTCAGGACTGCACCAGGACTTGTGAGCCATGTATTAT

CAGAGAAACAAAAAGGTTGTATTCAATCTTTCTTGGGTGATCCTTATATCCAAGGAGACTTCAATAAGCACCTTAAATCT

CTAACCTCTATTGAGAAAATCCTGGTAAATTGTGGTCTCTCGATCAATGGCATAAAAATCTGTAGAGATCTAATCCACCA

TGATATCGCCTCCGGTCCAGACGGCCTTATGAGCTCTACAATTATTTTATATAGGGAATTGGCTCATTTCAAAGATAATA

TAAGGAGTCAGCACGGAATGTTCCACCCCTACCCGGTACTGGCCAACAGCAGGCAACGTGAATTAATCCTTCGAATAGCC

AAGAAATTCTGGGGGTATGTCTTGCTATATTCTGATGACCCGGCACTAATCAGACAAACAATCAAGAACTTGAAACGGAA

TCATCTAACCTTTGACTTACATAGTAACCCATTTATTAAAGGGCTATCCAAGGCTGAGAAACTGCTGGTGCGGACGAGTT

CACTAAGAAGAGAATGGTTGTTCACTCTCGAAACGAAAGAAGTAAAAGAGTGGTTCAAATTGGTAGGTTACAGTGCACTC

ATCAGAGGCTAGTCAGCTATGCATCTGCCCCCTTCTCCTCCGCCATGAGACCCCACTGGCGATCCAGAAGATTAAAGAAA

ACTACATATTGGATAAGTATCTATTCCCAGCTTTGTCTGGT

>MF443339.1/goat/China/2014-04-01

ACCAAACAAAGTTGGGTAAGGATAGATCTTATAATAACTATGGACTGGCAAACTTAGGAGTAAAGATCCTACTGTCGGGG

GGAGGAGGAGGAGCAAGATCCTTGATCATGGCGACTCTCCTTAAAAGCTTAGCATTGTTCAAAAGGAACAAAGACAAAGC

GCCGACAGCATCAGGTTCAGGAGGGGCCATCCGGGGGATTAAGAATGTTATCATAGTCCCGATTCCCGGAGACTCGTCCA

TCACTACCCGTTCAAGACTGCTCGACAGGCTTGTCAGATTGGCCGGAGATCCTGACATCAACGGGTCAAAGCTGACCGGG

GTGATGATCAGCATGTTATCCTTGTTCGTAGAGTCACCCGGGCAATTGATACAGCGAATCACAGATGATCCGGATGTTAG

TATCCGCCTTGTTGAGGTAGTTCAAAGTACTAGGTCCCAGTCCGGGTTGACCTTTGCATCACGTGGTGCTGATTTAGACA

ACGAGGCAGACATGTATTTTTCAACTGAGGGGCCCTCGAGTGGAGGTAAGAAAAGGATCAACTGGTTTGAGAACAGAGAA

ATAATAGACATAGAGGTGCAGGATCCAGAAGAGTTCAATATGTTGTTAGCCTCCATACTAGCACAAGTCTGGATCCTTCT

GGCCAAGGCTGTTACGGCACCAGATACGGCAGCTGACTCAGAACTGAGAAGGTGGGTTAAATACACACAACAAAGGAGAG

TGATTGGGGAATTTCGCCTTGACAAGGGGTGGCTGGACGCAGTCCGCAACAGAATTGCAGAAGATCTATCACTCCGGCGG

TTCATGGTATCTCTCATACTTGACATCAAGAGGACCCCAGGCAACAAGCCAAGGATTGCAGAAATGATCTGCGACATTGA

CAACTATATTGTCGAAGCAGGACTCGCCAGTTTCATCCTTACTATCAAATTTGGTATCGAAACCATGTATCCTGCATTAG

GGCTCCACGAGTTTGCCGGAGAATTGTCCACTATTGAGTCCTTGATGAACTTGTATCAACAGCTAGGAGAGGTTGCACCC

TACATGGTAATTCTAGAGAATTCAATTCAGAACAAGTTTAGTGCAGGAGCTTATCCCCTCCTCTGGAGCTATGCGATGGG

TGTCGGAGTCGAGCTGGAGAACTCAATGGGGGGCTTGAACTTTGGCAGGTCATATTTTGACCCTGCCTATTTTCGTCTCG

GACAGGAGATGGTCAGAAGATCTGCAGGAAAGGTCAGCTCTGTGATTGCGGCTGAGCTCGGCATCACAGCAGAGGAAGCT

AAACTAGTCTCGGAAATCGCCTCGCAGGCTGGGGACGAAAGAACCGCCAGAGGGACTGGGCCTCGACAGGCGCAGGTTTC

CTTCCTTCAGCACCAAACAGGAGGGGGAGAGTCGTCCGCACCAGCGACCAGAGAAGGGGTCAAAGCTGCGATCCCAAACG

GAGCTGAAGAAAGGGACAGAAAGCAAACACGCCCAGGAAGGCCCAGAGGAGAGACCTCCGGCCAACTGCTCCTGGACATC

ATGCCAGAGGATGAGATCTTGCGAGAGTCTGGTCAAAACCCTCGTGAGGCTCAAAGATCGGCCGAGGCACTCTTCAGGCT

GCAGGCCATGGCCAAGATTCTGGAGGACCAGGAGGAGGGAGAAGACAACAGTCAGGTCTACAACGACAAGGATCTCCTCG

GCTAAGCAGACGCACCCTCTGTCGAAATCAGTGACAAGACATCCCCTACCAGTATTATAAAAAACTTAGGACCCAGGTCC

AAGCAAGCACACATCGACACTCCAACCAGTCGAGCGGAGACCACCGATGGCAGAAGAACAAGCATACCATGTCAACAAGG

GGCTGGAATGTATCAAGTCTCTCAAAGCCTCACCCCCGGATCTATCCACCATCAGAGATACCATCGAGAGCTGGAGAGAG

GGGCTTAGCCCCTCGGGCCGTGCAACACCGAACCCTGATACGTCCGAGGGAGATCATCAGAATATCAACCAATCATGCTC

ACCAGCAATCGGACCAAACAAAGTCTACTTGTCTCCTGAAGATAATCTCGGAGTTAGAGAGATCACTGGCAACGACTGTG

AGGCTGGGCTCGGAGGGGCCCAGAGAGAAGGATCCAACTCTCAAGTACAGCGTTACCATGTTTATAGTCACGGGGGTGAA

GAGATTGAAGGACTCGAGGATGCTGACTCTCTCGTGGTTCAAGCAGATCCTCCAGTTGCTAACGTCTTCAATGGAGGAGA

GGATGGATCTGACGACAGCGATGTGGACTCTGGCCCAGATGATCCCGGCAGAGATACTCTATATGACCGGGGATCTGTTG

CCGGCAATGGTATCGCTAGGTCCACAGATGTCGAAAAACTAGAAGGTGCTGATATTCAAGAAGTTCTTAACTCCCAGAAA

GGCAAAGGAGGAAGATTCCAGGGCGGGAAAACCTTGCGAGTCCCGGAAATACCCGATGTCAAGCACTCCAGACCATCAGC

CCAATCAATTAAAAAGGGCACAGACGGGAACTCAGTCTCATCTGGAACGGTGATAGAGTGTTTATCGATAAGTGGTGCAA

CCCAAACTGTGCCAGAGTCAAGATGGGAGTCATCAGAGCAAAATGCGTCTGTGGGGAGTGTCCTCAAGTCTGCGAGGAGT

GCAAAGACGATCCAGGGGTCGACACAAGAATCTGGTACCATAGCATCACTGACTCAGCCTAAAGAGAATGACTCCGAGTA

TGAGTATGAGGATGACCTATTTACAGAGATTCAGGACATCCGTGCAAGCATTGCCAAGATCCATGATGACAATAAAACTA

TCCTCTCAAAGCTTGATTCTATACTGTTATTGAAAGGGGAAGTCGACACTATCAAGAAACAAATCAGCAAGCAGAATATA

AGTATATCCACCATTGAGGGCCATCTCTCCAGTATAATGATAGCCATCCCTGGCTTTGGGAAGGACATCAAGGACCCAAC

ATCCGAAGTCGAGTTGAACCCAGATCTAAGACCTATAATAAGCCGTGACTCTGGCAGAGCTCTCGCGGAGGTCCTCAAGA

AACCCGCTGTGGATAGGTCTCAGAAAATTGGAACCAAAGCCAACTCCAGCTCAAAGGGTCAGCTTCTTAAGGATCTCCAG

CTAAAACCTGTCGACAAGCAGGCAAGCTCTGCAATCGGGTTTGTCCCGTCCGACCATGAATCATCCAGAAATGTCATCCG

CTCCATAATCAAGTCGAGCAAGCTAAATATTGATCACAAGGACTATCTTCTAGATTTACTAAATGATGTGAAAGGCTCCA

AGGATCTTAAGGAATTTCACAAGATGCTAACAGCAATTCTCGCCAAGCACCCGTAACACATCCTCCAGTCATCATCTCAT

ACTCGACTAAAAACATCCTTTCAATCAGGCTATTACAAAAAACTTAGGAGCAAGGGCAACTGAGCTTCGCAGACAGGATG

ACCGAGATCTACGACTTCGATAAATCAGCATGGGATGTCAAAGGGTCAATTGCCCGCATAGAACCCACCACCTACCACGA

CGGCCGACTGATACCCCAGGTGAGGGTCATCGATCCTGGTCTGGGAGACAGAAAAGATGAGTGCTTTATGTACCTGTTTC

TCCTAGGAGTGATTGAGGATAACGACCCCCTGTCTCCCCCCGTCGGAAGAACCTTTGGCTCTCTACCTCTAGGGGTTGGT

AGGTCAACTGCCAAGCCAGAAGAACTACTAAGGGAGGCCACAGAATTAGATATAGTGGTGAGGCGCACTGCAGGAGTAAA

TGAGAAACTGGTATTTTACAACAACACTCCGCTGTCCTTGTTAACGCCCTGGAAGAAAGTTCTGACAACCGGAAGTGTGT

TTAGCGCTAACCAGGTTTGCAATGCAGTCAACCTAGTCCCGCTTGATACTCCCCAGAGATTCAGGGTTGTGTACATGAGC

ATAACTAGATTATCAGACAATGGTTACTATAGTGTGCCCAGAAGAATGCTGGAGTTCCGCTCGGCCAATGCAGTAGCCTT

CAACATCTTGGTTACACTAAGAATTGAAAATGGCACAAACCCTAGAAGATACATAGTCGGCTCATGGGAGAATTCAGAGG

TCACATTTATGGTACACGTGGGCAACTTTAGGAGAAAGAAGAACGAAGTATACTCTGCTGATTACTGCAAAATGAAGATT

GAAAAGATGGGTTTAGTTTTTGCCCTGGGTGGAATAGGCGGAACAAGTCTCCATATTAGAAGCACCGGGAAAATGAGCAA

AACCCTCCATGCACAGCTGGGGTTCAAGAAAATCTTATGTTACCCCCTAATGGATGTCAATGAGGATCTTAACCGATATC

TCTGGCGGGCAGAGTGCCGAATAGTCAAAATCCAAGCTGTCTTACAGCCATCAGTACCCCAAGAGTTCCGTGTCTACGAT

GATGTCATCATCAACGATGATCAAGGCTTGTTCAAGATCTTGTAGTTCATTTGCAAAGGAACGAACCCACCCCCAGGAAG

GGGCCCCACCCTCCCCCG-CCAAAAAACACACTCCAAGGCTCCCCGGCACAGCGCCAACACAGCTACTCCCGCGATGGAA

CGAGGTCGGCC-CCCGAACAAATCCGGTCCAGGGGGGGGGGCCCCCCCCCTGACCAACCGGGCGCCGACCGAGCAGGCCG

GCGAGAAGCCCCCAAGAGGCCCCGACGAAGACCGAGCACCCCATCCCCCC-AAAAACGTCCCCC-AACACGCCCGAGGCT

AGCCACCGGACACCCCCAGACCCGCCACCGGCGAGACAGAACCCCAAGGGGGGACCCTCCAAACCAAAGGAATCACCGGG

GTAGGCTCCGCCGAACCAGCACAGACCCGACCCAAGCAAGGCGGACCCCCAACACGAAACTGCCCAGCACACCCTACTCA

TCATCAAACAAAACTTAGGGGCCAAGTCCATACGAGAGGCCCTAGTCCGTCAGGCGGAGGGACGCAGAAAGGAAGGAGAC

ACGACCGTCCCAGAAGGAGGACACCCAACCACCGAAACGCCCGCCCCCAGGGAAGGAG--CCA-CCCCCACGCAGCGAAG

GAGCGGGTGGGAGACCTCGGGGAGGGGCCGACCGGGACCCGGGGGCCGGCGGCGCCGGCGGTGAGCAGGCCGGGCGCATA

CCTGACCCCACCGGAGGATCCGAGGAGGGCCCGACCCCGCGTGGACCAAACGGCAGGGAGCCCAACCCCACCAAAAAGAG

AGGCGAGGCCCCCCCTCCCTCCCCCCCCGCACACAACCCGCAGACAGACCGAGGGAGGGCCATCCCCCAGCGAGACAGGC

GGCTCCCTCGCAGCCGAGCCGCAGCTCCCCAGAATAGGACACCCCCGCACACTCCGAACACCCCCTCGCCCCCCGCCACC

GGGACAGGGGCTCCACCACCACAGACCGGAGCCACCGACAGCCAGCCAGAGGTCCCCTCCAGTCCCGCCCGGCGCCCCCC

CGAAGAGGCCCCGGCAAGGCCCCCCAGCCCGACGTCCCCCGCACGCACCGAAAGAAGGACCCGCCCGAGAACGTGCGGCA

CATCCATGCGCAAACATCATGACGCGGGTCGCAATTTTGACATTTCTGTTTCTTTTCCCAAATGTTGTTGCGTGTCAGAT

TCACTGGGGCAATCTATCCAAGATCGGGATTGTAGGAACAGGGAGTGCCAGCTACAAGGTGATGACTAGGCCAAGCCACC

AGACTCTGGTTATAAAGTTAATGCCAAATATAACGGCCATCGACAATTGTACAAAGTCAGAGATTGCAGAGTACAAGAGA

TTGCTGATCACAGTGTTAAAGCCTGTAGAGGATGCTCTGTCGGTGATAACCAAGAATGTAAGACCAATTCAAACTCTAAC

ACCTGGGCGTAGAACCCGCCGTTTTGCTGGAGCTGTTCTGGCCGGGGTAGCACTTGGAGTTGCGACAGCCGCTCAGATAA

CTGCAGGAGTCGCCCTTCATCAATCATTGATGAACTCCCAAGCAATTGAGAGTTTAAAAACCAGTCTTGAGATGTCGAAT

CAGGCAATAGAAGAAATCAGACTTGCAAATAAGGAGACCATACTGGCAGTACAGGGCGTCCAGGATTATATCAACAATGA

GCTCGTCCCTTCTGTTCATAGAATGTCATGCGAGCTGGTAGGTCACAAGCTCGGCCTCAAGCTCCTTAGGTACTACACCG

AGATCCTGTCCATATTCGGGCCCAGTCTTCGAGACCCGATATCTGCCGAAATATCAATCCAGGCACTTAGTTATGCATTA

GGCGGAGACATTAATAAAATCCTGGACAAGCTTGGGTATAGCGGTGGGGATTTCCTTGCCATCCTAGAAAGCAAGGGAAT

AAAGGCCCGGGTTACATATGTGGACACAAGAGATTACTTTATAATCCTTAGCATCGCCTACCCAACCTTATCTGAGATCA

AGGGAGTGATAGTTCACAAGATAGAAGCTATAACATACAACATTGGGGCACAGGAGTGGTATACTACTATCCCTAAATAT

GTAGCCACTCAGGGGTATCTGATATCGAACTTTGATGAGACGTCATGCGTATTCACTCCAGAGGGGACAGTTTGCAGCCA

GAATGCGTTGTACCCAATGAGCCCATTGCTTCAGGAATGTTTCAGGGGGTCAACAAAATCGTGCGCCAGAACCTTAGTTT

CAGGGACCATAAGTAATAGATTTATCCTATCAAAAGGGAACCTGATTGCAAATTGTGCGTCAGTTTTGTGCAAATGTTAC

ACAACAGAGACAGTTATCAGCCAAGATCCTGACAAACTACTAACTGTTGTAGCATCCGACAAGTGTCCTGTAGTTGAGGT

GGATGGAGTGACAATACAGGTCGGCAGTCGAGAGTATCCGGATTCTGTATACTTACACAAAATAGACTTAGGTCCAGCCA

TCTCCCTAGAAAAACTGGATGTAGGCACCAATTTAGGCAATGCAGTCACAAGACTGGAGAATGCAAAGGAGCTCCTAGAT

GCATCAGACCAAATACTGAAGACTGTTAAAGGGGTACCTTTCAGTGGGAATATGTACATAGCACTGGCAGCTTGCATAGG

AGTATCCCTGGGCCTTGTCACATTAATATGCTGCTGTAAGGGGAGGTGTAAGAACAAGGAAATCCCTATCTCCAAAATCA

ACCCAGGGCTCAAACCCGACCTGACCGGGACCTCAAAGTCGTACGTAAGATCACTGTAGTCAGAATCACCTGGATCATCT

GGCATCACACACAGACATGCACGACACAGGCAGTCCGAGGACGCAAGAAACCCAGCCTCCGGTCACCCACCCGACCCCAC

TCCACGCTCCACCACACATTAGTCATCAAACAAAACTTAGGACGAAAGGTCAATCACCATGTCCGCACAAAGGGAGAGGA

TCAATGCCTTCTACAAAGACAATCCTCACAATAAGAACCATAGGGTGATCCTGGATAGAGAACGCTTGGTCATTGAAAGA

CCCTACATCTTGCTTGGAGTCCTGCTGGTAATGTTCCTGAGTCTAATCGGACTGCTGGCCATTGCAGGGATCAGGCTTCA

CCGGGCCACCGTTGGAACTTCAGAGATCCAGAGTCGGCTGAATACCAATATTAAGTTGACCGAATCTATTGATCACCAGA

CTAAGGATGTCTTAACTCCCCTTTTTAAAATCATTGGCGATGAAGTCGGCATCAGAATTCCACAGAAATTCAGTGATCTT

GTCAAGTTCATCTCCGATAAGATTAAATTCCTCAACCCTGATAGAGAGTATGATTTCAGAGATCTCCGGTGGTGTATGAA

TCCCCCCGAGAGAGTCAAAATTAATTTTGATCAGTTTTGTGAGTACAAGGCTGCGGTTAAGTCAATTGAACATATATTTG

AGTCACCACTCAACAAGTCAAAAAAGCTGCAATCTTTGACTCTCGGGCCCGGAACAGGCTGTCTAGGCAGGACAGTAACA

AGAGCCCATTTCTCAGAACTTACAATGACCTTAATGGACCTGGATCTAGAGATGAAGCACAACGTGTCCTCAGTGTTTAC

CGTAGTTGAAGAGGGATTATTCGGAAGAACATATACCGTCTGGAGATCCGATGCCAGGGATCCGAGCACCGATCTAGGTA

TCGGCCATTTTTTAAGAGTCTTCGAGATTGGACTGGTAAGAGATCTTGGGCTGGGTCCCCCTGTTTTTCATATGACCAAC

TATCTCACGGTGAACATGAGTGATGACTATCGGAGATGTCTTTTAGCGGTAGGGGAGTTGAAGTTGACAGCCCTATGCAC

CTCATCTGAGACTGTGACACTGAGTGAGAGAGGAGTTCCAAGGAGGGAACCTCTTGTGGTTGTGATACTTAATCTAGCTG

GACCCACTCTAGGGGGAGAGCTATACAGTGTCTTGCCTACCTCTGATCTCATGGTGGAGAAACTCTATTTGTCTTCACAT

AGAGGGATCATCAAAGATGATGAGGCCAATTGGGTAGTGCCGTCTACCGATGTTCGTGATCTTCAGAACAAAGGTGAATG

TCTGGTGGAAGCATGCAAGACTCGACCTCCTTCATTTTGCAATGGCACAGGATCAGGCCCGTGGTCAGAAGGGAGAATCC

CTGCCTACGGGGTGATCAGGGTCAGTCTTGACTTAGCGAGTGACCCGGATGTAGTTATCACTTCAGTGTTTGGCCCACTG

ATACCTCACCTATCCGGCATGGATCTTTACAACAACCCGTTTTCAAAAGCTGTATGGTTGGCTGTACCACCTTATGAGCA

GTCATTTCTAGGAATGATAAATACAATTGGATTCCCTAACAGAGCAGAGGTTATGCCGCACATTTTGACCACAGAGATCA

GAGGCCCTCGGGGTCGTTGCCATGTTCCCATAGAATTGTCCCGCCGGGTTGATGACGATATCAAGATCGGGTCCAACATG

GTCATATTGCCGACGATGGACCTGAGGTATATTACAGCCACTTATGATGTTTCCAGGAGCGAGCATGCAATCGTGTACTA

TATCTATGACACGAGTCGCTCATCATCTTACTTCTACCCAGTTCGACTGAATTTCAAAGGCAATCCTCTCTCTTTGAGGA

TAGAGTGTTTCCCCTGGCGTCATAAGGTGTGGTGCTACCATGATTGTCTTATATACAACACCATAACAGGTGAAGAGGTC

CATACGAGAGGGCTGACCGGCATAGAGGTAACATGCAATCCAGTCTGAGCAGAGCTACGACCGTCACTCAGGCAGTCCCT

TGAGTCGCCACCGAGTCCAAGCAGCACAGCCTGGGACAATCAACGGCACAGCCCAGCCAACAATGTTATAAAAAACTTAG

GAGCCAGGGTTGTAGGGGCCATGGACTCTCTATCAGTCAATCAGGTCTTGTACCCTGAGGTTCATCTAGACAGCCCTATT

GTCACAAACAAACTAGTTGCCATCCTTGAGTACTCGGGTATCGACCATAACTATGTTCTTGAAGACCAGACTCTTGTCAA

GAATATTAGGTATAGGCTGGGGTGCGGTTTTTCAAATCAAATGATCATCAATAACAGGGGGGTGGGTGAAACAGTCAATT

CTAAACTTAAAAGTTACCCCCATAATCGTCATATCATATACCCGGATTGCAATAAGGAGTTGTTTTGTATCAAGGATAGC

TGCATATCTAAGAAGCTCTCGGAGCTATTCAAGAAGGGTAATTCCTTGTACTCTAAGATAAGTCACCAGGTACTGGATTG

TCTTAAGAGAGTCAATGGGAAATTAGGTCTGGGCACAGATCTTACCCATGGTCTGAAGGAGGGTATCCTTGACCTAGGGT

TGCACATGCATAGCTCTCAATGGTTCGAGACTTTCCTGTTCTGGTTCACTATCAAGACAGAGATGAGATCAATGATCAAA

GAACAGTCCCATATATGCCACAAGAGGAGGTATAACCCGACTTTTGTGTCGGGGGATGCATTCGAGGTGCTCGTATCGCG

AGACCTCGTTGTGATAATTGATAAGAATACCCAGTATGTCTTCTACTTAACATTTGAGCTGGTCCTTATGTATTGTGATG

TCATAGAGGGTAGACTTATGACGGAGACAGCCATGGCCATAGACCAGAGATATTCAGAACTTCTAAGCCGGGTCAGATAC

TTGTGGGATCTTATTGATGGGTTCTTCCCAACACTAGGCAACACCACATACCAAATTGTTGCTCTGCTTGAACCATTATC

GTTGGCTTATCTTCAACTTCAGGATGTCACTCTGGAGTTAAGAGGTGCTTTTTTAGACCATTGCTTCAAAGAACTCTATG

AGATACTGGAACATTGCGGCATTGACACAGCAGGCACTTACAATTCCATCACTGAAGGATTGGATTACGTATTCATCACC

CACGATATACATCTAACTGGGGAGATTTTTTCATTTTTCCGGAGTTTCGGACACCCCCGTCTAGAAGCAGTCACCGCTGC

GGAAAATGTCAGAAAACATATGAACCAACCGAAGGTAATCAGTTATGAGACTATGATGAAAGGCCATGCGGTATTTTGCG

GGATAATCATAAATGGTTTTCGGGACCGACATGGCGGCAGCTGGCCCCCTGTTGCACTGCCAGAACATGCCTCTGCTGCG

ATCCGGAATGCGCAGGCATCAGGTGAGGGACTGACCCATGACCTGTGTATAGACAACTGGAAATCCTTTGTTGGATTCAA

ATTTGGCTGCTTTATGCCACTCAGCCTAGATAGTGATTTGACCATGTATCTCAAAGACAAGGCATTGGCTGCACTGAAGA

ATGAGTGGGATTCAGTTTACCCGAAAGAATACCTCCGTTATAACCCACCTAGAGGGACAGAGTCAAGGCGATTGGTAGAG

GTATTCCTGAATGACTCCAGCTTTGATCCTTATAACATGATAATGTACGTGGTGAATGGCTCCTACCTTAAAGACCCTGA

GTTTAACCTCTCATACAGCCTAAAAGAGAAGGAGATAAAGGAGACAGGGCGGTTGTTTGCCAAAATGACCTATAAGATGC

GGGCCTGTCAGGTAATCGCTGAAAATCTGATATCGAATGGTGTTGGGAAGTATTTCCGAGACAATGGGATGGCAAAGGAC

GAGCATGACCTAACAAAAGCCCTTCACACCCTGGCAGTCTCAGGTGTTCCCAAAAACAACAAAGATAACCACCGAGGTGG

GCCTCCCAGAAGGACAACAAACCGAGGGGTGAGATCAAGCCAAGGCACCAAAACACAAGATAGAGACAAGGTTCAAGGGG

GACCTATGTACAACTATTTGCGATGCCAACCGATCAGCCCTGATCAGGGTGAGTCATACGAGACTGTTAGTGCATTCATC

ACCGCTGACCTTAAGAAGTATTGCCTGAATTGGAGATACGAGACAATCAGCATATTTGCACAGAGGCTGAATGAAATATA

TGGACTACCATCCTTCTTTCAATGGTTACACAGGATATTGGAAAAATCCGTACTCTACGTCAGTGACCCACATTGTCCTC

CCGATCTAGATAATCATATCCCTCTGGACAGTGTCCCCAATGCCCAAATATTCATCAAGTACCCAATGGGCGGAATAGAA

GGTTATTGCCAAAAACTATGGACAATCAGCACTATACCATACTTGTATCTGGCAGCCTATGAGAGCGGAGTAAGAATCGC

CTCACTGGTGCAGGGTGACAATCAGACAATCGCAGTGACAAAAAGAGTTCCAAGTTCTTGGCCTTATTCACTAAAAAAGA

GGGAGGCCTCCAAGGCCGCTCAAAATTACTTCGTAGTCTTAAGGCAAAGATTGCACGATGTAGGTCATCACTTAAAGGCT

AATGAGACCATAGTATCTTCTCACTTTTTTGTATATTCCAAAGGGATTTATTATGACGGCCTGCTAGTCTCACAATCACT

AAAGAGCATCGCCAGATGTGTCTTCTGGTCCGAGACTATAGTGGATGAAACCAGAGCGGCGTGCAGTAATATTGCAACAA

CTGTCGCTAAGAGTATAGAGAGGGGTTATGATAGGTACCTTGCATATTCTTTGAATATCCTCAAGATTTTCCAACAGATC

CTTATATCCCTTAACTTCACTATTAACACAACAATGACTCAGGATGTCGTGGCACCGATCATAGAGAACGGTGATTTGCT

GATAAGGATGGCACTCTTGCCAGCACCCATCGGGGGTCTCAATTATCTTAACATGAGCAGGTTATTTGTGAGAAATATCG

GTGACCCGGTCACCTCCTCTATAGCCGACCTGAAGAGGATGATAGATGCCGGGCTAATGCCAGAAGAAACATTGCATCAA

GTGATGACCCAGACCCCGGGAGAATCATCCTACCTAGATTGGGCAAGTGACCCTTATTCTGCCAACCTACCCTGCGTACA

GAGTATCACTCGCCTTCTCAAAAACATCACTGCACGATATATTTTAATCAGCAGCCCAAACCCGATGCTGAAAGGGTTAT

TTCATGAGGGGAGTAGAGATGAAGACGAGGAGCTTGCAAGTTTCCTAATGGATCGGCATATAATTGTTCCAAGAGCTGCA

CATGAAATCTTAGACCATAGTATAACCGGAGCAAGAGAAGCTATAGCCGGGATGTTGGACACCACCAAGGGTCTGATTAG

AACAAGTATGAAACGGGGCGGCCTCACCCCTCGAGTCTTAGCCCGCCTTTCCAATTATGATTATGAACAATTCCGATCCG

GGATAACACTATTAACAAAGAAAGGGCAGTGTTATCTCATTGACAAGGACTCGTGCTCGGTGCAGCTCGCTATAGCTCTG

AGGGGCCACATGTGGGCTAAGTTGGCTCGCGGGAGACCAATCTATGGGTTGGAGGTGCCTGATGTATTAGAATCGATGAA

CGGCTACCTTATCAAACGCCACGAGTCGTGTGCTATCTGTGAAACGGGCTCAAGTCACTACGGATGGTTTTTCGTCCCTG

CAGGGTGCCAGCTTGACGATGTATCTAGAGAGACTTCGGCTCTTCGTGTACCTTATGTCGGATCAACAACTGAGGAAAGG

ACAGACATGAAACTTGCATTTGTTAGGTCTCCAAGTAGATCCCTCAAATCAGCAGTTAGAATTGCAACAGTTTACTCATG

GGCTTACGGGGATGATGAGAAATCATGGAGTGAAGCCTGGATGCTAGCTAGGCAGAGAGCCAATATCACCTTAGATGAAT

TGAGAATGATCACTCCAGTCTCCACATCCACCAACTTAGCCCACCGGTTGAGGGATCGGAGTACCCAGGTGAAATACTCG

GGGACATCCCTTGTGAGGGTTGCAAGATACACAACCATCTCCAATGATAACTTGTCGTTTGTGATATCTGAGAAAAAAGT

GGATACCAACTTCATTTATCAGCAAGGGATGTTGCTTGGTCTTGGGATCCTTGAGAACCTCTTTAGGCTAGAGGCCACCA

CCGGAGTATCCAACACAGTGCTACACCTGCACGTGGAAACAGAATGTTGTGTTGTACCTATGGTGGATCACCCAAGGATA

CCGAGTCTCCGTAATATAAAAGTTACGGATGAGCTATGCACAAACCCTTTGATCTACGACAGGTCCCCCATCATAGAGCA

CGATGCAACCCGGCTATATTCACAGAGCCACAGGAGACATTTGGTGGAGTTTGTTACCTGGTCAACAAGCCAGCTTTACC

ATATACTGGCTAAGTCTACAGCAATGTCCATGATTGAATTGATCACGAGATTCGAGAAAGATCACATGAATGAAATAGCC

GCCCTGATTGGCGATGACGACATCAACAGCTTCATCACAGAATTTTTGCTAGTGGAGCCTAGATTATTTATAGTTTACCT

TGGTCAGTGTGCTGCCATCAATTGGGCCTTTGATATACATTATCATCGGCCCTCGGGCAAGTACCAGATGGGGGAACTTC

TCTACTCTCTGCTCTCTCGGATGAGCAAAGGAGTATATAAGATCTTTACTAATGCTTTGAGCCACCCCAAAGTTTACAAG

AAATTTTGGCGAAGTGGCATCATTGAGCCGATTCATGGCCCATCCTTGGATACACAGAATTTACATGTCACTGTCTGTGA

CATGATATATGGGTCCTATGTCACCTATTTGGATCTTTTGCTGAATGATGAGCTAGATGATTACCCGTATTTGCTCTGCG

AGAGTGATGAGGACGTGGTCACAGACAGGTTCGACAACATTCAAGCCAAACATCTCTGTGTATTGGCCGATGTATATTGC

AGCTCCAAGAGGTGTCCTTCAATAATCGGGATGTCTCCTATAGAAAAATGTACCATTCTCACACATTACATCAAGGGAGA

ATCAGTACAATCCCCGTCTGGGACCTCATGGAACACTGATCCCCTTGTAGTAGATCATTACTCATGCTCTCTGACCTACC

TTCGCCGCGGTTCCATCAAACAAATCAGGTTGAGGGTGGATCCTGGGTTTGTATTCGAGGCGTTGACAGACGTCGACTTT

AAACAACCTCGCAAAGCTAAGTTAGATATATCGGTCGTTGGGTTGACTGATTTCTCTCCCCCTTACGACAACGTCGGTGA

TTTTTTAGGGACTATCAACACATTGAGGCACGATCTGCCTGTCACCGGAACCGGGGTCTCGAACTATGAAGTCCACGCTT

ATCGTAGAATTGGCCTGAATTCATCAGCATGTTATAAAGCCGTAGAGATCTCCACGTTAATCAAGCCATCCTTAGAAGTC

GGAGAGCATGGATTGTTCTTAGGAGAAGGTTCCGGTTCAATGCTGGCTGCGTACAAAGAAGTTCTTAAATTAGCAAATTG

TTACTACAACAGCGGAGTAACAGCGGAGGGCAGAGCCGGACAGAGGGAAATATCTCCCTATCCTTCAGAGATGAGCCTAG

TAGAGAATCAAATGGGGATAGAGCGGAGTGTTAAAGTGCTGTTCAACGGGAAACCTGAAGTAACTTGGGTGGGGACCACA

GATTGCTACAAGTACATAATCAGTAACATCCAAACCTCTAGTCTGGGTTTCATACATTCAGATATTGAGACGCTCCCAAC

CAAAGATGCTGTTGAGAAATTGGAGGAATTTGCCTCTATCCTATCCTTATCCTTGATTTTGGGGAAAATCGGCTCTATTA

CAGTTGTCAAAATTATGCCCATTAGCGGGGATTTTACCCAAGGCTTCATAGCCTATGCCATCCAATATTTTAGGGAGAGC

CTGCTTGCCTATCCGAGATATAGTAACTTCATCTCGACTGAGTGTTACCTTATTATGATAGGATTAAAGGCCAATCGATT

GATAAACCCAGAAGCCATTAAGCAAAGCATAATCAGAGCGGGGGTCAGGACTGCACCAGGACTTGTGAGCCATGTATTAT

CAGAGAAACAAAAAGGTTGTATTCAATCTTTCTTGGGTGATCCTTATATCCAAGGAGACTTCAATAAGCACCTTAAATCT

CTAACCTCTATTGAGAAAATCCTGGTAAATTGTGGTCTCTCGATCAATGGCATAAAAATCTGTAGAGATCTAATCCACCA

TGATATCGCCTCCGGTCCAGACGGCCTTATGAGCTCTACAATTATTTTATATAGGGAATTGGCTCATTTCAAAGATAATA

TAAGGAGTCAGCACGGAATGTTCCACCCCTACCCGGTACTGGCCAACAGCAGGCAACGTGAATTAATCCTTCGAATAGCC

AAGAAATTCTGGGGGTATGTCTTGCTATATTCTGATGACCCGGCACTAATCAGACAAACAATCAAGAACTTGAAACGGAA

TCATCTAACCTTTGACTTACATAGTAACCCATTTATTAAAGGGCTATCCAAGGCTGAGAAACTGCTGGTGCGGACGAGTT

CACTAAGAAGAGAATGGTTGTTCACTCTCGAAACGAAAGAAGTAAAAGAGTGGTTCAAATTGGTAGGTTACAGTGCACTC

ATCAGAGGCTAGTCAGCTATGCATCTGCCCCCTTCTCCTCCGCCATGAGACCCCACTGGCGATCCAGAAGATTAAAGAAA

ACTACATATTGGATAAGTATCTATTCCCAGCTTTGTCTGGT

>MF443346.1/goat/China/2014-03-31

ACCAAACAAAGTTGGGTAAGGATAGATCTTATAATAACTATGGACTGGCAAACTTAGGAGTAAAGATCCTACTGTCGGGG

GGAGGAGGAGGAGCAAGATCCTTGATCATGGCGACTCTCCTTAAAAGCTTAGCATTGTTCAAAAGGAACAAAGACAAAGC

GCCGACAGCATCAGGTTCAGGAGGGGCCATCCGGGGGATTAAGAATGTTATCATAGTCCCGATTCCCGGAGACTCGTCCA

TCACTACCCGTTCAAGACTGCTCGACAGGCTTGTCAGATTGGCCGGAGATCCTGACATCAACGGGTCAAAGCTGACCGGG

GTGATGATCAGCATGTTATCCTTGTTCGTAGAGTCACCCGGGCAATTGATACAGCGAATCACAGATGATCCGGATGTTAG

TATCCGCCTTGTTGAGGTAGTTCAAAGTACTAGGTCCCAGTCCGGGTTGACCTTTGCATCACGTGGTGCTGATTTAGACA

ACGAGGCAGACATGTATTTTTCAACTGAGGGGCCCTCGAGTGGAGGTAAGAAAAGGATCAACTGGTTTGAGAACAGAGAA

ATAATAGACATAGAGGTGCAGGATCCAGAAGAGTTCAATATGTTGTTAGCCTCCATACTAGCACAAGTCTGGATCCTTCT

GGCCAAGGCTGTTACGGCACCAGATACGGCAGCTGACTCAGAACTGAGAAGGTGGGTTAAATACACACAACAAAGGAGAG

TGATTGGGGAATTTCGCCTTGACAAGGGGTGGCTGGACGCAGTCCGCAACAGAATTGCAGAAGATCTATCACTCCGGCGG

TTCATGGTATCTCTCATACTTGACATCAAGAGGACCCCAGGCAACAAGCCAAGGATTGCAGAAATGATCTGCGACATTGA

CAACTATATTGTCGAAGCAGGACTCGCCAGTTTCATCCTTACTATCAAATTTGGTATCGAAACCATGTATCCTGCATTAG

GGCTCCACGAGTTTGCCGGAGAATTGTCCACTATTGAGTCCTTGATGAACTTGTATCAACAGCTAGGAGAGGTTGCACCC

TACATGGTAATTCTAGAGAATTCAATTCAGAACAAGTTTAGTGCAGGAGCTTATCCCCTCCTCTGGAGCTATGCGATGGG

TGTCGGAGTCGAGCTGGAGAACTCAATGGGGGGCTTGAACTTTGGCAGGTCATATTTTGACCCTGCCTATTTTCGTCTCG

GACAGGAGATGGTCAGAAGATCTGCAGGAAAGGTCAGCTCTGTGATTGCGGCTGAGCTCGGCATCACAGCAGAGGAAGCT

AAACTAGTCTCGGAAATCGCCTCGCAGGCTGGGGACGAAAGAACCGCCAGAGGGACTGGGCCTCGACAGGCGCAGGTTTC

CTTCCTTCAGCACCAAACAGGAGGGGGAGAGTCGTCCGCACCAGCGACCAGAGAAGGGGTCAAAGCTGCGATCCCAAACG

GAGCTGAAGAAAGGGACAGAAAGCAAACACGCCCAGGAAGGCCCAGAGGAGAGACCTCCGGCCAACTGCTCCTGGACATC

ATGCCAGAGGATGAGATCTTGCGAGAGTCTGGTCAAAACCCTCGTGAGGCTCAAAGATCGGCCGAGGCACTCTTCAGGCT

GCAGGCCATGGCCAAGATTCTGGAGGACCAGGAGGAGGGAGAAGACAACAGTCAGGTCTACAACGACAAGGATCTCCTCG

GCTAAGCAGACGCACCCTCTGTCGAAATCAGTGACAAGACATCCCCTACCAGTATTATAAAAAACTTAGGACCCAGGTCC

AAGCAAGCACACATCGACACTCCAACCAGTCGAGCGGAGACCACCGATGGCAGAAGAACAAGCATACCATGTCAACAAGG

GGCTGGAATGTATCAAGTCTCTCAAAGCCTCACCCCCGGATCTATCCACCATCAGAGATACCATCGAGAGCTGGAGAGAG

GGGCTTAGCCCCTCGGGCCGTGCAACACCGAACCCTGATACGTCCGAGGGAGATCATCAGAATATCAACCAATCATGCTC

ACCAGCAATCGGACCAAACAAAGTCTACTTGTCTCCTGAAGATAATCTCGGATTTAGAGAGATCACTGGCAACGACTGTG

AGGCTGGGCTCGGAGGGGTCCAGAGAGAAGGACCCAACTCTCAAGTACAGCGTTACCATGTTTATAGTCACGGGGGTGAA

GAGATTGAAGGACTCGAGGATGCTGACTCTCTCGTGGTTCAAGCAGATCCTCCAGTTGCTAACGTCTTCAATGGAGGAGA

GGATGGATCTGACGACAGCGATGTGGACTCTGGCCCAGATGATCCCGGCAGAGATACTCTATATGACCGGGGATCTGTTG

CCGGCAATGGTATCGCTAGGTCCACAGATGTCGAAAAACTAGAAGGTGCTGATATTCAAGAAGTTCTTAACTCCCAGAAA

GGCAAAGGAGGAAGATTCCAGGGCGGGAAAACCTTGCGAGTCCCGGAAATACCCGATGTCAAGCACTCCAGACCATCAGC

CCAATCAATTAAAAAGGGCACAGACGGGAACTCAGTCTCATCTGGAACGGTGATAGAGTGTTTATCGATAAGTGGTGCAA

CCCAAACTGTGCCAGAGTCAAGATGGGAGTCATCAGAGCAAAATGCGTCTGTGGGGAGTGTCCTCAAGTCTGCGAGGAGT

GCAAAGACGATCCAGGGGTCGACACAAGAATCTGGTACCATAGCATCACTGACTCAGCCTAAAGAGAATGACTCCGAGTA

TGAGTATGAGGATGACCTATTTACAGAGATTCAGGACATCCGTGCAAGCATTGCCAAGATCCATGATGACAATAAAACTA

TCCTCTCAAAGCTTGATTCTATACTGTTATTGAAAGGGGAAGTCGACACTATCAAGAAACAAATCAGCAAGCAGAATATA

AGTATATCCACCATTGAGGGCCATCTCTCCAGTATAATGATAGCCATCCCTGGCTTTGGGAAGGACATCAAGGACCCAAC

ATCCGAAGTCGAGTTGAACCCAGATCTAAGACCTATAATAAGCCGTGACTCTGGCAGAGCTCTCGCGGAGGTCCTCAAGA

AACCCGCTGTGGATAGGTCTCAGAAAATTGGAACCAAAGCCAACTCCAGCTCAAAGGGTCAGCTTCTTAAGGATCTCCAG

CTAAAACCTGTCGACAAGCAGGCAAGCTCTGCAATCGGGTTTGTCCCGTCCGACCATGAATCATCCAGAAATGTCATCCG

CTCCATAATCAAGTCGAGCAAGCTAAATATTGATCACAAGGACTATCTTCTAGATTTACTAAATGATGTGAAAGGCTCCA

AGGATCTTAAGGAATTTCACAAGATGCTAACAGCAATTCTCGCCAAGCACCCGTAACACATCCTCCAGTCATCATCTCAT

ACTCGACTAAAAACATCCTTTCAATCAGGCTATTACAAAAAACTTAGGAGCAAGGGCAACTGAGCTTCGCAGACAGGATG

ACCGAGATCTACGACTTCGATAAATCAGCATGGGATGTCAAAGGGTCAATTGCCCGCATAGAACCCACCACCTACCACGA

CGGCCGACTGATACCCCAGGTGAGGGTCATCGATCCTGGTCTGGGAGACAGAAAAGATGAGTGCTTTATGTACCTGTTTC

TCCTAGGAGTGATTGAGGACAACGACCCCCTGTCTCCCCCCGTCGGAAGAACCTTTGGCTCTCTACCTCTAGGGGTTGGT

AGGTCAACTGCCAAGCCAGAAGAACTACTAAGGGAGGCCACAGAATTAGATATAGTGGTGAGGCGCACTGCAGGAGTAAA

TGAGAAACTGGTATTTTACAACAACACTCCGCTGTCCTTGTTAACGCCCTGGAAGAAAGTTCTGACAACCGGAAGTGTGT

TTAGCGCTAACCAGGTTTGCAATGCAGTCAACCTAGTCCCGCTTGATACTCCCCAGAGATTCAGGGTTGTGTACATGAGC

ATAACTAGATTATCAGACAATGGTTACTATAGTGTGCCCAGAAGAATGCTGGAGTTCCGCTCGGCCAATGCAGTAGCCTT

CAACATCTTGGTTACACTAAGAATTGAAAATGGCACAAACCCTAGAAGATACATAGTCGGCTCATGGGAGAATTCAGAGG

TCACATTTATGGTACACGTGGGCAACTTTAGGAGAAAGAAGAACGAAGTATACTCTGCTGATTACTGCAAAATGAAGATT

GAAAAGATGGGTTTAGTTTTTGCCCTGGGTGGAATAGGCGGAACAAGTCTCCATATTAGAAGCACCGGGAAAATGAGCAA

AACCCTCCATGCACAGCTGGGGTTCAAGAAAATCTTATGTTACCCCCTAATGGATGTCAATGAGGATCTTAACCGATATC

TCTGGCGGGCAGAGTGCCGAATAGTCAAAATCCAAGCTGTCTTACAGCCATCAGTACCCCAAGAGTTCCGTGTCTACGAT

GATGTCATCATCAACGATGATCAAGGCTTGTTCAAGATCTTGTAGTTCATTTGCAAAGGAACGAACCCACCCCCAGGAAG

GGGCCCCACCCTCCCCCC-CCAAAAAACACACTCCAAGGCTCCCCGGCACAGCGCCAACACAGCTACTCCCGCGATGGAA

CGAGGTCGGCC-CCCGAACAAATCCGGTCCAGGGGGGGGGGCCCCCCCCCTGACCAACCGGGCGCCGACCGAGCAGGCCG

GCGAGAAGCCCCCAAGAGGCCCCGACGAAGACCGAGCACCCCATCCCCCC-AAAAACGTCCCCC-AACACGCCCGAGGCT

AGCCACCGGACACCCCCAGACCCGCCACCGGCGAGACAGAACCCCAAGGGGGGACCCTCCAAACCAAAGGAATCACCGGG

GTAGGCTCCGCCGAACCAGCACAGACCCGACCCAAGCAAGGCGGACCCCCAACACGAAACTGCCCAGCACACCCTACTCA

TCATCAAACAAAACTTAGGGGCCAAGTCCATACGAGAGGCCCTAGTCCGTCAGGCGGAGGGACGCAGAAAGGAAGGAGAC

ACGACCGTCCCAGAAGGAGGACACCCAACCACCGAAACGCCCGCCCCCAGGGAAGGAG--CCA-CCCCCACGCAGCGAAG

GAGCGGGTGGGAGACCTCGGGGAGGGGCCGACCGGGACCCGGGGGCCGGCGGCGCCGGCGGTGAGCAGGCCGGGCGCATA

CCTGACCCCACCGGAGGATCCGAGGAGGGCCCGACCCCGCGTGGACCAAACGGCAGGGAGCCCAACCCCACCAAAAAGAG

AGGCGAGGCCCCCCCTCCCTCCCCCCCCGCACACAACCCGCAGACAGACCGAGGGAGGGCCATCCCCCAGCGAGACAGGC

GGCTCCCTCGCAGCCGAGCCGCAGCTCCCCAGAATAGGACACCCCCGCACACTCCGAACACCCCCTCGCCCCCCGCCACC

GGGACAGGGGCTCCACCACCACAGACCGGAGCCACCGACAGCCAGCCAGAGGTCCCCTCCAGTCCCGCCCGGCGCCCCCC

CGAAGAGGCCCCGGCAAGGCCCCCCAGCCCGACGTCCCCCGCACGCACCGAAAGAAGGACCCGCCCGAGAACGTGCGGCA

CATCCATGCGCAAACATCATGACGCGGGTCGCAATTTTGACATTTCTGTTTCTTTTCCCAAATGTTGTTGCGTGTCAGAT

TCACTGGGGCAATCTATCCAAGATCGGGATTGTAGGAACAGGGAGTGCCAGCTACAAGGTGATGACTAGGCCAAGCCACC

AGACTCTGGTTATAAAGTTAATGCCAAATATAACGGCCATCGACAATTGTACAAAGTCAGAGATTGCAGAGTACAAGAGA

TTGCTGATCACAGTGTTAAAGCCTGTAGAGGATGCTCTGTCGGTGATAACCAAGAATGTAAGACCAATTCAAACTCTAAC

ACCTGGGCGTAGAACCCGCCGTTTTGCTGGAGCTGTTCTGGCCGGGGTAGCACTTGGAGTTGCGACAGCCGCTCAGATAA

CTGCAGGAGTCGCCCTTCATCAATCATTGATGAACTCCCAAGCAATTGAGAGTTTAAAAACCAGTCTTGAGATGTCGAAT

CAGGCAATAGAAGAAATCAGACTTGCAAATAAGGAGACCATACTGGCAGTACAGGGCGTCCAGGATTATATCAACAATGA

GCTCGTCCCTTCTGTTCATAGAATGTCATGCGAGCTGGTAGGTCACAAGCTCGGCCTCAAGCTCCTTAGGTACTACACCG

AGATCCTGTCCATATTCGGGCCCAGTCTTCGAGACCCGATATCTGCCGAAATATCAATCCAGGCACTTAGTTATGCATTA

GGCGGAGACATTAATAAAATCCTGGACAAGCTTGGGTATAGCGGTGGGGATTTCCTTGCCATCCTAGAAAGCAAGGGAAT

AAAGGCCCGGGTTACATATGTGGACACAAGAGATTACTTTATAATCCTTAGCATCGCCTACCCAACCTTATCTGAGATCA

AGGGAGTGATAGTTCACAAGATAGAAGCTATAACATACAACATTGGGGCACAGGAGTGGTATACTACTATCCCTAAATAT

GTAGCCACTCAGGGGTATCTGATATCGAACTTTGATGAGACGTCATGCGTATTCACTCCAGAGGGGACAGTTTGCAGCCA

GAATGCGTTGTACCCAATGAGCCCATTGCTTCAGGAATGTTTCAGGGGGTCAACAAAATCGTGCGCCAGAACCTTAGTTT

CAGGGACCATAAGTAATAGATTTATCCTATCAAAAGGGAACCTGATTGCAAATTGTGCGTCAGTTTTGTGCAAATGTTAC

ACAACAGAGACAGTTATCAGCCAAGATCCTGACAAACTACTAACTGTTGTAGCATCCGACAAGTGTCCTGTAGTTGAGGT

GGATGGAGTGACAATACAGGTCGGCAGTCGAGAGTATCCGGATTCTGTATACTTACACAAAATAGACTTAGGTCCAGCCA

TCTCCCTAGAAAAACTGGATGTAGGCACCAATTTAGGCAATGCAGTCACAAGACTGGAGAATGCAAAGGAGCTCCTAGAT

GCATCAGACCAAATACTGAAGACTGTTAAAGGGGTACCTTTCAGTGGGAATATGTACATAGCACTGGCAGCTTGCATAGG

AGTATCCCTGGGCCTTGTCACATTAATATGCTGCTGTAAGGGGAGGTGTAAGAACAAGGAAATCCCTATCTCCAAAATCA

ACCCAGGGCTCAAACCCGACCTGACCGGGACCTCAAAGTCGTACGTAAGATCACTGTAGTCAGAATCACCTGGATCATCT

GGCATCACACACAGACATGCACGACACAGGCAGTCCGAGGACGCAAGAAACCCAGCCTCCGGTCACCCACCCGACCCCAC

TCCACGCTCCACCACACATTAGTCATCAAACAAAACTTAGGACGAAAGGTCAATCACCATGTCCGCACAAAGGGAGAGGA

TCAATGCCTTCTACAAAGACAATCCTCACAATAAGAACCATAGGGTGATCCTGGATAGAGAACGCTTGGTCATTGAAAGA

CCCTACATCTTGCTTGGAGTCCTGCTGGTAATGTTCCTGAGTCTAATCGGACTGCTGGCCATTGCAGGGATCAGGCTTCA

CCGGGCCACCGTTGGAACTTCAGAGATCCAGAGTCGGCTGAATACCAATATTAAGTTGACCGAATCTATTGATCACCAGA

CTAAGGATGTCTTAACTCCCCTTTTTAAAATCATTGGCGATGAAGTCGGCATCAGAATTCCACAGAAATTCAGTGATCTT

GTCAAGTTCATCTCCGATAAGATTAAATTCCTCAACCCTGATAGAGAGTATGATTTCAGAGATCTCCGGTGGTGTATGAA

TCCCCCCGAGAGAGTCAAAATTAATTTTGATCAGTTTTGTGAGTACAAGGCTGCGGTTAAGTCAATTGAACATATATTTG

AGTCACCACTCAACAAGTCAAAAAAGCTGCAATCTTTGACTCTCGGGCCCGGAACAGGCTGTCTAGGCAGGACAGTAACA

AGAGCCCATTTCTCAGAACTTACAATGACCTTAATGGACCTGGATCTAGAGATGAAGCACAACGTGTCCTCAGTGTTTAC

CGTAGTTGAAGAGGGATTATTCGGAAGAACATATACCGTCTGGAGATCCGATGCCAGGGATCCGAGCACCGATCTAGGTA

TCGGCCATTTTTTAAGAGTCTTCGAGATTGGACTGGTAAGAGATCTTGGGCTGGGTCCCCCTGTTTTTCATATGACCAAC

TATCTCACGGTGAACATGAGTGATGACTATCGGAGATGTCTTTTAGCGGTAGGGGAGTTGAAGTTGACAGCCCTATGCAC

CTCATCTGAGACTGTGACACTGAGTGAGAGAGGAGTTCCAAGGAGGGAACCTCTTGTGGTTGTGATACTTAATCTAGCTG

GACCCACTCTAGGGGGAGAGCTATACAGTGTCTTGCCTACCTCTGATCTCATGGTGGAGAAACTCTATTTGTCTTCACAT

AGAGGGATCATCAAAGATGATGAGGCCAATTGGGTAGTGCCGTCTACCGATGTTCGTGATCTTCAGAACAAAGGTGAATG

TCTGGTGGAAGCATGCAAGACTCGACCTCCTTCATTTTGCAATGGCACAGGATCAGGCCCGTGGTCAGAAGGGAGAATCC

CTGCCTACGGGGTGATCAGGGTCAGTCTTGACTTAGCGAGTGACCCGGATGTAGTTATCACTTCAGTGTTTGGCCCACTG

ATACCTCACCTATCCGGCATGGATCTTTACAACAACCCGTTTTCAAAAGCTGTATGGTTGGCTGTACCACCTTATGAGCA

GTCATTTCTAGGAATGATAAATACAATTGGATTCCCTAACAGAGCAGAGGTTATGCCGCACATTTTGACCACAGAGATCA

GAGGCCCTCGGGGTCGTTGCCATGTTCCCATAGAATTGTCCCGCCGGGTTGATGACGATATCAAGATCGGGTCCAACATG

GTCATATTGCCGACGATGGACCTGAGGTATATTACAGCCACTTATGATGTTTCCAGGAGCGAGCATGCAATCGTGTACTA

TATCTATGACACGAGTCGCTCATCATCTTACTTCTACCCAGTTCGACTGAATTTCAAAGGCAATCCTCTCTCTTTGAGGA

TAGAGTGTTTCCCCTGGCGTCATAAGGTGTGGTGCTACCATGATTGTCTTATATACAACACCATAACAGGTGAAGAGGTC

CATACGAGAGGGCTGACCGGCATAGAGGTAACATGCAATCCAGTCTGAGCAGAGCTACGACCGTCACTCAGGCAGTCCCT

TGAGTCGCCACCGAGTCCAAGCAGCACAGCCTGGGACAATCAACGGCACAGCCCAGCCAACAATGTTATAAAAAACTTAG

GAGCCAGGGTTGTAGGGGCCATGGACTCTCTATCAGTCAATCAGGTCTTGTACCCTGAGGTTCATCTAGACAGCCCTATT

GTCACAAACAAACTAGTTGCCATCCTTGAGTACTCGGGTATCGACCATAACTATGTTCTTGAAGACCAGACTCTTGTCAA

GAATATTAGGTATAGGCTGGGGTGCGGTTTTTCAAATCAAATGATCATCAATAACAGGGGGGTGGGTGAAACAGTCAATT

CTAAACTTAAAAGTTACCCCCATAATCGTCATATCATATACCCGGATTGCAATAAGGAGTTGTTTTGTATCAAGGATAGC

TGCATATCTAAGAAGCTCTCGGAGCTATTCAAGAAGGGTAATTCCTTGTACTCTAAGATAAGTCACCAGGTACTGGATTG

TCTTAAGAGAGTCAATGGGAAATTAGGTCTGGGCACAGATCTTACCCATGGTCTGAAGGAGGGTATCCTTGACCTAGGGT

TGCACATGCATAGCTCTCAATGGTTCGAGACTTTCCTGTTCTGGTTCACTATCAAGACAGAGATGAGATCAATGATCAAA

GAACAGTCCCATATATGCCACAAGAGGAGGTATAACCCGACTTTTGTGTCGGGGGATGCATTCGAGGTGCTCGTATCGCG

AGACCTCGTTGTGATAATTGATAAGAATACCCAGTATGTCTTCTACTTAACATTTGAGCTGGTCCTTATGTATTGTGATG

TCATAGAGGGTAGACTTATGACGGAGACAGCCATGGCCATAGACCAGAGATATTCAGAACTTCTAAGCCGGGTCAGATAC

TTGTGGGATCTTATTGATGGGTTCTTCCCAACACTAGGCAACACCACATACCAAATTGTTGCTCTGCTTGAACCATTATC

GTTGGCTTATCTTCAACTTCAGGATGTCACTCTGGAGTTAAGAGGTGCTTTTTTAGACCATTGCTTCAAAGAACTCTATG

AGATACTGGAACATTGCGGCATTGACACAGCAGGCACTTACAATTCCATCACTGAAGGATTGGATTACGTATTCATCACC

CACGATATACATCTAACTGGGGAGATTTTTTCATTTTTCCGGAGTTTCGGACACCCCCGTCTAGAAGCAGTCACCGCTGC

GGAAAATGTCAGAAAACATATGAACCAACCGAAGGTAATCAGTTATGAGACTATGATGAAAGGCCATGCGGTATTTTGCG

GGATAATCATAAATGGTTTTCGGGACCGACATGGCGGCAGCTGGCCCCCTGTTGCACTGCCAGAACATGCCTCTGCTGCG

ATCCGGAATGCGCAGGCATCAGGTGAGGGACTGACCCATGACCTGTGTATAGACAACTGGAAATCCTTTGTTGGATTCAA

ATTTGGCTGCTTTATGCCACTCAGCCTAGATAGTGATTTGACCATGTATCTCAAAGACAAGGCATTGGCTGCACTGAAGA

ATGAGTGGGATTCAGTTTACCCGAAAGAATACCTCCGTTATAACCCACCTAGAGGGACAGAGTCAAGGCGATTGGTAGAG

GTATTCCTGAATGACTCCAGCTTTGATCCTTATAACATGATAATGTACGTGGTGAATGGCTCCTACCTTAAAGACCCTGA

GTTTAACCTCTCATACAGCCTAAAAGAGAAGGAGATAAAGGAGACAGGGCGGTTATTTGCCAAAATGACCTATAAGATGC

GGGCCTGTCAGGTAATCGCTGAAAATCTGATATCGAATGGTGTTGGGAAGTATTTCCGAGACAATGGGATGGCAAAGGAC

GAGCATGACCTAACAAAAGCCCTTCACACCCTGGCAGTCTCAGGTGTTCCCAAAAACAACAAAGATAACCACCGAGGTGG

GCCTCCCAGAAGGACAACAAACCGAGGGGTGAGATCAAGCCAAGGCACCAAAACACAAGATAGAGACAAGGTTCAAGGGG

GACCTATGTACAACTATTTGCGATGCCAACCGATCAGCCCTGATCAGGGTGAGTCATACGAGACTGTTAGTGCATTCATC

ACCGCTGACCTTAAGAAGTATTGCCTGAATTGGAGATACGAGACAATCAGCATATTTGCACAGAGGCTGAATGAAATATA

TGGACTACCATCCTTCTTTCAATGGTTACACAGGATATTGGAAAAATCCGTACTCTACGTCAGTGACCCACATTGTCCTC

CCGATCTAGATAATCATATCCCTCTGGACAGTGTCCCCAATGCCCAAATATTCATCAAGTACCCAATGGGCGGAATAGAA

GGTTATTGCCAAAAACTATGGACAATCAGCACTATACCATACTTGTATCTGGCAGCCTATGAGAGCGGAGTAAGAATCGC

CTCACTGGTGCAGGGTGACAATCAGACAATCGCAGTGACAAAAAGAGTTCCAAGTTCTTGGCCTTATTCACTAAAAAAGA

GGGAGGCCTCCAAGGCCGCTCAAAATTACTTCGTAGTCTTAAGGCAAAGATTGCACGATGTAGGTCATCACTTAAAGGCT

AATGAGACCATAGTATCTTCTCACTTTTTTGTATATTCCAAAGGGATTTATTATGACGGCCTGCTAGTCTCACAATCACT

AAAGAGCATCGCCAGATGTGTCTTCTGGTCCGAGACTATAGTGGATGAAACCAGAGCGGCGTGCAGTAATATTGCAACAA

CTGTCGCTAAGAGTATAGAGAGGGGTTATGATAGGTACCTTGCATATTCTTTGAATATCCTCAAGATTTTCCAACAGATC

CTTATATCCCTTAACTTCACTATTAACACAACAATGACTCAGGATGTCGTGGCACCGATCATAGAGAACGGTGATTTGCT

GATAAGGATGGCACTCTTGCCAGCACCCATCGGGGGTCTCAATTATCTTAACATGAGCAGGTTATTTGTGAGAAATATCG

GTGACCCGGTCACCTCCTCTATAGCCGACCTGAAGAGGATGATAGATGCCGGGCTAATGCCAGAAGAAACATTGCATCAA

GTGATGACCCAGACCCCGGGAGAATCATCCTACCTAGATTGGGCAAGTGACCCTTATTCTGCCAACCTACCCTGCGTACA

GAGTATCACTCGCCTTCTCAAAAACATCACTGCACGATATATTTTAATCAGCAGCCCAAACCCGATGCTGAAAGGGTTAT

TTCATGAGGGGAGTAGAGATGAAGACGAGGAGCTTGCAAGTTTCCTAATGGATCGGCATATAATTGTTCCAAGAGCTGCA

CATGAAATCTTAGACCATAGTATAACCGGAGCAAGAGAAGCTATAGCCGGGATGTTGGACACCACCAAGGGTCTGATTAG

AACAAGTATGAAACGGGGCGGCCTCACCCCTCGAGTCTTAGCCCGCCTTTCCAATTATGATTATGAACAATTCCGATCCG

GGATAACACTATTAACAAAGAAAGGGCAGTGTTATCTCATTGACAAGGACTCGTGCTCGGTGCAGCTCGCTATAGCTCTG

AGGGGCCACATGTGGGCTAAGTTGGCTCGCGGGAGACCAATCTATGGGTTGGAGGTGCCTGATGTATTAGAATCGATGAA

CGGCTACCTTATCAAACGCCACGAGTCGTGTGCTATCTGTGAAACGGGCTCAAGTCACTACGGATGGTTTTTCGTCCCTG

CAGGGTGCCAGCTTGACGATGTATCTAGAGAGACTTCGGCTCTTCGTGTACCTTATGTCGGATCAACAACTGAGGAAAGG

ACAGACATGAAACTTGCATTTGTTAGGTCTCCAAGTAGATCCCTCAAATCAGCAGTTAGAATTGCAACAGTTTACTCATG

GGCTTACGGGGATGATGAGAAATCATGGAGTGAAGCCTGGATGCTAGCTAGGCAGAGAGCCAATATCACCTTAGATGAAT

TGAGAATGATCACTCCAGTCTCCACATCCACCAACTTAGCCCACCGGTTGAGGGATCGGAGTACCCAGGTGAAATACTCG

GGGACATCCCTTGTGAGGGTTGCAAGATACACAACCATCTCCAATGATAACTTGTCGTTTGTGATATCTGAGAAAAAAGT

GGATACCAACTTCATTTATCAGCAAGGGATGTTGCTTGGTCTTGGGATCCTTGAGAACCTCTTTAGGCTAGAGGCCACCA

CCGGAGTATCCAACACAGTGCTACACCTGCACGTGGAAACAGAATGTTGTGTTGTACCTATGGTGGATCACCCAAGGATA

CCGAGTCTCCGTAATATAAAAGTTACGGATGAGCTATGCACAAACCCTTTGATCTACGACAGGTCCCCCATCATAGAGCA

CGATGCAACCCGGCTATATTCACAGAGCCACAGGAGACATTTGGTGGAGTTTGTTACCTGGTCAACAAGCCAGCTTTACC

ATATACTGGCTAAGTCTACAGCAATGTCCATGATTGAATTGATCACGAGATTCGAGAAAGATCACATGAATGAAATAGCC

GCCCTGATTGGCGATGACGACATCAACAGCTTCATCACAGAATTTTTGCTAGTGGAGCCTAGATTATTTATAGTTTACCT

TGGTCAGTGTGCTGCCATCAATTGGGCCTTTGATATACATTATCATCGGCCCTCGGGCAAGTACCAGATGGGGGAACTTC

TCTACTCTCTGCTCTCTCGGATGAGCAAAGGAGTATATAAGATCTTTACTAATGCTTTGAGCCACCCCAAAGTTTACAAG

AAATTTTGGCGAAGTGGCATCATTGAGCCGATTCATGGCCCATCCTTGGATACACAGAATTTACATGTCACTGTCTGTGA

CATGATATATGGGTCCTATGTCACCTATTTGGATCTTTTGCTGAATGATGAGCTAGATGATTACCCGTATTTGCTCTGCG

AGAGTGATGAGGACGTGGTCACAGACAGGTTCGACAACATTCAAGCCAAACATCTCTGTGTATTGGCCGATGTATATTGC

AGCTCCAAGAGGTGTCCTTCAATAATCGGGATGTCTCCTATAGAAAAATGTACCATTCTCACACATTACATCAAGGGAGA

ATCAGTACAATCCCCGTCTGGGACCTCATGGAACACTGATCCCCTTGTAGTAGATCATTACTCATGCTCTCTGACCTACC

TTCGCCGCGGTTCCATCAAACAAATCAGGTTGAGGGTGGATCCTGGGTTTGTATTCGAGGCGTTGACAGACGTCGACTTT

AAACAACCTCGCAAAGCTAAGTTAGATATATCGGTCGTTGGGTTGACTGATTTCTCTCCCCCTTACGACAACGTCGGTGA

TTTTTTAGGGACTATCAACACATTGAGGCACGATCTGCCTGTCACCGGAACCGGGGTCTCGAACTATGAAGTCCACGCTT

ATCGTAGAATTGGCCTGAATTCATCAGCATGTTATAAAGCCGTAGAGATCTCCACGTTAATCAAGCCATCCTTAGAAGTC

GGAGAGCATGGATTGTTCTTAGGAGAAGGTTCCGGTTCAATGCTGGCTGCGTACAAAGAAGTTCTTAAATTAGCAAATTG

TTACTACAACAGCGGAGTAACAGCGGAGGGCAGAGCCGGACAGAGGGAAATATCTCCCTATCCTTCAGAGATGAGCCTAG

TAGAGAATCAAATGGGGATAGAGCGGAGTGTTAAAGTGCTGTTCAACGGGAAACCTGAAGTAACTTGGGTGGGGACCACA

GATTGCTACAAGTACATAATCAGTAACATCCAAACCTCTAGTCTGGGTTTCATACATTCAGATATTGAGACGCTCCCAAC

CAAAGATGCTGTTGAGAAATTGGAGGAATTTGCCTCTATCCTATCCTTATCCTTGATTTTGGGGAAAATCGGCTCTATTA

CAGTTGTCAAAATTATGCCCATTAGCGGGGATTTTACCCAAGGCTTCATAGCCTATGCCATCCAATATTTTAGGGAGAGC

CTGCTTGCCTATCCGAGATATAGTAACTTCATCTCGACTGAGTGTTACCTTATTATGATAGGATTAAAGGCCAATCGATT

GATAAACCCAGAAGCCATTAAGCAAAGCATAATCAGAGCGGGGGTCAGGACTGCACCAGGACTTGTGAGCCATGTATTAT

CAGAGAAACAAAAAGGTTGTATTCAATCTTTCTTGGGTGATCCTTATATCCAAGGAGACTTCAATAAGCACCTTAAATCT

CTAACCTCTATTGAGAAAATCCTGGTAAATTGTGGTCTCTCGATCAATGGCATAAAAATCTGTAGAGATCTAATCCACCA

TGATATCGCCTCCGGTCCAGACGGCCTTATGAGCTCTACAATTATTTTATATAGGGAATTGGCTCATTTCAAAGATAATA

TAAGGAGTCAGCACGGAATGTTCCACCCCTACCCGGTACTGGCCAACAGCAGGCAACGTGAATTAATCCTTCGAATAGCC

AAGAAATTCTGGGGGTATGTCTTGCTATATTCTGATGACCCGGCACTAATCAGACAAACAATCAAGAACTTGAAACGGAA

TCATCTAACCTTTGACTTACATAGTAACCCATTTATTAAAGGGCTATCCAAGGCTGAGAAACTGCTGGTGCGGACGAGTT

CACTAAGAAGAGAATGGTTGTTCACTCTCGAAACGAAAGAAGTAAAAGAGTGGTTCAAATTGGTAGGTTACAGTGCACTC

ATCAGAGGCTAGTCAGCTATGCATCTGCCCCCTTCTCCTCCGCCATGAGACCCCACTGGCGATCCAGAAGATTAAAGAAA

ACTACATATTGGATAAGTATCTATTCCCAGCTTTGTCTGGT

>MF443344.1/sheep/China/2014-04-01

ACCAAACAAAGTTGGGTAAGGATAGATCTTATAATAACTATGGACTGGCAAACTTAGGAGTAAAGATCCTACTGTCGGGG

GGAGGAGGAGGAGCAAGATCCTTGATCATGGCGACTCTCCTTAAAAGCTTAGCATTGTTCAAAAGGAACAAAGACAAAGC

GCCGACAGCATCAGGTTCAGGAGGGGCCATCCGGGGGATTAAGAATGTTATCATAGTCCCGATTCCCGGAGACTCGTCCA

TCACTACCCGTTCAAGACTGCTCGACAGGCTTGTCAGATTGGCCGGAGATCCTGACATCAACGGGTCAAAGCTGACCGGG

GTGATGATCAGCATGTTATCCTTGTTCGTAGAGTCACCCGGGCAATTGATACAGCGAATCACAGATGATCCGGATGTTAG

TATCCGCCTTGTTGAGGTAGTTCAAAGTACTAGGTCCCAGTCCGGGTTGACCTTTGCATCACGTGGTGCTGATTTAGACA

ACGAGGCAGACATGTATTTTTCAACTGAGGGGCCCTCGAGTGGAGGTAAGAAAAGGATCAACTGGTTTGAGAACAGAGAA

ATAATAGACATAGAGGTGCAGGATCCAGAAGAGTTCAATATGTTGTTAGCCTCCATACTAGCACAAGTCTGGATCCTTCT

GGCCAAGGCTGTTACGGCACCAGATACGGCAGCTGACTCAGAACTGAGAAGGTGGGTTAAATACACACAACAAAGGAGAG

TGATTGGGGAATTTCGCCTTGACAAGGGGTGGCTGGACGCAGTCCGCAACAGAATTGCAGAAGATCTATCACTCCGGCGG

TTCATGGTATCTCTCATACTTGACATCAAGAGGACCCCAGGCAACAAGCCAAGGATTGCAGAAATGATCTGCGACATTGA

CAACTATATTGTAGAAGCAGGACTCGCCAGTTTCATCCTTACTATCAAATTTGGTATCGAAACCATGTATCCTGCATTAG

GGCTCCACGAGTTTGCCGGAGAATTGTCCACTATTGAGTCCTTGATGAACTTGTATCAACAGCTAGGAGAGGTTGCACCC

TACATGGTAATTCTAGAGAATTCAATTCAGAACAAGTTTAGTGCAGGAGCTTATCCCCTCCTCTGGAGCTATGCGATGGG

TGTCGGAGTCGAGCTGGAGAACTCAATGGGGGGCTTGAACTTTGGCAGGTCATATTTTGACCCTGCCTATTTTCGTCTCG

GACAGGAGATGGTCAGAAGATCTGCAGGAAAGGTCAGCTCTGTGATTGCGGCTGAGCTCGGCATCACAGCAGAGGAAGCT

AAACTAGTCTCGGAAATCGCCTCGCAGGCTGGGGACGAAAGAACCGCCAGAGGGACTGGGCCTCGACAGGCGCAGGTTTC

CTTCCTTCAGCACCAAACAGGAGGGGGAGAGTCGTCCGCACCAGCGACCAGAGAAGGGGTCAAAGCTGCGATCCCAAACG

GAGCTGAAGAAAGGGACAGAAAGCAAACACGCCCAGGAAGGCCCAGAGGAGAGACCTCCGGCCAACTGCTCCTGGACATC

ATGCCAGAGGATGAGATCTTGCGAGAGTCTGGTCAAAACCCTCGTGAGGCTCAAAGATCGGCCGAGGCACTCTTCAGGCT

GCAGGCCATGGCCAAGATTCTGGAGGACCAGGAGGAGGGAGAAGACAACAGTCAGGTCTACAACGACAAGGATCTCCTCG

GCTAAGCAGACGCACCCTCTGTCGAAATCAGTGACAAGACATCCCCTACCAGTATTATAAAAAACTTAGGACCCAGGTCC

AAGCAAGCACACATCGACACTCCAACCAGTCGAGCGGAGACCACCGATGGCAGAAGAACAAGCATACCATGTCAACAAGG

GGCTGGAATGTATCAAGTCTCTCAAAGCCTCACCCCCGGATCTATCCACCATCAGAGATACCATCGAGAGCTGGAGAGAG

GGGCTTAGCCCCTCGGGCCGTGCAACACCGAACCCTGATACGTCCGAGGGAGATCATCAGAATATCAACCAATCATGCTC

ACCAGCAATCGGACCAAACAAAGTCTACTTGTCTCCTGAAGATAATCTCGGACTTAGAGAGATCACTGGCAACGACTGTG

AGGCTGGGCTCGGAGGGGTCCAGAGAGAAGGATCCAACTCTCAAGTACAGCGTTACCATGTTTATAGTCACGGGGGTGAA

GAGATTGAAGGACTCGAGGATGCTGACTCTCTCGTGGTTCAAGCAGATCCTCCAGTTGCTAACGTCTTCAATGGAGGAGA

GGATGGATCTGACGACAGCGATGTGGACTCTGGCCCAGATGATCCCGGCAGAGATACTCTATATGACCGGGGATCTGTTG

CCGGCAATGGTATCGCTAGGTCCACAGATGTCGAAAAACTAGAAGGTGCTGATATTCAAGAAGTTCTTAACTCCCAGAAA

GGCAAGGGAGGAAGATTCCAGGGCGGGAAAACCTTGCGAGTCCCGGAAATACCCGATGTCAAGCACTCCAGACCATCAGC

CCAATCAATTAAAAAGGGCACAGACGGGAACTCAGTCTCATCTGGAACGGTGATAGAGTGTTTATCGATAAGTGGTGCAA

CCCAAACTGTGCCAGAGTCAAGATGGGAGTCATCAGAGCAAAATGCGTCTGTGGGGAGTGTCCTCAAGTCTGCGAGGAGT

GCAAAGACGATCCAGGGGTCGACACAAGAATCTGGTACCATAGCATCACTGACTCAGCCTAAAGAGAATGACTCCGAGTA

TGAGTATGAGGATGACCTATTTACAGAGATTCAGGACATCCGTGCAAGCATTGCCAAGATCCATGATGACAATAAAACTA

TCCTCTCAAAGCTTGATTCTATACTGTTATTGAAAGGGGAAGTCGACACTATCAAGAAACAAATCAGCAAGCAGAATATA

AGTATATCCACCATTGAGGGCCATCTCTCCAGTATAATGATAGCCATCCCTGGCTTTGGGAAGGACATCAAGGACCCAAC

ATCCGAAGTCGAGTTGAACCCAGATCTAAGACCTATAATAAGCCGTGACTCTGGCAGAGCTCTCGCGGAGGTCCTCAAGA

AACCCGCTGTGGATAGGTCTCAGAAAATTGGAACCAAAGCCAACTCCAGCTCAAAGGGTCAGCTTCTTAAGGATCTCCAG

CTAAAACCTGTCGACAAGCAGGCAAGCTCTGCAATCGGGTTTGTCCCGTCCGACCATGAATCATCCAGAAATGTCATCCG

CTCCATAATCAAGTCGAGCAAGCTAAATATTGATCACAAGGACTATCTTCTAGATTTACTAAATGATGTGAAAGGCTCCA

AGGATCTTAAGGAATTTCACAAGATGCTAACAGCAATTCTCGCCAAGCACCCGTAACACATCCTCCAGTCATCATCTCAT

ACTCGACTAAAAACATCCTTTCAATCAGGCTATTACAAAAAACTTAGGAGCAAGGGCAACTGAGCTTCGCAGACAGGATG

ACCGAGATCTACGACTTCGATAAATCAGCATGGGATGTCAAAGGGTCAATTGCCCGCATAGAACCCACCACCTACCACGA

CGGCCGACTGATACCCCAGGTGAGGGTCATCGATCCTGGTCTGGGAGACAGAAAAGATGAGTGCTTTATGTACCTGTTTC

TCCTAGGAGTGATTGAGGACAACGACCCCCTGTCTCCCCCCGTCGGAAGAACCTTTGGCTCTCTACCTCTAGGGGTTGGT

AGGTCAACTGCCAAGCCAGAAGAACTACTAAGGGAGGCCACAGAATTAGATATAGTGGTGAGGCGCACTGCAGGAGTAAA

TGAGAAACTGGTATTTTACAACAACACTCCGCTGTCCTTGTTAACGCCCTGGAAGAAAGTTCTGACAACCGGAAGTGTGT

TTAGCGCTAACCAGGTTTGCAATGCAGTCAACCTAGTCCCGCTTGATACTCCCCAGAGATTCAGGGTTGTGTACATGAGC

ATAACTAGATTATCAGACAATGGTTACTATAGTGTGCCCAGAAGAATGCTGGAGTTCCGCTCGGCCAATGCAGTAGCCTT

CAACATCTTGGTTACACTGAGAATTGAAAATGGCACAAACCCTAGAAGATACATAGTCGGCTCATGGGAGAATTCAGAGG

TCACATTTATGGTACACGTGGGCAACTTTAGGAGAAAGAAGAACGAAGTATACTCTGCTGATTACTGCAAAATGAAGATT

GAAAAGATGGGTTTAGTTTTTGCCCTGGGTGGAATAGGCGGAACAAGTCTCCATATTAGAAGCACCGGGAAAATGAGCAA

AACCCTCCATGCACAGCTGGGGTTCAAGAAAATCTTATGTTACCCCCTAATGGATGTCAATGAGGATCTTAACCGATATC

TCTGGCGGGCAGAGTGCCGAATAGTCAAAATCCAAGCTGTCTTACAGCCATCAGTACCCCAAGAGTTCCGTGTCTACGAT

GATGTCATCATCAACGATGATCAAGGCTTGTTCAAGATCTTGTAGTTCATTTGCAAAGGAACGAACCCACCCCCAGGAAG

GGGCCCCACCCTCCCCCG-CCAAAAAACACACTCCAAGGCTCCCCGGCACAGCGCCAACACAGCTACTCCCGCGATGGAA

CGAGGTCGGCC-CCCGAACAAATCCGGTCCAGGGGGGGGGGCCCCCCCCCTGACCAACCGGGCGCCGACCGAGCAGGCCG

GCGAGAAGCCCCCAAGAGGCCCCGACGAAGACCGAGCACCCCATCCCCCC-AAAAACGTCCCCC-AACACGCCCGAGGCT

AGCCACCGGACACCCCCAGACCCGCCACCGGCGAGACAGAACCCCAAGGGGGGACCCTCCAAACCAAAGGAATCACCGGG

GTAGGCTCCGCCGAACCAGCACAGACCCGACCCAAGCAAGGCGGACCCCCAACACGAAACTGCCCAGCACACCCTACTCA

TCATCAAACAAAACTTAGGGGCCAAGTCCATACGAGAGGCCCTAGTCCGTCAGGCGGAGGGACGCAGAAAGGAAGGAGAC

ACGACCGTCCCAGAAGGAGGACACCCAACCACCGAAACGCCCGCCCCCAGGGAAGGAG--CCA-CCCCCACGCAGCGAAG

GAGCGGGTGGGAGACCTCGGGGAGGGGCCGACCGGGACCCGGGGGCCGGCGGCGCCGGCGGTGAGCAGGCCGGGCGCATA

CCTGACCCCACCGGAGGATCCGAGGAGGGCCCGACCCCGCGTGGACCAAACGGCAGGGAGCCCAACCCCACCAAAAAGAG

AGGCGAGGCCCCCCCTCCCTCCCCCCCCGCACACAACCCGCAGACAGACCGAGGGAGGGCCATCCCCCAGCGAGACAGGC

GGCTCCCTCGCAGCCGAGCCGCAGCTCCCCAGAATAGGACACCCCCGCACACTCCGAACACCCCCTCGCCCCCCGCCACC

GGGACAGGGGCTCCACCACCACAGACCGGAGCCACCGACAGCCAGCCAGAGGTCCCCTCCAGTCCCGCCCGGCGCCCCCC

CGAAGAGGCCCCGGCAAGGCCCCCCAGCCCGACGTCCCCCGCACGCACCGAAAGAAGGACCCGCCCGAGAACGTGCGGCA

CATCCATGCGCAAACATCATGACGCGGGTCGCAATTTTGACATTTCTGTTTCTTTTCCCAAATGTTGTTGCGTGTCAGAT

TCACTGGGGCAATCTATCCAAGATCGGGATTGTAGGAACAGGGAGTGCCAGCTACAAGGTGATGACTAGGCCAAGCCACC

AGACTCTGGTTATAAAGTTAATGCCAAATATAACGGCCATCGACAATTGTACAAAGTCAGAGATTGCAGAGTACAAGAGA

TTGCTGATCACAGTGTTAAAGCCTGTAGAGGATGCTCTGTCGGTGATAACCAAGAATGTAAGACCAATTCAAACTCTAAC

ACCTGGGCGTAGAACCCGCCGTTTTGCTGGAGCTGTTCTGGCCGGGGTAGCACTTGGAGTTGCGACAGCCGCTCAGATAA

CTGCAGGAGTCGCCCTTCATCAATCATTGATGAACTCCCAAGCAATTGAGAGTTTAAAAACCAGTCTTGAGATGTCGAAT

CAGGCAATAGAAGAAATCAGACTTGCAAATAAGGAGACCATACTGGCAGTACAGGGCGTCCAGGATTATATCAACAATGA

GCTCGTCCCTTCTGTTCATAGAATGTCATGCGAGCTGGTAGGTCACAAGCTCGGCCTCAAGCTCCTTAGGTACTACACCG

AGATCCTGTCCATATTCGGGCCCAGTCTTCGAGACCCGATATCTGCCGAAATATCAATCCAGGCACTTAGTTATGCATTA

GGCGGAGACATTAATAAAATCCTGGAAAAGCTTGGGTATAGCGGTGGGGATTTCCTTGCCATCCTAGAAAGCAAGGGAAT

AAAGGCCCGGGTTACATATGTGGACACAAGAGATTACTTTATAATCCTTAGCATCGCCTACCCAACCTTATCTGAGATCA

AGGGAGTGATAGTTCACAAGATAGAAGCTATAACATACAACATTGGGGCACAGGAGTGGTATACTACTATCCCTAAATAT

GTAGCCACTCAGGGGTATCTGATATCGAACTTTGATGAGACGTCATGCGTATTCACTCCAGAGGGGACAGTTTGCAGCCA

GAATGCGTTGTACCCAATGAGCCCATTGCTTCAGGAATGTTTCAGGGGGTCAACAAAATCGTGCGCCAGAACCTTAGTTT

CAGGGACCATAAGTAATAGATTTATCCTATCAAAAGGGAACCTGATTGCAAATTGTGCGTCAGTTTTGTGCAAATGTTAC

ACAACAGAGACAGTTATCAGCCAAGATCCTGACAAACTACTAACTGTTGTAGCATCCGACAAGTGTCCTGTAGTTGAGGT

GGATGGAGTGACAATACAGGTCGGCAGTCGAGAGTATCCGGATTCTGTATACTTACACAAAATAGACTTAGGTCCAGCCA

TCTCCCTAGAAAAACTGGATGTAGGCACCAATTTAGGCAATGCAGTCACAAGACTGGAGAATGCAAAGGAGCTCCTAGAT

GCATCAGACCAAATACTGAAGACTGTTAAAGGGGTACCTTTCAGTGGGAATATGTACATAGCACTGGCAGCTTGCATAGG

AGTATCCCTGGGCCTTGTCACATTAATATGCTGCTGTAAGGGGAGGTGTAAGAACAAGGAAATCCCTATCTCCAAAATCA

ACCCAGGGCTCAAACCCGACCTGACCGGGACCTCAAAGTCGTACGTAAGATCACTGTAGTCAGAATCACCTGGATCATCT

GGCATCACACACAGACATGCACGACACAGGCAGTCCGAGGACGCAAGAAACCCAGCCTCCGGTCACCCACCCGACCCCAC

TCCACGCTCCACCACACATTAGTCATCAAACAAAACTTAGGACGAAAGGTCAATCACCATGTCCGCACAAAGGGAGAGGA

TCAATGCCTTCTACAAAGACAATCCTCACAATAAGAACCATAGGGTGATCCTGGATAGAGAACGCTTGGTCATTGAAAGA

CCCTACATCTTGCTTGGAGTCCTGCTGGTAATGTTCCTGAGTCTAATCGGACTGCTGGCCATTGCAGGGATCAGGCTTCA

CCGGGCCACCGTTGGAACTTCAGAGATCCAGAGTCGGCTGAATACCAATATTAAGTTGACCGAATCTATTGATCACCAGA

CTAAGGATGTCTTAACTCCCCTTTTTAAAATCATTGGCGATGAAGTCGGCATCAGAATTCCACAGAAATTCAGTGATCTT

GTCAAGTTCATCTCCGATAAGATTAAATTCCTCAACCCTGATAGAGAGTATGATTTCAGAGATCTCCGGTGGTGTATGAA

TCCCCCCGAGAGAGTCAAAATTAATTTTGATCAGTTTTGTGAGTACAAGGCTGCGGTTAAGTCAATTGAACATATATTTG

AGTCACCACTCAACAAGTCAAAAAAGCTGCAATCTTTGACTCTCGGGCCCGGAACAGGCTGTCTAGGCAGGACAGTAACA

AGAGCCCATTTCTCAGAACTTACAATGACCTTAATGGACCTGGATCTAGAGATGAAGCACAACGTGTCCTCAGTGTTTAC

CGTAGTTGAAGAGGGATTATTCGGAAGAACATATACCGTCTGGAGATCCGATGCCAGGGATCCGAGCACCGATCTAGGTA

TCGGCCATTTTTTAAGAGTCTTCGAGATTGGACTGGTAAGAGATCTTGGGCTGGGTCCCCCTGTTTTTCATATGACCAAC

TATCTCACGGTGAACATGAGTGATGACTATCGGAGATGTCTTTTAGCGGTAGGGGAGTTGAAGTTGACAGCCCTATGCAC

CTCATCTGAGACTGTGACACTGAGTGAGAGAGGAGTTCCAAGGAGGGAACCTCTTGTGGTTGTGATACTTAATCTAGCTG

GACCCACTCTAGGGGGAGAGCTATACAGTGTCTTGCCTACCTCTGATCTCATGGTGGAGAAACTCTATTTGTCTTCACAT

AGAGGGATCATCAAAGATGATGAGGCCAATTGGGTAGTGCCGTCTACCGATGTTCGTGATCTTCAGAACAAAGGTGAATG

TCTGGTGGAAGCATGCAAGACTCGACCTCCTTCATTTTGCAATGGCACAGGATCAGGCCCGTGGTCAGAAGGGAGAATCC

CTGCCTACGGGGTGATCAGGGTCAGTCTTGACTTAGCGAGTGACCCGGATGTAGTTATCACTTCAGTGTTTGGCCCACTG

ATACCTCACCTATCCGGCATGGATCTTTACAACAACCCGTTTTCAAAAGCTGTATGGTTGGCTGTACCACCTTATGAGCA

GTCATTTCTAGGAATGATAAATACAATTGGATTCCCTAACAGAGCAGAGGTTATGCCGCACATTTTGACCACAGAGATCA

GAGGCCCTCGGGGTCGTTGCCATGTTCCCATAGAATTGTCCCGCCGGGTTGATGACGATATCAAGATCGGGTCCAACATG

GTCATATTGCCGACGATGGACCTGAGGTATATTACAGCCACTTATGATGTTTCCAGGAGCGAGCATGCAATCGTGTACTA

TATCTATGACACGAGTCGCTCATCATCTTACTTCTACCCAGTTCGACTGAATTTCAAAGGCAATCCTCTCTCTTTGAGGA

TAGAGTGTTTCCCCTGGCGTCATAAGGTGTGGTGCTACCATGATTGTCTTATATACAACACCATAACAGGTGAAGAGGTC

CATACGAGAGGGCTGACCGGCATAGAGGTAACATGCAATCCAGTCTGAGCAGAGCTACGACCGTCACTCAGGCAGTCCCT

TGAGTCGCCACCGAGTCCAAGCAGCACAGCCTGGGACAATCAACGGCACAGCCCAGCCAACAATGTTATAAAAAACTTAG

GAGCCAGGGTTGTAGGGGCCATGGACTCTCTATCAGTCAATCAGGTCTTGTACCCTGAGGTTCATCTAGACAGCCCTATT

GTCACAAACAAACTAGTTGCCATCCTTGAGTACTCGGGTATCGACCATAACTATGTTCTTGAAGACCAGACTCTTGTCAA

GAATATTAGGTATAGGCTGGGGTGCGGTTTTTCAAATCAAATGATCATCAATAACAGGGGGGTGGGTGAAACAGTCAATT

CTAAACTTAAAAGTTACCCCCATAATCGTCATATCATATACCCGGATTGCAATAAGGAGTTGTTTTGTATCAAGGATAGC

TGCATATCTAAGAAGCTCTCGGAGCTATTCAAGAAGGGTAATTCCTTGTACTCTAAGATAAGTCACCAGGTACTGGATTG

TCTTAAGAGAGTCAATGGGAAATTAGGTCTGGGCACAGATCTTACCCATGGTCTGAAAGAGGGTATCCTTGACCTAGGGT

TGCACATGCATAGCTCTCAATGGTTCGAGACTTTCCTGTTCTGGTTCACTATCAAGACAGAGATGAGATCAATGATCAAA

GAACAGTCCCATATATGCCACAAGAGGAGGTATAACCCGACTTTTGTGTCGGGGGATGCATTCGAGGTGCTCGTATCGCG

AGACCTCGTTGTGATAATTGATAAGAATACCCAGTATGTCTTCTACTTAACATTTGAGCTGGTCCTTATGTATTGTGATG

TCATAGAGGGTAGACTTATGACGGAGACAGCCATGGCCATAGACCAGAGATATTCAGAACTTCTAAGCCGGGTCAGATAC

TTGTGGGATCTTATTGATGGGTTCTTCCCAACACTAGGCAACACCACATACCAAATTGTTGCTCTGCTTGAACCATTATC

GTTGGCTTATCTTCAACTTCAGGATGTCACTCTGGAGTTAAGAGGTGCTTTTTTAGACCATTGCTTCAAAGAACTCTATG

AGATACTGGAACATTGCGGCATTGACACAGCAGGCACTTACAATTCCATCACTGAAGGATTGGATTACGTATTCATCACC

CACGATATACATCTAACTGGGGAGATTTTTTCATTTTTCCGGAGTTTCGGACACCCCCGTCTAGAAGCAGTCACCGCTGC

GGAAAATGTCAGAAAACATATGAACCAACCGAAGGTAATCAGTTATGAGACTATGATGAAAGGCCATGCGGTATTTTGCG

GGATAATCATAAATGGTTTTCGGGACCGACATGGCGGCAGCTGGCCCCCTGTTGCACTGCCAGAACATGCCTCTGCTGCG

ATCCGGAATGCGCAGGCATCAGGTGAGGGACTGACCCATGACCTGTGTATAGACAACTGGAAATCCTTTGTTGGATTCAA

ATTTGGCTGCTTTATGCCACTCAGCCTAGATAGTGATTTGACCATGTATCTCAAAGACAAGGCATTGGCTGCACTGAAGA

ATGAGTGGGATTCAGTTTACCCGAAAGAATACCTCCGTTATAACCCACCTAGAGGGACAGAGTCAAGGCGATTGGTAGAG

GTATTCCTGAATGACTCCAGCTTTGATCCTTATAACATGATAATGTACGTGGTGAATGGCTCCTACCTTAAAGACCCTGA

GTTTAACCTCTCATACAGCCTAAAAGAGAAGGAGATAAAGGAGACAGGGCGGTTGTTTGCCAAAATGACCTATAAGATGC

GGGCCTGTCAGGTAATCGCTGAAAATCTGATATCGAATGGTGTTGGGAAGTATTTCCGAGACAATGGGATGGCAAAGGAC

GAGCATGACCTAACAAAAGCCCTTCACACCCTGGCAGTCTCAGGTGTTCCCAAAAACAACAAAGATAACCACCGAGGTGG

GCCTCCCAGAAGGACAACAAACCGAGGGGTGAGATCAAGCCAAGGCACCAAAACACAAGATAGAGACAAGGTTCAAGGGG

GACCTATGTACAACTATTTGCGATGCCAACCGATCAGCCCTGATCAGGGTGAGTCATACGAGACTGTTAGTGCATTCATC

ACCGCTGACCTTAAGAAGTATTGCCTGAATTGGAGATACGAGACAATCAGCATATTTGCACAGAGGCTGAATGAAATATA

TGGACTACCATCCTTCTTTCAATGGTTACACAGGATATTGGAAAAATCCGTACTCTACGTCAGTGACCCACATTGTCCTC

CCGATCTAGATAATCATATCCCTCTGGACAGTGTCCCCAATGCCCAAATATTCATCAAGTACCCAATGGGCGGAATAGAA

GGTTATTGCCAAAAACTATGGACAATCAGCACTATACCATACTTGTATCTGGCAGCCTATGAGAGCGGAGTAAGAATCGC

CTCACTGGTGCAGGGTGACAATCAGACAATCGCAGTGACAAAAAGAGTTCCAAGTTCTTGGCCTTATTCACTAAAAAAGA

GGGAGGCCTCCAAGGCCGCTCAAAATTACTTCGTAGTCTTAAGGCAAAGATTGCACGATGTAGGTCATCACTTAAAGGCT

AATGAGACCATAGTATCTTCTCACTTTTTTGTATATTCCAAAGGGATTTATTATGACGGCCTGCTAGTCTCACAATCACT

AAAGAGCATCGCCAGATGTGTCTTCTGGTCCGAGACTATAGTGGATGAAACCAGAGCGGCGTGCAGTAATATTGCAACAA

CTGTCGCTAAGAGTATAGAGAGGGGTTATGATAGGTACCTTGCATATTCTTTGAATATCCTCAAGATTTTCCAACAGATC

CTTATATCCCTTAACTTCACTATTAACACAACAATGACTCAGGATGTCGTGGCACCGATCATAGAGAACGGTGATTTGCT

GATAAGGATGGCACTCTTGCCAGCACCCATCGGGGGTCTCAATTATCTTAACATGAGCAGGTTATTTGTGAGAAATATCG

GTGACCCGGTCACCTCCTCTATAGCCGACCTGAAGAGGATGATAGATGCCGGGCTAATGCCAGAAGAAACATTGCATCAA

GTGATGACCCAGACCCCGGGAGAATCATCCTACCTAGATTGGGCAAGTGACCCTTATTCTGCCAACCTACCCTGCGTACA

GAGTATCACTCGCCTTCTCAAAAACATCACTGCACGATATATTTTAATCAGCAGCCCAAACCCGATGCTGAAAGGGTTAT

TTCATGAGGGGAGTAGAGATGAAGACGAGGAGCTTGCAAGTTTCCTAATGGATCGGCATATAATTGTTCCAAGAGCTGCA

CATGAAATCTTAGACCATAGTATAACCGGAGCAAGAGAAGCTATAGCCGGGATGTTGGACACCACCAAGGGTCTGATTAG

AACAAGTATGAAACGGGGCGGCCTCACCCCTCGAGTCTTAGCCCGCCTTTCCAATTATGATTATGAACAATTCCGATCCG

GGATAACACTATTAACAAAGAAAGGGCAGTGTTATCTCATTGACAAGGACTCGTGCTCGGTGCAGCTCGCTATAGCTCTG

AGGGGCCACATGTGGGCTAAGTTGGCTCGCGGGAGACCAATCTATGGGTTGGAGGTGCCTGATGTATTAGAATCGATGAA

CGGCTACCTTATCAAACGCCACGAGTCGTGTGCTATCTGTGAAACGGGCTCAAGTCACTACGGATGGTTTTTCGTCCCTG

CAGGGTGCCAGCTTGACGATGTATCTAGAGAGACTTCGGCTCTTCGTGTACCTTATGTCGGATCAACAACTGAGGAAAGG

ACAGACATGAAACTTGCATTTGTTAGGTCTCCAAGTAGATCCCTCAAATCAGCAGTTAGAATTGCAACAGTTTACTCATG

GGCTTACGGGGATGATGAGAAATCATGGAGTGAAGCCTGGATGCTAGCTAGGCAGAGAGCCAATATCACCTTAGATGAAT

TGAGAATGATCACTCCAGTCTCCACATCCACCAACTTAGCCCACCGGTTGAGGGATCGGAGTACCCAGGTGAAATACTCG

GGGACATCCCTTGTGAGGGTTGCAAGATACACAACCATCTCCAATGATAACTTGTCGTTTGTGATATCTGAGAAAAAAGT

GGATACCAACTTCATTTATCAGCAAGGGATGTTGCTTGGTCTTGGGATCCTTGAGAACCTCTTTAGGCTAGAGGCCACCA

CCGGAGTATCCAACACAGTGCTACACCTGCACGTGGAAACAGAATGTTGTGTTGTACCTATGGTGGATCACCCAAGGATA

CCGAGTCTCCGTAATATAAAAGTTACGGATGAGCTATGCACAAACCCTTTGATCTACGACAGGTCCCCCATCATAGAGCA

CGATGCAACCCGGCTATATTCACAGAGCCACAGGAGACATTTGGTGGAGTTTGTTACCTGGTCAACAAGCCAGCTTTACC

ATATACTGGCTAAGTCTACAGCAATGTCCATGATTGAATTGATCACGAGATTCGAGAAAGATCACATGAATGAAATAGCC

GCCCTGATTGGCGATGACGACATCAACAGCTTCATCACAGAATTTTTGCTAGTGGAGCCTAGATTATTTATAGTTTACCT

TGGTCAGTGTGCTGCCATCAATTGGGCCTTTGATATACATTATCATCGGCCCTCGGGCAAGTACCAGATGGGGGAACTTC

TCTACTCTCTGCTCTCTCGGATGAGCAAAGGAGTATATAAGATCTTTACTAATGCTTTGAGCCACCCCAAAGTTTACAAG

AAATTTTGGCGAAGTGGCATCATTGAGCCGATTCATGGCCCATCCTTGGATACACAGAATTTACATGTCACTGTCTGTGA

CATGATATATGGGTCCTATGTCACCTATTTGGATCTTTTGCTGAATGATGAGCTAGATGATTACCCGTATTTGCTCTGCG

AGAGTGATGAGGACGTGGTCACAGACAGGTTCGACAACATTCAAGCCAAACATCTCTGTGTATTGGCCGATGTATATTGC

AGCTCCAAGAGGTGTCCTTCAATAATCGGGATGTCTCCTATAGAAAAATGTACCATTCTCACACATTACATCAAGGGAGA

ATCAGTACAATCCCCGTCTGGGACCTCATGGAACACTGATCCCCTTGTAGTAGATCATTACTCATGCTCTCTGACCTACC

TTCGCCGCGGTTCCATCAAACAAATCAGGTTGAGGGTGGATCCTGGGTTTGTATTCGAGGCGTTGACAGACGTCGACTTT

AAACAACCTCGCAAAGCTAAGTTAGATATATCGGTCGTTGGGTTGACTGATTTCTCTCCCCCTTACGACAACGTCGGTGA

TTTTTTAGGGACTATCAACACATTGAGGCACGATCTGCCTGTCACCGGAACCGGGGTCTCGAACTATGAAGTCCACGCTT

ATCGTAGAATTGGCCTGAATTCATCAGCATGTTATAAAGCCGTAGAGATCTCCACGTTAATCAAGCCATCCTTAGAAGTC

GGAGAGCATGGATTGTTCTTAGGAGAAGGTTCCGGTTCAATGCTGGCTGCGTACAAAGAAGTTCTTAAATTAGCAAATTG

TTACTACAACAGCGGAGTAACAGCGGAGGGCAGAGCCGGACAGAGGGAAATATCTCCCTATCCTTCAGAGATGAGCCTAG

TAGAGAATCAAATGGGGATAGAGCGGAGTGTTAAAGTGCTGTTCAACGGGAAACCTGAAGTAACTTGGGTGGGGACCACA

GATTGCTACAAGTACATAATCAGTAACATCCAAACCTCTAGTCTGGGTTTCATACATTCAGATATTGAGACGCTCCCAAC

CAAAGATGCTGTTGAGAAATTGGAGGAATTTGCCTCTATCCTATCCTTATCCTTGATTTTGGGGAAAATCGGCTCTATTA

CAGTTGTCAAAATTATGCCCATTAGCGGGGATTTTACCCAAGGCTTCATAGCCTATGCCATCCAATATTTTAGGGAGAGC

CTGCTTGCCTATCCGAGATATAGTAACTTCATCTCGACTGAGTGTTACCTTATTATGATAGGATTAAAGGCCAATCGATT

GATAAACCCAGAAGCCATTAAGCAAAGCATAATCAGAGCGGGGGTCAGGACTGCACCAGGACTTGTGAGCCATGTATTAT

CAGAGAAACAAAAAGGTTGTATTCAATCTTTCTTGGGTGATCCTTATATCCAAGGAGACTTCAATAAGCACCTTAAATCT

CTAACCTCTATTGAGAAAATCCTGGTAAATTGTGGTCTCTCGATCAATGGCATAAAAATCTGTAGAGATCTAATCCACCA

TGATATCGCCTCCGGTCCAGACGGCCTTATGAGCTCTACAATTATTTTATATAGGGAATTGGCTCATTTCAAAGATAATA

TAAGGAGTCAGCACGGAATGTTCCACCCCTACCCGGTACTGGCCAACAGCAGGCAACGTGAATTAATCCTTCGAATAGCC

AAGAAATTCTGGGGGTATGTCTTGCTATATTCTGATGACCCGGCACTAATCAGACAAACAATCAAGAACTTGAAACGGAA

TCATCTAACCTTTGACTTACATAGTAACCCATTTATTAAAGGGCTATCCAAGGCTGAGAAACTGCTGGTGCGGACGAGTT

CACTAAGAAGAGAATGGTTGTTCACTCTCGAAACGAAAGAAGTAAAAGAGTGGTTCAAATTGGTAGGTTACAGTGCACTC

ATCAGAGGCTAGTCAGCTATGCATCTGCCCCCTTCTCCTCCGCCATGAGACCCCACTGGCGATCCAGAAGATTAAAGAAA

ACTACATATTGGATAAGTATCTATTCCCAGCTTTGTCTGGT

>MF443348.1/goat/China/2014-04-03

ACCAAACAAAGTTGGGTAAGGATAGATCTTATAATAACTATGGACTGGCAAACTTAGGAGTAAAGATCCTACTGTCGGGG

GGAGGAGGAGGAGCAAGATCTTTGATTATGGCGACTCTCCTTAAAAGCTTAGCATTGTTCAAAAGGAACAAAGACAAAGC

GCCGACAGCATCAGGTTCAGGAGGGGCCATCCGGGGGATTAAGAATGTTATCATAGTCCCGATTCCCGGAGACTCGTCCA

TCACTACCCGTTCAAGACTGCTCGACAGGCTTGTCAGATTGGCCGGAGATCCTGACATCAACGGGTCAAAGCTGACCGGG

GTGATGATCAGCATGTTATCCTTGTTCGTAGAGTCACCCGGGCAATTGATACAGCGAATCACAGATGATCCGGATGTTAG

TATCCGCCTTGTTGAGGTAGTTCAAAGTACTAGGTCCCAGTCCGGGTTGACCTTTGCATCACGTGGTGCTGATTTAGACA

ACGAGGCAGACATGTATTTTTCAACTGAGGGGCCCTCGAGTGGAGGTAAGAAAAGGATCAACTGGTTTGAGAACAGAGAA

ATAATAGACATAGAGGTGCAGGATCCAGAAGAGTTCAATATGTTGTTAGCCTCCATACTAGCACAAGTCTGGATCCTTCT

GGCCAAGGCTGTTACGGCACCAGATACGGCAGCTGACTCAGAACTGAGAAGGTGGGTTAAATACACACAACAAAGGAGAG

TGATTGGGGAATTTCGCCTTGACAAGGGGTGGCTGGACGCAGTCCGCAACAGAATTGCAGAAGATCTATCACTCCGGCGG

TTCATGGTATCTCTCATACTTGACATCAAGAGGACCCCAGGCAACAAGCCAAGGATTGCAGAAATGATCTGCGACATTGA

CAACTATATTGTCGAAGCAGGACTCGCCAGTTTCATCCTTACTATCAAATTTGGTATCGAAACCATGTATCCTGCATTAG

GGCTCCACGAGTTTGCCGGAGAATTGTCCACTATTGAGTCCTTGATGAACTTGTATCAACAGCTAGGAGAGGTTGCACCC

TACATGGTAATTCTAGAGAATTCAATTCAGAACAAGTTTAGTGCAGGAGCTTATCCCCTCCTCTGGAGCTATGCGATGGG

TGTCGGAGTCGAGCTGGAGAACTCAATGGGGGGCTTGAACTTTGGCAGGTCATATTTTGACCCTGCCTATTTTCGTCTCG

GACAGGAGATGGTCAGAAGATCTGCAGGAAAGGTCAGCTCTGTGATTGCGGCTGAGCTCGGCATCACAGCAGAGGAAGCT

AAACTAGTCTCGGAAATCGCCTCGCAGGCTGGGGACGAAAGAACCGCCAGAGGGACTGGGCCTCGACAGGCGCAGGTTTC

CTTCCTTCAGCACCAAACAGGAGGGGGAGAGTCGTCCGCACCAGCGACCAGAGAAGGGGTCAAAGCTGCGATCCCAAACG

GAGCTGAAGAAAGGGACAGAAAGCAAACACGCCCAGGAAGGCCCAGAGGAGAGACCTCCGGCCAACTGCTCCTGGACATC

ATGCCAGAGGATGAGATCTTGCGAGAGTCTGGTCAAAACCCTCGTGAGGCTCAAAGATCGGCCGAGGCACTCTTCAGGCT

GCAGGCCATGGCCAAGATTCTGGAGGACCAGGAGGAGGGAGAAGACAACAGTCAGGTCTACAACGACAAGGATCTCCTCG

GCTAAGCAGACGCACCCTCTGTCGAAATCAGTGACAAGACATCTCCTACCAGTATTATAAAAAACTTAGGACCCAGGTCC

AAGCAAGCACACATCGACACTCCAACCAGTCGAGCGGAGACCACCGATGGCAGAAGAACAAGCATACCATGTCAACAAGG

GGCTGGAATGTATCAAGTCTCTCAAAGCCTCACCCCCGGATCTATCCACCATCAGAGATACCATCGAGAGCTGGAGAGAG

GGGCTTAGCCCCTCGGGCCGTGCAACACCGAACCCTGATACGTCCGAGGGAGATCATCAGAATATCAACCAATCATGCTC

ACCAGCAATCGGACCAAACAAAGTCTACTTGTCTCCTGAAGATAATCTCGGATTTAGAGAGATCACTGGCAACGACTGTG

AGGCTGGGCTCGGAGGGGTCCAGAGAGAAGGATCCAACTCTCAAGTACAGCGTTACCATGTTTATAGTCACGGGGGTGAA

GAGATTGAAGGACTCGAGGATGCTGACTCTCTCGTGGTTCAAGCAGATCCTCCAGTTGCTAACGTCTTCAATGGAGGAGA

GGATGGATCTGACGACAGCGATGTGGACTCTGGCCCAGATGATCCCGGCAGAGATACTCTATATGACCGGGGATCTGTTG

CCGGCAATGGTATCGCTAGGTCCACAGATGTCGAAAAACTAGAAGGTGCTGATATTCAAGAAGTTCTTAACTCCCAGAAA

GGCAAAGGAGGAAGATTCCAGGGCGGGAAAACCTTGCGAGTCCCGGAAATACCCGATGTCAAGCACTCCAGACCATCAGC

CCAATCAATTAAAAAGGGCACAGACGGGAACTCAGTCTCATCTGGAACGGTGATAGAGTGTTTATCGATAAGTGGTGCAA

CCCAAACTGTGCCAGAGTCAAGATGGGAGTCATCAGAGCAAAATGCGTCTGTGGGGAGTGTCCTCAAGTCTGCGAGGAGT

GCAAAGACGATCCAGGGGTCGACACAAGAATCTGGTACCATAGCATCACTGACTCAGCCTAAAGAGAATGACTCCGAGTA

TGAGTATGAGGATGACCTATTTACAGAGATTCAGGACATCCGTGCAAGCATTGCCAAGATCCATGATGACAATAAAACTA

TCCTCTCAAAGCTTGATTCTATACTGTTATTGAAAGGGGAAGTCGACACTATCAAGAAACAAATCAGCAAGCAGAATATA

AGTATATCCACCATTGAGGGCCATCTCTCCAGTATAATGATAGCCATCCCTGGCTTTGGGAAGGACATCAAGGACCCAAC

ATCCGAAGTCGAGTTGAACCCAGATCTAAGACCTATAATAAGCCGTGACTCTGGCAGAGCTCTCGCGGAGGTCCTCAAGA

AACCCGCTGTGGATAGGTCTCAGAAAATTGGAACCAAAGCCAACTCCAGCTCAAAGGGTCAGCTTCTTAAGGATCTCCAG

CTAAAACCTGTCGACAAGCAGGCAAGCTCTGCAATCGGGTTTGTCCCGTCCGACCATGAATCATCCAGAAATGTCATCCG

CTCCATAATCAAGTCGAGCAAGCTAAATATTGATCACAAGGACTATCTTCTAGATTTACTAAATGATGTGAAAGGCTCCA

AGGATCTTAAGGAATTTCACAAGATGCTAACAGCAATTCTCGCCAAGCACCCGTAACACATCCTCCAGTCATCATCTCAT

ACTCGACTAAAAACATCCTTTCAATCAGGCTATTACAAAAAACTTAGGAGCAAGGGCAACTGAGCTTCGCAGACAGGATG

ACCGAGATCTACGACTTCGATAAATCAGCATGGGATGTCAAAGGGTCAATTGCCCGCATAGAACCCACCACCTACCACGA

CGGCCGACTGATACCCCAGGTGAGGGTCATCGATCCTGGTCTGGGAGACAGAAAAGATGAGTGCTTTATGTACCTGTTTC

TCCTAGGAGTGATTGAGGATAACGACCCCCTGTCTCCCCCCGTCGGAAGAACCTTTGGCTCTCTACCTCTAGGGGTTGGT

AGGTCAACTGCCAAGCCAGAAGAACTACTAAGGGAGGCCACAGAATTAGATATAGTGGTGAGGCGCACTGCAGGAGTAAA

TGAGAAACTGGTATTTTACAACAACACTCCGCTGTCCTTGTTAACGCCCTGGAAGAAAGTTCTGACAACCGGAAGTGTGT

TTAGCGCTAACCAGGTTTGCAATGCAGTCAACCTAGTCCCGCTTGATACTCCCCAGAGATTCAGGGTTGTGTACATGAGC

ATAACTAGATTATCAGACAATGGTTACTATAGTGTGCCCAGAAGAATGCTGGAGTTCCGCTCGGCCAATGCAGTAGCCTT

CAACATCTTGGTTACACTAAGAATTGAAAATGGCACAAACCCTAGAAGATACATAGTCGGCTCATGGGAGAATTCAGAGG

TCACATTTATGGTACACGTGGGCAACTTTAGGAGAAAGAAGAACGAAGTATACTCTGCTGATTACTGCAAAATGAAGATT

GAAAAGATGGGTTTAGTTTTTGCCCTGGGTGGAATAGGCGGAACAAGTCTCCATATTAGAAGCACCGGGAAAATGAGCAA

AACCCTCCATGCACAGCTGGGGTTCAAGAAAATCTTATGTTACCCCCTAATGGATGTCAATGAGGATCTTAACCGATATC

TCTGGCGGGCAGAGTGCCGAATAGTCAAAATCCAAGCTGTCTTACAGCCATCAGTACCCCAAGAGTTCCGTGTCTACGAT

GATGTCATCATCAACGATGATCAAGGCTTGTTCAAGATCTTGTAGTTCATTTGCAAAGGAACGAACCCACCCCCAGGAAG

GGGCCCCACCCTCCCCCG-CCAAAAAACACACTCCAAGGCTCCCCGGCACAGCGCCAACACAGCTACTCCCGCGATGGAA

CGAGGTCGGCC-CCCGAACAAATCCGGTCCAGGGGGGGGGGCCCCCCCCCTGACCAACCGGGCGCCGACCGAGCAGGCCG

GCGAGAAGCCCCCAAGAGGCCCCGACGAAGACCGAGCACCCCATCCCCCC-AAAAACGTCCCCC-AACACGCCCGAGGCT

AGCCACCGGACACCCCCAGACCCGCCACCGGCGAGACAGAACCCCAAGGGGGGACCCTCCAAACCAAAGGAATCACCGGG

GTAGGCTCCGCCGAACCAGCACAGACCCGACCCAAGCAAGGCGGACCCCCAACACGAAACTGCCCAGCACACCCTACTCA

TCATCAAACAAAACTTAGGGGCCAAGTCCATACGAGAGGCCCTAGTCCGTCAGGCGGAGGGACGCAGAAAGGAAGGAGAC

ACGACCGTCCCAGAAGGAGGACCCCCAACCACCGAAACGCCCGCCCCCAGGGAAGGAG--CCA-CCCCCACGCAGCGAAG

GAGCGGGTGGGAGACCTCGGGGAGGGGCCGACCGGGACCCGGGGGCCGGCGGCGCCGGCGGTGAGCAGGCCGGGCGCATA

CCTGACCCCACCGGAGGATCCGAGGAGGGCCCGACCCCGCGTGGACCAAACGGCAGGGAGCCCAACCCCCCCAAAAAGAG

AGGCGAGGCCCCCCCTCCCTCCCCCCCCGCACACAACCCGCAGACAGACCGAGGGAGGGCCATCCCCCAGCGAGACAGGC

GGCTCCCTCGCAGCCGAGCCGCAGCTCCCCAGAATAGGACACCCCCGCACACTCCGAACACCCCCTCGCCCCCCGCCACC

GGGACAGGGGCTCCACCACCACAGACCGGAGCCACCGACAGCCAGCCAGAGGTCCCCTCCAGTCCCGCCCGGCGCCCCCC

CGAAGAGGCCCCGGCAAGGCCCCCCAGCCCGACGTCCCCCGCACGCACCGAAAGAAGGACCCGCCCGAGAACGTGCGGCA

CATCCATGCGCAAACATCATGACGCGGGTCGCAATTTTGACATTTCTGTTTCTTTTCCCAAATGTTGTTGCGTGTCAGAT

TCACTGGGGCAATCTATCCAAGATCGGGATTGTAGGAACAGGGAGTGCCAGCTACAAGGTGATGACTAGGCCAAGCCACC

AGACTCTGGTTATAAAGTTAATGCCAAATATAACGGCCATCGACAATTGTACAAAGTCAGAGATTGCAGAGTACAAGAGA

TTGCTGATCACAGTGTTAAAGCCTGTAGAGGATGCTCTGTCGGTGATAACCAAGAATGTAAGACCAATTCAAACTCTAAC

ACCTGGGCGTAGAACCCGCCGTTTTGCTGGAGCTGTTCTGGCCGGGGTAGCACTTGGAGTTGCGACAGCCGCTCAGATAA

CTGCAGGAGTCGCCCTTCATCAATCATTGATGAACTCCCAAGCAATTGAGAGTTTAAAAACCAGTCTTGAGATGTCGAAT

CAGGCAATAGAAGAAATCAGACTTGCAAATAAGGAGACCATACTGGCAGTACAGGGCGTCCAGGATTATATCAACAATGA

GCTCGTCCCTTCTGTTCATAGAATGTCATGCGAGCTGGTAGGTCACAAGCTCGGCCTCAAGCTCCTTAGGTACTACACCG

AGATCCTGTCCATATTCGGGCCCAGTCTTCGAGACCCGATATCTGCCGAAATATCAATCCAGGCACTTAGTTATGCATTA

GGCGGAGACATTAATAAAATCCTGGACAAGCTTGGGTATAGCGGTGGGGATTTCCTTGCCATCCTAGAAAGCAAGGGAAT

AAAGGCCCGGGTTACATATGTGGACACAAGAGATTACTTTATAATCCTTAGCATCGCCTACCCAACCTTATCTGAGATCA

AGGGAGTGATAGTTCACAAGATAGAAGCTATAACATACAACATTGGGGCACAGGAGTGGTATACTACTATCCCTAAATAT

GTAGCCACTCAGGGGTATCTGATATCGAACTTTGATGAGACGTCATGCGTATTCACTCCAGAGGGGACAGTTTGCAGCCA

GAATGCGTTGTACCCAATGAGCCCATTGCTTCAGGAATGTTTCAGGGGGTCAACAAAATCGTGCGCCAGAACCTTAGTTT

CAGGGACCATAAGTAATAGATTTATCCTATCAAAAGGGAACCTGATTGCAAATTGTGCGTCAGTTTTGTGCAAATGTTAC

ACAACAGAGACAGTTATCAGCCAAGATCCTGACAAACTACTAACTGTTGTAGCATCCGACAAGTGTCCTGTAGTTGAGGT

GGATGGAGTGACAATACAGGTCGGCAGTCGAGAGTATCCGGATTCTGTATACTTACACAAAATAGACTTAGGTCCAGCCA

TCTCCCTAGAAAAACTGGATGTAGGCACCAATTTAGGCAATGCAGTCACAAGACTGGAGAATGCAAAGGAGCTCCTAGAT

GCATCAGACCAAATACTGAAGACTGTTAAAGGGGTACCTTTCAGTGGGAATATGTACATAGCACTGGCAGCTTGCATAGG

AGTATCCCTGGGCCTTGTCACATTAATATGCTGCTGTAAGGGGAGGTGTAAGAACAAGGAAATCCCTATCTCCAAAATCA

ACCCAAGGCTCAAACCCGACCTGACCGGGACCTCAAAGTCGTACGTAAGATCACTGTAGTCAGAATCACCTGGATCATCT

GGCATCGCACACATACATGCACGACACAGGCAGTCCGAGGACGCAAGAAACCCAGCCTCCGGTCACCCACCCGACCCCAC

TCCACGCTCCACCACACATTAGTCATCAAACAAAACTTAGGACGAAAGGTCAATCACCATGTCCGCACAAAGGGAGAGGA

TCAATGCCTTCTACAAAGACAATCCTCACAATAAGAACCATAGGGTGATCCTGGATAGAGAACGCTTGGTCATTGAAAGA

CCCTACATCTTGCTTGGAGTCCTGCTGGTAATGTTCCTGAGTCTAATCGGACTGCTGGCCATTGCAGGGATCAGGCTTCA

CCGGGCCACCGTTGGAACTTCAGAGATCCAGAGTCGGCTGAATACCAATATTAAGTTGACCGAATCTATTGATCACCAGA

CTAAGGATGTCTTAACTCCCCTTTTTAAAATCATTGGCGATGAAGTCGGCATCAGAATTCCACAGAAATTCAGTGATCTT

GTCAAGTTCATCTCCGATAAGATTAAATTCCTCAACCCTGATAGAGAGTATGATTTCAGAGATCTCCGGTGGTGTATGAA

TCCCCCCGAGAGAGTCAAAATTAATTTTGATCAGTTTTGTGAGTACAAGGCTGCGGTTAAGTCAATTGAACATATATTTG

AGTCACCACTCAACAAGTCAAAAAAGCTGCAATCTTTGACTCTCGGGCCCGGAACAGGCTGTCTAGGCAGGACAGTAACA

AGAGCCCATTTCTCAGAACTTACAATGACCTTAATGGACCTGGATCTAGAGATGAAGCACAACGTGTCCTCAGTGTTTAC

CGTAGTTGAAGAGGGATTATTCGGAAGAACATATACCGTCTGGAGATCCGATGCCAGGGATCCGAGCACCGATCTAGGTA

TCGGCCATTTTTTAAGAGTCTTCGAGATTGGACTGGTAAGAGATCTTGGGCTGGGTCCCCCTGTTTTTCATATGACCAAC

TATCTCACGGTGAACATGAGTGATGACTATCGGAGATGTCTTTTAGCGGTAGGGGAGTTGAAGTTGACAGCCCTATGCAC

CTCATCTGAGACTGTGACACTGAGTGAGAGAGGAGTTCCAAGGAGGGAACCTCTTGTGGTTGTGATACTTAATCTAGCTG

GACCCACTCTAGGGGGAGAGCTATACAGTGTCTTGCCTACCTCTGATCTCATGGTGGAGAAACTCTATTTGTCTTCACAT

AGAGGGATCATCAAAGATGATGAGGCCAATTGGGTAGTGCCGTCTACCGATGTTCGTGATCTTCAGAACAAAGGTGAATG

TCTGGTGGAAGCATGCAAGACTCGACCTCCTTCATTTTGCAATGGCACAGGATCAGGCCCGTGGTCAGAAGGGAGAATCC

CTGCCTACGGGGTGATCAGGGTCAGTCTTGACTTAGCGAGTGACCCGGATGTAGTTATCACTTCAGTGTTTGGCCCACTG

ATACCTCACCTATCCGGCATGGATCTTTACAACAACCCGTTTTCAAAAGCTGTATGGTTGGCTGTACCACCTTATGAGCA

GTCATTTCTAGGAATGATAAATACAATTGGATTCCCTAACAGAGCAGAGGTTATGCCGCACATTTTGACCACAGAGATCA

GAGGCCCTCGGGGTCGTTGCCATGTTCCCATAGAATTGTCCCGCCGGGTTGATGACGATATCAAGATCGGGTCCAACATG

GTCATATTGCCGACGATGGACCTGAGGTATATTACAGCCACTTATGATGTTTCCAGGAGCGAGCATGCAATCGTGTACTA

TATCTATGACACGAGTCGCTCATCATCTTACTTCTACCCAGTTCGACTGAATTTCAAAGGCAATCCTCTCTCTTTGAGGA

TAGAGTGTTTCCCCTGGCGTCATAAGGTGTGGTGCTACCATGATTGTCTTATATACAACACCATAACAGGTGAAGAGGTC

CATACGAGAGGGCTGACCGGCATAGAGGTAACATGCAATCCAGTCTGAGCAGAGCTACGACCGTCACTCAGGCAGTCCCT

TGAGTCGCCACCGAGTCCAAGCAGCACAGCCTGGGACAATCAACGGCACAGCCCAGCCAACAATGTTATAAAAAACTTAG

GAGCCAGGGTTGTAGGGGCCATGGACTCTCTATCAGTCAATCAGGTCTTGTACCCTGAGGTTCATCTAGACAGCCCTATT

GTCACAAACAAACTAGTTGCCATCCTTGAGTACTCGGGTATCGACCATAACTATGTTCTTGAAGACCAGACTCTTGTCAA

GAATATTAGGTATAGGCTGGGGTGCGGTTTTTCAAATCAAATGATCATCAATAACAGGGGGGTGGGTGAAACAGTCAATT

CTAAACTTAAAAGTTACCCCCATAATCGTCATATCATATACCCGGATTGCAATAAGGAGTTGTTTTGTATCAAGGATAGC

TGCATATCTAAGAAGCTCTCGGAGCTATTCAAGAAGGGTAATTCCTTGTACTCTAAGATAAGTCACCAGGTACTGGATTG

TCTTAAGAGAGTCAATGGGAAATTAGGTCTGGGCACAGATCTTACCCATGGTCTGAAGGAGGGTATCCTTGACCTAGGGT

TGCACATGCATAGCTCTCAATGGTTCGAGACTTTCCTGTTCTGGTTCACTATCAAGACAGAGATGAGATCAATGATCAAA

GAACAGTCCCATATATGCCACAAGAGGAGGTATAACCCGACTTTTGTGTCGGGGGATGCATTCGAGGTGCTCGTATCGCG

AGACCTCGTTGTGATAATTGATAAGAATACCCAGTATGTCTTCTACTTAACATTTGAGCTGGTCCTTATGTATTGTGATG

TCATAGAGGGTAGACTTATGACGGAGACAGCCATGGCCATAGACCAGAGATATTCAGAACTTCTAAGCCGGGTCAGATAC

TTGTGGGATCTTATTGATGGGTTCTTCCCAACACTAGGCAACACCACATACCAAATTGTTGCTCTGCTTGAACCATTATC

GTTGGCTTATCTTCAACTTCAGGATGTCACTCTGGAGTTAAGAGGTGCTTTTTTAGACCATTGCTTCAAAGAACTCTATG

AGATACTGGAACATTGCGGCATTGACACAGCAGGCACTTACAATTCCATCACTGAAGGATTGGATTACGTATTCATCACC

CACGATATACATCTAACTGGGGAGATTTTTTCATTTTTTCGGAGTTTCGGACACCCCCGTCTAGAAGCAGTCACCGCTGC

GGAAAATGTCAGAAAACATATGAACCAACCGAAGGTAATCAGTTATGAGACTATGATGAAAGGCCATGCGGTATTTTGCG

GGATAATCATAAATGGTTTTCGGGACCGACATGGCGGCAGCTGGCCCCCTGTTGCACTGCCAGAACATGCCTCTGCTGCG

ATCCGGAATGCGCAGGCATCAGGTGAGGGACTGACCCATGACCTGTGTATAGACAACTGGAAATCCTTTGTTGGATTCAA

ATTTGGCTGCTTTATGCCACTCAGCCTAGATAGTGATTTGACCATGTATCTCAAAGACAAGGCATTGGCTGCACTGAAGA

ATGAGTGGGATTCAGTTTACCCGAAAGAATACCTCCGTTATAACCCACCTAGAGGGACAGAGTCAAGGCGATTGGTAGAG

GTATTCCTGAATGACTCCAGCTTTGATCCTTATAACATGATAATGTACGTGGTGAATGGCTCCTACCTTAAAGACCCTGA

GTTTAACCTCTCATACAGCCTAAAAGAGAAGGAGATAAAGGAGACAGGGCGGTTGTTTGCCAAAATGACCTATAAGATGC

GGGCCTGTCAGGTAATCGCTGAAAATCTGATATCGAATGGTGTTGGGAAGTATTTCCGAGACAATGGGATGGCAAAGGAC

GAGCATGACCTAACAAAAGCCCTTCACACCCTGGCAGTCTCAGGTGTTCCCAAAAACAACAAAGATAACCACCGAGGTGG

GCCTCCCAGAAGGACAACAAACCGAGGGGTGAGATCAAGCCAAGGCACCAAAACACAAGATAGAGACAAGGTTCAAGGGG

GACCTATGTACAACTATTTGCGATGCCAACCGATCAGCCCTGATCAGGGTGAGTCATACGAGACTGTTAGTGCATTCATC

ACCGCTGACCTTAAGAAGTATTGCCTGAATTGGAGATACGAGACAATCAGCATATTTGCACAGAGGCTGAATGAAATATA

TGGACTACCATCCTTCTTTCAATGGTTACACAGGATATTGGAAAAATCCGTACTCTACGTCAGTGACCCACATTGTCCTC

CCGATCTAGATAATCATATCCCTCTGGACAGTGTCCCCAATGCCCAAATATTCATCAAGTACCCAATGGGCGGAATAGAA

GGTTATTGCCAAAAACTATGGACAATCAGCACTATACCATACTTGTATCTGGCAGCCTATGAGAGCGGAGTAAGAATCGC

CTCACTGGTGCAGGGTGACAATCAGACAATCGCAGTGACAAAAAGAGTTCCAAGTTCTTGGCCTTATTCACTAAAAAAGA

GGGAGGCCTCCAAGGCCGCTCAAAATTACTTCGTAGTCTTAAGGCAAAGATTGCACGATGTAGGTCATCACTTAAAGGCT

AATGAGACCATAGTATCTTCTCACTTTTTTGTATATTCCAAAGGGATTTATTATGACGGCCTGCTAGTCTCACAATCACT

AAAGAGCATCGCCAGATGTGTCTTCTGGTCCGAGACTATAGTGGATGAAACCAGAGCGGCGTGCAGTAATATTGCAACAA

CTGTCGCTAAGAGTATAGAGAGGGGTTATGATAGGTACCTTGCATATTCTTTGAATATCCTCAAGATTTTCCAACAGATC

CTTATATCCCTTAACTTCACTATTAACACAACAATGACTCAGGATGTCGTGGCACCGATCATCGAGAACGGTGATTTGCT

GATAAGGATGGCACTCTTGCCAGCACCCATCGGGGGTCTCAATTATCTTAACATGAGCAGGTTATTTGTGAGAAATATCG

GTGACCCGGTCACCTCCTCTATAGCCGACCTGAAGAGGATGATAGATGCCGGGCTAATGCCAGAAGAAACATTGCATCAA

GTGATGACCCAGACCCCGGGAGAATCATCCTACCTAGATTGGGCAAGTGACCCTTATTCTGCCAACCTACCCTGCGTACA

GAGTATCACTCGCCTTCTCAAAAACATCACTGCACGATATATTTTAATCAGCAGCCCAAACCCGATGCTGAAAGGGTTAT

TTCATGAGGGGAGTAGAGATGAAGACGAGGAGCTTGCAAGTTTCCTAATGGATCGGCATATAATTGTTCCAAGAGCTGCA

CATGAAATCTTAGACCATAGTATAACCGGAGCAAGAGAAGCTATAGCCGGGATGTTGGACACCACCAAGGGTCTGATTAG

AACAAGTATGAAACGGGGCGGCCTCACCCCTCGAGTCTTAGCCCGCCTTTCCAATTATGATTATGAACAATTCCGATCCG

GGATAACACTATTAACAAAGAAAGGGCAGTGTTATCTCATTGACAAGGACTCGTGCTCGGTGCAGCTCGCTATAGCTCTG

AGGGGCCACATGTGGGCTAAGTTGGCTCGCGGGAGACCAATCTATGGGTTGGAGGTGCCTGATGTATTAGAATCGATGAA

CGGCTACCTTATCAAACGCCACGAGTCGTGTGCTATCTGTGAAACGGGCTCAAGTCACTACGGATGGTTTTTCGTCCCTG

CAGGGTGCCAGCTTGACGATGTATCTAGAGAGACTTCGGCTCTTCGTGTACCTTATGTCGGATCAACAACTGAGGAAAGG

ACAGACATGAAACTTGCATTTGTTAGGTCTCCAAGTAGATCCCTCAAATCAGCAGTTAGAATTGCAACAGTTTACTCATG

GGCTTACGGGGATGATGAGAAATCATGGAGTGAAGCCTGGATGCTAGCTAGGCAGAGAGCCAATATCACCTTAGATGAAT

TGAGAATGATCACTCCAGTCTCCACATCCACCAACTTAGCCCACCGGTTGAGGGATCGGAGTACCCAGGTGAAATACTCG

GGGACATCCCTTGTGAGGGTTGCAAGATACACAACCATCTCCAATGATAACTTGTCGTTTGTGATATCTGAGAAAAAAGT

GGATACCAACTTCATTTATCAGCAAGGGATGTTGCTTGGTCTTGGGATCCTTGAGAACCTCTTTAGGCTAGAGGCCACCA

CCGGAGTATCCAACACAGTGCTACACCTGCACGTGGAAACAGAATGTTGTGTTGTACCTATGGTGGATCACCCAAGGATA

CCGAGTCTCCGTAATATAAAAGTTACGGATGAGCTATGCACAAACCCTTTGATCTACGACAGGTCCCCCATCATAGAGCA

CGATGCAACCCGGCTATATTCACAGAGCCACAGGAGACATTTGGTGGAGTTTGTTACCTGGTCAACAAGCCAGCTTTACC

ATATACTGGCTAAGTCTACAGCAATGTCCATGATTGAATTGATCACGAGATTCGAGAAAGATCACATGAATGAAATAGCC

GCCCTGATTGGCGATGACGACATCAACAGCTTCATCACAGAATTTTTGCTAGTGGAGCCTAGATTATTTATAGTTTACCT

TGGTCAGTGTGCTGCCATCAATTGGGCCTTTGATATACATTATCATCGGCCCTCGGGCAAGTACCAGATGGGGGAACTTC

TCTACTCTCTGCTCTCTCGGATGAGCAAAGGAGTATATAAGATCTTTACTAATGCTTTGAGCCACCCCAAAGTTTACAAG

AAATTTTGGCGAAGTGGCATCATTGAGCCGATTCATGGCCCATCCTTGGATACACAGAATTTACATGTCACTGTCTGTGA

CATGATATATGGGTCCTATGTCACCTATTTGGATCTTTTGCTGAATGATGAGCTAGATGATTACCCGTATTTGCTCTGCG

AGAGTGATGAGGACGTGGTCACAGACAGGTTCGACAACATTCAAGCCAAACATCTCTGTGTATTGGCCGATGTATATTGC

AGCTCCAAGAGGTGTCCTTCAATAATCGGGATGTCTCCTATAGAAAAATGTACCATTCTCACACATTACATCAAGGGAGA

ATCAGTACAATCCCCGTCTGGGACCTCATGGAACACTGATCCCCTTGTAGTAGATCATTACTCATGCTCTCTGACCTACC

TTCGCCGCGGTTCCATCAAACAAATCAGGTTGAGGGTGGATCCTGGGTTTGTATTCGAGGCGTTGACAGACGTCGACTTT

AAACAACCTCGCAAAGCTAAGTCAGATATATCGGTCGTTGGGTTGACTGATTTCTCTCCCCCTTACGACAACGTCGGTGA

TTTTTTAGGGACTATCAACACATTGAGGCACGATCTGCCTGTCACCGGAACCGGGGTCTCGAACTATGAAGTCCACGCTT

ATCGTAGAATTGGCCTGAATTCATCAGCATGTTATAAAGCCGTAGAGATCTCCACGTTAATCAAGCCATCCTTAGAAGTC

GGAGAGCATGGATTGTTCTTAGGAGAAGGTTCCGGTTCAATGCTGGCTGCGTACAAAGAAGTTCTTAAATTAGCAAATTG

TTACTACAACAGCGGAGTAACAGCGGAGGGCAGAGCCGGACAGAGGGAAATATCTCCCTATCCTTCAGAGATGAGCCTAG

TAGAGAATCAAATGGGGATAGAGCGGAGTGTTAAAGTGCTGTTCAACGGGAAACCTGAAGTAACTTGGGTGGGGACCACA

GATTGCTACAAGTACATAATCAGTAACATCCAAACCTCTAGTCTGGGTTTCATACATTCAGATATTGAGACGCTCCCAAC

CAAAGATGCTGTTGAGAAATTGGAGGAATTTGCCTCTATCCTATCCTTATCCTTGATTTTGGGGAAAATCGGCTCTATTA

CAGTTGTCAAAATTATGCCCATTAGCGGGGATTTTACCCAAGGCTTCATAGCCTATGCCATCCAATATTTTAGGGAGAGC

CTGCTTGCCTACCCGAGATATAGTAACTTCATCTCGACTGAGTGTTACCTTATTATGATAGGATTAAAGGCCAATCGATT

GATAAACCCAGAAGCCATTAAGCAAAGCATAATCAGAGCGGGGGTCAGGACTGCACCAGGACTTGTGAGCCATGTATTAT

CAGAGAAACAAAAAGGTTGTATTCAATCTTTCTTGGGTGATCCTTATATCCAAGGAGACTTCAATAAGCACCTTAAATCT

CTAACCTCTATTGAGAAAATCCTGGTAAATTGTGGTCTCTCGATCAATGGCATAAAAATCTGTAGAGATCTAATCCACCA

TGATATCGCCTCCGGTCCAGACGGCCTTATGAGCTCTACAATTATTTTATATAGGGAATTGGCTCATTTCAAAGATAATA

TAAGGAGTCAGCACGGAATGTTCCACCCCTACCCGGTACTGGCCAACAGCAGGCAACGTGAATTAATCCTTCGAATAGCC

AAGAAATTCTGGGGGTATGTCTTGCTATATTCTGATGACCCGGCACTAATCAGACAAACAATCAAGAACTTGAAACGGAA

TCATCTAACCTTTGACTTACATAGTAACCCATTTATTAAAGGGCTATCCAAGGCTGAGAAACTGCTGGTGCGGACGAGTT

CACTAAGAAGAGAATGGTTGTTCACTCTCGAAACGAAAGAAGTAAAAGAGTGGTTCAAATTGGTAGGTTACAGTGCACTC

ATCAGAGGCTAGTCAGCTATGCATCTGCCCCCTTCTCCTCCGCCATGAGACCCCACTGGCGATCCAGAAGATTAAAGAAA

ACTACATATTGGATAAGTATCTATTCCCAGCTTTGTCTGGT

>MF443343.1/goat/China/2014-04-02

ACCAAACTTTGTTTGGTAAGGATAGATCTTATAATAACTATGGACTGGCAAACTTAGGAGTAAAGATCCTACTGTCGGGG

GGAGGAGGAGGAGCAAGATCTTTGATTATGGCGACTCTCCTTAAAAGCTTAGCATTGTTCAAAAGGAACAAAGACAAAGC

GCCGACAGCATCAGGTTCAGGAGGGGCCATCCGGGGGATTAAGAATGTTATCATAGTCCCGATTCCCGGAGACTCGTCCA

TCACTACCCGTTCAAGACTGCTCGACAGGCTTGTCAGATTGGCCGGAGATCCTGACATCAACGGGTCAAAGCTGACCGGG

GTGATGATCAGCATGTTATCCTTGTTCGTAGAGTCACCCGGGCAATTGATACAGCGAATCACAGATGATCCGGATGTTAG

TATCCGCCTTGTTGAGGTAGTTCAAAGTACTAGGTCCCAGTCCGGGTTGACCTTTGCATCACGTGGTGCTGATTTAGACA

ACGAGGCAGACATGTATTTTTCAACTGAGGGGCCCTCGAGTGGAGGTAAGAAAAGGATCAACTGGTTTGAGAACAGAGAA

ATAATAGACATAGAGGTGCAGGATCCAGAAGAGTTCAATATGTTGTTAGCCTCCATACTAGCACAAGTCTGGATCCTTCT

GGCCAAGGCTGTTACGGCACCAGATACGGCAGCTGACTCAGAACTGAGAAGGTGGGTTAAATACACACAACAAAGGAGAG

TGATTGGGGAATTTCGCCTTGACAAGGGGTGGCTGGACGCAGTCCGCAACAGAATTGCAGAAGATCTATCACTCCGGCGG

TTCATGGTATCTCTCATACTTGACATCAAGAGGACCCCAGGCAACAAGCCAAGGATTGCAGAAATGATCTGCGACATTGA

CAACTATATTGTCGAAGCAGGACTCGCCAGTTTCATCCTTACTATCAAATTTGGTATCGAAACCATGTATCCTGCATTAG

GGCTCCACGAGTTTGCCGGAGAATTGTCCACTATTGAGTCCTTGATGAACTTGTATCAACAGCTAGGAGAGGTTGCACCC

TACATGGTAATTCTAGAGAATTCAATTCAGAACAAGTTTAGTGCAGGAGCTTATCCCCTCCTCTGGAGCTATGCGATGGG

TGTCGGAGTCGAGCTGGAGAACTCAATGGGGGGCTTGAACTTTGGCAGGTCATATTTTGACCCTGCCTATTTTCGTCTCG

GACAGGAGATGGTCAGAAGATCTGCAGGAAAGGTCAGCTCTGTGATTGCGGCTGAGCTCGGCATCACAGCAGAGGAAGCT

AAACTAGTCTCGGAAATCGCCTCGCAGGCTGGGGACGAAAGAACCGCCAGAGGGACTGGGCCTCGACAGGCGCAGGTTTC

CTTCCTTCAGCACCAAACAGGAGGGGGAGAGTCGTCCGCACCAGCGACCAGAGAAGGGGTCAAAGCTGCGATCCCAAACG

GAGCTGAAGAAAGGGACAGAAAGCAAACACGCCCAGGAAGGCCCAGAGGAGAGACCTCCGGCCAACTGCTCCTGGACATC

ATGCCAGAGGATGAGATCTTGCGAGAGTCTGGTCAAAACCCTCGTGAGGCTCAAAGATCGGCCGAGGCACTCTTCAGGCT

GCAGGCCATGGCCAAGATTCTGGAGGACCAGGAGGAGGGAGAAGACAACAGTCAGGTCTACAACGACAAGGATCTCCTCG

GCTAAGCAGACGCACCCTCTGTCGAAATCAGTGACAAGACATCTCCTACCAGTATTATAAAAAACTTAGGACCCAGGTCC

AAGCAAGCACACATCGACACTCCAACCAGTCGAGCGGAGACCACCGATGGCAGAAGAACAAGCATACCATGTCAACAAGG

GGCTGGAATGTATCAAGTCTCTCAAAGCCTCACCCCCGGATCTATCCACCATCAGAGATACCATCGAGAGCTGGAGAGAG

GGGCTTAGCCCCTCGGGCCGTGCAACACCGAACCCTGATACGTCCGAGGGAGATCATCAGAATATCAACCAATCATGCTC

ACCAGCAATCGGACCAAACAAAGTCTACTTGTCTCCTGAAGATAATCTCGGATTTAGAGAGATCACTGGCAACGACTGTG

AGGCTGGGCTCGGAGGGGTCCAGAGAGAAGGATCCAACTCTCAAGTACAGCGTTACCATGTTTATAGTCACGGGGGTGAA

GAGATTGAAGGACTCGAGGATGCTGACTCTCTCGTGGTTCAAGCAGATCCTCCAGTTGCTAACGTCTTCAATGGAGGAGA

GGATGGATCTGACGACAGCGATGTGGACTCTGGCCCAGATGATCCCGGCAGAGATACTCTATATGACCGGGGATCTGTTG

CCGGCAATGGTATCGCTAGGTCCACAGATGTCGAAAAACTAGAAGGTGCTGATATTCAAGAAGTTCTTAACTCCCAGAAA

GGCAAAGGAGGAAGATTCCAGGGCGGGAAAACCTTGCGAGTCCCGGAAATACCCGATGTCAAGCACTCCAGACCATCAGC

CCAATCAATTAAAAAGGGCACAGACGGGAACTCAGTCTCATCTGGAACGGTGATAGAGTGTTTATCGATAAGTGGTGCAA

CCCAAACTGTGCCAGAGTCAAGATGGGAGTCATCAGAGCAAAATGCGTCTGTGGGGAGTGTCCTCAAGTCTGCGAGGAGT

GCAAAGACGATCCAGGGGTCGACACAAGAATCTGGTACCATAGCATCACTGACTCAGCCTAAAGAGAATGACTCCGAGTA

TGAGTATGAGGATGACCTATTTACAGAGATTCAGGACATCCGTGCAAGCATTGCCAAGATCCATGATGACAATAAAACTA

TCCTCTCAAAGCTTGATTCTATACTGTTATTGAAAGGGGAAGTCGACACTATCAAGAAACAAATCAGCAAGCAGAATATA

AGTATATCCACCATTGAGGGCCATCTCTCCAGTATAATGATAGCCATCCCTGGCTTTGGGAAGGACATCAAGGACCCAAC

ATCCGAAGTCGAGTTGAACCCAGATCTAAGACCTATAATAAGCCGTGACTCTGGCAGAGCTCTCGCGGAGGTCCTCAAGA

AACCCGCTGTGGATAGGTCTCAGAAAATTGGAACCAAAGCCAACTCCAGCTCAAAGGGTCAGCTTCTTAAGGATCTCCAG

CTAAAACCTGTCGACAAGCAGGCAAGCTCTGCAATCGGGTTTGTCCCGTCCGACCATGAATCATCCAGAAATGTCATCCG

CTCCATAATCAAGTCGAGCAAGCTAAATATTGATCACAAGGACTATCTTCTAGATTTACTAAATGATGTGAAAGGCTCCA

AGGATCTTAAGGAATTTCACAAGATGCTAACAGCAATTCTCGCCAAGCACCCGTAACACATCCTCCAGTCATCATCTCAT

ACTCGACTAAAAACATCCTTTCAATCAGGCTATTACAAAAAACTTAGGAGCAAGGGCAACTGAGCTTCGCAGACAGGATG

ACCGAGATCTACGACTTCGATAAATCAGCATGGGATGTCAAAGGGTCAATTGCCCGCATAGAACCCACCACCTACCACGA

CGGCCGACTGATACCCCAGGTGAGGGTCATCGATCCTGGTCTGGGAGACAGAAAAGATGAGTGCTTTATGTACCTGTTTC

TCCTAGGAGTGATTGAGGATAACGACCCCCTGTCTCCCCCCGTCGGAAGAACCTTTGGCTCTCTACCTCTAGGGGTTGGT

AGGTCAACTGCCAAGCCAGAAGAACTACTAAGGGAGGCCACAGAATTAGATATAGTGGTGAGGCGCACTGCAGGAGTAAA

TGAGAAACTGGTATTTTACAACAACACTCCGCTGTCCTTGTTAACGCCCTGGAAGAAAGTTCTGACAACCGGAAGTGTGT

TTAGCGCTAACCAGGTTTGCAATGCAGTCAACCTAGTCCCGCTTGATACTCCCCAGAGATTCAGGGTTGTGTACATGAGC

ATAACTAGATTATCAGACAATGGTTACTATAGTGTGCCCAGAAGAATGCTGGAGTTCCGCTCGGCCAATGCAGTAGCCTT

CAACATCTTGGTTACACTAAGAATTGAAAATGGCACAAACCCTAGAAGATACATAGTCGGCTCATGGGAGAATTCAGAGG

TCACATTTATGGTACACGTGGGCAACTTTAGGAGAAAGAAGAACGAAGTATACTCTGCTGATTACTGCAAAATGAAGATT

GAAAAGATGGGTTTAGTTTTTGCCCTGGGTGGAATAGGCGGAACAAGTCTCCATATTAGAAGCACCGGGAAAATGAGCAA

AACCCTCCATGCACAGCTGGGGTTCAAGAAAATCTTATGTTACCCCCTAATGGATGTCAATGAGGATCTTAACCGATATC

TCTGGCGGGCAGAGTGCCGAATAGTCAAAATCCAAGCTGTCTTACAGCCATCAGTACCCCAAGAGTTCCGTGTCTACGAT

GATGTCATCATCAACGATGATCAAGGCTTGTTCAAGATCTTGTAGTTCATTTGCAAAGGAACGAACCCACCCCCAGGAAG

GGGCCCCACCCTCCCCCG-CCAAAAAACACACTCCAAGGCTCCCCGGCACAGCGCCAACACAGCTACTCCCGCGATGGAA

CGAGGTCGGCC-CCCGAACAAATCCGGTCCAGGGGGGGGGGCCCCCCCCCTGACCAACCGGGCGCCGACCGAGCAGGCCG

GCGAGAAGCCCCCAAGAGGCCCCGACGAAGACCGAGCACCCCATCCCCCC-AAAAACGTCCCCC-AACACGCCCGAGGCT

AGCCACCGGACACCCCCAGACCCGCCACCGGCGAGACAGAACCCCAAGGGGGGACCCTCCAAACCAAAGGAATCACCGGG

GTAGGCTCCGCCGAACCAGCACAGACCCGACCCAAGCAAGGCGGACCCCCAACACGAAACTGCCCAGCACACCGTACTCA

TCATCAAACAAAACTTAGGGGCCAAGTCCATACGAGAGGCCCTAGTCCGTCAGGCGGAGGGACGCAGAAAGGAAGGAGAC

ACGACCGTCCCAGAAGGAGGACCCCCAACCACCGAAACGCCCGCCCCCAGGGAAGGAG--CCA-CCCCCACGCAGCGAAG

GAGCGGGTGGGAGACCTCGGGGAGGGGCCGACCGGGACCCGGGGGCCGGCGGCGCCGGCGGTGAGCAGGCCGGGCGCATA

CCTGACCCCACCGGAGGATCCGAGGAGGGCCCGACCCCGCGTGGACCAAACGGCAGGGAGCCCAACCCCCCCAAAAAGAG

AGGCGAGGCCCCCCCTCCCTCCCCCCCCGCACACAACCCGCAGACAGACCGAGGGAGGGCCATCCCCCAGCGAGACAGGC

GGCTCCCTCGCAGCCGAGCCGCAGCTCCCCAGAATAGGACACCCCCGCACACTCCGAACACCCCCTCGCCCCCCGCCACC

GGGACAGGGGCTCCACCACCACAGACCGGAGCCACCGACAGCCAGCCAGAGGTCCCCTCCAGTCCCGCCCGGCGCCCCCC

CGAAGAGGCCCCGGCAAGGCCCCCCAGCCCGACGTCCCCCGCACGCACCGAAAGAAGGACCCGCCCGAGAACGTGCGGCA

CATCCATGCGCAAACATCATGACGCGGGTCGCAATTTTGACATTTCTGTTTCTTTTCCCAAATGTTGTTGCGTGTCAGAT

TCACTGGGGCAATCTATCCAAGATCGGGATTGTAGGAACAGGGAGTGCCAGCTACAAGGTGATGACTAGGCCAAGCCACC

AGACTCTGGTTATAAAGTTAATGCCAAATATAACGGCCATCGACAATTGTACAAAGTCAGAGATTGCAGAGTACAAGAGA

TTGCTGATCACAGTGTTAAAGCCTGTAGAGGATGCTCTGTCGGTGATAACCAAGAATGTAAGACCAATTCAAACTCTAAC

ACCTGGGCGTAGAACCCGCCGTTTTGCTGGAGCTGTTCTGGCCGGGGTAGCACTTGGAGTTGCGACAGCCGCTCAGATAA

CTGCAGGAGTCGCCCTTCATCAATCATTGATGAACTCCCAAGCAATTGAGAGTTTAAAAACCAGTCTTGAGATGTCGAAT

CAGGCAATAGAAGAAATCAGACTTGCAAATAAGGAGACCATACTGGCAGTACAGGGCGTCCAGGATTATATCAACAATGA

GCTCGTCCCTTCTGTTCATAGAATGTCATGCGAGCTGGTAGGTCACAAGCTCGGCCTCAAGCTCCTTAGGTACTACACCG

AGATCCTGTCCATATTCGGGCCCAGTCTTCGAGACCCGATATCTGCCGAAATATCAATCCAGGCACTTAGTTATGCATTA

GGCGGAGACATTAATAAAATCCTGGACAAGCTTGGGTATAGCGGTGGGGATTTCCTTGCCATCCTAGAAAGCAAGGGAAT

AAAGGCCCGGGTTACATATGTGGACACAAGAGATTACTTTATAATCCTTAGCATCGCCTACCCAACCTTATCTGAGATCA

AGGGAGTGATAGTTCACAAGATAGAAGCTATAACATACAACATTGGGGCACAGGAGTGGTATACTACTATCCCTAAATAT

GTAGCCACTCAGGGGTATCTGATATCGAACTTTGATGAGACGTCATGCGTATTCACTCCAGAGGGGACAGTTTGCAGCCA

GAATGCGTTGTACCCAATGAGCCCATTGCTTCAGGAATGTTTCAGGGGGTCAACAAAATCGTGCGCCAGAACCTTAGTTT

CAGGGACCATAAGTAATAGATTTATCCTATCAAAAGGGAACCTGATTGCAAATTGTGCGTCAGTTTTGTGCAAATGTTAC

ACAACAGAGACAGTTATCAGCCAAGATCCTGACAAACTACTAACTGTTGTAGCATCCGACAAGTGTCCTGTAGTTGAGGT

GGATGGAGTGACAATACAGGTCGGCAGTCGAGAGTATCCGGATTCTGTATACTTACACAAAATAGACTTAGGTCCAGCCA

TCTCCCTAGAAAAACTGGATGTAGGCACCAATTTAGGCAATGCAGTCACAAGACTGGAGAATGCAAAGGAGCTCCTAGAT

GCATCAGACCAAATACTGAAGACTGTTAAAGGGGTACCTTTCAGTGGGAATATGTACATAGCACTGGCAGCTTGCATAGG

AGTATCCCTGGGCCTTGTCACATTAATATGCTGCTGTAAGGGGAGGTGTAAGAACAAGGAAATCCCTATCTCCAAAATCA

ACCCAAGGCTCAAACCCGACCTGACCGGGACCTCAAAGTCGTACGTAAGATCACTGTAGTCAGAATCACCTGGATCATCT

GGCATCGCACACATACATGCACGACACAGGCAGTCCGAGGACGCAAGAAACCCAGCCTCCGGTCACCCACCCGACCCCAC

TCCACGCTCCACCACACATTAGTCATCAAACAAAACTTAGGACGAAAGGTCAATCACCATGTCCGCACAAAGGGAGAGGA

TCAATGCCTTCTACAAAGACAATCCTCACAATAAGAACCATAGGGTGATCCTGGATAGAGAACGCTTGGTCATTGAAAGA

CCCTACATCTTGCTTGGAGTCCTGCTGGTAATGTTCCTGAGTCTAATCGGACTGCTGGCCATTGCAGGGATCAGGCTTCA

CCGGGCCACCGTTGGAACTTCAGAGATCCAGAGTCGGCTGAATACCAATATTAAGTTGACCGAATCTATTGATCACCAGA

CTAAGGATGTCTTAACTCCCCTTTTTAAAATCATTGGCGATGAAGTCGGCATCAGAATTCCACAGAAATTCAGTGATCTT

GTCAAGTTCATCTCCGATAAGATTAAATTCCTCAACCCTGATAGAGAGTATGATTTCAGAGATCTCCGGTGGTGTATGAA

TCCCCCCGAGAGAGTCAAAATTAATTTTGATCAGTTTTGTGAGTACAAGGCTGCGGTTAAGTCAATTGAACATATATTTG

AGTCACCACTCAACAAGTCAAAAAAGCTGCAATCTTTGACTCTCGGGCCCGGAACAGGCTGTCTAGGCAGGACAGTAACA

AGAGCCCATTTCTCAGAACTTACAATGACCTTAATGGACCTGGATCTAGAGATGAAGCACAACGTGTCCTCAGTGTTTAC

CGTAGTTGAAGAGGGATTATTCGGAAGAACATATACCGTCTGGAGATCCGATGCCAGGGATCCGAGCACCGATCTAGGTA

TCGGCCATTTTTTAAGAGTCTTCGAGATTGGACTGGTAAGAGATCTTGGGCTGGGTCCCCCTGTTTTTCATATGACCAAC

TATCTCACGGTGAACATGAGTGATGACTATCGGAGATGTCTTTTAGCGGTAGGGGAGTTGAAGTTGACAGCCCTATGCAC

CTCATCTGAGACTGTGACACTGAGTGAGAGAGGAGTTCCAAGGAGGGAACCTCTTGTGGTTGTGATACTTAATCTAGCTG

GACCCACTCTAGGGGGAGAGCTATACAGTGTCTTGCCTACCTCTGATCTCATGGTGGAGAAACTCTATTTGTCTTCACAT

AGAGGGATCATCAAAGATGATGAGGCCAATTGGGTAGTGCCGTCTACCGATGTTCGTGATCTTCAGAACAAAGGTGAATG

TCTGGTGGAAGCATGCAAGACTCGACCTCCTTCATTTTGCAATGGCACAGGATCAGGCCCGTGGTCAGAAGGGAGAATCC

CTGCCTACGGGGTGATCAGGGTCAGTCTTGACTTAGCGAGTGACCCGGATGTAGTTATCACTTCAGTGTTTGGCCCACTG

ATACCTCACCTATCCGGCATGGATCTTTACAACAACCCGTTTTCAAAAGCTGTATGGTTGGCTGTACCACCTTATGAGCA

GTCATTTCTAGGAATGATAAATACAATTGGATTCCCTAACAGAGCAGAGGTTATGCCGCACATTTTGACCACAGAGATCA

GAGGCCCTCGGGGTCGTTGCCATGTTCCCATAGAATTGTCCCGCCGGGTTGATGACGATATCAAGATCGGGTCCAACATG

GTCATATTGCCGACGATGGACCTGAGGTATATTACAGCCACTTATGATGTTTCCAGGAGCGAGCATGCAATCGTGTACTA

TATCTATGACACGAGTCGCTCATCATCTTACTTCTACCCAGTTCGACTGAATTTCAAAGGCAATCCTCTCTCTTTGAGGA

TAGAGTGTTTCCCCTGGCGTCATAAGGTGTGGTGCTACCATGATTGTCTTATATACAACACCATAACAGGTGAAGAGGTC

CATACGAGAGGGCTGACCGGCATAGAGGTAACATGCAATCCAGTCTGAGCAGAGCTACGACCGTCACTCAGGCAGTCCCT

TGAGTCGCCACCGAGTCCAAGCAGCACAGCCTGGGACAATCAACGGCACAGCCCAGCCAACAATGTTATAAAAAACTTAG

GAGCCAGGGTTGTAGGGGCCATGGACTCTCTATCAGTCAATCAGGTCTTGTACCCTGAGGTTCATCTAGACAGCCCTATT

GTCACAAACAAACTAGTTGCCATCCTTGAGTACTCGGGTATCGACCATAACTATGTTCTTGAAGACCAGACTCTTGTCAA

GAATATTAGGTATAGGCTGGGGTGCGGTTTTTCAAATCAAATGATCATCAATAACAGGGGGGTGGGTGAAACAGTCAATT

CTAAACTTAAAAGTTACCCCCATAATCGTCATATCATATACCCGGATTGCAATAAGGAGTTGTTTTGTATCAAGGATAGC

TGCATATCTAAGAAGCTCTCGGAGCTATTCAAGAAGGGTAATTCCTTGTACTCTAAGATAAGTCACCAGGTACTGGATTG

TCTTAAGAGAGTCAATGGGAAATTAGGTCTGGGCACAGATCTTACCCATGGTCTGAAGGAGGGTATCCTTGACCTAGGGT

TGCACATGCATAGCTCTCAATGGTTCGAGACTTTCCTGTTCTGGTTCACTATCAAGACAGAGATGAGATCAATGATCAAA

GAACAGTCCCATATATGCCACAAGAGGAGGTATAACCCGACTTTTGTGTCGGGGGATGCATTCGAGGTGCTCGTATCGCG

AGACCTCGTTGTGATAATTGATAAGAATACCCAGTATGTCTTCTACTTAACATTTGAGCTGGTCCTTATGTATTGTGATG

TCATAGAGGGTAGACTTATGACGGAGACAGCCATGGCCATAGACCAGAGATATTCAGAACTTCTAAGCCGGGTCAGATAC

TTGTGGGATCTTATTGATGGGTTCTTCCCAACACTAGGCAACACCACATACCAAATTGTTGCTCTGCTTGAACCATTATC

GTTGGCTTATCTTCAACTTCAGGATGTCACTCTGGAGTTAAGAGGTGCTTTTTTAGACCATTGCTTCAAAGAACTCTATG

AGATACTGGAACATTGCGGCATTGACACAGCAGGCACTTACAATTCCATCACTGAAGGATTGGATTACGTATTCATCACC

CACGATATACATCTAACTGGGGAGATTTTTTCATTTTTTCGGAGTTTCGGACACCCCCGTCTAGAAGCAGTCACCGCTGC

GGAAAATGTCAGAAAACATATGAACCAACCGAAGGTAATCAGTTATGAGACTATGATGAAAGGCCATGCGGTATTTTGCG

GGATAATCATAAATGGTTTTCGGGACCGACATGGCGGCAGCTGGCCCCCTGTTGCACTGCCAGAACATGCCTCTGCTGCG

ATCCGGAATGCGCAGGCATCAGGTGAGGGACTGACCCATGACCTGTGTATAGACAACTGGAAATCCTTTGTTGGATTCAA

ATTTGGCTGCTTTATGCCACTCAGCCTAGATAGTGATTTGACCATGTATCTCAAAGACAAGGCATTGGCTGCACTGAAGA

ATGAGTGGGATTCAGTTTACCCGAAAGAATACCTCCGTTATAACCCACCTAGAGGGACAGAGTCAAGGCGATTGGTAGAG

GTATTCCTGAATGACTCCAGCTTTGATCCTTATAACATGATAATGTACGTGGTGAATGGCTCCTACCTTAAAGACCCTGA

GTTTAACCTCTCATACAGCCTAAAAGAGAAGGAGATAAAGGAGACAGGGCGGTTGTTTGCCAAAATGACCTATAAGATGC

GGGCCTGTCAGGTAATCGCTGAAAATCTGATATCGAATGGTGTTGGGAAGTATTTCCGAGACAATGGGATGGCAAAGGAC

GAGCATGACCTAACAAAAGCCCTTCACACCCTGGCAGTCTCAGGTGTTCCCAAAAACAACAAAGATAACCACCGAGGTGG

GCCTCCCAGAAGGACAACAAACCGAGGGGTGAGATCAAGCCAAGGCACCAAAACACAAGATAGAGACAAGGTTCAAGGGG

GACCTATGTACAACTATTTGCGATGCCAACCGATCAGCCCTGATCAGGGTGAGTCATACGAGACTGTTAGTGCATTCATC

ACCGCTGACCTTAAGAAGTATTGCCTGAATTGGAGATACGAGACAATCAGCATATTTGCACAGAGGCTGAATGAAATATA

TGGACTACCATCCTTCTTTCAATGGTTACACAGGATATTGGAAAAATCCGTACTCTACGTCAGTGACCCACATTGTCCTC

CCGATCTAGATAATCATATCCCTCTGGACAGTGTCCCCAATGCCCAAATATTCATCAAGTACCCAATGGGCGGAATAGAA

GGTTATTGCCAAAAACTATGGACAATCAGCACTATACCATACTTGTATCTGGCAGCCTATGAGAGCGGAGTAAGAATCGC

CTCACTGGTGCAGGGTGACAATCAGACAATCGCAGTGACAAAAAGAGTTCCAAGTTCTTGGCCTTATTCACTAAAAAAGA

GGGAGGCCTCCAAGGCCGCTCAAAATTACTTCGTAGTCTTAAGGCAAAGATTGCACGATGTAGGTCATCACTTAAAGGCT

AATGAGACCATAGTATCTTCTCACTTTTTTGTATATTCCAAAGGGATTTATTATGACGGCCTGCTAGTCTCACAATCACT

AAAGAGCATCGCCAGATGTGTCTTCTGGTCCGAGACTATAGTGGATGAAACCAGAGCGGCGTGCAGTAATATTGCAACAA

CTGTCGCTAAGAGTATAGAGAGGGGTTATGATAGGTACCTTGCATATTCTTTGAATATCCTCAAGATTTTCCAACAGATC

CTTATATCCCTTAACTTCACTATTAACACAACAATGACTCAGGATGTCGTGGCACCGATCATCGAGAACGGTGATTTGCT

GATAAGGATGGCACTCTTGCCAGCACCCATCGGGGGTCTCAATTATCTTAACATGAGCAGGTTATTTGTGAGAAATATCG

GTGACCCGGTCACCTCCTCTATAGCCGACCTGAAGAGGATGATAGATGCCGGGCTAATGCCAGAAGAAACATTGCATCAA

GTGATGACCCAGACCCCGGGAGAATCATCCTACCTAGATTGGGCAAGTGACCCTTATTCTGCCAACCTACCCTGCGTACA

GAGTATCACTCGCCTTCTCAAAAACATCACTGCACGATATATTTTAATCAGCAGCCCAAACCCGATGCTGAAAGGGTTAT

TTCATGAGGGGAGTAGAGATGAAGACGAGGAGCTTGCAAGTTTCCTAATGGATCGGCATATAATTGTTCCAAGAGCTGCA

CATGAAATCTTAGACCATAGTATAACCGGAGCAAGAGAAGCTATAGCCGGGATGTTGGACACCACCAAGGGTCTGATTAG

AACAAGTATGAAACGGGGCGGCCTCACCCCTCGAGTCTTAGCCCGCCTTTCCAATTATGATTATGAACAATTCCGATCCG

GGATAACACTATTAACAAAGAAAGGGCAGTGTTATCTCATTGACAAGGACTCGTGCTCGGTGCAGCTCGCTATAGCTCTG

AGGGGCCACATGTGGGCTAAGTTGGCTCGCGGGAGACCAATCTATGGGTTGGAGGTGCCTGATGTATTAGAATCGATGAA

CGGCTACCTTATCAAACGCCACGAGTCGTGTGCTATCTGTGAAACGGGCTCAAGTCACTACGGATGGTTTTTCGTCCCTG

CAGGGTGCCAGCTTGACGATGTATCTAGAGAGACTTCGGCTCTTCGTGTACCTTATGTCGGATCAACAACTGAGGAAAGG

ACAGACATGAAACTTGCATTTGTTAGGTCTCCAAGTAGATCCCTCAAATCAGCAGTTAGAATTGCAACAGTTTACTCATG

GGCTTACGGGGATGATGAGAAATCATGGAGTGAAGCCTGGATGCTAGCTAGGCAGAGAGCCAATATCACCTTAGATGAAT

TGAGAATGATCACTCCAGTCTCCACATCCACCAACTTAGCCCACCGGTTGAGGGATCGGAGTACCCAGGTGAAATACTCG

GGGACATCCCTTGTGAGGGTTGCAAGATACACAACCATCTCCAATGATAACTTGTCGTTTGTGATATCTGAGAAAAAAGT

GGATACCAACTTCATTTATCAGCAAGGGATGTTGCTTGGTCTTGGGATCCTTGAGAACCTCTTTAGGCTAGAGGCCACCA

CCGGAGTATCCAACACAGTGCTACACCTGCACGTGGAAACAGAATGTTGTGTTGTACCTATGGTGGATCACCCAAGGATA

CCGAGTCTCCGTAATATAAAAGTTACGGATGAGCTATGCACAAACCCTTTGATCTACGACAGGTCCCCCATCATAGAGCA

CGATGCAACCCGGCTATATTCACAGAGCCACAGGAGACATTTGGTGGAGTTTGTTACCTGGTCAACAAGCCAGCTTTACC

ATATACTGGCTAAGTCTACAGCAATGTCCATGATTGAATTGATCACGAGATTCGAGAAAGATCACATGAATGAAATAGCC

GCCCTGATTGGCGATGACGACATCAACAGCTTCATCACAGAATTTTTGCTAGTGGAGCCTAGATTATTTATAGTTTACCT

TGGTCAGTGTGCTGCCATCAATTGGGCCTTTGATATACATTATCATCGGCCCTCGGGCAAGTACCAGATGGGGGAACTTC

TCTACTCTCTGCTCTCTCGGATGAGCAAAGGAGTATATAAGATCTTTACTAATGCTTTGAGCCACCCCAAAGTTTACAAG

AAATTTTGGCGAAGTGGCATCATTGAGCCGATTCATGGCCCATCCTTGGATACACAGAATTTACATGTCACTGTCTGTGA

CATGATATATGGGTCCTATGTCACCTATTTGGATCTTTTGCTGAATGATGAGCTAGATGATTACCCGTATTTGCTCTGCG

AGAGTGATGAGGACGTGGTCACAGACAGGTTCGACAACATTCAAGCCAAACATCTCTGTGTATTGGCCGATGTATATTGC

AGCTCCAAGAGGTGTCCTTCAATAATCGGGATGTCTCCTATAGAAAAATGTACCATTCTCACACATTACATCAAGGGAGA

ATCAGTACAATCCCCGTCTGGGACCTCATGGAACACTGATCCCCTTGTAGTAGATCATTACTCATGCTCTCTGACCTACC

TTCGCCGCGGTTCCATCAAACAAATCAGGTTGAGGGTGGATCCTGGGTTTGTATTCGAGGCGTTGACAGACGTCGACTTT

AAACAACCTCGCAAAGCTAAGTCAGATATATCGGTCGTTGGGTTGACTGATTTCTCTCCCCCTTACGACAACGTCGGTGA

TTTTTTAGGGACTATCAACACATTGAGGCACGATCTGCCTGTCACCGGAACCGGGGTCTCGAACTATGAAGTCCACGCTT

ATCGTAGAATTGGCCTGAATTCATCAGCATGTTATAAAGCCGTAGAGATCTCCACGTTAATCAAGCCATCCTTAGAAGTC

GGAGAGCATGGATTGTTCTTAGGAGAAGGTTCCGGTTCAATGCTGGCTGCGTACAAAGAAGTTCTTAAATTAGCAAATTG

TTACTACAACAGCGGAGTAACAGCGGAGGGCAGAGCCGGACAGAGGGAAATATCTCCCTATCCTTCAGAGATGAGCCTAG

TAGAGAATCAAATGGGGATAGAGCGGAGTGTTAAAGTGCTGTTCAACGGGAAACCTGAAGTAACTTGGGTGGGGACCACA

GATTGCTACAAGTACATAATCAGTAACATCCAAACCTCTAGTCTGGGTTTCATACATTCAGATATTGAGACGCTCCCAAC

CAAAGATGCTGTTGAGAAATTGGAGGAATTTGCCTCTATCCTATCCTTATCCTTGATTTTGGGGAAAATCGGCTCTATTA

CAGTTGTCAAAATTATGCCCATTAGCGGGGATTTTACCCAAGGCTTCATAGCCTATGCCATCCAATATTTTAGGGAGAGC

CTGCTTGCCTACCCGAGATATAGTAACTTCATCTCGACTGAGTGTTACCTTATTATGATAGGATTAAAGGCCAATCGATT

GATAAACCCAGAAGCCATTAAGCAAAGCATAATCAGAGCGGGGGTCAGGACTGCACCAGGACTTGTGAGCCATGTATTAT

CAGAGAAACAAAAAGGTTGTATTCAATCTTTCTTGGGTGATCCTTATATCCAAGGAGACTTCAATAAGCACCTTAAATCT

CTAACCTCTATTGAGAAAATCCTGGTAAATTGTGGTCTCTCGATCAATGGCATAAAAATCTGTAGAGATCTAATCCACCA

TGATATCGCCTCCGGTCCAGACGGCCTTATGAGCTCTACAATTATTTTATATAGGGAATTGGCTCATTTCAAAGATAATA

TAAGGAGTCAGCACGGAATGTTCCACCCCTACCCGGTACTGGCCAACAGCAGGCAACGTGAATTAATCCTTCGAATAGCC

AAGAAATTCTGGGGGTATGTCTTGCTATATTCTGATGACCCGGCACTAATCAGACAAACAATCAAGAACTTGAAACGGAA

TCATCTAACCTTTGACTTACATAGTAACCCATTTATTAAAGGGCTATCCAAGGCTGAGAAACTGCTGGTGCGGACGAGTT

CACTAAGAAGAGAATGGTTGTTCACTCTCGAAACGAAAGAAGTAAAAGAGTGGTTCAAATTGGTAGGTTACAGTGCACTC

ATCAGAGGCTAGTCAGCTATGCATCTGCCCCCTTCTCCTCCGCCATGAGACCCCACTGGCGATCCAGAAGATTAAAGAAA

ACTACATATTGGATAAGTATCTATTCCCAGCTTTGTCTGGT

>MF443340.1/sheep/China/2014-02-17

ACCAAACAAAGTTGGGTAAGGATAGATCTTATAATAACTATGGACTGGCAAACTTAGGAGTAAAGATCCTACTGTCGGGG

GGAGGAGGAGGAGCAAGATCTTTGATTATGGCGACTCTCCTTAAAAGCTTAGCATTGTTCAAAAGGAACAAAGACAAAGC

GCCGACAGCATCAGGTTCAGGAGGGGCCATCCGGGGGATTAAGAATGTTATCATAGTCCCGATTCCCGGAGACTCGTCCA

TCACTACCCGTTCAAGACTGCTCGACAGGCTTGTCAGATTGGCCGGAGATCCTGACATCAACGGGTCAAAGCTGACCGGG

GTGATGATCAGCATGTTATCCTTGTTCGTAGAGTCACCCGGGCAATTGATACAGCGAATCACAGATGATCCGGATGTTAG

TATCCGCCTTGTTGAGGTAGTTCAAAGTACTAGGTCCCAGTCCGGGTTGACCTTTGCATCACGTGGTGCTGATTTAGACA

ACGAGGCAGACATGTATTTTTCAACTGAGGGGCCCTCGAGTGGAGGTAAGAAAAGGATCAACTGGTTTGAGAACAGAGAA

ATAATAGACATAGAGGTGCAGGATCCAGAAGAGTTCAATATGTTGTTAGCCTCCATACTAGCACAAGTCTGGATCCTTCT

GGCCAAGGCTGTTACGGCACCAGATACGGCAGCTGACTCAGAACTGAGAAGGTGGGTTAAATACACACAACAAAGGAGAG

TGATTGGGGAATTTCGCCTTGACAAGGGGTGGCTGGACGCAGTCCGCAACAGAATTGCAGAAGATCTATCACTCCGGCGG

TTCATGGTATCTCTCATACTTGACATCAAGAGGACCCCAGGCAACAAGCCAAGGATTGCAGAAATGATCTGCGACATTGA

CAACTATATTGTCGAAGCAGGACTCGCCAGTTTCATCCTTACTATCAAATTTGGTATCGAAACCATGTATCCTGCATTAG

GGCTCCACGAGTTTGCCGGAGAATTGTCCACTATTGAGTCCTTGATGAACTTGTATCAACAGCTAGGAGAGGTTGCACCC

TACATGGTAATTCTAGAGAATTCAATTCAGAACAAGTTTAGTGCAGGAGCTTATCCCCTCCTCTGGAGCTATGCGATGGG

TGTCGGAGTCGAGCTGGAGAACTCAATGGGGGGCTTGAACTTTGGCAGGTCATATTTTGACCCTGCCTATTTTCGTCTCG

GACAGGAGATGGTCAGAAGATCTGCAGGAAAGGTCAGCTCTGTGATTGCGGCTGAGCTCGGCATCACAGCAGAGGAAGCT

AAACTAGTCTCGGAAATCGCCTCGCAGGCTGGGGACGAAAGAACCGCCAGAGGGACTGGGCCTCGACAGGCGCAGGTTTC

CTTCCTTCAGCACCAAACAGGAGGGGGAGAGTCGTCCGCACCAGCGACCAGAGAAGGGGTCAAAGCTGCGATCCCAAACG

GAGCTGAAGAAAGGGACAGAAAGCAAACACGCCCAGGAAGGCCCAGAGGAGAGACCTCCGGCCAACTGCTCCTGGACATC

ATGCCAGAGGATGAGATCTTGCGAGAGTCTGGTCAAAACCCTCGTGAGGCTCAAAGATCGGCCGAGGCACTCTTCAGGCT

GCAGGCCATGGCCAAGATTCTGGAGGACCAGGAGGAGGGAGAAGACAACAGTCAGGTCTACAACGACAAGGATCTCCTCG

GCTAAGCAGACGCACTCTCTGTCGAAATCAGTGACAAGACATCTCCTACCAGTATTATAAAAAACTTAGGACCCAGGTCC

AAGCAAGCACACATCGACACTCCAACCAGTCGAGCGGAGACCACCGATGGCAGAAGAACAAGCATACCATGTCAACAAGG

GGCTGGAATGTATCAAGTCTCTCAAAGCCTCACCCCCGGATCTATCCACCATCAGAGATACCATCGAGAGCTGGAGAGAG

GGGCTTAGCCCCTCGGGCCGTGCAACACCGAACCCTGATACGTCCGAGGGAGATCATCAGAATATCAACCAATCATGCTC

ACCAGCAATCGGACCAAACAAAGTCTACTTGTCTCCTGAAGATAATCTCGGATTTAGAGAGATCACTGGCAACGACTGTG

AGGCTGGGCTCGGAGGGGTCCAGAGAGAAGGATCCAACTCTCAAGTACAGCGTTACCATGTTTATAGTCACGGGGGTGAA

GAGATTGAAGGACTCGAGGATGCTGACTCTCTCGTGGTTCAAGCAGATCCTCCAGTTGCTAACGTCTTCAATGGAGGAGA

GGATGGATCTGACGACAGCGATGTGGACTCTGGCCCAGATGATCCCGGCAGAGATACTCTATATGACCGGGGATCTGTTG

CCGGCAATGGTATCGCTAGGTCCACAGATGTCGAAAAACTAGAAGGTGCTGATATTCAAGAAGTTCTTAACTCCCAGAAA

GGCAAAGGAGGAAGATTCCAGGGCGGGAAACCCTTGCGAGTCCCGGAAATACCCGATGTCAAGCACTCCAGACCATCAGC

CCAATCAATTAAAAAGGGCACAGACGGGAACTCAGTCTCATCTGGAACGGTGATAGAGTGTTTATCGATAAGTGGTGCAA

CCCAAACTGTGCCAGAGTCAAGATGGGAGTCATCAGAGCAAAATGCGTCTGTGGGGAGTGTCCTCAAGTCTGCGAGGAGT

GCAAAGACGATCCAGGGGTCGACACAAGAATCTGGTACCATAGCATCACTGACTCAGCCTAAAGAGAATGACTCCGAGTA

TGAGTATGAGGATGACCTATTTACAGAGATTCAGGACATCCGTGCAAGCATTGCCAAGATCCATGATGACAATAAAACTA

TCCTCTCAAAGCTTGATTCTATACTGTTATTGAAAGGGGAAGTCGACACTATCAAGAAACAAATCAGCAAGCAGAATATA

AGTATATCCACCATTGAGGGCCATCTCTCCAGTATAATGATAGCCATCCCTGGCTTTGGGAAGGACATCAAGGACCCAAC

ATCCGAAGTCGAGTTGAACCCAGATCTAAGACCTATAATAAGCCGTGACTCTGGCAGAGCTCTCGCGGAGGTCCTCAAGA

AACCCGCTGTGGATAGGTCTCAGAAAATTGGAACCAAAGCCAACTCCAGCTCAAAGGGTCAGCTTCTTAAGGATCTCCAG

CTAAAACCTGTCGACAAGCAGGCAAGTTCTGCAATCGGGTTTGTCCCGTCCGACCATGAATCATCCAGAAATGTCATCCG

CTCCATAATCAAGTCGAGCAAGCTAAATATTGATCACAAGGACTATCTTCTAGATTTACTAAATGATGTGAAAGGCTCCA

AGGATCTTAAGGAATTTCACAAGATGCTAACAGCAATTCTCGCCAAGCACCCGTAACACATCCTCCAGTCATCATCTCAT

ACTCGACTAAAAACATCCTTTCAATCAGGCTATTACAAAAAACTTAGGAGCAAGGGCAACTGAGCTTCGCAGACAGGATG

ACCGAGATCTACGACTTCGATAAATCAGCATGGGATGTCAAAGGGTCAATTGCCCGCATAGAACCCACCACCTACCACGA

CGGCCGACTGATACCCCAGGTGAGGGTCATCGATCCTGGTCTGGGAGACAGAAAAGATGAGTGCTTTATGTACCTGTTTC

TCCTAGGAGTGATTGAGGATAACGACCCCCTGTCTCCCCCCGTCGGAAGAACCTTTGGCTCTCTACCTCTAGGGGTTGGT

AGGTCAACTGCCAAGCCAGAAGAACTACTAAGGGAGGCCACAGAATTAGATATAGTGGTGAGGCGCACTGCAGGAGTAAA

TGAGAAACTGGTATTTTACAACAACACTCCGCTGTCCTTGTTAACGCCCTGGAAGAAAGTTCTGACAACCGGAAGTGTGT

TTAGCGCTAACCAGGTTTGCAATGCAGTCAACCTAGTCCCGCTTGATACTCCCCAGAGATTCAGGGTTGTGTACATGAGC

ATAACTAGATTATCAGACAATGGTTACTATAGTGTGCCCAGAAGAATGCTGGAGTTCCGCTCGGCCAATGCAGTAGCCTT

CAACATCTTGGTTACACTAAGAATTGAAAATGGCACAAACCCTAGAAGATACATAGTCGGCTCATGGGAGAATTCAGAGG

TCACATTTATGGTACACGTGGGCAACTTTAGGAGAAAGAAGAACGAAGTATACTCTGCTGATTACTGCAAAATGAAGATT

GAAAAGATGGGTTTAGTTTTTGCCCTGGGTGGAATAGGCGGAACAAGTCTCCATATTAGAAGCACCGGGAAAATGAGCAA

AACCCTCCATGCACAGCTGGGGTTCAAGAAAATCTTATGTTACCCCCTAATGGATGTCAATGAGGATCTTAACCGATATC

TCTGGCGGGCAGAGTGCCGAATAGTCAAAATCCAAGCTGTCTTACAGCCATCAGTACCCCAAGAGTTCCGTGTCTACGAT

GATGTCATCATCAACGATGATCAAGGCTTGTTCAAGATCTTGTAGTTCATTTGCAAAGGAACGAACCCACCCCCAGGAAG

GGGCCCCACCCTCCCCCG-CCAAAAAACACACTCCAAGGCTCCCCGGCACAGCGCCAACACAGCTACTCCCGCGATGGAA

CGAGGTCGGCC-CCCGAACAAATCCGGTCCAGGGGGGGGGGCCCCCCCCCTGACCAACCGGGCGCCGACCGAGCAGGCCG

GCGAGAAGCCCCCAAGAGGCCCCGACGAAGACCGAGCACCCCATCCCCCC-AAAAACGTCCCCC-AACACGCCCGAGGCT

AGCCACCGGACACCCCCAGACCCGCCACCGGCGAGACAGAACCCCAAGGGGGGACCCTCCAAACCAAAGGAATCACCGGG

GTAGGCTCCGCCGAACCAGCACAGACCCGACCCAAGCAAGGCGGACCCCCAACACGAAACTGCCCAGCACACCCTACTCA

TCATCAAACAAAACTTAGGGGCCAAGTCCATACGAGAGGCCCTAGTCCGTCAGGCGGAGGGACGCAGAAAGGAAGGAGAC

ACGACCGTCCCAGAAGGAGGACACCCAACCACCGAAACGCCCGCCCCCAGGGAAGGAG--CCA-CCCCCACGCAGCGAAG

GAGCGGGTGGGAGACCTCGGGGAGGGCCCGACCGGGACCCGGGGGCCGGCGGCGCCGGCGGTGAGCAGGCCGGGCGCATA

CCTGACCCCACCGGAGGATCCGAGGAGGGCCCGACCCCGCGTGGACCAAACGGCAGGGAGCCCAACCCCACCAAAAAAAG

AGGCGAGGCCCCCCCTCCCTCCCCCCCCGCACACAACCCGCAGACAGACCGAGGGAGGGCCATCCCCCAGCGAGACAGGC

GGCCCCCTCGCAGCCGAGCCGCAGCTCCCCAGAATAGGACACCCCCGCACACTCCGAACACCCCCCCGCCCCCCGCCACC

GGGACAGGGGCTCCACCACCACAGACCGGAGCCACCGACAGCCAGCCAGAGGTCCCCTCCAGTCCCGCCCGGCGCCCCCC

CGAAGAGGCCCCGGCAAGGCCCCCCAGCCCGACGTCCCCCGCACGCACCGAAAGAAGGACCCGCCCGAGAACGTGCGGCA

CATCCATGCGCAAACATCATGACGCGGGTCGCAATTTTGACATTTCTGTTTCTTTTCCCAAATGTTGTTGCGTGTCAGAT

TCACTGGGGCAATCTATCCAAGATCGGGATTGTAGGAACAGGGAGTGCCAGCTACAAGGTGATGACTAGGCCAAGCCACC

AGACTCTGGTTATAAAGTTAATGCCAAATATAACGGCCATCGACAATTGTACAAAGTCAGAGATTGCAGAGTACAAGAGA

TTGCTGATCACAGTGTTAAAGCCTGTAGAGGATGCTCTGTCGGTGATAACCAAGAATGTAAGACCAATTCAAACTCTAAC

ACCTGGGCGTAGAACCCGCCGTTTTGCTGGAGCTGTTCTGGCCGGGGTAGCACTTGGAGTTGCGACAGCCGCTCAGATAA

CTGCAGGAGTCGCCCTTCATCAATCATTGATGAACTCCCAAGCAATTGAGAGTTTAAAAACCAGTCTTGAGATGTCGAAT

CAGGCAATAGAAGAAATCAGACTTGCAAATAAGGAGACCATACTGGCAGTACAGGGCGTCCAGGATTATATCAACAATGA

GCTCGTCCCTTCTGTTCATAGAATGTCATGCGAGCTGGTAGGTCACAAGCTCGGCCTCAAGCTCCTTAGGTACTACACCG

AGATCCTGTCCATATTCGGGCCCAGTCTTCGAGACCCGATATCTGCCGAAATATCAATCCAGGCACTTAGTTATGCATTA

GGCGGAGACATTAATAAAATCCTGGACAAGCTTGGGTATAGCGGTGGGGATTTCCTTGCCATCCTAGAAAGCAAGGGAAT

AAAGGCCCGGGTTACATATGTGGACACAAGAGATTACTTTATAATCCTTAGCATCGCCTACCCAACCTTATCTGAGATCA

AGGGAGTGATAGTTCACAAGATAGAAGCTATAACATACAACATTGGGGCACAGGAGTGGTATACTACTATCCCTAAATAT

GTAGCCACTCAGGGGTATCTGATATCGAACTTTGATGAGACGTCATGCGTATTCACTCCAGAGGGGACAGTTTGCAGCCA

GAATGCGTTGTACCCAATGAGCCCATTGCTTCAGGAATGTTTCAGGGGGTCAACAAAATCGTGCGCCAGAACCTTAGTTT

CAGGGACCATAAGTAATAGATTTATCCTATCAAAAGGGAACCTGATTGCAAATTGTGCGTCAGTTTTGTGCAAATGTTAC

ACAACAGAGACAGTTATCAGCCAAGATCCTGACAAACTACTAACTGTTGTAGCATCCGACAAGTGTCCTGTAGTTGAGGT

GGATGGAGTGACAATACAGGTCGGCAGTCGAGAGTATCCGGATTCTGTATACTTACACAAAATAGACTTAGGTCCAGCCA

TCTCCCTAGAAAAACTGGATGTAGGCACCAATTTAGGCAATGCAGTCACAAGACTGGAGAATGCAAAGGAGCTCCTAGAT

GCATCAGACCAAATACTGAAGACTGTTAAAGGGGTACCTTTCAGTGGGAATATGTACATAGCACTGGCAGCTTGCATAGG

AGTATCCCTGGGCCTTGTCACATTAATATGCTGCTGTAAGGGGAGGTGTAAGAACAAGGAAATCCCTATCTCCAAAATCA

ACCCAGGGCTCAAACCCGACCTGACCGGGACCTCAAAGTCGTACGTAAGATCACTGTAGTCAGAATCACCTGGATCATCT

GGCATCACACACATACATGCACGACACAGGCAGTCCGAGGACGCAAGAAACCCAGCCTCCGGTCACCCACCCGACCCCAC

TCCACGCTCCACCACACATTAGTCATCAAACAAAACTTAGGACGAAAGGTCAATCACCATGTCCGCACAAAGGGAGAGGA

TCAATGCCTTCTACAAAGACAATCCTCACAATAAGAACCATAGGGTGATCCTGGATAGAGAACGCTTGGTCATTGAAAGA

CCCTACATCTTGCTTGGAGTCCTGCTGGTAATGTTCCTGAGTCTAATCGGACTGCTGGCCATTGCAGGGATCAGGCTTCA

CCGGGCCACCGTTGGAACTTCAGAGATCCAGAGTCGGCTGAATACCAATATTAAGTTGACCGAATCTATTGATCACCAGA

CTAAGGATGTCTTAACTCCCCTTTTTAAAATCATTGGCGATGAAGTCGGCATCAGAATTCCACAGAAATTCAGTGATCTT

GTCAAGTTCATCTCCGATAAGATTAAATTCCTCAACCCTGATAGAGAGTATGATTTCAGAGATCTCCGGTGGTGTATGAA

TCCCCCCGAGAGAGTCAAAATTAATTTTGATCAGTTTTGTGAGTACAAGGCTGCGGTTAAGTCAATTGAACATATATTTG

AGTCACCACTCAACAAGTCAAAAAAGCTGCAATCTTTGACTCTCGGGCCCGGAACAGGCTGTCTAGGCAGGACAGTAACA

AGAGCCCATTTCTCAGAACTTACAATGACCTTAATGGACCTGGATCTAGAGATGAAGCACAACGTGTCCTCAGTGTTTAC

CGTAGTTGAAGAGGGATTATTCGGAAGAACATATACCGTCTGGAGATCCGATGCCAGGGATCCGAGCACCGATCTAGGTA

TCGGCCATTTTTTAAGAGTCTTCGAGATTGGACTGGTAAGAGATCTTGGGCTGGGTCCCCCTGTTTTTCATATGACCAAC

TATCTCACGGTGAACATGAGTGATGACTATCGGAGATGTCTTTTAGCGGTAGGGGAGTTGAAGTTGACAGCCCTATGCAC

CTCATCTGAGACTGTGACACTGAGTGAGAGAGGAGTTCCAAGGAGGGAACCTCTTGTGGTTGTGATACTTAATCTAGCTG

GACCCAATCTAGGGGGAGAGCTATACAGTGTCTTGCCTACCTCTGATCTCATGGTGGAGAAACTCTATTTGTCTTCACAT

AGAGGGATCATCAAAGATGATGAGGCCAATTGGGTAGTGCCGTCTACCGATGTTCGTGATCTTCAGAACAAAGGTGAATG

TCTGGTGGAAGCATGCAAGACTCGACCTCCTTCATTTTGCAATGGCACAGGATCAGGCCCGTGGTCAGAAGGGAGAATCC

CTGCCTACGGGGTGATCAGGGTCAGTCTTGACTTAGCGAGTGACCCGGATGTAGTTATCACTTCAGTGTTTGGCCCACTG

ATACCTCACCCATCCGGCATGGATCTTTACAACAACCCGTTTTCAAAAGCTGTATGGTTGGCTGTACCACCTTATGAGCA

GTCATTTCTAGGAATGATAAATACAATTGGATTCCCTAACAGAGCAGAGGTTATGCCGCACATTTTGACCACAGAGATCA

GAGGCCCTCGGGGTCGTTGCCATGTTCCCATAGAATTGTCCCGCCGGGTTGATGACGATATCAAGATCGGGTCCAACATG

GTCATATTGCCGACGATGGACCTGAGGTATATTACAGCCACTTATGATGTTTCCAGGAGCGAGCATGCAATCGTGTACTA

TATCTATGACACGAGTCGCTCATCATCTTACTTCTACCCAGTTCGACTGAATTTCAAAGGCAATCCTCTCTCTTTGAGGA

TAGAGTGTTTCCCCTGGCGTCATAAGGTGTGGTGCTACCATGATTGTCTTATATACAACACCATAACAGGTGAAGAGGTC

CATACGAGAGGGCTGACCGGCATAGAGGTAACATGCAATCCAGTCTGAGCAGAGCTACGACCGTCACTCAGGCAGTCCCT

TGAGTCGCCACCGAGTCCAAGCAGCACAGCCTGGGACAATCAACGGCACAGCCCAGCCAACAATGTTATAAAAAACTTAG

GAGCCAGGGTTGTAGGGGCCATGGACTCTCTATCAGTCAATCAGGTCTTGTACCCTGAGGTTCATCTAGACAGCCCTATT

GTCACAAACAAACTAGTTGCCATCCTTGAGTACTCGGGTATCGACCATAACTATGTTCTTGAAGACCAGACTCTTGTCAA

GAATATTAGGTATAGGCTGGGGTGCGGTTTTTCAAATCAAATGATCATCAATAACAGGGGGGTGGGTGAAACAGTCAATT

CTAAACTTAAAAGTTACCCCCATAATCGTCATATCATATACCCGGATTGCAATAAGGAGTTGTTTTGTATCAAGGATAGC

TGCATATCTAAGAAGCTCTCGGAGCTATTCAAGAAGGGTAATTCCTTGTACTCTAAGATAAGTCACCAGGTACTGGATTG

TCTTAAGAGAGTCAATGGGAAATTAGGTCTGGGCACAGATCTTACCCATGGTCTGAAGGAGGGTATCCTTGACCTAGGGT

TGCACATGCATAGCTCTCAATGGTTCGAGACTTTCCTGTTCTGGTTCACTATCAAGACAGAGATGAGATCAATGATCAAA

GAACAGTCCCATATATGCCACAAGAGGAGGTATAACCCGACTTTTGTGTCGGGGGATGCATTCGAGGTGCTCGTATCGCG

AGACCTCGTTGTGATAATTGATAAGAATACCCAGTATGTCTTCTACTTAACATTTGAGCTGGTCCTTATGTATTGTGATG

TCATAGAGGGTAGACTTATGACGGAGACAGCCATGGCCATAGACCAGAGATATTCAGAACTTCTAAGCCGGGTCAGATAC

TTGTGGGATCTTATTGATGGGTTCTTCCCAACACTAGGCAACACCACATACCAAATTGTTGCTCTGCTTGAACCATTATC

GTTGGCTTATCTTCAACTTCAGGATGTCACTCTGGAGTTAAGAGGTGCTTTTTTAGACCATTGCTTCAAAGAACTCTATG

AGATACTGGAACATTGCGGCATTGACACAGCAGGCACTTACAATTCCATCACTGAAGGATTGGATTACGTATTCATCACC

CACGATATACATCTAACTGGGGAGATTTTTTCATTTTTCCGGAGTTTCGGACACCCCCGTCTAGAAGCAGTCACCGCTGC

GGAAAATGTCAGAAAACATATGAACCAACCGAAGGTAATCAGTTATGAGACTATGATGAAAGGCCATGCGGTATTTTGCG

GGATAATCATAAATGGTTTTCGGGACCGACATGGCGGCAGCTGGCCCCCTGTTGCACTGCCAGAACATGCCTCTGCTGCG

ATCCGGAATGCGCAGGCATCAGGTGAGGGACTGACCCATGACCTGTGTATAGACAACTGGAAATCCTTTGTTGGATTCAA

ATTTGGCTGCTTTATGCCACTCAGCCTAGATAGTGATTTGACCATGTATCTCAAAGACAAGGCATTGGCTGCACTGAAGA

ATGAGTGGGATTCAGTTTACCCGAAAGAATACCTCCGTTATAACCCACCTAGAGGGACAGAGTCAAGGCGATTGGTAGAG

GTATTCCTGAATGACTCCAGCTTTGATCCTTATAACATGATAATGTACGTGGTGAATGGCTCCTACCTTAAAGACCCTGA

GTTTAACCTCTCATACAGCCTAAAAGAGAAGGAGATAAAGGAGACAGGGCGGTTGTTTGCCAAAATGACCTATAAGATGC

GGGCCTGTCAGGTAATCGCTGAAAATCTGATATCGAATGGTGTTGGGAAGTATTTCCGAGACAATGGGATGGCAAAGGAC

GAGCATGACCTAACAAAAGCCCTTCACACCCTGGCAGTCTCAGGTGTTCCCAAAAACAACAAAGATAACCACCGAGGTGG

GCCTCCCAGAAGGACAACAAACCGAGGGGTGAGATCAAGCCAAGGCACCAAAACACAAGATAGAGACAAGGTTCAAGGGG

GACCTATGTACAACTATTTGCGATGCCAACCGATCAGCCCTGATCAGGGTGAGTCATACGAGACTGTTAGTGCATTCATC

ACCGCTGACCTTAAGAAGTATTGCCTGAATTGGAGATACGAGACAATCAGCATATTTGCACAGAGGCTGAATGAAATATA

TGGACTACCATCCTTCTTTCAATGGTTACACAGGATATTGGAAAAATCCGTACTCTACGTCAGTGACCCACATTGTCCTC

CCGATCTAGATAATCATATCCCTCTGGACAGTGTCCCCAATGCCCAAATATTCATCAAGTACCCAATGGGCGGAATAGAA

GGTTATTGCCAAAAACTATGGACAATCAGCACTATACCATACTTGTATCTGGCAGCCTATGAGAGCGGAGTAAGAATCGC

CTCACTGGTGCAGGGTGACAATCAGACAATCGCAGTGACAAAAAGAGTTCCAAGTTCTTGGCCTTATTCACTAAAAAAGA

GGGAGGCCTCCAAGGCCGCTCAAAATTACTTCGTAGTCTTAAGGCAAAGATTGCACGATGTAGGTCATCACTTAAAGGCT

AATGAGACCATAGTATCTTCTCACTTTTTTGTATATTCCAAAGGGATTTATTATGACGGCCTGCTAGTCTCACAATCACT

AAAGAGCATCGCCAGATGTGTCTTCTGGTCCGAGACTATAGTGGATGAAACCAGAGCGGCGTGCAGTAATATTGCAACAA

CTGTCGCTAAGAGTATAGAGAGGGGTTATGATAGGTACCTTGCATATTCTTTGAATATCCTCAAGATTTTCCAACAGATC

CTTATATCCCTTAACTTCACTATTAACACAACAATGACTCAGGATGTCGTGGCACCGATCATCGAGAACGGTGATTTGCT

GATAAGGATGGCACTCTTGCCAGCACCCATCGGGGGTCTCAATTATCTTAACATGAGCAGGTTATTTGTGAGAAATATCG

GTGACCCGGTCACCTCCTCTATAGCCGACCTGAAGAGGATGATAGATGCCGGGCTAATGCCAGAAGAAACATTGCATCAA

GTGATGACCCAGACCCCGGGAGAATCATCCTACCTAGATTGGGCAAGTGACCCTTATTCTGCCAACCTACCCTGCGTACA

GAGTATCACTCGCCTTCTCAAAAACATCACTGCACGATATATTTTAATCAGCAGCCCAAACCCGATGCTGAAAGGGTTAT

TTCATGAGGGGAGTAGAGATGAAGACGAGGAGCTTGCAAGTTTCCTAATGGATCGGCATATAATTGTTCCAAGAGCTGCA

CATGAAATCTTAGACCATAGTATAACCGGAGCAAGAGAAGCTATAGCCGGGATGTTGGACACCACCAAGGGCCTGATTAG

AACAAGTATGAAACGGGGCGGCCTCACCCCTCGAGTCTTAGCCCGCCTTTCCAATTATGATTATGAACAATTCCGATCCG

GGATAACACTATTAACAAAGAAAGGGCAGTGTTATCTCATTGACAAGGACTCGTGCTCGGTGCAGCTCGCTATAGCTCTG

AGGGGCCACATGTGGGCTAAGTTGGCTCGCGGGAGACCAATCTATGGGTTGGAGGTGCCTGATGTATTGGAATCGATGAA

CGGCTACCTTATCAAACGCCACGAGTCGTGTGCTATCTGTGAAACGGGCTCAAGTCACTACGGATGGTTTTTCGTCCCTG

CAGGGTGCCAGCTTGACGATGTATCTAGAGAGACTTCGGCTCTTCGTGTACCTTATGTCGGATCAACAACTGAGGAAAGG

ACAGACATGAAACTTGCATTTGTTAGGTCTCCAAGTAGATCCCTCAAATCAGCAGTTAGAATTGCAACAGTTTACTCATG

GGCTTACGGGGATGATGAGAAATCATGGAGTGAAGCCTGGATGCTAGCTAGGCAGAGAGCCAATATCACCTTAGATGAAT

TGAGAATGATCACTCCAGTCTCCACATCCACCAACTTAGCCCACCGGTTGAGGGATCGGAGTACCCAGGTGAAATACTCG

GGGACATCCCTTGTGAGGGTTGCAAGATACACAACCATCTCCAATGATAACTTGTCGTTTGTGATATCTGAGAAAAAAGT

GGATACCAACTTCATTTATCAGCAAGGGATGTTGCTTGGTCTTGGGATCCTTGAGAACCTCTTTAGGCTAGAGGCCACCA

CCGGAGTATCCAACACAGTGCTACACCTGCACGTGGAAACAGAATGTTGTGTTGTACCTATGGTGGATCACCCAAGGATA

CCGAGTCTCCGTAATATAAAAGTTACGGATGAGCTATGCACAAACCCTTTGATCTACGACAGGTCCCCCATCATAGAGCA

CGATGCAACCCGGCTATATTCACAGAGCCACAGGAGACATTTGGTGGAGTTTGTTACCTGGTCAACAAGCCAGCTTTACC

ATATACTGGCTAAGTCTACAGCAATGTCCATGATTGAATTGATCACGAGATTCGAGAAAGATCACATGAATGAAATAGCC

GCCCTGATTGGCGATGACGACATCAACAGCTTCATCACAGAATTTTTGCTAGTGGAGCCTAGATTATTTATAGTTTACCT

TGGTCAGTGTGCTGCCATCAATTGGGCCTTTGATATACATTATCATCGGCCCTCGGGCAAGTACCAGATGGGGGAACTTC

TCTACTCTCTGCTCTCTCGGATGAGCAAAGGAGTATATAAGATCTTTACTAATGCTTTGAGCCACCCCAAAGTTTACAAG

AAATTTTGGCGAAGTGGCATCATTGAGCCGATTCATGGCCCATCCTTGGATACACAGAATTTACATGTCACTGTCTGTGA

CATGATATATGGGTCCTATGTCACCTATTTGGATCTTTTGCTGAATGATGAGCTAGATGATTACCCGTATTTGCTCTGCG

AGAGTGATGAGGACGTGGTCACAGACAGGTTCGACAACATTCAAGCCAAACATCTCTGTGTATTGGCCGATGTATATTGC

AGCTCCAAGAGGTGTCCTTCAATAATCGGGATGTCTCCTATAGAAAAATGTACCATTCTCACACATTACATCAAGGGAGA

ATCAGTACAATCCCCGTCTGGGACCTCATGGAACACTGATCCCCTTGTAGTAGATCATTACTCATGCTCTCTGACCTACC

TTCGCCGCGGTTCCATCAAACAAATCAGGTTGAGGGTGGATCCTGGGTTTGTATTCGAGGCGTTGACAGACGTCGACTTT

AAACAACCTCGCAAAGCTAAGTTAGATATATCGGTCGTTGGGTTGACTGATTTCTCTCCCCCTTACGACAACGTCGGTGA

TTTTTTAGGGACTATCAACACATTGAGGCACGATCTGCCTGTCACCGGAACCGGGGTCTCGAACTATGAAGTCCACGCTT

ATCGTAGAATTGGCCTGAATTCATCAGCATGTTATAAAGCCGTAGAGATCTCCACGTTAATCAAGCCATCCTTAGAAGTC

GGAGAGCATGGATTGTTCTTAGGAGAAGGTTCCGGTTCAATGCTGGCTGCGTACAAAGAAGTTCTTAAATTAGCAAATTG

TTACTACAACAGCGGAGTAACAGCGGAGGGCAGAGCCGGACAGAGGGAAATATCTCCCTATCCTTCAGAGATGAGCCTAG

TAGAGAATCAAATGGGGATAGAGCGGAGTGTTAAAGTGCTGTTCAACGGGAAACCTGAAGTAACTTGGGTGGGGACCACA

GATTGCTACAAGTACATAATCAGTAACATCCAAACCTCTAGTCTGGGTTTCATACATTCAGATATTGAGACGCTCCCAAC

CAAAGATGCTGTTGAGAAATTGGAGGAATTTGCCTCTATCCTATCCTTATCCTTGATTTTGGGGAAAATCGGCTCTATTA

CAGTTGTCAAAATTATGCCCATTAGCGGGGATTTTACCCAAGGCTTCATAGCCTATGCCATCCAATATTTTAGGGAGAGC

CTGCTTGCCTATCCGAGATATAGTAACTTCATCTCGACTGAGTGTTACCTTATTATGATAGGATTAAAGGCCAATCGATT

GATAAACCCAGAAGCCATTAAGCAAAGCATAATCAGAGCGGGGGTCAGGACTGCACCAGGACTTGTGAGCCATGTATTAT

CAGAGAAACAAAAAGGTTGTATTCAATCTTTCTTGGGTGATCCTTATATCCAAGGAGACTTCAATAAGCACCTTAAATCT

CTAACCTCTATTGAGAAAATCCTGGTAAATTGTGGTCTCTCGATCAATGGCATAAAAATCTGTAGAGATCTAATCCACCA

TGATATCGCCTCCGGTCCAGACGGCCTTATGAGCTCTACAATTATTTTATATAGGGAATTGGCTCATTTCAAAGATAATA

TAAGGAGTCAGCACGGAATGTTCCACCCCTACCCGGTACTGGCCAACAGCAGGCAACGTGAATTAATCCTTCGAATAGCC

AAGAAATTCTGGGGGTATGTCTTGCTATATTCTGATGACCCGGCACTAATCAGACAAACAATCAAGAACTTGAAACGGAA

TCATCTAACCTTTGACTTACATAGTAACCCATTTATTAAAGGGCTATCCAAGGCTGAGAAACTGCTGGTGCGGACGAGTT

CACTAAGAAGAGAATGGTTGTTCACTCTCGAAACGAAAGAAGTAAAAGAGTGGTTCAAATTGGTAGGTTACAGTGCACTC

ATCAGAGGCTAGTCAGCTATGCATCTGCCCCCTTCTCCTCCGCCATGAGACCCCACTGGCGATCCAGAAGATTAAAGAAA

ACTACATATTGGATAAGTATCTATTCCCAGCTTTGTCTGGT

>MF443352.1/goat/China/2014-05-15

ACCAAACAAAGTTGGGTAAGGATAGATCTTATAATAACTATAGACTGGCAAACTTAGGAGTAAAGATCCTACTGTCGGGG

GGAGGAGGAGGAGCAAGATCCTTGATCATGGCGACTCTCCTTAAAAGCTTAGCATTGTTCAAAAGGAACAAAGACAAAGC

GCCGACAGCATCAGGTTCAGGAGGGGCCATCCGGGGGATTAAGAATGTTATCATAGTCCCGATTCCCGGAGACTCGTCCA

TCACTACCCGTTCAAGACTGCTCGACAGGCTTGTCAGATTGGCCGGAGATCCTGACATCAACGGGTCAAAGCTGACCGGG

GTGATGATCAGCATGTTATCCTTGTTCGTAGAGTCACCCGGGCAATTGATACAGCGAATCACAGATGATCCGGATGTTAG

TATCCGCCTTGTTGAGGTAGTTCAAAGTACTAGGTCCCAGTCCGGGTTGACCTTTGCATCACGTGGTGCTGATTTAGACA

ACGAGGCAGACATGTATTTTTCAACTGAGGGGCCCTCGAGTGGAGGTACGAAAAGGATCAACTGGTTTGAGAACAGAGAA

ATAATAGACATAGAGGTGCAGGATCCAGAAGAGTTCAATATGTTGTTAGCCTCCATACTAGCACAAGTCTGGATCCTTCT

GGCCAAGGCTGTTACGGCACCAGATACGGCAGCTGACTCAGAACTGAGAAGGTGGGTTAAATACACACAACAAAGGAGAG

TGATTGGGGAATTTCGCCTTGACAAGGGGTGGCTGGACGCAGTCCGCAACAGAATTGCAGAAGATCTATCACTCCGGCGG

TTCATGGTATCTCTCATACTTGACATCAAGAGGACCCCAGGCAACAAGCCAAGGATTGCAGAAATGATCTGCGACATTGA

CAACTATATTGTCGAAGCAGGACTCGCCAGTTTCATCCTTACTATCAAATTTGGTATCGAAACCATGTATCCTGCATTAG

GGCTCCACGAGTTTGCCGGAGAATTGTCCACTATTGAGTCCTTGATGAACTTGTATCAACAGCTAGGAGAGGTTGCACCC

TACATGGTAATTCTAGAGAATTCAATTCAGAACAAGTTTAGTGCAGGAGCTTATCCCCTCCTCTGGAGCTATGCGATGGG

TGTCGGAGTCGAGCTGGAGAACTCAATGGGGGGCTTGAACTTTGGCAGGTCATATTTTGACCCTGCCTATTTTCGTCTCG

GACAGGAGATGGTCAGAAGATCTGCAGGAAAGGTCAGCTCTGTGATTGCGGCTGAGCTCGGCATCACAGCAGAGGAAGCT

AAACTAGTCTCGGAAATCGCCTCGCAGGCTGGGGACGAAAGAACCGCCAGAGGGACTGGGCCTCGACAGGCGCAGGTTTC

CTTCCTTCAGCACCAAACAGGAGGGGGAGAGTCGTCCGCACCAGCGACCAGAGAAGGGGTCAAAGCTGCGATCCCAAACG

GAGCTGAAGAAAGGGACAGAAAGCAAACACGCCCAGGAAGGCCCAGAGGAGAGACCTCCGGCCAACTGCTCCTGGACATC

ATGCCAGAGGATGAGATCTTGCGAGAGTCTGGTCAAAACCCTCGTGAGGCTCAAAGATCGGCCGAGGCACTCTTCAGGCT

GCAGGCCATGGCCAAGATTCTGGAGGACCAGGAGGAGGGAGAAGACAACAGTCAGGTCTACAACGACAAGGATCTCCTCG

GCTAAGCAGACGCACCCTCTGTCGAAATCAGTGACAAGACATCTCCTACCAGTATTATAAAAAACTTAGGACCCAGGTCC

AAGCAAGCACACATCGACACTCCAACCAGTCGAGCGGAGACCACCGATGGCAGAAGAACAAGCATACCATGTCAACAAGG

GGCTGGAATGTATCAAGTCTCTCAAAGCCTCACCCCCGGATCTATCCACCATCAGAGATACCATCGAGAGCTGGAGAGAG

GGGCTTAGCCCCTCGGGCCGTGCAACACCGAACCCTGATACGTCCGAGGGAGATCATCAGAATATCAACCAATCATGCTC

ACCAGCAATCGGACCAAACAAAGTCTACTTGTCTCCTGAAGATAATCTCGGATTTAGAGAGATCACTGGCAACGACTGTG

AGGCTGGGCTCGGAGGGGTCCAGAGAGAAGGATCCAACTCTCACGTACAGCGTTACCATGTTTATAGTCACGGGGGTGAA

GAGATTGAAGGACTCGAGGATGCTGACTCTCTCGTGGTTCAAGCAGATCCTCCAGTTGCTAACGTCTTCAATGGAGGAGA

GGATGGATCTGACGACAGCGATGTGGACTCTGGCCCAGATGATCCCGGCAGAGATACTCTATATGACCGGGGATCTGTTG

CCGGCAATGGTATCGCTAGGTCCACAGATGTCGAAAAACTAGAAGGTGCTGATATTCAAGAAGTTCTTAACTCCCAGAAA

GGCAAAGGAGGAAGATTCCAGGGCGGGAAAACCTTGCGAGTCCCGGAAATACCCGATGTCAAGCACTCCAGACCATCAGC

CCAATCAATTAAAAAGGGCACAGACGGGAACTCAGTCTCATCTGGAACGGTGATAGAGTGTTTATCGATAAGTGGTGCAA

CCCAAACTGTGCCAGAGTCAAGATGGGAGTCATCAGAGCAAAATGCGTCTGTGGGGAGTGTCCTCAAGTCTGCGAGGAGT

GCAAAGACGATCCAGGGGTCGACACAAGAATCTGGTACCATAGCATCACTGACTCAGCCTAAAGAGAATGACTCCGAGTA

TGAGTATGAGGATGACCTATTTACAGAGATTCAGGACATCCGTGCAAGCATTGCCAAGATCCATGATGACAATAAAACTA

TCCTCTCAAAGCTTGATTCTATACTGTTATTGAAAGGGGAAGTCGACACTATCAAGAAACAAATCAGCAAGCAGAATATA

AGTATATCCACCATTGAGGGCCATCTCTCCAGTATAATGATAGCCATCCCTGGCTTTGGGAAGGACATCAAGGACCCAAC

ATCCGAAGTCGAGTTGAACCCAGATCTAAGACCTATAATAAGCCGTGACTCTGGCAGAGCTCTCGCGGAGGTCCTCAAGA

AACCCGCTGTGGATAGGTCTCAGAAAATTGGAACCAAAGCCAACTCCAGCTCAAAGGGTCAGCTTCTTAAGGATCTCCAG

CTAAAACCTGTCGACAAGCAGGCAAGCTCTGCAATCGGGTTTGTCCCGTCCGACCATGAATCATCCAGAAATGTCATCCG

CTCCATAATCAAGTCGAGCAAGCTAAATATTGATCACAAGGACTATCTTCTAGATTTACTAAATGATGTGAAAGGCTCCA

AGGATCTTAAGGAATTTCACAAGATGCTAACAGCAATTCTCGCCAAGCACCCGTAACACATCCTCCAGTCATCATCTCAT

ACTCGACTAAAAACATCCTTTCAATCAGGCTATTACAAAAAACTTAGGAGCAAGGGCAACTGAGCTTCGCAGACAGGATG

ACCGAGATCTACGACTTCGATAAATCAGCATGGGATGTCAAAGGGTCAATTGCCCGCATAGAACCCACCACCTACCACGA

CGGCCGACTGATACCCCAGGTGAGGGTCATCGATCCTGGTCTGGGAGACAGAAAAGATGAGTGCTTTATGTACCTGTTTC

TCCTAGGAGTGATTGAGGATAACGACCCCCTGTCTCCCCCCGTCGGAAGAACCTTTGGCTCTCTACCTCTAGGGGTTGGT

AGGTCAACTGCCAAGCCAGAAGAACTACTAAGGGAGGCCACAGAATTAGATATAGTGGTGAGGCGCACTGCAGGATTAAA

TGAGAAACTGGTATTTTACAACAACACTCCGCTGTCCTTGTTAACGCCCTGGAAGAAAGTTCTGACAACCGGAAGTGTGT

TTAGCGCTAACCAGGTTTGCAATGCAGTCAACCTAGTCCCGCTTGATACTCCCCAGAGATTCAGGGTTGTGTACATGAGC

ATAACTAGATTATCAGACAATGGTTACTATAGTGTGCCCAGAAGAATGCTGGAGTTCCGATCGGCCAATGCAGTAGCCTT

CAACATCTTGGTTACACTAAGAATTGAAAATGGCACAAACCCTAGAAGATACATAGTCGGCTCATGGGAGAATTCAGAGG

TCACATTTATGGTACACGTGGGCAACTTTAGGAGAAAGAAGAACGAAGTATACTCTGCTGATTACTGCAAAATGAAGATT

GAAAAGATGGGTTTAGTTTTTGCCCTGGGTGGAATAGGCGGAACAAGTCTCCATATTAGAAGCACCGGGAAAATGAGCAA

AACCCTCCATGCACAGCTGGGGTTCAAGAAAATCTTATGTTACCCCCTAATGGATGTCAATGAGGATCTTAACCGATATC

TCTGGCGGGCAGAGTGCCGAATAGTCAAAATCCAAGCTGTCTTACAGCCATCAGTACCCCAAGAGTTCCGTGTCTACGAT

GATGTCATCATCAACGATGATCAAGGCTTGTTCAAGATCTTGTAGTTCATTTGCAAAGGAACGAACCCACCCCCAGGAAG

GGGCCCCACCCTCCCCCG-CCAAAAAACACACTCCAAGGCTCCCCGGCACAGCGCCAACACAGCTACTCCCGCGATGGAA

CGAGGTCGGCC-CCCGAACAAATCCGGTCCAGGGGGGGGGGCCCCCCCCCTGACCAACCGGGCGCCGACCGAGCAGGCCG

GCGAGAAGCCCCCAAGAGGCCCCGACGAAGACCGAGCACCCCATCCCCCC-AAAAACGTCCCCC-AACACGCCCGAGGCT

AGCCACCGGACACCCCCAGACCCGCCACCGGCGAGACAGAACCCCAAGGGGGGACCCTCCAAACCAAAGGAATCACCGGG

GTAGGCTCCGCCGAACCAGCACAGACCCGACCCAAGCAAGGCGGACCCCCAACACGAAACTGCCCAGCACACCCTACTCA

TCATCAAACAAAACTTAGGGGCCAAGTCCATACGAGAGGCCCTAGTCCGTCAGGCGGAGGGACGCAGAAAGGAAGGAGAC

ACGACCGTCCCAGAAGGAGGACACCCAACCACCGAAACGCCCGCCCCCAGGGAAGGAG--CCA-CCCCCACGCAGCGAAG

GAGCGGGTGGGAGACCTCGGGGAGGGCCCGACCGGGACCCGGGGGCCGGCGGCGCCGGCGGTGAGCAGGCCGGGCGCATA

CCTGACCCCACCGGAGGATCCGAGGAGGGCCCGACCCCGCGTGGACCAAACGGCAGGGAGCCCAACCCCACCAAAAAGAG

AGGCGAGGCCCCCCCTCCCTCCCCCCCCGCACACAACCCGCAGACAGACCGAGGGAGGGCCATCCCCCAGCGAGACAGGC

GGCTCCCTCGCAGCCGAGCCGCAGCTCCCCAGAATAGGACACCCCCGCACACTCCGAACACCCCCTCGCCCCCCGCCACC

GGGACAGGGGCTCCACCACCACAGACCGGAGCCACCGACAGCCAGCCAGAGGTCCCCTCCAGTCCCGCCCGGCGCCCCCC

CGAAGAGGCCCCGGCAAGGCCCCCCAGCCCGACGTCCCCCGCACGCACCGAAAGAAGGACCCGCCCGAGAACGTGCGGCA

CATCCATGCGCAAACATCATGACGCGGGTCGCAATTTTGACATTTCTGTTTCTTTTCCCAAATGTTGTTGCGTGTCAGAT

TCACTGGGGCAATCTATCCAAGATCGGGATTGTAGGAACAGGGAGTGCCAGCTACAAGGTGATGACTAGGCCAAGCCACC

AGACTCTGGTTATAAAGTTAATGCCAAATATAACGGCCATCGACAATTGTACAAAGTCAGAGATTGCAGAGTACAAGAGA

TTGCTGATCACAGTGTTAAAGCCTGTAGAGGATGCTCTGTCGGTGATAACCAAGAATGTAAGACCAATTCAAACTCTAAC

ACCTGGGCGTAGAACCCGCCGTTTTGCTGGAGCTGTTCTGGCCGGGGTAGCACTTGGAGTTGCGACAGCCGCTCAGATAA

CTGCAGGAGTCGCCCTTCATCAATCATTGATGAACTCCCAAGCAATTGAGAGTTTAAAAACCAGTCTTGAGATGTCGAAT

CAGGCAATAGAAGAAATCAGACTTGCAAATAAGGAGACCATACTGGCAGTACAGGGCGTCCAGGATTATATCAACAATGA

GCTCGTCCCTTCTGTTCATAGAATGTCATGCGAGCTGGTAGGTCACAAGCTCGGCCTCAAGCTCCTTAGGTACTACACCG

AGATCCTGTCCATATTCGGGCCCAGTCTTCGAGACCCGATATCTGCCGAAATATCAATCCAGGCACTTAGTTATGCATTA

GGCGGAGACATTAATAAAATCCTGGACAAGCTTGGGTATAGCGGTGGGGATTTCCTTGCCATCCTAGAAAGCAAGGGAAT

AAAGGCCCGGGTTACATATGTGGACACAAGAGATTACTTTATAATCCTTAGCATCGCCTACCCAACCTTATCTGAGATCA

AGGGAGTGATAGTTCACAAGATAGAAGCTATAACATACAACATTGGGGCACAGGAGTGGTATACTACTATCCCTAAATAT

GTAGCCACTCAGGGGTATCTGATATCGAACTTTGATGAGACGTCATGCGTATTCACTCCAGAGGGGACAGTTTGCAGCCA

GAATGCGTTGTACCCAATGAGCCCATTGCTTCAGGAATGTTTCAGGGGGTCAACAAAATCGTGCGCCAGAACCTTAGTTT

CAGGGACCATAAGTAATAGATTTATCCTATCAAAAGGGAACCTGATTGCAAATTGTGCGTCAGTTTTGTGCAAATGTTAC

ACAACAGAGACAGTTATCAGCCAAGATCCTGACAAACTACTAACTGTTGTAGCATCCGACAAGTGTCCTGTAGTTGAGGT

GGATGGAGTGACAATACAGGTCGGCAGTCGAGAGTATCCGGATTCTGTATACTTACACAAAATAGACTTAGGTCCAGCCA

TCTCCCTAGAAAAACTGGATGTAGGCACCAATTTAGGCAATGCAGTCACAAGACTGGAGAATGCAAAGGAGCTCCTAGAT

GCATCAGACCAAATACTGAAGACTGTTAAAGGGGTACCTTTCAGTGGGAATATGTACATAGCACTGGCAGCTTGCATAGG

AGTATCCCTGGGCCTTGTCACATTAATATGCTGCTGTAAGGGGAGGTGTAAGAACAAGGAAATCCCTATCTCCAAAATCA

ACCCAGGGCTCAAACCCGACCTGACCGGGACCTCAAAGTCGTACGTAAGATCACTGTAGTCAGAATCACCTGGATCATCT

GGCATCACACACATACATGCACGACACAGGCAGTCCGAGGACGCAAGAAACCCAGCCTCCGGTCACCCACCCGACCCCAC

TCCATGCTCCACCACACATTAGTCATCAAACAAAACTTAGGACGAAAGGTCAATCACCATGTCCGCACAAAGGGAGAGGA

TCAATGCCTTCTACAAAGACAATCCTCACAATAAGAACCATAGGGTGATCCTGGATAGAGAACGCTTGGTCATTGAAAGA

CCCTACATCTTGCTTGGAGTCCTGCTGGTAATGTTCCTGAGTCTAATCGGACTGCTGGCCATTGCAGGGATCAGGCTTCA

CCGGGCCACCGTTGGAACTTCAGAGATCCAGAGTCGGCTGAATACCAATATTAAGTTGACCGAATCTATTGATCACCAGA

CTAAGGATGTCTTAACTCCCCTTTTTAAAATCATTGGCGATGAAGTCGGCATCAGAATTCCACAGAAATTCAGTGATCTT

GTCAAGTTCATCTCCGATAAGATTAAATTCCTCAACCCTGATAGAGAGTATGATTTCAGAGATCTCCGGTGGTGTATGAA

TCCCCCCGAGAGAGTCAAAATTAATTTTGATCAGTTTTGTGAGTACAAGGCTGCGGTTAAGTCAATTGAACATATATTTG

AGTCACCACTCAACAAGTCAAAAAAGCTGCAATCTTTGACTCTCGGGCCCGGAACAGGCTGTCTAGGCAGGACAGTAACA

AGAGCCCATTTCTCAGAACTTACAATGACCTTAATGGACCTGGATCTAGAGATGAAGCACAACGTGTCCTCAGTGTTTAC

CGTAGTTGAAGAGGGATTATTCGGAAGAACATATACCGTCTGGAGATCCGATGCCAGGGATCCGAGCACCGATCTAGGTA

TCGGCCATTTTTTAAGAGTCTTCGAGATTGGACTGGTAAGAGATCTTGGGCTGGGTCCCCCTGTTTTTCATATGACCAAC

TATCTCACGGTGAACATGAGTGATGACTATCGGAGATGTCTTTTAGCGGTAGGGGAGTTGAAGTTGACAGCCCTATGCAC

CTCATCTGAGACTGTGACACTGAGTGAGAGAGGAGTTCCAAGGAGGGAACCTCTTGTGGTTGTGATACTTAATCTAGCTG

GACCCACTCTAGGGGGAGAGCTATACAGTGTCTTGCCTACCTCTGATCTCATGGTGGAGAAACTCTATTTGTCTTCACAT

AGAGGGATCATCAAAGATGATGAGGCCAATTGGGTAGTGCCGTCTACCGATGTTCGTGATCTTCAGAATAAAGGTGAATG

TCTGGTGGAAGCATGCAAGACTCGACCTCCTTCATTTTGCAATGGCACAGGATCAGGCCCGTGGTCAGAAGGGAGAATCC

CTGCCTACGGGGTGATCAGGGTCAGTCTTGACTTAGCGAGTGACCCGGATGTAGTTATCACTTCAGTGTTTGGCCCACTG

ATACCTCACCTATCCGGCATGGATCTTTACAACAACCCGTTTTCAAAAGCTGTATGGTTGGCTGTACCACCTTATGAGCA

GTCATTTCTAGGAATGATAAATACAATTGGATTCCCTAACAGAGCAGAGGTTATGCCGCACATTTTGACCACAGAGATCA

GAGGCCCTCGGGGTCGTTGCCATGTTCCCATAGAATTGTCCCGCCGGGTTGATGACGATATCAAGATCGGGTCCAACATG

GTCATATTGCCGACGATGGACCTGAGGTATATTACAGCCACTTATGATGTTTCCAGGAGCGAGCATGCAATCGTGTACTA

CATCTATGACACGAGTCGCTCATCATCTTACTTCTACCCAGTTCGACTGAATTTCAAAGGCAATCCTCTCTCTTTGAGGA

TAGAGTGTTTCCCCTGGCGTCATAAGGTGTGGTGCTACCATGATTGTCTTATATACAACACCATAACAGGTGAAGAGGTC

CATACGAGAGGGCTGACCGGCATAGAGGTAACATGCAATCCAGTCTGAGCAGAGCTACGACCGTCACTCAGGCAGTCCCT

TGAGTCGCCACCGAGTCCAAGCAGCACAGCCTGGGACAATCAACGGCACAGCCCAGCCAACAATGTTATAAAAAACTTAG

GAGCCAGGGTTGTAGGGGCCATGGACTCTCTATCAGTCAATCAGGTCTTGTACCCTGAGGTTCATCTAGACAGCCCTATT

GTCACAAACAAACTAGTTGCCATCCTTGAGTACTCGGGTATCGACCATAACTATGTTCTTGAAGACCAGACTCTTGTCAA

GAATATTAGGTATAGGCTGGGGTGCGGTTTTTCAAATCAAATGATCATCAATAACAGGGGGGTGGGTGAAACAGTCAATT

CTAAACTTAAAAGTTACCCCCATAATCGTCATATCATATACCCGGATTGCAATAAGGAGTTGTTTTGTATCAAGGATAGC

TGCATATCTAAGAAGCTCTCGGAGCTATTCAAGAAGGGTAATTCCTTGTACTCTAAGATAAGTCACCAGGTACTGGATTG

TCTTAAGAGAGTCAATGGGAAATTAGGTCTGGGCACAGATCTTACCCATGGTCTGAAGGAGGGTATCCTTGACCTAGGGT

TGCACATGCATAGCTCTCAATGGTTCGAGACTTTCCTGTTCTGGTTCACTATCAAGACAGAGATGAGATCAATGATCAAA

GAACAGTCCCATATATGCCACAAGAGGAGGTATAACCCGACTTTTGTGTCGGGGGATGCATTCGAGGTGCTCGTATCGCG

AGACCTCGTTGTGATAATTGATAAGAATACCCAGTATGTCTTCTACTTAACATTTGAGCTGGTCCTTATGTATTGTGATG

TCATAGAGGGTAGACTTATGACGGAGACAGCCATGGCCATAGACCAGAGATATTCAGAACTTCTAAGCCGGGTCAGATAC

TTGTGGGATCTTATTGATGGGTTCTTCCCAACACTAGGCAACACCACATACCAAATTGTTGCTCTGCTTGAACCATTATC

GTTGGCTTATCTTCAACTTCAGGATGTCACTCTGGAGTTAAGAGGTGCTTTTTTAGACCATTGCTTCAAAGAACTCTATG

AGATACTGGAACATTGCGGCATTGACACAGCAGGCACTTACAATTCCATCACTGAAGGATTGGATTACGTATTCATTACC

CACGATATACATCTAACTGGGGAGATTTTTTCATTTTTCCGGAGTTTCGGACACCCCCGTCTAGAAGCAGTCACCGCTGC

GGAAAATGTCAGAAAACATATGAACCAACCGAAGGTAATCAGTTATGAGACTATGATGAAAGGCCATGCGGTATTTTGCG

GGATAATCATAAATGGTTTTCGGGACCGACATGGCGGCAGCTGGCCCCCTGTTGCACTGCCAGAACATGCCTCTGCTGCG

ATCCGGAATGCGCAGGCATCAGGTGAGGGACTGACCCATGACCTGTGTATAGACAACTGGAAATCCTTTGTTGGATTCAA

ATTTGGCTGCTTTATGCCACTCAGCCTAGATAGTGATTTGACCATGTATCTCAAAGACAAGGCATTGGCTGCACTGAAGA

ATGAGTGGGATTCAGTTTACCCGAAAGAATACCTCCGTTATAACCCACCTAGAGGGACAGAGTCAAGGCGATTGGTAGAG

GTATTCCTGAATGACTCCAGCTTTGATCCTTATAACATGATAATGTACGTGGTGAATGGCTCCTACCTTAAAGACCCTGA

GTTTAACCTCTCATACAGCCTAAAAGAGAAGGAGATAAAGGAGACAGGGCGGTTGTTTGCCAAAATGACCTATAAGATGC

GGGCCTGTCAGGTAATCGCTGAAAATCTGATATCGAATGGTGTTGGGAAGTATTTCCGAGACAATGGGATGGCAAAGGAC

GAGCATGACCTAACAAAAGCCCTTCACACCCTGGCAGTCTCAGGTGTTCCCAAAAACAACAAAGATAACCACCGAGGTGG

GCCTCCCAGAAGGACAACAAACCGAGGGGTGAGATCAAGCCAAGGCACCAAAACACAAGATAGAGACAAGGTTCAAGGGG

GACCTATGTACAACTATTTGCGATGCCAACCGATCAGCCCTGATCAGGGTGAGTCATACGAGACTGTTAGTGCATTCATC

ACCGCTGACCTTAAGAAGTATTGCCTGAATTGGAGATACGAGACAATCAGCATATTTGCACAGAGGCTGAATGAAATATA

TGGACTACCATCCTTCTTTCAATGGTTACACAGGATATTGGAAAAATCCGTACTCTACGTCAGTGACCCACATTGTCCTC

CCGATCTAGATAATCATATCCCTCTGGACAGTGTCCCCAATGCCCAAATATTCATCAAGTACCCAATGGGCGGAATAGAA

GGTTATTGCCAAAAACTATGGACAATCAGCACTATACCATACTTGTATCTGGCAGCCTATGAGAGCGGAGTAAGAATCGC

CTCACTGGTGCAGGGTGACAATCAGACAATCGCAGTGACAAAAAGAGTTCCAAGTTCTTGGCCTTATTCACTAAAAAAGA

GGGAGGCCTCCAAGGCCGCTCAAAATTACTTCGTAGTCTTAAGGCAAAGATTGCACGATGTAGGTCATCACTTAAAGGCT

AATGAGACCATAGTATCTTCTCACTTTTTTGTATATTCCAAAGGGATTTATTATGACGGCCTGCTAGTCTCACAATCACT

AAAGAGCATCGCCAGATGTGTCTTCTGGTCTGAGACTATAGTGGATGAAACCAGAGCGGCGTGCAGTAATATTGCAACAA

CTGTCGCTAAGAGTATAGAGAGGGGTTATGATAGGTACCTTGCATATTCTTTGAATATCCTCAAGATTTTCCAACAGATC

CTTATATCCCTTAACTTCACTATTAACACAACAATGACTCAGGATGTCGTGGCACCGATCATCGAGAACGGTGATTTGCT

GATAAGGATGGCACTCTTGCCAGCACCCATCGGGGGTCTCAATTATCTTAACATGAGCAGGTTATTTGTGAGAAATATCG

GTGACCCGGTCACCTCCTCTATAGCCGACCTGAAGAGGATGATAGATGCCGGGCTAATGCCAGAAGAAACATTGCATCAA

GTGATGACCCAGACCCCGGGAGAATCATCCTACCTAGATTGGGCAAGTGACCCTTATTCTGCCAACCTACCCTGCGTACA

GAGTATCACTCGCCTTCTCAAAAACATCACTGCACGATATATTTTAATCAGCAGCCCAAACCCGATGCTGAAAGGGTTAT

TTCATGAGGGGAGTAGAGATGAAGACGAGGAGCTTGCAAGTTTCCTAATGGATCGGCATATAATTGTTCCAAGAGCTGCA

CATGAAATCTTAGACCATAGTATAACCGGAGCAAGAGAAGCTATAGCCGGGATGTTGGACACCACCAAGGGTCTGATTAG

AACAAGTATGAAACGGGGCGGCCTCACCCCTCGAGTCTTAGCCCGCCTTTCCAATTATGATTATGAACAATTCCGATCCG

GGATAACACTATTAACAAAGAAAGGGCAGTGTTATCTCATTGACAAGGACTCGTGCTCGGTGCAGCTCGCTATAGCTCTG

AGGGGCCACATGTGGGCTAAGTTGGCTCGCGGGAGACCAATCTATGGGTTGGAGGTGCCTGATGTATTAGAATCGATGAA

CGGCTACCTTATCAAACGCCACGAGTCGTGTGCTATCTGTGAAACGGGCTCAAGTCACTACGGATGGTTTTTCGTCCCTG

CAGGGTGCCAGCTTGACGATGTATCTAGAGAGACTTCGGCTCTTCGTGTACCTTATGTCGGATCAACAACTGAGGAAAGG

ACAGACATGAAACTTGCATTTGTTAGGTCTCCAAGTAGATCCCTCAAATCAGCAGTTAGAATTGCAACAGTTTACTCATG

GGCTTACGGGGATGATGAGAAATCATGGAGTGAAGCCTGGATGCTAGCTAGGCAGAGAGCCAATATCACCTTAGATGAAT

TGAGAATGATCACTCCAGTCTCCACATCCACCAACTTAGCCCACCGGTTGAGGGATCGGAGTACCCAGGTGAAATACTCG

GGGACATCCCTTGTGAGGGTTGCAAGATACACAACCATCTCCAATGATAACTTGTCGTTTGTGATATCTGAGAAAAAAGT

GGATACCAACTTCATTTATCAGCAAGGGATGTTGCTTGGTCTTGGGATCCTTGAGAACCTCTTTAGGCTAGAGGCCACCA

CCGGAGTATCCAACACAGTGCTACACCTGCACGTGGAAACAGAATGTTGTGTTGTACCTATGGTGGATCACCCAAGGATA

CCGAGTCTCCGTAATATAAAAGTTACGGATGAGCTATGCACAAACCCTTTGATCTACGACAGGTCCCCCATCATAGAGCA

CGATGCAACCCGGCTATATTCACAGAGCCACAGGAGACATTTGGTGGAGTTTGTTACCTGGTCAACAAGCCAGCTTTACC

ATATACTGGCTAAGTCTACAGCAATGTCCATGATTGAATTGATCACGAGATTCGAGAAAGATCACATGAATGAAATAGCC

GCCCTGATTGGCGATGACGACATCAACAGCTTCATCACAGAATTTTTGCTAGTGGAGCCTAGATTATTTATAGTTTACCT

TGGTCAGTGTGCTGCCATCAATTGGGCCTTTGATATACATTATCATCGGCCCTCGGGCAAGTACCAGATGGGGGAACTTC

TCTACTCTCTGCTCTCTCGGATGAGCAAAGGAGTATATAAGATCTTTACTAATGCTTTGAGCCACCCCAAAGTTTACAAG

AAATTTTGGCGAAGTGGCATCATTGAGCCGATTCATGGCCCATCCTTGGATACACAGAATTTACATGTCACTGTCTGTGA

CATGATATATGGGTCCTATGTCACCTATTTGGATCTTTTGCTGAATGATGAGCTAGATGATTACCCGTATTTGCTCTGCG

AGAGTGATGAGGACGTGGTCACAGACAGGTTCGACAACATTCAAGCCAAACATCTCTGTGTATTGGCCGATGTATATTGC

AGCTCCAAGAGGTGTCCTTCAATAATCGGGATGTCTCCTATAGAAAAATGTACCATTCTCACACATTACATCAAGGGAGA

ATCAGTACAATCCCCGTCTGGGACCTCATGGAACACTGATCCCCTTGTAGTAGATCATTACTCATGCTCTCTGACCTACC

TTCGCCGCGGTTCCATCAAACAAATCAGGTTGAGGGTGGATCCTGGGTTTGTATTCGAGGCGTTGACAGACGTCGACTTT

AAACAACCTCGCAAAGCTAAGTTAGATATATCGGTCGTTGGGTTGACTGATTTCTCTCCCCCTTACGACAACGTCGGTGA

TTTTTTAGGGACTATCAACACATTGAGGCACGATCTGCCTGTCACCGGAACCGGGGTCTCGAACTATGAAGTCCACGCTT

ATCGTAGAATTGGCCTGAATTCATCAGCATGTTATAAAGCCGTAGAGATCTCCACGTTAATCAAGCCATCCTTAGAAGTC

GGAGAGCATGGATTGTTCTTAGGAGAAGGTTCCGGTTCAATGCTGGCTGCGTACAAAGAAGTTCTTAAATTAGCAAATTG

TTACTACAACAGCGGAGTAACAGCGGAGGGCAGAGCCGGACAGAGGGAAATATCTCCCTATCCTTCAGAGATGAGCCTAG

TAGAGAATCAAATGGGGATAGAGCGGAGTGTTAAAGTGCTGTTCAACGGGAAACCTGAAGTAACTTGGGTGGGGACCACA

GATTGCTACAAGTACATAATCAGTAACATCCAAACCTCTAGTCTGGGTTTCATACATTCAGATATTGAGACGCTCCCAAC

CAAAGATGCTGTTGAGAAATTGGAGGAATTTGCCTCTATCCTATCCTTATCCTTGATTTTGGGGAAAATCGGCTCTATTA

CAGTTGTCAAAATTATGCCCATTAGCGGGGATTTTACCCAAGGCTTCATAGCCTATGCCATCCAATATTTTAGGGAGAGC

CTGCTTGCCTATCCGAGATATAGTAACTTCATCTCGACTGAGTGTTACCTTATTATGATAGGATTAAAGGCCAATCGATT

GATAAACCCAGAAGCCATTAAGCAAAGCATAATCAGAGCGGGGGTCAGGACTGCACCAGGACTTGTGAGCCATGTATTAT

CAGAGAAACAAAAAGGTTGTATTCAATCTTTCTTGGGTGATCCTTATATCCAAGGAGACTTCAATAAGCACCTTAAATCT

CTAACCTCTATTGAGAAAATCCTGGTAAATTGTGGTCTCTCGATCAATGGCATAAAAATCTGTAGAGATCTAATCCACCA

TGATATCGCCTCCGGTCCAGACGGCCTTATGAGCTCTACAATTATTTTATATAGGGAATTGGCTCATTTCAAAGATAATA

TAAGGAGTCAGCACGGAATGTTCCACCCCTACCCGGTACTGGCCAACAGCAGGCAACGTGAATTAATCCTTCGAATAGCC

AAGAAATTCTGGGGGTATGTCTTGCTATATTCTGATGACCCGGCACTAATCAGACAAACAATCAAGAACTTGAAACGGAA

TCATCTAACCTTTGACTTACATAGTAACCCATTTATTAAAGGGCTATCCAAGGCTGAGAAACTGCTGGTGCGGACGAGTT

CACTAAGAAGAGAATGGTTGTTCACTCTCGAAACGAAAGAAGTAAAAGAGTGGTTCAAATTGGTAGGTTACAGTGCACTC

ATCAGAGGCTAGTCAGCTATGCATCTGCCCCCTTCTCCTCCGCCATGAGACCCCACTGGCGATCCAGAAGATTAAAGAAA

ACTACATATTGGATAAGTATCTATTCCCAGCTTTGTCTGGT

>KM816619.1/goat/China/2014-04-09

ACCAAACAAAGTTGGGTAAGGATCGATCTTATAATAACTATGGACTGGCAAACTTAGGAGTAAAGATCCTACTGTCGGGG

GGAGGAGGAGGAGCAAGATCCTTGATCATGGCGACTCTCCTTAAAAGCTTAGCATTGTTCAAAAGGAACAAAGACAAAGC

GCCGACAGCATCAGGTTCAGGAGGGGCCATCCGGGGGATTAAGAATGTTATCATAGTCCCGATTCCCGGAGACTCGTCCA

TCACTACCCGTTCAAGACTGCTCGACAGGCTTGTCAGATTGGCCGGAGATCCTGACATCAACGGGTCAAAGCTGACCGGG

GTGATGATCAGCATGTTATCCTTGTTCGTAGAGTCACCCGGGCAATTGATACAGCGAATCACAGATGATCCGGATGTTAG

TATCCGCCTTGTTGAGGTAGTTCAAAGTACTAGGTCCCAGTCCGGGTTGACCTTTGCATCACGTGGTGCTGATTTAGACA

ACGAGGCAGACATGTATTTTTCAACTGAGGGGCCCTCGAGTGGAGGTAAGAAAAGGATCAACTGGTTTGAGAACAGAGAA

ATAATAGACATAGAGGTGCAGGATCCAGAAGAGTTCAATATGTTGTTAGCCTCCATACTAGCACAAGTCTGGATCCTTCT

GGCCAAGGCTGTTACGGCACCAGATACGGCAGCTGACTCAGAACTGAGAAGGTGGGTTAAATACACACAACAAAGGAGAG

TGATTGGGGAATTTCGCCTTGACAAGGGGTGGCTGGACGCAGTCCGCAACAGAATTGCAGAAGATCTATCACTCCGGCGG

TTCATGGTATCTCTCATACTTGACATCAAGAGGACCCCAGGCAACAAGCCAAGGATTGCAGAAATGATCTGCGACATTGA

CAACTATATTGTCGAAGCAGGACTCGCCAGTTTCATCCTTACTATCAAATTTGGTATCGAAACCATGTATCCTGCATTAG

GGCTCCACGAGTTTGCCGGAGAATTGTCCACTATTGAGTCCTTGATGAACTTGTATCAACAGCTAGGAGAGGTTGCACCC

TACATGGTAATTCTAGAGAATTCAATTCAGAACAAGTTTAGTGCAGGAGCTTATCCCCTCCTCTGGAGCTATGCGATGGG

TGTCGGAGTCGAGCTGGAGAACTCAATGGGGGGCTTGAACTTTGGCAGGTCATATTTTGACCCTGCCTATTTTCGTCTCG

GACAGGAGATGGTCAGAAGATCTGCAGGAAAGGTCAGCTCTGTGATTGCGGCTGAGCTCGGCATCACAGCAGAGGAAGCT

AAACTAGTCTCGGAAATCGCCTCGCAGGCTGGGGACGAAAGAACCGCCAGAGGGACTGGGCCTCGACAGGCGCAGGTTTC

CTTCCTTCAGCACCAAACAGGAGGGGGAGAGTCGTCCGCACCAGCGACCAGAGAAGGGGTCAAAGCTGCGATCCCAAACG

GAGCTGAAGAAAGGGACAGAAAGCAAACACGCCCAGGAAGGCCCAGAGGAGAGACCTCCGGCCAACTGCTCCTGGACATC

ATGCCAGAGGATGAGATCTTGCGAGAGTCTGGTCAAAACCCTCGTGAGGCTCAAAGATCGGCCGAGGCACTCTTCAGGCT

GCAGGCCATGGCCAAGATTCTGGAGGACCAGGAGGAGGGAGAAGACAACAGTCAGGTCTACAACGACAAGGATCTCCTCG

GCTAAGCAGACGCACCCTCTGTCGAAATCAGTGACAAGACATCTCCTACCAGTATTATAAAAAACTTAGGACCCAGGTCC

AAGCAAGCACACATCGACACTCCAACCAGTCGAGCGGAGACCACCGATGGCAGAAGAACAAGCATACCATGTCAACAAGG

GGCTGGAATGTATCAAGTCTCTCAAAGCCTCACCCCCGGATCTATCCACCATCAGAGATACCATCGAGAGCTGGAGAGAG

GGGCTTAGCCCCTCGGGCCGTGCAACACCGAACCCTGATACGTCCGAGGGAGATCATCAGAATATCAACCAATCATGCTC

ACCAGCAATCGGACCAAACAAAGTCTACTTGTCTCCTGAAGATAATCTCGGATTTAGAGAGATCACTGGCAACGACTGTG

AGGCTGGGCTCGGAGGGGTCCAGAGAGAAGGATCCAACTCTCAAGTACAGCGTTACCATGTTTATAGTCACGGGGGTGAA

GAGATTGAAGGACTCGAGGATGCTGACTCTCTCGTGGTTCAAGCAGATCCTCCAGTTGCTAACGTCTTCAATGGAGGAGA

GGATGGATCTGACGACAGCGATGTGGACTCTGGCCCAGATGATCCCGGCAGAGATACTCTATATGACCGGGGATCTGTTG

CCGGCAATGGTCTCGCTAGGTCCACAGATGTCGAAAAACTAGAAGGTGCTGATATTCAAGAAGTTCTTAACTCCCAGAAA

GGCAAAGGAGGAAGATTCCAGGGCGGGAAAACCTTGCGAGTCCCGGAAATACCCGATGTCAAGCACTCCAGACCATCAGC

CCAATCAATTAAAAAGGGCACAGACGGGAACTCAGTCTCATCTGGAACGGTGATAGAGTGTTTATCGATAAGTGGTGCAA

CCCAAACTGTGCCAGAGTCAAGATGGGAGTCATCAGAGCAAAATGCGTCTGTGGGGAGTGTCCTCAAGTCTGCGAGGAGT

GCAAAGACGATCCAGGGGTCGACACAAGAATCTGGTACCATAGCATCACTGACTCAGCCTAAAGAGAATGACTCCGAGTA

TGAGTATGAGGATGACCTATTTACAGAGATTCAGGACATCCGTGCAAGCATTGCCAAGATCCATGATGACAATAAAACTA

TCCTCTCAAAGCTTGATTCTATACTGTTATTGAAAGGGGAAGTCGACACTATCAAGAAACAAATCAGTAAGCAGAATATA

AGTATATCCACCATTGAGGGCCATCTCTCCAGTATAATGATAGCCATCCCTGGCTTTGGGAAGGACATCAAGGACCCAAC

ATCCGAAGTCGAGTTGAACCCAGATCTAAGACCTATAATAAGCCGTGACTCTGGCAGAGCTCTCGCGGAGGTCCTCAAGA

AACCCGCTGTGGATAGGTCTCAGAAAATTGGAACCAAAGCCAACTCCAGCTCAAAGGGTCAGCTTCTTAAGGATCTCCAG

CTAAAACCTGTCGACAAGCAGGCAAGCTCTGCAATCGGGTTTGTCCCGTCCGACCATGAATCATCCAGAAATGTCATCCG

CTCCATAATCAAGTCGAGCAAGCTAAATATTGATCACAAGGACTATCTTCTAGATTTACTAAATGATGTGAAAGGCTCCA

AGGATCTTAAGGAATTTCACAAGATGCTAACAGCAATTCTCGCCAAGCACCCGTAACACATCCTCCAGTCATCATCTCAT

ACTCGACTAAAAACATCCTTTCAATCAGGCTATTACAAAAAACTTAGGAGCAAGGGCAACTGAGCTTCGCAGACAGGATG

ACCGAGATCTACGACTTCGATAAATCAGCATGGGATGTCAAAGGGTCAATTGCCCGCATAGAACCCACCACCTACCACGA

CGGCCGACTGATACCCCAGGTGAGGGTCATCGATCCTGGTCTGGGAGACAGAAAAGATGAGTGCTTTATGTACCTGTTTC

TCCTAGGAGTGATTGAGGATAACGACCCCCTGTCTCCCCCCGTCGGAAGAACCTTTGGCTCTCTACCTCTAGGGGTTGGT

AGGTCAACTGCCAAGCCAGAAGAACTACTAAGGGAGGCCACAGAATTAGATATAGTGGTGAGGCGCACTGCAGGAGTAAA

TGAGAAACTGGTATTTTACAACAACACTCCGCTGTCCTTGTTAACGCCCTGGAAGAAAGTTCTGACAACCGGAAGTGTGT

TTAGCGCTAACCAGGTTTGCAATGCAGTCAACCTAGTCCCGCTTGATACTCCCCAGAGATTCAGGGTTGTGTACATGAGC

ATAACTAGATTATCAGACAATGGTTACTACAGTGTGCCCAGAAGAATGCTGGAGTTCCGCTCGGCCAATGCAGTAGCCTT

CAACATCTTGGTTACACTAAGAATTGAAAATGGCACAAACCCTAGAAGATACATAGTCGGCTCATGGGAGAATTCAGAGG

TCACATTTATGGTACACGTGGGCAACTTTAGGAGAAAGAAGAACGAAGTATACTCTGCTGATTACTGCAAAATGAAGATT

GAAAAGATGGGTTTAGTTTTTGCCCTGGGTGGAATAGGCGGAACAAGTCTCCATATTAGAAGCACCGGGAAAATGAGCAA

AACCCTCCATGCACAGCTGGGGTTCAAGAAAATCTTATGTTACCCCCTAATGGATGTCAATGAGGATCTTAACCGATATC

TCTGGCGGGCAGAGTGCCGAATAGTCAAAATCCAAGCTGTCTTACAGCCATCAGTACCCCAAGAGTTCCGTGTCTACGAT

GATGTCATCATCAACGATGATCAAGGCTTGTTCAAGATCTTGTAGTTCATTTGCAAAGGAACGAACCCACCCCCAGGAAG

GGGCCCCACCCTCCCCCG-CCAAAAAACACACTCCAAGGCTCCCCGGCACAGCGCCAACACAGCTACTCCCGCGATGGAA

CGAGGTCGGCC-CCCGAACAAATCCGGTCCAGGGGGGGGGGCCCCCCCCCTGACCAACCGGGCGCCGACCGAGCAGGCCG

GCGAGAAGCCCCCAAGAGGCCCCGACGAAGACCGAGCACCCCATCCCCCC-AAAAACGTCCCCC-AACACGCCCGAGGCT

AGCCACCGGACACCCCCAGACCCGCCACCGGCGAGACAGAACCCCAAGGGGGGACCCTCCAAACCAAAGGAATCACCGGG

GTAGGCTCCGCCGAACCAGCACAGACCCGACCCAAGCAAGGCGGACCCCCAACACGAAACTGCCCAGCACACCCTACTCA

TCATCAAACAAAACTTAGGGGCCAAGTCCATACGAGAGGCCCTAGTCCGTCAGGCGGAGGGACGCAGAAAGGAAGGAGAC

ACGACCGTCCCAGAAGGAGGACACCCAACCACCGAAACGCCCGCCCCCAGGGAAGGAG--CCA-CCCCCACGCAGCGAAG

GAGCGGGTGGGAGACCTCGGGGAGGGCCCGACCGGGACCCGGGGGCCGGCGGCGCCGGCGGTGAGCAGGCCGGGCGCATA

CCTGACCCCACCGGAGGATCCGAGGAGGGCCCGACCCCGCGTGGACCAAACGGCAGGGAGCCCAACCCCACCAAAAAGAG

AGGCGAGGCCCCCCCTCCCTCCCCCCCCGCACACAACCCGCAGACAGACCGAGGGAGGGCCATCCCCCAGCGAGACAGGC

GGCTCCCTCGCAGCCGAGCCGCAGCTCCCCAGAATAGGACACCCCCGCACACTCCGAACACCCCCTCGCCCCCCGCCACC

GGGACAGGGGCTCCACCACCACAGACCGGAGCCACCGACAGCCAGCCAGAGGTCCCCTCCAGTCCCGCCCGGCGCCCCCC

CGAAGAGGCCCCGGCAAGGCCCCCCAGCCCGACGTCCCCCGCACGCACCGAAAGAAGGACCCGCCCGAGAACGTGCGGCA

CATCCATGCGCAAACATCATGACGCGGGTCGCAATTTTGACATTTCTGTTTCTTTTCCCAAATGTTGTTGCGTGTCAGAT

TCACTGGGGCAATCTATCCAAGATCGGGATTGTAGGAACAGGGAGTGCCAGCTACAAGGTGATGACTAGGCCAAGCCACC

AGACTCTGGTTATAAAGTTAATGCCAAATATAACGGCCATCGACAATTGTACAAAGTCAGAGATTGCAGAGTACAAGAGA

TTGCTGATCACAGTGTTAAAGCCTGTAGAGGATGCTCTGTCGGTGATAACCAAGAATGTAAGACCAATTCAAACTCTAAC

ACCTGGGCGTAGAACCCGCCGTTTTGCTGGAGCTGTTCTGGCCGGGGTAGCACTTGGAGTTGCGACAGCCGCTCAGATAA

CTGCAGGAGTCGCCCTTCATCAATCATTGATGAACTCCCAAGCAATTGAGAGTTTAAAAACCAGTCTTGAGATGTCGAAT

CAGGCAATAGAAGAAATCAGACTTGCAAATAAGGAGACCATACTGGCAGTACAGGGCGTCCAGGATTATATCAACAATGA

GCTCGTCCCTTCTGTTCATAGAATGTCATGCGAGCTGGTAGGTCACAAGCTCGGCCTCAAGCTCCTTAGGTACTACACCG

AGATCCTGTCCATATTCGGGCCCAGTCTTCGAGACCCGATATCTGCCGAAATATCAATCCAGGCACTTAGTTATGCATTA

GGCGGAGACATTAATAAAATCCTGGACAAGCTTGGGTATAGCGGTGGGGATTTCCTTGCCATCCTAGAAAGCAAGGGAAT

AAAGGCCCGGGTTACATATGTGGACACAAGAGATTACTTTATAATCCTTAGCATCGCCTACCCAACCTTATCTGAGATCA

AGGGAGTGATAGTTCACAAGATAGAAGCTATAACATACAACATTGGGGCACAGGAGTGGTATACTACTATCCCTAAATAT

GTAGCCACTCAGGGGTATCTGATATCGAACTTTGATGAGACGTCATGCGTATTCACTCCAGAGGGGACAGTTTGCAGCCA

GAATGCGTTGTACCCAATGAGCCCATTGCTTCAGGAATGTTTCAGGGGGTCAACAAAATCGTGCGCCAGAACCTTAGTTT

CAGGGACCATAAGTAATAGATTTATCCTATCAAAAGGGAACCTGATTGCAAATTGTGCGTCAGTTTTGTGCAAATGTTAC

ACAACAGAGACAGTTATCAGCCAAGATCCTGACAAACTACTAACTGTTGTAGCATCCGACAAGTGTCCTGTAGTTGAGGT

GGATGGAGTGACAATACAGGTCGGCAGTCGAGAGTATCCGGATTCTGTATACTTACACAAAATAGACTTAGGTCCAGCCA

TCTCCCTAGAAAAACTGGATGTAGGCACCAATTTAGGCAATGCAGTCACAAGACTGGAGAATGCAAAGGAGCTCCTAGAT

GCATCAGACCAAATACTGAAGACTGTTAAAGGGGTACCTTTCAGTGGGAATATGTACATAGCACTGGCAGCTTGCATAGG

AGTATCCCTGGGCCTTGTCACATTAATATGCTGCTGTAAGGGGAGGTGTAAGAACAAGGAAATCCCTATCTCCAAAATCA

ACCCAGGGCTCAAACCCGACCTGACCGGGACCTCAAAGTCGTACGTAAGATCACTGTAGTCAGAATCACCTGGATCATCT

GGCATCACACACATACATGCACGACACAGGCAGTCCGAGGACGCAAGAAACCCAGCCTCCGGTCACCCACCCGACCCCAC

TCCACGCTCCACCACACATTAGTCATCAAACAAAACTTAGGACGAAAGGTCAATCACCATGTCCGCACAAAGGGAGAGGA

TCAATGCCTTCTACAAAGACAATCCTCACAATAAGAACCATAGGGTGATCCTGGATAGAGAACGCTTGGTCATTGAAAGA

CCCTACATCTTGCTTGGAGTCCTGCTGGTAATGTTCCTGAGTCTAATCGGACTGCTGGCCATTGCAGGGATCAGGCTTCA

CCGGGCCACCGTTGGAACTTCAGAGATCCAGAGTCGGCTGAATACCAATATTAAGTTGACCGAATCTATTGATCACCAGA

CTAAGGATGTCTTAACTCCCCTTTTTAAAATCATTGGCGATGAAGTCGGCATCAGAATTCCACAGAAATTCAGTGATCTT

GTCAAGTTCATCTCCGATAAGATTAAATTCCTCAACCCTGATAGAGAGTATGATTTCAGAGATCTCCGGTGGTGTATGAA

TCCCCCCGAGAGAGTCAAAATTAATTTTGATCAGTTTTGTGAGTACAAGGCTGCGGTTAAGTCAATTGAACATATATTTG

AGTCACCACTCAACAAGTCAAAAAAGCTGCAATCTTTGACTCTCGGGCCCGGAACAGGCTGTCTAGGCAGGACAGTAACA

AGAGCCCATTTCTCAGAACTTACAATGACCTTAATGGACCTGGATCTAGAGATGAAGCACAACGTGTCCTCAGTGTTTAC

CGTAGTTGAAGAGGGATTATTCGGAAGAACATATACCGTCTGGAGATCCGATGCCAGGGATCCGAGCACCGATCTAGGTA

TCGGCCATTTTTTAAGAGTCTTCGAGATTGGACTGGTAAGAGATCTTGGGCTGGGTCCCCCTGTTTTTCATATGACCAAC

TATCTCACGGTGAACATGAGTGATGACTATCGGAGATGTCTTTTAGCGGTAGGGGAGTTGAAGTTGACAGCCCTATGCAC

CTCATCTGAGACTGTGACACTGAGTGAGAGAGGAGTTCCAAGGAGGGAACCTCTTGTGGTTGTGATACTTAATCTAGCTG

GACCCACTCTAGGGGGAGAGCTATACAGTGTCTCGCCTACCTCTGATCTCATGGTGGAGAAACTCTATCTGTCTTCACAT

AGAGGGATCATCAAAGATGATGAGGCCAATTGGGTAGTGCCGTCTACCGATGTTCGTGATCTTCAGAATAAAGGTGAATG

TCTGGTGGAAGCATGCAAGACTCGACCTCCTTCATTTTGCAATGGCACAGGATCAGGCCCGTGGTCAGAAGGGAGAATCC

CTGCCTACGGGGTGATCAGGGTCAGTCTTGACTTAGCGAGTGACCCGGATGTAGTTATCACTTCAGTGTTTGGCCCACTG

ATACCTCACCTATCCGGCATGGATCTTTACAACAACCCGTTTTCAAAAGCTGTATGGTTGGCTGTACCACCTTATGAGCA

GTCATTTCTAGGAATGATAAATACAATTGGATTCCCTAACAGAGCAGAGGTTATGCCGCACATTTTGACCACAGAGATCA

GAGGCCCTCGGGGTCGTTGCCATGTTCCCATAGAATTGTCCCGCCGGGTTGATGACGATATCAAGATCGGGTCCAACATG

GTCATATTGCCGACGATGGACCTGAGGTATATTACAGCCACTTATGATGTTTCCAGGAGCGAGCATGCAATCGTGTACTA

TATCTATGACACGAGTCGCTCATCATCTTACTTCTACCCAGTTCGACTGAATTTCAAAGGCAATCCTCTCTCTTTGAGGA

TAGAGTGTTTCCCCTGGCGTCATAAGGTGTGGTGCTACCATGATTGTCTTATATACAACACCATAACAGGTGAAGAGGTC

CATACGAGAGGGCTGACCGGCATAGAGGTAACATGCAATCCAGTCTGAGCAGAGCTACGACCGTCACTCAGGCAGTCCCT

TGAGTCGCCACCGAGTCCAAGCAGCACAGCCTGGGACAATCAACGGCACAGCCCAGCCAACAATGTTATAAAAAACTTAG

GAGCCAGGGTTGTAGGGGCCATGGACTCTCTATCAGTCAATCAGGTCTTGTACCCTGAGGTTCATCTAGACAGCCCTATT

GTCACAAACAAACTAGTTGCCATCCTTGAGTACTCGGGTATCGACCATAACTATGTTCTTGAAGACCAGACTCTTGTCAA

GAATATTAGGTATAGGCTGGGGTGCGGTTTTTCAAATCAAATGATCATCAATAACAGGGGGTTGGGTGAAACAGTCAATT

CTAAACTTAAAAGTTACCCCCATAATCGTCATATCATATACCCGGATTGCAATAAGGAGTTGTTCTGTATCAAGGATAGC

TGCATATCTAAGAAGCTCTCGGAGCTATTCAAGAAGGGTAATTCCTTGTACTCTAAGATAAGTCACCAGGTACTGGATTG

TCTTAAGAGAGTCAATGGGAAATTAGGTCTGGGCACAGATCTTACCCATGGTCTGAAGGAGGGTATCCTTGACCTAGGGT

TGCACATGCATAGCTCTCAATGGTTCGAGACTTTCCTGTTCTGGTTCACTATCAAGACAGAGATGAGATCAATGATCAAA

GAACAGTCCCATATATGCCACAAGAGGAGGTATAACCCGACTTTTGTGTCGGGGGATGCATTCGAGGTGCTCGTATCGCG

AGACCTCGTTGTGATAATTGATAAGAATACCCAGTATGTCTTCTACTTAACATTTGAGCTGGTCCTTATGTATTGTGATG

TCATAGAGGGTAGACTTATGACGGAGACAGCCATGGCCATAGACCAGAGATATTCAGAACTTCTAAGCCGGGTCAGATAC

TTGTGGGATCTTATTGATGGGTTCTTCCCAACACTAGGCAACACCACATACCAAATTGTTGCTCTGCTTGAACCATTATC

GTTGGCTTATCTTCAACTTCAGGATGTCACTCTGGAGTTAAGAGGTGCTTTTTTAGACCATTGCTTCAAAGAACTCTATG

AGATACTGGAACATTGCGGCATTGACACAGCAGGCACTTACAATTCCATCACTGAAGGATTGGATTACGTATTCATCACC

CACGATATACATCTAACTGGGGAGATTTTTTCATTTTTCCGGAGTTTCGGACACCCCCGTCTAGAAGCAGTCACCGCTGC

GGAAAATGTCAGAAAACATATGAACCAACCGAAGGTAATCAGTTATGAGACTATGATGAAAGGCCATGCGGTATTTTGCG

GGATAATCATAAATGGTTTTCGGGACCGACATGGCGGCAGCTGGCCCCCTGTTGCACTGCCAGAACATGCCTCTGCTGCG

ATCCGGAATGCGCAGGCATCAGGTGAGGGACTGACCCATGACCTGTGTATAGACAACTGGAAATCCTTTGTTGGATTCAA

ATTTGGCTGCTTTATGCCACTCAGCCTAGATAGTGATTTGACCATGTATCTCAAAGACAAGGCATTGGCTGCACTGAAGA

ATGAGTGGGATTCAGTTTACCCGAAAGAATACCTCCGTTATAACCCACCTAGAGGGACAGAGTCAAGGCGATTGGTAGAG

GTATTCCTGAATGACTCCAGCTTTGATCCTTATAACATGATAATGTACGTGGTGAATGGCTCCTACCTTAAAGACCCTGA

GTTTAACCTCTCATACAGCCTAAAAGAGAAGGAGATAAAGGAGACAGGGCGGTTGTTTGCCAAAATGACCTATAAGATGC

GGGCCTGTCAGGTAATCGCTGAAAATCTGATATCGAATGGTGTTGGGAAGTATTTCCGAGACAATGGGATGGCAAAGGAC

GAGCATGACCTAACAAAAGCCCTTCACACCCTGGCAGTCTCAGGTGTTCCCAAAAACAACAAAGATAACCACCGAGGTGG

GCCTCCCAGAAGGACAACAAACCGAGGGGTGAGATCAAGCCAAGGCACCAAAACACAAGATAGAGACAAGGTTCAAGGGG

GACCTATGTACAACTATTTGCGATGCCAACCGATCAGCCCTGATCAGGGTGAGTCATACGAGACTGTTAGTGCATTCATC

ACCGCTGACCTTAAGAAGTATTGCCTGAATTGGAGATACGAGACAATCAGCATATTTGCACAGAGGCTGAATGAAATATA

TGGACTACCATCCTTCTTTCAATGGTTACACAGGATATTGGAAAAATCCGTACTCTACGTCAGTGACCCACATTGTCCTC

CCGATCTAGATAATCATATCCCTCTGGACAGTGTCCCCAATGCCCAAATATTCATCAAGTACCCAATGGGCGGAATAGAA

GGTTATTGCCAAAAACTATGGACAATCAGCACTATACCATACTTGTATCTGGCAGCCTATGAGAGCGGAGTAAGAATCGC

CTCACTGGTGCAGGGTGACAATCAGACAATCGCAGTGACAAAAAGAGTTCCAAGTTCTTGGCCTTACTCACTAAAAAAGA

GGGAGGCCTCCAAGGCCGCTCAAAATTACTTCATAGTCTTAAGGCAAAGATTGCACGATGTAGGTCATCACTTAAAGGCT

AATGAGACCATAGTATCTTCTCACTTTTTTGTATATTCCAAAGGGATTTATTATGACGGCCTGCTAGTCTCACAATCACT

AAAGAGCATCGCCAGATGTGTCTTCTGGTCCGAGACTATAGTGGATGAAACCAGAGCGGCGTGCAGTAATATTGCAACAA

CTGTCGCTAAGAGTATAGAGAGGGGTTATGATAGGTACCTTGCATATTCTTTGAATATCCTCAAGATTTTCCAACAGATC

CTTATATCCCTTAACTTCACTATTAACACAACAATGACTCAGGATGTCGTGGCACCGATCATCGAGAACGGTGATTTGCT

GATAAGGATGGCACTCCTGCCAGCACCCATCGGGGGTCTCAATTATCTTAACATGAGCAGGTTATTTGTGAGAAATATCG

GTGACCCGGTCACCTCCTCTATAGCCGACCTGAAGAGGATGATAGATGCCGGGCTAATGCCAGAAGAAACATTGCATCAA

GTGATGACCCAGACCCCGGGAGAATCATCCTACCTAGATTGGGCAAGTGACCCTTATTCTGCCAACCTACCCTGCGTACA

GAGTATCACTCGCCTTCTCAAAAACATCACTGCACGATATATTTTAATCAGCAGCCCAAACCCGATGCTGAAAGGGTTAT

TTCATGAGGGGAGTAGAGATGAAGACGAGGAGCTTGCAAGTTTCCTAATGGATCGGCATATAATTGTTCCAAGAGCTGCA

CATGAAATCTTAGACCATAGTATAACCGGAGCAAGAGAAGCTATAGCCGGGATGTTGGACACCACCAAGGGTCTGATTAG

AACAAGTATGAAACGGGGCGGCCTCACCCCTCGAGTCTTAGCCCGCCTTTCCAATTATGATTATGAACAATTCCGATCCG

GGATAACACTATTAACAAAGAAAGGGCAGTGTTATCTCATTGACAAGGACTCGTGCTCGGTGCAGCTCGCTATAGCTCTG

AGGGGCCACATGTGGGCTAAGTTGGCTCGCGGGAGACCAATCTATGGGTTGGAGGTGCCTGATGTATTAGAATCGATGAA

CGGCTACCTTATCAAACGCCACGAGTCGTGTGCTATCTGTGAAACGGGCTCAAGTCACTACGGATGGTTTTTCGTCCCTG

CAGGGTGCCAGCTTGACGATGTATCTAGAGAGACTTCGGCTCTTCGTGTACCTTATGTCGGATCAACAACTGAGGAAAGG

ACAGACATGAAACTTGCATTTGTTAGGTCTCCAAGTAGATCCCTCAAATCAGCAGTTAGAATTGCAACAGTTTACTCATG

GGCTTACGGGGATGATGAGAAATCATGGAGTGAAGCCTGGATGCTAGCTAGGCAGAGAGCCAATATCACCTTAGATGAAT

TGAGAATGATCACTCCAGTCTCCACATCCACCAACTTAGCCCACCGGTTGAGGGATCGGAGTACCCAGGTGAAATACTCG

GGGACATCCCTTGTGAGGGTTGCAAGATACACAACCATCTCCAATGATAACTTGTCGTTTGTGATATCTGAGAAAAAAGT

GGATACCAACTTCATTTATCAGCAAGGGATGTTGCTTGGTCTTGGGATCCTTGAGAACCTCTTTAGGCTAGAGGCCACCA

CCGGAGTATCCAACACAGTGCTACACCTGCACGTGGAAACAGAATGTTGTGTTGTACCTATGGTGGATCACCCAAGGATA

CCGAGTCTCCGTAATATAAAAGTTACGGATGAGCTATGCACAAACCCTTTGATCTACGACAGGTCCCCCATCATAGAGCA

CGATGCAACCCGGCTATATTCACAGAGCCACAGGAGACATTTGGTGGAGTTTGTTACCTGGTCAACAAGCCAGCTTTACC

ATATACTGGCTAAGTCTACAGCAATGTCCATGATTGAATTGATCACGAGATTCGAGAAAGATCACATGAATGAAATAGCC

GCCCTGATTGGCGATGACGACATCAACAGCTTCATCACAGAATTTTTGCTAGTGGAGCCTAGATTATTTATAGTTTACCT

TGGTCAGTGTGCTGCCATCAATTGGGCCTTTGATATACATTATCATCGGCCCTCGGGCAAGTACCAGATGGGGGAACTTC

TCTACTCTCTGCTCTCTCGGATGAGCAAAGGAGTATATAAGATCTTTACTAATGCTTTGAGCCACCCCAAAGTTTACAAG

AAATTTTGGCGAAGTGGCATCATTGAGCCGATTCATGGCCCATCCTTGGATACACAGAATTTACATGTCACTGTCTGTGA

CATGATATATGGGTCCTATGTCACCTATTTGGATCTTTTGCTGAATGATGAGCTAGATGATTACCCGTATTTGCTCTGCG

AGAGTGATGAGGACGTGGTCACAGACAGGTTCGACAACATTCAAGCCAAACATCTCTGTGTATTGGCCGATGTATATTGC

AGCTCCAAGAGGTGTCCTTCAATAATCGGGATGTCTCCTATAGAAAAATGTACCATTCTCACACATTACATCAAGGGAGA

ATCAGTACAATCCCCGTCTGGGACCTCATGGAACACTGATCCCCTTGTAGTAGATCATTACTCATGCTCTCTGACCTACC

TTCGCCGCGGTTCCATCAAACAAATCAGGTTGAGGGTGGATCCTGGGTTTGTATTCGAGGCGTTGACAGACGTCGACTTT

AAACAACCTCGCAAAGCTAAGTTAGATATATCGGTCGTTGGGTTGACTGATTTCTCTCCCCCTTACGACAACGTCGGTGA

TTTTTTAGGGACTATCAACACATTGAGGCACGATCTGCCTGTCACCGGAACCGGGGTCTCGAACTATGAAGTCCACGCTT

ATCGTAGAATTGGCCTGAATTCATCAGCATGTTATAAAGCCGTAGAGATCTCCACGTTAATCAAGCCATCCTTAGAAGTC

GGAGAGCATGGATTGTTCTTAGGAGAAGGTTCCGGTTCAATGCTGGCTGCGTACAAAGAAGTTCTTAAATTAGCAAATTG

TTACTACAACAGCGGAGTAACAGCGGAGGGCAGAGCCGGACAGAGGGAAATATCTCCCTATCCTTCAGAGATGAGCCTAG

TAGAGAATCAAATGGGGATAGAGCGGAGTGTTAAAGTGCTGTTCAACGGGAAACCTGAAGTAACTTGGGTGGGGACCACA

GATTGCTACAAGTACATAATCAGTAACATCCAAACCTCTAGTCTGGGTTTCATACATTCAGATATTGAGACGCTCCCAAC

CAAAGATGCTGTTGAGAAATTGGAGGAATTTGCCTCTATCCTATCCTTATCCTTGATTTTGGGGAAAATCGGCTCTATTA

CAGTTGTCAAAATTATGCCCATTAGCGGGGATTTTACCCAAGGCTTCATAGCCTATGCCATCCAATATTTTAGGGAGAGC

CTGCTTGCCTATCCGAGATATAGTAACTTCATCTCGACTGAGTGTTACCTTATTATGATAGGATTAAAGGCCAATCGATT

GATAAACCCAGAAGCCATTAAGCAAAGCATAATCAGAGCGGGGGTCAGGACTGCACCAGGACTTGTGAGCCATGTATTAT

CAGAGAAACAAAAAGGTTGTATTCAATCTTTCTTGGGTGATCCTTATATCCAAGGAGACTTCAATAAGCACCTTAAATCT

CTAACCTCTATTGAGAAAATCCTGGTAAATTGTGGTCTCTCGATCAATGGCATAAAAATCTGTAGAGATCTAATCCACCA

TGATATCGCCTCCGGTCCAGACGGCCTTATGAGCTCTACAATTATTTTATATAGGGAATTGGCTCATTTCAAAGATAATA

TAAGGAGTCAGCACGGAATGTTCCACCCCTACCCGGTACTGGCCAACAGCAGGCAACGTGAATTAATCCTTCGAATAGCC

AAGAAATTCTGGGGGTATGTCTTGCTATATTCTGATGACCCGGCACTAATCAGACAAACAATCAAGAACTTGAAACGGAA

TCATCTAACCTTTGACTTACATAGTAACCCATTTATTAAAGGGCTATCCAAGGCTGAGAAACTGCTGGTGCGGACGAGTT

CACTAAGAAGAGAATGGTTGTTCACTCTCGAAACGAAAGAAGTAAAAGAGTGGTTCAAATTGGTAGGTTACAGTGCACTC

ATCAGAGGCTAGTCAGCTATGCATCTGCCCCCTTCTCCTCCGCCATGAGACCCCACTGGCGATCCAGAAGATTAAAGAAA

ACTACATATTGGATAAGTATCTATTCCCAGCTTTGTCTGGT

>KM089831.1/goat/China/2014-05-08

ACCAAACAAAGTTGGGTAAGGATAGATCTTATAATAACTATGGACTGGCAAACTTAGGAGTAAAGATCCTACTGTCGGGG

GGAGGAGGAGGAGCAAGATCCTTGATCATGGCGACTCTCCTTAAAAGCTTAGCATTGTTCAAAAGGAACAAAGACAAAGC

GCCGACAGCATCAGGTTCAGGAGGGGCCATCCGGGGGATTAAGAATGTTATCATAGTCCCGATTCCCGGAGACTCGTCCA

TCACTACCCGTTCAAGACTGCTCGACAGGCTTGTCAGATTGGCCGGAGATCCTGACATCAACGGGTCAAAGCTGACCGGG

GTGATGATCAGCATGTTATCCTTGTTCGTAGAGTCACCCGGGCAATTGATACAGCGAATCACAGATGATCCGGATGTTAG

TATCCGCCTTGTTGAGGTAGTTCAAAGTACTAGGTCCCAGTCCGGGTTGACCTTTGCATCACGTGGTGCTGATTTAGACA

ACGAGGCAGACATGTATTTTTCAACTGAGGGGCCCTCGAGTGGAGGTAAGAAAAGGATCAACTGGTTTGAGAACAGAGAA

ATAATAGACATAGAGGTGCAGGATCCAGAAGAGTTCAATATGTTGTTAGCCTCCATACTAGCACAAGTCTGGATCCTTCT

GGCCAAGGCTGTTACGGCACCAGATACGGCAGCTGACTCAGAACTGAGAAGGTGGGTTAAATACACACAACAAAGGAGAG

TGATTGGGGAATTTCGCCTTGACAAGGGGTGGCTGGACGCAGTCCGCAACAGAATTGCAGAAGATCTATCACTCCGGCGG

TTCATGGTATCTCTCATACTTGACATCAAGAGGACCCCAGGCAACAAGCCAAGGATTGCAGAAATGATCTGCGACATTGA

CAACTATATTGTCGAAGCAGGACTCACCAGTTTCATCCTTACTATCAAATTTGGTATCGAAACCATGTATCCTGCATTAG

GGCTCCACGAGTTTGCCGGAGAATTGTCCACTATTGAGTCCTTGATGAACTTGTATCAACAGCTAGGAGAGGTTGCACCC

TACATGGTAATTCTAGAGAATTCAATTCAGAACAAGTTTAGTGCAGGAGCTTATCCCCTCCTCTGGAGCTATGCGATGGG

TGTCGGAGTCGAGCTGGAGAACTCAATGGGGGGCTTGAACTTTGGCAGGTCATATTTTGACCCTGCCTATTTTCGTCTCG

GACAGGAGATGGTCAGAAGATCTGCAGGAAAGGTCAGCTCTGTGATTGCGGCTGAGCTCGGCATCACAGCAGAGGAAGCT

AAACTAGTCTCGGAAATCGCCTCGCAGGCTGGGGACGAAAGAACCGCCAGAGGGACTGGGCCTCGACAGGCGCAGGTTTC

CTTCCTTCAGCACCAAACAGGAGGGGGAGAGTCGTCCGCACCAGCGACCAGAGAAGGGGTCAAAGCTGCGATCCCAAACG

GAGCTGAAGAAAGGGACAGAAAGCAAACACGCCCAGGAAGGCCCAGAGGAGAGACCTCCGGCCAACTGCTCCTGGACATC

ATGCCAGAGGATGAGATCTTGCGAGAGTCTGGTCAAAACCCTCGTGAGGCTCAAAGATCGGCCGAGGCACTCTTCAGGCT

GCAGGCCATGGCCAAGATTCTGGAGGACCAGGAGGAGGGAGAAGACAACAGTCAGGTCTACAACGACAAGGATCTCCTCG

GCTAAGCAGACGCACCCTCTGTCGAAATCAGTGACAAGACATCTCCTACCAGTATTATAAAAAACTTAGGACCCAGGTCC

AAGCAAGCACACATCGACACTCCAACCAGTCGAGCGGAGACCACCGATGGCAGAAGAACAAGCATACCATGTCAACAAGG

GGCTGGAATGTATCAAGTCTCTCAAAGCCTCACCCCCGGATCTATCCACCATCAGAGATACCATCGAGAGCTGGAGAGAG

GGGCTTAGCCCCTCGGGCCGTGCAACACCGAACCCTGATACGTCCGAGGGAGATCATCAGAATATCAACCAATCATGCTC

ACCAGCAATCGGACCAAACAAAGTCTACTTGTCTCCTGAAGATAATCTCGGATTTAGAGAGATCACTGGCAACGACTGTG

AGGCTGGGCTCGGAGGGGTCCAGAGAGAAGGATCCAACTCTCAAGTACAGCGTTACCATGTTTATAGTCACGGGGGTGAA

GAGATTGAAGGACTCGAGGATGCTGACTCTCTCGTGGTTCAAGCAGATCCTCCATTTGCTAACGTCTTCAATGGAGGAGA

GGATGGATCTGACGACAGCGATGTGGACTCTGGCCCAGATGATCCCGGCAGAGGTACTCTATATGACCGGGGATCTGTTG

CCGGCAATGGTATCGCTGGGTCCACAGATGTCGAAAAACTAGAAGGTGCTGATATTCAAGAAGTTCTTAACTCCCAGAAA

GGCAAAGGAGGAAGATTCCAGGGCGGGAAAACCTTGCGAGTCCCGGAAATACCCGATGTCAAGCACTCCAGACCATCAGC

CCAATCAATTAAAAAGGGCACAGAAGGGAACTCAGTCTCATCTGGAACGGTGATAGAGTGTTTATCGATAAGTGGTGCAA

CCCAAACTGTGCCAGAGTCAAGATGGGAGTCATCAGAGCAAAATGCGTCTGTGGGGAGTGTCCTCAAGTCTGCGAGGAGT

GCAAAGACGATCCAGGGGTCGACACAAGAATCTGGTACCATAGCATCACTGACTCAGCCTAAAGAGAATGACTCCGAGTA

TGAGTATGAGGATGACCTATTTACAGAGATTCAGGACGTCCGTGCAAGCATTGCCAAGATCCATGATGACAATAAAACTA

TCCTCTCAAAGCTTGATTCTATACTGTTATTGAAAGGGGAAGTCGACACTATCAAGAAACAAATCAGCAAGCAGAATATA

AGTATATCCACCATTGAGGGCCATCTCTCCAGTATAATGATAGCCATCCCTGGCTTTGGGAAGGACATCAAGGACCCAAC

ATCCGAAGTCGAGTTGAACCCAGATCTAAGACCTATAATAAGCCGTGACTCTGGCAGAGCTCTCGCGGAGGTCCTCAAGA

AACCCGCTGTGGATAGGTCTCAGAAAATTGGAACCAAAGCCAACTCCAGCTCAAAGGGTCAGCTTCTTAAGGATCTCCAG

CTAAAACCTGTCGACAAGCAGGCAAGCTCTGCAATCGGGTTTGTCCCGTCCGACCATGAATCATCCAGAAATGTCATCCG

CTCCATAATCAAGTCGAGCAAGCTAAATATTGATCACAAGGACTATCTTCTAGATTTACTAAATGATGTGAAAGGCTCCA

AGGATCTTAAGGAATTTCACAAGATGCTAACAGCAATTCTCGCCAAGCACCCGTAACACATCCTCCAGTCATCATCTCAT

ACTCGACTAAAAACATCCTTTCAATCAGGCTATTACAAAAAACTTAGGAGCAAGGGCAACTGAGCTTCGCAGACAGGATG

ACCGAGATCTACGACTTCGATAAATCAGCATGGGATGTCAAAGGGTCAATTGCCCGCATAGAACCCACCACCTACCACGA

CGGCCGACTGATACCCCAGGTGAGGGTCATCGATCCTGGTCTGGGAGACAGAAAAGATGAGTGCTTTATGTACCTGTTTC

TCCTAGGAGTGATTGAGGATAACGACCCCCTGTCTCCCCCCGTCGGAAGAACCTTTGGCTCTCTACCTCTAGGGGTTGGT

AGGTCAACTGCCAAGCCAGAAGAACTACTAAGGGAGGCCACAGAATTAGATATAGTGGTGAGGCGCACTGCAGGAGTAAA

TGAGAAACTGGTATTTTACAACAACACTCCGCTGTCCTTGTTAACGCCCTGGAAGAAAGTTCTGACAACCGGAAGTGTGT

TTAGCGCTAACCAGGTTTGCAATGCAGTCAACCTAGTCCCGCTTGATACTCCCCAGAGATTCAGGGTTGTGTACATGAGC

ATAACTAGATTATCAGACAATGGTTACTATAGTGTGCCCAGAAGAATGCTGGAGTTCCGCTCGGCCAATGCAGTAGCCTT

CAACATCTTGGTTACACTAAGAATTGAAAATGGCACAAACCCTAGAAGATACATAGTCGGCTCATGGGAGAATTCAGAGG

TCACATTTATGGTACACGTGGGCAACTTTAGGAGAAAGAAGAACGAAGTATACTCTGCTGATTACTGCAAAATGAAGATT

GAAAAGATGGGTTTAGTTTTTGCCCTGGGTGGAATAGGCGGAACAAGTCTCCATATTAGAAGCACCGGGAAAATGAGCAA

AACCCTCCATGCACAGCTGGGGTTCAAGAAAATCTTATGTTACCCCCTAATGGATGTCAATGAGGATCTTAACCGATATC

TCTGGCGGGCAGAGTGCCGAATAGTCAAAATCCAAGCTGTCTTACAGCCATCAGTACCCCAAGAGTTCCGTGTCTACGAT

GATGTCATCATCAACGATGATCAAGGCTTGTTCAAGATCTTGTAGTTCATTTGCAAAGGAACGAACCCACCCCCAGGAAG

GGGCCCCACCCTCCCCCG-CCAAAAAACACACTCCAAGGCTCCCCGGCACAGCGCCAACACAGCTACTCCCGCGATGGAA

CGAGGTCGGCC-CCCGAACAAATCCGGTCCAGGGGGGGGGGCCCCCCCCCTGACCAACCGGGCGCCGACCGAGCAGGCCG

GCGAGAAGCCCCCAAGAGGCCCCGACGAAGACCGAGCACCCCATCCCCCC-AAAAACGTCCCCC-AACACGCCCGAGGCT

AGCCACCGGACACCCCCAGACCCGCCACCGGCGAGACAGAACCCCAAGGGGGGACCCTCCAAACCAAAGGAATCACCGGG

GTAGGCTCCGCCGAACCAGCACAGACCCGACCCAAGCAAGGCGGACCCCCAACACGAAACTGCCCAGCACACCCTACTCA

TCATCAAACAAAACTTAGGGGCCAAGTCCATACGAGAGGCCCTAGTCCGTCAGGCGGAGGGACGCAGAAAGGAAGGAGAC

ACGACCGTCCCAGAAGGAGGACACCCAACCACCGAAACGCCCGCCCCCAGGGAAGGAG--CCA-CCCCCACGCAGCGAAG

GAGCGGGTGGGAGACCTCGGGGAGGGCCCGACCGGGACCCGGGGGCCGGCGGCGCCGGCGGTGAGCAGGCCGGGCGCATA

CCTGACCCCACCGGAGGATCCGAGGAGGGCCCGACCCCGGCGGGACCAAACGGCAGGGAGCCCAACCCCACCAAAAAGAG

AGGCGAGGCCCCCCCTCCCTCCCCCCCCGCACACAACCCGCAGACAGACCGAGGGAGGGCCATCCCCCAGCGAGACAGGC

GGCTCCCTCGCAGCCGAGCCGCAGCTCCCCAGAATAGGACACCCCCGCACACTCCGAACACCCCCTCGCCCCCCGCCACC

GGGACAGGGGCTCCACCACCACAGACCGGAGCCACCGACAGCCAGCCAGAGGTCCCCTCCAGTCCCGCCCGGCGCCCCCC

CGAAGAGGCCCCGGCAAGGCCCCCCAGCCCGACGTCCCCCGCACGCACCGAAAGAAGGACCCGCCCGAGAACGTGCGGCA

CATCCATGCGCAAACATCATGACGCGGGTCGCAATTTTGACATTTCTGTTTCTTTTCCCAAATGTTGTTGCGTGTCAGAT

TCACTGGGGCAATCTATCCAAGATCGGGATTGTAGGAACAGGGAGTGCCAGCTACAAGGTGATGACTAGGCCAAGCCACC

AGACTCTGGTTATAAAGTTAATGCCAAATATAACGGCCATCGACAATTGTACAAAGTCAGAGATTGCAGAGTACAAGAGA

TTGCTGATCACAGTGTTAAAGCCTGTAGAGGATGCTCTGTCGGTGATAACCAAGAATGTAAGACCAATTCAAACTCTAAC

ACCTGGGCGTAGAACCCGCCGTTTTGCTGGAGCTGTTCTGGCCGGGGTAGCACTTGGAGTTGCGACAGCCGCTCAGATAA

CTGCAGGAGTCGCCCTTCATCAATCATTGATGAACTCCCAAGCAATTGAGAGTTTAAAAACCAGTCTTGAGATGTCGAAT

CAGGCAATAGAAGAAATCAGACTTGCAAATAAGGAGACCATACTGGCAGTACAGGGCGTCCAGGATTATATCAACAATGA

GCTCGTCCCTTCTGTTCATAGAATGTCATGCGAGCTGGTAGGTCACAAGCTCGGCCTCAAGCTCCTTAGGTACTACACCG

AGATCCTGTCCATATTCGGGCCCAGTCTTCGAGACCCGATATCTGCCGAAATATCAATCCAGGCACTTAGTTATGCATTA

GGCGGAGACATTAATAAAATCCTGGACAAGCTTGGGTATAGCGGTGGGGATTTCCTTGCCATCCTAGAAAGCAAGGGAAT

AAAGGCCCGGGTTACATATGTGGACACAAGAGATTACTTTATAATCCTTAGCATCGCCTACCCAACCTTATCTGAGATCA

AGGGAGTGATAGTTCACAAGATAGAAGCTATAACATACAACATTGGGGCACAGGAGTGGTATACTACTATCCCTAAATAT

GTAGCCACTCAGGGGTATCTGATATCGAACTTTGATGAGACGTCATGCGTATTCACTCCAGAGGGGACAGTTTGCAGCCA

GAATGCGTTGTACCCAATGAGCCCATTGCTTCAGGAATGTTTCAGGGGGTCAACAAAATCGTGCGCCAGAACCTTAGTTT

CAGGGACCATAAGTAATAGATTTATCCTATCAAAAGGGAACCTGATTGCAAATTGTGCGTCAGTTTTGTGCAAATGTTAC

ACAACAGAAACAGTTATCAGCCAAGATCCTGACAAACTACTAACTGTTGTAGCATCCGACAAGTGTCCTGTAGTTGAGGT

GGATGGAGTGACAATACAGGTCGGCAGTCGAGAGTATCCGGATTCTGTATACTTACACAAAATAGACTTAGGTCCAGCCA

TCTCCCTAGAAAAACTGGATGTAGGCACCAATTTAGGCAATGCAGTCACAAGACTGGAGAATGCAAAGGAGCTCCTAGAT

GCATCAGACCAAATACTGAAGACTGTTAAAGGGGTACCTTTCAGTGGGAATATGTACATAGCACTGGCAGCTTGCATAGG

AGTATCCCTGGGCCTTGTCACATTAATATGCTGCTGTAAGGGGAGGTGTAAGAACAAGGAAATCCCTATCTCCAAAATCA

ACCCAGGGCTCAAACCCGACCTGACCGGGACCTCAAAGTCGTACGTAAGATCACTGTAGTCAGAATCACCTGGATCATCT

GGCATCACACACATACATGCACGACACAGGCAGTCCGAGGACGCAAGAAACCCAGCCTCCGGTCACCCACCCGACCCCAC

TCCACGCTCCACCACACATTAGTCATCAAACAAAACTTAGGACGAAAGGTCAATCACCATGTCCGCACAAAGGGAGAGGA

TCAATGCCTTCTACAAAGACAATCCTCACAATAAGAACCATAGGGTGATCCTGGATAGAGAACGCTTGGTCATTGAAAGA

CCCTACATCTTGCTTGGAGTCCTGCTGGTAATGTTCCTGAGTCTAATCGGACTGCTGGCCATTGCAGGGATCAGGCTTCA

CCGGGCCACCGTTGGAACTTCAGAGATCCAGAGTCGGCTGAATACCAATATTAAGTTGACCGAATCTATTGATCACCAGA

CTAAGGATGTCTTAACTCCCCTTTTTAAAATCATTGGCGATGAAGTCGGCATCAGAATTCCACAGAAATTCAGTGATCTT

GTCAAGTTCATCTCCGATAAGATTAAATTCCTCAACCCTGATAGAGAGTATGATTTCAGAGATCTCCGGTGGTGTATGAA

TCCCCCCGAGAGAGTCAAAATTAATTTTGATCAGTTTTGTGAGTACAAGGCTGCGGTTAAGTCAATTGAACATATATTTG

AGTCACCACTCAACAAGTCAAAAAAGCTGCAATCTTTGACTCTCGGGCCCGGAACAGGCTGTCTAGGCAGGACAGTAACA

AGAGCCCATTTCTCAGAACTTACAATGACCTTAATGGACCTGGATCTAGAGATGAAGCACAACGTGTCCTCAGTGTTTAC

CGTAGTTGAAGAGGGATTATTCGGAAGAACATATACCGTCTGGAGATCCGATGCCAGGGATCCGAGCACCGATCTAGGTA

TCGGCCATTTTTTAAGAGTCTTCGAGATTGGACTGGTAAGAGATCTTGGGCTGGGTCCCCCTGTTTTTCATATGACCAAC

TATCTCACGGTGAACATGAGTGATGACTATCGGAGATGTCTTTTAGCGGTAGGGGAGTTGAAGTTGACAGCCCTATGCAC

CTCATCTGAGACTGTGACACTGAGTGAGAGAGGAGTTCCAAGGAGGGAACCTCTTGTGGTTGTGATACTTAATCTAGCTG

GACCCACTCTAGGGGGAGAGCTATACAGTGTCTTGCCTACCTCTGATCTCATGGTGGAGAAACTCTATTTGTCTTCACAT

AGAGGGATCATCAAAGATGATGAGGCCAATTGGGTAGTGCCGTCTACCGATGTTCGTGATCTTCAGAATAAAGGTGAATG

TCTGGTGGAAGCATGCAAGACTCGACCTCCTTCATTTTGCAATGGCACAGGATCAGGCCCGTGGTCAGAAGGGAGAATCC

CTGCCTACGGGGTGATCAGGGTCAGTCTTGACTTAGCGAGTGACCCGGATGTAGTTATCACTTCAGTGTTTGGCCCACTG

ATACCTCACCTATCCGACATGGATCTTTACAACAACCCGTTTTCAAAAGCTGTATGGTTGGCTGTACCACCTTATGAGCA

GTCATTTCTAGGAATGATAAATACAATTGGATTCCCTAATAGAGCAGAGGTTATGCCGCACATTTTGACCACAGAGATCA

GAGGCCCTCGGGGTCGTTGCCATGTTCCCATAGAATTGTCCCGCCGGGTTGATGACGATATCAAGATCGGGTCCAACATG

GTCATATTGCCGACGATGGACCTGAGGTATATTACAGCCACTTATGATGTTTCCAGGAGCGAGCATGCAATCGTGTACTA

TATCTATGACACGAGTCGCTCATCATCTTACTTCTACCCAGTTCGACTGAATTTCAAAGGCAATCCTCTCTCTTTGAGGA

TAGAGTGTTTCCCCTGGCGTCGTAAGGTGTGGTGCTACCATGATTGTCTTATATACAACACCATAACAGGTGAAGAGGTC

CATACGAGAGGGCTGACCGGCATAGAGGTAACATGCAATCCAGTCTGAGCAGAGCTACGACCGTCACTCAGGCAGTCCCT

TGAGTCGCCACCGAGTCCAAGCAGCACAGCCTGGGACAATCAACGGCACAGCCCAGCCAACAATGTTATAAAAAACTTAG

GAGCCAGGGTTGTAGGGGCCATGGACTCTCTATCAGTCAATCAGGTCTTGTACCCTGAGGTTCATCTAGACAGCCCTATT

GTCACAAACAAACTAGTTGCCATCCTTGAGTACTCGGGTATCGACCATAACTATGTTCTTGAAGACCAGACTCTTGTCAA

GAATATTAGGTATAGGCTGGGGTGCGGTTTTTCAAATCAAATGATCATCAATAACAGGGGGGTGGGTGAAACAGTCAATT

CTAAACTTAAAAGTTACCCCCATAATCGTCATATCATATACCCGGATTGCAATAAGGAGTTGTTTTGTATCAAGGATAGC

TGCATATCTAAGAAGCTCTCGGAGCTATTCAAGAAGGGTAATTCCTTGTACTCTAAGATAAGTCACCAGGTACTGGATTG

TCTTAAGAGAGTCAATGGGAAATTAGGTCTGGGCACAGATCTTACCCATGGTCTGAAGGAGGGTATCCTTGACCCAGGGT

TGCACATGCATAGCTCTCAATGGTTCGAGACTTTCCTGTTCTGGTTCACTATCAAGACAGAGATGAGATCAATGATCAAA

GAACAGTCCCATATATGCCACAAGAGGAGGTATAACCCGACTTTTGTGTCGGGGGATGCATTCGAGGTGCTCGTATCGCG

AGACCTCGTTGTGATAATTGATAAGAATACCCAGTATGTCTTCTACTTAACATTTGAGCTGGTCCTTATGTATTGTGATG

TCATAGAGGGTAGACTTATGACGGAGACAGCCATGGCCATAGACCAGAGATATTCAGAACTTCTAAGCCGGGTCAGATAC

TTGTGGGATCTTATTGATGGGTTCTTCCCAACACTAGGCAACACCACATACCAAATTGTTGCTCTGCTTGAACCATTATC

GTTGGCTTATCTTCAACTTCAGGATGTCACTCTGGAGTTAAGAGGTGCTTTTTTAGACCATTGCTTCAAAGAACTCTATG

AGATACTGGAACATTGCGGCATTGACACAGCAGGCACTTACAATTCCATCACTGAAGGATTGGATTACGTATTCATCACC

CACGATATACATCTAACTGGGGAGATTTTTTCATTTTTCCGGAGTTTCGGACACCCCCGTCTAGAAGCAGTCACCGCTGC

GGAAAATGTCAGAAAACATATGAACCAACCGAAGGTAATCAGTTATGAGACTATGATGAAAGGCCATGCGGTATTTTGCG

GGATAATCATAAATGGTTTTCGGGACCGACATGGCGGCAGCTGGCCCCCTGTTGCACTGCCAGAACATGCCTCTGCTGCG

ATCCGGAATGCGCAGGCATCAGGTGAGGGACTGACCCATGACCTGTGTATAGACAACTGGAAATCCTTTGTTGGATTCAA

ATTTGGCTGCTTTATGCCACTCAGCCTAGATAGTGATTTGACCATGTATCTCAAAGACAAGGCATTGGCTGCACTGAAGA

ATGAGTGGGATTCAGTTTACCCGAAAGAATACCTCCGTTATAACCCACCTAGAGGGACAGAGTCAAGGCGATTGGTAGAG

GTATTCCTGAATGACTCCAGCTTTGATCCTTATAACATGATAATGTACGTGGTGAATGGCTCCTACCTTAAAGACCCTGA

GTTTAACCTCTCATACAGCCTAAAAGAGAAGGAGATAAAGGAGACAGGGCGGTTGTTTGCCAAAATGACCTATAAGATGC

GGGCCTGTCAGGTAATCGCTGAAAATCTGATATCGAATGGTGTTGGGAAGTATTTCCGAGACAATGGGATGGCAAAGGAC

GAGCATGACCTAACAAAAGCCCTTCACACCCTGGCAGTCTCAGGTGTTCCCAAAAACAACAAAGATAACCACCGAGGTGG

GCCTCCCAGAAGGACAACAAACCGAGGGGTGAGATCAAGCCAAGGCACCAAAACACAAGATAGAGACAAGGTTCAAGGGG

GACCTATGTACAACTATTTGCGATGCCAACCGATCAGCCCTGATCAGGGTGAGTCATACGAGACTGTTAGTGCATTCATC

ACCGCTGACCTTAAGAAGTATTGCCTGAATTGGAGATACGAGACAATCAGCATATTTGCACAGAGGCTGAATGAAATATA

TGGACTACCATCCTTCTTTCAATGGTTACACAGGATATTGGAAAAATCCGTACTCTACGCCAGTGACCCACATTGTCCTC

CCGATCTAGATAATCATATCCCTCTGGACAGTGTCCCCAATGCCCAAATATTCATCAAGTACCCAATGGGCGGAATAGAA

GGTTATTGCCAAAAACTATGGACAATCAGCACTATACCATACTTGTATCTGGCAGCCTATGAGAGCGGAGTAAGAATCGC

CTCACTGGTGCAGGGTGACAATCAGACAATCGCAGTGACAAAAAGAGTTCCAAGTTCTTGGCCTTATTCACTAAAAAAGA

GGGAGGCCTCCAAGGCCGCTCAAAATTACTTCGTAGTCTTAAGGCAAAGATTGCACGATGTAGGTCATCACTTAAAGGCT

AATGAGACCATAGTATCCTCTCACTTTTTTGTATATTCCAAAGGGATTTATTATGACGGCCTGCTAGTCTCACAATCACT

AAAGAGCATCGCCAGATGTGTCTTCTGGTCCGAGACTATAGTGGATGAAACCAGAGCGGCGTGCAGTAATATTGCAACAA

CTGTCGCTAAGAGTATAGAGAGGGGTTATGATAGGTACCTTGCATATTCTTTGAATATCCTCAAGATTTTCCAACAGATC

CTTATATCCCTTAACTTCACTATTAACACAACAATGACTCAGGATGTCGTGGCACCGATCATCGAGAACGGTGATTTGCT

GATAAGGATGGCACTCTTGCCAGCACCCATCGGGGGTCTCAATTATCTTAACATGAGCAGGTTATTTGTGAGAAATATCG

GTGACCCGGTCACCTCCTCTATAGCCGACCTGAAGAGGATGATAGATGCCGGGCTAATGCCAGAAGAAACATTGCATCAA

GTGATGACCCAGACCCCGGGAGAATCATCCTACCTAGATTGGGCAAGTGACCCTTACTCTGCCAACCTACCCTGCGTACA

GAGTATCACTCGCCTTCTCAAAAACATCACTGCACGATATATTTTAATCAGCAGCCCAAACCCGATGCTGAAAGGGTTAT

TTCATGAGGGGAGTAGAGATGAAGACGAGGAGCTTGCAAGTTTCCTAATGGATCGGCATATAATTGTTCCAAGAGCTGCA

CATGAAATCTTAGACCATAGTATAACCGGAGCAAGAGAAGCTATAGCCGGGATGTTGGACACCACCAAGGGTCTGATTAG

AACAAGTATGAAACGGGGCGGCCTCACCCCTCGAGTCTTAGCCCGCCTTTCCAATTATGATTATGAACAATTCCGATCCG

GGATAACACTATTAACAAAGGAAGGGCAGTGTTATCTCATTGACAAGGACTCGTGCTCGGTGCAGCTCGCTATAGCTCTG

AGGGGCCACATGTGGGCTAAGTTGGCTCGCGGGAGACCAATCTATGGGTTGGAGGTGCCTGATGTATTAGAATCGATGAA

CGGCTACCTTATCAAACGCCACGAGTCGTGTGCTATCTGTGAAACGGGCTCAAGTCACTACGGATGGTTTTTCGTCCCTG

CAGGGTGCCAGCTTGACGATGTATCTAGAGAGACTTCGGCTCTTCGTGTACCTTATGTCGGATCAACAACTGAGGAAAGG

ACAGACATGAAACTTGCATTTGTTAGGTCTCCAAGTAGATCCCTCAAATCAGCAGTTAGAATTGCAACAGTTTACTCATG

GGCTTACGGGGATGATGAGAAATCATGGAGTGAAGCCTGGATGCTAGCTAGGCAGAGAGCCAATATCACCTTAGATGAAT

TGAGAATGATCACTCCAGTCTCCACATCCACCAACTTAGCCCACCGGTTGAGGGATCGGAGTACCCAGGTGAAATACTCG

GGGACATCCCTTGTGAGGGTTGCAAGATACACAACCATCTCCAATGATAACTTGTCGTTTGTGATATCTGAGAAAAAAGT

GGATACCAACTTCATTTATCAGCAAGGGATGTTGCTTGGTCTTGGGATCCTTGAGAACCTCTTTAGGCTAGAGGCCACCA

CCGGAGTATCCAACACAGTGCTACACCTGCACGTGGAAACAGAATGTTGTGTTGTACCTATGGTGGATCACCCAAGGATA

CCGAGTCTCCGTAATATAAAAGTTACGGATGAGCTATGCACAAACCCTTTGATCTACGACAGGTCCCCCATCATAGAGCA

CGATGCAACCCGGCTATATTCACAGAGCCACAGGAGACATTTGGTGGAGTTTGTTACCTGGTCAACAAGCCAGCTTTACC

ATATACTGGCTAAGTCTACAGCAATGTCCATGATTGAATTGATCACGAGATTCGAGAAAGATCACATGAATGAAATAGCC

GCCCTGATTGGCGATGACGACATCAACAGCTTCATCACAGAATTTTTGCTAGTGGAGCCTAGATTATTTATAGTTTACCT

TGGTCAGTGTGCTGCCATCAATTGGGCCTTTGATATACATTATCATCGGCCCTCGGGCAAGTACCAGATGGGGGAACTTC

TCTACTCTCTGCTCTCTCGGATGAGCAAAGGAGTATATAAGATCTTTACTAATGCTTTGAGCCACCCCAAAGTTTACAAG

AAATTTTGGCGAAGTGGCATCATTGAGCCGATTCATGGCCCATCCTTGGATACACAGAATTTACATGTCACTGTCTGTGA

CATGATATATGGGTCCTATGTCACCTATTTGGATCTTTTGCTGAATGATGAGCTAGATGATTACCCGTATTTGCTCTGCG

AGAGTGATGAGGACGTGGTCACAGACAGGTTCGACAACATTCAAGCCAAACATCTCTGTGTATTGGCCGATGTATATTGC

AGCTCCAAGAGGTGTCCTTCAATAATCGGGATGTCTCCTATAGAAAAATGTACCATTCTCACACATTACATCAAGGGAGA

ATCAGTACAATCCCCGTCTGGGACCTCATGGAACACTGATCCCCTTGTAGTAGATCATTACTCATGCTCTCTGATCTACC

TTCGCCGCGGTTCCATCAAACAAATCAGGTTGAGGGTGGATCCTGGGTTTGTATTCGAGGCGTTGACAGACGTCGACTTT

AAACAACCTCGCAAAGCTAAGTTAGATATATCGGTCGTTGGGTTGACTGATTTCTCTCCCCCTTACGACAACGTCGGTGA

TTTTTTAGGGACTATCAACACATTGAGGCACGATCTGCCTGTCACCGGAACCGGGGTCTCGAACTATGAAGTCCACGCTT

ATCGTAGAATTGGCCTGAATTCATCAGCATGTTATAAAGCCGTAGAGATCTCCACGTTAATCAAGCCATCCTTAGAAGTC

GGAGAGCATGGATTGTTCTTAGGAGAAGGTTCCGGTTCAATGCTGGCTGCGTACAAAGAAGTTCTTAAATTAGCAAATTG

TTACTACAACAGCGGAGTAACAGCGGAGGGCAGAGCCGGACAGAGGGAAATATCTCCCTATCCTTCAGAGATGAGCCTAG

TAGAGAATCAAATGGGGATAGAGCGGAGTGTTAAAGTGCTGTTCAACGGGAAACCTGAAGTAACTTGGGTGGGGACCACA

GATTGCTACAAGTACATAATCAGTAACATCCAAACCTCTAGTCTGGGTTTCATACATTCAGATATTGAGACGCTCCCAAC

CAAAGATGCTGTTGAGAAATTGGAGGAATTTGCCTCTATCCTATCCTTATCCTTGATTTTGGGGAAAATCGGCTCTATTA

CAGTTGTCAAAATTATGCCCATTAGCGGGGATTTTACCCAAGGCTTCATAGCCTATGCCATCCAATATTTTAGGGAGAGC

CTGCTTGCCTATCCGAGATATAGTAACTTCATCTCGACTGAGTGTTACCTTATTATGATAGGATTAAAGGCCAATCGATT

GATAAACCCAGAAGCCATTAAGCAAAGCATAATCAGAGCGGGGGTCAGGACTGCACCAGGACTTGTGAGCCATGTATTAT

CAGAGAAACAAAAAGGTTGTATTCAATCTTTCTTGGGTGATCCTTATATCCAAGGAGACTTCAATAAGCACCTTAAATCT

CTAACCTCTATTGAGAAAATCCTGGTAAATTGTGGTCTCTCGATCAATGGCATAAAAATCTGTAGAGATCTAATCCACCA

TGATATCGCCTCCGGTCCAGATGGCCTTATGAGCTCTACAATTATTTTATATAGGGAATTGGCTCATTTCAAAGATAATA

TAAGGAGTCAGCACGGAATGTTCCACCCCTACCCGGTACTGGCCAACAGCAGGCAACGTGAATTAATCCTTCGAATAGCC

AAGAAATTCTGGGGGTATGTCTTGCTATATTCTGATGACCCGGCACTAATCAGACAAACAATCAAGAACTTGAAACGGAA

TCATCTAACCTTTGACTTACATAGTAACCCATTTATTAAAGGGCTATCCAAGGCTGAGAAACTGCTGGTGCGGACGAGTT

CACTAAGAAGAGAATGGTTGTTCACTCTCGAAACGAAAGAAGTAAAAGAGTGGTTCAAATTGGTAGGTTACAGTGCACTC

ATCAGAGGCTAGTCAGCTATGCATCTGCCCCCTTCTCCTCCGCCATGAGACCCCACTGGCGATCCAGAAGATTAAAGAAA

ACTACATATTGGATAAGTATCTATTCCCAGCTTTGTCTGGT

>KM089832.1/goat/China/2014-05-08

ACCAAACAAAGTTGGGTAAGGATAGATCTTATAATAACTATGGACTGGCAAACTTAGGAGTAAAGATCCTACTGTCGGGG

GGAGGAGGAGGAGCAAGATCTTTGATTATGGCGACTCTCCTTAAAAGCTTAGCATTGTTCAAAAGGAACAAAGACAAAGC

GCCGACAGCATCAGGTTCAGGAGGGGCCATCCGGGGGATTAAGAATGTTATCATAGTCCCGATTCCCGGAGACTCGTCCA

TCACTACCCGTTCAAGACTGCTCGACAGACTTGTCAGATTGGCCGGAGATCCTGACATCAACGGGTCAAAGCTGACCGGG

GTGATGATCAGCATGTTATCCTTGTTCGTAGAGTCACCCGGGCAATTGATACAGCGAATCACAGATGATCCGGATGTTAG

TATCCGCCTTGTTGAGGTAGTTCAAAGTACTAGGTCCCAGTCCGGGTTGACCTTTGCATCACGTGGTGCTGATTTAGACA

ACGAGGCAGACATGTATTTTTCAACTGAGGGGCCCTCGAGTGGAGGTAAGAAAAGGATCAACTGGTTTGAGAACAGAGAA

ATAATAGACATAGAGGTGCAGGATCCAGAAGAGTTCAATATGTTGTTAGCCTCCATACTAGCACAAGTCTGGATCCTTCT

GGCCAAGGCTGTTACGGCACCAGATACGGCAGCTGACTCAGAACTGAGAAGGTGGGTTAAATACACACAACAAAGGAGAG

TGATTGGGGAATTTCGCCTTGATAAGGGGTGGCTGGACGCAGTCCGCAACAGAATTGCAGAAGATCTATCACTCCGGCGG

TTCATGGTATCTCTCATACTTGACATCAAGAGGACCCCAGGCAACAAGCCAAGGATTGCAGAAATGATCTGCGACATTGA

CAACTATATTGTCGAAGCAGGACTCGCCAGTTTCATCCTTACTATCAAATTTGGTATCGAAACCATGTATCCTGCATTAG

GGCTCCACGAGTTTGCCGGAGAATTGTCCACTATTGAGTCCTTGATGAACTTGTATCAACAGCTAGGAGAGGTTGCACCC

TACATGGTAATTCTAGAGAATTCAATTCAGAACAAGTTTAGTGCAGGAGCTTATCCCCTCCTCTGGAGCTATGCGATGGG

TGTCGGAGTCGAGCTGGAGAACTCAATGGGGGGCTTGAACTTTGGCAGGTCATATTTTGACCCTGCCTATTTTCGTCTCG

GACAGGAGATGGTCAGAAGATCTGCAGGAAAGGTCAGCTCTGTGATTGCGGCTGAGCTCGGCATCACAGCAGAGGAAGCT

AAACTAGTCTCGGAAATCGCCTCGCAGGCTGGGGACGAAAGAACCGCCAGAGGGACTGGGCCTCGACAGGCGCAGGTTTC

CTTCCTTCAGCACCAAACAGGAGGGGGAGAGTCGTCCGCACCAGCGACCAGAGAAGGGGTCAAAGCTGCGATCCCAAACG

GAGCTGAAGAAAGGGACAGAAAGCAAACACGCCCAGGAAGGCCCAGAGGAGAGACCTCCGGCCAACTGCTCCTGGACATC

ATGCCAGAGGATGAGATCTTGCGAGAGTCTGGTCAAAACCCTCGTGAGGCTCAAAGATCGGCCGAGGCACTCTTCAGGCT

GCAGGCCATGGCCAAGATTCTGGAGGACCAGGAGGAGGGAGAAGACAACAGTCAGGTCTACAACGACAAGGATCTCCTCG

GCTAAGCAGACGCACCCTCTGTCGAAATCAGTGACAAGACATCTCTTACCAGTATTATAAAAAACTTAGGACCCAGGTCC

AAGCAAGCACACATCGACACTCCAACCAGTCGAGCGGAGACCACCGATGGCAGAAGAACAAGCATACCATGTCAACAAGG

GGCTGGAATGTATCAAGTCTCTCAAAGCCTCACCCCCGGATCTATCCACCATCAGAGATACCATCGAGAGCTGGAGAGAG

GGGCTTAGCCCCTCGGGCCGTGCAACACCGAACCCTGATACGTCCGAGGGAGATCATCAGAATATCAACCAATCATGCTC

ACCAGCAATCGGACCAAACAAAGTCTACTTGTCTCCTGAAGATAATCTCGGATTTAGAGAGATCACTGGCAACGACTGTG

AGGCTGGGCTCGGAGGGGTCCAGAGAGAAGGATCCAACTCTCAAGTACAGCGTTACCATGTTTATAGTCACGGGGGTGAA

GAGATTGAAGGACTCGAGGATGCTGACTCTCTCGTGGTTCAAGCAGATCCTCCAGTTGCTAACGTCTTCAATGGAGGAGA

GGATGGATCTGACGACAGCGATGTGGACTCTGGCCCAGATGATCCCGGCAGAGATACTCTATATGACCGGGGATCTGTTG

CCGGCAATGGTATCGCTAGGTCCACAGATGTCGAAAAACTAGAAGGTGCTGATATTCAAGAAGTTCTTAACTCCCAGAAA

GGCAAAGGAGGAAGATTCCAGGGCGGGAAAACCTTGCGAGTCCCGGAAATACCCGATGTCAAGCACTCCAGACCATCAGC

CCAATCAATTAAAAAGGGCACAGACGGGAACTCAGTCTCATCTGGAACGGTGATAGAGTGTTTATCGATAAGTGGTGCAA

CCCAAACTGTGCCAGAGTCAAGATGGGAGTCATCAGAGCAAAATGTGTCTGTGGGGAGTGTCCTCAAGTCTGCGAGGAGT

GCAAAGACGATCCAGGGGTCGACACAAGAATCTGGTACCATAGCATCACTGACTCAGCCTAAAGAGAATGACTCCGAGTA

TGAGTATGAGGATGACCTATTTACAGAGATTCAGGACATCCGCGCAAGCATTGCCAAGATCCATGATGACAATAAAACTA

TCCTCTCAAAGCTTGATTCTATACTGTTATTGAAAGGGGAAGTCGACACTATCAAGAAACAAATCAGCAAGCAGAATATA

AGTATATCCACCATTGAGGGCCATCTCTCCAGTATAATGATAGCCATCCCTGGCTTTGGGAAGGACATCAAGGACCCAAC

ATCCGAAGTCGAGTTGAACCCAGATCTAAGACCTATAATAAGCCGTGACTCTGGCAGAGCTCTCGCGGAGGTCCTCAAGA

AACCCGCTGTGGATAGGTCTCAGAAAATTGGAACCAAAGCCAACTCCAGCTCAAAGGGTCAGCTTCTTAAGGATCTCCAG

CTAAAACCTGTCGACAAGCAGGCAAGCTCTGCAATCGGGTTTGTCCCGTCCGACCATGAATCATCCAGAAATGTCATCCG

CTCCATAATCAAGTCGAGCAAGCTAAATATTGATCACAAGGACTATCTTCTAGATTTACTAAATGATGTGAAAGGCTCCA

AGGATCTTAAGGAATTTCACAAGATGCTAACAGCAATTCTCGCCAAGCACCCGTAACACATCCTCCAGTCATCATCTCAT

ACTCGACTAAAAACATCCTTTCAATCAGGCTATTACAAAAAACTTAGGAGCAAGGGCAACTGAGCTTCGCAGACAGGATG

ACCGAGATCTACGACTTCGATAAATCAGCATGGGATGTCAAAGGGTCAATTGCCCGCATAGAACCCACCACCTACCACGA

CGGCCGACTGATACCCCAGGTGAGGGTCATCGATCCTGGTCTGGGAGACAGAAAAGATGAGTGCTTTATGTACCTGTTTC

TCCTAGGAGTGATTGAGGATAACGACCCCCTGTCTCCCCCCGTCGGAAGAACCTTTGGCTCTCTACCTCTAGGGGTTGGT

AGGTCAACTGCCAAGCCGGAAGAACTACTAAGGGAGGCCACAGAATTAGATATAGTGGTGAGGCGCACTGCAGGAGTAAA

TGAGAAACTGGTATTCTACAACAACACTCCGCTGTCCTTGTTAACGCCCTGGAAGAAAGTTCTGACAACCGGAAGTGTGT

TTAGCGCTAACCAGGTTTGCAATGCAGTCAACCTAGTCCCGCTTGATACTCCCCAGAGATTCAGGGTTGTGTACATGAGC

ATAACTAGATTATCAGACAATGGTTACTATAGTGTGCCCAGAAGAATGCTGGAGTTCCGCTCGGCCAATGCAGTAGCCTT

CAACATCTTGGTTACACTAAGAATTGAAAATGGCACAAACCCTAGAAGATACATAGTCGGCTCATGGGAGAATTCAGAGG

TCACATTTATGGTACACGTGGGCAACTTTAGGAGAAAGAAGAACGAAGTATACTCTGCTGATTACTGCAAAATGAAGATT

GAAAAGATGGGTTTAGTTTTTGCCCTGGGTGGAATAGGCGGAACAAGTCTCCATATTAGAAGCACCGGGAAAATGAGCAA

AACCCTCCATGCACAGCTGGGGTTCAAGAAAATCTTATGTTACCCCCTAATGGATGTCAATGAGGATCTTAACCGATATC

TCTGGCGGGCAGAGTGCCGAATAGTCAAAATCCAAGCTGTCTTACAGCCATCAGTACCCCAAGAGTTCCGTGTCTACGAT

GATGTCATCATCAACGATGATCAAGGCTTGTTCAAGATCTTGTAGTTCATTTGCAAAGGAACGAACCCACCCCCAGGAAG

GGGCCCCACCCTCCCCCG-CCCAAAAACACACTCCAAGGCTCCCCGGCACAGCGCCAACACAGCTACTCCCGCGATGGAA

CGAGGTCGGCC-CCCGAACAAATCCGGTCCAGGGGGGGGGGCCCCCCCCCTGACCAACCGGGCGCCGACCGAGCAGGCCG

GCGAGAAGCCCCCAAGAGGCCCCGACGAAGACCGAGCACCCCATCCCCCC-AAAAACGTCCCCC-AACACGCCCGAGGCC

AGCCACCGGACACCCCCAGACCCGCCACCGGCGAGACAGAACCCCAAGGGGGGACCCTCCAAACCAAAGGAATCACCGGG

GTAGGCTCCGCCGAACCAGCACAGACCCGACCCAAGCAAGGCGGACCCCCAACACGAAACTGCCCAGCACACCCTACTCA

TCATCAAACAAAACTTAGGGGCCAAGTCCATACGAGAGGCCCTAGTCCGTCAGGCGGAGGGACGCAGAAAGGAAGGAGAC

ACGACCGTCCCAGAAGGAGGACACCCAACCACCGAAACGCCCGCCCCCAGGGAAGGAG--CCA-CCCCCACGCAGCGAAG

GAGCGGGTGGGAGACCTCGGGGAGGGCCCGACCGGGACCCGGGGGCCGGCGGCGCCGGCGGTGAGCAGGCCGGGCGCATA

CCTGACCCCACCGGAGGATCCGAGGAGGGCCCGACCCCGCGTGGACCAAACGGCAGGGAGCCCAACCCCACCAAAAAAAG

AGGCGAGGCCCCCCCTCCCTCCCCCCCCGCACACAACCCGCAGACAGACCGAGGGAGGGCCATCCCCCAGCGAGACAGGC

GGCTCCCTCGCAGCCGAGCCGCAGCTCCCCAGAATAGGACACCCCCGCACACTCCGAACACCCCCTCGCCCCCCGCCACC

GGGACAGGGGCTCCACCACCACAGACCGGAGCCACCGACAGCCAGCCAGAGGTCCCCTCCAGTCCCGCCCGGCGCCCCCC

CGAAGAGGCCCCGGCAAGGCCCCCCAGCCCGACGTCCCCCGCACGCACCGAAAGAAGGACCCGCCCGAGAACGTGCGGCA

CATCCATGCGCAAACATCATGACGCGGGTCGCAATTTTGACATTTCTGTTTCTTTTCCCAAATGTTGTTGCGTGTCAGAT

TCACTGGGGCAATCTATCCAAGATCGGGATTGTAGGAACAGGGAGTGCCAGCTACAAGGTGATGACTAGGCCAAGCCACC

AGACTCTGGTTATAAAGTTAATGCCAAATATAACGGCCATCGACAATTGTACAAAGTCAGAGATTGCAGAGTACAAGAGA

TTGCTGATCACAGTGTTAAAGCCTGTAGAGGATGCTCTGTCGGTGATAACCAAGAATGTAAGACCAATTCAAACTCTAAC

ACCTGGGCGTAGAACCCGCCGTTTTGCTGGAGCTGTTCTGGCCGGGGTAGCACTTGGAGTTGCGACAGCCGCTCAGATAA

CTGCAGGGGTCGCCCTTCATCAATCATTGATGAACTCCCAAGCAATTGAGAGTTTAAAAACCAGTCTTGAGATGTCGAAT

CAGGCAATAGAAGAAATCAGACTTGCAAATAAGGAGACCATACTGGCAGTACAGGGCGTCCAGGATTATATCAACAATGA

GCTCGTCCCTTCTGTTCATAGGATGTCATGCGAGCTGGTAGGTCACAAGCTCGGCCTCAAGCTCCTTAGGTACTACACCG

AGATCCTGTCCATATTCGGGCCCAGTCTTCGAGACCCGATATCTGCCGAAATATCAATCCAGGCACTTAGTTATGCATTA

GGCGGAGACATTAATAAAATCCTGGACAAGCTTGGGTATAGCGGTGGGGATTTCCTTGCCATCCTAGAAAGCAAGGGAAT

AAAGGCCCGGGTTACATATGTGGACACAAGAGATTACTTTATAATCCTTAGCATCGCCTACCCAACCTTATCTGAGATCA

AGGGAGTGATAGTTCACAAGATAGAAGCTATAACATACAACATTGGGGCACAGGAGTGGTATACTACTATCCCTAAATAT

GTAGCCACTCAGGGGTATCTGATATCGAACTTTGATGAGACGTCATGCGTATTCACTCCAGAGGGGACAGTTTGCAGCCA

GAATGCGTTGTACCCAATGAGCCCATTGCTTCAGGAATGTTTCAGGGGGTCAACAAAATCGTGCGCCAGAACCTTAGTTT

CAGGGACCATAAGTAATAGATTTATCCTATCAAAAGGGAACCTGATTGCAAATTGTGCGTCAGTTTTGTGCAAATGTTAC

ACAACAGAGACAGTTATCAGCCAAGATCCTGACAAACTACTAACTGTTGTAGCACCCGACAAGTGTCCTGTAGTTGAGGT

GGATGGAGTGACAATACAGGTCGGCAGTCGAGAGTATCCGGATTCTGTATACTTACACAAAATAGACTTAGGTCCAGCCA

TCTCCCTAGAAAAACTGGATGTAGGCACCAATTTAGGCAATGCAGTCACAAGACTGGAGAATGCAAAGGAGCTCCTAGAT

GCATCAGACCAAATACTGAAGACTGTTAAAGGGGTACCTTTCAGTGGGAATATGTACATAGCACTGGCAGCTTGCATAGG

AGTATCCTTGGGCCTTGTCACATTAATATGCTGCTGTAAGGGGAGGTGTAAGAACAAGGAAATCCCTATCTCCAAAATCA

ACCCAGGGCTCAAACCCGACCTGACCGGGACCTCAAAGTCGTACGTAAGATCACTGTAGTCAGAATCACCTGGATCATCT

GGCATCGCACACATACATGCACGACACAGGCAGTCCGAGGACGCAAGAAACCCAGCCTCCGGTCACCCACCCGACCCCAC

TCCACGCTCCACCACACATTAGTCATCAAACAAAACTTAGGACGAAAGGTCAATCACCATGTCCGCACAAAGGGAGAGGA

TCAATGCCTTCTACAAAGACAATCCTCACAATAAGAACCATAGGGTGATCCTGGATAGGGAACGCTTGGTCATTGAAAGA

CCCTACATCTTGCTTGGAGTCCTGCTGGTAATGTTCCTGAGTCTAATCGGACTGCTGGCCATTGCAGGGATCAGGCTTCA

CCGGGCCACCGTTGGAACTTCAGAGATCCAGAGTCGGCTGAATACCAATATTAAGTTGACCGAATCTATTGATCACCAGA

CTAAGGATGTCTTAACTCCCCTTTTTAAAATCATTGGCGATGAAGTCGGCATCAGAATTCCACAGAAATTCAGTGATCTT

GTCAAGTTCATCTCCGATAAGATTAAATTCCTCAACCCTGATAGAGAGTATGATTTCAGAGATCTCCGGTGGTGTATGAA

TCCCCCCGAGAGAGTCAAAATTAATTTTGATCAGTTTTGTGAGTACAAGGCTGCGGTTAAGTCAATTGAACATATATTTG

AGTCACCACTCAACAAGTCAAAAAAGCTGCAATCTTTGACTCTCGGGCCCGGAACAGGCTGTCTAGGCAGGACAGTAACA

AGAGCCCATTTCTCAGAACTTACAATGACCTTAATGGACCTGGATCTAGAGATGAAGCACAACGTGTCCTCAGTGTTTAC

CGTAGTTGAAGAGGGATTATTCGGAAGAACATATACCGTCTGGAGATCCGATGCCAGGGATCCGAGCACCGATCTAGGTA

TCGGCCATTTTTTAAGAGTCTTCGAGATTGGACTGGTAAGAGATCTTGGGCTGGGTCCCCCTGTTTTTCATATGACCAAC

TATCTCACGGTGAACATGAGTGATGACTATCGGAGATGTCTTTTAGCGGTAGGGGAGCTGAAGTTGACAGCCCTATGCAC

CTCATCTGAGACTGTGACACTGAGTGAGAGAGGAGTTCCAAGGAGGGAACCTCTTGTGGTTGTGATACTTAATCTAGCTG

GACCCACTCTAGGGGGAGAGCTATACAGTGTCTTGCCTACCTCTGATCTCATGGTGGAGAAACTCTATTTGTCTTCACAT

AGAGGGATCATCAAAGATGATGAGGCCAATTGGGTAGTGCCGTCTACCGATGTTCGTGATCTTCAGAACAAAGGTGAATG

TCTGGTGGAAGCATGCAAGACTCGACCTCCTTCATTTTGCAATGGCACAGGATCAGGCCCGTGGTCAGAAGGGAGAATCC

CTGCCTACGGGGTGATCAGGGTCAGTCTTGACTTAGCGAGTGACCCGGATGTAGTTATCACTTCAGTGTTTGGCCCACTG

ATACCTCACCTATCCGGCATGGATCTTTACAACAACCCGTTTTCAAAAGCTGTATGGTTGGCTGTACCACCCTATGAGCA

GTCATTTCTAGGAATGATAAATACAATTGGATTCCCTAACAGAGCAGAGGTTATGCCGCACATTTTGACCACAGAGATCA

GAGGCCCTCGGGGTCGTTGCCATGTTCCCATAGAATTGTCCCGCCGGGTTGATGACGATATCAAGATCGGGTCCAACATG

GTCATATTGCCGACGATGGACCTGAGGTATATTACAGCCACTTATGATGTTTCCAGGAGCGAGCATGCAATCGTGTACTA

TATCTATGACACGAGTCGCTCATCATCTTACTTCTACCCAGTTCGACTGAATTTCAAAGGCAATCCTCTCTCTTTGAGGA

TAGAGTGTTTCCCCTGGCGTCATAAGGTGTGGTGCTACCATGATTGTCTTATATACAACACCATAACAGGTGAAGAGGTC

CATACGAGAGGGCTGACCGGCATAGAGGTAACATGCAATCCAGTCTGAGCAGAGCTACGACCGTCACTCAGGCAGTCCCT

TGAGTCGCCACCGAGTCCAAGCAGCACAGCCTGGGACAATCAACGGCACAGCCCAGCCAACAATGTTATAAAAAACTTAG

GAGCCAGGGTTGTAGGGGCCATGGACTCTCTATCAGTCAATCAGGTCTTGTACCCTGAGGTTCATCTAGACAGCCCTATT

GTCACAAACAAACTAGTTGCCATCCTTGAGTACTCGGGTATCGACCATAACTATGTTCTTGAAGACCAGACTCTTGTCAA

GAATATTAGGTATAGGCTGGGGTGCGGTTTTTCAAATCAAATGATCATCAATAACAGGGGGGTGGGTGAAACAGTCAATT

CTAAACTTAAAAGTTACCCCCATAATCGTCATATCATATACCCGGATTGCAATAAGGAGTTGTTTTGTATCAAGGATAGC

TGCATATCTAAGAAGCTCTCGGAGCTATTCAAGAAGGGTAATTCCTTGTACTCTAAGATAAGTCACCAGGTACTGGATTG

TCTTAAGAGAGTCAATGGGAAATTAGGTCTGGGCACAGATCTTACCCATGGTCTGAAGGAGGGTATCCTTGACCTAGGGT

TGCACATGCATAGCTCTCAATGGTTCGAGACTTTCCTGTTCTGGTTCACTATCAAGACAGAGATGAGATCAATGATCAAA

GAACAGTCCCATATATGCCACAAGAGGAGGTATAACCCGACTTTTGTGTCGGGGGATGCATTCGAGGTGCTCGTATCGCG

AGACCTCGTTGTGATAATTGATAAGAATACCCAGTATGTCTTCTACTTAACATTTGAGCTGGTCCTTATGTATTGTGATG

TCATAGAGGGTAGACTTATGACGGAGACAGCCATGGCCATAGACCAGAGATATTCAGAACTTCTAAGCCGGGTCAGATAC

TTGTGGGATCTTATTGATGGGTTCTTCCCAACACTAGGCAACACCACATACCAAATTGTTGCTCTGCTTGAACCATTATC

GTTGGCTTATCTTCAACTTCAGGATGTCACTCTGGAGTTAAGAGGTGCTTTTTTAGACCATTGCTTCAAAGAACTCTATG

AGATACTGGAACATTGCGGCATTGACACAGCAGGCACTTACAATTCCATCACTGAAGGATTGGATTACGTATTCATCACC

CACGATATACATCTAACTGGGGAGATTTTTTCATTTTTCCGGAGTTTCGGACACCCCCGTCTAGAAGCAGTCACCGCTGC

GGAAAATGTCAGAAAACATATGAACCAACCGAAGGTAATCAGTTATGAGACTATGATGAAAGGCCATGCGGTATTTTGCG

GGATAATCATAAATGGTTTTCGGGACCGACATGGCGGCAGCTGGCCCTCTGTTGCACTGCCAGAACATGCCTCTGCTGCG

ATCCGGAATGCGCAGGCATCAGGTGAGGGACTGACCCATGACCTGTGTATAGACAACTGGAAATCCTTTGTTGGATTCAA

ATTTGGCTGCTTTATGCCACTCAGCCTAGATAGTGATTTGACCATGTATCTCAAAGACAAGGCATTGGCTGCACTGAAGA

ATGAGTGGGATTCAGTTTACCCGAAAGAATACCTCCGTTATAACCCACCTAGAGGGACAGAGTCAAGGCGATTGGTAGAG

GTATTCCTGAATGACTCCAGCTTTGATCCTTATAACATGATAATGTACGTGGTGAATGGCTCCTACCTTAAAGACCCTGA

GTTTAACCTCTCATACAGCCTAAAAGAGAAGGAGATAAAGGAGACAGGGCGGTTGTTTGCCAAAATGACCTATAAGATGC

GGGCCTGTCAGGTAATCGCTGAAAATCTGATATCGAATGGTGTTGGGAAGTATTTCCGGGACAATGGGATGGCAAAGGAC

GAGCATGACCTAACAAAAGCCCTTCACACCCTGGCAGTCTCAGGTGTTCCCAAAAACAACAAAGATAACCACCGAGGTGG

GCCTCCCAGAAGGACAACAAACCGAGGGGTGAGATCAAGCCAAGGCACCAAAACACAAGATAGAGACAAGGTTCAAGGGG

GACCTATGTACAACTATTTGCGATGCCAACCGATCAGCCTTGATCAGGGTGAGTCATACGAGACTGTTAGTGCATTCATC

ACCGCTGACCTTAAGAAGTATTGCCTGAATTGGAGATACGAGACAATCAGCATATTTGCACAGAGGCTGAATGAAATATA

TGGACTACCATCCTTCTTTCAATGGTTACACAGGATATTGGAGAAATCCGTACTCTACGTCAGTGACCCACATTGTCCTC

CCGATCTAGATAATCATATCCCTCTGGACAGTGTCCCCAATGCCCAAATATTCATCAAGTACCCAATGGGCGGAATAGAA

GGTTATTGCCAAAAACTATGGACAATCAGCACTATACCATACTTGTATCTGGCAGCCTATGAGAGCGGAGTAAGAATCGC

CTCACTGGTGCAGGGTGACAATCAGACAATCGCAGTGACAAAAAGAGTTCCAAGTTCTTGGCCTTATTCACTAAAAAAGA

GGGAGGCCTCCAAGGCCGCTCAAAATTACTTCGTAGTCTTAAGGCAAAGATTGCACGATGTAGGTCATCACTTAAAGGCT

AATGAGACCATAGTATCTCCTCACTTTTTTGTATATTCCAAAGGGATTTATTATGACGGCCTGCTAGTCTCACAATCACT

AAAGAGCATCGCCAGATGTGTCTTCTGGTCCGAGACTATAGTGGATGAAACCAGAGCGGCGTGCAGTAATATTGCAACAA

CTGTCGCTAAGAGTATAGAGAGGGGTTATGATAGGTACCTTGCATATTCTTTGAATATCCTCAAGATTTTCCAACAGATC

CTTATATCCCTCAACTTCACTATTAACACAACAATGACTCAGGATGTCGTGGCACCGATCATCGAGAACGGTGATTTGCT

GATAAGGATGGCACTCTTGCCAGCACCCATCGGGGGTCTCAATTATCTTAACATGAGCAGGTTATTTGTGAGAAATATCG

GTGACCCGGTCACCTCCTCTATAGCCGACCTGAAGAGGATGATAGATGCCGGGCTAATGCCAGAAGAAACATTGCATCAA

GTGATGACCCAGACCCCGGGAGAATCATCCTACCTAGATTGGGCAAGTGACCCTTATTCTGCCAACCTACCCTGCGTACA

GAGTATCACTCGCCTTCTCAAAAACATCACTGCACGATATATTTTAATCAGCAGCCCAAACCCGATGCTGAAAGGGTTAT

TTCATGAGGGGAGTAGAGATGAAGACGAGGAGCTTGCAAGTTTCCTAATGGATCGGCATATAATTGTTCCAAGAGCTGCA

CATGAAATCTTAGACCATAGTATAACCGGAGCAAGAGAAGCTATAGCCGGGATGTTGGACACCACCAAGGGTCTGATTAG

AACAAGTATGAAACGGGGCGGCCTCACCCCTCGAGTCTTAGCCCGCCTTTCCAATTATGATTATGAACAATTCCGATCCG

GGATAACACTATTAACAAAGAAAGGGCAGTGTTATCTCATTGACAAGGACTCATGCTCGGTGCAGCTCGCTATAGCTCTG

AGGGGCCACATGTGGGCTAAGTTGGCTCGCGGGAGACCAATCTATGGGTTGGAGGTGCCTGATGTATTAGAATCGATGAA

CGGCTACCTTATCAAACGCCACGAGTCGTGTGCTATCTGTGAAACGGGCTCAAGTCACTACGGATGGTTTTTCGTCCCTG

CAGGGTGCCAGCTTGACGATGTATCTAGAGAGACTTCGGCTCTTCGTGTACCTTATGTCGGATCAACAACTGAGGAAAGG

ACAGACATGAAACTTGCATTTGTTAGGTCTCCAAGTAGATCCCTCAAATCAGCAGTTAGAATTGCAACAGTTTACTCATG

GGCTTACGGGGATGATGAGAAATCATGGAGTGAAGCCTGGATGCTAGCTAGGCAGAGAGCCAATATCACCTTAGATGAAT

TGAGAATGATCACTCCAGTCTCCACATCCACCAACTTAGCCCACCGGTTGAGGGATCGGAGTACCCAGGTGAAATACTCG

GGGACATCCCTTGTGAGGGTTGCAAGATACACAACCATCTCCAATGATAACTTGTCGTTTGTGATATCTGAGAAAAAAGT

GGATACCAACTTCATTTATCAGCAAGGGATGTTGCTTGGTCTTGGGATCCTTGAGAACCTCTTTAGGCTAGAGGCCACCA

CCGGAGTATCCAACACAGTGCTACACCTGCACGTGGAAACAGAATGTTGTGTTGTACCTATGGTGGATCACCCAAGGATA

CCGAGTCTCCGTAATATAAAAGTTACGGATGAGCTATGCACAAACCCTTTGATCTACGACAGGTCCCCCATCATAGAGCA

CGATGCAACCCGGCTATATTCACAGAGCCACAGGAGACATTTGGTGGAGTTTGTTACCTGGTCAACAAGCCAGCTTTACC

ATATACTGGCTAAGTCTACAGCAATGTCCATGATTGAATTGATCACGGGATTCGAGAAAGATCACATGAATGAAATAGCC

GCCCTGATTGGCGATGACGACATCAACAGCTTCATCACAGAATTTTTGCTAGTGGAGCCTAGATTATTTATAGTTTACCT

TGGTCAGTGTGCTGCCATCAATTGGGCCTTTGATATACATTATCATCGGCCCTCGGGCAAGTACCAGATGGGGGAACTTC

TCTACTCTCTGCTCTCTCGGATGAGCAAAGGAGTATATAAGATCTTTACTAATGCTTTGAGCCACCCCAAAGTTTACAAG

AAATTTTGGCGAAGTGGCATCATTGAGCCGATTCATGGCCCATCCTTGGATACACAGAATTTACATGTCACTGTCTGTGA

CATGATATATGGGTCCTATGTCACCTATTTGGATCTTTTGCTGAATGATGAGCTAGATGATTACCCGTATTTGCTCTGCG

AGAGTGATGAGGACGTGGTCACAGACAGGTTCGACAACATTCAAGCCAAACATCTCTGTGTATTGGCCGATGTATATTGC

AGCTTCAAGAGGTGTCCTTCAATAATCGGGATGTCTCCTATAGAAAAATGTACCATTCTCACACATTACATCAAGGGAGA

ATCAGTACAATCCCCGTCTGGGACCTCATGGAACACTGATCCCCTTGTAGTAGATCATTACTCATGCTCTCTGACCTACC

TTCGCCGCGGTTCCATCAAACAAATCAGGTTGAGGGTGGATCCTGGGTTTGTACTCGAGGCGTTGACAGACGTCGACTTT

AAACAACCTCGCAAAGCTAAGTTAGATATATCGGTCGTTGGGTCGACTGATTTCTCTCCCCCTTACGACAACGTCGGTGA

TTTTTTAGGGACTATCAACACATTGAGGCACGATCTGCCTGTCACCGGAACCGGGGTCTCGAACTATGAAGTCCACGCTT

ATCGTAGAATTGGCCTGAATTCATCAGCATGTTATAAAGCCGTAGAGATCTCCACGTTAATCAAGCCATCCTTAGAAGTC

GGAGAGCATGGATTGTTCTTAGGAGAAGGTTCCGGTTCAATGCTGGCTGCGTACAAAGAAGTTCTTAAATTAGCAAATTG

TTACTACAACAGCGGAGTAACAGCGGAGGGCAGAGCCGGACAGAGGGAAATATCTCCCTATCCTTCAGAGATGAGCCTAG

TAGAGAATCAAATGGGGATAGAGCGGAGTGTTAAAGTGCTGTTCAACGGGAAACCTGAAGTAACTTGGGTGGGGACCACA

GATTGCTACAAGTACATAATCAGTAACATCCAAACCTCTAGTCTGGGTTTCATACATTCAGATATTGAGACGCTCCCAAC

CAAAGATGCTGTTGAGAAATTGGAGGAATTTGCCTCTATCCTATCCTTATCCTTGATTTTGGGGAAAATCGGCTCTATTA

CAGTTGTCAAAATTATGCCCATTAGCGGGGATTTTACCCAAGGCTTCATAGCCTATGCCATCCAATATTTTAGGGAGAGC

CTGCTTGCCTACCCGAGATATAGTAACTTCATCTCGACTGAGTGTTACCTTATTATGATAGGATTAAAGGCCAATCGATT

GATAAACCCAGAAGCCATTAAGCAAAGCATAATCAGAGCGGGGGTCAGGACTGCACCAGGACTTGTGAGCCATGTATTAT

CAGAGAAACAAAAAGGTTGTATTCAATCTTTCTTGGGTGATCCTTATATCCAAGGAGACTTCAATAAGCACCTTAAATCT

CTAACCTCTATTGAGAAAATCCTGGTAAATTGTGGTCTCTCGATCAATGGCATAAAAATCTGTAGAGATCTAATCCACCA

TGATATCGCCTCCGGTCCAGACGGCCTTATGAGCTCTACAATTATTTTATATAGGGAATTGGCTCATTTCAAAGATAATA

TAAGGAGTCAGCACGGAATGTTCCACCCCTACCCGGTACTGGCCAACAGCAGGCAACGTGAATTAATCCTTCGAATAGCC

AAGAAATTCTGGGGGTATGTCTTGCTATATTCTGATGACCCGGCACTAATCAGACAAACAATCAAGAACTTGAAACGGAA

TCATCTAACCTTTGACTTACATAGTAACCCATTTATTAAAGGGCTATCCAAGGCTGAGAAACTGCTGGTGCGGACGAGTT

CACTAAGAAGAGAATGGTTGTTCACTCTCGAAACGAAAGAAGTAAAAGAGTGGTTCAAATTGGTAGGTTACAGTGCACTC

ATCAGAGGCTAGTCAGCTATGCATCTGCCCCCTTCTCCTCCGCCATGAGACCCCACTGGCGATCCAGAAGATTAAAGAAA

ACTACATATTGGATAAGTATCTATTCCCAGCTTTGTCTGGT

>KX354359.1/goat/China/2015-12-15

ACCAAACAAAGTTGGGTAAGGATAGATCTTATAATAACTATGGACTGGCAAACTTAGGAGTAAAGATCCTACTGTCGGGG

GGAGGAGGAGGAGCAAGATCTTTGATTATGGCGACTCTCCTTAAAAGCTTAGCATTGTTCAAAAGGAACAAAGACAAAGC

GCCGACAGCATCAGGTTCAGGAGGGGCCATCCGGGGGATTAAGAATGTTATCATAGTCCCGATTCCCGGAGACTCGTCCA

TCACTACCCGTTCAAGACTGCTCGACAGGCTTGTCAGATTGGCCGGAGATCCTGACATCAACGGGTCAAAGCTGACCGGG

GTGATGATCAGCATGTTATCCTTGTTCGTAGAGTCACCCGGGCAATTGATACAGCGAATCACAGATGATCCGGATGTTAG

TATCCGCCTTGTTGAGGTAGTTCAAAGTACTAGGTCCCAATCCGGGTTGACCTTTGCATCACGTGGTGCTGATTTAGACA

ACGAGGCAGACATGTATTTTTCAACTGAGGGGCCCTCGAGTGGAGGTAAGAAAAGGATCAACTGGTTTGAGAACAGAGAA

ATAATAGACATAGAGGTGCAGGATCCAGAAGAGTTCAATATGTTGTTAGCCTCCATACTAGCACAAGTCTGGATCCTTCT

GGCCAAGGCTGTTACGGCACCAGATACGGCAGCTGACTCAGAACTGAGAAGGTGGGTTAAATACACACAACAAAGGAGAG

TGATTGGGGAATTTCGCCTTGACAAGGGGTGGCTGGACGCAGTCCGCAACAGAATTGCAGAAGATCTATCACTCCGGCGG

TTCATGGTATCTCTCATACTTGACATCAAGAGGACCCCAGGCAACAAGCCAAGGATTGCAGAAATGATCTGCGACATTGA

CAACTATATTGTCGAAGCAGGACTCGCCAGTTTCATCCTTACTATCAAATTTGGTATCGAAACCATGTATCCTGCATTAG

GGCTCCACGAGTTTGCCGGAGAATTGTCCACTATTGAGTCCTTGATGAACTTGTATCAACAGCTAGGAGAGGTTGCACCC

TACATGGTAATTCTAGAGAATTCAATTCAGAACAAGTTTAGTGCAGGAGCTTATCCCCTCCTCTGGAGCTATGCGATGGG

TGTCGGAGTCGAGCTGGAGAACTCAATGGGGGGCTTGAACTTTGGCAGGTCATATTTTGACCCTGCCTATTTTCGTCTCG

GACAGGAGATGGTCAGAAGATCTGCAGGAAAGGTCAGCTCTGTGATTGCGGCTGAGCTCGGCATCACAGCAGAGGAAGCT

AAACTAGTCTCGGAAATCGCCTCGCAGGCTGGGGACGAAAGAACCGCCAGAGGGACTGGGCCTCGACAGGCGCAGGTTTC

CTTCCTTCAGCACCAAACAGGAGGGGGAGAGTCGTCCGCACCAGCGACCAGAGAAGGGGTCAAAGCTGCGATCCCAAACG

GAGCTGAAGAAAGGGACAGAAAGCAAACACGCCCAGGAAGGCCCAGAGGAGAGACCTCCGGCCAACTGCTCCTGGACATC

ATGCCAGAGGATGAGATCTTGCGAGAGTCTGGTCAAAACCCTCGTGAGGCTCAAAGATCGGCCGAGGCACTCTTCAGGCT

GCAGGCCATGGCCAAGATTCTGGAGGACCAGGAGGAGGGAGAAGACAACAGTCAGGTCTACAACGACAAGGATCTCCTCG

GCTAAGCAGACGCACCCTCTGTCGAAATCAGTGACAAGACATCTCCTACCAGTATTATAAAAAACTTAGGACCCAGGTCC

AAGCAAGCACACATCGACACTCCAACCAGTCGAGCGGAGACCACCGATGGCAGAAGAACAAGCATACCATGTCAACAAGG

GGCTGGAATGTATCAAGTCTCTCAAAGCCTCACCCCCGGATCTATCCACCATCAGAGATACCATCGAGAGCTGGAGAGAG

GGGCTTAGCCCCTCGGGCCGTGCAACACCGAACCCTGATACGTCCGAGGGAGATCATCAGAATATCAACCAATCATGCTC

ACCAGCAATCGGACCAAACAAAGTCTACTTGTCTCCTGAAGATAATCTCGGATTTAGAGAGATCACTGGCAACGACTGTG

AGGCTGGGCTCGGAGGGGTCCAGAGAGAAGGATCCAACTCTCAAGTACAGCGTTACCATGTTTATAGTCACGGGGGTGAA

GAGATTGAAGGACTCGAGGATGCTGACTCTCTCGTGGTTCAAGCAGATCCTCCAGTTGCTAACGTCTTCAATGGAGGAGA

GGATGGATCTGACGACAGCGATGTGGACTCTGGCCCAGATGATCCCGGCAGAGATACTCTATATGACCGGGGATCTGTTG

CCGGCAATGGTATCGCTAGGTCCACAGATGTCGAAAAACTAGAAGGTGCTGATATTCAAGAAGTTCTTAACTCCCAGAAA

GGCAAAGGAGGAAGATTCCAGGGCGGGAAAACCTTGCGAGTCCCGGAAATACCCGATGTCAAGCACTCCAGACCATCAGC

CCAATCAATTAAAAAGGGCACAGACGGGAACTCAGTCTCATCTGGAACGGTGATAGAGTGTTTATCGATAAGTGGTGCAA

CCCAAACTGTGCCAGAGTCAAGATGGGAGTCATCAGAGCAAAATGCGTCTGTGGGGAGTGTCCTCAAGTCTGCGAGGAGT

GCAAAGACGATCCAGGGGTCGACACAAGAATCTGGTACCATAGCATCACTGACTCAGCCTAAAGAGAATGACTCCGAGTA

TGAGTATGAGGATGACCTATTTACAGAGATTCAGGACATCCGTGCAAGCATTGCCAAGATCCATGATGACAATAAAACTA

TCCTCTCAAAGCTTGATTCTATACTGTTATTGAAAGGGGAAGTCGACACTATCAAGAAACAAATCAGCAAGCAGAATATA

AGTATATCCACCATTGAGGGCCATCTCTCCAGTATAATGATAGCCATCCCTGGCTTTGGGAAGGACATCAAGGACCCAAC

ATCCGAAGTCGAGTTGAACCCAGATCTAAGACCTATAATAAGCCGTGACTCTGGCAGAGCTCTCGCGGAGGTCCTCAAGA

AACCCGCTGTGGATAGGTCTCAGAAAATTGGAACCAAAGCCAACTCCAGCTCAAAGGGTCAGCTTCTTAAGGATCTCCAG

CTAAAACCTGTCGACAAGCAGGCAAGCTCTGCAATCGGGTTTGTCCCGTCCGACCATGAATCATCCAGAAATGTCATCCG

CTCCATAATCAAGTCGAGCAAGCTAAATATTGATCACAAGGACTATCTTCTAGATTTACTAAATGATGTGAAAGGCTCCA

AGGATCTTAAGGAATTTCACAAGATGCTAACAGCAATTCTCGCCAAGCACCCGTAACACAYCCTCCAGTCATCATCTCAT

ACTCGACTAAAAACATCCTTTCAATCAGGCTATTACAAAAAACTTAGGAGCAAGGGCAACTGAGCTTCGCAGACAGGATG

ACCGAGATCTACGACTTCGATAAATCAGCATGGGATGTCAAAGGGTCAATTGCCCGCATAGAACCCACCACCTACCACGA

CGGCCGACTGATACCCCAGGTGAGGGTCATCGATCCTGGTCYGGGAGACAGAAAAGATGAGTGCTTTATGTACCTGTTTC

TCCYAGGAGTGATTGAGGATAACGACCCCCTGTCTCCCCCCGTCGGAAGAACCTTTGGCTCTCTACCTCTAGGGGTTGGT

AGGTCAACTGCCAAGCCAGAAGAACTACTAAGGGAGGCCACAGAATTAGATATAGTGGTGAGGCGCACTGCAGGAGTAAA

TGAGAAACTGGTATTTTACAACAACACTCCGCTGTCCTTGTTAACGCCCTGGAAGAAAGTTCTGACAACCGGAAGTGTGT

TTAGCGCTAACCAGGTTTGCAATGCAGTCAACCTAGTCCCGCTTGATACTCCCCAGAGATTCAGGGTTGTGTACATGAGC

ATAACTAGATTATCAGACAATGGTTACTATAGTGTGCCCAGAAGAATGCTGGAGTTCCGCTCGGCCAATGCAGTAGCCTT

CAACATCTTGGTTACACTAAGAATTGAAAATGGCACAAACCCTAGAAGATACATAGTCGGCTCATGGGAGAATTCAGAGG

TCACATTTATGGTACACGTGGGCAACTTTAGGAGAAAGAAGAACGAAGTATACTCTGCTGATTACTGCAAAATGAAGATT

GAAAAGATGGGTTTAGTTTTTGCCCTGGGTGGAATAGGCGGAACAAGTCTCCATATTAGAAGCACCGGGAAAATGAGCAA

AACCCTCCATGCACAGCTGGGGTTCAAGAAAATCTTATGTTACCCCCTAATGGATGTCAATGAGGATCCTAACCGATATC

TCTGGCGGGCAGAGTGCCGAATAGTCAAAATCCAAGCTGTCTTACAGCCATCAGTACCCCAAGAGTTCCGTGTCTACGAT

GATGTCATCATCAACGATGATCAAGGCTTGTTCAAGATCTTGTAGTTCATTTGCAAAGGAACGAACCCACCCCCAGGAAG

GGGCCCCACCC-CCCCCG-CCCAAAAACACACTCCAAGGCTCCCCGGCACAGCGCCAACACAGCTACTCCCGCGATGGAA

CGAGGTCGGCC-CCCGAACAAATCCGGTCCAGGGGGGGGGGCCCCCCCCCTGACCAACCGGGCGCCGACCGAGCAGGCCG

GCGAGAAGCCCCCAAGAGGCCCCGACGAAGACCGAGCACCCCATCCCCCC-AAAAACGTCCCCC-AACACGCCCGAGGCT

AGCCACCGGACACCCCCAGACCCGCCACCGGCGAGACAGAACCCCAAGGGGGGACCCTCCAAACCAAAGGAATCACCGGG

GTAGGCTCCGCCGAACCAGCACAGACCCGACCCAAGCAAGGCGGACCCCCAACACGAAACTGCCCAGCACACCCTACTCA

TCATCAAACAAAACTTAGGGGCCAAGTCCATACGAGAGGCCCCAGTCCGTCAGGCGGAGGGACGCAGAAAGGAAGGAGAC

ACGACCGTCCCAGAAGGAGGACACCCAACCACCGAAACGCCCGCCCCCAGGGAAGGAG--CCACCCCCCACGCAGCGAAG

GAGCGGGTGGGAGACCTCGGGGAGGGCCCGACCGGGACCCGGGGGCCGGCGGCGCCGGCGGTGAGCAGGCCGGGCGCATA

CCTGACCCCACCGGAGGATCCGAGGAGGGCCCGACCCCGCGTGGACCAAACGGCAGGGAGCCCAACCCCACCAAAAAAAG

AGGCGAGGCCCCC-------CCCCCCCCGCACACAACCCGCAGACAGACCGAGGGAGGGCCATCCCCCAGCGAGACAGGC

GGCTCCCTCGCAGCCGAGCCGCAGCTCCCCAGAATAGGACACCCCCGCACACTCCGAACACCCCCTCGCCCCCCGCCACC

GGGACAGGGGCTCCACCACCACAGACCGGAGCCACCGACAGCCAGCCAGAGGTCCCCTCCAGTCCCGCCCGGCGCCCCCC

CGAAGAGGCCCCGGCAAGGCCCCCCAGCCCGACGTCCCCCGCACGCACCGAAAGAAGGACCCGCCCGAGAACGTSCGGCA

CATCCATGYGYAAACATCATGACGCGGGTCGCAATTTTGACATTTCTGTTTCTTTTCCCAAATGTTGTTGCGTGTCAGAT

TCACTGGGGCAATCTATCCAAGATCGGGATTGTAGGAACAGGGAGTGCCAGCTACAAGGTGATGACTAGGCCAAGCCACC

AGACTCTGGTTATAAAGTTAATGCCAAATATAACGGCCATCGACAATTGTACAAAGTCAGAGATTGCAGAGTACAAGAGA

TTGCTGATCACAGTGTTAAAGCCTGTAGAGGATGCTCTGTCGGTGATAACCAAGAATGTAAGACCAATTCAAACTCTAAC

ACCTGGGCGTAGAACCCGCCGTTTTGCTGGAGCTGTTCTGGCCGGGGTAGCACTTGGAGTTGCGACAGCCGCTCAGATAA

CTGCAGGAGTCGCCCTTCATCAATCATTGATGAACTCCCAAGCAATTGAGAGTTTAAAAACCAGTCTTGAGATGTCGAAT

CAGGCAATAGAAGAAATCAGACTTGCAAATAAGGAGACCATACTGGCAGTACAGGGCGTCCAGGATTATATCAACAATGA

GCTCGTCCCTTCTGTTCATAGAATGTCATGCGAGCTGGTAGGTCACAAGCTCGGCCTCAAGCTCCTTAGGTACTACACCG

AGATCCTGTCCATATTCGGGCCCAGTCTTCGAGACCCGATATCTGCCGAAATATCAATCCAGGCACTTAGTTATGCATTA

GGCGGAGACATTAATAAAATCCTGGACAAGCTTGGGTATAGCGGTGGGGATTTCCTTGCCATCCTAGAAAGCAAGGGAAT

AAAGGCCCGGGTTACATATGTGGACACAAGAGATTACTTTATAATCCTTAGCATCGCCTACCCAACCTTATCTGAGATCA

AGGGAGTGATAGTTCACAAGATAGAAGCTATAACATACAACATTGGGGCACAGGAGTGGTATACTACTATCCCTAAATAT

GTAGCCACTCAGGGGTATCTGATATCGAACTTTGATGAGACGTCATGCGTATTCACTCCAGAGGGGACAGTTTGCAGCCA

GAATGCGTTGTACCCAATGAGCCCATTGCTTCAGGAATGTTTCAGGGGGTCAACAAAATCGTGCGCCAGAACCTTAGTTT

CAGGGACCATAAGTAATAGATTTATCCTATCAAAAGGGAACCTGATTGCAAATTGTGCGTCAGTTTTGTGCAAATGTTAC

ACAACAGAGACAGTTATCAGCCAAGATCCTGACAAACTACTAACTGTTGTAGCATCCGACAAGTGTCCTGTAGTTGAGGT

GGATGGAGTGACAATACAGGTCGGCAGTCGAGAGTATCCGGATTCTGTATACTTACACAAAATAGACTTAGGTCCAGCCA

TCTCCCTAGAAAAACTGGATGTAGGCACCAATTTAGGCAATGCAGTCACAAGACTGGAGAATGCAAAGGAGCTCCTAGAT

GCATCAGACCAAATACTGAAGACTGTTAAAGGGGTACCTTTCAGTGGGAATATGTACATAGCACTGGCAGCTTGCATAGG

AGTATCCCTGGGCCTTGTCACATTAATATGCTGCTGTAAGGGGAGGTGTAAGAACAAGGAAATCCCTATCTCCAAAATCA

ACCCAGGGCTCAAACCCGACCTGACCGGGACCTCAAAGTCGTACGTAAGATCACTGTAGTCAGAATCACCTGGATCATCT

GGCATCGCACACATACATGCACGACACAGGCAGTCCGAGGACGCAAGAAACCCAGCCTCCGGTCACCCACCCGACCCCAC

TCCACGCTCCACCACACATTAGTCATCAAACAAAACTTAGGACGAAAGGTCAATCACCATGTCCGCACAAAGGGAGAGGA

TCAATGCCTTCTACAAAGACAATCCTCACAATAAGAACCATAGGGTGATCCTGGATAGAGAACGCTTGGTCATTGAAAGA

CCCTACATCTTGCTTGGAGTCCTGCTGGTAATGTTCCTGAGTCTAATCGGACTGCTGGCCATTGCAGGGATCAGGCTTCA

CCGGGCCACCGTTGGAACTTCAGAGATCCAGAGTCGGCTGAATACCAATATTAAGTTGACCGAATCTATTGATCACCAGA

CTAAGGATGTCTTAACTCCCCTTTTTAAAATCATTGGCGATGAAGTCGGCATCAGAATTCCACAGAAATTCAGTGATCTT

GTCAAGTTCATCTCCGATAAGATTAAATTCCTCAACCCTGATAGAGAGTATGATTTCAGAGATCTCCGGTGGTGTATGAA

TCCCCCCGAGAGAGTCAAAATTAATTTTGATCAGTTTTGTGAGTACAAGGCTGCGGTTAAGTCAATTGAACATATATTTG

AGTCACCACTCAACAAGTCAAAAAAGCTGCAATCTTTGACTCTCGGGCCCGGAACAGGCTGTCTAGGCAGGACAGTAACA

AGAGCCCATTTCTCAGAACTTACAATGACCTTAATGGACCTGGATCTAGAGATGAAGCACAACGTGTCCTCAGTGTTTAC

CGTAGTTGAAGAGGGATTATTCGGAAGAACATATACCGTCTGGAGATCCGATGCCAGGGATCCGAGCACCGATCTAGGTA

TCGGCCATTTTTTAAGAGTCTTCGAGATTGGACTGGTAAGAGATCTTGGGCTGGGTCCCCCTGTTTTTCATATGACCAAC

TATCTCACGGTGAACATGAGTGATGACTATCGGAGATGTCTTTTAGCGGTAGGGGAGTTGAAGTTGACAGCCCTATGCAC

CTCATCTGAGACTGTGACACTGAGTGAGAGAGGAGTTCCAAGGAGGGAACCTCTTGTGGTTGTGATACTTAATCTAGCTG

GACCCACTCTAGGGGGAGAGCTATACAGTGTCTTGCCTACCTCTGATCTCATGGTGGAGAAACTCTATTTGTCTTCACAT

AGAGGGATCATCAAAGATGATGAGGCCAATTGGGTAGTGCCGTCTACCGATGTTCGTGATCTTCAGAACAAAGGTGAATG

TCTGGTGGAAGCATGCAAGACTCGACCTCCTTCATTTTGCAATGGCACAGGATCAGGCCCGTGGTCAGAAGGGAGAATCC

CTGCCTACGGGGTGATCAGGGTCAGTCTTGACTTAGCGAGTGACCCGGATGTAGTTATCACTTCAGTGTTTGGCCCACTG

ATACCTCACCTATCCGGCATGGATCTTTACAACAACCCGTTTTCAAAAGCTGTATGGTTGGCTGTACCACCTTATGAGCA

GTCATTTCTAGGAATGATAAATACAATTGGATTCCCTAACAGAGCAGAGGTTATGCCGCACATTTTGACCACAGAGATCA

GAGGCCCTCGGGGTCGTTGCCATGTTCCCATAGAATTGTCYCGCMGGGTTGATGACGATATCAAGATCGGGTCCAACATG

GTCATATTGCCGACGATGGACCTGAGGTATATTACAGCCACTTATGATGTTTCYAGGAGCGAGCATGCAATCGTGTACTA

TATCTATGACACGAGTCGCTCATCATCTTACTTCTACCCAGTTCGACTGAATTTCAAAGGCAATCCTCTCTCTTTGAGGA

TAGAGTGTTTCCCCTGGCGTCATAAGGTGTGGTGCTACCATGATTGTCTTATATACAACACCATAACAGGTGAAGAGGTC

CATACGAGAGGGCTGACCGGCATAGAGGTAACATGCAATCCAGTCTGAGCAGAGCTACGACCGTCACTCAGGCAGTCCCT

TGAGTCGCCACCGAGTCCAAGCAGCACAGCCTGGGACAATCAACGGCACAGCCCAGCCAACAATGTTATAAAAAACTTAG

GAGCCAGGGTTGTAGGGGCCATGGACTCTCTATCAGTCAATCAGGTCTTGTACCCTGAGGTTCATCTAGACAGCCCTATT

GTCACAAACAAACTAGTTGCCATCCTTGAGTACTCGGGTATCGACCATAACTATGTTCTTGAAGACCAGACTCTTGTCAA

GAATATTAGGTATAGGCTGGGGTGCGGTTTTTCAAATCAAATGATCATCAATAACAGGGGGGTGGGTGAAACAGTCAATT

CTAAACTTAAAAGTTACCCCCATAATCGTCATATCATATACCCGGATTGCAATAAGGAGTTGTTTTGTATCAAGGATAGC

TGCATATCTAAGAAGCTCTCGGAGCTATTCAAGAAGGGTAATTCCTTGTACTCTAAGATAAGTCACCAGGTACTGGATTG

TCTTAAGAGAGTCAATGGGAAATTAGGTCTGGGCACAGATCTTACCCATGGTCTGAAGGAGGGTATCCTTGACCTAGGGT

TGCACATGCATAGCTCTCAATGGTTCGAGACTTTCCTGTTCTGGTTCACTATCAAGACAGAGATGAGATCAATGATCAAA

GAACAGTCCCATATATGCCACAAGAGGAGGTATAACCCGACTTTTGTGTCGGGGGATGCATTCGAGGTGCTCGTATCGCG

AGACCTCGTTGTGATAATTGATAAGAATACCCAGTATGTCTTCTACTTAACATTTGAGCTGGTCCTTATGTATTGTGATG

TCATAGAGGGTAGACTTATGACGGAGACAGCCATGGCCATAGACCAGAGATATTCAGAACTTCTAAGCCGGGTCAGATAC

TTGTGGGATCTTATTGATGGGTTCTTCCCAACACTAGGCAACACCACATACCAAATTGTTGCTCTGCTTGAACCATTATC

GTTGGCTTATCTTCAACTTCAGGATGTCACTCTGGAGTTAAGAGGTGCTTTTTTAGACCATTGCTTCAAAGAACTCTATG

AGATACTGGAACATTGCGGCATTGACACAGCAGGCACTTACAATTCCATCACTGAAGGATTGGATCACGTATTCATCACC

CACGATATACATCTAACTGGGGAGATTTTTTCATTTTTCCGGAGTTTCGGACACCCCCGTCTAGAAGCAGTCACCGCTGC

GGAAAATGTCAGAAAACATATGAACCAACCGAAGGTAATCAGTTATGAGACTATGATGAAAGGCCATGCGGTATTTTGCG

GGATAATCATAAATGGTTTTCGGGACCGACATGGCGGCAGCTGGCCCCCTGTTGCAYTGCCAGAACATGCYTCTGCTGCG

ATCCGGAATGCGCAGGCATCAGGTGAGGGACTGACCCATGACCTGTGTATAGACAACTGGAAATCCTTTGTTGGATTCAA

ATTTGGCTGCTTTATGCCACTCAGCCTAGATAGTGATTTGACCATGTATCTCAAAGACAAGGCATTGGCTGCACTGAAGA

ATGAGTGGGATTCAGTTTACCCGAAAGAATACCTCCGTTATAACCCACCTAGAGGGACAGAGTCAAGGCGATTGGTAGAG

GTATTCCTGAATGACTCCAGCTTTGATCCTTATAACATGATAATGTACGTGGTGAATGGCTCCTACCTTAAAGACCCTGA

GTTTAACCTCTCATACAGCCTAAAAGAGAAGGAGATAAAGGAGACAGGGCGGTTGTTTGCCAAAATGACCTATAAGATGC

GGGCCTGTCAGGTAATCGCTGAAAATCTGATATCGAATGGTGTTGGGAAGTATTTCCGAGACAATGGGATGGCAAAGGAC

GAGCATGACCTAACAAAAGCCCTTCACACCCTGGCAGTCTCAGGTGTTCCCAAAAACAACAAAGATAACCACCGAGGTGG

GCCTCCCAGAAGGACAACAAACCGAGGGGTGAGATCAAGCCAAGGCACCAAAACACAAGATAGAGACAAGGTTCAAGGGG

GACCTATGTACAACTATTTGCGATGCCAACCGATCAGCCCTGATCAGGGTGAGTCATACGAGACTGTTAGTGCATTCATC

ACCGCTGACCTTAAGAAGTATTGCCTGAATTGGAGATACGAGACAATCAGCATATTTGCACAGAGGCTGAATGAAATATA

TGGACTACCATCCTTCTTTCAATGGTTACACAGGATATTGGAAAAATCCGTACTCTACGTCAGTGACCCACATTGTCCTC

CCGATCTAGATAATCATATCCCTCTGGACAGTGTCCCCAATGCCCAAATATTCATCAAGTACCCAATGGGCGGAATAGAA

GGTTATTGCCAAAAACTATGGACAATCAGCACTATACCATACTTGTATCTGGCAGCCTATGAGAGCGGAGTAAGAATCGC

CTCACTGGTGCAGGGTGACAATCAGACAATCGCAGTGACAAAAAGAGTTCCAAGTTCTTGGCCTTATTCACTAAAAAAGA

GGGAGGCCTCCAAGGCCGCTCAAAATTACTTCGTAGTCTTAAGGCAAAGATTGCACGATGTAGGTCATCACTTAAAGGCT

AATGAGACCATAGTATCTTCTCACTTTTTTGTATATTCCAAAGGGATTTATTATGACGGCCTGCTAGTCTCACAATCACT

AAAGAGCATCGCCAGATGTGTCTTCTGGTCCGAGACTATAGTGGATGAAACCAGAGCGGCGTGCAGTAATATTGCAACAA

CTGTCGCTAAGAGTATAGAGAGGGGTTATGATAGGTACCTTGCATATTCTTTGAATATCCTCAAGATTTTCCAACAGATC

CTTATATCCCTTAACTTCACTATTAACACAACAATGACTCAGGATGTCGTGGCACCGATCATCGAGAACGGTGATTTGCT

GATAAGGATGGCACTCTTGCCAGCACCCATCGGGGGTCTCAATTATCTTAACATGAGCAGGTTATTTGTGAGAAATATCG

GTGACCCGGTCACCTCCTCTATAGCCGACCTGAAGAGGATGATAGATGCCGGGCTAATGCCAGAAGAAACATTGCATCAA

GTGATGACCCAGACCCCGGGAGAATCATCCTACCTAGATTGGGCAAGTGACCCTTATTCTGCCAACCTACCCTGCGTACA

GAGTATCACTCGCCTTCTCAAAAACATCACTGCACGATATATTTTAATCAGCAGCCCAAACCCGATGCTGAAAGGGTTAT

TTCATGAGGGGAGTAGAGATGAAGACGAGGAGCTTGCAAGTTTCCTAATGGATCGGCATATAATTGTTCCAAGAGCTGCA

CATGAAATCTTAGACCATAGTATAACCGGAGCAAGAGAAGCTATAGCCGGGATGTTGGACACCACCAAGGGTCTGATTAG

AACAAGTATGAAACGGGGCGGCCTCACCCCTCGAGTCTTAGCCCGCCTTTCCAATTATGATTATGAACAATTCCGATCCG

GGATAACACTATTAACAAAGAAAGGGCAGTGTTATCTCATTGACAAGGACTCGTGCTCGGTGCAGCTCGCTATAGCTCTG

AGGGGCCACATGTGGGCTAAGTTGGCTCGCGGGAGACCAATCTATGGGTTGGAGGTGCCTGATGTATTAGAATCGATGAA

CGGCTACCTTATCAAACGCCACGAGTCGTGTGCTATCTGTGAAACGGGCTCAAGTCACTACGGATGGTTTTTCGTCCCTG

CAGGGTGCCAGCTTGACGATGTATCTAGAGAGACTTCGGCTCTTCGTGTACCTTATGTCGGATCAACAACTGAGGAAAGG

ACAGACATGAAACTTGCATTTGTTAGGTCTCCAAGTAGATCCCTCAAATCAGCAGTTAGAATTGCAACAGTTTACTCATG

GGCTTACGGGGATGATGAGAAATCATGGAGTGAAGCCTGGATGCTAGCTAGGCAGAGAGCCAATATCACCTTAGATGAAT

TGAGAATGATCACTCCAGTCTCCACATCCACCAACTTAGCCCACCGGTTGAGGGATCGGAGTACCCAGGTGAAATACTCG

GGGACATCCCTTGTGAGGGTTGCAAGATACACAACCATCTCCAATGATAACTTGTCGTTTGTGATATCTGAGAAAAAAGT

GGATACCAACTTCATTTATCAGCAAGGGATGTTGCTTGGTCTTGGGATCCTTGAGAACCTCTTTAGGCTAGAGGCCACCA

CCGGAGTATCCAACACAGTGCTACACCTGCACGTGGAAACAGAATGTTGTGTTGTACCTATGGTGGATCACCCAAGGATA

CCGAGTCTCCGTAATATAAAAGTTACGGATGAGCTATGCACAAACCCTTTGATCTACGACAGGTCCCCCATCATAGAGCA

CGATGCAACCCGRCTATATTCACAGAGCCACAGGAGACATTTGGTGGAGTTTGTTACCTGGTCAACAAGCCAGCTTTACC

ATATACTGGCTAAGTCTACAGCAATGTCCATGATTGAATTGATCACGAGATTCGAGAAAGATCACATGAATGAAATAGCC

GCCCTGATTGGCGATGACGACATCAACAGCTTCATCACAGAATTTTTGCTAGTGGAGCCTAGATTATTTATAGTTTACCT

TGGTCAGTGTGCTGCCATCAATTGGGCCTTTGATATACATTATCATCGGCCCTCGGGCAAGTACCAGATGGGGGAACTTC

TCTACTCTCTGCTCTCTCGGATGAGCAAAGGAGTATATAAGATCTTTACTAATGCTTTGAGCCACCCCAAAGTTTACAAG

AAATTTTGGCGAAGTGGCATCATTGAGCCGATTCATGGCCCATCCTTGGATACACAGAATTTACATGTCACTGTCTGTGA

CATGATATATGGGTCCTATGTCACCTATTTGGATCTTTTGCTGAATGATGAGCTAGATGATTACCCGTATTTGCTCTGCG

AGAGTGATGAGGACGTGGTCACAGACAGGTTCGACAACATTCAAGCCAAACATCTCTGTGTATTGGCCGATGTATATTGC

AGCTCCAAGAGGTGTCCTTCAATAATCGGGATGTCTCCTATAGAAAAATGTACCATTCTCACACATTACATCAAGGGAGA

ATCAGTACAATCCCCGTCTGGGACCTCATGGAACACTGATCCCCTTGTAGTAGATCATTACTCATGCTCTCTGACCTACC

TTCGCCGCGGTTCCATCAAACAAATCAGGTTGAGGGTGGATCCTGGGTTTGTATTCGAGGCGTTGACAGACGTCGACTTT

AAACAACCTCGCAAAGCTAAGTTAGATATATCGGTCGTTGGGTTGACTGATTTCTCTCCCCCTTACGACAACGTCGGTGA

TTTTTTAGGGACTATCAACACATTGAGGCACGATCTGCCTGTCACCGGAACCGGGGTCTCGAACTATGAAGTCCACGCTT

ATCGTAGAATTGGCCTGAATTCATCAGCATGTTATAAAGCCGTAGAGATCTCCACGTTAATCARGCCATCCTTAGAAGTC

GGAGAGCATGGATTGTTCTTAGGAGAAGGTTCCGGTTCAATGCTGGCTGCGTACAAAGAAGTTCTTAAATTAGCAAATTG

TTACTACAACAGCGGAGTAACAGCGGAGGGCAGAGCCGGACAGAGGGAAATATCTCCCTATCCTTCAGAGATGAGCCTAG

TAGAGAATCAAATGGGGATAGAGCGGAGTGTTAAAGTGCTGTTCAACGGGAAACCTGAAGTAACTTGGGTGGGGACCACA

GATTGCTACAAGTACATAATCAGTAACATCCAAACCTCTAGTCTGGGTTTCATACATTCAGATATTGAGACGCTCCCAAC

CAAAGATGCTGTTGAGAAATTGGAGGAATTTGCCTCTATCCTATCCTTATCCTTGATTTTGGGGAAAATCGGCTCTATTA

CAGTTGTCAAAATTATGCCCATTAGCGGGGATTTTACCCAAGGCTTCATAGCCTATGCCATCCAATATTTTAGGGAGAGC

CTGCTTGCCTACCCGAGATATAGTAACTTCATCTCGACTGAGTGTTACCTTATTATGATAGGATTAAAGGCCAATCGATT

GATAAACCCAGAAGCCATTAAGCAAAGCATAATCAGAGCGGGGGTCAGGACTGCACCAGGACTTGTGAGCCATGTATTAT

CAGAGAAACAAAAAGGTTGTATTCAATCTTTCTTGGGTGATCCTTATATCCAAGGAGACTTCAATAAGCACCTTAAATCT

CTAACCTCTATTGAGAAAATCCTGGTAAATTGTGGTCTCTCGATCAATGGCATAAAAATCTGTAGAGATCTAATCCACCA

TGATATCGCCTCCGGTCCAGACGGCCTTATGAGCTCTACAATTATTTTATATAGGGAATTGGCTCATTTCAAAGATAATA

TAAGGAGTCAGCACGGAATGTTCCACCCCTACCCGGTACTGGCCAACAGCAGGCAACGTGAATTAATCCTTCGAATAGCC

AAGAAATTCTGGGGGTATGTCTTGCTATATTCTGATGACCCGGCACTAATCAGACAAACAATCAAGAACTTGAAACGGAA

TCATCTAACCTTTGACTTACATAGTAACCCATTTATTAAAGGGCTATCCAAGGCTGAGAAACTGCTGGTGCGGACGAGTT

CACTAAGAAGAGAATGGTTGTTCACTCTCGAAACGAAAGAAGTAAAAGAGTGGTTCAAATTGGTAGGTTACAGTGCACTC

ATCAGAGGCTAGTCAGCTATGCATCTGCCCCCTTCTCCTCCGCCATGAGACCCCACTGGCGATCCAGAAGATTAAAGAAA

ACTACATATTGGATAAGTATCTATTCCCAGCTTTGTCTGGT

>KP868655.1/goat/China/2014-12-05

ACCAAACAAAGTTGGGTAAGGATAGATCTTATAATAACTATGGACTGGCAAACTTAGGAGTAAAGATCCTACTGTCGGGG

GGAGGAGGAGGAGCAAGATCTTTGATTATGGCGACTCTCCTTAAAAGCTTAGCATTGTTCAAAAGGAACAAAGACAAAGC

GCCGACAGCATCAGGTTCAGGAGGGGCCATCCGGGGGATTAAGAATGTTATCATAGTCCCGATTCCCGGAGACTCGTCCA

TCACTACCCGTTCAAGGCTGCTCGACAGGCTTGTCAGATTGGCCGGAGATCCTGACATCAACGGGTCAAAGCTGACCGGG

GTGATGATCAGCATGTTATCCTTGTTCGTAGAGTCACCCGGGCAATTGATACAGCGAATCACAGATGATCCGGATGTTAG

TATCCGCCTTGTTGAGGTAGTTCAAAGTACTAGGTCCCAGTCCGGGTTGACCTTTGCATCACGTGGTGCTGATTTAGACA

ACGAGGCAGACATGTATTTTTCAACTGAGGGGCCCTCGAGTGGAGGTAAGAAAAGGATCAACTGGTTTGAGAACAGAGAA

ATAATAGACATAGAGGTGCAGGATCCAGAAGAGTTCAATATGTTGTTAGCCTCCATACTAGCACAAGTCTGGATCCTTCT

GGCCAAGGCTGTTACGGCACCAGATACGGCAGCTGACTCAGAACTGAGAAGGTGGGTTAAATACACACAACAAAGGAGAG

TGATTGGGGAATTTCGCCTTGACAAGGGGTGGCTGGATGCAGTCCGCAACAGAATTGCAGAAGATCTATCACTCCGGCGG

TTCATGGTATCTCTCATACTTGACATCAAGAGGACCCCAGGCAACAAGCCAAGGATTGCAGAAATGATCTGCGACATTGA

CAACTATATTGTCGAAGCAGGACTCGCCAGTTTCATCCTTGCTATCAAATTTGGTATCGAAACCATGTATCCTGCATTAG

GGCTCCACGAGTTTGCCGGAGAATTGTCCACTATTGAGTCCTTGATGAACTTGTATCAACAGCTAGGAGAGGTTGCACCC

TACATGGTAATTCTAGAGAATTCAATTCAGAACAAGTTTAGTGCAGGAGCTTATCCCCTCCTCTGGAGCTATGCGATGGG

TGTCGGAGTCGAGCTGGAGAACTCAATGGGGGGCTTGAACTTTGGCAGGTCATATTTTGACCCTGCCTATTTTCGTCTCG

GACAGGAGATGGTCAGAAGATCTGCAGGAAAGGTCAGCTCTGTGATTGCGGCTGAGCTCGGCATCACAGCAGAGGAAGCT

AAACTAGTCTCGGAAATCGCCTCGCAGGCTGGGGACGAAAGAACCGCCAGAGGGACTGGGCCTCGACAGGCGCAGGTTTC

CTTCCTTCAGCACCAAACAGGAGGGGGAGAGTCGTCCGCACCAGCGACCAGAGAAGGGGTCAAAGCTGCGATCCCAAACG

GAGCTGAAGAAAGGGACAGAAAGCAAACACGCCCAGGAAGGCCCAGAGGAGAGACCTCCGGCCAACTGCTCCTGGACATC

ATGCCAGAGGATGAGATCTTGCGAGAGTCTGGTCAAAACCCTCGTGAGGCTCAAAGATCGGCCGAGGCACTCTTCAGGCT

GCAGGCCATGGCCAAGATTCTGGAGGACCAGGAGGAGGGAGAAGACAACAGTCAGGTCTACAACGACAAGGATCTCCTCG

GCTAAGCAGACGCACCCTCTGTCGAAATCAGTGACAAGACATCTCTTACCAGTATTATAAAAAACTTAGGACCCAGGTCC

AAGCAAGCACACATCGACACTCCAACCAGTCGAGCGGAGACCACCGATGGCAGAAGAACAAGCATACCATGTCAACAAGG

GGCTGGAATGTATCAAGTCTCTCAAAGCCTCACCCCCGGATCTATCCACCATCAGAGATACCATCGAGAGCTGGAGAGAG

GGGCTTAGCCCCTCGGGCCGTGCAACACCGAACCCTGATACGTCCGAGGGAGATCATCAGAATATCAACCAATCATGCTC

ACCGGCAATCGGACCAAACAAAGTCTACTTGTCTCCTGAAGATAATCTCGGATTTAGAGAGATCACTGGCAACGACTGTG

AGGCTGGGCTCGGAGGGGTCCAGAGAGAAGGATCCAACTCTCAAGTACAGCGTTACCATGTTTATAGTCACGGGGGTGAA

GAGATTGAAGGACTCGAGGATGCTGACTCTCTCGTGGTTCAAGCAGATCCTCCAGTTGCTAACGTCTTCAATGGAGGAGA

GGATGGATCTGACGACAGCGATGTGGACTCTGGCCCAGATGATCCCGGCAGAGATACTCTATATGACCGGGGATCTGTTG

CCGGCAATGGTATCGCTAGGTCCACAGATGTCGAAAAACTAGAAGGTGCTGATATTCAAGAAGTTCTTAACTCCCAGAAA

GGCAAAGGAGGAAGATTCCAGGGC-GGAAAACCTTGCGAGTCCCGGAAATACCCGATGTCAAGCACTCCAGACCATCAGC

CCAATCAATTAAAAAGGGCACAGACGGGAACTCAGTCTCATCTGGAACGGTGATAGAGTGTTTATCGATAAGTGGTGCAA

CCCAAACTGTGCCAGAGTCAAGATGGGAGTCATCAGAGCAAAATGCGTCTGTGGGGAGTGTCCTCAAGTCTGCGAGGAGT

GCAAAGACGATCCAGGGGTCGACACAAGAATCTGGTACCATAGCATCACCGACTCAGCCTAAAGAGAATGACTCCGAGTA

TGAGTATGAGGATGACCTATTTACAGAGATTCAGGACATCCGTGCAAGCATTGCCAAGATCCATGATGACAATAAAACTA

TCCTCTCAAAGCTTGATTCTATACTGTTATTGAAAGGGGAAGTCGACACTATCAAGAAACAAATCAGCAAGCAGAATATA

AGTATATCCACCATTGAGGGCCATCTCTCCAGTATAATGATAGCCATCCCTGGCTTTGGGAAGGACATCAAGGACCCAAC

ATCCGAAGTCGAGTTGAACCCAGATCTAAGACCTATAATAAGCCGTGACTCTGGCAGAGCTCTCGCGGAGGTCCTCAAGA

AACCCGCTGTGGATAGGTCTCAGAAAATTGGAACCAAAGCCAACTCCAGCTCAAAGGGTCAGCTTCTTAAGGATCTCCAG

CTAAAACCTGTCGACAAGCAGGCAAGCTCTGCAATCGGGTTTGTCCCGTCCGACCATGAATCATCCAGAAATGTCATCCG

CTCCATAATCAAGTCGAGCAAGCTAAATATTGATCACAAGGACTATCTTCTAGATTTACTAAATGATGTGAAAGGCTCCA

AGGATCTTAAGGAATTTCACAAGATGCTAACAGCAATTCTCGCCAAGCACCCGTAACACATCCTCCAGTCATCATCTCAT

ACTCGACTAAAAACATCCTTTCAATCAGGCTATTACAAAAAACTTAGGAGCAAGGGCAACTGAGCTTCGCAGACAGGATG

ACCGAGATCTACGACTTCGATAAATCAGCATGGGATGTCAAAGGGTCAATTGCCCGCATAGAACCCACCACCTACCACGA

CGGCCGACTGATACCCCAGGTGAGGGTCATCGATCCTGGTCTGGGAGACAGAAAAGATGAGTGCTTTATGTACCTGTTTC

TCCTAGGAGTGATTGAGGATAACGACCCCCTGTCTCCCCCCGTCGGAAGAACCTTTGGCTCTCTACCTCTAGGGGTTGGT

AGGTCAACTGCCAAGCCAGAAGAACTACTAAGGGAGGCCACAGAATTAGATATAGTGGTGAGGCGCACTGCAGGAGTAAA

TGAGAAACTGGTATTTTACAACAACACTCCGCTGTCCTTGTTAACGCCCTGGAAGAAAGTTCTGACAACCGGAAGTGTGT

TTAGCGCTAACCAGGTTTGCAATGCAGTCAACCTAGTCCCGCTTGATACTCCCCAGAGATTCAGGGTTGTGTACATGAGC

ATAACTAGATTATCAGACAATGGTTACTATAGTGTGCCCAGAAGAATGCTGGAGTTCCGCTCGGCCAATGCAGTAGCCTT

CAACATCTTGGTTACACTAAGAATTGAAAATGGCACAAACCCTAGAAGATACATAGTCGGCTCATGGGAGAATTCTGAGG

TCACATTTATGGTACACGTGGGCAACTTTAGGAGAAAGAAGAACGAAGTATACTCTGCTGATTACTGCAAAATGAAGATT

GAAAAGATGGGTTTAGTTTTTGCCCTGGGTGGAATAGGCGGAACAAGTCTCCATATTAGAAGCACCGGGAAAATGAGCAA

AACCCTCCATGCACAGCTGGGGTTCAAGAAAATCTTATGTTACCCCCTAATGGATGTCAATGAGGATCTTAACCGATATC

TCTGGCGGGCAGAGTGCCGAATAGTCAAAATCCAAGCTGTCTTACAGCCATCAGTACCCCAAGAGTTCCGTGTCTACGAT

GATGTCATCATCAACGATGATCAAGGCTTGTTCAAGATCTTGTAGTTCATTTGCAAAGGAACGAACCCACCCCCAGGAAG

GGGCCCCACCCTCCCCCG-CCCAAAAACACACTCCAAGGCTCCCCGGCACAGCGCCAACACAGCCACTCCCGCGATGGAA

CGAGGTCGGCC-CCCGAACAGATCCGGTCCAGGGGGGGGGGCCCCCCCCCTGACCAACCGGGCGCCGACCGAGCAGGCCG

GCGAGAAGCCCCCAAGAGGCCCCGACGAAGACCGAACACCCCATCCCCTC-AAAAACGTCCCCC-AACACGCCCGAGGCC

AGCCACCGGACACCCCCAGACCCGCCACCGGCGAGACAGAACCCCAAGGGGGGACCCTCCAAACCAAAGGAATCACCGGG

GTAGGCTCCGCCGAACCAGCACAGACCCGACCCAAGCAAGGCGGACCCCCAACACGAAACTGCCCAGCACACCCTACTCA

TCATCAAACAAAACTTAGGGGCCAAGTCCATACGAGAGGCCCTAGTCCGTCAGGCGGAGGGACGCAGAAAGGAAGGAGAC

ACGACCGTCCCAGAAGGAGGACACCCAACCACCGAAACGCCCGCCCCCAGGGAAGGAG--CCA-CCCCCACGCAGCGAAG

GAGCGGGTGGGAGACCTCGGGGAGGGCCCGACCGGGACCCGGGGGCCGGCGGCGCCGGCGGTGAGCAGGCCGGGCGCATA

CCTGACCCCACCGGAGGATCCGAGGAGGGCCCGACCCCGCGTGGACCAAACGGCAGGGAGCCCAACCCCACCAAAAAAAG

AGGCGAGGCCCCCCCTCCCTCCCCCCCCGCACACAACCCGCAGACAGACCGAGGGAGGGCCATCCCCCAGCGAGACAGGC

GGCTCCCTCGCAGCCGAGCCGCAGCTCCCCAGAATAGGACACCCCCGCACACTCCGAACACCCCCTCGCCCCCCGCCACC

GGGACAGGGGCTCCACCACCACAGACCGGAGCCACCGACAGCCAGCCAGAGGTCCCCTCCAGTCCCGCCCGGCGCCCCCC

CGAAGAGGCCCCGGCAAGGCCCCCCAGCCCGACGTCCCCCGCACGCACCGAAAGAAGGACCCGCCCGAGAACGTGCGGCA

CATCCATGCGCAAACATCATGACGTGGGTCGCAATTTTGACATTTCTGTTTCTTTTCCCAAATGTTGTTGCGTGTCAGAT

TCACTGGGGCAATCTATCCAAGATCGGGATTGTAGGAACAGGGAGTGCCAGCTACAAGGTGATGACTAGGCCAAGCCACC

AGACTCTGGTTATAAAGTTAATGCCAAATATAACGGCCATCGACAATTGTACAAAGTCAGAGATTGCAGAGTACAAGAGA

TTGCTGATCACAGTGTTAAAGCCTGTAGAGGATGCTCTGTCGGTGATAACCAAGAATGTAAGACCAATTCAAACTCTAAC

ACCTGGGCGTAGAACCCGCCGTTTTGCTGGAGCTGTTCTGGCCGGGGTAGCACTTGGAGTTGCGACAGCCGCTCAGATAA

CTGCAGGAGTCGCCCTTCATCAATCATTGATGAACTCCCAAGCAATTGAGAGTTTAAAAACCAGTCTTGAGATGTCGAAT

CAGGCAATAGAAGAAATCAGACTTGCAAATAAGGAGACCATACTGGCAGTACAGGGCGTCCAGGATTATATCAACAATGA

GCTCGTCCCTTCTGTTCATAGAATGTCATGCGAGCTGGTAGGTCACAAGCTCGGCCTCAAGCTCCTTAGGTACTACACCG

AGATCCTGTCCATATTCGGGCCCAGTCTTCGAGACCCGATATCTGCCGAAATATCAATCCAGGCACTTAGTTATGCATTA

GGCGGAGACATTAATAAAATCCTGGACAAGCTTGGGTATAGCGGTGGGGATTTCCTTGCCATCCTAGAAAGCAAGGGAAT

AAAGGCCCGGGTTACATATGTGGACACAAGAGATTACTTTATAATCCTTAGCATCGCCTACCCAACCTTATCTGAGATCA

AGGGAGTGATAGTTCACAAGATAGAAGCTATAACATACAACATTGGGGCACAGGAGTGGTATACTACTATCCCTAAATAT

GTAGCCACTCAGGGGTATCTGATATCGAACTTTGATGAGACGTCATGCGTATTCACTCCAGAGGGGACAGTTTGCAGCCA

GAATGCGTTGTACCCAATGAGCCCATTGCTTCAGGAATGTTTCAGGGGGTCAACAAAATCGTGCGCCAGAACCTTAGTTT

CAGGGACCATAAGTAATAGATTTATCCTATCAAAAGGGAACCTGATTGCAAATTGTGCGTCAGTTTTGTGCAAATGTTAC

ACAACAGAGACAGTTATCAGCCAGGATCCTGACAAACTACTAACTGTTGTAGCATCCGACAAGTGTCCTGTAGTTGAGGT

GGATGGAGTGACAATACAGGTCGGCAGTCGAGAGTATCCGGATTCTGTATACTTACACAAAATAGACTTAGGTCCAGCCA

TCTCCCTAGAAAAACTGGATGTAGGCACCAATTTAGGCAATGCAGTCACAAGACTGGAGAATGCAAAGGAGCTCCTAGAT

GCATCAGACCAAATACTGAAGACTGTTAAAGGGGTACCTTTCAGTGGGAATATGTACATAGCACTGGCAGCTTGCATAGG

AGTATCCCTGGGCCTTGTCACATTAATATGCTGCTGTAAGGGGAGGTGTAAGAACAAGGAAATCCCTATCTCCAAAATCA

ACCCAGGGCTCAAACCCGACCTGACCGGGACCTCAAAGTCGTACGTAAGATCACTGTAGTCAGAATCACCTGGATCATCT

GGCATCGCACACATACATGCACGACACAGGCAGTCCGAGGACGCAAGAAACCCAGCCTCCGGTCACCCACCCGACCCCAC

TCCACGCTCCACCACACATTAGTCATCAAACAAAACTTAGGACGAAAGGTCAATCACCATGTCCGCACAAAGGGAGAGGA

TCAATGCCTTCTACAAAGACAATCCTCACAATAAGAACCATAGGGTGATCCTGGATAGAGAACGCTTGGTCATTGAAAGA

CCCTACATCTTGCTTGGAGTCCTGCTGGTAATGTTCCTGAGTCTAATCGGACTGCTGGCCATTGCAGGGATCAGGCTTCA

CCGGGCCACCGTTGGAACTTCAGAGATCCAGAGTCGGCTGAATACCAATATTAAGTTGACCGAATCTATTGATCACCAGA

CTAAGGATGTCTTAACTCCCCTTTTTAAAATCATTGGCGATGAAGTCGGCATCAGAATTCCACAGAAATTCAGTGATCTT

GTCAAGTTCATCTCCGATAAGATTAAATTCCTCAACCCTGATAGAGAGTATGATTTCAGAGATCTCCGGTGGTGTATGAA

CCCCCCCGAGAGAGTCAAAATTAATTTTGATCAGTTTTGTGAGTACAAGGCTGCGGTTAAGTCAATTGAACATATATTTG

AGTCACCACTCAACAAGTCAAAAAAGCTGCAATCTTTGACTCTCGGGCCCGGAACAGGCTGTCTAGGCAGGACAGTAACA

AGAGCCCATTTCTCAGAACTTACAATGACCTTAATGGACCTGGATCTAGAGACGAAGCACAACGTGTCCTCAGTGTTTAC

CGTAGTTGAAGAGGGATTATTCGGAAGAACATATACCGTCTGGAGATCCGATGCCAGGGATCCGAGCACCGATCTAGGTA

TCGGCCATTTTTTAAGAGTCTTCGAGATTGGACTGGTAAGAGATCTTGGGCTGGGTCCCCCTGTTTTTCATATGACCAAC

TATCTCACGGTGAACATGAGTGATGACTATCGGAGATGTCTTTTAGCGGTAGGGGAGTTGAAGTTGACAGCCCTATGCAC

CTCATCTGAGACTGTGACACTGAGTGAGAGAGGAGTTCCAAGGAGGGAACCTCTTGTGGTTGTGATACTTAATCTAGCTG

GACCCACTCTAGGGGGAGAGCTATACAGTGTCTTGCCTACCTCTGATCTCATGGTGGAGAAACTCTATTTGTCTTCACAT

AGAGGGATCATCAAAGATGATGAGGCCAATTGGGTAGTGCCGTCTACCGATGTTCGTGATCTTCAGAACAAAGGTGAATG

TCTGGTGGAAGCATGCAAGACTCGACCTCCTTCATTTTGCAATGGCACAGGATCAGGCCCGTGGTCAGAAGGGAGAATCC

CTGCCTACGGGGTGATCAGGGTCAGTCTTGACTTAGCGAGTGACCCGGATGTAGTTATCACTTCAGTGTTTGGCCCACTG

ATACCTCACCTATCCGGCATGGATCTTTACAACAACCCGTTTTCAAAAGCTGTATGGTTGGCTGTACCACCTTATGAGCA

GTCATTTCTAGGAATGATAAATACAATTGGATTCCCTAACAGAGCAGAGGTTATGCCGCACATTTTGACCACAGAGATCA

GAGGCCCTCGGGGTCGTTGCCATGTTCCCATAGAATTGTCCCGCCGGGTTGATGACGATATCAAGATCGGGTCCAACATG

GTCATATTGCCGACGATGGACCTGAGGTATATTACAGCCACTTATGATGTTTCCAGGAGCGAGCATGCAATCGTGTACTA

TATCTATGACACGAGTCGCTCATCATCTTACTTCTACCCAGTTCGACTGAATTTCAAAGGCAATCCTCTCTCTTTGAGGA

TAGAGTGTTTCCCCTGGCGTCATAAGGTGTGGTGCTACCATGATTGTCTTATATACAACACCATAACAGGTGAAGAGGTC

CATACGAGAGGGCTGACCGGCATAGAGGTAACATGCAATCCAGTCTGAGCAGAGCTACGACCGTCACTCAGGCAGTCCCT

TGAGTCGCCACCGAGTCCAAGCAGCACAGCCTGGGACAATCAACGGCACAGCCCAGCCAACAATGTTATAAAAAACTTAG

GAGCCAGGGTTGTAGGGGCCATGGACTCTCTATCAGTCAATCAGGTCTTGTACCCTGAGGTTCATCTAGACAGCCCTATT

GTCACAAACAAACTAGTTGCCATCCTTGAGTACTCGGGTATCGACCATAACTATGTTCTTGAAGACCAGACTCTTGTCAA

GAATATTAGGTATAGGCTGGGGTGCGGTTTTTCAAATCAAATGATCATCAATAACAGGGGGTTGGGTGAAACAGTCAATT

CTAAACTTAAAAGTTACCCCCATAATCGTCATATCATATACCCGGATTGCAATAAGGAGTTGTCTTGTATCAAGGATAGC

TGCATATCTAAGAAGCTCTCGGAGCTATTCAAGAAGGGTAATTCCTTGTACTCTAAGATAAGTCACCAGGTACTGGATTG

TCTTAAGAGAGTCAATGGGAAATTAGGTCTGGGCACAGATCTTACCCATGGTCTGAAGGAGGGTATCCTTGACCTAGGGT

TGCACATGCATAGCTCTCAATGGTTCGAGACTTTCCTGTTCTGGTTCACTATCAAGACAGAGATGAGATCAATGATCAAA

GAACAGTCCCATATATGCCACAAGAGGAGGTATAACCCGACTTTTGTGTCGGGGGATGCATTCGAGGTGCTCGTATCGCG

AGACCTCGTTGTGATAATTGATAAGAATACCCAGTATGTCTTCTACTTAACATTTGAGCTGGTCCTTATGTATTGTGATG

TTATAGAGGGTAGACTTATGACGGAGACAGCCATGGCCATAGACCAGAGATATTCAGAACTTCTAAGCCGGGTCAGATAC

TTGTGGGATCTTATTGATGGGTTCTTCCCAACACTAGGCAACACCACATACCAAATTGTTGCTCTGCTTGAACCATTATC

GTTGGCTTATCTTCAACTTCAGGATGTCACTCTGGAGTTAAGAGGTGCTTTTTTAGACCATTGCTTCAAAGAACTCTATG

AGATACTGGAACATTGCGGCATTGACACAGCAGGCACTTACAATTCCATCACTGAAGGATTGGATTACGTATTCATCACC

CACGATATACATCTAACTGGGGAGATTTTTTCATTTTTCCGGAGTTTCGGACACCCCCGTCTAGAAGCAGTCACCGCTGC

GGAAAATGTCAGAAAACATATGAACCAACCGAAGGTAATCAGTTATGAGACTATGATGAAAGGCCATGCGGTATTTTGCG

GGATAATCATAAATGGTTTTCGGGACCGACATGGCGGCAGCTGGCCCCCTGTTGCACTGCCAGAACATGCCTCTGCTGCG

ATCCGGAATGCGCAGGCATCAGGTGAGGGACTGACCCATGACCTGTGTATAGACAACTGGAAATCCTTTGTTGGATTCAA

ATTTGGCTGCTTTATGCCACTCAGCCTAGATAGTGATTTGACCATGTATCTCAAAGACAAGGCATTGGCTGCACTGAAGA

ATGAGTGGGATTCAGTTTACCCGAAAGAATACCTCCGTTATAACCCACCTAGAGGGACAGAGTCAAGGCGATTGGTAGAG

GTATTCCTGAATGACTCCAGCTTTGATCCTTATAACATGATAATGTACGTGGTGAATGGCTCCTACCTTAAAGACCCTGA

GTTTAACCTCTCATACAGCCTAAAAGAGAAGGAGATAAAGGAGACAGGGCGGTTGTTTGCCAAAATGACCTATAAGATGC

GGGCCTGTCAGGTAATCGCTGAAAATCTGATATCGAATGGTGTTGGGAAGTATTTCCGAGACAATGGGATGGCAAAGGAC

GAGCATGACCTAACAAAAGCCCTTCACACCCTGGCAGTCTCAGGTGTTCCCAAAAACAACAAAGATAACCACCGAGGTGG

GCCTCCCAGAAGGACAACAAACCGAGGGTTGAGATCAAGCCAAGGCACCAAAACACAAGATAGAGACAAGGTTCAAGGGG

GACCTATGTACAACTATTTGCGATGCCAACCGATCAGCCCTGATCAGGGTGAGTCATACGAGACTGTTAGTGCATTCATC

ACCGCTGACCTTAAGAAGTATTGCCTGAATTGGAGATACGAGACAATCAGCATATTTGCACAGAGGCTGAATGAAATATA

TGGACTACCATCCTTCTTTCAATGGTTACACAGGATATTGGAAAAATCCGTACTCTACGTCAGTGACCCACATTGTCCCC

CCGATCTAGATAATCATATCCCTCTGGACAGTGTCCCCAATGCCCAAATATTCATCAAGTACCCAATGGGCGGAATAGAA

GGTTATTGCCAAAAACTATGGACAATCAGCACTATACCATACTTGTATCTGGCAGCCTATGAGAGCGGAGTAAGAATCGC

CTCACTGGTGCAGGGTGACAATCAGACAATCGCAGTGACAAAAAGAGTTCCAAGTTCTTGGCCTTATTCACTAAAAAAGA

GGGAGGCCTCCAAGGCCGCTCAAAATTACTTCGTAGTCTTAAGGCAAAGATTGCACGATGTAGGTCATCACTTAAAGGCT

AATGAGACCATAGTATCTTCTCACTTTTTTGTATATTCCAAAGGGATTTATTATGACGGCCTGCTAGTCTCACAATCACT

AAAGAGCATCGCCAGATGTGTCTTCTGGTCCGAGACTATAGTGGATGAAACCAGAGCGGCGTGCAGTAATATTGCAACAA

CTGTCGCTAAGAGTATAGAGAGGGGTTATGATAGGTACCTTGCATATTCTTTGAATATCCTCAAGATTTTCCAACAGATC

CTTATATCCCTTAACTTCACTATTAACACAACAATGACTCAGGATGTCGTGGCACCGATCATCGAGAACGGTGATTTGCT

GATAAGGATGGCACTCTTGCCAGCACCCATCGGGGGTCTCAATTATCTTAACATGAGCAGGTTATTTGTGAGAAATATCG

GTGACCCGGTCACCTCCTCTATAGCCGACCTGAAGAGGATGATAGATGCCGGGCTAATGCCAGAAGAAACATTGCATCAA

GTGATGACCCAGACCCCGGGAGAATCATCCTACCTAGATTGGGCAAGTGACCCTTATTCTGCCAACCTACCCTGCGTACA

GAGTATCACTCGCCTTCTCAAAAACATCACTGCACGATATATTTTAATCAGCAGCCCAAACCCGATGCTGAAAGGGTTAT

TTCATGAGGGGAGTAGAGATGAAGACGAGGAGCTTGCAAGTTTCCTAATGGATCGGCATATAATTGTTCCAAGAGCTGCA

CATGAAATCTTAGACCATAGTATAACCGGAGCAAGAGAAGCTATAGCCGGGATGTTGGACACCACCAAGGGTCTGATTAG

AACAAGTATGAAACGGGGCGGCCTCACCCCTCGAGTCTTAGCCCGCCTTTCCAATTATGATTATGAACAATTCCGATCCG

GGATAACACTATTAACAAAGAAAGGGCAGTGTTATCTCATTGACAAGGACTCGTGCTCGGTGCAGCTCGCTATAGCTCTG

AGGGGCCACATGTGGGCTAAGTTGGCTCGCGGGAGACCAATCTATGGGTTGGAGGTGCCTGATGTATTAGAATCGATGAA

CGGCTACCTTATCAAACGCCACGAGTCGTGTGCTATCTGTGAAACGGGCTCAAGTCACTACGGATGGTTTTTCGTCCCTG

CAGGGTGCCAGCTTGACGATGTATCTAGAGAGACTTCGGCTCTTCGTGTACCTTATGTCGGATCAACAACTGAGGAAAGG

ACAGACATGAAACTTGCATTTGTTAGGTCTCCAAGTAGATCCCTCAAATCAGCAGTTAGAATTGCAACAGTTTACTCATG

GGCTTACGGGGATGATGAGAAATCATGGAGTGAAGCCTGGATGCTAGCTAGGCAGAGAGCCAATATCACCTTAGATGAAT

TGAGAATGATCACTCCAGTCTCCACATCCACCAACTTAGCCCACCGGTTGAGGGATCGGAGTACCCAGGTGAAATACTCG

GGGACATCCCTTGTGAGGGTTGCAAGATACACAACCATCTCCAATGATAACTTGTCGTTTGTGATATCTGAGAAAAAAGT

GGATACCAACTTCATTTATCAGCAAGGGATGTTGCTTGGTCTTGGGATCCTTGAGAACCTCTTTAGGCTAGAGGCCACCA

CCGGAGTATCCAACACAGTGCTACACCTACACGTGGAAACAGAATGTTGTGTTGTACCTATGGTGGATCACCCAAGGATA

CCGAGTCTCCGTAATATAAAAGTTACGGATGAGCTATGCACAAACCCTTTGATCTACGACAGGTCCCCCATCATAGAGCA

CGATGCAACCCGGCTATATTCACAGAGCCACAGGAGACATTTGGTGGAGTTTGTTACCTGGTCAACAAGCCAGCTTTACC

ATATACTGGCTAAGTCTACAGCAATGTCCATGATTGAATTGATCACGAGATTCGAGAAAGATCACATGAATGAAATAGCC

GCCCTGATTGGCGATGACGACATCAACAGCTTCATCACAGAATTTTTGCTAGTGGAGCCTAGATTATTTATAGTTTACCT

TGGTCAGTGTGCTGCCATCAATTGGGCCTTTGATATACATTATCATCGGCCCTCGGGCAAGTACCAGATGGGGGAACTTC

TCTACTCTCTGCTCTCTCGGATGAGCAAAGGAGTATATAAGATCTTTACTAATGCTTTGAGCCACCCCAAAGTTTACAAG

AAATTTTGGCGAAGTGGCATCATTGAGCCGATTCATGGCCCATCCTTGGATACACAGAATTTACATGTCACTGTCTGTGA

CATGATATATGGGTCCTATGTCACCTATTTGGATCTTTTGCTGAATGATGAGCTAGATGATTACCCGTATTTGCTCTGCG

AGAGTGATGAGGACGTGGTCACAGACAGGTTCGACAACATTCAAGCCAAACATCTCTGTGTATTGGCCGATGTATATTGC

AGCTCCAAGAGGTGTCCTTCAATAATCGGGATGTCTCCTATAGAAAAATGTACCATTCTCACACATTACATCAAGGGAGA

ATCAGTACAATCCCCGTCTGGGACCTCATGGAACACTGATCCCCTTGTAGTAGATCATTACTCATGCTCTCTGACCTACC

TTCGCCGCGGTTCCATCAAACAAATCAGGTTGAGGGTGGATCCTGGGTTTGTATTCGAGGCGTTGACAGACGTCGACTTT

AAACAACCTCGCAAAGCTAAGTTAGATATATCGGTCGTTGGGCTGACTGATTTCTCTCCCCCTTACGACAACGTCGGTGA

TTTTTTAGGGACTATCAACACATTGAGGCACGATCTGCCTGTCACCGGAACCGGGGTCTCGAACTATGAAGTCCACGCTT

ATCGTAGAATTGGCCTGAATTCATCAGCATGTTATAAAGCCGTAGAGATCTCCACGTTAATCAAGCCATCCTTAGAAGTC

GGAGAGCATGGATTGTTCTTAGGAGAAGGTTCCGGTTCAATGCTGGCTGCGTACAAAGAAGTTCTTAAATTAGCAAATTG

TTACTACAACAGCGGAGTAACAGCGGAGGGCAGAGCCGGACAGAGGGAAATATCTCCCTATCCTTCAGAGATGAGCCTAG

TAGAGAATCAAATGGGGATAGAGCGGAGTGTTAAAGTGCTGTTCAACGGGAAACCTGAAGTAACTTGGGTGGGGACCACA

GATTGCTACAAGTACATAATCAGTAACATCCAAACCTCTAGTCTGGGTTTCATACATTCAGATATTGAGACGCTCCCAAC

CAAAGATGCTGTTGAGAAATTGGAGGAATTTGCCTCTATCCTATCCTTATCCTTGATTTTGGGGAAAATCGGCTCTATTA

CAGTTGTCAAAATTATGCCCATTAGCGGGGATTTTACCCAAGGCTTCATAGCCTATGCCATCCAATATTTTAGGGAGAGC

CTGCTTGCCTACCCGAGATATAGTAACTTCATCTCGACTGAGTGTTACCTTATTATGATAGGATTAAAGGCCAATCGATT

GATAAACCCAGAAGCCATTAAGCAAAGCATAATCAGAGCGGGGGTCAGGACTGCACCAGGACTTGTGAGCCATGTATTAT

CAGAGAAACAAAAAGGTTGTATTCAATCTTTCTTGGGTGATCCTTATACCCAAGGAGACTTCAATAAGCACCTTAAATCT

CTAACCTCTATTGAGAAAATCCTGGTAAATTGTGGTCTCTCGATCAATGGCATAAAAATCTGTAGAGATCTAATCCACCA

TGATATCGCCTCCGGTCCAGACGGCCTTATGAGCTCTACAATTATTTTATATAGGGAATTGGCTCATTTCAAAGATAATA

TAAGGAGTCAGCACGGAATGTTCCACCCCTACCCGGTACTGGCCAACAGCAGGCAACGTGAATTAATCCTTCGAATAGCC

AAGAAATTCTGGGGGTATGTCTTGCTATATTCTGATGACCCGGCACTAATCAGACAAACAATCAAGAACTTGAAACGGAA

TCATCTAACCTTTGACTTACATAGTAACCCATTTATTAAAGGGCTATCCAAGGCTGAGAAACTGCTGGTGCGGACGAGTT

CACTAAGAAGAGAATGGTTGTTCACTCTCGAAACGAAAGAAGTAAAAGAGTGGTTCAAATTGGTAGGTTACAGTGCACTC

ATCAGAGGCTAGTCAGCTATGCATCTGCCCCCTTCTCCTCCGCCATGAGACCCCACTGGCGATCCAGAAGATTAAAGAAA

ACTACATATTGGATAAGTATCTATTCCCAGCTTTGTCTGGT

>KM089830.1/goat/China/2014-05-08

ACCAAACAAAGTTGGGTAAGGATAGATCTTATAATAACTATGGACTGGCAAACTTAGGAGTAAAGATCCTACTGTCGGGG

GGAGGAGGAGGAGCAAGATCTTTGATTATGGCGACTCTCCTTAAAAGCTTAGCATTGTTCAAAAGGAACAAAGACAAAGC

GCCGACAGCATCAGGTTCAGGAGGGGCCATCCGGGGGATTAAGAATGTTATCATAGTCCCGATTCCCGGAGACTCGTCCA

TCACTACCCGTTCAAGACTGCTCGACAGGCTTGTCAGATTGGCCGGAGATCCTGACATCAACGGGTCAAAGCTGACCGGG

GTGATGATCAGCATGTTATCCTTGTTCGTAGAGTCACCCGGGCAATTGATACAGCGAATCACAGATGATCCGGATGTTAG

TATCCGCCTTGTTGAGGTAGTTCAAAGTACTAGGTCCCAGTCCGGGTTGACCTTTGCATCACGTGGTGCTGATTTAGACA

ACGAGGCAGACATGTATTTTTCAACTGAGGGGCCCTCGAGTGGAGGTAAGAAAAGGATCAACTGGTTTGAGAACAGAGAA

ATAATAGACATAGAGGTGCAGGATCCAGAAGAGTTCAATATGTTGTTAGCCTCCATACTAGCACAAGTCTGGATCCTTCT

GGCCAAGGCTGTTACGGCACCAGATACGGCAGCTGACTCAGAACTGAGAAGGTGGGTTAAATACACACAACAAAGGAGAG

TGATTGGGGAATTTCGCCTTGACAAGGGGTGGCTGGACGCAGTCCGCAACAGAATTGCAGAAGATCTATCACTCCGGCGG

TTCATGGTATCTCTCATACTTGACATCAAGAGGACCCCAGGCAACAAGCCAAGGATTGCAGAAATGATCTGCGACATTGA

CAACTATATTGTCGAAGCAGGACTCGCCAGTTTCATCCTTACTATCAAATTTGGTATCGAAACCATGTATCCTGCATTAG

GGCTCCACGAGTTTGCCGGAGAATTGTCCACTATTGAGTCCTTGATGAACTTGTATCAACAGCTAGGAGAGGTTGCACCC

TACATGGTAATTCTAGAGAATTCAATTCAGAACAAGTTTAGTGCAGGAGCTTATCCCCTCCTCTGGAGCTATGCGATGGG

TGTCGGAGTCGAGCTGGAGAACTCAATGGGGGGCTTGAACTTTGGCAGGTCATATTTTGACCCTGCCTATTTTCGTCTCG

GACAGGAGATGGTCAGAAGATCTGCAGGAAAGGTCAGCTCTGTGATTGCGGCTGAGCTCGGCATCACAGCAGAGGAAGCT

AAACTTGTCTCGGAAATCGCCTCGCAGGCTGGGGACGAAAGAACCGCCAGAGGGACTGGGCCTCGACAGGCGCAGGTTTC

CTTCCTTCAGCACCAAACAGGAGGGGGAGAGTCGTCCGCACCAGCGACCAGAGAAGGGGTCAAAGCTGCGATCCCAAACG

GAGCTGAAGAAAGGGACAGAAAGCAAACACGCCCAGGAAGGCCCAGAGGAGAGACCTCCGGCCAACTGCTCCTGGACATC

ATGCCAGAGGATGAGATCTTGCGAGAGTCTGGTCAAAACCCTCGTGAGGCTCAAAGATCGGCCGAGGCACTCTTCAGGCT

GCAGGCCATGGCCAAGATTCTGGAGGACCAGGAGGAGGGAGAAGACAACAGTCAGGTCTACAACGACAAGGATCTCCTCG

GCTAAGCAGACGCACCCTCTGTCGAAATCAGTGACAAGACATCTCTTACCAGTATTATAAAAAACTTAGGACCCAGGTCC

AAGCAAGCACACATCGACACTCCAACCAGTCGAGCGGAGACCACCGATGGCAGAAGAACAAGCATACCATGTCAACAAGG

GGCTGGAATGTATCAAGTCTCTCAAAGCCTCACCCCCGGATCTATCCACCATCAGAGATACCATCGAGAGCTGGAGAGAG

GGGCTTAGCCCCTCGGGCCGTGCAACACCGAACCCTGATACGTCCGAGGGAGATCATCAGAATATCAACCAATCATGCTC

ACCAGCAATCGGACCAAACAAAGTCTACTTGTCTCCTGAAGATAATCTCGGATTTAGAGAGATCACTGGCAACGACTGTG

AGGCTGGGCTCGGAGGGGTCCAGAGAGAAGGATCCAACTCTCAAGTACAGCGTTACCATGTTTATAGTCACGGGGGTGAA

GAGATTGAAGGACTCGAGGATGCTGACTCTCTCGTGGTTCAAGCAGATCCTCCGGTTGCTAACGTCTTCAATGGAGGAGA

GGACGGATCTGACGACAGCGATGTGGACTCTGGCCCAGATGATCCCGGCAGAGATACTCTATATGACCGGGGATCTGTTG

CCGGCAATGGTATCGCTAGGTCCACAGATGTCGAAAAACTAGAAGGTGCTGATATTCAAGAAGTTCTTAACTCCCAGAAA

GGCAAAGGAGGAAGATTCCAGGGCGGGAAAACCTTGCGAGTCCCGGAAATACCCGATGTCAAGCACTCCAGACCATCAGC

CCAATCAATTAAAAAGGGCACAGACGGGAACTCAGTCTCATCTGGAACGGTGATAGAGTGTTTATCGATAAGTGGTGCAA

CCCAAACTGTGCCAGAGTCAAGATGGGAATCATCAGAGCAAAATGCGTCTGTGGGGAGTGTCCTCAAGTCTGCGAGGAGT

GCAAAGACGATCCAGGGGTCGACACAAGAATCTGGTACCATAGCATCACTGACTCAGCCTAAAGAGAATGACTCCGAGTA

TGAGTATGAGGATGACCTATTTACAGAGATTCAGGACATCCGTGCAAGCATTGCCAAGATCCATGATGACAATAAAACTA

TCCTCTCAAAGCTTGATTCTATACTGTTATTGAAAGGGGAAGTCGACACTATCAAGAAACAAATCAGCAAGCAGAATATA

AGTATATCCACCATTGAGGGCCATCTCTCCAGTATAATGATAGCCATCCCTGGCTTTGGGAAGGACATCAAGGACCCAAC

ATCCGAAGTCGAGTTGAACCCAGATCTAAGACCTATAATAAGCCGTGACTCTGGCAGAGCTCTCGCGGAGGTCCTCAAGA

AACCCGCTGTGGATAGGTCTCAGAAAATTGGAACCAAAGCCAACTCCAGCTCAAAGGGTCAGCTTCTTAAGGATCTCCAG

CTAAAACCTGTCGACAAGCAGGCAAGCTCTGCAATCGGGTTTGTCCCGTCCGACCATGAATCATCCAGAAATGTCATCCG

CTCCATAATCAAGTCGAGCAAGCTAAATATTGATCACAAGGACTATCTTCTAGATTTACTAAATGATGTGAAAGGCTCCA

AGGATCTTAAGGAATTTCACAAGATGCTAACAGCAATTCTCGCCAAGCACCCGTAACACATCCTCCAGTCATCATCTCAT

ACTCGACTAAAAACATCCTTTCAATCAGGCTATTACAAAAAACTTAGGAGCAAGGGCAACTGAGCTTCGCAGACAGGATG

ACCGAGATCTACGACTTCGATAAATCAGCATGGGATGTCAAAGGGTCAATTGCCCGCATAGAACCCACCACCTACCACGA

CGGCCGACTGATACCCCAGGTGAGGGTCATCGATCCTGGTCTGGGAGACAGAAAAGATGAGTGCTTTATGTACCTGTTTC

TCCTAGGAGTGATTGAGGATAACGACCCCCTGTCTCCCCCCGTCGGAAGAACCTTTGGCTCTCTACCTCTAGGGGTTGGT

AGGTCAACTGCCAAGCCAGAAGAACTACTAAGGGAGGCCACAGAATTAGATATAGTGGTGAGGCGCACTGCAGGAGTAAA

TGAGAAACTGGTATTTTACAACAACACTCCGCTGTCCTTGTTAACGCCCTGGAAGAAAGTTCTGACAACCGGAAGTGTGT

TTAGCGCTAACCAGGTTTGCAATGCAGTCAACCTAGTCCCGCTTGATACTCCCCAGAGATTCAGGGTTGTGTACATGAGC

ATAACTAGATTATCAGACAATGGTTACTATAGTGTGCCCAGAAGAATGCTGGAGTTCCGCTCGGCCAATGCAGTAGCCTT

CAACATCTTGGTTACACTAAGAATTGAAAATGGCACAAACCCTAGAAGATACATAGTCGGCTCATGGGAGAATTCAGAGG

TCACATTTATGGTACACGTGGGCAACTTTAGGAGAAAGAAGAACGAAGTATACTCTGCTGATTACTGCAAAATGAAGATT

GAAAAGATGGGTTTAGTTTTTGCCCTGGGTGGAATAGGCGGAACAAGTCTCCATATTAGAAGCACCGGGAAAATGAGCAA

AACCCTCCATGCACAGCTGGGGTTCAAGAAAATCTTATGTTACCCCCTAATGGATGTCAATGAGGATCTTAACCGATATC

TCTGGCGGGCAGAGTGCCGAATAGTCAAAATCCAAGCTGTCTTACAGCCATCAGTACCCCAAGAGTTCCGTGTCTACGAT

GATGTCATCATCAACGATGATCAAGGCTTGTTCAAGATCTTGTAGTTCATTTGCAAAGGAACGAACCCACCCCCAGGAAG

GGGCCCCACCCTCCCCCG-CCCAAAAACACACTCCAAGGCTCCCCGGCACAGCGCCAACACAGCTACTCCCGCGATGGAA

CGAGGTCGGCC-CCCGAACAAATCCGGTCCAGGGGGGGGGGCCCCCCCCCTGACCAACCGGGCGCCGACCGAGCAGGCCG

GCGAGAAGCCCCCAAGAGGCCCCGACGAAGACCGAGCACCCCATCCCCCC-AAAAACGTCCCCC-AACACGCCCGAGGCT

AGCCACCGGACACCCCCAGACCCGCCACCGGCGAGACAGAACCCCAAGGGGGGACCCTCCAAACCAAAGGAATCACCGGG

GTAGGCTCCGCCGAACCAGCACAGACCCGACCCAAGCAAGGCGGACCCCCAACACGAAACTGCCCAGCACACCCTACTCA

TCATCAAACAAAACTTAGGGGCCAAGTCCATACGAGAGGCCCTAGTCCGTCAGGCGGAGGGACGCAGAAAGGAAGGAGAC

ACGACCGTCCCAGAAGGAGGACACCCAACCACCGAAACGCCCGCCCCCAGGGAAGGAG--CCA-CCCCCACGCAGCGAAG

GAGCGGGTGGGAGACCTCGGGGAGGGCCCGACCGGGACCCGGGGGCCGGCGGCGCCGGCGGTGAGCAGGCCGGGCGCATA

CCTGACCCCACCGGAGGATCCGAGGAGGGCCCGACCCCGGCGGGACCAAACGGCAGGGAGCCCAACCCCACCAAAAAAAG

AGGCGAGGCCCCCCCTCCCTCCCCCCCCGCACACAACCCGCAGACAGACCGAGGGAGGGCCATCCCCCAGCGAGACAGGC

GGCTCCCTCGCAGCCGAGCCGCAGCTCCCCAGAATAGGACACCCCCGCACACTCCGAACACCCCCTCGCCCCCCGCCACC

GGGACAGGGGCTCCACCACCACAGACCGGAGCCACCGACAGCCAGCCAGAGGTCCCCTCCAGTCCCGCCCGGCGCCCCCC

CGAAGAGGCCCCGGCAAGGCCCCCCAGCCCGACGTCCCCCGCACGCACCGAAAGAAGGACCCGCCCGAGAACGTGCGGCA

CATCCATGCGCAAACATCATGACGCGGGTCGCAATTTTGACATTTCTGTTTCTTTTCCCAAATGTTGTTGCGTGTCAGAT

TCACTGGGGCAATCTATCCAAGATCGGGATTGTAGGAACAGGGAGTGCCAGCTACAAGGTGATGACTAGGCCAAGCCACC

AGACTCTGGTTATAAAGTTAATGCCAAATATAACGGCCATCGACAATTGTACAAAGTCAGAGATTGCAGAGTACAAGAGA

TTGCTGATCACAGTGTTAAAGCCTGTAGAGGATGCTCTGTCGGTGATAACCAAGAATGTAAGACCAATTCAAACTCTAAC

ACCTGGGCGTAGAACCCGCCGTTTTGCTGGAGCTGTTCTGGCCGGGGTAGCACTTGGAGTTGCGACAGCCGCTCAGATAA

CTGCAGGAGTCGCCCTTCATCAATCATTGATGAACTCCCAAGCAATTGAGAGTTTAAAAACCAGTCTTGAGATGTCGAAT

CAGGCAATAGAAGAAATCAGACTTGCAAATAAGGAGACCATACTGGCAGTACAGGGCGTCCAGGATTATATCAACAATGA

GCTCGTCCCTTCTGTTCATAGAATGTCATGCGAGCTGGTAGGTCACAAGCTCGGCCTCAAGCTCCTTAGGTACTACACCG

AGATCCTGTCCATATTCGGGCCCAGTCTTCGAGACCCGATATCTGCCGAAATATCAATCCAGGCACTTAGTTATGCATTA

GGCGGAGACATTAATAAAATCCTGGACAAGCTTGGGTATAGCGGTGGGGATTTCCTTGCCATCCTAGAAAGCAAGGGAAT

AAAGGCCCGGGTTACATATGTGGACGCAAGAGATTACTTTATAATCCTTAGCATCGCCTACCCAACCTTATCTGAGATCA

AGGGAGTGATAGTTCACAAGATAGAAGCTATAACATACAACATTGGGGCACAGGAGTGGTATACTACTATCCCTAAATAT

GTAGCCACTCAGGGGTATCTGATATCGAACTTTGATGAGACGTCATGCGTATTCACTCCAGAGGGGACAGTTTGCAGCCA

GAATGCGTTGTACCCAATGAGCCCATTGCTTCAGGAATGTTTCAGGGGGTCAACAAAATCGTGCGCCAGAACCTTAGTTT

CAGGGACCATAAGTAATAGATTTATCCTATCAAAAGGGAACCTGATTGCAAATTGTGCGTCAGTTTTGTGCAAATGTTAC

ACAACAGAGACAGTTATCAGCCAAGACCCTGACAAACTACTAACTGTTGTAGCATCCGACAAGTGTCCTGTAGTTGAGGT

GGATGGAGTGACAATACAGGTCGGCAGTCGAGAGTATCCGGATTCTGTATACTTACACAAAATAGACTTAGGTCCAGCCA

TCTCCCTAGAAAAACTGGATGTAGGCACCAATTTAGGCAATGCAGTCACAAGACTGGAGAATGCAAAGGAGCTCCTAGAT

GCATCAGACCAAATACTGAAGACTGTTAAAGGGGTACCTTTCAGTGGGAATATGTACATAGCACTGGCAGCTTGCATAGG

AGTATCCCTGGGCCTTGTCACATTAATATGCTGCTGTAAGGGGAGGTGTAAGAACAAGGAAATCCCTATCTCCAAAATCA

ACCCAGGGCTCAAACCCGACCTGACCGGGACCTCAAAGTCGTACGTAAGATCACTGTAGTCAGAATCACCTGGATCATCT

GGCATCGCACACATACATGCACGACACAGGCAGTCCGAGGACGCAAGAAACCCAGCCTCCGGTCACCCACCCGACCCCAC

TCCACGCTCCACCACACATTAGTCATCAAACAAAACTTAGGACGAAAGGTCAATCACCATGTCCGCACAAAGGGAGAGGA

TCAATGCCTTCTACAAAGACAATCCTCACAATAAGAACCATAGGGTGATCCTGGATAGAGAACGCTTGGTCATTGAAAGA

CCCTACATCTTGCTTGGAGTCCTGCTGGTAATGTTCCTGAGTCTAATCGGACTGCTGGCCATTGCAGGGATCAGGCTTCA

CCGGGCCACCGTTGGAACTTCAGAGATCCAGAGTCGGCTGAATACCAATATTAAGTTGACCGAATCTATTGATCACCAGA

CTAAGGATGTCTTAACTCCCCTTTTTAAAATCATTGGCGATGAAGTCGGCATCAGAATTCCACAGAAATTCAGTGATCTT

GTCAAGTTCATCTCCGATAAGATTAAATTCCTCAACCCTGATAGAGAGTATGATTTCAGAGATCTCCGGTGGTGTATGAA

TCCCCCCGAGAGAGTCAAAATTAATTTTGATCAGTTTTGTGAGTACAAGGCTGCGGTTAAGTCAATTGAACATATATTTG

AGTCACCACTCAACAAGTCAAAAAAGCTGCAATCTTTGACTCTCGGGCCCGGAACAGGCTGTCTAGGCAGGACAGTAACA

AGAGCCCATTTCTCAGAACTTACAATGACCTTAATGGACCTGGATCTAGAGATGAAGCACAACGTGTCCTCAGTGTTTAC

CGTAGTTGAAGAGGGATTATTCGGAAGAACATATACCGTCTGGAGATCCGATGCCAGGGATCCGAGCACCGATCTAGGTA

TCGGCCATTTTTTAAGAGTCTTCGAGATTGGACTGGTAAGAGATCTTGGGCTGGGTCCCCCTGTTTTTCATATGACCAAC

TATCTCACGGTGAACATGAGTGATGACTATCGGAGATGTCTTTTAGCGGTAGGGGAGTTGAAGTTGACAGCCCTATGCAC

CTCATCTGAGACTGTGACACTGAGTGAGAGAGGAGTTCCAAGGAGGGAACCTCTTGTGGTTGTGATACTTAATCTAGCTG

GACCCACTCTAGGGGGAGAGCTATACAGTGTCTTGCCTACCTCTGATCTCATGGTGGAGAAACTCTATTTGTCTTCACAT

AGAGGGATCATCAAAGATGATGAGGCCAATTGGGTAGTGCCGTCTACCGATGTTCGTGATCTTCAGAACAAAGGTGAATG

TCTGGTGGAAGCATGCAAGACTCGACCTCCTTCATTTAGCAATGGCACAGGATCAGGCCCGTGGTCAGAAGGGAGAATCC

CTGCCTGCGGGGTGATCAGGGTCAGTCTTGACTTAGCGAGTGACCCGGATGTAGTTATCACTTCAGTGTTTGGCCCACTG

ATACCTCACCTATCCGGCATGGATCTTTACAACAACCCGTTTTCAAAAGCTGTATGGTTGGCTGTACCACCTTATGAGCA

GTCATTTCTAGGAATGATAAATACAATTGGATTCCCTAACAGAGCAGAGGTTATGCCGCACATTTTGACCACAGAGATCA

GAGGCCCTCGGGGTCGTTGCCATGTTCCCATAGAATTGTCCCGCCGGGTTGATGACGATATCAAGATCGGGTCCCACATG

GTCATATTGCCGACGATGGACCTGAGGTATATTACAGCCACTTATGATGTTTCCAGGAGCGAGCATGCAATCGTGTACTA

TATCTATGACACGAGTCGCTCATCATCTTACTTCTACCCAGTTCGACTGAATTTCAAAGGCAATCCTCTCTCTTTGAGGA

TAGAGTGTTTCCCCTGGCGTCATAAGGTGTGGTGCTACCATGATTGTCTTATATACAACACCATAACAGGTGAAGAGGTC

CATACGAGAGGGCTGACCGGCATAGAGGTAACATGCAATCCAGTCTGAGCAGAGCTACGACCGTCACTCAGGCAGTCCCT

TGAGTCGCCACCGAGTCCAAGCAGCACAGCCTGGGACAATCAACGGCACAGCCCAGCCAACAATGTTATAAAAAACTTAG

GAGCCAGGGTTGTAGGGGCCATGGACTCTCTATCAGTCAATCAGGTCTTGTACCCTGAGGTTCATCTAGACAGCCCTATT

GTCACAAACAAACTAGTTGCCATCCTTGAGTACTCGGGTATCGACCATAACTATGTTCTTGAAGACCAGACTCTTGTCAA

GAATATTAGGTATAGGCTGGGGTGCGGTTTTTCAAATCAAATGATCATCAATAACAGGGGGGTGGGTGAAACAGTCAATT

CTAAACTTAAAAGTTACCCCCATAATCGTCATATCATATACCCGGGTTGCAATAAGGAGTTGTTTTGTATCAAGGATAGC

TGCATATCTAAGAAGCTCTCGGAGCTATTCAAGAAGGGTAATTCCTTGTACTCTAAGATAGGTCACCAGGTACTGGATTG

TCTTAAGAGAGTCAATGGGAAATTAGGTCTGGGCACAGATCTTACCCATGGTCTGAAGGAGGGTATCCTTGACCTAGGGT

TGCACATGCATAGCTCTCAATGGTTCGAGACTTTCCTGTTCTGGTTCACTATCAAGACAGAGATGAGATCAATGATCAAA

GAACAGTCCCATATATGCCACAAGAGGAGGTATAACCCGACTTTTGTGTCGGGGGATGCATTCGAGGTGCTCGTATCGCG

AGACCTCGTTGTGATAATTGATAAGAATACCCAGTATGTCTTCTACTTAACATTTGAGCTGGTCCTTATGTATTGTGATG

TCATAGAGGGTAGACTTATGACGGAGACAGCCATGGCCATAGACCAGAGATATTCAGAACTTCTAAGCCGGGTCAGATAC

TTGTGGGATCTTATTGATGGGTTCTTCCCAACACTAGGCAACACCACATACCAAATTGTTGCTCTGCTTGAACCATTATC

GTTGGCTTATCTTCAACTTCAGGATGTCACTCTGGGGTTAAGAGGTGCTTTTTTAGACCATTGCTTCAAAGAACTCTATG

AGATACTGGAACATTGCGGCATTGACACAGCAGGCACTTACAATTCCATCACTGAAGGATTGGATTACGTATTCATCACC

CACGATATACATCTAACTGGGGAGATTTTTTCATTTTTCCGGAGTTTCGGACACCCCCGTCTAGAAGCAGTCACCGCTGC

GGAAAATGTCAGAAAACATATGAACCAACCGAAGGTAATCAGTTATGAGACTATGATGAAAGGCCATGCGGTATTTTGCG

GGATAATCATAAATGGTTTTCGGGACCGACATGGCGGCAGCTGGCCCCCTGTTGCACTGCCAGAACATGCCTCCGCTGCG

ATCCGGAATGCGCAGGCATCAGGCGAGGGACTGACCCATGACCTGTGTATAGACAACTGGAAATCCTTTGTTGGATTCAA

ATTTGGCTGCTTTATGCCACTCAGCCTAGATAGTGATTTGACCATGTATCTCAAAGACAAGGCATTGGCTGCACTGAAGA

ATGAGTGGGATTCAGTTTACCCGAAAGAATACTTCCGTTATAACCCACCTAGAGGGACAGAGTCAAGGCGATTGGTAGAG

GTATTCCTGAATGACTCCAGCTTTGATCCTTATAACATGATAATGTACGTGGTGAATGGCTCCTACCTTAAAGACCCTGA

GTTTAACCTCTCATACAGCCTAAAAGAGAAGGAGATAAAGGAGACAGGGCGGTTGTTTGCCAAAATGGCCTATAAGATGC

GGGCCTGTCAGGTAATCGCTGAAAATCTGATATCGAATGGTGTTGGGAAGTATTTCCGAGACAATGGGATGGCAAAGGAC

GAGCATGACCTAACAAAAGCCCTTCACACCCTGGCAGTCTCAGGTGTTCCCAAAAACAACAAAGATAACCACCGAGGTGG

GCCTCCCAGAAGGACAACAAACCGAGGGGTGAGATCAAGCCAAGGCACCAAAACACAAGATAGAGACAAGGTTCAAGGGG

GACCTATGTACAACTATTTGCGATGCCAACCGATCAGCCCTGATCAGGGTGAGTCATACGAGACTGTTAGTGCATTCATC

ACCGCTGACCTTAAGAAGTATTGCCTGAATTGGAGATACGAGACAATCAGCATATTTGCACAGAGGCTGAATGAAATATA

TGGACTACCATCCTTCTTTCAATGGTTACACAGGATATTGGAAAAATCCGTACTCTACGTCAGTGACCCACATTGTCCTC

CCGATCTAGATAATCATATCCCTCTGGACAGTGTCCCCAATGCCCAAATATTCATCAAGTACCCAATGGGCGGAATAGAA

GGTTATTGCCAAAAACTATGGACAATCAGCACTATACCATACTTGTATCTGGCAGCCTATGAGAGCGGAGTAAGAATCGC

CTCACTGGTGCAGGGTGACAATCAGACAATCGCAGTGACAAAAAGAGTTCCAAGTTCTTGGCCTTATTCACTAAAAAAGA

GGGAGGCCTCCAAGGCCGCTCAAAATTACTTCGTAGTCTTAAGGCAAAGATTGCACGATGTAGGTCATCACTTAAAGGCT

AATGAGACCATAGTATCTTCTCACTTTTTTGTATATTCCAAAGGGATTTATTATGACGGCCTGCTAGTCTCACAATCACT

AAAGAGCATCGCCAGATGTGTCTTCTGGTCCGAGACTATAGTGGATGAAACCAGAGCGGCGTGCAGTAATATTGCAACAA

CTGTCGCTAAGAGTATAGAGAGGGGTTATGATAGGTACCTTGCATATTCTTTGAATATCCTCAAGATTTTCCAACAGATC

CTTATATCCCTTAACTTCACTATTAACACAACAATGACTCAGGATGTCGTGGCACCGATCATCGAGAACGGTGATTTGCT

GATAAGGATGGCACTCTTGCCAGCACCCATCGGGGGTCTCAATTATCTTAACATGAGCAGGTTATTTGTGAGAAATATCG

GTGACCCGGTCACCTCCTCTATAGCCGACCTGAAGAGGATGATAGATGCCGGGCTAATGCCAGAAGAAACATTGCATCAA

GTGATGACCCAGACCCCGGGAGAATCATCCTACCTAGATTGGGCAAGTGACCCTTATTCTGCCAACCTACCCTGCGTACA

GAGTATCACTCGCCTTCTCAAAAACATCACTGCACGATATATTTCAATCAGCAGCCCAAACCCGATGCTGAAAGGGTTAT

TTCATGAGGGGAGTAGGGATGAAGACGAGGAGCTTGCAAGTTTCCTAATGGATCGGCATATAATTGTTCCAAGAGCTGCA

CATGAAATCTTAGACCATAGTATAACCGGAGCAAGAGAAGCTATAGCCGGGATGTTGGACACCACCAAGGGTCTGATTAG

AACAAGTATGAAACGGGGCGGCCTCACCCCTCGAGTCTTAGCCCGCCTTTCCAATTATGATTATGAACAATTCCGATCCG

GGATAACACTATTAACAAAGAAAGGGCAGTGTTATCTCATTGACAAGGACTCGTGCTCGGTGCAGCTCGCTATAGCTCTG

AGGGGCCACATGTGGGCTAAGTTGGCTCGCGGGAGACCAATCTATGGGTTGGAGGTGCCTGATGTATTAGAATCGATGAT

CGGCTACCTTATCAAACGCCACGAGTCGTGTGCTATCTGTGAAACGGGCTCAAGTCACTACGGATGGTTTTTCGTCCCTG

CAGGGTGCCAGCATGACGATGTATCTAGAGAGACTTCGGCTCTTCGTGTACCTTATGTCGGATCAACAACTGAGGAAAGG

ACAGACATGAAACTTGCATTTGTTAGGTCTCCAAGTAGATCCCTCAAATCAGCAGTTAGAATTGCAACAGTTTACTCATG

GGCTTACGGGGATGATGGGAAATCATGGAGTGAAGCCTGGATGCTAGCTAGGCAGAGAGCCAATATCACCTTAGATGAAT

TGAGAATGATCACTCCAGTCTCCACATCCACCAACTTAGCCCACCGGTTGAGGGATCGGAGTACCCAGGTGAAATACTCG

GGGACATCCCTTGTGAGGGTTGCAAGATACACAACCATCTCCAATGATAACTTGTCGTTTGTGATATCTGAGAAAAAAGT

GGATACCAACTTCATTTATCAGCAAGGGATGTTGCTTGGTCTTGGGATCCTTGAGAACCTCTTTAGGCTAGAGGCCACCA

CCGGAGTATCCAACACAGTGCTACACCTGCACGTGGAAACAGAATGTTGTGTTGTACCTATGGTGGATCACCCAAGGATA

CCGAGTCTCCGTAATATAAAAGTTACGGATGAGCTATGCACAAACCCTTTGATCTACGACAGGTCCCCCATCATAGAGCA

CGATGCAACCCGGCTATATTCACAGAGCCACAGGAGACATTTGGTGGAGTTTGTTACCTGGTCAACAAGCCAGCTTTACC

ATATACTGGCTAAGTCTACAGCAATGTCCATGATTGAATTGATCACGAGATTCGCGAAAGATCACATGAATGAAATAGCC

GCCCTGATTGGCGATGACGACATCAACAGCTTCATCACAGAATTTTTGCTAGTGGAGCCTAGATTATTTATAGTTTACCT

TGGTCAGTGTGCTGCCATCAATTGGGCCTTTGATATACATTATCATCGGCCCTCGGGCAAGTACCAGATGGGGGAACTTC

TCTACTCTCTGCTCTCTCGGATGAGCAAAGGAGTATATAAGATCTTTACTAATGCTTTGAGCCACCCCAAAGTTTACAAG

AAATTTTGGCGAAGTGGCATCATTGAGCCGATTCATGGCCCATCCTTGGATACACAGAATTTACATGTCACTGTCTGTGA

CATGATATATGGGTCCTATGTCACCTATTTGGATCTTTTGCTGAATGATGAGCTAGATGATTACCCGTATTTGCTCTGCG

AGAGTGATGAGGACGTGGTCACAGACAGGTTCGACAACATTCAAGCCAAACATCTCTGTGTATTGGCCGATGTATATTGC

AGCTCCAAGAGGTGTCCTTCAATAATCGGGATGTCTCCTATAGAAAAATGTACCATTCTCACACATTACATCAAGGGAGA

ATCAGTACAATCCCCGTCTGGGACCTCATGGAACACTGATCCCCTTGTAGTAGATCATTACTCATGCTCTCTGACCTACC

TTCGCCGCGGTTCCATCAAACAAATCAGGTTGAGGGTGGATCCTGGGTTTGTATTCGAGGCGTTGACAGACGTCGACTTT

AAACAACCTCGCAAAGCTAAGTTAGATATATCGGTCGTTGGGTTGACTGATTTCTCTCCCCCTTACGACAACGTCGGTGA

TTTTTTAGGGACTATCAACACATTGAGGCACGATCTGCCTGTCACCGGAACCGGGGTCTCGAACTATGAAGTCCACGCTT

ATCGTAGAATTGGCCTGAATTCATCAGCATGTTATAAAGCCGTAGAGATCTCCACGTTAATCAAGCCATCCTTAGAAGTC

GGAGAGCATGGATTGTTCTTAGGAGAAGGTTCCGGTTCAATGCTGGCTGCGTACAAAGAAGTTCTTAAATTAGCAAATTG

TTACTACAACAGCGGAGTAACAGCGGAGGGCAGAGCCGGACAGAGGGAAATATCTCCCTATCCTTCAGAGATGAGCCTAG

TAGAGAATCAAATGGGGATAGAGCGGAGTGTTAAAGTGCTGTTCAACGGGAAACCTGAAGTAACTTGGGTGGGGACCACA

GATTGCTACAAGTACATAATCAGTAACATCCAAACCTCTAGTCTGGGTTTCATACATTCAGATATTGAGACGCTCCCAAC

CAAAGATGCTGTTGAGAAATTGGAGGAATTTGCCTCTATCCTATCCTTATCCTTGATTTTGGGGAAAATCGGCTCTATTA

CAGTTGTCAAAATTATGCCCATTAGCGGGGATTTTACCCAAGGCTTCATAGCCTATGCCATCCAATATTTTAGGGAGAGC

CTGCTTGCCTACCCGAGATATAGTAACTTCATCTCGACTGAGTGTTACCTTATTATGATAGGATTAAAGGCCAATCGATT

GATAAACCCAGAAGCCATTAAGCAAAGCATAATCAGAGCGGGGGTCAGGACTGCACCAGGACTTGTGTGCCATGTATTAT

CAGAGAAACAAAAAGGTTGTATTCAATCTCTCTTGGGTGATCCTTATATCCCAGGAGACTTCAATAAGCACCTTAAATCT

CTAACCTCTATTGAGAAAATCCTGGTAAATTGTGGTCTCTCGATCAATGGCATAAAAATCTGTAGAGATCTAATCCACCA

TGATATCGCCTCCGGTCCAGACGGCCTTATGAGCTCTACAATTATTTTATATAGGGAATTGGCTCATTTCAAAGAGAATA

TAAGGAGTCAGCACGGAATGTTCCACCCCTACCCGGTACTGGCCGACAGCAGGCAACGTGAATTAATCCTTCGAATAGCC

AAGAAATTCTGGGGGTATGTCTTGCTATATTCTGATGACCCGGCACTAATCAGACAAACAATCAAGAACTTGAAACGGAA

TCATCTAACCTTTGACTTACATAGTAACCCATTTATTAAAGGGCTATCCAAGGCTGAGAAACTGCTGGTGCGGACGAGTT

CACTAAGAAGAGAATGGTTGTTCACTCTCGAAACGAAAGAAGTAAAAGAGTGGTTCAAATTGGTAGGTTACAGTGCACTC

ATCAGAGGCTAGTCAGCTATGCATCTGCCCCCTTCTCCTCCGCCATGAGACCCCACTGGCGATCCAGAAGATTAAAGAAA

ACTACATATTGGATAAGTATCTATTCCCAGCTTTGTCTGGT

>KY888168.1/goat/Mongolia/2016-09

ACCAAACAAAGTTGGGTAAGGATAGATCTTATAATAACTATGGACTGGCAAACTTAGGAGTAAAGATCCTACCGTCGGGG

GGAGGAGGAGGAGCAAGATCCTTGATCATGGCGACTCTCCTTAAAAGCTTAGCATTGTTCAAAAGGAACAAAGACAAAGC

GCCGACAGCATCAGGTTCAGGAGGGGCCATCCGGGGGATTAAGAATGTTATCATAGTCCCGATTCCCGGAGACTCGTCCA

TCACTACCCGTTCAAGACTGCTCGACAGGCTTGTCAGATTGGCCGGAGATCCTGACATCAACGGGTCAAAGCTGACCGGG

GTGATGATCAGCATGTTATCCTTGTTCGTAGAGTCACCCGGGCAATTGATACAGCGAATCACAGATGATCCGGATGTTAG

TATCCGCCTTGTTGAGGTAGTTCAAAGTACTAGGTCCCAGTCCGGGTTGACCTTTGCATCACGTGGTGCTGATTTAGACA

ACGAGGCAGACATGTATTTTTCAACTGAGGGGCCCTCGAGTGGAGGTAAGAAAAGGATCAACTGGTTTGAGAACAGAGAA

ATAATAGACATAGAGGTGCAGGATCCAGAAGAGTTCAATATGTTGTTAGCCTCAATACTAGCACAAGTCTGGATCCTTCT

GGCCAAGGCTGTTACGGCACCAGATACGGCAGCTGACTCAGAACTGAGAAGGTGGGTTAAATACACACAACAAAGGAGAG

TGATTGGGGAATTTCGCCTTGACAAGGGGTGGCTGGACGCAGTCCGCAACAGAATTGCAGAAGATCTATCACTCCGGCGG

TTCATGGTATCTCTCATACTTGACATCAAGAGGACCCCAGGCAACAAGCCAAGGATTGCAGAAATGATCTGCGACATTGA

CAACTATATTGTCGAAGCAGGACTCGCCAGTTTCATCCTTACTATCAAATTTGGTATCGAAACCATGTATCCTGCATTAG

GGCTCCACGAGTTTGCCGGAGAATTGTCCACTATTGAGTCCTTGATGAACTTGTATCAACAGCTAGGAGAGGTTGCACCC

TACATGGTAATTCTAGAGAATTCAATTCAGAACAAGTTTAGTGCAGGAGCTTATCCCCTCCTCTGGAGCTATGCGATGGG

TGTCGGAGTCGAGCTGGAGAACTCAATGGGGGGCTTGAACTTTGGCAGGTCATATTTTGACCCTGCCTATTTTCGTCTCG

GACAGGAGATGGTCAGAAGATCTGCAGGAAAGGTCAGCTCTGTGATTGCGGCTGAGCTCGGCATCACAGCAGAGGAAGCT

AAACTAGTCTCGGAAATCGCCTCGCAGGCCGGGGACGAAAGAACCGCCAGAGGGACTGGGCCTCGACAGGCGCAGGTTTC

CTTCCTTCAGCACCAAACAGGAGGGGGAGAGTCGTCCGCACCAGCGACCAGAGAAGGGGTCAAAGCTGCGATCCCAAACG

GAGCTGAAGAAAGGGACAGAAAGCAAACACGCCCAGGAAGGCCCAGAGGAGAGACCTCCGGCCAACTGCTCCTGGACATC

ATGCCAGAGGATGAGATCTTGCGAGAGTCTGGTCAAAAACCTCGTGAGGCTCAAAGATCGGCCGAGGCACTCTTCAGGCT

GCAGGCCATGGCCAAGATTCTGGAGGACCAGGAGGAGGGAGAAGACAACAGTCAGGTCTACAACGACAAGGATCTCCTCG

GCTAAGCAGACGCACCCTCTGTCGAAATCAGTGACAAGACATCTCCTACCAGTATTATAAAAAACTTAGGACCCAGGTCC

AAGCAAGCACACATCGACACTCCAACCAGTCGAGCGGAGACCACCGATGGCAGAAGAACAAGCATACCATGTCAACAAGG

GGCTGGAATGTATCAAGTCTCTCAAAGCCTCACCCCCGGATCTATCCACCATCAGAGATACCATCGAGAGCTGGAGAGAG

GGGCTTAGCCCCTCGGGCCGTGCAACACCGAACCCTGATACGTCCGAGGGAGATCATCAGAATATCAACCAATCATGCTC

ACCAGCAATCGGACCAAACAAAGTCTACTTGTCTCCTGAAGATAATCTCGGATTTAGAGAGATCACTGGCAACGACTGTG

AGGCTGGGCTTGGAGGGGTCCAGAGAGAAGGATCCAACTCTCAAGTACAGCGTTACCATGTTTATAGTCACGGGGGTGAA

GAGATTGAAGGACTCGAGGATGCTGACTCTCTCGTGGTTCAAGCAGATCCTCCAGCTGCTAACGTCTTCAATGGAGGAGA

GGATGGATCTGACGACAGCGATGTGGACTCTGGCCCAGATGATCCCGGCAGAGATACTCTATATGACCGGGGATCTGTTG

CCGGCAATGGTATCGCTAGGTCCACAGATGTCGAAAAACTAGAAGGTGCTGATATTCAAGAAGTTCTTAACTCCCAGAAA

GGCAAAGGAGGAAGATTCCAGGGCGGGAAAACCTTGCGAGTCCCGGAAATACCCGATGTCAAGCACTCCAGACCATCAGC

CCAATCAATTAAAAAGGGCACAGACGGGAACTCAGTCTCATCTGGAACGGTGATAGAGTGTTTATCGATAAGTGGTGCAA

CCCAAACTGTGCCAGAGTCAAGATGGGAGTCATCAGAGCAAAATGCGTCTGTGGGGAGTGTCCTCAAGTCTGCGAGGAGT

GCAAAGACGATCCAGGGGCCGACACAAGAATCTGGTACCATAGCATCACTGACTCAGCCTAAAGAGAATGACTCCGAGTA

TGAGTATGAGGATGACCTATTTACAGAGATTCAGGACATCCGTGCAAGCATTGCCAAGATCCATGATGACAATAAAACTA

TCCTCTCAAAGCTTGATTCTATACTGTTATTGAAAGGGGAAGTCGACACTATCAAGAAACAAATCAGCAAGCAGAATATA

AGTATATCCACCATTGAGGGCCATCTCTCCAGTATAATGATAGCCATCCCTGGCTTTGGGAAGGACATCAAGGACCCAAC

ATCCGAAGTCGAGTTGAACCCAGATCTAAGACCTATAATAAGCCGTGACTCTGGCAGAGCTCTCGCGGAGGTCCTCAAGA

AACCCGCTGTGGATAGGTCTCAGAAAATTGGAACCAAAGCCAACTCCAGCTCAAAGGGTCAGCTTCTTAAGGATCTCCAG

CTAAAACCTGTCGACAAGCAGGCAAGCTCTGCAATCGGGTTTGTCCCGTCCGACCATGAATCATCCAGAAATGTCATCCG

CTCCATAATCAAGTCGAGCAAGCTAAATATTGATCACAAGGACTATCTTCTAGATTTACTAAATGATGTGAAAGGCTCCA

AGGATCTTAAGGAATTTCACAAGATGCTAACAGCAATTCTCGCCAAGCACCTGTAACACATCCCCCAGTCATCATCTCAT

ACTCGACTAAAAACATCCTTTCAATCAGGCTATTACAAAAAACTTAGGAGCAAGGGCAACTGAGCTTCGCAGACAGGATG

ACCGAGATCTACGACTTCGATAAATCAGCATGGGATGTCAAAGGGTCAATTGCCCGCATAGAACCCACCACCTACCACGA

CGGCCGACTGATACCCCAGGTGAGGGTCATCGATCCTGGTCTGGGAGACAGAAAAGATGAGTGCTTTATGTACCTGTTTC

TCCTAGGAGTGATTGAGGATAACGACCCCCTGTCTCCCCCCGTCGGAAGAACCTTTGGCTCTCTACCTCTAGGGGTTGGT

AGGTCAACTGCCAAGCCAGAAGAACTACTAAGGGAGGCCACAGAATTAGATATAGTGGTGAGGCGCACTGCAGGAGTAAA

TGAGAAACTGGTATTTTACAACAACACTCCGCTGTCCTTGTTAACGCCATGGAAGAAAGTTCTGACAACCGGAAGTGTGT

TTAGCGCTAACCAGGTTTGCAATGCAGTCAACCTAGTCCCGCTTGATACTCCCCAGAGATTCAGGGTTGTGTACATGAGC

ATAACTAGATTATCAGACAATGGTTACTATAGTGTGCCCAGAAGAATGCTGGAGTTCCGCTCGGCCAATGCAGTAGCCTT

CAACATCTTGGTTACACTAAGAATTGAAAATGGCACAAACCCTAGAAGATACATAGTCGGCTCATGGGAGAATTCAGAGG

TCACATTTATGGTACACGTGGGCAACTTTAGGAGAAAGAAGAACGAAGTATACTCTGCTGATTACTGCAAAATGAAGATT

GAAAAGATGGGTTTAGTGTTTGCCCTGGGTGGAATAGGCGGAACAAGTCTCCATATTAGAAGCACCGGGAAAATGAGCAA

AACCCTCCATGCACAGCTGGGGTTCAAGAAAATCTTATGTTACCCCCTAATGGATGTCAATGAGGATCTTAACCGATATC

TCTGGCGGGCAGAGTGCCGAATAGTCAAAATCCAAGCTGTCTTACAGGTATCCATACCCCAAGAGTTCCGTGTTTACGAT

GATGTCATCATCAACGATGATCAAGGCTTGTTCAAGATCTTGTAGTTCATTTGCAAAGGAACGAACCCACCCCCAGGAAG

GGGCCCCGCCCTCCCCCG-CCAAAAAACACACTCCAAGGCTCCCCGGCACAGCGCCAACACAGCTACTCCCGCGATGGAA

CGAGGTCGGCC-CCCGAACAAATCCGGTCCAAAGAGGGGGGCCCCCCCCCTGACCAACCGGGCGCCGACCGAGCAGGCCG

GCGAGAAGCCCCCAAGAGGCTCCGACGAAGACCGGGCACCCCATCCCCCC-AAAAACGTCCCCC-AACACGCCCGAGGCT

AGCCACCGGACACCCCCAGACCCGCCACCGGCGAGACAGAACCCCAAGGGGGGACCCTCCAAACCAAAGGAACCACCGGG

GTAGGCTCCGCCGAACCAGCACAGACCCGACCCAAGCAAGGCGGACCCCCAACACGAAACTGCCCAGCACACCCTACTCA

TCATCAAACAAAACTTAGGGGCCAAGTCCACACGAGAGGCCCCAGTCCGTCAGGCGGAGGGACGCAGAAAGGAAGGAGAC

ACGACCGTCCCAGAAGGAGGACACCCAACCACCGAAACGCCCGCCCCCAGGGAAGGAG--CCA-CCCCCACGCAGCGAAG

GAGCGGGCGGGAGACCCCGGGGAGGGCCCGACCGGGACCCGGGGGCCGGCGGCGCCGGCGGTGAGCAGGCCGGGCGCACA

CCCGACCCCACCGGAGGATCCGAGGAGGGCCCGACCCCGCGCGGACCAAACGGCAGGGAGCCCAACCCCACCAAAAAGAG

AGGCGAGGCCCCCCCTCCCTCCCCCCCCGCACACAACCCGCAGACAGACCGAGGGAGGGCCATCCCCCAGCGAGACAGGC

TGCTCCCTCGCAGCCGAGCCGCAGCTCCCCAGAATAGGACACCCCCGCACACTCCGAACACCCCCCCGCCCCCCGCCACC

GGGACAAGGGCTCCACCACCACAGACCGGAGCCACCGACAGCCAGCCAGAGGTCCCCTCCAGTCCCGCCCGGCGCCCCCC

CGAAGAGGCCCCGGCAAGGCCCCCCAGCCCGACGTCCCCCGCACGCACCGAAAGAAGGACCCGCCCGAGAAGGTGCGGCA

CATCCATGCGCAAACATCATGACGCGGGTCGCAATTTTGACATTTCTGTTTCTTTTCCCAAATGTTGTTGCGTGCCAGAT

TCACTGGGGCAATCTATCCAAGATCGGGATTGTAGGAACAGGGAGTGCCAGCTACAAGGTGATGACTAGGCCAAGCCACC

AGACTCTGGTTATAAAGTTAATGCCAAATATAACGGCCATCGACAATTGTACAAGGTCAGAGATTGCAGAGTACAAGAGA

TTGCTGATCACAGTGTTAAAGCCTGTAGAGGATGCCCTGTCGGTGATAACCAAGAATGTAAGACCAATTCAAACTCTAAC

ACCTGGGCGTAGAACCCGCCGTTTTGCTGGAGCTGTTCTGGCCGGGGTAGCACTTGGAGTTGCGACAGCCGCTCAGATAA

CTGCAGGAGTCGCCCTTCATCAATCATTGATGAACTCCCAAGCAATTGAGAGTTTAAAAACCAGTCTTGAGATGTCGAAT

CAGGCAATAGAAGAAATCAGACTTGCAAATAAGGAGACCATACTGGCAGTACAGGGCGTCCAGGATTATATCAACAATGA

GCTCGTCCCTTCTGTTCATAGAATGTCATGCGAGCTGGTAGGTCACAAGCTCGGCCTCAAGCTCCTTAGGTACTACACCG

AGATCCTGTCCATATTCGGGCCCAGTCTTCGAGACCCGATATCTGCCGAAATATCAATCCAGGCACTTAGTTATGCATTA

GGCGGAGACATTAATAAAATCCTGGACAAGCTTGGGTATAGCGGTGGGGATTTCCTTGCCATCCTAGAAAGCAAGGGAAT

AAAGGCCCGGGTTACATATGTGGACACAAGAGATTACTTTATAATCCTTAGCATCGCCTACCCAACCTTATCTGAGATCA

AGGGAGTGATAGTTCACAAGATAGAAGCTATAACATACAACATTGGGGCACAGGAGTGGTATACTACTATCCCTAAATAT

GTAGCCACTCAGGGGTATCTGATATCGAACTTTGATGAGACGTCATGCGTATTCACTCCAGAGGGGACAGTTTGCAGCCA

GAATGCGTTGTACCCAATGAGCCCATTGCTTCAGGAATGTTTCAGGGGGTCAACAAAATCGTGCGCCAGAACCTTAGTTT

CAGGGACCATAAGTAATAGATTTATCCTATCAAAAGGGAACCTGATTGCAAATTGTGCGTCAGTTTTGTGCAAATGTTAC

ACAACAGAGACAGTTATCAGCCAAGATCCTGACAAACTACTAACTGTTGTAGCATCCGACAAGTGTCCTGTAGTTGAGGT

GGATGGAGTGACAATACAGGTCGGCAGTCGAGAGTATCCGGACTCTGTATACTTACACAAAATAGACTTAGGTCCAGCCA

TCTCCCTAGAAAAACTGGATGTAGGCACCAATTTAGGCAATGCAGTCACAAGACTGGAGAATGCAAAGGAGCTCCTAGAT

GCATCAGACCAAATACTGAAGACTGTTAAAGGGGTACCTTTCAGTGGGAATATGTACATAGCACTGGCAGCTTGCATAGG

AGTATCCCTGGGCCTTGTCACATTAATATGCTGCTGTAAGGGGAGGTGTAAGAACAAGGAAATCCCTATCTCCAAAATCA

ACCCAGGGCTCAAACCCGACCTGACCGGGACCTCAAAGTCGTACGTAAGATCACTGTAGTCAGAATCACCTGGATCATCT

GGCATCACACACATACATGCACGACACAGGCAGTCCGATGACGCAAGAAACCCAACCTCCGGTCACCCACCCGACCCCAC

TCCACGCTCCACCACACATTAGTCATCAAACAAAACTTAGGACGAAAGGTCAATCACCATGTCCGCACAAAGGGAGAGGA

TCAATGCCTTCTACAAAGACAATCCTCACAATAAGAACCATAGGGTGATCCTGGATAGAGAACGCTTGGTCATTGAAAGA

CCCTACATCTTGCTTGGAGTCCTGCTGGTAATGTTCCTGAGTCTAATCGGACTGCTGGCCATTGCAGGGATCAGGCTTCA

CCGGGCCACCGTTGGAACTTCAGAGATCCAGAGTCGGCTGAATACCAATATTAAGTTGACCGAATCTATTGATCACCAGA

CTAAGGATGTCTTAACTCCCCTTTTTAAAATCATTGGCGATGAAGTCGGCATCAGAATTCCACAGAAATTCAGTGATCTT

GTCAAGTTCATCTCCGATAAGATTAAATTCCTCAACCCTGATAGAGAGTATGATTTCAGAGATCTCCGGTGGTGTATGAA

TCCTCCCGAGAGAGTCAAAATTAATTTTGATCAGTTTTGTGAGTACAGGGCTGCGGTTAAGTCAATTGAACATATATTTG

AGTCACCACTCAACAAGTCAAAAAAGCTGCAATCTTTGACTCTCGGGCCCGGAACAGGCTGTCTAGGCAGGACAGTAACA

AGAGCCCATTTCTCAGAACTTACAATGACCTTAATGGACCTGGATCTAGAGATGAAGCACAACGTGTCCTCAGTGTTTAC

CGTAGTTGAAGAGGGATTATTCGGAAGAACATATACCGTCTGGAGATCCGATGCCAGGGATCCGAGCGCCGATCTAGGTA

TTGGCCATTTTTTAAGAGTCTTCGAGATTGGACTGGTAAGAGATTTTGGGCTGGGTCCCCCTGTTTTTCATATGACCAAC

TATCTCACGGTGAACATGAGTGATGACTATCGGAGATGTCTTTTAGCGGTAGGGGAGTTGAAGTTGACAGCCCTATGCAC

CTCATCTGAGACTGTGACACTGAGTGAGAGAGGAGTTCCAAGGAGGGAACCTCTTGTGGTTGTGATACTTAATCTAGCTG

GACCCACTCTAGGGGGAGAGCTATACAGTGTCTTGCCTACCTCTGATCTCATGGTGGAGAAACTCTATTTGTCTTCACAT

AGAGGGATCATCAAAGATGATGAGGCCAATTGGGTAGTGCCGTCTACCGATGTTCGTGATCTTCAGAACAAAGGTGAATG

TCTGGTGGAAGCATGCAAGACTCGACCTCCTTCATTTTGCAATGGCACAGGATCAGGCCCGTGGTCAGAAGGGAGAATCC

CTGCCTACGGGGTGATCAGGGTCAGTCTTGACTTAGCGAGTGACCCGGATGTAGTTATCACTTCAGTGTTTGGCCCACTG

ATACCTCACCTATCCGGCATGGATCTTTACAACAACCCGTTTTCAAAAGCTGTATGGTTGGCTGTACCACCTTATGAGCA

GTCATTTCTAGGAATGATAAATACAATTGGATTCCCTAACAGAGCAGAGGTTATGCCGCACATTTTGACCACAGAGATCA

GAGGCCCTCGGGGTCGTTGCCATGTTCCCATAGAATTGTCCCGCCGGGTTGATAACGATATCAAGATCGGGTCCAACATG

GTCATATTGCCGACGATGGACCTGAGGTATATTACAGCCACTTATGATGTTTCCAGGAGCGAGCATGCAATCGTGTACTA

TATCTATGACACGAGTCGCTCATCATCTTACTTCTACCCAGTTCGACTGAATTTCAAAGGCAATCCTCTCTCTTTGAGGA

TAGAGTGTTTCCCCTGGCGTCATAAGGTGTGGTGCTACCATGATTGTCTTATATACAACACCATAACAGGTGAAGAGGTC

CATACGAGAGGGCTGACCGGCATAGAGGTAACATGCAATCCAGTCTGAGCAGAGCTACGACCGTCACTCAGGCAGTCCCT

TGAGTCGCCACCGAGTCCAAGCAGCACAGCCTGGGACAATCAACGGCACAGCCCAGCCAACAATGTTATAAAAAACTTAG

GAGCCAGGGTTGTAGGGGCCATGGACTCTCTATCAGTCAATCAGGTCTTGTACCCTGAGGTTCATCTAGACAGCCCTATT

GTCACAAACAAACTAGTTGCCATCCTTGAGTACTCGGGTATCGACCATAACTATGTTCTTGAAGACCAGACTCTTGTCAA

GAATATTAGGTATAGGCTGGGGTGCGGTTTTTCAAATCAAATGATCATCAATAACAGGGGGGTGGGTGAAACAGTCAATT

CTAAACTTAAAAGTTACCCCCATAATCGTCATATCATATACCCGGATTGCAATAAGGAGTTGTTTTGTATCAAGGATAGC

TGCATATCTAAGAAGCTCTCGGAGCTATTCAAGAAGGCTAATTCCTTGTACTCTAAGATAAGTCACCAGGCACTGGATTG

TCTTAATAGAGTCAATGGGAAATTAGGTCTGGGCACAGATCTTACCCATGGTCTGAAGGAGGTTATCCTTGACCTAGGGT

TGCACATGCATAGCTCTCAATGGTTCGAGACTTTCCTGTTCTGGTTCACTATCAAGACAGAGATGAGATCAATGATCAAA

GAACAGTCCCATATATGCCACAAGAGGAGGTATAACCCGACTTTTGTGTCGGGGGATGCATTCGAGGTGCTCGTATCGCG

AGACCTCGTTGTGATAATTGATAAGAATACCCAGTATGTCTTCTACTTAACATTTGAGCTGGTCCTTATGTATTGTGATG

TCATAGAGGGTAGACTTATGACGGAGACAGCCATGGCCATAGACCAGAGATATTCAGAACTTCTAAGCCGGGTCAGATAC

TTGTGGGATCTTATTGATGGGTTCTTCCCAACACTAGGCAACACCACATGCCAAATTGTTGCTCTGCTTGAACCATTATC

GTTGGCTTATCTTCAACTTCAGGATGTCACTCTGGAGTTAAGAGGTGCTTTTTTAGACCATTGCTTCAAAGAACTCTATG

AGATACTGGAACATTGCGGCATTGACACAGCAGGCACTTACAATTCCATCACTGAAGGATTGGATTACGTATTCATCACC

CACGATATACATCTAACTGGGGAGATTTTTTCATTTTTCCGGAGTTTCGGACACCCCCGTCTAGAAGCAGTCACCGCTGC

GGAAAATGTCAGAAAACATATGAACCAACCGAAGGTAATCAGTTATGAGACTATGATGAAGGGCCATGCGGTATTTTGCG

GGATAATCATAAATGGTTTCCGGGACCGACATGGCGGCAGCTGGCCCCCTGTTGCACTGCCAGAACATGCCTCTGCTGCG

ATCCGGAATGCGCAGGCATCAGGTGAGGGACTGACCCATGACCTGTGTATAGACAACTGGAAATCCTTTGTTGGATTCAA

ATTTGGCTGCTTTATGCCACTCAGCCTAGATAGTGATTTGACCATGTATCTCAAAGACAAGGCATTGGCTGCACTGAAGA

ATGAGTGGGATTCAGTTTACCCGAAAGAATACCTCCGTTATAACCCACCTAGAGGGACAGAGTCAAGGCGATTGGTAGAG

GTATTCCTGAATGACTCCAGCTTTGATCCTTATAACATGATAATGTACGTGGTGAATGGCTCCTACCTTAAAGACCCTGA

GTTTAACCTCTCATACAGCCTAAAAGAGAAGGAGATAAAGGAGACAGGGCGGTTGTTTGCCAAAATGACCTATAAGATGC

GGGCCTGTCAGGTAATCGCTGAAAATCTGATATCGAATGGTGTTGGGAAGTATTTCCGAGACAATGGGATGGCAAAGGAC

GAGCATGACCTAACAAAAGCCCTTCACACCCTGGCAGTCTCAGGTGTTCCCAAAAACAACAAAGATAACCACCGAGGTGG

GCCTCCCAGAAGGACAACAAACCGAGGGGTGAGATCAAGCCAAGGCACCAAAACACAAGATAGAGACAAGGTTCAAGGGG

GACCTATGTACAACTATTTGCGATGCCAACCGATCAGCCCTGATCAGGGTGAGTCATACGAGACTGTTAGTGCATTCATC

ACCGCTGACCTTAAGAAGTATTGCCTGAATTGGAGATACGAGACAATCAGCATATTTGCACAGAGGCTGAATGAAATATA

TGGACTACCATCCTTCTTTCAATGGTTACACAGGATATTGGAAAAATCCGTACTCTACGTCAGTGACCCACATTGTCCTC

CCGATCTAGATAATCATATCCCTCTGGACAGTGTCCCCAATGCCCAAATATTCATCAAGTACCCAATGGGCGGAATAGAA

GGTTATTGCCAAAAACTATGGACAATCAGCACTATACCATACTTGTATCTGGCAGCCTATGAGAGCGGAGTAAGAATCGC

CTCACTGGTGCAGGGTGACAATCAGACAATCGCAGTGACAAAAAGAGTTCCAAGTTCTTGGCCTTATTCACTAAAAAAGA

GGGAGGCCTCCAAGGCCGCTCAAAATTACTTCGTAGTCTTAAGGCAAAGATTGCACGATGTAGGTCATCACTTAAAGGCT

AATGAGACCATAGTATCTTCTCACTTTTTTGTATATTCCAAAGGGATTTATTATGACGGCCTGCTAGTCTCACAATCACT

AAAGAGCATCGCCAGATGTGTCTTCTGGTCCGAGACTATAGTGGATGAAACCAGAGCGGCGTGCAGTAATATTGCAACAA

CTGTCGCTAAGAGTATAGAGAGGGGTTATGATAGGTACCTTGCATATTCTTTGAATATCCTCAAGATTTTCCAACAGATC

CTTATATCCCTTAACTTCACTATTAACACAACAATGACTCAGGATGTCGTGGCACCGATCATCGAGAACGGTGATTTGCT

GATAAGGATGGCACTCTTGCCAGCACCCATCGGGGGTCTCAATTATCTTAACATGAGCAGGTTATTTGTGAGAAATATCG

GTGACCCGGTCACCTCCTCTATAGCCGACCTGAAGAGGATGATAGATGCCGGGCTAATGCCAGAAGAAACATTGCATCAA

GTGATGACCCAGACCCCGGGAGAATCATCTTACCTAGATTGGGCAAGTGACCCTTATTCTGCCAACCTACCCTGCGTACA

GAGTATCACTCGCCTTCTCAAAAACATCACTGCACGATATATTTTAATCAGCAGCCCAAACCCGATGCTGAAAGGGTTAT

TTCATGAGGGGAGTAGAGATGAAGACGAGGAGCTTGCAAGTTTCCTAATGGATCGGCATATAATTGTTCCAAGAGCTGCA

CATGAAATCTTAGACCATAGTATAACCGGAGCAAGAGAAGCTATAGCCGGGATGTTGGACACCACCAAGGGTCTGATTAG

AACAAGTATGAAACGGGGCGGCCTCACCCCTCGAGTCTTAGCCCGCCTTTCCAATTATGATTATGAACAATTCCGATCCG

GGATAACACTATTAACAAAGAAAGGGCAGTGTTATCTCATTGACAAGGACTCGTGCTCGGTGCAGCTCGCTATAGCTCTG

AGGGGCCACATGTGGGCTAAGTTGGCTCGCGGGAGACCAATCTATGGGTTGGAGGTGCCTGATGTATTAGAATCGATGAA

CGGCTACCTTATTAAACGCCACGAGTCGTGTGCTATCTGTGAAACGGGCTCAAGTCACTACGGATGGTTTTTCGTCCCTG

CAGGGTGCCAGCTTGACGATGTATCTAGAGAGACTTCGGCTCTTCGTGTACCTTATGTCGGATCAACAACTGAGGAAAGG

ACAGACATGAAACTTGCATTTGTTAGGTCTCCAAGTAGATCCCTCAAATCAGCAGTTAGAATTGCAACAGTTTACTCATG

GGCTTACGGGGATGATGAGAAATCATGGAGTGAAGCCTGGATGCTAGCAGGGCAGAGAGCCAATATCACCTTAAATGAAT

TGAGAATGATCACTCCAGTCTCCACATCCACCAACTTAGCCCACCGGTTGAGGGATCGGAGTACCCAGGTGAAATACTCG

GGGACATCCCTTGTGAGGGTTGCAAGATACACAACCATCTCCAATGATAACTTGTCGTTTGTGATATCTGAGAAAAAAGT

GGATACCAACTTCATTTATCAGCAAGGGATGTTGCTTGGTCTTGGGATCCTTGAGAACCTCTTTAGGCTAGAGGCCACCA

CCGGAGTATCCAACACAGTGCTACACCTGCACGTGGAAACAGAATGTTGTGTTGTACCTATGTTGCATCACCCAAGGATA

CCGAGTCTCCGTAATATAAAAGTTACGGATGAACTATGCACAAACCCTTTGATCTACGACAGGTCCCCCATCATAGAGCA

CGATGCAACCCGGCTATATTCACAGAGCCACAGGAGACATTTGGTGGAGTTTGTTACCTGGTCAACAAGCCAGCTTTACC

ATATACTGGCTAAGTCTACAGCAATGTCCATGATTGAATTGATCACGAGATTCGAGAAAGATCACATGAATGAAATAGCC

GCCCTGATTGGCGATGACGACATCAACAGCTTCATCACAGAATTTTTGCTAGTGGAGCCTAGATTATTTATAGTTTACCT

TGGTCAGTGTGCTGCCATCAATTGGGCCTTTGATATACATTATCATCGGCCCTCGGGCAAGTACCAGATGGGGGAACTTC

TCTACTCTCTGCTCTCTCGGATGAGCAAAGGAGTATATAAGATCTTTACTAATGCTTTGAGCCACCCCAAAGTTTACAAG

AAATTTTGGCGAAGTGGCATCATTGAGCCGATTCATGGACCATCCTTGGATACACAGAATTTACATGTCACTGTCTGTGA

CATGATATATGGGTCCTATGTCACCTATTTGGATCTTTTGCTGAATGATGAGCTAGATGATTACCCGTATTTGCTCTGCG

AGAGTGATGAGGACGTGGTCACAGACAGGTTCGACAACATTCAAGCCAAACATCTCTGTGTATTGGCCGATGTATATTGC

AGCTCCAAGAGGTGTCCTTCAATAATCGGGATGTCTCCTATAGAAAAATGTACCATTCTCACACATTACATCAAGGGAGA

ATCAGTACAATCCCCGTCTGGGACCTCATGGAACACTGATCCCCTTGTAGTAAATCATTACTCATGCTCTCTGACCTACC

TTCGCCGCGGTTCCATCAAACAAATCAGGTTGAGGGTGGATCCTGGGTTTGTATTCGAGGCGTTGACAGACGTCGACTTT

AAACAACCTCGCAAAGCTAAGTTAGATATATCGGTCGTTGGGTTGACTGATTTCTCTCCCCCTTACGACAACGTCGGTGA

TTTTTTAGGGACTATCAACACATTGAGGCACGATCTGCCTGTCACCGGAACCGGGGTCTCGAACTATGAAGTCCACGCTT

ATCGTAGAATTGGCCTGAATTCATCAGCATGTTATAAAGCCGTAGAGATCTCCACGTTAATCAAGCCATCCTTAGAAGTC

GGAGAGCATGGATTGTTCTTAGGAGAAGGTTCCGGTTCAATGCTGGCTGCGTACAAAGAAGTTCTTAAATTAGCAAATTG

TTACTACAACAGCGGAGTAACAGCGGAGGGCAGAGCCGGACAGAGGGAAATATCTCCCTATCCTTCAGAGATGAGCCTAG

TAGAGAATCAAATGGGGATAGAGCGGAGCGTTAAAGTGCTGTTCAACGGGAAACCTGAAGTAACTTGGGTGGGGACCACA

GATTGCTACAAGTACATAATCAGTAACATCCAAACCTCTAGTCTGGGTTTCATACATTCAGATATTGAGACGCTCCCAAC

CAAAGATGCTGTTGAGAAATTGGAGGAATTTGCCTCTCTATCATCCTTATCCTTGATTTTGGGGAAAATCGGCTCTATTA

CAGTTGTCAAAATTATGCCCATTAGCGGGGATTTTACCCAAGGCTTCATAGCCTATGCCATCCAATATTTTAGGGAGAGC

CTGCTTGCCTATCCGAGATATAGTAACTTCATCTCGACTGAGTGTTACCTTATTATGATAGGATTAAAGGCCAATCGATT

AATAAACCCAGAAGCCATTAAGCAAAGCATAATCAGAGCGGGGGTCAGGACTGCACCAGGACTTGTGAGCCATGTATTAT

CAGAGAAACAAAAAGGTTGTATTCAATCTTTCTTGGGTGATCCTTATATCCAAGGAGACTTCAATAAGCACCTTAAATCT

CTAACCTCTATTGAGAAAATCCTGGTAAATTGTGGTCTCTCGATCAATGGCATAAAAATCTGTAGAGATCTAATCCACCA

TGATATCGCCTCCGGTCCAGACGGCCTTATGAGCTCTACAATTATTTTATATAGGGAATTGGCTCATTTCAAAGATAATA

TAAGGAGTCAGCACGGAATGTTCCACCCCTACCCGGTACTGGCCAACAGCAGGCAACGTGAATTAATCCTTCGAATAGCC

AAGAAATTCTGGGGGTATGTCTTGCTATATTCTGATGACCCGGCACTAATCAGACAAACAATCAAGAACTTGAAACGGAA

TCATCTAACCTTTGACTTACATAGTAACCCATTTATTAAAGGGCTATCCAAGGCTGAGAAACTGCTGGTGCGGACGAGTT

CACTAAGAAGAGAATGGTTGTTCACTCTCGAAACGAAAGAAGTAAAAGAGTGGTTCAAATTGGTAGGTTACAGTGCACTC

ATCAGAGGCTAGTCAGCTATGCATCTGCCCCCTTCTCCTCCGCCATGAGACCCCACTGGCGATCCAGAAGATTAAAGAAA

ACTACATATTGGATAAGTATCTATTCCCAGCTTTGTCTGGT

>KP260624.1/goat/China/2014-08-16

ACCAAACAAAGTTGGGTAAGGATAGATCTTATAATAACTATGGACTGGCAAACTTAGGAGTAAAGATCCTACTGTCGGGG

GGAGGAGGAGGAGCAAGATCCTTGATCATGGCGACTCTCCTTAAAAGCTTAGCATTGTTCAAAAGGAACAAAGACAAAGC

GCCGACAGCATCAGGTTCAGGAGGGGCCATCCGGGGGATTAAGAATGTTATCATAGTCCCGATTCCCGGAGACTCGTCCA

TCATTACCCGTTCAAGACTGCTCGACAGGCTTGTCAGATTGGCCGGAGATCCTGACATCAACGGGTCAAAGCTGACCGGG

GTGATGATCAGCATGTTATCCTTGTTCGTAGAGTCACCCGGGCAATTGATACAGCGAATCACAGATGATCCGGATGTTAG

TATCCGCCTTGTTGAGGTAGTTCAAAGTACTAGGTCCCAGTCCGGGTTGACCTTTGCATCACGTGGTGCTGATTTAGACA

ACGAGGCAGACATGTATTTTTCAACTGAGGGGCCCTCGAGTGGAGGTAAGAAAAGGATCAACTGGTTTGAGAACAGAGAA

ATAATAGACATAGAGGTGCAGGATCCAGAAGAGTTCAATATGTTGTTAGCCTCCATACTAGCACAAGTCTGGATCCTTCT

GGCCAAGGCTGTTACGGCACCAGATACGGCAGCTGACTCAGAACTGAGAAGGTGGGTTAAATACACACAACAAAGGAGAG

TGATTGGGGAATTTCGCCTTGACAAGGGGTGGCTGGACGCAGTCCGCAACAGAATTGCAGAAGATCTATCACTCCGGCGG

TTCATGGTATCTCTCATACTTGACATCAAGAGGACCCCAGGCAACAAGCCAAGGATTGCAGAAATGATCTGCGACATTGA

CAACTATATTGTCGAAGCAGGACTCGCCAGTTTCATCCTTACTATCAAGTTTGGTATCGAAACCATGTATCCTGCATTAG

GGCTCCACGAGTTTGCCGGAGAATTGTCCACTATTGAATCCTTGATGAACTTGTATCAACAGCTAGGAGAGGTTGCACCC

TACATGGTAATTCTAGAGAATTCAATTCAGAACAAGTTTAGTGCAGGAGCTTATCCCCTCCTCTGGAGCTATGCGATGGG

TGTCGGAGTCGAGCTGGAGAACTCAATGGGGGGCTTGAACTTTGGCAGGTCATATTTTGACCCTGCCTATTTTCGTCTCG

GACAGGAGATGGTCAGAAGATCTGCAGGAAAGGTCAGCTCTGTGATTGCGGCCGAGCTCGGCATCACAGCAGAGGAAGCT

AAACTAGTCTCGGAAATCGCCTCGCAGGCTGGGGACGAAAGAACCGCTAGAGGGACTGGGCCTCGACAGGCGCAGGTCTC

CTTCCTTCAGCACAAAACAGGAGAGGGAGAGTCGTCCGCACCAGCAACCAGAGAGGGGGTCAAAGCTGCGATCCCAAACG

GAGCCGAAGAAAGGGACAGGAAGCAAACACGCTCAGGAAGGCCCAGAGGAGAGACCTCCGGCCAACTGCTCCTGGACATC

ATGCCAGAGGATGAGATCTTGCGAGAGTCTGGTCAAAACCCTCGTGAGGCTCAAAGATCGGCCGAGGCACTCTTCAGGCT

GCAGGCCATGGCCAAGATTCTGGAGGACCAGGAGGAGGGAGAAGACAACAGTCAGGTCTACAACGACAAGGATCTCCTCG

GCTAAGCAGACGCACCCTCTGTCGAAATCAGTGACAAGACATCTCCTACCAGTATTATAAAAAACTTAGGACCCAGGTCC

AAGCAAGCACACATCGACACTCCAACCATTCGAGCGGAGACCACCGATGGCAGAAGAACAAGCATACCATGTCAACAAGG

GGCTGGAATGTATCAAGTCTCTCAAAGCCTCACCCCCGGATCTATCCACCATCAGAGATACCATCGAGAGCTGGAGAGAG

GGGCTTAGCCCCTCGGGCCGTGCAACACCGAACCCTGATACGTCCGAGGGAGATCATCAGAATATCAACCAATCATGCTC

ACCAGCAATCGGACCAAACAAAGTCTACTTGTCTCCTGAAGATAATCTCGGATTTAGAGAGATCACTGGCAACGACTGTG

AGGCTGGGCTCGGAGGGGTCCAGAGAGAAGGATCCAACTCTCAAGTACAGCGTTACCATGTTTATAGTCACGGGGGTGAA

GAGATTGAAGGACTCGAGGATGCTGACTCTCTCGTGGTTCAAGCAGATCCTCCAGTTGCTAACGTCTTCAATGGAGGAGA

GGATGGATCTGACGACAGCGATGTGGACTCTGGCCCAGATGATCCCGGCAGAGATACTCTATATGACCGGGGATCTGTTG

CCGGCAATGGCATCGCTAGGTCCACAGATGTCGAAAAACTAGAAGGTGCTGATATTCAAGAAGTTCTTAACTCCCAGAAA

GGCAAAGGAGGAAGATTCAAGGGCGGGAAAACCTTGCGAGTCCCGGAAATACCCGATGTCAAGCACTCCAGACCATCAGC

CCAATCAATTAAAAAGGGCACAGACGGGAACTCAGTCTCATCTGGAACGGTGATAGAGTGTTTATCGATAAGTGGTGCAA

CCCAAACTGTGCCAGAGTCAAGATGGGAGTCATCAGAGCAAAATGCGTCTGTGGGGAGTGTCCTCAAGTCTGCGAGGAGT

GCAAAGACGATCCAGGGGTCGACACAAGAATCTGGTACCATAGCATCACTGACTCAGCCTAAAGAGAATGACTCCGAGTA

TGAGTATGAGGATGACCTATTTACAGAGATTCAGGACATCCGTGCAAGCATTGCCAAGATCCATGATGACAATAAAACTA

TCCTCTCAAAGCTTGATTCTATACTGTTATTGAAAGGGGAAGTCGACACTATCAAGAAACAAATCAGCAAGCAGAATATA

AGTATATCCACCATTGAGGGCCATCTCTCCAGTATAATGATAGCCATCCCTGGCTTTGGGAAGGACATCAAGGACCCAAC

ATCCGAAGTCGAGTTGAACCCAGATTTAAGACCTATAATAAGCCGTGACTCTGGCAGAGCTCTCGCGGAGGTCCTCAAGA

AACCCGCTGTGGATAGGTCTCAGAAAATTGGAACCAAAGCCAACTCCAGCTCAAAGGGTCAGCTTCTTAAGGATCTCCAG

CTAAAACCTGTCGACAAGCAGGCAAGCTCTGCAATCGGGTTTGTCCCGTCCGACCATGAATCATCCAGAAATGTCATCCG

CTCCATAATCAAGTCGAGCAAGCTAAATATTGATCACAAGGACTATCTTCTAGATTTACTAAATGATGTGAAAGGCTCCA

AGGATCTTAAGGAATTTCACAAGATGCTAACAGCAATTCTCGCCAAGCACCCGTAACACATCCTCCAGTCATCATCTCAT

ACTCGACTAAAAACATCCTTTCAATCAGGCTATTACAAAAAACTTAGGAGCAAGGGCAACTGAGCTTCGCAGACAGGATG

ACCGAGATCTACGACTTCGATAAATCAGCATGGGATGTCAAAGGGTCAATTGCCCGCATAGAACCCACCACCTACCACGA

CGGCCGACTGATACCCCAGGTGAGGGTCATCGATCCTGGTCTGGGAGACAGAAAAGATGAGTGCTTTATGTACCTGTTTC

TCCTAGGAGTGATTGAGGATAACGACCCCCTGTCTCCCCCCGTCGGAAGAACCTTTGGCTCTCTACCTCTAGGGGTTGGT

AGGTCAACTGCCAAGCCAGAAGAACTACTAAGGGAGGCCACAGAATTAGATATAGTGGTGAGGCGCACTGCAGGAGTAAA

TGAGAAACTGGTATTTTACAACAACACTCCGCTGTCCTTGTTAACGCCCTGGAAGAAAGTTCTGACAACCGGAAGTGTGT

TTAGCGCTAACCAGGTTTGCAATGCAGTCAACCTAGTCCCGCTTGATACTCCCCAGAGATTCAGGGTTGTGTACATGAGC

ATAACTAGATTATCAGACAATGGTTACTATAGTGTGCCCAGAAGAATGCTGGAGTTCCGCTCGGCCAATGCAGTAGCCTT

CAACATCTTGGTTACACTAAGAATTGAAAATGGCACAAACCCTAGAAGATACATAGTCGGCTCATGGGAGAATTCAGAGG

TCACATTTATGGTACACGTGGGCAACTTTAGGAGAAAGAAGAACGAAGTATACTCTGCTGATTACTGCAAAATGAAGATT

GAAAAGATGGGTTTAGTTTTTGCCCTGGGTGGAATAGGCGGAACAAGTCTCCATATTAGAAGCACCGGGAAAATGAGCAA

AACCCTCCATGCACAGCTGGGGTTCAAGAAAATCTTATGTTACCCCCTAATGGATGTCAATGAGGATCTTAACCGATATC

TCTGGCGGGCAGAGTGCCGAATAGTCAAAATCCAAGCTGTCTTACAGCCATCAGTACCCCAAGAGTTCCGTGTCTACGAT

GATGTCATCATCAACGATGATCAAGGCTTGTTCAAGATCTTGTAGTTCATTTGCAAAGGAACGAACCCACCCCCAGGAAG

GGGCCCCACCCTCCCCCG-CCAAAAAACACACTCCAAGGCTCCCCGGCACAGCGCCAACACAGCTACTCCCGCGATGGAA

CGAGGTCGGCC-CCCGAACAAATCCGGTCCAGGGGGGGGGGCCCCCCCCCTGACCAACCGGGCGCCGACCGAGCAGGCCG

GCGAGAAGCCCCCAAGAGGCCCCGACGAAGACCGAGCACCCCATCCCCCC-AAAAACGTCCCCC-AACACGCCCGAGGCT

AGCCACCGGACACCCCCAGACCCGCCACCGGCGAGACAGAACCCCAAGGGGGGACCCTCCAAACCAAAGGAATCACCGGG

GTAGGCTCCGCCGAACCAGCACAGACCCGACCCAAGCAAGGCGGACCCCCAACACGAAACTGCCCAGCACACCCTACTCA

TCATCAAACAAAACTTAGGGGCCAAGTCCATACGAGAGGCCCTAGTCCGTCAGGCGGAGGGACGCAGAAAGGAAGGAGAC

ACGACCGTCCCAGAAGGAGGACACCCAACCACCGAAACGCCCGCCCCCAGGGAAGGAG--CCA-CCCCCACGCAGCGAAG

GAGCGGGTGGGAGACCTCGGGGAGGGCCCGACCGGGACCCGGGGGCCGGCGGCGCCGGCGGTGAGCAGGCCGGGCGCATA

CCTGACCCCACCGGAGGATCCGAGGGGGGCCCGACCCCGCGTGGACCAAACGGCAGGGAGTCCAACCCCACCAAAAAGAG

AGGCGAGGCCCCCCCTCCTTCCCCCCCCGCACACAACCCGCAGACAGACCGAGGGAGGGCCATCCCCCAGCGAGACAGGC

GGCTCCCTCGCAGCCGAGCCGCAGCTCCCCAGAATAGGACACCCCCGCACACTCCGAACACCCCCTCGCCCCCCGCCACC

GGGACAGGGGCTCCACCACCACAGACCGGAGCCACCGACAGCCAGCCAGAGGTCCCCTCCAGTCCCGCCCGGCGCCCCCC

CGAAGAGGCCCCGGCAAGGCCCCCCAGCCCGACGTCTCCCGCACGCACCGAAAGAAGGACCCGCCCGAGAACGTGCGGCA

CATCCATGCGCAAACATCATGACGCGGGTCGCAATTTTGACATTTCTGTTTCTTTTCCCAAATGTTGTTGCGTGTCAGAT

TCACTGGGGCAATCTATCCAAGATCGGGATTGTAGGAACAGGGAGTGCCAGCTACAAGGTGATGACTAGGCCAAGCCACC

AGACTCTGGTTATAAAGTTAATGCCAAATATAACGGCCATCGACAATTGTACAAAGTCAGAGATTGCAGAGTACAAGAGA

TTGCTGATCACAGTGTTAAAGCCTGTAGAGGATGCTCTGTCGGTGATAACCAAGAATGTAAGACCAATTCAAACTCTAAC

ACCTGGGCGTAGAACCCGCCGTTTTGCTGGAGCTGTTCTGGCCGGGGTAGCACTTGGAGTTGCGACAGCCGCTCAGATAA

CTGCAGGAGTCGCCCTTCATCAATCATTGATGAACTCCCAAGCAATTGAGAGTTTAAAAACCAGTCTTGAGATGTCGAAT

CAGGCAATAGAAGAAATCAGACTTGCAAATAAGGAGACCATACTGGCAGTACAGGGCGTCCAGGATTATATCAACAATGA

GCTCGTCCCTTCTGTTCATAGAATGTCATGCGAGCTGGTAGGTCACAAGCTCGGCCTCAAGCTCCTTAGGTACTACACCG

AGATCCTGTCCATATTCGGGCCCAGTCTTCGAGACCCGATATCTGCCGAAATATCAATCCAGGCACTTAGTTATGCATTA

GGCGGAGACATTAATAAAATCCTGGACAAGCTTGGGTATAGCGGTGGGGATTTCCTTGCCATCCTAGAAAGCAAGGGAAT

AAAGGCCCGGGTTACATATGTGGACACAAGAGATTACTTTATAATCCTTAGCATCGCCTACCCAACCTTATCTGAGATCA

AGGGAGTGATAGTTCACAAGATAGAAGCTATAACATACAACATTGGGGCACAGGAGTGGTATACTACTATCCCTAAATAT

GTAGCCACTCAGGGGTATCTGATATCGAACTTTGATGAGACGTCATGCGTATTCACTCCAGAGGGGACAGTTTGCAGCCA

GAATGCGTTGTACCCAATGAGCCCATTGCTTCAGGAATGTTTCAGGGGGTCAACAAAATCGTGCGCCAGAACCTTAGTTT

CAGGGACCATAAGTAATAGATTTATCCTATCAAAAGGGAACCTGATTGCAAATTGTGCGTCAGTTTTGTGCAAATGTTAC

ACAACAGAGACAGTTATCAGCCAAGATCCTGACAAACTATTAACTGTTGTAGCATCCGACAAGTGTCCTGTAGTTGAGGT

GGATGGAGTGACAATACAGGTCGGCAGTCGAGAGTATCCGGATTCTGTATACTTACACAAAATAGATTTAGGTCCAGCAA

TCTCCCTAGAAAAACTGGATGTAGGCACCAATTTAGGCAATGCAGTCACAAGACTGGAGAATGCAAAGGAGCTCCTAGAT

GCATCAGACCAAATACTGAAGACTGTTAAAGGGGTACCTTTCAGTGGGAATATGTACATAGCACTGGCAGCTTGCATAGG

AGTATCCCTGGGCCTTGTCACATTAATATGCTGCTGTAAGGGGAGGTGTAAGAACAAGGAAATCCCTATCTCCAAAATCA

ACCCAGGGCTCAAACCCGACCTGACCGGGACCTCAAAGTCGTACGTAAGATCACTGTAGTCAGAATCACCTGGATCATCT

GGCATCACACACATACATGCACGACACAGGCAGTCCGAGGACGCAAGAAACCCAGCCTCCGGTCACCCACCCGACCCCAC

TCCACGCTCCACCACACATTAGTCATCAAACAAAACTTAGGACGAAAGGTCAATCACCATGTCCGCACAAAGGGAGAGAA

TCAATGCCTTCTACAAAGACAATCCTCACAATAAGAACCATAGGGTGATCCTGGATAGAGAACGCTTGGTCATTGAAAGA

CCCTACATCTTGCTTGGAGTCCTGCTGGTAATGTTCCTGAGTCTAATCGGACTGCTGGCCATTGCAGGGATCAGGCTTCA

CCGGGCCACCGTTGGAACTTCAGAGATCCAGAGTCGGCTGAATACCAATATTAAGTTGACCGAATCTATTGATCACCAGA

CTAAGGATGTCTTAACTCCCCTTTTTAAAATCATTGGCGATGAAGTCGGCATCAGAATTCCACAGAAATTCAGTGATCTT

GTCAAGTTCATCTCCGATAAGATTAAATTCCTCAACCCTGATAGAGAGTATGATTTCAGAGATCTCCGGTGGTGTATGAA

TCCCCCCGAGAGAGTCAAAATTAATTTTGATCAGTTTTGTGAGTACAAGGCTGCGGTTAAGTCAATTGAACATATATTTG

AGTCACCACTCAACAAGTCAAAAAAGCTGCAATCTTTGACTCTCGGGCCCGGAACAGGCTGTCTAGGCAGGACAGTAACA

AGAGCCCATTTCTCAGAACTTACAATGACCTTAATGGACCTGGATCTAGAGATGAAGCACAACGTGTCCTCAGTGTTTAC

CGTAGTTGAAGAGGGATTATTCGGAAGAACATATACCGTCTGGAGATCCGATGCCAGGGATCCGAGCACCGATCTAGGTA

TCGGCCATTTTTTAAGAGTCTTCGAGATTGGACTGGTAAGTGATCTTGGGCTGGGTCCCCCTGTTTTTCATATGACCAAC

TATCTCACGGTGAACATGAGTGATGACTATCGGAGATGTCTTTTAGCGGTAGGGGAGTTGAAGTTGACAGCCCTATGCAC

CTCATCTGAGACTGTGACACTGAGTGAGAGAGGAGTTCCAAGGAGGGAACCTCTTGTGGTTGTGATACTTAATCTAGCTG

GACCCACTCTAGGGGGAGAGCTATACAGTGTCTTGCCTACCTCTGATCTCATGGTGGAGAAACTCTATTTGTCTTCACAT

AGAGGGATCATCAAAGATGATGAGGCCAATTGGGTAGTGCCGTCTACCGATGTTCGTGATCTTCAGAACAAAGGTGAATG

TCTGGTGGAAGCATGCAAGACTCGACCTCCTTCATTTTGCAATGGCACAGGATCAGGCCCGTGGTCAGAAGGGAGAATCC

CTGCCTACGGGGTGATCAGGGTCAGTCTTGACTTAGCGAGTGACCCGGATGTAGTTATCACTTCAGTGTTTGGCCCACTG

ATACCTCACCTATCCGGCATGGATCTTTACAACAACCCGTTTTCAAAAGCTGTATGGTTGGCTGTACCACCTTATGAGCA

GTCATTTCTAGGAATGATAAATACAATTGGATTCCCTAACAGAGCAGAGGTTATGCCGCACATTTTGACCACAGAGATCA

GAGGCCCTCGGGGTCGTTGCCATGTTCCCATAGAATTGTCCCGCCGGGTGGATGACGATATCAAGATCGGGTCCAACATG

GTCATATTGCCGACGATGGACCTGAGGTATATTACAGCCACTTATGATGTTTCCAGGAGCGAGCATGCAATCGTGTACTA

TATCTATGACACGAGTCGCTCATCATCTTACTTCTACCCAGTTCGACTGAATTTCAAAGGCAATCCTCTCTCTTTGAGGA

TAGAGTGTTTCCCCTGGCGTCATAAGGTGTGGTGCTACCATGATTGTCTTATATACAACACCATAACAGGTGAAGAGGTC

CATACGAGAGGGCTGACCGGCATAGAGGTAACATGCAATCCAGTCTGAGCAGAGCTACGACCGTCACTCAGGCAGTCCCT

TGAGTCGCCACCGAGTCCAAGCAGCACAGCCTGGGACAATCAACGGCACAGCCCAGCCAACAATGTTATAAAAAACTTAG

GAGCCAGGGTTGTAGGGGCCATGGACTCTCTATCAGTCAATCAGGTCTTGTACCCTGAGGTTCATCTAGACAGCCCTATT

GTCACAAACAAACTAGTTGCCATCCTTGAGTACTCGGGTATCGACCATAACTATGTTCTTGAAGACCAGACTCTTGTCAA

GAATATTAGGTATAGGCTGGGGTGCGGTTTTTCAAATCAAATGATCATCAATAACAGGGGGGTGGGTGAAACAGTCAATT

CTAAACTTAAAAGTTACCCCCATAATCGTCATATCATATACCCGGATTGCAATAAGGAGTTGTTTTGTATCAAGGATAGC

TGCATATCTAAGAAGCTCTCGGAGCTATTCAAGAAGGGTAATTCCTTGTACTCTAAGATAAGTCACCAGGTACTGGATTG

TCTTAAGAGAGTCAATGGGAAATTAGGTCTGGGCACAGATCTTACCCATGGTCTGAAGGAGGGTATCCTTGACCTAGGGT

TGCACATGCATAGCTCTCAATGGTTCGAGACTTTCCTGTTCTGGTTCACTATCAAGACAGAGATGAGATCAATGATCAAA

GAACAGTCCCATATATGCCACAAGAGGAGGTATAACCCGACTTTTGTGTCGGGGGATGCATTCGAGGTGCTCGTATCGCG

AGACCTCGTTGTGATAATTGATAAGAATACCCAGTATGTCTTCTACTTAACATTTGAGCTAGTCCTTATGTATTGTGATG

TCATAGAGGGTAGACTTATGACGGAGACAGCCATGGCCATAGACCAGAGATATTCAGAACTTCTAAGCCGGGTCAGATAC

TTGTGGGATCTTATTGATGGGTTCTTCCCAACACTAGGCAACACCACATACCAAATTGTTGCTCTGCTTGAACCATTATC

GTTGGCTTATCTTCAACTTCAGGATGTCACTCTGGAGTTAAGAGGTGCTTTTTTAGACCATTGCTTCAAAGAACTCTATG

AGATACTGGAACATTGCGGCATTGACACAGCAGGCACTTACAATTCCATCACTGAAGGATTGGATTACGTATTCATCACC

CACGATATACATCTAACTGGGGAGATTTTTTCATTTTTCCGGAGTTTCGGACACCCCCGTCTAGAAGCAGTCACCGCTGC

GGAAAATGTCAGAAAACATATGAACCAACCGAAGGTAATCAGTTATGAGACTATGATGAAAGGCCATGCGGTATTTTGCG

GGATAATCATAAATGGTTTTCGGGACCGACATGGCGGCAGCTGGCCCCCTGTTGCACTGCCAGAACATGCCTCTGCTGCG

ATCCGGAATGCGCAGGCATCAGGTGAGGGACTGACCCATGACCTGTGTATAGACAACTGGAAATCCTTTGTTGGATTCAA

ATTTGGCTGCTTTATGCCACTCAGCCTAGATAGTGATTTGACCATGTATCTCAAAGACAAGGCATTGGCTGCACTGAAGA

ATGAGTGGGATTCAGTTTACCCGAAAGAATACCTCCGTTATAACCCACCTAGAGGGACAGAGTCAAGGCGATTGGTAGAG

GTATTCCTGAATGACTCCAGCTTTGATCCTTATAACATGATAATGTACGTGGTGAATGGCTCCTACCTTAAAGACCCTGA

GTTTAACCTCTCATACAGCCTAAAAGAGAAGGAGATAAAGGAGACAGGGCGGTTGTTTGCCAAAATGACCTATAAGATGC

GGGCCTGTCAGGTAATCGCTGAAAATCTGATATCGAATGGTGTTGGGAAGTATTTCCGAGACAATGGGATGGCAAAAGAC

GAGCATGACCTAACAAAAGCCCTTCACACCCTGGCAGTCTCAGGTGTTCCCAAAAACAACAAAGATAACCACCGAGGTGG

GCCTCCCAGAAGGACAACAAACCGAGGGGTGAGATCAAGCCAAGGCACCAAAACACAAGATAGAGACAAGGTTCAAGGGG

GACCTATGTACAACTATTTGCGATGCCAACCGATCAGCCCTGATCAGGGTGAGTCATACGAGACTGTTAGTGCATTCATC

ACCGCTGACCTTAAGAAGTATTGCCTGAATTGGAGATACGAGACAATCAGCATATTTGCACAGAGGCTGAATGAAATATA

TGGACTACCATCCTTCTTTCAATGGTTACACAGGATATTGGAAAAATCCGTACTCTACGTCAGTGACCCACATTGCCCTC

CCGACCTAGATAATAAGGGCACTCTGCAGAGTGTCCCCAATGCCCAAATATTCATCAAGTACCCAATGGGCGGAATAGAA

GGTTATTGCCAAAAACTATGGACAATCAGCACTATACCATACTTGTATCTGGCAGCCTATGAGAGCGGAGTAAGAATCGC

CTCACTGGTGCAGGGTGACAATCAGACAATCGCAGTGACAAAAAGAGTTCCAAGTTCTTGGCCTTATTCACTAAAAAAGA

GGGAGGCCTCCAAGGCCGCTCAAAATTACTTCGTAGTCTTAAGGCAAAGATTGCACGATGTAGGTCATCACTTAAAGGCT

AATGAGACCATAGTATCTTCTCACTTTTTTGTATATTCCAAAGGGATTTATTATGACGGCCTGCTAGTCTCACAATCACT

AAAGAGCATCGCCAGATGTGTCTTCTGGTCCGAGACTATAGTGGATGAAACCAGAGCGGCGTGCAGTAATATTGCAACAA

CTGTCGCTAAGAGTATAGAGAGGGGTTATGATAGGTACCTTGCATATTCTTTGAATATCCTCAAGATTTTCCAACAGATC

CTTATATCCCTTAACTTCACTATTAACACAACAATGACTCAGGATGTCGTGGCACCGATCATCGAGAACGGTGATTTGCT

GATAAGGATGGCACTCTTGCCAGCACCCATCGGGGGTCTCAATTATCTTAACATGAGCAGGTTATTTGTGAGAAATATCG

GTGACCCGGTCACCTCCTCTATAGCCGACCTGAAGAGGATGATAGATGCCGGGCTAATGCCAGAAGAAACATTGCATCAA

GTGATGACCCAGACCCCGGGAGAATCATCCTACCTAGATTGGGCAAGTGACCCTTATTCTGCCAACCTACCCTGCGTACA

GAGTATCACTCGCCTTCTCAAAAACATCACTGCACGATATATTTTAATCAGCAGCCCAAACCCGATGCTGAAAGGGTTAT

TTCATGAGGGGAGTAGAGATGAAGACGAGGAGCTTGCAAGTTTCCTAATGGATCGGCATATAATTGTTCCAAGAGCTGCA

CATGAAATCTTAGACCATAGTATAACCGGAGCAAGAGAAGCTATAGCCGGGATGTTGGACACCACCAAGGGTCTGATTAG

AACAAGTATGAAACGGGGCGGCCTCACCCCTCGAGTCTTAGCCCGCCTTTCCAATTATGATTATGAACAATTCCGATCCG

GGATAACACTATTAACAAAGAAAGGGCAGTGTTATCTCATTGACAAGGACTCGTGCTCGGTGCAGCTCGCTATAGCTCTG

AGGGGCCACATGTGGGCTAAGTTGGCTCGCGGGAGACCAATCTATGGGTTGGAGGTGCCTGATGTATTAGAATCGATGAA

CGGCTACCTTATCAAACGCCACGAGTCGTGTGCTATCTGTGAAACGGGCTCAAGTCACTACGGATGGTTTTTCGTCCCTG

CAGGGTGCCAGCTTGACGATGTATCTAGAGAGACTTCGGCTCTTCGTGTACCTTATGTCGGATCAACAACTGAGGAAAGG

ACAGACATGAAACTTGCATTTGTTAGGTCTCCAAGTAGATCCCTCAAATCAGCAGTTAGAATTGCAACAGTTTACTCATG

GGCTTACGGGGATGATGAGAAATCATGGAGTGAAGCCTGGATGCTAGCTAGGCAGAGAGCCAATATCACCTTAGATGAAT

TGAGAATGATCACTCCAGTCTCCACATCCACCAACTTAGCCCACCGGTTGAGGGATCGGAGTACCCAGGTGAAATACTCG

GGGACATCCCTTGTGAGGGTTACAAGATACACAACCATCTCCAATGATAACTTGTCGTTTGTGATATCTGAGAAAAAAGT

GGATACCAACTTCATTTATCAGCAAGGGATGTTGCTTGGTCTTGGGATCCTTGAGAACCTCTTTAGGCTAGAGGCCACCA

CCGGAGTATCCAACACAGTGCTACACCTGCACGTGGAAACAGAATGTTGTGTTGTACCTATGGTGGATCACCCAAGGATA

CCGAGTCTCCGTAATATAAAAGTTACGGATGAGCTATGCACAAACCCTTTGATCTACGACAGGTCCCCCATCATAGAGCA

CGATGCAACCCGGCTATATTCACAGAGCCACAGGAGACATTTGGTGGAGTTTGTTACCTGGTCAACAAGCCAGCTTTACC

ATATACTGGCTAAGTCTACAGCAATGTCCATGATTGAATTGATCACGAGATTCGAGAAAGATCACATGAATGAAATAGCC

GCCCTGATTGGCGATGACGACATCAACAGCTTCATCACAGAATTTTTGCTAGTGGAGCCTAGATTATTTATAGTTTACCT

TGGTCAGTGTGCTGCCATCAATTGGGCCTTTGATATACATTATCATCGGCCCTCGGGCAAGTACCAGATGGGGGAACTTC

TCTACTCTCTGCTCTCTCGGATGAGCAAAGGAGTATATAAGATCTTTACTAATGCTTTGAGCCACCCCAAAGTTTACAAG

AAATTTTGGCGAAGTGGCATCATTGAGCCGATTCATGGCCCATCCTTGGATACACAGAATTTACATGTCACTGTCTGTGA

CATGATATATGGGTCCTATGTCACCTATTTGGATCTTTTGCTGAATGATGAGCTAGATGATTACCCGTATTTGCTCTGCG

AGAGTGATGAGGACGTGGTCACAGACAGGTTCGACAACATTCAAGCCAAACATCTCTGTGTATTGGCCGATGTATATTGC

AGCTCCAAGAGGTGTCCTTCAATAATCGGGATGTCTCCTATAGAAAAATGTACCATTCTCACACATTACATCAAGGGAGA

ATCAGTACAATCCCCGTCTGGGACCTCATGGAACACTGATCCCCTTGTAGTAGATCATTACTCATGCTCTCTGACCTACC

TTCGCCGCGGTTCCATCAAACAAATCAGGTTGAGGGTGGATCCTGGGTTTGTATTCGAGGCGTTGACAGACGTCGACTTT

AAACAACCTCGCAAAGCTAAGTTAGATATATCGGTCGTTGGGTTGACTGATTTCTCTCCCCCTTACGACAACGTCGGTGA

TTTTTTAGGGACTATCAACACATTGAGGCACGATCTGCCTGTCACCGGAACCGGGGTCTCGAACTATGAAGTCCACGCTT

ATCGTAGAATTGGCCTGAATTCATCAGCATGTTATAAAGCCGTAGAGATCTCCACGTTAATCAAGCCATCCTTAGAAGTC

GGAGAGCATGGATTGTTCTTAGGAGAAGGTTCCGGTTCAATGCTGGCTGCGTACAAAGAAGTTCTTAAATTAGCAAATTG

TTACTACAACAGCGGAGTAACAGCGGAGGGCAGAGCCGGACAGAGGGAAATATCTCCCTATCCTTCAGAGATGAGCCTAG

TAGAGAATCAAATGGGGATAGAGCGGAGTGTTAAAGTGCTGTTCAACGGGAAACCTGAAGTAACTTGGGTGGGGACCACA

GATTGCTACAAGTACATAATCAGTAACATCCAAACCTCTAGTCTGGGTTTCATACATTCAGATATTGAGACGCTCCCAAC

CAAAGATGCTGTTGAGAAATTGGAGGAATTTGCCTCTATCCTATCCTTATCCTTGATTTTGGGGAAAATCGGCTCTATTA

CAGTTGTCAAAATTATGCCCATTAGCGGGGATTTTACCCAAGGCTTCATAGCCTATGCCATCCAATATTTTAGGGAGAGC

CTGCTTGCCTATCCGAGATATAGTAACTTCATCTCGACTGAGTGTTACCTTATTATGATAGGATTAAAGGCCAATCGATT

GATAAACCCAGAAGCCATTAAGCAAAGCATAATCAGAGCGGGGGTCAGGACTGCACCAGGACTTGTGAGCCATGTATTAT

CAGAGAAACAAAAAGGTTGTATTCAATCTTTCTTGGGTGATCCTTATATCCAAGGAGACTTCAATAAGCACCTTAAATCT

CTAACCTCTATTGAGAAAATCCTGGTAAATTGTGGTCTCTCGATCAATGGCACAAAAATCTGTAGTGATCTAATCCACCA

TGATATCGCCTCCGGTCCAGACGGTCTTATGAGCTCTACAATTATTTTATATAGGGAATTGGCTCATTTCAAAGATAATA

TAAGGAGTCAGCACGGAATGTTCCACCCCTACCCGGTACTGGCCAACAGCAGGCAACGTGAATTAATCCTTCGAATAGCC

AAGAAATTCTGGGGGTATGTCTTGCTATATTCTGATGACCCGGCACTAATCAGACAAACAATCAAGAACTTGAAACGGAA

TCATCTAACCTTTGACTTACATAGTAACCCATTTATTAAAGGGCTATCCAAGGCTGAGAAACTGCTGGTGCGGACGAGTT

CACTAAGAAGAGAATGGTTGTTCACTCTCGAAACGAAAGAAGTAAAAGAGTGGTTCAAATTGGTAGGTTACAGTGCACTC

ATCAGAGGCTAGTCAACTATGCATCTGCCCCCTTCTCCTCCGCCATGAGACCCCACTGGCGATCCAGAAGATTAAAGAAA

ACTACATATTGGATAAGTATCTATTCCCAGCTTTGTCTGGT

>MN657232.1/sheep/Turkey/2018-09

ACCAAACAAAGTTGGGTAGGGATAGATCTTATAATAACTATGGACTGGCAAACTTAGGAGTAAAGATCCTACTGTCGAGG

AGAGGAGGAGGAGCAAGATCTTTGATTATGGCGACCCTTCTTAAAAGCTTAGCATTGTTCAAAAGGAACAAAGACAAAGC

GCCGACGGCATCAGGTTCAGGAGGAGCCATCCGGGGGATCAAGAATGTTATCATAGTCCCGATTCCCGGAGACTCGTCCA

TCATTACCCGTTCGAGACTGCTCGACAGGCTTGTCAGGTTGGCCGGAGATCCTGACATCAACGGGTCAAAGTTGACCGGC

GTGATGATCAGCATGTTATCTTTGTTCGTGGAGTCACCCGGGCAATTGATACAGCGAATCACAGATGATCCAGATGTTAG

TATCCGCCTTGTTGAGGTAGTTCAAAGTACTAGGTCTCAGTCCGGGTTGACCTTTGCATCACGTGGTGCTGATTTAGACA

ACGAGGCAGACATGTATTTTTCAACTGAGGGGCCCTCGAGTGGAGGTAAGAAAAGGATCAACTGGTTTGAGAACAGGGAA

ATAATAGACATAGAGGTGCAGGATCCAGAAGAGTTCAATATGCTGTTAGCCTCCATACTAGCACAAGTCTGGATCCTTCT

GGCCAAGGCTGTTACGGCACCGGATACAGCAGCTGACTCAGAATTGAGAAGGTGGGTTAAATATACACAACAAAGGAGAG

TGATTGGGGAATTCCGCCTTGACAAAGGATGGTTGGACGCAGTCCGCAACAGAATTGCAGAAGATCTATCACTTCGGCGG

TTCATGGTATCTCTCATACTTGACATCAAGAGGACCCCCGGCAACAAGCCAAGGATTGCAGAAATGATCTGCGACATTGA

CAACTATATTGTCGAAGCCGGACTCGCCAGTTTCATCCTTACTATCAAGTTTGGTATCGAAACCATGTATCCTGCATTGG

GGCTCCACGAGTTTGCCGGGGAATTGTCCACTATTGAGTCCTTGATGAACTTGTATCAACAGCTGGGAGAGGTTGCACCC

TACATGGTAATTCTCGAGAACTCAATTCAGAACAAGTTCAGTGCAGGAGCTTATCCCCTCCTCTGGAGCTATGCGATGGG

TGTCGGAGTTGAGCTGGAGAACTCAATGGGGGGCTTGAACTTCGGCAGGTCATATTTTGACCCTGCCTATTTTCGTCTCG

GACAGGAGATGGTCAGAAGATCTGCAGGAAAGGTCAGCTCTGTGATTGCGGCTGAGCTCGGCATCACAGCAGAGGAAGCC

AAGCTAGTCTCGGAAATCGCCTCGCAGGCTGGGGACGAAAGAACCGCCAGAGGGACTGGGCCTCGACAAGCGCAGGTCTC

CTTCCTCCAGCACAAAACAGGAGAGGGAGAGTCGTCCGCACCAGCAACCAGAGAAGGGGTCAAAGCTACGATCCCAAACG

GGTCCGAAGAAAGGGACAGAAAGCAAACGCGCCCGGGAAGGCCCAGAGGAGAGACCCCCAGCCAACTGCTCCTGGAAATC

ATGCCAGAGGATGAGGTCTCGCGAGAGTCTGGTCAAAACCCTCGTGAGGCTCAAAGATCGGCCGAGGCACTCTTCAGGCT

GCAGGCTATGGCCAAGATTCTGGAGGACCAGGAGGAAGGAGAAGACAACAGTCAGGTCTACAACGACAAGGATCTCCTCG

GCTAAGCAGACGCACCCTCCGTCGAAATCAGCGACAAGACATCTCCCACCAGTATTATAAAAAACTTAGGACCCAGGTCC

AAGCAAGCACACATCGACACTCCAATCAGTCGAGCGGAGACCACCGATGGCAGAAGAACAAGCATACCATGTCAACAAGG

GGCTGGAATGTATCAAGTCCCTCAAAGCCTCACCCCCGGATCTATCCACCATCAGAGATACCATTGAGAGCTGGAGAGAG

GGGCTCAGCCCCCCGGGCCGTGCAACACCGAACCCCGGTACGTCCGAGAGAGATCATCAGAATATCAACCAATCATGCTC

ACCAGCAATCGGACCAAACAAAGTCTACTTGTCTCCTGAAGATAATCTCGGATTTAGAGAGATCACTGGCAATGACTGTG

AGGCTGGGCTCGGAGGAGTCCAGGGAAAAGGATCCAACTCTCAAGTACAGCGTTATCATGTTTATAGCCACGGGGGTGAA

GAGATTGAAGGACTCGAGGATGCTGACTCTCTCGTGGTTCAGGCAGATCCTCCAGTTGCTAACATCTTCAATGGAGGAGA

GGATGGATCTGACGACAGCGATGTGGACTCTGGCCCAGATGATCCCAGCAGAGATACTATATATGACCGGGGATCTTTTG

CCGGCAATGATGTCGCTAGGTCCACAGATGTCGAAAAACTAGAAGGTGCTGATATTCAAGAGGTTCTTAACTCCCAGAAA

GGCAAAGGAGGAAGATTCCAAGGCGGGAAGACCTTACGAATCCCGGAAATACCCGATGTCAAGCACTCCAGACCATCATC

CCAATCAATTAAAAAGGGCACAGACGGGAACTCAGTCTCATCTGGAACGGTGACAGAGTGTTTATCGATAAGTGGTGCAA

CCCAAGCTGTGCCAGAGTCAAGATGGGAGTCATCAGAGCAAAATGCGTCTGTGGGGAGTGTCCTCAAGTCTGCGAGGAGT

GCAGAGACGATCCAGGGGTTGACACAAGAATCTGGTACCATAGCATCACTGACTCAGCCTAAAGAGAATGACTCCGAGTA

TGAGTATGAGGATGACCTATTTAAAGAGATTCAGGACATCCGTGCAAGCATTGCCAAGATCCATGATGACAACAAAACTA

TCCTCTCAAAGCTTGATTCTATACTGTTATTGAAAGGAGAAGTCGACACTATCAAGAAACAAATCAGCAAGCAGAATATA

AGTATATCCACCATTGAGGGCCATCTCTCCAGTATAATGATAGCTATCCCGGGCTTTGGGAAGGATATCAAGGACCCAAC

ATCCGAGGTTGAGTTGAACCCAGATTTAAGACCTATAATAAGCCGTGACTCTGGCAGAGCTCTCGCGGAGGTGCTCAAGA

AACCAGCTGTGGATAGGTCTCTGAAAATTGGAACCAAAGCCAATTCCAGCTCAAAGGGTCAGCTTCTTAAGGATCTCCAG

CTAAAACCTGTCGACAAGCAGGCAAGCTCTGCAATCGGGTTTGTCCCATCCGACCATGAATCATCCAGAAATGTCATCCG

CTCCATAATCAAGTCGAGCAAGCTAAACATTGATCACAAGGACTATCTTCTAGATTTACTAAATGATGTGAAAGGCTCTA

AGGATCTCAAGGAATTCCACAAGATGCTAACAGCAATTCTCGCCAAGCACCTGTAACACACCTTCCAGTCATCATCTCAT

ACTCGGCTGAAAACATCCTCTCAATCAGGCTATTACAAAAAACTTAGGAGCAAGGGTAACTGAGCTTCGCAGACAAGATG

ACCGAGATCTACGACTTCGATAAATCAGCATGGGATGTCAAAGGGTCAATTGCTCGCATAGAACCCACCACCTATCACGA

CGGTCGACTGATACCCCAGGTAAGGGTCATCGATCCTGGTCTGGGAGACAGAAAGGATGAGTGCTTCATGTACCTGTTTC

TCCTAGGAGTGATTGAGGATAACGACCCCCTGTCCCCCCCAGTTGGAAGAACCTTTGGTTCTTTACCTCTAGGGGTCGGT

AGGTCAACTGCTAAGCCGGAAGAACTACTAAGGGAGGCCACAGAACTAGATATAGTGGTGAGGCGCACTGCAGGATTAAG

TGAGAAACTGGTATTTTACAACAACACTCCGCTGTCCCTGTTAACGCCCTGGAAGAAGGTTCTGACAACCGGAAGTGTGT

TTAGTGCTAACCAGGTTTGCAATGCAGTCAACCTAGTCCCGCTTGATACTCCCCAGAGATTCAGGGTTGTGTACATGAGC

ATAACTAGATTATCAGACAATGGTTACTATAGTGTGCCCAGGAGAATGCTGGAGTTCCGCTCAGCCAATGCAGTAGCCTT

CAACATCCTGGTTACACTAAGAATTGAAAATGGCACAAGCCCTAGAAGATACATAGTCGGCTCATGGGAGAATCCAGAGG

TCACATTTATGGTACACGTGGGCAACTTTAGAAGAAAGAAGAACGAAGTATACTCAGCTGATTACTGCAAAATGAAAATT

GAAAAGATGGGTTTAGTTTTTGCTCTGGGAGGAATAGGTGGAACAAGTCTCCATATTAGAAGCACCGGGAAAATGAGCAA

AACCCTCCATGCACAGCTGGGGTTCAAGAAAATCTTATGTTACCCCCTAATGGATGTTAATGAGGATCTCAACCGATATC

TCTGGCGGGCAGAGTGCCGAATAGTCAAAATCCAAGCCGTCTTGCAGCCATCAGTACCCCAAGAATTCCGTGTCTACGAT

GATGTCATCATCAACGATGATCAAGGCCTATTCAAGATCCTGTAGTTCGCTTGCAAAGGAATGAATCCACCCCCAGGAAG

GGGCCCCACCCTCCCCCG-CCAGAAAACACACCCCAAGGCTCCCCGACACAACACCAACACAGCCACTCTCCCGATGGGA

CGAGGCCGGCC-TCCCAACAGACCCGGTCCAGGC-GGGGGGCCCCTCCCC-GACCCGCCGGGCGCCGACCGAGCAGGTCG

GCGAGAAGCCCCCAGGAGCCCCCGATGGAGACCGAGCACCCCATCCCCCCAAAAACGCCCCCCC-AACACGCCCGAGGCC

AGGTGCCGGGCACCCCCAGACCCGCCACCGGCGAGACAGAACCCCAAGGAGGGATCTCCTAAACCAAAGGGACCACCGGG

GAAGGACCCGCCGAACCAGCACAGACCCGACCCAAGCAAGGCGGGCCCCCAACACGAAACCGTCCAGCGCACCCCACCTA

TCATCAAACAAAACTTAGGGGCCAAGTCCATACGACAGGCCCCAGCCCGCCAGGCGAAGGGCCACAGAAAGGAAGGAGAC

ACGACCGCCCCAGAAGGAGGACACCCGACCACCGAAACACCCAGCCCCAGGGAAGGAG---CCCCCCCCACACGGCGAAG

GAGCAGGCGAGAGACCCCGAGGAGAACCCGACCGGAAGCCAGGAGCCGGCGACGCCGGCGGCAAACAGGCCGGGCGCACA

CCCGACTCCGCCAGAGGAGCCGAGGAGAGCCCTATCCCGCGTGGGCCAAACGGCAGGGAATCCAACCCCGCCAAGGAAGG

AGGTGGGGCCCCTTTT-----CCCCCCCACACACAACCTGCGGACAGACCGAGGGAGGACCACCTCCCAACTGGACAGGC

GACTCCCTCGCAGCCGAGCCGCATCTCCCCAGAAGAGGACACCCCCGCACACTCCGAACACCCCCCCGCCCCCCGCCACC

GGGACAGAGGTTCCGCCACCACAGACCAGAGCCACCGAGAGCCAACCAGAGGTCCTCTCCAGTCCCGTCCAGTGCCCCCC

CGAAGAGGCCCCGGCAAGCCCCCCAACCCCGACGACCCCCGCACGCACCGGAAACGAGACCCGCCCGAGAACGTCCGGCA

CACCCATGTGTAAACATCATGACACGGGTCGCAATCTTGACATTTCTCTTTCTTTTCCTAAATGCTGTTGCGTGTCAGAT

TCACTGGGGCAATCTATCCAAGATCGGGATTGTAGGGACAGGGAGTGCCAGCTACAAGGTGATGACTAGGCCAAGCCACC

AAACTCTGGTTATAAAGTTAATGCCAAATATAACAGCCATCGACAATTGTACAAAGTCAGAGATTGCAGAGTACAAGAGA

TTGCTGACCACAGTGTTAAAGCCTGTAGAGGATGCTCTATCAGTGATAACCAAGAATGTAAGACCAATTCAAACTCTGAC

ACCTGGGCGTAGAACTCGCCGTTTTGCTGGAGCTGTTCTAGCCGGAGTAGCACTTGGAGTTGCGACAGCCGCTCAGATAA

CTGCAGGAGTCGCCCTTCATCAATCATTGATGAACTCCCAAGCAATTGAGAGTTTAAAAACCAGTCTTGAGAAGTCGAAT

CAGGCAATAGAAGAAATCAGACTTGCAAATAAGGAGACCATACTAGCAGTACAGGGCGTCCAGGATTATATCAACAATGA

GCTTGTCCCTTCCGTTCATAGAATGTCATGCGAGCTGGTAGGTCACAAGCTCGGCCTCAAGCTCCTTAGGTACTACACTG

AGATCCTGTCCATATTCGGGCCCAGCCTTCGAGACCCGATAGCTGCCGAAATATCAATCCAGGCACTCAGTTATGCATTA

GGCGGAGACATTAATAAAATCCTGGACAAGCTTGGGTATAGCGGTGAGGATTTCCTTGCCATCCTAGAAAGCAAGGGAAT

AAAGGCCCGGGTCACATATGTGGACACANGAGATTACTTTATAATTCTTAGCATCGCCTACCCAACCTTATCTGAGATCA

AGGGAGTGATAGTTCACAAGATTGAAGCTATAACATACAACATTGGGGCACAGGAGTGGTATACTACTATCCCTAAATAT

GTAGCCACTCAGGGGTATCTGATATCGAACTTTGATGAGACGTCATGCGTATTCACTCCGGAGGGGACAGTTTGCAGCCA

GAATGCGTTGTACCCAATGAGCCCATTGCTCCAGGAATGTTTCCGGGGGTCAACAAAATCGTGCGCCAGAACCCTAGTTT

CAGGGACCATAAGTAATAGATTTATCCTATCAAAGGGGAACCTGATTGCAAATTGTGCGTCAGTCTTGTGCAAATGTTAC

ACAACAGAGACAGTTATTAGCCAAGATCCTGACAAACTACTAACTGTTGTAGCATCCGACAAGTGTCCTGTAGTTGAGGT

GGATGGAGTGACAATACAGGTCGGCAGTCGAGAGTATCCGGACTCTGTATACTTACACAAAATAGACTTAGGTCCAGCCA

TCTCCCTAGAAAAACTGGATGTAGGCACCAATTTAGGCAATGCAGTCACAAGACTGGAGAATGCAAGGGAGCTCCTAGAT

GCATCAGACCAAATACTGAAGACTGTTAAAGGGGTACCTTTCGGTGGGAATATGTACATAGCACTGGCGGCTTGCATTGG

GGTATCCCTAGGGCTTGTCACATTAATATGCTGTTGCAAGGGGAGGTGTAAGAACAAGGAAGTCTCCATCTCCAAAATCA

ATCCAGGGCTCAAACCCGACCTGACCGGGACCTCAAAGTCGTACGTGAGATCACTGTAGCCAGAATCACCCGAATCATCC

GGAATCACACACATACATACACGACACAAGCAGTCAGAGAGCGCAGGAAACCCAGCCCCCGATCACCCACCTGACCCCAC

CCCATGCTCCACCACACATTAGTCATCAAACAAAACTTAGGACGAAAGGTCAATCACCATGTCCGCACAAAGGGAAAGGA

TCAATGCCTTCTACAAAGATAATCCTCACAATAAGAACCATAGGGTGATCCTGGATAGAGAACGCTTGGTCATTGAAAGA

CCCTACATCTTGCTTGGAGTCCTGCTGGTAATGTTCCTGAGTCTAATCGTACTGCTGGCCATTGCAGGGATCAGGCTTCA

CCGGGCCACCGTTGGAACCTCAGAGATCCAGAGTCGATTGAATACCAATATCGAGTTGACCGAATCTATTGATCACCAAA

CTAAGGATGTCTTAACTCCCCTTTTCAAAATCATTGGCGATGAAGTCGGAATCAGAATTCCACAGAAATTCAGTGATCTT

GTCAAGTTCATCTCCGATAAGATTAAATTCCTCAACCCTGATAGAGAGTATGATTTCAGGGATCTCCGGTGGTGTATGAA

TCCCCCCGAGAGAGTCAAAATTAATTTTGATCAGTTTTGTGAGTACAAGGCTGCGGTTAAGTCAATTGAACATATATTTG

AGTCACCACTCAACAAGTCAAAAAAGCTGCAATCATTGACTCTTGGGCCCGGAACAGGCTGTCTAGGCAGGACAGTAACA

AGAGCTCATTTCTCGGAACTTACACTGACCTTAATGGACCTGGATCTAGAGATGAAGCACAACGTATCCTCAGTGTTTAC

CGTAGTTGAAGAGGGATTATTCGGGAGAACATATACCGTCTGGAGATCCGATGCCAGAGATCCGAGCACCGATCTAGGTA

TTGGTCACCTTTTAAGAGTCTTCGAGATTGGACTGGTAAGAGATCTCGGGCTGGGTCCCCCTGTTTTTCATATGACCAAC

TATCTCACAGTGAACATGAGTGATGACTATCGGAGATGTCTTTTAGCGGTGGGGGAGTTAAAGTTGACAGCCCTATGCAC

CTCATCTGAGACTGTGACACTGAGTGAGAGAGGAGTCCCAAAGAGGGAGCCTCTTGTAGTTGTGATACTCAACCTAGCTG

GGCCCACTCTAGGGGGCGAACTATACAGTGTCTTGCCTACCTCTGATCCCATGGTGGAGAAACTCTATTTATCTTCACAT

AGAGGGATTATCAAAGATGACGAAGCCAATTGGGTAGTACCGTCTACCGATGTTCGTGATCTTCAAAACAAAGGAGAATG

TCTGATAGAGGCATGCAAGACTCGACCTCCTTCATTTTGCAATGGCACAGGATCAGGCCCGTGGTCAGAGGGGAGAATCC

CTGCTTACGGGGTGATCAGGGTCAGTCTTGACCCAGCCAGTGATCCGGATGTAGTTATCACTTCAGTGTTTGGCCCACTA

ATACCTCACCTATCCGGCATGGATCTTTACAACAACCCGTTTTCAGGAGCTGTATGGTTGGCTGTACCACCTTATGAGCA

GTCATTTTTGGGAATGATTAATACGATTGGATTCCCTAACAGAGCAGAGGTTATGCCGCACATTTTGACCACAGAGATCA

GAGGCCCTCGGGGCCGTTGCCATGTTCCCATAGAATTGTCCCGCAGGGTTGATGACGACATCAAGATTGGGTCCAACATG

GTCATATTGCCGACGATGGACCTGAGATATATTACAGCCACTTATGATGTTTCCAGGAGCGAGCATGCAATCGTGTACTA

TATCTATGACACAGGTCGCTCATCATCTTACTTCTACCCAGTTCGGCTGAACTTCAAAGGCAATCCTCTCTCTCTGAGGA

TAGAGTGTTTCCCTTGGCGTCATAAGGTGTGGTGCTACCATGATTGTCTTATATACAACACCATAACAGATGAAGAAGTC

CATACGAGAGGACTGACCGGTATAGAGGTAACATGTAATCCAGTCTGAGCAGGGCTACGACCGCCACTCAGGCAATCCCT

TGAGTCGCCACCGAGTCCAAGCAGCACAGCCTGGGACACTCAATAGCACAGCCCAGTCAACAATGTTATAAAAAACTTAG

GAGCCAGGGCTGTGGGGGCCATGGACTCCCTATCAGTCAATCAGGTCTTGTACCCTGAGGTCCATTTAGATAGCCCTATT

GTCACAAACAAACTAGTTGCCATCCTTGAGTACTCGGGTATCGACCATAACTATGTTCTTGAAGACCAGACTCTTGTCAA

GAATATTAGGTATAGACTGGGGTGCGGTTTTTCAAATCAAATGATCATCAATAACAGGGGGGTGGGTGAAACAGTCAATT

CTAAACTCAAAAGTTACCCCCATAATCGTCATATCATATACCCGGATTGCAATAAGGAGTTGTTTTGTATCAAGGATAGC

TGCATATCTAAGAAGCTCTCGGAGCTATTCAAGAAAGGTAATTCCTTGTACTCTAAGATAAGTCACCAGGTACTGGATTG

TCTTAAGAGAGTCAATGGGAAATTAGGCCTGGGCACAGATCTTACCCATGGTCTGAAGGAGGGTATCCTTGACCTAGGGT

TACACATGCATAGCTCTCAATGGTTCGAGACTTTCCTGTTTTGGTTCACTATCAAGACAGAGATGAGATCAATGATCAAA

GAACAGTCCCATATATGCCACAAGAGGAGGTATAACCCGGTTTTTGTGTCGGGGGATGCATTCGAGGTGCTCGTATCACG

AGACCTCGTTGTGATAATTGATAAGAATACCCAATATGTCTTCTACTTAACATTTGAGCTGGTCCTTATGTATTGTGATG

TCATAGAGGGCAGACTTATGACGGAGACAGCCATGGCCATAGACCAGAGATATTCAGAACTTCTAAGCCGAGTCAGATAC

TTGTGGGATCTTATTGACGGGTTCTTCCCAACACTAGGCAACACCACATACCAAGTTGTTGCTCTGCTTGAACCATTATC

GTTGGCTTACCTTCAACTTCAGGATGTCACTCTGGAGTTAAGAGGTGCTTTTTTAGACCATTGCTTCAAAGAACTCTATG

AGATACTGGAGCACTGCGGCATTGACACAGAAGGCACCTACAATTCCATCACCGAAGGATTGGATTACGTATTCATCACC

CACGATATACATCTAACTGGGGAGATTTTTTCATTCTTTCGGAGTTTTGGACACCCCCGTCTAGAAGCAGTCACCGCTGC

AGAAAATGTCAGAAAACATATGAACCAACCGAAGGTAATCAGTTATGAGACTATGATGAAAGGGCATGCAGTATTTTGCG

GGATAATCATAAATGGTTTTAGGGACCGACATGGCGGCAGCTGGCCCCCTGTTGTATTGCCGGAACATGCTTCGGCTGCG

ATCCGGAATGCGCAGGCATCAGGTGAGGGACTGACCCATGACCTGTGTATAGACAATTGGAAGTCCTTTGTTGGATTCAA

ATTCGGCTGCTTTATGCCACTCAGCTTAGATAGTGATTTGACCATGTACCTCAAAGACAAGGCATTGGCTGCACTGAAGA

ATGAGTGGGATTCAGTTTACCCGAAAGAATACCTCCGTTATAACCCACCTAGAGGGACAGAGTCAAGACGATTGGTGGAG

GTATTCCTGAATGACTCCAGCTTTGATCCTTATAACATGATAATGTACGTGGTGAATGGCTCCTACCTTAAAGACCCTGA

GTTTAATCTCTCATACAGCCTAAAAGAGAAGGAGATAAAGGAGACAGGGCGGTTGTTTGCCAAAATGACCTATAGGATGA

GGGCCTGTCAGGTAATCGCTGAAAATCTGATATCGAATGGAGTTGGGAAGTATTTCAGAGACAATGGGATGGCAAAGGAC

GAGCATGACCTAACAAAAGCCCTTCACACTCTGGCAGTCTCAGGTGTTCCCAAGAACAACAAAGATAACCACCGAGGTGG

GCCTCCCAGAAGAACAACAAGCCGAGAGATGAGATCAAGCCAAGACACCAAAACACAAAATAGAGATAAGGTCCACGGGG

GGCCTATGTACAACTATTTGCGATGCCAACCAATCAACCCTGATCAAGGTGAGTCATACGAGACTGTTAGTGCATTCATC

ACCGCTGACCTTAAGAAGTATTGCCTTAATTGGAGATACGAGACAATCAGCATATTTGCACAGAGGCTGAATGAAATATA

TGGACTACCATCCTTCTTTCAATGGTTACACAGGATATTGGAAAAATCTGTACTCTACGTCAGTGACCCACATTGCCCTC

CCGACCTAGATAATCATATCCCTCTGGACAGTGTCCCTAATGCCCAGATATTCATTAAGTACCCAATGGGCGGAATAGAA

GGTTATTGCCAAAAACTATGGACAATCAGTACTATACCATACTTGTATCTGGCAGCCTATGAGAGCGGAGTGAGAATCGC

CTCACTGGTGCAGGGTGACAATCAGACAATCGCGGTGACAAAAAGAGTTCCAAGTTCTTGGCCTTATTCACTAAAAAAGA

GGGAGGCATCCAAAGCTGCTCAAAATTACTTCGTAGTCTTAAGGCAAAGATTGCACGATGTAGGTCACCACTTAAAGGCT

AATGAAACCATAGTATCTTCTCACTTTTTTGTATATTCTAAGGGGATTTATTATGACGGCCTGCTAGTCTCACAATCACT

AAAGAGCATCGCCAGATGTGTCTTCTGGTCCGAGACTATAGTGGATGAAACCAGAGCGGCTTGCAGCAATATTGCAACAA

CTGTCGCCAAGAGTATAGAGAGGGGTTATGATAGGTACCTTGCATATTCTTTGAATATCCTCAAGATTTTCCAACAGATC

CTTATATCCCTTAATTTCACTATTAACACAACAATGACTCAGGATGTCGTGGCACCGATCATCGAAAACGGGGATTTGCT

GATAAGGATGGCACTCTTGCCAGCACCCATCGGGGGTCTCAATTATCTTAACATGAGCAGGTTATTTGTGAGAAATATCG

GTGACCCGGTCACCTCCTCTATAGCTGACCTGAAGAGGATGATAAATGCTGGGCTAATGCCAGAAGAAACATTGCATCAA

GTGATGACTCAGACCCCGGGAGAATCATCCTACCTCGATTGGGCAAGTGACCCTTATTCCGCCAACCTACCCTGCGTACA

GAGTATAACTCGCCTTCTCAAAAACATCACTGCACGATATATTTTAATCAGCAGCCCAAACCCGATGCTGAAAGGATTGT

TTCACGAGGGGAGTAGAGATGAAGACGAGGAGCTTGCAAGTTTCTTAATGGATAGGCATATAATTGTCCCAAGAGCTGCA

CATGAAATCTTAGACCATAGCATAACCGGGGCAAGAGAAGCTATAGCCGGGATGTTGGACACCACCAAGGGTCTGATTAG

AACAAGTATGAAACGGGGTGGCCTCACCCCTCGAGTCTTAGCTCGCCTTTCCAATTATGATTATGAACAATTCCGATCCG

GGATAACACTATTGACAAAGAAAGGGCAGTGTTATCTCATTGACAAGGACTCATGCTCGGTGCAGCTCGCTATAGCTCTG

AGGAGCCACATGTGGGCTAGGTTGGCTCGAGGGAGACCAATCTATGGGTTGGAGGTGCCCGATATATTAGAATCTATGAA

CGGCTACCTTATCAAACGCCACGAGTCTTGTGCTATCTGTGAGACGGGCTCAAGTCACTACGGATGGTTTTTTGTCCCTG

CAGNGTGCCAGCTTGACGATGTATCTAGAGAGACTTCGGCTCTTCGTGTACCTTATGTCGGATCAACAACTGAGGAGAGG

ACAGACATGAAACTTGCTTTTGTTAGGTCTCCAAGTCGATCCCTTAAATCAGCAGTCAGAATTGCAACAGTTTACTCATG

GGCCTACGGGGATGATGAGAAATCGTGGGGTGAAGCCTGGATGCTAGCTAGGCAGAGAGCCAACATCACCTTAGATGAAT

TGAGAATGATCACTCCAGTCTCTACATCCACCAACCTGGCCCACCGGTTGAGGGATCGGAGCACCCAGGTGAAATACTCG

GGAACATCCCTTGTGAGGGTTGCAAGATACACAACCATCTCCAATGATAACCTGTCATTTGTGATATCTGAGAAAAAAGT

GGATACCAACTTCATTTATCAGCAAGGGATGCTGCTCGGTCTTGGGATCCTTGAGAACCTCTTTAGGCTAGAGGCCACCA

CCGGAGTATCCAACACAGTGCTACACCTGCATGTGGAAACAGAATGTTGTGTTGTACCTATGGTGGATCACCCAAGGATA

CCGAGTCTCCGTAATATAAAAGTCACGGATGAGCTATGCACAAACCCTTTGATCTACGACAGGTCTCCCATCATAGAGCA

CGATGCAACCCGACTGTACTCACAGAGCCACAGGAGACATCTAGTGGAGTTTGTTACCTGGTCAACAAGCCAGCTTTACC

ATATACTGGCTAAGTCTACAGCAATGTCCATGATTGAGTTGATCACGAGATTCGAGAAAGATCACATGAATGAAATAGCC

GCCCTGATTGGCGATGACGACATCAACAGCTTCATCACAGAATTTTTGCTAGTAGAGCCTAGATTATTTATAGTTTACCT

TGGTCAGTGTGCTGCCATAAATTGGGCCTTTGACATACATTATCATCGGCCCTCAGGCAAGCACCAGATGGGGGAACTCC

TCTACTCTCTGCTCTCTCGGATGAGCAAAGGAGTATATAAGATCTTCACCAATGCTTTGAGCCACCCCAAAGTTTACAAG

AAATTCTGGCGAAGTGGTGTCATTGAGCCGATTCATGGCCCATCCCTGGATACACAGAATTTACATATCACTGTCTGTGA

CATGATATATGGGTCCTATGTCACCTATTTGGATCTTTTACTGAATGATGAGCTAGATGATTACCCGTATTTACTCTGCG

AGAGTGATGAGGACGTGGTCACAGACAGGTTCGACAACATTCAAGCCAAACATCTCTGTGTATTGGCCGATGTATATTGC

AGCTCCAAGAGGTGTCCTTCGATAATCGGGATGTCTCCTATAGAAAAATGTACCATTCTCACACATTACATCAAAGGAGA

ATCAGTACAATCCCCGTCAGGGACCTCATGGAACACTGATCCCCTTGTAGTAGATCATTACTCATGCTCTCTGACCTACC

TTCGCCGCGGTTCCATCAAACAAATCAGGCTGAGAGTGGATCCTGGGTTTGTATTCGAGGCGTTGACAGACGTCGACTTT

AAACAGCCTCGCAAGGCTAAGTTAGATATATCGGTTGTTGGGTTGGCTGATTTCTCTCCCCCTTGCGACAACGTTGGTGA

TTTCCTGGGGACTATCAACACATTAAGGCACAATCTGCCTGTCACCGGAACCGGGGTCTCGAACTATGAAGTCCACGCTT

ATCGTAGAATTGGCCTAAATTCATCAGCATGTTATAAAGCCGTAGAGATCTCCACTTTAATCAAGCCATCCTTAGAAGTC

GGAGAGCACGGATTGTTCCTAGGAGAAGGTTCTGGTTCAATGCTGGCTGCGTACAAGGAAGTTCTTAAACTAGCAAATTG

TTATTACAACAGCGGGGTAACAGCGGAGGGTAGAGCCGGACAGAGGGAAATATCTCCCTATCCTTCAGAGATGAGCCTAG

TAGAGAATCAAATGGGGATAGAGCGGAGTGTTAAAGTGCTGTTCAACGGCAAACCTGAAGTAACTTGGGTGGGGACCACC

GATTGCTACAAGTACATAATCAGTAACATCCAAACCTCTAGTCTAGGTTTCATACATTCAGATATTGAGACACTCCCAAC

CAAAGACGCTGTTGAGAAATTGGAAGAATTTGCCTCTATCCTATCCTTATCCTTGATTTTAGGAAAAATCGGCTCTATCA

CAGTTGTCAAAATTATGCCCATTAGCGGGGATTTTACCCAAGGCTTCATAGCCTATGCTATCCAATATTTTAGGGAGAGC

CTGCTTGCCTATCCGAGATATAGTAACTTCATCTCGACTGAGTGTTACCTTGTTATGATAGGATTAAAGGCCAATCGATT

GATAAACCCAGAAGCTATTAAGCAAAGCATAATCAGAGCAGGGGTCAGGACTGCACCAGGACTTGTGAGCCATATATTAT

CAGAGAAACAGAAAGGTTGTATTCAATCTTTCTTGGGTGATCCTTATATCCAAGGGGACTTTAATAAGCACCTTAAATCT

CTAACCCCTATTGAGAAAATCCTGGTAAATTGTGGTCTCTCGATCAATGGCACAAAAATTTGTAGAGATCTAATCCACCA

TGATATAGCCTCCGGTCCAGACGGTCTTATGAGCTCTACAATTATTTTATATAGGGAGTTGGCTCATTTCAAAGATAATA

TAAGGAGTCAGCACGGAATGTTCCACCCCTACCCAGTACTGGCCAACAGCAGGCAACGTGAATTAATCCTTCGAATAGCC

AAGAAATTCTGGGGGTATGTCCTGCTATATTCTGATGACCCGGCACTAATCAGACAAACAATCAAGAACTTGAAACGGAA

TCATCTAACCTTTGACTTACATAGTAACCCGTTTATTAAAGGGCTATCCAAAGCTGAGAAACTGCTGGTGCGGACGAGTT

CACTAAGAAGGGAATGGTTGTTCACTCTCGATACGAAAGAAGTGAAAGAATGGTTCAAATTGGTAGGTTACAGTGCACTC

ATTAGAGGCTAGTTGGCTATACATCTGCCCCCTTTCCCTCCGCCATAAGACCCCACTGACGATCCAAAAGATTAAAGAAA

ACTACATATTGGATAAGTATCTATTCCCAGCTTTGTCTGGT

>MK408669.1/WildGoat/Iraq/2011-02

NNNNNNNNNNNNNNNNNNNNGATAGATCTTATAATAACTATGGACTGGCAAACTTAGGAGTAAAGATCCTACTGTCGGGG

GGAGGAGGAGGAGCAAGATCTTTGATTATGGCGACCCTTCTTAAAAGCTTAGCATTGTTCAAAAGGAACAAAGACAAAGC

GCCGACGGCATCAGGTTCAGGAGGGGCCATCCGGGGGATTAAGAATGTTATCATAGTCCCGATTCCCGGAGACTCGTCCA

TCATTACCCGTTCGAGACTGCTCGACAGGCTTGTCAGATTGGCCGGAGATCCTGACATCAACGGGTCAAAGTTGACCGGC

GTGATGATCAGCATGTTATCTTTGTTCGTAGAGTCACCCGGGCAATTGATACAGCGAATCACAGATGATCCAGATGTTAG

TATCCGCCTTGTTGAGGTAGTTCAAAGTACTAGGTCTCAGTCCGGGTTGACCTTTGCATCACGTGGTGCTGATTTAGACA

ACGAGGCAGACATGTATTTTTCAACTGAGGGGCCCTCAAGTGGAGGTAAGAAAAGGATCAACTGGTTTGAGAACAGAGAG

ATAATAGACATAGAGGTGCAGGATCCAGAAGAGTTCAATATGCTGTTAGCCTCCATACTAGCACAAGTCTGGATCCTTCT

GGCCAAGGCTGTTACGGCACCGGACACGGCAGCCGACTCAGAATTGAGAAGGTGGGTTAAATATACACAACAAAGGAGAG

TGATTGGGGAATTTCGCCTTGACAAAGGGTGGCTGGACGCAGTCCGCAACAGAATTGCAGAAGATCTATCACTTCGGCGG

TTCATGGTATCTCTCATACTTGACATCAAGAGGACCCCCGGCAACAAGCCAAGGATTGCAGAAATGATCTGCGACATTGA

CAACTATATTGTCGAAGCCGGACTCGCCAGTTTCATCCTTACTATCAAGTTTGGTATCGAAACTATGTATCCTGCATTAG

GGCTCCACGAGTTTGCCGGGGAATTGTCCACTATTGAATCCTTGATGAACTTGTATCAACAGCTGGGAGAGGTTGCACCC

TACATGGTAATTCTCGAGAACTCCATTCAGAACAAGTTCAGTGCAGGAGCTTATCCCCTCCTCTGGAGCTATGCGATGGG

TGTCGGAGTTGAGCTGGAGAACTCAATGGGGGGCTTGAACTTCGGCAGGTCATATTTTGACCCTGCCTATTTTCGTCTCG

GACAGGAGATGGTCAGAAGATCTGCAGGAAAGGTCAGCTCTGTGATTGCGGCCGAGCTCGGCATCACAGCAGAGGAAGCT

AAACTAGTCTCGGAAATCGCCTCGCAGGCTGGGGACGAAAGAACCGCTAGAGGGACTGGGCCTCGACAGGCGCAGGTCTC

CTTCCTCCAGCACAAAACAGGAGAGGGAGAGTCGTCCGCACCAGCAACCAGAGAAGGGGTCAAAGCTGCGATCCCAAACG

GATCCGAAGAAAGGGACAGAAAGCAAACACGCCCAGGAAGGCCCAGAGGAGAGACCCCCAGCCAACTGCTCCTGGAAATC

ATGCCAGAGGATGAGGTCTCGCGAGAGTCTGGTCAAAACCCTCGTGAGGCTCAAAGATCGGCCGAGGCACTCTTCAGGCT

GCAGGCCATGGCCAAGATTCTGGAGGACCAGGAGGAGGGAGAAGACAACAGTCAGGTCTACAATGACAAGGATCTCCTCG

GCTAAGCAGACGCACCCTCTGTCGAAATCAGTGACAAGACATCTCCTACCAGTATTATAAAAAACTTAGGACCCAGGTCC

AAGCAAGCACACATCGACACTCCAATCAGTCGAGCGGAGACCACCGATGGCAGAAGAACAAGCATACCATGTCAACAAGG

GGCTGGAATGTATCAAGTCCCTCAAAGCCTCTCCCCCGGATCTATCCATCATCAGAGATACCATTGAGAGCTGGAGAGAG

GGGCTCAGCCCCTCGGGCCGTGCAACACCGAACCCCGATACGTCCGAGGGAGATCATCAGAATATCAACCAATCATGCTC

ACCAGCAATCGGACCAAACAAAGTCTACTTGTCTCCTGAAGATAATCTCGGATTTAGAGAGATCACTGGCAACGACTGTG

AGGCTGGGCTCGGAGGAGTCCAGGGAAAAGGATCCAACTCTCAAGTACAGCGTTATCATGTTTATAGCCACGGGGGTGAA

GAGATTGAAGGACTCGAGGATGCTGACTCTCTCGTGGTTCAGGCAGATCCTCCAGTTGCTAACATCTTCAATGGAGGAGA

GGATGGATCTGACGACAGCGATGTGGACTCTGGCCCAGATGATCCCAGCAGAGATACTATATATGACCGGGGATTTGTTG

CCGGCAATGATGTCGCTAGATCCACAGATGTCGAAAAACTAGAAGGTGCTGATATTCAAGAGGTTCTTAACTCCCAGAAA

GGCAAAGGAGGAAGATTCCAAGGCGGGAAGACCTTACGAGTCCCGGAAATACCCGATGTCAAGCACTCCAGACCACCATC

CCAATCAATTAAAAAGGGCACAGACGGGAACTCAGTCTCATCTGGAACGGTGACAGAGTGTTTATCGATAAGTGGTGCAA

CCCAAGCTGTGCCAGAGTCAAGATGGGAGTCATCAGAGCAAAATGCGTCTGTGGGGAGTGTCCTCAAGTCTGCGAGGAGT

GCAGAGACGATCCAGGGGTTGACACAAGAATCTGGTACCATAGCATCACTGACTCAGCCTAAAGAGAACGACTCCGAGTA

TGAGTATGAGGATGATCTATTTAAAGAGATTCAGGACATCCGTGCAAGCATTGCCAAGATCCATGATGACAACAAAACTA

TCCTCTCAAAGCTTGATTCTATACTGTTATTGAAAGGAGAAGTCGACACTATCAAGAAACAAATCAGCAAGCAGAATATA

AGTATATCCACCATTGAGGGCCATCTCTCTAGTATAATGATAGCTATCCCGGGCTTTGGAAAGGATATCAAGGACCCAAC

ATCCGAGGTTGAGTTGAACCCAGATTTAAGACCTATAATAAGCCGTGACTCTGGCAGAGCTCTCGCGGAGGTCCTCAAGA

AACCCGCTGTGGATAGGTCTCAGAAAATTGGAACCAAAGCCAACTCCAGCTCAAAGGGTCAGCTTCTTAAGGATCTCCAG

CTAAAACCTGTCGACAAGCAGGCAAGCTCTGCAATCGGGTTTGTCCCATCCGACCATGAATCATCCAGAAATGTCATCCG

CTCCATAATCAAGTCGAGCAAGCTAAACATTGATCACAAGGACTATCTTCTAGATTTGCTAAATGATGTGAAAGGCTCTA

AGGATCTCAAGGAATTCCACAAGATGCTAACAGCAATTCTCGCCAAGCACCTGTAACACACCCTCCAGTCACCATCTCAT

ACTCGGCTGAAAACATCCTCTCAATCAGGCTATTACAAAAAACTTAGGAGCAAGGGTAACTGAGCTTCGCAGACAAGATG

ACCGAGATCTACGACTTCGATAAATCAGCATGGGATGTCAAAGGGTCAATTGCTCGCATAGAACCCACCACCTATCACGA

CGGTCGACTGATACCCCAGGTAAGGGTTATCGATCCTGGTCTGGGAGACAGAAAAGATGAGTGCTTCATGTACCTGTTTC

TCCTAGGAGTGATTGAGGATAACGACCCCCTGTCTCCCCCAGTTGGAAGAACCTTTGGTTCTTTACCTCTAGGGGTCGGT

AGGTCAACTGCTAAGCCAGAAGAACTACTAAGGGAGGCCACAGAACTAGATATAGTGGTGAGGCGCACTGCAGGATTAAG

TGAGAAACTGGTATTTTACAACAACACTCCGCTGTCCTTGCTAACGCCCTGGAAGAAGGTTCTGACAACCGGAAGTGTGT

TTAGTGCTAACCAGGTTTGCAATGCAGTCAACCTAGTCCCGCTTGATACTCCCCAGAGATTCAGGGTTGTGTACATGAGC

ATAACTAGATTATCAGACAATGGTTACTATAGTGTGCCCAGGAGAATGCTGGAGTTCCGCTCAGCCAATGCAGTAGCCTT

CAACATCTTGGTTACACTAAGAATTGAAAATGGCACAAGCCCTAGAAGATACATAGTCGGCTCATGGGAGAATCCAGAGG

TCACATTTATGGTACACGTGGGTAACTTTAGAAGAAAGAAGAACGAGGTATACTCAGCTGATTACTGCAAAATGAAAATT

GAAAAGATGGGTTTAGTTTTTGCCCTGGGAGGAATAGGTGGAACAAGTCTCCATATTAGAAGCACCGGGAAAATGAGCAA

AACCCTCCATGCACAGCTGGGGTTCAAGAAAATCTTATGTTACCCCCTAATGGATGTTAATGAGGATCTCAACCGATATC

TCTGGAGGGCAGAGTGCCGAATAGTCAAAATCCAAGCCGTCTTACAGCCATCAGTACCCCAAGAATTCCGTGTCTACGAT

GATGTCATCATCAACGATGATCAAGGCCTATTCAAGATCCTGTAGTTCATCTGCAAAGGAATGAACCCACCCCCAGGAAG

GGGCCCCACCTTCCCCCG-CCAGAAAACACACCCCAAGGCTCCCCGACACAACACCAACACAGCCACTCTCCCGATGGGA

CGAGGCCAGCC-CCACAACAAACCAGGTCCAGGG-GGGGGGCCCCCCCCC-GACCCGCCGGGCGCCGACCGAGCAGGTCG

GCGAGACGCCTCCAGGAGCCCCCGACGAAGACCGAGCACCCCATCCCCCCAAAAATGCCCCCCC-AACACGCCCGAGGCC

AGGCGCCGGGCACCCCCAGACCCGCCACCGGCGAGACAGAACCCCAAGGAGGGATCTCCCAAACCAAAGGGATCACCGAG

GAAAGCTCCGCCGAACCAGCACAGATCCGACCCAAGCAAGGCGGACCCCCAACACGAAACCGCCCAGCGCACCTCACCTA

TCATCAAACAAAACTTAGGGGCCAGGTCCATACGACAGGCCCCAGCCCGCCAGGCGAAGGGCCGCAGAAAGGAAGGAGAC

ACGACCGCCCCAGAAGGAGGACACCCAACCACCGAAACACCCAGCCCCAGGGAAGGAG---CCCCCCCCACACGGCGAAG

GAGCGGGTGAGAGACCCCGAGGAGAACCCGACCGGAAGCCAGGAGCCGGCGACGCCGGCGGCAAACAGGCCGGGCGCACA

CCCGACTCCGCCAGGGGAGCCGAGGAGAGCCCGACCCCGCGTGGACCAAACGGCAGGGAATCCAACCCCGCCAAGGAAGG

AGGTGGGCCCCTTTTT-----CCCCCCCGCACACAACCTGCGGACAGACCGAGGGAGGACCACCTCCCAACAAGACAGGC

GGCTCCCTCGCAGCCGAGCCGCATCTCCCCAGAAGAGGACACCCCCGCACACTCCGAACACCCCCCCGCCCCCCGCCACC

GGGACAGAGGCTCCGCCACTACAGACCAGAGCCACCGAGAGCCAGCCAGAGGTCCCCTCCAGTCCCGTCCAGTGCCCCCC

CGAAGAGGCCCCGGCAAGCCCCCCCAGCCCGACGACCCCTGCACGCACCGGAAGAGGGACCCGCCCGAGAACGTCCGGCA

CATCCATGTGTAAACATCATGACACGGGTCGCAATCTTGACATTTCTGTTTCTTTTCCCAAATGCTGTTGCGTGCCAGAT

TCACTGGGGCAATCTATCCAAAATCGGGATTGTAGGAACAGGGAGTGCCAACTACAAGGTGATGACTAGGCCAAGCCACC

AAACTCTGGTTATAAAGTTAATGCCAAATATAACAGCCATCGACAATTGTACAAAGTCAGAGATTGCAGAGTACAAGAGA

TTGCTGATCACAGTGTTAAAGCCTGTAGAGGATGCTCTATCAGTGATAACCAAGAATGTAAGACCAATTCAAACTCTGAC

ACCTGGGCGTAGAACTCGCCGTTTTGCTGGAGCTGTTCTAGCCGGAGTAGCACTTGGAGTTGCGACAGCCGCTCAGATAA

CTGCAGGAGTCGCCCTTCATCAATCATTGATGAACTCCCAAGCAATTGAGAGTTTAAAAACCAGTCTTGAGAAGTCGAAT

CAGGCAATAGAAGAAATCAGACTTGCAAATAAGGAGACCATACTAGCAGTACAGGGCGTCCAGGATTATATCAACAATGA

GCTTGTCCCTTCTGTTCATAGAATGTCATGCGAGCTGGTAGGTCACAAGCTCGGCCTCAAGCTCCTTAGGTACTACACTG

AGATCCTGTCCATATTCGGGCCCAGCCTTCGAGACCCGATAGCTGCCGAAATATCAATCCAAGCACTCAGTTATGCATTA

GGCGGGGACATTAATAAAATCCTGGACAAGCTTGGGTATAGCGGTGGGGATTTCCTTGCCATCCTAGAAAGCAAGGGAAT

AAAGGCCCGGGTCACATATGTGGATACAAGAGATTACTTTATAATTCTTAGCATCGCCTACCCAACCTTATCTGAGATCA

AGGGAGTGATAGTTCACAAGATAGAAGCTATAACATACAACATTGGGGCACAGGAGTGGTATACTACTGTCCCTAAGTAT

GTAGCCACTCAGGGGTATCTGATATCGAACTTTGATGAGACGTCATGCGTATTCACTCCGGAGGGGACAGTTTGCAGCCA

GAATGCGTTGTACCCAATGAGCCCATTGCTCCAGGAATGTTTCCGGGGGTCAACAAAATCGTGCGCCAGAACCCTAGTTT

CAGGGACCATAAGCAATAGATTTATCCTATCAAAAGGGAACCTGATTGCAAATTGTGCGTCAGTCTTGTGCAAATGTTAC

ACAACAGAGACAGTTATTAGCCAAGATCCTGACAAACTACTAACTGTTGTAGCATCCGACAAGTGTCCTGTAGTTGAGGT

GGATGGAGTGACAATACAGGTCGGCAGTCGAGAGTATCCGGATTCTGTATACTTACACAAAATAGACTTAGGTCCAGCCA

TCTCCCTAGAAAAACTGGATGTAGGCACCAATTTAGGCAATGCAGTCACAAGACTGGAAAATGCAAAGGAGCTCCTAGAT

GCATCAGACCAAATACTGAAGACTGTTAAAGGGGTACCTTTCGGTGGGAATATGTATATAGCACTGGCAACTTGCATTGG

GGTATCCCTAGGGCTTGTCACATTAATATGCTGCTGCAAGGGGAGGTGTAAGAACAAGGAAGTTCCCATCTCCAAAATCA

ACCCAGGGCTCAAACCCGACCTGACTGGGACCTCAAAGTCGTACGTGAGATCACTGTAGTCAGAATCACTTGAATCATCC

GGAATCACACACATACATGTACGACACAAGCAGTCAGAGGACGCAGGAAACCCAGCCCCCTATCACCCACCTGACCCCAC

CCCATGCTCCACCACACATTAGTCATCAAACAAAACTTAGGACGAAAGGTCAATCACCATGTCCGCACAAAGGGAAAGGA

TCAATGCCTTCTACAAAGACAATCCTCACAATAAGAACCATAGGGTGATCCTGGATAGAGAACGCTTGGTCATTGAAAGA

CCCTACATCTTGCTTGGAGTCCTGCTGGTAATGTTCCTGAGTCTAATCGGACTGCTGGCCATTGCAGGGATCAGGCTTCA

CCGGGCCACCGTTGGAACTTCAGAGATCCAGAGTCGACTGAATACCAATATCGAGTTGACCGAATCTATTGATCACCAAA

CTAAGGATGTCTTAACTCCCCTTTTCAAAATCATTGGCGATGAAGTCGGCATCAGAATTCCACAGAAATTCAGTGATCTT

GTCAAGTTCATCTCCGATAAGATTAGATTCCTCAACCCTGATAGAGAGTATGATTTCAGGGATCTCCGGTGGTGTATGAA

TCCCCCCGAGAGAGTCAAAATTAATTTTGATCAGTTTTGTGAGTACAAGGCTGCGGTTAAGTCAATTGACCATATATTTG

AGTCGCCACTCAATAAGTCAAAAAAGCTGCAATCATTGACTCTTGGGCCCGGAACAGGCTGTCTAGGCAGGACAGTAACA

AGAGCTCATTTCTCGGAACTTACACTGACCTTAATGGACCTGGATCTAGAGATGAAACACAACGTATCCTCAGTGTTTAC

CGTAGTTGAAGAGGGATTATTCGGAAGAACATATACCGTCTGGAGATCCGATGCCAGGGATCCGAGCACCGATCTAGGTA

TTGGTCACTTCTTAAGAGTCTTCGAGATTGGACTGGTAAGAGATCTCGGGCTGGGTCCCCCTGTTTTTCATATGACCAAC

TATCTCACAGTGAACATGAGTGATGACTATCGGAGATGTCTTTTAGCGGTAGGGGAGTTAAAGTTGACAGCCCTATGCAC

CTCATCTGAGACTGTGACACTGAGTGAGAGAGGAGTTCCAAAGAGGGAGCCTCTTGTAGTTGTGATACTCAACCTAGCTG

GGCCCACTCTAGGGGGCGAACTATACAGTGTCTTGCCTACCTCTGATCCCATGGTGGAGAAACTCTATTTATCTTCACAT

AGAGGGATTATCAAAGATGACGAAGCCAATTGGGTAGTGCCGTCTACCGATGTTCGTGATCTTCAAAACAAAGGAGAATG

TCTGGTAGAGGCATGCAAGACTCGACCTCCTTCATTTTGCAATGGCACAGGATCAGGCCCGTGGTCAGAGGGGAGAATCC

CTGCTTACGGGGTGATCAGGGTCAGTCTTGACTTAGCTAGTGACCCGGATGTAGTTATCACTTCAGTGTTTGGCCCACTG

ATACCTCACCTATCCGGCATGGATCTTTACAACAACCCGTTTTCAGGAGCTGTATGGTTGGCTGTACCACCTTATGAGCA

GTCATTTTTGGGAATGATTAATACGATTGGATTCCCTAACAGAGCAGAGGTTATGCCGCACATTTTGACCACAGAGATCA

GAGGCCCTCGGAGCCGTTGCCATGTTCCCATAGAATTGTCCCGCAGGGTTGATGACGATATCAAGATTGGGTCCAACATG

GTCATATTGCCGACGATGGACCTGAGATATATTACAGCCACTTATGATGTTTCCAGGAGCGAGCATGCAATCGTGTACTA

TATCTATGACACAGGTCGCTCATCATCTTACTTCTACCCAGTTCGGCTGAACTTCAAAGGCAATCCTCTCTCTCTGAGGA

TAGAGTGTTTCCCTTGGCGTCATAAGGTGTGGTGCTACCATGATTGTCTTATATACAACACCGTAACAGATGAAGAAGTC

CATACGAGAGGGCTGACCGGTATAGAGGTAACATGTAATCCAGTCTGAGCAGGGCTACGACCGCCACTCAGGCAGTCCCT

TGAGTCGCCACCGAGTCCAAGCAGCACAGCCTGGGACACTCAACAGCACAGCCCAGCCAACAATGTTATAAAAAACTTAG

GAGCCAGGGCTGTAGGGGCCATGGACTCCCTATCAGTCAATCAGGTCTTGTACCCTGAGGTCCATCTAGATAGCCCTATT

GTCACAAACAAACTAGTTGCCATCCTTGAGTACTCGGGTATCGACCATAACTATGTTCTTGAAGACCAGACCCTTGTCAA

GAATATTAGGTATAGACTGGGGTGCGGTTTTTCAAATCAAATGATCATCAATAACAGGGGGTTGGGTGAAACAGTCAATT

CTAAACTCAAAAGTTACCCCCATAATCGTCATATCATATACCCGGATTGCAATAAGGAGTTGTTTTGTATCAAAGATAGC

TGCATATCTAAGAAGCTCTCGGAGCTATTCAAGAAGGGTAATTCCTTGTACTCTAAGATAAGTCACCAGGTACTGGATTG

TCTTAAGAGAGTCAATGGGAAATTAGGCCTGGGCACAGATCTTACCCATGGTCTGAAGGAGGGTATCCTTGACCTAGGGT

TACACATGCATAGCTCTCAATGGTTCGAGACTTTCCTGTTCTGGTTCACTATCAAGACAGAGATGAGGTCAATGATCAAA

GAACAGTCCCATATATGCCACAAGAGGAGGTATAACCCGATTTTTGTGTCGGGGGATGCATTCGAGGTGCTCGTATCACG

AGACCTCGTTGTGATAATTGATAAGAATACCCAGTATGTCTTCTACTTAACATTTGAGCTGGTCCTTATGTATTGTGATG

TCATAGAGGGCAGACTTATGACGGAGACAGCCATGGCCATAGACCAGAGATATTCAGAACTTCTAAGCCGAGTCAGATAC

TTGTGGGATCTTATTGATGGGTTCTTCCCAACACTAGGCAACACCACATACCAAGTTGTTGCTCTGCTTGAACCATTATC

GTTGGCTTACCTTCAACTACAGGATGTCACTCTGGAGTTAAGAGGTGCTTTTTTAGACCATTGCTTCAAAGAACTCTATG

AGATACTGGAGCATTGCGGCATTGGCACAGAAGGCACCTACAATTCCATCACTGAAGGATTGGATTACGTATTCATCACC

CACGATATACATCTAACTGGGGAGATTTTTTCATTCTTTCGGAGTTTCGGACACCCCCGTCTAGAAGCAGTCACCGCTGC

AGAAAATGTCAGAAAGCATATGAACCAACCGAAGGTAATCAGTTATGAGACTATGATGAAAGGGCATGCAGTATTTTGCG

GGATAATCATAAATGGTTTTAGGGACCGACATGGCGGCAGCTGGCCCCCTGTTGTATTGCCGGAACATGCTTCTGCTGCG

ATCCGGAATGCACAGGCATCAGGTGAGGGACTGACCCATGACCTGTGTATAGACAATTGGAAGTCCTTTGTTGGATTCAA

ATTTGGCTGCTTTATGCCACTCAGCTTAGATAGTGATTTGACCATGTACCTCAAAGACAAGGCATTGGCTGCACTGAAGA

ATGAGTGGGATTCAGTTTACCCGAAAGAATACCTCCGTTATAACCCACCTAGAGGGACAGAGTCAAGGCGATTGGTGGAG

GTGTTCCTGAATGACTCCAGCTTTGATCCTTATAACATGATAATGTACGTGGTGAATGGCTCCTACCTTAAAGACCCTGA

GTTTAATCTCTCATACAGCCTAAAAGAGAAGGAGATAAAGGAGACAGGGCGGTTGTTCGCCAAAATGACCTATAAGATGC

GGGCCTGTCAGGTAATCGCTGAAAATCTGATATCGAATGGAGTTGGGAAATATTTCCGAGACAATGGGATGGCAAAGGAC

GAGCATGACCTAACAAAAGCCCTTCACACTCTGGCAGTCTCAGGTGTTCCCAAGAACAACAAAGACAACCACCGAGGTGG

GCCTCCCAGAAGAACAACAAACCGAGAGATGAGATCAAGCCAAGACACCAAAACACAAAATAGAGACAAGGTCCAAGGGG

GGCCTATGTACAACTATTTGCGATGCCAACCAATCAACCCTGATCAAGGTGAGTCATACGAGACTGTTAGTGCATTCATC

ACCGCTGACCTTAAGAAGTATTGCCTGAATTGGAGATACGAGACAATCAGCATATTTGCACAGAGGCTGAATGAAATATA

TGGACTACCATCCTTCTTTCAATGGTTACACAGGATATTGGAAAAATCCGTACTCTACGTCAGTGACCCACATTGCCCTC

CCGACCTAGATAATCATATCCCTCTGGACAGTGTCCCTAATGCCCAAATATTCATTAAGTACCCAATGGGCGGAATAGAA

GGTTATTGCCAAAAACTATGGACAATCAGTACTATACCATACTTGTATCTGGCAGCCTATGAGAGCGGAGTAAGAATCGC

CTCACTAGTGCAGGGTGACAATCAGACAATCGCGGTGACAAAAAGAGTTCCAAGTTCTTGGCCCTATTCACTAAAAAAGA

GGGAGGCATCCAAAGCTGCTCAAAATTACTTCGTAGTCTTAAGGCAAAGATTGCACGATGTAGGTCATCACTTAAAGGCT

AATGAAACCATAGTATCTTCTCACTTTTTTGTATATTCTAAGGGGATTTATTATGACGGCCTGCTAGTCTCACAATCACT

AAAGAGCATCGCCAGATGTGTCTTCTGGTCCGAGACTATAGTGGATGAAACCAGAGCGGCTTGCAGCAATATTGCAACAA

CTGTCGCCAAGAGTATAGAGAGGGGTTATGATAGGTACCTTGCATATTCTTTGAATATCCTCAAGATTCTCCAACAGATC

CTTATATCCCTTAATTTCACTATTAACACAACAATGACTCAGGATGTCGTGGCACCGATCATCGAAAACGGGGATTTGCT

GATAAGGATGGCACTCTTGCCAGCACCCATCGGGGGTCTCAATTATCTTAACATGAGCAGGTTATTTGTGAGAAATATCG

GTGACCCGGTCACCTCCTCTATAGCCGACCTGAAGAGGATGATAGATGCTGGGCTAATGCCAGAAGAAACATTGCATCAA

GTGATGACCCAGACCCCGGGAGAATCATCCTACCTCGATTGGGCAAGTGACCCTTATTCTGCCAACCTACCCTGCGTACA

GAGTATAACTCGCCTTCTCAAAAACATCACTGCACGATATATTTTAATCAGCAGCCCAAACCCGATGCTGAAAGGATTGT

TTCATGAGGGGAGTAGAGATGAAGACGAGGAGCTTGCAAGTTTCTTAATGGATCGGCATATAATTGTTCCAAGAGCTGCA

CATGAAATCTTAGACCATAGCATAACCGGGGCAAGAGAAGCTATAGCCGGGATGTTGGACACCACCAAGGGTCTGATTAG

AACAAGTATGAAACGGGGTGGCCTCACCCCTCGAGTCTTAGCCCGCCTTTCCAATTATGATTATGAACAATTCCGATCCG

GGATAACACTATTGACAAAGAAAGGGCAGTGTTATCTCATTGACAAGGACTCGTGCTCGGTGCAGCTCGCTATAGCTCTG

AGGAGCCACATGTGGGCTAGGTTGGCTCGCGGGAGACCAATCTATGGCTTGGAGGTGCCCGATATATTAGAATCTATGAA

CGGCTACCTTATCAAACGCCACGAGTCTTGTGCTATCTGTGAGACGGGCTCAAGTCACTACGGATGGTTTTTTGTCCCTG

CAGGGTGCCAGCTTGACGATGTATCTAGAGAGACTTCAGCTCTCCGTGTACCTTATGTCGGATCAACAACTGAGGAGAGG

ACAGACATGAAACTTGCTTTTGTTAGGGCTCCAAGTCGATCCCTTAAATCAGCAGTCAGAATTGCAACAGTTTACTCATG

GGCCTACGGGGATGATGAGAAATCATGGGGTGAAGCCTGGATGCTAGCTAGGCAGAGAGCCAACATCACCTTAGATGAAT

TGAGAATGATCACTCCAGTCTCTACATCCACCAACCTAGCCCACCGGTTGAGGGATCGGAGCACCCAGGTGAAATACTCG

GGGACATCCCTTGTGAGGGTTGCAAGATACACAACCATCTCCAATGATAACCTGTCATTTGTGATATCTGAGAAAAAAGT

GGATACCAACTTCATTTATCAGCAAGGGATGCTGCTCGGTCTTGGGATCCTTGAGAACCTCTTTAGGCTAGAGGCCACCA

CCGGAGTATCTAACACAGTGCTACACCTGCACGTGGAAACAGAATGTTGTGTTGTACCTATGGTGGATCACCCAAGGATC

CCGAGTCTCCGTAATATAAAAGTTACGGATGAGCTATGCACAAACCCTTTGATCTACGACAGGTCTCCCATCATAGAGCA

CGATGCAACCCGACTGTACTCACAGAGCCACAGGAGACATCTGGTGGAGTTTGTTACCTGGTCAACAAGCCAGCTTTACC

ATATACTGGCTAAGTCTACAGCTATGTCCATGATTGAGTTGATCACGAGATTCGAGAAAGATCACATGAATGAAATAGCC

GCCCTGATTGGCGATGACGACATCAACAGCTTCATCACAGAATTTTTGCTAGTAGAGCCTAGATTATTTATAGTTTACCT

TGGTCAGTGTGCTGCCATCAATTGGGCCTTCGATATACATTATCATCGGCCCTCAGGCAAGCACCAGATGGGGGAACTCC

TCTACTCTCTGCTCTCTCGGATGAGCAAAGGAGTATATAAGATCTTCACCAATGCTTTGAGCCACCCCAAAGTTTACAAG

AAATTCTGGCGAAGTGGTGTCATTGAGCCGATTCATGGCCCATCCCTGGATACACAGAATTTACATGTCACTGTCTGTGA

CATGATATATGGGTCCTACGTCACCTATTTGGATCTTTTGCTGAATGATGAGCTAGATGATTACCCGTATTTACTCTGCG

AGAGTGATGAGGACGTGGTCACAGACAGGTTCGACAACATTCAAGCCAAACATCTCTGTGTACTGGCCGATGTATATTGC

AGCTCCAAGAGGTGTCCTTCGATAATCGGGATGTCTCCTATAGAAAAATGTACCATTCTCACACATTACATCAAAGGAGA

ATCAGTACAATCCCCGTCAGGGACCTCATGGAACACTGATCCCCTTGTAGTAGATCATTACTCATGCTCTCTGACCTACC

TTCGCCGCGGTTCCATCAAACAAATCAGGTTGAGAGTGGATCCTGGGTTTGTATTCGAGGCGTTGACAGACGTCGACTTT

AAACAGCCTCGCAAGGCTAAGTTAGATATATCGGTTGTTGGGTTGGCTGATTTCTCTCCCCCTTGCGACAACGTTGGTGA

TTTCCTGGGGACTATCAACACATTGAGGCACAATCTGCCTGTCACCGGAACCGGGGTCTCGAACTATGAAGTCCACGCTT

ATCGTAGAATTGGCCTAAATTCATCAGCATGTTATAAAGCCGTAGAGATCTCCACGTTAATCAAGCCATCCTTAGAAGTC

GGAGAGCATGGATTGTTCCTAGGAGAAGGTTCCGGTTCAATGCTGGCTGCGTACAAGGAAGTTCTTAAATTAGCAAATTG

TTATTACAACAGCGGGGTAACAGCGGAGGGTAGAGCCGGACAGAGGGAAATATCTCCCTATCCTTCAGAGATGAGCCTAG

TAGAGAATCAAATGGGGATAGAGCGGAGTGTTAAAGTGCTGTTCAACGGCAAACCTGAAGTAACTTGGGTGGGGACCACC

GATTGCTACAAGTACATAATCAGTAACATCCAAACCTCTAGTCTAGGTTTCATACATTCAGATATTGAGACACTCCCAAC

CAAAGACGCTGTTGAGAAATTGGAAGAATTTGCCTCTATCCTATCCTTATCCTTGATTTTGGGAAAAATCGGCTCTATCA

CAGTTGTCAAAATTATGCCCATTAGCGGGGATTTTACCCAAGGCTTCATAGCCTATGCTATCCAATATTTTAGGGAGAGC

CTGCTTACCTATCCGAGATATAGTAACTTCATCTCGACTGAGTGTTACCTTGTTATGATAGGATTAAAGGCCAATCGATT

GATAAACCCAGAAGCCATTAAGCAAAGCATAATCAGAGCGGGGGTCAGGACTGCACCAGGACTTGTGAGCCATATATTAT

CAGAGAAACAGAAAGGTTGTATTCAATCTTTCTTGGGTGATCCTTATATCCAAGGAGACTTCAATAAGCACCTTAAATCT

CTAACCCCTATTGAGAAAATCCTGGTAAATTGTGGTCTCTCGATCAATGGCACAAAAATCTGTAGAGATCTAATCCACCA

TGATATAGCCTCCGGTCCAGACGGTCTTATGAGCTCTACAATTATTTTATATAGGGAGTTGGCTCATTTCAAAGATAATA

TAAGGAGTCAGCACGGAATGTTCCACCCCTACCCAGTACTGGCCAACAGCAGGCAACGTGAATTAATCCTTCGAATAGCC

AAGAAATTCTGGGGGTATGTCCTGCTATATTCTGATGACCCGGCACTAATCAGACAAACAATCAAGAACTTGAAACGGAA

TCATCTAACCTTTGACTTACATAGTAACCCGTTTATTAAAGGGCTATCCAAAGCTGAGAAACTGCTGGTGCGGACGAGTT

CACTAAGAAGGGAATGGTTGTTCACTCTCGATACGAAAGAAGTGAAAGAGTGGTTCAAATTGGTAGGTTACAGTGCACTC

ATTAGAGGCTAGTTGGCTATACATCTGCCCCCTTTCCCTCCGCCATAAGACCCCAATGACGATCCAAAAGATTAAAGAAA

ACTACATATTGGATAAGTATCTATTCCCANNNNNNNNNNNN

>KR140086.1/goat/India/1994

ACCAAACAAAGTTGGGTAAGGATAGATCTTATAATAACTATGGACTGGCAAACTTAGGAGTAAAGATCCTACTGTCGGGG

GGAGGAGGAGGAGCAAGATCTTTGATTATGGCGACTCTCCTTAAAAGCTTAGCATTGTTCAAAAGGAACAAAGACAAAGC

GCCGACGGCATCAGGTTCAGGAGGGGCCATCCGGGGGATTAAGAATGTTATCATAGTCCCGATTCCCGGAGACTCGTCCA

TCATTACCCGTTCAAGACTGCTCGACAGGCTTGTCAGATTGGCCGGAGATCCTGACATCAACGGGTCAAAGCTGACCGGC

GTGATGATCAGCATGTTATCTTTGTTCGTAGAGTCACCCGGGCAATTGATACAGCGAATCACAGATGATCCAGATGTTAG

TATCCGCCTTGTTGAGGTAGTTCAAAGTACTAGGTCCCAGTCCGGGTTGACCTTTGCATCACGTGGTGCTGATTTGGACA

ACGAGGCAGACATGTATTTTTCAACTGAGGGGCCCTCGAGTGGAGGTAAGAAAAGGATCAACTGGTTTGAGAACAGAGAA

ATAATAGACATAGAGGTGCAGGATCCAGAAGAGTTCAATATGTTGTTAGCCTCCATACTAGCACAAGTCTGGATCCTTCT

GGCCAAGGCTGTTACGGCACCGGATACGGCAGCTGACTCAGAACTGAGAAGGTGGGTTAAATACACACAACAAAGGAGAG

TGATTGGGGAATTTCGCCTTGACAAAGGGTGGCTGGATGCAGTCCGCAACAGAATTGCAGAAGATCTATCACTCCGGCGG

TTCATGGTATCTCTCATACTTGACATCAAGAGGACCCCCGGCAACAAGCCAAGGATTGCAGAAATGATCTGCGACATTGA

CAATTATATTGTCGAAGCCGGACTCGCCAGTTTCATCCTTACTATCAAGTTTGGTATTGAAACCATGTATCCTGCATTAG

GGCTCCACGAGTTTGCCGGGGAATTGTCCACTATTGAGTCCTTGATGAACTTGTATCAACAGCTAGGAGAGGTTGCACCC

TACATGGTAATTCTAGAGAACTCAATTCAGAACAAGTTTAGTGCAGGAGCTTATCCCCTCCTCTGGAGCTATGCGATGGG

TGTCGGAGTTGAGCTGGAGAACTCAATGGGGGGCTTGAACTTTGGCAGGTCATATTTTGACCCTGCCTATTTTCGTCTCG

GACAGGAGATGGTCAGAAGATCTGCAGGAAAGGTCAGCTCTGTGATTGCGGCTGAGCTCGGCATCACAGCAGAGGAAGCT

AAACTAGTCTCGGAAATCGCCTCGCAGGCTGGGGACGAAAGAACCGCTAGAGGGACTGGGCCTCGACAGGCGCAGGTCTC

CTTCCTCCAGCACAAAACAGGAGAGGGAGAGTCGTCCGCACCAGCGACCAGAGAAGGGGTCAAGGCTGCGATCCCAAACG

GATCCGAAGAAAGGGACAGGAAGCAAACACGCTCAGGAAGGCCCAGAGGAGAGACCCCCAGCCAACTGCTCCTGGAAATC

ATGCCAGAGGATGAGGTCYCGCGAGAGTCTGGCCAAAACCCTCGTGAGGCCCAGAGATCGGCCGAGGCACTCTTCAGGCT

GCAGGCCATGGCCAAGATTCTGGAGGACCAGGAGGAGGGAGAAGACAACAGTCAGGTCTACAACGACAAGGATCTCCTCG

GCTAAGCAGACGCACCCTCTGTCGAAATCAGTGACAAGACATCTCCTACCAGTATTATAAAAAACTTAGGACCCAGGTCC

AAGCAAGCACACATCGACACTCCAACCAGTCGAGCGGAGACCACCGATGGCAGAAGAACAAGCATACCATGTCAACAAGG

GGCTGGAATGTATCAAGTCTCTCAAAGCCTCTCCCCCGGATCTATCCACCATCAGAGATACCATCGAGAGCTGGAGAGAG

GGGCTTAGCCCCTCGGGCCGTGCAACACCGAACCCTGATACGTCCGAGGGAGATCATCAGAATATCAACCAATCATGCTC

ACCAGCAATCGGACCAAACAAAGTCTACTTGTCTCCTGAAGATAATCTCGGATTTAGAGAGATCACTGGCAACGACTGTG

AGGCTGGGCTCGGAGGAGTCCAGGGAAAAGGATCCAACTCTCAAGTACAGCGTTACTATGTTTATAGCCACGGGGGTGAA

GAGATTGAAGGACTCGAGGATGCTGACTCTCTCGTGGTTCAAGCAGATTCTCCAGTTGCTAACATCTTCAATGGAGGAGA

GGATGGATCTGACGACAGCGATGTGGACTCTGGCCCAGATGATCCCGGCAGAGATACTCTATATGACCGGGGATCTGTTG

CCGGCAATGATGTCGCTAGGTCCACAGATGTCGAGAAACTAGAAGGTGCTGATATTCAAGAAGTTCTTAACTCCCAGAAA

GGCAAAGGAGGAAGATTCCAAGGCGGGAAAACCTTACGAGTCCCGGAAATACCCGATGTCAAGCACTCCAGACCATCAGC

CCAATCAATTAAAAAGGGCACAGACGGGAACTCAGTCTCATCTGGAACGGTGACAGAGTGTTTATCGATAAGTGGTGCAA

CCCAAGCTGTGCCAGAGTCAAGATGGGAGTCATCAGAGCAAAATGCGTCTGTGGGGAGTGTCCTCAAGTCTGCGAGGAGT

GCAAAGACGATCCAGGAGTCGACACAAGAATCTGGTACCATAGCATCACTGACTCAGCCTAAAGAGAATGACTCCGAGTA

TGAGTATGAGGATGACCTATTTACAGAGATTCAGGACATCCGTGCAAGCATTGCCAAGATCCATGATGACAACAAAACTA

TCCTCTCAAAGCTTGATTCTATACTGTTATTGAAAGGAGAAGTCGACACTATCAAGAAACAAATCAGCAAGCAGAATATA

AGTATATCCACCATTGAGGGCCATCTCTCCAGTATAATGATAGCCATCCCGGGCTTTGGGAAGGATATCAAGGACCCAAC

ATCCGAGGTTGAGTTGAACCCAGATTTAAGACCTATAATAAGCCGTGACTCTGGCAGAGCTCTCGCGGAGGTCCTCAAGA

AACCCGCTGTGGATAGGTCTCAGAAAATTGGAACCAAAGCCAACTCCAGCTCAAAGGGTCAGCTTCTTAAGGATCTCCAG

CTAAAACCTGTCGACAAGCAGGCAAGCTCTGCAATCGGGTTTGTCCCATCCGACCATGAATCATCCAGAAATGTCATCCG

CTCCATAATCAAGTCGAGCAAGCTAAACATTGATCACAAGGACTATCTTCTAGATTTACTAAATGATGTGAAAGGCTCCA

AGGATCTTAAGGAATTCCACAAGATGCTAACAGCAATTCTCGCCAAACACCCGTAACACACCCTCCAGTCACCATCTCAG

ACTCGGCTGAAAACATCCTCTCAATCAGGCTATTACAAAAAACTTAGGAGCAAGGGTAACTGAGCTTCGCAGACAAGATG

ACCGAGATCTACGACTTCGATAAATCAGCATGGGATGTCAAAGGGTCAATTGCCCGCATAGAACCCACCACCTACCACGA

CGGCCGACTGATACCCCAGGTAAGGGTCATCGATCCTGGTCTGGGAGACAGAAAAGATGAGTGCTTCATGTACCTGTTTC

TCCTAGGAGTGATTGAGGATAACGACCCCCTGTCTCCCCCCGTCGGAAGAACCTTTGGCTCTTTACCTCTAGGGGTCGGT

AGGTCAACTGCTAAGCCAGAAGAACTACTAAGGGAGGCCACAGAACTAGATATAGTGGTGAGGCGCACTGCAGGAGTAAA

TGAGAAACTGGTATTTTACAACAACACTCCGCTGTCCTTGTTAACGCCCTGGAAGAAAGTTCTGACAACCGGAAGTGTGT

TTAGCGCTAACCAGGTTTGCAATGCAGTCAACCTAGTCCCGCTTGATACTCCCCAGAGATTCAGGGTTGTGTACATGAGC

ATAACTAGATTATCAGACAATGGTTACTATAGTGTGCCCAGAAGAATGCTGGAGTTCCGCTCAGCCAATGCAGTAGCCTT

CAACATCTTGGTTACACTAAGAATTGAAAATGGCACAAACCCTAGAAGATACATAGTCGGCTCATGGGAGAATCCAGAGG

TCACATTTATGGTACACGTGGGCAACTTTAGAAGAAAGAAGAACGAAGTATACTCTGCTGATTACTGCAAAATGAAGATT

GAAAAGATGGGTTTAGTTTTTGCCCTGGGAGGAATAGGTGGAACAAGTCTCCATATTAGAAGCACCGGGAAAATGAGCAA

AACCCTCCATGCACAGCTGGGGTTCAAGAAAATCTTATGTTACCCCCTAATGGATGTTAATGAGGATCTTAACCGATATC

TCTGGCGGGCAGAGTGCCGAATAGTCAAAATCCAAGCCGTCTTACAGCCATCAGTACCCCAAGAATTCCGTGTCTACGAT

GATGTCATCATCAACGATGATCAAGGCTTGTTCAAGATCCCGTAGTTCACCCTCAAAGGAATGAACCCACCCCCAGGAAG

GGGCCCCACCCTCCCCCG-CCAGAAAACACACTCCAAGGCTCCCCGACACAGCACCAACACAGCCACTCTCCCGATGGGA

CGAGGCCGGCC-CCCCAACAAACCCGGTCCAGGC-GGGGGGCCCCCCCCCTGACCTGCCGGGCGCCGACCGAGCAGGCCG

GCGAGAAGCCCCCAAGAGCCCCCGACGAAGACCGAGCACCCCATCCCCCC-AAAACGTCTCCCC-AACACGCCCGAGGCC

AGGCGCCGGGCACCCCCAGACCCGCCACCGGCGAGACAGAACCTCAAGGGGGGACCCCCCAAACCAAAGGGACCAACGGG

GGAGGCTCCGCCGAACCAGCACAGACCCGACCCAAGCAAGGCGGACCCCCAACACGAAATTGCCCAGCGCACCTCACTCA

TCATCAAACAAAACTTAGGGGCCAAGTCCATACGACAGGCCCCAGCCCGCCAGGCGGAGGGCCGCAGAAAGGAAGGAGAC

ACGATCGCCCCAGAAGGAGGACACCCAACCACCGAAACACCCAGCCCCAGGGAAGGAG--CCCCCCCCCACACGGCGAAG

GAACGGGTGAGAGACCCCGAGGAGAACCCGACCGGGAGCCAGGAGCCGGCGACGCCGGTGGCAAACAGGCCGGGCACACA

CCCGACTCCACCAGAGGAGCCGAGGAGAGCCTGACCCCGCGTGGACCAAACGGCAGGGAATCCAACCCCACCACAAAAAG

AGGTGAGGCCCC-CTT-----CCCCCCCGCACACAACCTGCAGACAGACCGAGGGAGGACCACCTCCCAACAAGACAGGC

GGCTCCCTCGCAGCCGAGCCGCAGCTCCCCAGAATAGGACACCCCCGCACACTCCGAACACCCCCTCGCCCCCCGCCACC

GGGACAGGGGCTCCGCCACCACAGACCAGAGCCACCGAGAGCCAGCCAGAGGTCCCCTCCAGTCCCGCCCGGTGCCCCCC

CGAAGAGGCCCCGGCAAGGCCCCCCAGCCCGACGACCCCTGCACGCACCGAAAGAGGGACCTGCCCGAGAACGTCCGGCA

CATCCATGCGTAAACATCATGACACGGGTCGCAATCTTGACATTTCTGTTTCTTTTCCCAAATGCTGTTGCGTGCCAGAT

TCACTGGGGCAATCTATCCAAGATCGGGATTGTAGGAACAGGGAGTGCCAGCTACAAGGTGATGACTAGGCCGAGCCACC

AAACTCTGGTTATAAAGTTAATGCCAAATATAACAGCTATCGACAATTGTACAAAGTCAGAGATTGCAGAGTACAAGAGA

TTGCTGATCACAGTGTTAAAGCCTGTAGAGGATGCTCTGTCAGTGATAACCAAGAATGTAAGACCAATTCAAACTCTGAC

ACCTGGGCGTAGAACTCGCCGTTTTGCTGGAGCTGTTCTGGCCGGAGTAGCACTTGGAGTTGCGACAGCCGCTCAGATAA

CTGCAGGAGTCGCCCTTCATCAATCATTGATGAACTCCCAAGCAATTGAGAGTTTAAAAACCAGTCTTGAGAAGTCGAAT

CAGGCAATAGAAGAAATCAGACTTGCAAATAAGGAGACCATACTGGCAGTACAGGGCGTCCAGGATTATATCAACAATGA

GCTTGTCCCTTCTGTTCATAGAATGTCATGCGAGCTGGTAGGTCACAAGCTCGGCCTCAAGCTCCTTAGGTACTACACCG

AGATCCTGTCCATATTCGGGCCCAGCCTTCGAGACCCGATAGCTGCCGAAATATCAATCCAGGCACTCAGTTATGCATTA

GGCGGAGACATTAATAAAATCCTGGACAAGCTTGGGTATAGCGGTGGGGATTTCCTTGCCATCCTAGAAAGCAAGGGAAT

AAAGGCCCGGGTCACATATGTGGACACAAGAGATTACTTTATAATTCTTAGCATCGCCTACCCAACCTTATCTGAGATCA

AGGGAGTGATAGTTCACAAGATAGAAGCTATAACATACAACATTGGGGCACAGGAGTGGTATACTACTATCCCTAAATAT

GTAGCCACTCAGGGGTATCTGATATCGAACTTTGATGAGACGTCATGCGTATTCACTCCAGAGGGGACAGTTTGCAGCCA

GAATGCGTTGTACCCAATGAGCCCATTGCTTCAGGAATGTTTCAGGGGGTCAACAAAATCGTGCGCCAGAACCCTAGTTT

CAGGGACCATAAGTAATAGATTTATCCTATCAAAAGGGAACCTGATTGCAAATTGTGCGTCAGTTTTGTGCAAATGTTAC

ACAACGGAGACAGTTATCAGCCAAGATCCTGACAAACTACTAACTGTTGTAGCATCCGACAAGTGTCCTGTAGTTGAGGT

GGATGGAGTGACAATACAGGTCGGCAGTCGAGAGTATCCGGATTCTGTATACTTACACAAAATAGACTTAGGTCCAGCCA

TCTCCCTAGAAAAACTGGATGTAGGCACCAATTTAGGCAATGCAGTCACAAGACTGGAGAATGCAAAGGAGCTCCTAGAT

GCATCAGACCAAATACTGAAGACTGTTAAAGGGGTACCTTTCGGTGGGAATATGTACATAGCACTGGCAGCTTGCATTGG

GGTATCCCTAGGGCTTGTCACATTAATATGCTGCTGTAAGGGGAGGTGTAAGAACAAGGAAATCCCTATCTCCAAAATCA

ACCCAGGGCTCAAACCCGACCTGACCGGGACCTCAAAGTCGTACGTGAGATCACTGTAGTCAGAATCACCTGAATCATCC

GGCATCACACACATACATGTACGACACAAGCAGTCTGAGGACGCAGGAAACCCAGCCTCCGATCACCCACCTGACCCTAC

TCCACACTCCACCACACATTAGTCATCAAACAAAACTTAGGACGAAAGGTCAATCACCATGTCCGCACAAAGAGAAAGGA

TCAATGCCTTCTACAAAGACAATCCTCACAATAAGAACCATAGGGTGATCCTGGATAGAGAACGCTTGGTCATTGAAAGA

CCCTACATCTTGCTTGGAGTCCTGCTGGTAATGTTCCTGAGTCTAATCGGACTGCTGGCCATTGCAGGGATCAGGCTTCA

CCGGGCCACCGTTGGAACTTTAGAGATCCAGAGTCGGCTGAATACCAATATTGAGTTGACCGAATCTATTGATCACCAAA

CTAAGGATGTCTTAACTCCCCTTTTTAAAATCATTGGCGATGAAGTCGGCATCAGAATTCCACAGAAATTCAGTGATCTT

GTCAAGTTCATCTCCGATAAGATTAAATTCCTCAACCCTGATAGAGAGTATGATTTCAGGGATCTCCGGTGGTGTATGAA

TCCCCCCGAGAGAGTCAAAATTAATTTTGATCAGTTTTGTGAGTACAAAGCTGCGGTTAAGTCAATTGAACATATATTTG

AGTCACCACTCAACAAGTCAAAAAAGCTGCAATCATTGACTCTTGGGCCCGGAACAGGCTGTCAAGGCAGGACAGTAACA

AGAGCTCATTTCTCAGAACTTACACTGACCTTAATGGACCTGGATCTAGAGATGAAGCACAACGTGTCCTCAGTGTTTAC

CGTAGTTGAAGAGGGATTATTCGGAAGAACATATACCGTCTGGAGATCCGATGCCAGGGATCCGAGCACCGATCCAGGTA

TCGGCCATTTTTTAAGAGTCTTCGAGATTGGACTGGTAAGAGATCTCGGGCTGGGTCCCCCTGTTTTCCATATGACCAAC

TATCTCACAGTGAACATGAGTGATGACTATCGGAGATGTCTTTTAGCGGTAGGGGAGTTGAAGTTGACAGCCCTATGCAC

CTCATCTGAGACTGTGACACTGGGCGAGAGAGGAGTTCCAAAGAGGAAGCCTCTTGTGGTTGTGATACTTAACCTGGCTG

GGCCCACTCTAGGGGGCGAACTATACAGTGTCTTGCCTACCTCTGATCTCATGGTGGAGAAACTCTATTTATCTTCACAT

AGAGGGATTATCAAAGATGACGAGGCCAATTGGGTAGTGCCGTCTACCGATGTTCGTGATCTTCAAAACAAAGGTGAATG

TCTGGTGGAAGCATGCAAGACTCGACCTCCTTCATTTTGCAATGGCACAGGATCAGGCCCGTGGTCAGAGGGGAGAATCC

CTGCTTACGGGGTGATCAGGGTCAGTCTTGACTTAGCTAGTGACCCGGGTGTAGTTATCACTTCAGTGTTTGGCCCACTG

ATACCTCACCTATCCGGCATGGATCTTTACAACAACCCGTTTTCAAGAGCTGTATGGTTGGCTGTGCCACCTTATGAGCA

GTCATTTCTAGGAATGATAAATACAATTGGATTCCCTAACAGAGCAGAGGTTATGCCGCACATTTTGACCACAGAGATCA

GAGGCCCTCGGGGTCGTTGCCATGTTCCCATAGAATTGTCCCGCAGGGTTGATGACGATATCAAGATCGGGTCCAACATG

GTCATATTGCCGACGATGGACCTGAGGTATATTACAGCCACTTATGATGTTTCCAGGAGCGAGCATGCAATCGTGTACTA

TATCTATGACACAGGTCGCTCATCATCTTACTTCTACCCAGTTCGACTGAATTTCAAAGGCAATCCTCTCTCTCTGAGGA

TAGAGTGTTTCCCTTGGCGTCATAAGGTGTGGTGCTACCATGATTGTCTTATATACAACACCATAACAGATGAAGAAGTC

CATACGAGAGGGCTGACCGGTATAGAGGTAACATGTAATCCAGTCTGAGCAGAGCTACGACCGTCACTCAGGCAGTCCCT

TGAGTCGCCACCGAGTCCAAGCAGCACAGCCTGGGACACTCAACAGCACAGCCCGGCCAACAATGTTATAAAAAACTTAG

GAGCCAGGGYTGTAGGGGCCATGGACTCCCTATCAGTCAATCAGGTCTTGTACCCTGAGGTTCATCTAGATAGCCCTATT

GTCACAAACAAACTAGTTGCCATCCTTGAGTACTCGGGTATCGACCATAACTATGTTCTTGAAGACCAGACCCTTGTCAA

GAATATTAGGTATAGACTGGGGTGCGGTTTCTCAAATCAAATGATCATCAATAACAGGGGGGTGGGTGAAACAGTCAATT

CTAAACTTAAAAGTTACCCCCATAATCGTCATATCATATACCCGGATTGCAATAAGGAGTTGTTTTGTATCAAAGATAGC

TGCATATCTAAGAAGCTCTCGGAGCTATTCAAGAAGGGTAATTCCTTGTACTCTAAGATAAGTCACCAGGTACTGGATTG

TCTTAAGAGAGTCAATGGGAAATTAGGCCTGGGCACAGATCTTACCCATGGTCTGAAGGAGGGTATTCTTGACCTAGGGT

TGCACATGCATAGCTCTCAATGGTTCGAGACTTTCCTGTTCTGGTTCACTATCAAGACAGAGATGAGATCAATGATCAAA

GAACAGTCCCATATATGCCACAAGAGGAGGTATAACCCGATTTTTGTGTCGGGGGATGCATTCGAGGTGCTCGTATCACG

AGACCTCGTTGTGATAATTGATAAGAATACCCAGTATGTCTTCTACTTAACATTTGAGCTGGTCCTTATGTATTGTGATG

TCATAGAGGGCAGACTTATGACGGAGACAGCCATGGCCATAGACCAGAGATATTCAGAACTTCTAAGCCGGGTCAGATAC

TTGTGGGATCTTATTGATGGGTTCTTCCCAACACTAGGCAACACCACATACCAAATTGTTGCTCTGCTTGAACCATTATC

GTTGGCTTATCTTCAACTTCAGGATGTCACTCTGGAGTTAAGAGGTGCTTTTTTAGACCATTGCTTCAAAGAACTCTATG

AGATACTGGAGCATTGCGGCATCGACACAGAAGGCACCTACAATTCCATCACTGAAGGATTGGATTACGTATTCATCACC

CACGATATACATCTAACTGGGGAGATTTTTTCATTTTTTCGGAGTTTCGGACACCCCCGTCTAGAAGCAGTCACCGCTGC

AGAAAATGTCAGGAAACATATGAACCAACCGAAGGTAATCAGTTATGAGACTATGATGAAAGGGCACGCAGTATTTTGCG

GGATAATCATAAATGGTTTTAGGGACCGACATGGCGGCAGCTGGCCCCCTGTTGCATTGCCGGAACATGCTTCTGCTGCG

ATCCGGAATGCGCAGGCATCAGGTGAGGGACTGACCCATGACCTGTGTATAGACAATTGGAAATCCTTTGTTGGATTCAA

ATTTGGCTGCTTTATGCCACTCAGCCTAGATAGTGATTTGACTATGTACCTCAAAGACAAGGCATTGGCTGCACTGAAGA

ATGAGTGGGATTCAGTTTACCCGAAAGAATACCTCCGTTATAACCCACCTAGGGGGACAGAGTCAAGGCGATTGGTGGAG

GTGTTCCTGAATGACTCCAGCTTTGATCCTTATAACATGATAATGTACGTGGTGAATGGCTCCTACCTTAAAGACCCTGA

GTTTAATCTCTCATACAGCCTAAAAGAGAAGGAGATAAAGGAGACAGGGCGGTTGTTTGCCAAAATGACCTATAAGATGC

GGGCCTGTCAGGTAATCGCTGAAAATCTGATATCGAATGGTGTTGGGAAGTATTTCCGAGACAATGGGATGGCAAAGGAC

GAGCATGACCTAACAAAAGCCCTTCACACTCTGGCAGTCTCAGGTGTTCCCAAGAACAACAAAGATAACCACCGAGGTGG

CCCTCCCAGAAGAACAACAAGCCGAGAGGTGAGATCAAGCCAAGACACCAAAACACAAAATAGAGACAAGGTCCAAGGGG

GGCCTATGTACAACTATTTGCGATGCCAACCAATCAGCCCTGATCAGGGTGAGTCATACGAGACTGTTAGTGCATTCATC

ACCGCTGACCTTAAGAAGTATTGCCTGAATTGGAGATACGAGACAATCAGCATATTTGCACAGAGGCTGAATGAAATATA

TGGACTACCATCCTTCTTTCAATGGTTACACAGGATATTGGAAAAATCCGTACTCTACGTCAGTGACCCACATTGCCCTC

CCGACCTAGATAATCATATCCCTCTGGACAGTGTCCCTAATGCCCAAATATTCATTAAGTACCCAATGGGCGGAATAGAA

GGTTATTGCCAAAAACTATGGACAATCAGTACTATACCATACTTGTATCTGGCAGCCTATGAGAGCGGAGTAAGAATCGC

CTCACTAGTGCAGGGTGACAATCAGACAATCGCAGTGACAAAAAGAGTTCCAAGTTCTTGGCCTTATTCACTAAAAAAGA

GGGAGGCATCCAAAGCCGCTCAAAATTACTTCGTAGTCTTAAGGCAAAGATTGCACGATGTAGGTCATCACTTAAAGGCT

AATGAAACCATAGTATCTTCTCACTTTTTTGTATATTCCAAAGGGATTTATTATGACGGCCTGCTAGTCTCACAATCACT

AAAGAGCATCGCCAGATGTGTCTTCTGGTCCGAGACTATAGTGGATGAAACCAGAGCGGCCTGCAGCAATATTGCAACAA

CTGTCGCTAAGAGTATAGAGAGGGGTTATGATAGGTACCTTGCATATTCTTTGAATATCCTCAAGATTTTCCAACAGATC

CTTATATCCCTTAACTTCACTATTAACACAACAATGACTCAGGATGTCGTGGCACCGATCATCGAGAACGGTGATTTGCT

GATAAGGATGGCACTCTTGCCAGCACCCATCGGGGGTCTCAATTATCTTAACATGAGCAGGTTATTTGTGAGAAATATCG

GTGACCCGGTCACCTCCTCTATAGCCGACCTGAAGAGGATGATAGATGCTGGGCTAATGCCAGAAGAAACATTGCATCAA

GTGATGACCCAGACCCCGGGAGAATCATCCTACCTCGATTGGGCAAGTGACCCTTATTCTGCCAACCTACCCTGCGTACA

GAGTATAACTCGCCTTCTCAAAAACATCACTGCACGATATATTTTAATCAGCAGCCCAAACCCGATGCTGAAAGGATTGT

TTCATGAGGGGAGTAGAGATGAAGACGAGGAGCTTGCAAGTTTCCTAATGGATCGGCATATAATTGTTCCAAGAGCTGCA

CATGAAATCTTAGACCATAGCATAACCGGGGCAAGAGAAGCTATAGCCGGGATGTTGGACACCACCAAGGGTCTGATTAG

AACAAGTATGAAACGGGGTGGCCTCACCCCTCGAGTCTTAGCCCGCCTTTCCAATTATGATTATGAACAATTCCGATCCG

GGATAACACTATTGACAAAGAAAGGGCAGTGTTATCTCATTGACAAGGACTCGTGCTCGGTGCAGCTCGCTATAGCTCTG

AGGAGCCACATGTGGGCTAGGTTGGCTCGCGGGAGACCAATCTATGGGTTGGAGGTGCCCGATATATTAGAATCGATGAA

CGGCTACCTTATCAAACGCCACGAGTCTTGTGCTATCTGTGAGACGGGCTCAAGTCACTACGGATGGTTTTTCGTCCCTG

CAGGGTGTCAGCTTGACGATGTATCTAGAGAGACTTCGGCTCTTCGTGTACCTTATGTCGGATCAACAACTGAGGAAAGG

ACAGACATGAAACTTGCATTTGTTAGGTCTCCAAGTCGATCCCTCAAGTCAGCAGTCAGAATTGCAACAGTTTACTCATG

GGCCTACGGGGATGATGAGAAATCATGGAGTGAAGCCTGGATGCTAGCTAGGCAGAGAGCCAATATCACCTTAGATGAAT

TAAGAATGATCACTCCAGTCTCTACATCCACCAACCTAGCCCACCGGTTGAGGGATCGGAGTACCCAGGTGAAATACTCG

GGGACATCCCTTGTGAGGGTTGCAAGATACACAACCATCTCCAATGATAACTTGTCATTTGTGATATCTGAGAAAAAAGT

GGATACCAACTTCATTTATCAGCAAGGGATGCTGCTCGGTCTTGGGATCCTTGAGAACCTCTTTAGGCTAGAGGCCACCA

CCGGAGTATCCAACACAGTGCTACACCTGCACGTGGAAACAGAATGTTGTGTTGTACCTATGGTGGATCACCCAAGGATA

CCGAGTCTCCGTAATATAAAAGTGACGGATGAGCTATGCACAAACCCTTTGATCTACGACAGGTCTCCCATCATAGAGCA

CGATGCAACCCGACTATACTCACAGAGCCACAGGAGACATTTGGTGGAGTTTGTTACCTGGTCAACAAGCCAGCTTTACC

ATATACTGGCTAAGTCTACAGCAATGTCCATGATTGAGTTGATCACGAGATTCGAGAAAGATCACATGAATGAAATAGCC

GCCCTGATTGGCGATGACGACATCAACAGCTTCATCACAGAATTTTTGCTAGTAGAGCCTAGATTATTTATAGTTTACCT

TGGTCAGTGTGCTGCCATCAATTGGGCCTTTGATATACATTATCATCGCCCCTCGGGCAAGTACCAGATGGGGGAACTCC

TCTACTCTCTGCTCTCTCGGATGAGCAAAGGAGTATATAAGATCTTCACCAATGCTTTGAGCCACCCCAAAGTTTACAAG

AAATTTTGGCGAAGTGGTGTCATTGAGCCGATTCATGGCCCATCCCTGGATACACAGAATTTACATGTCACTGTCTGTGA

CATGATATATGGGTCCTATGTCACCTATTTGGATCTTTTGCTGAATGATGAGCTAGATGATTATCCGTATTTGCTCTGCG

AGAGTGATGAGGACGTGGTCACAGACAGGTTCGACAACATTCAAGCCAAACATCTCTGTGTATTGGCCGATGTATATTGC

AGCTCCAAGAGGTGTCCTTCGATAATCGGGATGTCTCCTATAGAAAAATGTACCATTCTCACACATTACATCAAGGGAGA

ATCAGTACAATCCCCGTCTGGGACCTCATGGAACACTGATCCCCTTGTAGTAGATCATTACTCATGCTCTCTGACCTACC

TTCGCCGCGGTTCCATCAAACAAATCAGGTTGAGAGTGGATCCTGGGTTTGTATTCGAGGCGTTGACAGACGTCGACTTT

AAACAACCTCGCAAAGCTAAGTTAGATATATCGGTCATTGGGTTGACTGATTTCTCTCCCCCTTGCGACAACGTCGGTGA

TTTTCTAGGGACTATCAACACATTGAGGCACAATCTGCCTGTCACCGGAACCGGGGTCTCGAACTATGAAGTCCACGCTT

ATCGTAGAATTGGCCTGAATTCATCAGCATGTTATAAAGCCGTAGAGATCTCCACGTTAATCAAGCCATCCTTAGAAGTC

GGAGAGCATGGATTGTTCTTAGGAGAAGGTTCCGGTTCAATGCTGGCTGCGTACAAGGAAGTTCTTAAATTAGCAAATTG

TTATTACAACAGCGGGGTAACAGCGGAGGGCAGAGCCGGACAGAGGGAAATATCTCCCTATCCTTCAGAGATGAGCCTAG

TAGAGAATCAAATGGGGATAGAGCGGAGTGTTAAAGTGCTGTTCAACGGCAAACCTGAAGTAACTTGGGTGGGGACCACC

GATTGCTACAAGTACATAATTAGTAACATCCAAACCTCTAGTCTGGGTTTCATACATTCAGATATTGAGACACTCCCAAC

CAAAGATGCTGTTGAGAAATTAGAAGAATTTGCCTCTATCCTATCCTTATCCTTGATTTTGGGAAAAATCGGCTCTATTA

CAGTTGTCAAAATTATGCCCATTAGCGGGGATTTTACCCAAGGCTTCATAGCCTATGCCATCCAATATTTTAGGGAGAGC

CTGCTTGCCTATCCGAGATATAGTAACTTCATCTCGACTGAGTGTTACCTTATTATGATAGGATTAAAGGCCAATCGATT

GATAAACCCAGAAGCCATTAAGCAAAGCATAATCAGAGCGGGGGTCAGGACTGCACCAGGACTTGTGAGCCATATATTAT

CAGAGAAACAGAAAGGTTGTATTCAATCTTTCTTGGGTGATCCTTATATCCAAGGGGACTTCAATAAGCACCTTAAATCT

CTAACCCCTATTGAGAAAATCCTGGTAAATTGTGGTCTCTCGATCAATGGCACAAAAATCTGTAGAGATCTACTCCACCA

TGATATCGCCTCCGGTCCAGACGGTCTTATGAGCTCTACAATTATTTTATATAGGGAATTGGCTCATTTCAAAGATAATA

TAAGGAGTCAGCACGGAATGTTCCACCCCTACCCAGTACTGGCCAACAGCAGGCAACGTGAATTAATCCTTCGAATAGCC

AAGAAATTCTGGGGGTATGTCTTGCTATATTCTGATGACCCGGCACTAATCAGACAAACAATCAAGAACTTGAAACGGAA

TCATCTAACCTTTGACTTACATAGTAACCCGTTTATTAAAGGGCTATCCAAAGCTGAGAAACTGCTGGTGCGGACGAGTT

CACTAAGAAGGGAATGGTTGTTCACTCTCGATACGAAAGAAGTGAAAGAGTGGTTCAAATTGGTAGGTTACAGTGCACTC

ATCAGAGGCTAGTTGGCTATGCATCTGCCCCCTTTTCCTCCGCCATAAGACCCCACTGATGATCCAAAAGATTAAAGAAA

ACTACATATTGGATAAGTATCTATTCCCAGCTTTGTCTGGT

>AJ849636.2/sheep/Turkey/2000

ACCAAACAAAGTTGGGTAAGGATAGATCTTATAATAACTATGGACTGGCAAACTTAGGAGTAAAGATCCTACTGTCGGGG

GGAGGAGGAGGAGCAAGATCTTTGACTATGGCGACTCTTCTTAAAAGCTTAGCATTGTTCAAGAGGAACAAAGACAAAGC

GCCGACGGCATCAGGTTCAGGAGGGGCCATCCGGGGGATTAAGAATGTTATCATAGTCCCGATTCCCGGAGACTCGTCCA

TCATTACCCGTTCAAGACTGCTCGACAGGCTTGTCAGATTGGCCGGAGATCCTGACATCAACGGGTCAAAGTTAACCGGC

GTGATGATCAGCATGTTATCTTTGTTCGTAGAGTCACCCGGGCAATTGATACAGCGAATCACAGATGATCCAGATGTTAG

TATCCGCCTTGTTGAGGTAGTTCAAAGTACTAGGTCTCAGTCCGGGTTGACCTTTGCATCACGTGGTGCTGATTTAGACA

ACGAGGCAGACATGTATTTTTCAACTGAGGGGCCCTCGAGTGGAGGTAAGAAAAGGATCAACTGGTTTGAGAACAGAGAA

ATAATAGACATAGAAGTGCAGGATCCAGAAGAGTTCAATATGTTGTTAGCCTCCATACTAGCACAAGTCTGGATCCTTCT

GGCCAAGGCTGTTACGGCACCGGATACGGCAGCTGACTCAGAATTGAGAAGGTGGGTTAAATATACACAACAAAGGAGAG

TGATTGGGGAATTTCGCCTTGACAAAGGGTGGCTGGACGCAGTCCGCAACAGAATTGCAGAAGATCTATCACTTCGGCGG

TTCATGGTATCTCTCATACTTGACATCAAGAGGACCCCCGGCAACAAGCCAAGGATTGCAGAAATGATCTGCGACATTGA

CAACTATATTGTCGAAGCCGGACTCGCCAGTTTCATCCTTACTATCAAGTTTGGTATCGAAACCATGTATCCTGCATTAG

GGCTCCACGAGTTTGCCGGGGAATTGTCCACTATTGAATCCTTGATGAACTTGTATCAACAGCTAGGAGAGGTTGCACCC

TACATGGTAATTCTCGAGAACTCAATTCAGAACAAGTTTAGTGCAGGAGCTTATCCCCTCCTCTGGAGCTATGCGATGGG

TGTCGGAGTTGGGCTGGAGAACTCAATGGGGGGCTTGAACTTTGGCAGGTCATATTTTGACCCTGCCTATTTTCGTCTCG

GACAGGAGATGGTCAGAAGATCTGCAGGAAAGGTCAGCTCTGTGATTGCGGCCGAGCTCGGCATCACAGCAGAGGAAGCC

AAACTAGTCTCGGAAATCGCCTCGCAGGCTGGGGACGAAAGAACCGCTAGAGGGACTGGGCCTCGACAGGCGCAGGTCTC

CTTCCTCCAGCACAAAACAGGAGAGGGAGAGTCGTCCGCACCAGCAACCAGAGAGGGGGTCAAAGCTGCGATCCCAAACG

GATCCGAAGAAAGGGACAGGAAGCAAACACGCTCAGGAAGGCCCAGAGGAGAGACCCCCAGCCAACTGCTCCTGGAAATC

ATGCCAGAGGATGAGGTCTCGCGAGAGTCTGGTCAAAACCCTCGTGAGGCTCAAAGATCGGCCGAGGCACTCTTCAGGCT

GCAGGCCATGGCCAAGATTCTGGAGGACCAGGAGGAGGGAGAAGACAACAGTCAGGTCTACAACGACAAGGATCTCCTCG

GCTAAGCAGACGCACCCTCTGTCGAAATCAGTGACAAGACATCTCCTACCAGTATTATAAAAAACTTAGGACCCAGGTCC

AAGCAAGCACACATCGACACTCCAATCAGTCGAGCGGAGACCACCGATGGCAGAAGAACAAGCATACCATGTCAACAAGG

GGCTGGAATGTATCAAGTCCCTCAAAGCCTCTCCCCCGGATCTATCCACCATCAGAGACACCATTGAGAGCTGGAGAGAG

GGGCTCAGCCCCTCGGGCCGTGCAACACCGAACCCTGATACGTCCGAGGGAGATCATCAGAGTATCAACCAATCATGCTC

ACCAGCAATCGGACCAAACAAAGTCTACTTGTCTCCTGGAGATAATCTCGGATTTAGAGAGATCACTGGCAACGACTGTG

AGGCTGGGCTCGGAGGAGTCCAGGGGAAAAGATCCAACTCTCAAGTACAGCGTTACTATGTTTATAGCCACGGGGGTGAA

GAGATTGAAGGACTCGAGGATGCTGACTCTCTCGTGGTTCAAGCAGATCCTCCAGTTGCTAACGTCTTCAATGGAGGAGA

GGATGGATCTGACGACAGCGATGTGGACTCTGGCCCAGATGATCCCAGCAGAGATACTCTATATGACCGGGGATCTGTTG

CCGGCAATGATGTCGCTAGGTCCACAGATGTCGAAAAACTAGAAGGTGCTGATATTCAAGAGGTTCTTAACTCCCAGAAA

GGCAAAGGAGGAAGATTCCAAGGCGGGAAGACCTTACGAGTCCCGGAAATACCCGATGTCAAGCACTCCAGACCATCAGC

CCAATCAATTAAAAAGGGCACAGACGGGAACTCAGTCTCATCTGGAACGGTGACAGAGTGTTTATCGATAAGTGGTGCAA

CCCAAGCTGTGCCAGAGTCAAGATGGGAGTCATCAGAGCAAAATGCGTCTGTGGGGAGTGTCCTCAAGTCTGCGAGGAGT

GCAGAGACGATCCAGGGGTTGACACAAGAATCTGGTACCATAGCATCACTGACTCAGCCTAAAGAGAATGACTCCGAGTA

TGAGTATGAGGATGACCTATTTAAAGAGATTCAGGACATCCGTGCAAGCATTGCCAAGATCCATGATGACAACAAAACTA

TCCTCTCAAAGCTTGATTCTATACTGTTATTGAAAGGAGAAGTCGACACTATCAAGAAACAAATCAGCAAGCAGAATATA

AGTATATCCACCATTGAGGGCCATCTCTCCAGTATAATGATAGCTATCCCGGGCTTTGGGAAGGATATCAAGGACCCAAC

ATCCGAGGTTGAGTTGAACCCAGATTTAAGACCCATAATAAGCCGTGACTCTGGCAGAGCCCTCGCGGAGGTCCTCAAGA

AACCCGCTGTGGATAGGTCTCAGAAAATTGGAACCAAAGCCAACTCCAGCTCAAAGGGTCAGCTTCTTAAGGATCTCCAG

CTAAAACCTGTCGACAAGCAGGCAAGCTCTGCAATCGGGTTTGTCCCATCCGACCATGAATCATCCAGAAATGTCATCCG

CTCCATAATCAAGTCGAGCAAGCTAAACACTGATCACAAGGACTATCTTCTAGATTTACTAAATGATGTGAAAGGCTCCA

AGGATCTTAAGGAATTCCACAAGATGCTAACAGCAATTCTCGCCAAGCACCCGTAACACACCCTCCAGTCACCATCTCAT

ACTCGGCTGAAAACATCCTCTCAATCAGGCTATTACAAAAAACTTAGGAGCAAGGGTAACTGAGCTTCGCAGACAAGATG

ACCGAGATCTACGACTTCGATAAATCAGCATGGGATGTCAAAGGGTCAATTGCTCGCATAGAACCCACCATCCACCATGA

CGGTCGACTGATACCCCAGGTAAGGGTCATCGATCCTGGTCTGGGAGACAGAAAAGATGAGTGCTTCATGTACCTGTTTC

TCCTAGGAGTGATTGAGGATAACGACCCCCTGTCTCCCCCAGTTGGAAGAACCTTTGGTTCTTTACCTCTAGGGGTCGGT

AGGTCAACTGCTAAGCCAGAAGAACTACTAAGAGAGGCCACAGAACTAGATATAGTGGTGAGGCGCACTGCAGGATTAAG

TGAGAAACTGGTATTTTACAACAACACTCCGCTGTCCTTGTTAACGCCCTGGAAGAAGGTTCTGACAACCGGAAGTGTGT

TTAGTGCTAACCAGGTTTGCAATGCAGTCAACCTAGTCCCGCTTGATACTCCCCAGAGATTCAGGGTTGTGTACATGAGC

ATAACTAGATTATCAGACAATGGTTACTATAGTGTGCCCAGAAGAATGCTGGAGTTCCGCTCAGCCAATGCAGTAGCCTT

CAACATCTTGGTTACACTAAGAATTGAAAATGGCACAAGCCCTAGAAGATACATAGTTGGCTCATGGGAGAATCCAGAGG

TCACATTTATGGTACACGTGGGCAACTTTAGAAGAAAGAAGAACGAAGTATACTCAGCTGATTACTGCAAAATGAAGATT

GAAAAGATGGGTTTAGTTTTTGCCCTGGGAGGAATAGGTGGAACAAGTCTCCATATTAGAAGCACCGGGAAAATGAGCAA

AACCCTCCATGCACAGCTGGGGTTCAAGAAAATCTTATGTTACCCCCTAATGGATGTTAATGAGGATCTTAACCGATATC

TCTGGCGGGCAGAGTGCCGAATAGTCAAAATCCAAGCCGTCCTACAGCCATCAGTACCCCAAGAATTCCGTGTCTACGAT

GATGTCATCATCAACGATGATCAAGGCCCATTCAAGATCCCGTAGTTCACCCTCAAAGGAATGAACCCACCCCCAGGAAG

GGGCCCCACCCTCCCCCG-CCAGAAAACACACTCCAAGGCTCCCCGACACAGCACCAACACAGCCACTCTCCCGATGGGA

CGAGGCCGGCC-CCCCAACAAACCCGGTCCAGGC-GGGGGGCCCCCCCCC-GACCTGCCGGGCGCCGACCGAGCAGGCCG

GCGAGAAGCCCCCAAGAGCCCCCGACGAAGACCGAGCACCCCATCCCCCCAAAAACGTCCCCCC-AACACGCCCGAGGCC

AGGCGCCGGGCACCCCCAGACCCGCCACCGGCGAGACAGAACCCCAAGGGGGGACCCCCCAAACCAAAGGGACCACCGGG

GAAGGCTCCGCCGAACCAGCACAGACCCGACCCAAGCAAGGCGGACCCCCAACACGAAACTGCCCAGCGCACCTCACTCA

TCATCAAACAAAACTTAGGGGCCAAGTCCATACGACAGGCCCCAGCCCGCCAGACGGAGGGCCGCAGAAAGGAAGGAGAC

ACGACCGCCCCAGAAGGAGGACACCCAACCACCGAAACACCCAGCCCCAGGGAAGGAG--TCCCCCCCCACACGGCGAAG

GAACGGGTGAGAGACCCCGAGGAGGACCCGACCGGGAGCCAGGAGCCGGCGACGCCGGCGGCAAACAGGCCGGGCGCACA

CCAGACTCCACCAGAGGAGCCGAGGAGAGCCTGACCCCGCGTGGACCAAACGGCAGGGAATCCAACCCCGCCAAAAAAAG

AGGTGAGCCCCC-CTT-----CCCCCCCGCACACAACCTGCAGACAGACCGAGGGAGGACCACCTCCCAACAAGACAGGC

GGCTCCCTCGCAGCCGAGCCGCATCTCCCCAGAAGAGGACACCCCCGCACACTCCGAACACCCCCTCGCCCCCCGCCACC

GGGACAGGGGCTCCGCCACCACAGACCAGAGCCACCGAGAGCCAGCCAGAGGTCCCCTCCAGTCCCGTCCAGTGCCCCCC

CGAAGAGGCCCCGGCAAGGCCCCCCAGCCCGACGACCCCTGCACGCACCGGAAGAGGGACCCGCCCGAGAACGTCCGGCA

CATCCATGTGTAAACATCATGACACGGGTCGCAATCTTGACATTTCTGTTTCTTTTCCCAAATGCTGTTGCGTGCCAGAT

TCACTGGGGCAATCTATCCAAGATCGGGATTGTAGGAACAGGGAGTGCCAGCTACAAGGTGATGACTAGGCCAAGCCACC

AAACTCTGGTTATAAAGTTAATGCCAAATATAACAGCCATCGACAATTGTACAAAGTCAGAGATTGCAGAGTACAAGAGA

TTGCTGATCACAGTGTTAAAGCCTGTAGAGGATGCTCTGTCAGTGATAACCAAGAATGTAAGACCAATTCAAACTCTGAC

ACCTGGGCGTAGAACTCGCCGCTTTGCTGGAGCTGTTCTGGCCGGAGTAGCACTTGGAGTTGCGACAGCCGCTCAGATAA

CTGCAGGAGTCGCCCTTCATCAATCATTGATGAACTCCCAAGCAATTGAGAGTTTAAAAACCAGTCTTGAGAAGTCGAAT

CAGGCAATAGAGGAAATCAGACTTGCAAATAAGGAGACCATACTAGCAGTACAGGGCGTCCAGGATTATATCAACAATGA

GCTTGTCCCTTCTGTCCATAGAATGTCATGCGAGCTGGTAGGTCACAAGCTCGGCCTCAAGCTCCTTAGGTACTACACCG

AGATCCTGTCCATATTCGGGCCCAGCCTTCGAGACCCGATAGCTGCCGAAATATCAATCCAGGCACTCAGTTATGCATTA

GGCGGGGACATTAATAGAATCCTGGACAAGCTTGGGTATAGCGGTGGGGATTTCCTTGCCATCCTAGAAAGCAAGGGAAT

AAAGGCCCGGGTAACATATGTGGATACAAGAGATTACTTTATAATTCTTAGCATCGCCTACCCAACCTTATCTGAGATCA

AGGGAGTGATAGTTCACAAGATAGAAGCTATAACATACAACATTGGGGCACAGGAGTGGTATACTACTATCCCTAAATAT

GTAGCCACTCAGGGGTATCTGATATCGAACTTTGATGAGACGTCATGCGTATTCACTCCGGATGGGACAGTTTGCAGCCA

GAATGCGTTGTACCCAATGAGCCCATTGCTCCAGGAATGTTTCCAGGGGTCAACAAAATCGTGCGCCAGAACCCTAGTTT

CAGGGACCATAAGTAATAGATTTATCCTATCAAAAGGGAACCTGATTGCAAATTGTGCGTCAGTTCTGTGCAAATGCTAC

ACAACAGAGACAGTCATTAGCCAAGATCCCGACAAACTACTAACTGTTGTAGCATCCGACAAGTGTCCTGTAGTTGAGGT

GGATGGAGTGACAATACAGGTCGGCAGTCGAGAGTATCCGGATTCTGTATACCTACACAAAATAGACTTAGGTCCAGCCA

TCTCCCTAGAAAAACTGGATGTAGGCACCAATTTAGGCAATGCAGTCACAAGACTGGAGAATGCAAAGGAGCTCCTAGAT

GCATCAGACCAAATACTGAAGACTGTTAAAGGGGTACCTTTCGGTGGGAATATGTACATAGCACTGGCAGCTTGCATTGG

GGTATCCCTAGGGCTTGTCACATTAATATGCTGCTGCAAGGGGAGGTGTAAGAACAAGGAAGTTCCCATCTCCAAAATCA

ACCCAGGGCTCAAACCCGACCTGACCGGGACCTCAAAGTCGTACGTGAGATCACTGTAGTCAGAATCACCTGAATCATTC

GGCATAACACACATACATGTACGACACAAGCAGTCAGAGGACGCAGGAAACCCAGCCTCCGATCACCCACCTGACCCCAC

CCCACACTCCACCACACATTAGTCATCAAACAAAACTTAGGACGAAAGGTCAATCACCATGTCCGCACAAAGGGAAAGGA

TCAATGCCTTCTACAAAGACAATCCTCACAATAAGAACCATAGGGTGATCCTGGATAGAGAACGCTTGGTCATCGAAAGA

CCCTACATCTTGCTTGGAGTCCTGCTGGTAATGTTCCTGAGTCTAATCGGACTGCTGGCCATTGCAGGGATCAGGCTTCA

CCGGGCCACCGTTGGAACATCAGAGATCCAGAGTCGGCTGAATACCAATATTGAGTTGACCGAATCTATTGATCACCAAA

CTAAGGATGTCTTAACTCCCCTTTTCAAAATCATTGGCGATGAAGTCGGCATCAGAATTCCACAGAAATTCAGTGATCTT

GTCAAGTTCATCTCCGATAAGATTAAATTCCTCAACCCTGATAGAGAGTATGATTTCAGGGATCTCCGGTGGTGCATGAA

TCCCCCCGAGAGAGTCAAAATTAATTTTGATCAGTTTTGTGAGTACAAGGCTGCGGTTAAGTCAATTGAACATATATTTG

AATCACCACTCAACAAGTCAAAAAAGCTGCAATCATTGACTCTTGGGCCCGGAACAGGCTGTCTAGGCAGGACAGTAACA

AGAGCTCATTTCTCAGAACTTACACTGACCTTAATGGACCTGGATCTAGAGATGAAGCACAACGTATCCTCAGTGTTTAC

CGTAGTTGAAGAGGGATTATTCGGAAGAACATATACCGTCTGGAGATCCGATGCCAGGGATCCGAGCACCGATCTAGGTA

TTGGCCACTTTTTAAGAGTCTTCGAGATTGGACTGGTAAGAGATCTCGGGCTGGGTCCCCCTGTTTTTCATATGACCAAC

TATCTCACAGTGAACATGAGTGATGACTATCGGAGATGTCTTTTAGCGGTAGGGGAGTTAAAGTTGACAGCCCTATGCAG

CTCATCTGAGACTGTGACACTGGGCGAGAGAGGAGTTCCAAAGAGGGAGCCTCTTGTAGTTGTGATACTCAACCTAGCTG

GGCCCACTCTAGGGGGCGAACTATACAGTGTCTTGCCTACCTCTGATCTCATGGTGGAGAAACTCTATTTATCTTCACAT

AGAGGGATTATCAAAGATGACGAGGCCAATTGGGTAGTGCCGTCTACCGATGTTCGTGATCTTCAAAACAAAGGAGAATG

TTTGGTGGAGGCATGCAAGACTCGACCTCCTTCATTTTGCAATGGCACAGGATCAGGCCCGTGGTCAGAGGGGAGAATCC

CTGCTTACGGGGTGATCAGGGTCAGTCTTGACTTAGCTAGTGACCCGGGTGTAGTTATCACTTCAGTGTTTGGCCCACTG

ATACCTCACCTATCTGGCATGGATCTTTACAACAACCCGTTTTCTAGAGCTGTATGGTTGGCTGTACCACCTTATGAGCA

GTCATTTCTAGGAATGATTAATACGATTGGATTCCCTAACAGAGCAGAGGTTATGCCGCACATTTTGACCACAGAGATCA

GAGGCCCTCGGGGTCGTTGCCATGTTCCCATAGAATTGTCTCGCAGGGTTGATGACGACATCAAGATTGGGTCCAACATG

GTCATATTGCCGACGATAGACCTGAGATATATTACAGCCACTTATGATGTTTCTAGGAGCGAGCATGCTATCGTGTACTA

TATCTATGACACAGGTCGCTCATCATCTTACTTCTACCCAGTTCGGCTGAACTTCAAAGGCAATCCTCTCTCTCTGAGGA

TAGAGTGTTTCCCTTGGCGTCATAAGGTGTGGTGCTACCATGATTGTCTTATATACAACACCATAACAGATGAAGAAGTC

CATACGAGAGGGCTGACCGGTATAGAGGTAACATGTAATCCAGTCTGAGCAGAGCTACGACCGCCACTCAGGCAGTCCCT

TGAGTCGCCACCGAATCCAAGCAGCACAGCCTGGGACACTCAACAGCACAGCCCAGCCAACAATGTTATAAAAAACTTAG

GAGCCAGGGCTGTAGGGGCCATGGACTCCCTATCAGTCAATCAGGTCTTGTACCCTGAGGTCCATCTAGATAGCCCTATT

GTCACAAACAAACTAGTTGCCATCCTTGAGTACTCGGGTATCGACCATAACTATGTTCTTGAAGACCAGACCCTTGTCAA

GAATATTAGGTATAGACTGGGGTGCGGTTTTTCAAATCAAATGATCATCAATAACAGGGGGGTGGGTGAAACAGTCAATT

CTAAACTCAAAAGTTACCCCCATAATCGTCATATCATATACCCGGATTGCAATAAGGAGTTGTTTTGTATCAAAGATAGC

TGCATATCTAAGAAGCTCTCGGAGCTATTCAAGAAGGGTAATTCCTTGTACTCTAAGATAAGTCACCAGGTACTGGATTG

TCTTAAGAGAGTCAATGGGAAATTAGGCCTGGGCACAGATCTTACCTATGGTCTGAAGGAGGGTATCCTTGACCTAGGGT

TGCACATGCATAGCTCTCAATGGTTCGAGACTTTCCTGTTCTGGTTCACTATCAAGACAGAGATGAGATCAATGATCAAA

GAACAGTCCCATATATGCCACAAGAGGAGGTATAACCCGATTTTTGTGTCGGGGGATGCATTCGAGGTGCTTGTATCACG

AGACCTCGTTGTGATAATTGATAAGAATACCCAGTATGTCTTCTACTTAACATTTGAGCTGGTCCTTATGTATTGTGATG

TCATAGAGGGCAGACTTATGACTGAGACAGCCATGGCCATAGATCAGAGATATTCAGAACTTCTAAGCCGAGTCAGATAC

TTGTGGGATCTTATTGATGGGTTCTTCCCAACACTAGGCAACACCACATACCAAGTTGTTGCCCTGCTTGAACCATTATC

GTTGGCTTATCTTCAACTTCAGGATGTCACTCTGGAGTTAAGAGGTGCTTTTTTAGACCATTGCTTCAAAGAACTCTATG

AGATATTGGAGCATTGCGGCATTGACACAGAAGGCACCTACAATTCCATCACTGAAGGATTGGATTACGTATTCATCACC

CACGATATACATCTAACTGGGGAGATTTTTTCATTCTTTCGGAGTTTCGGACACCCCCGTCTAGAAGCAGTCACCGCTGC

AGAAAATGTCAGAAAACATATGAACCAACCGAAGGTAATCAGTTATGAGACTATGATGAAAGGGCACGCAGTATTTTGCG

GGATAATCATAAATGGTTTTAGGGACCGACATGGCGGCAGCTGGCCCCCTGTTGCATTGCCAGAACATGCTTCTGCTGCG

ATCCGGAATGCGCAGGCATCAGGTGAGGGACTGACCCATGACCTGTGTATAGACAATTGGAAGTCCTTTGTTGGATTCAA

ATTTGGCTGCTTTATGCCACTCAGCCTAGATAGTGATTTGACCATGTACCTCAAAGACAAGGCATTGGCTGCACTGAAGA

ATGAGTGGGATTCAGTTTACCCGAAGGAATACCTCCGTTATAACCCACCTAGGGGGACAGAGTCAAGGCGATTGGTGGAG

GTGTTCCTGAATGACTCCAGCTTTGATCCTTATAACATGATAATGTACGTGGTGAATGGCTCCTACCTTAAAGACCCTGA

GTTTAATCTCTCATACAGCCTAAAAGAGAAGGAGATAAAGGAGACAGGGCGGTTGTTTGCCAAAATGACCTATAAGATGC

GGGCCTGTCAGGTAATCGCTGAAAATCTGATATCGAATGGTGTTGGGAAGTATTTCCGAGACAATGGGATGGCAAAGGAC

GAGCATGACCTAACAAAAGCCCTTCACACTCTGGCAGTCTCAGGTGTTCCCAAGAACAACAAAGATAACCACCGAGGTGG

CCCTCCCAGAAGAACAACAAACCGAGAGGTGAGATCAAGCCAAGACACCAAAACACAAAATAGAGACAAGGTCCAAGGGG

GGCCTGTGTACAACTATTTGCGATGCCAACCAATCAGCCCTGATCAGGGCGAGTCATACGAGACTGTTAGTGCATTCATC

ACCGCTGACCTTAAGAAGTATTGCCTGAATTGGAGATACGAGACAATCAGCATATTTGCACAGAGGCTGAATGAAATATA

TGGACTACCATCCTTCTTTCAATGGTTACACAGGATTTTGGAAAAATCCGTACTCTACGTCAGTGACCCACATTGCCCTC

CCGACCTAGATAATCATATCCCTCTGGACAGTGTCCCTAATGCCCAAATATTCATTAAGTACCCCATGGGCGGAATAGAA

GGTTATTGCCAAAAACTATGGACAATCAGTACTATACCATACTTGTATCTGGCAGCCTATGAGAGCGGAGTAAGAATCGC

CTCACTAGTGCAGGGTGACAATCAGACAATCGCAGTGACAAAAAGAGTTCCAAGTTCTTGGCCTTATTCACTAAAAAAGA

GGGAGGCATCCAAAGCCGCTCAAAATTACTTCGTAGTCTTAAGGCAAAGATTGCACGATGTAGGTCATCACTTAAAGGCT

AATGAAACCATAGTATCTTCTCACTTTTTTGTATATTCCAAGGGGATTTATTATGACGGCCTGCTAGTCTCACAATCACT

AAAGAGCATCGCCAGATGTGTCTTCTGGTCCGAGACTATAGTGGATGAAACCAGAGCGGCTTGCAGCAATATTGCAACAA

CTGTCGCCAAGAGTATAGAGAGGGGTTATGATAGATACCTTGCATATTCTTTGAATATCCTCAAGATTTTCCAACAGATC

CTTATATCCCTTAACTTCACTATTAACACAACAATGACTCAGGATGTCGTGGCACCGATCATCGAAAACGGGGATTTGCT

GATAAGGATGGCACTCTTGCCAGCACCCATCGGGGGTCTCAATTATCTTAACATGAGCAGGTTATTTGTGAGAAATATCG

GTGACCCGGTCACCTCCTCTATAGCCGACCTGAAGAGGATGATAGATGCTGGGCTAATGCCAGAAGAAACATTGCATCAA

GTGATGACCCAGACCCCGGGAGAATCATCCTACCTCGATTGGGCAAGTGACCCTTATTCTGCCAACCTACCCTGCGTACA

GAGTATAACTCGCCTTCTCAAAAACATCACTGCACGATATATTTTAATCAGCAGCCCAAACCCGATGCTGAAAGGATTGT

TTCATGAGGGGAGTAGAGATGAAGACGAGGAGCTTGCAAGTTTCTTAATGGATCGGCATATAATTGTTCCAAGAGCTGCA

CATGAAATCTTAGACCATAGCATAACCGGGGCAAGAGAAGCTATAGCCGGGATGTTGGACACCACCAAGGGTCTGATTAG

AACAAGTATGAAACGGGGTGGCCTCACCCCTCGAGTCTTAGCCCGCCTTTCCAATTATGATTATGAACAATTCCGATCCG

GGATAACACTATTGACAAAGAAAGGGCAGTGTTATCTCATTGACAAGGACTCGTGCTCGGTGCAGCTCGCTATAGCTCTG

AGGAGCCACATGTGGGCTAGGTTGGCTCGCGGGAGACCAATCTATGGGTTGGAGGTGCCCGATATATTGGAATCGATGAA

CGGCTACCTTATCAAACGCCACGAGTCTTGTGCTATCTGTGAGACGGGCTCAAGTCACTACGGATGGTTTTTTGTCCCTG

CAGGGTGCCAGCTTGACGATGTATCTAGAGAGACTTCGGCTCTTCGTGTACCTTATGTCGGATCAACAACTGAGGAAAGG

ACAGACATGAAACTTGCTTTTGTTAGGTCTCCAAGTCGATCCCTCAAGTCAGCAGTCAGAATTGCAACAGTTTACTCATG

GGCCTACGGGGATGATGAGAAATCATGGGGTGAAGCCTGGATGCTAGCTAGGCAGAGAGCCAACATCACCTTAGATGAAT

TGAGAATGATCACTCCAGTCTCTACATCCACCAACCTAGCCCACCGGTTGAGGGATCGGAGCACCCAGGTGAAATACTCG

GGGACATCCCTTGTGAGGGTTGCAAGATACACAACCATCTCCAATGATAACCTGTCATTTGTGATATCTGAGAAAAAAGT

GGATACCAACTTCATTTATCAGCAAGGGATGCTGCTCGGTCTTGGGATCCTTGAGAACCTCTTTAGGCTAGAGGCCACCA

CCGGAGTATCCAACACAGTGCTACACCTGCACGTGGAAACAGAATGTTGTGTTGTACCTATGGTGGATCACCCAAGGATA

CCGAGTCTCCGTAATATAAAAGTTACGGATGAGCTATGCACAAACCCTTTGATCTACGACAGGTCTCCCATCATAGAGCA

CGATGCAACCCGACTATACTCACAGAGCCACAGGAGACATCTGGTGGAGTTTGTTACCTGGTCAACAAGCCAGCTTTACC

ATATACTGGCTAAGTCTACAGCAATGTCCATGATTGAGTTGATCACGAGATTCGAGAAAGATCACATGAATGAAATAGCC

GCCCTGATTGGCGATGACGACATCAACAGCTTCATCACAGAATTTTTGCTAGTAGAGCCTAGATTATTTATAGTTTACCT

TGGTCAGTGTGCTGCCATCAATTGGGCCTTTGATATACATTATCATCGGCCCTCGGGCAAGCACCAGATGGGGGAACTCC

TCTACTCTCTGCTCTCTCGGATGAGCAAAGGAGTATATAAGATCTTCACCAATGCTTTGAGCCACCCCAAAGTTTACAAG

AAATTCTGGCGAAGTGGTGTCATTGAGCCGATTCATGGCCCATCCCTGGATACACAGAATTTACATGTCACTGTCTGTGA

CATGATATATGGGTCCTATGTCACCTATTTGGATCTTTTGCTGAATGATGAGCTAGATGATTACCCGTATTTACTCTGCG

AGAGTGATGAGGACGTAGTCACAGACAGGTTCGACAACATTCAAGCCAAACATCTCTGTGTATTGGCCGATGTATATTGC

AGCTCCAAGAGGTGTCCTTCGATAATCGGGATGTCTCCTATAGAAAAATGTACCATTCTCACACATTACATCAAGGGAGA

ATCAGTACAATCCCCGTCAGGGACCTCATGGAACACTGATCCCCTTGTAGTAGATCATTACTCATGCTCTCTGACCTACC

TTCGCTGCGGTTCCATCAAACAAATCAGGTTGAGAGTGGATCCTGGGTTTGTATTCGAGGCGTTGACAGACGTCGACTTT

AAACAGCCTCGCAAGGCTAAGTTAGATATATCGGTTGTTGGGTTGACTGATTTCTCTCCCCCTTGCGACAACGTTGGTGA

TTTTCTAGGGACTATCAACACATTGAGGCACAATCTGCCTGTCACCGGAACCGGGGTCTCGAACTATGAAGTCCACGCTT

ATCGTAGAATTGGCCTGAATTCATCAGCATGTTATAAAGCCGTAGAGATCTCCACGTTAATCAGGCCATCCTTAGAAGTC

GGAGAGCATGGATTGTTCCTAGGAGAAGGTTCCGGTTCAATGCTGGCTGCGTACAAGGAAGTTCTTAAATTAGCAAATTG

TTATTACAACAGCGGGGTAACAGCGGAGGGCAGAGCCGGACAGAGGGAAATATCTCCCTATCCTTCAGAGATGAGCCTAG

TAGAGAATCAAATGGGGATAGAGCGGAGTGTTAAAGTGCTGTTCAACGGCAAACCTGAGGTAACTTGGGTGGGGACCACC

GATTGCTACAAGTACATAATCAGTAACATCCAAACCTCTAGTCTGGGTTTCATACATTCAGATATTGAGACACTCCCAAC

CAAAGACGCTGTTGAGAAATTGGAAGAATTTGCCTCTATCCTATCCTTATCCTTGATTTTGGGAAAAATCGGCTCTATTA

CAGTTGTCAAAGTTATGCCCATTAGCGGGGATTTTACCCAAGGTTTCATAGCCTATGCTATCCAATATTTTAGGGAGAGC

CTGCTTGCCTATCCGAGATATAGTAACTTCATCTCGACTGAGTGTTACCTTATTATGATAGGATTAAAGGCCAATCGATT

GATAAACCCAGAAGCCATTAAGCAAAGCATAATCAGAGCGGGGGTCAGGACTGCACCAGGACTTGTGAGCCATATATTAT

CAGGGAAACAGAAAGGTTGTATTCAATCTTTCTTGGGTGATCCTTATATCCAAGGAGACTTCAATAAGCACCTTAAATCT

CTAACCCCTATTGAGAAAATCCTGGTAAATTGTGGTCTCTCGATCAATGGCACAAAAATCTGTAGAGATCTAATCCACCA

TGATATCGCCTCCGGTCCAGACGGTCTTATGAGCTCTACAATTATTTTATATAGGGAGTTGGCTCATTTCAAAGATAATA

TAAGGAGTCAGCACGGAATGTTCCACCCCTACCCAGTACTGGCCAACAGCAGGCAGCGTGAATTAATTCTTCGAATAGCC

AAGAAATTCTGGGGGTATGTCTTGCTATATTCTGATGACCCGGCACTAATCAGACAAACAATCAAGAACTTGAAACGGAA

TCATCTAACCTTTGACTTACATAGTAACCCGTTTATTAAAGGGCTATCCAAAGCTGAGAAACTGCTGGTGCGGACGAGTT

CACTAAGAAGGGAATGGTTGTTCACCCTCGATACGAAAGAAGTGAAAGAGTGGTTCAAATTGGTAGGTTACAGTGCACTC

ATCAGAGGCTAGTTAGCTATACATCTGCCCCCTTTTCCTCCGCCATAAGACCCCACTGATGATCCAAAAGATTAAAGAAA

ACTACATATTGGATAAGTATCTATTCCCAGCTTTGTCTGGT

>KX033350.1/goat/India/2016-01-27

ACCAAACAAAGTTGGGTAAGGATAGATCTTATAATAACTATGGACTAGCAAACTTAGGAGTAAAGATCCTACTGTCGGGG

GGAGGAGGAGGAGCAAGACCTCTGATTATGGCGACTCTCCTTAAAAGCTTAGCATTGTTCAAAAGGAACAAAGACAAAGC

GCCGACGGCATCAGGTTCAGGAGGGGCCATCCGAGGGATTAAGAATGTTATCATAGTCCCGATTCCCGGAGACTCGTCCA

TCACTACCCGTTCAAGACTGCTCGACAGGCTTGTCAGATTGGCCGGAGATCCTGACATCAACGGGTCAAAGCTGACCGGC

GTGATGATCAGCATGTTATCTTTGTTCGTAGAGTCACCCGGGCAGTTGATACAGCGAATCACAGATGATCCGGATGTTAG

TATCCGCCTTGTTGAGGTAGTTCAAAGTACTAGGTCCCAGTCCGGGTTGACCTTTGCATCACGTGGTGCTGATTTGGATA

ACGAGGCAGACATGTATTTTTCAACTGAGGGGCCCTCGAGTGGAGGTAAGAAAAGGATCAACTGGTTTGAGAACAGAGAA

ATAATAGACATAGAGGTGCAGGATCCAGAAGAGTTCAATATGTTGTTAGCCTCCATACTAGCACAAGTCTGGATCCTTCT

GGCCAAGGCTGTCACGGCACCGGATACGGCAGCTGACTCAGAACTGAGAAGGTGGGTTAAATATACACAACAAAGGAGAG

TGATTGGGGAATTTCGCCTTGACAAAGGGTGGCTGGACGCAGTCCGCAACAGAATTGCAGAAGATCTATCACTCCGGCGG

TTCATGGTATCTCTCATACTTGACATCAAGAGGACCCCCGGCAACAAGCCAAGGATCGCAGAAATGATCTGCGACATTGA

CAACTATATTGTCGAAGCAGGACTCGCCAGTTTCATCCTTACTATCAAATTTGGTATAGAAACCATGTATCCTGCATTAG

GGCTCCACGAGTTTGCCGGGGAATTGTCCACTATTGAGTCCTTGATGAACTTGTATCAACAGCTAGGAGAGGTTGCACCC

TACATGGTAATTCTAGAGAACTCAATTCAGAACAAGTTTAGTGCAGGAGCTTATCCCCTCCTCTGGAGCTATGCGATGGG

TGTCGGAGTTGAGCTGGAGAACTCAATGGGGGGCTTGAACTTTGGCAGATCATATTTTGACCCTGCCTATTTTCGTCTCG

GACAGGAGATGGTCAGAAGATCTGCAGGAAAGGTCAGCTCTGTGATTGCGGCTGAGCTCGGCATCACAGCAGAGGAAGCT

AAACTAGTCTCGGAAATAGCCTCGCAGGCTGGGGACGAAAGAACCGCTAGAGGGACTGGGCCTCGACAGGCGCAGGTCTC

CTTCCTACAGCACCAAACAGGAGGGGGAGAGTCGTCCGCACCAGCGACCAGAGAAGGGGTCAAAGCTGCGATCCCAAACG

GATCCGAAGAAAGGGACAGAAAGCAAACACGCCCAGGGAGGCCCAGAGGAGAGACCCCCGGCCAACTGCTCCTGGAATTC

ATGCCAGAGGATGAGGTCTCGCGAGAATCTGGTCAAAACCCTCGTGAGGCTCAAAGGTCGGCCGAGGCACTCTTCAGGCT

GCAGGCCATGGCCAAGATTCTGGAGGACCAGGAGGAGGGAGAAGACGACAATCAGGTCTACAACGACAAGGATCTCCTCG

GCTAAGCAGACGCACCCTCTGTCGAAATCAGTGACAAGACATCTCCTACCAGTATTATAAAAAACTTAGGACCCAGGTCC

AAGCAAGCACACATCGACACTCCAACCAGTCGAGCGGAGGCCACCGATGGCAGAAGAACAAGCATACCATGTCAACAAGG

GGCTGGAATGTATCAAGTCCCTCAAAGCCTCTCCCCCGGATCTATCCACCATTAGAGATACCATCGCGAGCTGGAGAGAG

GGGCTTAGCCCCTCGGGCCGTGCAACACCGAACCCTGATACGTCCGAGGGAGATCATCAGAATATCAACCAATCATGCTC

ACCAGCAATCGGACCAAACAAAGTCTACTTGTCTCCTGAAGATAATCTCGGATTTAGAGAGATCACTGGCAACGACTGTG

GGGCTGAGCTCGGAGGAGTCCAGGGAAAAGGATCCAACTCTCAAGTACAGCGTTACTATGTTTATAGCCACGGGGGTGAA

GAGATTGAAGGACTCGAGGATGCTGACTCTCTCGTGGTTCAAGCAGATCCTCCAGTTGCTAACATATTCAATGGAGGAGA

GGATGGATCTGACGACAGCGATGTGGACTCTGGCCCAGATGATCCCAGCAGAGATACTCTATATGACCGGGGACCTGCTG

CCGGCAATGATGTCGCTAGGTCCACAGATGTTGAAAAACTAGAAGGTGCTGATATTCAAGAAGTCCTCAACTCCCAGAAA

AGCAGAGGAGGAAGATTCCAAGGCGGGAAAACCCTACGAGTCCCGGAAGCACCCGATGTCAAGAACTCCAGACCATCAGC

CCAATCAATTAAAAAGGGCACAGACGGGAACTCAGTCTCATCTGGAACGGTGACAGAGTGTTTATCGATAAGTGGTGCAA

CCCAAGCTGTGCCAGAGTCAAGATGGGAGTCATCAGAGCAAAATGCGTCTGTGGGGAGTGTCCTCAAGTCTGCGAAGAGT

GCAAAGACGATCCAGGGGTCGACACAAGAATCTGGTACCACAGCATCACCGACTCAGCCTAAAGAGAATGACTCCGAGTA

TGAGTATGAGGATGACCTATTTACAGAGATTCAGGACATCCGTGCAAGCATTGCCAAGATCCATGATGACAACAAGACTA

TCCTCTCAAAGCTTGATTCTATACTGTTATTGAAAGGAGAAGTCGACACTATCAAGAAACAAATCAGCAAGCAGAATATA

AGTATATCCACCATTGAGGGCCATCTCTCCAGTATAATGATAGCCATCCCGGGCTTTGGGAAGGATATCAAGGACCCAAC

ATCCGAGGTTGAGTTGAACCCAGATTTAAGGCCTATAATAAGCCGTGACTCTGGCAGAGCTCTCGCTGAGGTCCTCAAGA

AACCCGCTGTGGATAGGTCTCAGAAAACTGGAACCAAAGCCAACTCCAGCTCAAAGGGTCAGCTTCTTAAGGATCTCCAG

CTAAAACCTGTCGACAAGCAGGCAAGCTCTGCAATCGGGTTTGTCCCATCCGACCATGAATCATCCAGAAATGTCATCCG

CTCCATAATCAAGTCGAGCAAGCTAAACATTGATCACAAGGACTATCTTCTAGATTTACTGAATGATGTGAAAGGCTCAA

AGGATCTTAAGGAATTCCACAAGATGCTAACAGCAATTCTCGCCAAGCACCCGTAACACATCCCCCAGTCACCATCTCAT

ACTCGGCTGAAGACATCCTTTCAATCAGACTATTACAAAAAACTTAGGAGCAAGGGCAACTGAGCTTCGCAGACAAGATG

ACCGAGATCTACGACTTCGACAAATCAGCATGGGATGTCAAAGGGTCAATTGCCCGCATAGAACCCACCACCTACCACGA

CGGCCGACTGATACCCCAGGTAAGGGTCATCGATCCTGGTCTGGGAGACAGAAAAGATGAGTGCTTTATGTATCTGTTTC

TCCTGGGGGTGATTGAGGACAACGACCCCCTGTCTCCCCCCGTCGGAAGAACCTTTGGCTCTTTACCTCTAGGGGTCGGT

AGGTCAACTGCTAAGCCAGAAGAACTACTAAGGGAGGCCACAGAACTAGACATAGTGGTGAGGCGCACTGCAGGAGTAAA

TGAGAAACTGGTATTTTACAACAACACTCCGCTGTCCTTGTTAACGCCCTGGAAGAAAGTTCTGACAACCGGAAGTGTGT

TTAGCGCTAACCAGGTTTGCAATGCAGTCAACCTAGTCCCGCTTGATACTCCCCAGAGGTTCAGGGTTGTGTACATGAGC

ATAACTAGATTATCAGACAATGGTTACTATAGTGTGCCCAGGAGAATGCTAGAGTTCCGCTCAGCCAATGCAGTAGCCTT

CAACATCTTGGTTACACTAAGAATTGAGAATGGCACAAACCCTAGAAGATACATAGTCGGCTCATGGGAGAATTCAGAGG

TCACATTTATGGTACACGTGGGCAACTTTAGAAGAAAGAAGAACGAAGTATACTCTGCTGATTACTGCAAAATGAAGATT

GAAAAGATGGGTTTAGTTTTTGCCCTGGGAGGAATAGGTGGAACAAGTCTCCATATTAGAAGCACCGGGAAAATGAGCAA

AACCCTCCATGCACAGCTGGGGTTCAAGAAAATCTTATGTTACCCCCTAATGGATGTTAATGAGGATCTTAACCGATATC

TCTGGCGGGCAGAGTGCCGAATAGTCAAAATCCAAGCCGTCTTACAGCCATCAGTGCCCCAAGAATTCCGTGTCTACGAT

GATGTCATCATCAACGATGACCAAGGCTTGTTCAAGATCCTGTAGTTCATCTGCAAAGGAACGAACCCACCCCCAGGGGG

GGGCCCCCCCCTCCCCCG-CCAAAAAACACACTCCAAGGCTCCCCGACACAGCGCCAACACAGCCACCCTCCCCACTGGA

CGAGGCCGGCC-CCCCAACAAACCCGGTCCAGGC-GGGGGGCCCCCCCCCTGACCCGCCGGGCGCCGACCGAGCAGGCCG

GCGAGAAGCCCCCAAGAGGCCCCGACGAAGACCGAGCACCCCTCCCCCCC-AAACCGTTCCCCC-AACACGCCCGAGGCC

AGGCGCCGGGCACCCCCAGACCCGCCACCGGCGGGACAGAACCCCAGGGGGGAACCCCCCAAACCAAAGGGACCACCGGG

GAAGGATCCGCCGAACCAGCACAGACCCGACCCAAGCAAGGCGGACCCCCAACATGAAACCGCCCAGCACACCCCACTCA

TCATCAAACAAAACTTAGGGGCCAAGCCCATACGACAGGCCCCAGTGCGCCAGGCGGAGGGACGCAGAAAGGAAGGAGAC

AAGACCGCCCCAGAAGGAGGACACCCAACCACCGAAACACCCAGCCCCAGGGAAGGAG--CCCCCCCCCACATGGCGAAA

GAACGGGCAAGAGACCCCGAGAAGAACCCGACCGGGAGCCAGGAGCCGGCGACGCCGGCGGCAAACAAGCCGGGCAAACA

CCCGACCCCACCAGAGGAGCCGAGGAGAGCCCGACCCCGCACGAACCAAACAGCAGGGAGCCCAACCCCACCAAAAAAAG

AGGCGAGGCCCC-CCT-----CCTCCCCGCACACAACCCGCAGACAGACCGAGAGAGGACCACCTCCCAACAAGACAGGC

GGCTCCCTCGCAGCCGAGCCGCAGCTCCCCAGAACAGGACACCCCCGTACACTCAAAACACCCCCCCGCCCCCCGCCACC

GGGACAGGGGCTCCGCCACCACAGACCAAAGCCACCGAGAGCAAGCCAGAGGTCCCCCCCAGTCCCGCCCGGCGCCCCCC

CGAAAAGGCCCCGGCAAGGCCCCCCAACCCGACGTCCCCCGCACGCACCGAAAGAGGGACCCGCCCGAGAACGTGCGGCA

CATCCATGCGTAAACATTATGACACGGGTTGCAATCTTGACATCTCTGTTTCTTCTCCCAAATGTTGTTGCGTGCCAGAT

TCACTGGGGCAATCTATCCAAGATCGGGATTGTGGGAACAGGGAGTGCCAGCTACAAGGTGATGACTAGGCCGAGCCACC

AGACTCTGGTTATAAAGTTAATGCCAAATATAACAGCCATCGACAATTGTACAAAGTCAGAGATTGCAGAGTACAAGAGA

TTGCTGATCACAGTGTTAAAGCCTGTAGAGGATGCTCTGTCGGTTATAACCAAGAATGTAAGACCAATTCAAACCCTGAC

ACCTGGGCGTAGAACCCGTCGTTTTGCTGGAGCTGTTCTGGCCGGAGTAGCGCTTGGAGTTGCGACAGCCGCTCAGATAA

CTGCAGGAGTCGCTCTTCATCAATCATTGATGAACTCCCAAGCAATTGAGAGTTTAAAAACCAGTCTTGAGAAGTCGAAT

CAGGCAATAGAAGAAATCAGACTTGCAAATAAGGAGACCATACTGGCAGTACAGGGCGTCCAGGATTATATCAACAATGA

GCTTGTCCCTTCTGTTCATAGAATGTCATGCGAGCTGGTAGGCCACAAGCTCGGCCTCAAGCTCCTTAGGTACTACACCG

AGATCCTGTCCATATTCGGGCCCAGCCTTCGAGACCCGATAGCTGCCGAAATATCAATCCAGGCACTCAGTTATGCATTA

GGCGGAGACATTAATAAAATCCTGGACAAGCTTGGGTATAGCGGTGGGGATTTCCTTGCCATCCTAGAAAGCAAGGGAAT

AAAGGCCCGGGTTACATATGTGGACACAAGAGATTACTTTATAATGCTTAGCATCGCCTACCCAACCTTATCTGAGATCA

AGGGAGTGATAGTTCACAAGATAGAAGCTATAACATACAACATTGGGGCACAGGAGTGGTATACTACTATCCCTAAATAT

GTAGCCACTCAGGGATATCTGATATCGAACTTTGATGAGACGTCATGCGTATTCACTCCAGAGGGGACAGTTTGCAGCCA

GAATGCATTGTACCCAATGAGCCCATTGCTTCAGGAATGTTTCAGGGGGTCAACAAAATCGTGCGCCAGAACCCTAGTTT

CAGGTACCATAAGTAATAGATTTATCCTATCAAAAGGAAACCTGATCGCGAATTGTGCGTCAGTTTTGTGCAAATGTTAC

ACAACGGAGACAGTCATCAGCCAAGATCCTGACAAACTACTAACTGTTGTAGCATCCGACAAGTGTCCTGTAGTTGAGGT

GGATGGAGTGACAATACAGGTCGGCAGTCGAGAGTATCCGGATTCTGTATACTTACACAAAATAGACTTAGGCCCAGCCA

TCTCCCTAGAAAAACTAGATGTAGGTACCAATTTAGGCAATGCAGTTACAAGACTGGAGAATGCAAAGGAGCTCCTAGAT

GCATCAGACCAAATACTGAAGACTGTCAAAGGGGCACCTTTTGGTGGGAACATGTACATAGCACTGGCAGCTTGCATTGG

AGTATCCCTAGGGCTTGTCACATTAATATGCTGCTGTAAGGGGAGGTGTAAGAACAAGGAAATCCCTATCTCCAAAATCA

ACCCAGGGCTCAAACCCGATCTGACCGGGACCTCAAAGTCGTACGTGAGATCACTGTAGCCAGAATCACCTGAATCATCT

GGCATCACACACATACATGTACGACACAAGCAGTCTGAGGACGCAGGAAACCCAGCCCCCGATCACCCACCTGACCCCAC

TCCACGCTCCACTACACATTGGTCATTAAACAAAACTTAGGACGAAAGGTCAATCACCATGTCCGCACAAAGGGAAAGGA

TCAATGCCTTCTACAAAGACAATCCTCACAATAAGAACCATAGGGTAATCCTGGATAGAGAACGCTTGATCATTGAAAGA

CCCTACATCTTGCTTGGAGCCCTGCTGGTAATGTTCCTGAGTCTAATCGGACTGCTGGCCATTGCAGGGATCAGGCTTCA

CCGGGCCACCGTTGGAACTTCAGAGATCCAGAGTCGGCTGAATACCAATATTAAGTTGACCGAATCTATTGATCACCAGA

CTAAGGATGTCTTAACTCCCCTTTTTAAAATAATTGGCGATGAAGTCGGCATCAGGATTCCACAGAAATTCAGTGATCTT

GTCAAGTTCATCTCCGACAAGATTAAATTCCTCAACCCTGATAGAGAGTATGATTTCAGGGATCTCCGGTGGTGCATGAG

TCCCCCCGAGAGAGTCAAAATTAATTTTGATCAGTTTTGTGAGTACAAAGCTGCGGTTAATTCAATTGAACATATATTTG

AGTCACCGCTCAACAAGTCAAAAAAGCTGCAATCATTGACTCTCGGGCCCGGAACAAGTTGTCTAGGCAGGACAGTAACA

AGAGCTCATTTCTCAGAACTTACACTGACCTTAATGGACTTGGATCTAGAGATGAAGCACAACGTGTCCTCAGTGTTTAC

CGTAGTTGAAGAGGGGTTATTCGGAAGAACATATACCGTCTGGAGATCCGATGCCAGGGATCCGAGCACCGATCTAGGTA

TCGGCCATTTTTTAAGAGTCTTCGAGATTGGACTGGTAAGAGATCTCGGGCTGGGTCCCCCCGTTTTTCATATGACTAAC

TATCTCACAGTGAACATGAGTGATGTCTATCGGAGATGTCTTTTAGCGGTAGGGGAGTTGAAGTTGACAGCCCTATGCAC

CTCATCTGAGACTGTAACACTGAGTGAGAGAGGAGTTCCAAAGAGGGAGCCTCTTGTAGTTGTGATACTTAACCTGGCTG

GACCAACTCTAGGGGGCGAACTATACAGTGTCTTGCCTACCTCTGATCTCATGGTGGAGAAACTCTATTTATCTTCACAT

AGAGGGATCATCAAGGATGACGAGGCCAATTGGGTAGTGCCGTCTACCGATGTTCGTGATCTTCAGAACAAAGGTGAGTG

TCTAGTGGAAGCATGCAAGACTCGACCTCCTTCATTTTGCAATGGCACAGGATCAGGCCCGTGGTCAGAGGGGAGAATCC

CTGCTTACGGGGTGATCAGGGTCAGTCTTGACTTAGCTAGTGACCCGGATGTAGTTATCACTTCAGTGTTTGGCCCACTG

ATTCCTCACCTATCCGGCATGGATCTTTACAACAACCCGTTTTCAAGAGCTGTATGGTTGGCTGTACCACCTTACGAGCA

GTCATTTCTAGGAATGATAAATACAATTGGATTCCCTAACAGAACAGAGGTTATGCCGCACATTTTGACCACAGAGATCA

GAGGCCCTCGGGGTCATTGCCATGTCCCCATAGAATTGTCCCGCAGGGTTGATGACGATATCAAGATCGGGTCCAACATG

GTCATATTGCCGACGATGGACCTGAGATATATTACAGCCACTTATGATGTTTCCAGGAGCGAGCATGCAATCGTGTACTA

TATCTATGACACAGGTCGCTCATCATCTTACTTCTACCCAGTTCGACTGAATTTCAAAGGCAATCCTCTCTCTCTGAGGA

TAGAGTGTTTCCCTTGGCGTCATAAGGTGTGGTGCTACCATGATTGTCTTATATACAACACCATAACAGATGAAGAGGTC

CATATGAGAGGGCTGACCGGTATAGAGGTAACATGCAATCCAGTCTGAGCAGAGCTACGACCGTCACTCAAGCAGTCCCG

TGAGTCGCCACCGAGTCCAAGCAGCACAGCCTGGGACTCTCAACAGCACAGCCCAGCCAACAGTGTTATAAAAAACTTAG

GAGCCAGGGTTGTAGGGGCCATGGACTCCCTATCAGTCAATCAGGTCTTGTACCCTGAGGTTCATCTAGACAGCCCTATT

GTCACAAACAAACTAGTTGCCATCCTTGAGTACTCGGGTATCGACCATAACTATGTTCTTGAAGACCAGACCCTTGTCAA

GAATATTAGGTATAGACTGGGGTGCGGTTTTTCAAATCAAATGATCATCAATAACAGGGGGGTGGGTGAAACAGTCAATT

CTAAACTTAAAAGTTATCCCCATAATCGTCATATCATATACCCGGATTGCAATAAGGAGTTGTTTTGTATCAAAGATAGC

TGCATATCTAAGAAGCTCTCGGAGCTATTCAAGAAGGGTAATTCCTTGTACTCTAAGATAAGTCACCAGGTACTGGACTG

TCTTAAGAGAGTCAATGGGAAATTAGGCCTGGGCACAGATCTTACCCATGGTCTGAAGGAGGGTATCCTTGACCTAGGGT

TGCACATGCATAGCTCTCAATGGTTCGAGACTTTCCTGTTCTGGTTCACTATCAAGACAGAGATGAGATCAATGATCAAA

GAACAGTCCCATATATGCCACAAGAGGAGATATAACCCGATTTTTGTATCGGGGGATGCATTCGAGGTGCTCGTATCACG

AGACCTCGTTGTGATAATTGATAAGAATACCCAGTATGTCTTCTACTTAACATTTGAGCTGGTCCTTTTGTATTGTGATG

TCATAGAGGGCAGACTTATGACGGAGACAGCCATGGCCATAGACCAGAGATATTCAGAACTTCTAAGCCGGGTTAGATAC

CTGTGGGATCTTATTGATGGGTTCTTCCCAACACTAGGCAATACCACCTACCAAATTGTTGCTCTGCTTGAACCATTATC

GTTGGCTTATCTTCAACTTCAGGATGTCACGCTGGAGTTAAGAGGTGCTTTTTTAGACCATTGCTTCAAAGAACTCTATG

AGATACTGGAGCATTGCGGCATTGACACAGAAGGCACCTACAATTCCATCACTGAAGGATTGGATTACGTATTCATCACC

CACGATATACATCTAACTGGGGAGATTTTTTCATTTTTCCGGAGTTTCGGACACCCCCGTCTAGAAGCAGTCACTGCCGC

AGAAAATGTCAGGAAACATATGAATCAACCGAAGGTAATCAGTTATGAGACTATGATGAAAGGACATGCGGTATTCTGCG

GGATAATCATCAATGGTTTTAGAGACCGACATGGCGGCAGCTGGCCCCCTGTTGCATTGCCAGAACATGCTTCTGCTGCG

ATCCGGAATGCGCAAGCATCAGGTGAGGGACTGACCCATGACCTGTGTATAGACAACTGGAAATCCTTTGTTGGATTCAA

ATTTGGCTGCTTTATGCCACTCAGCCTAGATAGTGATTTGACCATGTATCTCAAAGACAAGGCATTGGCTGCACTGAAGA

ATGAGTGGGATTCAGTTTACCCGAAAGAATACCTCCGTTATAACCCACCTAGAGGGACAGAGTCAAGGCGACTGGTGGAG

GTGTTCCTGAATGACTCCAGCTTCGATCCTTATAACATGATAATGTACGTGGTGAATGGCTCCTATCTTAAAGACCCTGA

GTTTAATCTCTCATACAGCCTAAAAGAGAAGGAGATAAAGGAGACAGGGCGGTTGTTTGCCAAAATGACCTATAAGATGC

GGGCCTGTCAGGTAATTGCTGAGAATCTGATATCGAATGGTGTTGGGAAGTACTTCCGAGACAATGGGATGGCGAAGGAC

GAGCATGACCTAACAAAAGCCCTTCACACTTTGGCAGTCTCAGGTGTTCCCAAGAACAACAAAGATAACCACCGAGGTGG

GCCTCCCAGAAGGACAACAACCCGAGAGGTGAGATCAAGCCAGGACACCAGAACACAAAATAGAGACAAGGTCCAAGGGG

GACCTATGTACAACTATTTGCAATGCCAACCAATCAGCCCTGATCAAGGTGAGTCATACGAGACTGTTAGTGCATTTATC

ACCGCTGACCTTAAGAAGTATTGCCTGAATTGGAGATACGAGACAATCAGCATATTTGCACAGAGGCTGAATGAAATATA

TGGACTACCATCCTTCTTTCAATGGTTGCACAGGATATTGGAAAAATCCGTACTCTACGTCAGTGACCCACATTGCCCTC

CCGATCTAGATAATCATATCCCTCTGGACAGTGTCCCTAATGCCCAAATATTCATTAAGTACCCAATGGGCGGAATAGAA

GGTTATTGCCAAAAACTATGGACAATCAGTACTATACCATACTTGTATCTGGCAGCCTATGAGAGCGGAGTAAGAATCGC

CTCACTGGTGCAGGGTGACAATCAGACAATCGCAGTGACAAAAAGAGTTCCAAGTTCTTGGCCTTATTCACTAAAAAAGA

GGGAGGCATCCAAAGCCGCTCAAAATTACTTCGTAGTCTTAAGGCAAAGATTGCACGATGTAGGTCATCACTTGAAGGCT

AATGAAACCATAGTATCTTCTCACTTTTTTGTATATTCCAAAGGGATTTATTATGACGGCCTGCTAGTCTCACAATCACT

AAAGAGCATCGCCAGATGTGTCTTCTGGTCCGAGACTATAGTGGATGAAACCAGAGCGGCTTGCAGCAATATCGCAACAA

CTGTCGCTAAGAGTATAGAGAGGGGTTATGATAGGTACCTTGCATATTCTTTGAATATCCTCAAGATTTTCCAACAGATC

CTTATATCCCTTAACTTCACTATTAACACAACAATGACTCAGGATGTCGTGGCACCGATCGTCGAGAACGGTGATTTGCT

GATAAGGATGGCACTCTTGCCAGCACCCATCGGAGGTCTAAATTATCTTAACATGAGCAGGCTATTCGTGAGAAATATCG

GCGACCCGGTCACCTCCTCTATAGCCGACCTGAAGAGGATGATAGATGCCGGGCTCATGCCAGAAGAAACATTGCATCAA

GTGATGACCCAGACCCCGGGAGAATCATCCTACCTTGATTGGGCAAGTGACCCTTATTCTGCCAACCTACCCTGCGTACA

GAGTATAACTCGTCTTCTCAAAAACATCACTGCACGATATATTTTAATCAGCAGCCCAAATCCGATGCTGAAAGGATTGT

TTCATGAGGGGAGTAGAGATGAAGACGAGGAGCTTGCAAGTTTCCTAATGGATCGGCATATAATTGTTCCAAGAGCTGCA

CATGAAATCTTAGACCATAGTATAACCGGGGCAAGAGAGGCTATAGCCGGGATGTTGGACACCACCAAGGGTCTGATTAG

AACAAGTATGAAACGGGGTGGCCTCACCCCTCGAGTCTTAGCCCGCCTTTCCAATTATGATTATGAACAATTCCGATCCG

GGATAACACTATTGACAAAGAAAGGGCAGTGTTATCTCATTGACAAGGATTCGTGCTCGGTGCAGCTCGCTATAGCTCTG

AGGAGCCACATGTGGGCTAGGTTGGCTCGCGGGAGACCAATCTATGGGTTGGAGGTGCCCGATATATTAGAATCGATGAA

CGGCTACCTTATCAAACGCCACGAGTCTTGTGCTATCTGTGAAACGGGCTCAAGTCACTACGGATGGTTTTTCGTCCCTG

CAGGGTGCCAGCTTGACGATGTATCTAGAGAGACTTCGGCTCTCCGTGTACCTTATGTCGGATCAACAACTGAGGAAAGG

ACTGACATGAAACTTGCATTTGTTAGATCTCCAAGTAGATCCCTCAAATCAGCAGTTAGAATTGCAACAGTTTACTCATG

GGCCTACGGGGATGATGAGAAGTCATGGAGTGAAGCCTGGATGCTAGCTAGGCAGAGAGCCAATATCACCTTAGATGAAT

TGAGAATGATCACTCCAGTCTCTACATCCACCAACCTAGCCCACCGGTTGAGGGATCGGAGCACCCAGGTGAAATACTCG

GGGACATCCCTTGTGAGGGTTGCAAGATACACAACCATCTCCAATGATAACTTGTCATTTGTGATATCTGAGAAGAAAGT

AGATACCAACTTCATTTATCAGCAAGGGATGTTGCTTGGTCTCGGGATCCTTGAGAACCTCTTTAGGCTAGAGGCCACCA

CCGGAGTATCCAACACAGTGCTACACCTGCACGTGGAAACAGAATGTTGTGTTGTACCTATGGTGGATCACCCAAGGATA

CCGAGTCTCCGTAATATAAAAGTTACGGATGAGCTATGCACAAACCCTTTGATCTATGATAGGTCTCCCATCATAGAGCA

CGATGCAACCCGACTATACTCACAGAGCCACAGGAGACATTTGGTGGAGTTTGTTACCTGGTCCACAAGCCAGCTTTACC

ATATACTGGCTAAGTCTACAGCAATGTCCATGATTGAATTGATCACGAGATTCGAGAAAGATCACATGAATGAAATAGCC

GCCCTGATTGGCGATGACGACATCAACAGCTTCATCACAGAATTTTTGCTAGTGGAGCCTAGATTATTTATAGTTTACCT

TGGTCAGTGTGCTGCCATCAATTGGGCCTTTGATATACATTATCATCGGCCCTCGGGCAAGTACCAGATGGGGGAACTCC

TCTACTCTCTGCTCTCTCGGATGAGCAAAGGAGTATATAAGATCTTCACCAATGCTTTGAGCCACCCCAAAGTTTACAAG

AAATTTTGGCGAAGTGGTGTCATTGAGCCGATTCATGGCCCATCCCTGGATACACAGAATTTACATGTCACTGTCTGTGA

CATGATATATGGGTCCTATGTCACCTACCTGGATCTTTTGCTGAATGATGAGCTAGATGATTACCCGTATTTGCTCTGCG

AGAGTGATGAGGATGTGGTCACAGACAGGTTCGACAACATTCAAGCCAAACATCTCTGTGTATTAGCCGATGTATATTGC

AGCTCCAAGAGGTGTCCTTCCATAATCGGGATGTCTCCTATAGAAAAATGTACCATTCTCACACATTACATCAAGGGAGA

ATCAGTACAATCCCCGTCTGGAACCTCATGGAACACTGATCCCCTTGTAGTAGATCATTACTCATGCTCTCTGACCTACC

TTCGCCGAGGTTCCATCAAACAAATCAGGTTGAGGGTGGATCCTGGGTTTGTATTCGAGGCGTTGACAGACGTCGACTTT

AAACAACCTCGCAAAGCTAAGTTAGATATATCGGTCGTTGGGTTGACTGATTTCTCTCCCCCTTGCGACAACGTCGGTGA

TTTTCTAGGGACTATCAACACATTGAGGCACAATCTGCCTGTCACCGGAACCGGGGTCTCGAACTATGAAGTCCACGCTT

ATCGTAGAATTGGCCTGAATTCATCAGCATGTTACAAAGCCGTAGAGATCTCCACGTTAATCAAGCCATCCCTAGAAGTC

GGAGAGCATGGATTGTTCTTAGGAGAAGGTTCCGGTTCAATGCTGGCTGCGTACAAGGAAGTTCTTAAATTAGCAAATTG

TTATTACAACAGCGGGGTAACAGCGGAGGGCAGAGCCGGACAGAGGGAAATATCTCCCTATCCTTCAGAGATGAGCCTAG

TAGAGAATCAAATGGGGATAGAGCGGAGTGTTAAAGTGCTGTTCAACGGCAAACCTGAAGTAACTTGGGTGGGGACCACC

GATTGCTACAAGTACATAATCAGTAACATCCAAACCTCTAGTCTGGGTTTCATACATTCAGATATTGAGACGCTCCCAAC

CAAAGATGCTGTTGAGAAATTAGAAGAATTTGCCTCTATCCTATCCTTATCCTTGATTTTGGGAAAAATCGGCTCTATTA

CAGTTGTCAAAATTATGCCCATTAGCGGGGATTTTACCCAAGGCTTCATAGCCTATGCCATCCAATATTTTAGGGAGAGC

CTGCTTGCCTATCCGAGATATAGTAACTTCATCTCGACTGAGTGTTACCTTATCATGATAGGATTAAAGGCCAATCGATT

GATTAACCCAGAAGCCATTAAGCAAAGCATAATCAGAGCGGGGGTCAGGACTGCACCAGGACTTGTGAGCCATATATTAT

CAGAGAAACAGAAAGGTTGTATTCAATCTTTTTTGGGTGATCCCTATATCCAAGGAGACTTCAATAAGCACCTTAAATCT

CTAACCCCTATTGAGAAAATCCTGGTAAATTGTGGTCTCTCGATCAATGGCACAAAAATCTGTAGAGATCTAATCCACCA

TGATATCGCCTCCGGTCCAGACGGTCTTATGAGCTCTACAATTATTTTATATAGGGAATTGGCTCATTTCAAAGATAATA

TAAGGAGTCAGCACGGAATGTTCCACCCCTACCCAGTACTGGCCAATAGCCGGCAACGTGAATTAATCCTTCGAATAGCC

AAGAAATTCTGGGGGTATGTCTTGCTATATTCTGATGACCCGGCACTAATCAGACAAACAATCAAGAACTTGAAACGGAA

TCATCTAACCTTTGACTTACATAGTAACCCGTTTATTAAAGGGCTATCCAAAGCTGAGAAATTGCTGGTGCGGACGAGTT

CACTAAGAAGGGAATGGTTGTTCACTCTCGATACGAAAGAAGTGAAAGAGTGGTTCAAATTGGTAGGTTACAGTGCACTC

ATCAGAGGCTAATTAGTTATACATCTGCCCCCTTCTCCTCCGCCATGAGACCCCACTGACGATTCAGAAGATTAAAGAAA

ACTACATATTGGATAAGTATCTATTCCCAGCTTTGTCTGGT

>KT860063.1/goat/India/2014-11-23

ACCAAACAAAGTTGGGTAAGGATAGATCTTATAATAACTATGGACTAGCAAACTTAGGAGTAAAGATCCTACTGTCGGGG

GGAGGAGGAGGAGCAAGACCTCTGATTATGGCGACTCTCCTTAAAAGCTTAGCATTGTTCAAAAGGAACAAAGACAAAGC

GCCGACGGCATCAGGTTCAGGAGGGGCCATCCGAGGGATTAAGAATGTTATCATAGTCCCGATTCCCGGAGACTCGTCCA

TCACTACCCGTTCAAGACTGCTCGACAGGCTTGTCAGATTGGCCGGAGATCCTGACATCAACGGGTCAAAGCTGACCGGC

GTGATGATCAGCATGTTATCTTTGTTCGTAGAGTCACCCGGGCAGTTGATACAGCGAATCACAGATGATCCGGATGTTAG

TATCCGCCTTGTTGAGGTAGTTCAAAGTACTAGGTCCCAGTCCGGGTTGACCTTTGCATCACGTGGTGCTGATTTGGATA

ACGAGGCAGACATGTATTTTTCAACTGAGGGGCCCTCGAGTGGAGGTAAGAAAAGGATCAACTGGTTTGAGAACAGAGAA

ATAATAGACATAGAGGTGCAGGATCCAGAAGAGTTCAATATGTTGTTAGCCTCCATACTAGCACAAGTCTGGATCCTTCT

GGCCAAGGCTGTCACGGCACCGGATACGGCAGCTGACTCAGAACTGAGAAGGTGGGTTAAATACACACAACAAAGGAGAG

TGATTGGGGAATTTCGCCTTGACAAAGGGTGGCTGGACGCAGTCCGCAACAGAATTGCAGAAGATCTATCACTCCGGCGG

TTCATGGTATCTCTCATACTTGACATCAAGAGGACCCCCGGCAACAAGCCAAGGATCGCAGAAATGATCTGCGACATTGA

CAACTATATTGTCGAAGCAGGACTCGCCAGTTTCATCCTTACTATCAAATTTGGTATAGAAACCATGTATCCTGCATTAG

GGCTCCACGAGTTTGCCGGGGAATTGTCCACTATTGAGTCCTTGATGAACTTGTATCAACAGCTAGGAGAGGTTGCACCC

TACATGGTAATTCTAGAGAACTCAATTCAGAACAAGTTTAGTGCAGGAGCTTATCCCCTCCTCTGGAGCTATGCGATGGG

TGTCGGAGTTGAGCTGGAGAACTCAATGGGGGGCTTGAACTTTGGCAGATCATATTTTGACCCTGCCTATTTTCGTCTCG

GACAGGAGATGGTCAGAAGATCTGCAGGAAAGGTCAGCTCTGTGATTGCGGCTGAGCTCGGCATCACAGCAGAGGAAGCT

AAACTAGTCTCGGAAATAGCCTCGCAGGCTGGGGACGAAAGAACCGCTAGAGGAACTGGGCCTCGACAGGCGCAGGTCTC

CTTCCTACAGCACCAAACAGGAGGGGGAGAGTCGTCCGCACCAGCGACCAGAGAAGGGGTCAAAGCTGCGATCCCAAACG

GATCCGGAGAAAGGGACAGAAAGCAAACACGCCCAGGGAGGCCCAGAGGAGAGACCCCCGGCCAACTGCTCCTGGAATTC

ATGCCAGATGATGAGGTCTCGCGAGAATCTGGTCAAAACCCTCGTGAGGCTCAAAGGTCGGCCGAGGCACTCTTCAGGCT

GCAGGCCATGGCCAAGATTCTGGAGGACCAGGAGGAGGGAGAAGACGACAATCAGGTCCACAACGACAAGGATCTCCTCG

GCTAAGCAGACGCACCCTCTGTCGAAATCAGTGACGAGACATCTCCCACCAGTATTATAAAAAACTTAGGACCCAGGTCC

AAGCAAGCACACATCGACACTCCAACCAGTCGAGCGGAGGCCACCGATGGCAGAAGAACAGGCATACCATGTCAACAAGG

GGCTGGAATGTATCAAGTCCCTCAAAGCCTCTCCCCCGGATCTATCCACCATTAGAGATACCATCGCGAGCTGGAGAGAG

GGGCTTAGCCCCTCGGGCCGTGCAACACCGAACCCTGATACGTCCGAGGGAGATCATCAGAATATCAACCAATCATGCTC

ACCAGCAATCGGACCAAACAAAGTCTACTTGTCTCCTGAAGATAATCTCGGATTTAGAGAGATCACTGGCAACGACTGTG

GGGCTGAGCTCGGAGGAGTCCAGGGAAAAGGATCCAACTCTCAAGTACAGCGTTACTATGTTTATAGCCACGGGGGTGAA

GAGATTGAAGGACTCGAGGATGCTGACTCTCTCGTGGTTCAAGCAGATCCTCCAGTTGCTAACATATTCAATGGAGGAGA

GGATGGATCTGACGACAGCGATGTGGACTCTGGCCCAGATGATCCCAGCAGAGATACTCTATATGACCGGGGACCTGCTG

CCGGCAATGATGTCGCTAGGTCCACAGATGTCGAAAAACTAGAAGGTGCTGATATTCAAGAAGTTCTCAACTCCCAGAAA

AGCAGAGGAGGAAGATTCCAAGGCGGGAAAACCTTACGAGTCCCGGAAGTACCCGATGTCAAGAACTCCGGACCATCAGC

CCAATCAATTAAAAAGGGCACAGACGGGAACTCAGTCTCATCTGGAACGGTGACAGAGTGTTTATCGATAAGTGGTGCAA

CCCAAGCTGTGCCAGAGTCAAGATGGGAGTCATCAGAGCAAAATGCGTCTGTGGGGAGTGTCCTCAAGTCTGCGAAGAGT

GCAAAGACGATCCAGGGGTCGACACAAGAATCTGGTACCACAGCATCACCGACTCAGCCTAAAGAGAATGACTCCGAGTA

TGAGTATGAGGATGACCTATTTACAGAGATTCAGGACATCCGTGCAAGCATTGCCAAGATCCATGATGACAACAAGACTA

TCCTCTCAAAGCTTGATTCTATACTGTTATTGAAAGGAGAAGTCGACACTATCAAGAAACAAATCAGCAAGCAGAATATA

AGTATATCCACCATTGAGGGCCATCTCTCCAGTATAATGATAGCCATCCCGGGCTTTGGGAAGGATATCAAGGACCCAAC

ATCCGAGGTTGAGTTGAACCCAGATTTAAGACCTATAATAAGCCGTGACTCTGGCAGAGCTCTCGCGGAGGTCCTCAAGA

AACCCGCTGTGGATAGGTCTCAGAAAACTGAAACCAAAGCCAACTCCAGCTCAAAGGGTCAGCTTCTTAAGGATCTCCAG

CTAAAACCTGTCGACAAGCAGGCAAGCTCTGCAATCGGGTTTGTCCCATCCGACCATGAATCATCCAGAAATGTCATCCG

CTCCATAATCAAGTCGAGCAAGCTAAACATTGATCACAAGGACTATCTTCTAGATTTACTGAATGATGTGAAAGGCTCAA

AGGATCTTAAGGAATTCCACAAGATGCTAACAGCAATTCTCGCCAAGCACCCGTAACACATCCCCCAGTCACCATCTCAT

ACTCGGCTGAAGACATCCTTTCAATCAGACTATTACAAAAAACTTAGGAGCAAGGGCAACTGAGCTTCGCAGACAAGATG

ACCGAGATCTACGACTTCGACAAATCAGCATGGGATGTCAAAGGGTCAATTGCCCGCATAGAACCCACCACCTACCACGA

CGGCCGACTGATACCCCAGGTAAGGGTCATCGATCCTGGTCTGGGAGACAGAAAAGATGAGTGCTTTATGTACCTGTTTC

TCCTGGGAGTGATTGAGGATAACGACCCCCTGTCTCCCCCCGTCGGAAGAACCTTTGGCTCTTTACCTCTAGGGGTCGGT

AGGTCAACTGCTAAGCCAGAAGAACTACTAAGGGAGGCCACAGAACTAGACATAGTGGTGAGGCGCACTGCAGGAGTAAA

TGAGAAACTGGTATTTTACAACAACACTCCGCTGTCCTTGTTAACGCCCTGGAAGAAAGTTCTGACAACCGGAAGTGTGT

TTAGCGCTAACCAGGTTTGCAATGCAGTCAACCTAGTCCCGCTTGATACTCCCCAGAGGTTCAGGGTTGTGTACATGAGC

ATAACTAGATTATCAGACAATGGTTACTATAGTGTGCCCAGGAGAATGCTAGAGTTCCGCTCAGCCAATGCAGTAGCCTT

CAACATCTTGGTTACACTAAGAATTGAGAATGGCACAAACCCTAGAAGATACATAGTCGGCTCATGGGAGAATTCAGAGG

TCACATTTATGGTACACGTGGGCAACTTTAGAAGAAAGAAGAACGAAGTATACTCTGCTGATTACTGCAAAATGAAGATT

GAAAAGATGGGTTTAGTTTTTGCCCTGGGAGGAATAGGTGGAACAAGTCTCCATATTAGAAGCACCGGGAAAATGAGCAA

AACCCTCCATGCACAGCTGGGGTTCAAGAAAATCTTATGTTACCCCCTAATGGATGTTAATGAGGATCTTAACCGATATC

TCTGGCGGGCAGAGTGCCGAATAGTCAAAATCCAGGCCGTCTTACAGCCATCAGTGCCCCAAGAATTCCGTGTCTACGAT

GATGTCATCATCAACGATGACCAAGGCTTGTTCAAGATCCTGTAGTTCATCTGCAAAGGAACGAACCCACCCCCAGGGGG

GGGCCCCCCCCTCCCCCG-CCAAAAAACACACTCCAAGGCTCCCCGACACAGCGCCAACACAGCCACCCTCCCCACTGGA

CGAGGCCGGCC-CCCCAACAAACCCGGTCCAGGC-GGGGGGCCCCCCCCCTGACCCGCCGGGCGCCGACCGAGCAGGCCG

GCGAGAAGCCCCCAAGAGGCCCCGACGAAGACCGAGCACCCCTCCCCCCC-AAACCGTTCCCCC-AACACGCCCGAGGCC

AGGCGCCGGGCACCCCCAGACCCGCCACCGGCGGGACAGAACCCCAGGGGGGAACCCCCCAAACCAAAGGGACCACCGGG

GAAGGATCCGCCGAACCAGCACAGACCCGACCCAAGCAAGGCGGACCCCCAACATGAAACCGCCCAGCACACCCCACTCA

TCATCAAACAAAACTTAGGGGCCAAGCCCATACGACAGGCCCCAGTGCGCCAGGCGGAGGGACGCAGAAAGGAAGGAGAC

AAGACCGCCCCAGAAGGAGGACACCCAACCACCGAAACACCCAGCCCCAGGGAAGGAG--CCCCCCCCCACATGGCGAAA

GAACGGGCAAGAGACCCCGAGAAGAACCCGACCGGGAGCCAGGAGCCGGCGACGCCGGCGGCAAACAAGCCGGGCAAACA

CCCGACCCCACCAGAGGAGCCGAGGAGAGCCCGACCCCGCACGAACCAAACAGCAGGGAGCCCAACCCCACCAAAAAAAG

AGGCGAGGCCCC-CCT-----CCTCCCCGCACACAACCCGCAGACAGACCGAGAGAGGACCACCTCCCAACAAGACAGGC

GGCTCCCTCGCAGCCGAGCCGCAGCTCCCCAGAACAGGACACCCCCGTACACTCAAAACACCCCCCCGCCCCCCGCCACC

GGGACAGGGGCTCCGCCACCACAGACCAAAGCCACCGAGAGCAAGCCAGAGGTCCCCCCCAGTCCCGCCCGGCGCCCCCC

CGAAAAGGCCCCGGCAAGGCCCCCCAACCCGACGTCCCCCGCACGCACCGAAAGAGGGACCCGCCCGAGAACGTGCGGCA

CATCCATGCGTAAACATTATGACACGGGTTGCAATCTTGACATCTCTGTTTCTTCTCCCAAATGTTGTTGCGTGCCAGAT

TCACTGGGGCAATCTATCCAAGATCGGGATTGTGGGAACAGGGAGTGCCAGCTACAAGGTGATGACTAGGCCGAGCCACC

AGACTCTGGTTATAAAGTTAATGCCAAATATAACAGCCATCGACAATTGTACAAAGTCAGAGATTGCAGAGTACAAGAGA

TTGCTGATCACAGTGTTAAAGCCTGTAGAGGATGCTCTGTCGGTTATAACCAAGAATGTAAGACCAATTCAAACCCTGAC

ACCTGGGCGTAGAACCCGTCGTTTTGCTGGAGCTGTTCTGGCCGGAGTAGCGCTTGGAGTTGCGACAGCCGCTCAGATAA

CTGCAGGAGTCGCTCTTCATCAATCATTGATGAACTCCCAAGCAATTGAGAGTTTAAAAACCAGTCTTGAGAAGTCGAAT

CAGGCAATAGAAGAAATCAGACTTGCAAATAAGGAGACCATACTGGCAGTACAGGGCGTCCAGGATTATATCAACAATGA

GCTTGTCCCTTCTGTTCATAGAATGTCATGCGAGCTGGTAGGCCACAAGCTCGGCCTCAAGCTCCTTAGGTACTACACCG

AGATCCTGTCCATATTCGGGCCCAGCCTTCGAGACCCGATAGCTGCCGAAATATCAATCCAGGCACTCAGTTATGCATTA

GGCGGAGACATTAATAAAATCCTGGACAAGCTTGGGTATAGCGGTGGGGATTTCCTTGCCATCCTAGAAAGCAAGGGAAT

AAAGGCCCGGGTTACATATGTGGACACAAGAGATTACTTTATAATGCTTAGCATCGCCTACCCAACCTTATCTGAGATCA

AGGGAGTGATAGTTCACAAGATAGAAGCTATAACATACAACATTGGGGCACAGGAGTGGTATACTACTATCCCTAAATAT

GTAGCCACTCAGGGATATCTGATATCGAACTTTGATGAGACGTCATGCGTATTCACTCCAGAGGGGACAGTTTGCAGCCA

GAATGCATTGTACCCAATGAGCCCATTGCTTCAGGAATGTTTCAGGGGGTCAACAAAATCGTGCGCCAGAACCCTAGTTT

CAGGTACCATAAGTAATAGATTTATCCTATCAAAAGGAAACCTGATCGCGAATTGTGCGTCAGTTTTGTGCAAATGTTAC

ACAACGGAGACAGTCATCAGCCAAGATCCTGACAAACTACTAACTGTTGTAGCATCCGACAAGTGTCCTGTAGTTGAGGT

GGATGGAGTGACAATACAGGTCGGCAGTCGAGAGTATCCGGATTCTGTATACTTACACAAAATAGACTTAGGCCCAGCCA

TCTCCCTAGAAAAACTAGATGTAGGTACCAATTTAGGCAATGCAGTTACAAGACTGGAGAATGCAAAGGAGCTCCTAGAT

GCATCAGACCAAATACTGAAGACTGTCAAAGGGGCACCTTTTGGTGGGAACATGTACATAGCACTGGCAGCTTGCATTGG

AGTATCCCTAGGGCTTGTCACATTAATATGCTGCTGTAAGGGGAGGTGTAAGAACAAGGAAATCCCTATCTCCAAAATCA

ACCCAGGGCTCAAACCCGATCTGACCGGGACCTCAAAGTCGTACGTGAGATCACTGTAGCCAGAATCACCTGAATCATCT

GGCATCACACACATACATGTACGACACAAGCAGTCTGAGGACGCAGGAAACCCAGCCCCCGATCACCCACCTGACCCTAC

TCCACGCTCCACTACACATTAGTCATTAAACAAAACTTAGGACGAAAGGTCAATCACCATGTCCGCACAAAGGGAAAGGA

TCAATGCCTTCTACAAAGACAATCCTCACAATAAGAACCATAGGGTAATCCTGGATAGAGAACGCTTGATCATTGAAAGA

CCCTACATCTTGCTTGGAGCCCTGCTGGTAATGTTCCTGAGTCTAATCGGACTGCTGGCCATTGCAGGGATCAGGCTTCA

CCGGGCCACCGTTGGAACTTCAGAGATCCAGAGTCGGCTGAATACCAATATTAAGTTGACCGAATCTATTGATCACCAGA

CTAAGGATGTCTTAACTCCCCTCTTTAAAATAATTGGCGATGAAGTCGGCATCAGGATTCCACAGAAATTCAGTGATCTT

GTCAAGTTCATCTCCGACAAGATTAAATTCCTCAACCCCGATAGAGAGTATGATTTCAGGGATCTCCGGTGGTGCATGAG

TCCCCCCGAGAGAGTCAAAATTAATTTTGATCAGTTTTGTGAGTACAAAGCTGCGGTTAATTCAATTGAACATATATTTG

AGTCACCGCTCAACAAGTCAAAAAAGCTGCAATCATTGACTCTCGGGCCCGGAACAAGTTGTCTAGGCAGGACAGTAACA

AGAGCTCATTTCTCAGAACTTACACTGACCTTAATGGACTTGGATCTAGAGATGAAGCACAACGTGTCCTCAGTGTTTAC

CGTAGTTGAAGAGGGGTTATTCGGAAGAACATATACCGTCTGGAGATCCGATGCCAGGGATCCGAGCACCGATCTAGGTA

TCGGCCATTTTTTAAGAGTCTTCGAGATTGGACTGGTAAGAGATCTCGGGCTGGGTCCCCCCGTTTTTCATATGACTAAC

TATCTCACAGTGAACATGAGTGATGTCTATCGGAGATGTCTTTTAGCGGTAGGGGAGTTGAAGTTGACAGCCCTATGCAC

CTCATCTGAGACTGTAACACTGAGTGAGAGAGGAGTTCCAAAGAGGGAGCCTCTTGTAGTTGTGATACTTAACCTGGCTG

GACCAACTCTAGGGGGCGAACTATACAGTATCTTGCCTACCTCTGATCTCATGGTGGAGAAACTCTATTTATCTTCACAT

AGAGGGATCATCAAGGATGACGAGGCCAATTGGGTAGTGCCGTCTACCGATGTTCGTGATCTTCAGAACAAAGGTGAGTG

TCTAGTGGAAGCATGCAAGACTCGACCTCCTTCATTTTGCAATGGCACAGGATCAGGCCCGTGGTCAGAGGGGAGAATCC

CTGCTTACGGGGTGATCAGGGTCAGTCTTGACTTAGCTAGTGACCCGGATGTAGTTATCACTTCAGTGTTTGGCCCACTG

ATTCCTCACCTATCCGGCATGGATCTTTACAACAACCCGTTTTCAAGAGCTGTATGGTTGGCTGTACCACCTTACGAGCA

GTCATTTCTAGGAATGATAAATACAATTGGATTCCCTAACAGAACAGAGGTTATGCCGCACATTTTGACCACAGAGATCA

GAGGCCCTCGGGGTCATTGCCATGTCCCCATAGAATTGTCCCGCAGGGTTGATGACGATATCAAGATCGGGTCCAACATG

GTCATATTGCCGACGATGGACCTGAGATATATTACAGCCACTTATGATGTTTCCAGGAGCGAGCATGCAATCGTGTACTA

TATCTATGACACAGGTCGCTCATCATCTTACTTCTACCCAGTTCGACTGAATTTCAAAGGCAATCCTCTCTCTCTGAGGA

TAGAGTGTTTCCCTTGGCGTCATAAGGTGTGGTGCTACCATGATTGTCTTATATACAACACCATAACAGATGAAGAGGTC

CATATGAGAGGGCTGACCGGTATAGAGGTAACATGCAATCCAGTCTGAGCAGAGCTACGACCGTCACTCAAGCAGTCCCT

TGAGTCGCCACCGAGTCCAAGCAGCACAGCCTGGGACTCTCAACAGCACAGCCCAGCTAACAGTGTTATAAAAAACTTAG

GAGCCAGGGTTGTAGGGGCCATGGACTCCCTATCAGTCAATCAGGTCTTGTACCCTGAGGTTCATCTAGACAGCCCTATT

GTCACAAACAAACTAGTTGCCATCCTTGAGTACTCGGGTATCGACCATAACTATGTTCTTGAAGACCAGACCCTTGTCAA

GAATATTAGGTATAGACTGGGGTGCGGTTTTTCAAATCAAATGATCATCAATAACAGGGGGGTGGGTGAAACAGTCAATT

CTAAACTTAAAAGTTATCCCCATAATCGTCATATCATATACCCGGATTGCAATAAGGAGTTGTTTTGTATCAAAGATAGC

TGCATATCTAAGAAGCTCTCGGAGCTATTCAAGAAGGGTAATTCCTTGTACTCTAAGATAAGTCACCAGGTACTGGACTG

TCTTAAGAGAGTCAATGGGAAATTAGGCCTGGGCACAGATCTTACCCATGGTCTGAAGGAGGGTATCCTTGACCTAGGGT

TGCACATGCATAGCTCTCAATGGTTCGAGACTTTCCTGTTCTGGTTCACTATCAAGACAGAGATGAGATCAATGATCAAA

GAACAGTCCCATATATGCCACAAGAGGAGATATAACCCGATTTTTGTATCAGGGGATGCATTCGAGGTGCTCGTATCACG

AGACCTCGTTGTGATAATTGATAAGAATACCCAGTATGTCTTCTACTTAACATTTGAGCTGGTCCTTATGTATTGTGATG

TCATAGAGGGCAGACTTATGACGGAGACAGCCATGGTCATAGACCAGAGATATTCAGAACTTCTAAGCCGGGTTAGATAC

CTGTGGGATCTTATTGATGGGTTCTTCCCAACACTAGGCAATACCACCTACCAAATTGTTGCTCTGCTTGAACCATTATC

GTTGGCTTATCTTCAACTTCAGGATGTCACGCTGGAGTTAAGAGGTGCTTTTTTAGACCATTGCTTCAAAGAACTCTATG

AGATACTGGAGCATTGCGGCATTGACACAGAAGGCACCTATAATTCCATCACTGAAGGATTGGATTACGTATTCATCACC

CACGATATACATCTAACTGGGGAGATTTTTTCATTTTTCCGGAGTTTCGGACACCCCCGTCTAGAAGCAGTCACCGCCGC

AGAAAATGTCAGGAAACATATGAATCAACCGAAGGTAATCAGTTATGAGACTATGATGAAAGGACATGCGGTATTCTGCG

GGATAATCATCAATGGTTTTAGAGACCGACATGGCGGCAGCTGGCCCCCTGTTGCATTGCCAGAGCATGCTTCTGCTGCG

ATCCGGAATGCGCAAGCATCAGGTGAGGGACTGACCCATGACCTGTGTATAGACAACTGGAAATCCTTTGTTGGATTCAA

ATTTGGCTGCTTTATGCCACTCAGCCTAGACAGTGATTTGACCATGTATCTCAAAGACAAGGCATTGGCTGCACTGAAGA

ATGAGTGGGATTCAGTTTACCCGAAAGAATACCTCCGTTATAACCCACCTAGAGGGACAGAGTCAAGGCGACTGGTGGAG

GTGTTCCTGAATGACTCCAGCTTCGATCCTTATAACATGATAATGTACGTGGTGAATGGCTCCTATCTTAAAGACCCTGA

GTTTAATCTCTCATACAGCCTAAAAGAGAAGGAGATAAAGGAGACAGGGCGGTTGTTTGCCAAAATGACCTATAAGATGC

GGGCCTGTCAGGTAATTGCTGAGAATCTGATATCGAATGGTGTTGGGAAGTACTTCCGAGACAATGGGATGGCGAAGGAC

GAGCATGACCTAACAAAAGCCCTTCACACTTTGGCAGTCTCAGGTGTTCCCAAGAACAACAAAGATAACCGCCGAGGTGG

GCCTCCCAGAAGGACAACAACCCGAGAGGTGAGATCAAGCCGGGACACCAGAACACAAAATAGAGACAAGGTCCAAGGGG

GACCTATGTACAACTATTTGCAATGCCAACCAATCAGCCCTGATCAAGGTGAGTCATACGAGACTGTTAGTGCATTTATC

ACTGCTGACCTTAAGAAGTATTGCCTGAATTGGAGATACGAGACAATCAGCATATTTGCACAGAGGCTGAATGAAATATA

TGGACTACCATCCTTCTTTCAATGGTTGCACAGGATATTGGAAAAATCCGTACTCTACGTCAGTGACCCACATTGCCCTC

CCGATCTAGATAATCATATCCCTCTGGACAGTGTCCCTAATGCCCAAATATTCATTAAGTACCCAATGGGTGGAATAGAA

GGTTATTGCCAAAAACTATGGACAATCAGTACTATACCATACTTGTATCTGGCAGCCTATGAGAGCGGAGTAAGAATCGC

CTCACTGGTGCAGGGTGACAATCAGACAATCGCAGTGACAAAAAGAGTTCCAAGTTCTTGGCCTTATTCACTAAAAAAGA

GGGAGGCATCCAAAGCCGCTCAAAATTACTTCGTAGTCTTAAGGCAAAGATTGCACGATGTAGGTCATCACTTGAAGGCT

AATGAAACCATAGTATCTTCTCACTTTTTTGTATATTCCAAAGGGATTTATTATGACGGCCTGCTAGTCTCACAATCACT

AAAGAGCATCGCCAGATGTGTCTTCTGGTCCGAGACTATAGTGGATGAAACCAGAGCGGCTTGCAGCAATATCGCAACAA

CTGTCGCTAAGAGTATAGAGAGGGGTTATGATAGGTACCTTGCATATTCTTTGAATATCCTCAAGATTTTCCAACAGATC

CTTATATCCCTTAACTTCACTATTAACACAACAATGACTCAGGATGTCGTGGCACCGATCGTCGAGAACGGTGATTTGCT

GATAAGGATGGCACTCTTGCCAGCACCCATCGGAGGTCTAAATTATCTTAACATGAGCAGGCTATTCGTGAGAAATATCG

GCGACCCGGTCACCTCCTCTATAGCCGACCTGAAGAGGATGATAGATGCCGGGCTCATGCCAGAAGAAACATTGCATCAA

GTGATGACCCAGACCCCGGGAGAATCATCCTACCTTGATTGGGCAAGTGACCCTTATTCTGCCAACCTACCCTGCGTACA

GAGTATAACTCGTCTTCTCAAAAACATCACTGCACGATATATTTTAATCAGCAGCCCAAATCCGATGCTGAAAGGATTGT

TTCATGAGGGGAGTAGAGATGAAGACGAGGAGCTTGCAAGTTTCCTAATGGATCGGCATATAATTGTTCCAAGAGCTGCA

CATGAAATCTTAGACCATAGTATAACCGGGGCAAGAGAGGCTATAGCCGGGATGTTGGACACCACCAAGGGTCTGATTAG

AACAAGTATGAAACGGGGTGGCCTCACCCCTCGAGTCTTAGCCCGCCTTTCCAATTATGATTATGAACAATTCCGATCCG

GGATAACACTATTGACAAAGAAAGGGCAGTGTTATCTCATTGACAAGGATTCGTGCTCGGTGCAGCTCGCTATAGCTCTG

AGGAGCCACATGTGGGCTAGGTTGGCTCGCGGGAGACCAATCTATGGGTTGGAGGTGCCCGATATATTAGAATCGATGAA

CGGCTACCTTATCAAACGCCACGAGTCTTGTGCTATCTGTGAAACGGGCTCAAGTCACTACGGATGGTTTTTCGTCCCTG

CAGGGTGCCAGCTTGACGATGTATCTAGAGAGACTTCGGCTCTCCGTGTACCTTATGTCGGATCAACAACTGAGGAAAGG

ACTGACATGAAACTTGCATTTGTTAGATCTCCAAGTAGATCCCTCAAATCAGCAGTTAGAATTGCAACAGTTTACTCATG

GGCCTACGGGGATGATGAGAAGTCATGGAGTGAAGCCTGGATGCTAGCTAGGCAGAGAGCCAATATCACCTTAGATGAAT

TGAGAATGATCACTCCAGTCTCTACATCCACCAACCTAGCCCACCGGTTGAGGGATCGGAGCACCCAGGTGAAATACTCG

GGGACATCCCTTGTGAGGGTTGCAAGATACACAACCATCTCCAATGATAACTTGTCATTTGTGATATCTGAGAAGAAAGT

AGATACCAACTTCATTTATCAGCAAGGGATGTTGCTTGGTCTCGGGATCCTTGAGAACCTCTTTAGGCTAGAGGCCACCA

CCGGAGTATCCAACACAGTGCTACACCTGCACGTGGAAACAGAATGTTGTGTTGTACCTATGGTGGATCACCCAAGGATA

CCGAGTCTCCGTAATATAAAAGTTACGGATGAGCTATGCACAAACCCTTTGATCTATGATAGGTCTCCCATCATAGAGCA

CGATGCAACCCGACTATACTCACAGAGCCACAGGAGACATTTGGTGGAGTTTGTTACCTGGTCCACAAGCCAGCTTTACC

ATATACTGGCTAAATCTACAGCAATGTCCATGATTGAATTGATCACGAGATTCGAGAAAGATCACATGAATGAAATAGCC

GCCCTGATTGGCGATGACGACATCAACAGCTTCATCACAGAATTTTTGCTAGTGGAGCCCAGATTATTTATAGTTTACCT

TGGTCAGTGTGCTGCCATCAATTGGGCCTTTGATATACATTATCATCGGCCCTCGGGCAAGTACCAGATGGGGGAACTCC

TCTACTCTCTGCTCTCACGGATGAGCAAAGGAGTATATAAGATCTTCACCAATGCTTTGAGCCACCCCAAAGTTTACAAG

AAATTTTGGCGAAGTGGTGTCATTGAGCCGATTCATGGCCCATCCCTGGATACACAGAATTTACATGTCACTGTTTGTGA

CATGATATATGGGTCCTATGTCACCTACCTGGATCTTTTGCTGAATGATGAGCTAGATGATTACCCGTATTTGCTCTGCG

AGAGTGATGAGGATGTGGTCACAGACAGGTTCGACAACATTCAAGCCAAACATCTCTGTGTATTAGCCGATGTATATTGC

AGCTCCAAGAGGTGTCCTTCCATAATCGGGATGTCTCCTATAGAAAAATGTACCATTCTCACACATTACATCAAGGGAGA

ATCAGTACAATCCCCGTCTGGAACCTCATGGAACACTGATCCCCTTGTAGTAGATCATTACTCATGCTCTCTGACCTACC

TTCGCCGAGGTTCCATCAAACAAATCAGGTTGAGGGTGGATCCTGGGTTTGTATTCGAGGCGTTGACAGACGTCGACTTT

AAACAACCTCGCAAAGCTAAGTTAGATATATCGGTCGTTGGGTTGACTGATTTCTCTCCCCCTTGCGACAACGTCGGTGA

TTTTCTAGGGACTATCAACACATTGAGGCACAATCTGCCTGTCACCGGAACCGGGGTCTCGAACTATGAAGTCCACGCTT

ATCGTAGAATTGGCCTGAATTCATCAGCATGTTACAAAGCCGTAGAGATCTCCACGTTAATCAAGCCATCCCTAGAAGTC

GGAGAGCATGGATTGTTCTTAGGAGAAGGTTCCGGTTCAATGCTGGCTGCGTACAAGGAAGTTCTTAAATTAGCAAATTG

TTATTACAACAGCGGGGTAACAGCGGAGGGCAGAGCCGGACAGAGGGAAATATCTCCCTATCCTTCAGAGATGAGCCTAG

TAGAGAATCAAATGGGGATAGAGCGGAGTGTTAAAGTGCTGTTCAACGGCAAACCTGAAGTAACTTGGGTGGGGACCACC

GATTGCTACAAGTACATAATCAGTAACATCCAAACCTCTAGTCTGGGTTTCATACATTCAGATATTGAGACGCTCCCAAC

CAAAGATGCTGTTGAGAAATTAGAAGAATTTGCCTCTATCCTATCCTTATCCCTGATTTTGGGGAAAATCGGCTCTATTA

CAGTTGTCAAAATTATGCCCATTAGCGGGGATTTCACCCAAGGCTTCATAGCCTATGCCATCCAATATTTTAGGGAGAGC

CTGCTTGCCTATCCGAGATATAGTAACTTTATCTCGACTGAGTGTTACCTTATCATGATAGGATTAAAGGCCAATCGATT

GATTAACCCAGAAGCCATTAAGCAAAGCATAATCAGAGCGGGGGTCAGGACTGCACCAGGACTTGTGAGCCATATATTAT

CAGAGAAACAGAAAGGTTGTATTCAATCTTTTTTGGGTGATCCCTATATCCAAGGAGACTTCAATAAGCACCTTAAATCT

CTAACCCCTATTGAGAAAATCCTGGTAAATTGTGGTCTCTCGATCAATGGCACAAAAATCTGTAGGGATCTAATCCACCA

TGATATCGCCTCCGGTCCAGACGGTCTTATGAGCTCTACAATTATTTTATATAGGGAATTGGCTCATTTCAAAGATAATA

TAAGGAGTCAGCACGGAATGTTCCACCCCTACCCAGTACTGGCCAATAGCCGGCAACGTGAATTAATCCATCGAATAGCC

AAGAAATTCTGGGGGTATGTCTTGCTATATTCTGATGACCCGGCACTAATCAGACAAACAATCAAGAACTTGAAACGGAA

TCATCTAACCTTTGACTTACATAGTAACCCGTTTATTAAAGGGCTATCCAAAGCTGAGAAACTGCTGGTGCGGACGAGTT

CACTAAGAAGGGAATGGTTGTTCACTCTCGATACGAAAGAAGTGAAAGAGTGGTTCAAATTGGTAGGTTACAGTGCACTC

ATCAGAGGCTAATTGGTTATACATCTGCCCCCTTCTCCTCCGCCATGAGACCCCACTGACGATTCAGAAGATTAAAGAAA

ACTACATATTGGATAAGTATCTATTCCCAGCTTTGTCTGGT

>KT270355.1/goat/India/2014-09-25

ACCAAACAAAGTTGGGTAAGGATAGATCTTATAATAACTATGGACTAGCAAACTTAGGAGTAAAGATCCTACTGTCGGGG

GGAGGAGGAGGAGCAAGACCTCTGATTATGGCGACTCTCCTTAAAAGCTTAGCATTGTTCAAAAGGAACAAAGACAAAGC

GCCGACGGCATCAGGTTCAGGAGGGGCCATCCGAGGGATTAAGAATGTTATCATAGTCCCGATTCCCGGAGACTCGTCCA

TCACTACCCGTTCAAGACTGCTCGACAGGCTTGTCAGATTGGCCGGAGATCCTGACATCAACGGGTCAAAGCTGACCGGC

GTGATGATCAGCATGTTATCTTTGTTCGTAGAGTCACCCGGGCAGTTGATACAGCGAATCACAGATGATCCGGATGTTAG

TATCCGCCTTGTTGAGGTAGTTCAAAGTACTAGGTCCCAGTCCGGGTTGACCTTTGCATCACGTGGTGCTGATTTGGATA

ACGAGGCAGACATGTATTTTTCAACTGAGGGGCCCTCGAGTGGAGGTAAGAAAAGGATCAACTGGTTTGAGAACAGAGAA

ATAATAGACATAGAGGTGCAGGATCCAGAAGAGTTCAATATGTTGTTAGCCTCCATACTAGCACAAGTCTGGATCCTTCT

GGCCAAGGCTGTCACGGCACCGGATACGGCAGCTGACTCAGAACTGAGAAGGTGGGTTAAATACACACAACAAAGGAGAG

TGATTGGGGAATTTCGCCTTGACAAAGGGTGGCTGGACGCAGTCCGCAACAGAATTGCAGAAGATCTATCACTCCGGCGG

TTCATGGTATCTCTCATACTTGACATCAAGAGGACCCCCGGCAACAAGCCAAGGATCGCAGAAATGATCTGCGACATTGA

CAACTATATTGTCGAAGCAGGACTCGCCAGTTTCATCCTTACTATCAAATTTGGTATAGAAACCATGTATCCTGCATTAG

GGCTCCACGAGTTTGCCGGGGAATTGTCCACTATTGAGTCCTTGATGAACTTGTATCAACAGCTAGGAGAGGTTGCACCC

TACATGGTAATTCTAGAGAACTCAATTCAGAACAAGTTTAGTGCAGGAGCTTATCCCCTCCTCTGGAGCTATGCGATGGG

TGTCGGAGTTGAGCTGGAGAACTCAATGGGGGGCTTGAACTTTGGCAGATCATATTTTGACCCTGCCTATTTTCGTCTCG

GACAGGAGATGGTCAGAAGATCTGCAGGAAAGGTCAGCTCTGTGATTGCGGCTGAGCTCGGCATCACAGCAGAGGAAGCT

AAACTAGTCTCGGAAATAGCCTCGCAGGCTGGGGACGAAAGAACCGCTAGAGGAACTGGGCCTCGACAGGCGCAGGTCTC

CTTCCTACAGCACCAAACAGGAGGGGGAGAGTCGTCCGCACCAGCGACCAGAGAAGGGGTCAAAGCTGCGATCCCAAACG

GATCCGGAGAAAGGGACAGAAAGCAAACACGCCCAGGGAGGCCCAGAGGAGAGACCCCCGGCCAACTGCTCCTGGAATTC

ATGCCAGATGATGAGGTCTCGCGAGAATCTGGTCAAAACCCTCGTGAGGCTCAAAGGTCGGCCGAGGCACTCTTCAGGCT

GCAGGCCATGGCCAAGATTCTGGAGGACCAGGAGGAGGGAGAAGACGACAATCAGGTCTACAACGACAAGGATCTCCTCG

GCTAAGCAGACGCACCCTCTGTCGAAATCAGTGACGAGACATCTCCTACCAGTATTATAAAAAACTTAGGACCCAGGTCC

AAGCAAGCACACATCGACACTCCAACCAGTCGAGCGGAGGCCACCGATGGCAGAAGAACAGGCATACCATGTCAACAAGG

GGCTGGAATGTATCAAGTCCCTCAAAGCCTCTCCCCCGGATCTATCCACCATTAGAGATACCATCGCGAGCTGGAGAGAG

GGGCTTAGCCCCTCGGGCCGTGCAACACCGAACCCTGATACGTCCGAGGGAGATCATCAGAATATCAACCAATCATGCTC

ACCAGCAATCGGACCAAACAAAGTCTACTTGTCTCCTGAAGATAATCTCGGATTTAGAGAGATCACTGGCAACGACTGTG

GGGCTGAGCTCGGAGGAGTCCAGGGAAAAGGATCCAACTCTCAAGTACAGCGTTACTATGTTTATAGCCACGGGGGTGAA

GAGATTGAAGGACTCGAGGATGCTGACTCTCTCGTGGTTCAAGCAGATCCTCCAGTTGCTAACATATTCAATGGAGGAGA

GGATGGATCTGACGACAGCGATGTGGACTCTGGCCCAGATGATCCCAGCAGAGATACTCTATATGACCGGGGACCTGCTG

CCGGCAATGATGTCGCTAGGTCCACAGATGTCGAAAAACTAGAAGGTGCTGATATTCAAGAAGTTCTCAACTCCCAGAAA

AGCAGAGGAGGAAGATTCCAAGGCGGGAAAACCTTACGAGTCCCGGAAGTACCCGATGTCAAGAACTCCGGACCATCAGC

CCAATCAATTAAAAAGGGCACAGACGGGAACTCAGTCTCATCTGGAACGGTGACAGAGTGTTTATCGATAAGTGGTGCAA

CCCAAGCTGTGCCAGAGTCAAGATGGGAGTCATCAGAGCAAAATGCGTCTGTGGGGAGTGTCCTCAAGTCTGCGAAGAGT

GCAAAGACGATCCAGGGGTCGACACAAGAATCTGGTACCACAGCATCACCGACTCAGCCTAAAGAGAATGACTCCGAGTA

TGAGTATGAGGATGACCTATTTACAGAGATTCAGGACATCCGTGCAAGCATTGCCAAGATCCATGATGACAACAAGACTA

TCCTCTCAAAGCTTGATTCTATACTGTTATTGAAAGGAGAAGTCGACACTATCAAGAAACAAATCAGCAAGCAGAATATA

AGTATATCCACCATTGAGGGCCATCTCTCCAGTATAATGATAGCCATCCCGGGCTTTGGGAAGGATATCAAGGACCCAAC

ATCCGAGGTTGAGTTGAACCCAGATTTAAGACCTATAATAAGCCGTGACTCTGGCAGAGCTCTCGCGGAGGTCCTCAAGA

AACCCGCTGTGGATAGGTCTCAGAAAACTGAAACCAAAGCCAACTCCAGCTCAAAGGGTCAGCTTCTTAAGGATCTCCAG

CTAAAACCTGTCGACAAGCAGGCAAGCTCTGCAATCGGGTTTGTCCCATCCGACCATGAATCATCCAGAAATGTCATCCG

CTCCATAATCAAGTCGAGCAAGCTAAACATTGATCACAAGGACTATCTTCTAGATTTACTGAATGATGTGAAAGGCTCAA

AGGATCTTAAGGAATTCCACAAGATGCTAACAGCAATTCTCGCCAAGCACCCGTAACACATCCCCCAGTCACCATCTCAT

ACTCGGCTGAAGACATCCTTTCAATCAGACTATTACAAAAAACTTAGGAGCAAGGGCAACTGAGCTTCGCAGACAAGATG

ACCGAGATCTACGACTTCGACAAATCAGCATGGGATGTCAAAGGGTCAATTGCCCGCATAGAACCCACCACCTACCACGA

CGGCCGACTGATACCCCAGGTAAGGGTCATCGATCCTGGTCTGGGAGACAGAAAAGATGAGTGCTTTATGTACCTGTTTC

TCCTGGGAGTGATTGAGGATAACGACCCCCTGTCTCCCCCCGTCGGAAGAACCTTTGGCTCTTTACCTCTAGGGGTCGGT

AGGTCAACTGCTAAGCCAGAAGAACTACTAAGGGAGGCCACAGAACTAGACATAGTGGTGAGGCGCACTGCAGGAGTAAA

TGAGAAACTGGTATTTTACAACAACACTCCGCTGTCCTTGTTAACGCCCTGGAAGAAAGTTCTGACAACCGGAAGTGTGT

TTAGCGCTAACCAGGTTTGCAATGCAGTCAACCTAGTCCCGCTTGATACTCCCCAGAGGTTCAGGGTTGTGTACATGAGC

ATAACTAGATTATCAGACAATGGTTACTATAGTGTGCCCAGGAGAATGCTAGAGTTCCGCTCAGCCAATGCAGTAGCCTT

CAACATCTTGGTTACACTAAGAATTGAGAATGGCACAAACCCTAGAAGATACATAGTCGGCTCATGGGAGAATTCAGAGG

TCACATTTATGGTACACGTGGGCAACTTTAGAAGAAAGAAGAACGAAGTATACTCTGCTGATTACTGCAAAATGAAGATT

GAAAAGATGGGTTTAGTTTTTGCCCTGGGAGGAATAGGTGGAACAAGTCTCCATATTAGAAGCACCGGGAAAATGAGCAA

AACCCTCCATGCACAGCTGGGGTTCAAGAAAATCTTATGTTACCCCCTAATGGATGTTAATGAGGATCTTAACCGATATC

TCTGGCGGGCAGAGTGCCGAATAGTCAAAATCCAGGCCGTCTTACAGCCATCAGTGCCCCAAGAATTCCGTGTCTACGAT

GATGTCATCATCAACGATGACCAAGGCTTGTTCAAGATCCTGTAGTTCATCTGCAAAGGAACGAACCCACCCCCAGGGGG

GGGTCC------CCCCCG-CCAAAAAACACACTCCAAGGCTCCCCGACACAGCGCCAACACAGCCACCCTCCCCACTGGA

CGAGGCCGGCC-CCCCAACAAACCCGGTCCAGGC-GGGGGGGGGCCCCCCTGACCCGCCGGGCGCCGACCGAGCAGGCCG

GCGAGAAGCCCCCAAGAGGCCCCGACGAAGACCGAGCACCCCTCCCCCCC-AAACCGTYCCCCC-AACACGCCCGAGGCC

AGGCGCCGGGCACCCCCAGACCCGCCACCGGCGGGACAGAACCCCAGGGGGGAACCCCCCAAACCAAAGGGACCACCGGG

GAAGGATCCGCCGAACCAGCACAGACCCGACCCAAGCAAGGCGGACCCCCAACATGAAACCGCCCAGCACACCCCACTCA

TCATCAAACAAAACTTAGGGGCCAAGCCCATACGACAGGCCCCAGTGCGCCAGGCGGAGGGACGCAGAAAGGAAGGAGAC

AAGACCGCCCCAGAAGGAGGACACCCAACCACCGAAACACCCAGCCCCAGGGAAGGAG--CCCCCCCCCACATGGCGAAA

GAACGGGCAAGAGACCCCGAGAAGAACCCGACCGGGAGCCAGGAGCCGGCGACGCCGGCGGCAAACAAGCCGGGCAAACA

CCCGACCCCACCAGAGGAGCCGAGGAGAGCCCGACCCCGCACGAACCAAACAGCAGGGAGCCCAACCCCACCAAAAAAAG

AGGCGAGGCCCC-CCT-----CCTCCCCGCACACAACCCGCAGACAGACCGAGAGAGGACCACCTCCCAACAAGACAGGC

GGCTCCCTCGCAGCCGAGCCGCAGCTCCCCAGAACAGGACACCCCCGTACACTCAAAACACCCCCCCGCCCCCCGCCACC

GGGACAGGGGCTCCGCCACCACAGACCAAAGCCACCGAGAGCAAGCCAGAGGTCCCCCCCAGTCCCGCCCGGCGCCCCCC

CGAAAAGGCCCCGGCAAGGCCCCCCAACCCGACGTCCCCCGCACGCACCGAAAGAGGGACCCGCCCGAGAACGTGCGGCA

CATCCATGCGTAAACATTATGACACGGGTTGCAATCTTGACATCTCTGTTTCTTCTCCCAAATGTTGTTGCGTGCCAGAT

TCACTGGGGCAATCTATCCAAGATCGGGATTGTGGGAACAGGGAGTGCCAGCTACAAGGTGATGACTAGGCCGAGCCACC

AGACTCTGGTTATAAAGTTAATGCCAAATATAACAGCCATCGACAATTGTACAAAGTCAGAGATTGCAGAGTACAAGAGA

TTGCTGATCACAGTGTTAAAGCCTGTAGAGGATGCTCTGTCGGTTATAACCAAGAATGTAAGACCAATTCAAACCCTGAC

ACCTGGGCGTAGAACCCGTCGTTTTGCTGGAGCTGTTCTGGCCGGAGTAGCGCTTGGAGTTGCGACAGCCGCTCAGATAA

CTGCAGGAGTCGCTCTTCATCAATCATTGATGAACTCCCAAGCAATTGAGAGTTTAAAAACCAGTCTTGAGAAGTCGAAT

CAGGCAATAGAAGAAATCAGACTTGCAAATAAGGAGACCATACTGGCAGTACAGGGCGTCCAGGATTATATCAACAATGA

GCTTGTCCCTTCTGTTCATAGAATGTCATGCGAGCTGGTAGGCCACAAGCTCGGCCTCAAGCTCCTTAGGTACTACACCG

AGATCCTGTCCATATTCGGGCCCAGCCTTCGAGACCCGATAGCTGCCGAAATATCAATCCAGGCACTCAGTTATGCATTA

GGCGGAGACATTAATAAAATCCTGGACAAGCTTGGGTATAGCGGTGGGGATTTCCTTGCCATCCTAGAAAGCAAGGGAAT

AAAGGCCCGGGTTACATATGTGGACACAAGAGATTACTTTATAATGCTTAGCATCGCCTACCCAACCTTATCTGAGATCA

AGGGAGTGATAGTTCACAAGATAGAAGCTATAACATACAACATTGGGGCACAGGAGTGGTATACTACTATCCCTAAATAT

GTAGCCACTCAGGGATATCTGATATCGAACTTTGATGAGACGTCATGCGTATTCACTCCAGAGGGGACAGTTTGCAGCCA

GAATGCATTGTACCCAATGAGCCCATTGCTTCAGGAATGTTTCAGGGGGTCAACAAAATCGTGCGCCAGAACCCTAGTTT

CAGGTACCATAAGTAATAGATTTATCCTATCAAAAGGAAACCTGATCGCGAATTGTGCGTCAGTTTTGTGCAAATGTTAC

ACAACGGAGACAGTCATCAGCCAAGATCCTGACAAACTACTAACTGTTGTAGCATCCGACAAGTGTCCTGTAGTTGAGGT

GGATGGAGTGACAATACAGGTCGGCAGTCGAGAGTATCCGGATTCTGTATACTTACACAAAATAGACTTAGGCCCAGCCA

TCTCCCTAGAAAAACTAGATGTAGGTACCAATTTAGGCAATGCAGTTACAAGACTGGAGAATGCAAAGGAGCTCCTAGAT

GCATCAGACCAAATACTGAAGACTGTCAAAGGGGCACCTTTTGGTGGGAACATGTACATAGCACTGGCAGCTTGCATTGG

AGTATCCCTAGGGCTTGTCACATTAATATGCTGCTGTAAGGGGAGGTGTAAGAACAAGGAAATCCCTATCTCCAAAATCA

ACCCAGGGCTCAAACCCGATCTGACCGGGACCTCAAAGTCGTACGTGAGATCACTGTAGCCAGAATCACCTGAATCATCT

GGCATCACACACATACATGTACGACACAAGCAGTCTGAGGACGCAGGAAACCCAGCCCCCGATCACCCACCTGACCCTAC

TCCACGCTCCACTACACATTAGTCATTAAACAAAACTTAGGACGAAAGGTCAATCACCATGTCCGCACAAAGGGAAAGGA

TCAATGCCTTCTACAAAGACAATCCTCACAATAAGAACCATAGGGTAATCCTGGATAGAGAACGCTTGATCATTGAAAGA

CCCTACATCTTGCTTGGAGCCCTGCTGGTAATGTTCCTGAGTCTAATCGGACTGCTGGCCATTGCAGGGATCAGGCTTCA

CCGGGCCACCGTTGGAACTTCAGAGATCCAGAGTCGGCTGAATACCAATATTAAGTTGACCGAATCTATTGATCACCAGA

CTAAGGATGTCTTAACTCCCCTCTTTAAAATAATTGGCGATGAAGTCGGCATCAGGATTCCACAGAAATTCAGTGATCTT

GTCAAGTTCATCTCCGACAAGATTAAATTCCTCAACCCCGATAGAGAGTATGATTTCAGGGATCTCCGGTGGTGCATGAG

TCCCCCCGAGAGAGTCAAAATTAATTTTGATCAGTTTTGTGAGTACAAAGCTGCGGTTAATTCAATTGAACATATATTTG

AGTCACCGCTCAACAAGTCAAAAAAGCTGCAATCATTGACTCTCGGGCCCGGAACAAGTTGTCTAGGCAGGACAGTAACA

AGAGCTCATTTCTCAGAACTTACACTGACCTTAATGGACTTGGATCTAGAGATGAAGCACAACGTGTCCTCAGTGTTTAC

CGTAGTTGAAGAGGGGTTATTCGGAAGAACATATACCGTCTGGAGATCCGATGCCAGGGATCCGAGCACCGATCTAGGTA

TCGGCCATTTTTTAAGAGTCTTCGAGATTGGACTGGTAAGAGATCTCGGGCTGGGTCCCCCCGTTTTTCATATGACTAAC

TATCTCACAGTGAACATGAGTGATGTCTATCGGAGATGTCTTTTAGCGGTAGGGGAGTTGAAGTTGACAGCCCTATGCAC

CTCATCTGAGACTGTAACACTGAGTGAGAGAGGAGTTCCAAAGAGGGAGCCTCTTGTAGTTGTGATACTTAACCTGGCTG

GACCAACTCTAGGGGGCGAACTATACAGTATCTTGCCTACCTCTGATCTCATGGTGGAGAAACTCTATTTATCTTCACAT

AGAGGGATCATCAAGGATGACGAGGCCAATTGGGTAGTGCCGTCTACCGATGTTCGTGATCTTCAGAACAAAGGTGAGTG

TCTAGTGGAAGCATGCAAGACTCGACCTCCTTCATTTTGCAATGGCACAGGATCAGGCCCGTGGTCAGAGGGGAGAATCC

CTGCTTACGGGGTGATCAGGGTCAGTCTTGACTTAGCTAGTGACCCGGATGTAGTTATCACTTCAGTGTTTGGCCCACTG

ATTCCTCACCTATCCGGCATGGATCTTTACAACAACCCGTTTTCAAGAGCTGTATGGTTGGCTGTACCACCTTACGAGCA

GTCATTTCTAGGAATGATAAATACAATTGGATTCCCTAACAGAACAGAGGTTATGCCGCACATTTTGACCACAGAGATCA

GAGGCCCTCGGGGTCATTGCCATGTCCCCATAGAATTGTCCCGCAGGGTTGATGACGATATCAAGATCGGGTCCAACATG

GTCATATTGCCGACGATGGACCTGAGATATATTACAGCCACTTATGATGTTTCCAGGAGCGAGCATGCAATCGTGTACTA

TATCTATGACACAGGTCGCTCATCATCTTACTTCTACCCAGTTCGACTGAATTTCAAAGGCAATCCTCTCTCTCTGAGGA

TAGAGTGTTTCCCTTGGCGTCATAAGGTGTGGTGCTACCATGATTGTCTTATATACAACACCATAACAGATGAAGAGGTC

CATATGAGAGGGCTGACCGGTATAGAGGTAACATGCAATCCAGTCTGAGCAGAGCTACGACCGTCACTCAAGCAGTCCCT

TGAGTCGCCACCGAGTCCAAGCAGCACAGCCTGGGACTCTCAACAGCACAGCCCAGCTAACAGTGTTATAAAAAACTTAG

GAGCCAGGGTTGTAGGGGCCATGGACTCCCTATCAGTCAATCAGGTCTTGTACCCTGAGGTTCATCTAGACAGCCCTATT

GTCACAAACAAACTAGTTGCCATCCTTGAGTACTCGGGTATCGACCATAACTATGTTCTTGAAGACCAGACCCTTGTCAA

GAATATTAGGTATAGACTGGGGTGCGGTTTTTCAAATCAAATGATCATCAATAACAGGGGGGTGGGTGAAACAGTCAATT

CTAAACTTAAAAGTTATCCCCATAATCGTCATATCATATACCCGGATTGCAATAAGGAGTTGTTTTGTATCAAAGATAGC

TGCATATCTAAGAAGCTCTCGGAGCTATTCAAGAAGGGTAATTCCTTGTACTCTAAGATAAGTCACCAGGTACTGGACTG

TCTTAAGAGAGTCAATGGGAAATTAGGCCTGGGCACAGATCTTACCCATGGTCTGAAGGAGGGTATCCTTGACCTAGGGT

TGCACATGCATAGCTCTCAATGGTTCGAGACTTTCCTGTTCTGGTTCACTATCAAGACAGAGATGAGATCAATGATCAAA

GAACAGTCCCATATATGCCACAAGAGGAGATATAACCCGATTTTTGTATCAGGGGATGCATTCGAGGTGCTCGTATCACG

AGACCTCGTTGTGATAATTGATAAGAATACCCAGTATGTCTTCTACTTAACATTTGAGCTGGTCCTTATGTATTGTGATG

TCATAGAGGGCAGACTTATGACGGAGACAGCCATGGTCATAGACCAGAGATATTCAGAACTTCTAAGCCGGGTTAGATAC

CTGTGGGATCTTATTGATGGGTTCTTCCCAACACTAGGCAATACCACCTACCAAATTGTTGCTCTGCTTGAACCATTATC

GTTGGCTTATCTTCAACTTCAGGATGTCACGCTGGAGTTAAGAGGTGCTTTTTTAGACCATTGCTTCAAAGAACTCTATG

AGATACTGGAGCATTGCGGCATTGACACAGAAGGCACCTATAATTCCATCACTGAAGGATTGGATTACGTATTCATCACC

CACGATATACATCTAACTGGGGAGATTTTTTCATTTTTCCGGAGTTTCGGACACCCCCGTCTAGAAGCAGTCACCGCCGC

AGAAAATGTCAGGAAACATATGAATCAACCGAAGGTAATCAGTTATGAGACTATGATGAAAGGACATGCGGTATTCTGCG

GGATAATCATCAATGGTTTTAGAGACCGACATGGCGGCAGCTGGCCCCCTGTTGCATTGCCAGAGCATGCTTCTGCTGCG

ATCCGGAATGCGCAAGCATCAGGTGAGGGACTGACCCATGACCTGTGTATAGACAACTGGAAATCCTTTGTTGGATTCAA

ATTTGGCTGCTTTATGCCACTCAGCCTAGACAGTGATTTGACCATGTATCTCAAAGACAAGGCATTGGCTGCACTGAAGA

ATGAGTGGGATTCAGTTTACCCGAAAGAATACCTCCGTTATAACCCACCTAGAGGGACAGAGTCAAGGCGACTGGTGGAG

GTGTTCCTGAATGACTCCAGCTTCGATCCTTATAACATGATAATGTACGTGGTGAATGGCTCCTATCTTAAAGACCCTGA

GTTTAATCTCTCATACAGCCTAAAAGAGAAGGAGATAAAGGAGACAGGGCGGTTGTTTGCCAAAATGACCTATAAGATGC

GGGCCTGTCAGGTAATTGCTGAGAATCTGATATCGAATGGTGTTGGGAAGTACTTCCGAGACAATGGGATGGCGAAGGAC

GAGCATGACCTAACAAAAGCCCTTCACACTTTGGCAGTCTCAGGTGTTCCCAAGAACAACAAAGATAACCGCCGAGGTGG

GCCTCCCAGAAGGACAACAACCCGAGAGGTGAGATCAAGCCGGGACACCAGAACACAAAATAGAGACAAGGTCCAAGGGG

GACCTATGTACAACTATTTGCAATGCCAACCAATCAGCCCTGATCAAGGTGAGTCATACGAGACTGTTAGTGCATTTATC

ACTGCTGACCTTAAGAAGTATTGCCTGAATTGGAGATACGAGACAATCAGCATATTTGCACAGAGGCTGAATGAAATATA

TGGACTACCATCCTTCTTTCAATGGTTGCACAGGATATTGGAAAAATCCGTACTCTACGTCAGTGACCCACATTGCCCTC

CCGATCTAGATAATCATATCCCTCTGGACAGTGTCCCTAATGCCCAAATATTCATTAAGTACCCAATGGGTGGAATAGAA

GGTTATTGCCAAAAACTATGGACAATCAGTACTATACCATACTTGTATCTGGCAGCCTATGAGAGCGGAGTAAGAATCGC

CTCACTGGTGCAGGGTGACAATCAGACAATCGCAGTGACAAAAAGAGTTCCAAGTTCTTGGCCTTATTCACTAAAAAAGA

GGGAGGCATCCAAAGCCGCTCAAAATTACTTCGTAGTCTTAAGGCAAAGATTGCACGATGTAGGTCATCACTTGAAGGCT

AATGAAACCATAGTATCTTCTCACTTTTTTGTATATTCCAAAGGGATTTATTATGACGGCCTGCTAGTCTCACAATCACT

AAAGAGCATCGCCAGATGTGTCTTCTGGTCCGAGACTATAGTGGATGAAACCAGAGCGGCTTGCAGCAATATCGCAACAA

CTGTCGCTAAGAGTATAGAGAGGGGTTATGATAGGTACCTTGCATATTCTTTGAATATCCTCAAGATTTTCCAACAGATC

CTTATATCCCTTAACTTCACTATTAACACAACAATGACTCAGGATGTCGTGGCACCGATCGTCGAGAACGGTGATTTGCT

GATAAGGATGGCACTCTTGCCAGCACCCATCGGAGGTCTAAATTATCTTAACATGAGCAGGCTATTCGTGAGAAATATCG

GCGACCCGGTCACCTCCTCTATAGCCGACCTGAAGAGGATGATAGATGCCGGGCTCATGCCAGAAGAAACATTGCATCAA

GTGATGACCCAGACCCCGGGAGAATCATCCTACCTTGATTGGGCAAGTGACCCTTATTCTGCCAACCTACCCTGCGTACA

GAGTATAACTCGTCTTCTCAAAAACATCACTGCACGATATATTTTAATCAGCAGCCCAAATCCGATGCTGAAAGGATTGT

TTCATGAGGGGAGTAGAGATGAAGACGAGGAGCTTGCAAGTTTCCTAATGGATCGGCATATAATTGTTCCAAGAGCTGCA

CATGAAATCTTAGACCATAGTATAACCGGGGCAAGAGAGGCTATAGCCGGGATGTTGGACACCACCAAGGGTCTGATTAG

AACAAGTATGAAACGGGGTGGCCTCACCCCTCGAGTCTTAGCCCGCCTTTCCAATTATGATTATGAACAATTCCGATCCG

GGATAACACTATTGACAAAGAAAGGGCAGTGTTATCTCATTGACAAGGATTCGTGCTCGGTGCAGCTCGCTATAGCTCTG

AGGAGCCACATGTGGGCTAGGTTGGCTCGCGGGAGACCAATCTATGGGTTGGAGGTGCCCGATATATTAGAATCGATGAA

CGGCTACCTTATCAAACGCCACGAGTCTTGTGCTATCTGTGAAACGGGCTCAAGTCACTACGGATGGTTTTTCGTCCCTG

CAGGGTGCCAGCTTGACGATGTATCTAGAGAGACTTCGGCTCTCCGTGTACCTTATGTCGGATCAACAACTGAGGAAAGG

ACTGACATGAAACTTGCATTTGTTAGATCTCCAAGTAGATCCCTCAAATCAGCAGTTAGAATTGCAACAGTTTACTCATG

GGCCTACGGGGATGATGAGAAGTCATGGAGTGAAGCCTGGATGCTAGCTAGGCAGAGAGCCAATATCACCTTAGATGAAT

TGAGAATGATCACTCCAGTCTCTACATCCACCAACCTAGCCCACCGGTTGAGGGATCGGAGCACCCAGGTGAAATACTCG

GGGACATCCCTTGTGAGGGTTGCAAGATACACAACCATCTCCAATGATAACTTGTCATTTGTGATATCTGAGAAGAAAGT

AGATACCAACTTCATTTATCAGCAAGGGATGTTGCTTGGTCTCGGGATCCTTGAGAACCTCTTTAGGCTAGAGGCCACCA

CCGGAGTATCCAACACAGTGCTACACCTGCACGTGGAAACAGAATGTTGTGTTGTACCTATGGTGGATCACCCAAGGATA

CCGAGTCTCCGTAATATAAAAGTTACGGATGAGCTATGCACAAACCCTTTGATCTATGATAGGTCTCCCATCATAGAGCA

CGATGCAACCCGACTATACTCACAGAGCCACAGGAGACATTTGGTGGAGTTTGTTACCTGGTCCACAAGCCAGCTTTACC

ATATACTGGCTAAATCTACAGCAATGTCCATGATTGAATTGATCACGAGATTCGAGAAAGATCACATGAATGAAATAGCC

GCCCTGATTGGCGATGACGACATCAACAGCTTCATCACAGAATTTTTGCTAGTGGAGCCCAGATTATTTATAGTTTACCT

TGGTCAGTGTGCTGCCATCAATTGGGCCTTTGATATACATTATCATCGGCCCTCGGGCAAGTACCAGATGGGGGAACTCC

TCTACTCTCTGCTCTCACGGATGAGCAAAGGAGTATATAAGATCTTCACCAATGCTTTGAGCCACCCCAAAGTTTACAAG

AAATTTTGGCGAAGTGGTGTCATTGAGCCGATTCATGGCCCATCCCTGGATACACAGAATTTACATGTCACTGTTTGTGA

CATGATATATGGGTCCTATGTCACCTACCTGGATCTTTTGCTGAATGATGAGCTAGATGATTACCCGTATTTGCTCTGCG

AGAGTGATGAGGATGTGGTCACAGACAGGTTCGACAACATTCAAGCCAAACATCTCTGTGTATTAGCCGATGTATATTGC

AGCTCCAAGAGGTGTCCTTCCATAATCGGGATGTCTCCTATAGAAAAATGTACCATTCTCACACATTACATCAAGGGAGA

ATCAGTACAATCCCCGTCTGGAACCTCATGGAACACTGATCCCCTTGTAGTAGATCATTACTCATGCTCTCTGACCTACC

TTCGCCGAGGTTCCATCAAACAAATCAGGTTGAGGGTGGATCCTGGGTTTGTATTCGAGGCGTTGACAGACGTCGACTTT

AAACAACCTCGCAAAGCTAAGTTAGATATATCGGTCGTTGGGTTGACTGATTTCTCTCCCCCTTGCGACAACGTCGGTGA

TTTTCTAGGGACTATCAACACATTGAGGCACAATCTGCCTGTCACCGGAACCGGGGTCTCGAACTATGAAGTCCACGCTT

ATCGTAGAATTGGCCTGAATTCATCAGCATGTTACAAAGCCGTAGAGATCTCCACGTTAATCAAGCCATCCCTAGAAGTC

GGAGAGCATGGATTGTTCTTAGGAGAAGGTTCCGGTTCAATGCTGGCTGCGTACAAGGAAGTTCTTAAATTAGCAAATTG

TTATTACAACAGCGGGGTAACAGCGGAGGGCAGAGCCGGACAGAGGGAAATATCTCCCTATCCTTCAGAGATGAGCCTAG

TAGAGAATCAAATGGGGATAGAGCGGAGTGTTAAAGTGCTGTTCAACGGCAAACCTGAAGTAACTTGGGTGGGGACCACC

GATTGCTACAAGTACATAATCAGTAACATCCAAACCTCTAGTCTGGGTTTCATACATTCAGATATTGAGACGCTCCCAAC

CAAAGATGCTGTTGAGAAATTAGAAGAATTTGCCTCTATCCTATCCTTATCCCTGATTTTGGGGAAAATCGGCTCTATTA

CAGTTGTCAAAATTATGCCCATTAGCGGGGATTTCACCCAAGGCTTCATAGCCTATGCCATCCAATATTTTAGGGAGAGC

CTGCTTGCCTATCCGAGATATAGTAACTTTATCTCGACTGAGTGTTACCTTATCATGATAGGATTAAAGGCCAATCGATT

GATTAACCCAGAAGCCATTAAGCAAAGCATAATCAGAGCGGGGGTCAGGACTGCACCAGGACTTGTGAGCCATATATTAT

CAGAGAAACAGAAAGGTTGTATTCAATCTTTTTTGGGTGATCCCTATATCCAAGGAGACTTCAATAAGCACCTTAAATCT

CTAACCCCTATTGAGAAAATCCTGGTAAATTGTGGTCTCTCGATCAATGGCACAAAAATCTGTAGGGATCTAATCCACCA

TGATATCGCCTCCGGTCCAGACGGTCTTATGAGCTCTACAATTATTTTATATAGGGAATTGGCTCATTTCAAAGATAATA

TAAGGAGTCAGCACGGAATGTTCCACCCCTACCCAGTACTGGCCAATAGCCGGCAACGTGAATTAATCCATCGAATAGCC

AAGAAATTCTGGGGGTATGTCTTGCTATATTCTGATGACCCGGCACTAATCAGACAAACAATCAAGAACTTGAAACGGAA

TCATCTAACCTTTGACTTACATAGTAACCCGTTTATTAAAGGGCTATCCAAAGCTGAGAAACTGCTGGTGCGGACGAGTT

CACTAAGAAGGGAATGGTTGTTCACTCTCGATACGAAAGAAGTGAAAGAGTGGTTCAAATTGGTAGGTTACAGTGCACTC

ATCAGAGGCTAATTGGTTATACATCTGCCCCCTTCTCCTCCGCCATGAGACCCCACTGACGATTCAGAAGATTAAAGAAA

ACTACATATTGGATAAGTATCTATTCCCAGCTTTGTCTGGT

>KT860065.1/sheep/India/2015-07-10

ACCAAACAAAGTTGGGTAAGGATAGATCTTATAATAACTATGGACTAGCAAACTTAGGAGTAAAGATCCTACTGTCGGGG

GGAGGAGGAGGAGCAAGATCTCTGATTATGGCGACTCTCCTTAAAAGCTTAGCATTGTTCAAAAGGAACAAAGACAAAGC

GCCGACGGCATCAGGTTCAGGAGGGGCCATCCGAGGGATTAAGAATGTTATCATAGTCCCGATTCCCGGGGACTCGTCCA

TCACTACCCGTTCAAGACTGCTCGACAGGCTTGTCAGATTGGCCGGAGATCCTGACATCAACGGGTCAAAGCTGACCGGC

GTGATGATCAGCATGTTATCTTTGTTCGTAGAGTCACCCGGGCAGTTGATACAGCGAATCACAGATGATCCGGATGTTAG

TATCCGCCTTGTTGAGGTAGTTCAAAGTACTAGGTCCCAGTCCGGGTTGACCTTTGCATCACGTGGTGCTGATTTGGATA

ACGAGGCAGACATGTATTTTTCAACTGAGGGGCCCTCGAGTGGAGGTAAGAAAAGGATCAACTGGTTTGAGAACAGAGAA

ATAATAGACATAGAGGTGCAGGATCCAGAAGAGTTCAATATGTTGTTAGCCTCCATACTAGCACAAGTCTGGATCCTTCT

GGCCAAGGCTGTCACGGCACCGGATACGGCAGCTGACTCAGAACTGAGAAGGTGGGTTAAATACACACAACAAAGGAGAG

TGATTGGGGAATTTCGCCTTGACAAAGGGTGGCTGGACGCAGTCCGCAACAGAATTGCAGAAGATCTATCACTCCGGCGG

TTCATGGTATCTCTCATACTTGACATCAAGAGGACCCCCGGCAACAAGCCAAGGATCGCAGAAATGATCTGCGACATTGA

CAACTATATTGTCGAAGCAGGACTCGCCAGTTTCATCCTTACTATCAAATTTGGTATAGAAACCATGTATCCTGCATTAG

GGCTCCACGAGTTTGCCGGGGAATTGTCCACTATTGAGTCCTTGATGAACTTGTATCAACAGCTAGGAGAGGTTGCACCC

TACATGGTAATTCTAGAGAACTCAATTCAGAACAAGTTTAGTGCAGGAGCTTATCCCCTCCTCTGGAGCTATGCGATGGG

TGTCGGAGTTGAGCTGGAGAACTCAATGGGGGGCTTGAACTTTGGCAGATCATATTTTGACCCTGCCTATTTTCGTCTCG

GACAGGAGATGGTCAGAAGATCTGCAGGAAAGGTCAGCTCTGTGATTGCGGCTGAGCTCGGCATCACAGCAGAGGAAGCT

AAACTAGTCTCGGAAATAGCCTCGCAGGCTGGGGACGAAAGAACCGCTAGAGGGACTGGGCCTCGACAGGCGCAGGTCTC

CTTCCTACAGCACCAAACAGGAGGGGGAGAGTCGTCCGCACCAGCGACCAGAGAAGGGGTCAAAGCTGCGATCCCAAACG

GATCCGAAGAAAGGGACAGAAAGCAAACACGCCCAGGGAGGCCCAGAGGAGAGACCCCCGGCCAACTGCTCCTGGAATTC

ATGCCAGAGGATGAGGTCTCGCGAGAATCTGGTCAAAACCCTCGTGAGGCTCAAAGGTCGGCCGAGGCACTCTTCAGGCT

GCAGGCCATGGCCAAGATTCTGGAGGACCAGGAGGAGGGAGAAGACGACAATCAGGTCTACAACGACAAGGATCTCCTCG

GCTAAGCAGACGCACCCTCTGTCGAAATCAGTGACAAGACATCTCCTACCAGTATTATAAAAAACTTAGGACCCAGGTCC

AAGCAAGCACACATCGACACTTCAACCAGTCGAGCGGAGGCCACCGATGGCAGAAGAACAAGCATACCATGTCAACAAGG

GGCTGGAATGTATCAAGTCCCTCAAAGCCTCTCCCCCGGATCTATCCACCATTAGAGATACCATCGAGAGCTGGAGAGAG

GGGCTTAGCCCCTCGGGCCGTGCAACACCGAACCCTGATACGTCCGAGGGAGATCATCAGAATATCAACCAATCATGCTC

ACCAGCAATCGGACCAAACAAAGTCTACTTGTCTCCTGAAGATAATCTCGGATTTAGAGAGATCACTGGCAACGACTGTG

AGGCTGAGCTCGGAGGAGCCCAGGGAAAAGGATCCAACTCTCAAGTACAGCGTTACTATGTTTATAGCCACGGGGGTGAA

GAGATTGAAGGACTCGAGGATGCTGACTCTCTCGTGGTTCAAGCAGATCCTCCAGTTGCTAACATATTCAATGGAGGAGA

GGATGGATCTGACGACAGCGATGTGGACTCTGGCCCAGATGATCCCAGCAGAGATACTCTATATGACCGGGGACCTGCTG

CCGGCAATGATGTCGCTAGGTCCACAGATGTCGAAAAACTAGAAGGTGCCGATATTCAAGAAGTTCTCAACTCCCAGAAA

AGCAGAGGAGGAAGATTCCAAGGCGGGAAAACCTTACGAGTCCCGGAAGTACCCGATGTCAAGAACTCCAGACCATCAGC

CCAATCAATTAAAAAGGGCACAGACGGGAACTCAGTCTCATCTGGAACGGTGACAGAGTGTTTATCGATAAGTGGTGCAA

CCCAAGCTGTGCCAGAGTCAAGATGGGAGTCATCAGAGCAAAATGCGTCTGTGGGGAGTGTCCTCAAGTCTGCGAAGAGT

GCAAAGACGATCCAGGGGTCGACACAAGAATCTGGTACCACAGCATCACCGACTCAACCTAAAGAGAATGACTCCGAGTA

TGAGTATGAGGATGACCTATTTACAGAGATTCAGGACATCCGTGCAAGCATTGCCAAGATCCATGATGACAACAAGACTA

TCCTCTCAAAGCTTGATTCTATACTGTTATTGAAAGGAGAAGTCGACACTATCAAGAAACAAATCAGCAAGCAGAATATA

AGTATATCCACCATTGAGGGCCATCTCTCCAGTATAATGATAGCCATCCCGGGCTTTGGGAAGGATATCAAGGACCCAAC

ATCCGAGGTTGAGTTGAACCCAGATTTAAGACCTATAATAAGCCGTGACTCTGGCAGAGCTCTCGCGGAGGTCCTCAAGA

AACCCGCTGTGGATAGGTCTCAGAAAACTGGAACCAAAGCCAACTCCAGCTCAAAGGGTCAGCTTCTTAAGGATCTCCAG

CTAAAACCTGTCGACAAGCAGGCAAGCTCTGCAATCGGGTTTGTCCCGTCCGACCATGAATCATCCAGAAATGTCATCCG

CTCCATAATCAAGTCGAGCAAGCTAAACATTGATCACAAGGACTATCTTCTAGATTTACTGAATGATGTGAAAGGCTCCA

AGGATCTTAAGGAATTCCACAAGATGCTAACAGCAATTCTCGCCAAGCACCCGTAACACATCTCCCAGTCACCATCTCAT

ACTCGGCTGAAGACATCCTTTCAATCAGGCTATTACAAAAAACTTAGGAGCAAGGGCAACTGAGCTTCGCAGACAAGATG

ACCGAGATCTACGACTTCGACAAATCAGCATGGGATGTCAAAGGGTCAATTGCCCGCATAGAACCCACCACCTACCACGA

CGGCCGACTGATACCCCAGGTAAGAGTCATCGATCCTGGTCTGGGAGACAGAAAAGATGAGTGCTTTATGTACCTGTTTC

TCCTAGGAGTGATTGAGGATAACGACCCCCTGTCTCCCCCCGTCGGAAGAACCTTTGGCTCTTTACCTCTAGGGGTCGGT

AGGTCAACTGCTAAGCCAGAAGAACTACTAAGGGAGGCCACAGAACTAGACATAGTGGTGAGGCGCACTGCAGGAGTAAA

TGAGAAACTGGTATTTTACAACAACACTCCGCTGTCCTTGTTAACGCCCTGGAAGAAAGTTCTGACAACCGGAAGTGTGT

TTAGCGCTAACCAGGTTTGCAATGCAGTCAACCTAGTCCCGCTTGATACTCCCCAGAGGTTCAGGGTTGTGTACATGAGC

ATAACTAGATTATCAGACAATGGTTACTATAGTGTGCCCAGGAGAATGCTGGAGTTCCGCTCAGCCAATGCAGTAGCCTT

CAACATCTTGGTTACACTAAGAATTGAGAATGGCACAAACCCTAGAAGATACATAGTCGGCTCATGGGAGAATTCAGAGG

TCACATTTATGGTACACGTGGGCAACTTTAGAAGAAAGAAGAACGAAGTATACTCTGCTGATTACTGCAAAATGAAGATT

GAAAAGATGGGTTTAGTTTTTGCCCTGGGAGGAATAGGTGGAACAAGTCTCCATATTAGAAGCACCGGGAAAATGAGCAA

AACCCTCCATGCACAGCTGGGGTTCAAGAAAATCTTATGTTACCCCCTAATGGATGTTAATGAGGATCTTAACCGATATC

TCTGGCGGGCAGAGTGCCGAATAGTCAAAATCCAAGCCGTCTTACAGCCATCAGTGCCCCAAGAATTCCGTGTCTACGAT

GATGTCATCATCAACGATGACCAAGGCTTGTTCAAGATCCTGTAGTTCATCTGCAAAGGAACGAACCCACCCCCAGGAAG

GGGCCCTACCCTCCCCCG-CCAAAAAACACACCCCAAGGCTCCCCGACACAGCGCCAACACAGCCACTCTCCCGATGGGA

CGAGGTTGGCC-CCCCAACAAACCCGGTCCAGGC-GGGGGGCCCCCCCCCTGACCCGCCGGGCGCCGACCGAGCAGGCCG

GCGAGAAGCCCCCAAGAGGCCCCGACGAAGACCGAGCACCCCTCCCCCCC-AAACCGTTCCCCC-AACACGCCCGAGGCC

AGGCGCCGGGCACCCCCAGACCCGCCACCGGCGGGACAGAACCCCAGGGGGGGACCCCCCAAACCAAAGGGACCACCGGG

GAAGGATCCGCCGAACCAGCACAGACCCGACCCAAGCAAGGCGGACCCCCAACACGAAACCGCCCAGCACACCCCACTCA

TCATCAAACAAAACTTAGGGGCCAAGCCCATACGACAGGCCCCAGTGCGCCAGGCGGAGGGACGCAGAAAGGAAGGAGAC

AAGACCGCCCCAGAAGGAGGACACCCAACCACCGAAACACCCAGCCCCAGGGAAGGAG--CCCCCCCCCACATGGCGAAA

GAACGGGCGAGAGACCCCGAGAAGAACCCGACCGGGAGCCAGGAGCCGGCGACGCCGGCGGCAAACAAGCCGGGCAAACA

CCCGACCCCACCAGAGGAGCCGAGGAGAGCCCGACCCCGCACGAACCAAACAGCAGGGAGCCCAACCCCACCAAAAAAAG

AGGCGAGGCCCC-CCT-----CCTCCCCGCACACAACCCGCAGACAGACCGAGAGAGGACCACCTCCCAACAAGACAGGC

GGCTCCCTCGCAGCCGAGCCGCAGCTCCCCAGAACAGGACACCCCCGTACACTCAAAACACCCCCCCGCCCCCCGCCACC

GGGACAGGGGCTCCGCCACCACAGACCAAAGCCACCGAGAGCAAGCCAGAGGTCCCCCCCAGTCCCGCCCGGCGCCCCCC

CGAAAAGGCCCCGGCAAGGCCCCCCAACCCGACGTCCCCCGCACGCACCGAAAGAGGGACCCGCCCGAGAACGTGCGGCA

CATCCATGCGTAAACATTATGACACGGGTCGCAATCTTGACATCTCTGTTTCTTCTCCCAAATGTTGTTGCGTGCCAGAT

TCACTGGGGCAATCTATCCAAGATCGGGATTGTGGGAACAGGGAGTGCCAGCTACAAGGTGATGACTAGGCCGAGCCACC

AGACTCTGGTTATAAAGTTAATGCCAAATATAACAGCCATCGACAATTGTACAAAGTCAGAGATTGCAGAGTACAAGAGA

TTGCTGATCACAGTGTTAAAGCCTGTAGAGGATGCTCTGTCGGTGATAACCAAGAATGTAAGACCAATTCAAACTCTGAC

ACCTGGGCGTAGAACCCGTCGTTTTGCTGGAGCTGTTCTGGCCGGAGTAGCGCTTGGAGTTGCGACAGCCGCTCAGATAA

CTGCAGGAGTCGCTCTTCATCAATCATTGATGAACTCCCAAGCAATTGAGAGTTTAAAAACCAGTCTTGAGAAGTCGAAT

CAGGCAATAGAAGAAATCAGACTTGCAAATAAGGAGACCATACTGGCAGTACAGGGCGTCCAGGATTATATCAACAATGA

GCTTGTCCCTTCTGTTCATAGAATGTCATGCGAGCTGGTAGGCCACAAGCTCGGCCTCAAGCTCCTTAGGTACTACACCG

AGATCCTGTCCATATTCGGGCCCAGCCTTCGAGACCCGATAGCTGCCGAAATATCAATCCAGGCACTCAGTTATGCATTA

GGCGGAGACATTAATAAAATCCTGGACAAGCTTGGGTATAGCGGTGGGGATTTCCTTGCCATCCTAGAAAGCAAGGGAAT

AAAGGCCCGGGTTACATATGTGGACACAAGAGATTACTTTATAATGCTTAGCATCGCCTACCCAACCTTATCTGAGATCA

AGGGAGTGATAGTTCACAAGATAGAAGCTATAACATACAACATTGGGGCACAGGAGTGGTATACTACTATCCCTAAATAT

GTAGCCACTCAGGGATACCTGATATCGAACTTTGATGAGACGTCATGCGTATTCACTCCAGAGGGGACAGTTTGCAGCCA

GAATGCGTTGTACCCAATGAGCCCATTGCTTCAGGAATGTTTCAGGGGGTCAACAAAATCGTGCGCCAGAACCCTAGTTT

CAGGTACCATAAGTAATAGATTTATCCTATCAAAAGGGAACCTGATCGCAAATTGTGCGTCAGTTTTGTGCAAATGTTAC

ACAACGGAGACAGTCATCAGCCAAGATCCTGACAAACTACTAACTGTTGTAGCATCCGACAAGTGTCCTGTAGTTGAGGT

GGATGGAGTGACAATACAGGTCGGCAGTCGAGAGTATCCGGATTCTGTATACTTACACAAAATAGACTTAGGCCCAGCCA

TCTCCCTAGAAAAACTGGATGTAGGTACCAATTTAGGCAATGCAGTTACAAGACTGGAGAATGCAAAGGAGCTCCTAGAT

GCATCAGACCAAATACTGAAGACTGTCAAAGGGGCACCTTTTGGTGGGAACATGTACATAGCACTGGCAGCTTGCATTGG

AGTATCCCTAGGGCTTGTCACATTAATATGCTGCTGTAAGGGGAGGTGTAAGAACAAGGAAATCCCTATCTCCAAAATCA

ACCCAGGGCTCAAACCCGATCTGACCGGGACCTCAAAGTCGTACGTGAGATCACTGTAGCCAGAATCACCTGAATCATCT

GGCATCACACACATACATGTACGACACAAGCAGTCTGAGGACGCAGGAAACCCAGCCCCCGATCACCCACCTGACCCTAC

TCCACGCTCCACTACACATTAGTCATTAAACAAAACTTAGGACGAAAGGTCAATCACCATGTCCGCACAAAGGGAAAGGA

TCAATGCCTTCTACAAAGACAATCCTCACAATAAGAACCATAGGGTAATCCTGGATAGAGAACGCTTGATCATTGAAAGA

CCCTACATCTTGCTTGGAGCCCTGCTGGTAATGTTCCTGAGTCTAATCGGACTGCTGGCCATTGCAGGGATCAGGCTTCA

CCGGGCCACCGTTGGAACTTCAGAGATCCAGAGTCGGCTGAATACCAATATTAAGTTGACCGAATCTATTGATCACCAGA

CTAAGGATGTCTTAACTCCCCTCTTTAAAATAATTGGCGATGAAGTCGGCATCAGGATTCCACAGAAATTCAGTGATCTT

GTCAAGTTCATCTCCGACAAGATTAAATTCCTCAACCCCGATAGAGAGTATGATTTCAGGGATCTCCGGTGGTGCATGAG

TCCCCCCGAGAGAGTCAAAATTAATTTTGATCAGTTTTGTGAGTACAAAGCTGCGGTTAATTCAATTGAACATATATTTG

AGTCACCACTCAACAAGTCAAAAAAGCTGCAATCATTGACTCTCGGGCCCGGAACAAGTTGTCTAGGCAGGACAGTAACA

AGAGCTCATTTCTCAGAACTTACACTGACCTTGATGGACTTGGATCTAGAGATGAAGCACAACGTGTCCTCAGTGTTTAC

CGTAGTTGAAGAGGGGTTATTCGGAAGAACATATACCGTCTGGAGATCCGATGCCAGGGATCCGAGCACCGATCTAGGTA

TCGGCCATTTTTTAAGAGTCTTCGAGATTGGACTGGTAAGAGATCTCGGGCTGGGTCCCCCCGTTTTTCATATGACCAAC

TATCTCACAGTGAACATGAGTGATGTCTATCGGAGATGTATTTTAGCGGTAGGGGAGTTGAAGTTGACAGCCCTATGCAC

CTCATCTGAGACTGTAACACTGAGTGAGAGAGGAGTTCCAAAGAGGGAGCCTCTTGTAGTTGTGATACTTAACCTGGCTG

GACCAACTCTAGGGGGCGAACTATACAGTGTCTTGCCTACCTCTGATCTCATGGTGGAGAAACTCTATTTATCTTCACAT

AGAGGGATCATCAAGGATGACGAGGCCAATTGGGTAGTGCCGTCYACCGATGTTCGTGATCTTCAGAACAAAGGTGAGTG

TCTAGTGGAAGCATGCAAGACTCGACCTCCTTCATTTTGCAATGGCACAGGATCAGGCCCGTGGTCAGAGGGGAGAATCC

CTGCTTACGGGGTGATCAGGGTCAGTCTTGACTTAGCTAGTGACCCGGATGTAGTTATCACTTCAGTGTTTGGCCCACTG

ATTCCTCACCTATCCGGCATGGATCTTTACAACAACCCGTTTTCAAGAGCTGTATGGTTGGCTGTACCACCTTATGAGCA

GTCATTTCTAGGAATGATAAATACAATTGGATTCCCTAACAGAACAGAGGTTATGCCGCACATTTTGACCACAGAGATCA

GAGGCCCTCGGGGTCATTGCCATGTCCCCATAGAATTGTCCCGCAGGGTTGATGACGATATCAAGATCGGGTCCAACATG

GTCATATTGCCGACGATGGACCTGAGATATATTACAGCCACTTATGATGTTTCCAGGAGCGAGCATGCAATCGTGTACTA

TATCTATGACACAGGTCGCTCATCATCTTACTTCTACCCAGTTCGACTGAATTTCAAAGGCAATCCTCTCTCTCTGAGGA

TAGAGTGTTTCCCTTGGCGTCATAAGGTGTGGTGCTACCATGATTGTCTTATATACAACACCATAACAGATGAAGAGGTC

CATATGAGAGGGCTGACCGGTATAGAGGTAACATGCAATCCAGTCTGAGCAGAGCTACGACCGTCACTCAAGCAGTCCCG

TGAGTCGCCACCGAGTCCAAGCAGCACAGCCTGGGACTCTCAACAGCACAGCCCAGCCAACAGTGTTATAAAAAACTTAG

GAGCCAGGGTTGTAGGGGCCATGGACTCCCTATCAGTCAATCAGGTCTTGTACCCTGAGGTTCATCTAGACAGCCCTATT

GTCACAAACAAACTAGTTGCCATCCTTGAGTACTCGGGTATCGACCATAACTATGTTCTTGAAGACCAGACCCTTGTCAA

GAATATTAGGTATAGATTGGGGTGCGGTTTTTCAAATCAAATGATCATCAATAACAGGGGGGTGGGTGAAACAGTCAATT

CTAAACTTAAAAGTTATCCCCATAATCGTCATATCATATACCCGGATTGCAATAAGGAGTTGTTTTGTATCAAAGATAGC

TGCATATCTAAGAAGCTCTCGGAGCTATTCAAGAAGGGTAATTCCTTGTACTCTAAGATAAGTCACCAGATACTGGACTG

TCTTAAGAGAGTCAATGGGAAATTAGGCCTGGGCACAGATCTTACCCATGGTCTGAAGGAGGGTATCCTTGACCTAGGGT

TGCACATGCATAGCTCTCAATGGTTCGAGACTTTCCTGTTCTGGTTCACTATCAAGACAGAGATGAGATCAATGATCAAA

GAACAGTCCCATATATGCCACAAGAGGAGATATAACCCGATTTTTGTATCGGGGGATGCATTCGAGGTGCTCGTATCACG

AGACCTCGTTGTGATAATTGACAAGAATACCCAGTATGTCTTCTACTTAACATTTGAGCTGGTCCTTATGTATTGTGATG

TCATAGAGGGCAGACTTATGACGGAGACAGCCATGGCCATAGACCAGAGATATTCAGAACTTCTAAGCCGGGTTAGATAC

TTGTGGGATCTTATTGATGGGTTCTTCCCAACACTAGGCAATACCACCTACCAAATTGTTGCTCTGCTTGAACCATTATC

GTTGGCTTATCTTCAACTTCAGGATATCACGCTGGAGTTAAGAGGTGCTTTTTTAGACCATTGCTTCAAAGAACTCTATG

AGATACTGGAGCATTGCGGCATTGACACAGAAGGCACCTACAATTCCATCACTGAAGGATTGGATTACGTATTCATCACC

CACGATATACATCTAACTGGGGAGATTTTTTCATTTTTCCGGAGTTTCGGACACCCCCGTCTAGAAGCAGTCACCGCCGC

AGAAAATGTCAGGAAACATATGAATCAACCGAAGGTAATCAGTTATGAAACTATGATGAAAGGACATGCGGTATTTTGCG

GGATAATCATCAATGGTTTTAGAGACCGACATGGCGGCAGCTGGCCCCCTGTTGCATTGCCAGAACATGCTTCTGCTGCG

ATCCGGAATGCGCAAGCATCAGGTGAGGGACTGACCCATGACCTGTGTATAGACAACTGGAAATCCTTTGTTGGATTCAA

ATTCGGCTGCTTTATGCCACTCAGCCTAGATAGTGATTTGACCATGTATCTCAAAGACAAGGCATTGGCTGCACTGAAGA

ATGAGTGGGATTCAGTTTACCCGAAAGAATACCTCCGTTATAACCCACCTAGAGGGACAGAGTCAAGGCGACTGGTGGAG

GTGTTCCTGAATGACTCCAGCTTTGATCCTTATAACATGATAATGTACGTGGTGAATGGCTCCTATCTTAAAGACCCTGA

GTTTAATCTCTCATACAGCCTAAAAGAGAAGGAGATAAAGGAGACAGGGCGGTTGTTCGCCAAAATGACCTATAAGATGC

GGGCCTGTCAGGTAATTGCTGAGAATCTGATATCGAATGGTGTTGGGAAGTACTTCCGAGACAATGGGATGGCGAAGGAC

GAGCATGACCTAACAAAAGCCCTTCACACTTTGGCAGTCTCAGGTGTTCCCAAGAACAACAAAGATAACCACCGAGGTGG

GCCTCCCAGAAGGACAACAACCCGAGAGGTGAGGTCAAGCCGGGACACCAGAACACAAAATAGAGACAGGGTCCAAGGGG

GACCTATGTACAACTATTTGCAATGCCAACCAATCAGCCCTGATCAAGGTGAGTCATACGAGACTGTTAGTGCATTTATC

ACCGCTGACCTTAAGAAGTATTGCCTGAATTGGAGATACGAGACAATCAGCATATTTGCACAGAGGCTGAATGAAATATA

TGGACTACCATCCTTCTTTCAATGGTTGCACAGGATATTGGAAAAATCCGTACTCTACGTCAGTGACCCACATTGCCCTC

CCGATCTAGATAATCATATCCCTCTGGACAGTGTCCCTAATGCCCAAATATTCATTAAGTACCCAATGGGCGGAATAGAA

GGTTATTGCCAAAAACTATGGACAATCAGTACTATACCATACTTGTATCTGGCAGCCTATGAGAGCGGAGTAAGAATCGC

CTCACTGGTGCAGGGTGACAATCAGACAATCGCAGTGACAAAAAGAGTTCCAAGTTCTTGGCCTTATTCACTAAAAAAGA

GGGAGGCATCCAAAGCCGCTCAAAATTACTTCGTAGTCTTAAGGCAAAGATTGCACGATGTAGGTCATCACTTGAAGGCT

AATGAAACCATAGTATCTTCTCACTTTTTTGTGTATTCCAAAGGGATTTATTATGATGGCCTGCTAGTCTCACAATCACT

AAAGAGCATCGCCAGATGTGTCTTCTGGTCCGAGACTATAGTGGATGAAACCAGAGCGGCTTGCAGCAATATCGCAACAA

CTGTCGCTAAGAGTATAGAGAGGGGTTATGATAGGTACCTTGCATATTCTTTGAATATCCTCAAGATTTTCCAACAGATC

CTTATATCCCTTAACTTCACTATTAACACAACAATGACTCAGGATGTCGTGGCACCGATCGTCGAGAACGGTGATTTGCT

GATAAGGATGGCACTCTTGCCAGCACCCATCGGAGGTCTAAATTATCTTAACATGAGCAGGTTATTCGTGAGAAATATCG

GCGACCCGGTCACCTCCTCTATAGCCGACCTGAAGAGGATGATAGATGCCGGGCTCATGCCAGAAGAAACATTGCATCAA

GTGATGACCCAGACCCCGGGAGAATCATCCTACCTTGATTGGGCAAGTGACCCTTATTCTGCCAACCTACCCTGCGTACA

GAGTATAACTCGTCTTCTCAAAAACATCACTGCACGATATATTTTAATCAGCAGCCCAAACCCGATGCTGAAAGGATTGT

TTCATGAGGGGAGTAGAGATGAAGACGAGGAGCTTGCAAGTTTCCTAATGGATCGGCATATAATTGTTCCAAGAGCTGCA

CATGAAATCTTAGACCATAGTATAACCGGGGCAAGAGAAGCTATAGCCGGGATGTTGGACACCACCAAGGGTCTGATTAG

AACAAGTATGAAACGGGGTGGCCTCACCCCTCGAGTCTTAGCCCGCCTTTCCAATTATGATTATGAACAATTCCGATCCG

GGATAACACTATTGACAAAGAAAGGGCAGTGTTATCTCATTGACAAGGATTCGTGCTCGGTGCAGCTCGCTATAGCTCTG

AGGAGCCACATGTGGGCTAGGTTGGCTCGCGGGAGACCAATCTATGGGTTGGAGGTGCCCGATATATTAGAATCGATGAA

CGGCTACCTTATCAAACGCCACGAGTCTTGTGCTATCTGTGAAACGGGCTCAAGTCACTACGGATGGTTTTTCGTCCCTG

CAGGGTGCCAGCTTGACGATGTATCTAGAGAGACTTCGGCTCTCCGTGTACCTTATGTCGGATCAACAACTGAGGAAAGG

ACTGACATGAAACTTGCATTTGTTAGATCTCCAAGTAGATCCCTCAAATCAGCAGTTAGAATTGCAACAGTTTACTCATG

GGCCTACGGGGATGATGAGAAGTCATGGAGTGAAGCCTGGATGCTAGCTAGGCAGAGAGCCAATATCACCTTAGATGAAT

TGAGAATGATCACTCCAGTCTCTACATCCACCAACCTAGCCCACCGGTTGAGGGATCGGAGCACCCAGGTGAAATACTCG

GGGACATCCCTTGTGAGGGTTGCAAGATACACAACCATCTCCAATGATAACCTGTCATTTGTGATATCTGAGAAGAAAGT

AGATACCAACTTCATTTATCAGCAAGGGATGTTGCTTGGTCTTGGGATCCTTGAGAACCTCTTTAGGCTAGAGGCCACCA

CCGGAGTATCCAACACAGTGCTACACCTGCACGTGGAAACAGAATGTTGTGTTGTACCTATGGTGGATCACCCAAGGATA

CCGAGTCTCCGTAATATAAAAGTTACGGATGAGCTATGCACAAACCCTTTGATCTATGATAGGTCTCCCATCATAGAGCA

CGATGCAACCCGACTATACTCACAGAGCCACAGGAGACATTTGGTGGAGTTTGTTACCTGGTCCACAAGCCAGCTTTACC

ATATACTGGCTAAGTCTACAGCAATGTCCATGATTGAATTGATCACGAGATTCGAGAAAGATCACATGAATGAAATAGCC

GCCCTGATTGGCGATGACGACATCAACAGCTTCATCACAGAATTTTTGCTAGTGGAGCCTAGATTATTTATAGTTTACCT

TGGTCAGTGTGCTGCCATCAATTGGGCCTTTGATATACATTATCATCGGCCCTCGGGCAAGTACCAGATGGGGGAACTCC

TCTACTCTCTGCTCTCTCGGATGAGCAAAGGAGTATATAAGATCTTCACCAATGCTTTGAGCCACCCCAAAGTTTACAAG

AAATTTTGGCGAAGTGGTGTCATTGAGCCGATTCATGGCCCATCCCTGGATACACAGAATTTACATGTCACTGTCTGTGA

CATGATATATGGGTCCTATGTCACCTACCTGGATCTTTTGCTGAATGATGAGCTAGATGATTACCCGTATTTGCTCTGCG

AGAGTGATGAGGATGTGGTCACAGACAGGTTCGACAACATTCAAGCCAAACATCTCTGTGTATTAGCCGATGTATATTGC

AGCTCCAAGAGGTGTCCTTCAATAATCGGGATGTCTCCTATAGAAAAATGTACCATTCTCACACATTACATCAAGGGAGA

ATCAGTACAATCCCCGTCTGGAACCTCATGGAACACTGATCCCCTTGTAGTAGATCATTACTCATGCTCTCTGACCTACC

TTCGCCGAGGTTCCATCAAACAAATCAGGTTGAGGGTGGATCCTGGGTTTGTATTCGAGGCGTTGACAGACGTCGACTTT

AAACAACCTCGCAAAGCTAAGTTAGACATATCGGTCGTTGGGTTGACTGATTTCTCTCCCCCTTGCGACAACGTCGGTGA

TTTTCTAGGGACTATCAACACATTGAGGCACAATCTGCCTGTCACCGGAACCGGGGTCTCGAACTATGAAGTCCACGCTT

ATCGTAGAATTGGCCTGAATTCATCAGCATGTTACAAAGCCGTAGAGATCTCCACGTTAATCAAGCCATCCCTAGAAGTC

GGAGAGCATGGATTGTTCTTAGGGGAAGGTTCCGGTTCAATGCTGGCTGCGTACAAGGAAGTTCTTAAATTAGCAAATTG

TTATTACAACAGTGGGGTAACAGCGGAGGGCAGAGCCGGACAGAGGGAAATATCTCCCTATCCTTCAGAGATGAGCCTAG

TAGAGAATCAAATGGGGATAGAGCGGAGTGTTAAAGTGCTGTTCAACGGCAAACCTGAAGTAACTTGGGTGGGGACCACC

GATTGCTACAAGTACATAATCAGTAACATCCAAACCTCTAGTCTGGGTTTCATACATTCAGATATTGAGACGCTCCCAAC

CAAAGATGCTGTTGAGAAATTAGAAGAATTTGCCTCTATCCTATCCTTATCCTTGATTTTGGGGAAAATCGGCTCTATTA

CAGTTGTCAAAATTATGCCCATTAGCGGGGATTTTACCCAAGGCTTCATAGCCTATGCCATCCAATATTTTAGGGAGAGC

CTGCTTGCCTATCCGAGATATAGTAACTTCATCTCGACTGAGTGTTACCTTATCATGATAGGATTAAAGGCCAATCGATT

GATTAACCCAGAAGCCATTAAGCAAAGCATAATCAGAGCGGGGGTCAGGACTGCACCAGGACTTGTGAGCCATATATTAT

CAGAGAAACAGAAAGGTTGTATTCAATCTTTTTTGGGTGATCCCTATATCCAGGGAGACTTCAATAAGCACCTTAAATCT

CTAACCCCTATTGAGAAAATCCTGGTAAATTGTGGTCTCTCGATCAATGGCACAAAAATCTGTAGAGATCTAATCCACCA

TGATATCGCCTCCGGTCCAGACGGTCTTATGAGCTCTACAATTATTTTATATAGGGAATTGGCTCATTTCAAAGATAATA

TAAGGAGTCAGCACGGAATGTTCCACCCCTACCCAGTACTGGCCAATAGCCGGCAACGTGAATTAATCCTTCGAATAGCC

AAGAAATTCTGGGGGTATGTCTTGCTATATTCTGATGACCCGGCACTAATCAGACAAACAATCAAGAACTTGAAACGGAA

TCATCTAACCTTTGACTTACATAGTAACCCGTTTATTAAAGGGCTATCCAAAGCTGAGAAACTGCTGGTGCGGACGAGTT

CACTAAGAAGGGAATGGTTGTTCACTCTCGATACGAAAGAAGTGAAAGAGTGGTTCAAATTGGTAGGTTACAGTGCACTC

ATCAGAGGCTAGTTAGTTATACGTCTGCCCCCTTCTCCTCCGCCATGAGACCCCACTGACGATTCAGAAGATTAAAGAAA

ACTACATATTGGATAAGTATCTATTCCCAGCTTTGTCTGGT

>KT860064.1/sheep/India/2015-01-05

ACCAAACAAAGTTGGGTAAGGATAGATCTTATAATAACTATGGACTAGCAAACTTAGGAGTAAAGATCCTACTGTCGGGG

GGAGGAGGAGGAGCAAGATCTCTGATTATGGCGACTCTCCTTAAAAGCTTAGCATTGTTCAAAAGGAACAAAGACAAAGC

GCCGACGGCATCAGGTTCAGGAGGGGCCATCCGAGGGATCAAGAATGTTATCATAGTCCCGATTCCCGGAGACTCGTCCA

TCACTACCCGTTCAAGACTGCTCGACAGGCTTGTCAGATTGGCCGGAGATCCTGACATCAACGGGTCAAAGCTGACCGGC

GTGATGATCAGCATGTTATCTTTGTTCGTAGAGTCACCCGGGCAGTTGATACAGCGAATCACAGATGATCCGGATGTTAG

TATCCGCCTTGTTGAGGTAGTTCAAAGTACTAGGTCCCAGTCCGGGTTGACCTTTGCATCACGTGGTGCTGATTTGGATA

ACGAAGCAGACATGTATTTTTCAACTGAGGGGCCCTCGAGTGGAGGTAAGAAAAGGATCAACTGGTTTGAGAACAGAGAA

ATAATAGACATAGAGGTGCAGGATCCAGAAGAGTTCAATATGTTGTTAGCCTCCATACTAGCACAAGTCTGGATCCTTCT

GGCCAAGGCTGTCACGGCACCGGATACGGCAGCTGACTCAGAACTGAGAAGGTGGGTTAAATACACACAACAAAGGAGAG

TGATTGGGGAATTTCGCCTTGACAAAGGGTGGCTGGACGCAGTCCGCAACAGAATTGCAGAAGATCTATCACTCCGGCGG

TTCATGGTATCTCTCATACTTGACATCAAGAGGACCCCCGGCAACAAGCCAAGGATCGCAGAAATGATCTGCGACATTGA

CAACTATATTGTCGAAGCAGGACTCGCCAGTTTCATCCTTACTATCAAATTTGGTATAGAAACCATGTATCCTGCATTAG

GGCTCCACGAGTTTGCCGGGGAATTGTCCACTATTGAGTCCTTGATGAACTTGTATCAACAGCTAGGAGAGGTTGCACCC

TACATGGTAATTCTAGAGAACTCAATTCAGAACAAGTTTAGTGCAGGAGCTTATCCCCTCCTCTGGAGCTATGCGATGGG

TGTCGGAGTTGAGCTGGAGAACTCAATGGGGGGCTTGAACTTTGGCAGATCATATTTTGACCCTGCCTATTTTCGTCTCG

GACAGGAGATGGTCAGAAGATCTGCAGGAAAGGTCAGCTCTGTGATTGCGGCTGAGCTCGGCATCACAGCAGAGGAAGCT

AAACTAGTCTCGGAAATAGCCTCGCAGGCTGGGGACGAAAGAACCGCTAGAGGGACTGGGCCTCGACAGGCGCAGGTCTC

CTTCCTACAGCACCAAACAGGAGGGGGAGAGTCGTCCGCACCAGCGACCAGAGAAGGGGTCAAAGCTGCGATCCCAAACG

GATCCGAAGAAAGGGACAGAAAGCAAACACGCCCAGGGAGGCCCAGAGGAGAGACCCCCGGCCAACTGCTCCTGGAATTC

ATGCCAGAGGATGAGGTCTCGCGAGAATCTGGTCAAAACCCTCGTGAGGCTCAAAGGTCGGCCGAGGCACTCTTCAGGCT

GCAGGCCATGGCCAAGATTCTGGAAGACCAGGAGGAGGGAGAAGACGACAATCAGGTCTACAACGACAAGGATCTCCTCG

GCTAAGCAGACGCACCCTCTGTCGAAATCAGTGACGAGACATCTCCTACCAGTATTATAAAAAACTTAGGACCCAGGTCC

AAGCAAGCACACATCGACACTCCAACCAGTCGAGCGGAGGCCACCGATGGCAGAAGAACAAGCATACCATGTCAACAAGG

GGCTGGAATGTATCAAGTCCCTCAAAGCCTCTCCCCCGGATCTATCCACCATTAGAGATACCATCGAGAGCTGGAGAGAG

GGGCTTAGCCCCTCGGGCCGTGCAACACCGAACCCTGATACGTCCGAGGGAGATCATCAGAATATCAACCAATCATGCTC

ACCAGCAATCGGACCAAACAAAGTCTACTTGTCTCCTGAAGATAATCTCGGATTTAGAGAGATCACTGGCAACGACTGTG

AGGCTGAGTTCGGAGGAGTCCAGGGAAAAGGATCCAACTCTCAAGTACAGCGTTACTATGTTTATAGCCACGGGGGTGAA

GAGATTGAAGGACTCGAGGATGCTGACTCTCTCGTGGTTCAAGCAGATCCTCCAGTTGCTAACATATTCAATGGAGGAGA

GGATGGATCTGACGACAGCGATGTGGACTCTGGCCCAGATGATCCCAGCAGAGATACTCTATATGACCGGGGACCTGCTG

CCGGCAATGATGTCGCTAGGTCCACAGATGTCGAAAAACTAGAAGGTGCTGATATTCAAGAAGTCCTCAACTCCCAGAAA

AGCAGAGGAGGAAGATTCCAAGGCGGGAAAACCTTACGAGTCCCGGAAGTACCCGATGTCAAGAACTCCAGACCATCAGC

CCAATCAATTAAAAAGGGCACAGACGGGAACTCAGTCTCATCTGGAACGGTGACAGAGTGTTTATCGATAAGTGGTGCAA

CCCAAGCTGTGCCAGAGTCAAGATGGGAGTCATCAGAGCAAAATGCGTCTGTGGGGAGTGTCCTCAAGTCTGCGAAGAGT

GCAAAGACGATCCAGGGGTCGACACAAGAATCTGGTACCACAGCATCACCGACTCAGCCTAAAGAGAATGACTCCGAGTA

TGAGTATGAGGATGACCTATTTACAGAGATTCAGGACATCCGTGCAAGCATTGCCAAGATCCATGATGACAACAAGACTA

TCCTCTCAAAGCTTGATTCTATACTGTTATTGAAAGGAGAAGTCGACACTATCAAGAAACAAATCAGCAAGCAGAATATA

AGTATATCCACCATTGAGGGCCATCTCTCCAGTATAATGATAGCCATCCCGGGCTTTGGGAAGGATATCAAGGACCCAAC

ATCCGAGGTTGAGTTGAACCCAGATTTAAGACCTATAATAAGCCGTGACTCTGGCAGAGCTCTCGCGGAGGTCCTCAAGA

AACCCGCTGTGGATAGGTCTCAGAAAACTGGAACCAAAGCCAACTCCAGCTCAAAGGGTCAGCTTCTTAAGGATCTCCAG

CTAAAACCTGTCGACAAGCAGGCAAGCTCTGCAATCGGGTTTGTCCCATCCGACCATGAATCATCCAGAAATGTCATCCG

CTCCATAATCAAGTCGAGCAAGCTAAACATTGATCACAAGGACTATCTTCTAGATTTACTGAATGATGTGAAAGGCTCAA

AGGATCTTAAGGAATTCCACAAGATGCTAACAGCAATTCTCGCCAAGCACCCGTAACACATCCCCCAGTCACCATCTCAT

ACTCGGCTGAAGACATCCTTTCAATCAGGCTATTACAAAAAACTTAGGAGCAAGGGCAACTGAGCTTCGCAGACAAGATG

ACCGAGATCTACGACTTCGACAAATCAGCATGGGATGTCAAAGGGTCAATTGCCCGCATAGAACCCACCACCTACCACGA

CGGCCGACTGATACCCCAGGTAAGGGTCATCGATCCTGGTCTGGGAGACAGAAAAGATGAGTGCTTTATGTACCTGTTTC

TCCTGGGAGTGATTGAGGATAACGACCCCCTGTCTCCCCCCGTCGGAAGAACCTTTGGCTCTTTACCTCTAGGGGTCGGT

AGGTCAACTGCTAAGCCAGAAGAACTACTAAGGGAGGCCACAGAACTAGACATAGTGGTGAGGCGCACTGCAGGAGTAAA

TGAGAAACTGGTATTTTACAACAACACTCCGCTGTCCTTGTTAACGCCCTGGAAGAAAGTTCTGACAACCGGAAGTGTGT

TTAGCGCTAACCAGGTTTGCAATGCAGTCAACCTAGTCCCGCTTGATACTCCCCAGAGGTTCAGGGTTGTGTACATGAGC

ATAACTAGATTATCAGACAATGGTTACTATAGTGTGCCCAGGAGAATGCTGGAGTTCCGCTCAGCCAATGCAGTAGCCTT

CAACATCTTGGTCACACTAAGAATTGAGAATGGCACAAACCCTAGAAGATACATAGTCGGCTCATGGGAGAATTCAGAGG

TCACATTTATGGTACACGTGGGCAACTTTAGAAGAAAGAAGAACGAAGTATACTCTGCTGATTACTGCAAAATGAAGATT

GAAAAGATGGGTTTAGTTTTTGCCCTGGGAGGAATAGGTGGAACAAGTCTCCATATTAGAAGCACCGGGAAAATGAGCAA

AACCCTCCATGCACAGCTGGGGTTCAAGAAAATCTTATGTTACCCCCTAATGGATGTTAATGAGGATCTTAACCGATATC

TCTGGCGGGCAGAGTGCCGAATAGTCAAAATCCAAGCCGTCTTACAGCCATCAGTGCCCCAAGAATTCCGTGTCTACGAT

GATGTCATCATCAACGATGATCAAGGCTTGTTCAAGATCCTGTAGTTCATCTGCAAAGGAACGAACCCACCCCCAGGGGG

GGGCCCCCCCCTCCCCCG-CCAAAAAACACACTCCAAGGCTCCCCGACACAGCGCCAACACAGCCACCCTCCCCACTGGA

CGAGGCCGGCC-CCCCAACAAACCCGGTCCAGGC-GGGGGGGGGCCCCCCTGACCCGCCGGGCGCCGACCGAGCAGGCCG

GCGAGAAGCCCCCAAGAGGCCCCGACGAAGACCGAGCACCCCTCCCCCCC-AAACCGTTCCCCC-AACACGCCCGAGGCC

AGGCGCCGGGCACCCCCAGACCCGCCACCGGCGGGACAGAACCCCAGGGGGGGACCCCCCAAACCAAAGGGACCACCGGG

GAAGGATCCGCCGAACCAGCACAGACCCGACCCAAGCAAGGCGGACCCCCAACATGAAACCGCCCAGCACACCCCACTCA

TCATCAAACAAAACTTAGGGGCCAAGCCCATACGACAGGCCCCAGTGCGCCAGGCGGAGGGACGCAGAAAGGAAGGAGAC

AAGACCGCCCCAGAAGGAGGACACCCAACCACCGAAACACCCAGCCCCAGGGAAGGAG--CCCCCCCCCACATGGCGAAA

GAACGGGCAAGAGACCCCGAGAAGAACCCGACCGGGAGCCAGGAGCCGGCGACGCCGGCGGCAAACAAGCCGGGCAAACA

CCCGACCCCACCAGAGGAGCCGAGGAGAGCCCGACCCCGCACGAACCAAACAGCAGGGAGCCCAACCCCACCAAAAAAAG

AGGCGAGGCCCC-CCT-----CCTCCCCGCACACAACCCGCAGACAGACCGAGAGAGGACCACCTCCCAACAAGACAGGC

GGCTCCCTCGCAGCCGAGCCGCAGCTCCCCAGAACAGGACACCCCCGTACACTCAAAACACCCCCCCGCCCCCCGCCACC

GGGACAGGGGCTCCGCCACCACAGACCAAAGCCACCGAGAGCAAGCCAGAGGTCCCCCCCAGTCCCGCCCGGCGCCCCCC

CGAAAAGGCCCCGGCAAGGCCCCCCAACCCGACGTCCCCCGCACGCACCGAAAGAGGGACCCGCCCGAGAACGTGCGGCA

CATCCATGCGTAAACATTATGACACGGGTCGCAATCTTGACATCTCTGTTTCTTCTCCCAAATGTTGTTGCGTGCCAGAT

TCACTGGGGCAATCTATCCAAGATCGGGATTGTGGGAACAGGGAGTGCCAGCTACAAGGTGATGACTAGGCCGAGCCACC

AAACTCTGGTTATAAAGTTAATGCCAAATATAACAGCCATCGACAATTGTACAAAGTCAGAGATTGCAGAGTACAAGAGA

TTGCTGATCACAGTGTTAAAGCCTGTAGAGGATGCTCTGTCGGTGATAACCAAGAATGTAAGACCAATTCAAACTCTGAC

ACCTGGGCGTAGAACCCGTCGTTTTGCTGGAGCTGTTCTGGCCGGAGTAGCGCTTGGAGTTGCGACAGCCGCTCAGATAA

CTGCAGGAGTCGCTCTTCATCAATCATTGATGAACTCCCAAGCAATTGAGAGTTTAAAAACCAGTCTTGAGAAGTCGAAT

CAGGCAATAGACGAAATCGGACTTGCAAATAAGGAGACCATACTGGCAGTACAGGGCGTCCAGGATTATATCAACAATGA

GCTTGTCCCTTCTGTTCATAGAATGTCATGCGAGCTGGTAGGCCACAAGCTCGGCCTCAAGCTCCTTAGGTACTACACCG

AGATCCTGTCCATATTCGGGCCCAGCCTTCGAGACCCGATAGCTGCCGAAATATCAATCCAGGCACTCAGTTATGCATTA

GGCGGAGACATTAATAAAATCCTGGACAAGCTTGGGTATAGCGGTGGGGATTTCCTTGCCATCCTAGAAAGCAAGGGAAT

AAAGGCCCGGGTTACATATGTGGACACAAGAGATTACTTTATAATGCTTAGCATCGCCTACCCAACCTTATCTGAGATCA

AGGGGGTGATAGTTCACAAGATAGAAGCTATAACATACAACATTGGGGCACAGGAGTGGTATACTACTATCCCTAAATAT

GTAGCCACTCAGGGATATCTGATATCGAACTTTGATGAGACGTCATGCGTATTCACTCCAGAGGGGACAGTTTGCAGCCA

GAATGCGTTGTACCCAATGAGCCCATTGCTTCAGGAATGTTTCAGGGGGTCAACAAAATCGTGCGCCAGAACCCTAGTTT

CAGGTACCATAAGTAATAGATTTATCCTATCAAAAGGGAACCTGATCGCAAATTGTGCGTCAGTTTTGTGCAAATGTTAC

ACAACGGAGACAGTCATCAGCCAAGATCCTGACAAACTACTAACTGTTGTAGCATCCGACAAGTGTCCTGTAGTTGAGGT

GGATGGAGTGACAATACAGGTCGGCAGTCGAGAGTATCCGGATTCTGTATACTTACACAAAATAGACTTAGGCCCAGCCA

TCTCCCTAGAAAAACTGGATGTAGGTACCAATTTAGGCAATGCAGTTACAAGACTGGAGAATGCAAAGGAGCTCCTGGAT

GCATCAGACCAAATACTGAAGACTGTCAAAGGGGCACCTTTTGGTGGGAACATGTACATAGCATTGGCAGCTTGCATTGG

AGTATCCCTAGGGCTTGTCACATTAATATGCTGCTGTAAGGGGAGGTGTAAGAACAAGGAAATCCCTATCTCCAAAATCA

ACCCAGGGCTCAAACCCGATCTGACCGGGACCTCAAAGTCGTACGTGAGATCACTGTAGCCAGAATCACCTGAATCATCT

GGCATCACACACATACATGTACGACACAAGCAGTCTGAGGACGCAGGAAACCCAGCCCCCGATCACCCACCTGACCCTAC

TCCACGCTCCGCTACACATTAGTCATTAAACAAAACTTAGGACGAAAGGTCAATCACCATGTCCGCACAAAGGGAAAGGA

TCAATGCCTTCTACAAAGACAATCCTCACAATAAGAACCATAGGGTAATCCTGGATAGAGAACGCTTGGTCATTGAAAGA

CCCTACATCTTGCTTGGAGTCCTGCTGGTAATGTTCCTGAGTCTAATCGGACTGCTGGCCATTGCAGGGATCAGACTTCA

CCGGGCCACCGTTGGAACTTCAGAGATCCAGAGTCGGCTGAATACCAATATTAAGTTGACCGAATCTGTTGATCACCAGA

CTAAGGATGTCTTAACTCCCCTTTTTAAAATAATTGGCGATGAAGTCGGCATCAGGATTCCACAGAAATTCAGTGATCTT

GTCAAGTTCATCTCCGACAAGATTAAATTCCTCAACCCTGATAGAGAGTATGATTTCAGGGATCTCCGGTGGTGCATGAG

TCCCCCCGAGAGAGTCAAAATTAATTTTGATCAGTTTTGTGAGTACAAAGCTGCGGTTAATTCAATTGAACATATATTTG

AGTCACCACTCAACAAGTCAAAAAAGCTGCAATCATTGACTCTCGGGCCCGGAACAAGTTGTCTAGGCAGGACAGTAACA

AGAGCTCATTTCTCAGAACTTACACTGACCTTGATGGACTTGGATCTAGAGATGAAGCACAACGTGTCCTCAGTGCTTAC

CGTAGTTGAAGAGGGGTTATTCGGAAGAACATATACCGTCTGGAGATCCGATGCCAGGGATCCGAGCACCGATCTAGGTA

TCGGCCATTTTTTAAGAGTCTTCGAGATTGGACTGGTAAGAGATCTCGGGCTGGGTCCCCCCGTTTTTCATATGACCAAC

TATCTCACAGTGAACATGAGTGATGTCTATCGGAGATGTATTTTAGCGGTAGGGGAGTTGAAGTTGACAGCCCTATGCAC

CTCATCTGAGACTGTAACACTGAGTGAGAGAGGAGTTCCAAAGAGGGAGCCTCTTGTAGTTGTGATACTTAACCTGGCTG

GACCAACTCTAGGGGGCGAACTATACAGTGTCTTGCCTACCTCTGATCTCATGGTGGAGAAACTCTATTTATCTTCACAT

AGAGGGATCATCAAGGATGACGAGGCCAATTGGGTAGTGCCGTCTACCGATGTTCGTGATCTTCAGAACAAAGGTGAGTG

TCTAGTGGAAGCATGCAAGACTCGACCTCCTTCATTTTGCAATGGCACAGGATCAGGCCCGTGGTCAGAGGGGAGAATCC

CTGCTTACGGGGTGATCAGGGTCAGTCTTGACTTAGCTAGTGACCCGGATGTAGTTATCACTTCAGTGTTTGGCCCACTG

ATTCCTCACCTATCCGGCATGGATCTTTACAACAACCCGTTTTCAAGAGCTGTATGGTTGGCTGTACCACCTTATGAGCA

GTCATTTCTAGGAATGATAAATACAATTGGATTCCCTAACAGAACAGAGGTTATGCCGCACATTTTGACCACAGAGATCA

GAGGCCCTCGGGGTCATTGCCATGTCCCCATAGAATTGTCCCGCAGGGTTGATGACGATATCAAGATCGGGTCCAACATG

GTCATATTGCCGACGATGGACCTGAGATATATTACGGCCACTTATGATGTTTCCAGGAGCGAGCATGCAATCGTGTACTA

TATCTATGACACAGGTCGCTCATCATCTTACTTCTACCCAGTTCGACTGAATTTCAAAGGCAATCCCCTCTCTCTGAGGA

TAGAGTGTTTCCCTTGGCGTCATAAGGTGTGGTGCTACCATGATTGTCTTATATACAACACCATAACAGATGAAGAGGTC

CATATGAGAGGGCTGACCGGTATAGAGGTAACATGCAATCCAGTCTGAGCAGAGCTACGACCGTCACTCAAGCAGTCCCT

TGAGTCGCCACCGAGTCCAAGCAGCACAGCCTGGGACTCTCAACAGCACAGCCCAGCCAACAGTGTTATAAAAAACTTAG

GAGCCAGGGTTGTAGGGGCCATGGACTCTCTATCAGTCAATCAGGTCTTGTACCCTGAGGTTCACCTAGACAGCCCTATT

GTCACAAACAAACTAGTTGCCATCCTTGAGTACTCGGGTATCGACCATAACTATGTTCTTGAAGACCAGACCCTTGTCAA

GAATATTAGGTATAGACTGGGGTGCGGTTTTTCAAATCAAATGATCATCAATAACAGGGGGGTGGGTGAAACAGTCAATT

CTAAACTTAAAAGTTATCCCCATAATCGTCATATCATATACCCGGATTGCAATAAGGAGTTGTTTTGTATCAAAGATAGC

TGCGTATCTAAGAAGCTCTCGGAGCTATTCAAGAAGGGTAATTCCTTGTACTCTAAGATAAGTCACCAGGTACTGGACTG

TCTTAAGAGAGTCAATGGGAAATTAGGCCTGGGCACAGATCTTACCCATGGTCTGAAGGAGGGTATCCTTGACCTAGGGT

TGCACATGCATAGCTCTCAATGGTTCGAGACTTTCCTGTTCTGGTTCACTATCAAGACAGAGATGAGATCAATGATCAAA

GAACAGTCCCATATATGCCACAAGAGGAGATATAACCCGATTTTTGTATCGGGGGATGCATTCGAGGTGCTCGTATCACG

AGACCTCGTTGTGATAATTGATAAGAATACCCAGTATGTCTTCTACTTAACATTTGAGCTGGTCCTTATGTATTGTGATG

TCATAGAGGGCAGACTTATGACGGAGACAGCCATGGCCATAGACCAGAGATATTCAGAACTTCTAAGCCGGGTTAGATAC

TTGTGGGATCTTATTGATGGGTTCTTCCCAACACTAGGCAATACCACCTACCAAATTGTTGCTCTGCTTGAACCATTATC

GTTGGCTTATCTTCAACTTCAGGATGTCACGCTGGAGTTAAGAGGTGCTTTTTTAGACCATTGCTTCAAAGAACTCTATG

AGATACTGGAGCATTGCGGCATTGATACAGAAGGCACCTACAATTCCATCACTGAAGGATTGGATTACGTATTCATCACC

CACGATATACATCTAACTGGGGAGATTTTTTCATTTTTCCGGAGTTTCGGACACCCCCGTCTAGAAGCAGTCACCGCCGC

AGAAAATGTCAGGAAACATATGAATCAACCGAAGGTAATCAGTTATGAGACTATGATGAAAGGACATGCGGTATTTTGCG

GGATAATCATCAATGGTTTTAGAGACCGACATGGCGGCAGCTGGCCCCCTGTTGCATTGCCAGAACATGCTTCTGCTGCG

ATCCGGAATGCGCAAGCATCAGGTGAGGGACTGACCCATGACCTGTGTATAGACAACTGGAAATCCTTTGTTGGATTCAA

ATTTGGCTGCTTTATGCCACTCAGCCTAGATAGTGATTTGACCATGTATCTCAAAGACAAGGCATTGGCTGCACTGAAGA

ATGAGTGGGATTCAGTTTACCCGAAAGAATACCTCCGTTATAACCCACCTAGAGGGACAGAGTCAAGGCGACTGGTGGAG

GTGTTCCTGAATGACTCCAGCTTCGATCCTTATAACATGATAATGTACGTGGTGAATGGCTCCTATCTTAAAGACCCTGA

GTTTAATCTCTCATACAGCCTAAAAGAGAAGGAGATAAAGGAGACAGGGCGGTTGTTTGCCAAAATGACCTATAAGATGC

GGGCCTGTCAGGTAATTGCTGAGAATCTGATATCGAATGGTGTTGGGAAGTACTTCCGAGACAATGGGATGGCGAAGGAC

GAGCATGACCTAACAAAAGCCCTTCACACTTTGGCAGTCTCAGGTGTTCCCAAGAACAACAAAGATAACCACCGAGGTGG

GCCTCCCAGAAGGACAACAACCCGAGAGGTGAGATCAAGCCAGGACACCAGAACACAAAATAGAGACAAGGTCCAAGGGG

GACCTATGTACAACTATTTGCAATGCCAACCAATCAGCCCTGATCAAGGTGAGTCATACGAGACTGTTAGTGCATTTATC

ACCGCTGACCTTAAGAAGTATTGCCTGAATTGGAGATACGAGACAATCAGCATATTTGCACAGAGGCTGAATGAAATATA

TGGACTACCATCCTTCTTTCAATGGTTGCACAGGATATTGGAAAAATCCGTACTCTACGTCAGTGACCCACATTGCCCTC

CCGATCTAGATAATCATATCCCTCTGGACAGTGTCCCTAATGCCCAAATATTCATTAAGTACCCAATGGGCGGAATAGAA

GGTTATTGCCAAAAACTATGGACAATCAGTACTATACCATACTTGTATCTGGCAGCCTATGAGAGCGGAGTAAGAATCGC

CTCACTGGTGCAGGGTGACAATCAGACAATCGCAGTGACAAAAAGAGTTCCAAGTTCTTGGCCTTATTCACTAAAAAAGA

GGGAGGCATCCAAAGCCGCTCAAAATTACTTCGTAGTCTTAAGGCAAAGATTGCACGATGTAGGTCATCACTTGAAGGCT

AATGAAACCATAGTATCTTCTCACTTTTTTGTATATTCCAAAGGGATTTATTATGACGGCCTGCTAGTCTCACAATCACT

AAAGAGCATCGCCAGATGTGTCTTCTGGTCCGAGACTATAGTGGATGAAACCAGAGCGGCTTGCAGCAATATCGCAACAA

CTGTTGCTAAGAGTATAGAGAGGGGTTATGATAGGTACCTTGCATATTCTTTGAATATCCTCAAGATTTTCCAACAGATC

CTCATATCCCTTAACTTCACTATTAACACAACAATGACTCAGGATGTCGTGGCACCGATCGTCGAGAACGGTGATTTGCT

GATAAGGATGGCACTCTTGCCAGCACCCATCGGAGGTCTAAATTATCTTAACATGAGCAGGTTATTCGTGAGAAATATCG

GCGACCCGGTCACCTCCTCTATAGCCGACCTGAAGAGGATGATAGATGCCGGGCTCATGCCAGAAGAAACATTGCATCAA

GTGATGACCCAGACCCCGGGAGAATCATCCTACCTTGATTGGGCAAGTGACCCTTATTCTGCCAACCTACCCTGCGTACA

GAGTATAACTCGTCTTCTCAAAAACATCACTGCACGATATATTTTAATCAGCAGCCCAAACCCGATGCTGAAAGGATTGT

TTCATGAGGGGAGTAGAGATGAAGACGAGGAGCTTGCAAGTTTCCTAATGGATCGGCATATAATTGTTCCAAGAGCTGCA

CATGAAATCTTAGACCATAGTATAACCGGGGCAAGAGAAGCTATAGCCGGGATGTTGGACACCACCAAGGGTCTGATTAG

AACAAGTATGAAACGGGGTGGCCTCACCCCTCGAGTCTTAGCCCGCCTTTCCAATTATGATTATGAACAATTCCGATCCG

GGATAACACTATTGACAAAGAAAGGGCAGTGTTATCTCATTGACAAGGATTCGTGCTCGGTGCAGCTCGCTATAGCTCTG

AGGAGCCACATGTGGGCTAGGTTGGCTCGCGGGAGACCAATCTATGGGTTGGAGGTGCCCGATATATTAGAATCGATGAA

CGGCTACCTTATCAAACGCCACGAGTCTTGTGCTATCTGTGAAACGGGCTCAAGTCACTACGGATGGTTTTTCGTCCCTG

CAGGGTGCCAGCTTGACGATGTATCTAGAGAGACTTCGGCTCTCCGTGTACCTTATGTCGGATCAACAACTGAGGAAAGG

ACTGACATGAAACTTGCATTTGTTAGATCTCCAAGTAGATCCCTCAAATCAGCAGTTAGAATTGCAACAGTTTACTCATG

GGCCTACGGGGATGATGAGAAGTCATGGAGTGAAGCCTGGATGCTAGCTAGGCAGAGAGCCAATATCACCTTAGATGAAT

TGAGAATGATCACTCCAGTCTCTACATCCACCAACCTAGCCCACCGGTTGAGGGATCGGAGCACCCAGGTGAAATACTCG

GGGACATCCCTTGTGAGGGTTGCAAGATACACAACCATCTCCAATGATAACTTGTCATTTGTGATATCTGAGAAGAAAGT

AGATACCAACTTCATTTATCAGCAAGGGATGTTGCTTGGTCTTGGGATCCTTGAGAACCTCTTTAGGCTAGAGGCCACCA

CCGGAGTATCCAACACAGTGCTACACCTGCACGTGGAAACAGAATGTTGTGTTGTACCTATGGTGGATCACCCAAGGATA

CCGAGTCTCCGTAATATAAAAGTTACGGATGAGCTATGCACAAACCCTTTGATCTATGATAGGTCTCCCATCATAGAGCA

CGATGCAACCCGACTATACTCACAGAGCCACAGGAGACATTTGGTGGAGTTTGTTACCTGGTCCACAAGCCAGCTTTACC

ATATACTGGCTAAGTCTACAGCAATGTCCATGATTGAATTGATCACGAGATTCGAGAAAGATCACATGAATGAAATAGCC

GCCCTGATTGGCGATGACGACATCAACAGCTTCATCACAGAATTTTTGCTAGTGGAGCCTAGATTATTTATAGTTTACCT

TGGTCAGTGTGCTGCCATCAATTGGGCCTTTGATATACATTATCATCGGCCCTCGGGCAAGTACCAGATGGGGGAACTCC

TCTACTCTCTGCTCTCTCGGATGAGCAAAGGAGTATATAAGATCTTCACCAATGCTTTGAGCCACCCCAAAGTTTACAAG

AAATTTTGGCGAAGTGGTGTCATTGAGCCGATTCATGGCCCATCCCTGGATACACAGAATTTACATGTCACTGTCTGTGA

CATGATATATGGGTCCTATGTCACCTACCTGGATCTTTTGCTGAATGATGAGCTAGATGATTACCCGTATTTGCTCTGCG

AGAGTGATGAGGATGTGGTCACAGACAGGTTCGACAACATTCAAGCCAAACATCTCTGTGTATTAGCCGATGTATATTGC

AGCTCCAAGAGGTGTCCTTCAATAATCGGGATGTCTCCTATAGAAAAATGTACCATTCTCACACATTACATCAAGGGAGA

ATCAGTACAATCCCCGTCTGGAACCTCATGGAACACTGATCCCCTTGTAGTAGATCATTACTCATGCTCTCTGACCTACC

TTCGCCGAGGTTCCATCAAACAAATCAGGTTGAGGGTGGATCCTGGGTTTGTATTCGAGGCGTTGACAGACGTCGACTTT

AAACAACCTCGCAAAGCTAAGTTAGACATATCGGTCGTTGGGTTGACTGATTTCTCTCCCCCTTGCGACAACGTCGGTGA

TTTTCTAGGGACTATCAACACATTGAGGCACAATCTGCCTGTCACCGGAACCGGGGTCTCGAACTATGAAGTCCACGCTT

ATCGTAGAATTGGCCTGAATTCATCAGCATGTTACAAAGCCGTAGAGATCTCCACGTTAATCAAGCCATCCCTAGAAGTC

GGAGAGCATGGATTGTTCTTAGGGGAAGGTTCCGGTTCAATGCTGGCTGCGTACAAGGAAGTTCTTAAATTAGCAAATTG

TTATTACAACAGTGGGGTAACAGCGGAGGGCAGAGCCGGACAGAGGGAAATATCTCCCTATCCTTCAGAGATGAGCCTAG

TAGAGAATCAAATGGGGATAGAGCGGAGTGTTAAAGTGCTGTTCAACGGCAAACCTGAAGTAACTTGGGTGGGGACCACC

GATTGCTACAAGTACATAATCAGTAACATCCAAACCTCTAGTCTGGGTTTCATACATTCAGATATTGAGACGCTCCCAAC

CAAAGATGCTGTTGAGAAATTAGAAGAATTTGCCTCTATCCTATCCTTATCCTTGATTTTGGGGAAAATCGGCTCTATTA

CAGTTGTCAAAATTATGCCCATTAGCGGGGATTTTACCCAAGGCTTCATAGCCTATGCCATCCAATATTTTAGGGAGAGC

CTGCTTGCCTATCCGAGATATAGTAACTTCATCTCGACTGAGTGTTACCTTATCATGATAGGATTAAAGGCCAATCGATT

GATTAACCCGGAAGCCATTAAGCAAAGCATAATCAGAGCGGGGGTCAGGACTGCACCAGGACTTGTGAGCCATATATTAT

CAGAGAAACAGAAAGGTTGTATTCAATCTTTTTTGGGTGATCCCTATATCCAAGGAGACTTCAATAAGCACCTTAAATCT

CTAACCCCTATTGAGAAAATCCTGGTAAATTGTGGTCTCTCGATCAATGGCACAAAAATCTGTAGAGATCTAATCCACCA

TGATATCGCCTCCGGTCCAGACGGTCTTATGAGCTCTACAATTATTTTATATAGGGAATTAGCTCATTTCAAAGATAATA

TAAGGAGTCAGCACGGAATGTTCCACCCCTACCCAGTACTGGCCAATAGCCGGCAACGTGAATTAATCCTTCGAATAGCC

AAAAAATTCTGGGGGTATGTCTTGCTATATTCTGATGACCCGGCACTAATCAGACAAACAATCAAGAACTTGAAACGGAA

TCATCTAACCTTTGACTTACATAGTAACCCGTTTATTAAAGGGCTATCCAAAGCTGAGAAACTGCTGGTGCGGACGAGTT

CACTAAGAAGGGAATGGTTGTTCACTCTCGATACGAAAGAAGTGAAAGAGTGGTTCAAATTGGTAGGTTACAGTGCACTC

ATCAGAGGCTAGTTAGTTATACATCTGCCCCCTTCTCCTCCGCCATGAGACCCCACTGACGATTCAGAAGATTAAAGAAA

ACTACATATTGGATAAGTATCTATTCCCAGCTTTGTCTGGT

>KR261605.1/goat/India/2014-09-16

ACCAAACAAAGTTGGGTAAGGATCGATCTTAGAATAACTATGGACTGGCAAACTTAGGAGTAAAGATCCTACTGTCGGGG

GGAGGAGGAGGAGCAAGACCTCTGATTATGGCTACTCTCCTTAAAAGCTTAGCATTGTTCAAAAGGAACAAAGACAAAGC

GCCGACGGCATCAGGTTCAGGAGGGGCCATCCGGGGGATTAAGAATGTTATCATAGTCCCGATTCCCGGAGACTCGTCCA

TCACTACCCGTTCAAGACTGCTCGACAGGCTTGTCAGATTGGCCGGAGATCCTGACATCAACGGGTCAAAGCTGACCGGC

GTGATGATCAGCATGTTATCTTTGTTCGTAGAGTCACCCGGGCAATTGATACAGCGAATCACAGATGATCCAGATGTTAG

TATCCGCCTTGTTGAGGTAGTTCAAAGTACTAGGTCCCAGTCCGGGTTGACCTTTGCATCACGTGGTGCTGATTTGGACA

ACGAGGCAGACATGTATTTTTCAACTGAGGGGCCCTCGAGTGGAGGTAAGAAAAGGATCAACTGGTTTGAGAACAGAGAA

ATAATAGACATAGAGGTGCAGGATCCAGAAGAGTTCAATATGTTGTTAGCCTCCATACTAGCACAAGTCTGGATCCTTCT

GGCCAAGGCTGTTACGGCACCGGATACGGCAGCTGACTCAGAACTGAGAAGGTGGGTTAAATACACACAACAAAGGAGAG

TGACTGGGGAATTTCGCCTTGACAAAGGGTGGCTGGACGCAGTCCGCAACAGAATTGCAGAAGATCTATCACTCCGGCGG

TTCATGGTATCTCTCATACTTGACATCAAGAGGACCCCCGGCAACAAGCCAAGGATCGCAGAAATGATCTGCGACATTGA

CAACTATATTGTCGAAGCAGGACTCGCCAGTTTCATCCTTACTATCAAATTTGGTATAGAAACCATGTATCCTGCATTAG

GGCTCCACGAGTTTGCCGGGGAATTGTCCACTATTGAGTCCTTGATGAACTTGTATCAACAGCTAGGAGAGGTTGCACCC

TACATGGTAATTCTAGAGAACTCAATTCAGAACAAGTTTAGTGCAGGAGCTTATCCCCTCCTCTGGAGCTATGCGATGGG

TGTCGGAGTTGAGCTGGAGAACTCAATGGGGGGCTTGAACTTTGGCAGATCATATTTTGACCCTGCCTATTTTCGTCTCG

GACAGGAGATGGTCAGAAGATCTGCAGGAAAGGTCAGCTCTGTGATTGCGGCTGAGCTCGGCATCACAGCAGAGGAAGCT

AAACTAGTCTCGGAAATAGCCTCGCAGGCTGGGGACGAAAGAACCGCTAGAGGAACTGGGCCTCGACAGGCGCAGGTCTC

CTTCCTACAGCACCAAACAGGAGGGGGAGAGTCGTCCGCACCAGCGACCAGAGAAGGGGTCAAAGCTGCGATCCCAAACG

GATCCGGAGAAAGGGACAGAAAGCAAACACGCCCAGGGAGGCCCAGAGGAGAGACCCCCGGCCAACTGCTCCTGGAATTC

ATGCCAGATGATGAGGTCTCGCGAGAATCTGGTCAAAACCCTCGTGAGGCTCAAAGGTCGGCCGAGGCACTCTTCAGGCT

GCAGGCCATGGCCAAGATTCTGGAGGACCAGGAGGAGGGAGAAGACGACAATCAGGTCTACAACGACAAGGATCTCCTCG

GCTAAGCAGACGCACCCTCTGTCGAAATCAGTGACGAGACATCTCCTACCAGTATTATAAAAAACTTAGGACCCAGGTCC

AAGCAAGCACACATCGACACTCCAACCAGTCGAGCGGAGGCCACCGATGGCAGAAGAACAGGCATACCATGTCAACAAGG

GGCTGGAATGTATCAAGTCCCTCAAAGCCTCTCCCCCGGATCTATCCACCATTAGAGATACCATCGCGAGCTGGAGAGAG

GGGCTTAGCCCCTCGGGCCGTGCAACACCGAACCCTGATACGTCCGAGGGAGATCATCAGAATATCAACCAATCATGCTC

ACCAGCAATCGGACCAAACAAAGTCTACTTGTCTCCTGAAGATAATCTCGGATTTAGAGAGATCACTGGCAACGACTGTG

GGGCTGAGCTCGGAGGAGTCCAGGGAAAAGGATCCAACTCTCAAGTACAGCGTTACTATGTTTATAGCCACGGGGGTGAA

GAGATTGAAGGACTCGAGGATGCTGACTCTCTCGTGGTTCAAGCAGATCCTCCAGTTGCTAACATATTCAATGGAGGAGA

GGATGGATCTGACGACAGCGATGTGGACTCTGGCCCAGATGATCCCAGCAGAGATACTCTATATGACCGGGGACCTGCTG

CCGGCAATGATGTCGCTAGGTCCACAGATGTCGAAAAACTAGAAGGTGCTGATATTCAAGAAGTTCTCAACTCCCAGAAA

AGCAGAGGAGGAAGATTCCAAGGCGGGAAAACCTTACGAGTCCCGGAAGTACCCGATGTCAAGAACTCCGGACCATCAGC

CCAATCAATTAAAAAGGGCACAGACGGGAACTCAGTCTCATCTGGAACGGTGACAGAGTGTTTATCGATAAGTGGTGCAA

CCCAAGCTGTGCCAGAGTCAAGATGGGAGTCATCAGAGCAAAATGCGTCTGTGGGGAGTGTCCTCAAGTCTGCGAAGAGT

GCAAAGACGATCCAGGGGTCGACACAAGAATCTGGTACCACAGCATCACCGACTCAGCCTAAAGAGAATGACTCCGAGTA

TGAGTATGAGGATGACCTATTTACAGAGATTCAGGACATCCGTGCAAGCATTGCCAAGATCCATGATGACAACAAGACTA

TCCTCTCAAAGCTTGATTCTATACTGTTATTGAAAGGAGAAGTCGACACTATCAAGAAACAAATCAGCAAGCAGAATATA

AGTATATCCACCATTGAGGGCCATCTCTCCAGTATAATGATAGCCATCCCGGGCTTTGGGAAGGATATCAAGGACCCAAC

ATCCGAGATTGAGTTGAACCCAGATTTAAGACCTATAATAAGCCGTGACTCTGGCAGAGCTCTCGCGGAGGTCCTCAAGA

AACCCGCTGTGGATAGGTCTCAGAAAACTGAAACCAAAGCCAACTCCAGCTCAAAGGGTCAGCTTCTTAAGGATCTCCAG

CTAAAACCTGTCGACAAGCAGGCAAGCTCTGCAATCGGGTTTGTCCCATCCGACCATGAATCATCCAGAAATGTCATCCG

CTCCATAATCAAGTCGAGCAAGCTAAACATTGATCACAAGGACTATCTTCTAGATTTACTGAATGATGTGAAAGGCTCAA

AGGATCTTAAGGAATTCCACAAGATGCTAACAGCAATTCTCGCCAAGCACCCGTAACACATCCCCCAGTCACCATCTCAT

ACTCGGCTGAAGACATCCTTTCAATCAGACTATTACAAAAAACTTAGGAGCAAGGGCAACTGAGCTTCGCAGACAAGATG

ACCGAGATCTACGACTTCGACAAATCAGCATGGGATGTCAAAGGGTCAATTGCCCGCATAGAACCCACCACCTACCACGA

CGGCCGACTGATACCCCAGGTAAGGGTCATCGATCCTGGTCTGGGAGACAGAAAAGATGAGTGCTTTATGTACCTGTTTC

TCCTGGGAGTGATTGAGGATAACGACCCCCTGTCTCCCCCCGTCGGAAGAACCTTTGGCTCTTTACCTCTAGGGGTCGGT

AGGTCAACTGCTAAGCCAGAAGAACTACTAAGGGAGGCCACAGAACTAGACATAGTGGTGAGGCGCACTGCAGGAGTAAA

TGAGAAACTGGTATTTTACAACAACACTCCGCTGTCCTTGTTAACGCCCTGGAAGAAAGTTCTGACAACCGGAAGTGTGT

TTAGCGCTAACCAGGTTTGCAATGCAGTCAACCTAGTCCCGCTTGATACTCCCCAGAGGTTCAGGGTTGTGTACATGAGC

ATAACTAGATTATCAGACAATGGTTACTATAGTGTGCCCAGGAGAATGCTAGAGTTCCGCTCAGCCAATGCAGTAGCCTT

CAACATCTTGGTTACACTAAGAATTGAGAATGGCACAAACCCTAGAAGATACATAGTCGGCTCATGGGAGAATTCAGAGG

TCACATTTATGGTACACGTGGGCAACTTTAGAAGAAAGAAGAACGAAGTATACTCTGCTGATTACTGCAAAATGAAGATT

GAAAAGATGGGTTTAGTTTTTGCCCTGGGAGGAATAGGTGGAACAAGTCTCCATATTAGAAGCACCGGGAAAATGAGCAA

AACCCTCCATGCACAGCTGGGGTTCAAGAAAATCTTATGTTACCCCCTAATGGATGTTAATGAGGATCTTAACCGATATC

TCTGGCGGGCAGAGTGCCGAATAGTCAAAATCCAGGCCGTCTTACAGCCATCAGTACCCCAAGAATTCCGTGTCTACGAT

GATGTCATCATCAACGATGATCAAGGCTTGTTCAAGATCTTGTAGTTCATTTGCAAAGGAATGAACCCACCCCCAGGAAG

GGGCCCTACCCTCCCCCG-CCAAAAAACACACCCCAAGGCTCCCCGACACAGCGCCAACACAGCCACTCTCCCGATGGGA

CGAGGTTGGCC-CCCCAACAAACCCGGTCCAGGC-GGGGGGCCCCCCCCCTGACCCGCCGGGCGCCGACCGAGCAGGCCG

GCGAGAAGCCCCCAAGAGGCCCCGACGAAGACCGAGCACCCCTCCCCCCC-AAACCGTTCCCCC-AACACGCCCGAGGCC

AGGCGCCGGGCACCCCCAGACCCGCCACCGGCGGGACAGAACCCCAGGGGGGAACCCCCCAAACCAAAGGGACCACCGGG

GAAGGATCCGCCGAACCAGCACAGACCCGACCCAAGCAAGGCGGACCCCCAACACGAAACCGCCCAGCACACCCCACTCA

TCATCAAACAAAACTTAGGGGCCAAGCCCATACGACAGGCCCCAGTGCGCCAGGCGGAGGGACGCAGAAAGGAAGGAGAC

AAGACCGCCCCAGAAGGAGGACACCCAACCACCGAAACACCCAGCCCCAGGGAAGGAG--CCCCCCCCCACATGGCGAAA

GAACGGGCAAGATACCCCGAGAAGAACCCGACCGGGAGCCAGGAGCCGGCGACGCCGGCGGCAAACAAGCCGGGCAAACA

CCCGACCCCACCAGAGGAGCCGAGGAGAGCCCGACCCCGCACGAACCAAACAGCAGGGAGCCCAACCCCACCAAAAAAAG

AGGCGAGGCCCC-CCT-----CCTCCCCGCACACAACCCGCAGACAGACCGAGAGAGGACCACCTCCCAACAAGACAGGC

GGCTCCCTCGCAGCCGAGCCGCAGCTCCCCAGAACAGGACACCCCCGTACACTCAAAACACCCCCCCGCCCCCCGCCACC

GGGACAGGGGCTCCGCCACCACAGACCAAAGCCACCGAGAGCAAGCCAGAGGTCCCCCCCAGTCCCGCCCGGCGCCCCCC

CGAAAAGGCCCCGGCAAGGCCCCCCAACCCGACGTCCCCCGCACGCACCGAAAGAGGGACCCGCCCGAGAACGTGCGGCA

CATCCATGCGTAAACATTATGACACGGGTTGCAATCTTGACATCTCTGTTTCTTCTCCCAAATGTTGTTGCGTGCCAGAT

TCACTGGGGCAATCTATCCAAGATCGGGATTGTGGGAACAGGGAGTGCCAGCTACAAGGTGATGACTAGGCCGAGCCACC

AGACTCTGGTTATAAAGTTAATGCCAAATATAACAGCCATCGACAATTGTACAAAGTCAGAGATTGCAGAGTACAAGAGA

TTGCTGATCACAGTGTTAAAGCCTGTAGAGGATGCTCTGTCGGTTATAACCAAGAATGTAAGACCAATTCAAACCCTGAC

ACCTGGGCGTAGAACCCGTCGTTTTGCTGGAGCTGTTCTGGCCGGAGTAGCGCTTGGAGTTGCGACAGCCGCTCAGATAA

CTGCAGGAGTCGCTCTTCATCAATCATTGATGAACTCCCAAGCAATTGAGAGTTTAAAAACCAGTCTTGAGAAGTCGAAT

CAGGCAATAGAAGAAATCAGACTTGCAAATAAGGAGACCATACTGGCAGTACAGGGCGTCCAGGATTATATCAACAATGA

GCTTGTCCCTTCTGTTCATAGAATGTCATGCGAGCTGGTAGGCCACAAGCTCGGCCTCAAGCTCCTTAGGTACTACACCG

AGATCCTGTCCATATTCGGGCCCAGCCTTCGAGACCCGATAGCTGCCGAAATATCAATCCAGGCACTCAGTTATGCATTA

GGCGGAGACATTAATAAAATCCTGGACAAGCTTGGGTATAGCGGTGGGGATTTCCTTGCCATCCTAGAAAGCAAGGGAAT

AAAGGCCCGGGTTACATATGTGGACACAAGAGATTACTTTATAATGCTTAGCATCGCCTACCCAACCTTATCTGAGATCA

AGGGAGTGATAGTTCACAAGATAGAAGCTATAACATACAACATTGGGGCACAGGAGTGGTATACTACTATCCCTAAATAT

GTAGCCACTCAGGGATATCTGATATCGAACTTTGATGAGACGTCATGCGTATTCACTCCAGAGGGGACAGTTTGCAGCCA

GAATGCATTGTACCCAATGAGCCCATTGCTTCAGGAATGTTTCAGGGGGTCAACAAAATCGTGCGCCAGAACCCTAGTTT

CAGGTACCATAAGTAATAGATTTATCCTATCAAAAGGAAACCTGATCGCGAATTGTGCGTCAGTTTTGTGCAAATGTTAC

ACAACGGAGACAGTTATCAGCCAAGATCCTGACAAACTACTAACTGTTGTAGCATCCGACAAGTGTCCTGTAGTTGAGGT

GGATGGAGTGACAATACAGGTCGGCAGTCGAGAGTATCCGGATTCTGTATACTTACACAAAATAGACTTAGGCCCAGCCA

TCTCCCTAGAAAAACTAGATGTAGGTACCAATTTAGGCAATGCAGTTACAAGACTGGAGAATGCAAAGGAGCTCCTAGAT

GCATCAGACCAAATACTGAAGACTGTCAAAGGGGCACCTTTCGGTGGGAATATGTACATAGCACTGGCAGCTTGCATTGG

AGTATCCCTAGGGCTTGTCACATTAATATGCTGCTGTAAGGGGAGGTGTAAGAACAAGGAAATCCCTATCTCCAAAATCA

ACCCAGGGCTCAAACCCGATCTGACCGGGACCTCAAAGTCGTACGTGAGATCACTGTAGTCAGAATCACCTGAATCATCC

GGCATCACACACATACATGTACGACACAAGCAGTCTGAGGACGCAGGAAACCCAGCCCCCGATCACCCACCTGACCCTAC

TCCACGCTCCACTACACATTAGTCATTAAACAAAACTTAGGACGAAAGGTCAATCACCATGTCCGCACAAAGAGAAAGGA

TCAATGCCTTCTACAAAGGCAATCCTCACAATAAGAACCATAGGGTGATCCTGGATAGAGAACGCTTGATCATTGAAAGA

CCCTACATCTTGCTTGGAGCCCTGCTGGTAATGTTCCTGAGTCTAATCGGACTGCTGGCCATTGCAGGGATCAGGCTTCA

CCGGGCCACCGTTGGAACTTCAGAGATCCAGAGTCGGCTGAATACCAATATTAAGTTGACCGAATCTATTGATCACCAGA

CTAAGGATGTCTTAACTCCCCTTTTTAAAATCATTGGCGATGAAGTCGGCATCAGGATTCCACAGAAATTCAGTGATCTT

GTCAAGTTCATCTCCGACAAGATTAAATTCCTCAACCCCGATAGAGAGTATGATTTCAGGGATCTCCGGTGGTGCATGAG

TCCCCCCGAGAGAGTCAAAATTAATTTTGATCAGTTTTGTGAGTACAAAGCTGCGGTTAATTCAATTGAACATATATTTG

AGTCACCGCTCAACAAGTCAAAAAAGCTGCAATCATTGACTCTTGGGCCCGGAACAGGCTGTCAAGGCAGGACAGTAACA

AGAGCTCATTTCTCAGAACTTACACTGACCTTAATGGACTTGGATCTAGAGATGAAGCACAACGTGTCCTCAGTGTTTAC

CGTAGTTGAAGAGGGGTTATTCGGAAGAACATATACCGTCTGGAGATCCGATGCCAGGGATCCGAGCACCGATCTAGGTA

TCGGCCATTTTTTAAGAGTCTTCGAGATTGGACTGGTAAGAGATCTCGGGCTGGGTCCCCCTGTTTTCCATATGACCAAC

TATCTCACAGTGAACATGAGTGATGTCTATCGGAGATGTCTTTTAGCGGTAGGGGAGTTGAAGTTGACAGCCCTATGCAC

CTCATCTGAGACTGTAACACTGAGTGAGAGAGGAGTTCCAAAGAGGAAGCCTCTTGTGGTTGTGATACTTAACCTGGCTG

GGCCCACTCTAGGGGGCGAACTATACAGTATCTTGCCTACCTCTGATCTCATGGTGGAGAAACTCTATTTATCTTCACAT

AGAGGGATTATCAAGGATGACGAGGCCAATTGGGTAGTGCCGTCCACCGATGTTCGTGATCTTCAGAACAAAGGTGAATG

TCTGGTGGAAGCATGCAAGACTCGACCTCCTTCATTTTGCAATGGCACAGGATCAGGCCCGTGGTCAGAGGGGAGAATCC

CTGCTTACGGGGTGATCAGGGTCAGTCTTGACTTAGCTAGTGACCCGGATGTAGTTATCACTTCAGTGTTTGGCCCACTG

ATTCCTCACCTATCCGGCATGGATCTTTACAACAACCCGTTTTCAAGAGCTGTATGGTTGGCTGTACCACCTTACGAGCA

GTCATTTCTAGGAATGATAAATACAATTGGATTCCCTAACAGAACAGAGGTTATGCCGCACATTTTGACCACAGAGATCA

GAGGCCCTCGGGGTCATTGCCATGTCCCCATAGAATTGTCCCGCAGGGTTGATGACGATATCAAGATCGGGTCCAACATG

GTCATATTGCCGACGATGGACCTGAGATATATTACAGCCACTTATGATGTTTCCAGGAGCGAGCATGCAATCGTGTACTA

TATCTATGACACAGGTCGCTCATCATCTTACTTCTACCCAGTTCGACTGAATTTCAAAGGCAATCCTCTCTCTCTGAGGA

TAGAGTGTTTCCCTTGGCGTCATAAGGTGTGGTGCTACCATGATTGTCTTATATACAACACCATAACAGATGAAGAGGTC

CATATGAGAGGGCTGACCGGTATAGAGGTAACATGCAATCCAGTCTGAGCAGAGCTACGACCGTCACTCAAGCAGTCCCT

TGAGTCGCCACCGAGTCCAAGCAGCACAGCCTGGGACTCTCAACAGCACAGCCCAGCTAACAGTGTTATAAAAAACTTAG

GAGCCAGGGTTGTAGGGGCCATGGACTCCCTATCAGTCAATCAGGTCTTGTACCCTGAGGTTCATCTAGACAGCCCTATT

GTCACAAACAAACTAGTTGCCATCCTTGAGTACTCGGGTATCGACCATAACTATGTTCTTGAAGACCAGACCCTTGTCAA

GAATATTAGGTATAGACTGGGGTGCGGTTTTTCAAATCAAATGATCATCAATAACAGGGGGGTGGGTGAAACAGTCAATT

CTAAACTTAAAAGTTATCCCCATAATCGTCATATCATATACCCGGATTGCAATAAGGAGTTGTTTTGTATCAAAGATAGC

TGCATATCTAAGAAGCTCTCGGAGCTATTCAAGAAGGGTAATTCCTTGTACTCTAAGATAAGTCACCAGGTACTGGACTG

TCTTAAGAGAGTCAATGGGAAATTAGGCCTGGGCACAGATCTTACCCATGGTCTGAAGGAGGGTATCCTTGACCTAGGGT

TGCACATGCATAGCTCTCAATGGTTCGAGACTTTCCTGTTCTGGTTCACTATCAAGACAGAGATGAGATCAATGATCAAA

GAACAGTCCCATATATGCCACAAGAGGAGATATAACCCGATTTTTGTATCAGGGGATGCATTCGAGGTGCTCGTATCACG

AGACCTCGTTGTGATAATTGATAAGAATACCCAGTATGTCTTCTACTTAACATTTGAGCTGGTCCTTATGTATTGTGATG

TCATAGAGGGCAGACTTATGACGGAGACAGCCATGGTCATAGACCAGAGATATTCAGAACTTCTAAGCCGGGTTAGATAC

CTGTGGGATCTTATTGATGGGTTCTTCCCAACACTAGGCAATACCACCTACCAAATTGTTGCTCTGCTTGAACCATTATC

GTTGGCTTATCTTCAACTTCAGGATGTCACGCTGGAGTTAAGAGGTGCTTTTTTAGACCATTGCTTCAAAGAACTCTATG

AGATACTGGAGCATTGCGGCATTGACACAGAAGGCACCTATAATTCCATCACTGAAGGATTGGATTACGTATTCATCACC

CACGATATACATCTAACTGGGGAGATTTTTTCATTTTTCCGGAGTTTCGGACACCCCCGTCTAGAAGCAGTCACCGCCGC

AGAAAATGTCAGAAAACATATGAATCAACCGAAGGTGATCAGTTATGAGACTATGATGAAAGGGCATGCAGTATTTTGCG

GGATAATCATAAATGGTTTTAGGGACCGACATGGCGGCAGCTGGCCCCCTGTTGCATTGCCGGAACATGCTTCTGCTGCG

ATCCGGAATGCGCAGGCATCAGGTGAGGGACTGACCCATGACCTGTGTATAGACAACTGGAAATCCTTTGTTGGATTCAA

ATTTGGCTGCTTTATGCCACTCATCCTAGATAGTGATTTGACTATGTACCTCAAAGACAAGGCATTGGCTGCACTGAAGA

ATGAGTGGGATTCAGTTTACCCGAAAGAATACCTCCGTTATAACCCACCTAGAGGGACAGAGTCAAGGCGACTGGTGGAG

GTGTTCCTGAATGACTCCAGCTTTGATCCTTATAACATGATAATGTACGTGGTGAATGGCTCCTACCTTAAAGACCCTGA

GTTTAATCTCTCATACAGCCTAAAAGAGAAGGAGATAAAGGAGACAGGGCGGTTGTTTGCCAAAATGACCTATAAGATGC

GGGCCTGTCAGGTAATTGCTGAGAATCTGATATCGAATGGTGTTGGGAAGTACTTCCGAGACAATGGGATGGCGAAGGAC

GAGCATGACCTAACAAAAGCCCTTCACACTTTGGCAGTCTCAGGTGTTCCCAAGAACAACAAAGATAACCGCCGAGGTGG

GCCTCCCAGAAGGACAACAACCCGAGAGGTGAGATCAAGCCGGGACACCAGAACACAAAATAGAGACAAGGTCCAAGGGG

GACCTATGTACAACTATTTGCAATGCCAACCAATCAGCCCTGATCAAGGTGAGTCATACGAGACTGTTAGTGCATTTATC

ACTGCTGACCTTAAGAAGTATTGCCTGAATTGGAGATACGAGACAATCAGCATATTTGCACAGAGGCTGAATGAAATATA

TGGACTACCATCCTTCTTTCAATGGTTGCACAGGATATTGGAAAAATCCGTACTCTACGTCAGTGACCCACATTGCCCTC

CCGATCTAGATAATCATATCCCTCTGGACAGTGTCCCTAATGCCCAAATATTCATTAAGTACCCAATGGGTGGAATAGAA

GGTTATTGCCAAAAACTATGGACAATCAGTACTATACCATACTTGTATCTGGCAGCCTATGAGAGCGGAGTAAGAATCGC

CTCACTGGTGCAGGGTGACAATCAGACAATCGCAGTGACAAAAAGAGTTCCAAGTTCTTGGCCTTATTCACTAAAAAAGA

GGGAGGCATCCAAAGCCGCTCAAAATTACTTCGTAGTCTTAAGGCAAAGATTGCACGATGTAGGTCATCACTTGAAGGCT

AATGAAACCATAGTATCTTCTCACTTTTTTGTATATTCCAAAGGGATTTATTATGACGGCCTGCTAGTCTCACAATCACT

AAAGAGCATCGCCAGATGTGTCTTCTGGTCCGAGACTATAGTGGATGAAACCAGAGCGGCTTGCAGCAATATCGCAACAA

CTGTCGCTAAGAGTATAGAGAGGGGTTATGATAGGTACCTTGCATATTCTTTGAATATCCTCAAGATTTTCCAACAGATC

CTTATATCCCTTAACTTCACTATTAACACAACAATGACTCAGGATGTCGTGGCACCGATCGTCGAGAACGGTGATTTGCT

GATAAGGATGGCACTCTTGCCAGCACCCATCGGAGGTCTAAATTATCTTAACATGAGCAGGCTATTCGTGAGAAATATCG

GCGACCCGGTCACCTCCTCTATAGCCGACCTGAAGAGGATGATAGATGCCGGGCTCATGCCAGAAGAAACATTGCATCAA

GTGATGACCCAGACCCCGGGAGAATCATCCTACCTTGATTGGGCAAGTGACCCTTATTCTGCCAACCTACCCTGCGTACA

GAGTATAACTCGTCTTCTCAAAAACATCACTGCACGATATATTTTAATCAGCAGCCCAAATCCGATGCTGAAAGGATTGT

TTCATGAGGGGAGTAGAGATGAAGACGAGGAGCTTGCAAGTTTCCTAATGGATCGGCATATAATTGTTCCAAGAGCTGCA

CATGAAATCTTAGACCATAGTATAACCGGGGCAAGAGAGGCTATAGCCGGGATGTTGGACACCACCAAGGGTCTGATTAG

AACAAGTATGAAACGGGGTGGCCTCACCCCTCGAGTCTTAGCCCGCCTTTCCAATTATGATTATGAACAATTCCGATCCG

GGATAACACTATTGACAAAGAAAGGGCAGTGTTATCTCATTGACAAGGATTCGTGCTCGGTGCAGCTCGCTATAGCTCTG

AGGAGCCACATGTGGGCTAGGTTGGCTCGCGGGAGACCAATCTATGGGTTGGAGGTGCCCGATATATTAGAATCGATGAA

CGGCTACCTTATCAAACGCCACGAGTCTTGTGCTATCTGTGAAACGGGCTCAAGTCACTACGGATGGTTTTTCGTCCCTG

CAGGGTGCCAGCTTGACGATGTATCTAGAGAGACTTCGGCTCTCCGTGTACCTTATGTCGGATCAACAACTGAGGAAAGG

ACAGACATGAAACTTGCATTTGTTAGATCTCCAAGTAGATCCCTCAAATCAGCAGTTAGAATTGCAACAGTTTACTCATG

GGCCTACGGGGATGATGAGAAGTCATGGAGTGAAGCCTGGATGCTAGCTAGGCAGAGAGCCAATATCACCTTAGATGAAT

TGAGAATGATCACTCCAGTCTCTACATCCACCAACCTAGCCCACCGGTTGAGGGATCGGAGCACCCAGGTGAAATACTCG

GGGACATCCCTTGTGAGGGTTGCAAGATACACAACCATCTCCAATGATAACTTGTCATTTGTGATATCTGAGAAGAAAGT

AGATACCAACTTCATTTATCAGCAAGGGATGTTGCTTGGTCTCGGGATCCTTGAGAACCTCTTTAGGCTAGAGGCCACCA

CCGGAGTATCCAACACAGTGCTACACCTGCACGTGGAAACAGAATGTTGTGTTGTACCTATGGTGGATCACCCAAGGATA

CCGAGTCTCCGTAATATAAAAGTTACGGATGAGCTATGCACAAACCCTTTGATCTATGATAGGTCTCCCATCATAGAGCA

CGATGCAACCCGACTATACTCACAGAGCCACAGGAGACATTTGGTGGAGTTTGTTACCTGGTCCACAAGCCAGCTTTACC

ATATACTGGCTAAATCTACAGCAATGTCCATGATTGAATTGATCACGAGATTCGAGAAAGATCACATGAATGAAATAGCC

GCCCTGATTGGCGATGACGACATCAACAGCTTCATCACAGAATTTTTGCTAGTGGAGCCCAGATTATTTATAGTTTACCT

TGGTCAGTGTGCTGCCATCAATTGGGCCTTTGATATACATTATCATCGGCCCTCGGGCAAGTACCAGATGGGGGAACTCC

TCTACTCTCTGCTCTCACGGATGAGCAAAGGAGTATATAAGATCTTCACCAATGCTTTGAGCCACCCCAAAGTTTACAAG

AAATTTTGGCGAAGTGGTGTCATTGAGCCGATTCATGGCCCATCCCTGGATACACAGAATTTACATGTCACTGTTTGTGA

CATGATATATGGGTCCTATGTCACCTACCTGGATCTTTTGCTGAATGATGAGCTAGATGATTACCCGTATTTGCTCTGCG

AGAGTGATGAGGATGTGGTCACAGACAGGTTCGACAACATTCAAGCCAAACATCTCTGTGTATTAGCCGATGTATATTGC

AGCTCCAAGAGGTGTCCTTCCATAATCGGGATGTCTCCTATAGAAAAATGTACCATTCTCACACATTACATCAAGGGAGA

ATCAGTACAATCCCCGTCTGGAACCTCATGGAACACTGATCCCCTTGTAGTAGATCATTACTCATGCTCTCTGACCTACC

TTCGCCGAGGTTCCATCAAACAAATCAGGTTGAGGGTGGATCCTGGGTTTGTATTCGAGGCGTTGACAGACGTCGACTTT

AAACAACCTCGCAAAGCTAAGTTAGATATATCGGTCGTTGGGTTGACTGATTTCTCTCCCCCTTGCGACAACGTCGGTGA

TTTTCTAGGGACTATCAACACATTGAGGCACAATCTGCCTGTCACCGGAACCGGGGTCTCGAACTATGAAGTCCACGCTT

ATCGTAGAATTGGCCTGAATTCATCAGCATGTTACAAAGCCGTAGAGATCTCCACGTTAATCAAGCCATCCCTAGAAGTC

GGAGAGCATGGATTGTTCTTAGGAGAAGGTTCCGGTTCAATGCTGGCTGCGTACAAGGAAGTTCTTAAATTAGCAAATTG

TTATTACAACAGCGGGGTAACAGCGGAGGGCAGAGCCGGACAGAGGGAAATATCTCCCTATCCTTCAGAGATGAGCCTAG

TAGAGAATCAAATGGGGATAGAGCGGAGTGTTAAAGTGCTGTTCAACGGCAAACCTGAAGTAACTTGGGTGGGGACCACC

GATTGCTACAAGTACATAATCAGTAACATCCAAACCTCTAGTCTGGGTTTCATACATTCAGATATTGAGACGCTCCCAAC

CAAAGATGCTGTTGAGAAATTAGAAGAATTTGCCTCTATCCTATCCTTATCCCTGATTTTGGGGAAAATCGGCTCTATTA

CAGTTGTCAAAATTATGCCCATTAGCGGGGATTTCACCCAAGGCTTCATAGCCTATGCCATCCAATATTTTAGGGAGAGC

CTGCTTGCCTATCCGAGATATAGTAACTTTATCTCGACTGAGTGTTACCTTATCATGATAGGATTAAAGGCCAATCGATT

GATTAACCCAGAAGCCATTAAGCAAAGCATAATCAGAGCGGGGGTCAGGACTGCACCAGGACTTGTGAGCCATATATTAT

CAGAGAAACAGAAAGGTTGTATTCAATCTTTTCTTGGTGATCCTTATATCCAAGGAGACTTCAATAAGCACCTTAAATCT

CTAACCCCTATTGAGAAAATCCTGGTAAATTGTGGTCTCTCGATCAATGGCACAAAAATCTGTAGAGATCTAATCCACCA

TGATATCGCCTCCGGTCCAGACGGTCTTATGAGCTCTACAATTATTTTATATAGGGAATTGGCTCATTTCAAAGATAATA

TAAGGAGTCAGCACGGAATGTTCCACCCCTACCCAGTACTGGCCAATAGCCGGCAACGTGAATTAATCCTTCGAATAGCC

AAGAAATTCTGGGGGTATGTCTTGCTATATTCTGATGACCCGGCACTAATCAGACAAACAATCAAGAACTTGAAACGGAA

TCATCTAACCTTTGACTTACATAGTAACCCGTTTATTAAAGGGCTATCCAAAGCTGAGAAACTGCTGGTGCGGACGAGTT

CACTAAGAAGGGAATGGTTGTTCACTCTCGATACGAAAGAAGTGAAAGAGTGGTTCAAATTGGTAGGTTACAGTGCACTC

ATCAGAGGCTAATTGGTTATACATCTGCCCCCTTCTCCTCCGCCATGAGACCCCACTGACGATTCAGAAGATTAAAGAAA

ACTACATATTGGATAAGTATCTATTCCCAGCTTTGTCTGGT

>MN369542.1/MountainGazelle/UAE/2018-08-14

ACCAAACAAAGTTGGGTAAGGATCGATCTTATAATAACTATGAACTAGCAAACTTAGGAGTAAAGATCCTACTGTCGGGG

GGAGGAGGAGGAGCAAGATCTCTGATTATGGCGACTCTCCTTAAAAGCTTAGCATTGTTCAAAAGGAACAAAGACAAAGC

GCCGACGGCATCAGGTTCAGGAGGGGCCATCCGAGGGATTAAGAATGTTATCATAGTCCCGATTCCCGGAGACTCGTCCA

TCACTACCCGTTCAAGACTGCTCGACAGGCTTGTCAGATTGGCCGGAGATCCTGACATCAACGGGTCAAAGCTGACCGGC

GTGATGATCAGCATGTTATCTTTGTTCGTAGAGTCACCCGGGCAGTTGATACAGCGAATCACAGATGATCCGGATGTTAG

TATCCGCCTTGTTGAGGTAGTTCAAAGTACAAGGTCCCAGTCCGGGTTGACCTTTGCATCACGTGGTGCTGATTTGGATA

ACGAGGCAGACATGTATTTTTCAACTGAGGGGCCCTCGAGTGGAGGTAAGAAAAGGATCAACTGGTTTGAGAACAGAGAA

ATAATAGACATAGAGGTGCAGGATCCAGAAGAGTTCAATATGTTGTTAGCCTCCATACTAGCACAAGTCTGGATCCTTCT

GGCCAAGGCTGTCACGGCACCGGATACGGCAGCTGACTCAGAACTGAGAAGGTGGGTTAAATACACACAACAAAGGAGAG

TGATTGGGGAATTTCGCCTTGACAAAGGGTGGCTGGACGCAGTCCGCAACAGAATTGCAGAAGATCTATCACTCCGGCGG

TTCATGGTATCTCTCATACTTGACATCAAGAGGACCCCCGGCAACAAGCCGAGGATCGCAGAAATGATCTGCGACATTGA

CAACTATATTGTCGAAGCAGGACTCGCCAGTTTCATCCTTACTATCAAATTTGGTATAGAAACAATGTATCCTGCATTAG

GGCTCCACGAGTTTGCTGGGGAATTGTCCACTATTGAGTCCTTGATGAACTTGTATCAACAGCTAGGAGAGGTTGCACCC

TACATGGTAATTCTAGAGAACTCAATTCAGAACAAGTTTAGTGCAGGAGCTTATCCCCTCCTCTGGAGCTATGCGATGGG

TGTCGGAGTTGAGCTGGAGAACTCAATGGGGGGCTTGAACTTTGGCAGATCATATTTTGACCCTGCCTATTTTCGTCTCG

GACAGGAGATGGTCAGAAGATCTGCAGGAAAGGTCAGCTCTGTGATTGCGGCTGAGCTCGGCATCACAGCAGAGGAAGCT

AAACTAGTCTCGGAAATAGCCTCGCAGGCTGGGGACGAAAGAACCGCTAGAGGGACTGGGCCTCGACAGGCGCAGGTCTC

CTTCCTACAGCACCAAACAGGAGGGGGAGAGTCGTCCACACCAGCGACCAGAGAAGGGGTCAAAGCTGCGATCCCAAACG

GATCCGAAGAAAGGGACAGAAAGCAAACACGCTCAGGGAGGCCCAGAAGAGAGACCCCCGGCCAACTGCTCCTGGAATTC

ATGCCCGAGGATGAGGTCTCGCGAGAATCTGGTCAAAACCCTCGTGAGGCTCAAAGGTCGGCCGAGGCACTCTTTAGGCT

GCAGGCCATGGCCAAGATTCTGGAGGACCAGGAGGAGGGAGGAGACGACAATCAGGTCTACAACGACAAGGATCTCCTCG

GCTAAGCAGACGCACCCTCTGTCGAAATCAGTGATGAGACATCTCCTATCAGTATTATAAAAAACTTAGGACCCAGGTCC

AAGCAAGCACACATCGACACTCCAACCAGTCGAGCGGAGGCTACCGATGGCAGAAGAACAGGCATACCATGTCAACAAGG

GGCTGGAATGTATCAAGTCCCTCAAAGCCTCTCCCCCGGATCTATCCACCATTAGAGATACCATCGAGAGCTGGAGAGAG

GAGCTTAGCCCCTCGGGCCGTGCAACACCGAACCCTGATACGTCCGAGGGAGATCATCAGAATATCAACCAATCATGCTC

ACCAGCAATCGGACCAAACAAAGTCTACTTGTCTCCTGAAGATAATCTCGGATTTAGAGAGATCACTGGCAACGACTGTG

GGGCTGAGCTCGGAGGAGTCCAGGGGAAAGGATCCAACTCTCAAGTACAGCGTTACTATGTTTATAGCCACGGGGGTGAA

GAGATTGAAGGACTCGAGGATGCTGACTCTCTCGTGGTTCAAGCAGATCCTCCAGTTGCTAACATATTCAATGGAGGAGA

GGATGGATCTGACGACAGCGATGTGGACTCTGGCCCAGATGATCCCAGCAGAGATACTCTATATGACCGGGGACCTGCTG

CCGGCAATGATGTCGCTAGGTCCACAGATGTCGAAAAACTAGAAGGTGCTGATATTCAAGAAGTTCTTAACTCCCAGAAA

AGCAGAGGAGGAAGATTCCAAGGCGGGAAAACCTTACGAGTCCCGGAAGTACCCGATGTCAAGAACTCCAGACCATCAGC

CCAATCAATTAAAAAGGGCACAGACGGGAACTCAGTCTCATCTGGAACGGTGACAGAGTGTTTATCGATAAGTGGTGCAA

CCCAAGCTGTGCCAGAGTCAAGATGGGAGTCATCAGAGCAAAATGCGTCTGTGGGGAGTGTCCTCAAGTCTGCGAAGAGT

GCAAAGACGATCCAGGGGTCGACACAAGAATCTGGTACCACAGCATCACCGACTCAGCCTAAAGAGAATGACTCCGAGTA

TGAGTATGAGGATGACCTATTTACAGAGATTCAGGACATCCGTGCAAGCATTGCCAAGATCCATGATGACAACAAGACTA

TCCTCTCAAAGCTTGATTCTATACTGTTATTGAAAGGAGAAGTCGACACTATCAAGAAACAAATCAGCAAGCAGAATATA

AGTATATCCACCATTGAGGGCCATCTCTCCAGTATAATGATAGCCATCCCGGGCTTTGGGAAGGATATCAAGGACCCAAC

ATCCGAGGTTGAGTTGAACCCAGATTTAAGACCTATAATAAGCCGTGACTCTGGCAGAGCTCTCGCGGAGGTCCTCAAGA

AACCCGCTGTGGATAGGTCTCAGAAAACTGGAACAAAAGCCAACTCCAGCTCAAAGGGTCAGCTTCTTAAGGATCTCCAG

CTAAAACCTGTCGACAAGCAGGCAAGCTCTGCAATCGGGTTTGTCCCATCCGACCATGAATCATCCAGAAATGTCATCCG

CTCCATAATCAAGTCGAGCAAGCTAAACATTGATCACAAGGACTATCTTCTAGATTTACTGAATGATGTGAAAGGCTCCA

AGGATCTTAAGGAATTCCACAAGATGCTAACAGCAATTCTCGCCAAGCACCCGTAACACATCCCCAAGTCACCATCTCAT

ACTCGGCTGAAGACATCCTTTCAATCAGGCTATTACAAAAAACTTAGGAGCAAGGGCAACTGAGCTTCACAGACAAGATG

ACCGAGATCTACGACTTCGACAAATCAGCATGGGATGTCAAAGGGTCAATTGCCCGCATAGAACCCACCACCTACCACGA

CGGCCGACTGATACCCCAGGTAAGGGTCATCGATCCTGGTCTGGGAGACAGAAAAGATGAGTGCTTTATGTACCTGTTTC

TCCTGGGAGTGATTGAGGATAACGACCCCCTGTCTCCCCCCGTCGGAAGAACCTTTGGCTCTTTACCTCTAGGGGTCGGT

AGGTCAACTGCTAAGCCAGAAGAACTACTAAGAGAGGCCACAGAACTAGACATAGTGGTGAGGCGCACTGCAGGAGTAAA

TGAGAAACTGGTATTTTACAACAACACTCCGCTGTCCTTGTTAACGCCCTGGAAGAAAGTTCTGACAACCGGAAGTGTGT

TTAGCGCTAACCAGGTTTGCAATGCAGTCAACCTAATCCCGCTTGATACTCCCCAGAGGTTCAGGGTTGTGTACATGAGC

ATAACTAGATTATCAGACAATGGTTACTATAGTGTGCCCAGGAGAATGCTGGAGTTCCGCTCAGCCAATGCAGTAGCCTT

CAACATCTTGGTTACACTAAGAATTGAGAATGGCACAAACCCTAGAAGATACATAGTCGGCTCATGGGAGAATTCAGAGG

TCACATTTATGGTACACGTGGGCAACTTTAGAAGAAAGAAGAACGAAGTATACTCTGCTGATTACTGCAAAATGAAGATT

GAAAAGATGGGTTTAGTTTTTGCCTTGGGAGGAATAGGTGGAACAAGTCTCCATATTAGAAGCACCGGGAAAATGAGCAA

AACCCTCCATGCACAGCTGGGGTTCAAGAAAATCTTATGTTACCCCCTAATGGATGTCAATGAGGATCTTAACCGATATC

TCTGGCGGGCAGAGTGCCGAATAGTCAAAATCCAAGCCGTCTTACAGCCATCAGTGCCCCAAGAATTCCGTGTCTACGAT

GATGTCATCATCAATGATGATCAAGGCTTGTTCAAGATCCTGTAGTTCATCTGCAAAGGAACGAACCCACCCCCAGGGGG

GGGCCCC-----CCCCCG-CCAGAAAACACACTCCAAGGCTCCCCGACACAGCGCCAACACAGCCACCCCCCCCACAGGA

CGAGGCCGGCC-CCCCAACAAACCCGATCCAGGCGGGGGGGGCCCCCCCCCGACCCGCCGGGTGCCGACCGAGCAGGCCG

GCGAGAAGCCCCCAAGAGGCCCCGACGAAGACCGAGCACCCCTCCCTCCC-AAACCGTCCCCCC-AACACGCCCGAGGCC

AGGCGCCGGGGACCCCCAGACCCGCCACCGGCGGGACAGAACCCCAGGGGGGGACCCCCCAAACCAAAGGGACCACCGGG

GAAGGATCCGCCGAACCAGCACAGACCCGACCCAAGCAAGGCGGACCCCCAACACAAAACCGCCCAGCACACCCCACTCA

TCACCAAACAAAACTTAGGGGCCAAGCCCATACGACAGGCCCCAGTCCGCCAAGCGGAGGGACGCAGAAAGGAAGGAGAC

ACGACCGCCCCAGAAGGAAGACATCCAACCACCGAAACACCCAGCCCCAGGGAAGGAG--CCCCCCCTCACATGGCGAAA

GAACGAGTGAGAGACCCCGAGAAGAACCCGACCGGGAGCCAGGAGCTGGCGACGCCGGCGGCAAACAAGCCGGGCAAACA

CCCGACCCCACCAGAGGAGCCGAGGAGAGCCCGACCCCGCACGAACCAAACAGCAGGGAGCCCAATCCCACCAAAAAAAG

AGGCGAGGCCCC-CCT-----CCTCCCCGCACACAACCCGCGGACAGACAGAGAGAGGACCACCTCCCAACAAGACAGGC

GGCCCCCTCGCAGCCGAGCCGCAGCTCCCCAGAACAGGACACCCCCGCACACTCCGAACACCCCCCCGCCCCCCGCCACC

GGGACAGGGGCTCCGCCACCACAGACCAAAGCCACCGAGAGCAAGCCAGAGGTCCCCCCCAGTCCCGCCCGGCGCCCCCC

CGAAAAGGCCCCGGCAAGGCCCTCCAACCCGACGTCCCCCGCACGCACCGAAAGAGGGACCCGCCCGAGAACGTGCGGCA

CATCCATGCGCAACCGTTATGACACGGGTCGCAATCTTGACATCTCTGTTTCTTCTCCCAAATGTTGTTGCGTGCCAGAT

TCACTGGGGCAATCTATCCAAGATCGGGATTGTGGGAACAGGGAGTGCCAGCTACAAGGTGATGACTAGGCCAAGCCACC

AAACTCTGGTTATAAAGTTAATGCCAAATATAACAGCCATCGACAATTGTACAAAGTCAGAGATTGCAGAGTACAAGAGA

TTGCTGATCACAGTGTTAAAGCCTGTAGAGGATGCTCTGTCGGTGATAACCAAGAATGTAAGACCAATTCAAACTCTGAC

ACCTGGGCGTAGAACCCGTCGTTTTGCTGGAGCTGTTCTGGCCGGGGTAGCGCTTGGAGTTGCGACAGCCGCTCAGATAA

CTGCAGGAGTCGCTCTTCATCAATCATTGATGAACTCCCAAGCAATTGAGAGTTTAAAAACCAGTCTTGAGAAGTCGAAT

CAGGCAATAGAAGAAATCAGACTTGCAAATAAGGAGACTATACTGGCAGTACAGGGCGTCCAGGATTATATTAACAATGA

GCTTGTCCCTTCTGTTCATAGAATGTCATGCGAGCTGGTAGGCCACAAGCTCGGCCTCAAGCTCCTTAGGTACTACACCG

AGATCCTGTCCATATTCGGGCCCAGCCTTCGAGACCCGATATCTGCCGAAATATCAATCCAGGCACTCAGTTATGCATTA

GGCGGAGACATTAATAAAATCTTGGACAAGCTTGGGTATAGCGGTGGGGATTTCCTTGCCATCCTAGAAAGCAAGGGAAT

AAAGGCCCGGGTTACATATGTGGACACAAGAGATTACTTTATAATGCTTAGCATCGCCTACCCAACCTTATCTGAGATCA

AGGGAGTGATAGTTCACAAGATAGAAGCTATAACATACAACATTGGAGCACAGGAGTGGTATACTACTATCCCTAAATAT

GTAGCCACTCAGGGATATCTGATATCAAACTTTGATGAGACGTCATGCGTATTCACTCCAGAGGGGACAGTTTGCAGCCA

GAATGCGTTGTACCCAATGAGCCCGTTGCTTCAGGAATGTTTCAGGGGGTCAACAAGATCGTGCGCCAGAACCCTAGTTT

CAGGTACCATAAGTAATAGATTTATCCTATCAAAAGGGAACCTGATCGCAAATTGTGCGTCAGTTTTGTGCAAATGTTAC

ACAACGGAGACAGTCATCAGCCAAGATCCTGACAAACTACTAACTGTTGTAGCATCCGACAAGTGCCCTGTAGTTGAGGT

GGATGGAGTGACAATACAGGTCGGCAGCCGAGAGTATCCGGATTCTGTATACTTACACAAAATAGACTTAGGCCCAGCCA

TCTCCCTAGAAAAACTGGATGTAGGTACCAATTTAGGCAATGCAGTTACAAGACTGGATAATGCAAAGGAGCTCCTCGAT

GCATCAGACCAAATACTGAAGACTGTCAAAGGGGCACCTTTTGGTGGGAACATGTACATAGCACTGGCAGCTTGCATTGG

AGTATCCCTAGGGCTTGTCACATTAATATGCTGCTGTAAGGGGAGGTGTAAGAACAAGGAAATCCCTATCTCTAAAATCA

ACCCAGGGCTCAAACCTGACCTGACCGGGACCTCAAAGTCGTACGTGAGATCACTGTAGCCAGAATCACCTGAATCATCT

GGCTTCACACACATACATGTACGACACAAGCAGTCTGAGGACGCAGGAAACCCAGCCCCCGATCACCCACCTGACCCTAC

TCCACGCTCCACTACACATTAGTCATTAAACAAAACTTAGGACGAAAGGTCAATCACCATGTCCGCACAAAGGGAAAGGA

TCAATGCCTTCTACAAAGACAATCCTCACAATAAGAACCATAGGGTAATCCTGGATAGAGAACGCTTGGTCATTGAAAGA

CCCTACATCTTGCTTGGAGTCCTGCTGGTAATGTTCCTGAGTCTAATCGGACTGCTGGCCATTGCAGGGATCAGGCTTCA

CCGGGCCACCGTTGGAACTTCAGAGATCCAGAGTCGGCTGAATACCAATATTAAGTTGACCGAATCTATTGATCACCAGA

CTAAGGATGTCTTAACTCCCCTTTTTAAAATAATTGGCGATGAAGTCGGCATCCGGATTCCACAGAAATTCAGTGATCTT

GTCAAGTTCATCTCCGACAAGATTAAATTCCTCAACCCTGATAGAGAGTATGATTTCAGGGATCTCCGGTGGTGCATGAG

TCCCCCCGAGAGAGTCAAAATTAATTTTGATCAGTTTTGTGAGTACAAAGCTGCAGTTAATTCAATTGAACATATATTTG

AGTCACCACTCAACAAGTCAAAAAAGCTGCAATCATTGACTCTCGGGCCCGGAACAAGTTGTCTAGGCAGGACAGTAACA

AGAGCTCATTTCTCAGAACTTACACTGACCTTAATGGACTTGGATCTAGAGATGAAGCACAACGTGTCCTCAGTGTTTAC

CGTAATTGAAGAGGGGTTATTCGGAAGAACATATACCGTCTGGAGATCCGATGCCAGGGATCCGAGCACCGATCTGGGTA

TCGGCCATTTTTTAAGAGTCTTCGAGATTGGACTGGTAAGAGATCTCGGGCTGGGTCCCCCCGTTTTCCATATGACCAAC

TATCTCACAGTGAACATGAGTGATGTCTATCGGAGATGTCTTTTAGCGGTAGGGGAGTTGAAGTTGACAGCCCTATGCAC

CTCATCTGAGACTGTAACACTGAGTGAGAGAGGAGTTCCAAAGAGGGAGCCTCTTGTAGTTGTGATACTTAACCTGGCTG

GACCAACTCTAGGGGGCGAACTATACAGTGTCTTGCCTACCTCTGATCTCATGGTGGAGAAACTCTATTTATCTTCACAT

AGAGGGATCATCAAGGATGACGAGGCCAATTGGGTAGTGCCGTCTACCGATGTTCGTGATCTTCAGAACAAAGGTGAGTG

TCTAGTGGAAGCATGCAAGACTCGACCTCCTTCATTTTGCAATGGCACAGGATCAGGCCCGTGGTCAGAGGGGAGAATCC

CTGCTTACGGGGTGATCAGGGTCAGTCTTGACTTAGCTAGTGACCCGGATGTAGTTATTACTTCAGTGTTTGGCCCACTG

ATTCCTCACCTATCCGGCATGGATCTTTACAACAACCCGTTTTCAAGAGCTGTATGGTTAGCTGTACCACCTTATGAGCA

GTCATTTCTAGGAATGATAAATACAATTGGATTCCCTAACAGAACAGAGGTTATGCCGCACATTTTGACCACAGAGATCA

GAGGCCCTCGGGGTCATTGCCATGTTCCCATAGAATTGTCCCGCAGGGTTGATGACGATATCAGGATCGGGTCCAACATG

GTCATATTGCCGACGATGGACCTGAGATATATTACAGCCACTTATGATGTTTCCAGGAGCGAGCATGCAATCGTGTACTA

TATCTATGACACAGGTCGCTCATCATCTTACTTCTACCCAGTTCGACTGAATTTCAAAGGCAATCCTCTCTCTCTGAGGA

TAGAGTGTTTCCCTTGGCGTCATAAGGTGTGGTGCTACCATGATTGTCTTATATACAACACCACAACAGATGAAGAGGTC

CATATGAGAGGGCTGACCGGTATAGAGGTAACATGCAATCCAGTCTGAGCAGAGCTACGACCGTCACTCAAGCAGTCCCT

TGAGTCGCCACCGAGTCCAAGCAGCACAGCCTGGGACTCTCAACAGCACAGCCCAGCCAACAGTGTTATAAAAAACTTAG

GAGCCAGGGTTGTAGGGGCCATGGACTCCCTATCAGTCAATCAGGTCTTGTACCCTGAGGTTCATCTAGACAGCCCTATT

GTCACAAACAAACTAGTTGCCATCCTTGAGTACTCGGGTATCGACCATAACTATGTTCTTGAAGACCAGACCCTTGTCAA

GAATATTAGGTATAGACTGGGGTGCGGTTTTTCAAATCAAATGATCATCAATAACAGGGGAGTGGGTGAAACAGTCAATT

CTAAACTTAAAAGTTATCCCCATAATCGTCATATCATATACCCGGATTGCAATAAGGAGTTGTTTTGTATCAAAGATAGC

TGCATATCTAAGAAGCTCTCGGAGCTATTCAAGAAGGGTAATTCCTTGTACTCTAAGATAAGTCACCAGGTACTGGACTG

TCTTAAGAGAGTCAATGGGAAATTAGGCCTGGGCACAGATCTTACCCATGGTCTGAAGGAGGGTATCCTTGACCTAGGGT

TGCACATGCATAGCTCTCAATGGTTCGAGACTTTCCTGTTCTGGTTCACTATCAAGACAGAGATGAGATCAATGATCAAA

GAACAGTCCCATATATGCCACAAGAGGAGATATAACCCGATTTTTGTATCGGGGGATGCATTCGAGGTGCTCGTATCACG

AGACCTCGTTGTGATAATTGATAAGAATACCCAGTATGTCTTCTACTTAACATTTGAGCTGGTCCTTATGTATTGTGATG

TCATAGAGGGCAGACTCATGACGGAGACAGCCATGGCCATAGACCAGAGATATTCAGAACTTCTAAGCCGGGTTAGATAC

TTGTGGGATCTTATTGATGGGTTCTTCCCAACACTAGGCAATACCACCTACCAAATTGTTGCTCTGCTTGAACCATTATC

GTTGGCTTATCTTCAACTTCAGGATGTCACGCTGGAGTTAAGAGGTGCTTTTTTAGACCATTGCTTCAAAGAACTCTATG

AGATACTGGAGCATTGCGGCATTGACACAGAAGGCACCTACAATTCCATCACTGAAGGATTGGATTACGTATTCATCACC

CACGATATACATCTAACTGGGGAGATTTTTTCATTTTTCCGGAGTTTCGGACACCCCCGTCTAGAAGCAGTCACCGCCGC

AGAAAATGTCAGGAAACATATGAATCAACCGAAGGTAATCAGTTATGAGACTATGATGAAAGGACATGCGGTATTTTGCG

GGATAATCATCAATGGTTTTAGAGACCGACATGGCGGCAGCTGGCCCCCTGTTGCATTGCCAGAACATGCTTCTGCTGCG

ATCCGGAATGCGCAAGCATCAGGTGAGGGACTGACCCATGACCTGTGTATAGACAACTGGAAATCCTTTGTTGGTTTCAA

ATTTGGCTGCTTTATGCCACTCAGCCTAGATAGTGATTTGACCATGTATCTCAAAGACAAGGCATTGGCTGCACTGAAGA

ATGAGTGGGATTCAGTTTACCCGAAAGAATACCTCCGTTATAACCCACCTAGAGGGACAGAGTCAAGGCGACTGGTGGAG

GTGTTCCTGAATGACTCCAGCTTCGATCCTTATAACATGATAATGTACGTGGTGAATGGCTCCTATCTTAAAGACCCTGA

GTTTAATCTCTCATACAGCCTAAAAGAGAAGGAGATAAAGGAGACAGGGCGGTTGTTTGCCAAAATGACCTATAAGATGC

GGGCCTGTCAGGTAATTGCTGAGAATCTGATATCGAATGGTGTCGGGAAGTACTTCCGAGACAATGGGATGGCGAAGGAC

GAGCATGACCTAACAAAGGCCCTTCACACTTTGGCAGTCTCGGGTGTTCCCAAGAACAACAAAGATAACCACCGAGGTGG

GCCTCCCAGAAGGACAACAACCCGTGAGGTGAGATCAAGCCAGGACACCAGAACACAAAATAGAGACAAGGTCCAAGGGG

GACCTATGTACAACTATTTGCAATGCCAACCAATCAGCCCTGATCAAGGTGAGTCATACGAGACTGTTAGTGCATTTATC

ACCGCTGACCTTAAGAAGTATTGCCTGAATTGGAGATACGAGACAATCAGCATATTTGCACAGAGGCTGAATGAAATATA

TGGACTACCATCCTTCTTTCAATGGTTGCACAGGATATTGGAAAAGTCCGTACTCTACGTCAGTGACCCACATTGCCCTC

CCGATCTAGATAATCATATCCCTCTGGACAGTGTCCCTAATGCCCAAATATTCATTAAGTACCCAATGGGCGGAATAGAA

GGTTATTGCCAAAAACTATGGACAATCAGTACTATACCATACTTGTATCTGGCAGCCTATGAGAGCGGAGTAAGAATCGC

CTCACTGGTGCAGGGTGACAATCAGACAATCGCAGTGACAAAAAGAGTTCCAAGTTCTTGGCCTTATTCACTAAAAAAGA

GGGAAGCATCCAAAGCCGCTCAAAATTACTTCGTAGTCTTAAGGCAAAGATTGCACGATGTAGGTCACCACTTGAAGGCT

AATGAAACCATAGTATCTTCTCACTTTTTTGTATATTCCAAAGGGATTTATTATGACGGCCTGCTAGTCTCACAATCACT

AAAGAGCATCGCTAGATGTGTCTTCTGGTCCGAGACTATAGTGGATGAAACCAGAGCGGCTTGCAGCAATATCGCAACAA

CTGTCGCTAAGAGTATAGAGAGGGGTTATGATAGGTACCTCGCATATTCTTTGAATATCCTCAAGATTTTCCAACAGATC

CTTATATCCCTTAACTTCACTATTAACACAACAATGACTCAGGATGTCGTGGCACCTATTGTCGAGAACGGTGATTTGCT

GATAAGGATGGCACTCTTGCCAGCACCCATCGGAGGTCTAAATTATCTTAACATGAGCAGGTTATTCGTGAGAAATATCG

GCGACCCGGTCACCTCCTCTATAGCCGACCTGAAGAGGATGATAGATGCCGGGCTCATGCCAGAAGAAACATTGCATCAA

GTGATGACCCAGACCCCGGGAGAATCATCCTACCTTGATTGGGCAAGTGACCCTTATTCAGCCAACCTACCCTGCGTACA

GAGTATAACTCGTCTTCTCAAAAACATCACTGCACGATATATTTTAATTAGCAGCCCAAACCCGATGCTGAAAGGATTGT

TTCATGAGGGGAGTAGAGATGAAGACGAGGAGCTTGCAAGTTTCCTAATGGATCGGCATATAATTGTTCCAAGAGCTGCA

CATGAAATCTTAGACCATAGTATAACCGGGGCCAGAGAAGCTATAGCCGGGATGTTGGACACCACCAAGGGTCTGATTAG

AACAAGTATGAAACGGGGTGGCCTCACCCCTCGAGTCTTAGCCCGCCTTTCCAATTATGATTATGAGCAATTCCGATCCG

GGATAACACTATTGACAAAGAAAGGGCAGTGTTATCTCATTGACAAGGATTCGTGCTCGGTGCAGCTCGCTATAGCTCTG

AGGAGCCACATGTGGGCTAGGTTGGCTCGCGGGAGACCAATCTATGGGTTGGAGGTGCCCGATATATTAGAATCGATGAA

CGGCTACCTTATCAAACGCCACGAGTCTTGTGCTATCTGTGAAACGGGCTCAAGTCACTACGGATGGTTTTTCGTCCCTG

CAGGGTGCCAGCTTGACGATGTATCTAGAGAGACTTCGGCTCTCCGTGTACCTTATGTCGGGTCAACAACTGAGGAAAGG

ACTGACATGAAACTTGCATTTGTTAGATCTCCAAGTAGATCTCTCAAATCAGCAGTTAGAATTGCAACAGTTTACTCATG

GGCCTACGGGGATGATGAGAAGTCATGGAGTGAAGCCTGGATGCTAGCTAGGCAGAGAGCCAATATCACCTTAGATGAAT

TGAGAATGGTCACTCCAGTCTCTACATCCACCAACCTAGCCCACCGGTTGAGGGATCGGAGCACCCAGGTGAAATACTCG

GGGACATCCCTTGTGAGGGTTGCAAGATACACAACCATCTCCAATGATAACTTGTCATTTGTGATATCTGAGAAGAAAGT

AGATACCAACTTCATTTATCAGCAAGGGATGTTGCTTGGTCTTGGGATCCTTGAGAACCTCTTTAGGCTAGAGGCCACCA

CCGGAGTATCCAACACAGTGCTACACCTGCACGTGGAAACAGAATGTTGTGTTGTACCTATGGTGGATCACCCGAGGATA

CCGAGTCTCCGTAATATAAAAGTTACGGATGAGCTATGCACAAACCCTTTGATCTATGATAAGTCTCCCATCATAGAGCA

CGATGCAACCCGACTATACTCACAGAGCCACAGGAGACATTTGGTGGAGTTTGTTACCTGGTCCACAAGCCAGCTTTACC

ATATACTGGCTAAATCTACAGCAATGTCCATGATTGAATTGATCACGAGATTCGAGAAAGATCACATGAATGAAATAGCC

GCCCTGATTGGCGATGACGACATCAACAGCTTCATCACAGAATTTTTGCTAGTGGAGCCTAGATTATTTATAGTTTACCT

TGGTCAGTGTGCTGCCATCAATTGGGCCTTTGATATACATTATCATCGGCCCTCGGGCAAGTACCAGATGGGGGAACTCC

TCTACTCTCTTCTCTCTCGGATGAGCAAAGGAGTATATAAGATCTTCACCAATGCTTTGAGCCACCCCAAAGTTTACAAG

AAATTTTGGCGAAGTGGTGTCATTGAGCCGATTCATGGCCCATCCCTGGATACACAGAATTTACATGTCACTGTCTGTGA

CATGATATATGGGTCCTATGTAACCTACCTGGATCTTTTGCTGAATGATGAGCTAGATGATTACCCATATTTGCTCTGCG

AGAGTGATGAGGATGTGGTCACAGACAGGTTCGACAACATTCAAGCCAAACATCTCTGTGTATTAGCCGATGTATATTGC

AGCTCCAAGAGGTGTCCTTCAATAATCGGGATGTCTCCTATAGAAAAATGTACCATTCTCACACATTACATCAAGGGAGA

ATCAGTACAATCCCCGTCTGGAACCTCATGGAACACTGATCCCCTTGTAGTAGATCATTACTCATGCTCTCTGACCTACC

TTCGCCGAGGTTCCATCAAACAAATCAGGTTGAGGGTGGATCCTGGGTTTGTATTCGAGGCGTTGACAGACGTCGACTTT

AAACAACCTCGCAAAGCTAAGTTAGATATATCGGTCGTTGGGTTGACTGACTTCTCTCCCCCTTGCGACAATGTCGGTGA

TTTTCTAGGGACTATCAACACATTGAGGCACAATCTGCCTGTCACCGGAACCGGGGTCTCGAACTATGAAGTCCACGCTT

ATCGTAGAATTGGCCTGAATTCATCAGCATGTTACAAAGCCGTAGAGATCTCCACGTTAATCAAGCCATCCCTAGAAGTC

GGAGAGCATGGATTGTTCTTAGGAGAAGGTTCCGGTTCAATGCTGGCTGCGTACAAGGAAGTTCTTAAATTAGCAAATTG

TTATTACAACAGCGGGGTAACAGCGGAGGGCAGAGCCGGACAGAGGGAAATATCTCCCTATCCTTCAGAGATGAGCCTAG

TAGAGAATCAAATGGGGGTAGAGCGGAGTGTTAAAGTGCTGTTCAACGGCAAACCTGAAGTAACTTGGGTGGGGACCACC

GATTGCTACAAGTACATAATCAGTAACATCCAAACCTCTAGTCTAGGTTTCATACATTCAGATATTGAGACGCTCCCAAC

CAAAGATGCTGTTGAGAAATTAGAAGAATTTGCCTCTATCCTATCCTTATCCTTGATTTTGGGAAAAATCGGCTCTATTA

CAGTTGTCAAAATTATGCCCATTAGCGGGGATTTTACCCAAGGCTTCATAGCCTATGCCATCCAATATTTTAGGGAGAGC

CTGCTTGCCTATCCGAGATATAGTAACTTCATCTCGACTGAGTGTTACCTTATCATGATAGGATTAAAGGCCAATCGGTT

GATTAACCCAGAAGCCATTAAGCAAAGCATAATCAGAGCGGGGGTCAGGACTGCACCAGGACTTGTGAGCCATATATTAT

CAGAGAAACAGAAAGGTTGTATTCAATCTTTTTTGGGTGCTCCCTATATCCAAGGAGACTTCAATAAGCACCTTAAATCT

CTAACCCCTATTGAGAAAATCCTGGTAAATTGTGGTCTCTCGATCAATGGAACAAAAATCTGTAGAGATCTAATCCACCA

TGATATCGCCTCCGGTCCAGACGGTCTTATGAGCTCTACAATTATTTTATATAGAGAATTGGCTCATTTCAAAGATAATA

TAAGGAGTCAGCACGGAATGTTCCACCCCTACCCAGTACTGGCCAATAGCCGGCAACGTGAATTAATCCTTCGAATAGCC

AAGAAATTCTGGGGGTATGTCTTGCTATATTCTGATGACCCGGCACTAATCAGACAAACAATCAAGAACTTGAAACGGAA

TCATCTAACCTTTGACTTACATAGTAACCCGTTTATTAAAGGGCTATCCAAAGCTGAGAAACTGCTGGTGCGGACGAGTT

CACTAAGAAGGGAATGGTTGTTCACCTTCGATACGAAAGAAGTGAAAGAGTGGTTCAAATTGGTAGGTTACAGTGCACTC

ATCAGAGGCTAGTTAGTTATACATCTGCCCCCTTCTCCTCCGCCATGAGACCCCACTGACGATTCAGAAGATTAAAGAAA

ACTACATATTGGATAAGTATCTATTCCCAGCTTTGTCTGGT

>MG581412.1/goat/Bangladesh/2008-05

ACCAAACAAAGTTGGGTAAGGATAGATCTTATAATAACTATGGACTAGCAAACTTAGGAGTAAAGATCCTACTGTCGGGG

GGAGGAGGAGGAGCAAGATCTCTGATTATGGCGACTCTCCTTAAAAGCTTAGCATTGTTCAAAAGGAACAAAGACAAAGC

GCCGACGGCGTCAGGTTCAGGAGGGGCCATCCGAGGGATTAAGAATGTTATCATAGTCCCGATTCCCGGAGACTCGTCCA

TCACTACCCGTTCAAGACTGCTCGACAGGCTTGTCAGATTGGCCGGAGATCCTGACATCAACGGGTCAAAGCTGACCGGC

GTGATGATCAGCATGTTATCTTTGTTCGTAGAGTCACCCGGGCAATTGATACAGCGAATCACAGATGATCCAGATGTTAG

TATCCGCCTTGTTGAGGTAGTTCAAAGTACTAGGTCCCAGTCCGGGTTGACCTTTGCATCACGTGGTGCTGATTTGGATA

ATGAGGCAGACATGTATTTTTCAACTGAGGGGCCCTCGAGTGGGGGTAAAAAAAGGATCAACTGGTTTGAGAACAGAGAA

ATAATAGACATAGAGGTGCAGGATCCAGAAGAGTTCAATATGTTGTTAGCCTCCATACTAGCACAAGTCTGGATCCTTCT

GGCCAAGGCTGTTACGGCACCGGATACAGCAGCTGACTCAGAACTGAGGAGGTGGGTTAAATACACACAACAAAGGAGAG

TGATTGGGGAATTTCGCCTTGACAAAGGGTGGCTGGACGCAGTCCGCAACAGAATTGCAGAAGATCTATCACTCCGGCGG

TTCATGGTATCTCTCATACTTGACATCAAGAGGACCCCTGGCAACAAGCCAAGGATTGCAGAAATGATCTGCGACATTGA

CAACTATATCGTCGAAGCAGGACTCGCCAGTTTCATCCTTACTATCAAATTTGGTATAGAAACCATGTATCCTGCATTAG

GGCTCCACGAGTTTGCCGGGGAATTGTCCACTATTGAGTCCTTGATGAACTTGTATCAACAGCTAGGAGAGGTTGCACCC

TACATGGTAATTCTAGAGAACTCAATTCAGAACAAGTTTAGTGCAGGAGCTTATCCCCTCCTCTGGAGCTATGCGATGGG

TGTCGGAGTTGAGCTGGAGAACTCAATGGGGGGCTTGAACTTTGGCAGGTCATATTTTGACCCTGCCTATTTTCGTCTCG

GACAGGAGATGGTCAGAAGATCTGCAGGAAAGGTCAGCTCTGTGATTGCGGCTGAGCTCGGCATCACAGCAGAGGAAGCT

AAACTAGTCTCGGAAATAGCCTCGCAGACTGGGGACGAAAGAACCGCTAGAGGGACTGGGCCTCGACAGGCGCAGGTCTC

CTTCCTCCAGCACCAAACAGGAGGGGGAGAGTCGTCCGCACCAGTGACCAGAGAAGGGGTCAAAGCTGCGATCCCAAGCG

GATCCGAAGAAAGGGACAGAAAGCAAACACGCCCAGGAAGGCCCAGAGGAGAGACCCCCGGCCAACTGCTCCTGGATATC

ATGCCAGAGGATGAGGTCTCGCGAGAATCTGGTCGAAACCCTCGTGAGGCTCAAAGATCGGCTGAGGCACTCTTCAGGCT

GCAGGCCATGGCCAAGATTCTGGAGGACCAGGAGGAGGGAGAAGACAACAGTCAGGTCTACAACGACAAGGATCTCCTCG

GCTAAGCAGACGCACCCTCTGTCGAAATCAGTGACGAGACATCTCCTACCAGTATTATAAAAAACTTAGGACCCAGGTCC

AAGCAAGCACACGTCGACACTCCAACCAGTCAAGCGGAGGCCACCGATGGCAGAGGAACAAGCATACCATGTCAACAAGG

GGCTGGAATGTATCAAGTCCCTCAAAGCCTCTCCCCCGGATCTATCCACCATCAGAGATACCATCGAGAGCTGGAGAGAG

GGGCTTAGCCCCTCGGGCCGTGCAACACCGAACCCTGATACGTCCGAGGGAGATCATCAGAATATCAGCCAATCATGCTC

ACCAGCAATCGGACCAAACAAAGTCTACTTGTCTCCTGAAGATAATCTCGGATTTAGAGAGATCACTGGCAATGACTGTG

AGGCTGAGCTCGGAGGAGTCCAGGGAGAAGGATCCAACTCTCAAGTACAGCGTTACTATGTTTATAGCCACGGGGGTGAA

GAGATTGAAGGACTCGAGGATGCTGACTCTCTCGTGGTTCAAGCAAATCCTCCAGTTGCTAACATATTCAATGGAGGAGA

GGATGGATCTGACGACAGCGATGTGGACTCTGGCCCAGATGATCCCGGCAGAGATACTCTATATGACCGGGGATCTGTTG

CCAGCAATGATGTCGCTAGGTCCACAGATGTCGAGAAACTAGAAGGTGCTGATATTCAAGAAGTCCTCAACTCCCAGAAA

GGCAGAGGGGGAAGATTCCAAGGCGGGAAAACCTTACGAGTCCCGGAAGTACCCGATGTCAAGCACTCCAGACCATCAGC

CCAATCAATTAAAAAGGGCACAGACGGGAACTCAGTCTCATCTGGAACGGTGACAGAGTGTTTATCGATAAGTGGTGCAA

CCCAAGCTGTGCCAGAGTCAAGATGGGAGTCATCAGAGCAAAATGCGTCTGTGGGGAGTGTCCTCAAGTCTGCGAGGAGT

GCAAAGACGATCCAGGGGTCAACACAAGAATCTGGTACCATAGCATCACTGACTCAGCATAAAGAGAATGACTCCGAGTA

TGAGTATGAGGATGACCTATTTACAGAGATTCAGGACATCCGTGCAAGCATTGCCAAGATCCATGATGACAACAAGACTA

TCCTCTCAAAGCTTGATTCTATACTGTTATTGAAAGGAGAAGTCGACACTATCAAGAAACAAATCAGCAAGCAGAATATA

AGTATATCCACCATTGAGGGCCATCTCTCCAGTATAATGATAGCCATCCCGGGCTTTGGGAAGGATATCAAGGACCCAAC

ATCCGAGGTTGAGCTGAACCCAGATTTAAGACCTATAATAAGCCGTGACTCTGGCAGAGCTCTCGCAGAGGTCCTCAAGA

AACCCGCTGTGGATAGGTCTCAGAAAATTGGAGCCAAAGCCAACTCCAGCTCAAAGGGTCAGCTTCTTAAAGATCTCCAG

CTAAAACCTGTCGACAAGCAGGCAAGCTCTGCAATCGGGTTTGTCCCATCCGACCATGAATCATCCAGAAATGTCATCCG

CTCCATAATCAAGTCGAGCAAGCTAAACATTGATCACAAGGATTATCTTCTAGATTTACTGAATGATGTGAAAGGCTCCA

AGGATCTTAAGGAATTCCACAAGATGCTAACAGCAATTCTCGCCAAGCACCCGTAACACATCCTCCAGTCACCATCTCAT

ACTCGGCTGAAAACACCCTTTCAATCAGGCTATTACAAAAAACTTAGGAGCAAGGGCAACTGAGCTTCGCAGACAAGATG

ACCGAGATCTACGACTTCGATAAATCAGCATGGGATGTCAAAGGGTCAATTGCCCGCATAGAACCCACCACCTACCACGA

CGGCCGACTGATACCCCAAGTAAGGGTCATCGATCCCGGTCTGGGAGACAGAAAAGATGAGTGCTTTATGTACCTGTTTC

TCCTGGGAGTGATTGAGGATAACGACCCCCTATCTCCCCCCGTCGGAAGAACCTTTGGCTCTTTACCTCTAGGGGTCGGT

AGGTCAACTGCTAAGCCAGAAGAACTACTAAGGGAGGCCACAGAACTAGACATAGTGGTGAGGCGCACTGCAGGAGTAAA

TGAGAAACTGGTTTTTTACAACAACACTCCGCTGTCCTTGTTAACGCCCTGGAAGAAAGTTCTGACAACCGGAAGTGTGT

TTAGCGCTAACCAGGTTTGCAATGCAGTCAACCTAGTCCCGCTTGATACTCCCCAGAGGTTCAGGGTTGTGTACATGAGC

ATAACTAGATTATCAGACAATGGTTACTATAGTGTGCCCAGAAGAATGCTGGAGTTCCGCTCAGCCAATGCAGTAGCCTT

TAACATCTTGGTAACACTAAGAATTGAAAATGGCACAAACCCTAGAAGATACATAGTCGGCTCATGGGAGAATTCAGAAG

TCACATTTATGGTACACGTGGGCAACTTTAGAAGAAAGAAGAACGAAGTATACTCTGCTGATTACTGCAAAATGAAGATT

GAAAAGATGGGTTTAGTTTTTGCCCTGGGAGGAATAGGTGGAACAAGTCTCCATATTAGAAGCACCGGGAAAATGAGCAA

AACCCTCCATGCACAGCTGGGGTTCAAGAAAATCTTATGTTACCCCCTAATGGACGTTAATGAGGATCTTAACCGATACC

TCTGGCGGGCAGAGTGCCGAATAGTCAAAATCCAAGCCGTCTTACAGCCATCAGTGCCCCAAGAATTCCGTGTCTATGAT

GATGTCATCATCAACGATGATCAAGGCCTGTTCAAGATCTTGTAGTTCATTTGCAAAGGAACGAACCCACCCCCAGGAAG

GGGCCCCACCCCCTCCCG-CCAAAAAACACACTCCAAAGCTCCCCGACACAGCGCCAACACAGCCACCCCTCCGACTGGA

CGAGGCCGGCC-CCCCAATAAACCCGGTCCAGGCGGGGGG-CCCCCCCCCTGACCCGCCGGGCGCCGATCGAGCAGGCCG

GCGAGAAGCCCCCAAGAGGCCCCGACGAAGACCGAGCACCCCATCCTCCC-AAAGCGTCTCCCC-AACACGCCCGAGGCC

AGGCGCCGGGCACCCCCAGACCCGCCACCGGCGAGACAGAACCCCAGGGGGGGACCCCCCAAACCAAAGGGACCACCGAG

GAAGGCTCCGCCGAACCAGCACAGACCCGACCCAAGCAAGGCGGACCCCCAACACGAAACCGCCCAGCACACCCCACTCA

TCATCAAACAAAACTTAGGGGCCAAGCCCATACGACAGGCCCCAGTCCGCCAGGCGGAGGGACGCAGAAAGGAAGGAGAC

ATGACCGCCCCAGAAGGGGGACACCCAACCACCGCAACACCCAGCCCCAGGGAAGGAG--CCCCCCCCCACACGGCGAGG

GAACGGGTGAGAGACCCCGAGGAGAACCCGACCGGGAGCCAGGAGCCGGCGGCGCCGGCGGCAAACAAGCCGGGCAAACA

CCCGACCCCACCAGAGGAGCCGAGGAGAGCCCGACCCCGCACGAACCAAACGGCAGGGAGGCCAACCCCACCACAAAAAG

AGGCGAGGCCTC-CCT-----CCCCTCCGCACACAACCCGCAGACAGACCGAGAGAGGACCACCTCCCAACAAGACAGGC

GGCTCCCTCGCAGCCGAGCCGCAGCTCCCCAGAACAGGACACCCCCGCACACTCCGAACACCCCCCCGCCCCCCGCCACC

GGGACAGGGGCTCCGCCACCACAGACCAAAGCCACCGAGAGCCAGCCAGAGGTCCCCCCCAGTCCCGCCCGGTCCCCCCC

CGAAAAGGCCCCGGCAAGGCCCCCAAACCCGACGTCCCCCGCACGCACCGAAAGAGGGACCCGCCCGAGAACGTGCGGCA

CATCCATGCGTAAACATCATGACACGGGTCGCAATCTTGATATTTCTGTTTCTTCTCCCAAATGTTGTTGCGTGCCAGAT

TCACTGGGGCAATCTATCCAAGATCGGGATTGTAGGAACAGGGAGTGCCAGCTACAAGGTGATGACTAGGCCAAGCCACC

AAACTCTGGTTATAAAGTTAATGCCAAATATAACAGCTATCGACAATTGTACAAGGTCAGAGATTGCAGAGTACAAGAGA

TTGCTGATCACAGTGTTAAAGCCTGTAGAGGATGCTCTGTCGGTGATAACCAAGAATGTAAGACCAATTCAAACTCTGAC

ACCTGGGCGTAGAACCCGTCGTTTTGCTGGAGCTGTTCTGGCCGGAGTAGCACTTGGAGTTGCGACAGCCGCTCAGATAA

CTGCAGGAGTCGCTCTTCATCAATCATTGATGAACTCCCAAGCAATTGAGAGTTTAAAAACCAGTCTTGAGAAGTCGAAT

CAGGCAATAGAAGAAATCAGACTTGCAAATAAGGAGACCATACTGGCAGTACAGGGCGTCCAGGATTATATCAACAATGA

GCTTGTCCCTTCTGTTCATAGAATGTCATGCGAGCTGGTAGGCCACAAGCTCGGCCTCAAGCTCCTTAGGTACTACACCG

AGATCCTGTCCATATTCGGGCCCAGCCTTCGAGACCCGATAGCTGCCGAAATATCAATCCAGGCACTCAGTTATGCATTA

GGCGGAGACATTAATAAAATCCTGGACAAGCTTGGGTATAGCGGTGGGGATTTCCTTGCCATCCTAGAAAGCAAGGGAAT

AAAGGCCCGGGTTACATATGTGGACACAAGAGATTACTTTATAATCCTTAGCATCGCCTACCCAACCTTATCTGAGATCA

AGGGAGTGATAGTTCACAAGATAGAAGCTATAACATACAACATTGGGGCACAGGAGTGGTATACTACTATCCCTAAATAT

GTAGCCACTCAGGGGTATCTGATATCGAACTTTGATGAGACGTCATGCGTATTCACTCCAGAGGGGACAGTTTGCAGCCA

GAATGCGTTGTACCCAATGAGCCCATTGCTTCAGGAATGTTTCAGGGGGTCAACAAAATCGTGCGCCAGAACCTTAGTTT

CAGGGACCATAAGTAATAGATTTATCCTATCAAAAGGGAACCTGATTGCAAATTGTGCATCAGTTTTGTGCAAATGCTAC

ACAACGGAGACAGTCATCAGCCAAGATCCTGACAAACTACTAACTGTTGTAGCATCCGACAAGTGTCCTGTAGTTGAGGT

GGATGGAGTGACAATACAGGTCGGCAGTCGAGAGTACCCGGATTCTGTATACTTACACAAAATAGACTTAGGTCCAGCCA

TCTCCCTAGAAAAACTGGATGTAGGCACCAATTTAGGCAATGCAGTCGCAAGACTGGAGAATGCAAAGGAGCTCCTAGAT

GCATCAGACCAAATACTGAAGACTGTTAAAGGGGCACCTTTCGGTGGGAACATGTACATAGCACTGGCAGCTTGCATTGG

AGTATCCCTAGGGCTTGTCACATTAATATGCTGCTGTAAGGGGAGGTGTAAGAACAAGGAACTCCCTATCTCCAAAATCA

ACCCAGGGCTCAAACCCGATCTGACCGGGACCTCAAAGTCGTACGTGAGATCACTGTAGCCAGAATCACCTGAATCATCT

GGCATCACACACATACATGTACGACACAAGCAGTCTGAGGACGCAGGAAGCCCAGCCTCCGATCACCCACCTGACTCTAC

TCCACGCTCCACTACACATTAGTCATCAAACAAAACTTAGGACAAAAGGTCAATCACCATGTCCGCACAAAGGGAAAGGA

TCAATGCCTTCTACAAAGACAATCCTCACAATAAGAACCATAGGGTGATCCTGGATAGAGAACGCTTGGTCATTGAAAGA

CCCTACATCTTGCTTGGAGTCCTGCTGGTAATGTTCCTGAGTCTAATCGGACTGCTGGCCATTGCAGGGATCAGGCTTCA

CCGGGCCACCGTTGGAACTTCAGAGATCCAGAGTCGGCTAAATACCAATATTAAGTTGACCGAATCTATTGATCACCAGA

CTAAGGATGTCTTAACTCCCCTTTTTAAAATCATTGGCGATGAAGTCGGCATCAGAATTCCACAGAAATTCAGTGATCTT

GTCAAGTTCATCTCCGATAAGATTAAATTCCTCAACCCTGATAGAGAGTATGATTTCAGGGATCTCCGGTGGTGCATGAA

TCCCCCTGAGAGAGTCAAAATTAATTTTGATCAGTTTTGTGAGTACAAAGCTGCGGTTAAGTCAATTGAACATGTATTTG

AGTCACCACTCAACAAGTCAAAAAAGCTGCAATCATTGACTCTCGGGCCCGGAACAGGCTGTCTAGGCAGGACAGTAACA

AGAGCTCATTTCTCAGAACTTACATTGACCTTAATGGACCTGGATCTAGAGATGAAGCACAACGTGTCCTCAGTGTTTAC

CGTAGTTGAAGAGGGGTTATTCGGAAGAACATATACCGTCTGGAGATCCGATGCCAGGGATCCGAGCACCGATCTAGGTA

TCGGCCATTTTTTAAGAGTCTTCGAGATTGGACTGGTAAGAGATCTCGGGCTGGGTCCCCCTGTTTTTCATATGACCAAC

TATCTCACAGTGAACATGAGTGATGACTATCGGAGATGTCTTTTAGCGGTAGGGGAGTTGAAGTTGACAGCCCTATGCAC

CTCATCTGAGACTGTGACACTGAGTGAGAGAGGAGTTCCAAAGAGGGAGCCCCTTGTGGTTGTGATACTTAACCTGGCTG

GACCCACTCTAGGGGGCGAACTATACAGTGTCTTGCCTACCTCTGATCTCATGGTGGAGAAACTCTATTTATCTTCACAT

AGAGGGATCATCAAAGATGACGAGGCCAATTGGGTAGTGCCGTCTACCGATGTTCGTGATCTTCAGAACAAAGGTGAATG

TCTGGTGGAAGCATGCAAGACCCGACCTCCTTCATTTTGCAATGGCACAGGATCAGGCCCGTGGTCAGAGGGGAGAATCC

CTGCTTACGGGGTGATCAAGGTCAGTCTTGACTTAGCTAGTGACCCGGATGTAGTTATCACTTCAGTGTTTGGCCCACTG

ATTCCTCACCTATCCGGCATGGATCTTTACAACAACCCGTTTTCAAGAGCTGTATGGTTGGCTGTACCACCTTATGAGCA

GTCATTTCTAGGAATGATAAATACAATTGGATTCCCTAACAGAGCAGAGGTTATGCCGCACATTTTGACCACAGAGATCA

GAGGCCCTCGGGGTCGTTGCCATGTTCCCATAGAATTGTCCCGCAGGGTTGATGACGATATCAAGATCGGGTCCAACATG

GTCATATTGCCGACGATGGACCTGAGGTATATTACAGCCACTTATGATGTTTCCAGGAGCGAGCATGCAATCGTGTACTA

TATCTATGACACAGGTCGCTCATCATCTTACTTCTACCCAGTTAGACTGAATTTCAAAGGCAATCCTCTCTCTCTGAGGA

TAGAGTGTTTCCCTTGGCGTCATAAGGTGTGGTGCTACCATGATTGTCTTATATACAACACCATAACAGATGAAGAGGTC

CATACGAGGGGGCTGACCGGTATAGAGGTAACATGCAATCCGGTCTGAGCAGAGCTACGACCGTCACTCAAGCAGTCCCT

TGAGTCGCCACCGAGTCCAAGCAGCACAGCCTGGGACACTCAACAGCACAGCCTAGCCAACAATGTTATAAAAAACTTAG

GAGCCAGGGTTGTAGGGGCCATGGACTCCCTATCAGTCAATCAGGTCTTGTACCCTGAGGTTCATCTAGACAGCCCTATT

GTCACAAACAAACTAGTTGCCATCCTTGAGTACTCGGGTATCGACCATAACTATGTTCTTGAAGACCAGACCCTTGTCAA

AAATATTAGGTATAGACTGGGGTGCGGTTTTTCAAATCAAATGATCATCAATAACAGGGGGGTGGGTGAAACAGTCAATT

CTAAACTTAAAAGTTATCCCCATAATCGTCATATCATATACCCGGATTGCAATAAGGAGTTGTTTTGTATCAAAGATAGC

TGCATATCTAAGAAGCTCTCGGAGCTATTCAAGAAGGGTAACTCCTTGTACTCTAAGATAAGTCACCAGGTACTGGATTG

TCTTAAGAGAGTCAATGGGAAATTAGGCCTGGGCACAGATCTTACCCATGGTCTGAAGGAGGGTATCCTTGACCTAGGGT

TGTACATGCATAGCTCTCAATGGTTTGAGACTTTCTTGTTCTGGTTCACTATCAAGACAGAGATGAGATCAATGATCAAA

GAACAGTCCCATATATGCCACAAGAGGAAGTATAACCCGATTTTTGTATCGGGGGATGCATTCGAGGTGCTCGTATCACG

AGACCTCGTTGTGATAATTGATAAGAGTACCCAGTATGTCTTCTACTTAACATTTGAGCTGGTCCTTATGTATTGTGATG

TCATAGAGGGCAGACTTATGACGGAGACAGCCATGGCCATAGACCAGAGATATTCAGAACTTCTAAGCCGGGTTAGATAC

TTGTGGGATCTTATTGATGGGTTCTTCCCAACACTAGGCAACACCACATACCAAATTGTTGCTCTGCTTGAACCATTATC

GTTGGCTTATCTTCAACTTCAGGATGTCACGCTGGAGTTAAGAGGTGCTTTTTTAGACCATTGCTTCAAAGAACTCTATG

AGATACTGGAGCATTGCGGCATTGACACAGAAGGCACCTACAATTCCATCACTGAAGGATTGGATTACGTATTCATTACC

CACGATATACATCTAACTGGGGAGATTTTTTCATTTTTTCGAAGTTTCGGACACCCCCGTCTAGAAGCAGTCACCGCTGC

AGAAAATGTCAGGAAACATATGAACCAACCGAAGGTAATCAGTTATGAAACTATGATGAAAGGACATGCGGTATTTTGCG

GGATAATCATAAATGGTTTTAGGGACCGACATGGCGGCAGCTGGCCCCCTGTTGCATTGCCAGAACACGCCTCTGCTGCG

ATCCGGAATGCGCAGGCATCAGGTGAGGGACTGACCCATGACCTGTGTATAGACAACTGGAAATCCTTTGTTGGATTCAA

ATTTGGCTGCTTTATGCCACTCAGCCTAGATAGTGATTTGACCATGTACCTCAAAGACAAGGCATTGGCTGCACTAAAGA

ATGAGTGGGATTCAGTTTACCCGAAAGAATACCTCCGTTATAACCCACCTAGAGGGACAGAGTCAAGGCGACTGGTGGAG

GTGTTCCTGAATGACTCCAGCTTTGATCCTTATAACATGATAATGTACGTGGTGAATGGCTCCTACCTTAAAGACCCTGA

GTTTAATCTCTCATACAGCCTAAAAGAGAAGGAGATAAAGGAGACAGGGCGGTTGTTTGCCAAAATGACCTATAAGATGC

GGGCCTGTCAGGTAATTGCTGAAAATCTGATATCGAATGGTGTTGGGAAGTATTTCCGAGACAATGGGATGGCAAAGGAC

GAGCATGACCTAACAAAAGCCCTTCACACCCTGGCAGTCTCAGGTGTTCCCAAGAACAACAAAGATAACCACCGAGGTGG

GCCTCCCAGAAGGACAACAAGCCGAGAGGTGAGATCAAGCCAGGACACCAAAACACAAAATAGAGACAAGGTCCAAGGAG

GACCTATGTACAACTATTTGCGATGCCAACCAATCAGACCTGATCATGGTGAGTCATACGAGACTGTTAGTGCATTCATC

ACCGCTGACCTTAAGAAGTATTGCCTGAATTGGAGATACGAGACAATCAGCATATTTGCACAGAGGCTGAATGAAATATA

TGGACTACCATCCTTCTTTCAATGGTTACACAGGATATTGGAAAAATCCGTACTCTACGTCAGTGACCCACATTGCCCTC

CCGATCTAGATAATCATATCCCTCTGGACAGTGTCCCTAATGCCCAAATATTCATTAAGTACCCAATGGGCGGAATAGAA

GGTTATTGCCAAAAACTATGGACAATCAGTACTATACCATACTTGTATCTGGCAGCCTATGAGAGCGGAGTAAGAATCGC

CTCACTGGTGCAGGGTGACAATCAGACAATCGCAGTGACAAAAAGAGTTCCAAGCTCTTGGCCTTATTCACTAAAAAAGA

GGGAGGCATCCAAAGCCGCTCAAAATTACTTCGTAGTCTTAAGGCAAAGATTGCACGATGTAGGTCATCACTTAAAGGCT

AATGAAACCATAGTATCTTCTCACTTTTTTGTATATTCCAAAGGGATTTATTATGACGGCCTGCTAGTCTCACAATCACT

AAAGAGCATCGCTAGATGTGTCTTCTGGTCCGAGACTATAGTGGATGAAACCAGAGCGGCTTGCAGCAATATTGCAACAA

CTGTCGCTAAGAGTATAGAGAGGGGTTATGATAGGTACCTTGCATATTCTTTGAATATCCTCAAGATTTTCCAACAGATC

CTTATATCCCTTAACTTCACTATTAACACAACAATGACTCAGGATGTCGTGGCACCGATCATCGAGAACGGCGACTTGCT

GATAAGGATGGCACTCCTGCCAGCACCCATCGGAGGTCTAAATTATCTTAACATGAGCAGGTTATTTGTGAGAAATATCG

GTGACCCAGTCACCTCCTCTATAGCCGACCTGAAGAGGATGATAGATGCCGGGCTAATGCCAGAAGAAACATTGCATCAA

GTGATGACCCAGACCCCGGGAGAATCATCCTACCTCGATTGGGCAAGTGACCCTTATTCTGCCAACCTACCCTGCGTACA

GAGTATAACTCGCCTTCTCAAAAACATCACTGCACGATATATTTTAATCAGCAGCCCAAACCCGATGCTGAAAGGATTGT

TTCATGAGGGGAGTAGAGATGAAGACGAGGAGCTTGCAAGTTTCCTAATGGATCGGCATATAATTGTTCCAAGAGCTGCA

CATGAAATCTTAGACCATAGTATAACCGGGGCAAGAGAAGCTATAGCCGGGATGTTGGACACCACCAAGGGTCTGATTAG

AACAAGTATGAAACGGGGTGGCCTCACCCCTCGAGTCTTAGCCCGCCTTTCCAATTATGATTATGAACAATTCCGATCCG

GGATAACACTATTGACAAAGAAAGGGCAGTGTTATCTCATTGACAAGGATTCGTGCTCGGTGCAGCTCGCTATAGCTCTG

AGGAGCCACATGTGGGCTAGGTTGGCTCGCGGGAGACCAATCTATGGGTTGGAGGTGCCCGATATATTAGAATCGATGAA

CGGCTACCTTATCAAACGCCACGAGTCTTGTGCTATCTGTGAAACGGACTCAAGTCACTACGGATGGTTTTTCGTCCCTG

CAGGGTGCCAGCTTGACGATGTATCTAGAGAGACTTCGGCTCTTCGTGTACCTTATGTCGGATCAACAACTGAGGAAAGG

ACAGACATGAAACTTGCATTTGTTAGATCTCCAAGTAGATCCCTCAAATCAGCAGTTAGAATTGCAACAGTTTACTCATG

GGCCTACGGGGATGATGAGAAGTCATGGAGTGAAGCCTGGATGCTAGCTAGGCAGAGAGCCAATATCACCTTAGATGAAT

TGAGAATGATCACTCCAGTCTCTACATCCACCAACCTAGCCCACCGGTTGAGGGATCGGAGCACCCAGGTGAAATACTCG

GGGACATCCCTTGTGAGGGTTGCAAGATACACAACCATCTCCAATGATAACTTGTCATTTGTGATATCTGAGAAAAAAGT

GGATACCAACTTCATTTACCAGCAAGGGATGTTGCTTGGTCTTGGGATCCTTGAGAACCTCTTTAGGCTAGAGGCCACCA

CCGGAGTATCCAACACAGTGCTACACCTGCACGTGGAAACAGAATGTTGTGTTGTACCTATGGTGGATCACCCAAGGGTA

CCGAGTCTCCGTAATATAAAAGTCACGGATGAGCTATGCACAAACCCTTTGATCTACGACAGGTCTCCCATCATAGAGCA

CGATGCAACCCGACTATACTCACAGAGCCACAGGAGACATTTGGTGGAGTTTGTTACCTGGTCCACAAGCCAGCTTTACC

ATATACTGGCTAAGTCTACAGCTATGTCCATGATTGAATTGATCACGAGATTCGAGAAAGATCACATGAATGAAATAGCT

GCTCTGATTGGCGATGACGACATCAACAGCTTCATCACAGAATTTTTGCTAGTGGAGCCTAGATTATTTATAGTTTACCT

TGGTCAGTGTGCTGCCATCAATTGGGCCTTTGATATACATTATCATCGGCCCTCGGGCAAGTACCAGATGGGGGAACTCC

TCTACTCTCTGCTCTCTCGGATGAGCAAAGGAGTATATAAAATCTTCACCAATGCTTTGAGCCACCCCAAAGTTTATAAG

AAATTTTGGCGAAGTGGTGTCATTGAGCCGATTCATGGCCCATCCTTGGATACACAGAATTTACATGTCACTGTCTGTGA

CATGATATATGGGTCCTATGTCACCTATTTGGATCTTTTGCTGAATGATGAGCTAGATGATTACCCGTATTTGCTCTGCG

AGAGTGATGAGGACGTGGTCACAGACAGGTTCGACAACATTCAAGCCAAACATCTCTGTGTATTAGCCGATGTATATTGC

AGCTCCAAAAGGTGTCCTTCAATAATCGGGATGTCTCCTATAGAAAAATGTACCATTCTCACACATTACATCAAGGGAGA

ATCAGTACAATCCCCGTCTGGGACCTCATGGAACACTGATCCCCTTGTAGTAGATCATTACTCATGCTCCCTGACGTACC

TTCGCCGCGGTTCCATCAAACAAATCAGGTTGAGGGTGGATCCTGGGTTTGTATTCGAGGCGTTGACAGACGTCGACCTT

AAACAACCTCGCAAAGCTAAGTTAGATATATCGGTCGTTGGATTGACTGATTTCTCTCCCCCTTGCGACAACGTCGGTGA

TTTTCTAGGGACTATCAACACATTGAGGCACAATCTGCCTGTCACCGGAACCGGGGTCTCGAACTATGAAGTCCACGCTT

ATCGTAGAATTGGCCTGAATTCATCAGCATGTTACAAAGCCGTAGAGATCTCCACGTTAATCAAGCCATCCCTAGAAGTC

GGAGAGCATGGATTGTTCTTAGGAGAAGGTTCCGGTTCAATGCTGGCTGCGTACAAAGAAGTTCTTAAATTAGCAAATTG

TTATTACAACAGCGGGGTAACAGCGGAGGGCAGAGCCGGACAGAGGGAAATATCTCCCTATCCTTCAGAGATGAGCCTAG

TAGAGAATCAGATGGGGATAGAGCGGAGTGTTAAAGTGCTGTTCAACGGCAAACCTGAAGTAACTTGGGTGGGGACCACC

GATTGCTACAAGTACATAATCAGTAACATCCAAACCTCTAGTCTGGGTTTCATACATTCAGATATTGAGACGCTCCCAAC

CAAAGATGCTGTTGAGAAATTAGAAGAATTTGCCTCTATTCTATCCTTATCCTTGATTTTAGGAAAAATCGGCTCTATTG

CAGTTGTCAAAATTATGCCCATTAGCGGGGATTTTACCCAAGGCTTCATAGCCTATGCCATCCAATATTTTAGGGAGAGC

CTGCTTGCCTATCCGAGATATAGTAACTTCATCTCGACTGAGTGTTACCTTATTATGATAGGATTAAAGGCCAATCGATT

GATAAACCCAGAAGCCATTAAGCAAAGCATAATCAGAGCGGGGGTCAGGACTGCACCAGGACTTGTGAGCCATATATTAT

CAGAGAAACAGAAAGGTTGTATTCAATCTTTCTTGGGTGATCCTTATATCCAAGGAGATTTCAATAAGCACCTTAAATCT

CTAACCCCTATTGAGAAAATCCTGGTAAATTGTGGTCTCTCGATCAATGGCACAAAAATCTGTAGAGATCTAATCCACCA

TGATATCGCCTCCGGTCCAGACGGTCTTATGAGCTCTACAATTATTTTATATAGGGAATTGGCTCATTTCAAAGATAATA

TAAGGAGTCAGCACGGAATGTTCCACCCCTACCCAGTACTGGCCAACAGCCGGCAACGTGAATTAATCCTTCGAATAGCC

AAGAAATTCTGGGGGTATGTCTTGCTATATTCTGATGACCCGGCACTAATCAGACAAACAATCAAGAACTTGAAACGGAA

TCATCTAACCTTTGACTTACATAGTAACCCGTTTATTAAAGGGCTGTCCAAAGCTGAGAAACTGCTGGTGCGGACGAGTT

CACTAAGAAGGGAATGGTTGTTCACTCTCGATACGAAAGAAGTGAAAGAGTGGTTCAAATTGGTAGGTTACAGTGCACTC

ATCAGAGGCTAGTTAGCTATACATCTGCCCCCTTCTCCTCCGCCATGGGACCCCACTGACGATCCAGAAGATTAAAGAAA

ACTACATATTGGATAAGTATCTATTCCCAGCTTTGTCTGGT

>KX421388.1/goat/China/2007-08

ACCAAACAAAGTTGGGTAAGGATAGATCTTATAATAACTATGGACTGGCAAACTTAGGAGTAAAGATCCTACTGTCGGGG

GGAGGAGGAGGAGCAAGATCTCTGATTATGGCGACTCTCCTTAAAAGCTTAGCATTGTTCAAAAGGAACAAAGACAAAGC

GCCGACGGCATCGGGTTCAGGAGGGGCCATCCGAGGGATTAAGAATGTTATCATAGTCCCGATTCCCGGAGACTCGTCCA

TCACTACCCGTTCAAGACTGCTCGACAGGCTTGTCAGGTTGGCCGGAGATCCTGACATCAACGGGTCAAAGCTGACCGGC

GTGATGATCAGCATGCTATCCTTGTTCGTAGAGTCACCCGGGCAATTGATACAGCGAATCACAGATGATCCAGATGTTAG

TATCCGCCTTGTTGAGGTAGTTCAAAGTACTAGGTCCCAGTCCGGGTTGACCTTTGCATCACGTGGTGCTGATTTGGATA

ACGAGGCAGACATGTATTTTTCAACTGAGGGGCCCTCGAGTGGAGGTAAGAAAAGGATCAACTGGTTTGAGAACAGAGAA

ATAATAGACATAGAGGTGCAGGATCCAGAAGAGTTCAATATGTTGTTAGCCTCCATACTAGCACAAGTCTGGATCCTTCT

GGCCAAGGCTGTTACGGCACCGGATACGGCAGCTGACTCAGAACTGAGAAGGTGGGTTAAATACACACAACAAAGGAGAG

TGATTGGGGAATTTCGCCTTGACAAAGGGTGGCTGGACGCAGTCCGCAACAGAATTGCAGAAGATCTATCACTCCGGCGG

TTCATGGTATCTCTCATACTTGACATCAAGAGGACCCCCGGCAACAAGCCAAGGATTGCAGAAATGATCTGCGACATTGA

CAACTATATTGTCGAAGCAGGACTCGCCAGTTTCATCCTTACTATCAAATTTGGTATAGAAACCATGTATCCTGCATTAG

GGCTCCACGAGTTTGCCGGGGAATTGTCCACTATTGAGTCCTTGATGAACTTGTATCAACAGCTAGGAGAGGTTGCACCC

TACATGGTAATTCTAGAGAACTCAATTCAGAACAAGTTTAGTGCAGGAGCTTATCCCCTCCTCTGGAGCTATGCGATGGG

TGTCGGAGTTGAGCTGGAGAACTCAATGGGGGGCTTGAACTTTGGCAGGTCATATTTTGACCCTGCCTATTTTCGTCTCG

GACAGGAGATGGTCAGAAGATCTGCAGGAAAGGTCAGCTCTGTGATTGCGGCTGAGCTCGGCATCACAGCAGAGGAAGCT

AAACTAGTCTCGGAAATAGCCTCGCAGGCTGGGGACGAAAGAACCGCTAGAGGGACTGGGCCTCGACAGGCGCAGGTCTC

CTTCCTCCAGCACCAAACAGGAGGGGGAGAGTCGTCCGCACCAGCGACCAGAGAAGGGGTCAAAGCTGCGATCCCAAACG

GATCCGAAGAAAGGGACAGAAAGCAAACACGCCCAGGAAGGCCCAGAGGAGAGACCCCCGGCCAACTGCTCCTGGAAATC

ATGCCAGAGGATGAGGTTTCGCGAGAATCTGGTCAAAACCCTCGTGAGGCTCAAAGATCGGCCGAGGCACTCTTCAGGCT

GCAGGCCATGGCCAAGATTCTGGAGGACCAGGAGGAGGGAGAAGACAACAATCAGGTCTACAACGACAAGGATCTCCTCG

GCTAAGCAGACGCACCCTCTGTCGAAATCAGTGACGAGACATCTCCTACCAGTATTATAAAAAACTTAGGACCCAGGTCC

AAGCAAGCACACATCGACACTCCAACCAGTCGAGCGGAGGCCACCGATGGCAGAAGAACAAGCATACCATGTCAACAAGG

GGCTGGAATGTATCAAGTCCCTCAAAGCCTCTCCCCCGGATCTATCCACCATCAGAGATACCATCGAGAGCTGGAGAGAG

GGGCTTAGCCCATCGGGCCGTGCAACACCGAACCCTGATATGTCCGAGGGAGATCATCAGAATATCAACCAACCATGCTC

ACCAGCAATCGGACCAAACAAAGTCTACTTGTCTCCTGAAGATAATCTCGGATTTAGAGAGATCACTAGCAACGACTATG

AGGCTGAGCTCGGAGGAGTCCAGGGAAAAGGATCCAACTCTCAAGTACAGCGTTACTATGTTTATAGCCACGGGGGTGAA

GAGATTGAAGGACTCGAGGATGCTGACTCTCTCGTGGTTCAAGCAGATCCTCCAGTTGCTAACATATTCAATGGAGGAGA

GGATGGATCTGACGACAGCGATGTGGACTCTGGCCCAGATGATCCCGGCAGAGATACTCTATATGACCGGGGATCTGTTG

CCAGCAATGATGTCGCTAGGTCCACAGATGTCGAAAAACTAGAAGGTGCTGATATTCAAGAAGTTCTCAACTCCCAGAAA

GGCAGAGGAGGAAGATTCCAAGGCGGGAAAACCTTACGAGTCCCGGAAATACCCGATGTCAAGCACTCCAGACCATCAGC

CCAATCAATTAAAAAGGGCACAGACGGGAACTCAGTCTCATCTGGAACGGTGACAGAGTGTTTATCGATAAGTGGTGCAA

CCCAAGCTGTGCCAGAGTCAAGATGGGAGTCATCAGAGCAAAATGCGTCTGTGGGGAGTGTCCTCAAGTCTGCGGGGAGT

GCAAAGACGATCCAGGGGTCGACACAAGAATCTGGTACCATAGCATCACTGACTCAGCCTAAAGAGAATGACTCCGAGTA

TGAGTATGAGGATGACCTATTTACAGAGATTCAGGACATCCGTGCAAGCATTGCCAAGATCCATGATGACAACAAGGCTA

TCCTCTCAAAGCTTGATTCTATACTGTTATTGAAAGGAGAAGTCGACACTATCAAGAAACAAATCAGCAAGCAGAATATA

AGTATATCCACCATTGAGGGCCATCTCTCCAGTATAATGATAGCCATCCCGGGCTTTGGGAAGGATATCAAGGACCCAAC

ATCCGAGGTTGAGTTGAACCCAGATTTAAGACCTATAATAAGCCGTGACTCTGGCAGAGCTCTCGCGGAGGTCCTCAAGA

AACCCGCTGTGGATAGGTCTCAGAAAATTGGAACCAAAGCCAACTCCAGCTCAAAGGGTCAGCTTCTTAAGGATCTCCAG

CTAAAACCTGTCGACAAGCAGGCAAGCTCTGCAATCGGGTTTGTCCCATCCGACCATGAATCATCCAGAAATGTCATCCG

CTCCATAATCAAGTCGAGCAAGCTCAACATTGATCACAAGGACTATCTTCTAGATTTACTGAATGATGTGAAAGGCTCCA

AGGATCTTAAGGAATTCCACAAGATGCTAACAGCAATTCTCGCCAAGCACCCGTAACACATCCTCCAGTCACCACCCCAT

ACTCGGCTGAAAACATCCTTTCAATCAGGCTATTACAAAAAACTTAGGAGCAAGGGCAACTGAGCTTCGCAGACAAGATG

ACCGAGATCTACGACTTCGATAAATCAGCATGGGATGTCAAAGGGTCAATTGCCCGCATAGAACCCACCACCTACCACGA

CGGCCGACTGATACCCCAGGTAAGGGTCATCGATCCTGGTCTGGGAGACAGAAAAGATGAGTGCTTTATGTACCTGTTTC

TCCTGGGAGTGATTGAGGATAACGACCCCCTGTCTCCCCCCGTCGGAAGAACCTTTGGCTCTTTACCTCTAGGGGTCGGT

AGGTCAACTGCTAAGCCAGAGGAACTACTAAGGGAGGCCACAGAACTAGACATAGTGGTGAGGCGCACTGCAGGAGTAAA

TGAGAAACTGGTATTTTACAACAACACTCCGCTGTCCTTGTTAACGCCCTGGAAGAAAGTTCTGACAACCGGAAGTGTGT

TTAGCGCTAACCAGGTTTGCAATGCAGTCAACCTAGTCCCGCTTGATACTCCCCAGAGGTTCAGGGTTGTGTACATGAGC

ATAACTAGATTATCAGACAATGGTTACTATAGTGTGCCCAGAAGAATGCTGGAGTTCCGCTCAGCCAATGCAGTAGCCTT

CAACATCTTGGTTACACTAAAAATTGAAAATGGCACAAACCCTAGAAGATACATAGTCGGCTCATGGGAGAATTCAGAGG

TCACATTTATGGTACACGTGGGCAACTTTAGAAGAAAGAAGAACGAAGTATACTCTGCTGATTACTGCAAAATGAAGATT

GAAAAGATGGGTTTAGTTTTTGCCCTGGGAGGAATAGGTGGAACAAGTCTCCATATTAGAAGCACCGGGAAAATGAGCAA

AACCCTCCATGCACAGCTGGGGTTCAAGAAAATCTTATGTTACCCCCTAATGGATGTTAATGAGGATCTTAACCGATATC

TCTGGCGGGCAGAGTGCCGAATAGTCAAAATCCAAGCCGTCCTACAGCCATCAGTGCCCCAAGAATTCCGTGTCTACGAT

GATGTCATCATCAACGATGATCAAGGCTTGTTCAAGATCTTGTAGTTCATTTGCAAAGGAACGCACCCACCCCCAGGAAG

GGGCCCCA-CCCCCCCCG-CCACAAAACACACTCCAAGGCTCCCCGACACAGCGCCAACACAGCCACCCCCCCGACTGGA

CGAGGCCGGCC-CCCCAACAAACCCGGTCCAGGCGGGGGGCCCCCCCCCCTGACCCGCCGGGCGCCGACCGAGCAGGCCG

GCGAGAAGCCCCCAAGAGGCCCCGACAAAGACCGAGCACCCCATCCCCCC-AAAACGTCTCTCC-AACACGCTCGAGGCC

AGGCGCCGGGCACCCCCAGATCCGCCACCGGCGAGACAGAACCCCAGGGGGGGACCTCCCAAACCAAAGGAACCACCGGG

GAAGGCTCCGCCGAACCAGCACAGACCCGACCCAAGCAAGGCGGACCCCCAACACGAAACCGCCCAGCACACCCCACTCA

TCATCAAACAAAACTTAGGGGCCAAGCCCATACGACAGGCCCCAGTCCGCCAGGCGGAGGGACGCAGAAAGGAAGGAGAC

ATGACCGCCCCAGAAGGGGGACACCCAACCACCGAAACACCCAGCCCCAGGGAAGGAG--CCCCCCCCCACACGGCGAAG

GAACGGGTGAGAGACCCCGAGGAGAACACGACCGGGAGCCAGGAGCCGGCGGCGCCGGCAGCAAACAAGCCGGGCAAACA

CCCGACCCCACCAGAGGAGCCGAGGAGAGCCCGACCCCGCACGAACCAAACGGCAGGGAGCCCAACCCCACCACAAAAAG

AGGCGAGGCCCC-CCT-----CCCCCCCGCACACAACCCGCAGACAGACCGAGGGAGGACCACCCCCCAACCAGACAGGC

GGCCCCCCCGCAGCCAAGCTGCAGCTCCTCGGAACAGGACACCCCCGCACACTCCGAACACCCCCCCTCCCCCCGCCACC

GGGACAGGGGCTCCGCCACCACAGACCAAAGCCACAGAGAGCCAGCCAGAGGTCCCCCCCAGTCTCGCCCGGTGCCCCCC

CGAAAAGGCCCCGGCAAGGCCCCCCAACCCGACGTCCCCCGCACGCACCGAAAGAAGGGCCCGCCCGAGAACGTGCGGCA

CATCCATGCGCGAACATCATGACACGGGTCGCAATCTTGACATTTCTGTTTCTTCTCCCAAATGTTGTTGCGTGCCAGAT

TCACTGGGGCAATCTATCCAAGATCGGGATTGTAGGAACAGGGAGTGCCAGCTACAAGGTGATGACTAGGCCAAGCCACC

AAACTCTGGTTATAAAGTTAATGCCAAATATAACAGCCATCAACAATTGTACAAAGTCAGAGATTGCAGAGTACAAGAGA

TTGCTGATCACAGTGTTAAAACCTGTAGAGGATGCTCTGTCGGTGATAACCAAGAATGTAAGACCAATTCAAACCCTGAC

ACCTGGGCGTAGAACCCGTCGTTTTGCTGGAGCTGTTCTGGCCGGAGTAGCACTTGGAGTTGCGACAGCCGCTCAGATAA

CTGCAGGAGTCGCTCTTCATCAATCATTGATGAACTCTCAAGCAATTGAGAGTTTAAAAACCAGTCTTGAGAAGTCGAAT

CAGGCAATAGAAGAAATCAGACTTGCAAATAAGGAGACCATACTGGCAGTACAGGGCGTCCAGGATTATATCAACAATGA

GCTTGTCCCTTCTGTTCATAGAATGTCATGCGAGCTGGTAGGCCACAAGCTCGGCCTCAAGCTCCTTAGGTACTACACCG

AGATCCTGTCCATATTCGGGCCCAGCCTTCGAGACCCGATAGCTGCCGAAATATCGATCCAGGCACTCAGTTATGCATTA

GGCGGAGACATTAATAAAATCCTGGACAAGCTTGGGTATAGCGGTGGGGATTTCCTTGCCATCCTAGAGAGCAAGGGAAT

AAAGGCCCGGGTTACATATGTGGACACAAGAGATTACTTTATAATCCTTAGCATCGCCTACCCAACCTTATCTGAGATCA

AGGGAGTGATAGTTCACAAGATAGAAGCTATAACATACAACATTGGGGCACAGGAGTGGTATACTACTATCCCTAAATAT

GTAGCCACTCAGGGGTATCTGATATCGAACTTTGATGAGACGTCATGCGTATTCACTCCAGAGGGGACAGTTTGCAGCCA

GAATGCGTTGTACCCAATGAGCCCATTGCTTCAGGAATGTTTCAGGGGATCAACAAAATCGTGCGCCAGAACCCTAGTTT

CAGGGACCATAAGTAATAGATTTATCCTATCAAAAGGGAACCTGATTGCAAATTGTGCGTCAGTTTTGTGCAAATGTTAC

ACAACGGAGACAGTCATCAGCCAAGATCCTGACAAACTACTAACTGTTGTAGCATCCGACAAGTGTCCTGTAGTTGAGGT

GGATGGAGTGACAATACAGGTCGGCAGTCGAGAGTATCCGGATTCTGTATACTTACACAAAATAGACTTAGGTCCAGCCA

TCTCCCTAGAAAAACTGGATGTAGGCACCAATTTAGGCAATGCAGTCACAAGACTGGAGAATGCAAAGGAGCTCCTAGAT

GCATCAGACCAAATACTGAAGACTGTCAAAGGGGCACCTTTCGGCGGGAACATGTACATAGCACTGGCAGCTTGCATTGG

AGTATCCCTAGGGCTTGTCACATTAATATGCTGCTGTAAGGGGAGGTGTAAGAACAAGGAAATCCCTATCTCCAAAATCA

ACCCAGGGCTCAAACCCGATCTGACCGGGACCTCAAAGTCGTACGTGAGATCACTGTAGCCAGAATCACCTGAATCATCT

GGCATCACACACATACATGTACGACACAAGCAGTCTGAGGACGCAGGAAACCCAGCCTCCGATCACCTACCTGACCCTAC

TCCACGCTCCACTACACATTAGTCATCAAACAAAACTTAGGACGAAAGGTCAATCACCATGTCCGCACAAAGGGAAAGGA

TCAATGCCTTCTACAAAGACAATCCTCACAATAAGAACCATAGGGTGATCCTGGATAGAGAACGCTTGGTCATTGAAAGA

CCCTACATCTTGCTTGGAGTCCTGCTGGTAATGTTCCTGAGTCTAATCGGACTGCTGGCCATTGCAGGGATCAGGCTTCA

CCGGGCCACCGTTGGAACTTCAGAGATCCAGAGTCGGCTGAATACCAATATTAAGTTGACCGAATCTATTGATCACCAGA

CTAAGGATGTCTTAACTCCCCTTTTTAAAATCATTGGCGATGAAGTCGGCATCAGAATTCCACAGAAATTCAGTGATCTT

GTCAAGTTCATCTCCGATAAGATTAAATTCCTCAACCCTGATAGAGAGTATGATTTCAGGGATCTCCGGTGGTGCATGAA

TCCCCCCGAGAGAGTCAAAATTAATTTTGATCAGTTTTGTGAGTACAAAGCTGCGGTTAAGTCAATTGAACATATATTTG

AGTCACCACTCAACAAGTCAAAAAAGCTGCAATCATTGACTCTCGGGCCCGGAACAGGCTGTCTAGGCAGGACAGTAACA

AAAGCTCATTTCTCAGAACTTACACTGACCTTAATGGACCTGGATCTAGAGATGAAGCACAACGTGTCCTCAGTGTTTAC

CGTAGTTGAAGAGGGGTTATTCGGAAGAACATATACCGTCTGGAGATCCGATGCCAGGGATCCGAGCACCGATCTAGGTA

TCGGCCATTTTTTAAGAGTCTTCGAGATTGGACTGATAAGAGATCTCGGGCTGGGTCCCCCTGTTTTTCATATGACCAAC

TATCTCACAGTGAACATGAGTGATGACTATCGGAGATGTCTTTTAGCGGTAGGGGAGTTGAAGTTGACAGCCCTATGCAC

CTCATCTGAGACTGTGACACTGAGTGAGAGAGGAGCTCCAAAGAGGGAGCCTCTTGTGGTTGTGATACTTAATCTGGCTG

GACCCACTCTAGGGGGCGAACTATACAGTGTCTTGCCTACCTCCGATCTCATGGTGGAGAAACTCTATTTATCTTCACAT

AGAGGGATCATCAAGGATGACGAGGCCAATTGGGTAGTGCCGTCTACCGATGTTCGTGATCTTCAGAACAAAGGTGAATG

TCTGGTGGAAGCATGCAAGACTCGACCTCCTTCATTTTGCAATGGCACAGGATCAGGCCCGTGGTCAGAGGGGAGAATCC

CTGCTTACGGGGTGATCAGGGTCAGTCTTGACTTAGCTAGTGACCCGGATGTAGTTATCACTTCAGTGTTTGGCCCACTG

ATTCCTCACCTATCCGGCATGGATCTTTACAACAACCCGTTTTCAAGAGCTATATGGTTGGCTGTACCACCTTATGAGCA

GTCATTTCTAGGAATGATAAATACAATTGGATTCCCTAACAGAGCAGAGGTTATGCCGCACATTTTGACCACAGAGATCA

GAGGCCCTCGGGGTCGTTGCCATGTTCCCATAGAATTGTCCCGCAGGGTTGATGACGATATCAAGATCGGGTCCAACATG

GTCATATTGCCGACGATGGACCTGAGGTATATTACAGCCACTTATGATGTTTCCAGGAGCGAGCATGCAATCGTGTACTA

TATCTATGACACAGGTCGCTCATCATCTTACTTCTACCCAGTTCGACTGAATTTCAAAGGCAATCCTCTCTCTCTGAGGA

TAGAGTGTTTCCCTTGGCGTCATAAGGTGTGGTGCTACCATGATTGTCTTATATACAACACCATAACAGATGAAGAGGTC

CATACGAGAGGGCTGACCGGTATAGAGGTAACATGCAATCCAGTTTGAGCAAAGCTACGACCGTCACTCAAGCAGTCCCT

TGAGTCGCCACCGAGTCCAAGCAGCATAGCCTGGGACACTCAACAGCACAGCCCAGCCAACAATGTTATAAAAAACTTAG

GAGCCAGGGTTGTAGGGGCCATGGACTCCCTATCAGTCAATCAGGTCTTGTACCCTGAGGTTCATCTAGACAGCCCTATT

GTCACAAACAAACTAGTTGCCATCCTTGAGTACTCGGGTATCGACCATAACTATGTTCTTGAAGACCAGACCCTTGTCAA

GAATATTAGGTATAGACTGGGGTGCGGTTTTTCAAATCAAATGATCATCAATAACAGGGGGGTGGGTGAAACAGTCAATT

CTAAACTTAAAAGTTATCCCCATAATCGTCATATCATATACCCGGATTGCAATAAGGAGTTGTTTGGTATCAAAGATAGC

TGCATATCTAAGAAGCTCTCGGAGCTATTCAAGAAGGGTAATTCCTTGTACTCTAAGATAAGTCACCAGGTACTGGATTG

TCTTAAGAGAGTCAATGGGAAATTAGGCCTGGGCACAGATCTTACCCATGGTCTGAAGGAAGGTATCCTTGACCTAGGGT

TGCACATGCATAGCTCTCAATGGTTCGAGACTTTCCTGTTCTGGTTCACTATCAAGACAGAGATGAGATCAATGATCAAA

GAACAGTCCCATATATGCCACAAGAGGAGGTATAACCCGATTTTTGTATCGGGGGATGCATTCGAGGTGCTCGTATCACG

AGACCTCGTTGTGATAATTGATAAGAATACCCAGTATGTCTTCTACTTAACATTTGAGCTGGTCCTTATGTATTGTGATG

TCATAGAGGGCAGACTTATGACGGAGACAGCCATGGCCATAGACCAGAGATATTCAGAACTTCTAAGCCGGGTTAGATAC

TTGTGGGATCTTATTGATGGGTTCTTCCCAACACTAGGCAACACCACATACCAAATTGTTGCTCTGCTTGAACCATTATC

GTTGGCTTATCTTCAACTTCAGGATGTCACGCTGGAGTTAAGAGGTGCTTTTTTAGACCATTGCTTCAAAGAACTCTATG

AGATACTGGAGCATTGCGGCATTGACACAGAAGGCACCTACAATTCCATCACTGAAGGACTGGATTACGTATTCATCACC

CACGATATACATCTAACTGGGGAGATTTTTTCATTTTTCCGGAGTTTCGGACACCCCCGTCTAGAAGCAGTCACCGCTGC

AGAAAATGTCAGGAAACATATGAACCAACCGAAGGTAATCAGTTATGAGACTATGATGAAAGGACATGCGGTATTTTGCG

GGATAATCATAAATGGTTTTAGGGACCGACATGGCGGCAGCTGGCCCCCTGTTGCATTGCCAGAACATGCTTCTGCTGCG

ATCCGGAATGCGCAGGCATCAGGTGAGGGACTGACCCATGACCTGTGTATAGACAACTGGAAATCCTTTGTTGGATTCAA

ATTTGGCTGCTTTATGCCACTCAGCCTAGATAGTGATTTGACCATGTATCTCAAAGACAAGGCATTGGCTGCACTGAAGA

ATGAGTGGGATTCAGTTTACCCGAAAGAATACCTCCGTTATAACCCACCTAGAGGGACAGAGTCAAGGCGACTGGTGGAG

GTGTTCCTGAATGACTCCAGCTTTGATCCTTATAACATGATAATGTACGTGGTGAATGGCTCCTACCTTAAAGACCCTGA

GTTTAATCTCTCATACAGCCTAAAAGAGAAGGAGATAAAGGAGACAGGGCGGTTGTTTGCCAAAATGACTTATAAGATGC

GGGCCTGTCAGGTAATTGCTGAAAATCTGATATCGAATGGTGTTGGGAAGTATTTCCGAGACAATGGGATGGCAAAGGAC

GAGCATGACCTAACAAAAGCCCTTCACACCCTGGCAGTCTCAGGTGTTCCCAAGAACAACAAAGATAACCACCGAGGTGG

GCCTCCCAGAAGGACAACAAGCCGAGAGGTGAGATCAAGCCAAGACACCAAAACACAAAATAGAGACAAGGTCCAAGGAG

GACCTATGTACAACTATTTGCGATGCCAACCAATCAGCCCTGATCAGGGTGAGTCATACGAGACTGTTAGTGCATTCATC

ACCGCTGACCTTAAGAAGTATTGCCTGAATTGGAGATACGAGACAATCAGCATATTTGCACAGAGGCTGAATGAAATATA

TGGACTACCATCCTTCTTTCAATGGTTACACAGGATATTGGAAAAATCCGTACTCTACGTCAGTGACCCACATTGCCCTC

CCGATCTAGATAATCATATCCCTCTGGACAGTGTCCCTAATGCCCAAATATTCATTAAGTACCCAATGGGCGGAATAGAA

GGTTATTGCCAAAAACTATGGACAATCAGTACTATACCATACTTGTATCTGGCAGCCTATGAGAGCGGAGTAAGAATCGC

CTCACTGGTGCAGGGTGACAATCAGACAATCGCAGTGACAAAAAGAGTTCCAAGCTCTTGGCCTTATTCACTAAAAAAGA

AGGAGGCATCCAGAGCCGCTCAAAATTACTTCGTAGTCTTAAGGCAAAGATTGCACGATGTAGGTCATCACTTAAAGGCT

AATGAAACCATAGTATCTTCTCACTTTTTTGTATATTCCAAAGGGATTTATTATGACGGCCTGCTAGTCTCACAATCACT

GAAGAGCATCGCCAGATGTGTCTTCTGGTCCGAGACTATAGTGGATGAAACCAGAGCGGCTTGCAGCAATATTGCAACAA

CTGTCGCTAAGAGTATAGAGAGGGGTTATGATAGGTACCTTGCATATTCTTTGAATATCCTCAAGATTTTCCAACAGATC

CTTATATCCCTTAACTTCACTATTAACACAACAATGACTCAGGATGTCGTGGCACCGATCATCGAGAACGGTGATTTGCT

GATAAGGATGGCACTCTTGCCAGCACCCATCGGAGGTCTAAATTATCTTAACATGAGCAGGTTATTTGTGAGAAATATCG

GTGACCCGGTCACCTCCTCTATAGCCGACCTGAAGAGGATGATAGATGCCGGGCTAATGCCAGAAGAAACATTGCATCAA

GTGATGACCCAGACCCCGGGAGAATCATCCTACCTCGATTGGGCAAGTGACCCTTATTCTGCCAACCTACCCTGCGTACA

GAGTATAACTCGCCTTCTCAAAAACATCACTGCACGATATATTTTAATCAGCAGCCCAAACCCGATGCTGAGGGGATTGT

TTCATGAGGGGAGTAGAGATGAAGACGAGGAGCTTGCAAGTTTCCTAATGGATCGGCATATAATTGTTCCAAGAGCTGCA

CATGAAATCTTAGACCATAGTATAACCGGGGCAAGAGAAGCTATAGCCGGGATGTTGGACACCACCAAAGGTCTGATTAG

AACAAGTATGAAACGGGGTGGCCTCACCCCTCGAGTCTTAGCCCGCCTTTCCAATTATGATTATGAACAATTCCGATCCG

GGATAACACTATTGACAAAGAAAGGGCAGTGTTATCTCATTGACAAGGATTCGTGCTCGGTGCAGCTCGCTATAGCTCTG

AGGAGCCACATGTGGGCTAGGTTGGCTCGCGGGAGACCAATCTATGGGTTGGAGGTGCCCGATATATTAGAATCGATGAA

CGGCTACCTTATCAAACGCCACGAGTCTTGTGCTATCTGTGAAACGGGCTCAAGTCACTACGGATGGTTTTTCGTCCCTG

CAGGGTGCCAGCTTGACGATGTATCTAGAGAGACTTCGGCTCTTCGTGTACCTTATGTCGGATCAACAACTGAGGAAAGG

ACAGACATGAAACTTGCATTTGTTAGATCTCCAAGTAGATCCCTCAAATCAGCAGTTAGAATTGCAACAGTTTACTCATG

GGCCTACGGGGATGATGAGAAGTCATGGAGTGAAGCCTGGATGCTAGCTAGGCAGAGAGCCAATATCACCTTAGATGAAT

TGAGAATGATAACTCCAGTCTCTACATCCACCAACCTAGCCCACCGGTTGAGGGATCGGAGCACCCAGGTGAAATACTCG

GGGACATCCCTTGTGAGGGTTGCAAGATACACAACCATCTCCAATGATAACTTGTCATTTGTGATATCTGAGAAAAAAGT

GGATACCAACTTCATTTATCAGCAAGGGATGTTGCTTGGTCTTGGGATCCTTGAGAACCTCTTTAGGCTAGAGGCCACCA

CCGGAGTATCCAACACAGTGCTACACCTGCACGTGGAAACAGAATGTTGTGTTGTACCTATGGTGGATCACCCAAGGATA

CCGAGTCTCCGTAATATAAAAGTTACGGATGAGCTATGCACAAACCCTTTGATCTACGACAGGTCTCCCATCATAGAGCA

CGATGCAACCCGACTATACTCACAGAGCCACAGGAGACATTTGGTGGAGTTTGTTACCTGGTCCACAAGCCAGCTTTACC

ATATACTGGCTAAGTCTACAGCAATGTCCATGATTGAATTGATCACGAGATTCGAGAAAGATCACATGAATGAAATAGCC

GCCCTGATTGGCGATGACGACATCAACAGCTTCATCACAGAATTTTTGCTAGTGGAGCCTAGATTATTTATAGTTTACCT

TGGTCAGTGTGCTGCCATCAATTGGGCCTTTGATATACATTATCATCGGCCCTCGGGCAAGTACCAGATGGGGGAACTCC

TCTACTCTTTGCTCTCTCGGATGAGCAAAGGAGTATATAAGATCTTCACCAATGCTTTGAGCCACCCCAAAGTTTACAAG

AAATTTTGGCGAAGTGGTGTCATTGAGCCGATTCATGGCCCATCCCTGGATACACAGAATTTACATGTCACTGTCTGTGA

CATGATATATGGGTCCTATGTCACCTATTTGGATCTTTTGCTGAATGATGAGCTAGATGATTACCCGTATTTGCTCTGCG

AGAGTGATGAGGACGTGGTCACAGACAGGTTCGACAACATTCAAGCCAAACATCTCTGTGTATTAGCCGATGTATATTGC

AGCTCCAAGAGGTGTCCTTCAATAATCGGGATGTCTCCTATAGAAAAATGTACCATTCTCACACATTACATCAAGGGAGA

ATCAGTACAATCCCCGTCTGGGACCTCATGGAACACTGATCCCCTTGTAGTAGATCATTACTCATGCTCTCTGACCTACC

TTCGCCGCGGTTCCATCAAACAAATCAGGTTGAGGGTGGATCCTGGGTTTGTATTCGAGGCGTTGACAGACGTCGACCTT

AAACAACCTCGCAAAGCTAAGTTAGATGTATCGGTCGTTGGGTTGACTGATTTCTCTCCCCCTTGCGACAACGTCGGTGA

TTTTCTAGGGACTATCAACACATTGAGGCACAATCTGCCTGTCACCGGAACCGGGGTCTCGAACTATGAAGTCCACGCTT

ATCGTAGAATTGGCCTAAATTCATCAGCATGTTACAAAGCCGTAGAGATCTCCACGTTAATCAAGCCATCCCTAGAAGTC

GGAGAGCATGGATTGTTCTTAGGAGAAGGTTCCGGTTCAATGCTGGCTGCGTACAAGGAAGTTCTTAAATTAGCAAATTG

TTATTACAACAGCGGGGTAACAACGGAGGGCAGAGCCGGACAGAGGGAAATATCTCCCTATCCTTCAGAGATGAGCCTAG

TAGAGAATCAAATGGGGATAGAGCGGAGTGTTAAAGTGCTGTTCAACGGCAAACCTGAAGTAACTTGGGTGGGGACCACC

GATTGCTACAAGTACATAATCAGTAACATCCAAACCTCTAGTCTGGGTTTCATACATTCAGATATTGAGACGCTCCCAAC

CAAAGATGCTGTTGAGAAATTAGAAGAATTTGCCTCTATCCTATCCTTATCCTTGATTTTGGGGAAAATCGGCTCTATTA

CAGTTGTCAAAATTATGCCCATTAGCGGGGATTTTACCCAAGGCTTTATAGCCTATGCCATCCAATATTTTAGGGAGAGC

CTGCTTGCCTATCCGAGATATAGTAACTTCATCTCGACTGAGTGTTACCTTATTATGATAGGATTAAAGGCCAATCGACT

GATAAACCCAGAAGCCATTAAGCAAAGCATAATCAGAGCGGGGGTCAGGACTGCACCAGGACTTGTGAGCCATATATTAT

CAGAGAAACAGAAAGGTTGTATTCAATCTTTCTTGGGTGATCCTTATATCCAAGGAGACTTCAATAAGCACCTTAAATCT

CTGACCCCTATTGAGAAAATCCTGGTAAATTGTGGTCTCTCGATCAATGGCACAAAAATCTGTAGAGATCTAATCCACCA

TGATATCGCCTCCGGTCCAGACGGTCTTATGAGCTCTACAATTATTTTATATAGGGAATTGGCTCATTTCAAAGATAATA

TAAGGAGTCAGCACGGAATGTTCCACCCCTACCCAGTACTGGCCAACAGCCGGCAACGTGAATTAATCCTTCGAATAGCC

AAGAAATTCTGGGGGTATGTCTTGCTATATTCTGATGACCCGGCACTAATCAGACAAACAATCAAGAACTTGAAACGGAA

TCATCTAACCTTTGACTTACATAGTAACCCGTTTATTAAAGGGCTATCCAAAGCTGAGAAACTGCTGGTGCGGACGAGTC

CACTAAGAAGGGAATGGTTGTTCGCTCTCGATACGAAAGAAGTGAAAGAGTGGTTCAAATTGGTAGGTTACAGTGCACTC

ATCAGAGGCTAGTTAGCTATACATCTGCCCCCTTCTCCTCCGCCATGAGACCCCACTGACGATCCAGAAGATTAAAGAAA

ACTACATATTGGATAAGTATCTATTCCCAGCTTTGTCTGGT

>JF939201.1/goat/China/2007-12

ACCAAACAAAGTTGGGTAAGGATAGATCTTATAATAACTATGGACTGGCAAACTTAGGAGTAAAGATCCTACTGTCGGGG

GGAGGAGGAGGAGCAAGATCTCTGATTATGGCGACTCTCCTTAAAAGCTTAGCATTGTTCAAAAGGAACAAAGACAAAGC

GCCGACGGCATCGGGTTCAGGAGGGGCCATCCGAGGGATTAAGAATGTTATCATAGTCCCGATTCCCGGAGACTCGTCCA

TCACTACCCGTTCAAGACTGCTCGACAGGCTTGTCAGGTTGGCCGGAGATCCTGACATCAACGGGTCAAAGCTGACCGGC

GTGATGATCAGCATGCTATCCTTGTTCGTAGAGTCACCCGGGCAATTGATACAGCGAATCACAGATGATCCAGATGTTAG

TATCCGCCTTGTTGAGGTAGTTCAAAGTACTAGGTCCCAGTCCGGGTTGACCTTTGCATCACGTGGTGCTGATTTGGATA

ACGAGGCAGACATGTATTTTTCAACTGAGGGGCCCTCGAGTGGAGGTAAGAAAAGGATCAACTGGTTTGAGAACAGAGAA

ATAATAGACATAGAGGTGCAGGATCCAGAAGAGTTCAATATGTTGTTAGCCTCCATACTAGCACAAGTCTGGATCCTTCT

GGCCAAGGCTGTTACGGCACCGGATACGGCAGCTGACTCAGAACTGAGAAGGTGGGTTAAATACACACAACAAAGGAGAG

TGATTGGGGAATTTCGCCTTGACAAAGGGTGGCTGGACGCAGTCCGCAACAGAATTGCAGAAGATCTATCACTCCGGCGG

TTCATGGTATCTCTCATACTTGACATCAAGAGGACCCCCGGCAACAAGCCAAGGATTGCAGAAATGATCTGCGACATTGA

CAACTATATTGTCGAAGCAGGACTCGCCAGTTTCATCCTTACTATCAAATTTGGTATAGAAACCATGTATCCTGCATTAG

GGCTCCACGAGTTTGCCGGGGAATTGTCCACTATTGAGTCCTTGATGAACTTGTATCAACAGCTAGGAGAGGTTGCACCC

TACATGGTAATTCTAGAGAACTCAATTCAGAACAAGTTTAGTGCAGGAGCTTATCCCCTCCTCTGGAGCTATGCGATGGG

TGTCGGAGTTGAGCTGGAGAACTCAATGGGGGGCTTGAACTTTGGCAGGTCATATTTTGACCCTGCCTATTTTCGTCTCG

GACAGGAGATGGTCAGAAGATCTGCAGGAAAGGTCAGCTCTGTGATTGCGGCTGAGCTCGGCATCACAGCAGAGGAAGCT

AAACTAGTCTCGGAAATAGCCTCGCAGGCTGGGGACGAAAGAACCGCTAGAGGGACTGGGCCTCGACAGGCGCAGGTCTC

CTTCCTCCAGCACCAAACAGGAGGGGGAGAGTCGTCCGCACCAGCGACCAGAGAAGGGGTCAAAGCTGCGATCCCAAACG

GATCCGAAGAAAGGGACAGAAAGCAAACACGCCCAGGAAGGCCCAGAGGAGAGACCCCCGGCCAACTGCTCCTGGAAATC

ATGCCAGAGGATGAGGTTTCGCGAGAATCTGGTCAAAACCCTCGTGAGGCTCAAAGATCGGCCGAGGCACTCTTCAGGCT

GCAGGCCATGGCCAAGATTCTGGAGGACCAGGAGGAGGGAGAAGACAACAATCAGGTCCACAACGACAAGGATCTCCTCG

GCTAAGCAGACGCACCCTCTGTCGAAATCAGTGACGAGACATCTCCTACCAGTATTATAAAAAACTTAGGACCCAGGTCC

AAGCAAGCACACATCGACACTCCAACCAGTCGAGCGGAGGCCACCGATGGCAGAAGAACAAGCATACCATGTCAACAAGG

GGCTGGAATGTATCAAGTCCCTCAAAGCCTCTCCCCCGGATCTATCCACCATCAGAGATACCATCGAGAGCTGGAGAGAG

GGGCTTAGCCCATCGGGCCGTGCAACACCGAACCCTGATATGTCCGAGGGAGATCATCAGAATATCAACCAACCATGCTC

ACCAGCAATCGGACCAAACAAAGTCTACTTGTCTCCTGAAGATAATCTCGGATTTAGAGAGATCACTAGCAACGACTATG

AGGCTGAGCTCGGAGGAGTCCAGGGAAAAGGATCCAACTCTCAAGTACAGCGTTACTATGTTTATAGCCACGGGGGTGAA

GAGATTGAAGGACTCGAGGATGCTGACTCTCTCGTGGTTCAAGCAGATCCTCCAGTTGCTAACATATTCAATGGAGGAGA

GGATGGATCTGACGACAGCGATGTGGACTCTGGCCCAGATGATCCCGGCAGAGATACTCTATATGACCGGGGATCTGTTG

CCAGCAATGATGTCGCTAGGTCCACAGATGTCGAAAAACTAGAAGGTGCTGATATTCAAGAAGTTCTCAACTCCCAGAAA

GGCAGAGGAGGAAGATTCCAAGGCGGGAAAACCTTACGAGTCCCGGAAATACCCGATGTCAAGCACTCCAGACCATCAGC

CCAATCAATTAAAAAGGGCACAGACGGGAACTCAGTCTCATCTGGAACGGTGACAGAGTGTTTATCGATAAGTGGTGCAA

CCCAAGCTGTGCCAGAGTCAAGATGGGAGTCATCAGAGCAAAATGCGTCTGTGGGGAGTGTCCTCAAGTCTGCGGGGAGT

GCAAAGACGATCCAGGGGTCGACACAAGAATCTGGTACCATAGCATCACTGACTCAGCCTAAAGAGAATGACTCCGAGTA

TGAGTATGAGGATGACCTATTTACAGAGATTCAGGACATCCGTGCAAGCATTGCCAAGATCCATGATGACAACAAGGCTA

TCCTCTCAAAGCTTGATTCTATACTGTTATTGAAAGGAGAAGTCGACACTATCAAGAAACAAATCAGCAAGCAGAATATA

AGTATATCCACCATTGAGGGCCATCTCTCCAGTATAATGATAGCCATCCCGGGCTTTGGGAAGGATATCAAGGACCCAAC

ATCCGAGGTTGAGTTGAACCCAGATTTAAGACCTATAATAAGCCGTGACTCTGGCAGAGCTCTCGCGGAGGTCCTCAAGA

AACCCGCTGTGGATAGGTCTCAGAAAATTGGAACCAAAGCCAACTCCAGCTCAAAGGGTCAGCTTCTTAAGGATCTCCAG

CTAAAACCTGTCGACAAGCAGGCAAGCTCTGCAATCGGGTTTGTCCCATCCGACCATGAATCATCCAGAAATGTCATCCG

CTCCATAATCAAGTCGAGCAAGCTCAACATTGATCACAAGGACTATCTTCTAGATTTACTGAATGATGTGAAAGGCTCCA

AGGATCTTAAGGAATTCCACAAGATGCTAACAGCAATTCTCGCCAAGCACCCGTAACACATCCTCCAGTCACCACCCCAT

ACTCGGCTGAAAACATCCTTTCAATCAGGCTATTACAAAAAACTTAGGAGCAAGGGCAACTGAGCTTCGCAGACAAGATG

ACCGAGATCTACGACTTCGATAAATCAGCATGGGATGTCAAAGGGTCAATTGCCCGCATAGAACCCACCACCTACCACGA

CGGCCGACTGATACCCCAGGTAAGGGTCATCGATCCTGGTCTGGGAGACAGAAAAGATGAGTGCTTTATGTACCTGTTTC

TCCTGGGAGTGATTGAGGATAACGACCCCCTGTCTCCCCCCGTCGGAAGAACCTTTGGCTCTTTACCTCTAGGGGTCGGT

AGGTCAACTGCTAAGCCAGAGGAACTACTAAGGGAGGCCACAGAACTAGACATAGTGGTGAGGCGCACTGCAGGAGTAAA

TGAGAAACTGGTATTTTACAACAACACTCCGCTGTCCTTGTTAACGCCCTGGAAGAAAGTTCTGACAACCGGAAGTGTGT

TTAGCGCTAACCAGGTTTGCAATGCAGTCAACCTAGTCCCGCTTGATACTCCCCAGAGGTTCAGGGTTGTGTACATGAGC

ATAACTAGATTATCAGACAATGGTTACTATAGTGTGCCCAGAAGAATGCTGGAGTTCCGCTCAGCCAATGCAGTAGCCTT

CAACATCTTGGTTACACTAAAAATTGAAAATGGCACAAACCCTAGAAGATACATAGTCGGCTCATGGGAGAATTCAGAGG

TCACATTTATGGTACACGTGGGCAACTTTAGAAGAAAGAAGAACGAAGTATACTCTGCTGATTACTGCAAAATGAAGATT

GAAAAGATGGGTTTAGTTTTTGCCCTGGGAGGAATAGGTGGAACAAGTCTCCATATTAGAAGCACCGGGAAAATGAGCAA

AACCCTCCATGCACAGCTGGGGTTCAAGAAAATCTTATGTTACCCCCTAATGGATGTTAATGAGGATCTTAACCGATATC

TCTGGCGGGCAGAGTGCCGAATAGTCAAAATCCAAGCCGTCCTACAGCCATCAGTGCCCCAAGAATTCCGTGTCTACGAT

GATGTCATCATCAACGATGATCAAGGCTTGTTCAAGATCTTGTAGTTCATTTGCAAAGGAACGCACCCACCCCCAGGAAG

GGGCCCCA-CCCCCCCCG-CCACAAAACACACTCCAAGGCTCCCCGACACAGCGCCAACACAGCCACCCCCCCGACTGGA

CGAGGCCGGCC-CCCCAACAAACCCGGTCCAGGCGGGGGGCCCCCCCCCCTGACCCGCCGGGCGCCGACCGAGCAGGCCG

GCGAGAAGCCCCCAAGAGGCCCCGACAAAGACCGAGCACCCCATCCCCCC-AAAACGTCTCTCC-AACACGCTCGAGGCC

AGGCGCCGGGCACCCCCAGATCCGCCACCGGCGAGACAGAACCCCAGGGGGGGACCTCCCAAACCAAAGGAACCACCGGG

GAAGGCTCCGCCGAACCAGCACAGACCCGACCCAAGCAAGGCGGACCCCCAACACGAAACCGCCCAGCACACCCCACTCA

TCATCAAACAAAACTTAGGGGCCAAGCCCATACGACAGGCCCCAGTCCGCCAGGCGGAGGGACGCAGAAAGGAAGGAGAC

ATGACCGCCCCAGAAGGGGGACACCCAACCACCGAAACACCCAGCCCCAGGGAAGGAG--CCCCCCCCCACACGGCGAAG

GAACGGGTGAGAGACCCCGAGGAGAACACGACCGGGAGCCAGGAGCCGGCGGCGCCGGCAGCAAACAAGCCGGGCAAACA

CCCGACCCCACCAGAGGAGCCGAGGAGAGCCCGACCCCGCACGAACCAAACGGCAGGGAGCCCAACCCCACCACAAAAAG

AGGCGAGGCCCC-CCT-----CCCCCCCGCACACAACCCGCAGACAGACCGAGGGAGGACCACCCCCCAACCAGACAGGC

GGCCCCCCCGCAGCCAAGCTGCAGCTCCTCGGAACAGGACACCCCCGCACACTCCGAACACCCCCCCTCCCCCCGCCACC

GGGACAGGGGCTCCGCCACCACAGACCAAAGCCACAGAGAGCCAGCCAGAGGTCCCCCCCAGTCTCGCCCGGTGCCCCCC

CGAAAAGGCCCCGGCAAGGCCCCCCAACCCGACGTCCCCCGCACGCACCGAAAGAAGGGCCCGCCCGAGAACGTGCGGCA

CATCCATGCGCGAACATCATGACACGGGTCGCAATCTTGACATTTCTGTTTCTTCTCCCAAATGTTGTTGCGTGCCAGAT

TCACTGGGGCAATCTATCCAAGATCGGGATTGTAGGAACAGGGAGTGCCAGCTACAAGGTGATGACTAGGCCAAGCCACC

AAACTCTGGTTATAAAGTTAATGCCAAATATAACAGCCATCAACAATTGTACAAAGTCAGAGATTGCAGAGTACAAGAGA

TTGCTGATCACAGTGTTAAAACCTGTAGAGGATGCTCTGTCGGTGATAACCAAGAATGTAAGACCAATTCAAACCCTGAC

ACCTGGGCGTAGAACCCGTCGTTTTGCTGGAGCTGTTCTGGCCGGAGTAGCACTTGGAGTTGCGACAGCCGCTCAGATAA

CTGCAGGAGTCGCTCTTCATCAATCATTGATGAACTCTCAAGCAATTGAGAGTTTAAAAACCAGTCTTGAGAAGTCGAAT

CAGGCAATAGAAGAAATCAGACTTGCAAATAAGGAGACCATACTGGCAGTACAGGGCGTCCAGGATTATATCAACAATGA

GCTTGTCCCTTCTGTTCATAGAATGTCATGCGAGCTGGTAGGCCACAAGCTCGGCCTCAAGCTCCTTAGGTACTACACCG

AGATCCTGTCCATATTCGGGCCCAGCCTTCGAGACCCGATAGCTGCCGAAATATCGATCCAGGCACTCAGTTATGCATTA

GGCGGAGACATTAATAAAATCCTGGACAAGCTTGGGTATAGCGGTGGGGATTTCCTTGCCATCCTAGAGAGCAAGGGAAT

AAAGGCCCGGGTTACATATGTGGACACAAGAGATTACTTTATAATCCTTAGCATCGCCTACCCAACCTTATCTGAGATCA

AGGGAGTGATAGTTCACAAGATAGAAGCTATAACATACAACATTGGGGCACAGGAGTGGTATACTACTATCCCTAAATAT

GTAGCCACTCAGGGGTATCTGATATCGAACTTTGATGAGACGTCATGCGTATTCACTCCAGAGGGGACAGTTTGCAGCCA

GAATGCGTTGTACCCAATGAGCCCATTGCTTCAGGAATGTTTCAGGGGATCAACAAAATCGTGCGCCAGAACCCTAGTTT

CAGGGACCATAAGTAATAGATTTATCCTATCAAAAGGGAACCTGATTGCAAATTGTGCGTCAGTTTTGTGCAAATGTTAC

ACAACGGAGACAGTCATCAGCCAAGATCCTGACAAACTACTAACTGTTGTAGCATCCGACAAGTGTCCTGTAGTTGAGGT

GGATGGAGTGACAATACAGGTCGGCAGTCGAGAGTATCCGGATTCTGTATACTTACACAAAATAGACTTAGGTCCAGCCA

TCTCCCTAGAAAAACTGGATGTAGGCACCAATTTAGGCAATGCAGTCACAAGACTGGAGAATGCAAAGGAGCTCCTAGAT

GCATCAGACCAAATACTGAAGACTGTCAAAGGGGCACCTTTCGGCGGGAACATGTACATAGCACTGGCAGCTTGCATTGG

AGTATCCCTAGGGCTTGTCACATTAATATGCTGCTGTAAGGGGAGGTGTAAGAACAAGGAAATCCCTATCTCCAAAATCA

ACCCAGGGCTCAAACCCGATCTGACCGGGACCTCAAAGTCGTACGTGAGATCACTGTAGCCAGAATCACCTGAATCATCT

GGCATCACACACATACATGTACGACACAAGCAGTCTGAGGACGCAGGAAACCCAGCCTCCGATCACCTACCTGACCCTAC

TCCACGCTCCACTACACATTAGTCATCAAACAAAACTTAGGACGAAAGGTCAATCACCATGTCCGCACAAAGGGAAAGGA

TCAATGCCTTCTACAAAGACAATCCTCACAATAAGAACCATAGGGTGATCCTGGATAGAGAACGCTTGGTCATTGAAAGA

CCCTACATCTTGCTTGGAGTCCTGCTGGTAATGTTCCTGAGTCTAATCGGACTGCTGGCCATTGCAGGGATCAGGCTTCA

CCGGGCCACCGTTGGAACTTCAGAGATCCAGAGTCGGCTGAATACCAATATTAAGTTGGCCGAATCTATTGATCACCAGA

CTAAGGATGTCTTAACTCCCCTTTTTAAAATCATTGGCGATGAAGTCGGCATCAGAATTCCACAGAAATTCAGTGATCTT

GTCAAGTTCATCTCCGATAAGATTAAATTCCTCAACCCTGATAGAGAGTATGATTTCAGGGATCTCCGGTGGTGCATGAA

TCCCCCCGAGAGAGTCAAAATTAATTTTGATCAGTTTTGTGAGTACAAAGCTGCGGTTAAGTCAATTGAACATATATTTG

AGTCACCACTCAACAAGTCAAAAAAGCTGCAATCATTGACTCTCGGGCCCGGAACAGGCTGTCTAGGCAGGACAGTAACA

AAAGCTCATTTCTCAGAACTTACACTGACCTTAATGGACCTGGATCTAGAGATGAAGCACAACGTGTCCTCAGTGTTTAC

CGTAGTTGAAGAGGGGTTATTCGGAAGAACATATACCGTCTGGAGATCCGATGCCAGGGATCCGAGCACCGATCTAGGTA

TCGGCCATTTTTTAAGAGTCTTCGAGATTGGACTGATAAGAGATCTCGGGCTGGGTCCCCCTGTTTTTCATATGACCAAC

TATCTCACAGTGAACATGAGTGATGACTATCGGAGATGTCTTTTAGCGGTAGGGGAGTTGAAGTTGACAGCCCTATGCAC

CTCATCTGAGACTGTGACACTGAGTGAGAGAGGAGCTCCAAAGAGGGAGCCTCTTGTGGTTGTGATACTTAATCTGGCTG

GACCCACTCTAGGGGGCGAACTATACAGTGTCTTGCCTACCTCCGATCTCATGGTGGAGAAACTCTATTTATCTTCACAT

AGAGGGATCATCAAGGATGACGAGGCCAATTGGGTAGTGCCGTCTACCGATGTTCGTGATCTTCAGAACAAAGGTGAATG

TCTGGTGGAAGCATGCAAGACTCGACCTCCTTCATTTTGCAATGGCACAGGATCAGGCCCGTGGTCAGAGGGGAGAATCC

CTGCTTACGGGGTGATCAGGGTCAGTCTTGACTTAGCTAGTGACCCGGATGTAGTTATCACTTCAGTGTTTGGCCCACTG

ATTCCTCACCTATCCGGCATGGATCTTTACAACAACCCGTTTTCAAGAGCTATATGGTTGGCTGTACCACCTTATGAGCA

GTCATTTCTAGGAATGATAAATACAATTGGATTCCCTAACAGAGCAGAGGTTATGCCGCACATTTTGACCACAGAGATCA

GAGGCCCTCGGGGTCGTTGCCATGTTCCCATAGAATTGTCCCGCAGGGTTGATGACGATATCAAGATCGGGTCCAACATG

GTCATATTGCCGACGATGGACCTGAGGTATATTACAGCCACTTATGATGTTTCCAGGAGCGAGCATGCAATCGTGTACTA

TATCTATGACACAGGTCGCTCATCATCTTACTTCTACCCAGTTCGACTGAATTTCAAAGGCAATCCTCTCTCTCTGAGGA

TAGAGTGTTTCCCTTGGCGTCATAAGGTGTGGTGCTACCATGATTGTCTTATATACAACACCATAACAGATGAAGAGGTC

CATACGAGAGGGCTGACCGGTATAGAGGTAACATGCAATCCAGTTTGAGCAAAGCTACGACCGTCACTCAAGCAGTCCCT

TGAGTCGCCACCGAGTCCAAGCAGCATAGCCTGGGACACTCAACAGCACAGCCCAGCCAACAATGTTATAAAAAACTTAG

GAGCCAGGGTTGTAGGGGCCATGGACTCCCTATCAGTCAATCAGGTCTTGTACCCTGAGGTTCATCTAGACAGCCCTATT

GTCACAAACAAACTAGTTGCCATCCTTGAGTACTCGGGTATCGACCATAACTATGTTCTTGAAGACCAGACCCTTGTCAA

GAATATTAGGTATAGACTGGGGTGCGGTTTTTCAAATCAAATGATCATCAATAACAGGGGGGTGGGTGAAACAGTCAATT

CTAAACTTAAAAGTTATCCCCATAATCGTCATATCATATACCCGGATTGCAATAAGGAGTTGTTTGGTATCAAAGATAGC

TGCATATCTAAGAAGCTCTCGGAGCTATTCAAGAAGGGTAATTCCTTGTACTCTAAGATAAGTCACCAGGTACTGGATTG

TCTTAAGAGAGTCAATGGGAAATTAGGCCTGGGCACAGATCTTACCCATGGTCTGAAGGAAGGTATCCTTGACCTAGGGT

TGCACATGCATAGCTCTCAATGGTTCGAGACTTTCCTGTTCTGGTTCACTATCAAGACAGAGATGAGATCAATGATCAAA

GAACAGTCCCATATATGCCACAAGAGGAGGTATAACCCGATTTTTGTATCGGGGGATGCATTCGAGGTGCTCGTATCACG

AGACCTCGTTGTGATAATTGATAAGAATACCCAGTATGTCTTCTACTTAACATTTGAGCTGGTCCTTATGTATTGTGATG

TCATAGAGGGCAGACTTATGACGGAGACAGCCATGGCCATAGACCAGAGATATTCAGAACTTCTAAGCCGGGTTAGATAC

TTGTGGGATCTTATTGATGGGTTCTTCCCAACACTAGGCAACACCACATACCAAATTGTTGCTCTGCTTGAACCATTATC

GTTGGCTTATCTTCAACTTCAGGATGTCACGCTGGAGTTAAGAGGTGCTTTTTTAGACCATTGCTTCAAAGAACTCTATG

AGATACTGGAGCATTGCGGCATTGACACAGAAGGCACCTACAATTCCATCACTGAAGGACTGGATTACGTATTCATCACC

CACGATATACATCTAACTGGGGAGATTTTTTCATTTTTCCGGAGTTTCGGACACCCCCGTCTAGAAGCAGTCACCGCTGC

AGAAAATGTCAGGAAACATATGAACCAACCGAAGGTAATCAGTTATGAGACTATGATGAAAGGACATGCGGTATTTTGCG

GGATAATCATAAATGGTTTTAGGGACCGACATGGCGGCAGCTGGCCCCCTGTTGCATTGCCAGAACATGCTTCTGCTGCG

ATCCGGAATGCGCAGGCATCAGGTGAGGGACTGACCCATGACCTGTGTATAGACAACTGGAAATCCTTTGTTGGATTCAA

ATTTGGCTGCTTTATGCCACTCAGCCTAGATAGTGATTTGACCATGTATCTCAAAGACAAGGCATTGGCTGCACTGAAGA

ATGAGTGGGATTCAGTTTACCCGAAAGAATACCTCCGTTATAACCCACCTAGAGGGACAGAGTCAAGGCGACTGGTGGAG

GTGTTCCTGAATGACTCCAGCTTTGATCCTTATAACATGATAATGTACGTGGTGAATGGCTCCTACCTTAAAGACCCTGA

GTTTAATCTCTCATACAGCCTAAAAGAGAAGGAGATAAAGGAGACAGGGCGGTTGTTTGCCAAAATGACTTATAAGATGC

GGGCCTGTCAGGTAATTGCTGAAAATCTGATATCGAATGGTGTTGGGAAGTATTTCCGAGACAATGGGATGGCAAAGGAC

GAGCATGACCTAACAAAAGCCCTTCACACCCTGGCAGTCTCAGGTGTTCCCAAGAACAACAAAGATAACCACCGAGGTGG

GCCTCCCAGAAGGACAACAAGCCGAGAGGTGAGATCAAGCCAAGACACCAAAACACAAAATAGAGACAAGGTCCAAGGAG

GACCTATGTACAACTATTTGCGATGCCAACCAATCAGCCCTGATCAGGGTGAGTCATACGAGACTGTTAGTGCATTCATC

ACCGCTGACCTTAAGAAGTATTGCCTGAATTGGAGATACGAGACAATCAGCATATTTGCACAGAGGCTGAATGAAATATA

TGGACTACCATCCTTCTTTCAATGGTTACACAGGATATTGGAAAAATCCGTACTCTACGTCAGTGACCCACATTGCCCTC

CCGATCTAGATAATCATATCCCTCTGGACAGTGTCCCTAATGCCCAAATATTCATTAAGTACCCAATGGGCGGAATAGAA

GGTTATTGCCAAAAACTATGGACAATCAGTACTATACCATACTTGTATCTGGCAGCCTATGAGAGCGGAGTAAGAATCGC

CTCACTGGTGCAGGGTGACAATCAGACAATCGCAGTGACAAAAAGAGTTCCAAGCTCTTGGCCTTATTCACTAAAAAAGA

AGGAGGCATCCAGAGCCGCTCAAAATTACTTCGTAGTCTTAAGGCAAAGATTGCACGATGTAGGTCATCACTTAAAGGCT

AATGAAACCATAGTATCTTCTCACTTTTTTGTATATTCCAAAGGGATTTATTATGACGGCCTGCTAGTCTCACAATCACT

GAAGAGCATCGCCAGATGTGTCTTCTGGTCCGAGACTATAGTGGATGAAACCAGAGCGGCTTGCAGCAATATTGCAACAA

CTGTCGCTAAGAGTATAGAGAGGGGTTATGATAGGTACCTTGCATATTCTTTGAATATCCTCAAGATTTTCCAACAGATC

CTTATATCCCTTAACTTCACTATTAACACAACAATGACTCAGGATGTCGTGGCACCGATCATCGAGAACGGTGATTTGCT

GATAAGGATGGCACTCTTGCCAGCACCCATCGGAGGTCTAAATTATCTTAACATGAGCAGGTTATTTGTGAGAAATATCG

GTGACCCGGTCACCTCCTCTATAGCCGACCTGAAGAGGATGATAGATGCCGGGCTAATGCCAGAAGAAACATTGCATCAA

GTGATGACCCAGACCCCGGGAGAATCATCCTACCTCGATTGGGCAAGTGACCCTTATTCTGCCAACCTACCCTGCGTACA

GAGTATAACTCGCCTTCTCAAAAACATCACTGCACGATATATTTTAATCAGCAGCCCAAACCCGATGCTGAGGGGATTGT

TTCATGAGGGGAGTAGAGATGAAGACGAGGAGCTTGCAAGTTTCCTAATGGATCGGCATATAATTGTTCCAAGAGCTGCA

CATGAAATCTTAGACCATAGTATAACCGGGGCAAGAGAAGCTATAGCCGGGATGTTGGACACCACCAAAGGTCTGATTAG

AACAAGTATGAAACGGGGTGGCCTCACCCCTCGAGTCTTAGCCCGCCTTTCCAATTATGATTATGAACAATTCCGATCCG

GGATAACACTATTGACAAAGAAAGGGCAGTGTTATCTCATTGACAAGGATTCGTGCTCGGTGCAGCTCGCTATAGCTCTG

AGGAGCCACATGTGGGCTAGGTTGGCTCGCGGGAGACCAATCTATGGGTTGGAGGTGCCCGATATATTAGAATCGATGAA

CGGCTACCTTATCAAACGCCACGAGTCTTGTGCTATCTGTGAAACGGGCTCAAGTCACTACGGATGGTTTTTCGTCCCTG

CAGGGTGCCAGCTTGACGATGTATCTAGAGAGACTTCGGCTCTTCGTGTACCTTATGTCGGATCAACAACTGAGGAAAGG

ACAGACATGAAACTTGCATTTGTTAGATCTCCAAGTAGATCCCTCAAATCAGCAGTTAGAATTGCAACAGTTTACTCATG

GGCCTACGGGGATGATGAGAAGTCATGGAGTGAAGCCTGGATGCTAGCTAGGCAGAGAGCCAATATCACCTTAGATGAAT

TGAGAATGATAACTCCAGTCTCTACATCCACCAACCTAGCCCACCGGTTGAGGGATCGGAGCACCCAGGTGAAATACTCG

GGGACATCCCTTGTGAGGGTTGCAAGATACACAACCATCTCCAATGATAACTTGTCATTTGTGATATCTGAGAAAAAAGT

GGATACCAACTTCATTTATCAGCAAGGGATGTTGCTTGGTCTTGGGATCCTTGAGAACCTCTTTAGGCTAGAGGCCACCA

CCGGAGTATCCAACACAGTGCTACACCTGCACGTGGAAACAGAATGTTGTGTTGTACCTATGGTGGATCACCCAAGGATA

CCGAGTCTCCGTAATATAAAAGTTACGGATGAGCTATGCACAAACCCTTTGATCTACGACAGGTCTCCCATCATAGAGCA

CGATGCAACCCGACTATACTCACAGAGCCACAGGAGACATTTGGTGGAGTTTGTTACCTGGTCCACAAGCCAGCTTTACC

ATATACTGGCTAAGTCTACAGCAATGTCCATGATTGAATTGATCACGAGATTCGAGAAAGATCACATGAATGAAATAGCC

GCCCTGATTGGCGATGACGACATCAACAGCTTCATCACAGAATTTTTGCTAGTGGAGCCTAGATTATTTATAGTTTACCT

TGGTCAGTGTGCTGCCATCAATTGGGCCTTTGATATACATTATCATCGGCCCTCGGGCAAGTACCAGATGGGGGAACTCC

TCTACTCTTTGCTCTCTCGGATGAGCAAAGGAGTATATAAGATCTTCACCAATGCTTTGAGCCACCCCAAAGTTTACAAG

AAATTTTGGCGAAGTGGTGTCATTGAGCCGATTCATGGCCCATCCCTGGATACACAGAATTTACATGTCACTGTCTGTGA

CATGATATATGGGTCCTATGTCACCTATTTGGATCTTTTGCTGAATGATGAGCTAGATGATTACCCGTATTTGCTCTGCG

AGAGTGATGAGGACGTGGTCACAGACAGGTTCGACAACATTCAAGCCAAACATCTCTGTGTATTAGCCGATGTATATTGC

AGCTCCAAGAGGTGTCCTTCAATAATCGGGATGTCTCCTATAGAAAAATGTACCATTCTCACACATTACATCAAGGGAGA

ATCAGTACAATCCCCGTCTGGGACCTCATGGAACACTGATCCCCTTGTAGTAGATCATTACTCATGCTCTCTGACCTACC

TTCGCCGCGGTTCCATCAAACAAATCAGGTTGAGGGTGGATCCTGGGTTTGTATTCGAGGCGTTGACAGACGTCGACCTT

AAACAACCTCGCAAAGCTAAGTTAGATGTATCGGTCGTTGGGTTGACTGATTTCTCTCCCCCTTGCGACAACGTCGGTGA

TTTTCTAGGGACTATCAACACATTGAGGCACAATCTGCCTGTCACCGGAACCGGGGTCTCGAACTATGAAGTCCACGCTT

ATCGTAGAATTGGCCTAAATTCATCAGCATGTTACAAAGCCGTAGAGATCTCCACGTTAATCAAGCCATCCCTAGAAGTC

GGAGAGCATGGATTGTTCTTAGGAGAAGGTTCCGGTTCAATGCTGGCTGCGTACAAGGAAGTTCTTAAATTAGCAAATTG

TTATTACAACAGCGGGGTAACAACGGAGGGCAGAGCCGGACAGAGGGAAATATCTCCCTATCCTTCAGAGATGAGCCTAG

TAGAGAATCAAATGGGGATAGAGCGGAGTGTTAAAGTGCTGTTCAACGGCAAACCTGAAGTAACTTGGGTGGGGACCACC

GATTGCTACAAGTACATAATCAGTAACATCCAAACCTCTAGTCTGGGTTTCATACATTCAGATATTGAGACGCTCCCAAC

CAAAGATGCTGTTGAGAAATTAGAAGAATTTGCCTCTATCCTATCCTTATCCTTGATTTTGGGGAAAATCGGCTCTATTA

CAGTTGTCAAAATTATGCCCATTAGCGGGGATTTTACCCAAGGCTTTATAGCCTATGCCATCCAATATTTTAGGGAGAGC

CTGCTTGCCTATCCGAGATATAGTAACTTCATCTCGACTGAGTGTTACCTTATTATGATAGGATTAAAGGCCAATCGACT

GATAAACCCAGAAGCCATTAAGCAAAGCATAATCAGAGCGGGGGTCAGGACTGCACCAGGACTTGTGAGCCATATATTAT

CAGAGAAACAGAAAGGTTGTATTCAATCTTTCTTGGGTGATCCTTATATCCAAGGAGACTTCAATAAGCACCTTAAATCT

CTGACCCCTATTGAGAAAATCCTGGTAAATTGTGGTCTCTCGATCAATGGCACAAAAATCTGTAGAGATCTAATCCACCA

TGATATCGCCTCCGGTCCAGACGGTCTTATGAGCTCTACAATTATTTTATATAGGGAATTGGCTCATTTCAAAGATAATA

TAAGGAGTCAGCACGGAATGTTCCACCCCTACCCAGTACTGGCCAACAGCCGGCAACGTGAATTAATCCTTCGAATAGCC

AAGAAATTCTGGGGGTATGTCTTGCTATATTCTGATGACCCGGCACTAATCAGACAAACAATCAAGAACTTGAAACGGAA

TCATCTAACCTTTGACTTACATAGTAACCCGTTTATTAAAGGGCTATCCAAAGCTGAGAAACTGCTGGTGCGGACGAGTC

CACTAAGAAGGGAATGGTTGTTCGCTCTCGATACGAAAGAAGTGAAAGAGTGGTTCAAATTGGTAGGTTACAGTGCACTC

ATCAGAGGCTAGTTAGCTATACATCTGCCCCCTTCTCCTCCGCCATGAGACCCCACTGACGATCCAGAAGATTAAAGAAA

ACTACATATTGGATAAGTATCTATTCCCAGCTTTGTCTGGT

>FJ905304.1/goat/China/2007-08

ACCAAACAAAGTTGGGTAAGGATAGATCTTATAATAACTATGGACTGGCAAACTTAGGAGTAAAGATCCTACTGTCGGGG

GGAGGAGGAGGAGCAAGATCTCTGATTATGGCGACTCTCCTTAAAAGCTTAGCATTGTTCAAAAGGAACAAAGACAAAGC

GCCGACGGCATCGGGTTCAGGAGGGGCCATCCGAGGGATTAAGAATGTTATCATAGTCCCGATTCCCGGAGACTCGTCCA

TCACTACCCGTTCAAGACTGCTCGACAGGCTTGTCAGGTTGGCCGGAGATCCTGACATCAACGGGTCAAAGCTGACCGGC

GTGATGATCAGCATGCTATCCTTGTTCGTAGAGTCACCCGGGCAATTGATACAGCGAATCACAGATGATCCAGATGTTAG

TATCCGCCTTGTTGAGGTAGTTCAAAGTACTAGGTCCCAGTCCGGGTTGACCTTTGCATCACGTGGTGCTGATTTGGATA

ACGAGGCAGACATGTATTTTTCAACTGAGGGGCCCTCGAGTGGAGGTAAGAAAAGGATCAACTGGTTTGAGAACAGAGAA

ATAATAGACATAGAGGTGCAGGATCCAGAAGAGTTCAATATGTTGTTAGCCTCCATACTAGCACAAGTCTGGATCCTTCT

GGCCAAGGCTGTTACGGCACCGGATACGGCAGCTGACTCAGAACTGAGAAGGTGGGTTAAATACACACAACAAAGGAGAG

TGATTGGGGAATTTCGCCTTGATAAAGGGTGGCTGGACGCAGTCCGCAACAGAATTGCAGAAGATCTATCACTCCGGCGG

TTCATGGTATCTCTCATACTTGACATCAAGAGGACCCCCGGCAACAAGCCAAGGATTGCAGAAATGATCTGCGACATTGA

CAACTATATTGTCGAAGCAGGACTCGCCAGTTTCATCCTTACTATCAAATTTGGTATAGAAACCATGTATCCTGCATTAG

GGCTCCACGAGTTTGCCGGGGAATTGTCCACTATTGAGTCCTTGATGAACTTGTATCAACAGCTAGGAGAGGTTGCACCC

TACATGGTAATTCTAGAGAACTCAATTCAGAACAAGTTTAGTGCAGGAGCTTATCCCCTCCTCTGGAGCTATGCGATGGG

TGTCGGAGTTGAGCTGGAGAACTCAATGGGGGGCTTGAACTTTGGCAGGTCATATTTTGACCCTGCCTATTTTCGTCTCG

GACAGGAGATGGTCAGAAGATCTGCAGGAAAGGTCAGCTCTGTGATTGCGGCTGAGCTCGGCATCACAGCAGAGGAAGCT

AAACTAGTCTCGGAAATAGCCTCGCAGGCTGGGGACGAAAGAACCGCTAGAGGGACTGGGCCTCGACAGGCGCAGGTCTC

CTTCCTCCAGCACCAAACAGGAGGGGGAGAGTCGTCCGCACCAGCGACCAGAGAAGGGGTCAAAGCTGCGATCCCAAACG

GATCCGAAGAAAGGGACAGAAAGCAAACACGCCCAGGAAGGCCCAGAGGAGAGACCCCCGGCCAACTGCTCCTGGAAATC

ATGCCAGAGGATGAGGTTTCGCGAGAATCTGGTCAAAACCCTCGTGAGGCTCAAAGATCGGCCGAGGCACTCTTCAGGCT

GCAGGCCATGGCCAAGATTCTGGAGGACCAGGAGGAGGGAGAAGACAACAATCAGGTCTACAACGACAAGGATCTCCTCG

GCTAAGCAGACGCACCCTCTGTCGAAATCAGTGACGAGACATCTCCTACCAGTATTATAAAAAACTTAGGACCCAGGTCC

AAGCAAGCACACATCGACACTCCAACCAGTCGAGCGGAGGCCACCGATGGCAGAAGAACAAGCATACCATGTCAACAAGG

GGCTGGAATGTATCAAGTCCCTCAAAGCCTCTCCCCCGGATCTATCCACCATCAGAGATACCATCGAGAGCTGGAGAGAG

GGGCTTAGCCCATCGGGCCGTGCAACACCGAACCCTGATATGTCCGAGGGAGATCATCAGAATATCAACCAACCATGCTC

ACCAGCAATCGGACCAAACAAAGTCTACTTGTCTCCTGAAGATAATCTCGGATTTAGAGAGATCACTAGCAACGACTATG

AGGCTGAGCTCGGAGGAGTCCAGGGAAAAGGATCCAACTCTCAAGTACAGCGTTACTATGTTTATAGCCACGGGGGTGAA

GAGATTGAAGGACTCGAGGATGCTGACTCTCTCGTGGTTCAAGCAGATCCTCCAGTTGCTAACATATTCAATGGAGGAGA

GGATGGATCTGACGACAGCGATGTGGACTCTGGCCCAGATGATCCCGGCAGAGATACTCTATATGACCGGGGATCTGTTG

CCAGCAATGATGTCGCTAGGTCCACAGATGTCGAAAAACTAGAAGGTGCTGATATTCAAGAAGTTCTCAACTCCCAGAAA

GGCAGAGGAGGAAGATTCCAAGGCGGGAAAACCTTACGAGTCCCGGAAATACCCGATGTCAAGCACTCCAGACCATCAGC

CCAATCAATTAAAAAGGGCACAGACGGGAACTCAGTCTCATCTGGAACGGTGACAGAGTGTTTATCGATAAGTGGTGCAA

CCCAAGCTGTGCCAGAGTCAAGATGGGAGTCATCAGAGCAAAATGCGTCTGTGGGGAGTGTCCTCAAGTCTGCGGGGAGT

GCAAAGACGATCCAGGGGTCGACACAAGAATCTGGTACCATAGCATCACTGACTCAGCCTAAAGAGAATGACTCCGAGTA

TGAGTATGAGGATGACCTATTTACAGAGATTCAGGACATCCGTGCAAGCATTGCCAAGATCCATGATGACAACAAGGCTA

TCCTCTCAAAGCTTGATTCTATACTGTTATTGAAAGGAGAAGTCGACACTATCAAGAAACAAATCAGCAAGCAGAATATA

AGTATATCCACCATTGAGGGCCATCTCTCCAGTATAATGATAGCCATCCCGGGCTTTGGGAAGGATATCAAGGACCCAAC

ATCCGAGGTTGAGTTGAACCCAGATTTAAGACCTATAATAAGCCGTGACTCTGGCAGAGCTCTCGCGGAGGTCCTCAAGA

AACCCGCTGTGGATAGGTCTCAGAAAATTGGAACCAAAGCCAACTCCAGCTCAAAGGGTCAGCTTCTTAAGGATCTCCAG

CTAAAACCTGTCGACAAGCAGGCAAGCTCTGCAATCGGGTTTGTCCCATCCGACCATGAATCATCCAGAAATGTCATCCG

CTCCATAATCAAGTCGAGCAAGCTCAACATTGATCACAAGGACTATCTTCTAGATTTACTGAATGATGTGAAAGGCTCCA

AGGATCTTAAGGAATTCCACAAGATGCTAACAGCAATTCTCGCCAAGCACCCGTAACACATCCTCCAGTCACCGCCCCAT

ACTCGGCTGAAAACATCCTTTCAATCAGGCTATTACAAAAAACTTAGGAGCAAGGGCAACTGAGCTTCGCAGACAAGATG

ACCGAGATCTACGACTTCGATAAATCAGCATGGGATGTCAAAGGGTCAATTGCCCGCATAGAACCCACCACCTACCACGA

CGGCCGACTGATACCCCAGGTAAGGGTCATCGATCCTGGTCTGGGAGACAGAAAAGATGAGTGCTTTATGTACCTGTTTC

TCCTGGGAGTGATTGAGGATAACGACCCCCTGTCTCCCCCCGTCGGAAGAACCTTTGGCTCTTTACCTCTAGGGGTCGGT

AGGTCAACTGCTAAGCCAGAGGAACTACTAAGGGAGGCCACAGAACTAGACATAGTGGTGAGGCGCACTGCAGGAGTAAA

TGAGAAACTGGTATTTTACAACAACACTCCGCTGTCCTTGTTAACGCCCTGGAAGAAAGTTCTGACAACCGGAAGTGTGT

TTAGCGCTAACCAGGTTTGCAATGCAGTCAACCTAGTCCCGCTTGATACTCCCCAGAGGTTCAGGGTTGTGTACATGAGC

ATAACTAGATTATCAGACAATGGTTACTATAGTGTGCCCAGAAGAATGCTGGAGTTCCGCTCAGCCAATGCAGTAGCCTT

CAACATCTTGGTTACACTAAAAATTGAAAATGGCACAAACCCTAGAAGATACATAGTCGGCTCATGGGAGAATTCAGAGG

TCACATTTATGGTACACGTGGGCAACTTTAGAAGAAAGAAGAACGAAGTATACTCTGCTGATTACTGCAAAATGAAGATT

GAAAAGATGGGTTTAGTTTTTGCCCTGGGAGGAATAGGTGGAACAAGTCTCCATATTAGAAGCACCGGGAAAATGAGCAA

AACCCTCCATGCACAGCTGGGGTTCAAGAAAATCTTATGTTACCCCCTAATGGATGTTAATGAGGATCTTAACCGATATC

TCTGGCGGGCAGAGTGCCGAATAGTCAAAATCCAAGCCGTCCTACAGCCATCAGTGCCCCAAGAATTCCGTGTCTACGAT

GATGTCATCATCAACGATGATCAAGGCTTGTTCAAGATCTTGTAGTTCATTTGCAAAGGAACGCACCCACCCCCAGGAAG

GGGCCCCA-CCCCCCCCG-CCACAAAACACACTCCAAGGCTCCCCGACACAGCGCCAACACAGCCACCCCCCCGACTGGA

CGAGGCCGGCC-CCCCAACAAACCCGGTCTAGGCGGGGGGCCCCCCCCCCTGACCCGCCGGGCGCCGACCGAGCAGGCCG

GCGAGAAGCCCCCAAGAGGCCCCGACAAAGACCGAGCACCCCATCCCCCC-AAAACGTCTCTCC-AACACGCTCGAGGCC

AGGCGCCGGGCACCCCCAGATCCGCCACCGGCGAGACAGAACCCCAGGGGGGGACCTCCCAAACCAAAGGAACCACCGGG

GAAGGCTCCGCCGAACCAGCACAGACCCGACCCAAGCAAGGCGGACCCCCAACACGAAACCGCCCAGCACACCCCACTCA

TCATCAAACAAAACTTAGGGGCCAAGCCCATACGACAGGCCCCAGTCCGCCAGGCGGAGGGACGCAGAAAGGAAGGAGAC

ATGACCGCCCCAGAAGGGGGACACCCAACCACCGAAACACCCAGCCCCAGGGAAGGAG--CCCCCCCCCACACGGCGAAG

GAACGGGTGAGAGACCCCGAGGAGAACACGACCGGGAGCCAGGAGCCGGCGGCGCCGGCAGCAAACAAGCCGGGCAAACA

CCCGACCCCACCAGAGGAGCCGAGGAGAGCCCGACCCCGCACGAACCAAACGGCAGGGAGCCCAACCCCACCACAAAAAG

AGGCGAGGCCCC-CCT-----CCCCCCCGCACACAACCCGCAGACAGACCGAGGGAGGACCACCCCCCAACCAGACAGGC

GGCCCCCCCGCAGCCAAGCTGCAGCTCCTCGGAACAGGACACCCCCGCACACTCCGAACACCCCCCCTCCCCCCGCCACC

GGGACAGGGGCTCCGCCACCACAGACCAAAGCCACAGAGAGCCAGCCAGAGGTCCCCCCCAGTCTCGCCCGGTGCCCCCC

CGAAAAGGCCCCGGCAAGGCCCCCCAACCCGACGTCCCCCGCACGCACCGAAAGAAGGGCCCGCCCGAGAACGTGCGGCA

AATCCATGCGCAAACATCATGACACGGGTCGCAATCTTGACATTTCTGTTTCTTCTCCCAAATGTTGTTGCGTGCCAGAT

TCACTGGGGCAATCTATCCAAGATCGGGATTGTAGGAACAGGGAGTGCCAGCTACAAGGTGATGACTAGGCCAAGCCACC

AAACTCTGGTTATAAAGTTAATGCCAAATATAACAGCCATCAACAATTGTACAAAGTCAGAGATTGCAGAGTACAAGAGA

TTGCTGATCACAGTGTTAAAACCTGTAGAGGATGCTCTGTCGGTGATAACCAAGAATGTAAGACCAATTCAAACCCTGAC

ACCTGGGCGTAGAACCCGTCGTTTTGCTGGAGCTGTTCTGGCCGGAGTAGCACTTGGAGTTGCGACAGCCGCTCAGATAA

CTGCAGGAGTCGCTCTTCATCAATCATTGATGAACTCTCAAGCAATTGAGAGTTTAAAAACCAGTCTTGAGAAGTCGAAT

CAGGCAATAGAAGAAATCAGACTTGCAAATAAGGAGACCATACTGGCAGTACAGGGCGTCCAGGATTATATCAACAATGA

GCTTGTCCCTTCTGTTCATAGAATGTCATGCGAGCTGGTAGGCCACAAGCTCGGCCTCAAGCTCCTTAGGTACTACACCG

AGATCCTGTCCATATTCGGGCCCAGCCTTCGAGACCCGATAGCTGCCGAAATATCGATCCAGGCACTCAGTTATGCATTA

GGCGGAGACATTAATAAAATCCTGGACAAGCTTGGGTATAGCGGTGGGGATTTCCTTGCCATCCTAGAGAGCAAGGGAAT

AAAGGCCCGGGTTACATATGTGGACACAAGAGATTACTTTATAATCCTTAGCATCGCCTACCCAACCTTATCTGAGATCA

AGGGAGTGATAGTTCACAAGATAGAAGCTATAACATACAACATTGGGGCACAGGAGTGGTATACTACTATCCCTAAATAT

GTAGCCACTCAGGGGTATCTGATATCGAACTTTGATGAGACGTCATGCGTATTCACTCCAGAGGGGACAGTTTGCAGCCA

GAATGCGTTGTACCCAATGAGCCCATTGCTTCAGGAATGTTTCAGGGGATCAACAAAATCGTGCGCCAGAACCCTAGTTT

CAGGGACCATAAGTAATAGATTTATCCTATCAAAAGGGAACCTGATTGCAAATTGTGCGTCAGTTTTGTGCAAATGTTAC

ACAACGGAGACAGTCATCAGCCAAGATCCTGACAAACTACTAACTGTTGTAGCATCCGACAAGTGTCCTGTAGTTGAGGT

GGATGGAGTGACAATACAGGTCGGCAGTCGAGAGTATCCGGATTCTGTATACTTACACAAAATAGACTTAGGTCCAGCCA

TCTCCCTAGAAAAACTGGATGTAGGCACCAATTTAGGCAATGCAGTCACAAGACTGGAGAATGCAAAGGAGCTCCTAGAT

GCATCAGACCAAATACTGAAGACTGTCAAAGGGGCACCTTTAGGCGGGAACATGTACATAGCACTGGCAGCTTGCATTGG

AGTATCCCTAGGGCTTGTCACATTAATATGCTGCTGTAAGGGGAGGTGTAAGAACAAGGAAATCCCTATCTCCAAAATCA

ACCCAGGGCTCAAACCCGATCTGACCGGGACCTCAAAGTCGTACGTGAGATCACTGTAGCCAGAATCACCTGAATCATCT

GGCATCACACACATACATGTACGACACAAGCAGTCTGAGGACGCAGGAAACCCAGCCTCCGATCACCTACCTGACCCTAC

TCCACGCTCCACTACACATTAGTCATCAAACAAAACTTAGGACGAAAGGTCAATCACCATGTCCGCACAAAGGGAAAGGA

TCAATGCCTTCTACAAAGACAATCCTCACAATAAGAACCATAGGGTGATCCTGGATAGAGAACGCTTGGTCATTGAAAGA

CCCTACATCTTGCTTGGAGTCCTGCTGGTAATGTTCCTGAGTCTAATCGGACTGCTGGCCATTGCAGGGATCAGGCTTCA

CCGGGCCACCGTTGGAACTTCAGAGATCCAGAGTCGGCTGAATACCAATATTAAGTTGGCCGAATCTATTGATCACCAGA

CTAAGGATGTCTTAACTCCCCTTTTTAAAATCATTGGCGATGAAGTCGGCATCAGAATTCCACAGAAATTCAGTGATCTT

GTCAAGTTCATCTCCGATAAGATTAAATTCCTCAACCCTGATAGAGAGTATGATTTCAGGGATCTCCGGTGGTGCATGAA

TCCCCCCGAGAGAGTCAAAATTAATTTTGATCAGTTTTGTGAGTACAAAGCTGCGGTTAAGTCAATTGAACATATATTTG

AGTCACCACTCAACAAGTCAAAAAAGCTGCAATCATTGACTCTCGGGCCCGGAACAGGCTGTCTAGGCAGGACAGTAACA

AAAGCTCATTTCTCAGAACTTACACTGACCTTAATGGACCTGGATCTAGAGATGAAGCACAACGTGTCCTCAGTGTTTAC

CGTAGTTGAAGAGGGGTTATTCGGAAGAACATATACCGTCTGGAGATCCGATGCCAGGGATCCGAGCACCGATCTAGGTA

TCGGCCATTTTTTAAGAGTCTTCGAGATTGGACTGATAAGAGATCTCGGGCTGGGTCCCCCTGTTTTTCATATGACCAAC

TATCTCACAGTGAACATGAGTGATGACTATCGGAGATGTCTTTTAGCGGTAGGGGAGTTGAAGTTGACAGCCCTATGCAC

CTCATCTGAGACTGTGACACTGAGTGAGAGAGGAGCTCCAAAGAGGGAGCCTCTTGTGGTTGTGATACTTAATCTGGCTG

GACCCACTCTAGGGGGCGAACTATACAGTGTCTTGCCTACCTCCGATCTCATGGTGGAGAAACTCTATTTATCTTCACAT

AGAGGGATCATCAAGGATGACGAGGCCAATTGGGTAGTGCCGTCTACCGATGTTCGTGATCTTCAGAACAAAGGTGAATG

TCTGGTGGAAGCATGCAAGACTCGACCTCCTTCATTTTGCAATGGCACAGGATCAGGCCCGTGGTCAGAGGGGAGAATCC

CTGCTTACGGGGTGATCAGGGTCAGTCTTGACTTAGCTAGTGACCCGGATGTAGTTATCACTTCAGTGTTTGGCCCACTG

ATTCCTCACCTATCCGGCATGGATCTTTACAACAACCCGTTTTCAAGAGCTATATGGTTGGCTGTACCACCTTATGAGCA

GTCATTTCTAGGAATGATAAATACAATTGGATTCCCTAACAGAGCAGAGGTTATGCCGCACATTTTGACCACAGAGATCA

GAGGCCCTCGGGGTCGTTGCCATGTTCCCATAGAATTGTCCCGCAGGGTTGATGACGATATCAAGATCGGGTCCAACATG

GTCATATTGCCGACGATGGACCTGAGGTATATTACAGCCACTTATGATGTTTCCAGGAGCGAGCATGCAATCGTGTACTA

TATCTATGACACAGGTCGCTCATCATCTTACTTCTACCCAGTTCGACTGAATTTCAAAGGCAATCCTCTCTCTCTGAGGA

TAGAGTGTTTCCCTTGGCGTCATAAGGTGTGGTGCTACCATGATTGTCTTATATACAACACCATAACAGATGAAGAGGTC

CATACGAGAGGGCTGACCGGTATAGAGGTAACATGCAATCCAGTTTGAGCAAAGCTACGACCGTCACTCAAGCAGTCCCT

TGAGTCGCCACCGAGTCCAAGCAGCATAGCCTGGGACACTCAACAGCACAGCCCAGCCAACAATGTTATAAAAAACTTAG

GAGCCAGGGTTGTAGGGGCCATGGACTCCCTATCAGTCAATCAGGTCTTGTACCCTGAGGTTCATCTAGACAGCCCTATT

GTCACAAACAAACTAGTTGCCATCCTTGAGTACTCGGGTATCGACCATAACTATGTTCTTGAAGACCAGACCCTTGTCAA

GAATATTAGGTATAGACTGGGGTGCGGTTTTTCAAATCAAATGATCATCAATAACAGGGGGGTGGGTGAAACAGTCAATT

CTAAACTTAAAAGTTATCCCCATAATCGTCATATCATATACCCGGATTGCAATAAGGAGTTGTTTGGTATCAAAGATAGC

TGCATATCTAAGAAGCTCTCGGAGCTATTCAAGAAGGGTAATTCCTTGTACTCTAAGATAAGTCACCAGGTACTGGATTG

TCTTAAGAGAGTCAATGGGAAATTAGGCCTGGGCACAGATCTTACCCATGGTCTGAAGGAAGGTATCCTTGACCTAGGGT

TGCACATGCATAGCTCTCAATGGTTCGAGACTTTCCTGTTCTGGTTCACTATCAAGACAGAGATGAGATCAATGATCAAA

GAACAGTCCCATATATGCCACAAGAGGAGGTATAACCCGATTTTTGTATCGGGGGATGCATTCGAGGTGCTCGTATCACG

AGACCTCGTTGTGATAATTGATAAGAATACCCAGTATGTCTTCTACTTAACATTTGAGCTGGTCCTTATGTATTGTGATG

TCATAGAGGGCAGACTTATGACGGAGACAGCCATGGCCATAGACCAGAGATATTCAGAACTTCTAAGCCGGGTTAGATAC

TTGTGGGATCTTATTGATGGGTTCTTCCCAACACTAGGCAACACCACATACCAAATTGTTGCTCTGCTTGAACCATTATC

GTTGGCTTATCTTCAACTTCAGGATGTCACGCTGGAGTTAAGAGGTGCTTTTTTAGACCATTGCTTCAAAGAACTCTATG

AGATACTGGAGCATTGCGGCATTGACACAGAAGGCACCTACAATTCCATCACTGAAGGACTGGATTACGTATTCATCACC

CACGATATACATCTAACTGGGGAGATTTTTTCATTTTTCCGGAGTTTCGGACACCCCCGTCTAGAAGCAGTCACCGCTGC

AGAAAATGTCAGGAAACATATGAACCAACCGAAGGTAATCAGTTATGAGACTATGATGAAAGGACATGCGGTATTTTGCG

GGATAATCATAAATGGTTTTAGGGACCGACATGGCGGCAGCTGGCCCCCTGTTGCATTGCCAGAACATGCTTCTGCTGCG

ATCCGGAATGCGCAGGCATCAGGTGAGGGACTGACCCATGACCTGTGTATAGACAACTGGAAATCCTTTGTTGGATTCAA

ATTTGGCTGCTTTATGCCACTCAGCCTAGATAGTGATTTGACCATGTATCTCAAAGACAAGGCATTGGCTGCACTGAAGA

ATGAGTGGGATTCAGTTTACCCGAAAGAATACCTCCGTTATAACCCACCTAGAGGGACAGAGTCAAGGCGACTGGTGGAG

GTGTTCCTGAATGACTCCAGCTTTGATCCTTATAACATGATAATGTACGTGGTGAATGGCTCCTACCTTAAAGACCCTGA

GTTTAATCTCTCATACAGCCTAAAAGAGAAGGAGATAAAGGAGACAGGGCGGTTGTTTGCCAAAATGACTTATAAGATGC

GGGCCTGTCAGGTAATTGCTGAAAATCTGATATCGAATGGTGTTGGGAAGTATTTCCGAGACAATGGGATGGCAAAGGAC

GAGCATGACCTAACAAAAGCCCTTCACACCCTGGCAGTCTCAGGTGTTCCCAAGAACAACAAAGATAACCACCGAGGTGG

GCCTCCCAGAAGGACAACAAGCCGAGAGGTGAGATCAAGCCAAGACACCAAAACACAAAATAGAGACAAGGTCCAAGGAG

GACCTATGTACAACTATTTGCGATGCCAACCAATCAGCCCTGATCAGGGTGAGTCATACGAGACTGTTAGTGCATTCATC

ACCGCTGACCTTAAGAAGTATTGCCTGAATTGGAGATACGAGACAATCAGCATATTTGCACAGAGGCTGAATGAAATATA

TGGACTACCATCCTTCTTTCAATGGTTACACAGGATATTGGAAAAATCCGTACTCTACGTCAGTGACCCACATTGCCCTC

CCGATCTAGATAATCATATCCCTCTGGACAGTGTCCCTAATGCCCAAATATTCATTAAGTACCCAATGGGCGGAATAGAA

GGTTATTGCCAAAAACTATGGACAATCAGTACTATACCATACTTGTATCTGGCAGCCTATGAGAGCGGAGTAAGAATCGC

CTCACTGGTGCAGGGTGACAATCAGACAATCGCAGTGACAAAAAGAGTTCCAAGCTCTTGGCCTTATTCACTAAAAAAGA

AGGAGGCATCCAGAGCCGCTCAAAATTACTTCGTAGTCTTAAGGCAAAGATTGCACGATGTAGGTCATCACTTAAAGGCT

AATGAAACCATAGTATCTTCTCACTTTTTTGTATATTCCAAAGGGATTTATTATGACGGCCTGCTAGTCTCACAATCACT

GAAGAGCATCGCCAGATGTGTCTTCTGGTCCGAGACTATAGTGGATGAAACCAGAGCGGCTTGCAGCAATATTGCAACAA

CTGTCGCTAAGAGTATAGAGAGGGGTTATGATAGGTACCTTGCATATTCTTTGAATATCCTCAAGATTTTCCAACAGATC

CTTATATCCCTTAACTTCACTATTAACACAACAATGACTCAGGATGTCGTGGCACCGATCATCGAGAACGGTGATTTGCT

GATAAGGATGGCACTCTTGCCAGCACCCATCGGAGGTCTAAATTATCTTAACATGAGCAGGTTATTTGTGAGAAATATCG

GTGACCCGGTCACCTCCTCTATAGCCGACCTGAAGAGGATGATAGATGCCGGGCTAATGCCAGAAGAAACATTGCATCAA

GTGATGACCCAGACCCCGGGAGAATCATCCTACCTCGATTGGGCAAGTGACCCTTATTCTGCCAACCTACCCTGCGTACA

GAGTATAACTCGCCTTCTCAAAAACATCACTGCACGATATATTTTAATCAGCAGCCCAAACCCGATGCTGAGGGGATTGT

TTCATGAGGGGAGTAGAGATGAAGACGAGGAGCTTGCAAGTTTCCTAATGGATCGGCATATAATTGTTCCAAGAGCTGCA

CATGAAATCTTAGACCATAGTATAACCGGGGCAAGAGAAGCTATAGCCGGGATGTTGGACACCACCAAAGGTCTGATTAG

AACAAGTATGAAACGGGGTGGCCTCACCCCTCGAGTCTTAGCCCGCCTTTCCAATTATGATTATGAACAATTCCGATCCG

GGATAACACTATTGACAAAGAAAGGGCAGTGTTATCTCATTGACAAGGATTCGTGCTCGGTGCAGCTCGCTATAGCTCTG

AGGAGCCACATGTGGGCTAGGTTGGCTCGCGGGAGACCAATCTATGGGTTGGAGGTGCCCGATATATTAGAATCGATGAA

CGGCTACCTTATCAAACGCCACGAGTCTTGTGCTATCTGTGAAACGGGCTCAAGTCACTACGGATGGTTTTTCGTCCCTG

CAGGGTGCCAGCTTGACGATGTATCTAGAGAGACTTCGGCTCTTCGTGTACCTTATGTCGGATCAACAACTGAGGAAAGG

ACAGACATGAAACTTGCATTTGTTAGATCTCCAAGTAGATCCCTCAAATCAGCAGTTAGAATTGCAACAGTTTACTCATG

GGCCTACGGGGATGATGAGAAGTCATGGAGTGAAGCCTGGATGCTAGCTAGGCAGAGAGCCAATATCACCTTAGATGAAT

TGAGAATGATAACTCCAGTCTCTACATCCACCAACCTAGCCCACCGGTTGAGGGATCGGAGCACCCAGGTGAAATACTCG

GGGACATCCCTTGTGAGGGTTGCAAGATACACAACCATCTCCAATGATAACTTGTCATTTGTGATATCTGAGAAAAAAGT

GGATACCAACTTCATTTATCAGCAAGGGATGTTGCTTGGTCTTGGGATCCTTGAGAACCTCTTTAGGCTAGAGGCCACCA

CCGGAGTATCCAACACAGTGCTACACCTGCACGTGGAAACAGAATGTTGTGTTGTACCTATGGTGGATCACCCAAGGATA

CCGAGTCTCCGTAATATAAAAGTTACGGATGAGCTATGCACAAATCCTTTGATCTACGACAGGTCTCCCATCATAGAGCA

CGATGCAACCCGACTATACTCACAGAGCCACAGGAGACATTTGGTGGAGTTTGTTACCTGGTCCACAAGCCAGCTTTACC

ATATACTGGCTAAGTCTACAGCAATGTCCATGATTGAATTGATCACGAGATTCGAGAAAGATCACATGAATGAAATAGCC

GCCCTGATTGGCGATGACGACATCAACAGCTTCATCACAGAATTTTTGCTAGTGGAGCCTAGATTATTTATAGTTTACCT

TGGTCAGTGTGCTGCCATCAATTGGGCCTTTGATATACATTATCATCGGCCCTCGGGCAAGTACCAGATGGGGGAACTCC

TCTACTCTTTGCTCTCTCGGATGAGCAAAGGAGTATATAAGATCTTCACCAATGCTTTGAGCCACCCCAAAGTTTACAAG

AAATTTTGGCGAAGTGGTGTCATTGAGCCGATTCATGGCCCATCCCTGGATACACAGAATTTACATGTCACTGTCTGTGA

CATGATATATGGGTCCTATGTCACCTATTTGGATCTTTTGCTGAATGATGAGCTAGATGATTACCCGTATTTGCTCTGCG

AGAGTGATGAGGACGTGGTCACAGACAGGTTCGACAACATTCAAGCCAAACATCTCTGTGTATTAGCCGATGTATATTGC

AGCTCCAAGAGGTGTCCTTCAATAATCGGGATGTCTCCTATAGAAAAATGTACCATTCTCACACATTACATCAAGGGAGA

ATCAGTACAATCCCCGTCTGGGACCTCATGGAACACTGATCCCCTTGTAGTAGATCATTACTCATGCTCTCTGACCTACC

TTCGCCGCGGTTCCATCAAACAAATCAGGTTGAGGGTGGATCCTGGGTTTGTATTCGAGGCGTTGACAGACGTCGACCTT

AAACAACCTCGCAAAGCTAAGTTAGATATATCGGTCGTTGGGTTGACTGATTTCTCTCCCCCTTGCGACAACGTCGGTGA

TTTTCTAGGGACTATCAACACATTGAGGCACAATCTGCCTGTCACCGGAACCGGGGTCTCGAACTATGAAGTCCACGCTT

ATCGTAGAATTGGCCTAAATTCATCAGCATGTTACAAAGCCGTAGAGATCTCCACGTTAATCAAGCCATCCCTAGAAGTC

GGAGAGCATGGATTGTTCTTAGGAGAAGGTTCCGGTTCAATGCTGGCTGCGTACAAGGAAGTTCTTAAATTAGCAAATTG

TTATTACAACAGCGGGGTAACAACGGAGGGCAGAGCCGGACAGAGGGAAATATCTCCCTATCCTTCAGAGATGAGCCTAG

TAGAGAATCAAATGGGGATAGAGCGGAGTGTTAAAGTGCTGTTCAACGGCAAACCTGAAGTAACTTGGGTGGGGACCACC

GATTGCTACAAGTACATAATCAGTAACATCCAAACCTCTAGTCTGGGTTTCATACATTCAGATATTGAGACGCTCCCAAC

CAAAGATGCTGTTGAGAAATTAGAAGAATTTGCCTCTATCCTATCCTTATCCTTGATTTTGGGGAAAATCGGCTCTATTA

CAGTTGTCAAAATTATGCCCATTAGCGGGGATTTTACCCAAGGCTTTATAGCCTATGCCATCCAATATTTTAGGGAGAGC

CTGCTTGCCTATCCGAGATATAGTAACTTCATCTCGACTGAGTGTTACCTTATTATGATAGGATTAAAGGCCAATCGACT

GATAAACCCAGAAGCCATTAAGCAAAGCATAATCAGAGCGGGGGTCAGGACTGCACCAGGACTTGTGAGCCATATATTAT

CAGAGAAACAGAAAGGTTGTATTCAATCTTTCTTGGGTGATCCTTATATCCAAGGAGACTTCAATAAGCACCTTAAATCT

CTGACCCCTATTGAGAAAATCCTGGTAAATTGTGGTCTCTCGATCAATGGCACAAAAATCTGTAGAGATCTAATCCACCA

TGATATCGCCTCCGGTCCAGACGGTCTTATGAGCTCTACAATTATTTTATATAGGGAATTGGCTCATTTCAAAGATAATA

TAAGGAGTCAGCACGGAATGTTCCACCCCTACCCAGTACTGGCCAACAGCCGGCAACGTGAATTAATCCTTCGAATAGCC

AAGAAATTCTGGGGGTATGTCTTGCTATATTCTGATGACCCGGCACTAATCAGACAAACAATCAAGAACTTGAAACGGAA

TCATCTAACCTTTGACTTACATAGTAACCCGTTTATTAAAGGGCTATCCAAAGCTGAGAAACTGCTGGTGCGGACGAGTC

CACTAAGAAGGGAATGGTTGTTCACTCTCGATACGAAAGAAGTGAAAGAGTGGTTCAAATTGGTAGGTTACAGTGCACTC

ATCAGAGGCTAGTTAGCTATACATCTGCCCCCTTCTCCTCCGCCATGAGACCCCACTGACGATCCAGAAGATTAAAGAAA

ACTACATATTGGATAAGTATCTATTCCCAGCTTTGTCTGGT

>JX217850.1/bharal/Tibet/2008-01

ACCAAACAAAGTTGGGTAAGGATAGATCTTATAATAACTATGGACTGGCAAACTTAGGAGTAAAGATCCTACTGTCGGGG

GGAGGAGGAGGAGCAAGATCTCTGATTATGGCGACTCTCCTTAAAAGCTTAGCATTGTTCAAAAGGAACAAAGACAAAGC

GCCGACGGCATCGGGTTCAGGAGGGGCCATCCGAGGGATTAAGAATGTTATCATAGTCCCGATTCCCGGAGACTCGTCCA

TCACTACCCGTTCAAGACTGCTCGACAGGCTTGTCAGGTTGGCCGGAGATCCTGACATCAACGGGTCAAAGCTGACCGGC

GTGATGATCAGCATGCTATCCTTGTTCGTAGAGTCACCCGGGCAATTGATACAGCGAATCACAGATGATCCAGATGTTAG

TATCCGCCTTGTTGAGGTAGTTCAAAGTACTAGGTCCCAGTCCGGGTTGACCTTTGCATCACGTGGTGCTGATTTGGATA

ACGAGGCAGACATGTATTTTTCAACTGAGGGGCCCTCGAGTGGAGGTAAGAAAAGGATCAACTGGTTTGAGAACAGAGAA

ATAATAGACATAGAGGTGCAGGATCCAGAAGAGTTCAATATGTTGTTAGCCTCCATACTAGCACAAGTCTGGATCCTTCT

GGCCAAGGCTGTTACGGCACCGGATACGGCAGCTGACTCAGAACTGAGAAGGTGGGTTAAATACACACAACAAAGGAGAG

TGATTGGGGAATTTCGCCTTGACAAAGGGTGGCTGGACGCAGTCCGCAACAGAATTGCAGAAGATCTATCACTCCGGCGG

TTCATGGTATCTCTCATACTTGACATCAAGAGGACCCCCGGCAACAAGCCAAGGATTGCAGAAATGATCTGCGACATTGA

CAACTATATTGTCGAAGCAGGACTCGCCAGTTTCATCCTTACTATCAAATTTGGTATAGAAACCATGTATCCTGCATTAG

GGCTCCACGAGTTTGCCGGGGAATTGTCCACTATTGAGTCCTTGATGAACTTGTATCAACAGCTAGGAGAGGTTGCACCC

TACATGGTAATTCTAGAGAACTCAATTCAGAACAAGTTTAGTGCAGGAGCTTATCCCCTCCTCTGGAGCTATGCGATGGG

TGTCGGAGTTGAGCTGGAGAACTCAATGGGGGGCTTGAACTTTGGCAGGTCATATTTTGACCCTGCCTATTTTCGTCTCG

GACAGGAGATGGTCAGAAGATCTGCAGGAAAGGTCAGCTCTGTGATTGCGGCTGAGCTCGGCATCACAGCAGAGGAAGCT

AAACTAGTCTCGGAAATAGCCTCGCAGGCTGGGGACGAAAGAACCGCTAGAGGGACTGGGCCTCGACAGGCGCAGGTCTC

CTTCCTCCAGCACCAAACAGGAGGGGGAGAGTCGTCCGCACCAGCGACCAGAGAAGGGGTCAAAGCTGTGATCCCAAACG

GATCCGAAGAAAGGGACAGAAAGCAAACACGCCCAGGAAGGCCCAGAGGAGAGACCCCCGGCCAACTGCTCCTGGAAATC

ATGCCAGAGGATGAGGTTTCGCGAGAATCTGGTCAAAACCCTCGTGAGGCTCAAAGATCGGCCGAGGCACTCTTCAGGCT

GCAGGCCATGGCCAAGATTCTGGAGGACCAGGAGGAGGGAGAAGACAACAATCAGGTCTACAACGACAAGGATCTCCTCG

GCTAAGCAGACGCACCCTCTGTCGAAATCAGTGACGAGACATCTCCTACCAGTATTATAAAAAACTTAGGACCCAGGTCC

AAGCAAGCACACATCGACACTCCAACCAGTCGAGCGGAGGCCACCGATGGCAGAAGAACAAGCATACCATGTCAACAAGG

GGCTGGAATGTATCAAGTCCCTCAAAGCCTCTCCCCCGGATCTATCCACCATCAGAGATACCATCGAGAGCTGGAGAGAG

GGGCTTAGCCCATCGGGCCGTGCAACACCGAACCCTGGTATGTCCGAGGGAGATCATCAGAATATCAACCAACCATGCTC

ACCAGCAATCGGACCAAACAAAGTCTACTTGTCTCCTGAAGATAATCTCGGATTTAGAGAGATCACTAGCAACGACTATG

AGGCTGAGCTCGGAGGAGTCCAGGGAAGAGGATCCAACTCTCAAGTACAGCGTTACTATGTTTATAGCCACGGGGGTGAA

GAGATTGAAGGACTCGAGGATGCTGACTCTCTCGTGGTTCAAGCAGATCCTCCAGTTGCTAACATATTCAATGGAGGAGA

GGATGGATCTGACGACAGCGATGTGGACTCTGGCCCAGATGATCCCGGCAGAGATACTCTATATGACCGGGGATCTGTTG

CCAGCAATGATGTCGCTAGGTCCACAGATGTCGAAAAACTAGAAGGTGCTGATATTCAAGAAGTTCTCAACTCCCAGAAA

GGCAGAGGAGGAAGATTCCAAGGCGGGAAAACCTTACGAGTCCCGGAAATACCCGATGTCAAGCACTCCAGACCATCAGC

CCAATCAATTAAAAAGGGCACAGACGGGAACTCAGTCTCATCTGGAACGGTGACAGAGTGTTTATCGATAAGTGGTGCAA

CCCAAGCTGTGCCAGAGTCAAGATGGGAGTCATCAGAGCAAAATGCGTCTGTGGGGAGTGTCCTCAAGTCTGCGGGGAGT

GCAAAGACGATCCAGGGGTCGACACAAGAATCTGGTACCATAGCATCACTGACTCAGCCTAAAGAGAATGACTCCGAGTA

TGAGTATGAGGATGACCTATTTACAGAGATTCAGGACATCCGTGCAAGCATTGCCAAGATCCATGATGACAACAAGGCTA

TCCTCTCAAAGCTTGATTCTATACTGTTATTGAAAGGAGAAGTCGACACTATCAAGAAACAAATCAGCAAGCAGAATATA

AGTATATCCACCATTGAGGGCCATCTCTCCAGTATAATGATAGCCATCCCGGGCTTTGGGAAGGATATCAAGGACCCAAC

ATCCGAGGTTGAGTTGAACCCAGATTTAAGACCTATAATAAGCCGTGACTCTGGCAGAGCTCTCGCGGAGGTCCTCAAGA

AACCCGCTGTGGATAGGTCTCAGAAAATTGGAACCAAAGCCAACTCCAGCTCAAAGGGTCAGCTTCTTAAGGATCTCCAG

CTAAAACCTGTCGACAAGCAGGCAAGCTCTGCAATCGGGTTTGTCCCATCCGACCATGAATCATCCAGAAATGTCATCCG

CTCCATAATCAAGTCGAGCAAGCTCAACATTGATCACAAGGACTATCTTCTAGATTTACTGAATGATGTGAAAGGCTCCA

AGGATCTTAAGGAATTCCACAAGATGCTAACAGCAATTCTCGCCAAGCACCCGTAACACATCCTCCAGTCACCACCCCAT

ACTCGGCTGAAAACATCCTTTCAATCAGGCTATTACAAAAAACTTAGGAGCAAGGGCAACTGAGCTTCGCAGACAAGATG

ACCGAGATCTACGACTTCGATAAATCAGCATGGGATGTCAAAGGGTCAATTGCCCGCATAGAACCCACCACCTACCACGA

CGGCCGACTGATACCCCAGGTAAGGGTCATCGATCCTGGTCTGGGAGACAGAAAAGATGAGTGCTTTATGTACCTGTTTC

TCCTGGGAGTGATTGAGGATAACGACCCCCTGTCTCCCCCCGTCGGAAGAACCTTTGGCTCTTTACCTCTAGGGGTCGGT

AGGTCAACTGCTAAGCCAGAGGAACTACTAAGGGAGGCCACAGAACTAGACATAGTGGTGAGGCGCACTGCAGGAGTAAA

TGAGAAACTGGTATTTTACAACAACACTCCGCTGTCCTTGTTAACGCCCTGGAAGAAAGTTCTGACAACCGGAAGTGTGT

TTAGCGCTAACCAGGTTTGCAATGCAGTCAACCTAGTCCCGCTTGATACTCCCCAGAGGTTCAGGGTTGTGTACATGAGC

ATAACTAGATTATCAGACAATGGTTACTATAGTGTGCCCAGAAGAATGCTGGAATTCCGCTCAGCCAATGCAGTAGCCTT

CAACATCTTGGTTACACTAAAAATTGAAAATGGCACAAACCCTAGAAGATACATAGTCGGCTCATGGGAGAATTCAGAGG

TCACATTTATGGTACACGTGGGCAACTTTAGAAGAAAGAAGAACGAAGTATACTCTGCTGATTACTGCAAAATGAAGATT

GAAAAGATGGGTTTAGTTTTTGCCCTGGGAGGAATAGGTGGAACAAGTCTCCATATTAGAAGCACCGGGAAAATGAGCAA

AACCCTCCATGCACAGCTGGGGTTCAAGAAAATCTTATGTTACCCCCTAATGGATGTTAATGAGGATCTTAACCGATATC

TCTGGCGGGCAGAGTGCCGAATAGTCAAAATCCAAGCCGTCCTACAGCCATCAGTGCCCCAAGAATTCCGTGTCTACGAT

GATGTCATCATCAACGATGATCAAGGCTTGTTCAAGATCTTGTAGTTCATTTGCAAAGGAACGCACCCACCCCCAGGAAG

GGGCCCCACCCCCCCCCG-CCACAAAACACACTCCAAGGCTCCCCGACACAGCGCCAACACAGCCACCCCCCCGACTGGA

CGAGGCCGGCC-CCCCAACAAACCCGGTCCAGGCGAGGGG-CCCCCCCCCTGACCCGCCGGGCGCCGACCGAGCAGGCCG

GCGAGAAGCCCCCAAGAGGCCCCGACAAAGACCGAGCACCCCATCCCCCC-AAAACGTCTCTCC-AACACGCTCGAGGCC

AGGCGCCGGGCACCCCCAGATCCGCCACCGGCGAGACAGAACCCCAGGGGGGGACCTCCCAAACCAAAGGGACCACCGGG

GAAGGCTCCGCCGAACCAGCACAGACCCGACCCAAGCAAGGCGGACCCCCAACACGAAACCGCCCAGCACACCCCACTCA

TCATCAAACAAAACTTAGGGGCCAAGCCCATACGACAGGCCCCAGTCCGCCAGGCGGAGGGACGCAGAAAGGAAGGAGAC

ATGACCGCCCCAGAAGGGGGACACCCAACCACCGAAACACCCAGCCCCAGGGAAGGAG--CCCCCCCCCACACGGCGAAG

GAACGGGTGAGAGACCCCGAGGAGAACACGACCGGGAGCCAGGAGCCGGCGGCGCCGGCAGCAAACAAGCCGGGCAAACA

CCCGACCCCACCAGAGGAGCCGAGGAGAGCCCGACCCCGCACGAACCAAACGGCAGGGAGCCCAACCCCACCACAAAAAG

AGGCGAGGCCCC-CCT-----CCCCCCCGCACACAACCCGCAGACAGACCGAGGGAGGACCACCCCCCAACCAGACAGGC

GGCCCCCCCGCAGCCAAGCCGCAGCTCCTCGGAACAGGACACCCCAGCACACTCCGAGCAACCCCCCTCCCCCCGCCACC

GGGACAGGGGCTCCGCCACCACAGACCAAAGCCACAGAGAGCCAGCCAGAGGTCCCCCCCAGTCTCGCCCGGTGCCCCCC

CGAAAAGGCCCCGGCAAGGCCCCCCAACCCGACGTCCCCCGCACGCACCGAAAGAAGGGCCCGCCCGAGAACGTGCGGCA

CATCCATGCGCAAACATCATGACACGGGTCGCAATCTTGACATTTCTGTTTCTTCTCCCAAATGTTGTTGCGTGCCAGAT

TCACTGGGGCAATCTATCCAAGATCGGGATTGTAGGAACAGGGAGTGCCAGCTACAAGGTGATGACTAGGCCAAGCCACC

AAACTCTGGTTATAAAGTTAATGCCAAATATAACAGCCATCAACAATTGTACAAAGTCAGAGATTGCAGAGTACAAGAGA

TTGCTGATCACAGTGTTAAAACCTGTAGAGGATGCTCTGTCGGTGATAACCAAGAATGTAAGACCAATTCAAACCCTGAC

ACCTGGGCGTAGAACCCGTCGTTTTGCTGGAGCTGTTCTGGCCGGAGTAGCACTTGGAGTTGCGACAGCCGCTCAGATAA

CTGCAGGAGTCGCTCTTCATCAATCATTGATGAACTCTCAAGCAATTGAGAGTTTAAAAACCAGTCTTGAGAAGTCGAAT

CAGGCAATAGAAGAAATCAGACTTGCAAATAAGGAGACCATACTGGCAGTACAGGGCGTCCAGGATTATATCAACAATGA

GCTTGTCCCTTCTGTTCATAGAATGTCATGCGAGCTGGTAGGCCACAAGCTCGGCCTCAAGCTCCTTAGGTACTACACCG

AGATCCTGTCCATATTCGGGCCCAGCCTTCGAGACCCGATAGCTGCCGAAATATCGATCCAGGCACTCAGTTATGCATTA

GGCGGGGACATTAATAAAATCCTGGACAAGCTTGGGTATAGCGGTGGGGATTTCCTTGCCATCCTAGAGAGCAAGGGAAT

AAAGGCCCGGGTTACATATGTGGACACAAGAGATTACTTTATAATCCTTAGCATCGCCTACCCAACCTTATCTGAGATCA

AGGGAGTGATAGTTCACAAGATAGAAGCTATAACATACAACATTGGGGCACAGGAGTGGTATACTACTATCCCTAAATAT

GTAGCCACTCAGGGGTATCTGATATCGAACTTTGATGAGACGTCATGCGTATTCACTCCAGAGGGGACAGTTTGCAGCCA

GAATGCGTTGTACCCAATGAGCCCATTGCTTCAGGAATGTTTCAGGGGATCAACAAAATCGTGCGCCAGAACCCTAGTTT

CAGGGACCATAAGTAATAGATTTATCCTATCAAAAGGGAACCTGATTGCAAATTGTGCGTCAGTTTTGTGCAAATGTTAC

ACAACGGAGACAGTCATCAGCCAAGATCCTGACAAACTACTAACTGTTGTAGCATCCGACAAGTGTCCTGTAGTTGAGGT

GGATGGAGTGACAATACAGGTCGGCAGTCGAGAGTATCCGGATTCTGTATACTTACACAAAATAGACTTAGGTCCAGCCA

TCTCCCTAGAAAAACTGGATGTAGGCACCAATTTAGGCAATGCAGTCACAAGACTGGAGAATGCAAAGGAGCTCCTAGAT

GCATCAGACCAAATACTGAAGACTGTCAAAGGGGCACCTTTCGGCGGGAACATGTACATAGCACTGGCAGCTTGCATTGG

AGTATCCCTAGGGCTTGTCACATTAATATGCTGCTGTAAGGGGAGGTGTAAGAACAAGGAAATCCCTATCTCCAAAATCA

ACCCAGGGCTCAAACCCGATCTGACCGGGACCTCAAAGTCGTACGTGAGATCACTGTAGCCAGAATCACCTGAATCATCT

GGCATCACACACATACATGTACGACACAAGCAGTCTGAGGACGCAGGAAACCCAGCCTCCGATCACCTACCTGACCCTAC

TCCACGCTCCACTACACATTAGTCATCAAACAAAACTTAGGACGAAAGGTCAATCACCATGTCCGCACAAAGGGAAAGGA

TCAATGCCTTCTACAAAGACAATCCTCACAATAAGAACCATAGGGTGATCCTGGATAGAGAACGCTTGGTCATTGAAAGA

CCCTACATCTTGCTTGGGGTCCTGCTGGTAATGTTCCTGAGTCTAATCGGACTGCTGGCCATTGCAGGGATCAGGCTTCA

CCGGGCCACCGTTGGAACTTCAGAGATCCAGAGTCGGCTGAATACCAATATTAAGTTGACCGAATCTATTGATCACCAGA

CTAAGGATGTCTTAACTCCCCTTTTTAAAATCATTGGCGATGAAGTCGGCATCAGAATTCCACAGAAATTCAGTGATCTT

GTTAAGTTCATCTCCGATAAGATTAAATTCCTCAACCCTGATAGAGAGTATGATTTCAGGGATCTCCGGTGGTGCATGAA

TCCCCCCGAGAGAGTCAAAATTAATTTTGATCAGTTTTGTGAGTACAAAGCTGCGGTTAAGTCAATTGAACATATATTTG

AGTCACCACTCAACAAGTCAAAAGAGCTGCAATCATTGACTCTCGGGCCCGGAACAGGCTGTCTAGGCAGGACAGTAACA

AAAGCTCATTTCTCAGAACTTACACTGACCTTAATGGACCTGGATCTAGAGATGAAGCACAACGTGTCCTCAGTGTTTAC

CGTAGTTGAAGAGGGGTTATTCGGAAGAACATATACCGTCTGGAGATCTGATGCCAGGGATCCGAGCACCGATCTAGGTA

TCGGCCATTTTTTAAGAGTCTTCGAGATTGGACTGATAAGAGATCTCGGGCTGGGTCCCCCTGTTTTTCATATGACCAAC

TATCTCACAGTGAACATGAGTGATGACTATCGGAGATGTCTTTTAGCGGTAGGGGAGTTGAAGTTGACAGCCCTATGCAC

CTCATCTGAGACTGTGACACTGAGTGAGAGAGGAGCTCCAAAGAGGGAGCCTCTTGTGGTTGTGATACTTAATCTGGCTG

GACCCACTCTAGGGGGCGAACTATACAGTGTCTTGCCTACCTCCGATCTCATGGTGGAGAAACTCTATTTATCTTCACAT

AGAGGGATCATCAAGGATGACGAGGCCAATTGGGTAGTGCCGTCTACCGATGTTCGTGATCTTCAGAACAAAGGTGAATG

TCTGGTGGAAGCATGCAAGACTCGACCTCCTTCATTTTGCAATGGCACAGGATCAGGCCCGTGGTCAGAGGGGAGAATCC

CTGCTTACGGGGTGATCAGGGTCAGTCTTGACTTAGCTAGTGACCCGGATGTAGTTATCACTTCAGTGTTTGGCCCACTG

ATTCCTCACCTATCCGGCATGGATCTTTACAACAACCCGTTTTCAAGAGCTATATGGTTGGCTGTACCACCTTATGAGCA

GTCATTTCTAGGAATGATAAATACAATTGGATTCCCTAACAGAGCAGAGGTTATGCCGCACATTTTGACCACAGAGATCA

GAGGCCCTCGGGGTCGTTGCCATGTTCCCATAGAATTGTCCCGCAGGGTTGATGACGATATCAAGATCGGGTCCAACATG

GTCATATTGCCGACGATGGACCTGAGGTATATTACAGCCACTTATGATGTTTCCAGGAGCGAGCATGCAATCGTGTACTA

TATCTATGACACAGGTCGCTCATCATCTTACTTCTACCCAGTTCGACTGAATTTCAAAGGCAATCCTCTCTCTCTGAGGA

TAGAGTGTTTCCCTTGGCGTCATAAGGTGTGGTGCTACCATGATTGTCTTATATACAACACCATAACAGATGAAGAGGTC

CATACGAGAGGGCTGACCGGTATAGAGGTAACATGCAATCCAGTTTGAGCAAAGCTACGACCGTCACTCAAGCAGTCCCT

TGAGTCGCCACCGAGTCCAAGCAGCATAGCCTGGGACACTCAACAGCACAGCCCAGCCAACAATGTTATAAAAAACTTAG

GAGCCAGGGTTGTAAGGGCCATGGACTCCCTATCAGTCAATCAGGTCTTGTACCCTGAGGTTCATCTAGACAGCCCTATT

GTCACAAACAAACTAGTTGCCATCCTTGAGTACTCGGGTATCGACCATAACTATGTTCTTGAAGACCAGACCCTTGTCAA

GAATATTAGGTATAGACTGGGGTGCGGTTTTTCAAATCAAATGATCATCAATAACAGGGGGGTGGGTGAAACAGTCAATT

CTAAACTTAAAAGTTATCCCCATAATCGTCATATCATATACCCGGATTGCAATAAGGAGTTGTTTGGTATCAAAGATAGC

TGCATATCTAAGAAGCTCTCGGAGCTATTCAAGAAGGGTAATTCCTTGTACTCTAAGATAAGTCACCAGGTACTGGATTG

TCTTAAGAGAGTCAATAGGAAATTAGGCCTGGGCACAGATCTTACCCATGGTCTGAAGGAAGGTATCCTTGACCTAGGGT

TGCACATGCATAGCTCTCAATGGTTCGAGACTTTCCTGTTCTGGTTCACTATCAAGACAGAGATGAGATCAATGATCAAA

GAACAGTCCCATATATGCCACAAGAGGAGGTATAACCCGATTTTTGTATCGGGGGATGCATTCGAGGTGCTCGTATCACG

AGACCTCGTTGTGATAATTGATAAGAATACCCAGTATGTCTTCTACTTAACATTTGAGCTGGTCCTTATGTATTGTGATG

TCATAGAGGGCAGACTTATGACGGAGACAGCCATGGCCATAGACCAGAGATATTCAGAACTTCTAAGCCGGGTTAGATAC

TTGTGGGATCTTATTGATGGGTTCTTCCCAACACTAGGCAACACCACATACCAAATTGTTGCTCTGCTTGAACCATTATC

GTTGGCTTATCTTCAACTTCAGGATGTCACGCTGGAGTTAAGAGGTGCTTTTTTAGACCATTGCTTCAAAGAACTCTATG

AGATACTGGAGCATTGCGGCATTGACACAGAAGGCACCTACAATTCCATCACTGAAGGACTGGATTACGTATTCATCACC

CACGATATACATCTAACTGGGGAGATTTTTTCATTTTTCCGGAGTTTCGGACACCCCCGTCTAGAAGCAGTCACCGCTGC

AGAAAATGTCAGGAAACATATGAACCAACCGAAGGTAATCAGTTATGAGACTATGATGAAAGGACATGCGGTATTTTGCG

GGATAATCATAAATGGTTTTAGGGACCGACATGGCGGCAGCTGGCCCCCTGTTGCATTGCCAGAACATGCTTCTGCTGCG

ATCCGGAATGCGCAGGCATCAGGTGAGGGACTGACCCATGACCTGTGTATAGACAACTGGAAATCCTTTGTTGGATTCAA

ATTTGGCTGCTTTATGCCACTCAGCCTAGATAGTGATTTGACCATGTATCTCAAAGACAAGGCATTGGCTGCACTGAAGA

ATGAGTGGGATTCAGTTTACCCGAAAGAATACCTCCGTTATAACCCACCTAGAGGGACAGAGTCAAGGCGACTGGTGGAG

GTGTTCCTGAATGACTCCAGCTTTGATCCTTATAACATGATAATGTACGTGGTGAATGGCTCCTACCTTAAAGACCCTGA

GTTTAATCTCTCATACAGCCTAAAAGAGAAGGAGATAAAGGAGACAGGGCGGTTGTTTGCCAAAATGACTTATAAGATGC

GGGCCTGTCAGGTAATTGCTGAAAATCTGATATCGAATGGTGTTGGGAAGTATTTCCGAGACAATGGGATGGCAAAGGAC

GAGCATGACCTAACAAAAGCCCTTCACACCCTGGCAGTCTCAGGTGTTCCCAAGAACAACAAAGATAACCACCGAGGTGG

GCCTCCCAGAAGGACAACAAGCCGAGAGGTGAGATCAAGCCAAGACACCAAAACACAAAATAGAGACAAGGTCCAAGGAG

GACCTATGTACAACTATTTGCGATGCCAACCAATCAGCCCTGATCAGGGTGAGTCATACGAGACTGTTAGTGCATTCATC

ACCGCTGACCTTAAGAAGTATTGCCTGAATTGGAGATACGAGACAATCAGCATATTTGCACAGAGGCTGAATGAAATATA

TGGACTACCATCCTTCTTTCAATGGTTACACAGGATATTGGAAAAATCCGTACTCTACGTCAGTGACCCACATTGCCCTC

CCGATCTAGATAATCATATCCCTCTGGACAGTGTCCCTAATGCTCAAATATTCATTAAGTACCCAATGGGCGGAATAGAA

GGTTATTGCCAAAAACTATGGACAATCAGTACTATACCATACTTGTATCTGGCAGCCTATGAGAGCGGAGTAAGAATCGC

CTCACTGGTGCAGGGTGACAATCAGACAATCGCAGTGACAAAAAGAGTTCCAAGCTCTTGGCCTTATTCACTAAAAAAGA

AGGAGGCATCCAGAGCCGCTCAAAACTACTTCGTGGTCTTAAGGCAAAGATTGCACGATGTAGGTCATCACTTAAAGGCT

AATGAAACCATAGTATCTTCTCACTTTTTTGTATATTCCAAAGGGATTTATTATGACGGCCTGCTAGTCTCACAATCACT

GAAGAGCATCGCCAGATGTGTCTTCTGGTCCGAGACTATAGTGGATGAAACCAGAGCGGCTTGCAGTAATATTGCAACAA

CTGTCGCTAAGAGTATAGAGAGGGGTTATGATAGGTACCTTGCATATTCTTTGAATATCCTCAAGATTTTCCAACAGATC

CTTATATCCCTTAACTTCACTATTAACACAACAATGACTCAGGATGTTGTGGCACCGATCATCGAGAACGGTGATTTGCT

GATAAGGATGGCACTCTTGCCAGCACCCATCGGAGGTCTAAATTATCTTAACATGAGCAGGTTATTTGTGAGAAATATCG

GTGACCCGGTCACCTCCTCTATAGCCGACCTGAAGAGGATGATAGATGCCGGGCTAATGCCAGAAGAAACATTGCATCAA

GTGATGACCCAGACCCCGGGAGAATCATCCTACCTCGATTGGGCAAGTGACCCTTATTCTGCCAACCTACCCTGCGTACA

GAGTATAACTCGCCTTCTCAAAAACATCACTGCACGATATATTTTGATCAGCAGCCCAAACCCGATGCTGAGAGGATTGT

TTCATGAGGGGAGTAGAGATGAAGACGAGGAGCTTGCAAGTTTCCTAATGGATCGGCATATAATTGTTCCAAGAGCTGCA

CATGAAATCTTAGACCATAGTATAACCGGGGCAAGAGAAGCTATAGCCGGGATGTTGGACACCACCAAAGGTCTGATTAG

AACAAGTATGAAACGGGGTGGCCTCACCCCTCGAGTCTTAGCCCGCCTTTCCAATTATGATTATGAACAATTCCGATCCG

GGATAACACTATTGACAAAGAAAGGGCAGTGTTATCTCATTGACAAGGATTCGTGCTCGGTGCAGCTCGCTATAGCTCTG

AGGAGCCACATGTGGGCTAGGTTGGCTCGCGGGAGACCAATCTATGGGTTGGAGGTGCCCGATATATTAGAATCGATGAA

CGGCTACCTTATCAAACGCCACGAGTCTTGTGCTATCTGTGAAACGGGCTCAAGTCACTACGGATGGTTTTTCGTCCCTG

CAGGGTGCCAGCTTGACGATATATCTAGAGAGACTTCGGCTCTTCGTGTACCTTATGTCGGATCAACAACTGAGGAAAGG

ACAGACATGAAACTTGCATTTGTTAGATCTCCAAGTAGATCCCTCAAATCAGCAGTTAGAATTGCAACAGTTTACTCATG

GGCCTACGGGGATGATGAGAAGTCATGGAGTGAAGCCTGGATGCTAGCTAGGCAGAGAGCCAATATCACCTTAGATGAAT

TGAGAATGATAACTCCAGTCTCTACATCCACCAACCTAGCCCACCGGTTGAGGGATCGGAGCACCCAGGTGAAATACTCG

GGGACATCCCTTGTGAGGGTTGCAAGATACACAACCATCTCCAATGATAACTTGTCATTTGTGATATCTGAGAAAAAAGT

GGATACCAACTTCATTTATCAGCAAGGGATGTTGCTTGGTCTTGGGATCCTTGAGAACCTCTTTAGGCTAGAGGCCACCA

CCGGAGTATCCAACACAGTGCTACACCTGCACGTGGAAACAGAATGTTGTGTTGTACCTATGGTGGATCACCCAAGGATA

CCGAGTCTCCGTAATATAAAAGTTACGGATGAGCTATGCACAAACCCTTTGATCTACGACAGGTCTCCCATCATAGAGCA

CGATGCAACCCGACTATACTCACAGAGCCACAGGAGACATTTGGTGGAGTTTGTTACCTGGTCCACAAGCCAGCTTTACC

ATATACTGGCTAAGTCTACAGCAATGTCCATGATTGAATTGATCACGAGATTCGAGAAAGATCACATGAATGAAATAGCC

GCCCTGATTGGCGATGACGACATCAACAGCTTCATCACAGAATTTTTGCTAGTGGAGCCTAGATTACTTATAGTTTACCT

TGGTCAGTGTGCTGCCATCAATTGGGCCTTTGATATACATTATCATCGGCCCTCGGGCAAGTACCAGATGGGGGAACTCC

TCTACTCTTTGCTCTCTCGGATGAGCAAAGGAGTATATAAGATCTTCACCAATGCTTTGAGCCACCCCAAAGTTTACAAG

AAATTTTGGCGAAGTGGTGTCATTGAGCCGATTCATGGCCCATCCCTGGATACACAGAATTTACATGTCACTGTCTGTGA

CATGATATATGGGTCCTATGTCACCTATTTGGATCTTTTGCTGAATGATGAGCTAGATGATTACCCGTATTTGCTCTGCG

AGAGTGATGAGGACGTGGTCACAGACAGGTTCGACAACATTCAAGCCAAACATCTCTGTGTATTAGCCGATGTATATTGC

AGCTCCAAGAGGTGTCCTTCAATAATCGGGATGTCTCCTATAGAAAAATGTACCATTCTCACACATTACATCAAGGGAGA

ATCAGTACAATCCCCGTCTGGGACCTCATGGAACACTGATCCCCTTGTAGTAGATCATTACTCATGCTCTCTGACCTACC

TTCGCCGCGGTTCCATCAAACAAATCAGGTTGAGGGTGGATCCTGGGTTTGTATTCGAGGCGTTGACAGACGTCGACCTT

AAACAACCTCGCAAAGCTAAGTTAGATATATCGGTCGTTGGGTTGACTGATTTCTCTCCCCCTTGCGACAACGTCGGTGA

TTTTCTAGGGACTATCAACACATTGAGGCACAATCTGCCTGTCACCGGAACCGGGGTCTCGAACTATGAAGTCCACGCTT

ATCGTAGAATTGGCCTAAATTCATCAGCATGTTACAAAGCCGTAGAGATCTCCACGTTAATCAAGCCATCCCTAGAAGTC

GGAGAGCATGGATTGTTCTTAGGAGAAGGTTCCGGTTCAATGCTGGCTGCGTACAAGGAAGTTCTTAAATTAGCAAATTG

TTATTACAACAGCGGGGTAACAACGGAGGGCAGAGCCGGACAGAGGGAAATATCTCCCTATCCTTCAGAGATGAGCCTAG

TAGAGAATCAAATGGGGATAGAGCGGAGTGTTAAAGTGCTGTTCAACGGCAAACCTGAAGTAACTTGGGTGGGGACCACC

GATTGCTACAAGTACATAATCAGTAACATCCAAACCTCTAGTCTGGGTTTCATACATTCAGATATTGAGACGCTCCCAAC

CAAAGATGCTGTTGAGAAATTAGAAGAATTTGCCTCTATCCTATCCTTATCCTTGATTTTGGGGAAAATCGGCTCTATTA

CAGTTGTCAAAATTATGCCCATTAGCGGGGATTTTACCCAAGGCTTTATAGCCTATGCCATCCAATATTTTAGGGAGAGC

CTGCTTGCCTATCCGAGATATAGTAACTTCATCTCGACTGAGTGTTACCTTATTATGATAGGATTAAAGGCCAATCGACT

GATAAACCCAGAAGCCATTAAGCAAAGCATAATCAGAGCGGGGGTCAGGACTGCACCAGGACTTGTGAGCCATATATTAT

CAGAGAAACAGAAAGGTTGTATTCAATCTTTCTTGGGTGATCCTTATATCCAAGGAGACTTCAATAAGCACCTTAAATCT

CTGACCCCTATTGAGAAAATCCTGGTAAATTGTGGTCTCTCGATCAATGGCACAAAAATCTGTAGAGATCTAATCCACCA

TGATATCGCCTCCGGTCCAGACGGTCTTATGAGCTCTACAATTATTTTATATAGGGAATTGGCTCATTTCAAAGATAATA

TAAGGAGTCAGCACGGAATGTTCCACCCCTACCCAGTACTGGCCAACAGCCGGCAACGTGAATTAATCCTTCGAATAGCC

AAGAAATTCTGGGGGTATGTCTTGCTATATTCTGATGACCCGGCACTAATCAGACAAACAATCAAGAACTTGAAACGGAA

TCATCTAACCTTTGACTTACATAGTAACCCGTTTATTAAAGGGCTATCCAAAGCTGAGAAACTGCTGGTGCGGACGAGTC

CACTAAGAAGGGAATGGTTGTTCACTCTCGATACGAAAGAAGTGAAAGAGTGGTTCAAATTGGTAGGTTACAGTGCACTC

ATCAGAGGCTAGTTAGCTATACATCTGCCCCCTTCTCCTCCGCCATGAGACCCCACTGACGATCCAGAAGATTAAAGAAA

ACTACATATTGGATAAGTATCTATTCCCAGCTTTGTCTGGT

>KY967610.1/sheep/Pakistan/2015

ACCAAACAAAGTTGGGTAAGGATAGATCTTATAATAACTATGGACTAGCAAACTTAGGAGTAAAGATCCTACTGTCGGGG

GGAGGAGGAGGAGCAAGATCTTTGATTATGGCGACTCTCCTTAAAAGCTTGGCATTGTTCAAAAGGAACAAAGACAAAGC

GCCGACGGCATCAGGTTCAGGGGGGGCCATCCGAGGGATTAAGAATGTTATCATAGTCCCGATTCCCGGGGACTCGTCCA

TCACTACCCGTTCAAGACTGCTCGACAGGCTTGTCAGATTGGCCGGAGATCCTGACATCAACGGGTCAAAGCTGACCGGC

GTGATGATCAGCATGTTATCTTTGTTCGTAGAGTCACCCGGGCAGTTGATACAGCGAATCACAGATGATCCGGATGTTAG

TATCCGCCTTGTTGAGGTAGTTCAAAGTACTAGGTCCCAGTCCGGGTTGACCTTTGCATCACGTGGTGCTGATTTGGATA

ACGAGGCAGACATGTATTTTTCAACTGAGGGGCCCTCGAGTGGAGGTAAGAAAAGGATCAACTGGTTTGAGAACAGAGAA

ATAATAGACATAGAGGTGCAGGATCCAGAAGAGTTCAATATGTTGTTAGCCTCCATACTAGCACAAGTCTGGATCCTTCT

GGCCAAGGCTGTTACGGCACCGGATACGGCAGCTGACTCAGAACTGAGGAGGTGGGTTAAATACACACAACAAAGGAGAG

TGATTGGGGAATTTCGCCTTGACAAAGGGTGGCTGGACGCAGTCCGTAACAGAATTGCAGAAGATCTATCACTCCGGCGG

TTCATGGTATCTCTCATACTTGACATCAAGAGGACTCCCGGCAACAAGCCAAGGATTGCAGAAATGATCTGCGACATTGA

CAACTATATTGTCGAAGCCGGACTCGCCAGTTTCATCCTTACTATCAAGTTTGGTATCGAAACCATGTATCCTGCATTAG

GGCTCCACGAGTTTGCCGGGGAATTGTCCACTATTGAGTCCTTGATGAACTTGTATCAACAGCTAGGAGAGGTTGCACCC

TACATGGTAATTCTAGAGAACTCAATTCAGAACAAGTTTAGTGCAGGAGCTTATCCCCTCCTCTGGAGCTATGCGATGGG

TGTCGGAGTTGAGCTAGAGAACTCAATGGGGGGCTTGAACTTTGGCAGGTCATATTTTGACCCTGCCTATTTTCGTCTCG

GACAGGAGATGGTCAGAAGATCTGCAGGAAAGGTCAGCTCTGTGATTGCGGCTGAGCTCGGCATCACAGCAGAGGAAGCT

AAACTAGTCTCGGAAATAGCCTCGCAGGCTGGGGACGAAAGAACCGCTAGAGGGACTGGGCCTCGACAGGCGCAGGTCTC

CTTCCTCCAGCACAAAACAGGAGAGGGAGAGTCGCCCGCACCAGCGACTAGAGAAGGGGTCAAAGCTGCGATCTCGAACG

GATCTGAAGAGAGGGACAGAAAGCAAACACGCCCAGGAAGGTCCAGAGGAGAGACCCCCGGCCAACTGCTCCTAGAAATC

ATGCCAGAGGATGAGGTCTCGCGAGAGTCTGGTCAAAACCCTCGTGAGGCTCAAAGATCGGCTGAGGCACTCTTCAGGCT

GCAGGCCATGGCCAAGATCCTGGAGGACCAGGAGGAGGGAGAAGATAACAGTCAGATCTACAATGACAAGGATCTCCTCG

GCTAAGCAGACGCGCCCTCTGTCGAAATCAGTGACAATACATCTCCTACCAGTATTATAAAAAACTTAGGACCCAGGTCC

AGGCAAGCACACACCGACACTCCAACCTGTCGAGCGGAGACCACCGATGGCAGAAGAACAAGCATACCATGTCAACAAGG

GGCTGGAATGTATCAAGTCTCTCAAAGCCTCTCCCCCGGATCTATCCACCATCAGAGACACCATCGAGAGCTGGAGAGAG

GGGCTTAGCCCCTCGGGCCGTGCAACACCGGATCCTGATACGTCCGAGGGAGATCATCAGAATATCAACCAATCATGCTC

ACCAGCAATCGGACCAAGCAAAGTCTACCTGTCTCCTGAAGATAATCTCGGATTTAGAGAGATCACTGGCAACGACTGTG

AGGCTGGGCTCGGAGGAGTCCAGGGAAAAGGATCCAACTCTCAAGTACAGCGTTACTATGTTTATAGCCACGGGGGTGAA

GAGATTGAAGGACTCGAGGATGCTGACTCTCTCGTGGTTCAAGCAGACCCTCCAGTTGCTAACATCTTCGATGGAGGAGA

GGATGGATCTGACGACAGCGATGTGGACTCTGGCCCAGATGATCCCGGCAGAGATACTCTACATGACCGGGGATCTGTTG

CCGGCAATGATGTCGCTAGGTCCACGGATGTCGAAAAGCTAGAAGGTGCTGACCTTCAAGAAGTTCTTAACTCCCAGAAA

GGCAAAGGAGGAAGATTCCAAGGCGGGAAAACCTTACGAGTCCCGGAAATACCCGATGTCAAGCACTCCAGACCATCAGC

TCAATCAATTAAAAAGGGCACAGACGGGAGCTCAGTCTCATCTGGAACGGTGACAGAGTGTTTATCGATAAGTGGTGCAA

CCCAAGCTGTGCCAGAGTCAGGATGGGAGTCATCAGAGCAAAATGCGTCTGTGGGGAGTGTCCTCAAATCTGCGAGGAGT

GCAAAGACGATCCAGGGGTCGACACAAGAATCTGGTACCATAGCATCACTGACTCAGCCTAAAGAGAATGACTCCGAGTA

TGAGTATGAGGATGACCTATTTACAGAGATTCAGGACATCCGTGCAAGCATTGCCAAGATCCATGATGACAACAAGACTA

TCCTCTCAAAGCTTGATTCTATACTGTTATTGAAAGGAGAAGTCGACACTATCAAGAAACAAATCAGCAAGCAGAATATA

AGTATATCCACCATTGAGGGCCATCTCTCCAGTATAATGATAGCCATCCCGGGCTTTGGGAAGGATATCAAGGACCCAAC

ATCCGAGGTTGAGTTGAACCCAGATTTAAGACCTATAATAAGCCGTGACTCTGGCAGAGCTCTCGCGGAGGTCCTCAAGA

AACCCGCTGTGGATAGGTCTCAGAAAACTGGAACCAAAGCCAACTCCAGCTCAAAGGGTCAGCTTCTTAAGGATCTCCAG

CTAAAACCTGTCGACAAGCAGGCAAGCTCTGCAATCGGGTTTGTCCCGTCCGACCATGAATCATCCAGAAATGTCATCCG

CTCCATAATCAAGTCGAGCAAGCTAAACATTGATCACAAGGACTATCTTCTAGATTTACTGAATGATGTGAAAGGCTCCA

AGGATCTTAAGGAATTCCACAAGATGCTAACAGCAATTCTCGCCAAGCACCCGTAACACATCTCCCAGTCACCATCTCAT

ACTCGGCTGAAGACATCCTTTCAATCAGGCTATTACAAAAAACTTAGGAGCAAGGGCAACTGAGCTTCGCAGACAAGATG

ACAGAGATCTACGACTTCGATAAATCAGCATGGGATGTCAAAGGGTCAATTGCCCGCATAGAACCCACCACCTACCACGA

TGGCCGCCTGATACCCCAGGTAAGAGTCATCGATCCTGGTCTGGGAGACAGAAAAGATGAGTGCTTCATGTACCTGTTTC

TCCTAGGAGTGATTGAGGATAACGACCCCCTGTCTCCCCCCGTCGGGAGAACCTTTGGCTCTTTACCTCTAGGGGTCGGT

AGGTCAACTGCTAAGCCAGAAGAACTACTAAGGGAGGCCACAGAACTAGATATAGTGGTGAGGCGCACTGCAGGAGTAAA

TGAGAAACTGGTATTTTACAACAACACACCGCTGTCCCTGTTAACGCCCTGGAAGAAAGTTCTGACAGCCGGAAGTGTGT

TTAGCGCTAACCAGGTTTGCAATGCAGTCAACCTAGTCCCGCTTGATACTCCCCAGAGATTCAGGGTTGTGTACATGAGC

ATAACTAGATTATCAGACAATGGTTACTATAGTGTGCCCAGAAGAATGCTGGAGTTCCGCTCAGCCAATGCAGTAGCCTT

CAATATCTTGGTTACACTAAGAATTGAAAATGGCACAAACCCTAAAAGATACATAGTCGGCTCATGGGAGAATCCAGAGG

TCACATTTATGGTACACGTGGGGAACTTTAGAAGAAAGAAGAACGAAGTATACTCTGCTGATTACTGCAAAATGAAGATT

GAAAAGATGGGTTTAGTTTTTGCCCTGGGAGGAATAGGTGGAACAAGTCTCCATATTAGGAGCACCGGGAAAATGAGCAA

AACCCTCCATGCACAGCTGGGGTTCAAGAAAATCTTATGTTACCCCCTAATGGATGTTAATGAGGATCTTAACCGATATC

TCTGGCGGGCAGAGTGCAGAATAGTCAAAATCCAAGCCGTCTTACAGCCATCAGTACCCCAAGAATTCCGTGTCTACGAT

GATGTCATCATCAACGATGATCAAGGCTTGTTCAAGATCTTGTAGTTCATTTGCAAAGGAATGAACCCACCCCTAGGATG

GGGCCCCACCCTCCCCCG-TCAAAAAACACACTCCAAGGCTCCTCGGCACAGCGCCAACACAGCTACTCCCGCGATGGAA

CGAGGTCGGCC-CCCCAACAAATCCGGTCCAGGT-GGGGGGCCCCCCCCCTGACCAACCGGGCGCTGACCGAGCAGGCCG

GCGAGAACCCCCCAAGAGGCCCCGACAAAAACCGACCACCCCATCCCCCC-AAAAATGTCCCCC-AACACGCCCGAGGCT

AGCCACCGGGCACCCCCAGACCCGCCACCGGCGAGACAGAACCCCAAGGGGGGACCCTCCAAACCAAAGGGATCACCGGG

GTAGGCTCCGCTGAACCAGCACAGACCCGACCCAAGCAAGACGGACCCCCAACACAAATCTGCCCAGCACACCCCACTCA

TCATCAAACAAAATTTAGGGGCCAAGTCCATACAAAAGGCCCCATTCCCTCAGGCGAAGGGACGCAAAAGGGAAGGAAAC

AAGACCGCCCCAGAAGGAGGACACCCAACCACCGAAACACCCAGCCCCAGGGAAGGAG--CCCCCCCCCACATGGCGAAA

GAACGGGCGAGAGACCCCGAGAAGAACCCGACCGGGAGCCAGGAGCCGGCGACGCCGGCGGCAAACAAGCCGGGCAAACA

CCCGACCCCACCAGAGGAGCCGAGGAGAGCCCGACCCCGCACGAACCAAACAGCAGGGAGCCCAACCCCACCAAAAAAAG

AGGCGAGGCCCC-CCT-----CCTCCCCGCACACAACCCGCAGACAGACCGAGAGAGGACCACCTCCCAACAAGACAGGC

GGCTCCCTCGCAGCCGAGCCGCAGCTCCCCAGAACAGGACACCCCCGTACACTCAAAACACCCCCCCGCCCCCCGCCACC

GGGACAGGGGCTCCGCCACCACAGACCAAAGCCACCGAGAGCAAGCCAGAGGTCCCCCCCAGTCCCGCCCGGCGCCCCCC

CGAAAAGGCCCCGGCAAGGCCCCCCAACCCGACGTCCCCCGCACGCACCGAAAGAGGGACCCGCCCGAGAACGTGCGGCA

CATCCATGCGTAAACATTATGACACGGGTCGCAATCTTGACATCTCTGTTTCTTCTCCCAAATGTTGTTGCGTGCCAGAT

TCACTGGGGCAATCTATCCAAGATCGGGATTGTGGGAACAGGGAGTGCCAGCTACAAGGTGATGACTAGGCCGAGCCACC

AGACTCTGGTTATAAAGTTAATGCCAAATATAACAGCCATCGACAATTGTACAAAGTCAGAGATTGCAGAGTACAAGAGA

TTGCTGATCACAGTGTTAAAGCCTGTAGAGGATGCTCTGTCAGTGATAACCAAGAATGTGAGACCAATTCAAACTCTGAC

ACCTGGGCGTAGAACTCGCCGTTTTGCTGGAGCTGTTCTGGCCGGAGTAGCACTTGGAGTTGCGACAGCCGCTCAGATAA

CTGCAGGAGTCGCCCTTCATCAATCACTGATGAACTCTCAAGCAATTGAGAGTTTAAAAACCAGTCTCGAGAAGTCGAAT

CAGGCAATAGAAGAAATCAGACTTGCCAATAAGGAGACCATACTGGCAGTACAGGGCGTCCAGGACTATATCAACAATGA

GCTTGTCCCTTCTGTTCATAGAATGTCATGCGAGCTTGTAGGTCACAAGCTCGGCCTCAAGCTCCTTAGGTACTACACCG

AGATCCTGTCCATATTCGGGCCCAGCCTTCGAGACCCGATAGCTGCCGAAATATCAATCCAGGCACTCAGTTATGCATTA

GGTGGAGACATTAATAAAATCCTGGACAAGCTTGGGTATAGCGGTGGGGATTTCCTTGCCATCCTAGAAAGCAAGGGAAT

AAAGGCCCGGGTCACATATGTGGATACGAGAGATTACTTTATAATTCTTAGCATCGCCTACCCAACCTTATCTGAGATCA

AGGGAGTGATAGTTCACAAGATAGAAGCTATAACATACAACATTGGGGCACAGGAGTGGTATACTACTATCCCTAAATAT

GTAGCCACTCAGGGGTATCTGATATCGAACTTTGATGAGACGTCATGCGTATTCACTCCAGAGGGGACAGTTTGCAGCCA

GAATGCGTTGTACCCAATGAGCCCATTGCTCCAGGAATGTTTTAGGGGGTCAACAAAATCGTGCGCCAGAACCCTAGTTT

CAGGGACCATAAGTAATAGATTTATCCTATCGAAAGGGAACCTGATTGCAAATTGTGCGTCAGTTTTGTGCAAATGCTAC

ACAACGGAGACAGTTATCAGCCAAGATCCTGACAAACTACTAACTGTTGTAGCATCCGACAAGTGTCCTGTAGTTGAGGT

GGACGGAGTGACAATACAGGTCGGCAGTCGAGAATATCCGGATTCTGTATACTTACACAAAATAGACTTAGGTCCAGCCA

TCTCCCTAGAAAAACTGGATGTAGGCACCAATTTAGGCAATGCAGTCACAAGACTGGAGAATGCAAAGGAGCTCCTAGAT

GCATCAGACCAAATACTGAAGACTGTTAAAGGGGCACCTTTCGGTGGGAATATGTACATAGCACTGGCAGCTTGCATTGG

AGTATCCCTAGGGCTTGTCACATTAATATGCTGCTGTAAGGGGAGGTGTAAGAACAAGGAAATCCCTATCTCCAAAATCA

ACCCAGGGCTCAAACCCGACCTGACCGGGACCTCAAAGTCGTACGTGAGATCACTGTAGTCTGAATCACTTGAATCATCC

AGCATCACACACATACAGGTACGACGCAGGCAGTCTGAGGACGCAGGAAACCCAGCCTCCGATCACCCACCTGACCCCAC

TCCACGCTTCACCACACATTAGTCATCAAACAAAACTTAGGACGAAAGGTCAATCACCATGTCCGCACAAAGGGAAAGGA

TCAATGCTTTCTACAAAGACAATCCTCACAATAAGAACCATAGGGTGATCCTGGATAGAGAACGCCTGGTCATTGAAAGA

CCTTACATCTTGCTTGGAGTCCTGCTGGTAATGTTCCTGAGTCTAATCGGACTGCTGGCCATCGCAGGGATCAGGCTTCA

CCGGGCCACCGTTGGGACTTCAGAGATCCAGAGTCGGCTGAATACCAATATTGAGTTGACCGAATCTATTGATCACCAAA

CTAAGGATGTCTTAACTCCCCTTTTTAAAATCATTGGCGATGAAGTCGGCATCAGGATTCCACAGAAATTCAGTGATCTT

GTCAAGTTCATCTCCGATAAGATTAAATTCCTTAACCCTGATAGAGAGTATGATTTCAGGGATCTCCGGTGGTGTATGAA

TCCCCCAGAGAGAGTCAAAATTAATTTTGATCAGTTTTGTGAGTACAAAGCTGCGGTTAAGTCAATTAACCATATATTTG

AGTCACCACTCAACAAGTCAAAAAAGCTGCAATCATTGACTCTTGGGCCCGGAACAGGCTGTCTAGGCAGGACAGTAACA

AGAGCTCATTTCTCAGAACTTACACTGACCTTAATGGACCTGGATCTAGAGATGAAGCACAATGTGTCCTCAGTGTTAAC

CGTTGTTGAAGAGGGATTATTCGGAAGAACGTATACCGTCTGGAGATCCGATGCCAGGGATCCGGGCACCGATCTAGGTA

TCGGCCATTTTTTAAGAGTCTTTGAGATTGGATTGGTAAGAGATCTCGGGCTGGGTCCCCCTGTTTTCCATATGACCAAC

TATCTCACAGTGAACATGAGTGATGACTATCGGAGATGTCTTTTAGCGGTAGGGGAGTTGAAGTTGACAGCCCTATGCAC

CTCATCTGAGACTGTGACACTGAGTGAGAGAGGAGTTCCAAAGAGGGAGCCTCTTGTGGTTGTGATACTTAACCTGGCTG

GGCCCACTCTAGGGGGCGAACTATACAGTGTCTTGCCTACCTCTGATCTCATGGTGGAGAAACTCTACCTATCTTCACAT

AGAGGGATTATCAAAGATGACGAGGCCAATTGGGTAGTGCCGTCTACCGATGTTCGTGATCTTCAAAACAAAGGTGAATG

TCTGGTGGAAGCATGCAAGACCCGACCTCCTTCATTCTGCAATGGCACAGGATCAGGCCCGTGGTCAGAGGGGAGAATCC

CTGCTTACGGGGTGATCAGGGTCAGTCTTGACTTAGCTAGTGACCCTGGTGTATTTATCACTTCAGTGTTTGGCCCACTG

ATACCTCACCTATCCGGCATGGATCTTTACAACAACCCGTTTTCAAGAGCTATATGGTTGGCTGTACCACCTTATGAACA

GTCATTTCTAGGAATGATAAATACCATTGGATTCCCTAACAGAACAGAGGTTATGCCGCACATTTTGACCACAGAGATCA

GAGGCCCTCGGGGTCGTTGCCATGTTCCCATAGAATTGTCCCGCAGGGTTGATGACGATATCAAGATCGGGTCCAACATG

GTCATATTGCCGACGATGGATTTGAGGTATATTACAGCCACTTATGATGTTTCCAGGAGCGAGCATGCAATCGTGTACTA

TATCTATGACACAGGTCGCTCATCATCTTACTTCTACCCAGTTAGACTAAATTTCAAAGGCAATCCTCTCTCTCTGAGGA

TAGAGTGCTTCCCTTGGCGTCATAAGGTGTGGTGCTACCATGATTGTCTTATATACAACACCATAACAGATGAAGAAGTC

CATACGAGAGGGCTGACCGGTATAGAGGTAACATGTAATCCAGTCTGAGCAGAGCTACGACCGTCACTCAGGCAGTCCCT

TGGGTCGCCACCGAGCCCAAGCAGCACAGCCTGGGACACTCAACAGCACAGCCCAGCCAACAACGTTATAAAAAACTTAG

GAGCCAGGGTTGTAGGGGCCATGGACTCTCTATCAGTCAATCAGGTCTTGTACCCTGAGGTTCATCTAGACAGCCCTATT

GTCACAAACAAACTAGTTGCCATCCTTGAGTACTCGGGTATCGACCATAACTATGTTCTTGAAGACCAGACTCTTGTCAA

GAATATTAGGTATAGGCTGGGGTGCGGTTTTTCAAATCAAATGATCATCAATAACAGGGGGGTGGGTGAAACAGTCAATT

CTAAACTTAAAAGTTACCCCCATAATCGTCATATCATATACCCGGATTGCAATAAGGAGTTGTTTTGTATCAAGGATAGC

TGCATATCTAAGAAGCTCTCGGAGCTATTCAAGAAGGGTAATTCCTTGTACTCTAAGATAAGTCACCAGGTACTGGATTG

TCTTAAGAGAGTCAATGGGAAATTAGGTCTGGGCACAGATCTTACCCATGGTCTGAAGGAGGGTATCCTTGACCTAGGGT

TGCACATGCATAGCTCTCAATGGTTCGAGACTTTCCTGTTCTGGTTCACTATCAAGACAGAGATGAGATCAATGATCAAA

GAACAGTCCCATATATGCCACAAGAGGAGGTATAACCCGACTTTTGTGTCGGGGGATGCATTCGAGGTGCTCGTATCACG

AGACCTCGTTGTGATAATTGATAAGAATACCCAGTATGTCTTCTACTTAACATTTGAGCTGGTCCTTATGTACTGTGATG

TCATAGAGGGCAGACTTATGACGGAGACAGCCATGGCCATAGACCAGAGATATTCAGAACTTCTAAGCCGGGTCAGATAC

TTGTGGGATCTTATTGATGGGTTCTTCCCAACACTAGGTAACACCACATACCAAATTGTTGCTCTGCTTGAACCATTATC

GTTGGCTTATCTTCAACTTCAGGATGTTACTCTGGAGTTAAGGGGTGCTTTTTTAGACCATTGCTTCAAAGAACTCTATG

AGATACTGGAGCATTGCGGCATTGACACAGAAGGCACCTACAATTCCATCACTGAAGGATTGGATTACGTATTCATCACC

CACGATATACATCTAACTGGGGAGATTTTTTCATTTTTTCGGAGTTTCGGACACCCCCGTCTAGAAGCAGTCACCGCTGC

AGAAAATGTCAGGAAACATATGAACCAACCGAAGGTGATCAGCTATGAGACTATGATGAAAGGGCATGCAGTATTTTGCG

GGATAATCATAAATGGTTTTAGGGACCGACATGGCGGCAGCTGGCCCCCTGTTGCATTGCCAGAACATGCTTCTGCTGTG

ATCCGGAATGCGCAGGCATCAGGTGAGGGACTGACTCATGACCTGTGTATAGACAACTGGAAATCTTTTGTTGGATTCAA

ATTTGGCTGCTTTATGCCACTCAGCCTAGATAGTGATTTGACCATGTACCTCAAAGACAAGGCATTGGCTGCACTGAAGA

ATGAGTGGGATTCAGTTTACCCGAAAGAATACCTCCGTTATAACCCACCTAGAGGGACAGAGTCAAGGCGACTGGTGGAG

GTGTTCCTGAATGACTCCAGCTTTGATCCTTATAACATGATAATGTACGTGGTGAATGGCTCCTACCTTAAAGACCCTGA

GTTTAATCTCTCATACAGCCTAAAAGAGAAGGAGATAAAGGAAACAGGGCGGTTGTTTGCCAAAATGACCTATAAGATGC

GGGCCTGTCAGGTAATCGCTGAAAATCTGATATCGAATGGTGTTGGGAAGTATTTCCGAGACAATGGGATGGCAAAGGAC

GAGCATGACCTAACAAAAGCCCTTCACACCCTGGCAGTCTCAGGTGTTCCCAAGAACAACAAAGATAACCACCGAGGTGG

GCCCCCCAGAAGGACAACAAGCCGAGGGGTGAGATCAAGCCAAGACACCAAAACACAAAATCGAGACAATGCCCAAGGGG

GGCCTGTGTACAACTATTTGCGATGCCAACCAATCAGCCCTGATCAGGGTGAGTCATACGAGACGGTTAGTGCTTTCATC

ACTGCCGACCTTAAGAAGTATTGCCTGAATTGGAGATACGAGACAATCAGCATATTTGCACAGAGGCTGAATGAAATATA

TGGACTACCATCCTTCTTTCAATGGTTACACAGGATATTGGAAAAATCCGTACTCTACGTCAGTGACCCACATTGCCCTC

CCGACCTAGATAATCATATTCCTCTGGACAGTGTCCCTAATTCCCAAATATTCATTAAGTACCCAATGGGCGGAATAGAA

GGTTATTGCCAAAAACTATGGACAATCAGTACTATACCATACTTGTATCTAGCAGCCTATGAGAGCGGAGTAAGAATCGC

CTCACTAGTACAGGGTGACAATCAGACAATCGCAGTGACAAAAAGAGTTCCAAGTTCTTGGCCTTATTCACTAAAAAAGA

GGGAGGCATCCAAAGCCGCTCAAAATTACTTCGTAGTCTTAAGGCAAAGATTGCACGATGTAGGTCATCACTTAAAGGCT

AATGAAACCATAGTATCTTCTCACTTTTTTGTGTATTCCAAAGGGATTTATTATGACGGGCTGCTAGTCTCACAATCACT

AAAGAGCATCGCCAGATGTGTCTTCTGGTCCGAGACTATAGTGGATGAAACCAGAGCGGCTTGCAGCAATATTGCAACAA

CTGTCGCTAAGAGTATAGAGAGGGGTTATGATAGGTACCTTGCATATTCTTTGAATATCCTCAAGATTTTCCAACAGATC

CTTATATCCCTTAACTTCACTATTAACACAACAATGACTCAGGATGTCGTGGCACCAATCATCGAGAACGGTGATTTGCT

GATAAGAATGGCACTCTTGCCAGCACCCATCGGAGGTCTCAATTATCTTAACATGAGCAGGTTATTTGTGAGAAATATCG

GTGACCCGGTCACCTCCTCTATAGCAGACCTGAAGAGGATGATAGATGCTGGGCTAATGCCAGAAGAAACATTGCATCAA

GTAATGACCCAGACCCCGGGAGAATCATCCTACCTCGATTGGGCAAGTGACCCTTATTCTGCCAACCTACCCTGCGTACA

GAGTATAACTCGCCTTCTCAAAAACATCACTGCACGATATATTTTAATCAGCAGCCCAAACCCGATGCTGAAAGGACTGT

TTCATGAGGGGAGTAGAGATGAAGACGAGGAGCTTGCAAGTTTCCTAATGGATCGGCATATAATTGTTCCAAGAGCTGCA

CATGAAATTTTAGACCATAGCATAACCGGGGCAAGAGAAGCTATAGCCGGGATGTTGGACACCACCAAGGGTCTTATTAG

AACAAGTATGAAACGGGGTGGCCTCACCCCTCGAGTCTTGGCCCGCCTTTCCAATTATGATTATGAACAATTCAGATCCG

GGATAACACTATTAACAAAGAAAGGGCTGTGTTATCTCATTGACAAGGACTCATGCTCGGTGCAGCTCGCTATAGCTCTG

AGGAGCCACATGTGGGCTAGGTTAGCTCGCGGGAGACCAATCTATGGATTGGAGGTGCCCGATATATTAGAATCGATGAA

TGGCTACCTTATCAAACGCCACGAGTCTTGTGCTATCTGTGAAACGGGCTCAAGTCACTACGGATGGTTTTTCGTCCCTG

CAGGGTGCCAGCTTGACGATGTGTCTAGAGAGACTTCGGCTCTTCGTGTACCTTATGTCGGATCAACAACTGAGGAAAGG

ACAGACATGAAACTTGCATTTGTTAGGTCCCCAAGTAGATCCCTCAAATCAGCAGTTAGAATTGCAACAGTTTACTCATG

GGCTTACGGGGATGATGAGAAATCATGGAGTGAAGCCTGGATGCTAGCTAGGCAGAGAGCCAATATCACTTTAGATGAAT

TGAGAATGATCACTCCAGTCTCCACATCCACTAACTTAGCCCACCGGTTGAGGGATCGGAGTACCCAGGTGAAATACTCG

GGGACATCCCTTGTGAGGGTTGCAAGATACACAACCATCTCCAATGATAACTTGTCATTTGTGATATCTGAGAAAAAAGT

GGATACCAACTTCATTTATCAGCAAGGGATGCTGCTCGGTCTTGGGATCCTTGAGAACCTCTTTAGGCTAGAGGCCACCA

CCGGAGTATCCAACACAGTGCTACACCTGCACGTGGAAACAGAATGTTGTGTTGTACCTATGGTGGATCACCCAAGGATT

CCGAGTCTCCGTAATATAAAAGTTACGGATGAGCTATGCACAAACCCTTTGATCTACGACAGGTCTCCCATCATAGAGCA

CGATGCAACCCGACTATACTCACAGAGCCACAGGAGACATTTGGTGGAGTTTGTTACCTGGTCAACAAGCCAGCTTTACC

ATATACTGGCTAAGTCTACAGCAATGTCCATGATTGAATTGATCACGAGATTCGAGAAAGATCACATGAATGAAATAGCC

GCCCTGATTGGCGATGACGACATCAACAGCTTCATCACAGAATTTTTGCTAGTGGAGCCTAGATTATTTATAGTTTACCT

TGGTCAGTGTGCTGCCATCAATTGGGCCTTTGATATACATTATCATCGGCCCTCGGGCAAGTACCAGATGGGGGAACTCC

TCTACTCTCTGCTCTCTCGGATGAGCAAAGGAGTATATAAGATCTTCACCAATGCTTTGAGCCACCCCAAAGTTTACAAG

AAATTTTGGCGAAGTGGTGTCATTGAGCCGATTCATGGCCCATCCCTGGATACACAGAATTTACATGTCACTGTCTGTGA

CATGATATATGGGTCCTATGTCACCTACCTGGATCTTTTGCTGAATGATGAGCTAGATGATTACCCGTATTTGCTCTGCG

AGAGTGATGAGGATGTGGTCACAGACAGGTTCGACAACATTCAAGCCAAACATCTCTGTGTATTAGCCGATGTATATTGC

AGCTCCAAGAGGTGTCCTTCAATAATCGGGATGTCTCCTATAGAAAAATGTACCATTCTCACACATTACATCAAGGGAGA

ATCAGTACAATCCCCGTCTGGAACCTCATGGAACACTGATCCCCTTGTAGTAGATCATTACTCATGCTCTCTGACCTACC

TTCGCCGAGGTTCCATCAAACAAATCAGGTTGAGGGTGGATCCTGGGTTTGTATTCGAGGCGTTGACAGACGTCGACTTT

AAACAACCTCGCAAAGCTAAGTTAGACATATCGGTCGTTGGGTTGACTGATTTCTCTCCCCCTTGCGACAACGTCGGTGA

TTTTCTAGGGACTATCAACACATTGAGGCACAATCTGCCTGTCACCGGAACCGGGGTCTCGAACTATGAAGTCCACGCTT

ATCGTAGAATTGGCCTGAATTCATCAGCATGTTACAAAGCCGTAGAGATCTCCACGTTAATCAAGCCATCCTTAGAAGTC

GGAGAACACGGATTGTTCTTAGGAGAAGGTTCCGGTTCAATGCTGGCTGCGTACAAGGAAGTTCTTAAATTAGCAAACTG

TTATTACAACAGCGGGGTAACAGCGGAGGGCAGAGCCGGACAGAGGGAAATATCTCCCTATCCTTCAGAGATGAGCCTAG

TAGAGAATCAAATGGGGATAGAGCGGAGCGTTAAAGTGCTGTTCAACGGCAAACCTGAAGTGACTTGGGTGGGGTCCACC

GATTGCTACAAGTACATAATCAGTAACATCCAAACCTCTAGTCTAGGTTTCATACATTCAGATATTGAGACACTCCCAAC

CAAAGATGCTGTTGAGAAATTAGAAGAATTTGCCTCTATCCTATCCTTATCCTTGATTTTGGGGAAAATCGGCTCTATTA

CAGTTGTCAAAATTATGCCCATTAGCGGGGATTTTACCCAAGGCTTCATAGCCTATGCTATCCAATATTTCAGGGAGAGT

CTGCTTGCCTACCCGAGATACAGTAACTTCATCTCGACTGAGTGTTACCTTATTATGATAGGATTAAAGGCCAATCGATT

GATAAACCCAGAGGCCATTAAGCAAAGCATAATCAGAGCGGGGGTCAGGACTGCACCAGGACTTGTGAGCCATATATTAT

CAGAGAAACAGAAAGGTTGTATTCAATCTTTCTTGGGTGATCCTTATATCCAAGGAGACTTCAACAAGCACCTTAAATCT

CTAACCCCTATTGAGAAAATCCTGGTAAATTGTGGTCTCTCGATCAATGGCACAAAAATCTGTAGAGATCTAATCCACCA

TGATATCGCCTCCGGTCCAGACGGTCTTATGAGCTCTACAATTATTTTATATAGGGAATTGGCTCATTTCAAAGATAATA

TAAGGAGTCAGCACGGAATGTTCCACCCCTACCCGGTACTGGCCAACAGCAGGCAACGTGAATTAATCCTTCGAATAGCC

AAGAAATTCTGGGGGTATGTCTTGCTATATTCTGATGACCCGGCACTAATCAGACAAACAATCAAGAACTTGAAACGGAA

TCATCTAACCTTTGACTTACATAGTAACCCGTTTATTAAAGGGCTATCCAAAGCTGAGAAACTGCTGGTGCGGACAAGTT

CACTAAGAAGGGAATGGTTGTTCACTCTAGATACGAAAGAAGTGAAAGAGTGGTTCAAATTGGTAGGTTACAGTGCACTC

ATCAGAGGCTAGTTAGCTATACATCTGCCCCCTCCTCCTCCACCATGAGACCCTACTGACGATCCAGAAGATTAAAGAAA

ACTACATATTGGATAAGTATCTATTCCCAGTTTTGTCTGGT

>KY967609.1/goat/Pakistan/2015

ACCAAACAAAGTTGGGTAAGGATAGATCTTACAATAACTATGGACTGGCAAACTTAGGAGTAAAGATCCTACTGTCGGGG

GGAGGAGGAGGAGCAAGATCTCTGATTATGGCGACTCTCCTTAAAAGCTTGGCATTGTTCAAGAGGAACAAAGACAAAGC

GCCGACGGCATCAGGTTCAGGAGGGGCCATCCGGGGGATTAAGAATGTTATCATAGTCCCGATTCCCGGAGACTCGTCCA

TCATTACCCGTTCAAGACTGCTCGACAGGCTTGTCAGATTGGCCGGAGATCCTGACATCAACGGGTCAAAGCTGACCGGC

GTGATGATCAGCATGTTATCTTTGTTCGTAGAGTCACCCGGACAACTGATACAGCGAATCACAGATGATCCAGATGTTAG

TATCCGCCTTGTTGAGGTAGTTCAAAGTACTAGGTCCCAGTCCGGGTTGACCTTTGCATCACGTGGTGCTGACTTGGACA

ACGAGGCAGACATGTATTTTTCAACTGAGGGGCCCTCGAGTGGAGGTAAGAAAAGGATCAACTGGTTTGAGAACAGAGAA

ATAATAGACATAGAGGTGCAGGATCCAGAAGAGTTCAATATGTTGTTAGCCTCCATACTAGCACAAGTCTGGATCCTTCT

AGCCAAGGCTGTTACGGCACCGGATACGGCAGCTGACTCAGAACTGAGAAGGTGGGTTAAATACACACAACAAAGGAGAG

TGATTGGGGAATTTCGCCTTGACAAAGGGTGGCTGGACGCAGTCCGCAACAGAATTGCAGAAGATCTATCACTCCGGCGG

TTCATGGTATCTCTCATACTTGACATCAAGAGGACTCCCGGCAACAAGCCAAGGATTGCAGAAATGATCTGCGACATTGA

CAACTATATTGTCGAAGCCGGACTCGCCAGTTTCATCCTTACTATCAAGTTTGGTATCGAAACCATGTATCCTGCATTAG

GGCTCCACGAGTTTGCCGGGGAATTGTCCACTATTGAGTCCTTGATGAACTTGTATCAACAGCTAGGAGAGGTTGCACCC

TACATGGTAATTCTAGAGAACTCAATTCAGAACAAGTTTAGTGCAGGAGCTTATCCCCTCCTCTGGAGCTATGCGATGGG

TGTCGGGGTTGAGCTAGAGAACTCAATGGGGGGCTTGAACTTTGGCAGGTCATATTTTGACCCTGCCTATTTTCGTCTCG

GACAGGAGATGGTCAGAAGATCTGCGGGAAAGGTCAGCTCTGTGATTGCGGCTGAGCTCGGCATCACAGCAGAGGAAGCT

AAACTAGTCTCGGAAATCGCCTCGCAGGCTGGGGACGAAAGAACCGCTAGAGGGACTGGGCCTCGACAGGCGCAGGTCTC

CTTCCTCCAGCACAAAACAGGAGAGGGAGAGTCGCCCGCACCAGCGACCAGAGAAGGGGTCAAAGCTGCGATCTCGAACG

GATCTGAAGAGAGGGACAGAAAGCAAACACGCCCAGGAAGGTCCAGAGGAGAGACCCCCGGCCAACTGCTCCTAGAAATC

ATGCCAGAGGATGAGGTCTCGCGAGAGTCTGGTCAAAACCCTCGTGAGGCTCAAAGATCGGCTGAGGCACTCTTCAGGCT

GCAGGCCATGGCCAAGATCCTGGAGGACCAGGAGGAGGGAGAAGACAACAGTCAGATCTACAACGACAAGGATCTCCTCG

GCTAAGCAGACGCGCCCTCTATCGAAATTAGTGACAATACATCTCCTGCCAGTATTATAAAAAACTTAGGACCCAGGTCC

AGGCAAGCACACACCGACACTCCAACCTGTCGAGCGGAGACCACCGATGGCAGAAGAACAAGCATACCATGTCAACAAGG

GGCTGGAATGTATCAAGTCTCTCAAAGCCTCTCCCCCGGATCTATCCACCATCAGAGACACCATCGAGAGCTGGAGAGAG

GGGCTTAGCCCCTCGGGCCGTGCAACACCGGATCCTGATACGTCCGAGGGAGATCATCAGAATATCAACCAATCATGCTC

ACCAGCAATCGGACCAAGCAAAGTCTACCTGTCTCCTGAAGATAATCTCGGATTTAGAGAGATCACTGGCAACGACTGTG

AGGCTGGGCTCGGAGGAGTCCAGGGAAAAGGATCCAACTCTCAAGTACAGCGTTACTATGTTTATAGCCACGGGGGTGAA

GAGATTGAAGGACTCGAGGATGCTGACTCTCTCGTGGTTCAAGCAGACCCTCCAGTTGCTAACATCTTCGATGGAGGAGA

GGATGGATCTGACGACAGCGATGTGGACTCTGGCCCAGATGATCCCGGCAGAGATACTCTACATGACCGGGGATCTGTTG

CCGGCAATGATGTCGCTAGGTCCACGGATGTCGAAAAGCTAGAAGGTGCTGACCTTCAAGAAGTTCTTAACTCCCAGAAA

GGCAAAGGAGGAAGATTCCAAGGCGGGAAAACCTTACGAGTCCCGGAAATACCCGATGTCAAGCACTCCAGACCATCAGC

TCAATCAATTAAAAAGGGCACAGACGGGAGCTCAGTCTCATCTGGAACGGTGACAGAGTGTTTATCGATAAGTGGTGCAA

CCCAAGCTGTGCCAGAGTCAGGATGGGAGTCATCAGAGCAAAATGCGTCTGTGGGGAGTGTCCTCAAATCTGCGAGGAGT

GCAAAGACGATCCAGGGGTCGACACAAGAATCTGGTACCATAGCATCACTGACTCAGCCTAAAGAGAATGACTCCGAGTA

TGAGTATGAGGATGACCTATTTACAGAGATTCAGGACATCCGTGCAAGCATTGCCAAGATCCATGATGACAACAAGACTA

TCCTCTCAAAGCTTGATTCTATACTGTTATTGAAAGGAGAAGTCGACACTATCAAGAAACAAATCAGCAAGCAGAATATA

AGTATATCCACCATTGAGGGCCATCTCTCCAGTATAATGATAGCCATCCCGGGCTTTGGGAAGGATATCAAGGACCCAAC

ATCCGAGGTTGAGTTGAACCCAGATTTAAGACCTATAATAAGCCGTGACTCTGGCAGAGCTCTCGCGGAGGTCCTCAAGA

AACCCGCTGTGGATAGGTCTCAGAAAACTGGAACCAAAGCCAACTCCAGCTCAAAGGGTCAGCTTCTTAAGGATCTCCAG

CTAAAACCTGTCGACAAGCAGGCAAGCTCTGCAATCGGGTTTGTCCCGTCCGACCATGAATCATCCAGAAATGTCATCCG

CTCCATAATCAAGTCGAGCAAGCTAAACATTGATCACAAGGACTATCTTCTAGATTTACTGAATGATGTGAAAGGCTCCA

AGGATCTTAAGGAATTCCACAAGATGCTAACAGCAATTCTCGCCAAGCACCCGTAACACATCTCCCAGTCACCATCTCAT

ACTCGGCTGAAGACATCCTTTCAATCAGGCTATTACAAAAAACTTAGGAGCAAGGGCAACTGAGCTTCGCAGACAAGATG

ACAGAGATCTACGACTTCGATAAATCAGCATGGGATGTCAAAGGGTCAATTGCCCGCATAGAACCCACCACCTACCACGA

TGGCCGCCTGATACCCCAGGTAAGAGTCATCGATCCTGGTCTGGGAGACAGAAAAGATGAGTGCTTCATGTACCTGTTTC

TCCTAGGAGTGATTGAGGATAACGACCCCCTGTCTCCCCCCGTCGGGAGAACCTTTGGCTCTTTACCTCTAGGGGTCGGT

AGGTCAACTGCTAAGCCAGAAGAACTACTAAGGGAGGCCACAGAACTAGATATAGTGGTGAGGCGCACTGCAGGAGTAAA

TGAGAAACTGGTATTTTACAACAACACACCGCTGTCCCTGTTAACGCCCTGGAAGAAAGTTCTGACAGCCGGAAGTGTGT

TTAGCGCTAACCAGGTTTGCAATGCAGTCAACCTAGTCCCGCTTGATACTCCCCAGAGATTCAGGGTTGTGTACATGAGC

ATAACTAGATTATCAGACAATGGTTACTATAGTGTGCCCAGAAGAATGCTGGAGTTCCGCTCAGCCAATGCAGTAGCCTT

CAATATCTTGGTTACACTAAGAATTGAAAATGGCACAAACCCTAAAAGATACATAGTCGGCTCATGGGAGAATCCAGAGG

TCACATTTATGGTACACGTGGGGAACTTTAGAAGAAAGAAGAACGAAGTATACTCTGCTGATTACTGCAAAATGAAGATT

GAAAAGATGGGTTTAGTTTTTGCCCTGGGAGGAATAGGTGGAACAAGTCTCCATATTAGGAGCACCGGGAAAATGAGCAA

AACCCTCCATGCACAGCTGGGGTTCAAGAAAATCTTATGTTACCCCCTAATGGATGTTAATGAGGATCTTAACCGATATC

TCTGGCGGGCAGAGTGCAGAATAGTCAAAATCCAAGCCGTCTTACAGCCATCAGTACCCCAAGAATTCCGTGTCTACGAT

GATGTCATCATCAACGATGATCAAGGCTTGTTCAAGATCTTGTAGTTCATTTGCAAAGGAATGAACCCACCCCTAGGATG

GGGCCCCACCCTCCCCCG-TCAAAAAACACACTCCAAGGCTCCTCGGCACAGCGCCAACACAGCTACTCCCGCGATGGAA

CGAGGTCGGCC-CCCCAACAAATCCGGTCCAGGT-GGGGGGCCCCCCCCCTGACCAACCGGGCGCTGACCGAGCAGGCCG

GCGAGAACCCCCCAAGAGGCCCCGACAAAAACCGACCACCCCATCCCCCC-AAAAATGTCCCCC-AACACGCCCGAGGCT

AGCCACCGGGCACCCCCAGACCCGCCACCGGCGAGACAGAACCCCAAGGGGGGACCCTCCAAACCAAAGGGATCACCGGG

GTAGGCTCCGCTGAACCAGCACAGACCCGACCCAAGCAAGACGGACCCCCAACACAAATCTGCCCAGCACACCCCACTCA

TCATCAAACAAAATTTAGGGGCCAAGTCCATACAAAAGGCCCCATTCCCTCAGGCGAAGGGACGCAAAAGGGAAGGAAAC

AAGACCGCCCCAGAAGGAGGACACCCAACCACCGAAACACCCAGCCCCAGGGAAGGAG--CCCCCCCCCACATGGCGAAA

GAACGGGCGAGAGACCCCGAGAAGAACCCGACCGGGAGCCAGGAGCCGGCGACGCCGGCGGCAAACAAGCCGGGCAAACA

CCCGACCCCACCAGAGGAGCCGAGGAGAGCCCGACCCCGCACGAACCAAACAGCAGGGAGCCCAACCCCACCAAAAAAAG

AGGCGAGGCCCC-CCT-----CCTCCCCGCACACAACCCGCAGACAGACCGAGAGAGGACCACCTCCCAACAAGACAGGC

GGCTCCCTCGCAGCCGAGCCGCAGCTCCCCAGAACAGGACACCCCCGTACACTCAAAACACCCCCCCGCCCCCCGCCACC

GGGACAGGGGCTCCGCCACCACAGACCAAAGCCACCGAGAGCAAGCCAGAGGTCCCCCCCAGTCCCGCCCGGCGCCCCCC

CGAAAAGGCCCCGGCAAGGCCCCCCAACCCGACGTCCCCCGCACGCACCGAAAGAGGGACCCGCCCGAGAACGTGCGGCA

CATCCATGCGTAAACATTATGACACGGGTCGCAATCTTGACATCTCTGTTTCTTCTCCCAAATGTTGTTGCGTGCCAGAT

TCACTGGGGCAATCTATCCAAGATCGGGATTGTGGGAACAGGGAGTGCCAGCTACAAGGTGATGACTAGGCCGAGCCACC

AGACTCTGGTTATAAAGTTAATGCCAAATATAACAGCCATCGACAATTGTACAAAGTCAGAGATTGCAGAGTACAAGAGA

TTGCTGATCACAGTGTTAAAGCCTGTAGAGGATGCTCTGTCAGTGATAACCAAGAATGTGAGACCAATTCAAACTCTGAC

ACCTGGGCGTAGAACTCGCCGTTTTGCTGGAGCTGTTCTGGCCGGAGTAGCACTTGGAGTTGCGACAGCCGCTCAGATAA

CTGCAGGAGTCGCCCTTCATCAATCACTGATGAACTCTCAAGCAATTGAGAGTTTAAAAACCAGTCTCGAGAAGTCGAAT

CAGGCAATAGAAGAAATCAGACTTGCCAATAAGGAGACCATACTGGCAGTACAGGGCGTCCAGGACTATATCAACAATGA

GCTTGTCCCTTCTGTTCATAGAATGTCATGCGAGCTTGTAGGTCACAAGCTCGGCCTCAAGCTCCTTAGGTACTACACCG

AGATCCTGTCCATATTCGGGCCCAGCCTTCGAGACCCGATAGCTGCCGAAATATCAATCCAGGCACTCAGTTATGCATTA

GGTGGAGACATTAATAAAATCCTGGACAAGCTTGGGTATAGCGGTGGGGATTTCCTTGCCATCCTAGAAAGCAAGGGAAT

AAAGGCCCGGGTCACATATGTGGATACGAGAGATTACTTTATAATTCTTAGCATCGCCTACCCAACCTTATCTGAGATCA

AGGGAGTGATAGTTCACAAGATAGAAGCTATAACATACAACATTGGGGCACAGGAGTGGTATACTACTATCCCTAAATAT

GTAGCCACTCAGGGGTATCTGATATCGAACTTTGATGAGACGTCATGCGTATTCACTCCAGAGGGGACAGTTTGCAGCCA

GAATGCGTTGTACCCAATGAGCCCATTGCTCCAGGAATGTTTTAGGGGGTCAACAAAATCGTGCGCCAGAACCCTAGTTT

CAGGGACCATAAGTAATAGATTTATCCTATCGAAAGGGAATCTGATTGCAAATTGTGCGTCAGTTTTGTGCAAATGCTAC

ACAACGGAGACAGTTATCAGCCAAGATCCTGACAAACTACTAACTGTTGTAGCATCCGACAAGTGTCCTGTAGTTGAGGT

GGACGGAGTGACAATACAGGTCGGCAGTCGAGAGTATCCGGATTCTGTATACTTACACAAAATAGACTTAGGTCCAGCCA

TCTCCCTAGAAAAACTGGATGTAGGCACCAATTTAGGCAATGCAGTCACAAGACTGGAGAATGCAAAGGAGCTCCTAGAT

GCATCAGACCAAATACTGAAGACTGTTAAAGGGGCACCTTTCGGTGGGAATATGTACATAGCACTGGCAGCTTGCATTGG

AGTATCCCTAGGGCTTGTCACATTAATATGCTGCTGTAAGGGGAGGTGTAAGAACAAGGAAATCCCTATCTCCAAAATCA

ACCCAGGGCTCAAACCCGACCTGACCGGGACCTCAAAGTCGTACGTGAGATCACTGTAGTCTGAATCACTTGAATCATCC

AGCATCACACACATACAGGTACGACGCAGGCAGTCTGAGGACGCAGGAAGCCCAGCCTCCGATCACCCACCTGACCCTAC

TCCACGCTCCACCACACATTAGTCATCAAACAAAACTTAGGACGAAAGGTCAATCACCATGTCCGCACAAAGGGAAAGGA

TCAATGCTTTCTACAAAGACAATCCTCACAATAAGAACCATAGGGTGATCCTGGATAGAGAACGCCTGGTCATTGAAAGA

CCCTATATCTTGCTTGGAGTCCTGCTGGTAATGTTCCTGAGTCTAATCGGACTGCTGGCCATCGCAGGGATCAGGCTTCA

CCGGGCCACCGTTGGGACTTCAGAGATCCAGAGTCGGCTGAATACCAATATTGAGTTGACCGAATCTATTGATCACCAAA

CTAAGGATGTCTTAACCCCCCTTTTTAAAATCATTGGCGATGAAGTCGGCATCAGAATTCCACAGAAATTCAGTGATCTT

GTCAAGTTCATCTCCGATAAGATTAAATTCCTTAACCCTGATAGAGAGTATGATTTCAGGGATCTCCGGTGGTGTATGAA

TCCCCCAGAGAGAGTCAAAATTAATTTTGATCAGTTTTGTGAGTACAAAGCTGCGGTTAAGTCAATTGACCATATATTTG

AGTCACCACTCAACAAGTCAAAAAAGCTGCAATCACTGACTCTTGGGCCCGGAACAGGCTGTCTAGGCAGGACAGTAACA

AGAGCTCATTTCTCAGAACTTACACTGACCTTAATGGACCTGGATCTAGAGATGAAGCACAACGTGTCCTCAGTGTTTAC

CGTTGTTGAAGAGGGATTATTCGGAAGAACGTATACCGTCTGGAGATCCGATGCCAGGGATCCGGGCACCGATCTAGGTA

TCGGCCATTTTTTAAGAGTATTTGAGATTGGATTGGTAAGAGATCTCGGGCTGGGTCCCCCTGTTTTCCATATGACCAAC

TATCTCACAGTGAACATGAGTGATGACTATCGGAGATGTCTTTTAGCGGTAGGGGAGTTGAAGTTGACAGCCCTATGCAC

CTCATCTGAGACTGTGACACTGAGTGAGAGAGGAGTTCCAAAGAGGGAGCCTCTTGTGGTTGTGATACTTAACCTGGCTG

GGCCCACTCTAGGGGGCGAACTATACAGTGTCTTGCCKACCTCTGATCTCATGGTGGAGAAACTCTACCTATCTTCACAT

AGAGGGATTATCAAAGATGACGAGGCCAATTGGGTAGTGCCGTCTACCGATGTTCGTGATCTTCAAAACAAAGGTGAATG

TCTGGTGGAAGCATGCAAGACCCGACCTCCTTCATTCTGCAATGGCACAGGATCAGGCCCGTGGTCAGAGGGGAGAATCC

CTGCTTACGGGGTGATCAGGGTCAGTCTTGACTTAGCTAGTGACCCGGGTGTATTTATCACTTCAGTGTTTGGCCCACTG

ATACCTCACCTATCCGGCATGGATCTTTACAACAACCCGTTTTCAAGAGCTATATGGTTGGCTGTACCTCCTTATGAACA

GTCATTTCTAGGAACGATAAATACCATTGGATTCCCCAACAGAACAGAGGTTATGCCGCACATCTTGACCACAGAGATCA

GAGGCCCTCGGGGTCGTTGCCATGTTCCCATAGAATTGTCCCGCAGGGTTGATGACGATATCAAGATCGGGTCCAACATG

GTCATATTGCCGACGATGGATTTGAGGTATATTACAGCCACTTATGATGTTTCCAGGAGCGAGCATGCAATCGTGTACTA

TATCTATGACACAGGTCGCTCATCATCTTACTTCTACCCAGTTCGACTGAATTTCAAAGGCAATCCTCTCTCTTTGAGGA

TAGAGTGCTTCCCTTGGCGTCATAAGGTGTGGTGCTACCATGATTGTCTTATATACAACACCATAACAGATGAAGAAGTC

CATACGAGAGGGCTGACCGGTATAGAGGTAACATGTAATCCAGTCTGAGCAGAGCTACGACCGTCACTCAGGCAGTCCCT

TGAGTCGCCACCGAGCCCAAGCAGCACAGCCTGGGACACTCAACAGCACAGCCCAGCCAACAACGTTATAAAAAACTTAG

GAGCCAGGGTTGTAGGGGCCATGGACTCTCTATCAGTCAATCAGGTCTTGTACCCTGAGGTTCATCTAGACAGCCCTATT

GTCACAAACAAACTAGTTGCCATCCTTGAGTACTCGGGTATCGACCATAACTATGTTCTTGAAGACCAGACTCTTGTCAA

GAATATTAGGTATAGGCTGGGGTGCGGTTTTTCAAATCAAATGATCATCAATAACAGGGGGGTGGGTGAAACAGTCAATT

CTAAACTTAAAAGTTACCCCCATAATCGTCATATCATATACCCGGATTGCAATAAGGAGTTGTTTTGTATCAAGGATAGC

TGCATATCTAAGAAGCTCTCGGAGCTATTCAAGAAGGGTAATTCCTTGTACTCTAAGATAAGTCACCAGGTACTGGATTG

TCTTAAGAGAGTCAATGGGAAATTAGGTCTGGGCACAGATCTTACCCATGGTCTGAAGGAGGGTATCCTTGACCTAGGGT

TGCACATGCATAGCTCTCAATGGTTCGAGACTTTCCTGTTCTGGTTCACTATCAAGACAGAGATGAGATCAATGATCAAA

GAACAGTCCCATATATGCCACAAGAGGAGGTATAACCCGACTTTTGTGTCGGGGGATGCATTCGAGGTGCTCGTATCACG

AGACCTCGTTGTGATAATTGATAAGAATACCCAGTATGTCTTCTACTTAACATTTGAGCTGGTCCTTATGTACTGTGATG

TCATAGAGGGCAGACTTATGACGGAGACAGCCATGGCCATAGACCAGAGATATTCAGAACTTCTAAGCCGGGTCAGATAC

TTGTGGGATCTTATTGATGGGTTCTTCCCAACACTAGGTAACACCACATACCAAATTGTTGCTCTGCTTGAACCATTATC

GTTGGCTTATCTTCAACTTCAGGATGTTACTCTGGAGTTAAGGGGTGCTTTTTTAGACCATTGCTTCAAAGAACTCTATG

AGATACTGGAGCATTGCGGCATTGACACAGAAGGCACCTACAATTCCATCACTGAAGGATTGGATTACGTATTCATCACC

CACGATATACATCTAACTGGGGAGATTTTTTCATTTTTTCGGAGTTTCGGACACCCCCGTCTAGAAGCAGTCACCGCTGC

AGAAAATGTCAGGAAACATATGAACCAACCGAAGGTGATCAGCTATGAGACTATGATGAAAGGGCATGCAGTATTTTGCG

GGATAATCATAAATGGTTTTAGGGACCGACATGGCGGCAGCTGGCCCCCTGTTGCATTGCCAGAACATGCTTCTGCTGTG

ATCCGGAATGCGCAGGCATCAGGTGAGGGACTGACTCATGACCTGTGTATAGACAACTGGAAATCTTTTGTTGGATTCAA

ATTTGGCTGCTTTATGCCACTCAGCCTAGATAGTGATTTGACCATGTACCTCAAAGACAAGGCATTGGCTGCACTGAAGA

ATGAGTGGGATTCAGTTTACCCGAAAGAATACCTCCGTTATAACCCACCTAGAGGGACAGAGTCAAGGCGACTGGTGGAG

GTGTTCCTGAATGACTCCAGCTTTGATCCTTATAACATGATAATGTACGTGGTGAATGGCTCCTACCTTAAAGACCCTGA

GTTTAATCTCTCATACAGCCTAAAAGAGAAGGAGATAAAGGAAACAGGGCGGTTGTTTGCCAAAATGACCTATAAGATGC

GGGCCTGTCAGGTAATCGCTGAAAATCTGATATCGAATGGTGTTGGGAAGTATTTCCGAGACAATGGGATGGCAAAGGAC

GAGCATGACCTAACAAAAGCCCTTCACACCCTGGCAGTCTCAGGTGTTCCCAAGAACAACAAAGATAACCACCGAGGTGG

GCCCCCCAGAAGGACAACAAGCCGAGGGGTGAGATCAAGCCAAGACACCAAAACACAAAATCGAGACAATGCCCAAGGGG

GGCCTGTGTACAACTATTTGCGATGCCAACCAATCAGCCCTGATCAGGGTGAGTCATACGAGACGGTTAGTGCTTTCATC

ACTGCCGACCTTAAGAAGTATTGCCTGAATTGGAGATACGAGACAATCAGCATATTTGCACAGAGGCTGAATGAAATATA

TGGACTACCATCCTTCTTTCAATGGTTACACAGGATATTGGAAAAATCCGTACTCTACGTCAGTGACCCACATTGCCCTC

CCGACCTAGATAATCATATTCCTCTGGACAGTGTCCCTAATTCCCAAATATTCATTAAGTACCCAATGGGCGGAATAGAA

GGTTATTGCCAAAAACTATGGACAATCAGTACTATACCATACTTGTATCTAGCAGCCTATGAGAGCGGAGTAAGAATCGC

CTCACTAGTACAGGGTGACAATCAGACAATCGCAGTGACAAAAAGAGTTCCAAGTTCTTGGCCTTATTCACTAAAAAAGA

GGGAGGCATCCAAAGCCGCTCAAAATTACTTCGTAGTCTTAAGGCAAAGATTGCACGATGTAGGTCATCACTTAAAGGCT

AATGAAACCATAGTATCTTCTCACTTTTTTGTGTATTCCAAAGGGATTTATTATGACGGGCTGCTAGTCTCACAATCACT

AAAGAGCATCGCCAGATGTGTCTTCTGGTCCGAGACTATAGTGGATGAAACCAGAGCGGCTTGCAGCAATATTGCAACAA

CTGTCGCTAAGAGTATAGAGAGGGGTTATGATAGGTACCTTGCATATTCTTTGAATATCCTCAAGATTTTCCAACAGATC

CTTATATCCCTTAACTTCACTATTAACACAACAATGACTCAGGATGTCGTGGCACCAATCATCGAGAACGGTGATTTGCT

GATAAGAATGGCACTCTTGCCAGCACCCATCGGAGGTCTCAATTATCTTAACATGAGCAGGTTATTTGTGAGAAATATCG

GTGACCCGGTCACCTCCTCTATAGCAGACCTGAAGAGGATGATAGATGCTGGGCTAATGCCAGAAGAAACATTGCATCAA

GTAATGACCCAGACCCCGGGAGAATCATCCTACCTCGATTGGGCAAGTGACCCTTATTCTGCCAACCTACCCTGCGTACA

GAGTATAACTCGCCTTCTCAAAAACATCACTGCACGATATATTTTAATCAGCAGCCCAAACCCGATGCTGAAAGGACTGT

TTCATGAGGGGAGTAGAGATGAAGACGAGGAGCTTGCAAGTTTCCTAATGGATCGGCATATAATTGTTCCAAGAGCTGCA

CATGAAATTTTAGACCATAGCATAACCGGGGCAAGAGAAGCTATAGCCGGGATGTTGGACACCACCAAGGGTCTTATTAG

AACAAGTATGAAACGGGGTGGCCTCACCCCTCGAGTCTTGGCCCGCCTTTCCAATTATGATTATGAACAATTCAGATCCG

GGATAACACTATTAACAAAGAAAGGGCTGTGTTATCTCATTGACAAGGACTCATGCTCGGTGCAGCTCGCTATAGCTCTG

AGGAGCCACATGTGGGCTAGGTTAGCTCGCGGGAGACCAATCTATGGATTGGAGGTGCCCGATATATTAGAATCGATGAA

TGGCTACCTTATCAAACGCCACGAGTCTTGTGCTATCTGTGAAACGGGCTCAAGTCACTACGGATGGTTTTTCGTCCCTG

CAGGGTGCCAGCTTGACGATGTGTCTAGAGAGACTTCGGCTCTTCGTGTACCTTATGTCGGATCAACAACTGAGGAAAGG

ACAGACATGAAACTTGCATTTGTTAGGTCCCCAAGTAGATCCCTCAAATCAGCAGTTAGAATTGCAACAGTTTACTCATG

GGCTTACGGGGATGATGAGAAATCATGGAGTGAAGCCTGGATGCTAGCTAGGCAGAGAGCCAATATCACTTTAGATGAAT

TGAGAATGATCACTCCAGTCTCCACATCCACTAACTTAGCCCACCGGTTGAGGGATCGGAGTACCCAGGTGAAATACTCG

GGGACATCCCTTGTGAGGGTTGCAAGATACACAACCATCTCCAATGATAACTTGTCATTTGTGATATCTGAGAAAAAAGT

GGATACCAACTTCATTTATCAGCAAGGGATGCTGCTCGGTCTTGGGATCCTTGAGAACCTCTTTAGGCTAGAGGCCACCA

CCGGAGTATCCAACACAGTGCTACACCTGCACGTGGAAACAGAATGTTGTGTTGTACCTATGGTGGATCACCCAAGGATT

CCGAGTCTCCGTAATATAAAAGTTACGGATGAGCTATGCACAAACCCTTTGATCTACGACAGGTCTCCCATCATAGAGCA

CGATGCAACCCGACTATACTCACAGAGCCACAGGAGACATTTGGTGGAGTTTGTTACCTGGTCAACAAGCCAGCTTTACC

ATATACTGGCTAAGTCTACAGCAATGTCCATGATTGAATTGATCACGAGATTCGAGAAAGATCACATGAATGAAATAGCC

GCCCTGATTGGCGATGACGACATCAACAGCTTCATCACAGAATTTTTGCTAGTGGAGCCTAGATTATTTATAGTTTACCT

TGGTCAGTGTGCTGCCATCAATTGGGCCTTTGATATACATTATCATCGGCCCTCGGGCAAGTACCAGATGGGGGAACTCC

TCTACTCTCTGCTCTCTCGGATGAGCAAAGGAGTATATAAGATCTTCACCAATGCTTTGAGCCACCCCAAAGTTTACAAG

AAATTTTGGCGAAGTGGTGTCATTGAGCCGATTCATGGCCCATCCCTGGATACACAGAATTTACATGTCACTGTCTGTGA

CATGATATATGGGTCCTATGTCACCTACCTGGATCTTTTGCTGAATGATGAGCTAGATGATTACCCGTATTTGCTCTGCG

AGAGTGATGAGGATGTGGTCACAGACAGGTTCGACAACATTCAAGCCAAACATCTCTGTGTATTAGCCGATGTATATTGC

AGCTCCAAGAGGTGTCCTTCAATAATCGGGATGTCTCCTATAGAAAAATGTACCATTCTCACACATTACATCAAGGGAGA

ATCAGTACAATCCCCGTCTGGAACCTCATGGAACACTGATCCCCTTGTAGTAGATCATTACTCATGCTCTCTGACCTACC

TTCGCCGAGGTTCCATCAAACAAATCAGGTTGAGGGTGGATCCTGGGTTTGTATTCGAGGCGTTGACAGACGTCGACTTT

AAACAACCTCGCAAAGCTAAGTTAGACATATCGGTCGTTGGGTTGACTGATTTCTCTCCCCCTTGCGACAACGTCGGTGA

TTTTCTAGGGACTATCAACACATTGAGGCACAATCTGCCTGTCACCGGAACCGGGGTCTCGAACTATGAAGTCCACGCTT

ATCGTAGAATTGGCCTGAATTCATCAGCATGTTACAAAGCCGTAGAGATCTCCACGTTAATCAAGCCATCCTTAGAAGTC

GGAGAACACGGATTGTTCTTAGGAGAAGGTTCCGGTTCAATGCTGGCTGCGTACAAGGAAGTTCTTAAATTAGCAAACTG

TTATTACAACAGCGGGGTAACAGCGGAGGGCAGAGCCGGACAGAGGGAAATATCTCCCTATCCTTCAGAGATGAGCCTAG

TAGAGAATCAAATGGGGATAGAGCGGAGCGTTAAAGTGCTGTTCAACGGCAAACCTGAAGTGACTTGGGTGGGGTCCACC

GATTGCTACAAGTACATAATCAGTAACATCCAAACCTCTAGTCTAGGTTTCATACATTCAGATATTGAGACACTCCCAAC

CAAAGATGCTGTTGAGAAATTAGAAGAATTTGCCTCTATCCTATCCTTATCCTTGATTTTGGGGAAAATCGGCTCTATTA

CAGTTGTCAAAATTATGCCCATTAGCGGGGATTTTACCCAAGGCTTCATAGCCTATGCTATCCAATATTTCAGGGAGAGT

CTGCTTGCCTACCCGAGATACAGTAACTTCATCTCGACTGAGTGTTACCTTATTATGATAGGATTAAAGGCCAATCGATT

GATAAACCCAGAGGCCATTAAGCAAAGCATAATCAGAGCGGGGGTCAGGACTGCACCAGGACTTGTGAGCCATATATTAT

CAGAGAAACAGAAAGGTTGTATTCAATCTTTCTTGGGTGATCCTTATATCCAAGGAGACTTCAACAAGCACCTTAAATCT

CTAACCCCTATTGAGAAAATCCTGGTAAATTGTGGTCTCTCGATCAATGGCACAAAAATCTGTAGAGATCTAATCCACCA

TGATATCGCCTCCGGTCCAGACGGTCTTATGAGCTCTACAATTATTTTATATAGGGAATTGGCTCATTTCAAAGATAATA

TAAGGAGTCAGCACGGAATGTTCCACCCCTACCCGGTACTGGCCAACAGCAGGCAACGTGAATTAATCCTTCGAATAGCC

AAGAAATTCTGGGGGTATGTCTTGCTATATTCTGATGACCCGGCACTAATCAGACAAACAATCAAGAACTTGAAACGGAA

TCATCTAACCTTTGACTTACATAGTAACCCGTTTATTAAAGGGCTATCCAAAGCTGAGAAACTGCTGGTGCGGACAAGTT

CACTAAGAAGGGAATGGTTGTTCACTCTAGATACGAAAGAAGTGAAAGAGTGGTTCAAATTGGTAGGTTACAGTGCACTC

ATCAGAGGCTAGTTAGCTATACATCTGCCCCCTCCTCCTCCACCATGAGACCCTACTGACGATCCAGAAGATTAAAGAAA

ACTACATATTGGATAAGTATCTATTCCCAGTTTTGTCTGGT

>KY967608.1/goat/Pakistan/2015

ACCAAACAAAGTTGGGTAAGGATAGATCTTACAATAACTATGGACTGGCAAACTTAGGAGTAAAGATCCTACTGTCGGGG

GGAGGAGGAGGAGCAAGATCTCTGATTATGGCGACTCTCCTTAAAAGCTTGGCATTGTTCAAAAGGAACAAAGACAAAGC

GCCGACGGCATCAGGTTCAGGAGGGGCCATCCGGGGGATTAAGAATGTTATCATAGTCCCGATTCCCGGAGACTCGTCCA

TCATTACCCGTTCAAGACTGCTCGACAGGCTTGTCAGATTGGCCGGAGATCCTGACATCAACGGGTCAAAGCTGACCGGC

GTGATGATCAGCATGTTATCTTTGTTCGTAGAGTCACCCGGGCAACTGATACAGCGAATCACAGATGATCCAGATGTGAG

TATCCGCCTTGTTGAGGTAGTTCAAAGTACTAGGTCCCAGTCCGGGTTGACCTTTGCATCACGTGGTGCTGACTTGGACA

ACGAGGCAGACATGTATTTTTCAACTGAGGGGCCCTCGAGTGGAGGTAAGAAAAGGATCAACTGGTTTGAGAACAGAGAA

ATAATAGACATAGAGGTGCAGGATCCAGAAGAGTTCAATATGTTGTTAGCCTCTATACTAGCACAAGTCTGGATCCTTCT

GGCCAAGGCTGTTACGGCACCGGATACGGCAGCTGACTCAGAACTGAGAAGGTGGGTTAAATACACACAACAAAGGAGAG

TGATTGGGGAATTTCGCCTTGACAAAGGGTGGCTGGACGCAGTCCGCAACAGAATTGCAGAAGATCTATCACTCCGGCGG

TTCATGGTATCTCTCATACTTGACATCAAGAGGACTCCCGGCAACAAGCCAAGGATTGCAGAAATGATCTGCGACATTGA

CAACTATATTGTCGAAGCCGGACTCGCCAGTTTTATCCTTACTATCAAGTTTGGTATCGAAACCATGTATCCTGCATTAG

GGCTCCACGAGTTTGCCGGGGAATTGTCCACTATTGAGTCCTTGATGAACTTGTATCAACAGCTAGGAGAGGTTGCACCC

TACATGGTAATTCTAGAGAACTCAATTCAGAACAAGTTTAGTGCAGGAGCTTATCCCCTCCTCTGGAGCTATGCGATGGG

TGTCGGGGTTGAGCTAGAGAACTCAATGGGGGGCTTGAACTTTGGCAGGTCATATTTTGACCCTGCCTATTTTCGTCTCG

GACAGGAGATGGTCAGAAGATCTGCAGGAAAGGTCAGCTCTGTGATTGCGGCTGAGCTCGGCATCACAGCAGAGGAAGCT

AAACTAGTCTCGGAAATCGCCTCGCAGGCTGGGGACGAAAGAACCGCTAGAGGGACTGGGCCTCGACAGGCGCAGGTCTC

CTTCCTCCAGCACAAAACAGGAGAGGGAGAGTCGCCCGCACCAGCGACCAGAGAAGGGGTCAAAGCTGCGATCTCGAACG

GATCCGAAGAGAGGGACAGAAAGCAAACACGCCCAGGAAGGTCCAGAGGAGAGACCCCCGGCCAACTGCTCCTAGAAATC

ATGCCAGAGGATGAGGTCTCGCGAGAGTCTGGTCAAAACCCTCGTGAGGCTCAAAGATCGGCTGAGGCACTCTTCAGGCT

GCAGGCCATGGCCAAGATCCTGGAGGACCAGGAGGAGGGAGAAGACAACAGTCAGATCTACAACGACAAGGATCTCCTCG

GCTAAGCAGACGCGCCCTCTATCGAAATTAGTGACAATACATCTCCTGCCAGTATTATAAAAAACTTAGGACCCAGGTCC

AGGCAAGCACACACCGACACTCCAACCTGTCGAGCGGAGACCACCGATGGCAGAAGAACAAGCATACCATGTCAACAAGG

GGCTGGAATGTATCAAGTCTCTCAAAGCCTCTCCCCCGGATCTATCCACCATCAGAGACACCATCGAGAGCTGGAGAGAG

GGGCTTAGCCCCTCGGGCCGTGCAACACCGGATCCTGATACGTCCGAGGGAGATCATCAGAATATCAACCAATCATGCTC

ACCAGCAATCGGACCAAGCAAAGTCTACCTGTCTCCTGAAGATAATCTCGGATTTAGAGAGATCACTGGCAACGACTGTG

AGGCTGGGCTCGGAGGAGTCCAGGGAAAAGGATCCAACTCTCAAGTACAGCGTTACTATGTTTATAGCCACGGGGGTGAA

GAGATTGAAGGACTCGAGGATGCTGACTCTCTCGTGGTTCAAGCAGACCCTCCAGTTGCTAACATCTTCGATGGAGGAGA

GGATGGATCTGACGACAGCGATGTGGACTCTGGCCCAGATGATCCCGGCAGAGATACTCTACATGACCGGGGATCTGTTG

CCGGCAATGATGTCGCTAGGTCCACGGATGTCGAAAAGCTAGAAGGTGCTGACCTTCAAGAAGTTCTTAACTCCCAGAAA

GGCAAAGGAGGAAGATTCCAAGGCGGGAAAACCTTACGAGTCCCGGAAATACCCGATGTCAAGCACTCCAGACCATCAGC

TCAATCAATTAAAAAGGGCACAGACGGGAGCTCAGTCTCATCTGGAACGGTGACAGAGTGTTTATCGATAAGTGGTGCAA

CCCAAGCTGTGCCAGAGTCAGGATGGGAGTCATCAGAGCAAAATGCGTCTGTGGGGAGTGTCCTCAAATCTGCGAGGAGT

GCAAAGACGATCCAGGGGTCGACACAAGAATCTGGTACCATAGCATCACTGACTCAGCCTAAAGAGAATGACTCCGAGTA

TGAGTATGAGGATGACCTATTTACAGAGATTCAGGACATCCGTGCAAGCATTGCCAAGATCCATGATGACAACAAGACTA

TCCTCTCAAAGCTTGATTCTATACTGTTATTGAAAGGAGAAGTCGACACTATCAAGAAACAAATCAGCAAGCAGAATATA

AGTATATCCACCATTGAGGGCCATCTCTCCAGTATAATGATAGCCATCCCGGGCTTTGGGAAGGATATCAAGGACCCAAC

ATCCGAGGTTGAGTTGAACCCAGATTTAAGACCTATAATAAGCCGTGACTCTGGCAGAGCTCTCGCGGAGGTCCTCAAGA

AACCCGCTGTGGATAGGTCTCAGAAAACTGGAACCAAAGCCAACTCCAGCTCAAAGGGTCAGCTTCTTAAGGATCTCCAG

CTAAAACCTGTCGACAAGCAGGCAAGCTCTGCAATCGGGTTTGTCCCGTCCGACCATGAATCATCCAGAAATGTCATCCG

CTCCATAATCAAGTCGAGCAAGCTAAACATTGATCACAAGGACTATCTTCTAGATTTACTGAATGATGTGAAAGGCTCCA

AGGATCTTAAGGAATTCCACAAGATGCTAACAGCAATTCTCGCCAAGCACCCGTAACACATCTCCCAGTCACCATCTCAT

ACTCGGCTGAAGACATCCTTTCAATCAGGCTATTACAAAAAACTTAGGAGCAAGGGCAACTGAGCTTCGCAGACAAGATG

ACCGAGATCTACGACTTCGACAAATCAGCATGGGATGTCAAAGGGTCAATTGCCCGCATAGAACCCACCACCTACCACGA

CGGCCGACTGATACCCCAGGTAAGAGTCATCGATCCTGGTCTGGGAGACAGAAAAGATGAGTGCTTTATGTACCTGTTTC

TCCTAGGAGTGATTGAGGATAACGACCCCCTGTCTCCCCCCGTCGGAAGAACCTTTGGCTCTTTACCTCTAGGGGTCGGT

AGGTCAACTGCTAAGCCAGAAGAACTACTAAGGGAGGCCACAGAACTAGACATAGTGGTGAGGCGCACTGCAGGAGTAAA

TGAGAAACTGGTATTTTACAACAACACTCCGCTGTCCTTGTTAACGCCCTGGAAGAAAGTTCTGACAACCGGAAGTGTGT

TTAGCGCTAACCAGGTTTGCAATGCAGTCAACCTAGTCCCGCTTGATACTCCCCAGAGGTTCAGGGTTGTGTACATGAGC

ATAACTAGATTATCAGACAATGGTTACTATAGTGTGCCCAGAAGAATGCTGGAGTTTCGCTCAGCCAATGCAGTAGCCTT

CAATATCTTGGTTACACTAAGAATTGAAAATGGCACAAACCCTAAAAGATACATAGTCGGCTCATGGGAGAATCCAGAGG

TCACATTTATGGTACACGTGGGGAACTTTAGAAGAAAGAAGAACGAAGTATACTCTGCTGATTACTGCAAAATGAAGATT

GAAAAGATGGGTTTAGTTTTTGCCCTGGGAGGAATAGGTGGAACAAGTCTCCATATTAGGAGCACCGGGAAAATGAGCAA

AACCCTCCATGCACAGCTGGGGTTCAAGAAAATCTTATGTTACCCCCTAATGGATGTTAATGAGGATCTTAACCGATATC

TCTGGCGGGCAGAGTGCCGAATAGTCAAAATCCAAGCCGTCTTACAGCCATCAGTACCCCAAGAATTCCGTGTCTACGAT

GATGTCATCATCAACGATGACCAGGGCTTGTTCAAGATCCTGTAGTTCATCTGCAAAGGAACGAACCCACCCCCAGGAAG

GGGCCCTACCCTCCCCCG-CCAAAAAACACACCCCAAGGCTCCCCGACACAGCGCCAACACAGCCACTCTCCCGATGGGA

CGAGGTTGGCC-CCCCAACAAACCCGGTCCAGGC-GGGGGGCCCCCCCCCTGACCCGCCGGGCGCCGACCGAGCAGGCCG

GCGAGAAGCCCCCAAGAGGCCCCGACGAAGACCGAGCACCCCTCCCCCCC-AAACCGTTCCCCC-AACACGCCCGAGGCC

AGGCGCCGGGCACCCCCAGACCCGCCACCGGCGGGACAGAACCCCAGGGGGGGACCCCCCAAACCAAAGGGACCACCGGG

GAAGGATCCGCCGAACCAGCACAGACCCGACCCAAGCAAGGCGGACCCCCAACACGAAACCGCCCAGCACACCCCACTCA

TCATCAAACAAAACTTAGGGGCCAAGCCCATACGACAGGCCCCAGTGCGCCAGGCGGAGGGACGCAGAAAGGAAGGAGAC

AAGACCGCCCCAGAAGGAGGACACCCAACCACCGAAACACCCAGCCCCAGGGAAGGAG--CCCCCCCCCACATGGCGAAA

GAACGGGCGAGAGACCCCGAGAAGAACCCGACCGGGAGCCAGGAGCCGGCGACGCCGGCGGCAAACAAGCCGGGCAAACA

CCCGACCCCACCAGAGGAGCCGAGGAGAGCCCGACCCCGCACGAACCAAACAGCAGGGAGCCCAACCCCACCAAAAAAAG

AGGCGAGGCCCC-CCT-----CCTCCCCGCACACAACCCGCAGACAGACCGAGAGAGGACCACCTCCCAACAAGACAGGC

GGCTCCCTCGCAGCCGAGCCGCAGCTCCCCAGAACAGGACACCCCCGTACACTCAAAACACCCCCCCGCCCCCCGCCACC

GGGACAGGGGCTCCGCCACCACAGACCAAAGCCACCGAGAGCAAGCCAGAGGTCCCCCCCAGTCCCGCCCGGCGCCCCCC

CGAAGAGGCCCCGAAAAGGCCCCCCAGCCCGACGACCCCTGCACGCACCGAAAGAGGGACCCGCCCGAGAACGTCCGGTA

CATCCATGCGTAGACAGCATGACACGGGTCGCAATCTTGGCATTTCTGTTTCTTTTTCTAAATGCTGTTGCGTGCCAGAT

TCACTGGGGCAATCTATCCAAGATCGGGATTGTAGGAACAGGGAGTGCCAGCTACAAGGTGATGACTAGGCCAAGCCACC

AAACTCTGGTTATAAAGTTAATGCCAAATATAACAGCCATCGACAATTGTACAAAGTCAGAGATTGCAGAGTACAAGAGA

TTGCTGATCACAGTGTTAAAGCCTGTAGAGGATGCTCTGTCGGTGATAACCAAGAATGTAAGACCAATTCAAACTCTGAC

ACCTGGGCGTAGAACCCGTCGTTTTGCTGGAGCTGTTCTGGCCGGAGTAGCGCTTGGAGTTGCGACAGCCGCTCAGATAA

CTGCAGGAGTCGCTCTTCATCAATCATTGATGAACTCCCAAGCAATTGAGAGTTTAAAAACCAGTCTTGAGAAGTCGAAT

CAGGCAATAGAAGAAATCAGACTTGCAAATAAGGAGACCATACTGGCAGTACAGGGCGTCCAGGATTATATCAACAATGA

GCTTGTCCCTTCTGTTCATAGAATGTCATGCGAGCTGGTAGGCCACAAGCTCGGCCTCAAGCTCCTTAGGTACTACACCG

AGATCCTGTCCATATTCGGGCCCAGCCTTCGAGACCCGATAGCTGCCGAAATATCAATCCAGGCACTCAGTTATGCATTA

GGCGGAGACATTAATAAAATCCTGGACAAGCTTGGGTATAGCGGTGGGGATTTCCTTGCCATCCTAGAAAGCAAGGGAAT

AAAGGCCCGGGTTACATATGTGGACACAAGAGATTACTTTATAATGCTTAGCATCGCCTACCCAACCTTATCTGAGATCA

AGGGAGTGATAGTTCACAAGATAGAAGCTATAACATACAACATTGGGGCACAGGAGTGGTATACTACTATCCCTAAATAT

GTAGCCACTCAGGGATACCTGATATCGAACTTTGATGAGACGTCATGCGTATTCACTCCAGAGGGGACAGTTTGCAGCCA

GAATGCGTTGTACCCAATGAGCCCATTGCTTCAGGAATGTTTCAGGGGGTCAACAAAATCGTGCGCCAGAACCCTAGTTT

TCAGGACCATAAGTAATAGATTTATCCTATCGAAAGGGAACCTGATTGCAAATTGTGCGTCAGTTTTGTGCAAATGCTAC

ACAACGGAGACAGTTATCAGCCAAGATCCTGACAAACTACTAACTGTTGTAGCATCCGACAAGTGTCCTGTAGTTGAGGT

GGACGGAGTGACAATACAGGTCGGCAGTCGAGAGTATCCGGATTCTGTATACTTACACAAAATAGACTTAGGTCCAGCCA

TCTCCCTAGAAAAACTGGATGTAGGCACCAATTTAGGCAATGCAGTCACAAGACTGGAGAATGCAAAGGAGCTCCTAGAT

GCATCAGACCAAATACTGAAGACTGTTAAAGGGGCACCTTTCGGTGGGAATATGTACATAGCACTGGCAGCTTGCATTGG

AGTATCCCTAGGGCTTGTCACATTAATATGCTGCTGTAAGGGGAGGTGTAAGAACAAGGAAATCCCTATCTCCAAAATCA

ACCCAGGGCTCAAACCCGACCTGACCGGGACCTCAAAGTCGTACGTGAGATCACTGTAGTCTGAATCACTTGAATCATCC

AGCATCACACACATACAGGTACGACGCAGGCAGTCTGAGGACGCAGGAAGCCCAGCCTCCGATCACCCACCTGACCCTAC

TCCACGCTCCACCACACATTAGTCATCAAACAAAACTTAGGACGAAAGGTCAATCACCATGTCCGCACAAAGGGAAAGGA

TCAATGCTTTCTACAAAGACAATCCTCACAATAAGAACCATAGGGTGATCCTGGATAGAGAACGCCTGGTCATTGAAAGA

CCCTATATCTTGCTTGGAGTCCTGCTGGTAATGTTCCTGAGTCTAATCGGACTGCTGGCCATCGCAGGGATCAGGCTTCA

CCGGGCCACCGTTGGGACTTCAGAGATCCAGAGTCGGCTGAATACCAATATTGAGTTGACCGAATCTATTGATCACCAAA

CTAAGGATGTCTTAACCCCCCTTTTTAAAATCATTGGCGATGAAGTCGGCATCAGAATTCCACAGAAATTCAGTGATCTT

GTCAAGTTCATCTCCGATAAGATTAAATTCCTTAACCCTGATAGAGAGTATGATTTCAGGGATCTCCGGTGGTGTATGAA

TCCCCCAGAGAGAGTCAAAATTAATTTTGATCAGTTTTGTGAGTACAAAGCTGCGGTTAAGTCAATTGACCATATATTTG

AGTCACCACTCAACAAGTCAAAAAAGCTGCAATCATTGACTCTTGGGCCCGGAACAGGCTGTCTAGGCAGGACAGTAACA

AGAGCTCATTTCTCAGAACTTACACTGACCTTAATGGACCTGGATCTAGAGATGAAGCACAATGTGTCCTCAGTGTTTAC

CGTTGTTGAAGAGGGATTATTCGGAAGAACGTATACCGTCTGGAGATCCGATGCCAGGGATCCGGGCACCGATCTAGGTA

TCGGCCATTTTTTAAGAGTCTTTGAGATTGGATTGGTAAGAGATCTCGGGCTGGGTCCCCCTGTTTTCCATATGACCAAC

TATCTCACAGTGAACATGAGTGATGACTATCGGAGATGTCTTTTAGCGGTAGGGGAGTTGAAGTTGACAGCCCTATGCAC

CTCATCTGAGACTGTGACACTGAGTGAGAGAGGAGTTCCAAAGAGGGAGCCTCTTGTGGTTGTGATACTTAACCTGGCTG

GGCCCACTCTAGGGGGCGAACTATACAGTGTCTTGCCTACCTCTGATCTCATGGTGGAGAAACTCTACCTATCTTCACAT

AGAGGGATTATCAAAGATGACGAGGCCAATTGGGTAGTGCCGTCTACCGATGTTCGTGATCTTCAAAACAAAGGTGAATG

TCTGGTGGAAGCATGCAAGACCCGACCTCCTTCATTCTGCAATGGCACAGGATCAGGCCCGTGGTCAGAGGGGAGAATCC

CTGCTTACGGGGTGATCAGGGTCAGTCTTGACTTAGCTAGTGACCCGGGTGTATTTATCACTTCAGTGTTTGGCCCACTG

ATACCTCACCTATCCGGCATGGATCTTTACAACAACCCGTTTTCAAGAGCTATATGGTTGGCTGTACCACCTTATGAACA

GTCATTTCTAGGAATGATAAATACCATTGGATTCCCTAACAGAACAGAGGTTATGCCGCATATTTTGACCACAGAGATCA

GAGGCCCTCGGGGTCGTTGCCATGTTCCCATAGAATTGTCCCGCAGGGTTGATGACGATATCAAGATCGGGTCCAACATG

GTCATATTGCCGACGATGGATTTGAGGTATATTACAGCCACTTATGATGTTTCCAGGAGCGAGCATGCAATCGTGTACTA

TATCTATGACACAGGTCGCTCATCATCTTACTTCTACCCAGTTCGACTAAATTTCAAAGGCAATCCTCTCTCTCTGAGGA

TAGAGTGCTTCCCTTGGCGTCATAAGGTGTGGTGCTACCATGATTGTCTTATATACAACACCATAACAGATGAAGAAGTC

CATACGAGAGGGCTGACCGGTATAGAGGTAACATGTAATCCAGTCTGAGCAGAGCTACTACCGTCACTCAGGCAGTCCCT

TGAGTCGCCACCGAGCCCAAGCAGCACAGCCTGGGACACTCAACAGCACAGCCCAGCCAACAACGTTATAAAAAACTTAG

GAGCCAGGGTCGTAGGGGCCATGGACTCCCTATCAGTCAATCAGGTCTTGTACCCTGAGGTTCATCTAGATAGCCCTATT

GTCACAAACAAACTAGTTGCCATCCTTGAGTACTCGGGTATCGACCATAACTATGTTCTTGAAGACCAGACCCTTGTCAA

GAATATTCGGTATAGACTGGGATGCGGTTTTTCAAATCAAATGATCGTCAATAACAGGGGGGTGGGAGAAACAGTCAATT

CTAAACTTAAAAGTTACCCCCATAATCGTCATATCATATACCCGGATTGCAATAAGGAGTTGTTTTGTATCAAAGATAGC

TGCATATCTAAGAAGCTCTCGGAGCTATTCAAAAAGGGTAATTCCTTGTACTCTAAAATAAGTCACCAGGTACTGGATTG

TCTTAAGAGAGTCAATGGGAAATTAGGCCTGGGCACAGACCTTAATCATGGTCTGAAGGAGGGTATCCTTGACCTAGGGT

TGCACATGCATAGCTCTCAATGGTTCGAGACTTTCCTGTTCTGGTTCACTATCAAGACAGAGATGAGATCAATGATCAAA

GAACAGTCCCATATATGCCACAAGAGGAGGTATAACCCGACTTTTGTATCGGGGGATGCATTCGAGGTGCTCGTATCACG

AGACCTCGTTGTGATAATTGATAAGAATACCCAGTATGTCTTCTACTTAACATTTGAGCTGGTCCTTATGTACTGTGATG

TCATAGAGGGCAGACTTATGACGGAGACAGCCATGGCCATAGACCAGAGATATTCAGAACTTCTAAGCCGGGTCAGATAC

TTGTGGGATCTTATTGATGGGTTCTTCCCAACACTAGGTAACACCACATACCAAATTGTTGCTCTGCTTGAACCATTATC

GTTGGCTTATCTTCAACTTCAGGATGTCACTCTGGAGTTAAGGGGTGCTTTTTTAGACCATTGCTTCAAAGAACTCTATG

AGATACTGGAGCATTGCGGCATTGACACAGAAGGCACCTACAATTCCATCACTGAAGGATTGGATTACGTATTCATCACC

CACGATATACATCTAACTGGGGAGATTTTTTCATTTTTCCGGAGTTTCGGACACCCCCGTCTAGAAGCAGTCACCGCTGC

AGAAAATGTCAGGAAACATATGAACCAACCGAAGGTGATCAGCTATGAGACTATGATGAAAGGGCATGCAGTATTTTGCG

GGATAATCATAAATGGTTTTAGGGACCGACATGGCGGCAGCTGGCCCCCTGTTGCATTGCCAGAACATGCTTCTGCTGCG

ATCCGGAATGCGCAGGCATCAGGTGAGGGACTGACTCATGACCTGTGTATAGACAACTGGAAATCTTTTGTTGGATTCAA

ATTTGGCTGCTTTATGCCACTCAGCCTAGATAGTGATTTGACCATGTACCTCAAAGACAAGGCATTGGCTGCACTGAAGA

ATGAGTGGGATTCAGTTTACCCGAAAGAATACCTCCGTTATAACCCACCTAGAGGGACAGAGTCAAGGCGACTGGTGGAG

GTGTTCCTGAATGACTCCAGCTTTGATCCTTATAACATGATAATGTACGTGGTGAATGGCTCCTACCTTAAAGACCCTGA

GTTTAATCTCTCATACAGCCTAAAAGAGAAGGAGATAAAGGAAACAGGGCGGTTGTTTGCCAAAATGACCTATAAGATGC

GGGCCTGTCAGGTAATCGCTGAAAATCTGATATCGAATGGTGTTGGGAAGTATTTCCGAGACAATGGGATGGCAAAGGAC

GAGCATGACCTAACAAAAGCCCTTCACACCCTGGCAGTCTCAGGTGTTCCCAAGAACAACAAAGATAACCACCGAGGTGG

GCCCCCCAGAAGGACAACAAGCCGAGGGGTGAGATCAAGCCAAGACACCAAAACACAAAATMGAGACAATGTCCAAGGGG

GGCCTGTGTACAACTATTTGCGATGCCAACCAATCAGCCCTGATCAGGGTGAGTCATACGAGACTGTTAGTGCATTCATC

ACTGCCGACCTTAAGAAGTATTGCCTGAATTGGAGATACGAGACAATCAGCATATTTGCACAGAGGCTGAATGAAATATA

TGGACTACCATCCTTCTTTCAATGGTTACACAGGATATTGGAAAAATCCGTACTCTACGTCAGTGACCCACATTGCCCTC

CCGACCTAGATAATCATATCCCTCTGGACAGTGTCCCTAATTCCCAAATATTCATTAAGTACCCAATGGGCGGAATAGAA

GGTTATTGCCAAAAACTATGGACAATCAGTACTATACCATACTTGTATCTAGCAGCCTATGAGAGCGGAGTAAGAATCGC

CTCACTAGTACAGGGTGACAATCAGACAATCGCAGTGACAAAAAGAGTTCCAAGTTCTTGGCCTTATTCACTAAAAAAGA

GGGAGGCATCCAAAGCCGCTCAAAATTACTTCGTAGTCTTAAGGCAAAGATTGCACGATGTAGGTCATCACTTAAAGGCT

AATGAAACCATAGTATCTTCTCACTTTTTTGTGTATTCCAAAGGGATTTATTATGACGGGCTGCTAGTCTCACAATCACT

AAAGAGCATCGCCAGATGTGTCTTCTGGTCCGAGACTATAGTGGATGAAACCAGAGCGGCTTGCAGCAATATTGCAACAA

CTGTCGCTAAGAGTATAGAGAGGGGTTATGATAGGTACCTTGCATATTCTTTGAATATCCTCAAGATTTTCCAACAGATC

CTTATATCCCTTAACTTCACTATTAACACAACAATGACTCAGGATGTCGTGGCACCAATCATCGAGAACGGTGATTTGCT

GATAAGAATGGCACTCTTGCCAGCACCCATCGGAGGTCTCAATTATCTTAACATGAGCAGGTTGTTTGTGAGAAATATCG

GTGACCCGGTCACCTCCTCTATAGCAGACCTGAAGAGGATGATAGATGCTGGGCTAATGCCAGAAGAAACATTGCATCAA

GTAATGACCCAGACCCCAGGAGAATCATCCTACCTCGATTGGGCAAGTGACCCTTATTCTGCCAACCTACCCTGCGTACA

GAGTATAACTCGCCTTCTCAAAAACATCACTGCACGATATATTTTAATCAGCAGCCCAAACCCGATGCTGAAAGGACTGT

TTCATGAGGGGAGTAGAGATGAAGACGAGGAGCTTGCAAGTTTCCTAATGGATCGGCATATAATTGTTCCAAGAGCTGCA

CATGAAATTTTAGACCATAGCATAACCGGGGCAAGAGAAGCTATAGCCGGGATGTTGGACACCACCAAGGGTCTTATTAG

AACAAGTATGAAACGGGGTGGCCTCACCCCTCGAGTCTTGGCCCGCCTTTCCAATTATGATTATGAACAATTCAGATCCG

GGATAACACTATTAACAAAGAAAGGGCAGTGTTATCTCATTGACAAGGACTCATGCTCGGTGCAGCTCGCTATAGCTCTG

AGGAGCCACATGTGGGCTAGGTTGGCTCGCGGGAGACCAATCTATGGATTGGAGGTGCCCGATATATTAGAATCGATGAA

TGGCTACCTTATCAAACGCCACGAGTCTTGTGCTATCTGTGAAACGGGCTCAAGTCACTACGGATGGTTTTTCGTCCCTG

CAGGGTGCCAGCTTGACGATGTATCTAGAGAGACTTCGGCTCTTCGTGTACCTTATGTCGGATCAACAACTGAGGAAAGG

ACAGACATGAAACTTGCATTTGTTAGATCCCCAAGTAGATCCCTCAAATCAGCAGTTAGAATTGCTACAGTTTACTCATG

GGCATACGGGGATGATGAGAAATCATGGAGTGAAGCCTGGATGCTAGCTAGGCAGAGAGCCAATATCACTTTAGATGAAT

TGAGAATGATCACTCCAGTCTCCACATCCACTAACTTAGCCCACCGGTTGAGGGATCGGAGTACCCAGGTGAAATACTCG

GGGACATCCCTTGTGAGGGTTGCAAGATACACAACCATCTCCAATGATAACTTGTCATTTGTGATATCTGAGAAAAAAGT

GGATACCAACTTCATTTATCAGCAAGGGATGCTGCTCGGTCTTGGGATCCTTGAGAACCTCTTTAGGCTAGAGGCCACCA

CCGGAGTATCCAACACAGTGCTACACCTGCACGTGGAAACAGAATGTTGTGTTGTACCTATGGTGGATCACCCAAGGATT

CCGAGTCTCCGTAATATAAAAGTTACGGATGAGCTATGCACAAACCCTTTGATCTACGACAGGTCTCCCATCATAGAGCA

CGATGCAACCCGACTATACTCACAGAGCCACAGGAGACATTTAGTGGAGTTTGTTACCTGGTCAACAAGCCAGCTTTACC

ATATACTGGCTAAGTCCACAGCAATGTCCATGATTGAGTTGATCACGAGATTCGAGAAAGATCACATGAATGAAATAGCC

GCCCTGATTGGCGATGACGATATCAACAGCTTCATCACTGAATTTTTGCTAGTAGAGCCTAGATTATTCATAGTTTACCT

TGGTCAGTGTGCTGCCATCAATTGGGCCTTTGATATACATTATCATCGGCCCTCGGGCAAGTACCAGATGGGGGAACTCC

TCTACTCTCTGCTCTCTCGGATGAGCAAAGGAGTATATAAGATCTTCACCAATGCTTTGAGCCACCCCAAAGTTTACAAG

AAATTTTGGCGAAGTGGTGTCATTGAGCCGATTCATGGCCCATCCCTGGATACACAGAATTTACATGTCACTGTCTGTGA

CATGATATATGGGTCCTATGTCACCTATTTGGATCTTTTGTTGAATGATGAGCTAGATGATTATCCGTATTTGCTCTGCG

AGAGTGATGAGGACGTGGTCACAGACAGGTTCGACAACATCCAAGCCAAACATCTCTGTGTATTGGCCGATGTATATTGC

AGCTCCAAGAGGTGTCCTTCGATAATCGGGATGTCTCCTATAGAAAAATGTACCATTCTCACACATTACATCAAGGGAGA

ATCAGTACAATCCCCGTCCGGGACCTCATGGAACACTGATCCTCTTGTAGTAGACCATTACTCATGCTCTCTGACCTACC

TTCGCCGCGGTTCCATCAAACAAATCAGGTTGAGAGTGGATCCTGGGTTTGTATTCGAGGCGTTGACAGATGTCGACTTT

AAACAACCTCGCAAAGCTAAATTAGATATATCGGTCGTTGGGTTGACTGATTTCTCTCCCCCTTTCGACAACGTCGGTGA

TTTTCTAGGGACTATCAACACATTGAGGCACAATCTGCCTGTCACTGGAACCGGGGTCTCGAACTATGAAGTCCACGCTT

ATCGTAGAATTGGCCTGAATTCATCAGCATGTTATAAAGCCGTAGAGATCTCCACGTTAATCAAGCCATCCTTAGAAGTC

GGAGAACACGGATTGTTCTTAGGAGAAGGTTCCGGTTCAATGCTGGCTGCGTACAAGGAAGTTCTTAAATTAGCAAATTG

TTATTACAACAGTGGGGTAACAGCGGAGGGCAGAGCCGGACAGAGGGAAATATCTCCCTATCCTTCAGAGATGAGCCTAG

TAGAGAATCAAATGGGGATAGAGCGGAGTGTTAAAGTGCTGTTCAACGGCAAACCTGAAGTAACTTGGGTGGGGACCACC

GATTGCTACAAGTACATAATCAGTAACATCCAAACCTCTAGTCTGGGTTTCATACATTCAGATATTGAGACGCTCCCAAC

CAAAGATGCTGTTGAGAAATTAGAAGAATTTGCCTCTATCCTATCCTTATCCTTGATTTTGGGGAAAATCGGCTCTATTA

CAGTTGTCAAAATTATGCCCATTAGCGGGGATTTTACCCAAGGCTTCATAGCCTATGCCATCCAATATTTTAGGGAGAGC

CTGCTTGCCTATCCGAGATATAGTAACTTCATCTCGACTGAGTGTTACCTTATCATGATAGGATTAAAGGCCAATCGATT

GATTAACCCAGAAGCCATTAAGCAAAGCATAATCAGAGCGGGGGTCAGGACTGCACCAGGACTTGTGAGCCATATATTAT

CAGAGAAACAGAAAGGTTGTATTCAATCTTTTTTGGGTGATCCCTATATCCAGGGAGACTTCAATAAGCACCTTAAATCT

CTAACCCCTATTGAGAAAATCCTGGTAAATTGTGGTCTCTCGATCAATGGCACAAAAATCTGTAGAGATCTAATCCACCA

TGATATCGCCTCCGGTCCAGACGGTCTTATGAGCTCTACAATTATTTTATATAGGGAATTGGCTCATTTCAAAGATAATA

TAAGGAGTCAGCACGGAATGTTCCACCCCTACCCGGTACTGGCCAACAGCAGGCAACGTGAATTAATCCTTCGAATAGCC

AAGAAATTCTGGGGGTATGTCTTGCTATATTCTGATGACCCGGCACTAATCAGACAAACAATCAAGAACTTGAAACGGAA

TCATCTAACCTTTGACTTACATAGTAACCCGTTTATTAAAGGGCTATCCAAAGCTGAGAAACTGCTGGTGCGGACAAGTT

CACTAAGAAGGGAATGGTTGTTCACTCTAGATACGAAAGAAGTGAAAGAGTGGTTCAAATTGGTAGGTTACAGTGCACTC

ATCAGAGGCTAGTTAGCTATACATCTGCCCCCTCCTCCTCCGCCATGAGACCCTACTGACGATCCAGAAGATTAAAGAAA

ACTACATATTGGATAAGTATCTATTCCCAGCTTTGTCTGGT

>MF737202.1/sheep/Georgia/2016-01-14

ACCAAACAAAGTTGGGTAAGGATAGATCTTATAATAACCATAGACTGGCAAACTTAGGAGTAAAGACCCTACTGTCGGGG

GGAGGAGGAGGAGCAAGATCTTTGATTATGGCGACTCTCCTTAAAAGCTTAGCATTGTTCAAAAGGAACAAAGACAAAGC

GCCGACGGCATCAGGTTCAGGGGGGGCCATCCGGGGGATTAAGAATGTTATCATAGTCCCGATTCCCGGAGACTCTTCCA

TCATTACCCGTTCAAGACTGCTAGACAGGCTTGTCAGATTGGCCGGAGACCCTGACATCAACGGGTCAAAGCTGACCGGC

GTGATGATCAGCATGTTATCTTTGTTCGTAGAGTCTCCCGGGCAATTGATACAGCGAATCACAGATGATCCAGATGTTAG

TATCCGCCTTGTTGAGGTAGTTCAAAGTACTAGGTCCCAGTCCGGGTTGACCTTTGCATCACGTGGTGCTGATTTGGACA

ACGAGGCAGACATGTATTTTTCAACTGAGGGGCCCTCGAGTGGGGGTAAGAAAAGGATCAACTGGTTTGAGAACAGAGAA

ATAATAGACATAGAGGTGCAGGATCCAGAAGAGTTCAATATGTTGTTGGCCTCCATACTAGCACAAGTCTGGATCCTTCT

GGCCAAGGCTGTTACGGCACCGGATACGGCAGCTGACTCAGAACTGAGAAGGTGGGTTAAATACACACAACAGAGGAGAG

TGATTGGGGAATTTCGCCTTGACAAAGGGTGGCTGGACGCAGTCCGCAACAGAATTGCAGAAGATCTATCACTTCGGCGG

TTCATGGTGTCTCTCATACTTGACATCAAGAGGACCCCCGGCAACAAGCCAAGGATTGCGGAAATGATCTGCGACATTGA

CAACTATATTGTCGAAGCCGGACTCGCCAGTTTCATCCTTACTATCAAGTTTGGTATCGAAACCATGTATCCTGCATTAG

GGCTCCACGAGTTTGCCGGGGAATTGTCCACTATTGAGTCCTTGATGAACTTGTATCAACAGCTAGGAGAGGTTGCACCC

TATATGGTAATTCTAGAGAACTCAATTCAGAACAAGTTTAGTGCAGGAGCTTATCCCCTCCTCTGGAGCTATGCGATGGG

TGTCGGAGTTGAGCTGGAGAACTCAATGGGGGGCTTGAACTTTGGCAGGTCATATTTTGACCCTGCCTATTTTCGTCTCG

GACAGGAGATGGTCAGAAGATCTGCTGGAAAGGTCAGCTCTGTGATTGCGGCTGAGCTCGGCATCACAGCAGAGGAAGCT

AAACTAGTCTCGGAAATCGCCTCGCAGACTGGGGACGAAAGAATTGCCAGAGGGACTGGGCCTCGACAGGCGCAGGTCTC

CTTCCTCCAGCACAAAGCAGGAGAGGGAGATTCGCCCGCACCAGCGACCAGAGAAGGGGTCAAAGCTGCGATCTCAAACG

GATCTGAAGAAAGGGACAGAAAGCAAACACGCCCAGGAAGGCCCAGAGGAGAGACCCCCGGTCAACTGCTCCTGGAAATC

ATGCCAGAGGATGAGGTCTCGCGAGAGTCTGGTCAAAACCCTCGTGAGGCTCAAAGATCAGCCGAGGCACTCTTCAGGCT

GCAGGCCATGGCCAAGATTCTGGAGGACCAAGAGGAGGAAGAAGACAACAGTCAGGTCTACAACGACAAGGATCTCCTCG

GCTAAGCAGACGCACCTTCTGTCGAAATCAGTGACAAGACATCTCCTACCAGTATTATAAAAAACTTAGGACCCAGGTCC

AAGCAAGCACACATCGACACTCCAATCAGTCGAGCGGAGACCACCGATGGCAGAAGAACAAGCATACCATGTCAACAAGG

GGCTGGAATGCATCAAGTCTCTCAAAGCCTCTCCCCCGGATCTATCCACCATCAGAGATACCATCGAAAGCTGGAGAGAG

GGGCTTAGCTCCCCGGGCCGTGCAACACCGAACCCTGATACGTCCGAGGGAGATCATCAGAATATCAACCAATCATGCTC

ACCAGCAATCGGACCAAACAAAGTCTACCTATCTTCTGAAGATAATCTCGGATTTAGAGAGATCACTGGCAACGACTGTG

AGGCTGGGCCCGGAGGAGTCCAGGGGAAAGGATCCAACCCTCAAGTACAGCGTTACTATGTTTATAGCCACAGGGGTGAA

GAGATTGAAGGACTCGAGGATGCTGACTCTCTCGTGGTTCAAGCAGATCCTCCAGTTGCTAACATCTTCAATGGAGGAGA

GGATGGATCTGACGACAGCGATGTGGACTCTGGCCCAGATGATCCCGGCAGAGATACTCTATATGACCGGGGATCTATTG

CCGGCAATGATGTCGCTAGGTCCACAGATGTCGAAAAACTAGAAGGTGCTGATATTCAAGAAGTTCTTAACTCCCAGAAA

GGCAAAGGAGGGAGATTCCAAGGCGGGAAAACCTTACGAGTCCCGGAAATACCCGATGTTAAGCACTCCAGACCATCAGC

CCAATCAATTAAAAAGGGCACAGACGAGAACTCAGTCTCATCTGGAACGGTGACAGAGTGTTTATCGATAAGTGGTGCAA

TCCAAGCTGTGCCAGAGTCAAGATGGGAGTCATCAGAGCAAAATGCGTCTGTGGGGAGTGTCCTCAAGTCTGCGAAGAGT

GCAAAGACGATCCAGGGGTTGACACAAGAATCTGGTACCATAGCATCACTGACTCAGCCTAAAGAAAATGACTCCGAGTA

TGAGTATGAGGACGACCTATTTACAGAGATTCAGGACATCCGTGCAAGCATTGCCAAGATCCATGATGACAACAAAACTA

TCCTCTCAAAGCTTGATTCTATACTGTTATTGAAAGGAGATGTTGACACTATTAAGAAACAAATCAGCAAGCAGAATATA

AGTATATCCACCATTGAGGGCCATCTCTCCAGTATAATGATAGCCATCCCGGGCTTTGGGAAGGATATCAAGGACCCAAC

ATCCGAGGTTGAGTTGAACCCGGACTTAAGACCCATAATAAGCCGTGACTCTGGCAGAGCTCTCGCGGAGGTCCTCAAGA

AACCCGCCGTGGATCGGTCCCAGAAAATTGGAACCAAAACCAACTCCAGCTCAAAGGGTCAGCTTCTTAAGGATCTCCAG

CTAAAACCTGTCGACAAGCAAGCAAGCTCTGCAATCGGGTTCGTCCCATCCGACCATGAATCATCCAGAAATGTCATCCG

CTCCATCATCAAGTCGAGCAAGCTAAACATTGATCACAAGGACTATCTTCTAGATTTACTAACTGATGTGAAAGGCTCCA

AGGATCTTAAGGAATTCCACAAGATGCTAACAGCAATTCTCGCCAAGCACCCGTAACACACCCTCCAGTCACCATCTCAT

ACTCGGCTGAAAGCGTCCTCTCAATCAGGCTATTACAAAAAACTTAGGAGCAAGGGCAACTGAGCTTCGCAGACAAGATG

ACCGAGATCTACGACTTTGATAAATCAGCATGGGATGTCAAAGGGTCAATTGCCCGCATAGAACCCACCACCTATCACGA

CGGCCGGCTGATACCCCAGGTAAGGGTCATCGATCCTGGTCTGGGAGACAGAAAAGATGAGTGCTTCATGTACCTGTTTC

TCCTAGGAGTGATTGAAGATAACGACCCCCTGTCTCCCCCAGTCGGAAGAACCTTTGGTTCTTTACCTCTAGGGGTCGGT

AGGTCAACTGCTAAGCCAGAAGAACTACTAAGGGAGGCCACGGAACTAGATATAGTGGTGAGGCGCACTGCAGGATTAAA

TGAGAAACTGGTATTTTACAACAACACTCCACTGTCCTTGTTAACGCCCTGGAAGAAAGTTCTGACAACCGGAAGTGTGT

TTAGCGCTAACCAGGTTTGCAATGCGGTCAACCTAGTCCCGCTTGATACTCCCCAGAGATTCAGGGTTGTGTACATGAGC

ATAACTAGATTATCAGACAATGGTTACTATAGTGTGCCCAGGAGAATGCTGGAGTTCCGCTCAGCCAATGCAGTAGCCTT

CAACATCCTGGTCACATTAAGAATTGAAAATGGCACAAACTCTAGAAGATACATAGCCGGCTCATGGGAGAATCCAGAGG

TCACATTTATGGTACACGTGGGCAACTTTAGAAGAAAGAAGAACGAAGTATACTCTGCTGATTACTGCAAAATGAAGATT

GAAAAGATGGGTTTAGTTTTTGCCCTAGGAGGAATAGGTGGAACAAGTCTCCATATTAGAAGCACCGGGAAAATGAGCAA

AACCCTCCATGCACAGCTAGGGTTCAAGAAAATCTTATGTTACCCCCTAATGGATGTTAATGAGGATCTTAACCGATATC

TCTGGCGGGCAGAGTGCCGAATAGTCAAAATCCAAGCCGTCTTACAGCCATCAGTACCCCAAGAATTCCGTGTCTACGAT

GATGTCATCATCAACGATGATCAAGGCTTGTTTAAGATCCTGTAGTTCATCTGCAAAGGAATGAACCCACCCCCAGGAAG

GAGCCTCACCCTCCCCCG-CCAAAAAACACACTCCAAGGCTCCCCGACACAGCGCCAACACAGCCACTCTCCCGATGGGA

CGAGGCTGGCT-CCCCAACAAACCCAGTCCAGGTGGGGGG-ACCCCCAAACGACCCGCCGGGCGCCGACCGATCAGGCCG

GCAAGAAGCCCCCAACAGGCCCCGACGACGACCGAGCACCCCACCCCCCC-AAACCGTCTCCCG-AACACGCCCGAGGCC

AGGCGCCGAGCACCCCCAGACCCGCCACCGGCGAGACAGAGCCCCAAGGGGGGACCCTCCAAACCAAAGGGACCACCGAG

GAACGCTCCGCCGAACCAGCACAGACCCGACCCAAGCAAGCCGGACCCCCAACACGAGACCGCCCAGCGCACCCCACCCA

TCACCAAACAAAACTTAGGGGCCAAGTCCGCACAACAGGCCCCGGCCCGCCAGGCGGAGGGCCGCAGAAAGGAAGGAGAC

ACGACCGTCCCAGAAGGAAGACACCCAACCACCGAAACACCCAGCCCCAGGGAAGGAG--CCCCCCCCCACACGGCGAAG

GAACGGGCGAGAGACCCCAAGGAGAACCCCACCGGAAGCCAAGAGCCGGCGACGCCGGCGGCAAACAGGCAGGGCGCACA

CCCGACTCCGCCAGAGGAACCGAGGAGAGCCAGACCCCGCGTGGAACAAGCAGCAGGGAATCCAACCCCGCCACAAAAAG

AGGTGAGGCCCT-CCT-----CTCTCTCGCACACAACCCACAGACAGGCCAAGGGAGGACCACCTCCCAACAAGACAGGC

GGCTCCCTCGCAGCCGAGCCGCAGCTCCCCAGAAGAGGACACCCCCGCACACTCCGAGCACCCCCCCACCCCCCGCCCCC

CGGACAGGAGCCCCGCCACCACTAACCAGAGCCATCGAGAGCCAGCCAGAGGCCCCCCCCAGTCCCGCCCAGCGCCTCYC

CGAAGGGGCCCCGGCAAGGCCCCCCAGCCCGACGACCCCTGCACGCACCGAAAGAGGGACTCGCTCAAGAACGTCCTGCA

CATCCATGCGCAAACATCATGACGCGGGTCGCAATCTTGAAATTTCTGTTTCTTTTTCTAAATGCTGTTGCGTGCCAGAT

TCACTGGGGCAATCTATCCAGGATCGGGATCGTAGGAACAGGAAGTGCCAGCTACAAGGTGATGACTAGGCCAAGCCACC

AAACTCTGGTTATAAAGTTAATGCCAAATATAACAGCCATCGACAATTGTACAAAGTCAGAGATTGCAGAGTACAAGAGA

CTGCTGATCACAGTGTTAAAGCCTGTAGAGGATGCTCTGTCAGTGATAACCAAAAATGTAAGACCAATTCAAACTTTGAC

ACCTGGGCGTAGAACTCGCCGTTTTGCTGGTGCTGTTCTGGCTGGAGTAGCACTTGGAGTTGCGACAGCCGCTCAGATAA

CTGCAGGAGTCGCCCTTCATCAATCATTGATGAACTCCCAAGCAATTGAGAGTTTAAAAACCAGTCTTGAGAAGTCGAAT

CAGGCAATAGAAGAAATCAGACTTGCAAATAAGGAGACCATACTGGCAGTACAGGGCGTCCAGGATTATATCAACAATGA

GCTTGTCCCTTCTGTTCACAGAATGTCATGCGAGCTGGTCGGTCACAAGCTCGGCCTCAAGCTCCTTAGGTACTACACCG

AGATCCTGTCCATATTCGGGCCCAGCCTTCGAGACCCGATAGCTGCAGAAATATCAATCCAGGCACTCAGTTATGCATTA

GGCGGAGACATTAATAAAATTCTGGACAAGCTTGGGTATAGCGGTGGGGATTTCCTTGCCATCCTGGAAAGCAAGGGAAT

AAAGGCTCGAGTCACATATGTGGACACAAGAGATTACTTCATAATTCTTAGCATCGCCTACCCAACCTTATCTGAGATCA

AGGGAGTGATAATTCACAAGATAGAAGCTATAACATACAACATTGGGGCACAGGAGTGGTATACTACTATCCCTAAATAT

GTAGCCACTCAGGGGTATCTGATATCGAACTTTGATGAGACATCATGCGTATTCACTCCAGAGGGGACAGTTTGCAGCCA

GAATGCGTTGTACCCAATGAGCCCATTGCTTCAGGAATGTTTCAGGGGGTCAACAAAATCGTGCGCCAGAACCCTAGTTT

CAGGGACCATAAGCAATAGATTCATCCTATCAAAAGGGAATCTGATTGCAAATTGCGCGTCAGTTTTGTGCAAATGTTAC

ACAACGGGGACAGTTATTAGCCAAGATCCTGACAAACTACTAACTGTTGTAGCATCTGACAAGTGTCCTGTAGTTGAGGT

GGACGGAGTGACAATACAGGTCGGTAGTCGAGAGTATCCGGATTCTGTATACTTACACAAGATAGACTTAGGTCCAGCCA

TCTCCCTAGAAAAACTGGATGTAGGCACCAATTTAGGCAATGCAGTCACAAGACTGGAGAATGCAAAGGAGCTTCTAGAT

GCATCAGACCAAATACTGAAGACTGTTAAAGGGGTACCTTTCAGTGGGAATATGTACATAGCACTGGCAGCTTGCATTGG

GGTATCCCTAGGGCTTGTCACATTAATATGCTGCTGTAAGGGGAGGTGCAAGAACAAGGAAATCCCGAACTCCAAAATCA

ACCCAGGGCTCAAACCCGATCTGACCGGGACCTCAAAGTCGTACGTGAGATCATTGTAGTCAGAATCACCCGAATCATCC

GGCATCACACACACACATGCACGACACAAGCAGTCAGAGGATGCAGGAAGCCCAGTCTCCGATCACCAACCTGACCCTAC

TCCAAGCTCCACCAGACATTATTCATCAAACAAAACTTAGGACGAAAGGTCAATCACCATGTCCGCACAAAGGGAAAGGA

TCAATGCCTTCTACAAGGACAATCCTCACAATAAGAACCATAGGGTGATCCTGGATAGAGAACGCTTGGTCATTGAAAGA

CCCTACATCTTGCTTGGAGTCCTGCTGGTAATGTTCCTGAGTCTAATCGGACTGCTGGCAATTGCGGGGATCAGGCTTCA

CCGGGCCACCGTTGGAACTTCAGAGATCCAGAGTCGGCTGAATACCAACATTGAGTTGACCGAATCTATTGATCACCAAA

CTAAGGATGTCTTAACTCCCCTTTTTAAAATCATTGGCGATGAAGTTGGCATCAGAATTCCACAGAAATTCAGTGATCTT

GTCAAGTTCATTTCCGATAAGATTAAATTCCTCAACCCTGATAGAGAGTATGATTTCAGGGATCTCCGGTGGTGTATGAA

TCCCCCCGAGAGAGTCAAAATTAATTTTGATCAGTTTTGTGAGTACAAAGCTGCGGCTAAGTCAATTGAACATATATTTG

AGTCACCACTCAACAAGTCAAAAAAGCTGCAATCACTGATTCTTGGGCCCGGATCAGGCTGTCTAGGCAGGACAGTAACA

AGAGCTCATTTCTCAGAACTTACACTGACCTTAATGGACCTAGATCTAGAGAAGAAGCACAACGTGTCCTCAGTGTTTAC

CGTAGTTGAAGAGGGATTATTCGGAAGAACATATACAGTCTGGAGATTCGATGCCAGAGATCCAAGCACTGATCTAGGTA

TTGGCCATTTTTTAAGAGTCTTCGAGATTGGACTGGTAAGGGATCTCGGGCTTGGTCCCCCTGTTTTTCAGATGACCAAC

TATCTCACAGTTAACATGAGCGATGACTATCGGAGATGTCTTTTAGCGGTAGGGGAGTTGAAGTTGACAGCTCTGTGCAC

CTCATCTGAGACTGTGACACTAAGTGAGGGAGGAGTCCCAAAGAGGGAGCCTCTTGCGGTTGTGATACTTAACCTAGCTG

GGCACACTCTAGGGGGCGAACTATACAGTGTCTTGCCTACTTCTGATCTCATGGTGGAGAAACTCTATTTATCTTCTCAT

AGAGGGGTTATCAAAGATGACGAGGCCAATTGGGTGGTGCCGTCTACCGATGTCCGTGATCTTCAAAACAAAGGAGAATG

TCTGGTGGAAGCATGCAAAACTCGACCTCCTTCATTTTGCAATGGCACAGGATCAGGCCCGTGGTCAGAGGGGAGAATCC

CTGCTTACGGGGTGATCAGGGTCAGTCTTGACTTAGCCAGTGACCCGGGTGTAGTTATCACTTCAGTGTTTGGCCCGCTG

ATACCTCACCTATCCGGCATGGATCTTTACAATAACCCGTTTTCAAGAGCTGTATGGTTGGCTGTACCACCTTATGAGCA

GTCATTTCTAGGAATGATAAATACAATTGGGTTCCCCGACAGAGGAGAGGTTATGCCGTACATTTTGACCACAGAGATCA

GAGGCCCTCGAGGTCGTTGCCATGTTCCCATAGAATTGTCCCGCAGGGTTGATGACGATATCAAGATCGGGTCCAACATG

GTCATATTGCCGACGATGGACCTGAGGTATATTACAGCCACTTATGATGTTTCCAGGAGCGAGCATGCAATCGTGTATTA

TATCTATGACACAGGTCGCTCATCATCTTACTTCTACCCTGTTCGACTGAATTTCAAAGGCAATCCCCTCTCTCTGAGGA

TAGAGTGTTTCCCTTGGCGTCATAAGGTGTGGTGCTACCATGATTGTCTTATATACAACACCATGACAGATGAAGAAGTC

CATGCAAGAGGACTGACCGGTATAGAGGTAACATGTAATCCAGCCTGAGCAGAGCCACGACCGTCACTCAGGCAGTCCCT

CGAGTCGTCACTGAGTCCAAGCAGCATAGCCTGGGATACTCAACAGCACAGCCCAGCCAATAATGTTATAAAAAACTTAG

GAGCCAGGGTTGTAGAGGCCATGGACTCCCTATCAGTCAATCAGGTCTTGTACCCTGAGGTTCATCTAGATAGCCCTATT

GTCACAAACAAACTAGTTGCCATCCTTGAGTACTCGGGTATCGACCATAACTATGTTCTTGAAGACCAGACTCTTGTCAA

GAATATTAGGTATAGACTGGGGTGCGGTTTTTCAAATCAAATGATCATCAATAACAGGGGGGTGGGTGAAACAGTCAATT

CTAAACTTAAAGGCTACCCCCATAATCGTCATATCATATACCCGGATTGCAATAAGGAGTTGTTTTGTATCAAAGATAGC

TGCATATCTAAGAAGCTCTCGGAGCTATTCAAGAAGGGTAATTCCTTGTACTCTAAGATAAGTCACCAGGTACTGGATTG

TCTCAAGAGAGTCAATGAGAAATTAGGCCTGGGCACAGATCTTACCCATGGTCTGAAGGAGGGTATCCTTGACCTAGGGT

TGCACATGCATAGCTCTCAGTGGTTCGAGACTTTCCTGTTCTGGTTCACTATCAAGACAGAGATGAGATCAATGATCAAA

GAACAGTCCCATATATGCCACAAGAAGAGGTATAACCCGATTTTTGTGTCGGGGGCTGCATTCGAGGTGCTCGTATCACG

GGACCTCGTTGTGATAATTGATAAGAATACCCAGTATGTCTTCTACTTAACATTTGAGCTGGTCCTTATGTATTGTGATG

TCATAGAGGGCAGACTTATGACGGAGACAGCCATGGCCATAGACCAGAGATATTCAGAACTTCTAAGCCGGGTCAGATAC

TTGTGGGATCTTATTGATGGGTTCTTCCCAACACTAGGCAACACCACATACCAAATTGTTGCTCTGCTTGAACCATTATC

GTTGGCTTATCTTCAACTTCAGGATGTCACTCTGGAGTTAAGAGGTGCTTTTTTAGACCATTGCTTCAAAGAACTCTATG

AGATACTGGAGCATTGCGGCATTGACACAGAAGGCACCTACAATTCCATCACTGAAGGATTGGATTACGTATTCATCACC

CACGATATACATCTAACTGGGGAGATCTTTTCATTTTTCCGGAGTTTCGGACACCCCCGTCTAGAAGCAGTCACCGCTGC

AGAAAATGTCAGGAAACATATGAACCAACCGAAGGTAATCAGTTATGAGACTATGATGAAAGGGCATGCAGTATTTTGCG

GGATAATCATAAATGGTTTTAGGGACCGACATGGCGGCAGCTGGCCCCCTGTTGCATTGCCAGAACATGCTTCTGCTGCG

ATCCGGAATGCGCAGGCATCAGGTGAGGGATTGACCCATGACCTGTGTATAGACAATTGGAAATCCTTTGTTGGATTCAA

ATTCGGCTGCTTTATGCCACTCAGCCTAGATAGTGATTTGACCATGTACCTCAAAGACAAGGCATTGGCTGCACTGAAGA

ATGAGTGGGATTCAGTTTACCCGAAAGAATACCTCCGTTATAACCCACCTAGAGGGACAGAGTCAAGGCGACTGGTGGAG

GTGTTCCTAAATGACTCCAGCTTTGATCCTTATAACATGATAATGTACGTGGTGAATGGCTCCTACCTTAAAGACCCTGA

GTTTAATCTTTCATACAGCCTAAAGGAGAAGGAGATAAAGGAGACAGGGCGGTTGTTTGCTAAAATGACCTATAAGATGC

GGGCCTGTCAGGTAATCGCTGAAAATCTGATATCAAATGGTGTTGGGAAGTATTTCCGAGACAATGGGATGGCAAAAGAC

GAGCATGACCTAACAAAAGCCCTTCACACCCTGGCAGTCTCAGGTGTTCCCAAGAACAACAAAGATAACCACCGAGGTGG

CCCTCCCAGAAGGACAACAAACCGAGATGTGGGATCAAGCCAGGACACCAAAACACAAAATAGAGATAAGGTCCAAGGGG

GGCCTATGTACAACTATTTGCAATGCCAACCAATCAGCCCTGATCAGGGTGAGTCATACGAGACTGTTAGCGCATTCATC

ACCGCTGACCTTAAGAAGTATTGCCTGAATTGGAGATACGAGACAATCAGCATATTTGCACAGAGGCTGAATGAAATATA

TGGACTACCATCCTTCTTTCAATGGTTACACAGGATATTGGAAAAATCCGTGCTCTACGTCAGTGACCCACATTGTCCTC

CCGACCTAGATAATCATATCCCTCTGGACAGTGTCCCTAATGCCCAAATATTCATTAAGTACCCGATGGGCGGAATAGAA

GGTTACTGCCAAAAACTATGGACAATCAGTACTATACCATACTTGTATCTAGCAGCCTATGAGAGCGGAGTAAGAATCGC

CTCACTAGTGCAGGGTGACAATCAGACAATTGCAGTGACAAAAAGAGTTCCAAGTTCTTGGCCTTATTCACTAAAAAAGA

GGGAGGCATCTAAAGCCGCTCAAAATTACTTCGTAGTCTTAAGGCAGAGATTGCACGATGTAGGTCATCACTTAAAGGCT

AATGAAACCATAGTATCTTCTCACTTTTTTGTATATTCCAAGGGGATTTATTATGACGGCCTGCTAGTTTCACAATCACT

AAAGAGCATCGCCAGATGTGTCTTCTGGTCCGAGACTATAGTGGATGAAACCAGAGCGGCTTGCAGCAATATTGCAACAA

CTGTCGCTAAGAGTATAGAGAGGGGTTATGATAGGTACCTTGCATATTCTTTGAATATCCTCAAGATTTTCCAACAGATC

CTCATATCCCTTAACTTCACTATTAACACAACAATGACTCAGGATGTCGTGGCACCGATCATCGAGAACGGGGATCTGCT

GATAAGGATGGCACTCTTGCCAGCACCCATCGGGGGTCTCAATTATCTTAACATGAGCAGGTTATTTGTGAGAAATATTG

GTGACCCGGTCACCTCCTCTATAGCCGACCTGAAGAGGATGATAGATGCTGGGCTAATGCCAGAAGAAACACTGCATCAA

GTGATGACCCAGACCCCGGGAGAATCATCCTACCTCGATTGGGCAAGTGACCCTTATTCTGCCAACCTACCCTGCGTACA

GAGTATAACTCGCCTTCTCAAAAACATCACTGCACGATATATTTTAATCAGCAGTCCAAACCCAATGCTGAGAGGATTGT

TTCATGAGGGGAGTAGAGATGAGGACGAAGAGCTTGCAAGTTTCTTAATGGATCGGCATATAATTGTTCCAAGAGCTGCA

CATGAAATCTTAGACCATAGCATAACTGGGGCAAGAGAAGCTATAGCCGGGATGTTGGACACCACCAAGGGTCTGATTAG

AACAAGTATGAGACGGGGTGGCCTCACCCCTCGAGTCTTAGCCCGCCTTTCCAACTATGATTACGAACAATTCCGATCCG

GGATAACACTATTGACAAAGAAAGGGCAGTGTTATCTCATTGACAAAGACTCGTGCTCGGTGCAGCTCGCTATAGCTCTG

AGGAGCCACATGTGGGCTAGGTTGGCTCGCGGGAGACCAATCTATGGGTTGGAGGTGCCCGATATATTAGAATCGATGAA

CGGCTACCTTATCAAACGCCACGAGTCTTGTGCTATCTGTGAAACGGGCTCAAGTCACTACGGATGGTTCTTTGTCCCTG

CAGGGTGCCAACTTGATGATGTATCTAGAGAGACTTCGGCTCTTCGTGTACCTTATGTCGGATCAACAACTGAAGAGAGG

ACGGACATGAAACTTGCTTTTGTTAGGTCTCCAAGTCGATCCCTCAAATCAGCAGTCAGAATTGCAACAGTTTACTCATG

GGCCTACGGGGATGATGAGAAATCATGGAATGAGGCCTGGATGCTAGCTAGGCAGAGAGCCAATATCACCTTAGATGAAT

TAAGAATGATCACTCCAGTCTCTACATCCACCAACCTAGCCCACCGGTTGAGGGATCGGAGCACCCAGGTGAAATACTCG

GGGACATCCCTTGTGAGGGTTGCAAGATACACAACCATCTCCAATGATAACCTGTCATTTGTGATATCTGAGAAAAAAGT

GGACACCAACTTCATTTATCAACAAGGGATGCTGCTCGGTCTTGGGATCCTTGAGAACCTCTTTAGGCTAGAGGCCACCA

CCGGAGTGTCCAACACAGTGCTACACCTGCACGTGGAAACAGAATGTTGTGTTGTACCTATGGTGGATCACCCAAGGATA

CCGAGTCTCCGTAATATAAAAGTTACGGATGAGCTATGCACAAACCCTTTGATCTACGACCGGTCTCCCATTATAGAGCA

CGATGCAACCCGACTATACTCACAGAGCCACAGGAGACATTTGGTGGAGTTTGTTACATGGTCAACAAGCCAGCTTTACC

ATATATTGGCTAAGTCTACAGCAATGTCCATGATCGAGTTGATCACGAGATTCGAGAAAGATCACATGAATGAAATAGCC

GCCCTGATTGGCGATGACGACATCAATAGCTTCATCACAGAATTCTTGCTAGTAGAGCCTAGATTATTTATAGTTTACCT

TGGTCAGTGTGCTGCCATCAATTGGGCCTTTGATATACATTATCATCGGCCCTCAGGCAAGTACCAGATGGGGGAACTCC

TCTACTCTCTGCTCTCTCGTATGAGCAAAGGAGTATATAAGATCTTCACCAATGCTTTGAGCCACCCCAAAGTTTACAAG

AAATTTTGGCGAAGTGGTGTCATTGAGCCGATTCATGGCCCGTCTTTGGATACACAGAATTTACATGTCACTGTCTGTGA

CATGATATATGGGTCCTATGTCACCTATTTGGATCTTTTGCTGAATGATGAGCTAGATGATTATCCGTATTTGCTCTGCG

AGAGTGATGAGGACGTGGTCACAGACAGGTTCGACAACATTCAAGCCAAACATCTCTGTGTATTGGCCGATGTATATTGC

AGCTCCAAGAGGTGTCCTTCGATAATCGGGATGTCTCCTATAGAAAAATGTACCATTCTCACACATTACATCAAGGGAGA

ATCAGTACAATCCCCGTCTGGGACCTCATGGAACATTGATCCCCTTGTAGTAGATCATTACTCATGCTCTCTGACCTACC

TTCGACGCGGTTCCATCAAACAAATCAGGTTGAGAGTTGATCCGGGGTTTGTATTCGAGGCGTTGACAGACATCGACTTT

AAACAGCCTCGCAAAGCTAAGTTAGATATATCGGTCGTTGGGTTGACTGATTTCTCTCCCCCTTGCGACAACGTCGGTGA

TTTTCTAGGGACTATCAACACATTGAGGCACAATCTGCCTGTCACCGGAACCGGGGTCTCGAACTATGAAGTCCACGCTT

ATCGCAGAATTGGCCTGAATTCATCAGCATGTTATAAAGCCGTAGAGATCTCCACGTTAATCAAGCCATCCTTAGAAGTC

GGAGAGCATGGATTGTTCTTAGGAGAAGGTTCCGGTTCAATGCTGGCTGCGTACAAGGAAGTTCTTAAGTTAGCAAATTG

TTATTACAACAGCGGGGTAACAGCGGAGGGCAGAGCCGGACAGAGGGAAATATCTCCCTATCCCTCAGAGATGAGCCTAG

TAGAGAATCAGATGGGGATAGAGCGGAGTGTTAAAGTGCTGTTCAACGGCAAACCTGAAGTAACCTGGGTGGGGACCACC

GATTGCTACAAGTACATAATCAGTAACATCCAAACCTCTAGTCTAGGTTTCATACATTCAGATATTGAGACACTCCCAAC

CAAAGATGCTGTTGAGAAATTAGAAGAGTTTGCCTCTATCCTATCCTTATCCTTGATTTTGGGAAAAATCGGCTCTATCA

CAGTTGTCAAAATTATGCCCATTAGCGGGGATTTTACCCAAGGCTTCATAGCCTATGCCATCCAATATTTTAGGGAGAGC

CTGCTTGCCTATCCGAGATATAGTAACTTCATCTCGACTGAGTGTTACCTTATTATGATAGGATTAAAGGCCAATCGATT

GATAAACCCAGAAGCCATTAAGCAAAGCATAATCAGGGCAGGAGTCAGGACTGCACCAGGACTTGTGAGCCATATATTAT

CAGAGAAACAGAAAGGTTGTATTCAATCTTTCTTGGGTGATCCTTATATCCAAGGAGACTTCAATAAGCACCTTAAATCT

CTAACCCCTATTGAGAAAATCCTGGTAAATTGTGGTCTCTCTATCAATGGCACAAAAATCTGTAGAGATCTAATCCACCA

TGATATCGCCTCCGGTCCAGACGGTCTTATGAGCTCTACAATCATTTTGTATAGGGAACTGGCTCATTTCAAAGATAATA

TAAGGAGTCAGCACGGAATGTTCCACCCCTACCCAGTACTGGCCAACAGCAGGCAACGTGAATTAATCCTTCGAATAGCC

AAGAAATTCTGGGGGTATGTCTTGCTATATTCTGATGACCCGGCACTAATCAGACAAACAATCAAGAACTTGAAACGGAA

TCATCTAACCTTTGACTTACATAGTAACCCGTTTATTAAAGGGCTATCCAAAGCTGAGAAACTGCTGGTGCGGACGAGTT

CACTAAGAAGGGAATGGTTGTTCACTCTCGATACGAAAGAAGTGAAAGAGTGGTTCAAATTGGTGGGTTACAGTGCACTC

ATCAGAGGCTAGTTAGCTATACATCTGCTCCCTTCTCCTCCGCCATGAGCCCCCACTGACGATCCAAAAGATTAAAGAAA

ACTACATATTGGATAAGTTTCTATTCCCAGCTTTGTCTGGT

>KY885100.1/goat/Algeria/2015-11-11

ACCAAACAAAGTTGGGTAAGGATAGATCTTATAATAACTATAGACTGGCAAACTTAGGAGTAAAGATCCTACTGTCGGGG

GGAGGAGGAGGAGCAAGATCTCTGATTATGGCGACTCTCCTTAAAAGCTTAGCATTGTTCAAAAGGAACAAAGACAAAGC

GCCGACGGCATCAGGTTCAGGGGGGGCCATCCGGGGGATTAAGAATGTTATCATAGTCCCGATTCCCGGAGACTCTTCCA

TCATTACCCGTTCAAGACTGCTCGACAGGCTTGTCAGATTGGCCGGAGATCCTGACATCAACGGGTCAAAGCTGACCGGC

GTGATGATCAGCATGTTATCTTTGTTCGTAGAGTCTCCCGGGCAATTGATACAGCGAATCACAGATGATCCAGATGTTAG

TATCCGCCTTGTTGAGGTAGTTCAAAGTACTAGGTCCCAGTCCGGGTTGACCTTTGCATCACGTGGTGCTGATTTGGACA

ACGAGGCAGACATGTATTTTTCAACTGAGGGGCCCTCGAGTGGAGGTAAGAAAAGGATCAACTGGTTTGAGAACAGAGAA

ATAATAGACATAGAGGTGCAGGATCCAGAAGAGTTCAATATGTTGTTGGCCTCCATACTAGCACAAGTCTGGATCCTTCT

GGCCAAGGCTGTTACGGCACCGGATACGGCAGCTGACTCAGAATTGAGAAGGTGGGTTAAATACACACAACAGAGGAGAG

TGATTGGGGAATTTCGCCTTGACAAAGGGTGGCTGGACGCAGTCCGCAACAGAATTGCAGAAGATCTATCACTTCGGCGG

TTCATGGTGTCTCTCATACTTGACATCAAGAGGACCCCCGGCAACAAGCCAAGGATTGCAGAAATGATCTGCGACATTGA

CAACTATATTGTCGAAGCCGGACTCGCCAGTTTCATCCTTACTATCAAGTTTGGTATCGAAACCATGTATCCTGCATTAG

GGCTCCACGAGTTTGCCGGGGAATTGTCCACTATTGAATCCTTGATGAACTTGTATCAACAGCTAGGAGAGGTTGCACCC

TATATGGTAATTCTAGAGAACTCAATTCAGAACAAGTTTAGCGCAGGAGCTTATCCCCTCCTCTGGAGCTATGCGATGGG

TGTCGGAGTTGAGCTGGAGAACTCAATGGGGGGCTTGAACTTTGGCAGGTCATATTTTGACCCTGCCTATTTTCGTCTCG

GACAGGAGATGGTCAGAAGATCTGCTGGAAAGGTCAGCTCTGTGATTGCGGCTGAGCTCGGCATCACAGCAGAGGAAGCT

AAACTATTCTCGGAAATCGCCTCACAGACTGGGGACGAAAGAACCGCTAGAGGGACTGGGCCTCGACAGGCGCAGGTCTC

CTTCCTCCAGCACAAAACAGGAGAGGGAGATTCGCCCGCACCAACGACCAGAGAAGGGGTCAAAGCTGTGATCTCAAACG

GATCTGAAGAAAGGGATAGAAAGCAAACACGCCCAGGAAGGCCCAGAGGAGAGACCCCCGGTCAACTGCTCCTGGAAATC

ATGCCAGAGGACGAGGCTTCGCGAGAGTCTGGTCAAAACCCTCGTGAGGCTCAAAGATCAGCCGAGGCACTCTTCAGGCT

GCAGGCCATGGCCAAGATTCTGGAGGACCAGGAGGAGGGAGAAGACAACAGTCAGGTCTACAACGACAAGGATCTCCTCG

GCTAAGCAGACGCACCCTCTGTCGAAATCAGTGACAAGACATCTCCTACCAGTATTATAAAAAACTTAGGACCCAGGTCC

AAGCAGGCACACACCGACACTCCAATCAGTCGAGCGGAGACCACCGATGGCAGAAGAACAAGCATACCATGTCAACAAGG

GGCTGGAATGTATCAAATCTCTCAAAGCCTCTCCCCCGGATCTATCCACCATCAGAGATACCATCGAGAGCTGGAGAGAG

GGGCTTAGCCCCTCGGGCCGTGCAACACCGAACCCTGATGCGTCCGAGGGAGATCATCAGAATCTCAACCAATCATGCTC

ACCAGCAATCGGACCAAACAAAGTCTACCTTTCTTCTGAAGATAATCTCGGATTTAGAGAGATCACTGGCAACGACTGTG

AGGCTGGGCTCGGAGGAGTCCAGGGAAAAGGATCCAACTCTCAAGTACAGCGTTACTATGTTTATAGCCACGGGGGTGAA

GAGATTGAAGGACTCGAGGATGCTGACTCTCTCGTGGTTCAAGCAGATCCTCCAGTTGCTAACATCTTCAATGGAGGAGA

GGATGGATCTGACGACAGCGATGTGGACTCTGGCCCAGATGATCCCGGCAGAGATACTCTATATGACCGGGGATCTGTTG

CCGGCAATGATGTCGCTAGGTCCACAGATGTCGAAAAACTAGAAGGTGCTGACATTCAAGAAGTTCTTAACTCCCAGAAA

AGCAAAGGAGGAAGATTCCAAGGCGGGAAAACCTTACGAGTCCCGGAAATACCCGATGTCAAGCACTCCAGACCATCTGC

CCAATCAATTAAAAAGGGCACAGACGGGAGCTCAGTCTCATCTGGAACGGTGACAGAGTGTTTATCGATAAGTGGTGCAA

CCCAAGCTGTGCCAGAGTCAAGATGGGAGTCATCAGAGCAAAATGCGTCTGTGGGGAGTGTCCTCAAGTCTGCGAAGAGT

GCAAAGACGATCCAGGGGTTGACACAAGAATCTGGTACCATAGCATCACTGACTCAGCCTAAAGAGAATGATTCCGAGTA

TGAGTATGAGGATGACCTATTTACAGAGATTCAGGACATCCGTGCAAGCATTGCCAAGATCCATGATGACAACAAAACTA

TCCTCTCAAAGCTTGATTCTATACTGCTATTGAAAGGAGAAGTCGACACTATCAAGAAACAAATCAGCAAGCAGAATATA

AGTATATCCACCATTGAGGGCCATCTCTCCAGTATAATGATAGCCATCCCGGGCTTTGGGAAGGATATCAAGGACCCAAC

ATCCGAGGTTGAGTTGAACCCAGATTTAAGACCTATAATAAGCCGTGACTCCGGCAGAGCTCTCGCGGAGGTCCTCAAGA

AACCCGCTGTGGACAGGTCTCAGAAAATTGGAACCAAAGCCAACTCCAGCTCAAAGGGTCAGCTTCTTAAGGATCTCCAG

CTAAAACCTGTCGACAAGCAAGCAAGCTCTGCAATCGGGTTTGTCCCATCCGACCATGAATCATCCAGAAATGTCATCCG

CTCCATAATCAAGTCGAGCAAGCTAAACATTGATCACAAGGACTATCTTCTAGATTTACTAAATGATGTGAAAGGCTCCA

AGGATCTTAAGGAATTCCACAAGATGCTAACAGCAATTCTCGCCAAGCACCCGTAACACACCCTCCAGTCACCATCTCAT

ACTCGGCTGAAAACACCCTCTCAATCAGGCTATTACAAAAAACTTAGGAGCAAGGGCAACTGAGCTTCGCAGACAAGATG

ACCGAGATCTACGACTTCGATAAATCAGCATGGGATGTCAAAGGGTCAATTGCCCGCATAGAACCCACCACCTATCACGA

CGGCCGACTGATACCCCAGGTAAGGGTCATCGATCCTGGTCTGGGAGACAGAAAAGATGAGTGCTTCATGTACCTGTTTC

TCCTAGGAGTGATTGAGGATAACGACCCCCTGTCTCCCCCAGTCGGAAGAACCTTTGGTTCTTTACCTCTAGGGGTCGGT

AGGTCAACTGCTAAGCCAGAAGAACTACTAAGGGAGGCCACAGAACTAGATATAGTGGTGAGGCGCACTGCAGGATTAAA

TGAGAAACTGGTATCTTACAACAACACTCCGCTGTCCTTGTTAACGCCCTGGAAGAAGGTTCTGACAACCGGAAGTGTGT

TTAGCGCTAACCAGGTTTGCAATGCAGTCAACCTAGTCCCGCTTGATACTCCCCAGAGATTCAGGGTTGTGTACATGAGC

ATAACTAGATTATCAGACAATGGTTACTATAGTGTGCCCAGAAGAATGCTGGAGTTCCGCTCCGCCAATGCAGTAGCCTT

CAACATCTTGGTTACACTAAGAATTGAAAATGGCACAAACCCTAGAAGATACATAGTCGGCTCATGGGAGAACCCAGAGG

TCACATTTATGGTACACGTGGGCAACTTTAGAAGAAAGAAGAACGAAGTATACTCTGCTGATTACTGCAAAATGAAGATT

GAAAAGATGGGTTTAGTTTTTGCCCTGGGAGGAATAGGTGGAACAAGTCTCCATATTAGAAGCACCGGGAAAATGAGCAA

AACCCTCCATGCACAGCTGGGGTTCAAGAAAATCTTATGTTACCCCCTAATGGATGTTAATGAGGATCTTAACCGATATC

TTTGGCGGGCAGAGTGCCGAATAGTCAAAATCCAAGCCGTCTTACAGCCATCAGTGCCCCAAGAATTCCGTGTCTACGAT

GATGTCATCATCAACGATGATCAAGGCCTGTTCAAGATCTTGTAGTTCATTTGCAAAGGAATGAACCCACCCCCAGGAAG

GGGCCCTACCCTCCTCCG-CCAAAAAACACACTCCAAGGCTCCCCGACATAGCGCCAACACAGCCACTCTCCCGATGGGA

CGAGGCTGG-G-CCCCGAGCAAACCCGCCCCAGCCGGGGGCCCCCCCCCCCGACCCGCTGGGCGCCGACCGAGCAGGTCG

GCGAAAAGCCCACAACAGGCCCCGACGAAGGTGGAGCACCCCATCCCCCC-AACACCCCTCCTC-AACACGCCCGAGGCC

AGGCGCCGGGCACCCCCAGACCCGCCACCGGCGAGACAGAACCCCATGGGGGGACCTCCCAAACCAAAGGGGCCACCGGG

GAAGGCTCCGCCGAACCAGCACGGACCCGACCCAAGCAAGTCGGACCTCCAACACGAAACCGCCCCGCGCACCACACTCA

TCATCAAACAAAACTTAGGGGCCAAGTCCACACGACAGGCCCCAGCCCGCCAGGCGGAGGGCCGCAGAAAGGAAGGAGAC

ACGACCGTCCCAGAAGGAAGACACCCAACCACCGAAACACCCAGCCCCAGGGAAGGAG--CCCCCCCCCACACGGCGAAG

GAACGGGTGAGAGACCCCGAGGAGAACCCGACCGGAAGCCAGGAGCCGGCGACGCCGGTGGCAAACAGGCAGGGCATACA

CCCGACTCCACCAGAGGAGCCGAGGAGAGCCTGACCCCGCGTGGACCAAACGGCAGGGAATCCAACCCCGCCACAAAAAG

AAGTGAGGCTCT-CCT-----CCCCCCCGCACACAACCCGCAGACAGACCAAGGGAGGACCACCTTCCAACAAGACAGGC

GGCTCCCTCGCAGCCGAGCCGCAGCTCCCCAGAAGAGGACACTCCCGCACACTCCGAACACCCCCCCGCCCCCCGCCACC

GGGACAGGGGCTCCGCCACCACAGACCAGAGCCACCGAGAGCCAGCCAGGGGTCCCCTCCAGTCCCGCTCAGCACCCCCC

CGAGGAGGCCCCGGCGAGGCCCCCAAGCCCGACGACCCCTGCACGCACCGAAAGAGGGACCCGCCCGAGAACGTCCTGCA

CATCCATGCGTAAACATCATGACACGGGTCGCAACCTTGACATTTCTGTTTCTCTCCCTAAATGCTGTTGCGTGCCAGAT

TCACTGGGGCAATCTATCCAAGATCGGGATTGTAGGCACAGGGAGTGCCAGCTACAAGGTGATGACTAGGCCAAGCCACC

AAACTCTGGTTATAAAGTTAATGCCAAATATAACAGCCATCGACAATTGTACAAAGTCAGAGATTGCAGAGTACAAGAGA

TTGCTGATCACAGTGTTAAAGCCTGTAGAGGATGCTCTGTCAGTGATAACCAAGAATGTAAGACCAATTCAAACTCTGAC

ACCTGGGCGTAGAACCCGCCGTTTTGCTGGAGCTGTTCTGGCCGGAGTAGCACTTGGAGTTGCGACAGCCGCTCAGATAA

CTGCAGGAGTCGCCCTTCATCAATCATTGATGAACTCCCAAGCAATTGAGAGTTTAAAAACCAGTCTTGAGAAGTCGAAT

CAGGCAATAGAAGAAATCAGACTTGCAAATAAGGAGACCATACTGGCAGTACAGGGCATCCAGGATTATATCAACAATGA

GCTTGTCCCTTCTGTCCATAGAATGTCATGCGAGCTGGTAGGCCACAAGCTCGGCCTCAAGCTCCTTAGGTACTACACCG

AGATCCTGTCCATATTCGGGCCCAGCCTTCGAGACCCGATAGCTGCCGAAATATCAATCCAGGCACTCAGTTATGCATTA

GGCGGAGACATCAATAAAATCCTGGACAAGCTTGGGTATAGCGGTGGGGATTTCCTTGCCATCCTAGAAAGTAAGGGAAT

AAAGGCCCGAGTCACATATGTGGACACAAGAGATTACTTTATAATTCTTAGCATCGCCTACCCAACCTTATCTGAGATCA

AGGGAGTGATAGTTCACAAGATAGAAGCTATAACATACAACATTGGGGCACAGGAGTGGTATACTACTATCCCTAAATAT

GTAGCTACTCAGGGGTATCTGATATCGAACTTTGATGAGACGTCATGCGTATTCACTCCAGAGGGGACAGTTTGCAGCCA

GAATGCGTTGTACCCAATGAGCCCGTTGCTTCAGGAATGTTTCAGGGGGTCAACAAAATCGTGCGCCAGAACCCTAGTTT

CAGGGACCATAAGTAATAGATTTATCCTATCAAAAGGGAACCTGATTGCCAATTGCGCGTCAGTTTTGTGCAAATGTTAC

ACAACGGAGACAGTTATCAGCCAAGATCCTGACAAACTACTAACTGTTGTAGCATCCGACAAGTGTCCTGTAGTTGAGGT

GGACGGAGTGACAATACAGGTCGGCAGTCGAGAGTATCCGGATTCTGTATACTTACACAAGATAGATTTAGGTCCAGCCA

TCTCCCTAGAAAAACTGGATGTAGGCACCAATTTAGGCAATGCAGTCACAAGACTGGAGAATGCAAAGGAGCTCCTAGAT

GCATCAGACCAAATACTGAAGACTGTTAAAGGGGTACCTTTCAGCGGGAATATGTACATAGCACTGGCAGCTTGCATTGG

GGTATCCCTAGGGCTTGTCACATTAATATGCTGCTGTAAGGGGAGGTGCAAGAACAAGGAAATCCCCATCTCCAAAATCA

ACCCAGGGCTCAAGCCCGACCTGACCGGGACCTCAAAGTCGTACGTGAGATCACTGTAGTCAGAATCACCCGAATCATCC

GGCATCACACACATACATGCACGACACAAGCAGTCAGAGGACGCAGGAAGCCCAGTCTCCGATCACCCACCTGACCCTAC

TCCACGCTCCACTACACATTAGTCATCAAACAAAACTTAGGACGAAAGGTCAATCACCATGTCCGCACAAAGGGAAAGGA

TCAATGCCTTCTATAAGGACAATCCTCACAATAAGAACCATAGGGTGATCCTGGATAGAGAACGCTTGGTCATTGAAAGA

CCCTACATCTTGCTAGGAGTCCTGCTGGTAATGTTCCTGAGTCTAATCGGACTGCTGGCCATTGCAGGGATCAGGCTTCA

CCGGGCCACCGTTGGAACTTCAGAGATCCAGAGTCGGCTGAATACCAATATTGAGTTGACTGAATCTATTGATCACCAAA

CTAAGGATGTCTTAACTCCCCTTTTTAAAATCATTGGCGATGAAGTCGGCATCAGAATTCCACAGAAATTCAGTGATCTT

GTCAAGTTCATCTCCGATAAGATTAAATTCCTCAACCCTGATAGAGAGTATGATTTCAGGGATCTCCGGTGGTGTATGAA

TCCCCCCGAAAGAGTCAAAATTAATTTTGATCAGTTCTGTGAGTACAAAGCTGCGGTTAAGTCAATTGAACATATATTTG

AGTCACCACTCAACAAGTCAAAAAAGTTGCAATCATTGACTCTTGGGCCCGGAACAGGCTGTCTAGGCAGGACAGTAACA

AGAGCTCGTTCCTCAGAACTTACACTGACCTTAATGGACCTGGATCTAGAGATGAAGCACAACGTGTCCTCAGTGTTTAC

CGTAGTTGAAGGGGGATTATTCGGAAGAACGTATACAGTCCGGAGATCCGATGCCAGGGATCCTAGCACCGATCTAGGTA

TTGGCCATTTTTTAAGAGTCTTCGAGATTGGACTGGTAAGAGATCTCGGGCTTGGTCCCCCTGTTTTTCAGATGACCAAC

TATCTCACAGTGAACATGAGTGATGACTATCGGAGATGTCTCTTAGCGGTAGGGGAGTTGAAGTTGACAGCCCTATGCAC

CTCATCTGAGACTGTGACACTGAGTGAGAGAGGAGTTCCAAAGAGGGAGCCTCTTGTGGTTGTGATACTTAACCTAGCTG

GGCCCACTCTAGGGGGCGAACTATACAGTGTATTGCCTACCTCTGATCTCATGGTGGAGAAACTCTATTTATCTTCTCAT

AGAGGGGTTATCAAAGATGACGAGGCCAATTGGGTAGTGCCGTCTACCGATGTTCGTGATCTTCAAAACAAAGGAGAATG

TCTGGTGGAAGCATGCAAGACTCGACCTCCCTCATTTTGCAATGGCACGGGATCAGGCCCGTGGTCAGAGGGGAGAATCC

CTGCTTACGGGGTGATCAGGGTCAGTCTTGACTTAGCTACTGACCCGGGTGTAGTTATCACTTCAGTGTTTGGCCCACTG

ATACCTCACCTATCCGGCATGGATCTTTACAACAACCCGTTTTCAAGAGCTGTATGGTTGGCTGTACCACCTTATGAGCA

GTCATTTCTAGGAATGATAAATACAATTGGATTCCCCAACAGAGCAGAGGTTATGCCGCACATTTTGACCACAGAGATCA

GAGGCCCTCGGGGTCGTTGCCATGTTCCCATAGAATTGTCCCGCAGGGTTGATGACGATATCAAGATCGGGTCCAACATG

GTCATATTGCCGACGATGGACCTGAGGTATATTACAGCCACTTATGATGTTTCCAGGAGCGAGCATGCAATCGTGTATTA

TATATATGACACAGGTCGCTCATCATCTTACTTCTACCCAGTTCGACTGAATTTCAAAGGCAATCCTCTCTCTCTGAGGA

TAGAGTGTTTCCCTTGGCGTCATAAGGTGTGGTGCTACCATGATTGTCTCATATACAACACCATAACAGATGAAGAAGTC

CATACGAGAGGGCTGACCGGTATAGAGGTAACATGTAATCCAGTCTGAGCAGAGCTACGACCGTCACTCAGGCAGTCCCT

TGAGTCGTCACTGAGTCCAAGCAGCACAGCCTGGGATACTCAACAGCACAGCCCAGCCAACAATGTTATAAAAAACTTAG

GAGCCAGGGTTGTAGGGGCCATGGACTCTCTATCAGTCAATCAGGTCCTGTACCCTGAGGTTCATCTAGATAGCCCTATT

GTCACAAACAAACTAGTTGCCATCCTTGAGTACTCGGGTATCGACCATAACTATGTTCTCGAAAACCAGACCCTTGTCAA

GAATATTAGGTATAGACTGGGGTGCGGTTTTTCAAATCAAATGATCATCAATAACAGGGGGGTGGGTGAAACAGTCAATT

CTAAACTTAAAAGTTACCCCCATAATCGTCATATCATATACCCGGATTGCAATAAGGAGTTGTTTTGTATCAAAGATAGC

TGCATATCTAAGAAGCTCTCGGAGCTATTCAAGAAGGGTCATTCCTTGTACTCTAAGATAAGTCACCAGGTACTGGACTG

TCTCAAGAGAGTCAATGGGAAATTAGGCCTGGGCACAGATCTTACCCATGGTCTGAAGGAGGGTATCCTTGACCTAGGGT

TGCACATGCATAGCTCTCAATGGTTCGAGACTTTCCTGTTCTGGTTCACTATCAAGACAGAGATGAGATCAATGATCAAA

GAACAGTCCCATATATGCCACAAGAGGAGGTATAACCCGATTTTTGTGTCAGGGGATGCATTCGAGGTGCTCGTATCACG

GGACCTCGTTGTGATAATTGATAAGAATACCCAGTATGTCTTCTACTTAACATTTGAGCTGGTCCTTATGTATTGTGATG

TCATAGAGGGCAGACTTATGACGGAGACAGCCATGGCCATAGACCAGAGATATTCAGAACTTCTAAGCCGGGTCAGATAC

TTGTGGGATCTTATTGATGGGTTCTTCCCAACACTAGGCAACACCACATACCAAGTTGTTGCTCTGCTTGAACCATTATC

GTTGGCTTATCTTCAACTTCAGGATGTCACTCTGGAGTTAAGAGGTGCTTTTTTAGACCATTGCTTCAAAGAACTCTATG

AGATACTGGAGCATTGCGGCATTGACACAGAAGGCACCTACAATTCCATCACTGAAGGAATAGATTACGTATTCATCACC

CACGATATACATCTAACTGGGGAGATTTTTTCATTTTTTCGGAGTTTCGGACACCCCCGTCTAGAAGCAGTCACCGCTGC

AGAAAATGTCAGGAAACATATGAACCACCCGAAGGTAATCAGTTATGAGACTATGATGAAAGGGCATGCAGTATTTTGCG

GGATAATCATAAATGGTTTTAGGGACCGACATGGCGGCAGCTGGCCCCCTGTTGCATTGCCAGAGCATGCTTCTGCTGCG

ATTCGGAATGCGCAGGCATCAGGTGAGGGGCTGACCCATGACCTGTGTATAGACAATTGGAAATCCTTTGTTGGATTCAA

ATTTGGCTGCTTTATGCCACTCAGCCTAGATAGTGATTTGACCATGTACCTCAAAGACAAGGCGTTGGCTGCACTGAAGA

ATGAGTGGGATTCAGTTTACCCGAAAGAATACCTCCGTTATAACCCACCTAGAGGGACAGAGTCAAGGCGACTGGTGGAG

GTGTTCCTAAATGACTCCAGCTTTGATCCTTATAACATGATAATGTACGTGGTGAATGGCTCCTACCTTAAAGACCCTGA

GTTTAATCTCTCATACAGCCTAAAAGAGAAGGAGATAAAGGAGACAGGGCGGTTGTTTGCTAAAATGACTTATAAGATGC

GGGCCTGTCAGGTAATCGCTGAAAATCTGATATCGAATGGTGTTGGGAAGTATTTCCGAGACAATGGGATGGCAAAGGAT

GAGCACGACCTAACAAAAGCCCTTCACACCCTGGCAGTCTCAGGTGTTCCCAAGAACAACAAAGATAACCACCGAGGTGG

GCCTCCCAGAAGGACAACAAGCCGAGAGGTGAGATCAAACCAAGACACCAAAACACAGAATAGAGACAAGGTCCAAGGGG

GGCCTATGTACAACTATTTGCGATGCCAACCAATCAGCCCTGATCAGGGTGAGTCATACGAGACTGTTAGTGCATTTATC

ACCGCTGACCTTAAGAAGTATTGCCTGAATTGGAGATACGAGACAATCAGCATATTTGCACAGAGGTTGAATGAAATATA

TGGACTACCATCCTTCTTTCAATGGTTACACAGGATATTGGAAAAATCCGTACTCTACGTCAGTGACCCACATTGCCCTC

CCGACCTAGATAATCATATCCCTCTGGACAGTGTCCCTAATGCCCAAATATTCATTAAGTACCCAATGGGCGGAATAGAA

GGTTATTGCCAAAAACTATGGACAATCAGTACTATACCATACTTGTATCTGGCAGCCTATGAGAGCGGAGTAAGAATCGC

CTCACTAGTGCAGGGTGACAATCAAACAATTGCAGTGACAAAAAGAGTTCCAAGTTCTTGGCCTTATTCACTAAAAAAGA

GGGAGGCATCCAAAGCCGCTCAAAACTACTTCGTAGTCTTAAGGCAAAGATTGCACGATGTAGGTCATCACTTGAAGGCT

AATGAAACCATAGTATCTTCTCACTTTTTTGTATATTCCAAGGGGATTTATTATGACGGCCTGCTAGTCTCACAATCACT

GAAGAGCATCGCCAGATGTGTCTTCTGGTCCGAGACTATAGTGGATGAAACCAGAGCGGCTTGCAGCAATATTGCAACAA

CTGTCGCTAAGAGTATAGAGAGGGGTTATGATAGGTACCTTGCATATTCTTTGAATATCCTCAAGATTTTCCAACAGATC

CTTATATCCCTTAACTTCACTATTAACACAACAATGACTCAGGATGTCGTGGCACCGATCATCGAGAACGGTGATTTGCT

GATAAGGATGGCACTCTTGCCAGCACCCATCGGGGGTCTCAATTATCTTAACATGAGCAGGTTATTTGTGAGAAATATCG

GTGACCCGGTCACCTCCTCTATAGCCGACCTGAAGAGGATGATAGATGCTGGGCTAATGCCAGAAGAAACACTGCATCAA

GTGATGACCCAGACCCCGGGAGAATCATCCTACCTAGATTGGGCAAGTGACCCTTATTCTGCCAACCTACCCTGCGTACA

GAGTATAACTCGCCTTCTCAAAAACATCACTGCACGATATATTTTAATCAGCAGCCCAAACCCAATGCTGAAAGGATTGT

TTCATGAGGGGAGTAGAGATGAGGACGAGGAGCTTGCAAGTTTCTTAATGGATCGGCATATAATTGTTCCAAGAGCTGCA

CATGAAATCTTAGACCATAGCATAACCGGGGCAAGAGAAGCTATAGCCGGGATGTTGGATACCACCAAGGGTCTGATTAG

AACAAGTATGAGACGGGGTGGCCTCACCCCTCGAGTCTTAGCCCGCCTTTCCAATTATGATTATGAACAATTCCGATCCG

GGATAACACTATTGACAAAGAAAGGGCAGTGTTATCTCATTGACAAGGACTCGTGCTCGGTGCAGCTCGCTATAGCTCTG

AGGAGCCACATGTGGGCTAGGTTGGCTCGCGGGAGACCAATCTATGGGTTGGAGGTGCCCGATATATTAGAATCGATGAA

CGGCTACCTTATCAAACGCCACGAGTCTTGTGCTATCTGTGAAACGGGCTCAAGTCACTACGGATGGTTTTTCGTCCCTG

CAGGGTGCCAGCTTGACGATGTATCTAGAGAGACTTCGGCTCTTCGTGTACCTTATGTCGGATCAACAACTGAGGAAAGG

ACAGACATGAAACTTGCTTTTGTTAGATCTCCAAGTCGATCCCTCAAATCAGCAGTCAGAATTGCAACAGTTTACTCATG

GGCCTACGGGGATGATGAGAAATCATGGAGTGAAGCCTGGATGCTAGCTAGGCAGAGAGCCAATATCACCTTAGATGAAT

TAAGAATGATCACTCCAGTCTCTACATCCACCAACCTAGCCCACCGGTTGAGGGATCGGAGCACCCAGGTGAAATACTCG

GGGACATCCCTTGTGAGGGTTGCAAGATACACAACCATCTCCAATGATAACCTGTCATTTGTGATATCTGAGAAAAAAGT

GGATACCAACTTCATTTATCAGCAAGGGATGCTGCTCGGTCTTGGGATCCTTGAGAACCTCTTTAGGCTAGAGGCCACCA

CCGGAGTGTCCAACACAGTGCTACACCTGCACGTGGAAACAGAATGTTGCGTTGTACCTATGGTGGATCACCCTAGGATA

CCGAGTCTCCGTAATATAAAAGTTACGGATGAGCTATGCACAAACCCTTTGATCTACGACCGGTCTCCCATCATAGAGCA

CGATGCAACCCGACTATACTCACAGAGCCACAGGAGACATTTGGTGGAGTTTGTTACCTGGTCAACAAGCCAGCTTTACC

ATATACTGGCTAAGTCTACAGCAATGTCCATGATCGAGTTGATCACGAGATTCGAGAAAGATCACATGAATGAAATAGCC

GCCCTGATTGGCGATGACGACATCAACAGCTTCATCACAGAATTTTTGCTAGTAGAGCCTAGATTATTTATAGTTTACCT

TGGTCAGTGTGCTGCCATCAATTGGGCCTTTGATATACACTATCATCGGCCCTCGGGCAAGTACCAGATGGGGGAACTCC

TCTACTCTCTGCTCTCTCGGATGAGCAAAGGAGTATATAAGATCTTCACCAATGCTCTGAGCCACCCCAAAGTTTACAAG

AAATTTTGGCGAAGTGGTGTCATTGAGCCGATTCATGGCCCATCCCTGGATACACAGAATCTACATGTCACTGTCTGTGA

CATGATATATGGGTCCTATGTCACCTATTTGGATCTTTTGCTGAATGATGAGCTAGATGATTATCCGTATTTGCTCTGCG

AGAGTGATGAGGACGTGGTCACAGACAGGTTCGACAACATTCAAGCCAAACATCTCTGTGTATTGGCCGATGTATATTGC

AGCTCCAAGAGGTGTCCTTCGATAATCGGGATGTCTCCTATAGAAAAATGTACCATTCTCACACATTACATCAAGGGAGA

ATCAGTACAATCCCCGTCTGGGACCTCATGGAACACTGATCCCCTTGTAGTAGATCATTACTCATGCTCTCTGACCTACC

TTCGCCGCGGTTCCATCAAACAAATCAGGTTGAGAGTGGATCCGGGGTTTGTATTCGAGGCGTTGACAGACGTCGACTTT

AAACAGCCTCGCAAAGCTAAGCTAGGTATATCGGTCGTTGGGTTGACTGATTTCTCTCCCCCTTGCGACAACGTCGGTGA

TTTTCTAGGGACTATCAACACATTGAGGCACAATCTGCCTGTCACCGGAACCGGGGTCTCGAACTATGAAGTCCACGCTT

ATCGTAGAATTGGCCTGAATTCATCAGCATGTTATAAAGCCGTAGAGATCTCCACGTTAATCAAGCCATACTTAGAAGTC

GGAGAGCATGGATTGTTCTTAGGAGAAGGTTCCGGTTCAATGCTGGCTGCGTACAAGGAAGTTCTTAAATTAGCAAATTG

TTATTACAACAGCGGGGTAACAGCGGAGGGCAGAGCCGGACAGAGGGAAATATCTCCCTATCCCTCAGAGATGAGCCTAG

TAGAGAATCAAATGGGGATAGAGCGGAGTGTTAAAGTGCTGTTCAACGGCAAACCTGAAGTAACTTGGGTGGGGACCACC

GATTGCTACAAGTACATAATCAGTAACATCCAAACCTCTAGTCTGGGTTTCATACATTCAGATATTGAGACACTCCCAAC

CAAAGATGCTGTTGAGAAATTAGAAGAATTTGCCTCTATCCTATCCTTATCCTTGATTTTGGGAAAAATCGGCTCTATTA

CAGTTGTCAAAATTATGCCCATTAGCGGGGATTTTACCCAAGGCTTCATAGCCTATGCCATCCAATATTTTAGGGAGAGC

CTGCTTGCCTATCCGAGATATAGTAACTTCATCTCGACTGAGTGTTACCTTATTATGATAGGATTAAAGGCCAATCGATT

GATGAACCCAGAAGCCATTAAGCAAAGCATAATCAGAGCGGGGGTCAGGACTGCACCAGGACTTGTGAGCCATATATTAT

CAGCGAAACAGAAAGGTTGTATTCAATCTTTCTTGGGTGATCCTTATATCCAAGGAGACTTCAATAAGCACCTTAAATCT

CTAACCCCTATTGAGAAGATCCTGGTAAATTGTGGTCTCTCTATCAATGGCACAAAAATCTGTAGAGATCTAATCCACCA

TGATATCGCCTCCGGTCCAGACGGTCTCATGAGCTCTACAATCATTTTATATAGGGAATTGGCTCATTTCAAAGATAATA

TAAGGAGTCAGCACGGAATGTTCCACCCCTACCCAGTACTGGCCAACAGCAGGCAACGTGAATTAATCCTTCGAATAGCC

AAGAAATTCTGGGGGTATGTCTTGCTATATTCTGATGACCCGGCACTAATCAGACAAACAATCAAGAACTTGAAACGGAA

TCATCTAACCTTTGACTTACATAGTAACCCGTTTATTAAAGGGCTATCCAAAGCTGAGAAACTGCTGGTGCGGACGAGTT

CACTAAGAAGGGAATGGTTGTTCACTCTCGATACGAAAGAAGTGAAAGAGTGGTTCAAATTGGTGGGTTACAGTGCACTC

ATCAGAGGCTAGTTAGCTATACATCTGCCCCCTTCTCCTCCGCCATGAGACCCCACTGACGATCCAAAAGATTAAAGAAA

ACTTCATATTGGATAAGTATCTATTCCCAGCTTTGTCTGGT

>KC594074.1/goat/Morocco/2008

ACCAAACAAAGTTGGGTAAGGATAGATCTTATAATAACTATAGACTGGCAAACTTAGGAGTAAAGATCCTACTGTCGGGG

GGAGGAGGAGGAGGAAGATCTTTGATTATGGCGACTCTCCTTAAAAGCTTAGCATTGTTCAAAAGGAACAAAGACAAAGC

GCCGACGGCATCAGGTTCAGGGGGGGCCATCCGGGGGATTAAGAATGTTATCATAGTCCCGATTCCCGGAGACTCGTCCA

TCATTACCCGTTCAAGACTGCTCGACAGGCTAGTCAGATTGGCCGGAGATCCTGACATCAACGGGTCAAAGCTGACCGGC

GTGATGATCAGCATGTTATCTTTGTTCGTAGAGTCACCCGGGCAATTGATACAGCGAATCACAGATGATCCAGATGTTAG

TATCCGCCTTGTTGAGGTAGTTCAAAGTACTAGGTCCCAGTCCGGGTTGACCTTTGCATCACGTGGTGCTGATTTGGACA

ACGAGGCAGACATGTATTTTTCAACTGAGGGGCCCTCGAGTGGAAGTAAGAAAAGGATCAACTGGTTTGAGAACAGAGAA

ATAATAGACATAGAGGTGCAGGATCCAGAAGAGTTCAATATGTTGTTGGCCTCCATACTAGCACAAGTCTGGATCCTTCT

GGCCAAGGCTGTTACGGCACCGGATACGGCAGCTGACTCAGAACTGAGAAGGTGGGTTAAATACACACAACAGAGGAGAG

TGATTGGGGAATTTCGCCTTGACAAAGGGTGGCTGGACGCAGTCCGCAACAGAATTGCAGAAGATCTATCACTTCGGCGG

TTCATGGTGTCTCTCATACTTGACATCAAGAGGACCCCCGGCAACAAGCCAAGGATTGCAGAAATGATCTGCGACATTGA

CAACTATATCGTCGAAGCCGGACTCGCCAGTTTCATCCTGACTATCAAGTTTGGTATCGAAACCATGTATCCTGCATTAG

GGCTCCACGAGTTTGCCGGGGAATTGTCCACTATCGAGTCCTTGATGAACTTGTATCAACAGCTAGGAGAGGTTGCACCC

TACATGGTAATTCTAGAGAACTCAATTCAGAACAAGTTTAGTGCAGGAGCTTATCCCCTCCTCTGGAGCTATGCGATGGG

TGTCGGAGTTGAGCTGGAGAACTCAATGGGGGGCCTGAACTTTGGCAGGTCATATTTTGACCCTGCTTATTTTCGTCTCG

GACAGGAGATGGTCAGAAGATCTGCAGGAAAGGTCAGCTCTGTGATTGCGGCTGAGCTCGGCATCACAGCAGAGGAAGCT

AAACTAGTCTCGGAAATCGCCTCGCAGGCCGGGGACGAAAGAACCGCCAGAGGGACTGGGCCTCGACAGGCGCAGGTCTC

CTTCCTCCAGCACAAAACAGGAGAGGGAGATTCGCCCACACCAGCGACCAGAGAAGGGGTCAAAGCTGCGATCTCAAACG

GACCTGAAGAAAGGGATAGAAAGCAAACACGCCCAGGAAGGCCCAGAGGAGAGACCCCCGGTCAACTGCTCCTGGAAATC

ATGCCAGAGGATGAGGTTCCGCGAGAGTCTGGTCAAAACCCTCGTGAGGCTCAAAGATCAGCCGAGGCACTCTTCAGGCT

GCAGGCCATGGCCAAGATTCTGGAGGACCAGGAGGAGGGAGAAGACAACAGTCAGGTCTACAACGACAAGGATCTCCTCG

GCTAAGCAGACGCACCCTCTGTCGAAATCAGTGATAAGACATCTTCTACCAGTATTATAAAAAACTTAGGACCCAGGTCC

AAGCAAGCACACACCGACACTCCAATCAGTCGAGCGGAGACCACCGATGGCAGAAGAACAAGCATACCATGTCAACAAGG

GGCTGGAATGTATCAAATCTCTCAAAGCCTCTCCCCCGGATCTATCCACCATCAGAGATACCATCGAGAGCTGGAGAGAG

GGGCTTAGCCCCTCGGGCCGTGCAACACCGAACCCTGATGCGTCCGAGGGAGATCATCAGAATCTCAACCAATCATGCTC

ACCAGCAATCGGACCAAACAAAGTCTACTTTTCTTCTGAAGATAATCTCGGATTTAGAGAGATCACTGGCAACGACTGTG

AGGCTGGGATCGGAGGAGTCCAGGGAAAAGGATCCAACTCTCAAGTACAGCGTTACTATGTTTATAGCCACGGGGGTGAA

GAGATTGAAGGACTCGAGGATGCTGACTCTCTCGTGGTTCAAGCAGATCCTCCAGTTGCTAACATCTTCAATGGAGGAGA

GGATGGATCTGACGACAGCGATGTGGACTCTGGCCCAGATGATCCCGGCAGAGATACTCTATATGACCGGGGATCTGTTG

CCGGCAATGATGTCGCTAGGTCCACAGATGTCGAAAAACTAGAAGGTGCTGACATTCAAGAAGTTCTTAACTCCCAGAAA

AGCAAAGGAGGAAGATTCCAAGGCGGGAAAACCTTACGAGTCCCGGAAATACCCGATGTCAAGCACTCCAGACCATCTGC

CCAATCAATTAAAAAGGGCACAGACGGGAACTCAGTCTCATCTGGAACGGTGACAGAGTGTTTATCGATAAGTGGTGCAA

CCCAAGCTGTGCCAGAGTCAAGATGGGAGTCATCAGAGCAAAATGCGTCTGTGGGGAGTGTCCTCAAGTCTGCGAAGAGT

GCAAAGACGATCCAGGGGTTGACACAAGAATCTGGTACCATAGCATCACTGACTCAGCCTAAAGAGAATGATTCCGAGTA

TGAGTATGAGGATGACCTATTTACAGAGATTCAGGACATCCGTGCAAGCATTGCCAAGATCCATGATGACAACAAAACTA

TCCTCTCAAAGCTTGACTCCATACTGCTATTGAAAGGAGAAGTCGACACTATCAAGAAACAAATCAGCAAGCAGAATATA

AGTATATCCACCATTGAGGGCCATCTCTCCAGTATAATGATAGCCATCCCGGGCTTTGGGAAGGATATCAAGGACCCAAC

ATCCGAGGTTGAGTTGAACCCAGATTTAAGACCTATAATAAGCCGTGACTCTGGCAGAGCTCTCGCGGAGGTCCTCAAGA

AACCCGCTGTGGATAGGTCTCAGAAAATTGGAACTAAAGCCAACTCCAGCTCAAAGGGTCAGCTTCTTAAGGATCTCCAG

CTAAAACCTGTCGACAAGCAAGCAAGCTCTGCAATCGGGTTTGTCCCATCCGACCATGAATCATCCAGAAATGTCATCCG

CTCCATAATCAAGTCGAGCAAGCTAAACATTGATCACAAGGACTATCTTCTAGATTTGCTAAATGATGTGAAAGGCTCCA

AGGATCTTAAGGAATTCCACAAGATGCTAACAGCAATTCTCGCCAAGCACCCGTAACACACCCTCCAGTCACCATCCCGC

ACTCGGCTGAAAACACCCTCTCAATCAGGCTATTACAAAAAACTTAGGAGCAAGGGCAACTGAGCTTCGCAGACAAGATG

ACCGAGATCTACGACTTCGATAAATCAGCATGGGATGTCAAAGGGTCAATTGCCCGCATAGAACCCACCACCTATCACGA

CGGCCGACTGATACCCCAGGTAAGGGTCATCGACCCTGGTCTGGGAGACAGGAAAGATGAGTGCTTCATGTACCTGTTTC

TCCTAGGAGTGATTGAGGATAACGACCCCCTGTCTCCCCCAGTCGGAAGAACCTTTGGTTCTTTACCTCTAGGGGTAGGT

AGGTCAACTGCTAAGCCAGAAGAACTACTAAGGGAGGCCACAGAACTAGATATAGTGGTGAGGCGCACTGCAGGATTAAA

TGAGAAACTGGTATTTTACAACAACACTCCGCTGTCCTTATTAACGCCCTGGAAGAAAGTTCTGACAACTGGAAGTGTGT

TTAGCGCTAACCAGGTTTGCAATGCAGTCAACCTAGTCCCGCTTGATACTCCCCAGAGATTCAGGGTTGTGTACATGAGC

ATAACTAGATTATCAGACAATGGTTACTATAGTGTGCCCAGAAGAATGCTGGAGTTCCGCTCCGCCAATGCAGTAGCCTT

CAACATCTTGGTTACACTAAGAATTGAAAATGGCACAAACCCTAGACGATACATAGTCGGCTCATGGGAGAATCCAGAGG

TCACATTTATGGTACACGTGGGCAACTTTAGAAGAAAGAAGAACGAAGTATACTCTGCTGATTACTGCAAAATGAAGATT

GAAAAGATGGGTTTAGTTTTTGCCCTGGGAGGAATAGGTGGAACAAGTCTCCATATTAGAAGCACCGGGAAAATGAGCAA

AACCCTCCATGCACAGCTGGGGTTCAAGAAAATCTTATGTTACCCCCTAATGGATATTAATGAGGATCTTAACCGATATC

TTTGGCGGGCAGAGTGCCGAATAGTCAAAATCCAAGCCGTCTTACAGCCATCAGTACCCCAAGAATTCCGTGTCTACGAT

GATGTCATCATCAACGATGATCAAGGCCTGTTCAAGATCTTGTAGTTCATTTGCAAAGGAATGAACCCACCCCCAGGAAG

GGGCCCTACCCTCCTCCG-CCAAAAAACACACTCCAAGGCTCCCCGACATAGCGCCAACACAGCCACTCTCCCGATGGGA

CGAGGCTGG-G-CCCCAGCAAACCCGGTCCAGGCGGGGGGCCCCCCCCCCCGACCCGCTGGGCGCCGACCGAGCAGGTCG

GCGAGAAGCCCACAACAGGCCCCGACGAAGACCGAGCACCCCATCCCCCC-AACACCCCTCCTC-AACACGCCCGAGGCC

AGGCGCCGGGCACCCCCAGACCCGCCACCGGCGAGACAGAACCCCATGGGGGGACCTCCCAAACCAAAGGGGCCACCGGG

GAAGGCTCCGCCGAACCAGCACGGACCCGACCCAAGCAAGTCGGACCTCCAACACGAAACCGCCCGGCGCACCACACTCA

TCATCAAACAAAACTTAGGGGCCAAGTCCACACGACAGGCCCCAGCCCGCCAGGCGGAGGGCCGCAGAAAGGAAGGAGAC

ACGACCGTCCCAGAAGGAAGACACCCAACCACCGAAACACCCAGCCCCAGGGAAGGAG--CCCCCCCCCACACGGCGAAG

GAACGGGTGAGAGACCCCGAGGAGAACCCGACCGGAAGCCAGGAGCCGGCGACGCCGGTGGCAAACAGGCAGGGCATACA

CCCGACTCCACCAGAGGAGCCGAGGAGAGCCTGACCCCGCGTGGACCAAACGGCAGGGAATCCAACCCCGCCACAAAAAG

AAGTGAGGCTCT-CCT-----CCCCCCCGCACACAACCCGCAGACAGACCAAGGGAGGACCACCTTCCAACAAGACAGGC

GGCTCCCTCGCAGCCGAGCCGCAGCTCCCCAGAAGAGGACACTCCCGCACACTCCGAACACCCCCCCGCCCCCCGCCACC

GGGACAGGGGCTCCGCCACCACAGACCAGAGCCACCGAGAGCCAGCCAGGGGTCCCCTCCAGTCCCGCTCAGCACCCCCC

CGAGGAGGCCCCGGCGAGGCCCCCAAGCCCGACGACCCCTGCACGCACCGAAAGAGGGACCCGCCCGAGAACGTCCTGCA

CATCCATGCGTAAACATCATGACGCGGGTCGCAATCTTGACATTTCTGTTTCTCTCCCTAAATGCTATTGCGTGCCAGAT

TCACTGGGGCAATCTATCCAAGATCGGGATTGTAGGAACAGGGAGTGCCAGCTACAAGGTGATGACTAGGCCAAGCCACC

AAACTTTGGTTATAAAGTTAATGCCAAATATAACAGCCATCGACAATTGTACAAAGTCAGAGATTGCAGAGTACAAGAGA

TTGCTGATCACAGTGTTAAAGCCTGTAGAGGATGCTCTGTCAGTGATAACCAAGAATGTAAGACCAATTCAAACTCTGAC

ACCTGGGCGTAGAACTCGCCGTTTTGCTGGAGCTGTTCTGGCCGGAGTAGCACTTGGAGTTGCGACAGCCGCTCAGATAA

CTGCAGGAGTCGCCCTTCATCAATCATTGATGAACTCCCAAGCAATTGAGAGTTTAAAAACCAGTCTTGAGAAGTCGAAT

CAGGCAATAGAAGAAATCAGACTTGCAAATAAGGAGACCATACTGGCAGTACAGGGCATCCAGGATTATATCAACAATGA

GCTTGTCCCTTCTGTTCATAGAATGTCATGCGAGCTAGTAGGTCACAAGCTCGGCCTCAAGCTCCTTAGGTACTACACCG

AGATCCTGTCCATATTCGGGCCTAGCCTTCGAGACCCGATAGCTGCCGAAATATCAATCCAGGCACTCAGTTATGCATTA

GGCGGAGACATTAATAAAATCCTGGACAAGCTTGGGTATAGCGGTGGGGATTTCCTTGCCATCCTAGAAAGTAAGGGAAT

AAAGGCCCGGGTCACATATGTGGACACAAGAGATTACTTTATAATTCTTAGCATCGCCTACCCAACCTTATCTGAGATCA

AGGGAGTGATAGTTCACAAGATAGAAGCTATAACATACAACATTGGGGCACAGGAGTGGTACACTACTATCCCTAAATAT

GTAGCCACTCAGGGGTATCTGATATCGAACTTTGATGAAACGTCATGCGTATTCACTCCAGAGGGGACAGTTTGCAGCCA

GAATGCGTTGTACCCAATGAGCCCGTTGCTTCAGGAATGTTTCAGGGGGTCAACAAAATCGTGCGCCAGAACCCTAGTTT

CAGGGACCATAAGTAATAGATTTATCCTATCAAAAGGGAACCTGATTGCAAATTGCGCGTCAGTTTTGTGCAAATGTTAC

ACAACGGAGACAGTTATCAGCCAGGATCCTGACAAACTACTAACTGTTGTAGCATCCGACAAGTGTCCTGTAGTTGAGGT

GGACGGAGTGACAATACAGGTCGGCAGTCGAGAGTATCCGGATTCTGTATACTTACACAAAATAGATTTAGGTCCAGCCA

TCTCCCTAGAAAAACTGGATGTAGGCACCAATTTAGGCAATGCAGTCACAAGACTGGAGAATGCAAAGGAGCTCCTAGAT

GCATCAGACCAAATACTGAAGACTGTTAACGGGGTACCTTTCGGCGGGAATATGTACATAGCACTGGCAGCTTGCATTGG

GGTATCCCTAGGGCTTGTCACATTAATATGCTGCTGTAAGGGGAGGTGTAAGAACAAGGAAATCCCTATCTCCAAAATCA

ACCCAGGGCTCAAACCCGACCTGACCGGGACCTCAAAGTCGTACGTGAGATCACTGTAGTCAGAATCACCCGAATCATCC

GGCATCACACACACACATGCACGACACAAGCAGTCAGAGGAGGCAGGAGACCCAGCCTCCGATCACCAACCTGACTCCAC

TCCACGCTCCACCAGACATCAGTCATCAAACAAAACTTAGGACGAAAGGTCAATCACCATGTCCGCACAAAGGGAAAGAA

TCAATGCCTTCTACAAAGACAATCCTCACAATAAGAACCATAGGGTGATTCTCGATAGAGAACGCTTGGTCATTGAAAGA

CCCTACATCTTGCTTGGAGTCCTGCTGGTAATGTTCCTGAGTCTAATCGGACTGCTGGCCATTGCAGGGATCAGGCTTCA

CCGGGCCACCGTTGGAACTTCAGAGATCCAGAGTCGGCTGAATACCAATATTGAGTTGACCGAATCTATTGATCACCAAA

CTAAGGATGTCTTAACTCCCCTTTTTAAAATCATTGGCGATGAAGTCGGCATCAGAATTCCACAGAAATTCAGTGATCTT

GTCAAGTTCATCTCCGATAAGATTAAATTCCTCAACCCTGATAGAGAGTATGATTTCAGGGATCTCCGGTGGTGTATGAA

TCCCCCTGAGAGAGTCAAAATTGATTTTGATCAGTTTTGTGGGTACAAAGCTGCGGTTAAGTCAATTGAACATATATTTG

AGTCACCACTCAACAAGTCAAAAAAGCTGCAATCATTGACTCTTGGGCCCGGAACAGGCTGTCTAGGCAGGACAGTAACA

AGAGCTCGTTCCTCAGAACTTACACTGACCTTAATGGACCTGGATATAGAGATGAAGCACAACGTGTCCTCAGTGTTTAC

CGTAGTTGAAGAGGGATTATTCGGAAGAACATATACAGTCTGGAGATTCGATGCCAGGGATCCTAGCACCGATCTAGGTA

TTGGCCATTTTTTAAGAGTCTTCGAGATTGGACTGGTAAGAGATCTCGGGCTGGGTCCCCCTGTTTTTCATATGACCAAC

TATCTCACAGTGAACATGAGTGATGACTATCGGAGATGTCTTTTAGCGGTAGGGGAGTTGAAGTTGACAGCCTTATGCAC

CTCATCTGAGACTGTGACACTGAGTGAGAGAGGAGTTCCAAAGAGGGAGCCTCTTGTGGTTGTGATACTTAACCTAGCTG

GGCCCACTCTAGGGGGAGAACTATACAGTGTCTTGCCTACCTCTGATCTCATGGTGGAGAAACTCTATTTATCTTCTCAT

AGAGGGGTTATCAAAGATGACGAGGCCAATTGGGTAGTGCCGTCAACCGATGTTCGTGATCTTCAGAACAAAGGAGAATG

TCTAGTGGAAGCATGTAAGACCCGACCTCCCTCATTTTGCAATGGCACAGGATCAGGCCCGTGGTCAGAGGGGAGAATCC

CTGCTTACGGGGTGATCAGGGTCAGTCTTGACTTAGCTAGTGACCCGGGTGTAGTTATCACCTCAGTGTTTGGCCCACTG

ATACCTCACCTATCCGGCATGGATCTTTACAATAACCCGTTTTCAAGAGATGTATGGTTGGCTGTACCACCTTATGAGCA

GTCATTTCTAGGAATGATAAATACAGTTGGATTCCCCAACAGGGCAGAGGTTATGCCGCACATTTTGACCACAGAGATCA

GGGGCCCTCGGGGTCGTTGCCATGTTCCCATAGAATTGTCCCGCAGGGTTGATGACGATATCAAGATCGGGTCCAACATG

GTCATATTGCCGACGATGGACCTGAGGTATATTACAGCCACTTATGATGTTTCCAGGAGCGAGCATGCAATCGTGTATTA

TATATATGACACAGGTCGCTCATCATCTTACTTCTACCCAGTTCGACTGAATTTCAAAGGCAATCCTCTCTCTCTGAGGA

TAGAGTGTTTCCCTTGGCGTCATAAGGTGTGGTGCTACCATGATTGTCTCATATACAACACCATAACAGATGAAGAAGTC

CACACGAGAGGGCTGACCGGTATAGAGGTAACATGTAATCCAGTCTGAGCAGAGCTACGACCGTCACTCAGGCAGTCCCT

TGAGTCGTCACTGAGTCCAAGCAGCACAGCCTGGGATACTCAACAGCACAGCCCAGCCAATAATGTTATAAAAAACTTAG

GAGCCAGGGTTGTAGGGGCCATGGATTCCCTATCAGTCAATCAGGTCCTGTACCCTGAGGTTCATCTAGATAGCCCTATT

GTCACAAACAAACTAGTTGCCATCCTTGAGTACTCGGGTATCGACCATAACTATGTTCTTGAAGACCAGACCCTTGTCAA

GAATATTAGGTATAGACTGGGGTGCGGTTTTTCAAATCAAATGATCATCAATAACAGGGGGGTGGGTGAAACAGTCAATT

CTAAACTTAAAAGTTACCCCCATAATCGTCATATCATATACCCGGATTGCAATAAGGAGTTGTTTTGTATCAAAGATAGC

TGCATATCTAAGAAGCTCTCAGAGCTATTCAAGAAGGGTAATTCCTTGTACTCTAAGATAAGTCACCAGGTACTGGACTG

TCTCAAGAGAGTCAATGGGAAATTAGGCCTGGGCACAGATCTTACCCATGGTCTGAAGGAGGGTATCCTTGACCTAGGGT

TGCACATGCATAGCTCTCAATGGTTCGAGACTTTCCTGTTCTGGTTCACTATCAAGACAGAGATGAGATCAATGATCAAA

GAACAGTCCCATATATGCCACAAGAAGAGGTATAACCCGATTTTTGTGTCGGGGGATGCATTCGAGGTGCTCGTATCACG

GGACCTCGTTGTGATAATTGATAAGAATACCCAGTATGTCTTCTACTTAACATTTGAGTTGGTCCTCATGTATTGTGATG

TCATAGAGGGCAGACTTATGACGGAGACAGCCATGGCCATAGACCAGAGATATTCAGAACTTCTAAGCCGGGTCAGATAC

TTGTGGGATCTTATTGATGGGTTCTTCCCAACACTAGGCAACACCACATACCAAATTGTTGCTCTGCTTGAACCGTTATC

GTTGGCTTATCTTCAACTTCAGGATGTCACTCTGGAGTTAAGAGGTGCTTTTTTAGACCATTGTTTCAAAGAACTCTATG

AGATACTGGAGCATTGCGGCATTGACACAGAAGGCACCTACAATTCCATCACTGAAGGATTAGATTACGTATTCATCACC

CACGATATACATCTAACTGGGGAGATTTTTTCATTTTTTCGGAGTTTCGGACACCCCCGTCTAGAAGCAGTCACCGCTGC

AGAAAATGTCAGGAAACATATGAACCAACCGAAGGTAATCAGTTATGAGACTATGATGAAAGGGCATGCAGTATTTTGCG

GGATAATCATAAATGGTTTTAGGGACCGACATGGCGGCAGCTGGCCCCCTGTTGCATTGCCAGAGCATGCTTCTGCTGCG

ATTCGGAATGCGCAGGCATCAGGTGAGGGGCTGACCCATGACCTGTGTATAGACAATTGGAAATCCTTTGTTGGATTCAA

ATTTGGCTGCTTTATGCCACTCAGCCTAGATAGTGATTTGACCATGTACCTCAAAGACAAGGCGTTGGCTGCACTGAAGA

ATGAGTGGGATTCAGTTTACCCGAAAGAATACCTCCGTTATAACCCACCTAGAGGGACAGAGTCAAGGCGACTGGTGGAG

GTGTTCCTGAATGACTCCAGCTTTGATCCTTATAACATGATAATGTACGTGGTGAATGGCTCCTACCTTAAAGACCCTGA

GTTTAATCTTTCATACAGCCTGAAAGAGAAGGAGATAAAAGAGACAGGGCGGTTGTTTGCTAAAATGACCTATAAGATGC

GGGCCTGTCAGGTAATCGCTGAAAATCTGATATCGAATGGTGTTGGGAAGTATTTCCGAGACAATGGGATGGCAAAGGAC

GAGCACGACCTAACAAAAGCCCTTCACACCCTGGCAGTCTCAGGTGTTCCCAAGAACAACAAAGATAACCACCGAGGTGG

GCCTCCCAGAAGGAAAACAAGCCGAGATGTGAGACCAAACCAAGACACCAAAACACAAGATAGAGACAAGGTCCAAGGGG

GGCCTATGTACAACTATTTGCGATGCCGACCAATCAGCCCTGATCAGGGTGAGTCATACGAGACTGTTAGTGCATTCATC

ACCGCTGACCTTAAGAAGTATTGCCTGAATTGGAGATACGAGACAATCAGCATATTTGCACAGAGGCTGAATGAAATATA

TGGACTACCATCCTTCTTTCAATGGTTACACAGGATATTGGAAAAATCCGTACTCTACGTCAGTGACCCACATTGCCCTC

CCGACCTAGATAATCATATCCCTCTGGACAGTGTCCCTAATGCCCAAATATTCATTAAGTACCCAATGGGCGGAATAGAA

GGTTATTGCCAAAAACTATGGACAATCAGTACTATACCATATTTGTATCTGGCAGCCTATGAGAGCGGAGTAAGAATCGC

CTCACTAGTGCAGGGTGACAATCAGACAATCGCAGTGACAAAAAGAGTTCCAAGTTCTTGGCCTTATTCACTAAAAAAGA

GGGAGGCATCCAAAGCCGCTCGAAATTACTTCGTAGTCTTAAGGCAAAGATTGCACGATGTAGGTCATCACTTAAAGGCT

AATGAAACCATAGTATCTTCTCACTTTTTTGTATATTCCAAGGGGATTTATTATGACGGCCTGCTAGTCTCACAATCACT

AAAGAGCATCGCCAGATGTGTCTTCTGGTCCGAGACTATAGTGGATGAAACCAGAGCGGCTTGCAGCAATATTGCAACAA

CTGTCGCTAAGAGTATAGAGAGGGGTTATGATAGGTACCTTGCATATTCTTTGAATATCCTCAAGATTTTCCAACAGATC

CTTATATCCCTTAACTTCACTATTAACACAACAATGACTCAGGATGTCGTGGCACCGATCATCGAGAACGGTGATTTGCT

GATAAGGATGGCACTCTTGCCGGCACCCATCGGGGGTCTCAATTATCTTAACATGAGCAGGTTATTTGTGAGAAATATCG

GTGACCCGGTCACCTCCTCTATAGCTGACCTGAAGAGGATGATAGATGCTGGGCTAATGCCAGAAGAAACACTGCATCAA

GTGATGACCCAGACCCCGGGAGAATCATCCTACCTCGATTGGGCAAGTGACCCTTATTCTGCCAACTTACCCTGCGTACA

GAGTATAACTCGCCTTCTCAAAAACATCACTGCACGATATATTTTAATCAGCAGCCCAAACCCGATGCTGAAAGGATTGT

TTCATGAGGGGAGTAGGGATGAGGACGAGGAGCTTGCAAGTTTCTTAATGGATCGGCATATAATTGTTCCAAGAGCTGCA

CATGAAATCTTAGACCATAGCATAACCGGGGCAAGAGAAGCTATAGCCGGGATGTTGGATACTACCAAGGGTCTGATTAG

AACAAGTATGAGACGGGGTGGCCTCACCCCTCGAGTCTTAGCCCGCCTTTCCAATTATGATTATGAACAATTCCGATCCG

GGATAGCACTATTGACAAAGAAAGGGCAGTGTTATCTCATTGACAAGGACTCGTGCTCGGTGCAGCTCGCTATAGCTCTG

AGGAGCCACATGTGGGCTAGGTTGGCTCGCGGGAGGCCAATCTATGGGTTGGAGGTGCCCGATATATTAGAATCGATGAA

CGGCTACCTTATCAAACGCCACGAGTCTTGTGCTATCTGTGAAACGGGCTCAAGTCACTACGGATGGTTTTTCGTCCCTG

CAGGGTGCCAGCTTGACGATGTATCTAGAGAGACTTCGGCTCTTCGTGTACCTTATGTCGGATCAACAACTGAGGAAAGG

ACAGACATGAAACTTGCTTTTGTTAGGTCTCCAAGTCGATCCCTCAAATCAGCAGTCAGAATTGCAACAGTTTACTCATG

GGCCTACGGGGATGATGAGAAATCATGGAGTGAAGCCTGGATGCTAGCTAGGCAGAGAGCCAATATCACCTTAGATGAAT

TAAGAATGATCACTCCAGTCTCTACATCCACCAACCTAGCCCACCGGTTGAGGGATCGGAGCACCCAGGTGAAATACTCG

GGGACATCCCTTGTGAGGGTTGCAAGATACACAACCATCTCCAATGATAACCTGTCATTTGTGATATCTGAGAAAAAAGT

GGATACCAACTTCATTTATCAGCAAGGGATGCTGCTCGGTCTTGGGATCCTTGAGAACCTCTTTAGGCTAGAGGCCACCA

CCGGAGTATCCAACACAGTGCTACACCTGCACGTGGAAACAGAGTGTTGTGTTGTACCTATGGTGGATCACCCAAGGATA

CCGAGTCTCCGTAATATAAAAGTTACGGATGAGCTATGCACAAACCCTTTGATCTACGACCGGTCTCCCATCATAGAGCA

CGATGCAACCCGACTATACTCACAGAGCCACAGGAGACATTTGGTGGAGTTTGTTACCTGGTCAACAAGCCAGCTTTACC

ATATACTGGCTAAGTCTACAGCAATGTCCATGATCGAGTTGATCACGAGATTCGAGAAAGATCACATGAATGAAATAGCC

GCCCTGATAGGCGATGACGACATCAACAGCTTCATCACAGAATTTTTGCTAGTAGAGCCTAGATTATTTATAGTTTACCT

TGGTCAGTGTGCTGCCATCAATTGGGCCTTTGATATACATTATCACCGGCCCTCGGGCAAGTATCAGATGGGGGAACTCC

TCTACTCTCTGCTCTCTCGGATGAGCAAAGGAGTATATAAGATCTTCACCAATGCTCTGAGCCACCCCAAAGTTTACAAG

AAATTTTGGCGAAGTGGTGTCATTGAGCCGATTCATGGTCCATCCCTGGATACACAGAATTTGCATGTCACTGTCTGTGA

CATGATATATGGGTCCTATGTCACCTATTTAGATCTTTTGCTGAATGATGAGCTAGATGATTATCCGTATTTGCTCTGCG

AGAGTGATGAGGACGTGGTCACAGACAGGTTCGACAACATTCAGGCCAAACATCTCTGTGTATTGGCCGATGTATATTGC

AGCTCCAAGAGGTGTCCTTCGATAATCGGAATGTCTCCTATAGAAAAATGTACCATTCTCACACATTACATCAAGGGAGA

ATCAGTACAATCCCCGTCTGGGACCTCATGGAACACGGATCCTCTTGTAGTAGATCATTACTCATGCTCTCTGACCTACC

TTCGCCGCGGTTCCATCAAACAAATCAGGTTGAGAGTGGATCCGGGGTTTATATTCGAGGCGTTGACAGACGTCGACTTT

AAACAGCCTCGCAAAGCTAAGTTAGATATATCGGTCGTTGGGTTGACTGATTTCTCTCCCCCTTGCGACAACGTCGGTGA

TTTTCTAGGGACTATCAACACATTGAGGCACAATCTGCCTGTCACCGGAACCGGGGTCTCGAACTATGAAGTCCACGCTT

ATCGTAGAATTGGCCTGAATTCATCAGCATGTTATAAAGCCGTAGAGATCTCCACGTTAATCAAGCCATACTTAGAAGTC

GGAGAGCATGGATTGTTCTTAGGAGAAGGTTCCGGTTCAATGCTGGCTGCGTACAAGGAAGTTCTTAAATTAGCAAATTG

TTATTACAACAGCGGGGTAACAGCGGAGGGCAGAGCCGGACAGAGGGAAATATCTCCCTATCCTTCAGAGATGAGCCTAG

TAGAGAATCAAATGGGGATAGAGCGGAGTGTTAAAGTGCTGTTCAACGGTAAACCTGAAGTAACTTGGGTGGGGACCACC

GATTGCTACAAGTACATAATCAGTAACATCCAAACCTCTAGTCTGGGTTTCATACATTCAGATATTGAGACACTCCCAAC

CAAAGATGCTGTTGAGAAATTAGAAGAATTTGCCTCTATCCTATCCTTATCCTTGATTTTGGGAAAAATCGGCTCTATTA

CAGTTGTCAAAATTATGCCCATTAGCGGGGATTTTACCCAAGGCTTCATAGCCTATGCCATCCAATATTTTAGGGAGAGT

CTGCTTGCCTATCCGAGATATAGTAACTTCATCTCGACTGAGTGTTACCTTATTATGATAGGATTAAAGGCCAATCGATT

GATGAACCCAGAAGCCATTAAGCAAGGCATAATCAGAGCGGGGGTCAGGACTGCACCAGGACTTGTGAGCCACATATTAT

CAGAGAAACAGAAAGGTTGTATTCAATCTTTCCTGGGTGATCCTTATATCCAAGGAGACTTCAATAAGCACCTTAAATCT

CTTACCCCTATTGAGAAAATCCTGGTAAATTGTGGTCTCTCGATCAATGGCACAAAAATCTGTAGAGATCTAATCCACCA

TGATATCGCCTCCGGTCCAGACGGTCTTATGAGCTCTACAATCATTTTATATAGGGAACTGGCTCATTTCAAAGATAATA

TAAGGAGTCAGCACGGAATGTTCCACCCCTACCCAGTACTGGCCAACAGCAGGCAACGTGAATTAATCCTTCGAATAGCC

AAGAAATTCTGGGGGTATGTCTTGCTGTATTCTGATGACCCGGCACTAATCAGACAAACAATCAAGAACTTGAAACGGAA

TCATCTAACCTTTGACTTACATAATAACCCGTTTATTAAAGGGCTATCCAAAGCTGAGAAACTGCTGGTGCGGACGAGTT

CACTAAGAAGGGAATGGTTGTTCACTCTCGATACGAAAGAAGTGAAAGAGTGGTTCAAATTGGTGGGTTACAGTGCACTC

ATCAGAGGCTAGTTAGCTATACATCTGCCCCCTTCTCCTCCGCCATGAGACCCCACTGACGATCCAAAAGATTAAAGAAA

ACTACATATTGGATAAGTATCTATTCCCAGCTTTGTCTGGT

>MF678816.1/NubianIbex/Israel/2017-01

NNNNNNNNNNNNNGGGTAAGGATAGATCTTACAATAACTATGGACTGGCAAACTTAGGAGTAAAGATCCTACTGTCGGGG

GGAGGAGGAGGAACAAGATCTTTGATTATGGCGACTCTTCTTAAAAGCTTAGCATTGTTCAAAAGGAACAAAGACAAAGC

GCCGACGGCATCAGGTTCAGGAGGGGCCATCCGGGGGATTAAGAATGTTATCATAGTCCCGATTCCCGGAGACTCGTCCA

TCATTACCCGTTCAAGACTGCTCGACAGGCTTGTCAGATTGGCCGGAGATCCTGACATCAACGGGTCAAAGTTGACCGGC

GTGATGATCAGCATGTTATCTTTGTTCGTAGAGTCACCCGGGCAATTGATACAGCGAATCACAGATGATCCAGATGTCAG

TATCCGCCTTGTTGAGGTAGTTCAAAGTACTAGGTCTCAGTCCGGGTTGACCTTTGCATCACGCGGTGCTGATTTGGACA

ACGAGGCAGACATGTATTTTTCAACTGAGGGGCCCTCGAATGGAGGCAAGAAAAGGATCAACTGGTTTGAGAACAGAGAG

ATAATAGACATAGAGGTGCAGGATCCAGAAGAGTTCAATATGTTGTTAGCCTCCATACTAGCACAAGTCTGGATCCTTCT

GGCCAAGGCTGTTACGGCACCGGATACGGCAGCTGACTCAGAATTGAGAAGGTGGGTTAAATATACACAACAAAGGAGAG

TGATTGGGGAGTTTCGCCTTGACAAAGGGTGGCTGGACGCAGTCCGCAACAGAATTGCAGAAGATCTATCACTTCGGCGG

TTCATGGTGTCTCTCATACTTGACATCAAGAGGACCCCCGGCAACAAGCCAAGGATTGCAGAAATGATATGCGACATTGA

CAACTATATTGTCGAAGCCGGACTCGCCAGTTTCATCCTTACTATCAAGTTTGGTATCGAAACCATGTATCCTGCATTAG

GGCTCCACGAGTTTGCCGGGGAATTGTCCACTATTGAATCCTTGATGAACTTGTATCAACAGCTAGGAGAGGTTGCACCC

TACATGGTAATTCTCGAGAACTCAATTCAGAACAAGTTTAGTGCAGGAGCTTATCCCCTCCTCTGGAGCTATGCGATGGG

TGTCGGAGTTGAGCTGGAGAACTCAATGGGGGGCTTGAACTTTGGCAGGTCATATTTTGACCCTGCCTATTTTCGTCTCG

GACAGGAGATGGTCAGAAGATCTGCAGGAAAGGTCAGCTCTGTGATTGCGGCCGAGCTCGGCATCACAGCAGAGGAAGCT

AAACTAGTCTCGGAAATCGCCTCGCAGGCTGGGGACGAACGAACCGCCAGAGGGACTGGGCCTCGACAGGCGCAGGTCTC

CTTCCTCCAGCATAAAACAGGAGAGGGAGAGTCGTCCGCACCAGCAGCCAGAGAAGGGGTCAAAGCTTCGATCCCAAACA

GATCCGAAGAAAGGGACAGAAAGCAAACACGCCCAGGAAGGCCCAGAGGAGAGACCCCCAGCCAACTGCTCCTGGAAATC

ATGCCAGAGGATGAGGTCTCGCGAGAGTCTGGTCAAAACCCTCGCGAGGCTCAAAGATCGGCCGAGGCACTCTTCAGGCT

GCAGGCCATGGCCAAGATTCTGGAGGACCAGGAGGAGGGAGAAGACAATAGTCAGGTCTACAACGACAAGGATCTCCTCG

GCTAAGCAAACGCACCCTCTGTCGAAGTCAGTGATAAGACATCTCCTACCAGTATTATAAAAAACTTAGGACCCAGGTCC

AAGCAAGCACACATCGACACTCCAATTAGTCGAGCGGAGACCACCGATGGCAGAAGAACAAGCATACCATGTCAACAAGG

GGCTGGAATGTATCAAGTCCCTCAAAGCCTCTCCCCCGGATCTATCCACCATCAGAGATACCATTGAGAGCTGGAGAGAG

GGGCTTAGCCCCCCGGGCCGTGCAACACCGAACCCTGATACGTCCGAGGGAGATCATCAGAATATCAACCAATCATGCTC

ACCAGCAATCGGACCAGACAAAGTCTACTTGTCTCCTGAAGATAATCTCGGATTTAGAGAGATCACTGGCAACGACTGTG

AGCCTGGGCTCGGAGGAGTCCAGGGAAAAGGATCCAACTCTCAAGTACAGCGTTACTATGTTTATAGCCACGGGGGTGAA

GAGATTGAAGGACTCGAGGATGCTGACTCTCTCGTGGTTCAAGCAGATCCTCCAGTTGCTAACATCTTCAATGGAGGAGA

GGATGGATCTGACGACAGCGATGTGGAATCTGGCTCAGATGATCCCAGCAGAGATACTCTATATGACCGGGGATCTGTTG

CCGGCAATGATGTCGCTAGGTCCACAGATGTTGAGAAACTAGAAGGTGCTGACATTCAAGAGGTTCTTAACTCCCAGAAA

GGCAGAGGAGGAAGATTCCAAGGCGGGAAGACCTTACGAGTCCCGGATATACCCGATGTCAAGCACCCCAGACCATCAGC

CCAATCAATTAAAAAGGGCACAGACGGGAACTCAGTCTCATCTGGAACGGTGACAGAGTGTTTATCGATAAGTGGTGCAA

CCCGAGCTGTGCCAGAGTCAAGATGGGAGTCATCAGAGCAAAATGCGTCTGTGGAGAGTGTCCTCAAGTCTGCGAGGAGT

GCAGAGACGATCCAGGGGTTGACACAAGAATCTGGTACCATAGCATCACTGACTCAGCCTAAGGAGAATGACTCCGAGTA

TGAGTATGAGGACGACTTATTCAAAGAGATCCAGGACATCCGTGCAAGCATTGCCAAGATCCATGATGACAACAAAACAA

TCCTCTCAAAGCTTGATTCTATACTGTTATTGAAAGGAGAAGTCGACACTATCAAGAAACAAATCAGCAAGCAAAATATA

AGTATATCCACCATTGAGGGCCATCTCTCCAGTATAATGATAGCTATCCCGGGCTTTGGGAAAGATACCAAGGACCCAAC

ATCCGAGGTTGAGTTGAACCCAGATTTAAGACCTATAATAAGCCGTGACTCTGGCAGAGCTCTCGCGGAGGTCCTCAAGA

AACCTGCTGTGGATAGGTCTCAGAAAATTGGAACCAAAGCCAACTCCAGCTCAAAGGGTCAGCTTCTTAAGGATCTCCAG

CTAAAACCTGTCGACAAGCAGGCAAGCTCTGCAATCGGGTTTGTCCCATCCGACCATGAATCATCCAGAAATGTCATCCG

CTCCATAATCAAGTCGAGCAAGCTAAACATTGATCACAAGGACTATCTCCTAGATTTACTAAATGATGTGAAAGGCTCCA

AGGATCTTAAGGAATTCCACAAGATGCTAACAGCAATTCTCGCCAAGCACCCGTAACACACCCTCCAGTCACCATCCTAC

ACTCGGCTGAAAACATCCTCTCAATCAGGCTATTACAAAAAACTTAGGAGCAAGGGTAACTGAGCTTCGCAGACAAGATG

ACCGAGATCTACGACTTCGATAAATCAGCATGGGATGTCAAAGGGTCAATTGCTCGCATAGAACCCACCACCTATCACGA

CGGTCGACTGATACCCCAGGTAAGGGTCATCGATCCTGGTCTGGGGGACAGAAAAGATGAGTGCTTCATGTACCTGTTTC

TCCTGGGAGTGATTGAGGATAATGACCCCCTGTCTCCCCCAGTTGGAAGAACTTTTGGTTCTTTACCTCTAGGGGTCGGT

AGGTCAACTGCTAAGCCAGAAGAACTACTAAGGGAGGCCACAGAACTAGATATAGTGGTGAGGCGCACTGCAGGATTAAG

TGAGAAACTGGTATTTTACAACAACACTCCGCTGTCCTTGTTAACGCCCTGGAAGAAGGTTCTGACAACCGGAAGTGTGT

TTAGTGCTAACCAGGTTTGCAATGCAGTCAACCTAGTCCCGCTTGATACTCCCCAGAGATTCAGGGTTGTGTACATGAGC

ATAACTAGATTATCAGACAATGGTTACTATAGTGTGCCCAGGAGAATGCTGGAGTTCCGCTCAGCCAATGCAGTAGCCTT

CAACATCTTGGTTACACTAAGAATTGAAAACGGCACAAGCCCTAGAAGATACATAGTCGGCTCATGGGAGAATCCAGAGG

TCACATTTATGGTACACGTGGGCAACTTTAGAAGAAAGAAGAACGAAGTCTACTCAGCTGATTACTGCAAAATGAAGATT

GAAAAGATGGGTTTAGTTTTTGCCCTGGGAGGAATAGGTGGAACAAGTCTCCACATTAGAAGCACCGGGAAAATGAGCAA

AACCCTCCATGCACAGCTGGGGTTCAAGAAAATCTTATGTTACCCCCTAATGGATGTTAATGAGGATCTTAACCGATATC

TCTGGCGGGCAGAGTGCCGAATAGTCAAAATCCAAGCCGTCTTACAGCCATCAGTACCCCAAGAATTTCGTGTCTACGAT

GATGTCATCATCAACGATGATCAAGGCCTATTCAAGATCTTGTAGGCCATTTGCAAAAGAATGAACCCACCCCCAGGAAG

GGGCCCAATCTTCCCCCG-CCAGAAAACACACTCCAAGGCACCCCAACACAGCACCAACACAGCCACTCTCCCGATGGGA

CGAGGTCGGCC-CCCCAACAAACCCGGTCCAGGCGGGGGG--CCCCCCCCCGACCCGCCGGGCGCCGACCGAGCAGGCCG

ACGAGAAGCCCCCAAGAGCCCCCGACGAAGACCGAGCCCCCCACCCCCCCAAGAGCGTCCCTCC-AGCACGCCCAGGGCC

AGGCGCCGGGCACCCCCAGATCCGCCATCGGCAAGACAGAACCCCAAGGGGGGACCCCCAAAAACAAAGGGACCACCGGG

GAGGGCCCCGCCGAACCAGCACAGACCCGACCCAAGCAAGGCGGACCCCCAACACGAGACTGCCCAGCCCACCCCACCCA

TCACCAAACAAAACTTAGGGGCCAAGTCCATACGACAGGCCCCAGCCCGCCAGGCGGAGGGCCGCAGAAAGGAAGGAGAC

ACAACCGCCCCAGAAGAAGGACACCCAACCCCCGAAACACCCAGCCCCAGGAAAGGAG--CCCCCCCCCACACGGCGAAG

GAACGGGTGAGAGACCCCGGAGAGAACCCGACCGGAAGCCAGGAGCCGGCGACGCCGGCGGCAAACAGGCCGGGCGCACA

CCCGACTCCACCAGAGGAGCCGGGGAGAGCCTAACCCCGCGTGGACCAAACGGCAGGGAATCCAACCCCGCCGAAAAAAG

AGGTGAGGCCCC-CTC-----CCCCCCCGCACACGACCTGTAGACAGACCGAGGGAGGACCACCTCCCAACAAGACAGGC

GGCTCCCTCGCAGCCGAGCCGCATCTCCCCAGAAGGGGACACCCCCGCATACCCCGAACACCCCCCCGCCCCCCGCCACC

GGGACAGGGGCTCCGCCACCACAGACCAGAGCCACCGAGAGCCAGCCAGAGGTCCCCTCCAGTCCCGTCCGGTGCCCCCC

CGAAGAGGCCCCGGCAAGGCCCCCCAGCCCGACGACCCCTGCACGCACCGGGAGAGGGACCCGCCCGAGAACGTCCGGCA

CATCCATGTGTAAACATCATGACACGGGTCGCAATCTTGACATTTCTGTTTCTTTTCCCAAATGCTGTTGCGTGCCAGAT

TCACTGGGGCAATCTATCCAAGATCGGGATTGTAGGAACAGGGAGTGCCAGTTACAAGGTGATGACTAGGCCAAGCCACC

AAACTCTGGTTATAAAGTTAATGCCAAATATAACAGCCATCGACAATTGTACAAAGTTAGAGATTGCAGAGTACAAGAGA

TTGCTGATCACAGTGTTAAAGCCTGTAGAGGATGCTCTATCAGTGATAACCAAGAATGTAAGACCAATTCAAACTCTGAC

AACTGGGCGTAGAACTCGCCGTTTTGCTGGAGCTGTTCTGGCCGGAGTAGCACTTGGAGTTGCGACAGCCGCTCAGATAA

CTGCAGGAGTCGCCCTTCATCAATCATTGATGAACTCCCAAGCAATTGAGAGTTTAAAGACCAGTCTTGAGAAGTCGAAT

CAGGCAATAGAAGAAATCAGACTTGCAAATAAGGAGACCATACTAGCAGTGCAGGGCGTCCAGGATTATATCAACAATGA

GCTTGTCCCTTCTGTTCATAGAATGTCATGCGAGCTGGTAGGTCACAAGCTCGGCCTCAAGCTCCTTAGATACTACACCG

AGATCCTGTCCATATTCGGGCCCAGCCTTCGGGACCCGATAGCTGCTGAAATATCAATCCAGGCACTCAGTTATGCATTA

GGCGGGGACATTAATAAAATCCTGGAAAAGCTTGGGTATAGCGGTGGGGATTTCCTTGCCATCCTAGAAAGCAAGGGAAT

AAAGGCCCGGGTCACATATGTGGATACAAGAGATTACTTTATAATTCTTAGCATCGCCTACCCAACCTTATCTGAGATCA

AGGGAGTGATAGTTCACAAGATAGAAGCTATAACATACAATATTGGGGCACAGGAGTGGTATACTACTATCCCCAAATAT

GTAGCCACTCAGGGGTATCTGATATCGAACTTTGATGAGACGTCATGCGTATTCACTCCGGAGGGGACAGTTTGCAGCCA

GAATGCGTTGTACCCAATGAGCCCATTGCTCCAGGAATGTTTCCGGGGGTCAACAAAATCGTGCGCCAGAACCCTAGTTT

CAGGGACCATAAGCAATAGATTTATCCTATCAAAAGGGAACCTGATTGCAAATTGCGCGTCAGTTTTGTGCAAATGTTAC

ACAACAGAGACAGTTATTAGCCAAGATCCTGATAAGCTACTAACTGTTGTAGCATCCGACAAGTGTCCTGTAGTTGAGGT

GGATGGAGTGACAATACAGGTCGGCAGTCGAGAGTATCCGGATTCTGTATACTTACACAAGATAGACTTAGGTCCAGCCA

TCTCCCTAGAAAAACTGGATGTAGGTACCAATTTAGGCAATGCAGTCACAAGACTGGAGAATGCAAAGGAGCTCCTAGAT

GCATCAGACCAAATACTGAAGACTGTTAAAGGGGTACCTTTCGGTGGGAATATGTACATAGCACTGGCAGCTTGCATTGG

GGTATTCCTAGGGCTTGTCACATTAATATGCTGCTGCAGGGGGAGGTGTAAGAAAAAGGAAATTCCCATCCCCAAAATCA

ACCCGGGGCTCAAACCCGACCTGACCGGGACCTCAAAGTCGTACGTGAGATCACTGTAGTCAGAATCACCTGAATCATCC

GGCATCACACACATACATGTACGACACAAGCAGTCAGAGGACGCAGGAAACCCAGCCTCCGATCACCCACCTGAACCCAC

CCCACGCTCAACCACACATTAGTCATCAAACAAAACTTAGGACGAAAGGTCAATCACCATGTCCGCACAAAGGGAAAGGA

TCAACGCCTTCTACAAAGACAATCCTCACAATAAGAACCATAGGGTGATCCTGGATAGAGAACGATTGGTCATTGAAAGA

CCCTACATCTTGCTTGGGGTCCTGCTGGTAATGTTCCTGAGTCTAATCGGACTGCTGGCCATTGCAGGGATCAGGCTTCA

CCGGGCCACCGTCGGAACTTCAGAAATCCAGAGTCGGCTGAATACCAATATTGAGTTGACCGAATCTATTGATCACCAAA

CTAAGGATGTCTTGACCCCCCTTTTCAAAATCATTGGCGATGAAGTCGGCATCAGAATTCCACAGAAATTCAGTGATCTT

GTCAAGTTCATCTCCGATAAGATTAAATTCCTCAACCCTGATAGAGAGTATGATTTCAGGGATCTCCGGTGGTGTATGAA

TCCCCCCGAGAGAGTCAAAATTAATTTTGATCAGTTTTGTGAATACAAGGCTGCGGTTAAGTCAATCGAACATATATTTG

AGTCACCACTCAACAAGTCAAAAAAGCTGCAATCATTGACTCTTGGGCCCGGAACAGGCTGTCTAGGCAGGACAGTAACA

AGAGCTCATTTCTCAGAACTTACACTGACCTTAATGGACCTGGATCTAGAGATGAAGCACAACGTATCCTCAGTGCTTAC

CGTAGTTGAAGAAGGATTATTCGGAAGAACATATACCGTCTGGAGATCCGATGCCAGGGATCCGAGCACCGATCTAGGTA

TTGGCCACTTTTTAAGAGTCTTCGAGATTGGACTGGTAAGAGATCTCGGGCTGGGTCCCCCTGTTTTTCATATGACCAAC

TATCTCACAGTGAACATGAGTGATGACTATCAGAGATGTCTTTTAGCGGTAGGGGAGTTAAAGTTGACAGCCTTATGCAC

CTCATCTGAGACTGTGACACTGAGTGAGAGAGGAGTTCCAAAGAGGGAGCCTCTTGTAGTTGTGATACTCAACCTAGCTG

GGCCCACTCTAGGGGGCGAACTATACAGTGTCTTGCCTACCTCTGATCTCATGGTGGAGAAACTCTATTTATCTTCGCAT

AGAGGGATTATCAAAGATGACGAGGCCAATTGGGTAGTGCCGTCTACCGATGTTCGTGATCTTCAAAACAAAGGAGAATG

TCTGGTGGAGGCATGCAAGACTCGACCCCCTTCATTTTGCAATGGCACAGGATCAGGCCCGTGGTCAGAGGGGAGAATCC

CTGCTTACGGGGTGATCAGGGTCAGTCTTGACTTGGCTAGTGACCCGGGTGTAGTTATCACTTCAGTGTTTGGTCCACTG

ATACCTCACCTATCCGGCATGGATCTTTACAACAATCTGTTTTCAAGAGCTGTATGGTTGGCTGTACCACCTTATGAGCA

GTCATTTCTAGGAATGATTAATACGATTGAGTTCCCTAACAGAGCAGAGGTTATGCCGCACATTTTGACCACAGAGATCA

GAGGCCCTCGGGGTCGTTGCCATGTTCCCATAGAATTGTCCCGCAGGGTTGATGACGATATCAAGATTGGGTCCAACATG

GTCATATTGCCGACGATGGACCTGAGATATATTACAGCCACTTATGATGTTTCCAGGAGCGAGCATGCAATCGTGTACTA

TATCTATGACACAGGTCGCTCATCATCTTACTTCTACCCAGTTCGGCTGAACTTCAAGGGCAATCCTCTCTCTCTGAGGA

TAGAGTGTTTCCCTTGGCGTCATAAGGTGTGGTGCTACCATGATTGTCTTATATACAACACCATAACAGATGAAGAAGTC

CATACGAGAGGGCTGACCGGTATAGAGGTAACATGTAATCCAGTCTGAGCAGAGCTACGACCGCCATTCAGGCAGTTCCT

TGAGTCGCCACCGAGTCCAAGCAGCACAGCCTGGGACACCCAACAGCACAGCCCAGCCAACAATGTTATAAAAAACTTAG

GAGCCAGGGCTGTAGGGGCCATGGACTCTCTATCAGTCAATCAGGTCTTGTACCCTGAGGTCCACCTAGATAGCCCTATT

GTCACAAACAAACTAGTTGCCATACTTGAGTACTCGGGCATCGACCATAACTATGTTCTTGAAGACCAGACCCTTGTCAA

GAATATTAGGTATAGACTGGGGTGCGGTTTTTCAAATCAAATGATCATCAATAACAGGGGGGTGGGTGAAACAGTCAATT

CTAAACTCAAAAGTTACCCCTATAATCGTCATATCATATACCCAGATTGCAATAAGGAGTTATTTTGTATCAAAGATAGC

TGCATATCAAAGAAGCTCTCGGAGCTATTCAAGAAGGGTAATTCCTTGTACTCTAAGATAAGTCACCAAGTACTGGATTG

CCTCAAGAGAGTCAATGGGAAATTAGGCCTGGGCACAGATCTTACCCATGGTATGAAGGAGGGTATCCTTGATCTAGGGT

TGCACATGCATAGCTCTCAATGGTTCGAGACTTTCCTGTTCTGGTTCACTATCAAGACAGAGATGAGATCAATGATCAAA

GAACAGTCCCATATATGCCACAAGAGGAGGTATAACCCGATTTTTGTGTCGGGGGATGCATTCGAGGTGCTCGTATCACG

AGACCTCGTTGTGATAATTGATAAGAATACCCAGTATGTCTTCTATTTAACATTTGAGCTAGTCCTTATGTATTGCGATG

TCATAGAGGGCAGACTTATGACGGAGACAGCCATGGCCATAGACCAGAGATATTCAGAACTTCTAAGCCGAGTCAGATAC

TTGTGGGATCTTATTGATGGGTTCTTCCCAACACTAGGCAACACCACATACCAAGTTGTTGCTCTGCTTGAACCATTATC

GTTGGCTTATCTTCAACTTCAGGATGTCACTCTGGAGTTAAGAGGTGCTTTTTTAGACCATTGCTTCAAAGAACTCTATG

AGATACTGGAGCATTGCGGCATTGACACAGAAGGCACCTACAATTCCATCACTGAAGGATTGGATTACGTATTCATCACC

CACGATATACATCTAACTGGGGAGATTTTTTCATTCTTTCGGAGTTTCGGACACCCCCGTCTAGAAGCAGTCACTGCCGC

AGAAAATGTCAGAAAACATATGAACCAACCGAAGGTAATCAGTTATGAGACTATGATGAAAGGGCATGCAGTATTTTGCG

GGATAATCATAAATGGTTTTAGGGACCGACATGGCGGCAGCTGGCCCCCTGTTACATTGCCAGAACATGCTTCTGCTGCG

ATCCGGAATGCGCAGGCATCAGGTGAGGGACTGACCCATGACCTGTGTATAGACAATTGGAAGTCTTTTGTTGGATTCAA

ATTTGGCTGCTTTATGCCACTCAGCCTAGATAGTGATTTGACCATGTACCTCAAAGACAAGGCATTGGCTGCACTGAAGA

ATGAGTGGGATTCAGTTTACCCGAAAGAATACCTCCGTTATAACCCACCTAGAGGGACAGAGTCAAGGCGATTGGTGGAG

GTGTTCCTGAATGACTCCAGCTTTGATCCTTATAACATGATAATGTACGTGGTGAATGGCTCCTACCTTAAAGACCCTGA

GTTTAATCTCTCATACAGCCTAAAAGAGAAGGAGATAAAGGAGACAGGGCGGTTATTTGCCAAAATGACCTATAAGATGC

GGGCCTGTCAGGTAATCGCTGAAAATCTGATATCGAATGGTGTTGGGAAGTATTTTCGAGACAATGGGATGGCAAAGGAC

GAGCATGACCTAACAAAAGCCCTTCACACGCTGGCAGTCTCAGGTGTTCCCAAGAACAACAAGGATAACTACCGAGGTGG

GCCTCCCAGGAGAACAACAAACCGAGAGGTGAGATCAAGCCAGGACATCAAAACACAAAATAGAGACAAGGTCCAAGGGG

GGCCTATGTACAACTATTTGCGATGCCAACCAATCAGCCCTGATCAGGGTGAGTCATACGAGACTGTTAGTGCATTCATC

ACCGCTGACCTTAAGAAGTATTGCCTGAATTGGAGATACGAGACAATCAGCATATTTGCACAGAGGCTGAATGAAATATA

TGGACTACCATCCTTCTTTCAATGGTTACACAGGATATTGGAAAAATCCGTACTCTATGTCAGTGACCCACATTGCCCTC

CCGACCTAGATAATCATATCCCTCTGGACAGTGTCCCTAATGCCCAAATATTCATAAAGTACCCAATGGGCGGAATAGAA

GGTTATTGCCAAAAACTATGGACAATCAGTACCATACCGTACTTGTATCTGGCAGCCTATGAGAGCGGAGTAAGAATCGC

CTCTCTAGTGCAGGGTGACAATCAGACAATCGCAGTGACAAAAAGAGTTCCAAGTTCTTGGCCTTATTCACTAAAAAAGA

GAGAGGCATCCAAAGCCGCTCAAAATTACTTCGTAGTCTTAAGGCAAAGATTGCACGATGTAGGTCATCACTTAAAGGCT

AATGAAACCATAGTATCCTCTCACTTTTTTGTATATTCCAAGGGGATTTATTATGACGGCCTGCTAATCTCACAGTCACT

CAAGAGCATCGCTAGATGTGTCTTCTGGTCCGAGACTATAGTGGATGAAACCAGAGCGGCTTGCAGCAATATTGCAACAA

CTGTCGCCAAGAGTATAGAGAGGGGTTATGATAGGTACCTTGCATATTCTCTAAATATCCTCAAAATTTTCCAACAGATC

CTTATATCCCTTAACTTCACTATTAACACAACAATGACTCAGGATGTCGTGGCACCGATCATCGAAAACGGGGATTTGCT

GATAAGGATGGCACTCTTGCCAGCACCCATTGGGGGTCTCAATTATCTTAACATGAGCAGGTTATTTGTGAGAAATATCG

GTGATCCGGTCACCTCCTCTATAGCCGACCTGAAGAGGATGATAGATGCCGGGCTAATGCCAGAAGAAACATTGCATCAA

GTGATGACCCAGACCCCGGGAGAATCATCCTACCTCGATTGGGCAAGTGACCCTTATTCTGCCAACCTACCCTGCGTACA

GAGTATAACTCGCCTTCTCAAAAACATCACTGCACGATATATTTTAATCAGCAGCCCTAATCCGATGCTGAAAGGACTAT

TTCATGAGGGGAGTAGAGATGAAGACGAGGAGCTTGCAAGTTTCTTAATGGATAGGCATATAATTGTTCCAAGAGCTGCA

CATGAAATCTTAGACCATAGCATAACCGGGGCAAGAGAAGCTATAGCCGGGATGTTGGACACCACCAAGGGTCTGATTAG

AACAAGTATGAGACGGGGTGGCCTCACCCCTCGAGTCTTAGCCCGCCTTTCCAATTATGATTATGAACAATTCCGATCCG

GGATAACACTATTGACAAAGAAAGGGCAGTGTTATCTCATTGACAAGGACTCGTGCTCGGTGCAGCTAGCTATAGCTCTG

AGGAGCCATATGTGGGCTAGGTTGGCTCGCGGGAGACCAATCTATGGGTTGGAGGTGCCCGATATATTAGAATCGATGAA

CGGCTACCTTATCAAACGCCACGAGTCTTGTGCTATCTGTGAGACGGGCTCAAGTCACTACGGATGGTTCTTTGTCCCTG

CAGGGTGCCAGCTTGACGATGTATCTAGGGAGACTTCGGCTCTTCGTGTACCTTATGTCGGATCAACAACTGAGGAAAGG

ACAGACATGAAACTTGCTTTTGTTAGGTCTCCAAGTCGATCCCTCAAATCAGCAGTCAGAATTGCAACAGTTTACTCATG

GGCCTACGGGGATGATGAGAAATCATGGAGTGAAGCCTGGATGCTAGCTAGGCAGAGAGCCAACATCACCTTAGATGAAT

TGAGAATGATTACTCCAGTCTCTACATCCACCAACCTAGCCCACCGGTTGAGGGATCGGAGCACCCAGGTGAAGTACTCG

GGGACATCCCTTGTGAGGGTTGCAAGATACACAACCATCTCCAATGATAACCTGTCATTTGTGATATCTGAGAAAAAAGT

GGACACCAACTTCATTTATCAGCAAGGGATGCTGCTCGGTCTTGGGATCCTTGAGAACCTCTTTAGGCTAGAGGCCACCA

CCGGAGTATCCAACACAGTGCTACACCTGCACGTGGAAACAGAGTGTTGTGTTGTACCTATGGTGGATCACCCAAGGATA

CCGAGTCTCCGTAATATAAAAGTTACGGATGAGTTATGCACAAACCCTTTGATCTACGACAGATCTCCCATCATAGAGCA

CGATGCAACCCGACTATACTCACAGAGCCACAGGAGACATCTGGTGGAGTTTGTTACCTGGTCAACAAGCCAGCTTTACC

ATATACTGGCTAAGTCTACAGCAATGTCCATGATTGAGTTGATCACGAGATTCGAGAAAGATCACATGAATGAAATAGCC

GCCCTGATTGGCGATGACGACATCAACAGCTTCATCACAGAATTTTTGCTAGTAGAGCCTAGATTATTTATAGTTTACCT

TGGTCAGTGTGCTGCCATCAATTGGGCCTTTGATATACATTATCATCGGCCCTCGGGCAAGCACCAGATGGGGGAACTCC

TCTACTCTCTGCTCTCTCGGATGAGCAAAGGAGTGTATAAGATCTTCACCAATGCTTTGAGCCACCCCAAAGTTTACAAG

AAATTTTGGCGAAGTGGTGTCATTGAGCCGATCCATGGCCCATCCCTGGATACACAGAATTTACATGTCACTGTCTGTGA

CATGATATATGGGTCCTATGTCACCTATTTGGATCTTTTGCTGAATGATGAGCTAGATGATTACCCGTATTTACTCTGCG

AGAGTGATGAGGACGTGGTCACAGACAGATTCGACAACATTCAAGCCAAACATCTCTGTGTATTGGCCGATGTATATTGC

AGCTCCAAGAGGTGTCCTTCGATAATCGGGATGTCTCCTATAGAAAAATGTACCATTCTCACACATTACATCAAGGGAGA

ATCAGTACAATCCCCGTCAGGGACCTCATGGAACACTGATCCTCTTGTAGTAGATCATTACTCATGCTCTCTGACCTACC

TTCGCCGCGGTTCCATCAAACAAATCAGGTTGAGAGTGGATCCTGGGTTTGTATTTGAGGCGTTGACAGACGTCGACTTT

AAACAGCCTAGCAAGGCTAAGTTAGATATATCGGTTGTTGGGTTAACTGATTTCTCTCCTCCTTGCGACAACGTTGGTGA

TTTTCTAGGGACTATCAACACATTGAGGCACGATCTGCCTGTCACCGGAACCGGGGTCTCGAACTATGAAGTCCACGCTT

ATCGTAGAATTGGCCTGAATTCATCAGCATGTTATAAAGCCGTAGAGATCTCCACGTTAATCAAGCCATCCTTAGAAGTC

GGAGAGCATGGATTGTTCCTAGGAGAAGGTTCCGGTTCAATGCTGGCTGCGTACAAGGAAGTTCTTAAATTAACAAATTG

TTATTACAACAGCGGGGTAACAGCGGAGGGCAGAGCCGGACAGAGGGAAATATCTCCCTATCCTTCAGAGATGAGCCTAG

TAGAGAATCAAATGGGGATAGAGCGAAGTGTTAAAGTGTTGTTCAACGGCAAACCTGAAGTAACTTGGGTGGGGACCACC

GATTGCTACAAGTATATAATCAGTAACATCCAAACCTCTAGTCTGGGTTTTATACATTCAGATATTGAGACACTCCCAAC

CAAAGACGCTGTTGAGAAATTGGAAGAATTTGCCTCTATCCTATCCTTATCCTTGATTTTGGGAAAAATCGGCTCTATTA

CAGTTGTCAAAATTATGCCCATTAGTGGGGATTTTACCCAAGGCTTCATAGCCTATGCTATCCAGTATTTTAGGGAGAGC

CTGCTTGCCTATCCGAGATATAGCAACTTCATCTCGACTGAGTGTTACCTTATTATGATAGGATTAAAGGCCAATCGATT

GATAAACCCAGAAGCCATTAAACAAAGCATAATCAGAGGGGGGGTCAGGACTGCACCAGGACTTGTGAGCCATATATTAT

CAGAGAAACAGAAAGGTTGTATTCAATCTTTCTTGGGTGATCCTTATATCCAAGGAGACTTCAATAAGCACCTTAAATCC

CTAACCCCTATTGAGAAAATCCTGGTAAATTGTGGTCTCTCGATCAATGGCACAAAAATCTGTAGAGATCTAATCCACCA

TGATATTGCCTCCGGTCCAGACGGTCTTATGAGCTCTACAATTATTTTGTATAGGGAATTGGCTCATTTCAAAGATAATA

TAAGGAGTCAGCACGGAATGTTCCACCCCTACCCAGTACTGGCCAACAGCAGGCAACGTGAATTAATCCTTCGAATAGCC

AAGAAGTTCTGGGGGTATGTCCTGCTATATTCTGATGACCCGGCACTAATCAGACAAACAATCAAGAACTTGAAACAGAA

TCATCTAACCTTTGACTTACATAGTAACCTGTTTATTAAAGGGCTATCCAAAGCTGAGAAACTGCTGGTGCGGACGAGTT

CACTAAGAAGGGAATGGTTGTTCACTCTCGATACGAAAGAAGTGAAAGAGTGGTTCAAATTGGTAGGTTACAGTGCACTC

ATCAGGGGCTAGTTAGCTATATATCTGCTCCCTTTTCCTCCGCCAGAAGACCCCACTGACGATCCAAAAGATTAAAGAAA

ACTACATATTGGATAAGTATCTATTCCCAGCNNNNNNNNNN

>KR828813.1/goat/Nigeria/2013-02-15

ACCAAACAAAGTTGGGTAAGGATAGATCTTATAATAACTATGAACTGACAAACTTAGGAGTAAAGATCCTATTGTCGGGG

AGAGGAGGAGGAGCAAGATCTTCGATTATGGCAACTCTCCTTAAAAGCTTAGCATTGTTCAAAAGGAACAAAGACAAAGC

GCCGACGGCATCAGGTTCAGGAGGGGCCATCCGGGGGATTAAGAATGTTATCATAGTCCCGATTCCCGGAGACTCGTCCA

TCATTACCCGTTCAAGACTGCTCGACAGGCTTGTCAGATTGGCCGGAGATCCTGACATCAACGGGTCAAAGCTGACCGGC

GTGATGATCAGCATGTTATCTTTGTTCGTAGAGTCACCCGGGCAATTGATACAGCGAATCACAGATGATCCAGATGTTAG

TATCCGCCTTGTTGAGGTAGTTCAAAGTACTAGGTCTCAGTCCGGGCTGACCTTTGCATCACGCGGTGCTGATTTGGACA

GCGAGGCAGACATGTATTTTTCAACTGAGGGGTCCTCGAGTGGAGGCAAGAAAAGGATCAACTGGTTTGAGAACAGAGAA

ATAATAGACATAGAGGTGCAGGATCCAGAAGAGTTCAATATGTTGTTAGCTTCCATACTAGCACAAGTCTGGATCCTCCT

GGCCAAGGCGGTTACGGCACCAGATACGGCAGCCGACTCAGAGCTGAGAAGGTGGGTTAAATACACACAGCAAAGGAGAG

TGATTGGGGAATTTCGCTTTGACAAAGGATGGCTGGACGCAGTTCGTAACAGGATTGCAGAAGATCTATCACTTCGGCGG

TTCATGGTATCTCTGATACTTGACATCAAGAGGACCCCCGGCAACAAGCCAAGGATTGCAGAAATGATCTGCGACATTGA

CAACTATATTGTCGAGGCCGGACTCGCCAGTTTCATTCTTACTATCAAGTTTGGTATCGAAACCATGTATCCTGCACTAG

GCCTTCACGAGTTTGCAGGGGAATTGTCCACTATTGAATCCTTGATGAACTTGTATCAACAGCTAGGAGAAGTTGCACCC

TACATGGTAATTCTAGAGAACTCAATTCAGAACAAGTTTAGTGCAGGAGCTTATCCTCTCCTCTGGAGCTATGCGATGGG

TGTCGGAGTTGAGCTGGAGAACTCAATGGGGGGCTTGAACTTTGGCAGGTCATATTTTGACCCGGCCTATTTTCGTCTCG

GACAGGAGATGGTCAGAAGATCTGCAGGAAAGGTCAGCTCTGTAATTGCGGCTGAGCTTGGCATCACAGCAGAGGAAGCT

AAACTGGTCTCGGAAATCGCCTCGCAGACTGGGGACGAAAGAACCGCTAGGGGGACTGGGCCTCGACAAGCGCAGGTCTC

CTTCCTCCAGCACAGACCGGGAGAGGGAGAGTCGCCCACACCAGCGACCAGAGAAGGGGTCAAAGCTGCGATCTCAAACG

GATCTGAAGAGAGGGAAAGAAAGCAAACACGCTCAGGAAGGCCTAGAGGGGAGACCCCTGGCCAACTGCTCCTGGAAATC

ATGCCAGAGGATGAAGTCTCGCGAGAGTCTGGTCAAAACCCTCGGGAGGCTCAAAGATCGGCCGAGGCACTCTTCAGGCT

GCAGGCCATGGCCAAGATTCTAGAGGACCAGGAGGAGGGAGAAGACAACAGTCAGATCCACAACGACAAAGATCTCCTCA

GCTGAGCAGACGCACCCTCTGTCGAAATCAGTGAAGAGACATCGCCCACCAGTATTATAAAAAACTTAGGACCCAGGTCC

AAGCGATCACACATCGACACTCCAGTCAGTCGAGCGGAGACCACCGATGGCAGAAGAACAAGCATACCATGTCAACAAGG

GGCTGGAATGCATCAAGTCTCTCAAAGCCTCTCCCCCGGATCTATCCACCATCAGGTACACCATTGAGAGCTGGAGAGAG

GGGCTTAGCCCCTCGGGTCGTGCAACACCGAGCCCCGATACGTCCGAGGGAGATCATCAGAATATCAGCCAATCATGCTT

GCCAGCAATCGGATCAAACAAAGTCTACTTGTCTCCTGAAGATAATCTCGGATTTAGAGAGATCACTGGCAACGACTGTG

AGGTTGGGCTCGGAGGAGTCCAGGGGAAAGGATCCAACTCTCAAGTACAGCGTTACTATGTTTATAGTCACGGGGGTGAA

GAGATTGAGGGACTCGAGGATGCTGACTCTCTCGTGGTTCAAGCAGATCCTCCAGTTGCTGACATCCTCAATGGAGGAGA

GGATGGATCTGACGACAGCGATGTGGAATCTGGCCCAGATGATCCCGGCAGAGATACTCTATATGACCGGGGATCTGTTG

CCGGCAATGGTGTCGCTAGATCCACAGATGTCGAAAAACTAGAAGGTGCTGATATTCAAGAAGTTCTTAACTCCCAGAAA

AGCAAAGGAGGAAGATTCCAAGGCGGGAAAACCTTGAGGGTCCCGGAAGTACCCGATGTCAAGCACTCCAGACCATCAGC

CCAATCAATTAAAAAGGGCACAGACGGGAACTCAGTCTCATCTGGAACGGGGACAGAGTGTTTATCGATAAGTGGTGCAA

CCCAAGCTGTGCCAGAGTCAAGATGGGAGTTATCAGAGCAAAATGCGTCTGTGGGGAGTGTCCTCAAGTCTGCGAGGAGT

GCAAAGACGATCCAGGAGTTGACACAAGAATCTGGTACCATAGCATCACTGACTCAGCCTAAAGAGAATGACTCCGAGTA

TGAGTATGAGGATGATCTATTTACAGAGATTCAGGACATCCGTGCAAGCATTGCCAAGATCCATGATGACAACAAAACTA

TCCTCTCAAAGCTTGATTCTATACTGTTATTGAAAGGAGAGATCGACACTATCAAGAAACAAATCAGCAAGCAGAATATA

AGTATATCCACCATTGAGGGCCATCTCTCCAGCATAATGATAGCCATCCCGGGCTTTGGGAAGGACATCAAGGACCCAAC

ATCCGAGGTAGAGCTGAACCCAGATTTAAGACCTATAATAAGCCGTGACTCTGGCAGAGCTCTCGCGGAGGTCCTCAAGA

AACCCGCTGTGGATAGATCTCAGAAAAATGGAATCAAAGTCAACTCCAGCTCAAAGGGTCAGCTTCTTAAGGATCTCCAG

CTAAAACCTGTCGACAAGCAGGTAAGCTCTGCAATCGGATTTGTCCCATCCGACCATGAATCATCCAGAAATGTCATCCG

CTCCATAATCAAGTCGAGCAAGCTAAACATCGATCACAAGGACTATCTTCTAGATTTACTAAATGATGTGAAAGGCTCCA

AGGATCTTAAGGAATTCCACAAGATGCTAACAGCTATTCTCGCCAAGCACCCGTAACGTACCCTCCAATCACCATCTCAC

ACTCGGTTGAAAACATCCTCTCAATCAGGCTATTACAAAAAACTTAGGAGCAAGGGCAACTGAGCTTCGCAGACAAGATG

ACTGAGATCTACGACTTCGATAAATCAGCATGGGATGTCAAAGGATCAATCGCTCGCATAGAACCCACCACCTATCACGA

CGGCCGACTGATACCTCAGGTAAGGGTCATTGATCCTGGTCTGGGAGACAGAAAAGATGAGTGCTTCATGTATCTGTTTC

TCCTAGGAGTGATTGAGGATAACGACCCCCTGTCTCCCCCAGTCGGAAGAACCTTTGGTTCTTTACCTCTAGGGGTTGGT

AGGTCAACTGCTAAGCCGGAAGAACTACTAAGGGAGGCCACAGAACTAGATATAGTGGTGAGGCGCACTGCAGGACTAAA

TGAGAAACTGGTATTTTACAACAACACTCCGCTGTCCTTGTTAACGCCCTGGAGGAAAGTCCTGACAACCGGAAGTGTGT

TTAGCGCTAACCAGGTTTGCAATGCAGTCAACCTAGTCCCACTTGATACTCCCCAGAGATTCAGGGTTGTGTACATGAGC

ATAACTAGATTATCAGACAATGGTTACTATAGTGTCCCCAGAAAAATGCTGGAGTTCCGCTCAGCCAATGCAGTAGCCTT

CAACATCTTGGTTACACTAAGAATTGAAAATGGCACAAACCCCAGAAGATACATAGTCGGCTCATGGGAGAATCCAGAAG

TCACATTTATGGTACACGTGGGCAATTTCAGAAGAAAGAAGAACGAAGTATACTCTGCGGATTACTGCAAAATGAAGATT

GAAAAGATGGGTTTAGTTTTTGCCCTGGGAGGAATAGGCGGAACAAGTCTCCATATTAGAAGCACCGGGAAAATGAGCAA

AACCCTTCATGCACAGCTGGGGTTCAAGAAAATCTTATGTTACCCCCTAATGGATGTTAACGAGGATCTTAACCGATATC

TCTGGCGGGCAGAGTGCCGAATAGTCAAAATCCAAGCTGTCTTACAGCCATCAGTACCCCAAGAATTCCGTGTCTACGAT

GATGTCATCATCAACGATGATCAAGGCTTGTTCAAGATCTTGTAATTCATTTGCAAAGGAATGAACCCACTTCCGTGAAG

AGACCCCACCCTCTCCCA-CCAAGAAGCACACTCCAAGGATCCCCGACACAACGCCAACACAGCCACTCTCCCGATGGGA

CAAGGTCGGCC-CCCCAACGAACCCGGTCCAAGC-TGGGGGCCCCCCCCACGACCCGCCGGGCGCCGACAGAGCAGGCCG

GCGAGAAGCCCCCAAGAGGCCCAGACGGAGACCGAGCACCCCACCCCCCC-AAAACGCCCCCCA-AACACGCCCGAGGCC

AAGCGCCGGGCACCCCCAGGCCCGCCACCGGCGAGATAGAACCCCCAAGGGGGACCCCCCAAACCAAAGGGACCACCGGG

GAAGGCTCCGCCGAACCAGCACAGACCCGGCCCAAGCACGGCGGACCCCCAACACGAAATCTCCCAGAGCACCCTTTTCA

TCATCAAACAAAACTTAGGGGCCAAGTCCACACGACAGACCCCAGCCTGCCAGGCGGAAGGCCACAGAAGGGAAGGAGAC

ACGACCGGCCCAGAAGGAGGACACCCAACCACAGAAACACCCAGTCCCAGGGAAGGAG--CCCCCCCTCCACAGGGGGAG

GAACGAGCGAGAGACCCCGAGGAGAACCCGACCGGGAGCCAGGAGCCGGCGACGCCGGCGGCAAACAGGCCGGGCACACA

CCCGGCACCACCAGAGGAGCAGAGGAGAGCCCAACCCCACGTGGACCCAACAGCAGGGAATCCAACCTCGCCACAAAAAG

AGGCGAGGACCC-CCT-----TCCCCCCGCACACAACCGGCAGACAGACCGAGGGAGGACCACCTCCCAACCAGACAGGC

GGCTCCCCCGCAGCCGAGCTGCAGCTCCTCAGAACAGGACACCCCCGCACACTCCGAACACCCCTCCTCCCCCGGCCACC

GGGACAGGGGCTCCGCCACCACAGACCAAAACCACCGAGAGTCAGCCAGAGGTCCCCTCCGGTCCCGTCCAGCGCCCCCC

CGAAGAGCCCCCGACGGGGTCCCCCAGGCCGACAACCCATGCACGCACCGAAAGAGGAACCCGCCCGAGAACATCCGGCA

CATCCATGCGCAAACATCATGACACGGGTCGCAATCTTGACATTTCTGATTCTTTTCCCGAATGCTGTTATGTGCCAGAT

TCACTGGGGCAATCTATCCAAGATCGGGATTGTAGGAACAGGGAGTGCCAGCTATAAGGTGATGACTAGGCCAAGCCACC

AAACTCTGGTTATAAAGTTAATGCCAAATATAACAGCCATAGACAATTGTACAACATCGGAGATTGCAGAGTACAAGAGA

TTGCTGATCACAGTTTTAAAGCCTATAGAGGATGCTCTGTCAGTGATAACCAAGAACGTAAGACCAATTCAAACTCTGAC

ATCTGGGCGCAGAACTCGCCGTTTTGCTGGAGCTGTTCTGGCCGGAGTAGCACTTGGGGTAGCGACAGCTGCTCAGATAA

CTGCCGGAGTCGCCCTTCATCAATCATTGATGAATTCCCAAGCAATCGAGAGTTTAAAAACCAGTCTTGAGAAGTCGAAT

CAGGCAATAGAAGAAATCAGACTTGCCAATAAGGAGACCATACTGGCGGTACAGGGTGTCCAGGATTATATCAACAATGA

GCTTGTCCCTTCTGTCCATAGAATGTCATGCGAGCTGGTAGGTCACAAGCTCGGCCTCAAGCTCCTTAGGTACTACACCG

AGATCCTGTCCATCTTCGGGCCCAGCCTTCGAGACCCGATAGCTGCCGAAATATCAATCCAGGCACTCAGTTATGCATTA

GGCGGAGACATTAATAAAATCCTGGAAAAGCTTGGGTATAGCGGTGGGGATTTCCTTGCTATTCTAGAAAGCAAGGGAAT

AAAGGCCCGGGTCACATATGTGGACACAAGAGATTACTTTATAATTCTTAGCATCGCCTACCCAACCTTATCTGAGATCA

AGGGAGTGATAGTTCACAAGATAGAAGCTATAACATACAACATCGGGGCACAGGAGTGGTATACTACTATCCCTAAATAT

GTAGCCACTCAGGGGTATCTGATATCAAACTTTGATGAGACGTCATGTGTATTCACTCCGGAAGGGACAGTTTGCAGCCA

AAATGCGTTGTACCCAATGAGCCCATTGCTTCAGGAATGTTTCAGGGGGTCGACAAAATCATGCGCCAGAACCCTCGTTT

CAGGGACCATAAGTAATAGATTCATCCTATCAAAAGGGAACCTGATTGCAAATTGTGCGTCAGTCTTGTGCAAATGTTAC

ACAACGGAGACAGTTATCAGCCAAGATCCTGATAAACTACTAACTGTTGTAGCATCCGATAAGTGTCCTGTAGTTGAGGT

GGATGGAGTGACAATACAGGTCGGCAGTCGAGAGTATCCGGACTCTGTATACTTACACAAAATAGACTTAGGTCCAGCCA

TCTCCCTAGAAAAACTGGATGTAGGCACCAATTTAGGCAATGCAGTTACAAGACTGGAGAATGCAAAGGAGCTCCTAGAT

GCATCAGACCAAATACTGAAGACTATTAAAGGGGTACCTTTCAGTGGGAATGTGTACATAGCACTGGCAGCTTGCATTGG

GGTATCCCTAGGCCTCGTCACATTAATATGCTGCTGTAAGGGGAGGTGTAAGAATAAAGAAATCCCTATCTCCAAAATCA

AACCAGGGCTCAAACCCGACCTGACCGGGACCTCAAAGTCGTACGTGAGATCACTGTAGTCAGAATCAACTGAACCATCC

GGCATCACACACATACATGTGCGACACAAGCAGTCGGAGGACGCAGGAAACCCAGCTTATGATCACCCACCTGACCCCAT

TCCACGCTCCACTACACATTAGTCATCAAACAAAACTTAGGACGAAAGGTCAATCACCATGTCCGCACAAAGGGAAAGGA

TCAATGCCTTCTACAAAGACAATCCTCATAATAAGAACCATAGGGTGATCCTGGATAGAGAACGCTTGATCATTGAAAGA

CCCTACATCTTGCTAGGAGTCCTGCTAGTAATGTTCCTGAGTCTAATCGGACTGCTGGCCATTGCAGGGATCAGGCTTCA

CCGGGCCACCGTTGGAACTTCAGAGATCCAGAGTCGGCTGAATACCAATATTGAGTTGACCGAATCTATTGATCACCAAA

CTAAGGATGTCTTAACTCCCCTGTTTAAAATCATTGGTGATGAAGTCGGCATCAGAATTCCACAGAAATTCAGTGATCTT

GTCAAGTTCATCTCCGATAAGATTAAGTTCCTCAACCCTGATAGAGAGTATGATTTCAGGGATCTCCGGTGGTGTATGAA

CCCCCCCGAGAGAGTCAAAATTAATTTTGATCAGTTTTGTGAGTACAAAGCCGCGGTTAAGTCAATTGAACATATATTTG

AGTCACCACTCAGCAAGTCAAAAGAGTTGCGATCATTGACTCTTGGGCCCAGAACAGGTTGTCTAGGCAGAACAGTAACA

AGAGCTCACTTCTCAGAACTTACACTGACCCTAATGGACCTGGACTTAGAGATGAAGCACAACGTGTCCTCAGTGTTTAC

CGTAGTTGAAGAGGGGTTATTCGGAAGAACATATACCGTCTGGAGATCAGATGCCAGGGATTTGAGCACCGATATAGATA

CTGGCCATTTTTTAAGGGTCTTCGAGATCGGACTGGTAAGAGATCTTGGGCTGGGTCCTCCTGTTTTCCATATGACCAAC

TATCTCACAGTGAACATGAGTGATGACTATCGGAGATGTCTTTTAGCGGTAGGGGGGTTGAAGCTGACAGCCCTATGCAC

CCCGTCCGAGACTGTGACACTGAGTGAGAGAGGGGTTCCAAAGAGGGAGCCTCTTGTGGTTGTGATACTTAACCTCGCTG

GGCCCACTCTAGGGGGTGAACTATACAGTGTCTTGCCTACCTCTGACCTCATGGTGGAGAAACTCTATTTATCTTCACAT

AGAGGGATTATCAAAGATGATGAGGCCAGTTGGGTAGTGCCGTCTACCGATGTTCGTGATCTTCAAAACAAAGGAGAATG

TCTGGTGGAAGCGTGCAAGACTCGACCTCCCTCATTTTGCAATGGCACAGGATCTGGCCCGTGGTCAGAGGGGAGAATCC

CTGCTTACGGGGTGATCAGAGTCAGTCTTGACTTGGCTAGGGACCCAGGTGTAGTTATCAATTCAGTGTTTGGCCCATTG

ATACCTCACCTATCCGGCATGGATCTTTACAACAATCCGTTCTCAAGAGCTGTATGGTTGGCTGTACCACCTTATGAGCA

GTCATTTCTAGGAATGATAAATACAATTGGATTCCCGAACAGAGCAGAGGTTATGCCGCACATTTTGACCACAGAGATCA

GAGGCCCTCGGGGTCGTTGCCACGTTCCCATAGAATTGTCCCGCAGGGTTGATGACGATATCAAGATCGGGTCCAACATG

GTCATATTACCGACGAAGGACCTGAGGTACATTACAGCCACTTATGATGTTTCCAGGAGCGAGCATGCGATCGTGTACTA

TATCTATGACACAGGCCGTTCATCATCTTACTTCTACCCGGTTCGACTGAATTTCAAAGGCAATCCTCTCTCTCTGAGGA

TAGAGTGTTTCCCTTGGCATCATAAGGTGTGGTGCTATCATGATTGTCTGATATACAACACCACATCAGGTGAAGAGGTC

CATACGAGAGGGCTAACCGGTATAGAGGTAACATGTAATCCAGCCTGAGCAGAGCTACAACCATCACTCAGGCAGACCCT

TGAGTCGTCACCGAGTCCAAGCAGCACAGCCTGGGACACTCAACAGCACAGCTCAGCCAACAATGTTATAAAAAACTTAG

GAGCCAAGGTTTTAGAGGCCATGGACTCCCTATCAGTCAATCAGGTCTTGTACCCTGAGGTTCATCTAGATAGCCCTATT

GTCACAAACAAACTAGTTGCCATCCTTGAGTACTCGGGGATCGACCATAACTATGTTCTTGAGGACCAGACTCTTATCAA

AAATATTAGGTATAGACTGGGATGCGGCTTTTCAAATCAAATGATCATCAATAATAGGGGGGTGGGTGAAACAGTCAACT

CTAAACTTAAAAGTTATCCTCATAATCGTCATATCATATACCCAGATTGCAATAAGGAGTTATTTTGTATCAAAGATAGC

TGCATATCTAAGAAGCTCTCGGAGCTATTCAAGAAGGGTAATTCCTTGTACTCTAAGATAAGTCACCGGGTACTGGATTG

TCTTAAGAGAGTCAACGGGAAATTAGGGCTGGGCACAGATCTTACTCATGGTCTGAAGGAGGGTATCCTTGACCTAGGGT

TGCACATGCATAGCTCTCAATGGTTCGAGACTTTTCTTTTCTGGTTCACTATCAAGACAGAGATGAGATCAATGATCAAA

GAACAGTCCCACATATGCCACAAGAGGAGGTATAACCCAATTTTTGTGTCAGGGGATGCATTCGAGGTGCTCGTATCACG

AGACCTCGTTGTGATAATTGATAAAAATACCCAGTATGTCTTCTACTTAACATTTGAGCTGGTCCTTATGTACTGTGATG

TCATAGAGGGCAGACTTATGACGGAGACAGCCATGGCCATAGACCAGAGATATTCAGAACTTCTAAGCCGGGTCAGATAC

TTGTGGGATCTTATTGATGGGTTCTTCCCAACACTAGGCAACACCACATACCAAGTTGTTGCTCTGCTTGAACCATTATC

GTTAGCTTATCTTCAACTTCAGGATGTCACTCTGGAGTTAAGAGGTGCTTTTTTAGACCATTGTTTCAAAGAACTCTATG

AAATACTGGAGCATTGCGGCATTGACACAGAAGGCACCTACAATTCCATCACTGAAGGATTGGATTACGTATTCATCACC

CATGATATACATTTAACTGGGGAGATTTTTTCATTTTTCCGGAGTTTCGGACACCCCCGTCTGGAAGCGGTCACCGCTGC

AGAAAATGTCAGAAAACATATGAACCAACCGAAGGTAATCAGTTATGAGACCATGATGAAAGGGCATGCAGTATTTTGCG

GGATAATCATAAATGGTTTTAGGGACCGACATGGTGGCAGCTGGCCCCCTGTTGCATTGCCAGAACATGCTTCTGCTGCG

ATCCGAAATGCGCAGGCATCAGGTGAGGGACTGACCCATGACCTGTGTATAGACAATTGGAAGTCTTTTGTTGGATTCAG

ATTTGGCTGCTTCATGCCACTCAGCCTAGATAGTGATTTGACCATGTACCTCAAAGACAAGGCCTTGGCTGCACTGAAGA

ATGAGTGGGATTCGGTTTACCCGAAAGAGTACCTCCGTTACAACCCACCTAGAGGGACAGAGTCAAGGCGACTGGTGGAG

GTGTTCCTGAATGACTCCAGCTTTGATCCTTATAACATGATAATGTACGTGGTGAATGGCTCCTACCTTAAAGACCCTGA

GTTTAATCTCTCATACAGCCTAAAAGAGAAAGAGATAAAAGAGACAGGGCGGTTGTTTGCCAAAATGACCTATAAGATGC

GGGCCTGTCAGGTAATCGCCGAAAATCTGATATCGAATGGTGTTGGGAAATATTTCCGAGACAATGGGATGGCAAAAGAC

GAGCATGACCTAACAAAAGCCCTTCACACCCTGGCAGTCTCAGGTGTTCCCAAGAATAACAAAGACAACCACCGAGGTGG

GCCTCCCAGAAGGACAACCAGCCGAGAGTTGAGATCGAGCCAATACACCAAAAAACAGAATAGAGACAAGGTCCAAGGAG

GGCCTATGTACAACTATTTGCGATGCCAACCAATCAGCCCTGATCAGGGTGAGTCATACGAGACTGTCAGTGCATTCATC

ACCGCTGACCTTAAGAAGTATTGCCTGAATTGGAGGTACGAGACAATCAGCATATTTGCACAGAGACTGAATGAGATATA

TGGACTACCATCCTTCTTTCAATGGTTACACAGAATATTGGAAAAATCCGTACTTTATGTCAGTGACCCACATTGCCCTC

CCGACCTAGATAATCATATCCCTCTGGACAGTGTCCCTAATGCCCAAATATTCATTAAATACCCAATGGGCGGAATAGAA

GGTTATTGTCAAAAACTATGGACAATCAGCACTATACCATACTTGTACCTGGCAGCCTATGAGAGCGGAGTAAGAATCGC

CTCACTAGTGCAGGGTGACAATCAGACAATCGCAGTGACAAAAAGAGTTCCAAGTTCTTGGCCTTATTCACTAAAAAAGA

GAGAGGCATCCAAAGCTGCTCAAAATTACTTCGTAGTCTTGAGGCAAAGATTGCACGATGTAGGTCATCACTTAAAGGCT

AATGAAACCATAGTATCTTCTCACTTTTTTGTATATTCTAAAGGGATTTATTATGATGGCCTGCTAGTCTCACAATCACT

AAAGAGCATCGCCAGGTGTGTCTTCTGGTCCGAGACTATAGTGGATGAAACCAGAGCGGCCTGCAGCAATATTGCAACAA

CTATCGCTAAGAGTATAGAGAGGGGTTATGATAGGTACCTTGCATATTCTTTGAATATCCTAAAGATCTTCCAGCAGATC

CTTATATCCCTTAACTTCACTATTAAGACAACAATGACTCAAGATGTCGTGGCACCGATCATCGAGAACGGTGATTTGCT

GACAAGGATGGCACTCTTGCCAGCACCCATCGGGGGTCTCAATTATCTTAACATGAGCAGGGTATTTGTGAGAAATATCG

GTGACCCGGTCACCTCATCTATAGCCGACCTGAAGAGGATGATAGACGCTGGGCTAATGCCAGAAGAAACATTGCACCAA

GTGATGACCCAGACCCCGGGAGAATCGTCCTATCTCGATTGGGCAAGTGACCCTTATTCTGCCAACCTACCCTGCGTACA

GAGTATAACTCGCCTTCTCAAAACCATCACTGCACGATATATTTTAATCAGCAGCCCAAAACCGATGCTGAGAGGACTGT

TTCATGAGGGAGGTAAAAATGAAGATGAGGAGCTTGCAAGTTTCTTGATGGATCGGCATATAATTGTTCCAAGAGCTGCA

CATGAAATTCTAGACCATAGCATAACCGAGGCAAGAAAGGCTATAGCCGGGATGTTGGATACCACCAAAGGTCTGATAAG

AACAAGTATGAAACGGGGTGGCCTCACCCCTCGAGTCTTAGCCCGCCTTTCCAATTATGATTATGAACAATTCCGATCCG

GGATAATACTATTGACAAAGAAAGGGCAGTGTTATCTCATTGACAAGGACTCGTGCTCGGTGCAGCTAGCTATAGCTCTG

AGGAGCCACATGTGGGCTAGGTTGGCTCGCGGGAGAACTATCTATGGATTGGAGGTGCCCGATATATTGGAATCGATGAA

CGGCTACCTCATCAAACGCCACGAGTCTTGTGCTATCTGTGAAACGGGCTCAAATCACTACGGATGGTTTTTCGTCCCTG

CAGGGTGCCAGCTTGACGATGTATCTAGAGAGACCTCGGCTCTTCGTGTACCTTATGTCGGGTCAACCACTGAGGAAAGG

ACAGATATGAAACTTGCTTTTGTTAGGTCTCCAAGTCGATCCCTTAAATCAGCAGTCAGGATCGCAACAGTTTACTCATG

GGCCTACGGGGATGATGAGAAATCATGGAATGAAGCCTGGATGCTAGCTAGGCAGAGAGCCAATATCACCTTAGATGAAT

TAAGAATGATCACTCCAGTCTCTACATCCACCAACCTAGCCCACCGGTTGAGGGATCGGAGCACCCAGGTGAAATACTCG

GGGACATCTCTTGTGAGGGTTGCAAGATACACAACCATCTCCAATGATAACTTGTCATTTGTGATATCTGAGAAAAAAGT

GGATACCAACTTCATTTACCAGCAAGGGATGCTGCTTGGTCTTGGGATCCTTGAGAACCTCTTCAGGCTAGAGGCCACCA

CCGGAGTATCCAACACAGTGTTACACCTGCATGTGGAAACAGAATGTTGTGTTGTACCTATGGTTGATCACCCAAGGATA

CCGAGTCTCCGTAATATAAAAGTTACGAATGAGCTATGCACAAACCCTTTGATCTACGACAGGTCTCCCATCATAGAACA

CGATGCAACCCGACTATACTCACAGAGCCACAGGAGACACTTGGTGGAGTTTGTTACCTGGTCAACAAGCCAGCTTTACC

ATATACTAGCCAAGTCTACAGCAATGTCCATGATTGAGTTGATCACGAGATTCGAGAAAGATCACATGAATGAAATAGCC

GCCCTGATTGGCGATGACGACATCAACAGCTTCATCACAGAATTCTTGCTAGTAGAGCCTAGATTATTTATAGTTTACCT

TGGTCAGTGTGCTGCCATCAATTGGGCTTTTGATATACATTATCATCGGCCCTCGGGCAAGTACCAGATGGGGGAACTCC

TCTACTCTCTACTCTCTCGGATGAGCAAAGGGGTATATAAGATCTTCACTAATGCTTTGAGCCACCCCAAAGTTTACAAG

AAATTTTGGCGAAGTGGTGTCATCGAGCCGATTCATGGACCGTCCCTGGATACACAGAATTTACATGTCACTGTCTGTGA

CATGATATATGGGTCCTACGTCACCTATCTAGATCTTTTGCTGAATGATGAGCTCGATGATTATCCGTATTTGCTCTGCG

AGAGTGATGAGGACGTGGTCACAGACAGGTTCGACAACATTCAAGCCAAACATCTCTGTGTATTGGCCGATGTATATTGC

AGCTCTAAGAGGTGTCCTTCGATAATCGGGATGTCTCCCATAGAAAAATGTACCATTCTCACACATTACATCAAGGGAGA

ATCAATACAAACCCCGTCTGGGACCTCATGGAACACTGATCCTCTTGTAGTAGATCATTACTCATGCTCTCTGACCTACC

TTCGCCGCGGTTCTATCAAACAAATCAGGTTGAGAGTGGATCCTGGGTTTGTATTCGAGGCATTGACAGACGTCGACTTT

AAACAGCCTCGCAAAGCCAAGTTGGATATATCGGTTGCGGGGTTGACTGATTTCTCTCCCCCTTGGGACAACATCGGTGA

TTTTCTAGGGACTATCAATACACTGAGGCATAATCTGCCCGTCACCGGAACCGGGGTCTCGAACTATGAAGTCCACGCTT

ATCGTAGAATTGGCCTGAATTCATCAGCATGTTATAAAGCCGTAGAGATCTCCACGTTAATCAAGCCATCCTTAGAAGTC

GGAGAGCATGGATTGTTCTTAGGAGAAGGTTCCGGTTCAATGCTGGCTGCGTACAAGGAAGTTCTTAAATTGGCAAACTG

TTATTACAACAGCGGGGTAACAGCGGAGGGCAGAGCCGGACAGAGGGAAATATCTCCCTATCCTTCAGAGATGAGCCTAG

TAGAGAATCAAATGGGGATAGAGAGGAGTGTTAAAGTACTATTCAACGGCAAACCTGAAGTAACTTGGGTGGGGACCACC

GATTGCTACAAGTACATAATCAGTAACATCCAAACCTCTAGTCTAGGTTTCATACACTCAGATATTGAGACACTCCCAAC

CAAGGATGCTGTTGAGAAATTAGAAGAATTTGCCTCTATCCTATCCTTATCCCTCATTTTGGGAAAAATCGGCTCTATCA

CAGTTGTCAAAATTATGCCCATTAGCGGGGATTTTACCCAAGGCTTCATAGCCTATGCAATCCAATATTTTAGGGAGAGC

CTGCTTGCCTATCCGAGATATAGTAACTTCATCTCGACTGAGTGTTACCTTATTTTGATAGGATTAAAGGCCAATCGATT

GATAAACCCGGAAGCCATTAAGCAAAGCATAATCAGAGCGGGGGCCAGGACTGCACCAGGACTTGTAAGCCATATATTAT

CAGAGAAACAGAAAGGTTGTATCCAATCTTTCCTGGGTGATCCTTATATCCAAGGAGACTTCAACAAGCACCTTAAATCT

CTAACCCCTATTGAGAAAATCCTGGTAAATTGTGGTCTCTCGATCAATGGCACAAAAATCTGTAGAGATCTAATCCACCA

TGATATCGCCTCCGGTCCAGACGGTCTTATGAGCTCTACAATTATTTTATATAGGGAATTGGCTCATTTCAAAGACAATA

TAAGGAGTCAGCACGGAATGTTTCACCCCTACCCAGTACTGGCCAGTAGCCGGCAACGTGAATTAATCCTTCGAATAGCC

AAGAAATTCTGGGGTTATGTCTTGCTATATTCTGATGACCCGGCACTAATCAGGCAAACAATCAAGAACTTGAAGCGGAA

TCATTTAACCTTTGACTTACACAGTAACCCGTTTATTAAAGGGCTATCCAAAGCTGAGAAACTGCTGGTGCGTACGAGTT

CACTCAGAAGGGAATGGCTGTTTACTCTCGATACGAAAGAAGTGAAAGAGTGGTTCAAATTAGTGGGTTACAGTGCACTC

ATCAGAGGCTAGTTAGCTATACGTCTGCCCCCTTCTCCTCTGCCATGAGACCCTACTGACGGTCTAAAAGATTAAAGAAA

ACTACATATTGGATAAGTATCTATTCCCAGCTTTGTCTGGT

>KP789375.1/goat/Senegal/1969-09-03

ACCAAACAAAGTTGGGTAAGGATAGTTCTTATAATAACTATGGATCAGGAAACTTAGGAGTAAAGATCCTACTGTCGGGG

GGAGGAGGAGGAGCAAGATCTTTGACCATGGCGACTCTCCTTAAAAGCTTGGCATTGTTCAAGAGGAACAAAGATAAAGC

GCCGACAGCATCAGGTTCAGGAGGGGCCATCCGGGGGATTAAGAATGTTATCATAGTCCCAATTCCCGGAGATTCATCTA

TCATTACCCGTTCAAGACTGCTTGACAAGCTTGTCAGATTGGCTGGAGATCCTGACATCAATGGGTCAAAGCTGACCGGC

GTGATGATCAGCATGTTATCTTTGTTCGTAGAGTCACCCGGGCAATTGATACAGCGGATCACAGATGACCCAGATGTCAG

TATCCGCCTTGTTGAGGTGGTTCAAAGTACTAGGTCTCAGTCAGGGTTGACCTTTGCATCACGTGGTGCTGATTTGGACA

ATGAGGCAGACATGTATTTTTCAACTGAAGGACCCTCGAGTGGAAGTAAGAAAAGGATCAACTGGTTTGAGAACAGAGAA

ATAATAGACATAGAGGTCCAGGATGCAGAAGAGTTCAATATGTTATTAGCATCCATACTAGCACAAGTTTGGATCCTCCT

GGCCAAGGCAGTTACGGCACCGGATACGGCAGCTGACTCAGAACTGAGAAGGTGGGTTAAATACACACAACAGAGGAGAG

TGATTGGGGAATTTCGCCTTGACAAAGGTTGGCTGGACGCAGTCCGCAATAGGATTGCAGAAGATCTATCACTTCGGCGG

TTCATGGTATCTCTCATACTTGACATCAAGAGGACCCCCGGGAACAAGCCAAGGATTGCAGAAATGATCTGCGACATCGA

CAACTATATTGTCGAAGCCGGACTTGCCAGTTTCATCCTTACCATCAAGTTTGGTATTGAAACCATGTATCCTGCATTAG

GTCTTCACGAGTTTGCCGGGGAATTGTCCACTATAGAATCCCTGATGAACTTGTATCAACAACTAGGGGAGGTTGCACCC

TACATGGTAATTCTAGAGAACTCAATTCAGAACAAGTTTAGTGCAGGAGCTTATCCCCTCCTGTGGAGCTATGCCATGGG

TGTCGGAGTTGAGCTGGAGAACTCAATGGGGGGCCTGAACTTTGGCAGGTCATATTTTGACCCGGCCTATTTTCGTCTTG

GACAGGAGATGGTCAGAAGATCTGCAGGGAAGGTCAGCTCTGTAATCGCGGCTGAGCTTGGCATCACAGCAGAGGAAGCT

AAACTAGTCTCGGAAATCGCCTCGCAGACTGGGGACGAAAGGACCGTTAGGGGGACTGGGCCACGACAGGCGCAGGTTTC

TTTCCTCCAGCATAAAACAGGAGAGGGAGAGTCATCTGCACCAGTGACCAGAGAAGAAGTCAAGGCTGCGATTCCAAATG

GGTCCGAAGAAAGGGACAAAAGGCGAACCCGCTCAGGAAGACCCAGAGGAGAGACCCCGGGCCAACCGCTCTTGGAAATC

ATGCCAGAGGATGAGGTCTCGCGAGAATCTGGTCAAACCCCTCGTGAGGCTCAGAGGTCGGCCGAGGCACTCTTCAGGCT

GCAAGCTATGGCCAAGATTCTAGAGGGCCAGGAGGAGGGAGAAGATAACAGTCAGATCTACAACGACAAGGATCTCCTCA

GTTGAGCAGAAGCTCCCTCTGTCAACATCAGCAGCAAAATATCGCCCATCAGTGTTATAAAAAACTTAGGACCCAGGTCC

AAGCAGTCACACATCGAAACTCTAGCCAACCGAGCAGAGACCACCGATGGCAGAGGAACAAGCATACCATGTCAACAAGG

GACTGGAATGTATCAAGTCCCTCAAAACCTCTCCCCCAGATCTATCCACCATCAGAGATACCCTTGAGAGCTGGAGAGAG

GGGCTTGGCCCCTCGGATCGTGCAACACCGAACCCTGATGCGTCCGAGGGAGACCATCAGGATATCAACCAATCATGCTC

ACCAGCAGTCGGATCAGGCAAAATCGACGTGTCTCCTGAAGGTAATCTCGGATTTAGAGAGATCACTTACGATGACAGTG

AGGCAGGGCTCAGAGGGATTCAGGACAAAGGATCCAACTCTCAAGTACAGCGTTACCATGTTTATAGCCACGGGGGTGAA

GAGATTGAAGGACTCGAGGATGCTGACTCTCTCGTGGTTCAAGCAGATCCTCCAGTTGCTAACACCTTCAGTGGAGGAGA

AGATGGATCTGACGACAGCGATGTGGACTCTGGCCCAGATGATCCCAGCAGAGATCCTCTATATGACTGGGGATCTGTTA

CCGGCCATGGTGTCGCTAAGTCCACAGATGTCGAAAAATTAGAAGGTGATGACATTCAAGCAGTTCTTAACTCCCAGAAG

AATAAAGGAGGAAGATTCCAAGGCGGGAAAACCTTGCGAGTCCCGGAAATACCCGATGTCAAGCACTCCGGACCATCAGC

CCAATCAATTAAAAAGGGCACAGACGGGAACTCAGTCTTATCTGGAACGGCGACAGAGTGTTTATTGATAAATGGTGCAA

CCCGAGCTGTGCCAGAGTCAAGATGGGAGTCGTCAGAGCGAAATGCATCTGTGGAGAGTGTCCCCAAATCTGCGAGGAGT

GCAAAGACGATCCAGGAGTTGACACAAGAATCTGGTACCATAGCATCACTGACTCAGCCTAAAGAGAATGACTCCGAGTA

TGAGTATGAGGATGGTCTATTTACAGAGATACAGGACATTCGTGCAAGCATTGCCAAGATCCATGAAGACAATAAAACTA

TCCTCTCAAAGCTTGAATCTATACTGTTATTGAAAGGAGAAATAGATACTATCAAGAAACAAATCAGCAAGCAGAATATA

AGTATATCTACCATTGAGGGCCATCTCTCCAGTATAATGATAGCAATCCCGGGTTTTGGGAAGGATATCAAGGACCCAAC

ATCCGAGGTTGAGTTGAACCCAGATTTAAGACCTATAATCAGCCGTGATTCTGGCAGAGCTCTCGCGGAGGTACTCAAGA

AACCCGCTGTTGATAGGTCTCAGAAAAATGGAATCAAAGCCAACTCAAGTTCAAAGGGTCAGCTCCTTAAGGACCTCCAG

CTAAAACCTGTTGACAAACAAGCAAGCTCTGCAATAGGGTTTATCCCGTCTGACCATGAGTCATCCAGAAGTGTCATCCG

CTCTATAATCAAGTCGAGCAAGCTAAACATTGATCACAAAGACTACCTTCTAGACTTACTGAATGATGTGAAAGGCTCCA

AGGATCTTAAGGAATTCCACAAAATGCTAACAACCATTCTTGCCAAGCAACCGTAACACAACCTCCAAACAACATCCCAT

ACTCGGCTGAACACATCTTCTCAACCAGGCTATTACAAAAAACTTAGGAGCAAGGGCAACCGAGCTTCACAGACAAGATG

ACCGAGATCTACGACTTCGATAAATCAGCATGGGATGTCAAAGGGTCAATTGCTCCCATCGAACCCACCACCTATCACGA

CGGCCGACTGATACCCCAAGTGAGAGTCATCGACCCTGGGCTAGGAGATAGAAAAGATGAGTGCTTCATGTACCTGTTTC

TTTTAGGAGTGATTGAGGATAACGACCCCCTATCCCCCCCTGTCGGAAGAACCTTTGGTTCTTTACCTCTAGGGGTCGGT

AGGTCAACTGCTAAGCCAGAGGAACTACTAAGGGAGGCCACGGAGCTAGACATAGTGGTGAGGCGCACGGCAGGATTAAA

CGAGAAGCTGGTATTTTACAACAACACTCCACTGTCCTTGCTAACACCCTGGAGGAAAGTCTTGACAACCGGAAGTGTGT

TCAGCGCTAACCAGGTTTGCAATGCAGTCAATCTGGTCCCACTTGATAGTCCCCAGAGATTCAGGGTAGTGTACATGAGC

ATCACTAGATTGTCAGATAATGGTTATTACAGTGTCCCCAGGAGAATGTTGGAGTTCCGCTCAGCCAATGCAGTCGCCTT

CAATATCTTGGTTACACTGAGAATCGAAAATGGCACAAACCCTAGAAGATACATAGTCGGCTCATGGGAGAATCCAGAGG

TCACATTTATGGTACACGTGGGCAACTTTAGAAGAAAGAAGAATGAAGTATATTCTGCTGATTACTGCAAAATGAAGATT

GAAAAGATGGGTCTAGTTTTTGCCCTGGGAGGAATAGGAGGAACAAGTCTCCATATTAGAAGCACCGGGAAAATGAGCAA

GACTCTCCATGCACAGCTAGGGTTCAAAAAAATCTTATGCTACCCCCTAATGGATATTAATGAGGATCTTAACCGATATC

TTTGGCGGGCAGAGTGCCGAATAGTCAAAATCCAAGCCGTCTTACAGCCATCAGTACCCCAAGAATTCCGTGTCTACGAT

GATGTCATCATCAACGATGACCAAGGCTTGTTCAAGATCTTGTAATTCACTCGCCGGACAATGGACCCACCTCCAAGAAG

TGGCTCCTCCTTCCCCCGTCCAAAAGACACACTCCAAGGCTCCCCAAAACAGCTCCAACATGGCCATCCTCCCAACGGGA

CGGGGCCAACCTCCCCCATGAACCCAGTCCGG--GGGGGGGCCCCCCCCGTGACCCACCGGGCGCCGACCGCGTGGGCCG

GCAGGACGCCCCCAGGAGGCCCCGACCAGGACCGGGCATCCTCCCCCCCC-CAAACGCTCCCCC-GACACATCCGAGGCC

AGGCGCCAGGCACTCCCAACCCCGCCACCGGGGGGACAGGCAGCCAAGACGGGGCCCCCCCACCCAAACGGACCGCCAGA

GGAGGCCCCGCCGACTCAGCACAGACCCGGCCCAAACAAGGGAGATTCCAAAGACGAAACCGCCCAGCGCACCCTACTCA

TCATCAAACAAAACTTAGGGGCCAAGTCCAGACGACAGACCCCAGCCCGCAACCCAGCAGGCCACAGAAAGGAAGCAGAC

AAGACCGGCCCGGAAGGCGGACACCCACCCACAGAAGCACCCAGGCCCCGGGGAGAAGCCTC-CCCTCCACACAGCGAGG

GAGCGGGCGAGAAACACCAGGGGGAATTCGATCGGGAGTCATGAGCCGGTGACGCCAACGGCAGCCAGACCGGGCACACA

CCCGACACCACCAGCAGAGCAGAGGAGAGCCCAGACCCGTGCGGACTCGACGGCAGGGGAGCCAACCCCGCCACAAAGAG

AGACGAGGGGCTC--------CCCCCAAGCACACAACCGGCAGACAGACCGAAAGAGGACCACTTCCTGGGAGGACAAGC

GGCTCCCCCGCAGCCGAGCCGCAACTCCCCAGGAGAGGACACCCCCGACGCCTCCGAACACCCTCCCGCCCCCGGCCACC

GGGAGAAGGGCTCCGCTGCCAAAGACCAGAGCCACCGAGAGCCAACCAGAGGCCCCCCCCAGTCCCGCCTGGGGCCCCCC

CGAAGAAGCCCCAGCAGGGTCCCCTGGCTCGACGACCCCTGCATTCGCCAAAAGAGAAACCCACCAGAGGACAACCAGCA

CATCCATGTATAAACATCATGACACGGGTCGCAACCTTGACATTTCTGTTTCTTTTCCCAAATACCATTGCATGCCAGAT

TCACTGGGGCAATCTATCCAAGATCGGGATTGTGGGAACAGGGAGTGCCAGTTACAAGGTGATGACTAGGCCAAGCCACC

AAACTCTGGTTATAAAGCTGATGCCAAATATAACAGCCATCGACAATTGTACGAAATCAGAGATTTCAGAGTACAAAAGA

TTGCTGATCACAGTGTTAAAGCCTGTAGAGGATGCTCTGTCAGTGATAACCAAGAATGTAAAACCAATTCAGACTCTAAC

GCCTGGGCGCAGGACCCGCCGTTTTGCCGGGGCTGTTCTGGCCGGAGTGGCACTTGGAGTCGCGACGGCTGCTCAGATAA

CTGCCGGAGTCGCACTTCATCAATCCTTGATGAACTCCCAAGCGATTGAAAGTTTAAAGACCAGTCTTGAGAAGTCGAAT

CAGGCAATTGAAGAAATAAGGCTTGCAAATAAGGAGACCATACTGGCAGTACAGGGTGTCCAAGACTATATCAACAGTGA

ACTTGTCCCTTCTGTCCATAAAATGTCATGTGAGCTGGTAGGTCACAAACTCAGCCTCAAGCTCCTTAGGTATTACACCG

AGATCTTGTCTATATTCGGGCCTAGCCTCCGAGACCCGATAGCTGCTGAAATATCAATCCAGGCACTCAGTTATGCATTA

GGTGGAGATATTAATAAAATCCTGGATAAGCTTGGGTATAGCGGCGGGGATTTCCTTGCTATCCTAGAAAGCAAGGGGAT

AAAGGCTCGAGTCACATATGTGGATACAAGAGATTACTTTATAATTCTAAGCATAGCCTACCCAACTTTATCTGAAATCA

AGGGGGTGATAGTTCATAAAATAGAAGCCATATCATACAACATTGGGGCACAGGAATGGTACACTACTATCCCTAAATAT

GTAGCCACTCAGGGGTATTTGATATCGAATTTTGATGAGACGTCGTGCGTATTCACTCCAGAGGGGACAGTCTGTAGTCA

GAATGCTTTGTACCCAATGAGCCCATTGCTTCAGGAATGTTTCAGGGGGTCGACAAAATCGTGCGCCAGAACTCTAGTTT

CAGGGACTACAAGCAATAGATTTATCCTGTCAAAAGGGAACTTGATTGCAAATTGTGCATCAGTTTTGTGCAAGTGTTAC

ACAACGGAGACAGTTATCAGCCAAGATCCTGATAAATTACTAACTGTTATAGCCTCCGATAAGTGTCCTGTAGTTGAAGT

GGATGGAGTGACAATACAGGTCGGCAGTCGTGAGTACCCAGACTCCGTGTACCTACACAAAATAGACTTAGGCCCAGCCA

TCTCTCTGGAAAAACTGGACGTAGGCACCAATTTAGGTAATGCAGTTACAAGACTGGAGAATGCAAAGGAGCTCTTAGAT

GCATCAGACCAGATATTGAAGACTGTTAAAGGGGTACCCTTCGGTGGCAATCTATACATAGGACTGGCAGCTTGCATCGG

AGTATCCCTAGGGCTTGTCACATTAATATGCTGCTGTAAGGGGAGATGTAGGAACAAAGAGATTCCTACGTCCAAAATCA

ACCCAGGGCTCAAACCTGACCTGACCGGGACCTCAAAGTCATACGTGAGATCACTGTAGTCAGAATAACCCGAATCACCA

AACATCACACACATACATGTGTGATACAAGCGGTCAGAGGACGCAGAAGATTCAACTTCCGATCACCAACCTGGCCCCGC

TCTATGCTCCATTACACATTGATCATCAAACAAAACTTAGGACGAAAGGTCAGTCACCATGTCCGCACAAAGGGAAAGAA

TCAATGCCTTCTACAAAGACAATCTTCATAACAAGAATCATAGGGTAATCCTGGATAGAGAACGCTTGACTATTGAAAGA

CCCTACATCTTACTTGGGGTCCTGCTGGTAATGTTTCTGAGCCTAATCGGGCTGCTGGCCATTGCAGGGATCAGACTTCA

CCGAGCCACCGTCGGAACTGCAGAGATCCAGAGTCGGCTTAATACCAACATTGAGTTAACTGAATCTATTGATCACCAAA

CCAAGAATGTCCTAACTCCGCTGTTTAAAGTCATTGGTGATGAAGTCGGCATAAGAATCCCACAAAAATTCAGTGATCTT

GTCAAGTTCATCTCCGATAAGATTAAGTTCCTCAACCCTGACAGAGAATATGATTTTAGGGATCTCCGGTGGTGTATGAA

TCCCCCCGAGAGAGTCAAAATTAATTTTGATCAGTTTTGTGAATACAAGGCTGCAGATAAGTCAATTGAACATATATTTG

AGTCATCACTCAACAGATCAAGAAGGTTGCGATTGTTGACCCTTGGGCCTGGAACAAGCTGTCTCGGTAGGGCAGTAACA

AGAGCTCAGTTCTCAAAGCTTACTCTGACCCTGTTGGATCTGGATCTCGAGATGAAACACAACGTGTCCTCAGTGTTCAC

CGTAGTTGAAGAGGGATTATTCGGAAGAACGTACACTGTCTGGAGATCTGATGCCGGGAATCCGAGCACCGACCCAGGTA

CTGGCCATTTTTTAAGAGTCTTCGAGATCGGGCTGGTAAGAGATCTCGGGCTGAGTGCTCCTGTTTTCCATATGACCAAT

CATCTCACAGTGAACATGAGCGATGGCTATCGGAGCTGTCTTTTGGCAGTAGGGGAGTTGAAGCTGGCAGCCCTATGCAC

CCCATCCGAGACTGTGACACTGAGTGAGAGAGGAGTTCCAAAGAGAGAGCCTCTTGTGGTTGTGATACTTAACCTAGCCG

GGCCTACTCTAGGTGGCGAACTATACAGTGTCTTGCCTACCTCTGACCTCACGGTAGAGAAACTTCATCTATCCTCACAC

AGAGGGATTATCAAAGATAACGAGGCCAATTGGGTAGTACCGTCCACCGATGTTCGTGATCTTCAAAACAAAGGAGAATG

TTTAGTGGAAGCATGTAAGACTCGACCTCCTTCATTTTGCAATGGCACAGGAGTGGGCCCATGGTCAGAGGGGAGAATCC

CTGCCTACGGGGTGATCAGGGTCAGTCTTGACTTAGCCAGTGACCCTGACGTAGTTATAACTTCAGTATTTGGCCCATTG

ATACCTCACCTGTCCGGCATGGATCTTTACAACAACCCGTTTTCAAGCGCTGTGTGGTTGGCGGTACCACCTTATGAGCA

GTCATATCTGGGAATGATAAATACAATTGGCCTCCCAAATAGAGCAGAGGTTATGCCGCACATTTTGACCACAGAGATCA

AGGGGCCTCGGGGTCGTTGTCATGTTCCCATAGAATTGTCCCGCAGGATTGATGATGATATCAAGATTGGGTCCAACATG

GTTGTATTGCCGACGAAGGATCTGAGGTACATAACAGCCACTTATGATGTTTCCAGGAGCGAGCATGCAATCGTGTACTA

TATCTATGACACAGGTCGCTCATCATCTTACTTCTACCCGGCTCGATTGAAGTTCAAAGGCAATCCTCTCTCTCTGAGGA

TAGAGTGTTTCCCCTGGCATCATAAGGTATGGTGCTACCATGATTGTCTTATATACAACACCATAACAAACGAAGAAGTC

CATAAGAGAGGACTGATCGGCATAGAGGTAACATGTAATCCAGCCTGAGCGGAGCTGCGACCATCACTCAAGCAGGCCTC

CGAGCTAATCCCAAGCCCAGGCAGCACAGCTCGGGACACTCGACAACACAACCCAGCCAACAATGTTATAAAAAACTTAG

GGGCCAAGGCTATAGGAGCCATGGACTCCCTGTCAGTCAATCAGGTCTTGTACCCTGAGGTCCATCTGGATAGCCCTATT

GTCACAAACAAGCTGGTTGCCATCCTCGAGTACTCAGGCATTGACCACAACTATGTCCTTGAAGACCAGACCCTTATCAA

GAATATTAGATATAGACTAGGGTGCGGTTTCTCAAATCAAATGATCATCAACAACAGGGGGGTAGGTGAAACAGTCAACT

CTAAACTTAAAAGTTACCCCCGTAATTGTCATATCATATACCCAGATTGCAATAAGGATCTGTTTTGTATCAAAGACAGC

TGCATATCTAGGAAGCTCTCTGAGCTATTCAAGAAGGGTAATTCCCTATACTCGAAGATAAGTCACCAGGTACTGGATTG

TCTTACAAGAGTCAACGGTAAATTAGGCTTGGGCACAGATCTTACTCATGGCCTGAAGGAGGGTATCCTCGACTTGGGGT

TACACATGCATAGCTCTCAATGGTTCGAGACTTTCCTGTTCTGGTTCACTATCAAGACAGAGATGAGATCAATGATAAAA

GAACAGTCCCATATATGCCACAAGAGGAGGTATAACCCGGTTTTTGTGTCGGGGGATGCATTCGAGGTGCTCGTATCACG

AGACCTCGTTGCGATAATTGACAAGAATACCCAGTATGTCTACTATTTGACATTTGAGCTGGTCCTTATGTATTGTGATG

TCATAGAGGGCAGACTTATGACGGAGACAGCCATGGCTATAGACCAGAGATATTCAGAACTTCTAAGCCGGGTCAGATAT

TTGTGGGATCTTATTGACGGGTTCTTCCCAACACTAGGTAACACCACATACCAGGTTGTTGCTCTGCTTGAACCATTGTC

GTTGGCTTATCTTCAACTTCAGGATGTCACCCTGGAGTTAAGAGGTGCTTTTTTAGATCACTGCTTCAAGGAACTCTATG

AGATACTAGAGCATTGCGGCATTGACACGGAAGGCACCTACAATTCCATCACTGAAGGATTGGACTACGTATTTATCACC

CACGATATCCATTTAACTGGGGAGATATTTTCGTTTTTCCGGAGTTTCGGACACCCCCGCCTCGAAGCGGTCACCGCCGC

AGAGAATGTCAGAAAACATATGAACCAGCCGAAGGTAATTAGTTATGAGACTATGATGAAAGGGCATGCAGTGTTTTGCG

GGATAATCATAAATGGGTTTAGAGACCGGCATGGCGGCAGCTGGCCCCCCGTTGCATTGCCAGAACATGCTTCTGCTGCG

ATTCGGAATGCGCAGGCATCCGGTGAGGGGCTGACCCATGACTTGTGTATAGACAACTGGAAGTCCTTTGTTGGATTCAG

ATTTGGCTGTTTCATGCCGCTCAGCCTAGATAGTGATTTGACCATGTACCTTAAAGACAAAGCACTGGCTGCACTGAAGA

ATGAGTGGGATTCAGTTTACCCGAAAGAATACCTCCGCTATAATCCACCCAGAGGGACAGAGTCAAGGCGGCTGGTGGAG

GTGTTCCTGAATGACTCCAGCTTTGATCCTTATAACATGATAATGTACGTGGTGAATGGCTCTTACCTTAAAGACCCTGA

GTTTAATCTCTCATATAGCTTAAAGGAGAAGGAGATAAAAGAGACAGGGCGGTTGTTTGCAAAAATGACTTACAAGATGC

GGGCCTGTCAGGTAATTGCGGAGAATCTGATATCAAATGGGGTCGGGAAGTATTTCCGAGACAATGGAATGGCGAAAGAC

GAGCATGACCTAACAAAAGCCCTCCACACTCTGGCAGTCTCAGGTGTTCCCAAGAACAATAAAGATAACCACCGAGGTGG

CCCTCCTAGAAGGACCACAAGCCGAGAGATGAGATCAAACCAAGACATCAACAGACAAACTAGAGACAAGGTCCAGGGGG

AGCCCATGTACAACTATTTGCGGTGCCAACCAATCAGCCCTGACCAGAGTGAGTCATACGAGACTGTTAGTGCATTCATC

ACCGCTGACCTTAAGAAGTATTGCCTGAATTGGAGATACGAGACAATCAGCATATTTGCACAAAGACTTAATGAAATTTA

CGGACTGCCATCCTTCTTTCAATGGTTACACAGGGTATTGGAAAAATCTGTGCTCTACGTCAGTGACCCACATTGCCCCC

CCGACTTAGATGATCACGTCCCCCTGGACAGTGTCCCTAATGCCCAAATATTCATCAAGTACCCAATGGGCGGAATAGAA

GGTTATTGTCAAAAACTATGGACAATCAGTACTATACCATATTTGTATCTGGCAGCCTATGAGAGCGGAGTAAGAATTGC

CTCACTAGTGCAAGGTGATAATCAGACAATTGCAGTGACAAAAAGAGTCCCGAGTTCTTGGCCTTACTCACTAAAAAAGA

GGGAGGCATCCAAAGCAGCTCAGAATTACTTCGTGGTTTTAAGGCAGAGGTTGCACGATGTAGGCCATCACTTAAAGGCT

AATGAAACCATAGTATCTTCTCACTTTTTTGTATACTCGAAAGGGATTTATTATGATGGCCTATTAGTCTCACAATCACT

AAAGAGCATTGCCAGGTGTGTTTTCTGGTCCGAGACTATTGTAGACGAAACCAGAGCGGCCTGTAGCAATATTGCAACAA

CTATTGCCAAGAGTATAGAGAGGGGTTATGATAGGTACCTTGCATACTCTTTGAATATCCTCAAGATTTTCCAACAGATC

CTTATATCCCTCAACTTCACTATTAACACGACAATGACTCAAGATGTTGTGGCACCTATCATCGAGAATGGTGATCTACT

AATAAGGATGGCACTTTTGCCAGCACCCATTGGGGGTCTCAACTATCTTAACATGAGCAGATTATTTGTGAGAAATATCG

GTGACCCGGTCACTTCCTCTATAGCCGACCTGAAGAGGATGATAGACGCTGGGCTAATGCCAGAGGAAACATTGCATCAA

GTGATGACCCAGACCCCTGGAGAATCGTCTTACCTTGATTGGGCAAGTGACCCTTATTCTGCTAACCTGACCTGCGTACA

GAGTATAACTCGCCTTCTCAAGAACATCACTGCTCGATATATTTTAATCAGCAGCCCTAATCCGATGCTAAAAGGATTGT

TCCATGAGGGGAGTAGAGATGAAGACGAAGAGCTTGCGAGTTTCTTGATGGATAGGCATATAATTGTTCCTAGAGCTGCA

CATGAAATCTTAGACCATAGCATAACCGGAGCAAGAGAAGCTATAGCTGGGATGTTGGATACCACCAAGGGTCTGATTAG

AACAAGTATGAAACGAGGTGGCCTCACCCCTCGAGTCTTAGCTCGCCTTTCCAATTATGATTATGAACAATTCAGATCCG

GAATAACATTATTGACCAAGAAAGGGCAGTGTTATCTCATTGACAAGGACTCGTGCTCGGTGCAGCTCGCTATAGCTCTG

AGGGGCCATATGTGGGCTAGGTTAGCTCGCGGGAGACCTATCTATGGATTAGAGGTGCCCGATATACTAGAATCGATGAA

CGGCTACCTTATCAAACGCCACGAGTCTTGTGCCATTTGCGAAACGGGCTCAAATCACTACGGGTGGTTCTTCGTCCCTG

CTGGGTGCCAGCTTGACGATGTCTCTAGAGAGACTTCTGCCCTTCGTGTACCTTATGTCGGATCGACCACTGAGGAAAGG

ACAGATATGAAACTTGCTTTCGTTAGATCTCCAAGTCGATCCCTCAAGTCAGCAGTCAGAATTGCTACAGTTTACTCATG

GGCCTACGGGGATGATGAAAAATCATGGAATGAAGCTTGGATGCTAGCCAGGCAGAGAGCTGATATCACCTTAGATGAAT

TGAGAATGATCACTCCAATCTCTACATCCACCAACCTAGCCCATCGGCTGAGGGATAGGAGTACCCAGGTGAAATATTCG

GGGACATCCCTTGTGAGGGTAGCAAGATACACAACCATTTCCAATGATAATTTGTCATTTGTGATATCTGAGAAGAAAGT

AGATACTAACTTCATTTACCAGCAAGGGATGCTGCTCGGTCTTGGGATCCTTGAGAATCTTTTCAGGTTGGAGGCCACCA

CAGGGGTATCCAACACAGTGTTACATCTGCACGTGGAAACAGAATGCTGTGTTGTACCTATGGTTGATCACCCAAGGATA

CCGAGTCTCCGTAACATTAAAGTTACGAGTGAGCTATGCACAAACCCCTTGATCTACGACAAGTCTCCCATCATAGATCA

CGATGCAACCCGATTATACTCACAGAGCCACAGGAGACATTTGGTGGAGTTTGTGACCTGGTCAACAAGCCAGCTTTACC

ATATACTGGCTAAGTCTACAGCAATGTCCATGATTGAGTTGATCACAAGATTCGAGAAAGATCACATGAATGAAATAGCC

GCCCTGATTGGAGATGACGACATCAACAGTTTCATCACAGAATTTTTGCTAGTAGAGCCCAGATTGTTTATAGTCTACCT

CGGCCAGTGTGCCGCCATCAATTGGGCTTTTGACATACATTATCATCGGCCCTCGGGCAAGTACCAGATGGGGGAGCTCC

TCTACTCTTTACTCTCTAGGATGAGCAAAGGGGTATATAAGATCTTCACCAATGCTTTGAGCCACCCCAAAGTTTACAAG

AAATTTTGGCGAAGCGGTATCATTGAGCCGACTCATGGCCCATCCCTAGATACACAGAACTTACATGTCACTGTCTGTGA

CATGATATATGGATCCTACGTCACCTATCTGGATCTTTTACTGAATGATGAGCTAGACGATTACACGTATTTGCTCTGCG

AGAGTGATGAGGACGTGGTCACAGACAGGTTCGATAACATTCAAGCCAAGCATCTCTGTGTATTGGCCGACTTATATTGC

ACCTCCAAAAGGTGCCCCTCGATAATCGGGATGTCCCCTATAGAAAAATGTACCGTCCTCACACATTACATCAAGGGAGA

ATCAATCCAATCCCCGTCTGGGACCTCATGGAACACTGGTCCCCTTGTAGTAGATCATTACTCGTGCTCTCTGACCTACC

TTCGCCGAGGTTCCATTAAACAAATCAGGTTGAGAGTGGATCCTGGGTTTGTATTCGAGGCGTTGACAGACATCGACATC

AAACAGCCTCGCAAGGCTAAACTGGATGTATCGGTTGTGGGGTTGACTGATTTCTCTCCCCCTTGGGACAACGTCGGTGA

TTTTCTAGGGACTATCAACACATTGAGACACAATCTGCCCGTCACCGGAACCGGGGTCTTGAACTATGAAGTCCATGCTT

ATCGTAGAATTGGCCTGAATTCATCAGCATGTTATAAGGCTGTAGAAATTTCCACGTTAATCAAGTCATCATTAGAAGTC

GGAGAGCATGGATTGTTCTTAGGGGAAGGTTCCGGTTCCATGCTGGCTGCGTACAAGGAAGTTCTCAAATTAGCAAACTG

TTATTACAACAGCGGGGTGACAGCGGAGGACAGAGCCGGACAGAGGGAAATATCCCCCTATCCTTCAGAGATGAGCCTAG

TAGAGAACCAGATGGGGATAGAGAGGAGTGTTAAGGTGCTGTTCAACGGCAAACCCGAAGTAACTTGGGTAGGGACCACC

GATTGCTATAAGTATATAATCAGTAACATTCAAACCTCTAGTTTAGGTCTCATACACTCAGATATTGAGACACTCCCAAA

CAAAGATGCTGTTGAAAAATTAGAAGAATTTGCCTCTATCCTATCCTTATCCCTAATTTTAGGAAAAATCGGCTCTATTA

CGGTTATCAAAATTATGCCCATTAGCGGAGACTTTACTCAAGGCTTCATAGGCTATGCCATTCAATATTTTAGGGAGAGC

CTGCTTGCCTATCCGAGATATAGCAACTTCATCTCCACTGAGTGTTACCTTATTATGATAGGATTGAAAGCCAATCGGTT

GATAAACCCGGAGGCCATTAAGCAAAGCATAGTCAGAGTGGGGACCAGGACTGCACCAGGACTAGTGAGCCACATATTAT

CAGAGAAACAGAAAGGCTGTATTCAATCTTTTCTGGGTGATCCTTATATCCAAGGAGACTTCAACAAACACCTTAAAGCT

CTAACCCCTATTGAGAGAATCCTAGTAAATTGTGGTCTCTCGATCAATGGTACAAAAATCTGTAGGGATCTAATCCACCA

TGATATCGCCTCCGGTCCAGACGGTCTGATGAGCTCCACAATCATTTTATATAGGGAGTTGGCTCATTTCAAAGACAACA

TAAGAAGTCAGCACGGGATGTTCCACCCCTACCCCGTACTGGCCAGTAGCAGGCAACGTGAATTAATCCTCCGAATAGCC

AAGAAATTCTGGGGGTATGTCTTGCTATATTCTGATGACCCGAGACTGATCGGACAAACAATCAAGAACTTAAAGAGGAA

CCACCTAACCTTTGACTTACACAGTAATCCGTTTATTAAGGGCCTATCCAAAGCTGAGAAACTACTGGTGCGGACAAGCT

CACTCAGGAGAGAATGGTTGTTCACTCTCGAGACGAAAGAAGTGAAAGAGTGGTTCAAATTGGTGGGTTACAGTGCACTC

GTCAGAGGTTGATTAGCGATACATCTGCCCCCTTCTCCTCCACCAGGTGACTCTACTGGCAATCTAAAAGATTAAAGAAA

ACTACATATTGGATAAGTATCTATTCCCAGCTTTGTCTGGT

>EU267273.1/goat/CotedIvoire/1989

ACCAAACAAAGTTGGGTAAGGATAGATCTTATAATAACTATGGATCAGGAAACTTAGGAGTAAAGATCCTACTGTCGGGG

GGAGGAGGAGGAGCAAGATCTATCGATATGGCGACTCTCCTTAAAAGCTTGGCATTGTTCAAGAGGAACAAAGATAAAGC

GCCGACAGCATCAGGTTCAGGAGGGGCCATCCGGGGGATTAAGAATGTTATCATAGTCCCAATTCCCGGAGATTCATCTA

TCATTACCCGTTCAAGACTGCTTGACAAGCTTGTCAGATTGGCTGGAGATCCTGACATCAATGGGTCAAAGCTGACCGGC

GTGCTGATCAGCATGTTATCTTTGTTCGTAGAGTCACCCGGGCAATTGATCCAGCGGATCACAGATGACCCAGATGTCAG

TATCCGCCTTGTTGAGGTGGTTCAAAGTACTAGGTCCCAGTCCGGGTTGACCTTTGCATCACGTGGTGCTGATTTGGACA

ATGAGGCAGACATGTATTTTTCAACTGAAGGACCCTCGAGTGGAAGTAAGAAAAGGATCAACTGGTTTGAGAACAGAGAA

ATAATAGACATAGAGGTCCAGGACGCAGAAGAGTTCAATATGTTATTAGCATCCATACTAGCACAAGTTTGGATCCTCCT

GGCCAAGGCAGTTACGGCACCGGATACGGCAGCTGACTCAGAACTGAGAAGGTGGGTTAAATACACACAACAGAGGAGAG

TGATTGGGGAATTCCGCCTTGACAAAGGGTGGCTGGACGCAGTCCGCAATAGGATTGCAGAAGATCTCTCACTTCGGCGG

TTCATGGTATCTCTCATACTTGACATCAAGAGGACCCCCGGGAACAAGCCAAGGATTGCAGAAATGATCTGCGACATCGA

CAACTATATTGTCGAAGCCGGACTTGCCAGTTTCATCCTTACCATCAAGTTTGGTATTGAAACCATGTATCCTGCATTAG

GTCTTCACGAGTTTGCCGGGGAATTGTCCACTATAGAATCCCTGATGAACTTGTATCAACAACTAGGGGAGGTTGCACCT

TACATGGTAATTCTAGAGAACTCAATTCAGAACAAGTTTAGTGCAGGTGCTTATCCCCTCCTGTGGAGCTATGCCATGGG

TGTTGGAGTTGAGCTGGAGAACTCAATGGGGGGCCTGAACTTTGGCAGATCATATTTTGACCCGGCTTATTTTCGTCTTG

GACAGGAGATGGTCAGAAGATCTGCAGGGAAGGTCAGCTCTGTAATCGCGGCTGAGCTTGGCATCACAGCAGAGGAAGCT

AAACTAGTCTCGGAAATCGCCTCGCAGACTGGGGACGAAAGGACCGTTAGGGGGACTGGGCCACGACAGGCGCAGGTTTC

TTTCCTCCAGCACAAAACAGGAGAGGGAGAGTCATCTGCACCAGTGACCAGAGAAGAAGTCAAGGCTGCGATTCCGAATG

GGTCCGAAGAAAGGGACAAAAGGCGAACCCGCCCAGGAAGGTCCAGAGGAGAAATCCCGAGCCAACCGCTCCTGGAAATC

ATGCCAGAGGATGAGGCCTCGAGAGAATCCGGCCAAACCCCTCGTGAGGCTCAGAGGTCGGCTGAGGCACTCTTCAGGCT

GCAAGCTATGGCCAAGATTCTAGAGGGCCAGGAGGGGGGAGAAGACAACAATCAGATCTACAACGACAAAGATCTCCTCA

GTTGAGCAGATGCTCCCTCAATCAACATCATTAATTAAATATCGCCCATCAGTGTTATAAAAAACTTAGGACCCAGGTCC

GAGCAATCGCACATCGAAACTCTAGCCAACCGAGCAGAGACCACCGATGGCAGAGGAACAAGCATACCATGTCAACAAGG

GACTGGAATGTATCAAGTCCCTCAAAACCTCTCCCCCAGATCTATCCATCATCAGAGATACCCTTGAGAGCTGGAGAGAG

GGGCTTGGCCCCTCGGATTGTGCAACGCCGAGCCCTGGTGCGTCCGAGGGAGTCCATCAGGATATCAACCAATCATGCTC

ATCAGCAGTCGGATCAGGCAAAATTGACGTGTCTCCTGAAGGTAATCTCGGATTTAGAGAGATCACTTACGATGACAGTG

AGGCAGGGCCCCGGGGGATTCAGGACAAAGGATCCGACTCTCAAATACAGCGTTACCATGTTTATAGCCACGGGGGTAAA

GAGATTGAAGGACTCGAGGATGCTGACTCTCTCGTGGTTCAAGCAGATCCTCCAGTTGCTAACACCTTCAGTGGAGGAGA

AGATGGATCTGACGACAGCGATGTGGACTCTGGCCCAGATGATCCCAGCAGAGATCCTCTATATGACTGGGGATCTGTTA

CCGGCCATGATGTCGCTAGGTCCACAGATGTCGAAAAATTAGAAGGTGATGACATTCAAGCAGTTCTTAACTCCCAGAAG

AATAAAGGAGGAAGGTTCCAAGGGGGGAAAACCTTGCGAGTCCCGGAAATACCCGATGTCAAGCACTTCGGACCATCAGC

CCAATCAATTAAAAAGGGCACAGACGGGAACTCAGTCTCATCTGGAACGGCGACAGAGTGTTTATTGATAAATGGTGCAA

CCCAAGCTGTGCCAGAGTCAAGATGGGAGTCGTCAGAGCGAAATGCATCTGTGGAGAGTGTCCCCAAATCTGCGAGGAGT

GCAAAGACGATCCAGGAGTTGACACCAGAATCTGGTACCATAGCATCACAGACTCAGCCTAAAGAGAATGACTCCGAGTA

TGAGTATGAGGATGGTCTATTTACAGAGATACAGGACATTCGTGCAAGCATTGCCAAAATCCATGAAGACAACAAAACCA

TCCTCTCAAAGCTTGAATCCATACTGTTATTGAAAGGAGAAATAGATACCATCAAGAAGCAAATCAGCAAGCAGAATATA

AGTATATCCACCATTGAGGGCCATCTCTCCAGTATAATGATAGCAATCCCGGGTTTTGGGAAGGACATCAAGGATCCAAC

ATCCGAGGTTGAGTTGAACCCAGATTTAAGACCTATAATCAGCCGTGACTCCGGCAGAGCTCTCGCGGAGGTACTCAAGA

AACCTGCTGTTGATAGGTCTCAGAAAAATGGAATCAAATCCAACTCAAGTTCAAAGGGTCAGCTCCTTAAGGACCTCCAG

CTAAAACCTGTTGACAAACAAGCAAGCTCTGCAATAGGGTTTATCCCGTCTGACCATGAGTCATCCAGAAGTGTCATCCG

CTCTATAATTAAGTCGAGCAAGCTCAACATTGATCACAAAGACTACCTTCTAGACCTTCTGAATGATGTGAGAGGCTCCA

AGGATCTTAAGGAATTCCACAAAATGCTAACAACCATTCTTGCCAAGCAACCGTAACACAACCTCCAACCAACATCCCAT

ACTCGGCTGAACACATCCCCTCAACCAGGCTATTACAAAAAACTTAGGAGCAAGGGCAACCGAGCTTCACAGACAAGATG

ACCGAGATCTACGACTTCGATAAATCAGCATGGGATGTCAAAGGGTCAATTGCTCCCATCGAACCCACCACTTATCACGA

TGGCCGACTGATACCCCAGGTGAGAGTCATCGACCCTGGGCTAGGAGATAGAAAAGATGAGTGCTTCATGTACCTGTTTC

TTTTAGGAGTGATTGAGGATAACGACCCCCTATCCCCCCCTGTCGGAAGAACGTTTGGTTCTTTACCTCTAGGGGTCGGT

AGGTCAACTGCTAAGCCAGAGGAACTACTAAGGGAAGCCACGGAACTAGACATAGTGGTGAGGCGCACGGCAGGATTAAA

CGAGAAGCTGGTATTTTATAACAACACTCCACTGTCCTTGCTAACACCCTGGAGGAAAGTCTTGACAACCGGAAGTGTGT

TTAGCGCCAACCAGGTTTGCAATGCAATCAATCTGGTCCCACTTGATATTCCCCAGAGATTCAGAGTAGTGTACATGAGC

ATAACTAGATTGTCAGATAATGGTTATTACAGTGTCCCCAGGAGAATGTTGGAGTTCCGCTCAGCCAATGCAGTCGCCTT

CAATATCCTGGTTACACTTAGAATTGAGAATGGCACAAACCCTAGAAGATACATAGTCGGCTCATGGGAGAATCCAGAGG

TCACATTTATGGTACACGTGGGCAACTTTAGACGAAAGAAGAATGAAGTATATTCTGCTGATTACTGCAAAATGAAGATT

GAAAAGATGGGTCTAGTTTTTGCCCTGGGAGGAATAGGAGGAACAAGTCTCCATATTAGAAGCACCGGGAAAATGAGCAA

GACTCTTCATGCACAGCTAGGGTTCAAAAAAATCTTATGCTACCCCTTAATGGACATTAATGAGGATCTTAACCGATATC

TTTGGCGGGCAGAGTGCCGAATAGTCAAAATCCAAGCCGTCTTACAGCCATCAGTACCCCAAGAATTCCGTGTCTACGAT

GATGTCATCATCAACGATGACCAAGGCTTGTTCAAGATCCTGTAAACCGGTCGCCGGACAATAGACCCACCTTCAAGAAG

TGGCTCCCCCTTCCCCCGTCCAAAAGACACACCCCAAGACCCCCCAAAACAGCTCCAACATGGCCATCCTCCCAACGGGA

CGGGGCCGACCTCCCCCACGAACCCGGTCCGGCAGGAGGGGCCCCCCCCGCAACCCACCGGGCGCCGACCGCGTGGGCCA

GCAGGACGCCCCCAAGAGGGCCCGACCGGGACCGGGCACCCT--CCCCCC-AAAAAAACCCCCC-GACACATCCGAGGCC

AGGCGCCAGGCACTCCCAATCCCGCCACCGGGGGGACAAGCAGCCAAGACAGGGCCCCCCCACCCAAACGGACCGCCAGG

GGAGGCCCCACCGACCCAGCACAGACCCGGCCCAAACAAAGGAGACTCCAAAGACGAAACCGCCCAGCGCACCCTACTCA

TCATCAAACAAAACTTAGGGGTCAAGTCCAGACGACAGACCCCAGCCCGCGACCCAGCAGGCCACAGAAGGGAAGCAGGC

ACGACGGACCTGGTAGGCGGACACCCACCCACAGAAGCACCCAGGCCCCGGGGAGAAGCCCCTCCCTCCACACAGCGAGG

GAGCGGGCGAGAAAAACCAGGAGGAACCCGACCGGGAGCCAGGAGCCGGCGACGCCAACGGCGGCCAGACCGGGCACACA

CTCGACACCACCAGCAGAGCAGAGGAGAGCCCAGACCCCCGCGGACTCGACGGTAGGGGAGCCAACCCCGTCACAAGGAG

AGACGAGGGGATC--------CCCCCAAGCACACAACCGGCAGGTAGACCGAGAGAGGACCACTTCCTGGGAGGACAAGC

GGCTCCCCTGCAGCCGAGCCGCAACTCCCCAGGAGAGGACACCCCCGACGCCTCCGAACACCCTCCCGCCCCCGGCCATC

GGGAGAAGGGCTCCGCTGCCAAAGACCAGAGCCCCTGAGAGCCAACCAGAGG-CCCCCCTAGTCTCGCCTGGAACCCCCC

CGAAGAAGCCCCGGCAGGGTCCCCTGGCTCGACGACCCCTGCACGCGCCAAAAGGGAAACCCACCAGAGGGCGATCGCCC

AATCCATTTA-AACCATCATGACAACGATCCCAACCTTGAAATTTTTGTTTTTTTCCCCGATTACTATTGCATGCCAGAT

TCACTGGGGCAATCTATCCAAGATAGGGATTGTTGGAACAGGGAGTGCCAGTTACAGGGTGATGACTAGGCAAGCCCACC

AAACCCTGGTTATAAAGCTAATGCCAAATATAACAGCCATCGACAATTGTACGAAATCAGAAATTTCAGAATACAAAAGA

TTGCTGATCACAGTGTTAAAGCCTGTAGAGGATGCTCTGTCAGTGATAACCAAGAATGTAAAACCAATTCAGACTCTAAC

GCCTGGGCGCAGGACCCGCCGTTTTGCCGGGGCTGTTCTGGCCGGAGTGGCACTTGGAGTCGCGACGGCTGCTCAGATAA

CTGCCGGAGTCGCACTTCATCAATCCTTGATGAACTCCCAAGCGATTGAGAGTTTAAAGACCAGTCTCGAGAAGTCGAAT

CAGGCAATTGAAGAAATAAGGCTTGCAAATAAAGAGACCGTACTGGCAGTGCAGGGTGTCCAAGACTATATCAACAGTGA

ACTTGTCCCTTCTGTCCATAAAATGTCATGTGAGCTTGTAGGTCACAAACTCAGTCTCAAGCTCCTTAGGTATTACACCG

AGATCTTGTCTATATTCGGGCCTAGCCTCCGAGACCCGATAGCTGCTGAAATATCAATCCAGGCACTCAGTTATGCATTA

GGTGGAGATATTAATAAAATCCTGGATAAGCTTGGGTATAGCGGCGAGGATTTCCTTGCTATCCTAGAAAGCAAGGGGAT

AAAGGCTCGAGTTACATATGTGGATACAAGAGATTACTTTATAATTCTAAGTATAGCCTACCCAACTTTATCTGAAATCA

AGGGGGTGATAGTTCATAAAATAGAAGCCATATCATACAACATTGGGGCACAGGAATGGTACACTACTATCCCTAAATAT

GTAGCCACTCAGGGGTATTTGATATCGAATTTTGATGAGACGTCGTGCGTATTCACTCCAGAGGGGACAGTCTGCAGTCA

GAATGCTTTGTACCCAATGAGCCCATTGCTTCAGGAGTGTTTCAGGGGGTCGACAAAGTCGTGCGCCAGAACTCTAGTCT

CAGGGACTACAAGCAACAGATTTATCCTGTCAAAAGGGAACTTGATTGCAAATTGTGCATCAGTTTTGTGCAAGTGTTAC

ACAACGGAGACAGTTATCAGCCAAGATCCTGATAAATTACTAACTGTTATAGCCTCCGATAAGTGGCCTGTAGTTGAGGT

GGATGGGGTGACAATACAGGTCGGCAGTCGTGAGTACCCAGACTCCGTGTACCTACACAAAATAGACCTAGGCCCAGCCA

TCTCTCTGGAAAAACTGGACGTAGGCACCAATTTAGGTAATGCAGTTACAAGACTGGAGAATGCAAAGGAGCTCTTAGAT

GCATCAGACCAGATATTGAAGACTGTTAAAGGAGTACCCTTAGGTGGCAATCTGTACATAGGACTGGCAGCTTGCATCGG

AGTATCCCTAGGGCTTGTCACATTAATATGCTGCTGTAAGGGGAGATGTAGGAACAAAGAGATTCCTACGTCCAAAATCA

ACCCAGGGCTCAAACCTGATCTGACCGGGACCTCAAAGTCATACGTGAGATCACTGTAGTCAGAATAACCCGAATCACTA

AGCATCACACACACACATGTGCAACACAAGCGGTCAGAGGATGCAGAAGATTCAACTTCCGATCACCAACCTGGCCCCGC

TCTATGCTTCACTATACATTGGTCATCAAACAAAACTTAGGACGAAAGGTCGGTCACCATGTCCGCACAAAGGGAAAGAA

TCAATGCCTTCTACAAAGACAATCCTCATAACAAGAATCATAGGGTAATCCTGGATAGAGAACGCTTGACTATTGAAAGA

CCCTACATCTTACTAGGGGTCCTGCTGGTAATGTTTCTGAGTCTAATCGGACTGCTGGCCATTGCAGGGATCAGACTTCA

CCGAGCCACCGTTGGAACTGCAGAGATCCAGAGTCGGCTTAATACCAACATTGAGTTAACTGAATCTGTTGATCACCAAA

CCAAGAATGTCCTAACTCCGCTGTTTAAAGTCATTGGTGATGAAGTCGGCATAAGAATCCCGCAGAAATTCAGTGATCTT

GTCAAGTTTATCTCCGATAAGATTAAGTTCCTCAACCCTGACAGAGAATATGATTTTAGGGATCTCCGGTGGTGTATGAA

TCCCCCCGAGAGAGTCAAAATTAATTTTGATCAGTTTTGTGAATACAAGGCCGCAGATAAGTCAATTGAACATATATTTG

AGTCATCACTCAACAGGTCAAGAAGGTTGCAATTGTTGACCCTTGGGCCTGGAACGAGCTGTCTCGGTAGGGCAGTAACA

AGAGCTCAGTTCTCAGAGCTGACTCTGACCCTGTTGGATCTGGATCTCGAGATGAAAAACAACGTGTCCTCAGTGTTCAC

CGTAGTTGAAGAGGGATTATTCGGAAGAACATACATTGTCTGGAGACCTGATGTCGGGAATCCGAGCATCGACTCAGGTG

CTGGCCATTTTTTAAGAGTCTTCGAGATCGGGCTGGTAAGAGATCTCGGGCTGGGTGCTCCTGTTCTCCATATGACCAAT

CATCTCGCAGTGAACATGAGTGATGGCTATCGGAGCTGTCTTTTAGCAGTAGGGGAGCTGAAGCTGGCAGCCCTATGCAC

CCCATCCGAGACTGTGACACTGAGTGAGAGAGGAGTTCCAAAGAGAGAGCCTCTTGTGGTTGTGATACTCAACCTAGCCG

GGCCTACTCTAGGTGGCGAACTATACAGTGTCTTGCCTACCTCTGACCTCACGGTGGAGAAACTTCATCTATCCTCACAC

AGAGGGATTATCAAAGATAACGAGGCCAATTGGGTAGTACCGTCCACCGATGTTCGTGATCTTCAAAACAAAGGAGAATG

TTTAGTGGAAGCATGTAAGACTCGACCTCCTTCATTTTGCAATGGCACGGGAGTGGGCCCATGGTCAGAGGGGAGAATCC

CTGCCTACGGGGTGATCAGGGTCAGTCTTGACTTAGCCAGTGACCCTGACATAGTTATAACTTCAGTCTTTGGCCCACTG

ATACCTCACCTGTCTGGCATGGATCTTTACAACAACCCGTTCTCAAGCGCTGTGTGGTTGGCGGTACCACCTTATGAGCA

GTCATTTTTGGGGATGATAAATACTATTGGCCTCCCAAATAGAGCAGAGGTTATGCCACACATTTTGACCACAGAGATCA

AGGGGCCTCGGGGTCGTTGTCATGTTCCCATAGAATTGTCCCGCAGGATTGATGATGACATCAAGATTGGGTCCAACATG

GTTGTATTGCCGACGAAGGACCTGAAGTACATAACAGCCACTTATGATGTTTCCAGGAGCGAGCATGCAATCGTGTACTA

TATCTATGATACAGGCCGCTCATCATCTTACTTCTACCCGGCTCGATTGAATTTCAAAGGCAATCCTCTCTCTCTGAGGA

TAGAGTGTTTCCCCTGGCATCATAAGGTATGGTGCTACCATGATTGTCTTATATACAACACCATAACAAACGAAGAAGTC

CATAAGAGAGGATTGATCGGCATAGAGGTAACATGTAATTCAGCCTGAGCGGAGCTGCGGCCGCCACTCAAGCAGGCCTC

CGAGCTAATCCCAAGCCCAGGCAAGATAGCTCGGGACACTCGACAACACAACCCAGTCAACAATGTTATAAAAAACTTAG

GGGCCAAGGCTATAGGAGCCATGGACTCCCTGTCAGTCAATCAGGTCTTGTACCCTGAGGTCCATCTGGATAGCCCTATT

GTCACAAACAAGCTGGTTGCCATCCTCGAGTACTCAGGCATTGACCACAACTATGTCCTTGAAGACCAGACCCTCATCAA

GAATATTAGATATAGACTGGGGTGTGGTTTCTCAAATCAAATGATCATCAACAATAGGGGGGTCGGGGAAACAGTCAACT

CTAAACTTAAAAGTTACCCCCGTAATTGTCATATCATATACCCAGATTGCAATAAGGATCTGTTTTGTATCAAAGACAGC

TGCATATCTAGGAAGCTCTCTGAGCTCTTCAAGAAGGGTAATTCCCTATACTCGAAGGTAAGTCACCAGGTACTGGATTG

TCTTACAAGAGTCAACGGTAAATTAGGCTTGGGTACAGATCTTACTCATGGCCTGAAGGAGGGTATCCTTGACTTGGGGT

TACACATGCACAGCTCTCAATGGTTCGAGACTTTCCTGTTCTGGTTCACTATCAAGACAGAGATGAGATCAATGATAAAA

GAACAGTCCCATATATGCCACAAGAGGAGATACAACCCGGTTTTTGTGTCGGGGGACGCATTCGAGGTGCTCGTATCACG

AGACCTCGTTGCGATAATTGGCAAGAATACCCAGTATGTCTACTATTTGACATTTGAGCTGGTCCTTATGTATTGTGATG

TCATAGAGGGGAGACTTATGACGGAGACAGCCATGGCTATAGACCAGAGATATTCAGAACTTCTAAGCCGGGTCAGATAT

TTGTGGGATCTTATTGACGGGTTCTTCCCAACACTAGGTAACACCACATACCAGGTTGTTGCTCTGCTTGAACCGCTGTC

GTTGGCTTATCTCCAACTTCAGGATGTCACCCTGGAGTTAAGAGGTGCTTTTTTAGATCACTGCTTCAAGGAACTCTATG

AGATACTAGAGCATTGCGGCATTGACACGGAAGGCACCTACAATTCCATCACTGAAGGATTGGACTACGTATTTATCACC

CACGATATACATTTAACTGGGGAGATATTTTCATTTTTCCGGAGTTTCGGACACCCCCGCCTCGAAGCGGTCACCGCCGC

AGAGAATGTCAGAAAACATATGAACCAGCCGAAGGTAATCAGTTATGAGACTATGATGAAAGGGCATGCAGTGTTTTGCG

GGATAATCATAAATGGGTTTAGAGACCGGCATGGCGGCAGCTGGCCCCCCGTTGCATTGCCAGAACATGCCTCTGCTGCA

ATTCGGAATGCTCAGGCATCCGGTGAGGGGCTGACCCATGACTTGTGTATAGACAACTGGAAGTCCTTTGTCGGATTCAG

ATTTGGCTGTTTCATGCCGCTCAGCCTAGATAGTGATTTGACCATGTACCTTAAAGACAAAGCACTGGCTGCACTGAAGA

ATGAGTGGGATTCAGTTTACCCGAAAGAATACCTCCGCTATAATCCACCCAGGGGGACAGAGTCAAGGCGGCTGGTGGAG

GTGTTCCTGAATGACTCCAGCTTTGATCCTTATAACATGATAATGTACGTGGTGAATGGCTCTTACCTTAAAGACCCTGA

GTTTAATCTCTCATATAGCTTAAAGGAGAAGGAGATAAAAGAGACAGGGCGGTTGTTTGCAAAAATGACTTACAAGATGC

GGGCCTGTCAGGTAATTGCGGAGAATCTGATATCAAATGGAGTCGGGAAGTATTTCCGAGACAATGGAATGGCGAAAGAC

GAGCATGACCTAACAAAAGCCCTCCACACTCTGGCAGTCTCAGGTGTTCCCAAGAACAATAAAGATAGCCACCGAGGTGG

CCCTCCTCGAAGGACCACAAGCCGAGAGATGAGATCAAACCAGGACATCAACAGACAAAATAGAGACAAGGTCCAAGGAG

AGGCTATGTACAACTATTTGCGGTGCCAACCAATCAGCCCTGACCAGAGTGAGTCATACGAGACTGTTAGTGCATTCATC

ACCGCTGACCTTAAGAAGTATTGCCTGAATTGGAGATACGAGACAATCAGCATATTTGCACAGAGACTTAATGAAATTTA

CGGATTGCCATCCTTCTTTCAATGGTTACACAGGGTATTGGAAAAATCTGTGCTCTACGTCAGTGACCCACATTGCCCCC

CCGACTTGGATGATCACGTCCCCCTGGACAATGTCCCTAATGCCCAAATATTCATCAAGTACCCAATGGGCGGAATAGAA

GGTTATTGTCAGAAACTATGGACAATCAGTACTATACCATATTTGTACCTGGCAGCCTATGAGAGCGGAGTAAGAATTGC

CTCACTAGTGCAAGGTGATAATCAGACAATTGCAGTGACAAAAAGAGTCCCGAGTTCTTGGCCTTACTCACTAAAAAAGA

GGGAGGCATCCAAAGCAGCTCAGAATTACTTCGTGGTTTTAAGGCAGAGGTTGCACGATGTGGGCCATCATTTAAAAGCT

AATGAAACCATAGTATCCTCTCACTTTTTTGTATACTCGAAAGGGATTTATTATGATGGCTTATTAGTCTCACAATCACT

AAAGAGCATTGCCAGGTGTGTCTTCTGGTCCGAGACTATTGTGGACGAAACCAGAGCGGCCTGTAGCAATATTGCAACAA

CTATCGCCAAGAGTATAGAGAGGGGTTATGATAGGTACCTTGCATACTCTTTGAATATCCTCAAGATTTTCCAACAGATC

CTTATATCCCTCAACTTCACCATTAACACGACGATGACCCAAGATGTTGTGGCACCTATCATTGAGAATGGCGATCTACT

AATAAGGATGGCACTTTTGCCAGCACCCATTGGGGGTCTCAACTATCTTAACATGAGCAGATTATTTGTGAGAAATATCG

GTGACCCGGTCACTTCCTCTATAGCCGACCTGAAGAGGATGATAGACGCTGGGCTAATGCCAGAGGAAACATTGCATCAA

GTGATGACCCAGACCCCTGGAGAATCGTCTTACCTTGATTGGGCAAGTGACCCTTATTCTGCCAACCTGACCTGCGTACA

GAGTATAACTCGCCTTCTCAAAAACATCACTGCTCGGTATATTTTAATCAGCAGCCCTAATCCGATGCTAAAAGGATTGT

TCCATGAGGGGAGCAGAGATGAAGACGAAGAGCTTGCGAGCTTCTTGATGGATAGGCATATAATTGTTCCTAGAGCTGCA

CATGAAATCTTAGACCATAGCATAACCGGAGCAAGAGAAGCTATAGCTGGGATGTTGGATACCACCAAGGGTCTGATTAG

AACAAGTATGAAACGAGGTGGCCTCACCCCTCGAGTCCTAGCTCGCCTTTCCAATTATGATTATGAACAATTCAGATCCG

GAATAACATTACTGACCCAAAAAGGGAAGTGTTACCTCATTGACAAGGACTCGTGCTCGGTGCAGCTCGCTATAGCTCTG

AGGGGCCATATGTGGGCTAGGTTAGCTCGTGGGAGACCTATCTATGGATTAGAGGTGCCCGATATACTAGAATCGATGAA

CGGCTACCTTATCAAACGCCACGAGTCTTGTGCCATTTGCGAAACGGGCTCAAATCACTACGGGTGGTTCTTCGTCCCTG

CTGGGTGCCAGCTTGACGATGTCTCTAGAGAGACTTCTGCCCTCCGTGTACCTTATGTCGGATCGACAACTGAGGAAAGG

ACAGATATGAAACTTGCTTTCGTTAGATCTCCAAGTCGATCCCTCAAGTCAGCAGTCAGAATTGCTACAGTTTACTCATG

GGCCTACGGGGATGATGAAAAATCATGGAATGAAGCTTGGATGCTAGCCAGGCAGAGAGCTGATATCACCTTAGATGAAT

TGAGAATGATCACTCCAGTCTCTACATCCACCAACCTAGCCCATCGGCTGAGGGACAGGAGTACCCAGGTGAAATATTCA

GGGACATCCCTCGTGAGGGTAGCGAGATACACAACCATTTCCAATGATAATTTGTCATTTGTAATATCTGAGAAGAAAGT

AGATACTAACTTCATTTACCAGCAAGGGATGTTGCTCGGTCTTGGGATCCTTGAAAATCTTTTCAGATTGGAGGCCACCA

CAGGGGTATCCAACACAGTGTTACATTTGCACGTGGAAACAGAATGCTGTGTTGTACCTATGGTTGATCACCCAAGGATA

CCGAGTCTCCGTGACATTAAAGTTACGAGTGAGCTATGCACAAACCCCTTGATCTACGACAAGTCTCCCATCATAGATCA

CGATGCAACCCGATTATACTCACAGAGCCACAGGAGACATTTGGTGGAGTTTGTTACCTGGTCAACAAGCCAGCTTTACC

ATATACTGGCTAAGTCTACAGCAATGTCCATGATTGAGTTGATCACAAGATTCGAGAAAGATCACATGAATGAAATAGCC

GCCCTGATTGGAGATGACGACATCAATAGTTTCATCACAGAATTTTTGCTAGTAGAGCCCAGACTGTTTATAGTCTACCT

CGGCCAGTGTGCCGCCATCAATTGGGCTTTTGACATACATTATCATCGGCCCTCGGGCAAGTACCAGATGGGGGAGCTCC

TCTACTCTTTACTCTCTAGGATGAGCAAAGGGGTATATAAGATCTTCACCAATGCTTTGAGCCACCCCAAAGTTTACAAG

AAATTCTGGCGAAGCGGTATCATTGAGCCGACTCATGGCCCGTCCCTAGATACACAGAACTTACATGTCACTGTCTGTGA

CATGATATATGGATCCTACGTCACTTACCTGGATCTTTTACTGAATGATGAGCTAGACGATTACACGTATTTGCTCTGTG

AAAGTGATGAGGACGTGGTCACAGACAGGTTCGATAACATTCAAGCCAAGCATCTCTGTGTATTGGCCGATTTATATTGC

ACCTCCAAAAGATGCCCCTCGATAATCGGGATGTCCCCTATAGAAAAATGTACTGTCCTCACACATTACATCAAGGGAGA

ATCAATCCAATCCCCGTCTGGGACCTCATGGAACACTGGTCCCCTCGTAGTAGATCATTACTCGTGCTCTCTGACCTACC

TTCGCCGAGGTTCCATTAAACAAATCAGGTTGAGAGTGGATCCCGGGTTTGTATTCGAGGCGTTGACAGACATCGACATC

AAACAGCCTTGCAAGGCTAAACTGGATGTATCTGTTGTGGGGTTGACTGACTTCTCTCCCCCTTGGGACAACGTCGGTGA

CTTTCTAGGGACTATCAACACATTGAGACACAATCTGCCCGTCACCGGAACCGGGGTCTCGAACTATGAAGTCCATGCTT

ATCGTAGGATTGGCCTGAATTCATCAGCATGTTATAAGGCTGTTGAAATCTCCACGTTAATCAAGTCATCATTAGAAGTC

GGAGAGCATGGATTGTTTTTAGGGGAAGGTTCCGGTTCTATGCTGGCTGCATACAAGGAAGTTCTCAAATTAGCAAACTG

TTATTACAACAGCGGGGTGACAGCGGAGGACAGAGCTGGACAGAGGGAAATATCCCCCTATCCTTCAGAGATGAGCCTAG

TAGAGAACCAGATGGGGATAGAGAGGAGTGTTAAGGTGCTGTTCAACGGCAAACCCGAAGTAACTTGGGTGGGGACCACC

GATTGCTATAAGTATATAATCAGTAACATTCAAACCTCTAGTTTAGGTCTCATACACTCAGATATTGAGACACTCCCAAA

CAAAGATGCTGTTGAAAAATTAGAAGAATTTGCCTCTATTCTATCCTTATCCCTAATTTTAGGGAAAATCGGCTCTATTA

CAGTTATCAAAATTATGCCCATTAGTGGGGACTTTACTCAAGGCTTCATAGGATATGCCATTCAATATTTTAGAGAGAGC

CTGCTTGCCTATCCGAGATATAGCAACTTCATCTCCACTGAGTGTTACCTTATTATGATAGGATTGAAAGCCAATCGGTT

GATAAACCCGGAGGCAATTAAGCAAAGCATAATTAGAGTGGGGACTAGGACTGCACCAGGACTAGTTAGCCACATATTAT

CAGAGAAACAGAAAGGTTGTATTCAATCTTTTCTGGGTGATCCTTATATCCAGGGAGACTTCAATAAACACCTTAAAGCT

CTAACACCTATTGAGAGAATCCTAGTAAATTGTGGTCTCTCGATCAATGGTACAAAAATCTGTAAGGATCTAATCCACCA

TGATGTCGCCTCCGGTCCAGACGGTCTGATGAGCTCCACAATCATTTTATATAGGGAGTTGGCTCATTTCAAAGACAATA

TAAGAAGTCAGCACGGGATGTTCCACCCCTACCCAGTGCTGGCCAGTAGCAGGCAACGTGAATTAATCCTCCGAATAGCC

AAGAAATTCTGGGGGTATGTCTTGCTGTACTCTGATGACCCGAGACTGATCGGACAAGCAATCAAGAACTTAAAGAGGAA

CCATCTGACTTTTGACTTACACAGTAACCCGTTTATTAAGGGCCTATCCAAAGCTGAGAAACTGCTGGTGCGAACAAGCT

CACTCAGGAGAGAATGGTTGTTCACTATCGAGACGAAAGAAGTGAAAGAGTGGTTCAAATTGGTGGGTTACAGTGCACTC

GTCAGAGGTTGATTAGCGAGCGCGCTGCCCCCTTCTCCTCCACCAGGTGACTCTACTGGCAATCTAAAAAATTAAAGAAA

ACTACATATCGGATAAGTATCTATTCCCAGCTTTGTCTGGT

>KM463083.1/goat/Kenya/2011-05

ACCAAACAAAGTTGGGTAAGGATAGATCTTATAATAACTATGAATTGCCACGCTTAGGATTAAAGATCCTACTGCCGGGG

GGAGGAGTAGGAGCAAGATCCCTGACTATGGCGACTCTCCTCAAGAGCTTGGCATTGTTCAAGAAGAACAGGGACAAAGC

GCCGACGGCATCAGGTTCAGGAGGAGCCATCCGGGGGATTAAGAATGTTATCATAGTCCCGATTCCCGGAGACTCATCCA

TCATTACCCGTTCAAGATTGCTCGACAGGCTTGTCAGATTGGCCGGAGATCCAGATATCAACGGGTCAAAGCTGACCGGC

GTGATGATCAGCATGCTGTCTTTGTTCGTAGAGTCACCCGGGCAACTGATCCAGCGGATCACAGATGATCCAGATGTCAG

TATCCGCCTTGTTGAGGTGGTCCAAAGTACTAGATCCCAGTCCGGATTGACCTTTGCATCACGTGGTGCTGATTTGGACA

ATGAGGCAGACATGTATTTTGCAACTGAAGGGCCCTCGAGTGGGAGCAGGAAAAGGATCAACTGGTTTGAGAACAGGGAA

ATAATAGATATAGAAGTACAGGATGCAGAAGAGTTCAATATGTTATTAGCCTCCATACTGGCACAAGTTTGGATTCTCCT

GGCCAAGGCAGTTACAGCACCAGACACTGCAGCTGACTCAGAATTGAGAAGGTGGGTTAAGTACACACAACAAAGAAGGG

TGATTGGGGAATTCCGCTTTGACAAAGGGTGGCTAGATGCGGTCCGGAACAGGATTGCAGAAGATCTATCACTCCGGCGG

TTCATGGTATCTCTTATACTTGACATCAAAAGGACCCCTGGGAACAAGCCAAGGATTGCAGAAATGATCTGTGACATTGA

CAACTATATTGTCGAAGCCGGGCTCGCCAGTTTCATCCTTACTATCAAGTTTGGTATTGAAACCATGTATCCTGCGCTAG

GTCTTCACGAGTTTGCCGGAGAGTTGTCCACTATAGAATCCTTGATGAATCTATATCAACAGCTAGGCGAAGTTGCACCC

TACATGGTGATTCTGGAGAACTCAGTTCAGAACAAGTTTAGTGCAGGAGCCTATCCACTTCTCTGGAGCTATGCAATGGG

TGTTGGAGTCGAGCTGGAGAACTCAATGGGAGGGCTGAACTTTGGTAGATCATATTTTGACCCAGCTTATTTTCGTCTCG

GACAGGAGATGGTCAGAAGATCCGCAGGAAAGGTCAGCTCTGCAATCGCAGCTGAGCTCGGTATCACAGCAGAGGAAGCT

AAACTAGTCTCGGAAATTGCCTCTCAGACTGGGGACGAAAGGACCGCTAGAGGGACCGGGCCCAGACAGGCGCAGGTTTC

CTTCCTCCAGCATAAAATAGGAGAGGGAGAGTCACATGCATCGGCGACCGGGGAAGAAGTCAAAGCTGCGACCCCAAATG

GGCCCGACGAAAAGGACAAAACTCGGGCGCGCTCAGGAAAGCCAAGAGGAGGAACCCCTGACCAACTGCTCCTGGAAATC

ATGCCTGAAGACGAGGTCCCGCGAGGGTCTGGACAAAACCCTCGTGAGGCTCAAAGATCGGCCGAGGCACTCTTTAGACT

GCAGGCCATGGCCAAGATCCTAGAGGGCCAAGAGGAGGGGGAAGACAACAGCCAGACATACAACGACAAGGACCTCCTTA

GCTGAGAGGAAGAGCCCTCCATCAAAACCAGCAACAAGACACCGCCCTTCAGTATTATAAAAAACTTAGGACCCAGGTCC

AAGCAGTCACACTCCGGCACCCCAGCCGACCGAGCGGAGACCACCGATGGCAGAGGAACAAGCATACCATGTCAACAAGG

GACTGGAATGTATCAAGTCCCTCAAAGCCTCTCCCCCGGATCTCTCCACCATCAGAGATACCCTTGAAAGCTGGAGGGAG

GGGCTTGACACATCGGACCGTGCAGCACCGAACCCTGATATATCCAAGGGGGACCATCAGAATATCAACCAATCATGCCC

ACCAGCAATCGGATCAGGCGAAATCGACGTGTCTACTGAAAGTAATCTCGGATATAGAGAGATCAATCACGATGACAGTG

AGGCTGGACTCAGAGGAGTTCAAAGCAGAGGACCCAACCCTCAAATACAGCGTTACCATGTTTATAGCCACGGGGGTGAA

GAGATTGAAGGACTCGAGGATGCTGACTCTCTCATGGTTCAAGCAGATCCTCCACTTGCTAACACCTTCAGTAGAGGAGA

GGGTGGATCTGACGACAGCGATGTGGACTCTGGCCCAGATGATCCCGACAGAGATCCTCTATATGACCGGGGATCTGTTG

CCGACAATGATGTCGTTAAGTCCACAGATGTCGAGAAATTAGAAGGTGACAACATTCAAGAAGTTCTTAACTCTCAGAAG

AGCAAGAGGGGAAAATTCCAAGGCGGGAAAACCTTGCGAGTCCCTGAAACGCCCGACGTCAAGCACCCCAGACCATCAGC

CCAATCAATTAAAAAGGGCACAGACGGGAACTCAGTCTTATCTGGAACGGTGACAGAGTGTTCATCGATAAATGGTGCAA

CCCAAGCTGTGCCAGAGTCAAGATGGGAGTCATCAGAGCGAAATGCATTTGTGGAGAGTGCCCCCAAATCTGCGAGGAGT

GCAAAGACGATCCAGGAGTTGCCACAAGAATCTGGTACCATAGCATCACCGACTCAGCCTAAAGAAAATGACTCCGAGTA

TGAGTATGAGGATGACCTATTTATAGAGATGCAGGACATTCGTGCAAGCATTGCCAAGATCCATGATGACAACAAAACCA

TCCTCTCAAAGCTTGACTCTATACTATTATTGAAAGGAGAAATCGATACTATCAAGAAACAAATCAGCAAACAGAATATA

AGTATATCTACCATTGAGGGCCATCTTTCCAGTATAATGATAGCCATCCCGGGTTTCGGGAAGGAGGTCAAGGACCCAAC

ATCCGAGGTTGAGTTAAACCCGGATTTAAGACCTATAATCAGCCGTGATTCTGGCAGGGCTCTCGCGGAGGTCCTCAAGA

AACCTGCTGTTGATAGGTCTCCTAAAACCGGATTCAAAGTCAACTCCGGTTCAAAGGGTCAGCTCCTCAAGGATCTCCAA

CTGAAACCCGTTGACAAACAGGCAAGTTCTGCCATAGGGTTTGTCCCGTCTGACCATGAATCATCCAGAAGTGTCATCCG

CTCTATAATCAAGTCAAGCAAGCTCAACATTGATCACAAAGACTATCTTTTAGATTTACTGAATGATGTGAAAGGCTCCA

AGGATCTCAAAGAATTCCACAAGATGCTAACAGCTATTCTTGCTAAGCAACCGTAATGCACCGCCTAACCAATTTCTCAT

GCTCAGCTGAAAACAGCCCCACAACCAGGCTATTACAAAAAACTTAGGAGCAAGGGCAACCGAGCTTCGCAGACAAGATG

ACCGAGATCTATGACTTCGACAAATCAGCATGGGACGTCAAAGGATCAATAGCTCCCATAGAGCCCACCACTTATCACGA

TGGTCGACTGATACCCCAGGTGAGAGTCATCGATCCTGGGCTAGGAGACAGGAAAGATGAATGCTTCATGTACTTGTTTC

TCCTAGGGGTGATTGAGGATAATGACCCCCTGTCTCCCCCAGTAGGAAGAACCTTTGGTTCCTTACCTCTAGGTGTCGGT

AGGTCAACTGCTAAGCCAGAAGAACTATTAAGGGAGGCCACTGAACTTGACATAGTGGTGAGACGCACAGCCGGACTAAA

CGAGAAGCTGGTATTTTACAACAACACTCCGCTGTCTTTGTTAACACCTTGGAGGAGAGTCCTGACAACCGGAAGTGTGT

TCAGCGCTAACCAGGTTTGTAACGCAGTCAATCTAATCCCACTCGATACTCCCCAGAGATTCAGGGTTGTGTACATGACC

ATCACTAGATTGTCTGACAATGGTTATTATAGTGTCCCCAAGAGAATGTTAGAGTTCCGCTCAGCCAATGCCGTCGCCTT

CAATATCCTGGTGACACTGAGAATTGAGAATGGGACAAACCCTAGACGATACATAGTCGGCTCATGGGAGAACCCAGAGG

TCACATTCATGATACATGTGGGCAACTTTAGAAGAAAGAAGAATGACGTATATTCCGCTGATTACTGCAAAATGAAGATT

GAGAAGATGGGTCTGGTTTTTGCTTTGGGAGGAATAGGGGGAACAAGTCTTCACATTAGAAGTACCGGGAAAATGAGCAA

GACCCTCCATGCACAGCTGGGGTTCAAGAAAATCTTATGCTACCCCTTAATGGATATTAATGAGGATCTCAATCGATATC

TTTGGCGGGCAGAGTGCCGAATAGTCAAAATTCAAGCTGTCTTACAGCCTTCAGTACCCCAAGAATTCCGTGTCTACGAT

GATGTCATCATTAATGATGACCAAGGCTTGTTCAAGATCCTGTAACTCACTCACAGAACAATGGACCCGCCTCCAACAAA

GGACCACCCCCTCCCTCTCCCAGAAAACACACCCCGAGCCCCCCCCAAAAAACGCCGACATGGTCATCCTCCCGGTAGGA

CAAGGATGACCCCCTCATCGGACCCGAGTCTGC-GGGGGGGCCCCCCCCGTGGCCCGCCGAGCACCGACCGCGCAGGGCA

CCGAGGCCCTACTGGGAGGCCCCACGGCAGACCAGGCACCCTCCTTCCTCCGCGTCGCGCCCCA-AACATGCCCAAGGCC

AGGAGCCGAGCACCCCCAGCCAAGCCAACGACAGGACAGACACCCAAGTAGGGGTCCTC--GACCGAAGAGAATACCAGG

GTGGGCTCCGCCGACCCAGCACAGATTTGGCCCTAACAAGGCAGATCCCCAAAACGGAATCTCCCAGTGCACTCCACTCA

TCATCAAACAAAACTTAGGGGCCAAGGTCCGACGACTGACCCCACTCCGCCAAGCAGCAGGCCGCACGAAGGAAGCAGAC

AAGACAGGCCCGGAAGGCAGACACCCGACCACAGAGACATCCAGGCCTCGGGGAGGAGGCCTCCCCCCT-CAAGGCGGAG

GAGCGGGAGGGAAACACCAGGGACAACCCGACCGGGAGCCGGGACCCGGAGGCTCCAACGACAACCGGGCCCGGCACACA

CCCGGCACCCCCGGCGAGACAGAGGAGAGCCCGAGCCCGCTTGGACCCGACGGCAGAGGGTCCAATCCTGCCACAAAGAG

AAGCGAGGAGATC--------TCCCGCCGCACACAGTCGGCCGACAACCCACAGGGGGACCACCTCCCGGGAGGATAAGC

GGCCGTCCCACAACTGAGCCGCCACCCCCCAGAGGAGGACGCCCCGGAGGTCTCAAGATATCCCCCCGTCCTCGACCACC

GGGGGAGGGGCCACACCGACAAAGCCCAGAGCCACCGGGAGCCGACCAAAGGTCCCCCCCGGCCCCGCCCGGCACCCCCC

CGACGAGACCCTAACAAGACCCCTCGACCCAACAACCCCCACACGCGCCAAGACAGAAACCTGCCAGTGGACATCCGGCA

CATTCATGCACAAACATCATGACAAAAGTCGCAATCTTGACATCCCTGTTTCTCCTCACCAGCACTGTCGTATCCCAGAT

TCACTGGGGCAACCTATCCAAGATCGGGATTGTGGGGACAGGAAGTGCCAGTTATAAGGTGATGACTAGGCCGAGCCAAC

AGACCCTGGTTATAAAGCTAATGCCAAATATAACAGCCATCGACAATTGTACGAAATCAGAGATTTCGGAATACAAAAGA

CTGCTGATCACGGTGTTAAAACCTGTCGAGGATGCTCTGTCAGTGATAACCAAGAATGTAAGACCAATTCAAGCTCTAAC

ACCCGGTCGCAGGACTCGCCGTTTTGCCGGGGCTGTCCTGGCCGGAGTGGCACTTGGGGTCGCAACGGCTGCTCAGATAA

CTGCCGGAGTCGCACTCCATCAATCTTTGATGAACTCCCAAGCAATTGAAAGTCTAAAAGCCAGTCTTGAGAAGTCGAAT

CAAGCAATAGAAGAAATCAGACTTGCAAATAAGGAAACCATACTGGCAGTACAGGGTGTCCAAGATTACATCAACAACGA

GCTAATCCCTTCTGTTCATAAGATGTCATGTGAGTTGGTAGGTCACAAACTCAGCCTCAAGCTCCTTAGGTACTACACCG

AAATCTTGTCTGTATTCGGGCCCAGCCTTCGAGACCCGATAGCTGCTGAAATATCAATCCAGGCACTCAGTTATGCACTA

GGTGGGGATATCAATAAGATCCTGGATAAGCTTGGATATAGCGGCGAGGACTTCCTTGCTATCTTGGAAAGCAAGGGGAT

AAAGGCCCGAGTCACCTATGTGGATACAAGAGATTACTTTATAATTCTTAGCATAGCCTACCCGACCTTATCTGAGATCA

AAGGGGTGATAGTTCATAAAATGGAAGCTGTGTCATACAACATCGGGGCACAAGAATGGTACACTACTATCCCTAAATAT

GTAGCCACTCAGGGATACTTGATATCAAATTTTGACGAGACGTCATGCGTGTTCACTCCAGGAGGGACAGTCTGCAGTCA

GAATGCTTTATACCCAATGAGCCCGTTGCTTCAGGAATGTTTCAGGGGGTCTACAAAATCGTGCGCCAGAACCCTAGTTT

CAGGGACTATAAGTAATCGATTTATCCTGTCAAAAGGGAACCTGATTGCCAATTGTGCATCAGTCCTGTGCAAATGTTAC

ACAACGGAGACGATTATCAGCCAAGATCCTGATAAATTACTAACTGTTATAGCCTCTGATAAATGTCCTGTAGTTGAAGT

GGATGGAGTGACGATACAGGTCGGCAGTAGGGAGTACCCAGATTCTGTGTACCTACATAAAATAGACTTGGGCCCAGCCA

TCTCTCTAGAAAAACTGGATGTAGGCACCAATTTAGGCAATGCAGTTACTAGGCTGGAGAATGCAAAGGAACTCCTAGAT

GCATCAGATCAGATATTGAAGACTGTTAAGGGGGTACCCTTCAGCGGCAATTTGTACATAGGACTGGCAGCTTGCATCGG

GGTATCTCTAGGGCTTGTCACATTAATATGCTGCTGTAGGGGGCGGTGTAGGAACAAGGAGATTCCTACCCCTAAGATCA

ACCCAGGGCTTAAACCAGACCTGACCGGGACCTCAAAGTCGTACGTAAGATCACTGTAATCAGAATAGCCCGAATCATCC

AGCGTCACACACAGACATGTGCAACACAAGCAGTCAGGGGACTCAGAGGGTTCAACTTCCGATCCTCAACCTGATCCCAT

TCCAAGCCCCATTGCACATCGGTTATTAAATAAAACTTAGGACGAAAGGTTAGTCACCATGTCCGCACAAAGAGAGAGGA

TCAATGCCTTCTACAAAGACACTCCTCACAACAAGAACCATAGGGTGACCCTGGATAAGGAGCGTCTGACTATTGAAAGA

CCCTATATCCTGCTTGGGGTCCTACTGGTAATGTTTCTGAGTCTGATTGGGTTGCTGGCCATTGCAGGGATTAGACTTCA

CCGAGCCACCGTCGGAACTGCAGAGATCCAGGGCCGGCTTAATACCAACATTGAGTTGACTGAATCTATTGATCATCAAA

CTAAGGATGTCCTAACTCCCCTATTTAAAATCATCGGTGATGAAGTCGGCATAAGGATTCCACAGAGATTCAGTGACCTT

GTCAAGTTCATCTCCGATAAGATTAAATTCTTCAACCCTGACAGAGAATATGATTTCAGGGACCTCCGGTGGTGTATGAA

TCCTCCAGAGAGAGTTAAAATAGATTTTGATCAGTTTTGTGAATACAAAGCCGCGGATAAGTCGATTGAACATATATTTG

AGTTACCACTCAACAGGTCAAAAGAGTTGCGATCGTTGACTCTTGGGCCTGGAACGGGCTGTCCCGGCAGGGCAGTAACA

AGGGCCCAATTCTCAGAGCTTACTCTGACTCTGTTGGATCTGGATCTCAATATGAAGCACAACGTGTCCTCAGTGTTTAC

CATAGTTGGAGAGGGATTATTCGGAAGAACATATACCGTCTGGAGATCATATGCCGGAAATCTGAGTACCGCCTCAGGTC

TCGGCCATTTTTTAAGAGTCTTCGAGATCGGACTGGTAAGAGATTTTGGGCTAAGTGCCCCTGTTTTCCATATGACTAAC

TATCTCACAGTGAACATGAGTGATGACTACCGGAGCTGTCTATTAGCAGTGGGGGAATTGAAGCTGACAGCTCTATGCAC

CCATTCTGAGACTGTGACTCTGAGTGAGAGAGGAGTTCCTAAGAGAGAGCCTCTTGCGGTTGTAATACTTAATCTAGCCG

GGCCTACTCTAGGGGGCGAACTATACAGTATTTTGCCTTCCTCTGACCTCATGGTAGAAAAACTCTATTTCTCATCAAAT

AGGGGGATTATCAAAGACAACGAGGCCAATTGGGTAGTACCATCTACCGATGTACGTGATCTTCAAAATAAAGGAGAATG

CCTGGTGGAGGCATGCAAAACTCGACCTCCTTCATTTTGCAATGGTACAGGGTTAGGCCCATGGTCAGAGGGGAGAATCC

CTGCCTACGGGGTGATCAGAGTCAGTCTTGACGTAGCTAGTGATCCAGATGTGGTTATTACCTCAGTGTTTGGCCCCTTG

ATACCTCATCTATCTGGCATGGATCTTTACAACAACCCATTCTCCAGAGCTGTGTGGTTGGCTGTACCACCTTATAAGCA

GTCATTTCTGGGAATGATAAATACAATTGCCTTCCCAAGTAGAGCAGAGGTTATGCCGCATATTTTGACCACAGAGATCA

GAGGGCTTCGGGGTCGTTGTCATGTTCCCATAGAATTGTCCCGCAAGACTGATGATGATATTAAGATTGGCTCCAACATG

GTTGTATTACCGACGAAGGACCTGAGGTACATTACAGCCACTTACGATGTTTCCAGGAGCGATCATGCAATCGTGTATTA

TATCTATGACACAGGTCGCTCGTCATCTTACTTCTTCCCCGCTCGATTAAAGCTCAAAGGTAACCCTCTCTCTATGAGGA

TAGAGTGTTTCCCCTGGCATCATAAGGTATGGTGCTACCATGACTGTCTCATATACAACACCACCACAAACGAAGAAGTT

CATACGAGAGGGCTAACTGGCATAGAGGTAACATGTAATCCAGTCTGAGCAGAGCTGCGACCATCAATCAAGCAAGCTTC

CGAGCCATTCCCAAGCCAAAGCATCACACCTAGGGGCACTCAACAGCACAACTCAGCCGACAGTGTTATAAAAAACTTAG

GAGCCAGGGCTATTGGAGCCATGGACTCCCTGTCAGTCAACCAGGTCCTATACCCTGAGGTCCATTTAGACAGCCCCATT

GTCACAAACAAGCTAGTTGCCATCCTTGAATACTCGGGCATCGACCACAACTATGTTCTTGAGGACCAGACTCTTATCAC

GAATATTAGATATAGGTTGGGATGCGGTTTTTCAAATCAAATGATCATCAATAACAGGGGGGTAGGTGAAACAGTCAATT

CCAAACTTAAAAGTTACCCCCGTAATTGTCATGTCATATATCCAGATTGCAATAAGGATCTGTTTTATATCAGGGACGGC

TGCATATCTAGGAAGCTCTCTGAGCTATTCAAGAAAGGTAATTCCCTGTACTCTAAGATAAGTCACCAGGTACTAGATTG

TCTCAGAAGAGTCAACGGCAAAATGGGCCTGGGCACAGATCTTAGCCATGGCCTGAAGGAGGGTATCCTCGATTTAGGGT

TGCACATGCATAGCTCTCAGTGGTTTGAGACCTTCCTGTTCTGGTTCACTATCAAGACTGAAATGAGGTCAATGATCAAA

GAACAATCCCATATATGCCACAAGAGGAGGTATAACCCAACCTTTGTCTCAGGAGATGCACTCGAGGTGCTTGTATCACG

GGACCTCGTTGTGATAATTGATAAAACCACTCAGTGTGTCTATTACCTGACATTCGAGCTGGTCCTTATGTATTGCGATG

TCATAGAGGGCAGACTTATGACGGAGACTGCCATGGCTATAGACCAGAGGTATTCAGAACTGCTAAGCCGGGTCAAATAC

TTATGGGATCTTATTGATGGGTTCTTCCCAACACTAGGTAACGCCACATACCAAGTTGTTGCTCTGCTTGAACCGTTATC

GTTGGCTTACCTTCAACTTCAGGATGTCACTCTGGAGTTAAGAGGTGCATTCTTGGATCACTGCTTCAAAGAACTCTACG

AGATACTGGAACATTGTGGCGTTGACACGGAAAGCACCTACGATTCCATCATCGAAGGATTGGATTACGTATTTATCACC

CACGATATACACTTAACTGGGGAGATTTTCTCCTTTTTCCGGAGTTTCGGACATCCCCGCCTTGAGGCGGTCACTGCCGC

AGAGAATGTCAGGAAACATATGAACCAACCTAAGGTAATCAGTTATGAGACTATGATGAAAGGGCACGCAGTGTTTTGCG

GGATCATCATAAATGGGTTTAGGGACCGCCATGGCGGCAGCTGGCCCCCCGTTGTGCTGCCAGAACATGCCTCTGCTGCA

ATTCGGAATGCGCAGGCATCCGGTGAAGGGTTGACCCATGACCTGTGTATAGACAACTGGAAGTCCTTCGTTGGATTCAG

GTTTGGCTGTTTTATGCCGCTCAGCTTAGACAGTGACCTGACCATGTATCTCAAAGACAAAGCACTAGCTGCACTGAAAA

ATGAGTGGGATTCAGTTTATCCGAAAGAATACCTCCGCTATAACCCACCCAGAGGGACGGAGTCAAGGCGACTGGTGGAG

GTGTTCCTGAATGACTCCAGCTTTGATCCTTATAACTTGATAATGTACGTTGTGAATGGCTCCTACCTTGAAGACCATGA

GTTTAATCTCTCATACAGCTTGAAAGAGAAAGAGATAAAAGAGACAGGGCGACTGTTTGCAAAAATGACTTACAAAATGC

GGGCCTGCCAGGTCATTGCGGAGAATCTGATATCAAATGGGGTAGGGAAGTATTTCCGAGACAATGGAATGGCGAAAGAT

GAGCACGACCTAACAAAGGCCCTTCATACCTTGGCAGTCTCAGGTGTTCCTAAAAACAACAAGGATAATCACCGGGGTGG

GCCTCCTAGAAGGACCACAAGCCAAAAGATGAGGTCAGATCGGGGCATCACCAGTCAAGATAGGGACAAAATACAGAGAG

AGCCCATGTATAACTATCTGCGATGCCAACCAGTCAGCCCTGATCAGGGTGAGTCATACGAGACTGTTAGTGCATTCATC

ACCGCTGACCTGAAGAAATATTGCCTGAACTGGAGATACGAGACAATCAGCATATTTGCACAGAGACTAAATGAGATTTA

TGGACTACCATCTTTCTTCCAATGGTTACACAGGGTATTAGAAAAATCTGTGCTGTACGTCAGTGACCCACATTGCCCTC

CCGACTTGGACGATCATATCCCTCTGGACAGTGTCCCCAATGCCCAAATATTCATCAAGTACCCGATGGGCGGGATAGAA

GGTTATTGTCAAAAATTATGGACAATCAGTACTATACCATACTTGTATCTAGCAGCCCATGAGAGCGGAGTAAGAATCGC

CTCTCTAGTGCAAGGTGATAATCAAACAATTGCAGTGACAAAGAGAGTCCCGAGTTCCTGGCCTTATTCACTAAAAAAGA

GGGAGGCATCTAGAGCAGCTCAGAATTACTTCGTGGTTTTAAGACAGAGGTTGCATGATGTAGGTCATCACTTAAAGGCC

AATGAAACTATAGTATCTTCTCATTTTTTTGTATACTCAAAAGGGATTTATTATGATGGCCTGTTAGTATCACAATCACT

AAAGAGCATTGCCAGATGTGTCTTCTGGTCCGAGACTATTGTGGATGAAACCAGAGCCGCCTGCAGCAATATTGCAACAA

CTGTCGCCAAGAGTATAGAGAGGGGTTATGACAGGTACCTTGCTTACTCCTTGAACATACTCAAAATTCTCCAACAGATC

CTTATATCCCTTGACTTCACTATCAACACAACAATGACTCAAGATGTCGTGGCACCGATCATCGAGAACGGTGACCTGCT

AACAAGGATGGCACTTTTGCCAGCACCCATCGGGGGCCTTAACTACCTCAACATGAGCAGGTTATTTGTGAGAAATATCG

GTGACCCAGTCACTTCCTCTATAGCCGACCTGAAGAGGATGATAGACGCTGGACTAATGCCAGAGGAAACATTGCACCAG

GTGATGACCCAGACCCCGGGAGAATCATCTTACCTTGATTGGGCAAGTGACCCTTATTCTGCCAACCTAACCTGTGTACA

GAGTATAACTCGTCTTCTCAAGAACATCACTGCAAGGTACATTTTAACCAGCAGCCCAAATCCGATGCTAAAAGGGTTAT

TCCATGAAGGGAGCAGAGATGAGGATGAAGAGCTTGCGAGTTTCCTGATGGATCGGCATATAATTGTTCCCAGAGCTGCA

CATGAAATCTTAGACCATAGTATAACTGGGGCAAGAGAAGCTATTGCCGGTATGTTAGATACCACCAAGGGTCTGATTAG

AACAAGTATGAAACGAGGTGGCCTTACGCCACGGGTCTTAGCTCGCCTTTCCAATTATGATTATGAACAATTTAGATCCG

GTATAACATTGCTGACAAAGAAGGGACAGTGTTATCTCATTGACAAAGATTCATGCTCGGTACAGCTTGCTATAGCCCTG

AGGAGTCATATGTGGGCTAGGTTGGCCCGCGGGAGACCTATCTATGGGTTGGAAGTGCCTGATGTACTGGAATCGATGAA

TGGCTATCTTATCAAACGCCATGAGGCATGTGCTATTTGCGAGACGGGCTCAGGTCATTACGGGTGGTTCTTCGTCCCTG

CAGGGTGCCAGCTTGACGATGTCTCGAAAGAGACTTCAGCTCTTCGTGTGCCTTATGTTGGATCAACCACAGAGGAAAGG

ACAGATATGAAACTTGCCTTCGTTAGATCTCCGAGTCGATCCCTCAAATCAGCAGTCAGAATTGCTACCGTTTATTCATG

GGCCTATGGGGATGATGAAAAATCATGGAGTGAAGCCTGGACGTTGGCAAGGCAGAGAGCCGACATCACCCTAGATGAGT

TGAGGATGATCACCCCAGTCTCTACGTCGACCAACCTAGCCCATCGGTTGAGGGATCGGAGCACCCAGGTGAAATACTCG

GGGACATCCCTTGTGAGGGTTGCGAGATACACAACCATTTCCAATGATAATTTGTCATTTGTGATATCTGAGAAGAAAGT

AGATACCAACTTCATTTACCAGCAAGGGATGCTCCTCGGTCTTGGGATCCTTGAGAATCTTTTCAGGTTGGAGACCACCA

CGGGGGTATCCAACACAGTGCTACATCTGCACGTGGAGACAGAATGTTGTGTTGTACCTATGATTGATCACCCAAGAATA

CCAAGTCTCCGTAACATTAAGGTTACGAATGAGCTATGTACAAACCCCTTGATCTACGACAAGTCTCCTATCATAGAACA

CGATGCAACTCGATTATACTCACAAAGCCACAGAAGACATCTGGTGGAATTTGTTACCTGGTCAACAAGTCAGCTTTACC

ATATACTAGCTAAGTCTACAGCAATGTCCATGATTGAGTTGATTACAAGATTCGAGAAGGATCACATGAATGAAATAACC

GCCTTGATTGGCGATGACGACATCAACAGTTTTATAACAGAGTTCCTGCTAGTAGAACCTAGACTGTTTATAGTCTACCT

TGGCCAGTGTGCTGCCATCAATTGGGCTTTTGATATACATTATCATCGCCCATCGGGCAAATACCAGATGGGGGAACTTC

TCTACTCTTTACTCTCTAGGATGAGCAAAGGAGTATATAAGATCTTTACCAATGCTCTGAGCCACCCCAAAGTTTACAAG

AAATTTTGGCGAAGCGGTGTCATCGAGCCGATTCATGGCCCGTCCTTAGACACACAGAATCTACATGTCACTGTCTGTGA

TATGATATATGGATCCTACGTCACATATCTGGATCTTTTGCTGAATGATGAGCTAGATGATTATACGTATCTGCTCTGCG

AAAGTGATGAGGACGTGGTCACAGACAGGTTTGATAACATCCAAGCCAAACATCTCTGTGTATTGGCCGATGTGTATTGC

AGCTCCAAAAGGTGTCCCTCAATAATCGGGATGTCTCCTATAGAAAAATGTACCATCCTCACACATTACATAAAAGGGGA

ATCAATACAGTCTCCGTCTGGGGCCTCATGGAACACTGATCCTCTTGTAGTAGATCATTACTCATGCTCTCTGACATACC

TTCGTCGCGGTTCCATCAAACAAATCAGGTTGAGAGTAGATCCTGGGTTTGTATTTGAGGCATTGACAGACATCGACGTC

AAACAGCCTCGCAAGGCTAACACTGGTGTATCGGTTGTGGGGTTGACTGATTTCTCTCCTCCTTGGGATAACATTGGTGT

TTTTCTAAGGACTATCAACACCTTGAGACACAATCTGCCAATCACCGGGACCGGAGTCTTAAACTATGAGGTCCACGCTT

ATCGTAGAATTGGTTTGAATTCATCAGCATGTTATAAAGCTGTAGAGATCTCCACGTTAATCAAGCCATCCTTAGAAGTC

GGAGAGCATGGGTTGTTCTTAGGGGAAGGTTCCGGTTCTATGCTGGCTGCGTACAAGGAAATTCTTAAATTAGCCAACTG

TTATTACAACAGCGGAGTAATAGCTGAGGGCAGAGCCGGACAAAGAGAAATATCCCCTTACCCATCAGAGATGAGCCTAG

TAGAGAATCAAATGGGGTTAGAAAAGAGTGTTAAAGTGCTGTTCAACGGCAAACCTGAAGTAACCTGGGTAGGGACTACC

GATTGTTATGAATATATAATCAGTAACGTCCATACCTCTAGTCTGGGTTTCATACACTCGGATATTGAGACTCTCCCAAC

CAAAGATGCTGTTGAGAAATTAGAAGAATTTGCCTCTATCCTTTCCTTATCCCTAGTTTTAGGAAAAATCGGCTCTATTA

CAGTTATCAAAGTTATGCCTATTAGCGGAGATTTTACTCAGGGCTTCATAAGCTATGCCATCCAGTATTTTAGGGAGAGC

CTGCTTGTCTATCCGAGATATAGTAATTTCATTTCAACTGAGTGTTACCTTATTATGATAGGATTGAAAGCTAATCGGTT

GATAAACCCAGAAGCCATTAAGCAAAGCATAATCAGAGCAGGGACTAGAACTGCACCAGGACTAGTGAGCCATATATTAT

CAGAGAAACAAAAAGGGTGTATCCAATCTTATCTGGGTGATCCTTACATCCAAGGAGACTTCAATAAACACCTTAAATCT

CTAACCCCTATCGAGAAAATCCTAATTAATTGTGGTCTCTCGGTCAATGGCACAAAAATCTGTAGGGATCTAATCCACCA

TGATATCGCCTCTGGTCCGGACGGTCTGATGAGCTCCACAATAATCTTATACAGGGAGCTGGCTCATTTCAAAGACAACG

TAAGAAGTCAGCACGGGATGTTCCACCCCTACCCAGTACTGGCCAGTAGCAGGCAACGCGAATTAATTCTTCGAATCGCC

AAGAAATTCTGGGGTTATGTCTTGCTATATTCTGATGATCCGGCACTGATCAGGCAAACAATCAAGAATTTGAAAAGGAA

CCACTTAACCTTTGACCTACACAATAACCCGTTTATTAAGGGCCTGTCCAGAGCTGAGAAACTGCTAGTGCGTACGAGTT

CACTCAGAAGAGAGTGGTTGTTCACTCTCGAAACGAAGGAAGTGAAAGAATGGTTTAAACTGGTGGGCTACAGTGCACTC

GTCAGAGGTTAATTAGCTATACATCTGTCCCCTTCTCCTCCACCATGAAATTCCACTGGCAATCTAATAGATTAAAGAAA

ACTACAGATCAGATAAGTATCTATTCCCAGCTTTGTCTGGT

>KJ867545.1/DorcasGazelle/UAE/1986

ACCAAACAAAGTTGGGTAAGGATAGTTCTTATAATAACTATGAATTGCCACGCTTAGGATTAAAGATCCTACTGCCGGGG

GGAGGAGTAGGAGCAAGATCCTTGACTATGGCGACTCTCCTCAAAAGCTTGGCACTGTTCAAGAAGAACAAAGACAAAGC

GCCGACGGCATCAGGTTCAGGTGGGGCCATCCGGGGGATTAAGAATGTTATCATAGTCCCGATTCCCGGAGACTCATCCA

TCATTACCCGTTCAAGATTGCTCGACAGGCTTGTCAGATTGGCCGGAGATCCGGATATCAACGGGTCAAAGCTGACCGGC

GTGATGATCAGCATGCTGTCTTTGTTCGTAGAGTCACCCGGGCAACTGATCCAGCGGATCACAGATGATCCAGATGTCAG

TATCCGCCTTGTTGAGGTGGTCCAAAGTACTAGATCTCAGTCCGGGTTGACCTTTGCATCACGTGGTGCTGATTTGGATA

ATGAGGCAGACATGTATTTTTCAACTGAGGGGTCCCCGAGTGGGAGCAAAAAAAGGATCAACTGGTTTGAGAACAGGGAA

ATAATAGACATAGAAGTGCAGGATGCAGAAGAGTTCAATATGTTATTAGCCTCCATACTGGCACAAGTTTGGATTCTCCT

GGCCAAAGCGGTCACGGCACCGGACACTGCAGCTGACTCAGAATTGAGAAGGTGGGTTAAATACACACAACAAAGAAGAG

TGATTGGGGAATTTCGCCTTGACAAAGGGTGGCTAGATGCGGTCCGCAATAGGATTGCAGAAGATCTATCACTCCGGCGG

TTCATGGTATCTCTTATACTTGACATCAAAAGGACCCCTGGCAACAAGCCAAGGATTGCAGAGATGGTCTGTGACATTGA

TAACTATATTGTCGAAGCCGGGCTCGCCAGTTTCATCCTTACCATCAAGTTTGGTATTGAAACCATGTATCCTGCACTAG

GTCTTCACGAGTTTGCCGGGGAGTTGTCCACTATAGAATCCTTGATGAATCTGTATCAACAGCTAGGCGAAGTTGCACCC

TACATGGTTATTCTAGAGAACTCAGTTCAGAATAAGTTTAGTGCAGGAGCCTATCCTCTTCTCTGGAGCTATGCGATGGG

TGTTGGAGTCGAGCTGGAGAACTCGATGGGGGGGCTGAACTTTGGTAGATCATATTTTGACCCGGCTTATTTTCGTCTCG

GACAGGAGATGGTCAGAAGATCAGCAGGAAAGGTCAGCTCTGTAATCGCAGCTGAGCTCGGCATCACAGCAGAGGAGGCC

AAACTAGTCTCGGAAATCGCCTCTCAGACTGGGGACGGAAGGACCACTAGAGGGACTGGGCCCAGACAGGCGCAGGTTTC

CTTCCTCCAGCATAAAACAGGAGAGGGAGAGTCACATGCATCGGTGACCAGGGAAGAAGTCACAGCTGAGACCCCAAATG

GGCCCGACGAGAAGGACAAGAAACGAGCACGCCCAGGAAGGCCAAGAGGAGGAACCCCCGACCAACTGCTCCTGGAGATC

ATGCCTGAAGACGAGGTCCCGCGAGGGCCTGGACAAACCCCTCGTGAGGCTCAACGATCGGCAGAGGCACTCTTTAGACT

ACAGGCCATGGCCAAGATTCTAGAGGGCCAGGAGGAGGGAGAAGACAACAGTCAGATATACAACGACAAGGATCTCCTCA

GCTGAGCAGAAGCGCCCTCCATCGAAACCAGTGACAAGACACCGCCCATCAGTATTATAAAAAACTTAGGACCCAGGTCC

AAGCGGTCACACCCCGGCACCCCAGCCGACCGAGCGGAGACCACCGATGGCAGAGGAACAAGCATACCATGTCAACAAGG

GACTGGAATGTATCAAGTCCCTCAAAGCCTCTCCCCCGGATCTATCCGCCATCAGAGATACCCTTGAAAACTGGAGAGAG

GGGCTTGACCCATCGGACCGTGCAACACCGAACCCTGATATGTCCGAGGGGAACCATCAGAATATCAACCAATCATGCTC

ACCAGCAATCGGATCAGGCAAGATCGACGTGTCTACTGAAGGTAATCTCAGATATAGAGAGATCAATTACGATGACAGTG

AGGCTGGACTCAGAGGAGTTCAAGACAGAGGATCCGACTCTCAAGTACAGCGTTACCATGTTTATAGCCACGGGGGTGAA

GAGATTGAAGGACTCGAGGATGCTGACTCTCTCGTGGTTCAAGCAGATCCTCCACTTGCTAACACCTTCAGTAGAGGAGA

GGATGGATCTGACGACAGCGATGTGGACTCTGGCCCAGATGATCCCGACAGAGATCCTCTATATGACCGGGGATCTGTTG

CCGGCAATGATGTCGTTAAGTCCACCGATGTCGAAAAATTAGAAGGTGACAACATTCAAGAAGTTCTTAACTCTCAGAAG

AGTAAAAGAGGAAGATTCCAAGGCGGGAAAACTTTGCGAGTCCCTGAAACACCCGATGTCAACCACCCCAGACCATCGGC

CCAATCAATTAAAAAGGGCACAGACGGGAACTCAGTCTTATCTGGAACGGTGACAGAGTGTTCATCGATAAGTGGTGCAA

CCCAAGCTGTGCCAGAGTCAAGATGGGAGTCATCAGAGCGAAATGCATTTGTGGAGAGTGTCCCCAAATCTGCGAGGAGT

GCAAAGACGACCCAGGGGTTGACACAAGAATCTGGTACCATAACACCACCGACTCAGCCTAAAGAGAATGACTCCGAGTA

TGAGTATGAGGATGATCTATTTACAGAAATGCAGGACATTCGTGCAAGCATTGCCAAGATCCATGATGACAACAAAACTA

TCCTCTCAAAGCTTGACTCTATACTATTACTGAAAGGAGAAATCGATACTATCAAGAAGCAAATCAGCAAACAGAATATA

AGTATATCTACCATTGAGGGTCATCTTTCCAGTATAATGATAGCCATCCCGGGTTTTGGAAAGGAGGTCAAGGACCCAAC

ATCTGAGGTTGAGTTGAACCCGGATTTAAGACCTATAATCAGCCGTGATTCTGGCAGGGCTCTTGCGGAGGTCCTCAAGA

AACCTGCTGTTGATAGGTCTCCCAAAACCGGAATCAAGGTCAACTCTGGTTCAAAGGGTCAGCTCCTCAAGGATCTCCAA

CTAAAACCCGTTGACAAACAGGCAAGCTCTGCCATAGGGTTTGTCCCGTCTGACCATGAATCATCCAGAAGTGTCATCCG

CTCTATAATCAAGTCAAGCAAGCTCAACATTGATCACAAAGACTATCTTTTAGACTTACTGAATGATGTGAAAGGCTCCA

AGGATCTTAAAGAATTCCACAAGATGCTAACAGCTATTCTTGCCAAGCAACCGTAATGCACCGCCCAATCAACTTCCCAT

GCTCGGCTGAAAACAGCTCCTCAACCAGGCTATTACAAAAAACTTAGGAGCAAGGGCAACCGAGCTTCGCAGACAAGATG

ACCGAGATCTATGACTTCGACAAATCAGCATGGGACGTCAAAGGATCAATAGCTCCCATAGAGCCCACCACTTATCACGA

TGGTCGACTGGTACCCCAGGTGAGAGTCATCGATCCCGGGCTAGGAGATAGGAAAGATGAGTGCTTCATGTACCTGTTCC

TCCTAGGGGTGATTGAGGATAATGACCCCCTGTCTCCCCCAGTGGGAAGAACCTTTGGTTCTTTACCTCTGGGGGTCGGC

AGGTCAACTGCTAAGCCAGAAGAACTATTAAGGGAGGCCACTGAACTTGACATAGTGGTGAGACGCACGGCTGGACTAAA

CGAGAAGCTGGTATTTTACAACAACACTCCGCTGTCCTTGTTAACACCTTGGAGGAGAGTCCTGACAACCGGAAGTGTGT

TCAGCGCTAACCAGGTTTGTAACGCAGTCAATCTAGTCCCACTCGATACTCCCCAGAGATTTAGGGTTGTGTACATGACC

ATCACTAGATTGTCGGACAACGGTTATTATAGTGTTCCCAGGAGAATGTTAGAGTTTCGCTCAGCCAATGCCGTCGCCTT

CAATATCTCGGTTACACTGAGAATTGAGAATGGGACAAACCCTAGAAGGTACATAGTCGGCTCATGGGAGAATCCAGAGG

TCACATTCATGATACACGTGGGCAACTTTAGAAGAAAGAAGAACGACGTATATTCTGCTGATTACTGCAAAATGAAGATT

GAGAAGATGGGTCTGGTTTTTGCTTTGGGAGGAATAGGGGGAACAAGTCTTCACATTAGAAGCACCGGGAAAATGAGCAA

GACCCTCCATGCACAGCTGGGGTTCAAGAAAATCTTATGCTACCCCTTAATGGATATCAATGAGGATCTCAATCGATATC

TTTGGCGGGCAGAGTGCCGAATAGTCAAAATTCAAGCTGTCTTACAGCCTTCAGTACCCCAAGAATTCCGTGTCTACGAT

GATGTTATCATTAACGATGACCAAGGCTTGTTTAAGATCCTGTAATTCACTCACAGAGCAATGGACCCGCCTCCGACAAA

GGACCACCCCTCCTCTCTTCCAAAAAACATACTCCGAGCCCCCCCAAAAAAACGCCGACATGGTCATCCCCCCGGTCGGA

CGAGGACGACCCCCCCAGCGGACCCGAGTCTGCGGGGGGGGCCCCCCTCGCGACCCGCCGAGCACCGACCGCGCGGGGCA

GCAGGACTCCCCTGGGAGGCCCCAAGGTAGACCGGGCACCCCCTCCCCCCCCCGTCGC-CCCCC-AACACGCCCGAGGCC

AGGAGCCGAGCACCCCCAGCCCTGCCACCGACAGGACA-ACAACCATGCCAGGGCCCCC--GACCAAAGAGACCACCAGG

GAGGGCTCTGTCGACCCAACACAGACTTGGCCCTAACAAGGCAGATCCCCAAAACGAAATCGCCCAGTGCACTCCACTCA

TCATCAAACAAAACTTAGGGGCCAAGCTCAGACGACAGACCCCACCCCGCCAAGCAGCAGGCCGCACAAAGGAAGCACAC

AAGACCGGCTCGGAAGGCAGACACCCGACCACAGAAACACTCAAGCCCCGGGGAGGGGGCCCCCCCCCCAAAAAGCGGAG

GGGCGGGAGAGAAACACCGAGGACAACCCGACCGGGAGCCGGGACCCGGAGGCCCCAGCGGCAACCGGGCCAGGCACACA

CCCGACACCCCCGGCGGAGCAGAGGAGAGCCCGACCCCGCGTGGCCCCGACGGCAGAAGATCCAACCCTGCCACAAAGAG

AAGCGAGGAGATC--------CTCCACCGCACACAGCCGGCAGACAAACCATAGGGGGACCACCTCCCGGGAGGACAAGC

GGCCCTCCCACAACTGAGCCGCCATCCCCCAGAGGAGGACACCCCGGAAGCCCCAGGACACCTCCCTGCCCTCGACCACC

GGGAGAGGGACCACACCGACAAAGACTAGAGCCACCGGGAGCCAACCAAGGGTCCCCCTCGGACTCGCCCGGCGCCCCCC

CGAAGAGGCCCCCACAGGACCTCTCGGCCCAACAACCCCCGCACGCGCCAAAACAAAAACCCACCAGAGGATATCCAGCA

CATTCATACACAAACATCATGACACGGGTCGCAATCTTGACATCTCTGTTTCTCCTCACCAACACTGTCGCATCCCAGAT

TCACTGGGGCAACCTATCCAAGATCGGGATTGTGGGAACAGGAAGTGCCAGTTATAAGGTGATGACTAGGCCGAGCCAAC

AAACCCTGGTTATAAAGCTAATGCCAAATGTAACAGCCATAGACAATTGTACGAAATCAGAGATTTCGGAATACAAAAGA

TTGCTGATCACAGTGTTAAAACCTGTCGAAGATGCTCTGTCAGTGATAACCAAGAATGTAAGACCAATTCAAGCTCTAAC

ACCCGGTCGCAGGACCCGCCGTTTTGCCGGGGCTGTCCTGGCCGGAGTGGCACTTGGAGTCGCAACTGCTGCTCAGATAA

CTGCCGGAGTCGCACTTCATCAATCTTTGATGAACTCCCAAGCAATTGAAGGGTTAAAAGCTAGTCTTGAGAAGTCAAAT

CAGGCAATAGAAGAAATCAGACTTGCAAATAAGGAAACCATACTGGCAGTACAGGGTGTCCAAGATTATATCAACAATGA

GCTAGTCCCTTCTGTTCATAAAATGTCATGTGAGTTGGTAGGTCACAAACTCAGCCTCAAGCTTCTTAGGTATTACACCG

AGATCTTGTCTATATTCGGGCCCAGCCTTCGAGACCCGATAGCTGCTGAAATATCAATCCAAGCACTCAGTTATGCACTA

GGTGGAGATATCAATAAGATCCTGGATAAGCTTGGGTATAGCGGCGGGGACTTCCTTGCTATCCTAGAAAGTAAGGGGAT

AAAGGCCCGAGTCACCTATGTGGATACAAGAGATTACTTTATAATTCTTAGCATAGCTTACCCGACCTTATCTGAGATCA

AGGGGGTGATAGTTCATAAAATGGAAGCTGTGTCATATAACATCGGGGCACAAGAATGGTACACTACTATCCCTAAATAT

GTAGCCACTCAGGGATACCTGATATCAAATTTTGATGAGACGTCATGCGTGTTCACTCCAGGAGGGACAGTCTGCAGTCA

GAATGCTTTATACCCAATGAGCCCGTTGCTTCAGGAATGTTTCAGGGGGTCTACAAAATCGTGCGCTAGAACCCTAGTTT

CAGGGACCATAAGCAATCGATTTATCCTATCAAAAGGGAACTTGATTGCCAATTGTGCATCAGTCCTGTGCAAATGTTAC

ACAACGGAGACAATTATCAACCAAGATCCTGATAAACTACTAACTGTTATAGCCTCCGATAAATGTCCTGTAGTTGAAGT

GGATGGAGTGACGATACAGGTCGGCAGTAGAGAGTACCCAGATTCTGTGTACCTACATAAAATAGACTTGGGCCCAGCCA

TCTCTCTAGAAAAACTGGACGTAGGCACCAATTTAGGCAATGCAGTTACAAGACTGGAGAATGCAAAGGAGCTCCTAGAT

GCATCAGATCAGATATTGAAGACTGTTAAGGGGGTACCCCTCAGCGGCAACTTGTACATAGGACTGGCAGCTTGCATTGG

GGTATCTCTAGGGCTTGTCACATTAATATGCTGCTGTAAGGGGAGGTGTAGGAACAAGGAGATTCCTACCCCTAAAATCA

ACCCAGGGCTCAAACCAGACCTGACAGGGACCTCAAAGTCGTACGTGAGATCACTGTAGTCAGAATAACCCGAATCATCT

ATCGTCACACACAGACATGTGCAACACAAGCAGTCAAGGGTCGTAGAAGATTCAACTTCCGATCATCAACCTGGCCTTAC

TCTACGCCCCATTGCACATTGGTTATCAAACAAAACTTAGGACGAAAGGTTAGTCACCATGTCCGCACAAAGAGAGAGGA

TCAATGCTTTCTACAAAGACACTCCTCACAACAAGAACCATAGGGTGACCCTGGATAAGGAACGTTTGACTATTGAAAGA

CCCTATATCCTGCTCGGGGTCCTACTGGTAATGTTTCTGAGCCTAATTGGGCTGCTGGCCATTGCAGGGATTAGACTTCA

CCGAGCCACCGTCGGAACTGCAGAGATCCAGAGCCGGCTTAATACCAATATTGAGTTAACTGAATCCATTGATCATCAAA

CTAAGGATGTCCTAACTCCCCTGTTTAAAATCATCGGTGATGAAGTCGGCATAAGGATTCCACAGAGGTTCAGTGACCTT

GTCAAGTTCATCTCCGATAAGATTAAATTCCTCAACCCTGACAAAGAATATGATTTCAGGGATCTCCGGTGGTGTATGAA

TCCCCCCGAGAGAGTTAAAATAGATTTTGATCAGTTTTGTGAATACAAAGCCGCGGGTAAGTCGATTGAACATATATTTG

AGTCACCACTCAACAGGTCAAAAAGGTTGCGATCGTTAACTCTAGGGCCTGGAACGGGCTGTCCTGGCAAGGCAGTAACA

AGGGCCCAGTTCTCAGAGCTTACTCTGACCCTGTTGGATCTGGATCTCAATATGAAGCACAACGTGTCCTCAGTGTTTAC

CATAGTTGAAGAGGGATTATTCGGAAGAACATACACCGTCTGGAGATCAGATGCCGGAAATCTGAGCACCGGCTCAGGTA

TTGGCAATTTTCTGAGAGTCTTCGAGATCGGACTGGTAAGAGATTTCGGGCTAAGTGCCCCTGTTTTCCATATGACTAAC

TATCTCACAGTGAACATGAGTGATGACTATCGGAGCTGTCTATTAGCAGTAGGGGAGTTGAAGCTGGCAGCTCTATGCAC

TCGTTCTGAGACTGTGACTCTGAGTGATAGAGGAGTTCCTAAGAGAGAGCCTCTTGTGGTTGTAATACTTAATCTGGCCG

GGCCTACTCTAGGGGGCGAACTATATAGTATTTTGCCTGCCTCTGACCTCATGGTAGAGAAACTCTATTTGTCATCACAT

AGAGGGATTATCAAAGACAACGAGGCCAATTGGGTAGTACCGTCTACCGATGTTCGTGATCTTCAAACCAAAGGAGAATG

CTTGGTGGAAGCATGCAAAACTCGACCTCCTTCATTTTGCAATGGTACAGGGTTAGGCCCATGGTCAGAGGGGAGAATCC

CCGCCTACGGGGTGATCAGGGTCAGTCTTGACGTAGCTAGTGATCCAGATGTGGTTATTACCTCAGTATTTGGCCCCTTG

ATACCTCATCTATCTGGCATGGATCTTTACAACAACCCATTCTCAAGGGCTGTGTGGTTGGCTGTACCACCTTATGAGCA

GTCATTTCTGGGAATGATAAATACAATTGCCTTCCCAGATAGAGCAGAGGTTACGCCGCACATTCTGACCACAGAGATCA

GAGGGCTTCGGGGTCGTTGTCATGTTCCTATAGAATTGTCCCGCAGGATTGATGATGATATTAAGATTGGCTCCAACATG

GTTGTACTACCGACGAAGGACCTGAGGTACATTACAGCCACTTATGATGTTTCCAGGAGCGATCATGCCATTGTGTACTA

TATCTATGACACAGGTCGCTCATCATCTTACTTCTTCCCCGCCCGATTAAAGATCAAAGGCAACCCTCTCTCCCTGAGAA

TAGAGTGTTTCCCCTGGTATCATAAGGTGTGGTGCTACCATGACTGCCTCATATACAACACCATCACAAACGAAGAAGCC

CATACGAGAGGGCTAACCGGCATAGAGGTAACATGTAATCCAGTCTGAGCAGAGCTGCGACCATCAATCAAGCAGGCTTC

CGAGCCATTCCCAAGCCAAAGTAGCACAGCTAGGGGCACCCAACAGCACAACTCAGCCGACAGTGTTATAAAAAACTTAG

GAGCCAGGGCTATAGGAGCCATGGACTCCCTGTCAGTCAATCAGGTCCTATACCCTGAGGTCCATTTAGACAGCCCCATT

GTCACAAACAAGCTAGTTGCCATCCTTGAGTATTCGGGCATCGACCACAACTATGTTCTTGAAGACCAGACCCTTATCAA

GAATATTAGATATAGACTGGGGTGCGGTTTTTCAAATCAAATGATCATCAATAACAGGGGGGTAGGTGAAACAGTCAATT

CCAAGCTTAAAAGTTACCCCCGTAGTTGTCATGTCATATATCCGGATTGCAATAAGGATCTGTTTTGTATCAGGGACGGC

TGCATATCTAGAAAGCTCTCTGAGCTATTCAAGAAGGGTAATTCCCTGTACTCCAAGATAAGTCACCAGGTACTAGATTG

TCTCAGGAGAGTCAACGGCAAAATGGGCCTGGGCACAGATCTTAACCATGGCCTGAAGGAGGGTATCCTCGATTTAGGGT

TGCACATGCATAGCTCTCAGTGGTTTGAGACTTTCCTGTTCTGGTTCACTATCAAGACTGAAATGAGATCCATGATCAAA

GAACAATCCCATATATGCCACAAGAGGAGGTATAACCCAACCTTTGTCTCAGGAGATGCATTCGAGGTGCTTGTATCACG

AGACCTCGTTGTGATAATTGACAAGAATACTCAGTGTGTCTATTACCTGACATTCGAGCTGGTCCTTATGTATTGTGATG

TCATAGAGGGCAGACTTATGACGGAGACAGCTATGGCTATAGACCAGAGGTATTCAGAACTGTTAAGCCGGGTCAAATAC

TTGTGGGATCTTATTGATGGGTTCTTCCCAACACTAGGTAACGCCACATACCAAGTCGTTGCTCTGCTTGAACCATTATC

GTTGGCTTACCTTCAACTTCAGGATGTCACTCTGGAGTTAAGAGGCGCATTTTTGGATCACTGCTTCAAAGAGCTCTATG

AGATACTGGAACATTGTGGCGTTGACACGGAAAGCACCTACGATTCCATCATCGAAGGATTGGATTACGTATTTATCACC

CACGATATCCACTTAACTGGGGAGATTTTCTCCTTTTTCCGGAGTTTCGGACACCCCCGCCTCGAAGCGGTCACCGCTGC

AGAGAATGTCAGAAAACACATGAATCAACCTAAGGTAATCAGTTATGAGACTATGATGAAAGGGCACGCAGTATTTTGCG

GAATCATCATAAATGGGTTTAGGGACCGCCATGGCGGCAGCTGGCCCCCCGTTGTGCTGCCAGAACATGCCTCTGCTGCA

ATTCGGAATGCGCAGGCATCCGGTGAGGGGCTGACCCATGACCTGTGTATAGACAACTGGAAGTCCTTCGTTGGATTCAG

ATTTGGCTGTTTTATGCCGCTCAGCCTAGATAGTGACCTGACCATGTACCTCAAAGACAAAGCACTAGCTGCACTGAAAA

ATGAGTGGGATTCAGTCTACCCGAAAGAATACCTCCGCTATAACCCACCTAGAGGGACAGAGTCAAGGCGACTGGTGGAG

GTGTTCCTGAATGACTCCAGCTTTGATCCTTATAACATGATAATGTACGTGGTGAACGGTTCCTACCTTGAAGACCACGA

GTTTAATCTCTCATACAGCTTGAAAGAGAAAGAGATAAAAGAGACAGGGCGATTGTTTGCAAAAATGACTTACAAAATGC

GGGCCTGCCAGGTCATTGCGGAAAATCTGATATCAAATGGGGTAGGGAAGTATTTCCGAGACAATGGAATGGCGAAAGAT

GAGCATGACCTAACAAAGGCTCTTCATACCCTGGCAGTCTCAGGCGTTCCCAAAAACAACAAAGATAATCACCGGGGTGG

GCCTCCCAGAAGGACCACAAGCCAAAAGATGAGGTCAGATCGAGGCATCACCAGACAAGATAGGGACAAGGTACAGGGAG

AGCCCATGTATAACTATCTGCGATGCCAACCAGTCAGCCCTGATCAAGGTGAGTCATACGAGACTGTTAGTGCATTCATC

ACCGCTGATCTTAAGAAATATTGCCTGAACTGGAGATACGAGACAATCAGCATATTTGCACAGAGACTGAATGAAATTTA

CGGACTACCATCTTTCTTTCAGTGGTTACACAGGGTATTAGAAAAATCTGTGCTGTACGTCAGTGACCCGCATTGCCCTC

CCGACTTAGATGATCACATCCCTCTGGACAGTGTCCCCAATGCCCAAATATTCATCAAGTACCCAATGGGCGGAATAGAA

GGTTATTGCCAAAAATTATGGACAATCAGTACTATACCGTACTTGTATCTAGCAGCCCATGAGAGCGGAGTAAGAATCGC

CTCTCTAGTGCAAGGTGACAATCAAACAATTGCAGTGACAAAAAGAGTCCCGAGTTCTTGGCCTTATTCACTAAAAAAGA

GGGAGGCATCCAGAGCAGCTCAAAATTACTTCGTGGTTTTAAGGCAGAGATTGCACGATGTAGGTCATCACTTAAAGGCC

AATGAAACCATTGTATCTTCCCACTTTTTTGTATACTCCAAAGGGATTTATTATGATGGCCTGTTAGTCTCACAGTCACT

AAAGAGCATTGCCAGATGTGTCTTCTGGTCCGAGACTATTGTGGATGAAACCAGGGCCGCCTGCAGCAATATTGCAACAA

CTGTCGCCAAGAGTATAGAGAGGGGTTATGATAGGTACCTTGCTTACTCTTTGAATATACTCAAAATTCTCCAACAGATC

CTTATATCCCTTAACTTCACTATCAATACAACAATGACTCAAGATGTCGTGGCACCGATCATCGAGAACGGTGATCTACT

AACAAGGATGGCACTTTTGCCAGCACCCATCGGGGGCCTTAATTACCTTAACATGAGCAGGTTATTTGTGAGAAATATCG

GTGACCCAGTCACTTCCTCTATAGCCGACCTGAAGAGGATGATAGACGCTGGGCTAATGCCAGAAGAAACATTGCACCAG

GTGATGACCCAAACCCCGGGAGAATCATCTTACCTTGATTGGGCAAGTGATCCTTATTCTGCCAACCTAACTTGTGTACA

GAGTATAACTCGCCTTCTCAAGAACATCACTGCAAGGTATATTTTAACCAGCAGCCCAAATCCGATGCTAAGAGGGTTAT

TCCATGAAGGGAGCAGAGATGAAGATGAAGAGCTTGCGAGTTTCTTGATGGATCGGCATATAATTGTTCCCAGAGCTGCA

CATGAAATCTTAGACCATAGCATAACTGGAGCAAGAGAAGCTATTGCCGGTATGTTAGATACCACCAAGGGTCTGATTAG

AACAAGTATGAAACGAGGTGGCCTTACCCCCCGGGTCTTAGCTCGCCTTTCCAATTATGATTATGAACAATTCAGATCCG

GTATAACATTATTGACGAAGAAGGGACAGTGTTATCTCATTGACAAAGATTCGTGCTCGGTGCAGCTTGCTATAGCCCTG

AGGAGCCATATGTGGGCTAGGTTGGCCCGCGGGAGACCTATCTATGGGTTGGAGGTGCCTGATGTACTAGAATCGATGAA

TGGCTATCTTATCAAACGCCACGAAGCTTGTGCCATTTGCGAAACAGGCTCAAGTCATTACGGGTGGTTCTTCGTCCCTG

CAGGGTGCCAGCTTGACGATGTCTCGAAAGAGACTTCAGCTCTTCGTGTGCCTTATGTCGGGTCAACCACAGAGGAAAGG

ACAGATATGAAACTTGCCTTCGTTCGATCTCCGAGTCGATCCCTCAAATCAGCAGTCAGAATTGCTACCGTTTATTCATG

GGCCTACGGGGATGATGAAAAATCATGGAGTGAAGCCTGGATGCTGGCTAGGCAGAGAGCCGATATCACCCTCGATGAGT

TGAGAATGATCACTCCAATCTCTACATCGACCAACCTAGCCCATCGGTTGAGGGATCGGAGCACCCAGGTGAAATACTCG

GGGACATCCCTTGTGAGGGTTGCGAGATACACAACCATTTCCAATGATAATTTGTCATTTGTGATATCTGAGAAGAAGGT

AGATACCAACTTCATTTACCAGCAAGGGATGCTCCTCGGTCTTGGGATCCTTGAGAATCTTTTCAGGTTGGAGGCTACCA

CAGGGGTATCCAACACAGTGCTACATCTGCACGTGGAGACAGAATGCTGTGTTGTACCTATGGTTGACCACCCAAGGATA

CCGAGTCTCCGTAACATTACGGTTACGAATGAGCTATGCACAAACCCCTTGATCTACGACAAGTCTCCTATCATAGAACA

CGATGCAACCCGATTATACTCACAAAGCCACAGAAGACATCTGGTGGAGTTTGTTACCTGGTCAACAAGTCAGCTTTACC

ATATACTGGCTAAGTCTACAGCGATGTCCATGATTGAGTTGATTACAAGATTCGAGAAGGATCACATGAATGAAATAACA

GCCTTGATTGGCGATGACGACATCAACAGTTTCATCACAGAATTCCTGTTAGTAGAACCTAGACTGTTTATAGTCTACCT

TGGCCAGTGTGCTGCCATCAATTGGGCTTTTGATATACATTATCATCGCCCATCGGGCAAATACCAGATGGGAGAGCTTC

TCTACTCTTTGCTCTCTAGGATGAGCAAAGGAGTATACAAGATCTTTACCAATGCTTTGAGCCACCCCAAGGTTTACAAG

AAATTTTGGAGAAGTGGTGTCATCGAGCCGATTCATGGCCCGTCCTTAGATACACAGAATCTACATGTCACTGTCTGTGA

CATGATATATGGATCCTACGTCACCTATCTGGATCTTTTACTGAATGATGAGCTAGATGATTATACGTATCTGCTCTGCG

AGAGTGATGAGGACGTGGTCACAGACAGGTTCGATAACATTCAAGCCAAACATCTCTGTGTATTAGCCGATGTTTATTGC

AGCTCCAAAAAGTGTCCCTCGATAATCGGGATGTCTCCTATAGAAAAATGTACCATCCTCACACATTACATCAAGGGAGA

ATCAATACAATCCCCGTCTGGGACCTCATGGAACACTGAACCTCTTGTAGTAGATCATTACTCATGCTCTCTGACATACC

TTCGTCGCGGTTCCATCAAACAAATCAGGTTGAGAGTAGATCCTGGATTTGTATTCGAGGCACTGACAGACATCGACACC

AAACTGCCTCGAAAGGCTAATACTGATGTATCGGTTGTAGGGTTGACTGATTTCTCCCCTCCTTGGGATAACATTGGTGA

TTTTCTAAGGACTATCAACACCTTGAGACACAATCTGCCTGTCACCGGGACCGGGGTCTCTAACTATGAGGTCCACGCTT

ATCGTAGGATTGGTTTGAATTCATCAGCATGTTATAAAGCTGTAGAAATCTCCACGTTAATCAAACCATCCTTAGAAGTC

GGAGAGCATGGGTTGTTCTTAGGGGAAGGTTCCGGTTCTATGCTGGCTGCGTATAAGGAAGTTCTTAAATTAGCCAACTG

TTATTACAACAGCGGAGTAATAGCTGAGGGGAGAGCCGGACAGAGAGAAATATCCCCTTACCCTTCAGAGATGAGCCTAG

TAGAGAACCAAATGGGGTTAGAAAGGAGTGTTAAAGTGCTGTTCAACGGCAAACCTGAAGTAACTTGGGTCGGGACCACC

GATTGCTACGAATATATCATCAGTAACATCCAAACCTCTAGTCTGGGTTTCATACACTCAGATATTGAGACACTCCCAAC

CAAAGATGCTGTTGAGAAATTAGAAGAATTTGCCTCTATCTTATCCTTATCCCTAATTTTGGGAAAAATTGGCTCTATTA

CCGTTATCAAAGTTATGCCTATTAGCGGAGATTTTACTCAAGGCTTCATAAGTTATGCCATCCAATATTTTAGGGAGAGC

CTGCTTGTCTATCCGAGATATAGTAATTTCATTTCAACTGAGTGTTACCTTATTATGATAGGATTGAAAGCCAATCGGTT

GATAAACCCAGAAGCCATTAAGCAAAACATAATCAGAGCGGGGACTAGAACTGCACCAGGACTAGTGAGCCATATATTAT

CAGAGAAACAAAAAGGGTGTATCCAATCTTATCTGGGTGATCCTTATATCCAAGGAGACTTCAATAAACACCTTAAATCT

CTAACCCCTATTGAGAAAATCCTAGTTAATTGTGGTCTCTCGATCAATGGCACAAAAATCTGTAGGGATCTAATCCACCA

TGATATCGCCTCTGGTCCAGACGGTCTGATGAGCTCCACAATTATTTTGTATAGGGAGCTGGCACATTTCAAAGACAACA

TAAGAAGTCAGCACGGGATGTTCCACCCCTACCCAGTACTGGCTAGTAGCAGGCAACGTGAATTAATCCTTCGAATCGCC

AAGAAATTCTGGGGTTATGTCTTGCTATATTCTGATGATCCGGCACTGATCAGGCAAACAATTAAGAACTTGAAGAGAAA

CCACCTAACCTTTGACTTACACAGTAACCCGTTTATTAAGGGTCTGTCCAGAGCTGAGAAAATGCTAGTGCGGACAAGTT

CACTCAGAAGAGAGTGGTTGTTCACTCTCGAAACGAAGGAAGTGAAAGAGTGGTTCAAATTGGTGGGTTACAGTGCTCTC

GTCAGAGGTTAATTAGCTATACATCTGCCCCCTTCTCCTCCACCATGAAATTCTACTGGCAATCTAATAGATTAAAGAAA

ACTACAGATCGGATAAGTATCTATTCCCAGCTTTGTCTGGT

>KJ867544.1/goat/Oman/1983

ACCAAACAAAGTTGGGTAAGGATAGATCTTATAATAACTATGAATTGTCACGCTTAGGATTAAAGATCCTACTGCCGGGG

GGAGGAGTAGGAGCAAGATCCTTGACTATGGCGACTCTCCTCAAAAGCTTGGCATTGTTCAAGAAGAACAAAGACAAAGC

GCCGACGGCATCAGGTTCAGGTGGGGCCATCCGGGGGATTAAGAATGTTATCATAGTCCCGATTCCCGGAGACTCATCCA

TCATTACCCGTTCAAGATTGCTCGACAGGCTTGTCAAATTGGCCGGAGATCCGGATATCAACGGGTCAAAGCTGACCGGC

GTGATGATCAGCATGCTGTCTTTGTTCGTAGAGTCACCCGGGCAACTGATCCAGCGGATCACAGATGATCCAGATGTCAG

TATCCGCCTTGTTGAGGTGGTCCAAAGTACTAGATCTCAGTCCGGGTTGACCTTTGCATCACGTGGTGCTGATTTGGATA

ATGAGGCAGACATGTATTTTTCAACTGAGGGGTCCCCGAGTGGGAGCAAAAAAAGGATCAACTGGTTTGAGAACAGGGAA

ATAATAGACATAGAAGTGCAGGATGCAGAAGAGTTCAATATGTTATTAGCCTCCATACTGGCACAAGTTTGGATTCTCCT

GGCCAAAGCGGTCACGGCACCGGACACTGCAGCTGACTCAGAATTGAGAAGGTGGGTTAAATACACACAACAAAGAAGAG

TGATTGGGGAATTTCGCCTTGACAAAGGGTGGCTAGATGCGGTCCGCAATAGGATTGCAGAAGATCTATCACTCCGGCGG

TTCATGGTATCTCTTATACTTGACATCAAAAGGACCCCTGGCAATAAGCCAAGGATTGCAGAGATGGTCTGTGACATTGA

TAACTATATTGTCGAAGCCGGGCTCGCAAGTTTCATCCTTACCATCAAGTTTGGTATTGAAACCATGTATCCTGCACTAG

GTCTTCACGAGTTTGCCGGGGAGTTGTCCACTATAGAATCCTTGATGAATCTGTATCAACAGCTAGGCGAAGTTGCACCC

TACATGGTTATTCTAGAGAACTCAGTTCAGAATAAGTTTAGTGCAGGAGCCTATCCTCTTCTCTGGAGCTATGCGATGGG

TGTTGGAGTCGAGCTGGAGAACTCGATGGGGGGGCTGAACTTTGGTAGATCATATTTTGACCCGGCTTATTTTCGTCTCG

GACAGGAGATGGTCAGAAGATCAGCAGGAAAGGTCAGCTCTGTAATCGCAGCTGAGCTCGGCATCACAGCAGAGGAGGCC

AAACTAGTCTCGGAAATCGCCTCTCAGACTGGGGACGGAAGGACCACTAGAGGGACTGGGCCCAGACAGGCGCAGGTTTC

CTTCCTCCAGCATAAAACAGGAGAGGGAGAGTCACATGCATCGGTGACCAGGGAAGAAGTCACAGCTGAGACCCCAAATG

GGCCCGACGAGAAGGACAAGAAACGAGCACGCCCAGGAAGGCCAAGAGGAGGAACCCCCGACCAACTGCTCCTGGAGATC

ATGCCTGAAGACGAGGTCCCGCGAGGGCCTGGACAAACCCCTCGTGAGGCTCAACGATCGGCGGAGGCACTCTTTAGACT

ACAGGCCATGGCCAAGATTCTAGAGGGCCAGGAGGAGGGAGAAGACAACAGTCAGATATACAACGACAAGGATCTCCTCA

GCTGAGCAGAAGCGCCCTCCATCGAAACCAGTGACAAGACACCGCCCATCAGTATTATAAAAAACTTAGGACCCAGGTCC

AAGCGGTCACACCCCGGCACCCCAGCCGACCGAGCGGAGACCACCGATGGCAGAGGAACAAGCATACCATGTCAACAAGG

GACTGGAATGTATCAAGTCCCTCAAAGCCTCTCCCCCGGATCTATCCGCCATCAGAGATACCCTTGAAAACTGGAGAGAG

GGGCTTGACCCATCGGACCGTGCAACACCGAACCCTAATATGTCCGAGGGGAACCATCAGAATATCAACCAATCATGCTC

ACCAGCAATCGGATCAGGCAAGATCGACGTGTCTACTGAAGGTAATCTCAGATATAGAGAGATCAATTACGATGACAGTG

AGGCTGGACTCAGAGGAGTTCAAGACAGAGGATCCGACTCTCAAGTACAGCGTTACCATGTTTATAGCCACGGGGGTGAA

GAGATTGAAGGACTCGAGGATGCTGACTCTCTCGTGGTTCAAGCAGATCCTCCACTTGCTAACACCTTCAGTAGAGGAGA

GGATGGATCTGACGACAGCGATGTGGACTCTGGCCCAGATGATCCCGACAGAGATCCTCTATATGACCGGGGATCTGTTG

CCGGCAATGATGTCGTTAAGTCCACCGATGTCGAAAAATTAGAAGGTGACAACATTCAAGAAGTTCTTAACTCTCAGAAG

AGTAAAAGAGGAAGATTCCAAGGCGGGAAAACTTTGCGAGTCCCTGAAACACCCGATGTCAACCACCCCAGACCATCGGC

CCAATCAATTAAAAAGGGCACAGACGGGAACTCAGTCTTATCTGGAACGGTGACAGAGTGTTCATCGATAAGTGGTGCAA

CCCAAGCTGTGCCAGAGTCAAGATGGGAGTCATCAGAGCGAAATGCATTTGTGGAGAGTGTCCCCAAATCTGCGAGGAGT

GCAAAGACGACCCAGGGGTTGACACAAGAATCTGGTACCATAGCACCACCGACTCAGCCTAAAGAGAATGACTCCGAGTA

TGAGTATGAGGATGATCTATTTACAGAAATGCAGGACATTCGTGCAAGCATTGCCAAGATCCATGATGACAACAAAACTA

TCCTCTCAAAGCTTGACTCTATACTATTACTGAAAGGAGAAATCGATACTATCAAGAAGCAAATCAGCAAACAGAATATA

AGTATATCTACCATTGAGGGTCATCTTTCCAGTATAATGATAGCCATCCCGGGTTTTGGAAAGGAGGTCAAGGACCCAAC

ATCTGAGGTTGAGTTGAACCCGGATTTAAGACCTATAATCAGCCGTGATTCTGGCAGGGCTCTTGCGGAGGTCCTCAAGA

AACCTGCTGTTGATAGGTCTCCCAAAACCGGAATCAAGGTCAACTCTGGTTCAAAGGGTCAGCTCCTCAAGGATCTCCAA

CTAAAACCCGTTGACAAACAGGCAAGCTCTGCCATAGGGTTTGTCCCGTCTGACCATGAATCATCCAGAAGTGTCATCCG

CTCTATAATCAAGTCAAGCAAGCTCAACATTGATCACAAAGACTATCTTTTAGACTTACTGAATGATGTGAAAGGCTCCA

AGGATCTTAAAGAATTCCACAAGATGCTAACAGCTATTCTTGCCAAGCAACCGTAATGCACCGCCCAATCAACTTCCCAT

GCTCGGCTGAAAACAGCTCCTCAACCAGGCTATTACAAAAAACTTAGGAGCAAGGGCAACCGAGCTTCGCAGACAAGATG

ACCGAGATCTATGACTTCGACAAATCAGCATGGGACGTCAAAGGATCAATAGCTCCCATAGAGCCCACCACTTATCACGA

TGGTCGACTGGTACCCCAGGTGAGAGTCATCGATCCCGGGCTAGGAGATAGGAAAGATGAGTGCTTCATGTACCTGTTCC

TCCTAGGGGTGATTGAGGATAATGACCCCCTGTCTCCCCTAGTGGGAAGAACCTTTGGTTCTTTACCTCTGGGGGTCGGC

AGGTCAACTGCTAAGCCAGAAGAACTATTAAGGGAGGCCACTGAACTTGACATAGTGGTGAGACGCACGGCTGGACTAAA

CGAGAAGCTGGTATTTTACAACAACACTCCGCTGTCCTTGTTAACACCTTGGAGGAGAGTCCTGACAACCGGAAGTGTGT

TCAGCGCTAACCAGGTTTGTAACGCAGTCAATCTAGTCCCACTCGATACTCCCCAGAGATTTAGGGTTGTGTACATGACC

ATCACTAGATTGTCGGACAACGGTTATTATAGTGTTCCCAGGAGAATGTTAGAGTTTCGCTCAGCCAATGCCGTCGCCTT

CAATATCTTGGTTACACTGAGAATTGAGAATGGGACAAACCCTAGAAGGTACATAGTCGGCTCATGGGAGAATCCAGAGG

TCACATTCATGATACACGTGGGCAACTTTAGAAGAAAGAAGAACGACGTATATTCTGCTGATTACTGCAAAATGAAGATT

GAGAAGATGGGTCTGGTTTTTGCTTTGGGAGGAATAGGGGGAACAAGTCTTCACATTAGAAGCACCGGGAAAATGAGCAA

GACCCTCCATGCACAGCTGGGGTTCAAGAAAATCTTATGCTACCCCTTAATGGATATTAATGAGGATCTCAATCGATATC

TTTGGCGGGCAGAGTGCCGAATAGTCAAAATTCAAGCTGTCTTACAGCCTTCAGTACCCCAAGAATTCCGTGTCTACGAT

GATGTCATCATTAACGATGACCAAGGCTTGTTTAAGATCCTGTAATTCACTCACAGAGCAATGGACCCGCCTCCGACAAA

GGACCACCCCTCCTCTCTTCCAAAAAACATACTCCGAGCCCCCCCAAAAAAACGCCGACATGGTCATCCCCCCGGTCGGA

CGAGGACGACCCCCCCAGCGGACCCGAGTCTGCGGGGGGGGCCCCCCTCGCGACCCGCCGAGCACCGACCGCGCGGGGCA

GCAGGACTCCCCTGGGAGGCCCCAAGGTAGACCGGGCACCCCCTCCCCCCCCCGTCGC-CCCCC-AACACGCCCGAGGCC

AGGAGCCGAGCACCCCCAGCCCTGCCACCGACAGGACA-ACAACCATGCCAGGGCCCCC--GACCAAAGAGACCACCAGG

GAGGGCTCTGTCGACCCAACACAGACTTGGCCCTAACAAGGCAGATTCCCAAAACGAAATCGCCCAGTGCACTCCACTCA

TCATCAAACAAAACTTAGGGGCCAAGCTCAGACGACAGACCCCACCCCGCCAAGCAGCAGGCCGCACAAAGGAAGCACAC

AAGACCGGCTCGGAAGGCAGACACCCGACCACAGAAACACTCAAGCCCCGGGGAGGGGGCCCCCCCCCCAAAAAGCGGAG

GGGCGGGAGAGAAACACCGAGGACAACCCGACCGGGAGCCGGGACCCGGAGGCCCCAGCGGCAACCGGGCCAGGCACACA

CCCGACACCCCCGGCGGAGCAGAGGAGAGCCCGACCCCGCGTGGCCCCGACGGCAAAAGATCCAACCCTGCCATAAAGAG

AAGCGAGGAGATC--------CTCCACCGCACACAGCCGGCAGACAAACCATAGGGGGACCACCTCCCGGGAGGACAAGC

GGCCCTCCCACAACTGAGCCGCCATCCCCCAGAGGAGGACACCCCGGAAGCCCCAGGACACCTCCCTGCCCTCGACCACC

GGGAGAGGGACCACACCGACAAAGACTAGAGCCACCGGGAGCCAACCAAGGGTCCCTCTCGGACTCGCCCGGCGCCCCCC

CGAAGAGGCCCCTACAGGACCTCTCGGCCCAACAACCCCCGCACGCGCCAAAATAAAAACCCACCAGAGGATATCCAGCA

CATTCATACACAAACATCATGACAAGAGTCGCAATCTTGACATCTCTGTTTCTCCTCACCAACACTGTCGCATCCCAGAT

TCACTGGGGCAACCTATCCAAGATCGGGATTGTGGGAACAGGAAGTGCCAGTTATAAGGTGATGACTAGGCCGAGCCAAC

AAACCCTGGTTATAAAGCTAATGCCAAATGTAACAGCCATAGACAATTGTACGAAATCAGAGATTTCGGAATACAAAAGA

TTGCTGATCACAGTGTTAAAACCTGTCGAAGATGCTCTGTCAGTGATAACCAAGAATGTAAGACCAATTCAAGCTCTAAC

ACCCGGTCGCAGGACCCGCCGTTTTGCCGGGGCTGTCCTGGCCGGAGTGGCACTTGGAGTCGCAACTGCTGCTCAGATAA

CTGCCGGAGTCGCACTTCATCAATCTTTGATGAACTCCCAAGCAATTGAAGGGTTAAAAGCTAGTCTTGAGAAGTCAAAT

CAGGCAATAGAAGAAATCAGACTTGCAAATAAGGAAACCATACTGGCAGTACAGGGTGTCCAAGATTATATCAACAATGA

GCTAGTCCCTTCTGTTCATAAAATGTCATGTGAGTTGGTAGGTCACAAACTCAGCCTCAAGCTTCTTAGGTATTACACCG

AGATCTTGTCTATATTCGGGCCCAGCCTTCGAGACCCGATAGCTGCTGAAATATCAATCCAAGCACTCAGTTATGCACTA

GGTGGAGATATCAATAAGATCCTGGATAAGCTTGGGTATAGCGGCGGGGACTTCCTTGCTATCCTAGAAAGTAAGGGGAT

AAAGGCCCGAGTCACCTATGTGGATACAAGAGATTACTTTATAATTCTTAGCATAGCTTACCCGACCTTATCTGAGATCA

AGGGGGTGATAGTTCATAAAATGGAAGCTGTGTCATATAACATCGGGGCACAAGAATGGTACACTACTATCCCTAAATAT

GTAGCCACTCAGGGATACCTGATATCAAATTTTGATGAGACGTCATGCGTGTTCACTCCAGGAGGGACAGTCTGCAGTCA

GAATGCTTTATACCCAATGAGCCCGTTGCTTCAGGAATGTTTCAGGGGGTCTACAAAATCGTGCGCTAGAACCCTAGTTT

CAGGGACCATAAGCAATCGATTTATCCTATCAAAAGGGAACTTGATTGCCAATTGTGCATCAGTCCTGTGCAAATGTTAC

ACAACGGAGACAATTATCAGCCAAGATCCTGATAAACTACTAACTGTTATAGCCTCCGATAAATGTCCTGTAGTTGAAGT

GGATGGAGTGACGATACAGGTCGGCAGTAGAGAGTACCCAGATTCTGTGTACCTACATAAAATAGACTTGGGCCCAGCCA

TCTCTCTAGAAAAACTGGACGTAGGCACCAATTTAGGCAATGCAGTTACAAGACTGGAGAATGCAAAGGAGCTCCTAGAT

GCATCAGATCAGATATTGAAGACTGTTAAGGGGGTACCCCTCAGCGGCAACTTGTACATAGGACTGGCAGCTTGCATTGG

GGTATCTCTAGGGCTTGTCACATTAATATGCTGCTGTAAGGGGAGGTGTAGGAACAAGGAGATTCCTACCCCTAAAATCA

ACCCAGGGCTCAAACCAGACCTGACAGGGACCTCAAAGTCGTACGTGAGATCACTGTAGTCAGAATAACCCGAATCATCC

ATCGTCACACACAGACATGTGCAACACAAGCAGTCAAGGGTCGTAGAAGATTCAACTTCCGATCATCAACCTGGCCTTAC

TCTACGCCCCATTGCACATTGGTTATCAAACAAAACTTAGGACGAAAGGTTAGTCACCATGTCCGCACAAAGAGAGAGGA

TCAATGCTTTCTACAAAGACACTCCTCACAACAAGAACCATAGGGTGACCCTGGATAAGGAACGTTTGACTATTGAAAGA

CCCTATATCCTGCTCGGGGTCCTACTGGTAATGTTTCTGAGCCTAATTGGGCTGCTGGCCATTGCAGGGATTAGACTTCA

CCGAGCCACCGTCGGAACTGCAGAGATCCAGAGCCGGCTTAATACCAATATTGAGTTAACTGAATCCATTGATCATCAAA

CTAAGGATGTCCTAACTCCCCTGTTTAAAATCATCGGTGATGAAGTCGGCATAAGGATTCCACAGAGGTTCAGTGACCTT

GTCAAGTTCATCTCCGATAAGATTAAATTCCTCAACCCTGACAAAGAATATGATTTCAGGGATCTCCGGTGGTGTATGAA

TCCCCCCGAGAGAGTTAAAATAGATTTTGATCAGTTTTGTGAATACAAAGCCGCGGGTAAGTCGATTGAACATATATTTG

AGTCACCACTCAACAGGTCAAAAAGGTTGCGATCGTTAACTCTAGGGCCTGGAACGGGCTGTCCTGGCAAGGCAGTAACA

AGGGCCCAGTTCTCAGAGCTTACTCTGACCCTGTTGGATCTGGATCTCAATATGAAGCACAACGTGTCCTCAGTGTTTAC

CATAATTGAAGAGGGATTATTCGGAAGAACATACACCGTCTGGAGATCAGATGCCGGAAATCTGAGCACCGGCTCAGGTA

TTGGCAATTTTCTGAGAGTCTTCGAGATCGGACTGGTAAGAGATTTCGGGCTAAGTGCCCCTGTTTTCCATATGACTAAC

TATCTCACAGTGAACATGAGTGATGACTATCGGAGCTGTCTATTAGCAGTAGGGGAGTTGAAGCTGGCAGCTCTATGCAC

TCGTTCTGAGACTGTGACTCTGAGTGATAGAGGAGTTCCTAAGAGAGAGCCTCTTGTGGTTGTAATACTTAATCTGGCCG

GGCCTACTCTAGGGGGCGAACTATATAGTATTTTGCCTGCCTCTGACCTCATGGTAGAGAAACTCTATTTGTCATCACAT

AGAGGGATTATCAAAGACAACGAGGCCAATTGGGTAGTACCGTCTACCGATGTTCGTGATCTTCAAACCAAAGGAGAATG

CTTGGTGGAAGCATGCAAAACTCGACCTCCTTCATTTTGCAATGGTACAGGGTTAGGCCCATGGTCAGAGGGGAGAATCC

CCGCCTACGGGGTGATCAGAGTCAGTCTTGACGTAGCTAGTGATCCAGATGTGGTTATTACCTCAGTATTTGGCCCCTTG

ATACCTCATCTATCTGGCATGGATCTTTACAACAACCCATTCTCAAGGGCTGTGTGGTTGGCTGTACCACCTTATGAGCA

GTCATTTCTGGGAATGATAAATACAATTGCCTTCCCAGATAGAGCAGAGGTTACGCCGCACATTCTGACCACAGAGATCA

GAGGGCTTCGGGGTCGTTGTCATGTTCCTATAGAATTGTCCCGCAGGATTGATGATGATATTAAGATTGGCTCCAACATG

GTTGTACTACCGACGAAGGACCTGAGGTACATTACAGCCACTTATGATGTTTCCAGGAGCGATCACGCCATTGTGTACTA

TATCTATGACACAGGTCGCTCATCATCTTACTTCTTCCCCGCCCGATTAAAGATCAAAGGCAACCCTCTCTCCCTGAGAA

TAGAGTGTTTCCCCTGGTATCATAAGGTGTGGTGCTACCATGACTGCCTCATATACAACACCATCACAAACGAAGAAGTC

CATACGAGAGGGCTAACCGGCATAGAGGTAACATGTAATCCAGTCTGAGCAGAGCTGCGACCATCAATCAAGCAGGCTTC

CGAGCCATTCCCAAGCCAAAGTAGCACAGCTAGGGGCACACAACAGCACAACTCAGCCGACAGTGTTATAAAAAACTTAG

GAGCCAGGGCTATAGGAGCCATGGACTCCCTGTCAGTCAATCAGGTCCTATACCCTGAGGTCCATTTAGACAGCCCCATT

GTCACAAACAAGCTAGTTGCCATCCTTGAGTATTCGGGCATCGACCACAACTATGTTCTTGAAGACCAGACCCTTATCAA

GAATATTAGATATAGACTGGGGTGCGGTTTTTCAAATCAAATGATCATCAATAACAGGGGGGTAGGTGAAACAGTCAATT

CCAAGCTTAAAAGCTACCCCCGTAGTTGTCATGTCATATATCCGGATTGCAATAAGGATCTGTTTTGTATCAAGGACGGC

TGCATATCTAGAAAGCTCTCTGAGCTATTCAAGAAGGGTAATTCCCTGTACTCCAAGATAAGTCACCAGGTACTAGATTG

TCTCAGGAGAGTCAACGGCAAAATGGGCCTGGGCACAGATCTTAACCATGGCCTGAAGGAGGGTATCCTCGATTTAGGGT

TGCACATGCATAGCTCTCAGTGGTTTGAGACTTTCCTGTTCTGGTTCACTATCAAGACTGAAATGAGATCCATGATCAAA

GAACAATCCCATATATGCCACAAGAGGAGGTATAACCCAACCTTTGTCTCAGGAGATGCATTCGAGGTGCTTGTATCACG

AGACCTCGTTGTGATAATTGACAAGAATACTCAGTGTGTCTATTACCTGACATTCGAGCTGGTCCTTATGTATTGTGATG

TCATAGAGGGCAGACTTATGACGGAGACAGCTATGGCTATAGACCAGAGGTATTCAGAACTGTTAAGCCGGGTCAAATAC

TTGTGGGATCTTATTGATGGGTTCTTCCCAACACTAGGTAACGCCACATACCAAGTCGTTGCTCTGCTTGAACCATTATC

GTTGGCTTACCTTCAACTTCAGGATGTCACTCTGGAGTTAAGAGGCGCATTTTTGGATCACTGCTTCAAAGAGCTCTATG

AGATACTGGAACATTGTGGCGTTGACACGGAAAGCACCTACGATTCCATCATCGAAGGATTGGATTACGTATTTATCACC

CACGATATCCACTTAACTGGGGAGATTTTCTCCTTTTTCCGGAGTTTCGGACACCCCCGCCTCGAAGCGGTCACCGCTGC

AGAGAATGTCAGAAAACACATGAATCAACCTAAGGTAATCAGTTATGAGACTATGATGAAAGGGCACGCAGTATTTTGCG

GAATCATCATAAATGGGTTTAGGGACCGCCATGGCGGCAGCTGGCCCCCCGTTGTGCTGCCAGAACATGCCTCTGCTGCA

ATTCGGAATGCGCAGGCATCCGGTGAGGGGCTGACCCATGACCTGTGTATAGACAACTGGAAGTCCTTCGTTGGATTCAG

ATTTGGCTGTTTTATGCCGCTCAGCCTAGATAGTGACCTGACCATGTACCTCAAAGACAAAGCACTAGCTGCACTGAAAA

ATGAGTGGGATTCAGTCTACCCGAAAGAATACCTCCGCTATAACCCACCTAGAGGGACAGAGTCAAGGCGACTGGTGGAG

GTGTTCCTGAATGACTCCAGCTTTGATCCTTATAACATGATAATGTACGTGGTGAACGGTTCCTACCTTGAAGACCACGA

GTTTAATCTCTCATACAGCTTGAAAGAGAAAGAGATAAAAGAGACAGGGCGATTGTTTGCAAAAATGACTTACAAAATGC

GGGCCTGCCAGGTCATTGCGGAAAATCTGATATCAAATGGGGTAGGGAAGTATTTCCGAGACAATGGAATGGCGAAAGAT

GAGCATGACCTAACAAAGGCTCTTCATACCCTGGCAGTCTCAGGCGTTCCCAAAAACAACAAAGATAATCACCGGGGTGG

GCCTCCCAGAAGGACCACAAGCCAAAAGATGAGGTCAGATCGAGGCATCACCAGACAAGATAGGGACAAGGTACAGGGAG

AGCCCATGTATAACTATCTGCGATGCCAACCAGTCAGCCCTGATCAAGGTGAGTCATACGAGACTGTTAGTGCATTCATC

ACCGCTGATCTTAAGAAATATTGCCTGAACTGGAGATACGAGACAATCAGCATATTTGCACAGAGACTGAATGAAATTTA

CGGACTACCATCTTTCTTTCAGTGGTTACACAGGGTATTAGAAAAATCTGTGCTGTACGTCAGTGACCCGCATTGCCCTC

CCGACTTAGATGATCACATCCCTCTGGACAGTGTCCCCAATGCCCAAATATTCATCAAGTACCCAATGGGCGGAATAGAA

GGTTATTGCCAAAAATTATGGACAATCAGTACTATACCGTACTTGTATCTAGCAGCCCATGAGAGCGGAGTAAGAATCGC

CTCTCTAGTGCAAGGTGACAATCAAACAATTGCAGTGACAAAAAGAGTCCCGAGTTCTTGGCCTTATTCACTAAAAAAGA

GGGAGGCATCCAGAGCAGCTCAAAATTACTTCGTGGTTTTAAGGCAGAGATTGCACGATGTAGGTCATCACTTAAAGGCC

AATGAAACCATTGTATCTTCCCACTTTTTTGTATACTCCAAAGGGATTTATTATGATGGCCTGTTAGTCTCACAGTCACT

AAAGAGCATTGCCAGATGTGTCTTCTGGTCCGAGACTATTGTGGATGAAACCAGGGCCGCCTGCAGCAATATTGCAACAA

CTGTCGCCAAGAGTATAGAGAGGGGTTATGATAGGTACCTTGCTTACTCTTTGAATATACTCAAAATTCTCCAACAGATC

CTTATATCCCTTAACTTCACTATCAATACAACAATGACTCAAGATGTCGTGGCACCGATCATCGAGAACGGTGATCTACT

AACAAGGATGGCACTTTTGCCAGCACCCATCGGGGGCCTTAATTACCTTAACATGAGCAGGTTATTTGTGAGAAATATCG

GTGACCCAGTCACTTCCTCTATAGCCGACCTGAAGAGGATGATAGACGCTGGGCTAATGCCAGAAGAAACATTGCACCAG

GTGATGACCCAAACCCCGGGAGAATCATCTTACCTTGATTGGGCAAGTGATCCTTATTCTGCCAACCTAACTTGTGTACA

GAGTATAACTCGCCTTCTCAAGAACATCACTGCAAGGTATATTTTAACCAGCAGCCCAAATCCGATGCTAAGAGGGTTAT

TCCATGAAGGGAGCAGAGATGAAGATGAAGAGCTTGCGAGTTTCTTGATGGATCGGCATATAATTGTTCCCAGAGCTGCA

CATGAAATCTTAGACCATAGCATAACTGGAGCAAGAGAAGCTATTGCCGGTATGTTAGATACCACCAAGGGTCTGATTAG

AACAAGTATGAAACGAGGTGGCCTTACCCCCCGGGTCTTAGCTCGCCTTTCCAATTATGATTATGAACAATTCAGATCCG

GTATAACATTATTGACGAAGAAGGGACAGTGTTATCTCATTGACAAAGATTCGTGCTCGGTGCAGCTTGCTATAGCCCTG

AGGAGCCATATGTGGGCTAGGTTGGCCCGCGGGAGACCTATCTATGGGTTGGAGGTGCCTGATGTACTAGAATCGATGAA

TGGCTATCTTATCAAACGCCACGAAGCTTGTGCCATTTGCGAAACAGGCTCAAGTCATTACGGGTGGTTCTTCGTCCCTG

CAGGGTGCCAGCTTGACGATGTCTCGAAAGAGACTTCAGCTCTTCGTGTGCCTTATGTCGGGTCAACCACAGAGGAAAGG

ACAGATATGAAACTTGCCTTCGTTCGATCTCCGAGTCGATCCCTCAAATCAGCAGTCAGAATTGCTACCGTTTATTCATG

GGCCTACGGGGATGATGAAAAATCATGGAGTGAAGCCTGGATGCTGGCTAGGCAGAGAGCCGATATCACCCTCGATGAGT

TGAGAATGATCACTCCAATCTCTACATCGACCAACCTAGCCCATCGGTTGAGGGATCGGAGCACCCAGGTGAAATACTCG

GGGACATCCCTTGTGAGGGTTGCGAGATACACAACCATTTCCAATGATAATTTGTCATTTGTGATATCTGAGAAGAAGGT

AGATACCAACTTCATTTACCAGCAAGGGATGCTCCTCGGTCTTGGGATCCTTGAGAATCTTTTCAGGTTGGAGGCTACCA

CAGGGGTATCCAACACAGTGCTACATCTGCACGTGGAGACAGAATGCTGCGTTGTACCTATGGTTGACCACCCAAGGATA

CCGAGTCTCCGTAACATTACGGTTACGAATGAGCTATGCACAAACCCCTTGATCTACGACAAGTCTCCTATCATAGAACA

CGATGCAACCCGATTATACTCACAAAGCCACAGAAGACATCTGGTGGAGTTTGTTACCTGGTCAACAAGTCAGCTTTACC

ATATACTGGCTAAGTCTACAGCGATGTCCATGATTGAGTTGATTACAAGATTCGAGAAGGATCACATGAATGAAATAACA

GCCTTGATTGGCGATGACGACATCAACAGTTTCATCACAGAATTCCTGTTAGTAGAACCTAGACTGTTTATAGTCTACCT

TGGCCAGTGTGCTGCCATCAATTGGGCTTTTGATATACATTATCATCGCCCATCGGGCAAATACCAGATGGGAGAGCTTC

TCTACTCTTTGCTCTCTAGGATGAGCAAAGGAGTATACAAGATCTTTACCAATGCTTTGAGCCACCCCAAGGTTTACAAG

AAATTTTGGAGAAGTGGTGTCATCGAGCCGATTCATGGCCCGTCCTTAGATACACAGAATCTACATGTCACTGTCTGTGA

CATGATATATGGATCCTACGTCACCTATCTGGATCTTTTACTGAATGATGAGCTAGATGATTATACGTATCTGCTCTGCG

AGAGTGATGAGGACGTGGTCACAGACAGGTTCGATAACATTCAAGCCAAACATCTCTGTGTATTAGCCGATGTTTATTGC

AGCTCCAAAAAGTGTCCCTCGATAATCGGGATGTCTCCTATAGAAAAATGTACCATCCTCACACATTACATCAAGGGAGA

ATCAATACAATCCCCGTCTGGGACCTCATGGAACACTGAACCTCTTGTAGTAGATCATTACTCATGCTCTCTGACATACC

TTCGTCGCGGTTCCATCAAACAAATCAGGTTGAGAGTAGATCCTGGATTTGTATTCGAGGCACTGACAGACATCGACACC

AAACTGCCTCGAAAGGCTAATACTGATGTATCGGTTGTAGGGTTGACTGATTTCTCCCCTCCTTGGGATAACATTGGTGA

TTTTCTAAGGACTATCAACACCTTGAGACACAATCTGCCTGTCACCGGGACCGGGGTCTCTAACTATGAGGTCCACGCTT

ATCGTAGGATTGGTTTGAATTCATCAGCATGTTATAAAGCTGTAGAAATCTCCACGTTAATCAAACCATCCTTAGAAGTC

GGAGAGCATGGGTTGTTCTTAGGGGAAGGTTCCGGTTCTATGCTGGCTGCGTATAAGGAAGTTCTTAAATTAGCCAACTG

TTATTACAACAGCGGAGTAATAGCTGAGGGGAGAGCCGGACAGAGAGAAATATCCCCTTACCCTTCAGAGATGAGCCTAG

TAGAGAACCAAATGGGGTTAGAAAGGAGTGTTAAAGTGCTGTTCAACGGCAAACCTGAAGTAACTTGGGTCGGGACCACC

GATTGCTACGAATATATCATCAGTAACATCCAAACCTCTAGTCTGGGTTTCATACACTCAGATATTGAGACACTCCCAAC

CAAAGATGCTGTTGAGAAATTAGAAGAATTTGCCTCTATCTTATCCTTATCCCTAATTTTGGGAAAAATTGGCTCTATTA

CCGTTATCAAAGTTATGCCTATTAGCGGAGATTTTACTCAAGGCTTCATAAGTTATGCCATCCAATATTTTAGGGAGAGC

CTGCTTGTCTATCCGAGATATAGTAATTTCATTTCAACTGAGTGTTACCTTATTATGATAGGATTGAAAGCCAATCGGTT

GATAAACCCAGAAGCCATTAAGCAAAACATAATCAGAGCGGGGACTAGAACTGCACCAGGACTAGTGAGCCATATATTAT

CAGAGAAACAAAAAGGGTGTATCCAATCTTATCTGGGTGATCCTTATATCCAAGGAGACTTCAATAAACACCTTAAATCT

CTAACCCCTATTGAGAAAATCCTAGTTAATTGTGGTCTCTCGATCAATGGCACAAAAATCTGTAGGGATCTAATCCACCA

TGATATCGCCTCTGGTCCAGACGGTCTGATGAGCTCCACAATTATTTTGTATAGGGAGCTGGCACATTTCAAAGACAACA

TAAGAAGTCAGCACGGGATGTTCCACCCCTACCCAGTACTGGCTAGTAGCAGGCAACGTGAATTAATCCTTCGAATCGCC

AAGAAATTCTGGGGTTATGTCTTGCTATATTCTGATGATCCGGCACTGATCAGGCAAACAATTAAGAACTTGAAGAGAAA

CCACCTAACCTTTGACTTACACAGTAACCCGTTTATTAAGGGTCTGTCCAGAGCTGAGAAAATGCTAGTGCGGACAAGTT

CACTCAGAAGAGAGTGGTTGTTCACTCTCGAAACGAAGGAAGTGAAAGAGTGGTTCAAATTGGTGGGTTACAGTGCTCTC

GTCAGAGGTTAATTAGCTATACATCTGCCCCCTTCTCCTCCACCATGAAATTCTACTGGCAATCTAATAGATTAAAGAAA

ACTACAGATCGGATAAGTATCTATTCCCAGCTTTGTCTGGT

>KJ867540.1/goat/Ethiopia/1994

ACCAGACAAAGCTGGGTAAGGATAGATCTTATAATAACTATGAATTGCCACGCTTAGGATTAAAGATCCTACTGCCGGGG

GGAGGAGGAGGAGCAAGATCCCTGACTATGGCGACTCTCCTCAAAAGCTTGGCATTGTTCAAGAAGAACAAAGACAAAGC

GCCGACGGCATCAGGTTCAGGAGGGGCCATCCGGGGGATTAAGAATGTTATCATAGTCCCGATTCCCGGAGACTCATCCA

TCATTACCCGTTCAAGATTGCTCGACAGGCTTGTCAGATTGGCCGGAGATCCAGATATCAACGGGTCAAAGCTGACCGGC

GTGATGATCAGCATGCTTTCTTTGTTCGTAGAGTCACCCGGGCAACTGATCCAGCGGATCACAGATGATCCAGATGTCAG

TATCCGCCTTGTTGAGGTGGTCCAAAGTACTAGATCTCAGTCCGGGTTGACCTTTGCATCACGTGGTGCTGATTTGGACA

ATGAGGCAGACATGTATTTCTCAACTGAAGGGTCCTCGAGTGGGAGCAAGAAAAGGATCAACTGGTTTGAGAACAGGGAA

ATAATAGACATAGAAGTGCAGGATGCGGAAGAGTTCAATATGTTATTAGCCTCCATACTGGCACAAGTTTGGATTCTCCT

GGCCAAAGCGGTTACGGCACCGGACACTGCAGCTGACTCAGAATTGAGAAGGTGGGTTAAATACACACAACAAAGAAGAG

TGATTGGGGAATTTCGCCTTGACAAAGGGTGGCTAGATGCGGTCCGCAACAGGATTGCAGAAGATCTATCACTCCGGCGG

TTCATGGTATCTCTTATACTTGACATCAAAAGGACCCCTGGCAACAAGCCAAGGATTGCAGAAATGATCTGTGACATTGA

TAACTATATTGTCGAAGCCGGGCTCGCCAGTTTCATCCTTACCATCAAGTTTGGTATTGAAACCATGTATCCTGCACTAG

GTCTTCACGAGTTTGCCGGGGAGTTGTCCACTATAGAATCCTTGATGAATCTGTATCAACAGCTAGGCGAAGTTGCACCC

TACATGGTAATTCTAGAGAACTCAGTTCAGAACAAGTTTAGTGCAGGAGCCTATCCTCTTCTCTGGAGCTATGCGATGGG

TGTTGGAGTCGAGCTGGAGAACTCAATGGGGGGGTTGAACTTTGGTAGATCATATTTTGACCCGGCTTATTTTCGTCTCG

GACAGGAGATGGTCAGAAGATCCGCAGGAAAGGTCAGCTCTGTAATCGCAGCTGAGCTCGGCATCACAGCAGAGGAAGCT

AAACTAGTCTCGGAAATCGCCTCTCAGACTGGGGACGAAAGGACCGCTAGAGGGACCGGGCCCAGACAGGCGCAGGTTTC

CTTCCTCCAGCATAAAATAGGAGAGGGAGAGTCACATGCATCGGCGACCAGGGAAGAAGTCAAAGCTGCGACCCCAAATG

GGCCCGACGAAAAGGACAAAAAACGAGCACGCTCAGGAAGGCCAAGAGGAGGAACCCCCGACCAACTGCTCCTGGAAATC

ATGCCTGAAGACGAGGTCCCGCGAGGGTCTGGACAAAACCCTCGTGAGGCTCAACGATCGGCCGAGGCACTCTTTAGACT

GCAGGCCATGGCCAAGATTCTAGAGGGCCAGGAGGAGGGAGAAGACAACAGTCAGATATATAACGACAAGGATCTCCTCA

GCTGAGCAGAAGCGCCATCCATCGAAACCAGCGACAAGACACCGCCCATCAGTATTATAAAAAACTTAGGACCCAGGTCC

AAGCGGTCACACCCCGGCACCCCAGCCGATCTAGCGGAGACCACCGATGGCAGAGGAACTAGCTTACCATGTCGTGGAGG

GACTGGCATGTCTCAAGTCCCTCTTTGCCTCTCCCCCGGATCTTTCCCCCATCCGAGATACCCTTGAAAACTGGAGAGAG

GGGCTTGACCCATCCCACCGTGGAACCCCGAACCCTGATATGTCCGAGGGGAACCATCAGAATATCAACCAATCATGCCC

ACCAGCCATCGGATCAGGCGAAATCGACTTGTCTACTGAAGGTAATCTCGGATATAGAGAGATCAATTACGATGACAGTG

AGGCTGGACTCAGAGGAGTTCAGGTCTACGGATCCCACCCTCAAATACAGCGTTACCATGTTTATAGCCACGGGCGTGAA

GAGATTGAAGGACTCCAGGATGCTGACTCTCTCATGGGTCAAGCAGATCCACCACTTGCTAACACCTTCAGTACAGGAGA

GGATGGATCTGACGACAGCGATGTGGACTCTGGCCCAGATGATCCCGACAGACATCCTCTATATGACCGGGGATCTGTTG

CCGACAATGATGACGTTAAGTCCACAGATGTCGAAAAGTTAGAAGGTGACGACATTCAAGAAGTTCTTAACTCTCAGAAG

AGTAAAAGAGGAAGATTCCAAGGCGGGAAAACCTTGCGGGTCCCTGAAACACCCGACGTCAAGCACCCCAGACCATCAGC

CCAATCAATTAAAAAGGGCACAGACGGGAACTCAGTCTTATCTGGAACGGTGACAGAGTGTTCATCGATAAGTGGTGCAA

CCCAAGCTGTGCCAGAGTCAAGATGCGAGTCATCAGAGAGAAATGCATTTGTGGAGAGTGTCCCCAAATCTGCGAGGAGT

GCAAAGACGATCCAGGAGTTGACACAAGAATCTGGTACCATAGCATCACCGACTCAGCCTAAAGAAAATGACTCCGAGTA

TGAGTATGAGGATGACCTATTTACAGAAATGCAGGACATTCGTGCAAGCATTGCCAAGATCCATGAAGACAACAAAACTA

TCCTCTCAAAGCTTGACTCTATACTATTATTGAAAGGAGAAATCGATACTATCAAGAAGCAAATCAGCAAACAGAATATA

AGTATATCTACCATTGAGGGCCATCTTTCCAGTATAATGATAGCCATCCCGGGTTTTGGAAAGGAGGTCAAGGACCCAAC

ATCCGAGGTTGAGTTGAACCCGGATTTAAGACCTATAATCAGCCGTGATTCTGGCAGGGCTCTCGCGGAGGTCCTCAAGA

AACCTGCTGTTGATAGGTCTCCCAAAACCGGATTCAAAGTCAACTCTGGTTCAAAGGGTCAGCTCCTCAAGGATCTCCAA

CTAAAACCCGTTGACAAACAGGCAAGCTCTGCCATAGGGTTTGTCCCGTCTGACCATGAATCATCCAAAAGTGTCATCCG

CTCTATAATTAAGTCAAGCAAGCTCAACATTGAACACAAAGACTATCTTTTAGATTTGGTGAATGATGTGAAAGGCTCCA

AGGATCGTAAAGAATTCCACAAGATGCTAACAGCTATTCTTGCCAAGCAACCGTAACACCGCCCCTAATCAACTTCTCAT

GCTCGGCTGAAAACAGCCCCACAACCAGGCTATTACAAAAAACTTAGGAGCAAGGGCAACCGAGCTTCGCAGACAAGATG

ACCGAGATCTATGACTTTGACAAATCAGCATGGGACGTCAAAGGATCAATAGCTCCCATAGAGCCTACCACTTATCACGA

TGGTCGACTGATACCCCAGGTGAGAGTCATCGATCCTGGGCTAGGAGATAGGAAAGATGAGTGCTTCATGTACTTGTTTC

TCCTAGGAGTGATGGAGGATAATGACCCCCTGTCTCCCCCAGTGGGAAGAACCTTTGGTTCTTTACCTCTAGGGGTCGGT

AGGTCAACTGCTAAGCCAGAAGAACTATTAAGGGAGGCCACTGAACTTGACATAGTGGTGAGACGCACGGCTGGACTAAA

CGAGAAGCTGGTATTTTACAACAACACTCCGCTGTCTTTGTTAACACCTTGGAGGAGAGTCCTGACAACCGGAAGTGTGT

TTAGCGCTAACCAGGTTTGTAACGCAGTCAATCTAGTCCCACTCGATACTCCCCAGAGATTCAGGGTTGTGTACATGACC

ATCACTAGATTGTCTGACAATGGTTATTATAGTGTCCCCGGGAGAATGTTAGAGTTCCGCTCAGCCAATGCCGTCGCCTT

CAATATCTTGGTTACACTGAGAATTGAGAATGGGACAAACCCTAGAAGGTACATAGTCGGCTCATGGGAGAATCCAGAGG

TCACATTCATGATACATGTGGGCAACTTTAGAAGAAAGAAGAACGACGTATATTCTGCTGATTACTGCAAAATGAAGATT

GAGAAGATGGGTCTGGTTTTTGCTTTAGGAGGAATAGGGGGAACAAGTCTTCACATTAGAAGCACCGGGAAAATGAGCAA

GACCCTCCATGCACAGCTGGGGTTCAAGAAAATCTTATGCTACCCCTTAATGGATATTAATGAGGATCTCAATCGATATC

TTTGGCGGGCAGAGTGCCGAATAGTCAAAATTCAAGCTGTCTTACAGCCTTCAGTACCCCAAGAATTCCGTGTCTACGAT

GATGTCATCATTAACGATGACCAAGGCTTGTTCAAGATCCTGTAATTCACTTACAGAACAATGGACCCGCCTCCAATGAA

AGGCCACCCCTTCTCTCGTCCAGAAAACACACCCCGAGCCCCCCCAAAAAAACGCCGACATGGTCATCCCCCCGGTAGGA

CGAGGACGACCCCCCCAGCGGACCCGA-TCTGT-GGGGGGGCCCCCCCCGCGACCCGCTGAGCACCGACCGCGCGGGGCA

ACAAGACCCTCCTGGGAGGCCCCAAGGTAGACCGGGCACCCCCCCCCCCGCGT-CGGC-CCACA-AACACGCCCGAGGCC

AGGAGCCGAGCACCCCCAGCCCTACCACCGACAGGACAGACACCCAAGCCAGGGTCCCTCGAACCAAAGAGACCACCAGG

GTGGGCTCCGCCGACCCAGCACAGACTTGGCCCCAACAAGGCAGATCCTCAAAACGAAATCGCCCAGTGCACTCCACTCA

TCATCAAACAAAACTTAGGGGCCAAGGTCAGACGACAGACCCCATCCCGCCAAGCAGCAGGCCGCACGAAGGAAGCAGAC

AAGACAGGCTCGGAAGGCAGACACCCCACCACAGAAACACCCAGGCCTCGGGGAGGAGGCCCCTCCTCCACAAGGCGGAG

GAGCGGGAGGGAAACACCAGGGACAACCCGACCGGGAGCCAGGACCCGGAGGTTCCAGCGGCAACCGGGCCGGGCACACA

CCCGACACCCCCGGCGGAGCAGAGGAGAGCCCGACCCCGCGTGGACCCGACGGCAGAAGATCCAACCCTGCCACAAAGAG

AAGCGAGGAGATC--------CTCCACCGCACATAGCCGGCAGACAACCCATAGGGGGACCACCTCCCGGGAGGACAAGC

GGCCCTCCCACAACTGAGCCGCCACCCCCCAGAGGAGGACACCCCGGAAGCCCCAAGCCATCCCCCCGCCCTCGACCACC

GGGGGAGGGGGTACACCGATAAAGACTAGAGCCACCGGGAGCCAACTAAGGGTCCCCCCCGGCCCCGCCCGGCGCCCCCC

CGACGAGGCCCTAACAGGACCCCCCGGCCCAACAACCCCCGCACGCGCCAAGACAGAAAGAGAGCAGAGGTTATCCAGCA

CATTCATGCACAAACATCATGACAAAGGTCGCAATCTTGACATCTCTGTTTCTCTTACCAAACACTGTCGCATGCCAGAT

TCACTGGGGCAACCTATCCAAGATCGGGATTGTGGGAACAGGAAGTGCCAGTTATAAGGTGATGACTAGGCCGAGCCAAC

AAACCCTGGTTATAAAGCTAATGCCAAATATAACAGCCATCGACAATTGTACGAAATCAGAGATTTCGAAATACAAAAGA

TTGCTGATCACAGTGTTAAAACCTGTAGAGGATGCCCTGTCAGTGATAACCAAGAATGTAAGACCAATTCAAGCTCTAAC

ACCCGGTCGCAGGACTCGCCGTTTTGCCGGGGCTGTCCTGGCCGGAGTGGCACTTGGAGTCGCAACGGCTGCTCAAATAA

CTGCCGGAGTCGCACTCCATCAATCTTTGATGAACTCCCAAGCAATTGAAAGTTTAAAAGCCAGTCTTGAGAAGTCGAAC

CAGGCAATAGAAGAAATCAGACTTGCAAATAAGGAAACCATACTGGCAGTACAGGGTGTCCAAGATTATATCAACAATGA

GCTTATCCCTTCTGTTCATAAGATGTCATGTGAGCTTGTAGGTCACACACTCAGTCTCAAGCTTCTTAGGTATTATACCG

AGATCTTGTCTATATTCGGGCCCAGCCTTCGAGACCCGATAGCTGCTGAAATATCAATCCAGGCACTCAGTTATGCACTA

GGTGGAGATATCAATAAGATCCTGGATAAGCTTGGGTATAGCGGCGAGGACTTCCTTGCTATCCTAGAAAGCAAGGGGAT

AAAGGCCCGAGTCACCTATGTGGATACAAGAGATTACTTTATAATTCTTAGCATAGCTTACCCGACCTTATCTGAGATCA

AGGGGGTGATAGTTCATAAGATGGAAGCTGTGTCATACAACATCGGGGCACAAGAATGGTACACTACTATCCCTAAATAT

GTAGCCACTCAGGGATATTTGATATCAAATTTTGACGAGACGTCATGCGTGTTCACTCCAGGGGGGACAGTCTGCAGTCA

GAATGCTTTATACCCAATGAGCCCGTTGCTTCAGGAATGTTTCAGGGGGTCTACAAAATCGTGCGCTAGAACCCTAGTTT

CAGGGACCATAAGTAATCGATTTATCCTATCAAAAGGGAACTTGATTGCCAATTGTGCATCAGTCCTGTGCAAATGTTAC

ACAACGGAGACAATTATCAGCCAAGATCCTGATAAATTACTAACTGTTATAGCCTCTGATAAATGTCCTGTAGTTGAAGT

GGATGGAGTGACGATACAGGTCGGCAGTAGGGAGTACCCAGATTCTGTGTACCTACATAAAATAGACTTGGGCCCAGCCA

TCTCTCTAGAAAAACTGGATGTAGGCACCAATTTAGGCAATGCAATTACAAGACTGGAGAATGCAAAGGAGCTCCTAGAT

GCATCAGATCAGATATTGAAGACTGTTAAGGGGGTACCCTTCAGCGGCAATTTGTACATAGGACTGGCAGCTTGCATTGG

GGTATCTCTAGGGCTTGTCACATTAATATGCTGCTGTAGGGGGAGGTGTAGGAACAAGGAGATTCCTACCCCTAAAATCA

ACCCAGGGCTCAAACCAGACCTGACCGGGACCTCAAAGTCGTACGTGAGATCACTGTAGTCAGAATAACCCGAATCATCC

AGCGTCACACACAGACATGTGCAACACAAGCAGTCAGGGGACGTAGAAGATTCAACTTCCGATCATCAACCTGGTCTTAC

TCTACGCCCCATTGCACATTGGTTATCAAACAAAACTTAGGACGAAAGGTTAGTCACCATGTCCGCACAAAGAGAGAGGA

TCAATGCTTTCTACAAAGACACTCCTCACAACAAGAACCATAGGGTGACCCTGGATAAGGAACGTTTGACTATTGAAAGA

CCCTATATCCTGCTCGGGGTCCTACTGGTAATGTTTCTGAGTCTAATTGGGTTGCTGGCCATTGCAGGGATTAGACTTCA

CCGAGCCACCGTCGGAACTGCAGAGATCCAGGGCCGGCTTAATACCAATATTGAGTTAACTGAATCTATTGATCATCAAA

CTAAGGATGTCCTAACTCCCCTGTTTAAAATCATCGGTGATGAAGTCGGCATAAGGATTCCACAGAGGTTCAGTGACCTT

GTCAAGTTCATCTCCGATAAGATTAAATTCCTCAACCCTGACAGAGAATATGATTTCAGGGATCTCCGGTGGTGTATGAA

TCCCCCCGAGAGAGTTAAAATAGATTTTGATCAGTTTTGTGAATACAAAGCCGCGGATAAGTCGATTGAACATATATTTG

AGTCACCACTCAACAGGTCAAAAAAGTTGCGATCGTTGACTCTTGGGCCTGGAGCGGGCTGTCCCGGCAGGGCAGTAACA

AGGGCCCAGTTCTCAGAGCTTACTCTGACCCTGTTGGATCTGGATCTCAATATGAAGCACAACGTGACCTCAGTGTTTAC

CATAGTTGAAGAGGGATTATTCGGAAGAACATACACCGTCTGGAGATCAGATGCCGGAAATCTGAGCACCCGACCAGGTA

TCGGCCATTTTCTAAGAGTCTTCGAGATCGGACTGGTAAGAGATTTCGGGCTAAGTGCCCCTGTTTCCCATATGACTAAC

TATCTCACAGTGAACATGAGTGATGACTATCGGAGCTGTCTATTAGCAGTGGGGGAGTTGAAGCTGACAGCTCTATGCAC

CCATTCTGAGACTGTGACTCTGAGTGAGAGAGGAGTTCCTAAGAGAGAGCCTCTTGCGGTTGTAATACTTAATCTAGCTG

GGCCTACTCTAGGGGGCGAACTATACAGTATTTTGCCTGCCTCTGACCTCATGGTAGAGAAACTCTATTTCTCATCACAT

AGAGGGATTATCAAAGACAACGAGGCCAATTGGGTAGTACCGTCTACCGATGTACGTGATCTTCAAAACAAAGGAGAATG

CTTGGTGGAAGCATGCAAAACTCGACCTCCTTCATTTTGCAATGGTACAGGGTTAGGCCCATGGTCAGAGGGGAGAATCC

CTGCCTACGGGGTGATCAGAGTCAGTCTTGACGTAGCTAGTGATCCAGATGTGGTTATTACCTCAGTATTTGGCCCCTTG

ATACCTCATCTATCTGGCATGGATCTTTACAACAACCCATTCTCAAGAGCCGTGTGGTTGGCTGTACCACCTTATGAGCA

GTCATTTCTGGGAATGATAAATACAATTGCATTCCCAAATAGAGCAGAGGTTATGCCGCACATTCTGACCACAGAGATCA

GAGGGCTTCGGGGTCGTTGTCATGTTCCCATAGAGTTGTCCCGCAGGATTGATGATGATATTAAGATTGGCTCCAACATG

GTTGTACTACCGACGAAGGACCTGAGGTACATTACAGCCACTTATGATGTTTCCAGGAGCGATCATGCAATCGTGTACTA

TATCTATGACACAGGTCGCTCATCATCTTACTTCTTCCCCGCTCGATTAAAGCTCAAAGGCAACCCTCTCTCTCTGAGGA

TAGAGTGTTTCCCCTGGTATCATAAGGTGTGGTGCTACCATGACTGCCTCATATACAACACCACCACAAACGAAGAAGTC

CATACGAGAGGGCTAACCGGCATAGAGGTAACATGTAATCCAGTCTGAGCAGAGCTGCGACCATCAATCAAGCAAGCTTC

CGAGCCATTCCCAAGCCAAAGCAGCACAGCTAGGGGCACTCAACAGCACAACTCAGCCGACAGTGTTATAAAAAACTTAG

GAGCCAGGGCTATAGGAGCCATGGACTCCCTGTCAGTCAATCAAGTCCTATACCCTGAGGTCCATTTAGACAGCCCCATT

GTCACAAACAAACTAGTTGCCATCCTTGAATATTCGGGCATCGACCACAACTATGTTCTTGAGGACCAGACTCTTATCAC

GAATATTAGATATAGACTGGGGTGCGGTTTTTCAAATCAAATGATCATCAATAACAGGGGGGTAGGTGAAACAGTCAATT

CCAAACTTAAAAGTTACCCCCGTAGTTGTCATGTCATATATCCGGATTGCAATAAAGATCTGTTTTGTATCAGGGACGGC

TGCATATCTAGGAAGCTCTCTGAGCTATTCAAGAAGGGTAATTCCCTGTACTCTAAGATAAGTCACCAGGTACTAGATTG

TCTCAGAAGAGTCAACGGCAAAATGGGCCTGGGCACAGATCTTAGCCATGGCCTGAAGGAGGGTATCCTCGATTTAGGGT

TGCACATGCATAGCTCTCAGTGGTTTGAGACCTTCCTATTCTGGTTCACTATCAAGACTGAAATGAGATCAATGATCAAA

GAACAATCCCATATATGCCACAGGAGGAGGTATAACCCAACCTTTGTCTCAGGAGATGCATTCGAGGTGCTTGTATCACG

AGACCTCGTTGTGATAATTGACAAGAATACTCAGTGTGTCTATTACCTGACATTCGAGCTGGTCCTTATGTATTGTGATG

TCATAGAGGGCAGGCTTATGACGGAGACCGCTATGGCTATAGACCAGAGGTATTCAGAACTGCTAAGCCGGGTCAAATAC

TTGTGGGATCTTATTGATGGGTTCTTCCCAACACTAGGTAACGCCACATACCAAGTCGTTGCTCTGCTTGAACCATTATC

GTTGGCTTACCTTCAACTTCAGGATGTCACTCTGGAGTTAAGAGGTGCATTCTTGGATCACTGCTTCAAAGAACTCTATG

AGATACTGGAACATTGTGGCGTTGACACGGAAAGCACCTACGATTCCATCATCGAAGGATTGGATTACGTATTTATCACC

CACGATATCCACTTAACTGGGGAGATTTTCTCCTTTTTCCGGAGTTTCGGACACCCCCGCCTCGAAGCGGTCACCGCTGC

AGAGAATGTCAGAAAACATATGAATCAACCTAAGGTAATCAGTTATGAGACTATGATGAAAGGGCACGCAGTGTTTTGCG

GAATCATCATAAATGGGTTTAGGGACCGCCATGGCGGCAGCTGGCCCCCCGTTGTGCTGCCAGAACATGCCTCTGCTGCA

ATTCGGAATGCGCAGGCATCCGGTGAGGGACTGACCCATGACCTGTGTATAGACAACTGGAAGTCCTTCGTTGGATTCAG

GTTTGGCTGTTTTATGCCGCTCAGCCTAGATAGTGACTTGACCATGTACCTCAAAGACAAAGCACTAGCTGCACTGAAAA

ATGAGTGGGATTCAGTTTATCCGAAAGAATACCTCCGCTATAACCCACCTAGAGGGACAGAGTCAAGGCGACTGGTAGAG

GTGTTCCTGAATGACTCCAGCTTTGATCCTTATAACATGATAATGTACGTGGTGAATGGATCCTACCTTGAAGACCATGA

GTTTAATCTCTCATACAGCTTGAAAGAGAAAGAGATAAAAGAGACAGGGCGATTGTTTGCAAAAATGACTTACAAAATGC

GGGCCTGCCAGGTCATTGCGGAAAATCTGATATCAAATGGAGTAGGGAAGTATTTCCGAGACAATGGAATGGCGAAAGAT

GAGCATGACCTAACAAAGGCCCTTCATACCTTGGCAGTCTCAGGCGTTCCCAAAAACAACAAAGATAATCACCGGGGTGG

GCCTCCTAGAAGGACCACAAGCCAAAAGATGAGGTCAGATCGAGGCATCACCAGACAAGATAGGGACAAGGTACAGGAAG

AGCCCATGTATAACTATCTGCGATGCCAACCAGTCAGCCCTGATCAGGGTGAGTCATACGAGACTGTTAGTGCATTCATC

ACCGCTGACCTTAAGAAATATTGCCTGAACTGGAGATACGAGACAATCAGCATATTTGCACAGAGACTGAATGAGATTTA

CGGACTACCATCTTTCTTTCAGTGGTTACACAGGGTATTAGAAAAATCTGTGCTGTACGTCAGTGACCCACATTGCCCTC

CCGACTTAGATGATCATATCCCTCTGGACAGTGTCCCCAATGCCCAAATATTCATTAAGTACCCAATGGGCGGAATAGAA

GGTTATTGTCAAAAATTATGGACAATCAGTACTATACCATACTTGTATCTAGCAGCCCATGAGAGCGGAGTAAGAATCGC

CTCTCTAGTGCAAGGTGACAATCAAACAATTGCAGTGACAAAAAGAGTCCCGAGTTCTTGGCCTTATTCACTAAAAAAGA

GGGAGGCATCCAGAGCAGCTCAAAATTACTTCGTGGTTTTAAGACAGAGGTTGCACGATGTAGGTCATCACTTAAAGGCC

AATGAAACTATAGTATCTTCCCACTTTTTTGTATACTCTAAAGGGATTTATTATGATGGCCTGTTAGTCTCACAATCACT

AAAGAGCATTGCCAGATGTGTCTTCTGGTCCGAGACTATTGTGGATGAAACCAGAGCCGCCTGCAGCAATATTGCAACAA

CTGTCGCCAAGAGTATAGAGAGGGGTTATGATAGGTACCTTGCTTACTCCTTGAATATACTCAAAATTCTCCAACAGATC

CTTATATCCCTTAACTTCACTATCAACACGACAATGACTCAAGATGTCGTGGCACCGATCATCGAGAACGGTGATCTACT

AACAAGGATGGCACTTTTGCCAGCACCCATCGGGGGCCTTAACTACCTTAACATGAGCAGGTTATTTGTGAGAAATATCG

GTGACCCAGTCACTTCCTCTATAGCCGACCTGAAGAGGATGATAGACGCTGGGCTAATGCCAGAAGAAACATTGCACCAG

GTGATGACCCAAACCCCGGGAGAATCATCTTACCTTGATTGGGCAAGTGACCCTTATTCTGCCAACCTAACCTGTGTACA

GAGTATAACTCGCCTTCTCAAGAACATCACTGCAAGGTACATTTTAACCAGCAGCCCAAATCCGATGCTAAAAGGGTTAT

TCCATGAGGGGAGCAGAGATGAAGATGAAGAGCTTGCGAGTTTCTTGATGGATCGGCATATAATTGTTCCCAGAGCTGCA

CATGAAATCTTAGACCATAGCATAACTGGAGCAAGAGAAGCTATCGCCGGTATGTTAGATACCACCAAGGGTCTGATTAG

AACAAGTATGAAACGAGGTGGCCTTACCCCACGGGTCTTAGCTCGCCTTTCCAATTATGATTATGAACAATTCAGATCCG

GTATAACATTATTGACGAAGAAGGGACAGTGTTATCTCATTGACAAAGATTCGTGCTCGGTGCAGCTTGCTATAGCCCTG

AGGAGCCATATGTGGGCTAGGTTGGCCCGCGGGAGACCTATCTATGGGTTGGAAGTGCCTGATGTACTGGAATCGATGAA

TGGCTATCTTATCAAACGCCATGAGGCTTGTGCCATTTGCGAGACGGGCTCAAGTCATTACGGGTGGTTCTTCGTCCCTG

CAGGGTGCCAGCTTGACGATGTCTCGAAAGAGACTTCAGCTCTTCGTGTGCCTTATGTCGGGTCAACCACAGAGGAAAGG

ACAGATATGAAACTTGCCTTCGTTAGATCTCCAAGTCGATCCCTCAAATCAGCAGTCAGAATTGCTACCGTTTATTCATG

GGCCTATGGGGATGATGAAAAATCATGGAGTGAAGCCTGGATGCTGGCAAGGCAGAGAGCCGATATCACCCTAGATGAGT

TGAGAATGATCACTCCAGTCTCTACATCGACCAACCTAGCCCACCGGTTGAGGGATCGGAGCACCCAGGTGAAATATTCG

GGGACATCCCTTGTGAGGGTTGCAAGATACACAACCATTTCCAATGATAATTTGTCATTTGTGATATCTGAGAAGAAAGT

AGATACCAACTTCATTTACCAGCAAGGGATGCTCCTCGGTCTTGGGATCCTTGAGAATCTTTTCAGGTTGGAGGCCACCA

CAGGGGTATCCAACACAGTGCTACATCTGCACGTGGAGACAGAATGTTGTGTTGTACCTATGGTTGATCACCCAAGAATA

CCGAGTCTCCGTAACATTAAGGTTACGAATGAGCTATGCACAAACCCCTTGATCTACGACAAGTCTCCTATCATAGAACA

CGATGCAACCCGATTATACTCACAAAGCCACAGAAGACATCTGGTGGAGTTTGTTACCTGGTCAACAAGTCAGCTTTATC

ATATACTGGCTAAGTCTACAGCAATGTCCATGATTGAGTTGATTACAAGATTCGAGAAGGATCACATGAATGAAATAACC

GCCTTGATTGGCGATGACGACATCAACAGTTTTATCACAGAATTCCTGCTAGTAGAACCTAGACTGTTTATAGTCTACCT

TGGCCAGTGTGCTGCCATCAATTGGGCTTTTGATATACATTATCATCGCCCATCGGGCAAATACCAGATGGGAGAGCTTC

TCTACTCTTTACTCTCTAGGATGAGCAAAGGAGTATATAAGATCTTTACCAATGCTTTGAGCCACCCCAAAGTTTACAAG

AAATTTTGGCGAAGTGGTGTCATCGAGCCGATTCATGGCCCGTCCTTAGATACACAGAATCTACATGTCACTGTCTGTGA

TATGATATATGGATCCTACGTCACCTATCTGGATCTTTTGCTGAATGATGAGCTAGATGATTATACGTATCTGCTCTGCG

AGAGTGATGAGGACGTGGTCACAGACAGGTTCGATAACATTCAAGCCAAACATCTCTGTGTATTAGCTGATGTGTATTGC

AGCTCCAAAAGGTGTCCCTCGATAATCGGGATGTCTCCTATAGAAAAATGTACCATCCTCACACATTACATCAAGGGAGA

ATCAATACAATCTCCGTCTGGGACCTCATGGAACACTGATCCTCTTGTAGTAGATCATTACTCATGCTCTCTGACATACC

TTCGTCGCGGTTCCATCAAACAAATCAGGTTGAGAGTAGATCCTGGGTTTGTATTTGAGGCATTGACAGACATCGACATC

AAACAGCCTCGCAAGGCTAATACTGATGTATCGGTTGTGGGGTTGACTGATTTCTCTCCTCCTTGGGATAACATTGGTGA

TTTTCTAAGGACTATCAACACCTTGAGACACAATCTGCCTGTCACCGGGACCGGGGTCTCTAACTATGAGGTCCACGCTT

ATCGTAGAATTGGTTTGAATTCATCAGCATGTTATAAAGCTGTAGAGATCTCCACGTTAATCAAGCCATCCTTAGAAGTC

GGAGAGCATGGGTTGTTCTTAGGGGAAGGTTCCGGTTCTATGCTGGCTGCGTACAAGGAAGTTCTTAAATTAGCCAACTG

TTATTACAACAGCGGAGTAATAGCTGAGGGCAGAGCCGGACAAAGAGAAATATCCCCTTACCCCTCAGAGATGAGCTTAG

TAGAGAATCAAATGGGGTTAGAAAAGAGTGTTAAAGTGCTGTTCAACGGCAAACCTGAAGTAACCTGGGTAGGGACTACC

GATTGCTATGAATATATAATCAGTAACATCCAAACCTCTAGTCTGGGTTTCATACACTCAGATATTGAGACACTCCCAAC

CAAAGATGCTGTTGAGAAATTAGAAGAATTTGCCTCGATCTTATCCTTATCCCTAATTCTGGGAAAAGTTGGCTCTATTA

CAGTTATCAAAGTTATGCCTATTAGCGGAGATTTTACTCAGAGCTTCATAAGCTATGCCATCCAATATTTTAGGGAGAGC

CTGCTTGTCTATCCGAGATATAGTAATTTCATTTCAACTGAGTGTTACCTTATTATGATAGGATTGAAAGCCAATCGGTT

GATAAACCCAGAAGCCATTAAGCAAAGCATAATCAGAGCAGGGACTAGAACTGCACCAGGACTAGTGAGCCATATATTAT

CAGAGAAACAAAAAGGGTGTATCCAATCTTATCTGGGTGATCCTTATATCCAAGGAGACTTCAATAAACACCTTAAATCT

CTAACCCCTATTGAGAAAATCCTAATTAATTGTGGTCTCTCGGTCAATGGCACAAAAATCTGTAGGGATCTAATCCACCA

TGATATCGCCTCTGGTCCAGACGGTCTGATGAGTTCCACAATTATTTTATACAGGGAGCTGGCTCATTTCAAAGACAACA

TAAGAAGTCAGCACGGGATGTTCCACCCCTACCCAGTACTGGCCAGTAGCAGGCAACGTGAATTAATTCTTCGAATCGCC

AAGAAATTCTGGGGTTATGTCTTGCTATATTCTGATGATCCGGCACTGATCAGGCAAACAATTAAGAACTTGAAGAGGAA

CCACCTAACCTTTGACTTACACAATAACCCGTTTATTAAGGGTCTGTCCAGAGCTGAGAAATTGCTAGTGCGGACAAGTT

CACTCAGAAGAGAGTGGTTGTTCACTCTCGAAACGAAGGAAGTGAAAGAATGGTTCAAACTGGTGGGTTACAGTGCTCTC

GTCAGAGGTTAATTAGCTATACATCTGCCCCCTTCTCCTCCACCATGAAATTCTACTGGCAATCTAATAGATTAAAGAAA

ACTACAGATCGGATAAGTATCTATTCCGAGTTTTTTCTGGT

>MN121838.1/PrzewalskisGazelle/China/2018-12

ACCAAACAAAGTTGGGTAAGGATAGATCTTATAATAACTATGGACTGGCAAACTTAGGAGTAAAGATCCTACTGTCGGGG

GGAGGAGGAGGAGCAAGATCCTTGATCATGGCGACTCTCCTTAAAAGCTTAGCATTGTTCAAAAGGAACAAAGACAAAGC

GCCGACAGCATCAGGTTCAGGAGGGGCCATCCGGGGGATTAAGAATGTTATCATAGTCCCGATTCCCGGAGACTCGTCCA

TCACTACCCGTTCAAGACTGCTCGACAGGCTTGTCAGATTGGCCGGAGATCCTGACATCAACGGGTCAAAGCTGACCGGG

GTGATGATCAGCATGTTATCCTTGTTCGTAGAGTCACCCGGGCAATTGATACAGCGAATCACAGATGATCCGGATGTTAG

TATCCGCCTTGTTGAGGTAGTTCAAAGTACTAGGTCCCAGTCCGGGTTGACCTTTGCATCACGTGGTGCTGATTTAGACA

ACGAGGCAGACATGTATTTTTCAACTGAGGGGCCCTCGAGTGGAGGCAAGAAAAGGATTAACTGGTTTGAGAACAGAGAA

ATAATAGACATAGAGGTGCAGGATCCAGAAGAGTTCAATATGTTGTTAGCCTCCATACTAGCACAAGTCTGGATCCTTCT

GGCCAAGGCTGTTACGGCACCAGATACGGCAGCTGACTCAGAACTGAGAAGGTGGGTTAAATACACACAACAAAGGAGAG

TGATTGGGGAATTTCGCCTTGACAAGGGGTGGCTGGACGCAGTCCGCAACAGAATTGCAGAAGATCTATCACTCCGGCGG

TTCATGGTATCTCTCATACTTGACATCAAGAGGACCCCAGGCAACAAGCCAAGGATTGCAGAAATGATCTGTGACATTGA

CAACTATATTGTCGAAGCAGGACTCGCCAGTTTCATCCTTACTATCAAATTTGGTATCGAAACCATGTATCCTGCATTAG

GGCTCCACGAGTTTGCCGGAGAATTATCCACTATTGAGTCCTTGATGAACTTGTATCAACAGCTAGGAGAGGTTGCACCC

TACATGGTAATTCTAGAGAATTCAATTCAGAACAAGTTTAGTGCAGGGGCTTATCCCCTCCTCTGGAGCTATGCGATGGG

TGTCGGAGTCGAGCTGGAGAACTCAATGGGGGGCTTGAACTTTGGCAGGTCATATTTTGACCCTGCCTATTTTCGTCTCG

GACAGGAGATGGTCAGAAGATCTGCAGGAAAGGTCAGCTCTGTGATTGCGGCTGAGCTCGGCATCACAGCAGAGGAAGCT

AAACTAGTCTCGGAAATCGCCTCGCAGGCTGGGGACGAAAGAACCGCCAGAGGGACTGGGCCTCGACAGGCGCAGGTTTC

CTTCCTTCAGCACCAAACAGGAGGGGGAGAGTCGTCCGCACCAGCGACCAGAGAAGGGGTCAAAGCTGCGATCCCAAACG

GAGCTGAAGAAAGGGACAGAAAGCAAACACGCCCAGGAAGGCCCAGAGGAGAGACCTCCGGCCAACTGCTCCTGGACATC

ATGCCAGAGGATGAGATCTTGCGAGAGTCTGGTCAAAACCCTCGTGAGGCTCAAAGATCGGCCGAGGCACTCTTCAGGCT

GCAGGCCATGGCCAAGATTCTGGAGGACCAGGAGGAGGGAGAAGACAACAGTCAGGTCTACAACGACAAGGATCTCCTCG

GCTAAGCAGACGCACCCTCTGTCGAAATCAGTGACAAGATATCCCCTACCAGTATTATAAAAAACTTAGGACCCAGGTCC

AAGCAAGCACACATCGACACTCCAACCAGTCGAGCGGAGACCACCGATGGCAGAAGAACAAGCATACCATGTCAACAAGG

GGCTGGAATGTATCAAGTCTCTCAAAGCCTCACCCCCGGATCTATCAACCATCAGAGATACCATCGAGAGCTGGAGAGAG

GGGCTTAGCCCCTCGGGCCGTGCAACACCGAACCCTGATACGTCCGAGGGAGATCATCAGAATATCAACCAATCATGCTC

ACCAGCAATCGGACCAAACAAAGTCTACTTGTCTCCTGAAGATAATCTCGGATTTAGAGAGATCACTGGCAACGACTGTG

AGGCTGGGCTCGGAGGGGTCCAGAGAGAAGGATCCAACTCTCAAGTACAGCGTTACCATGTTTATAGTCACGGGGGTGAA

GAGATTGAAGGACTCGAGGATGCTGACTCTCTCGTGGTTCAAGCAGATCCTCCAGTTGCTAACGTCTTCAATGGAGGAGA

GGATGGATCTGACGACAGCGATGTGGACTCTGGCCCAGATGATCCCGGCAGAGATACTTTATATGACCGGGGATCTGTTG

CCGGCAATGGTATCGCTAGGTCCACAGATGTCGAAAAACTAGAAGGTGCTGATATTCAAGAAGTTCTTAACTCCCAGAAA

GGCAAAGGAGGAAGATTCCAGGGCGGGAAAACCTTGCGAGTCCCGGAAATACCCGATGTCAAGCACTCCAGACCATCAGC

CCAATCAATTAAAAAGGGCACAGACGGGAACTCAGTCTCATCTGGAACGGTGATAGAGTGTTTATCGATAAGTGGTGCAA

CCCAAACTGTGCCAGAGTTAAGATGGGAGTCATCAGAGCAAAATGCGTCTGTGGGGAGTGTCCTCAAGTCTGCGAGGAGT

GCAAAGACGATCCAGGGGTCGACACAAGAATCTGGTACCATAGCATCACTGACTCAGCCTAAAGAGAATGACTCCGAGTA

TGAGTATGAGGATGACCTATTTACAGAGATTCAGGACATCCGTGCAAGCATTGCCAAGATTCATGATGACAATAAAACTA

TCCTCTCAAAGCTTGATTCTATACTGTTATTGAAAGGGGAAGTCGACACTATCAAGAAACAAATCAGCAAGCAGAATATA

AGTATATCCACCATTGAGGGCCATCTCTCCAGTATCATGATAGCCATCCCTGGCTTTGGGAAGGACATCAAGGACCCAAC

ATCCGAAGTCGAGTTGAACCCAGATCTAAGACCTATAATAAGCCGTGACTCTGGCAGAGCTCTCGCGGAGGTCCTCAAGA

AACCCGCTGTGGATAGGTCTCAGAAAATTGGAACCAAAGCCAACTCCAGCTCAAAGGGTCAGCTTCTTAAGGATCTCCAG

CTAAAACCTGTCGACAAGCAGGCAAGCTCTGCAATCGGGTTTGTCCCGTCCGACCATGAATCATCCAGAAATGTCATCCG

CTCCATAATCAAGTCGAGCAAGCTAAATATTGATCACAAGGACTATCTTCTAGATTTACTAAATGATGTGAAAGGCTCCA

AGGATCTTAAGGAATTTCACAAGATGCTAACAGCAATTCTCGCCAAGCACCCGTAACACATCCTCCAGTCATCATCTCAT

ACTCGACTAAAAACATCCTTTCAATCAGGCTATTACAAAAAACTTAGGAGCAAGGGCAACTGAGCTTCGCAGACAGGATG

ACCGAGATCTACGACTTCGATAAATCAGCATGGGATGTCAAAGGGTCAATTGCCCGCATAGAACCCACCACCTACCACGA

CGGCCGACTGATACCCCAGGTGAGGGTCATCGATCCTGGTCTGGGAGACAGAAAAGATGAGTGCTTTATGTACCTGTTTC

TCCTAGGAGTGATTGAGGATAACGACCCCCTGTCTCCCCCCGTCGGAAGAACCTTTGGCTCTCTACCTCTAGGGGTTGGT

AGGTCAACTGCCAAGCCAGAAGAACTACTAAGGGAGGCCACAGAATTAGATATAGTGGTGAGGCGCACTGCAGGAGTAAA

TGAGAAACTGGTATTTTACAACAACACTCCGCTGTCCTTGTTAACGCCCTGGAAGAAAGTTCTGACAACCGGAAGTGTGT

TTAGCGCTAACCAGGTTTGCAATGCAGTCAACCTAGTCCCGCTTGATACTCCCCAGAGATTCAGGGTTGTGTACATGAGC

ATAACTAGATTATCAGACAATGGTTACTATAGTGTGCCCAGAAGAATGCTGGAGTTCCGCTCGGCCAATGCAGTAGCCTT

CAACATCTTGGTTACACTAAGAATTGAAAATGGCACAAACCCTAGAAGATACATAGTCGGCTCATGGGAGAATTCAGAGG

TCACATTTATGGTACACGTGGGCAACTTTAGGAGAAAGAAGAACGAAGTATACTCTGCTGATTACTGCAAAATGAAGATT

GAGAAGATGGGTTTAGTTTTTGCCCTGGGTGGAATAGGCGGAACAAGTCTCCATATTAGAAGCACCGGGAAAATGAGCAA

AACCCTCCATGCACAGCTGGGGTTCAAGAAAATCTTATGTTACCCCCTAATGGATGTCAATGAGGATCTCAACCGATATC

TCTGGCGGGCAGAGTGCCGAATAGTCAAAATCCAAGCTGTCTTACAGCCATCAGTACCCCAAGAGTTCCGTGTCTACGAT

GATGTCATCATCAACGATGATCAAGGCCTGTTCAAGATCTTGTAGTTCATTTGCAAAGGAACGAACCCACCCCCAGGAAG

GGGCCCCACCCTCCCCCGCC--AAAAACACACTCCAAGGCTCCCCGGCACAGCGCCAACACAGCTACTTCCGCGATGGAA

CGAGGTCTG-CCCCCTAACAAATCCGGTCCAGGGGGGGGGGCCCCCCCCTTGACCAACCGGGCGCCGACCGAGCAGGCCG

GCGAGAAGCCCCCAAGAGGCCCCGACGAAGACCGAGCACCCCATCCCCCC-AAAAACGTCCCCC-AACACGCCCGAGGCT

AGCCACCGGACACCCCCAGACCCGTCACCGGCGAGACAGAACCCCAAGGGGGGACCCTCCAAACCAAAGGAATCACCGGG

GTAGGCTCCGCCGAACCAGCACAGACCCGACCCAAGCAAGGCGGACCCCCAACACGAAACTGCCCAGCACACCCCACTCA

TCATCAAACAAAACTTAGGGGCCAAGTCCATACGAAAGGCCCTAGTCCGTCAGGCGGAGGGACGCAGAAAGGAAGGAGAC

ACGACCGTCCCAGAAGGAGGACACCCAACCACCGAAACGCCCGCCCCCAGGGAAGGAG---CCACCCCCACGCAGCGAAG

GAGCGGGTGGGAGACCTCGGGGAGGGGCCGACCGGGATCCGGGGGCCGGCGGCGCCGGCGGTGAGCAGGCCGGGCGCACA

CCCGACCCCACCGGAGGATCCGAGGAGGGCCCGACCCCGCGCGGACCAAACGGCAGGGAGCCCAACCCCACCAAAAAGAG

AGGCGAGGCCCCCCCTCTCCCCCCCCCCGCACACAACCCGCAGACAGACCGAGGGAGGGCCATCCCCCAGCGAGACAGGC

GGCCCCCCCGCAGCCGAGCCGCAGCTCCCCAGAATAGGACACCCCCGCACACTCCGAACACCCCCCCGCCCCCCGCCACC

GGGACAGGGGCTCCACCACCACAGACTGGAGCCACCGACAGCCAGCCAGAGGTCCCCTCCAGTCCCGCCCGGCGCCCCCC

CGAAGAGGCCCCGGCAAGGCCCCCCAGCCCGACGTCCCCCGCACGCACCGAAAGAAGGACCCGCCCGAGAACGTGCGGCA

CATCCATGCGCAAACATCATGACGCGGGTCGCAATTTTGACATTTCTGTTTCTTTTCCCAAATGTTGTTGCGTGTCAGAT

TCACTGGGGCAATCTATCCAAGATCGGGATTGTAGGAACAGGGAGTGCCAGCTACAAGGTGATGACTAGGCCAAGCCACC

AGACTCTGGTTATAAAGTTAATGCCAAATATAACGGCTATCGACAATTGTACAAAGTCAGAGATTGCAGAGTACAAGAGA

TTGCTGATCACAGTGTTAAAGCCTGTAGAGGATGCTCTGTCGGTGATAACCAAGAATGTAAGACCAATTCAAACTCTAAC

ACCTGGGCGTAGAACCCGCCGTTTTGCTGGAGCTGTTCTGGCCGGGGTAGCACTTGGAGTTGCGACAGCCGCTCAGATAA

CTGCAGGAGTCGCCCTTCATCAATCATTGATGAACTCCCAAGCAATTGAGAGTTTAAAAACCAGTCTTGAGATGTCGAAT

CAGGCAATAGAAGAAATCAGACTTGCAAATAAGGAGACCATACTGGCAGTACAGGGCGTCCAGGATTATATCAACAATGA

GCTCGTCCCTTCTGTTCATAGAATGTCATGCGAGCTGGTAGGTCACAAGCTCGGCCTCAAGCTCCTTAGGTACTACACCG

AGATCCTGTCCATATTCGGGCCCAGTCTTCGAGACCCGATATCTGCCGAAATATCAATCCAGGCACTTAGTTATGCATTA

GGCGGAGACATTAATAAAATCCTGGACAAGCTTGGGTATAGCGGTGGGGATTTCCTTGCCATCCTAGAAAGCAAGGGAAT

AAAGGCCCGGGTTACATATGTGGACACAAGAGATTACTTTATAATCCTTAGCATTGCCTACCCAACCTTATCTGAGATCA

AGGGAGTGATAGTTCACAAGATAGAAGCTATAACATACAACATTGGGGCACAGGAGTGGTATACTACTATCCCTAAATAT

GTAGCCACTCAGGGGTATCTGATATCGAACTTTGATGAGACGTCATGCGTATTCACTCCAGAGGGGACAGTTTGCAGCCA

GAATGCGTTGTACCCAATGAGCCCATTGCTTCAGGAATGTTTCAGGGGGTCAACAAAATCGTGCGCCAGAACCTTAGTTT

CAGGGACCATAAGTAATAGATTTATCCTATCAAAAGGGAACCTGATTGCAAATTGTGCGTCAGTTTTGTGCAAATGTTAC

ACAACAGAGACAGTTATCAGCCAAGATCCTGACAAACTACTAACTGTTGTAGCATCCGACAAGTGTCCTGTAGTTGAGGT

GGATGGAGTGACAATACAGGTCGGCAGTCGAGAGTATCCGGATTCTGTATACTTACACAAAATAGACTTAGGTCCGGCCA

TCTCCCTAGAAAAACTGGATGTAGGCACCAATTTAGGCAATGCAGTCACAAGACTGGAGAATGCAAAGGAGCTCCTAGAT

GCATCAGACCAAATACTGAAGACTGTTAAGGGGGTACCTTTCAGTGGGAATATGTACATAGCACTGGCAGCTTGCATAGG

AGTATCCCTGGGCCTTGTCACATTAATATGCTGCTGTAAGGGGAGGTGTAAGAACAAGGAAATCCCTATCTCCAAAATCA

ACCCAGGGCTCAAACCCGACCTGACCGGGACCTCAAAGTCGTACGTAAGATCACTGTAGTCAGAATCACCTGGATCATCT

GGCATCACACACAGACATGCACGACACAGGCAGTCCGAGGACGCAAGAAACCCAGCCTCCGGTCACCCACCCGACCCCAC

TCCACGCTCCACCACACATTAGTCATCAAACAAAACTTAGGACGAAAGGTCAATCACCATGTCCGCACAAAGGGAGAGGA

TCAATGCCTTCTACAAAGACAATCCTCACAATAAGAACCATAGGGTGATCCTGGATAGAGAACGCTTGGTCATTGAAAGA

CCCTACATCTTGCTTGGAGTCCTGCTGGTAATGTTCCTGAGTCTAATCGGACTGCTGGCCATTGCAGGGATCAGGCTTCA

CCGGGCCACCGTTGGAACTTCAGAGATCCAGAGTCGGCTGAATACCAATATTAAGTTGACCGAATCTATTGATCACCAGA

CTAAGGATGTCTTAACTCCCCTTTTTAAAATCATTGGCGATGAAGTCGGCATCAGAATTCCACAGAAATTCAGTGATCTT

GTCAAGTTCATCTCCGATAAGATTAAATTCCTCAACCCTGATAGAGAGTATGATTTCAGAGATCTCCGGTGGTGTATGAA

TCCCCCCGAGAGAGTCAAAATTAATTTTGATCAGTTTTGTGAGTACAAGGCTGCGGTTAAGTTAATTGAACATATATTTG

AGTCACCACTCAACAAGTCAAAAAAGCTGCAATCTTTGACTCTCGGGCCCGGAACAGGCTGTCTAGGCAGGACAGTAACA

AGAGCCCATTTCTCAGAACTTACAATGACCTTAATGGACCTGGATCTAGAGATGAAGCACAACGTGTCCTCAGTGTTTAC

CGTAGTTGAAGAGGGATTATTCGGAAGAACATATACCGTCTGGAGATCCGATGCCAGGGATCCGAGCACCGATCTAGGTA

TCGGCCATTTTTTAAGAGTCTTCGAGATCGGACTGGTAAGAGATCTTGGGCTGGGTCCCCCTGTTTTTCATATGACCAAC

TATCTCACGGTGAACATGAGTGATGACTATCGGAGATGTCTTTTAGCGGTAGGGGAGTTGAAGTTGACAGCCCTATGCAC

CTCATCTGAGACTGTGACACTGAGTGAGAGAGGAGTTCCAAGGAGGGAACCTCTTGCGGTTGTGATACTTAATCTAGCTG

GACCCACTCTAGGGGGAGAGCTATACAGTGTCTTGCCTACCTCTGATCTCATGGTGGAGAAACTCTATTTGTCTTCACAT

AGAGGGATCATCAAAGATGATGAGGCCAATTGGGTAGTGCCGTCTACCGATGTTCGTGATCTTCAGAACAAAGGTGAATG

TCTGGTGGAAGCATGCAAGACTCGACCTCCTTCATTTTGCAATGGCACAGGATCAGGCCCGTGGTCAGAAGGGAGAATCC

CTGCCTACGGGGTGATCAGGGTCAGTCTTGACTTAGCGAGTGACCCGGATGTAGTTATCACTTCAGTGTTTGGCCCACTG

ATACCCCACCTATCCGGCATGGATCTTTACAACAACCCGTTTTCAAAAGCTGTATGGTTGGCTGTACCACCTTATGAGCA

GTCATTTCTAGGAATGATAAATACAATTGGATTCCCTAACAGAGCAGAGGTTATGCCGCACATTTTGACCACAGAGATCA

GAGGCCCTCGGGGTCGTTGCCATGTTCCCATAGAATTGTCCCGCCGGGTTGATGACGATATCAAGATCGGGTCCAATATG

GTCATATTGCCGACGATGGACCTGAGGTATATTACAGCCACTTATGATGTCTCCAGGAGCGAGCATGCAATCGTGTACTA

TATCTATGACACGAGTCGCTCATCATCTTACTTCTACCCAGTTCGACTGAATTTCAAAGGCAATCCTCTCTCTTTGAGGA

TAGAGTGTTTCCCCTGGCGTCATAAGGTGTGGTGCTACCATGATTGTCTTATATACAACACCATAATAGGTGAAGAGGTC

CATACGAGAGGGCTGACCGGCATAGAGGTAACATGCAATCCAGTCTGAGCAGAGCTACGACCGTCACTCAGGCAGTCCCT

TGAGTCGCCACCGAGTCCAAGCAGCACAGCTTGAGACAATCAACGGCACAGCCCAGCCAACAATGTTATAAAAAACTTAG

GAGCCAGGGTTGTAGGGGCCATGGACTCTCTATCAGTCAATCAGGTCTTGTACCCTGAGGTTCATCTAGACAGCCCTATT

GTCACAAACAAACTAGTTGCCATCCTTGAGTACTCGGGTATCGACCATAACTATGTTCTTGAAGACCAGACTCTTGTCAA

GAATATTAGGTATAGGCTGGGGTGCGGTTTTTCAAACCAAATGATCATCAATAACAGGGGGGTGGGTGAAACAGTCAATT

CTAAACTTAAAAGTTACCCCCATAATCGTCATATCATATACCCGGATTGCAATAAGGAGTTGTTTTGTATCAAGGATAGC

TGCATATCTAAGAAGCTCTCGGAGCTATTCAAGAAGGGTAATTCCTTGTACTCTAAGATAAGTCACCAGGTACTGGATTG

TCTTAAGAGAGTCAATGGGAAATTAGGTCTGGGCACAGATCTTACCCATGGTCTGAAGGAGGGTATCCTTGACCTAGGGT

TGCACATGCATAGCTCTCAATGGTTCGAGACTTTCCTGTTCTGGTTCACTATCAAGACAGAGATGAGATCAATGATCAAA

GAACAGTCCCATATATGCCACAAGAGGAGGTATAACCCGACTTTTGTGTCGGGGGATGCATTCGAGGTGCTCGTATCGCG

AGACCTCGTTGTGATAATTGATAAGAATACCCAGTATGTCTTCTACTTAACATTTGAGCTGGTCCTTATGTATTGTGATG

TCATAGAGGGTAGACTTATGACGGAGACAGCCATGGCCATAGACCAGAGATATTCAGAACTTCTAAGCCGGGTCAGATAC

TTGTGGGATCTTATTGATGGGTTCTTCCCAACACTAGGCAACACCACATACCAAATTGTTGCTCTGCTTGAACCATTATC

GTTGGCTTATCTTCAACTTCAGGATGTCACTCTGGAGTTAAGAGGTGCTTTTTTAGACCATTGCTTCAAAGAACTCTATG

AGATACTGGAACATTGCGGCATTGACACAGCAGGCACTTACAATTCTATCACTGAAGGATTGGATTACGTATTCATCACC

CACGATATACATCTAACTGGGGAGATCTTTTCATTTTTCCGGAGTTTCGGACACCCCCGTCTGGAAGCAGTCACCGCTGC

GGAAAATGTCAGAAAACATATGAACCAACCGAAGGTAATCAGTTATGAGACTATGATGAAAGGCCATGCGGTATTTTGCG

GGATAATCATAAATGGTTTTCGGGACCGACATGGCGGCAGCTGGCCCCCTGTTGCACTGCCAGAACATGCCTCTGCTGCG

ATCCGGAATGCGCAGGCATCAGGTGAGGGACTGACCCATGACCTGTGTATAGACAACTGGAAATCCTTTGTTGGATTCAA

ATTTGGCTGCTTTATGCCACTCAGCCTAGATAGTGATTTGACCATGTATCTCAAGGACAAGGCATTGGCTGCACTGAAGA

ATGAGTGGGATTCAGTTTACCCGAAAGAATACCTCCGTTATAACCCACCTAGAGGGACAGAGTCAAGGCGATTGGTAGAG

GTATTCCTGAATGACTCCAGCTTTGATCCTTATAACATGATAATGTACGTGGTGAATGGCTCCTACCTTAAAGACCCTGA

GTTTAACCTCTCATACAGCCTAAAAGAGAAGGAGATAAAGGAGACAGGGCGGTTGTTTGCCAAAATGACCTATAAGATGC

GGGCCTGTCAGGTAATCGCTGAAAATCTGATATCGAATGGTGTTGGGAAGTATTTCCGAGACAATGGGATGGCAAAGGAC

GAGCATGACCTAACAAAAGCCCTTCACACCCTGGCAGTCTCAGGTGTTCCCAAAAACAACAAAGATAACCACCGAGGTGG

GCCTCCCAGAAGGACAACAAACCGAGGGGTGAGATCAAGCCAAGGCACCAAAACACAAGATAGAGACAAGGTTCAAGGGG

GACCTATGTACAACTATTTGCGATGCCAACCGATCAGCCCTGATCAGGGTGAGTCATACGAGACTGTTAGTGCATTCATC

ACCGCTGACCTTAAGAAGTATTGCCTGAATTGGAGATACGAGACAATCAGCATATTTGCACAGAGGCTGAATGAAATATA

TGGACTACCATCCTTCTTTCAATGGTTACACAGGATATTGGAAAAATCCGTACTCTACGTCAGTGACCCACATTGTCCTC

CCGATCTAGATAATCATATCCCTCTGGACAGTGTCCCCAATGCCCAAATATTCATCAAGTACCCAATGGGCGGAATAGAA

GGTTATTGCCAAAAACTATGGACAATCAGCACTATACCATACTTGTATCTGGCAGCCTATGAGAGCGGAGTAAGAATCGC

CTCACTGGTGCAGGGTGACAATCAGACAATCGCAGTGACGAAAAGAGTTCCAAGTTCTTGGCCTTATTCACTAAAAAAGA

GGGAGGCCTCCAAAGCCGCTCAAAATTACTTCGTAGTCTTAAGGCAAAGATTGCACGATGTAGGTCATCATTTAAAGGCT

AATGAGACCATAGTATCTTCTCACTTTTTTGTATATTCCAAAGGGATTTATTATGACGGCCTGCTAGTCTCACAATCACT

AAAGAGCATCGCCAGATGTGTCTTCTGGTCCGAGACTATAGTGGATGAAACCAGAGCGGCGTGCAGTAATATTGCAACAA

CTGTCGCTAAGAGTATAGAGAGGGGTTATGATAGGTACCTTGCATATTCTTTGAATATCCTCAAGATTTTCCAACAGATC

CTTATATCCCTTAACTTCACTATTAACACAACAATGACTCAGGATGTCGTGGCACCGATCATAGAGAACGGTGATTTGCT

GATAAGGATGGCACTCTTGCCAGCACCCATCGGGGGTCTCAATTATCTTAACATGAGCAGGTTATTTGTGAGAAATATCG

GTGACCCGGTCACCTCCTCTATAGCCGACCTGAAGAGGATGATAGATGCCGGGCTAATGCCAGAAGAAACATTGCATCAA

GTGATGACCCAGACCCCGGGAGAATCATCCTACCTAGATTGGGCAAGTGACCCTTATTCTGCCAACCTACCCTGCGTACA

GAGTATCACTCGCCTTCTCAAAAACATTACTGCACGATATATTTTAATCAGCAGCCCAAACCCGATGCTGAGAGGGTTAT

TTCATGAGGGGAGTAGAGATGAAGACGAGGAGCTTGCAAGTTTCCTAATGGATCGGCATATAATTGTTCCAAGAGCTGCA

CATGAAATCTTAGACCATAGTATAACCGGAGCAAGAGAAGCTATAGCCGGGATGTTGGACACCACCAAGGGTCTGATTAG

AACAAGTATGAAACGGGGTGGCCTCACCCCTCGAGTCTTAGCCCGCCTTTCCAATTATGATTATGAACAATTCCGATCCG

GGATAACACTATTAACAAAGAAAGGGCAGTGTTATCTCATTGACAAGGACTCGTGCTCGGTGCAGCTCGCGATAGCTCTG

AGGGGCCACATGTGGGCTAAGTTGGCTCGCGGGAGACCAATCTATGGGTTGGAGGTGCCTGATGTATTAGAATCGATGAA

CGGCTATCTTATCAAACGCCATGAGTCGTGTGCTATCTGTGAAACGGGCTCAAGTCACTACGGATGGTTTTTCGTCCCTG

CAGGGTGCCAGCTTGACGATGTATCTAGAGAGACTTCGGCTCTTCGTGTACCTTATGTCGGATCAACAACTGAGGAAAGG

ACAGACATGAAACTTGCATTTGTTAGGTCTCCAAGTAGATCCCTCAAATCAGCAGTTAGAATTGCAACAGTTTACTCATG

GGCTTACGGGGATGATGAGAAATCATGGAGTGAAGCCTGGATGCTAGCTAGGCAGAGAGCCAATATCACCTTAGATGAAT

TGAGAATGATCACTCCAGTCTCCACATCCACCAACTTAGCCCACCGGTTGAGGGATCGGAGTACCCAGGTGAAATACTCG

GGGACATCCCTTGTGAGGGTTGCAAGATACACAACCATCTCCAATGATAACTTGTCGTTTGTGATATCTGAGAAAAAAGT

GGATACCAACTTCATTTATCAGCAAGGGATGTTGCTTGGTCTTGGGATCCTTGAGAACCTCTTTAGGCTAGAGGCCACCA

CCGGAGTATCCAACACAGTGCTACACCTGCACGTGGAAACAGAATGTTGTGTTGTACCTATGGTGGATCACCCAAGGATA

CCGAGTCTCCGTAATATAAAAGTTACGGATGAGCTATGCACAAACCCTTTGATCTACGACAGGTCCCCCATCATAGAGCA

CGATGCAACCCGGCTATATTCACAGAGCCACAGGAGACATTTGGTGGAGTTTGTTACCTGGTCAACAAGCCAGCTTTACC

ATATACTGGCTAAGTCTACAGCAATGTCCATGATTGAATTGATCACGAGATTCGAGAAAGATCACATGAATGAAATAGCC

GCCCTGATTGGTGATGACGACATCAACAGCTTCATCACAGAATTTTTGCTAGTGGAGCCTAGATTATTTATAGTTTACCT

TGGTCAGTGCGCTGCCATCAATTGGGCCTTTGATATACATTATCATCGGCCCTCGGGCAAGTACCAGATGGGGGAACTTC

TCTACTCTCTGCTCTCTCGTATGAGCAAAGGAGTATATAAGATCTTTACTAATGCTTTGAGCCACCCCAAAGTTTACAAG

AAATTTTGGCGAAGTGGCATCATTGAGCCGATTCATGGCCCATCCTTGGATACACAGAATTTACATGTCACTGTCTGTGA

CATGATATATGGGTCCTATGTCACCTATTTGGATCTTTTGCTGAATGATGAGCTAGATGATTACCCGTATTTGCTCTGCG

AGAGTGATGAGGACGTGGTCACAGACAGGTTCGACAACATTCAAGCCAAACATCTCTGTGTATTGGCCGATGTATATTGC

AGCTCCAAGAGGTGTCCTTCAATAATCGGGATGTCTCCTATAGAAAAATGTACCATTCTCACACATTACATCAAGGGAGA

ATCAGTACAATCCCCGTCTGGGACCACATGGAACACTGATCCCCTGGTAGTAGATCATTACTCATGCTCTCTGACCTACC

TTCGCCGCGGTTCCATCAAACAAATCAGGTTGAGGGTGGATCCTGGGTTTGTATTCGAGGCGTTAACAGACGTCGACTTT

AAACAACCTCGCAAAGCTAAGTTAGATATATCGGTCGTTGGGTTGACTGATTTCTCTCCCCCTTACGACAACGTCGGTGA

TTTTTTAGGGACTATCAACACATTGAGGCACGATCTGCCTGTCACCGGAACCGGGGTCTCGAACTATGAAGTCCACGCTT

ATCGTAGAATTGGCCTGAATTCATCAGCATGTTATAAAGCCGTAGAGATCTCCACGTTAATCAAGCCATCCTTAGAAGTC

GGAGAGCATGGATTGTTCTTAGGAGAAGGTTCCGGTTCAATGCTGGCTGCGTACAAAGAAGTTCTTAAATTAGCAAATTG

TTACTACAACAGCGGAGTAACAGCGGAGGGCAGAGCCGGACAGAGGGAAATATCTCCCTATCCTTCGGAGATGAGCCTAG

TAGAGAATCAAATGGGGATAGAGCGGAGTGTTAAAGTGCTGTTCAACGGGAAACCTGAAGTAACTTGGGTGGGGACCACA

GATTGCTACAAGTACATAATCAGTAACATCCAAACCTCTAGTCTGGGTTTCATACATTCAGATATTGAGACGCTCCCAAC

CAAAGATGCTGTTGAGAAATTGGAGGAATTTGCCTCTATCCTATCCTTATCCTTGATTTTGGGGAAAATCGGCTCTATTA

CAGTTGTCAAAATTATGCCCATTAGCGGGGATTTTACCCAAGGCTTCATCGCCTATGCCATCCAATATTTTAGGGAGAGC

CTGCTTGCCTATCCGAGATATAGTAACTTCATCTCGACTGAGTGTTACCTTATTATGATAGGATTAAAGGCCAATCGATT

GATAAACCCAGAAGCCATTAAGCAAAGCATAATCAGAGCGGGGGTCAGGACTGCACCAGGACTTGTGAGCCATGTATTAT

CAGAGAAACAAAAAGGTTGTATTCAATCTTTCTTGGGTGATCCTTATATCCAAGGAGACTTCAATAAGCACCTTAAATCT

CTAACCTCTATTGAGAAAATCCTGGTAAATTGTGGTCTCTCGATCAATGGCATAAAAATCTGTAGAGATCTAATCCACCA

TGATATCGCCTCCGGTCCAGACGGCCTTATGAGCTCTACAATTATTTTATATAGGGAATTGGCTCATTTCAAAGATAATA

TAAGGAGTCAGCACGGAATGTTCCACCCCTACCCGGTACTGGCCAACAGCAGGCAACGTGAATTAATCCTTCGAATAGCC

AAGAAATTCTGGGGGTATGTCTTGCTATATTCTGATGACCCGGCACTAATCAGACAAACAATCAAGAACTTGAAACGGAA

TCATTTAACCTTTGACTTACATAGTAACCCATTTATTAAAGGGCTATCCAAGGCTGAGAAACTGCTGGTGCGGACGAGTT

CACTAAGAAGAGAATGGTTGTTCACTCTCGAAACGAAAGAAGTAAAAGAGTGGTTCAAATTGGTAGGTTACAGTGCACTC

ATCAGAGGCTAGTCAGCTATGCATCTGCCCCCTTCTCCTCCGCCATGAGACCCCACTGGCGATCCAGAAGATTAAAGAAA

ACTACATATTGGATAAGTATCTATTCCCAGCTTTGTCTGGT

>MK991798.1/goat/Ethiopia/2011-05

ACCAAACAAAGTTGGGTAAGGATAGATCTTATAATAACCATAGACTGGCAAACTTAGGAGTAAAGATCCTACTGTCGGGG

GGAGGAGGAGGAGCAAGATCTTTGATTATGGCGACTCTCCTTAAAAGCTTAGCATTGTTCAAAAGGAACAAAGACAAAGC

GCCGACGGCATCAGGTTCAGGGGGGGCCATCCGGGGGATTAAGAATGTTATCATAGTCCCGATTCCCGGAGACTCTTCCA

TCATTACCCGTTCAAGACTGCTCGACAGGCTTGTCAGATTGGCCGGAGATCCTGACATCAACGGGTCAAAGCTGACCGGC

GTGATGATCAGCATGTTATCTTTGTTCGTAGAGTCTCCCGGGCAATTGATACAGCGAATCACAGATGATCCAGATGTTAG

TATCCGCCTTGTTGAGGTAGTTCAAAGTACTAGGTCCCAGTCCGGGTTGACCTTTGCATCACGTGGTGCTGATTTGGACA

ACGAGGCAGACATGTATTTTTCAACTGAGGGGCCCTCGAGTGGAGGTAAGAAAAGGATCAACTGGTTTGAGAACAGAGAA

ATAATAGACATAGAGGTGCAGGATCCAGAAGAGTTCAATATGTTGTTGGCCTCCATACTAGCACAAGTCTGGATCCTTCT

GGCCAAGGCTGTTACGGCACCGGATACGGCAGCTGACTCAGAACTGAGGAGGTGGGTTAAATACACACAACAGAGGAGAG

TGATTGGGGAATTTCGCCTTGACAAAGGGTGGCTGGACGCAGTCCGCAACAGAATTGCAGAAGATCTATCACTTCGGCGG

TTCATGGTGTCTCTCATACTTGACATCAAGAGGACCCCCGGCAACAAGCCAAGGATTGCGGAAATGATCTGCGACATTGA

CAACTATATTGTCGAAGCCGGACTCGCCAGTTTCATCCTTACTATCAAGTTTGGTATCGAAACCATGTATCCTGCATTAG

GGCTCCACGAGTTTGCCGGGGAATTGTCCACTATTGAGTCCTTGATGAACTTGTATCAACAGCTAGGAGAGGTTGCACCC

TATATGGTAATTCTAGAGAACTCAATTCAGAACAAGTTTAGCGCAGGAGCTTATCCCCTCCTCTGGAGCTATGCGATGGG

TGTCGGAGTTGAGCTGGAGAACTCAATGGGGGGCTTGAACTTTGGCAGGTCATATTTTGACCCTGCCTATTTTCGTCTCG

GACAGGAGATGGTCAGAAGATCTGCTGGAAAGGTCAGCTCTGTGATTGCGGCTGAGCTCGGCATCACAGCAGAGGAAGCT

AAACTAGTCTCGGAAATCGCCTCGCAGACTGGGGACGAAAGAATCGCCAGAGGGACTGGGCCTCGACAGGCGCAGGTCTC

CTTCCTGCAGCACAAAGCAGGAGAGGGAGATTCGCCCGCACCAGCGACCAGAGAAGGGGTCAAAGCTGCGATCTCAAACG

GATCTGAAGAAAGGGACAGAAAGCAAACACGCCCAGGAAGGCCCAGAGGAGAGACCCCCGGTCAACTGCTCCTGGAAATC

ATGCCAGAGGATGAGGTCTCGCGAGAGTCTGGTCAAAACCCTCGTGAGGCTCAAAGATCAGCCGAGGCACTCTTCAGGCT

GCAGGCCATGGCCAAGATTCTGGAGGACCAGGAGGAGGGAGAAGACAACAGTCAGGTCTACAACGACAAGGATCTCCTCG

GCTAAGCAGACGCACCTTCTGTCGAAATCAGTGACAAGACATCTCCTACCAGTATTATAAAAAACTTAGGACCCAGGTCC

AAGCAAGCACACATCGACACTCCAATCAGTCGAGCGGAGACCACCGATGGCAGAAGAACAAGCATACCATGTCAACAAGG

GGCTGGAATGTATCAAGTCTCTCAAAGCCTCTCCCCCGGATCTATCCACCATCAGAGATACCATCGAAAGCTGGAGAGAG

GGGCTTAGCCCCCCGGGCCGTGCAACACCGAACCCTGATACGTCCGAGGGAGATCATCAGAATATCAACCAATCATGCTC

ACCAGCAATCGGACCAAACAAAGTCTACCTGTCTCCTGAAGATAATCTCGGATTTAGAGAGATCACTGGCAACGACTGTG

AGGCTGGGCTCGGAGGAGTCCAGGGGAAAGGATCCAACCCTCAAGTACAGCGTTACTATGTTTATAGCCACGGGGGTGAA

GAGATTGAAGGACTCGAGGATGCTGACTCTCTCGTGGTTCAAGCAGATCCTCCAGTTGCTAACATCTTCAATAGAGGAGA

GGATGGATCTGACGACAGCGATGTGGACTCTGGCCCGGATGATCCCGACAGAGATACTCTATATGACCGGGGATCTATTG

CCGGCAATGATGTCGCTAGGTCCACAGATGTCGAAAAACTAGAAGGTGCTGATATTCAAGAAGTTCTTAACTCCCAGAAA

GGCAAAGGAGGGAGATTCCAAGGCGGGAAAACCTTACGAGTCCCGGAAATACCCGATGTTAAGCACTCCAGACCATCAGC

CCAATCAATTAAAAAGGGCACAGACGGGAACTCAGTCTCATCTGGAACGGTGACAGAGTGTTTATCGATAAGTGGTGCAA

CCCAAGCTGTGCCAGAGTCAAGATGGGAGTCATCAGAGCAAAATGCGTCTGTGGGGAGTGTCCTCAAGTCTGCGAAGAGT

GCAAAGACGATCCAGGGGTTGACACAAGAATCTGGTACCATAGCATCACTGACTCAGCCTAAAGAAAATGACTCCGAGTA

TGAGTATGAGGATGATCTATTTACAGAGATTCAGGACATCCGTGCAAGCATTGCCAAGATCCATGATGACAACAAAACTA

TCCTCTCAAAGCTTGATTCTATACTGTTATTGAAAGGAGAAGTTGACACTATCAAGAAACAAATCAGCAAGCAGAATATA

AGTATATCCACCATTGAGGGCCATCTCTCCAGTATAATGATAGCCATCCCGGGCTTTGGGAAGGATATCAAGGACCCAAC

ATCCGAGGTTGAGTTGAACCCAGATTTAAGACCTATAATAAGCCGTGACTCTGGCAGAGCTCTCGCGGAGGTCCTCAAGA

AACCCGCTGTGGATCGGTCTCAGAAAATTGGAACCAGAACCAACTCCAGCTCAAAGGGTCAGCTTCTTAAGGATCTCCAG

CTAAAACCTGTCGACAAGCAAGCAAGCTCTGCAATCGGGTTCGTCCCATCCGACCATGAATCATCCAGAAATGTCATCCG

CTCCATAATCAAGTCGAGCAAGCTAAACATTGATCACAAGGACTATCTTCTAGACTTACTAACTGATGTGAAAGGCTCCA

AGGATCTTAAGGAATTCCACAAGATGCTAACAGCAATTCTCGCCAAGCACCCGTAACACACCCTCCAGTCACCATCTCAT

ACTCGGCTGAAAGCGTCCTCTCAATCAGGCTATTACAAAAAACTTAGGAGCAAGGGCAACTGAGCTTCGCAGACAAGATG

ACCGAGATCTACGACTTTGATAAATCAGCATGGGATGTCAAAGGGTCAATTGCCCGCATAGAACCCACCACCTATCACGA

CGGCCGACTGATACCCCAGGTAAGGGTCATCGATCCTGGTCTGGGAGACAGAAAAGATGAGTGCTTCATGTACCTGTTTC

TCCTAGGAGTGATTGAAGATAACGACCCCCTGTCTCCCCCAGTCGGAAGAACCTTTGGTTCTTTACCTCTAGGGGTCGGT

AGGTCAACTGCTAAGCCAGAAGAACTACTAAGGGAGGCCACGGAACTAGATATAGTGGTGAGGCGCACTGCAGGATTAAA

TGAGAAACTGGTATTTTACAACAACACTCCGCTGTCCTTGTTAACGCCCTGGAAGAAAGTTCTGACAACCGGAAGTGTGT

TTAGCGCTAACCAGGTTTGCAATGCGGTCAACCTAGTCCCACTTGATACTCCCCAGAGATTCAGGGTTGTGTACATGAGC

ATAACTAGATTATCAGACAATGGTTACTATAGTGTGCCCAGGAGAATGCTGGAGTTCCGCTCAGCCAATGCAGTAGCTTT

CAACATCTTGGTTACACTAAGAATTGAAAATGGCACAAACTCTAGAAGATACATAGTCGGCTCATGGGAGAATCCAGAGG

TCACATTTATGGTACACGTGGGCAACTTTAGGAGAAAGAAGAACGAAGTATACTCTGCTGATTACTGCAAAATGAAGATT

GAAAAGATGGGTTTAGTTTTTGCCCTAGGAGGAATAGGTGGAACAAGTCTCCATATTAGAAGCACCGGGAAAATGAGCAA

AACCCTCCATGCACAGCTAGGGTTCAAGAAAATCTTATGTTACCCCCTAATGGATGTTAATGAAGATCTTAACCGATATC

TCTGGCGGGCAGAGTGCCGAATAGTCAAAATCCAAGCCGTCTTACAGCCATCAGTGCCCCAAGAATTCCGTGTCTACGAT

GATGTCATCATCAACGATGATCAAGGCTTGTTCAAGATCCTGTAGTTCATTTGCAAAGGAATGAACCCACCCCCAGGAAG

GAGCCCCACCCTCCCCCGCC-AAAAAACACACTCCAAGGCCCCCCGACACAGCGCCAACACAGCCACTCTCCCGATGGGA

CGAGGCCGG-CTCCCCAACAAACCCAGTCCAGGTCGGGGGA-CCCCCAAACGACCCGCCGGGCGCTGACCGATCAGGCCG

GCAAGAAGCCCCCGACAGGCCCCGACGACGACCGAGCACCCCATCCCCCC--AAACCGTCTCCCGAAAACGCCCGAGGCC

AGGCGCCGAGCACCCCCAGACCCGCCACCGGCGAGACAGACCCCCAAGGGGGGACCCTCCAAACCAAAGGGACCACCGAG

GAAAGCTCCGCCGAACCAGCACAGACCCGACCCAAGCAAGCCGGACCCCCAACACGAGACCGCCCAGCGCACCCCACCCA

TCACCAAACAAAACTTAGGGGCCAAGTCCGCACGACAGGCCCCAGCCCGCCAGGCGGAGGGCCGCAGAAAGGAAGGAGAC

ACGACCGTCCCAGAAGGAAGACACCCAACCACTGAAACACCCAGCCCCAGGGAAGGAG--CCCCCCCCCACACGGCGAAG

GAACGGGTGAGAGACCCCAAGGAGAACCCGACCGGAAGCCAAGAGCCGGCGACGCCGGCGGCAAACAGGCAGGGCACACA

CCCGACTCCACCAGAGGAGCCGAGGAGAGCCAGACCCCGCGTGGACCAATCGGCAGAGAATCCAACCCCGCCACAAAAAG

AGGCGAGGCCCT-CCT-----CCCCCTCGCACACAACCCACAGACAGGCCAAGGGAGGACCACCTCCCAACAAGACAGGC

GGCTCCCTCGCAGCCGAGCCGCAGCTCCCCAGAAGAGAACACCCCCGCACGCTCCGAGCACCCCCCCACCCCTCGCCGCC

GGGACAGGAGCTCCGCCACCACTGACCAGAGCCACCGAGAGCCAGCCAGAGGTCCCCTCCAGTCCCGCCCAGCGCCCCCC

CGAAGAGCCCCCGGCAAGGCCCCCCAGCCCGACGACCCCTGCACGCACCGAAAGAGGGACTCGCCCGAGAACGTCCTGCA

CATCCATGCGTAAACATCATGACGCGGGTCGCAATCTTGAAATTTCTGTTTCTTTTCCTAAATGCTGTTGCGTGCCAGAT

TCACTGGGGCAATCTATCCAAGATCGGGATCGTAGGAACAGGGAGTGCCAGCTACAAGGTGATGACTAGGCCAAGCCACC

AAACTCTGGTTATAAAGTTAATGCCAAATATAACAGCCATCGACAATTGTACAAAGTCAGAGATTGCAGAGTACAAGAGA

TTGCTGATCACAGTGTTAAAGCCTGTAGAGGATGCTCTGTCAGTGATAACCAAGAATGTAAGACCAATTCAAACCCTGAC

ACCTGGGCGTAGAACTCGCCGTTTCGCTGGTGCTGTTCTGGCCGGAGTAGCACTTGGAGTTGCGACAGCCGCTCAGATAA

CTGCAGGAGTCGCCCTTCATCAATCATTGATGAACTCCCAAGCAATTGAGAGTTTAAAAACCAGTCTTGAGAAGTCGAAT

CAGGCAATAGAAGAAATCAGACTTGCAAATAAGGAGACCATACTGGCAGTACAGGGCGTCCAGGATTATATCAACAATGA

GCTTGTCCCTTCTGTTCATAGAATGTCATGCGAGCTGGTCGGTCACAAGCTCGGCCTCAAGCTCCTTAGGTACTACACCG

AGATCCTGTCCATATTCGGGCCCAGCCTTCGAGACCCGATAGCTGCAGAAATATCAATCCAGGCACTCAGTTATGCATTA

GGCGGGGACATTAATAAAATTCTGGACAAGCTTGGGTATAGCGGTGGGGATTTCCTTGCCATCCTGGAAAGCAAGGGAAT

AAAGGCTCGAGTCACATATGTGGACACAAGAGATCACTTCATAATTCTTAGCATCGCCTACCCAACCTTATCTGAGATCA

AGGGAGTGATAGTTCACAAGATAGAAGCTATAACATACAACATTGGGGCACAGGAGTGGTATACTACTATCCCTAAGTAT

GTAGCCACTCAGGGGTATCTGATATCGAACTTTGATGAGACGTCATGCGTATTCACTCCGGAGGGGACAGTTTGCAGCCA

GAATGCGTTGTACCCAATGAGCCCATTGCTTCAGGAATGTTTCAGGGGGTCAACAAAATCGTGCGCCAGAACCCTAGTTT

CAGGGACCATAAGCAATAGATTTATCCTATCAAAAGGGAACCTGATTGCAAATTGCGCGTCAGTTTTGTGCAAATGTTAC

ACAACGGGGACAGTTATCAGCCAAGATCCTGACAAACTACTAACTGTTGTAGCATCTGACAAGTGTCCTGTAGTTGAGGT

GGACGGAGTGACAATACAGGTCGGCAGTCGAGAGTATCCGGATTCTGTATACTTACACAAGATAGACTTAGGTCCAGCCA

TCTCCCTAGAAAAACTGGATGTGGGCACCAATTTAGGCAATGCAGTCACAAGACTGGAGAATGCAAAGGAGCTTCTAGAT

GCATCAGACCAAATACTGAAGACTGTTAAAGGGGTACCTTTCAGTGGGAATATGTTCATAGCACTGGCAGTTTGCATTGG

GGTATCCCTAGGGCTTGTCACATTAATATGCTGCTGTAAGGGGAGGTGCAAGAACAAGGAAATCCCGATCTCCAAAATCA

AACCAGGGCTCAAACCCGATCTGACCGGGACCTCAAAGTCGTACGTGAGATCACTGTAGTCAGAATCACCCGAATCATCC

GGCATCACACACACACATGCACGACACAAGCAGTCAGAGGACGCAGGAAGCCCAGTCTCCGATCACCAACCTGACCCTAC

TCCAAGCTCCACCAGACATTATTCATCAAACAAAACTTAGGACGAAAGGTCAATCACCATGTCCGCACAAAGGGAAAGGG

TCAATGCCTTCTACAAGGACAATCCTCACAATAAGAACCATAGGGTGATCCTGGATAGAGAACGCTTGGTCATTGAAAGA

CCCTACATCTTGCTTGGAGTCCTGCTGGTAATGTTCCTGAGTCTAATCGGACTGCTGGCAATTGCAGGGATCAGGCTTCA

CCGGGCCACCGTTGGAACTTCAGAGATCCAGAGTCGGCTGAATACCAACATTGAGTTGACCGAATCCATTGATCACCAAA

CTAAGGATGTCTTAACTCCCCTTTTTAAAATCATTGGCGATGAAGTCGGCATCAGAATTCCACAGAAATTCAGTGATCTT

GTCAAGTTCATCTCCGATAAGATTAAATTCCTCAACCCTGATAGAGAGTATGATTTCAGGGATCTCCGGTGGTGTATGAA

TCCCCCCGAGAGAGTCAAAATTAATTTTGATCAGTTTTGTGAGTACAAAGCTGCGGCTAAGTCAATTGAACATATATTTG

AGTCACCACTCAACAAGTCAAAAAAGCTGCAATCACTGACTCTTGGGCCCGGATCAGGCTGTCTAGGCAGGACAGTAACA

AGAGCTCATTTCTCAGAACTTACACTGACCTTAATGGACCTAGATCTAGAGAAGAAGCACAACGTGTCCTCAGTGTTTAC

CGTAGTTGAAGAGGGATTATTCGGAAGAACATATACAGTCTGGAGATTCGATGCCAGAGATCCAAGCACCGATCTAGGTA

TTGGCCATTTTTTAAGAGTCTTCGAGATTGGACTGGTAAGGGATCTCGGGCTTGGTCCCCCTGTTTTTCAGATGACCAAC

TACCTCACAGTGAACATGAGTGATGACTATCGGAGATGTCTTTTAGCGGTAGGGGAGTTGAAGTTGACAGCTCTATGCAC

CTCATCTGAGACTGTGACACTAAATGAGGGAGGAGTTCCAAAGAGGGAGTCTCTTGCGGTTGTGATACTTAACCTAGCTG

GGCCCACTCTAGGGGGCGAACTATACAGTGTCTTGCCTATCTCTGATCTCATGGTGGAGAAACTCTATTTATCTTCTCAT

AGAGGGGTTATCAAAGATGACGAGGCCAATTGGGTAGTGCCGTCTACCGACGTTCGTGACCTTCAAAACAAAGGAGAATG

TCTGGTGGAAGCATGCAAAACTCGACCTCCTTCATTTTGCAATGGCACAGGATCAGGCCCGTGGTCAGAGGGGAGAATCC

CTGCTTACGGGGTGATCAGGGTCAGTCTTGACTTAGCCAGTGACCCGGGTGTAGTTATCACTTCAGCGTTTGGCCCGCTG

ATACCTCACCTATCCGGCATGGATCTTTACAATAACCCGTTTTCAAGAGCTGTATGGTTGGCTGTACCACCTTATGAGCA

GTCATTTCTAGGAATGATAAATACAATTGGATTCCCCGACAGAGCAGAGGTTATGCCGCACATTTTGACCACAGAGATCA

GAGGCCCTCGAGGTCGTTGCCATGTTCCCATAGAATTGTCCCACAGGGTTGATGACGATATCAAGATCGGGTCCAACATG

GTCATATTGCCGACGATGGACCTGAGGTATATTACAGCCACTTATGATGTTTCCAGGAGCGAGCATGCAATCGTGTATTA

TATCTATGACACAGGTCGCTCATCATCTTACTTCTACCCAGTTCGACTGAATTTCAAAGGCAATCCTCTCTCTCTGAGGA

TAGAGTGTTTCCCTTGGCGTCATAAGGTGTGGTGCTACCATGATTGTCTTATATACAACACCATAACAGATGAAGAAGTC

CATGCGAGAGGGCTGACCGGTATAGAGGTAACATGTAATCCAGTCTGAGCAGAGCTACGACCGTCACTCAGGCAGTCCCT

TGAGTCGTCACTGAGTCCAAGCAGCATAGCCTGGGATACTCAACAGCACAGCCCAGCCAATAATGTTATAAAAAACTTAG

GAGCCAGGGTTGTAGGGGCCATGGACTCCCTATCAGTCAATCAGGTCTTGTACCCTGAGGTTCATCTAGATAGCCCTATT

GTCACAAACAAACTAGTTGCCATCCTTGAGTACTCGGGTATCGACCATAACTATGTTCTTGAAGACCAGACCCTTGTCAA

GAATATTAGGTATAGGCTGGGGTGCGGTTTTTCAAATCAAATGATCATCAATAACAGGAGGGTAGGTGAAACAGTCAATT

CTAAACTTAAAGGCTACCCCCATAATCGTCATATCATATACCCGGATTGCAATAAGGAGTTGTTTTGTATCAAAGATAGC

TGCATATCTAAGAAGCTCTCGGAGCTATTCAAGAAGGGTAATTCCTTGTACTCTAAGATAAGTCACCAGGTACTGGACTG

TCTCAAGAGAGTCAATGGGAAATTAGGCCTGGGCACAGATCTTACCCATGATCTGAAGGAGGGTATCCTTGACCTAGGGT

TGCACATGCATAGCTCTCAATGGTTCGAGACTTTCCTGTTCTGGTTCACTATCAAGACAGAGATGAGATCAATGATCAAA

GAACAGTCCCATATATGCCACAAGAAGAGGTATAACCCAATTTTTGTGTCGGGGGCTGCATTCGAGGTGCTCGTATCACG

GGACCTCGTTGTGATAATTGATAAGAATACCCAGTATGTCTTCTACTTAACATTTGAGCTAGTCCTTATGTATTGTGATG

TCATAGAGGGCAGACTTATGACGGAGACAGCCATGGCCATAGACCAGAGATATTCAGAACTTCTAAGCCGGGTCAGATAC

TTGTGGGATCTTATTGATGGGTTCTTCCCAACACTAGGCAACACCACATACCAAATTGTTGCTCTGCTTGAACCATTATC

GTTGGCTTATCTTCAACTTCAGGATGTCACTCTGGAGTTAAGAGGTGCTTTTTTAGACCATTGCTTCAAAGAACTCTATG

AGATACTGGAGCATTGCGGCATTGACACAGAAGGCACCTACAATTCCATCACTGAAGGATTGGATTACGTATTCATCACC

CACGATATACATCTAACTGGGGAGATTTTTTCATTTTTCCGGAGTTTCGGACACCCCCGTCTAGAAGCAGTCACCGCCGC

AGAAAATGTCAGGAAACATATGAACCAACCGAAGGTAATCAGTTATGAGACTATGATGAAAGGGCATGCAGTATTTTGCG

GGATAATCATAAATGGTTTTAGGGACCGACATGGCGGCAGCTGGCCCCCTGTTGCATTGCCAGAACATGCTTCTGCTGCG

ATCCGGAATGCGCAGGCATCAGGTGAGGGATTGACCCATGACCTGTGTATAGACAATTGGAAATCCTTTGTTGGATTCAA

ATTCGGCTGTTTTATGCCACTCAGCCTAGATAGTGATTTGACCATGTACCTCAAAGACAAGGCATTGGCTGCACTGAAAA

ATGAGTGGGACTCAGTTTACCCGAAAGAATACCTCCGTTATAACCCACCTAGAGGGACAGAGTCAAGGCGACTGGTGGAG

GTGTTCCTAAATGACTCCAGCTTTGATCCTTATAACATGATAATGTACGTGGTGAATGGCTCCTACCTTAAAGACCCTGA

GTTTAATCTTTCATACAGCCTAAAAGAGAAGGAGATAAAGGAGACAGGGCGGTTGTTTGCTAAAATGACCTATAAGATGC

GGGCCTGTCAGGTAATCGCTGAAAATCTGATATCGAATGGTGTTGGGAAGTATTTCCGAGACAATGGGATGGCAAAAGAC

GAGCATGACCTAACAAAAGCCCTTCACACCCTGGCAGTCTCAGGTGTTCCCAAGAACAACAAAGATAACCACCGAGGTGG

GCCTCCCAGAAGGACAACAAGCCGAGGTGTGAGATCAAGCCAAGACACCAAAACACAAAATAGAGATAAGGTCCAAGGGG

GGCCTATGTACAACTATTTGCGATGCCAACCAATCAGCCCTGATCAGGGTGAGTCATACGAGACTGTTAGCGCATTCATC

ACCGCTGACCTTAAGAAGTATTGCCTGAATTGGAGATACGAGACAATCAGCATATTTGCACAGAGGCTGAATGAAATATA

TGGACTACCATCCTTCTTTCAATGGTTACACAGGATATTGGAAAAATCCGTACTCTACGTCAGTGACCCACATTGCCCTC

CCGACCTAGATAATCATATCCCTCTGGACAGTGTCCCTAACGCCCAAATATTCATTAAGTACCCAATGGGCGGAATAGAA

GGTTATTGCCAAAAACTATGGACAATTAGTACTATACCATACTTGTATCTAGCAGCCTATGAGAGCGGAGTAAGAATCGC

CTCACTAGTGCAGGGTGACAATCAGACAATTGCAGTGACAAAAAGAGTTCCAAGTTCTTGGCCTTATTCACTCAAAAAGA

GGGAGGCATCCAAAGCCGCTCAAAATTACTTCGTAGTCTTAAGGCAGAGATTGCACGATGTAGGTCATCACTTAAAGGCT

AATGAAACCATAGTATCTTCTCACTTTTTTGTATATTCCAAGGGGATTTATTATGACGGCCTGCTAGTTTCACAATCACT

AAAGAGCATCGCCAGATGTGTCTTCTGGTCCGAGACTATAGTGGATGAAACCAGAGCGGCTTGCAGCAATATTGCAACAA

CTGTCGCTAAGAGTATAGAGAGGGGTTATGATAGGTACCTTGCATATTCTTTGAATATCCTCAAGATTTTCCAACAGATC

CTTATATCCCTTAACTTCACTATTAACACAACAATGACTCAGGATGTCGTGGCACCGATCATCGAGAACGGTGATTTGCT

GATAAGGATGGCACTCTTGCCAGCACCCATCGGGGGTCTCAATTATCTTAACATGAGCAGGTTATTTGTGAGAAATATTG

GTGACCCGGTCACCTCCTCTATAGCCGACCTGAAGAGGATGATAGATGCTGGGCTAATGCCAGAAGAAACACTGCATCAA

GTGATGACCCAGACCCCGGGAGAATCATCCTACCTCGATTGGGCAAGTGACCCTTATTCTGCCAACCTACCCTGCGTACA

GAGTATAACTCGCCTTCTCAAAAACATCACTGCACGATATATTTTAATCAGCAGTCCAAACCCAATGCTGAGAGGATTGT

TTCATGAGGGGAGTAGAGATGAGGACGAAGAGCTTGCAAGTTTCTTAATGGATCGGCATATAATAGTTCCAAGAGCTGCA

CATGAAATCTTAGACCATAGCATAACCGGGGCAAGAGAAGCTATAGCCGGGATGTTGGACACCACCAAGGGTCTGATTAG

AACAAGTATGAGACGGGGTGGCCTCACCCCTCGAGTCTTAGCCCGCCTTTCCAATTATGATTATGAACAATTCCGATCCG

GGATAACACTATTGACAAAGAAAGGGCAGTGTTATCTCATTGACAAGGACTCGTGCTCGGTGCAGCTCGCTATAGCTCTG

AGGAGCCACATGTGGGCTAGGTTGGCTCGCGGGAGACCAATCTATGGGTTGGAGGTGCCCGATATATTAGAATCGATGAA

CGGCTACCTTATCAAACGCCACGAGTCTTGTGCTATCTGTGAAACGGGCTCAAGTCACTACGGATGGTTTTTCGTCCCTG

CAGGGTGCCAGCTTGACGATGTATCTAGAGAGACTTCGGCTCTTCGTGTACCTTATGTCGGATCAACAACTGAAGAAAGG

ACGGACATGAAACTTGCTTTTGTTAGGTCTCCGAGTCGATCCCTCAAATCAGCAGTCAGAATTGCAACAGTTTACTCATG

GGCCTACGGGGATGATGAGAAATCATGGAGTGAAGCCTGGATGCTAGCTAGGCAGAGAGCCAATATCACCTTAGATGAAT

TAAGAATGATCACTCCAGTCTCTACATCCACCAACCTAGCCCACCGGTTGAGGGATCGGAGCACCCAGGTGAAATACTCG

GGGACATCCCTTGTGAGGGTTGCAAGATACACAACCATCTCCAATGATAACCTGTCATTTGTGATATCTGAGAAAAAAGT

GGATACCAACTTCATTTATCAGCAAGGGATGCTGCTCGGTCTTGGGATCCTTGAGAACCTCTTTAGGCTAGAGGCCACCA

CCGGAGTGTCCAACACAGTGCTACACCTGCACGTGGAAACAGAATGTTGTGTTGTACCTATGGTGGATCACCCAAGGATA

CCGAGTCTCCGTAATATAAAAGTTACGGATGAGCTATGCACAAACCCTTTGATCTACGACCGGTCTCCCATTATAGAGCA

CGATGCAACCCGACTATACTCACAGAGCCACAGGAGACATTTGGTGGAGTTTGTTACATGGTCAACAAGCCAGCTTTACC

ATATACTGGCTAAGTCTACAGCAATGTCCATGATCGAGTTGATCACGAGATTCGAGAAAGATCACATGAATGAAATAGCC

GCCCTGATTGGCGATGACGACATCAACAGCTTCATCACAGAATTCTTGCTAGTAGAGCCTAGATTATTTATAGTTTACCT

TGGTCAGTGTGCTGCCATCAATTGGGCCTTTGATATACATTATCATCGGCCCTCAGGCAAGTACCAGATGGGGGAACTCC

TCTACTCTCTGCTCTCTCGTATGAGCAAAGGAGTATATAAGATCTTCACCAATGCTTTGAGCCACCCCAAAGTTTACAAG

AAATTTTGGCGAAGTGGTGTCATTGAGCCGATTCATGGCCCGTCTTTGGATACACAGAATTTACATGTCACTGTCTGTGA

CATGATATATGGGTCCTATGTCACCTATTTGGATCTTTTGCTGAATGATGAGCTAGATGATTATCCGTATTTGCTCTGCG

AGAGTGATGAGGACGTGGTCACAGACAGGTTCGACAACATTCAAGCCAAACATCTCTGTGTATTGGCCGATGTATATTGC

AGCTCCAAAAGGTGTCCTTCGATAATCGGGATGTCTCCTATAGAAAAATGTACCATTCTCACACATTACATCAAGGGAGA

ATCAGTACAATCCCCGTCTGGGACCTCATGGAACATTGATCCCCTTGTAGTAGATCATTACTCATGCTCTCTGACCTACC

TTCGCCGCGGTTCCATCAAACAAATCAGGTTGAGAGTTGATCCGGGGTTTGTATTCGAGGCGTTGACAGACGTCGACTTT

AAACAGCCTCGCAAAGCTAAGTTAGATATATCGGTCGTTGGGTTGACTGATTTCTCTCCCCCTTGCGACAACGTCGGTGA

TTTTCTAGGGACTATCAACACATTGAGGCACAATCTGCCTGTCACCGGAACCGGGGTCTCGAACTATGAAGTCCACGCTT

ATCGCAGAATTGGCCTGAATTCATCAGCATGTTATAAAGCCGTAGAGGTCTCCACGTTAATCAAGCCATCCTTAGAAGTC

GGAGAGCATGGATTGTTCTTAGGAGAAGGTTCCGGTTCAATGCTGGCTGCGTACAAGGAAGTTCTTAAATTAGCAAATTG

TTATTACAACAGCGGGGTAACAGCGGAGGGCAGAGCCGGACAGAGGGAAATATCTCCCTATCCCTCAGAGATGAGCCTAG

TAGAGAATCAAATGGGGATAGAGCGGAGTGTTAAAGTGCTGTTCAACGGCAAACCTGAAGTAACCTGGGTGGGGACCACC

GATTGCTACAAGTACATAATCAGTAACATCCAAACCTCTAGTCTAGGTTTCATACATTCAGATATTGAGACACTCCCAAC

CAAAGATGCTGTTGAGAAATTAGAAGAGTTTGCCTCCATCCTATCCTTATCCTTGATTTTGGGAAAAATCGGCTCTATTA

CAGTTGTCAAAATTATGCCCATTAGCGGGGATTTTACCCAAGGCTTCATAGCCTATGCCATCCAATATTTTAGGGAGAGC

CTGCTTGCCTATCCGAGATATAGTAACTTCATCTCGACTGAGTGTTACTTTATTATGATAGGATTAAAGGCCAATCGATT

GATAAACCCAGAAGCCATTAAGCAAAGCATAATCAGGGCAGGAGTCAGGACTGCACCAGGACTTGTGAGCCATATATTAT

CAGAGAAACAGAAAGGTTGTATTCAATCTTTCTTGGGTGATCCTTATATCCAAGGAGACTTCAATAAGCACCTTAAATCT

CTAACCCCTATTGAGAAGATCCTGGTAAATTGTGGTCTCTCTATCAATGGCACAAAAGTCTGTAGAGATCTAATCCACCA

TGATATCGCCTCTGGTCCAGACGGTCTTATGAGCTCTACAATCATTTTATATAGGGAATTGGCTCATTTCAAAGATAATA

TAAGGAGTCAGCACGGAATGTTCCACCCCTACCCAGTACTGGCCAACAGCAGGCAACGTGAATTAATCCTTCGAATAGCC

AAGAAATTCTGGGGGTATGTCTTGCTATATTCTGATGACCCGGCACTAATCAGACAAACAATCAAGAATTTGAAACGGAA

TCATCTAACCTTTGACTTACATAGTAACCCGTTTATTAAAGGGCTATCCAAAGCTGAGAAACTGCTGGTGCGGACGAGTT

CACTAAGAAGGGAATGGTTGTTCACTCTCGATACGAAAGAAGTGAAAGAGTGGTTCAAATTGGTGGGTTACAGTGCACTC

ATCAGAGGCTAGTTAGCTATACATCTGCCCCCTTCTCCTCCGCCATGAGCCCCCACTGACGATCCAAAAGATTAAAGAAA

ACTACATATTGGATAAGTATCTATTCCCAGCTTTGTCTGGT

>MK991799.1/goat/Ethiopia/2014

ACCAAACAAAGTTGGGTAAGGATCGATCTTATAATAACCATAGACTGGCAAACTTAGGAGTAAAGATCCTACTGTCGGGG

GGAGGAGGAGGAGCAAGATCTTTGATTATGGCGACTCTCCTTAAAAGCTTAGCATTGTTCAAAAGGAACAAAGACAAAGC

GCCGACGGCATCAGGTTCAGGGGGGGCCATCCGGGGGATTAAGAATGTTATCATAGTCCCGATTCCCGGAGACTCTTCCA

TCATTACCCGTTCAAGACTGCTCGACAGGCTTGTCAGATTGGCCGGAGATCCTGACATCAACGGGTCAAAGCTGACCGGC

GTGATGATCAGCATGTTATCTTTGTTCGTAGAGTCTCCCGGGCAATTGATACAGCGAATCACAGATGATCCAGATGTTAG

TATCCGCCTTGTTGAGGTAGTTCAAAGTACTAGGTCCCAGTCCGGGTTGACCTTTGCATCACGTGGTGCTGATTTGGACA

ACGAGGCAGACATGTATTTTTCAACTGAGGGGCCCTCGAGTGGAGGTAAGAAAAGGATCAACTGGTTTGAGAACAGAGAA

ATAATAGACATAGAGGTGCAGGATCCAGAAGAGTTCAATATGTTGTTGGCCTCCATACTGGCACAAGTCTGGATCCTTCT

GGCCAAGGCTGTTACGGCACCGGATACGGCAGCTGACTCAGAACTGAGAAGGTGGGTTAAATACACACAACAGAGGAGAG

TGATTGGGGAATTTCGCCTTGACAAAGGGTGGCTGGACGCAGTCCGCAACAGAATTGCAGAAGATCTATCACTTCGGCGG

TTCATGGTGTCTCTCATACTTGACATCAAGAGGACCCCCGGCAACAAGCCAAGGATTGCGGAAATGATCTGCGACATTGA

CAACTATATTGTCGAAGCCGGACTCGCCAGTTTCATCCTTACTATCAAGTTTGGTATCGAAACCATGTATCCTGCATTAG

GGCTCCACGAGTTTGCCGGGGAATTGTCCACTATTGAGTCCTTGATGAACTTGTATCAACAGCTAGGAGAGGTTGCACCC

TATATGGTAATTCTAGAGAACTCAATTCAGAACAAGTTTAGCGCAGGAGCTTATCCCCTCCTCTGGAGCTATGCGATGGG

TGTCGGAGTTGAGCTGGAGAACTCAATGGGGGGCTTGAACTTTGGCAGGTCATATTTTGACCCTGCCTATTTTCGTCTTG

GACAGGAGATGGTCAGAAGATCTGCTGGAAAGGTCAGCTCTGTGATTGCGACTGAGCTCGGCATCACAGCAGAGGAAGCT

AAATTAGTCTCGGAAATCGCCTCGCAGACTGGGGACGAAAGAATCGCCAGAGGGACTGGGCCTCGACAGGCGCAGGTCTC

CTTCCTCCAGCACAAAGCAGGAGAGGGGGATTCGCCCGCACCAGCGACCAGAGAAGGGGTCAAAGCTGCGGTCTCGAACG

GATCTGAAGAAAGGGACAGAAAGCAAACACGCCCAGGAAGGCCCAGAGGAGAGACCCCCGGTCAAATGCTCCTGGAAATC

ATGCCAGAGGATGAGGTCTCGCGAGAGTCTGGTCAAAACCCTCGTGAGGCTCAAAGATCAGCCGAGGCACTCTTCAGGCT

GCAGGCCATGGCCAAGATTCTGGAGGACCAGGAGGAGGGAGAAGACAACAGTCAGGTCTATAACGACAAGGATCTCCTCG

GCTAAGCAGACGCACCTTCTGTCGAAATCAGTGACAAGACATCTCCTACCAGTATTATAAAAAACTTAGGACCCAGGTCC

AAGCAAGCACACATCGACACTCCAATCAGTCGAGCGGAGACCACCGATGGCAGAAGAACAAGCATACCATGTCAACAAGG

GGCTGGAATGTATCAAGTCTCTCAAAGCCTCTCCCCCGGATCTATCCACCATCAGAGATACCATCGAAAGCTGGAGAGAG

GGGCTTAGCCCCCCGGGCCGTGCAACACCGAACCCCGATACGTCCGAGGGAGATCATCAGAATATCAACCAATCATGCTC

ACCAACAATCGGACCAAACAAAGTCTACCTGTCTCCTGAATATAATCTCGGATTTAGAAAGATCACTGGCAACGACTGTG

AGGCTGGGCTCGGAGGAGTCCAGGGGAAAGGATCCAACCCTCAAGTACAGCGTTGCTATGTTTATAGCCACGGGGGTGAA

GAGATTGAAGGACTCGAGGATGCTGACTCTCTCGTGGTTCAAGCAGATCCTCCAGTTGCTAACATCTTCAATGGAGGAGA

GGATGGATCTGACGACAGCGATGTGGACTCTGGCCCGGATGATCCCGGCAGAGATACTCTATATGACCGGGGATCTGTTG

CCGGCAATGATGTCGCTAGGTCCACAGATGTCGAAAAACTAGAAGGTGCTGATATTCAAGAAGTTCTTAACTCCCAGAAA

GGCAAAGGAGGGAGATTCCAAGGCGGGAAAACCTTACGAGTCCCGGAAATACCCGATGTCAAGCACTCCAGACCATCAGC

CCAATCAATTAAAAAGGGCACAGACGGGAGCTCAGTCTCATCTGGAACGGTGACAGAGTGTTTATCGATAAGTGGTGCAA

CCCAAGCTGTGCCAGAGTCAAGATGGGAGTCATCAGAGCAAAATGCGTCTGTGGGGAGTGTCCTCAAGTCTGCGAAGAGT

GCAAAGACGATCCAGGGGTTGACACAAGAATCTGGTACCATAGCATCACTGACTCAGCCTAAAGAAAATGACTCCGAGTA

TGAGTGTGAGGATGACCTATTTACAGAGATTCAGGACATCCGTGCAAGCATTGCCAAGATCCATGAAGACAACAAAACTA

TCCTCTCAAAGCTTGATTCTATACTGTTATTGAAAGGAGAAGTCGACACTATCAAGAAACAAATCAGCAAGCAGAATATA

AGTATGTCCACCATTGAGGGCCATCTCTCCAGTATAATGATAGCCATCCCGGGCTTTGGGAAGGATATCAGGGACCCAAC

ATCCGAGGTTGAGCTGAACCCAGATTTAAGACCTATAATAAGCCGTGACTCTGGCAGAGCTCTCGCGGAGGTCCTCAAGA

AACCCGCTGTGGATCGGTCTCAGAAAATTGGAACCAAAACCAACTCCAGCTCAAAGGGCCAGCTTCTTAAGGATCTCCAG

CTAAAACCTGTCGACAAGCAAGCAAGCTCTGCAATCGGGTTTGTCCCATCCGACCATGAATCATCCAGGAATGTCATCCG

CTCCATAATCAAGTCGAGCAAGCTAAGCATTGATCACAAGGACTATCTTCTAGATTTACTAACTGATGTGAAAGGCTCCA

AGGATCTTAAGGAATTCCACAAGATGCTAACAGCAATTCTCGCCAAGCACCCGTAACACACCCTCCAGTCACCATCTCAT

ACTCGGCTGAAAGCGTCCTCTCAATCAGGCTATTACAAAAAACTTAGGAGCAAGGGCAACTGAGCTTCGCAGACAAGATG

ACCGAGATCTACGACTTTGATAAATCAGCATGGGATGTCAAAGGGTCAATTGCCCGCATAGAACCCACCACCTATCACGA

CGGCCGACTGATACCCCAGGTAAGGGTCATCGATCCTGGTCTGGGAGACAGAAAAGATGAGTGCTTCATGTACCTGTTTC

TCCTAGGAGTGATTGAAGATAACGACCCCCTGTCTCCCCCAGTCGGAAGAACTTTTGGTTCTTTACCTCTAGGGGTCGGT

AGGTCAACTGCTAAGCCAGAAGAACTACTAAGGGAGGCCACGGAACTAGACATAGTGGTGAGGCGCACTGCAGGATTAAA

TGAGAAACTGGTATTTTACAACAACACTCCGCTGTCCTTGTTGACGCCCTGGAAGAAGGTTCTGACAACCGGAAGTGTGT

TTAGCGCCAACCAGGTTTGCAATGCGGTCAACCTAGTCCCGCTTGATACTCCCCAGAGATTCAGGGTTGTGTACATGAGC

ATAACTAGATTATCAGACAATGGTTACTATAGTGTGCCCAGGAGAATGCTGGAGTTCCGCTCAGCCAATGCAGTAGCCTT

CAATATCTTGGTTACACTAAGAATTGAAAATGGCACAAACTCTAGAAGATACATAGTCGGCTCATGGGAGAATCCAGAGG

TCACATTTATGGTACACGTGGGCAACTTTAGAAGAAAGAAGAACGAAGTATACTCTGCTGATTACTGCAAAATGAAGATT

GAAAAGATGGGTTTAGTTTTTGCCCTAGGAGGAATAGGTGGAACAAGTCTCCATATTAGAAGCACCGGGAAAATGAGCAA

AACCCTCCATGCACAGCTAGGGTTCAAGAAAATCTTATGTTATCCCCTAATGGATGTTAATGAGGATCTTAACCGATATC

TCTGGCGGGCAGAGTGCCGAATAGTCAAAATCCAAGCCGTCTTACAGCCATCAGTGCCCCAAGAATTCCGTGTCTACGAT

GATGTCATCATCAACGATGATCAGGGCTTGTTCAAGATCCTGTAGTTCATTTGCAAATGAATGAACCCACCCCCAGGAAG

GAGCCCCACCCTCCCCTGCC-AAAAAACACACTCCAAGGCTCCCCGACACAGCGCCAACACAGCCACTCTCCCGATGGGA

CGAGGCTGG-CTCCCCAAAAAACCCAGTCCAAGCGGGGGGG-CCCCCCAACGACCCGCCGGGCGCCGACCGATTAGGCCG

GCAAGTAGCCCCCAACAGGCCCCGACGAAGACCGAGCACCCCATCCCCCC--ACACCGTCTCCCGAACACGCCCAAGGCC

AGAAGCCGGGCACTCCCAGACCCGCCACCGGCGAGACAGAGCCCCAAGGGGGGACCCTCTAAACCAAAGGGACCACCGAG

GAAAGCTCCGCCGAACCAGCACAGACCCGACCCAAGCAAGCCGGACCCCCAACACGAGACCGCCCAGCGCACCCCACCCA

TCATCAAACAAAACTTAGGGGCCAAGTCCGCACGACAGGCCCCAGCCCGCCAGGCGGAGGGCCGCAGAAAGGAAGGAGAC

ACGACCGTCCCAGAGGGAAGACACCCAACCACCGAAACACCCAGCCCCAGGGAAGGAG--CCCCCCCCCACACGGCGAGG

GAACGGGTGAGAGACCCCAAGGAGAACCCGACCGAAAGCCAAGAGCCGGCGACGCCGGCGGCAAACAGGCAGGGCACACA

CCCGACTCCACCAGAGGAGCCGAGGAGAGCCCGACCCCGCGTGGGCCAAACGGCAGGGAATCCAACCCCGCCGCAAGAAG

AGGTGAGGCCCT-CCT-----CCCCCCCGCACACAACCCACAGACAGACCAAGGGAGGACCACCCCCCAACAAGACAGGC

GGCTCTCTCGCAGCCGAGCCGCAGCTCCCCAGAAGAGGACACCCCCGCACGCTCCGAGCACCCCCCCACCCCCCACCACC

GGGACAGGAGCTCCACCACCACTGACCAGAGCCACCGAGAGCCAGCCAGAGGTCCTCTCCAGTCCCGCCCAGCGCCCCCC

CGAAGAGGCCCCGGCAAGGCCCCCCAGCCCGACGACCCCTGCACGCACCGAAAGAGGGACTCGCCCGAGAAAGTCCTGCA

CATCCGTGCGTAAACATCATGACGCGGGTCGCAATCTTGAAATTTCTGTTTCTTTTCCTAAATGCTGTTGCGTGCCAGAT

TCACTGGGGCAATCTATCCAAAATCGGGATCGTAGGAACAGGGAGTGCCAGCTACAAGGTGATGACTAGGCCAAGCCGCC

AAACTCTGGTTATAAAGTTAATGCCAAATATAACAGCCATCGACAATTGTACAAAGTCAGAGATTGCAGAGTACATGAGG

TTGCTGATCACAGTGTTAAAGCCTGTAGAGGATGCTCTGTCAGTGATAACCAAGAATGTAAGACCAGTTCAAACTCTGAC

ACCTGGGCGTAGAACTCGCCGTTTTGCTGGTGCTGTTCTGGCCGGAGTAGCACTTGGAGTTGCGACAGCCGCTCAGATAA

CTGCAGGAGTCGCCCTTCATCAATCATTGATGAACTCCCAAGCAATTGAGAGTTTAAAAACCAGTCTTGAGAAGTCGAAT

CAGGCAATAGAAGAAATCAGACTTGCAAATAAGGAGACCATACTGGCAGTACAGGGCGTCCAGGATTATATCAACAATGA

ACTTGTCCCTTCTGTCCATAGAATGTCATGCGAGCTGGTCGGTCACAAGCTTGGCCTCAAGCTCCTTAGGTACTACACCG

AGATCCTGTCCATATTCGGGCCCAGCCTTCGAGACCCGATAGCTGCAGAAATATCAATCCAGGCACTCAGTTATGCATTA

GGCGGAGACATTAATAAAATTCTGGACAAGCTTGGGTATAGCGGTGGGGATTTCCTTGCCATCCTGGAAAGCAAGGGAAT

AAAGGCGCGAGTCACATATGTGGACACAAGAGATTACTTCATAATTCTTAGCATCGCCTACCCAACCTTATCTGAGATCA

AGGGAGTGATAGTTCATAAGATAGAAGCTATAACATACAACATTGGGGCACAGGAGTGGTATACTACTATCCCTAAATAT

GTAGCCACTCAGGGGTATCTGATATCGAACTTTGATGAGACGTCTTGCGTATTCACTCCAGAGGGGACAGTTTGCAGCCA

GAATGCGTTGTACCCAATGAGCCCATTGCTTCAGGAATGTTTCAGGGGGTCAACAAAATCGTGCGCTAGAACCCTAGTTT

CAGGGACCATAAGTAATAGATTTATCCTATCAAAAGGGAACCTGATTGCAAATTGCGCGTCAGTTTTGTGCAAATGTTAC

ACAACGGAGACAGTTATCAGCCAAGATCCTGACAAACTACTAACTGTTGTAGCATCCGACAAGTGTCCTGTAGTTGAGGT

GGACGGAGTGACAATACAGGTCGGCAGTCGAGAGTATCCGGATTCAGTATACTTACACAAGATAGACTTAGGTCCAGCCA

TCTCCCTAGAAAAACTGGATGTAGGCACCAATTTAGGCAATGCAGTCACAAGACTGGAGAATGCAAAGGAGCTTCTAGAT

GCATCAGACCAAATACTGAAGACTGTTAAAGGGGTACCTTTCAGTGGGAATATGTACATAGCACTGGCAGCTTGCATTGG

GGTATCCCTAGGGCTTGTCACATTAATATGCTGCTGTAAGGGGAGGTGCAAGAACAAGGAAATCCCGATCTCCAAAATCA

ACCCAGGGCTCAAACCCGACCTGACCGGGACCTCAAAGTCGTACGTGAGATCACTGTAGTCAGAATCACCCGAATCATCC

GGCATCACACACATACATGTACGACACAAGCAGTCAGAGGATGCAGGAAGCCCAGTCTCCGATCACCGACCTGACCCTAC

TCCAAGCTCCACCCGACATTATTCATCAAACAAAACTTAGGACGAAAGGTCAATCACCATGTCCGCACAAAGGGAAAGGA

TCAATGCCTTCTACAAGGACAATCCTCACAATAAGAACCATAGGGTGATCCTGGATAGAGAACGCTTGGTCATTGAAAGA

CCCTACATCTTGCTCGGAGTCCTGCTGGTAATGTTCCTGAGTCTAATCGGACTGCTGGCAATTGCAGGGATCAGGCTTCA

CCGGGCCACCGTTGGAACTTCAGAGATCCAGAATCGGCTGAATACCAATATTGAGTTGACCGAATCTATTGATCCCCAAA

CTAAGGATGTCTTAACCCCCCTTTTTAAAATCATTGGCGATGAAGTCGGCATCAGAATTCCACAGAAATTCAGTGATCTT

GTCAAGTTCATCTCCGATAAGATTAAATTCCTCAACCCTGACAGAGAGTATGATTTCAGGGATCTCCGGTGGTGTATGAA

TCCCCCCGAGAGAGTCAAAATTAATTTTGATCAGTTTTGTGAGTACAAAGCTGCGGCTAAGTCAATTGAACATATATTTG

AGTCACCACTCAACAAGTCAAAAAAGCTGCAATCACTGATTCTTGGGCCCGGATCAGGCTGTCTAGGCAGGACAGTAACA

AGAGCTCATTTCTCAGAACTTACACTGACCTTAATGGACCTAGATCTAGAGAAGAAGCACAACGTGTCCTCAGTGTTTAC

CGTAGTTGAAGAGGGATTATTCGGAAGAACGTATACAGTCTGGAGATTCGATGCCAGAGATCCAAGCACCGATCTAGGTA

TTGGCCATTTTTTAAGAGTCTTCGAGATTGGACTGGTAAGAGATCTCGGGCTTGGTCCCCCTGTTTTCCAGATGACCAAC

TATCTCACAGTGAACATGAGTGATGACTATCGGAGATGCCTTTTAGCGGTAGGGGAGTTGAAGTTGACAGCTCTATGCAC

CTCATCTGAGACTGTGACACTGAGTGAGGGAGGAGTTCCAAAGAGGGAGCCTCTTGCGGTTGTGATACTCAACCTAGCCG

GGCCCACTCTAGGGGGCGAACTATACAGTGTCTTGCCTACCTCTGATCTCATGGTGGAGAAACTCTATTTATCTTCTCAT

AGAGGGGTTATCAAAGATGACGAAGCCAATTGGGTAGTGCCGTCTACCGATGTTCGTGATCTTCAAAACAAAGGAGAATG

TCTGGTGGAAGCATGCAAGACTCGACCTCCTTCATTTTGCAATGCCACAGGATCAGGCCCGTGGTCAGAGGGGAGAATCC

CTGCCTACGGGGTGATCAGGGTCAGTCTTGACTTAGCCAGTGACCCGGGTGTAGTTATCACTTCAGTGTTTGGCCCGCTG

ATACCTCACCTATCCGGCATGGATCTTTACAATAACCCGTTTTCAAGAGCTGTATGGTTGGCTGTACCACCTTATGAGCA

GTCATTTCTAGGAATGATAAATACAATTGGATTCCCCAACAGAGCAGAGATTATGCCGCACATTTTGACCACAGAGATTA

GAGGTCCTCGAGGTCGTTGCCATGTTCCCATAGAATTATCCCGCAGGGTTGATGACGATATCAAGATCGGGTCCAACATG

GTCATATTGCCGACGATGGACTTGAGGTATATTACAGCCACTTATGATGTTTCCAGGTGCGAGCATGCAATCGTGTATTA

TATCTATGACACCGGTCGCTCATCATCTTACTTCTACCCAGTTCGACTGAATTTCAAAGGCAATCCTCTCTCTCTGAGGA

TAGAGTGTTTCCCTTGGCGTCATAAGGTGTGGTGCTACCATGATTGTCTTATATACAACACCGTAACAGATGAAGAAGTC

CATGCGAGAGGGCTGACCGGTATAGAGGTAACATGTAATCCAGTCTGAGCAGAGCTACGACCGTCACTCAGGCAGTCCCT

TGAGTCGTCACTGAGTCCAAGCAGCATAGCCTGGGATACTCAATAGCACAGCCCAGCCAATAATGTTATAAAAAACTTAG

GAGCCAGGGTTGTAGGGGCCATGGACTCCCTATCAGTCAATCAGGTCTTGTACCCTGAGGTTCATCTAGATAGCCCTATT

GTCACAAACAAACTAGTTGCCATCCTTGAGTACTCGGGTATCAACCATAACTATGTTCTTGAAGACCAGACCCTTGTCAA

GAATATTAGGTATAGACTGGGGTGCGGTTTTTCAAATCAAATGATCATCAATAACAGGGGGGTGGGTGAAACAGTCAATT

CTAAACTTAAAGGCTACCCCCACAATCGTCATATCATATACCCGGATTGCAATAAGGAGTTGTTTTGTATCAAAGATAGC

TGCATATCTAAGAAGCTCTCGGAGCTATTCAAGAAGGGTAATTCCTTGTACTCTAAGATAAGTCACCAGGTACTGGACTG

TCTCAAGAGAGTCAATGGGAAATTAGGCCTGGGCACAGATCTTACCCATGGTCTGAAGGAGGGTATCCTTGACCTAGGGT

TGCACATGCATAGCTCCCAATGGTTCGAGACTTTCCTGTTCTGGTTCACCATCAAGACAGAGATGAGATCAATGATCAAA

GAACAGTCCCATATATGCCACAAGAAGAGGTATAACCCGATTTTTGTGTCGGGGGATGCATTCCAGGTGCTCGTATCACG

GGACCTCGTTGTGATAATTGATAAGAATACCCAGTATGTCTTCTACTTAACATTTGAGCTGGTCCTTATGTATTGTGATG

TCATAGAGGGCAGACTTATGACGGAGACAGCCATGGCCATAGACCAGAGATATTCAGAACTTCTAAGCCGAGTCAGATAC

TTGTGGGATCTTATTGATGGGTTCTTCCCAACACTAGGCAACACCACATATCAAATTGTTGCTCTGCTTGAACCGTTATC

GTTGGCTTATCTTCAACTTCAGGATGTCACTCTGGAGTTAAGAGGTGCTTTTTTAGACCATTGCTTCAAAGAACTCTATG

AGATACTGGAGCATTGCGGCATTGACACAGAAGGCACCTACAATTCCATCACTGAAGGAATGGATTACGTATTCATCACC

CACGATATACATCTAACTGGGGAGATTTTTTCATTTTTCCGGAGTTTCGGACACCCCCGTCTAGAAGCAGTCACCGCTGC

AGAAAATGTCAGGAAACATATGAACCAACCGAAGGTAATCAGTTATGAGACTATGATGAAAGGGCATGCAGTATTTTGCG

GGATAATCATAAATGGTTTTAGGGACCGACATGGCGGTAGCTGGCCCCCTGTTGCATTGCCAGAACATGCTTCTGCTGCG

ATCCGGAATGCGCAGGCATCAGGTGAGGGATTGACCCATGACCTGTGTATAGACAATTGGAAATCCTTTGTTGGATTCAA

ATTTGGCTGCTTTATGCCACTCAGCCTAGATAGTGATTTGACCATGTACCTCAAAGACAAGGCATTGGCTGCACTGAAGA

ATGAGTGGGATTCAGTTTACCCGAAAGAATATCTCCGTTATAACCCACCTAGAGGGACAGAGTCAAGGCGACTGGTGGAG

GTGTTCCTAAATGACTCCAGCTTTGATCCTTATAACATGATAATGTACGTGGTGAATGGCTCCTACCTTAAAGACCCTGA

GTTTAATCTTTCATACAGCCTAAAAGAGAAGGAGATAAAGGAGACAGGGCGGTTGTTTGCTAAAATGACCTATAAGATGC

GGGCCTGTCAGGTAATCGCTGAAAATCTGATATCGAATGGTGTTGGGAAGTATTTCCGAGACAATGGGATGGCAAAAGAT

GAGCATGATCTAACAAAAGCCCTTCACACCCTGGCAGTCTCAGGTGTTCCCAAGAACAACAAAGATAACCACCGAGGTGG

GCCTCCCAGAAGGACAACAAGCCGAGATGTGAGATCAAGCCAAGACACCAAAACACAGAATAGAGATAAGGTCCAAGGGG

GGCCTATGTACAACTATTTGCGATGCCAACCAATCAGCCCTGATCAGGGTGAGTCATACGAGACTGTCAGCGCATTCATC

ACCGCTGACCTTAAGAAGTATTGCCTGAATTGGAGATATGAGACAATCAGCATATTTGCACAGAGGCTGAATGAAATATA

TGGACTACCATCCTTCTTTCAATGGTTACACAGGATATTGGAAAAATCCGTACTCTACGTCAGTGACCCACATTGCCCTC

CCGACCTAGATAATCATATCCCTCTGGACAATGTCCCTAATGCCCAAATATTCATTAAGTACCCAATGGGCGGAATAGAA

GGTTATTGCCAAAAACTATGGACAATCAGTACTATACCATACTTGTATCTAGCAGCCTATGAGAGCGGAGTAAGAATCGC

CTCACTAGTGCAGGGTGACAATCAGACAATTGCAGTGACAAAAAGAGTTCCAAGTTCTTGGCCTTATTCACTAAAAAAGA

GGGAGGCATCCAAAGCCGCTCAAAATTACTTCGTAGTCTTAAGGCAGAGATTGCACGATGTAGGTCATCACTTAAAGGCT

AATGAAACCATAGTATCTTCTCACTTTTTTGTATATTCCAAGGGGATTTATTATGACGGCCTGCTAGTTTCACAATCACT

AAAGAGCATCGCCAGATGTGTCTTCTGGTCCGAGACTATAGTGGATGAAACCAGAGCGGCTTGCAGCAATATTGCAACAA

CTGTCGCTAAGAGTATAGAGAGGGGTTATGATAGGTACCTTGCATATTCTTTGAATATCCTCAAGATTTTCCAACAGATC

CTTATATCCCTTAACTTCACTATTAACACAACAATGACTCAGGATGTCGTGGCACCGATCATCGAGAACGGTGATTTGCT

GATAAGGATGGCACTCTTGCCAGCACCCATCGGGGGTCTCAATTATCTTAACATGAGCAGGTTATTTGTGAGAAATATCG

GTGACCCGGTCACCTCCTCTATAGCCGACCTGAAGAGGATGATAGATGCTGGGCTAATGCCAGAAGAAACACTGCATCAA

GTGATGACCCAGACCCCGGGAGAATCATCCTACCTCGATTGGGCAAGTGACCCTTATTCTGCCAACCTACCCTGTGTACA

GAGTATAACTCGCCTTCTCAAAAACATCACTGCACGATATATTTTAATCAGCAGTCCAAACCCAATGCTGAGAGGATTGT

TTCATGAGGGGAGTAGAGATGAGGACGAAGAGCTTGCAAGTTTCTTAATGGATCGGCATATAATTGTTCCAAGAGCTGCA

CATGAAATCTTAGACCATAGCATAACCGGGGCAAGAGAAGCTATAGCCGGGATGTTGGACACCACCAAGGGTCTGATTAG

AACAAGTATGAGACGGGGTGGCCTAACCCCTAGAGTCTTAGCCCGCCTTTCCAATTATGATTATGAACAATTCCGATCCG

GGATAACACTATTGACAAAGAAAGGGCAGTGTTATCTCATTGACAAGGACTCGTGCTCGGTGCAGCTCGCTATAGCTCTA

AGGAGCCACATGTGGGCTAGGTTGGCTCGCGGGAGACCAATCTATGGGTTGGAGGTGCCCGATATATTAGAATCGATGAC

CGGCTACCTTATCAAACGCCACGAGTCTTGTGCTATCTGTGAAACGGGCTCAAGTCACTACGGATGGTTTTTCGTCCCTG

CAGGGTGCCAGCTTGACGATGTATCTAGAGAGACTTCGGCTCTTCGTGTACCTTATGTCGGATCAACAACTGAAGAAAGG

ACGGACATGAAACTTGCTTTTGTTAGATCTCCAAGTCGATCCCTCAAATCAGCAGTCAGAATTGCAACAGTTTACTCATG

GGCCTACGGGGATGATGAGAAATCATGGCGTGAAGCCTGGATGCTAGCTAGCCAGAGAGCCAATATCACCTTAGATGAAT

TAAGAATGATCACTCCAGTCTCTACATCCACCAACCTAGCCCACCGGTTGAGGGATCGGAGCACCCAGGTGAAATACTCG

GGGACATCCCTTGTGAGGGTTGCAAGATACACAACCATCTCCAATGATAACCTGTCATTCGTGATATCTGAGAAAAAAGT

GGACACCAACTTCATTTATCAGCAAGGGATGCTGCTCGGTCTTGGGATCCTTGAGAACCTCTTTAGGCTAGAGGCCACCA

CCGGAGTGTCCAACACAGTGCTACACCTGCACGTGGAAACAGAATGTTGCGTTGTACCCATGGTGGATCACCCTAGGATA

CCGAGTCTCCGTAATATAAAAGTTACGGATGAGCTATGCACAAACCCTTTGATCTACGACCGGTCTCCCATTATAGAGCA

CGATGCAACCCGACTATACTCACAGAGCCACAGGAGACATTTGGTGGAGTTTGTTACATGGTCAACAAGCCAGCTTTACC

ATATACTGGCTAAGTCTACAGCCATGTCCATGATCGAATTGATCACGAGATTCGAGAAAGATCACATGAATGAAATAGCC

GCCCTGATTGGCGATGACGACATCAACAGCTTCATCACAGAATTTTTGCTAGTAGAGCCTAGATTATTTATAGTTTACCT

TGGTCAGTGTGCTGCCATCAATTGGGCCTTTGATATACATTATCATCGGCCCTCAGGCAAGTACCAGATGGGGGAACTCC

TCTACTCTATGCTCTCTCGTATGAGCAAAGGAGTATATAAGATCTTCACCAATGCTTTGAGCCACCCCAAAGTTTACAAG

AAATTTTGGCGAAGTGGTGTCATTGAGCCGATTCATGGCCCGTCTCTGGATACACAGAATTTACATGTCACTGTCTGTGA

CATGATATATGGGTCCTATGTCACCTATTTGGATCTTTTGCTGAATGATGAGCTAGATGACTATCCGTATTTGCTCTGCG

AGAGTGATGAGGACGTGGTCACAGACAGGTTCGACAACATTCAAGCCAAACATCTCTGTGTATTGGCTGATGTATATTGC

AGCTCCAAGAGGTGTCCTTCGATAATCGGGATGTCTCCTATAGAAAAATGTACCATTCTCACACATTACATCAAGGGAGA

ATCAGTACAATCCCCGTCTGGGACCTCATGGAACATTGATCCCCTTGTAGTAGATCATTACTCATGTTCTCTGACCTACC

TTCGCCGCGGTTCCATCAAACAAATCAGGTTGAGAGTTGATCCGGGGTTTGTATTCGAGGCGTTGATCGACGTCGACTTT

AAACAGCCTCGCAAAGCTAAGTTAGATATATCGGTCGTTGGGTTGACTGATTTCTCTCCCCCTTGCGACAACGTCGGTGA

TTTTCTAGGGACTATCAACACATTGAGGCACAATCTGCCTGTCACCGGAACCGGGGTCTCGAACTATGAAGTCCACGCTT

ATCGTAGAATTGGCCTGAATTCATCAGCATGTTATAAAGCCGTAGAGATCTCCACGTTGATCAAGCCATCCTTAGAAGTC

GGAGAGCATGGATTGTTCTTAGGAGAAGGTTCCGGTTCAATGCTGGCTGCGTACAAGGAAGTTCTTAAATTAGCAAATTG

TTATTACAACAGCGGGGTAACAGCGGAGGGCAGAGCCGGACAGAGGGAAATATCTCCCTATCCCTCAGAGATGAGCCTAG

TAGAGAATCAAATGGGGATAGAGCGGAGTGTTAAAGTGCTGTTCAACGGCAAACCTGAAGTAACCTGGGTGGGGACCACC

GATTGCTACAAGTACATAATCAGTAACATCCGAACCTCTAGTCTAGGTTTCATACATTCAGATAATGAGACACTCCCAAC

CAAAGATGCTGTTGAGAAATTAGAAGAGTTTGCCTCTATCCTATCCTTATCCTTGATTTTGGGAAAAATCGGCTCTATTA

CAGTTGTCAAAATTATGCCCATTAGCGGGGATTTTACCCAAGGCTTCATAGCCTATGCCATCCAATATTTTAGGGAGAGC

CTGCTTGCCTATCCGAGATATAGTAACTTCATCTCGACTGAGTGTTACCTTATTATGATAGGATTAAAGGCCAATCGATT

GATAAACCCAGAAGCCATTAAGCAAGGCATAATCAGGGCAGGAGTCAGGACTGCACCAGGACTTGTGAGCCATATATTAT

CAGAGAAACAGAAAGGTTGTATTCAATCTTTCTTGGGTGATCCTTATATCCAAGGAGACTTCAATAAGCACCTTAAATCT

CTAACCCCTATTGAGAAGATCCTGGTAAATTGTGGTCTCTCTATCAATGGCACAAAAATCTGTAGAGATCTAATCCACCA

TGATATCGCCTCCGGTCCAGACGGTCTCATGAGCTCTACAATCATTTTATACAGGGAATTGGTTCATTTCAAAGATAATA

TAAGGAGTCAGCACGGAATGTTCCACCCCTACCCAGTACTGGCCAACAGCAGGCAACGTGAATTAATCCTTCGAATAGCC

AAGAAATTCTGGGGGTATGTCTTGCTATATTCTGATGACCCGGCACTAATCAGACAGATAATCAAGAACTTGAAACGGAA

TCATCTAACCTTTGACTTACATAGTAACCCGTTTATTAAAGGGCTATCCAAAGCTGAGAAACTGCTGGTGCGGACGAGTT

CACTAAGAAGGGAATGGTTGTTCACTCTCGATACGAAAGAAGTAAAAGAGTGGTTCAAATTGGTGGGTTACAGTGCACTC

ATCAGAGGCTAGTTAGCTATACATCTGCCCCCTTCTCCTCCGCCATGAGCCCCCACTGACGATCCAAAAGATTAAAGAAA

ACTACATATTGGATAAGTATCTATTCCCAGCTTTGTCTGGT

>MK991800.1/goat/Ethiopia/2017

ACCAAACAAAGTTGGGTAAGGATAGATCTTATAATAACCATAGACTGGCAAACTTAGGAGTAAAGATCCTACTGTCGGGG

GGAGGAGGAGGAGCAAGATCTTTGATTATGGCGACTCTCCTTAAAAGCTTAGCATTGTTCAAAAGGAACAAAGACAAAGC

GCCGACGGCATCAGGCTCAGGGGGGGCCATCCGGGGGATTAAGAATGTTATCATAGTCCCGATTCCCGGAGACTCTTCCA

TCATTACTCGTTCAAGACTGCTCGACAGGCTCGTCAGATTGGCCGGAGATCCTGACATCAACGGGTCAAAGCTGACCGGC

GTGATGATCAGCATGTTATCTTTGTTCGTAGAGTCTCCCGGGCAATTGATACAGCGAATCACAGATGATCCAGATGTTAG

TATCCGCCTTGTTGAGGTAGTTCAAAGTACTAGGTCCCAGTCCGGGTTGACCTTTGCATCACGTGGTGCTGATTTGGACA

ACGAGGCAGACATGTATTTTTCAACTGAGGGGCCCTCGAGTGGAGGTAAGAAAAGGATCAACTGGTTTGAGAACAGAGAA

ATAATAGACATAGAGGTGCAGGATCCAGAAGAGTTCAATATGTTGTTGGCCTCCATACTGGCACAAGTCTGGATCCTTCT

GGCCAAGGCTGTTACGGCACCGGATACGGCAGCTGATTCAGAACTGAGAAGGTGGGTTAAATACACACAACAGAGGAGAG

TGATTGGGGAATTTCGCCTTGACAAAGGGTGGCTGGACGCAGTCCGCAACAGAATTGCAGAAGATCTATCACTTCGGCGG

TTCATGGTGTCTCTCATACTTGACATCAAGAGGACCCCCGGCAACAAGCCAAGGATTGCGGAAATGATCTGTGACATTGA

CAACTATATTGTCGAAGCCGGACTCGCCAGTTTCATCCTTACTATCAAGTTTGGTATCGAAACCATGTATCCTGCATTAG

GGCTCCACGAGTTTGCCGGGGAATTGTCCACTATTGAGTCCTTGATGAACTTGTATCAACAGCTAGGAGAGGTTGCACCC

TATATGGTAATTCTAGAGAACTCAATTCAGAACAAGTTTAGCGCAGGAGCTTATCCCCTCCTCTGGAGCTATGCGATGGG

TGTCGGAGTTGAGCTGGAGAACTCAATGGGGGGCTTGAACTTTGGCAGGTCATATTTTGACCCTGCCTATTTTCGTCTCG

GACAGGAGATGGTCAGAAGATCTGCTGGAAAGGTCAGCTCTGTGATTGCGGCTGAGCTCGGCATCACAGCAGAAGAAGCT

AAACTAGTCTCGGAGATCGCCTCGCAGACTGGGGACGAAAGAATCGCCAGAGGGACTGGGCCTCGACAGGCGCAGGTCTC

CTTCCTCCAGCACAAAGCAGGAGAGGGAGATTCGCCCGCACCAACGACCAGAGAAGGGGTCAAGGCTGCGATCTCGAACG

GATCTGAAGAAAGGGACAGAAAGCAAACACGCCCAGGAAGGCCCAGAGGAGAGACCCCCGGTCAAATGCTCCTGGAAATC

ATGCCAGAGGATGAGGTCTCGCGAGAGTCTGGTCAAAACCCTCATGAGGCTCAAAGATCAGCCGAGGCACTCTTCAGGCT

GCAGGCCATGGCCAAGATTCTGGAGGACCAGGAGGAAGGAGAAGACAACAGTCAGGTCTACAACGACAAGGATCTCCTCG

GCTAAGCAGACGCACCATCTGTCGAAATCAGTGACAAGACATCTCCTACCAGTATTATAAAAAACTTAGGACCCAGGTCC

AAGCAAGCACACATCGACACTCCAATCAGTCGAGCGGAGACCACCGATGGCAGAAGAACAAGCATACCATGTCAACAAGG

GGCTGGAATGTATCAAGTCCCTCAAAGCCTCTCCCCCGGATCTATCCACCATCAAAGATACCATCGAAAGCTGGAGAGAG

GGGCTTAGCCCCCCGGGCCATGCAACACCGGACCCTGATACGTCCGAGGGAGATCATCAGAATATCAACCAATCATGCTC

ACCAACAATCGGACCAAACAAAGTCTACCTGTCTCCTGAAGATAATCTCGGATTTAGAAAGATCACTGGCAACGACTGTG

AGGCTGGGCTCGGAGGAGTCCAGGGGAAAGGATCCAACCCTCAAGTACAGCGTTGCTATGTTTATAGCCACGGGGGTGAA

GAGATTAAAGGACTCGAGGATGCTGACTCTCTCGTGGTTCAAGCAGATCCTCCAGTTGCTAACATCTTCAATGGAGGAGA

GGATGGATCTGACGACAGCGATGTGGACTCTGGCCCAGATGATCCCGGCAGAGATACTCTATATGACCGGGGATCTGTTG

CCAGCAATGATGTCGCTAGGTCCACAGATGTCGAAAAACTAGAAGGTGCTGATATTCAAGAAGTTCTGAACTCCCAGAAA

GGCAAAGGAGGGAGATTCCAAGGCGGGAAAACCTTACGAGTCCCGGAAATACCCGATGTCAAGCACTCCAGACCATCAGC

CCAATCAATTAAAAAGGGCACAGACGGGAGCTCAGTCTCATCTGGAACGGTGACAGAGTGTTTATCGATAAGTGGTGCAA

CCCAAGCTGTGCCAGAGTCAAGATGGGAGTCATCAGAGCAAAATGCGTCTGTGGGGAGTGTCCTCAAGTCTGCGAAGAGT

GCAAAGACGATCCAGGGGTTGACACAAGAATCTGGTACCATAGCATCACTGACTCAGCCTAAAGAAAATGACACCGAGTA

TGAGTATGAGGATGACCTATTTACAGAGATTCAGGACATCCGTGCAAGCATTGCCAAGATCCATGATGACAACAAAACTA

TCCTCTCAAAGCTTGATTCTATACTGTTATTGAAAGGAGAAGTCGACACTATCAAGAAACAAATCAGCAAGCAGAATATA

AGTATATCCACCATTGAGGGCCATCTCTCCAGTATAATGATAGCCATCCCGGGCTTTGGGAAGGATATCAGGGACCCAAC

ATCCGAGGTTGAGTTGAACCCAGATTTAAGACCTATAATAAGCCGTGACTCTGGCAGAGCTCTCGCGGAGGTCCTCAAGA

AACCTGCTGTGGATCGGTCTCAGAAAATTGGAACCAAAACCAACTCCAGCTCAAAAGGCCAGCTTCTTAAGGATCTCCAG

CTAAAACCTGTCGACAAGCAAGCAAGCTCTGCAATCGGGTTTGTCCCATCCGACCATGAATCATCCAGGAATGTCATCCG

CTCCATAATCAAGTCGAGCAAGCTAAGCATTGATCACAAGGACTATCTTCTAGATTTACTAACTGATGTGAAAGGCTCCA

AGGATCTTAAGGAATTCCACAAGATGCTAACAGCAATTCTCGCCAAGCACCCGTAACACACCCTCCAGTCACCATATCAT

ACTCGGCTGAAAGCGTCCTCTCAATCAGGCTATTACAAAAAACTTAGGAGCAAGGGCAACTGAGCTTCGCAGACAAGATG

ACCGAGATCTACGACTTTGATAAATCAGCATGGGATGTGAAAGGGTCAATTGCCCGCATAGAACCCACCACCTATCACGA

CGGCCGACTGATACCCCAGGTAAGGGTCATCGATCCTGGTCTGGGAGACAGAAAAGATGAGTGCTTCATGTACCTGTTTC

TCCTAGGAGTGATTGAAGATAACGACCCCCTGTCTCCCCCAGTCGGAAGAACCTTTGGTTCTTTACCTCTAGGGGTCGGT

AGGTCAACTGCTAAGCCAGAAGAACTACTAAGGGAGGCCACGGAACTAGATATAGTGGTGAGGCGCACTGCAGGATTAAA

TGAGAAACTGGTATTTTACAACAACACTCCGCTGTCCTTGTTAACGCCCTGGAAGAAGGTTCTGACAACCGGAAGTGTGT

TTAGCGCCAACCAGGTTTGCAATGCGGTCAACCTAGTCCCGCTTGATACTCCCCAGAGATTCAGGGTTGTGTACATGAGC

ATAACTAGATTATCAGACAATGGTTACTATAGTGTGCCCAGGAGAATGCTGGAGTTCCGCTCAGCCAATGCAGTAGCCTT

CAATATCTTGGTTACACTAAGAATTGAAAATGGCACAAACTCTAGAAGATACATAGTCGGCTCATGGGAGAATCCAGAGG

TCACATTTATGGTACACGTGGGCAACTTTAGAAGAAAGAAGAACGAAGTATACTCTGCTGATTACTGCAAAATGAAGATT

GAAAAGATGGGTTTAGTTTTTGCCCTAGGAGGAATAGGTGGAACAAGTCTCCATATTAGAAGCACCGGGAAAATGAGCAA

AACCCTCCATGCACAGCTAGGGTTCAAGAAAATCTTATGTTATCCCCTAATGGATGTTAATGAGGATCTTAACCGATATC

TCTGGCGGGCAGAGTGCCGAATAGTCAAAATCCAAGCCGTCTTACAGCCATCAGTACCCCAAGAATTCCGTGTCTACGAT

GATGTCATCATCAACGATGATCAAGGCTTGTTCAAGATCCTGTAGTTCATTTGCAAATGAATGAACCCACCCCCAGGAAG

GAGCCCCCCCCTCCCCTGCC-CAAAAACACACTCCAAGGCTCCCCGACACAGCGCCAACACAGCCACTCTCCCGATGGGA

CGAGGCTGG-CTCCCCAAAAAACCCAGTCCAGGCGGGGGGG-CCCCCCAACGACCCACCGGGCGCCGACCAATCAGGCCG

GCAAGCAGCCCCCAACAGGCCCCGACGAAGACCGAGCACCCCATCACCCC--AAACCGTCTCCCGACCACGCCCAAGGCC

AGAAGCCGGGCACTCCCAGACCCACCACCGGCGAGACGGACCCCCAAGGGGGGACCCTCTAAACCAAAGGGACCACCGAG

GAAAACTCCGCCTAACCAGCACAGACCCGACCCAAGCAAGCCGGACCCCAAACACGAGACCGCCCAGCGCACCCCACCCA

TCATCAAACAAAACTTAGGGGCCAAGTCTGCACGACAGGCCCCAGCCCGCCAGGCGGAGGGCCGCAGAAAGGAAGGAGAC

ACGACCGTCCCAGAAGGAAGACACCCAACCACCGAAACACCCAGCCCCAGGGAAGGAG--CCCCCCCCCACACGGCGAAG

GAACGGGTGAGAGACCCCAAGGAGAACCCGACCGAGAGCCAAGAGCCGGCGACGCCGGCGGCAAACAGGCAGGGCACACA

CCCTACTCCACCAGAGGAGCCGAGGAGAGCCCGACCTCGCGTGGGCCAAACGGCAGGGAATCCAACCCCGCCATAAGAAG

AGGTGAGGCCCT-CTT-----CCCCCCCGCATACAACCCACAGACAGGCCAAGGGAGGACCACCTCCCAACAAGACAGGC

GGCTCCCTCGCAGCCGAGCCGCAGCTCCCCAGAAGAGGACACCCCCGCACGCTCCGAGCACCCCCCCACTCCCCACCACC

GGGACAGGAGCTCCACCACCACTGAGCAGAATCACCGAGAGCCAGACAGAGGTCCTCTCCAGTCCCGCCCAGCGCCCCCC

CGAAGAGGCCCCGGCAAGGCCCCCCAGCCCGACGACCCCTGCACGCACCGAAAGAGGGACTCGCCCGAGAACGTCCTGCA

CATCCATGCGTAAACATCATGACGCGGGTCGCAATCTTGAAATTTCTGTTTCTTTTCCTAAATCCTGTTGCGTGCCAGAT

TCACTGGGGCAATCTATCCAAAATCGGGATCGTAGGAACAGGGAGTGCCAGCTACAAGGTGATGACTAGGCCAAGCCGCC

AAACTCTGGTTATAAAGTTAATGCCAAATATAACAGCCATCGACAATTGTACAAAGTCAGAGATTGCAGAGTACAAGAGA

TTGCTGATCACAGTGTTAAAGCCTGTAGAGGATGCTCTGTCAGTGATAACCAAGAATGTAAGACCAGTTCAAACTCTGAC

ACCTGGGCGTAGAACTCGCCGTTTTGCTGGTGCTGTTCTGGCCGGAGTAGCACTTGGAGTTGCGACAGCCGCTCAGATAA

CTGCAGGAGTTGCCCTTCATCAATCATTGATGAACTCCCAAGCAATTGAGAGTTTAAAAACCAGTCTTGAGAAGTCGAAT

CAGGCAATAGAAGAAATCAGACTTGCAAATAGGGAGACCATACTGGCAGTACAGGGCGTCCAGGATTATATCAACAATGA

ACTTGTCCCTTCTGTCCATAGAATGTCATGCGAGCTGGTCGGTCACAAGCTTGGCCTCAAGCTCCTTAGGTACTACACCG

AGATCCTGTCCATATTCGGGCCCAGCCTTCGAGACCCGATAGCTGCAGAAATATCAATCCAGGCACTCAGTTATGCATTA

GGCGGAGACATTAATAAAATTCTGGACAAGCTTGGGTATAGCGGTGGGGATTTCCTTGCCATCCTGGAAAGCAAGGGAAT

AAAGGCTCGAGTCACATATGTGGACACAAGAGATTACTTCATAATTCTTAGCATCGCCTACCCAACCTTATCTGAGATCA

AGGGAGTGATAGTTCACAAGATAGAAGCTATAACATACAACATTGGGGCACAGGAGTGGTATACTACTATCCCTAAATAT

GTAGCCACTCAGGGGTATCTCATATCGAACTTCGATGAGGCGTCTTGTGTGTTCACTCCAGAGGGGACAGTTTGCAGCCA

GAATGCGTTGTACCCAATGAGCCCATTGCTTCAGGAATGTTTCAGGGGGTCAACAAAATCGTGCGCCAGAACCCTAGTTT

CAGGGACCATAAGTAATAGATTTATCCTATCCAAAGGGAACCTGATTGCAAATTGCGCGTCAGTTTTGTGCAAATGTTAC

ACAACGGGGACAGTTATCAGCCAAGATCCTGACAAGCTACTAACTGTTGTAGCATCCGACAAGTGTCCTGTAGTCGAGGT

GGACGGAGTGACAATACAGGTCGGCAGTCGAGAGTATCCGGATTCTGTATACTTACACAAGATAGACTTAGGTCCAGCCA

TCTCCCTAGAAAAACTGGATGTAGGCACCAATTTAGGCAATGCAGTCACAAGACTGGAGAATGCAAAGGAGCTTCTAGAT

GCATCAGACCAAATACTGAAGACTGTTAAAGGGGTACCTTTCAGTGGGAATATGTACATAGCACTGGCAGCTTGCATTGG

GGTATCCCTAGGGCTTGTCACATTAATATGCTGCTGTAAGGGGAGGTGCAAGAACAAGGAAATCCCGATCTCCAAAATCA

ACCCAGGGCTCAAACCCGACCTGACCGGGACCTCAAAGTCGTACGTGAGATCACTGTAATCAGAATCACCCGAATCATCC

GGCATCACACACATACATGTACGACACAAGCAGTCAGAGGACGCAGGAAGCCCAGTCTCCGATCACCGACCTGACCCTAC

TCCAAGCTCCACCCGACATCATTCATCAAACAAAACTTAGGACGAAAGGTCAATCACCATGTCCGCACAAAGGGAAAGGA

TCAATGCCTTCTACAAGGACAATCCTCACAATAAGAACCATAGGGTGATCCTGGATAGAGAACGCTTGGTCATTGAAAGA

CCCTACATCTTGCTTGGAGTCCTGCTGGTAATGTTCCTGAGTCTAATCGGACTGCTGGCAATTGCAGGGATCAGGCTTCA

CCGGGCCACCGTTGGAACTTCAGAGATCCAGAATCGGCTGAATACCAATATTGAGTTGACCGAATCTATTGATCACCAAA

CTAAGGACGTCTTGACCCCCCTTTTTAAAATCATTGGCGATGAAGTCGGCATCAGAATTCCACAGAAATTCAGTGATCTT

GTCAAGTTCATCTCCGATAAGATTAAATTCCTCAACCCTGACAGAGAGTATGATTTCAGGGATCTCCGGTGGTGTATGAA

CCCCCCCGAGAGAGTCAAAATTAATTTTGATCAGTTTTGTGAGTACAAAGCTGCGGCTAAGTCAATTGAACATATATTTG

AGTCACCACTCAACATGTCAAAAAAGCTGCAATCACTGATTCTTGGGCCCGGATCAGGCTGTCTAGGCAGGACAGTAACA

AGAGCTCATTTCTCAGAACTTACACTGACCTTAATGGACCTAGATCTAGAGAAGAAGCACAACGTGTCCTCAGTGTTTAC

CGTAGTTGAAGAGGGATTATTCGGAAGAACGTATACAGTCTGGAGATTCGATGCCAGAGATCCAAGCACCGATCTAGGTA

TTGGCCATTTTTTAAGAGTCTTCGAGATTGGACTGGTAAGAGATCTCGGGCTTGGTCCCCCTGTTTTCCAGATGACCAAC

TATCTCACAGTGAACATGAGTGATGACTATCGGAGGTGTCTTTTAGCGGTAGGGGAGTTGAAGTTGACAGCTCTATGCAC

CTCATCTGAGACTGTGACACTGAGTGAGGGAGGAGTTCCAAAGAGGGAGCCTCTTGCGGTTGTGATACTCAACCTAGCTG

GGCCCACTCTAGGGGGCGAACTATACAGTGTCTTGCCTACCTCTGATCTCATGGTGGAGAAACTCTATTTATCTTCTCAT

AGAGGGGTTATCAAAGATGACGAGGCCAATTGGGTAGTGCCGTCTACCGATGTTCGTGATCTTCAAACCAAAGGAGAATG

TCTGGTGGAAGCATGCAAGACTCGACCTCCTTCATTTTGCAATGGCACAGGATCAGGCCCATGGTCAGAGGGGAGAATCC

CTGCTTACGGGGTGATCAGGGTCAGTCTTGACTTAGCCAGTGACCCGGGTGTAGTTATCACTTCAGTGTTTGGCCCGCTG

ATACCTCACCTATCCGGCATGGATCTTTACAACAACCCGTTTTCAAGAGCTGTATGGTTGGCTGTACCACCTTATGAGCA

GTCATTTCTAGGAATGATAAATACAATTGGATTCCCCAACAGAGCAGAGGTTATGCCGCACATTTTGACCACAGAGATCA

GAGGTCCTCGAGGTCGTTGCCATGTTCCCATAGAGTTGTCCCGCAGGGTTGATGACGATATCAAGATCGGGTCCAACATG

GTCATATTGCCGACGATGGACTTGAGGTATATTACAGCCACTTATGATGTTTCCAGGAGCGAGCATGCAATCGTGTATTA

TATCTATGACACCGGTCGCTCATCATCTTACTTCTACCCAGTTCGACTGAATTTCAAAGGCAATCCTCTCTCTCTGAGGA

TAGAGTGTTTCCCTTGGCGTCATAAGGTGTGGTGCTACCATGATTGTCTTATATACAACACCATAACAGATGAAGAAGTC

CATGCGGGAGGGCTGACCGGTATAGAGATAACATGTAATCCAGTCTGAGCAGAGCTACGACCGTCACTCAGGCAGTCTCT

TGAGTCGTCACTGAGCCCTAGCAGCATAGCCTGGGATACTCAACAGCACAGCCCAGCCAATAATGTTATAAAAAACTTAG

GAGCCAGGGTTGTAGGGGCCATGGACTCCCTATCAGTCAATCAGGTCTTGTACCCTGAGGTTCATCTAGATAGCCCTATT

GTCACAAACAAACTAGTTGCCATCCTTGAGTACTCGGGTATCAACCATAACTATGTCCTTGAAGACCAGACCCTTGTCAA

GAATATTAGGTATAGACTGGGGTGCGGTTTTTCAAATCAAATGATCATCAATAACAGGGGGGTGGGTGAAACAGTCAATT

CTAAACTTAAAGGCTACCCCCACAATCGTCATATCATATACCCGGATTGCAATAGGGAGTTGTTTTGTATCAAAGATAGC

TGCATATCTAAGAAGCTCTCGGAGCTATTCAAGAAGGGTAATTCCTTGTACTCTAAGATAAGTCACCAGATACTGGACTG

TCTCAAGAGAGTCAATGGGAAATTAGGCCTGGGCACAGATCTTACCCATGGTTTGAAGGAGGGGATCCTTGACCTAGGGT

TGCACATGCATAGCTCTCAATGGTTCGAGACTTTCCTGTTCTGGTTCACTATTAAGACAGAGATGAGATCAATGATCAAA

GAACAGTCCCATATATGCCACAAGAAGAGGTATAACCCGATTTTTGTGTCGGGGGATGCATTCGAGGTGCTCGTATCACG

GGACCTCGTTGTGATAATTGATAAGAATACCCAGTATGTCTTCTACTTAACATTTGAGCTGGTCCTTATGTATTGTGATG

TAATAGAGGGCAGACTTATGACGGAGACAGCCATGGCCATAGACCAGAGATATTCAGAACTTCTAAGCAGGGTCAGATAC

TTGTGGGACCTTATTGATGGGTTCTTCCCAACACTAGGCAACACCACATATCAAATTGTTGCTCTGCTTGAACCATTATC

ATTGGCTTATCTTCAACTTCAGGATGTCACTCTGGAGTTAAGAGGTGCTTTTTTAGACCATTGCTTCAAAGAACTCTATG

AGATACTGGAGCATTGCGGCATTGACACAGAAGGCACCTACAATTCCATCACTGAAGGAATGGATTACGTATTCATCACC

CACGATATACATCTAACTGGGGAGATTTTTTCATTTTTCCGGAGTTTCGGACACCCCCGTCTAGAAGCAGTCACCGCTGC

AGAAAATGTCAGGAAACATATGAACCAACCGAAGGTAATCAGTTATGAGACTATGATGAAAGGGCATGCAGTATTTTGCG

GGATAATCATAAATGGTTATAGGGACCGACATGGCGGTAGCTGGCCCCCTGTTGCACTGCCAGAACATGCTTCTGCTGCG

ATCCGGAATGCGCAGGCATCAGGTGAGGGATTGACCCATGACCTGTGTATAGACAATTGGAAATCCTTTGTTGGATTCAA

ATTTGGCTGCTTTATGCCACTCAGCCTAGATAGTGATTTGACCATGTACCTCAAAGACAAGGCATTGGCTGCACTGAAGA

ATGAGTGGGATTCAGTTTACCCGAAAGAATACCTCCGTTATAACCCACCTAGAGGGACAGAGTCAAGGCGACTGGTGGAG

GTGTTCCTAAATGACTCCAGCTTTGATCCTTATAACATGATAATGTACGTGGTGAATGGCTCCTACCTTAAAGACCCTGA

GTTTAATCTTTCATACAGCCTAAAAGAGAAGGAGATAAAGGAGACAGGGCGGTTGTTTGCTAAAATGACCTATAAGATGC

GGGCCTGTCAGGTAATCGCTGAAAATCTGATATCGAATGGTGTTGGGAAGTATTTCCGAGACAATGGGATGGCAAAAGAT

GAGCATGATCTAACAAAAGCCCTTCACACCCTGGCAGTCTCAGGTGTTCCCAAGAACAACAAAGATAACCACCGAGGTGG

GCCTCCCAGAAAGACAACAAGCCGAGATGTGAGATCAAGCCAAGACACCAAAACACAGAATAGAGATAAGGTCCAAGGGG

GGCCTATGTACAACTATTTGCGATGCCAACCAATCAGCCCTGATCAGGGTGAGTCATACGAGACTGTTAGCGCATTCATC

ACCGCTGACCTTAAGAAGTATTGCCTGAATTGGAGATACGAGACAATCAGCATATTTGCACAGAGGCTGAATGAAATATA

TGGACTACCATCCTTCTTTCAATGGTTACACAGGATATTGGAAAAATCCGTACTCTACGTCAGTGACCCACATTGCCCTC

CCGACCTAGATAATCATATCCCTCTGGACAGTGTCCCCAATGCCCAAATATTCATTAAGTACCCTATGGGCGGAATAGAA

GGTTATTGCCAAAAACTATGGACAATCAGTACTATACCATACTTGTATCTAGCAGCCTATGAGAGCGGAGTAAGAATCGC

CTCACTAGTGCAGGGTGACAATCAGACAATTGCAGTGACAAAAAGAGTTCCAAGTTCTTGGCCTTATTCACTAAAAAAGA

GGGAGGCATCCAAGGCCGCTCAAAATTACTTCGTAGTCTTAAGGCAGAGATTGCACGATGTAGGTCATCACTTAAAGGCT

AATGAAACCATAGTATCTTCTCACTTTTTTGTATATTCCAAGGGGATTTATTATGACGGCCTGCTAGTTTCACAATCACT

AAAGAGCATCGCCAGATGTGTCTTCTGGTCCGAGACTATAGTGGATGAAACCAGAGCGGCTTGCAGCAATATTGCAACAA

CTGTCGCTAAGAGTATAGAGAGGGGTTATGATAGGTACCTTGCATATTCTTTGAATATCCTCAAGATTTTCCAACAGATC

CTTATATCCCTTAACTTCACTATTAACACAACAATGACTCAGGATGTCGTGGCACCGATCATCGAGAACGGTGATTTGCT

GATAAGGATGGCACTCTTGCCAGCACCCATCGGGGGTCTCAATTATCTTAACATGAGCAGGTTATTTGTGAGAAATATCG

GTGACCCGGTCACCTCCTCTATAGCCGACCTGAAGAGGATGATAGATGCTGGGCTAATGCCAGAAGAAACACTGCATCAA

GTGATGACCCAGACCCCGGGAGAATCATCCTACCTCGATTGGGCAAGTGACCCTTATTCTGCCAACCTACCCTGCGTACA

GAGTATAACTCGCCTTCTCAAAAACATCACTGCACGATATATTTTAATCAGCAGTCCAAACCCAATGCTGAGAGGATTGT

TTCATGAGGGGAGTAGAGATGAGGACGAAGAGCTTGCAAGTTTCTTAATGGATCGGCATATAATTGTTCCAAGAGCTGCA

CATGAAATCTTAGACCATAGCATAACCGGGGCAAGAGAAGCCATAGCCGGGATGTTGGACACCACCAAGGGTCTGATTAG

AACAAGTATGAGACGGGGTGGCCTCACCCCTAGAGTCTTAGCCCGCCTTTCCAATTATGATTATGAACAATTCCGATCCG

GGATAACACTATTGACAAAGAAAGGGCAGTGTTATCTCATTGACAAGGACTCGTGCTCGGTGCAGCTCGCTATAGCTCTA

AGGAGCCACATGTGGGCTAGGTTGGCTCGCGGGAGACCAATCTATGGGTTGGAGGTGCCCGATATATTAGAATCGATGAC

TGGCTACCTTATCAAACGCCACGAGTCTTGTGCTATCTGTGAAACGGGCTCAAGTCACTACGGATGGTTTTTCGTCCCTG

CAGGGTGCCAGCTTGACGATGTATCTAGAGAGACTTCGGCTCTTCGTGTACCTTATGTCGGATCAACAACTGAAGAAAGG

ACGGACATGAAACTTGCTTTTGTTAGATCTCCAAGCCGATCCCTCAAATCAGCAGTCAGAATTGCAACAGTTTACTCATG

GGCCTACGGGGATGATGAGAAATCATGGAGTGAAGCCTGGATGCTAGCTAGCCAGAGAGCCAATATCACCTTAGATGAAT

TAAGAATGATCACTCCAGTCTCTACATCCACCAACCTAGCCCACCGGTTGAGGGATCGGAGCACCCAGGTGAAATACTCG

GGGACATCCCTTGTGAGGGTTGCCAGATACACAACCATCTCCAATGATAACCTGTCATTCGTGATATCTGAGAAAAAAGT

GGACACCAACTTCATTTATCAGCAAGGGATGCTGCTCGGTCTTGGGATCCTTGAGAACCTCTTTAGGCTAGAGGCCACCA

CCGGAGTGTCCAACACAGTGCTACACCTGCACGTGGAAACAGAATGTTGCGTTGTACCCATGGTGGATCACCCTAGGATA

CCTAGTCTCCGTAATATAAAAGTTACGGATGAGCTATGCACAAACCCTTTGATCTACGACCGGTCTCCCATTATAGAGCA

CGATGCAACCCGACTATACTCACAGAGCCACAGGAGACATTTGGTGGAGTTTGTTACATGGTCAACAAGCCAGCTTTACC

ATATACTGGCTAAGTCTACAGCCATGTCCATGATCGAATTGATCACGAGATTCGAGAAAGATCACATGAATGAAATAGCC

GCCCTGATTGGCGATGACGACATCAACAGCTTCATCACAGAATTTTTGCTAGTAGAGCCTAGATTATTTATAGTTTGCCT

TGGTCAGTGTGCTGCCATCAATTGGGCCTTTGATATACATTATCATCGGCCCTCAGGCAAGTACCAGATGGGGGAACTCC

TCTACTCTCTGCTCTCTCGTATGAGCAAAGGAGTATATAAGATCTTCACCAATGCTTTGAGCCACCCCAAAGTTTACAAG

AAATTTTGGCGAAGTGGTGTCATTGAGCCGATTCATGGCCCGTCTTTGGATACACAGAATTTACATGTCACTGTCTGTGA

CATGATATATGGGTCCTATGTCACCTATTTGGATCTTTTGCTGAATGATGAGCTAGATGACTATCCGTATTTGCTCTGCG

AGAGTGATGAGGACGTGGTCACAGACAGGTTCGACAACATTCAAGCCAAACATCTCTGTGTATTGGCCGATGTATATTGC

AACTCCAAGAGGTGTCCTTCGATAATCGGGATGTCTCCTATAGAAAAATGTACCATTCTCACACATTACATCAAGGGAGA

ATCAGTACAATCCCCGTCTGGGACCTCATGGAACATTGATCCCCTTGTAGTAGATCATTACTCATGCTCTCTGACCTACC

TTCGCCGCGGTTCCATTAAACAAATCAGGTTGAGAGTTGATCCGGGGTTTGTATTCGAGGCGTTGATCGACGTCGACTTT

AAACATCCTAGCAAAGCTAAGTTAGATATATCAGTCGTTGGGTTGACTGATTTCTCTCCCCCTTGCGACAACGTCGGTGA

TTTTCTAGGGACTATCAACACATTGAGGCACAATCTGCCTGTCACCGGAACCGGGGTCTCGAACTATGAAGTCCACGCTT

ATCGTAGAATTGGCCTGAATTCATCAGCATGTTATAAAGCCGTAGAGATCTCCACGTTGATCAAGCCATCCTTAGAAGTC

GGAGAGCATGGATTGTTCTTAGGAGAAGGTTCCGGTTCAATGCTGGCTGCGTACAAGGAAGTTCTTAAATTAGCAAATTG

TTATTACAACAGCGGGGTAACAGCGGAGGGCAGAGCCGGACAGAGGGAAATATCTCCCTATCCCTCAGAGATGAGCCTAG

TAGAGAATCAAATGGGGATAGAGCGGAGTGTTAAAGTGCTGTTCAACGGCAAACCTGAAGTAACCTGGGTGGGGACCACC

GATTGCTACAAGTACATAATCAGTAACATCCAAACCTCTAGTCTAGGTTTCATACATTCAGATATTGAGACACTCCCAAC

CAAAGATGCTGTTGAGAAATTAGAAGAGTTTGCCTCTATCCTATCCTTATCCTTGATTTTGGGAAAAATCGGCTCTATTA

CAGTTGTCAAAACTATGCCCATTAGCGGGGATTTTACCCAAGGCTTCATAGCCTATGCCATCCAATATTTTAGGGAGAGC

CTGCTTGCCTATCCGAGATATAGTAACTTCATCTCGACTGAGTGTTACCTTATTATGATAGGATTGAAGGCCAATCGATT

GATAAACCCAGAAGCCATTAAGCAAAGTATAATCAGGGCAGGAGTCAGGACTGCACCAGGACTTGTGAGCCATATATTAT

CAGAGAAACAGAAAGGTTGTATTCAATCTTTCTTGGGTGATCCTTATATCCAAGGAGACTTCAATAAGCACCTTAAATCT

CTAACCCCTATTGAGAAGATCCTGGTAAATTGTGGTCTCTCTATCAATGGAACAAAAATCTGTAGAGATCTAATCCACCA

TGATATCGCCTCCGGTCCAGACGGTCTCATGAGCTCTACAATCATTTTATATAGGGAATTGGCTCATTTCAAAGATAATA

TAAGGAGTCAGCACGGAATGTTCCACCCCTACCCAGTACTGGCCAACAGCAGGCAACGTGAATTAATCCTTCGAATAGCC

AAAAAATTCTGGGGGTATGTCTTGCTATATTCTGATGACCCGGCACTAATCAGACAGACAATCAAGAACTTGAAACGGAA

TCATCTAACCTTTGACTTACATAGTAACCCGTTTATTAAAGGGCTATCCAAAGCTGAGAAACTGCTGGTGCGGACGAGTT

CACTAAGAAGGGAATGGTTGTTCACTCTCGATACGAAAGAAGTAAAAGAGTGGTTCAAATTGGTGGGTTACAGTGCACTC

ATCAGAGGCTAGTTAGCTAGACATCTGACCCCTTCTCCTCCGCCATGAGCTCCCACTGACGATCCAAAAGATTAAAGAAA

ACTACATATTGGATAAGTATCTATTCCCAGCTTTGTCTGGT

>MK686066.1/goat/Burundi/2017-12

ACCAAACAAAGTTGGGTAAGGATAGATCTTATAATAACTATGAATTGCCACGCTTAGGATTAAAGACCCTACTGCCGGGG

GGAGGAGTAGGAGCAAGATCCCTGACTATGGCGACTCTCCTCAAGAGCTTGGCATTGTTCAAGAAGAACAGGGACAAAGC

GCCGACGGCATCAGGTTCAGGAGGAGCCATCCGGGGGATTAAGAATGTTATCATAGTCCCGATTCCCGGAGACTCATCCA

TCATTACCCGTTCAAGATTGCTCGACAGGCTTGTCAGATTGGCCGGAGATCCAGATATCAACGGGTCAAAGCTGACCGGC

GTGATGATCAGCATGTTGTCTTTGTTCGTAGAGTCACCTGGGCAACTGATCCAGCGGATCACAGATGATCCAGATGTCAG

TATCCGCCTTGTTGAGGTGGTCCAAAGTACTAGATCCCAGTCCGGATTGACCTTTGCATCACGTGGTGCTGATTTGGACA

ATGAGGCAGACATGTATTTTGCAACTGAAGGGCCCTCGAGTGGGAGCAGGAAAAGGATCAACTGGTTTGAGAACAGGGAA

ATAATAGATATAGAAGTACAGGATGCAGAAGAGTTCAATATGTTATTAGCCTCCATACTGGCACAAGTTTGGATTCTCCT

GGCCAAGGCAGTCACAGCACCAGACACTGCAGCTGACTCAGAGTTGAGAAGGTGGGTTAAGTACACACAACAAAGAAGGG

TGATTGGGGAATTCCGCTTTGACAAAGGGTGGCTAGATGCGGTCCGGAACAGGATTGCAGAAGATCTATCACTCCGGCGG

TTCATGGTATCTCTTATACTTGACATCAAAAGGACCCCTGGGAACAAGCCAAGGATTGCAGAAATGATCTGTGACATTGA

CAACTATATTGTCGAAGCCGGGCTCGCCAGTTTCATCCTTACTATCAAATTTGGTATTGAAACCATGTATCCTGCGCTAG

GTCTTCACGAGTTTGCCGGAGAGTTGTCCACTATAGAATCCTTGATGAATCTATATCAACAGCTAGGCGAAGTTGCACCC

TACATGGTGATTCTGGAGAACTCAGTTCAGAACAAGTTTAGTGCAGGAGCCTATCCACTTCTCTGGAGCTATGCAATGGG

TGTTGAGTTCGAGCTGGAGAACTCAATGGGAGGGCTGAACTTTGGTAGATCATATTTTGACCCAGCTTATTTTCGTCTCG

GACAGGAGATGGTCAGAAGATCCGCAGGAAAGGTCAGCTCTGCAATCGCAGCTGAGCTTGGTATCACAGCAGAGGAAGCT

AAACTAGTCTCGGAAATTGCCTCTCAGACTGGGGACGAAAGGACCGCTAGGGGGACCGGGCCCAGACAGGCGCAGGTTTC

CTTCCTCCAGCATAAAATAGGAGAGGGAGAGTCACATGCATCGGCGACCGGGGAAGAAGTCAAAGCTGCCACCCCAAATG

GGCCCGACGAAAAGGAAAAAACTCGGGCGCGCTCAGGAAAGCCAAGAGGAGGAACCCCTGACCAACTGCTCCTGGAAATC

ATGCCTGAAGACGAGGTCCCGCGAGGGTCTGGACAAAACCCTCGTGAGGCTCAAAGATCGGCCGAGGCACTCTTTAGACT

GCAGGCCATGGCCAAGATCCTAGAGAGCCAAGAGGAGGGAGAAGACAACAGCCAGACATACAACGACAAGGACCTCCTTA

GCTGAGAGGAAGAGCCCTCCATCAAAACCAGCAACAAGACAACGCCCTTCAGTATTATAAAAAACTTAGGACCCAGGTCC

AAGCAGTCACACTCCGGCACCCCAGCCGACCGAGCGGAGACAACCGATGGCAGAGGAACAAGCATACCATGTCAACAAGG

GACTGGAATGTATCAAGTCCCTCAAAGCCTCTCCCCCGGATCTCTCCACCATCAGAGATACCCTTGAAAGCTGGAGGGAG

GGGCTTGACACATCGGACCGTGCAGCACCGAACCCTGATATATCCAAGGGGGACCATCAGAATATCAACCAATCATGCCC

ACCAGCAATCGGATCAGGCGAAATCGACGTGTCTACTGAAAGTAATCTCGGATATAGAGAGATCAATCACGATGACAGTG

AGGCTAGACTCAGAGGAGTTCAAAGCAGAGGACCCAACCCTCAAATACAGCGTTACCATGTTTATAGCCACGGGGGTGAA

GAGATTGAAGGACTCGAGGATGCTGACTCTCTCATGGTTCAAGCAGATCCTCCACTTACTAACACCTTCAGTAGAGGAGA

GGATGGATCTGACGACAGCGATGTGGACTCTGGCCCAGATGATCCCGACAGAGATCCTCTATATGACCGGGGATCTGTTG

CCGACAATGATATCGTTAAGTCCACAGATGTCGAGAAATTAGAAGGTGACAACATTCAAGAAGTTCTTAACTCTCAGAAG

AGCAAGAGGGGAAAACTCCAGGGCGGGAAAACCTTGCGAGTCCCTGAAACGCCCGACATCAAGCACCCCAGACCATCAGC

CCAATCAATTAAAAAGGGCACAGACGGGAACTCAGTCTTATCTGGAACGGTGACAGAGTGTTCATCGATAAATGGTGCAA

CCCAAGCTGTGCCAGAGTCAAGATGGGAGTCATCAGAGCGAAATGCATTTGTGGAGAGTGCCCCCAAATCTGCGAGGAGT

GCAAAGACGATCCAGGAGTTGCCACAAGAATCTGGTACCATAGCATCACCGACTCAGCCTAAAGAAAATGACTCCGAGTA

TGAGTATGAGGATGACCTATTTATAGAGATGCAGGACATTCGTGCAAGCATTGCCAAGATCCATGATGACAACAAAACCA

TCCTCTCAAAGCTTGACTCTATACTATTATTGAAAGGAGAAATCGATACTATCAAGAAACAAATCAGCAAACAGAATATA

AGTATATCTACCATTGAGGGCCATCTTTCCAGTATAATGATAGCCATCCCGGGTTTCGGGAAGGAGGCCAAGGACCCAAC

ATCCGAGGTTGAGTTAAACCCGGATTTAAGACCTATAATCAGCCGTGATTCTGGCAGGGCTCTCGCGGAGGTCCTCAAGA

AACCTGCTGTTGATAGGTCTCCTAAAACCGGATTCAAAGTCAACTCCGGTTCAAAGGGTCAGCTCCTCAAGGATCTCCAA

CTGAAACCCGTTGACAAACAGGCAAGTTCTGCCATAGGGTTTGTCCCGTCTGACCATGGATCATCCAGAAGTGTCATCCG

CTCTATAATCAAGTCAAGCAAGCTCAACACTGATCACAAAGACTATCTTTTAGATTTACTGAATGATGTGAAAGGCTCCA

AGGATCTCAAAGAATTCCACAAGATGCTAACAGCTATTCTTGCTAAGCAACCGTAATGCACCGCCTAACCAATTTCTCAT

GCTCAGCTGAAAACAGCCCCACAACCAGGCTATTACAAAAAACTTAGGAGCAAGGGCAACCGAGCTTCGCAGACAAGATG

ACCGAGATCTATGACTTCGACAAATCAGCATGGGACGTCAAGGGATCAATAGCTCCCATAGAGCCCACCACTTATCACGA

TGGTCGACTGATACCCCAGGTGAGAGTCATCGATCCTGGGCTAGGAGACAGGAAAGATGAATGCTTCATGTACTTGTTTC

TTCTAGGGGTGATTGAGGATAATGACCCCCTGTCTCCCCCAGTAGGAAGAACCCTTGGTTCCTTACCTCTAGGGGTCGGT

AGGTCAACTGCTAAGCCAGAAGAACTATTAAGGGAGGCCACTGAACTTGACATAGTGGTGAGACGCACAGCCGGACTAAA

CGAGAAGCTGGTATTTTACAACAACACTCCGCTGTCTTTGTTAACACCTTGGAGGAGAGTCCTGACAACCGGAAGTGTGT

TCAGCGCTAACCAGGTTTGTAACGCAGTCAATCTAATCCCACTCGATACTCCCCAGAGATTCAGGGTTGTGTACATGACC

ATCACTAGATTGTCTGACAATGGTTATTATAGTGTCCCCAAGAGAATGTTAGAGTTCCGCTCAGCCAATGCCGTCGCCTT

CAATATCCTGGTGACACTGAGAATTGAGAATGGGACAAACCCTAGACGATACATAGTCGGCTCATGGGAGAACCCAGAGG

TCACATTCATGATACATGTGGGCAACTTTAGAAGAAAGAAGAACGACGTATATTCTGCTGATTACTGCAAAATGAAGATT

GAGAAGATGGGTCTGGTTTTTGCTTTGGGAGGAATAGGGGGAACAAGTCTTCACATTAGAAGCACCGGGAAAATGAGCAA

GACCCTCCATGCACAGCTGGGGTTCAAGAAAATCTTATGCTACCCCTTAATGGATATTAATGAGGATCTCAATCGATATC

TTTGGCGGGCAGAGTGCCGAATAGTCAAAATTCAAGCTGTCTTACAGCCTTCAGTACCCCAAGAATTCCGTGTTTACGAT

GATGTCATCATTAATGATGACCAAGGCTTGTTCAAGATCCTGTAACTCACTCACAGAACAATGGACCCGCCTCCAACAAA

GGACCACCCCCTCCCTCTCCCAGAAAACACACCCCGAGCCCCCCCCAAAAAACGCCGACATAGTCATCCTCCCGGTAGGA

CAAGGATGACCCCCTCATCGGACCCAGTCT--GCGGGGGGGCCCCCCCCGTGGCCCGCCGAGCGCCGACCGCGCAGGGCA

CCGAGGCCCCACTGGGAGGCTCCACGGCAGACCAGGCACCCCCTTCCTCC--GCGTCACGCCCCAAACATGTCCAAGGCC

AGGAGCCGAGCACCCCCAGCCAAGCCAACGACAGGACAGACACCCAAGTAGGGGTCCTC--GACCGAAAAGAATACCAGA

GTGGGCTCCGCCGACCCAGCACAGATTTGGCCCCAACAAGGCAGATCCCCAAAACGGAATCTCCCAGTGCACTCCACTCA

TCATCAAACAAAACTTAGGGGCCAAGGTCCGACGACTGACCCCACTCCGCCAAGCAGCAGGCCGCACGAAGGAAGCAGAC

AAGACAGGCCCGGAAGGCAGACACCCGACCACAGAGACATCCAAGCCTCGGGGAGGAGG-CCTCCCCCCTCAAAGCGGAG

GAGCGGGAGGGAAACACCAGGGACAACCCGACCGGGAGCCGGGACCCGGAGGCTCCAACGACAACCGGGCCCGGCACACA

CCCGGCACCCCCGGCGAGACAGAGGAGAGCCCGAGCCCGCCTGGACCCGACGGCAGAGGATCCAATCTCGCCACAAAGAG

AAGCGAGGAGATT--------TCCCGCCGCACACAGTCGGCCGACAACTCACAGGAGGACCATCTCCCGGGAGGATAAGC

GGCCGTCCCACAACTGGGCCGCCACCCCCCAGAGGAGGACGCCCCGGAGGTCTCAAGATATCCCCCCGTCCTCGACCACC

GGGGGAGGGGCCACACCGACAAAGCCCAGAGCCACCGGGAGCCGACCAAGGGTCCCCCCCGGCCCCGCTCGGCACCCCCC

CGACGGGACCCTAACAAGACCTCTCGACCCAACAACCCCCACACGCGCCAAGCCAGAAACCTGCCAGTGGACATCCTGCA

CATTCATGCACAAACATCATGACAAAAGTCGCAATCTTGACATCCCTGTTTCTCCTCACCAGCACTGTCGTATCCCAGAT

TCACTGGGGCAACTTATCCAAGATCGGGATTGTGGGGACAGGAAGTGCCAGTTATAAGGTGATGACTAGGCCGAGCCAAC

AGACCCTGGTTATAAAGCTAATGCCAAATATAACAGCCATCGACAATTGTACGAAATCAGAGATTTCGGAATACAAAAGA

CTGCTGATCACGGTGTTAAAACCTGTCGAGGATGCTCTGTCAGTGATAACCAAGAATGTAAGACCAATTCAAGCTCTAAC

ACCCGGTCGCAGGACTCGCCGTTTTGCCGGGGCTGTCCTGGCCGGAGTGGCGCTTGGGGTTGCAACGGCTGCTCAGATAA

CTGCCGGAGTCGCACTCCATCAATCTTTGATGAACTCCCAAGCAATTGAAAGTCTAAAAGCCAGTCTTGAGAAGTCTAAT

CAAGCAATAGAAGAAATCAGACTTGCAAATAAGGAAACCATACTGGCAGTACAGGGTGTCCAAGATTACATCAACAACGA

GCTAATCCCTTCTGTTCATAAGATGTCATGTGAGTTGGTAGGTCACAAACTCAGTCTCAAGCTCCTTAGGTACTACACCG

AGATCTTGTCTGTATTCGGGCCCAGCCTTCGAGACCCGATAGCTGCTGAAATATCAATCCAGGCACTCAGTTATGCACTA

GGTGGAGATATCAACAAGATCCTGGATAAGCTTGGATATAGCGGCGAGGACTTCCTTGCTATCTTGGAAAGCAAGGGGAT

AAAGGCCCGAGTCACCTATGTGGATACAAGAGATTACTTTATAATCCTTAGCATAGCCTACCCGACCTTATCTGAGATCA

AAGGGGTGATAGTTCATAAAATGGAAGCTGTGTCATACAACATCGGGGCACAAGAATGGTACACTACTATCCCTAAATAT

GTAGCCACTCAGGGATACTTGATATCAAATTTTGACGAGACGTCATGCGTGTTCACTCCAGGAGGGACAGTCTGCAGTCA

GAATGCTTTATACCCAATGAGCCCGCTGCTTCAGGAGTGTTTCAGGGGGTCTACAAAATCGTGCGCCAGAACCCTAGTTT

CAGGGACTATAAGTAATCGATTTATCCTGTCAAAAGGGAACCTGATTGCCAATTGTGCATCAGTCCTGTGCAAATGTTAC

ACAACGGAGACGATTATCAGCCAAGATCCTGATAAATTACTAACTGTTATAGCCTCTGATAAATGTCCTGTAGTTGAAGT

GGATGGAGTGACGATACAGGTCGGCAGTAGGGAGTACCCAGATTCTGTGTACCTACATAAAATAGACTTGGGCCCAGCCA

TCTCTCTAGAAAAACTGGATGTAGGCACCAATTTAGGCAATGCAGTTACTAGGCTGGAAAATGCAAAGGAACTCCTAGAT

GCATCAGATCAGATATTGAAGACTGTTAAGGGGGTACCCTTCAGCGGCAATTTGTACATAGGACTGGCAGCTTGCATCGG

AGTATCTCTAGGGCTTGTCACATTAATATGCTGCTGTAGGGGGCGGTGTAGGAACAAGGAGATTACTACCCCTAAGATCA

ACCCAGGGCTCAAACCAGACCTGACCGGGACCTCAAAGTCGTACGTAAGATCACTGTAATCAGAATAGCCCGAATCATCC

AGCGTCACACACAGACATGTGCAACCCAAGCAGTCAGGGGACTCAGAGGGTTCAACCTCCGATCCTCAACCTGATCCCAT

TCCAAGCCCCATTGCACATCGGTTATTAAATAAAACTTAGGACGAAAGGTTAGTCACCATGTCCGCACAAAGAGAGAGGA

TCAATGCCTTCTACAAAGACACTCCTCACAACAAGAACCATAGGGTGACCCTAGACAAGGAGCGTCTGACTATTGAAAGA

CCCTATATCCTGCTTGGGGTCCTACTGGTAATGTTTCTGAGTCTGATTGGGTTGCTGGCCATTGCAGGGATTAGACTTCA

CCGAGCCACCGTGGGAACTGCAGAGATCCAGGGCCGGCTTAATACAAACATTGAGTTGACTGAATCTATTGATCATCAAA

CTAAGGATGTCCTAACTCCCCTATTTAAAATCATCGGTGATGAAGTCGGCATAAGGATTCCACAGAGATTCAGTGACCTT

GTCAAGTTCATCTCCGATAAGATTAAGTTCTTCAACCCTGACAGAGAATATGATTTCAGGGACCTCCGGTGGTGTATGAA

TCCTCCAGAGAGAGTTAAAATAGATTTTGATCAGTTTTGTGAATACAAAGCCGCGGATAAGTCGATTGAACATATATTTG

AATTACCACTCAACAGGTCAAAAGAGTTGCGATCGTTGACTCTTGGGCCTGGAACGGGCTGTCCCGGCAGGGCAGTAACA

AGGGCCCAATTCTCAGAGCTTACTCTGACTCTGTTGGATCTGGATCTCAATATGAAGCACAACGTGTCCTCAGTGTTTAC

CATAGTTGGAGAGGGATTATTCGGAAGAACATATACTGTCTGGAGATCATATGCCGGAAATCTGAGTACCGCCTCAGGTC

TCGGCCATTTTTTAAGAGTCTTCGAGATCGGACTGGTAAGAGATTTTGGGCTAAGTGCCCCTGTTTTCCATATGACTAAC

TATCTCACGGTGAACATGAGTGATGACTACCGGAGCTGTCTATTAGCAGTAGGGGAATTGAAGCTGACAGCTCTATGCAC

CCATTCTGAGACTGTGACTCTGAGTGAGAGAGGAGTTCCTAAAAGAGAGCCTCTTGCGGTTGTAATACTTAATCTAGCCG

GGCCTACTCTAGGGGGCGAACTATACAGTATTTTGCCTTCCTCTGACCTCATGGTAGAAAAACTCTATTTCTCATCAAAT

AGGGGGATTATCAAAGACAACGAGGCCAATTGGGTAGTACCATCTACCGATGTACGTGATCTTCAAAATAAAGGAGAATG

CCTGGTAGAGGCATGCAAAACTCGACTTCCTTCATTTTGCAATGGTACAGGGTTAGGCCCATGGTCAGAGGGGAGAATCC

CTGCCTACGGGGTGATCAGAGTCAGTCTTGACGTAGCTAGTGATCCAGATGTGGTTATTACCTCAGTGTTTGGCCCCTTG

ATACCTCATCTATCTGGCATGGATCTTTACAACAACCCATTCTCCAGAGCTGTGTGGTTGGCTGTACCACCTTATAAGCA

GTCATTTCTGGGAATGATAAATACAATTGCTTTCCCAAGTAGAGCAGAGGTTATGCCGCATATTTTGACCACAGAGATCA

GAGGGCTTCGGGGTCGTTGTCATGTTCCCATAGAATTGTCCCGCAAGACTGATGATGATATTAAGATTGGCTCCAACATG

GTTGTATTACCGACGAAGGACCTGAGGTACATTACAGCCACTTACGATGTTTCCAGGAGCGATCATGCAATCGTGTATTA

TATCTATGACACAGGTCGCTCATCATCTTACTTCTTCCCCGCTCGATTGAAGCTCAAAGGTAACCCTCTCTCTATGAGGA

TAGAGTGTTTCCCCTGGCATCATAAGGTATGGTGCTACCATGACTGTCTCCTATACAACACCACCACAAACGAAGAAGTT

CATACGAGAGGGCTAACTGGCATAGAGGTAACATGTAATCCAGTCTGAGCAGAGCTGCGACCATCAATCAAGCAAGCTTC

CGAGCCATTCTCAAGCCAAAGCATCACACCTAGGGGCACTCAACAGCACAACCCAGCCGACAGTGTTATAAAAAACTTAG

GAGCCAGGGCTATTGGAGCCATGGACTCCCTGTCAGTCAACCAGGTCCTATACCCTGAGGTCCATTTAGACAGCCCCATT

GTCACAAACAAGCTAGTTGCCATCCTTGAATACTCGGGCATCGACCACAACTATGTTCTTGAGGACCAGACTCTTATCAC

GAATATTAGATATAGGTTGGGATGCGGTTTTTCAAATCAAATGATCATCAATAACAAGGGGGTAGGTGAAACAGTCAATT

CCAAACTTAAAAGTTACCCCCGTAATTGTCATGTCATATATCCAGATTGCAATAAGGATCTGTTTTATATCAGGGACGGC

TGCATATCTAGGAAGCTCTCTGAGCTATTCAAGAAAGGTAATTCCCTGTACTCTAAGATAAGTCACCAGGTACTAGATTG

TCTCAGAAGAGTTAACGGCAAAATGGGCCTGGGCACAGATCTTAGCCATGGCCTGAAGGAGGGTATCCTCGATTTAGGGT

TGCACATGCATAGCTCTCAGTGGTTTGAGACCTTCCTGTTCTGGTTCACTATCAAGACTGAAATGAGGTCAATGATCAAA

GAACAATCCCATATATGCCACAAGAGGAGGTATAACCCAACCTTTGTCTCAGGAGACCCACTCGAGGTGCTTGTATCACG

GGACCTCGTTGTGATAATTGATAAAACCACTCAGTGTGTCTATTACCTGACATTCGAGCTGGTCCTCATGTATTGCGATG

TCATAGAGGGCAGACTTATGACGGAGACTGCCATGGCTATAGATCAGAGGTATTCAGAATTGCTAAGCCGGGTCAAATAC

TTATGGGATCTTATTGATGGGTTCTTCCCAACACTAGGCAACGCCACATACCAAGTTGTTGCTCTGCTTGAACCGTTATC

GTTGGCTTACCTTCAACTTCAAGATGTCACTCTGGAGTTAAGAGGTGCATTCTTGGATCACTGCTTCAAAGAACTCTACG

AGATACTGGAACATTGTGGCGTTGACACGGAAAGAACCTACGATTCCATCATCGAAGGATTGGATTACGTATTTATCACC

CACGATATACACTTAACTGGGGAGATTTTCTCCTTTTTCCGGAGTTTCGGACATCCCCGCCTTGAGGCGGTCACTGCCGC

AGAGAATGTCAGGAAACATATGAACCAACCTAAGGTAATCAGTTATGAGACTATGATGAAAGGGCACGCAGTGTTTTGCG

GGATCATCATAAATGGGTTTAGGGACCGCCATGGCGGCAGCTGGCCCCCCGTTGTGTTGCCAGAACATGCCTCTGCTGCA

ATTCGGAATGCGCAGGCATCCGGTGAAGGGTTGACCCATGACCTGTGTATAGACAACTGGAAGTCCTTCGTTGGATTCAG

GTTTGGCTGTTTTATGCCGCTCAGCTTAGACAGTGACCTGACCATGTATCTCAAAGACAAAGCACTAGCTGCACTGAAAA

ATGAGTGGGATTCAGTTTATCCAAAAGAATACCTCCGCTATAACCCACCCAGAGGGACGGAGTCAAGGCGACTGGTGGAG

GTGTTCCTGAATGACTCCAGCTTTGATCCTTATAACTTGATAATGTACGTGGTGAATGGCTCCTACCTTGAAGACCATGA

GTTTAATCTCTCATACAGCTTGAAAGAGAAAGAGATAAAAGAGACAGGGCGACTGTTTGCAAAAATGACTTACAAGATGC

GGGCCTGCCAGGTCATTGCGGAGAATCTGATATCAAATGGGGTTGGGAAGTATTTCCGAGACAATGGAATGGCGAAAGAT

GAGCACGACCTGACAAAGGCCCTTCATACCTTGGCAGTCTCAGGTGTTCCTAAAAACAACAAGGATAATCACCGGGGTGG

GCCTCCTAGAAGGACCACAAGCCAAAAGATGAGATCAGATCGGGGCATCACCAGACAAGATAGGGACAAAATACAGAGAG

AGCCCATGTATAACTATCTGCGATGCCAACCAGTCAGCCCTGACCAGGGTGAGTCATACGAGACTGTTAGTGCATTCATC

ACCGCTGACCTGAAGAAATATTGCCTGAACTGGAGATACGAGACAATCAGCATATTTGCACAGAGACTAAATGAGATTTA

TGGACTACCATCTTTCTTCCAATGGTTACACAGGGTATTAGAAAAATCTGTGCTGTACGTCAGTGACCCACATTGCCCTC

CCGACTTAGACGATCATATCCCTCTGGACAGTGTCCCCAATGCCCAAATATTCATCAAGTACCCGATGGGCGGGATAGAA

GGTTATTGTCAAAAATTATGGACAATCAGTACTATACCATACTTGTATCTAGCAGCCCATGAGAGCGGAGTAAGAATCGC

CTCTCTAGTGCAAGGTGATAATCAAACAATTGCAGTGACAAAGAGAGTCCCGAGTTCCTGGCCTTATTCACTAAAAAAGA

GGGAGGCATCTAGAGCAGCTCAGAATTACTTCGTGGTTTTAAGACAGAGGTTGCATGATGTAGGTCATCACTTAAAGGCC

AATGAAACTATAGTATCTTCTCATTTTTTTGTATACTCAAAAGGGATTTATTATGATGGCCTGTTAGTATCACAATCACT

AAAGAGCATTGCCAGATGTGTCTTCTGGTCCGAGACTATTGTGGATGAAACCAGAGCCGCCTGCAGCAATATTGCAACAA

CTGTCGCCAAGAGTATAGAGAGGGGTTATGACAGGTACCTCGCTTACTCCTTGAACATACTCAAAATTCTCCAACCCATT

CTTATATCCCTTGACTTCACTATCAACACAACAATGACTCAAGATGTCGTGGCACCGATCATCGAGAACGGTGACCTGCT

AACAAGGATGGCACTTTTGCCAGCACCCATCGGAGGCCTTAACTACCTCAACATGAGCAGGTTATTTGTGAGAAATATCG

GTGACCCAGTCACTTCCTCTATAGCCGACCTGAAGAGGATGATAGGCGCTGGACTAATGCCAGAAGAAACATTGCACCAG

GTGATGACCCAGACCCCGGGAGAATCATCTTACCTTGATTGGGCAAGTGACCCTTATTCTGCCAACCTAACCTGTGTACA

GAGTATAACTCGTCTTCTCAAGAACATCACTGCAAGGTACATTTTAACCAGCAGCCCAAATCCGATGCTAAAAGGGTTAT

TCCATGAAGGGAGCAGAGATGAGGATGAAGAGCTTGCGAGTTTCCTGATGGATCGGCATATAATTGTTCCCAGAGCTGCA

CATGAAATCTTAGACCATAGTATAACTGGGGCAAGAGAAGCTATTGCCGGTATGTTAGATACCACCAAGGGTCTGATTAG

AACAAGTATGAAACGAGGTGGCCTTACGCCACGGGTCTTAGCTCGCCTTTCCAATTATGATTATGAACAGTTTAGATCCG

GTATAACATTGCTGACAAAGAAGGGACAGTGTTATCTCATTGACAAAGATTCATGCTCGGTACAGCTTGCTATAGCCCTG

AGGAGTCATATGTGGGCTAGGTTGGCCCGCGGGAGACCTATCTATGGGTTGGAAGTGCCTGATGTACTGGAATCGATGAA

TGGCTATCTTATCAAACGCCATGAGGCATGTGCTATTTGCGAGACGGGCTCAGGTCATTACGGGTGGTTCTTCGTCCCTG

CAGGGTGCCAGCTTGACGATGTCTCGAAAGAGACTTCAGCTCTTCGTGTGCCTTACGTTGGATCAACCACAGAGGAAAGG

ACAGACATGAAACTTGCCTTCGTTAGATCTCCGAGTCGATCCCTCAAATCAGCAGTCAGAATTGCTACCGTTTATTCATG

GGCCTATGGGGATGATGAAAAATCATGGAGTGAAGCCTGGACGTTGGCAAGGCAGAGAGCCGACATCACCCTAGATGAGT

TGAGGATGATCACCCCAGTCTTTACGTCGACCAACCTAGCCCATCGGTTGAGGGATCGGAGCACCCAGGTGAAATACTCG

GGGACATCCCTTGTGAGGGTTGCGAGATACACAACCATTTCCAATGATAATTTGTCATTTGTGATATCTGAGAAGAAAGT

AGATACCAACTTCATTTACCAGCAAGGGATGCTCCTCGGTCTTGGGATCCTTGAGAATCTTTTCAGGTTGGAGACCACCA

CGGGGGTATCCAACACAGTGCTACATCTGCACGTGGAGACAGAATGTTGTGTTGTACCTATGATTGATCACCCAAGAATA

CCAAGTCTCCGTAACATTAAGGTTACGAATGAGCTATGTACAAACCCCTTGATCTACGACAAGTCTCCTATCATAGAACA

CGATGCAACTCGATTATACTCACAAAGCCACAGAAGACATCTGGTGGAATTTGTTACCTGGTCAACAAGTCAGCTTTACC

ATATATTAGCTAAGTCTACAGCAATGTCCATGATTGAGTTGATTACAAGATTCGAGAAGGATCACATGAATGAAATAACC

GCCTTGATTGGCGATGACGACATCAACAGTTTTATAACAGAGTTCCTGCTAGTAGAACCTAGACTGTTTATAGTCTACCT

TGGCCAGTGTGCTGCCATTAATTGGGCTTTTGATATACATTATCATCGCCCATCGGGCAAATACCAGATGGGGGAGCTTC

TCTACTCTTTACTCTCTAGGATGAGCAAAGGAGTATATAAGATCTTTACCAATGCTCTGAGCCACCCCAAAGTTTACAAG

AAATTTTGGCGAAGCGGTGTCATCGAGCCGATTCATGGCCCGTCGTTGGACACACAGAATCTACATGTCACTGTCTGTGA

TATGATATATGGATCCTACGTCACATATCTGGATCTTTTGCTGAATGATGAGCTAGATGATTATACTTATCTGCTCTGCG

AAAGTGATGAGGACGTGGTCACAGACAGGTTTGATAACATCCAAGCCAAACATCTCTGTGTATTGGCCGATGTGTATTGC

AGCTCCAAAAGGTGTCCCTCAATAATCGGGATGTCTCCTATAGAAAAATGTACCATCCTCACACATTACATAAAAGGGGA

ATCAATACAGTCTCCGTCTGGGGCCTCATGGAACACTGATCCTCTTGTAGTAGATCATTACTCATGCTCTCTGACATACC

TTCGTCGCGGTTCCATCAAACAAATCAGGTTGAGAGTAGATCCTGGGTTTGTATTTGAGGCATTGACAGACATCGACGTC

AAACAGCCTCGCAAGGCTAACACTGATGTATCGGTTGTGGGGTTGACTGATTTCTCTCCCCCTTGGGATAATATTGGTGT

TTTTCTAAGGACTATCAACACCTTGAGACACAATCTGCCAATCACTGGGACCGGGGTCTTAAACTATGAGGTCCACGCTT

ATCGTAGAATTGGTTTGAATTCATCAGCATGTTATAAAGCTGTAGAGATCTCCACGTTAATCAAGCCATCCTTAGAAGTC

GGAGAGCATGGGTTGTTCTTAGGGGAAGGTTCCGGTTCTATGCTGGCTGCGTACAAGGAAATTCTTAAATTAGCCAACTG

TTATTACAACAGCGGAGTAATAGCAGAGGGCAGAGCCGGACAAAGAGAAATATCCCCTTACCCATCAGAGATGAGCCTAG

TAGAGAATCAAATGGGGTTAGAAAAGAGTGTTAAAGTGCTGTTCAACGGCAAACCTGAAGTAACCTGGGTAGGGACTACC

GATTGCTATGAATATATAATCAGTAACGTCCATACCTCTAGTCTGGGTTTCATACACTCGGATATTGAGACTCTCCCAAC

CAAAGATGCTGTTGAGAAATTAGAAGAATTTGCCTCTATCCTATCCTTATCCCTAGTTTTGGGAAAAATTGGCTCTATTA

CAGTTATCAAAGTTATGCCTATTAGCGGAGATTTTACTCAGGGCTTCATAAGCTATGCCACCCAGTATTTTAGGGAGAGC

CTGCTTGTCTATCCGAGATATAGTAATTTCATTTCAACTGAGTGTTACCTTATTATGATAGGATTGAAAGCTAATCGGTT

GATAAACCCAGAAGCCATTAAGCAAACCATAATCAGAGCAGGGACTAGAACTGCACCGGGACTAGTGAGCCATATATTAT

CAGAGAAACAAAAAGGGTGTATCCAAGCTTATCTGGGTGATCCTTACATCCAAGGAGACTTCAATAAACACCTTAAATCT

CTAACCCCTATCGAGAAAATCCTAATTAATTGTGGTCTATCGGTCAATGGCACAAAAATCTGTAGGGATCTAATCCACCA

TGATATCGCCTCTGGTCCGGATGGTCTGATGAGCTCCACAATAATCTTATACAGGGAGCTGGCTCATTTCAAAGACAATG

TGAGGAGTCAGCACGGGATGTTCCACCCCTACCCAGTACTGGCCAGTAGCAGGCAACGCGAATTAATTCTTCGAATCGCC

AAGAAATTCTGGGGTTATGTCTTGCTATATTCTGATGATCCGGCACTGATCAGGCAAACAATCAAGAATTTGAAAAGGAA

CCACTTAACCTTTGACTTACACAATAACCCGTTTATTAAGGGCCTGTCCAGAGCTGAGAAATTGCTAGTGCGGACAAGTT

CACTCAGAAGAGAGTGGTTGTTCACTCTCGAAACGAAGGAATTGAAAGAATGGTTTAAACTGGTGGGCTACAGTGCACTT

GTCAGAGGTTAATTAGCTATACATCTGTCCCCTTCTCCTCCACCATGAAATTCCACTGGCAATCTAATAGATTAAAGAAA

ACTACAGATCAGATAAGTATCTATTCCCAGCTTTGTCTGGT
